# Terminal regions of a protein are a hotspot for low complexity regions (LCRs) and selection

**DOI:** 10.1101/2023.07.05.547895

**Authors:** Lokdeep Teekas, Sandhya Sharma, Nagarjun Vijay

## Abstract

A majority of the protein-coding genes consist of low-complexity regions (LCRs) in eukaryotes. Volatile LCRs are a novel source of adaptive variation, functional diversification, and evolutionary novelty. LCRs contribute to a wide range of neurodegenerative disorders. Conversely, these regions also play a pivotal role in critical cellular functions, such as morphogenesis, signaling, and transcriptional regulation. An interplay of selection and mutation governs the composition and length of LCRs. High %GC and mutations provide length variability because of mechanisms like replication slippage. The selection is nearly neutral for expansion/contraction within the normal range and purifying above a critical length. Because of the complex dynamics between selection and mutation, we need a better understanding of the coexistence and mechanisms of the two. Our findings indicate that site-specific positive selection and LCRs prefer the terminal regions of a gene and co-occur in most of the Tetrapoda clades. Interestingly, positively selected sites (PSS) are significantly favored in LCRs in eight of the twelve clades studied. We also observed a significant favor of PSSs in the polyQ region of MAML2 in five clades. We also found that PSSs in a gene have position-specific roles. Terminal-PSS genes are enriched for adenyl nucleotide binding, while central-PSS genes are involved in glycosaminoglycan binding. Moreover, central-PSS genes mainly participate in defense responses, but terminal-PSS genes are non-specific. LCR-containing genes have a significantly higher %GC and lower ω (dN/dS) than genes without repeats across the Tetrapoda clade. A lower ω suggests that even though LCRs provide rapid functional diversity, LCR-containing genes face intense purifying selection.

## Introduction

Proteins do not contain amino acids in equal proportions as the theoretical 5% usage would suggest. Taxonomic dependence yields species-specific amino acid frequencies in the proteome. The regions containing fewer amino acid types than expected, defined by a set of compositionally biased residues and having lower amino acid residues diversity that differs from proteins’ average amino acid composition, are known as low-complexity regions (LCRs) (Golding 1999; Radó-Trilla and Albà 2012; Peng et al. 2015; Mier and Andrade-Navarro 2020; Lee et al. 2022). LCRs exhibit a range of amino acid compositions ranging from an aperiodic accumulation of limited amino acids to stretches consisting of a single amino acid residue (homopeptide repeat) (Haerty and Brian Golding 2010; Radó-Trilla and Albà 2012; Chaudhry et al. 2018). Low complexity regions are prevalent in eukaryotic proteins, with approximately 18-20% of human proteins featuring at least one amino acid repeat of length five residues or more (Karlin et al. 2002; Albà et al. 2002; Radó-Trilla and Albà 2012). The frequency of repeat-containing proteins is much higher in mammals compared to birds, fishes, or amphibians (Faux et al. 2005). LCRs have a lower propensity to form structured domains and are much less evolutionarily conserved than globular domains, which are the most common protein domains (Mier and Andrade-Navarro 2020; Kastano et al. 2021). Even though the LCRs vary in composition/purity of the stretch, studies are primarily focused on single amino acid (homopeptide) repeats, probably owing to their ease of search in the protein datasets (Albà and Guigó 2004; Faux et al. 2005; Radó-Trilla and Albà 2012).

LCRs of various types participate in essential biochemical functions like cell signaling and protein kinases (Hancock and Simon 2005), RNA processing (Kobe and Deisenhofer 1995), transcription regulation (King et al. 1997; Kashi and King 2006; Lynch and Wagner 2008), neurogenesis, DNA repair, recombination (Karlin et al. 2002), and cell adhesion (Myers et al. 1994). A series of studies about the consequences of LCRs in neurodegenerative diseases shed light on their functional implications (Deininger and Batzer 1999; Marcotte et al. 1999; Karlin et al. 2002; Delucchi et al. 2020). Later studies revealed their role not only in diseases but also in evolutionary contexts. Repeats of gene *RUNX2* are linked with facial structure and morphology across vertebrates and are intensively studied in dogs (Fondon and Garner 2004; Newton and Pask 2020).

Processes like replication-slippage, unequal crossing-over, and gene conversion (recombinational mechanisms) make LCRs unstable and lead to length expansion and contraction (Hancock and Simon 2005). This volatility leads to numerous disorders but opens avenues for evolutionary and morphological diversity within a short evolutionary time (Newton and Pask 2020; Deryusheva et al. 2021). Variable regions work as ’tuning ’knobs’ that create frequent, reversible, and site-specific genetic variations, which helps genomes and genes for efficient adaptation (King et al. 1997). Variable LCRs are known to participate in immunological pathways also. Leucine-rich repeats are a key structural component of Toll-like receptors in animals’ innate immunity and adaptive immunity in jawless fishes (Pancer and Cooper 2006; Deng et al. 2013; Berglund et al. 2015; Persi et al. 2016). Moreover, low-complexity sequences evolve more rapidly than the remaining gene sequence and thus promote evolutionary novelty (Huntley and Golding 2000; Albà et al. 2007). Low-complexity regions also generate polymorphisms within species and play an equally relevant role in microevolution (Wren et al. 2000; Lavoie et al. 2003; Albà et al. 2007). In an interesting study on the Great Pyrenees dog, the authors found an association between the polydactyly of the bilateral rear first digit and deletion within a P/Q repeat in the *ALX4* gene (Fondon and Garner 2004).

Although amino acid simple repeats are a subset of LCRs, the mechanisms of origin and maintenance vary. The “slippage” model is one of the possible mechanisms for the emergence of amino acid repeats (Levinson and Gutman 1987). This model posits that errors occurring during DNA replication or repair can trigger DNA polymerases or other repair enzymes to slip, leading to the insertion or deletion of a few nucleotides. Such events can alter a gene’s coding region, potentially inducing the expansion or contraction of short nucleotide repeats. Ultimately, these changes may culminate in the emergence of amino acid repeats within the protein sequence (Hancock and Simon 2005). The abovementioned mechanism will give rise to homogenous DNA sequence repeats due to slippage in triplets in the coding region. However, heterogeneous DNA sequences also contribute to conserved amino acid homopeptide repeats (Alba et al. 1999; Mularoni et al. 2007). The existence of homopeptide repeats through heterogeneous DNA sequence imply origin mechanisms other than slippage and maintenance by selective forces for the preservation of homopeptide repeat (Rorick and Wagner 2010; Radó-Trilla and Albà 2012).

In contrast to homopeptide repeats, most low-complexity regions are maintained by a complex interplay of mutations and selective forces (Radó-Trilla and Albà 2012). In a recent study, authors compared low-complexity regions in protein sequences to simulated proteomes and found that repetitive regions that have simple, repeating patterns could have evolved through neutral processes. However, regions that have cryptic, compositionally biased regions could not have evolved through these processes alone. The study suggests that other biological factors and processes are likely at play in the evolution and maintenance of these complex repetitive regions (Enright et al. 2023).

Conservation of LCRs across species suggests they are under natural selection and contribute to adaptation (Mularoni et al. 2010; Persi et al. 2016). Studies also show repeats require a minimum length to create a functional impact, and longer repeats are more prone to disruption of function (Salichs et al. 2009; Mularoni et al. 2010). In an intriguing study on nuclear proteins, researchers found that protein translocation requires a minimum repeat of 6 histidine residues (Salichs et al. 2009; Mularoni et al. 2010). Repeats are functionally efficient in a medium range, with small-sized repeats being less functional and longer ones forming protein aggregates and reducing function as proposed for RUNX2 QA repeat (Newton and Pask 2020). LCR length variability leads to morphological variability, as demonstrated in *ALX4* and *RUNX2* genes (Fondon and Garner 2004). Thus, LCRs generate rapid genetic variability for selection to act on it. This interaction between selection and LCR length variability can give rise to morphological and evolutionary novelty in a short time.

Even though some studies have focused on genome-wide patterns of LCRs across clades (Albà and Guigó 2004; Teekas et al. 2022; Lee et al. 2022), most of the studies either focus on limited gene sets, specific diseases, or limited species (Huntley and Clark 2007; Lobanov et al. 2016; Newton and Pask 2020). These focused studies provide a detailed mechanism of protein repeat function in particular species or gene sets. Still, they fail to dispense a general mechanism for the evolution and expansion of LCRs across clades. Despite the variation in origin, maintenance, and selection pressure based on the purity of composition of low-complexity regions, there is a shortage of studies that specifically address this. Furthermore, LCRs and selection work in a tightly entangled dynamics but, empirical studies to show an overlap of selection and LCRs are scarce. In the study, we first investigate the distribution pattern of LCRs as the purity of the LCR changes across the Tetrapoda clade. We achieve this by generating results for a gradient of LCR purity from 0 to 100%, with purity binned at intervals of 10. However, for our comparative phylogenetic analysis of LCRs and positively selected sites (PSSs), we only selected LCRs with a purity greater than 70%. This is done to include degenerated LCRs across different species while reducing the likelihood of generating false positives due to an overabundance and overlap of LCRs and PSSs. Later, we focus on the prevalence of positively selected sites and LCRs occurrence across the genes. We observed that the abundance pattern of PSSs and LCRs in genes is consistent across all clades. Our study also found seven of the twelve clades show significant co-occurrence of LCRs and PSSs. In two-thirds of the considered clades, the overlap between positively selected sites (PSSs) and low-complexity regions (LCRs) in a gene is higher than what could be expected by chance. We expanded our investigation by examining the genes containing positively selected sites that are favored in low-complexity regions across clades. Interestingly, we find a significant favor of positively-selected sites in the poly-glutamine repeat of MAML2 protein in five out of seven clades containing LCRs and PSSs. We also find the ω (dN/dS: non-synonymous substitution rate/synonymous substitution rate) of LCR-containing genes is significantly lower than genes without LCRs across all the clades considered from Tetrapoda.

## Materials and methods

### Dataset preparation

In this study, we retrieved genes from NCBI datasets (Sasyers et al. 2020) based on a list of human protein-coding genes obtained from BioMart of Ensembl release 105 (Howe et al. 2021). Using a customized script, orthologous genes were selected based on their similar length amino acid sequences in each clade. We grouped these genes into 13 clades, encompassing 308 species, as presented in **Supplementary Table 1**. The species are selected to represent a wide range of life forms within the vertebrate subphylum. Additionally, these species are categorized into various taxonomic ranks, including class (Aves and Amphibia), order (Primates, Rodentia, Lagomorpha, Chiroptera, Artiodactyla, Perissodactyla, Carnivora, Testudines, and Squamata), infraclass (Marsupialia), and superorder (Afrotheria), based on their shared morphology, common ancestry, and similar environmental conditions.

**Supplementary Table 2** contains a comprehensive list of protein-coding genes used in this study, including each species’ gene and amino acid accession number. We excluded LINCs, mitochondrial, and read-through genes from the list of 18,407 protein-coding genes and identified low-complexity regions in 17,940 genes (**Supplementary Table 3**). To infer the phylogenetic relationships of non-avian clades, we utilized the species tree from TimeTree (Kumar et al. 2017), while the tree for birds was obtained from the BirdTree website (Jetz et al. 2012).

### Protein LCR identification

We used fLPS 2.0 (Harrison 2021) to identify stretches of amino-acid low-complexity regions. We selected LCRs based on the following criteria: (a) the region should be longer than three amino acids, (b) the stretch should not contain X, and (c) the stretch should not contain more than five unique amino acids in the composition. We investigated the distribution and abundance patterns of LCRs across a purity gradient of composition. However, we restricted our analyses of co-occurrence or overlap of LCRs and positively selected sites to those LCRs that exhibited more than 70% purity. The combinations of multi-amino acid LCRs are clubbed to only one combination (e.g., ILA, LIA, AIL, IAL, LAI, and ALI are clubbed to AIL) to facilitate the comparison of orthologous LCRs across species within each clade. The above criteria enhance the identification of orthologous LCRs and minimize artifacts. We provided a list of LCR-containing genes (5768) with more than 70% purity in **Supplementary Table 3**.

### Comparison of orthologous sequences

We removed the coding sequences without complete ORF (Open Reading Frame), i.e., if they have premature stop codon, absence of start codon, presence of a non-nucleotide character (any character other than A, T, G, or C), or the sequence length is not a multiple of three.

Subsequently, we removed the gene sequences with less than three species in a clade. We use the GUIDANCE (Sela et al. 2015) program with MUSCLE (Edgar 2004) aligner with 100 iterations for multiple sequence alignment of the orthologous gene sequences. The coordinates of repeats identified by fLPS 2.0 are mapped to their respective aligned locations to identify the orthologous repeats across species in a clade. The positional overlap of repeats of different species in the alignment is considered an indicator of orthologous repeats.

### Analyses and visualization

We used the site model (M7 and M8) of PAML (Yang 2007) to identify positively selected sites. A likelihood ratio test is conducted on the compared models to identify signatures of site-specific positive selection. The Bayes empirical Bayes (BEB) method is used to calculate the posterior probabilities of each detected codon under positive selection. Sites with posterior probability > 0.95 are considered under significant positive Selection (Yang 2007). We calculated lineage-specific ω (dN/dS: non-synonymous substitution rate/synonymous substitution rate) by implementing the “free ratio” model (Model=1) in the PAML package.

### Positional preference of LCRs in a gene

To analyze the occurrence frequency of homopolymeric low-complexity regions (LCRs) consisting of a single type of amino acid, we calculated their abundance per unit length of the gene sequence. Specifically, we extracted all LCRs composed of a specific amino acid from all clades and determined their mid-point position to the normalized gene length. We divided the normalized LCR positions into twenty equal-sized bins. We then calculated the proportion of LCRs present in each bin and expected a proportion of approximately 5% in each bin under a completely random amino acid distribution. To identify the potential over-abundance or under-abundance of LCRs in each bin, we set thresholds of 8% and 2%, respectively. Moreover, we analyzed the distribution of each amino acid LCR across the purity gradient to gain insight into the changes in selection pressure.

### Co-occurrence of PSS and LCR

Using Fisher’s test, we investigated the co-occurrence and overlap patterns between LCRs and PSSs at three different scales. Specifically, we examined the overlap between LCRs and PSSs at a gene-wise and clade-wise level and evaluated their co-occurrence at the clade level. To perform these analyses, we used the fisher function in R to assess co-occurrence and the bedtools fisher tool to examine overlap, taking into account the coordinates of the LCRs and PSSs. We performed all the necessary analyses and visualizations using R (R Core Team 2021). The **Supplementary text** describes the methods used in producing the results in detail.

## Results

### Protein low-complexity regions show variable lengths across clades

Protein repeats can have varying length distribution according to clades and purity. In order to examine the possible divergence in length distribution of low complexity regions (LCRs) among diverse clades and levels of purity, we visualized the length distribution for each clade, utilizing a purity gradient of 10%. The resulting figures, presented in both **Fig. 1** and **Figures S1-10**, enabled us to carefully scrutinize the possible variations in the skewness of the length distribution for each clade and purity level to gain further insight into the potential clade and purity-specific impacts on the distribution of LCR lengths. Our study demonstrates that low-purity LCRs (0-60% purity) exhibit larger length variation across different clades. However, the distribution of length becomes more constrained at higher purity levels. Specifically, when the purity level is at or above 70%, the majority of LCRs in all clades are found in the lower range of the length distribution, and only a few regions are exceptionally long (greater than 2 Kb). The exceptionally large-length LCRs can be an artifact as their orthologs are scarcely present. The mean length of LCRs at >=70% purity across clades varies between 63-72 bp.

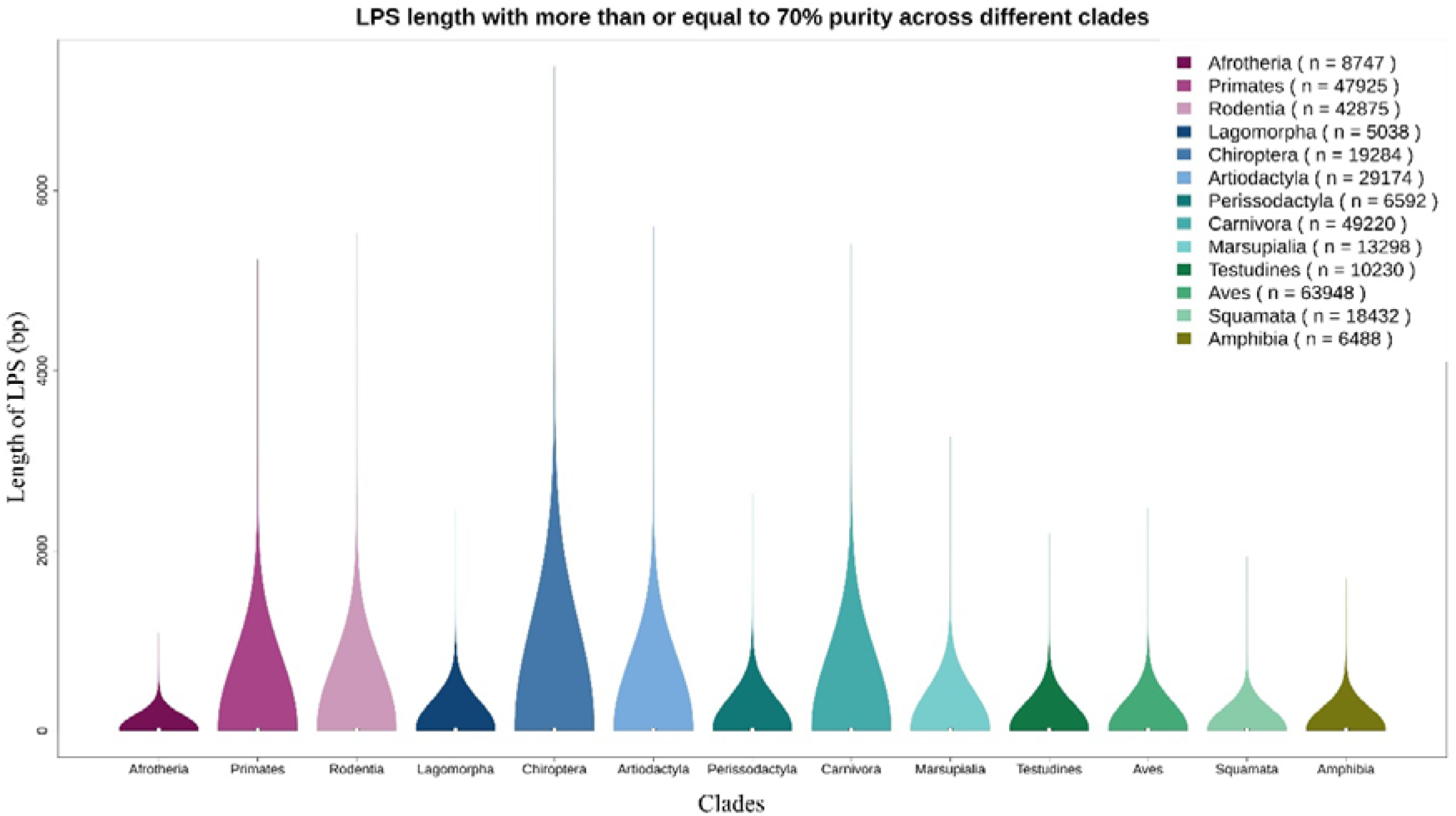
Length distribution of protein LCRs across clades. The violin plot shows the amino acid LCR length distribution in each clade, and the plot width represents the density of LCR length in that class interval. Most LCRs have a length in the lower range, while some are exceptionally large.

### LCRs show a heterogeneous distribution with purity

We detected ∼85% of protein-coding genes to have at least one LCR in all the clades (**Figure S11**). The percentage of LCR-containing genes shows a gradual decline with increasing purity of the LCR composition, with ∼9-15% at 70% purity and only ∼1-3% at 100% purity. For our comparative phylogenetic analysis of LCRs and positively selected sites (PSSs), we selectively analyzed LCRs with a purity greater than 70%. This was done to include degenerated LCRs across various species while reducing the likelihood of false positives arising from an overabundance and overlap of LCRs and PSSs.

### LCRs participate in a variety of biological processes

Studies highlighting the major pathways and processes of protein LCRs are scarce and are limited to specific genes (Andrade et al. 2001; Deryusheva et al. 2021; Teekas et al. 2022). To evaluate the major pathways and biological processes of protein LCRs, we used gene enrichment analyses using ShinyGO 0.76 server (Ge et al. 2020) (**Supplementary Table 4**). The analysis identified enrichment in biological processes related to morphogenesis, regulation of transcription by RNA, neurogenesis, and embryo development (**Figure S12**). The cellular component analysis highlights enrichment for RNA polymerase transcription regulator complex, basement membrane, ER lumen, chromatin, and components related to neurons (**Figure S13**). The molecular function is enriched mainly for molecular binding activities (histone, DNA, RNA, chromatin, protein kinase binding) (**Figure S14**). Similarly, the enrichment analyses of PSS-containing genes showed enrichment for biological processes like ion transport and neuron development and differentiation, cellular components like basement membrane and receptor complex, and molecular functions like protein tyrosine kinase and transferase activities (**Figure S15-17** and **Supplementary Table 5**).

We extended our study to examine position-specific enrichment of pathways for PSS and LCR-containing genes (**Figure S18-25** and **Supplementary Table 6**). We performed the analyses by classifying the PSS and LCR genes into two classes: terminal and central. Central PSS genes are enriched only for glycosaminoglycan binding (**Figure S24**), while terminal PSS genes are mainly for ATP and nucleotide binding (**Figure S25**). The biological processes of central-PSS genes are primarily associated with defense responses (**Figure S22**). Additionally, central PSS genes exhibit enrichment in components such as collagen-containing extracellular matrix and secretory granules (**Figure S23**). Interestingly, GO enrichment analyses for terminal PSS genes showed non-significant results for biological processes and cellular components despite having a large abundance of genes. We can attribute the non-significance of the results to the fact that when the background and foreground gene sets become very similar for GO enrichment, the results become non-significant.

### Repeat diversity varies across clades

To investigate the rate of detection of unique LCRs relative to the total number of LCRs in each clade, we employed the species-area relationship curve as a conceptual framework. The slope of the curve represents the rate of change in LCRs richness per unit increase in the number of LCRs. A steeper slope indicates a faster increase in the number of unique LCRs with the increasing number of LCRs, while a flatter slope indicates a slower increase in LCR richness. We calculated the slope of richness for each clade (**Figure S26-40**). Furthermore, we also calculated the richness and slope of richness species-wise for each clade and compared the distribution (**Figure S41-46**). Aves show a high variation in the richness of LCRs and the slope of the curve between species. Moreover, Aves have the highest LCR (>70% purity) richness compared to other clades, while Lagomorpha and Perissodactyla exhibit a low richness (**Figure S45**). After normalizing the LCR richness to the number of initial sequences, the richness across all the clades is comparable (**Figure S46**).

To overcome the effect of an unequal number of species and the number of complete open reading frames (ORFs) in clades, we use the Shannon diversity index (H) and Simpson’s diversity index (D). Simpson’s diversity index considers richness (the number of unique repeat types defined by the set of amino acid residues present) and abundance (the proportion of each repeat type). If all repeat types have the same abundance, the D will be 1, while the over-abundance of limited repeat types will lead to a lower D value (close to 0). We compare the distribution of species-specific D across all the clades (**Fig. 2**). Our result depicts that the diversity index (D) remains very high across all the clades (> 0.94). The overall diversity is highest in Amphibia while lowest in the Squamata clade. The Shannon-Wiener diversity index also shows a similar pattern of repeat diversity distribution (**Figure S47-51**). Similarly, we calculated richness, the slope of the curve, and diversity indices for LCRs of 100% purity and found a similar pattern (**Figure S52-56**).

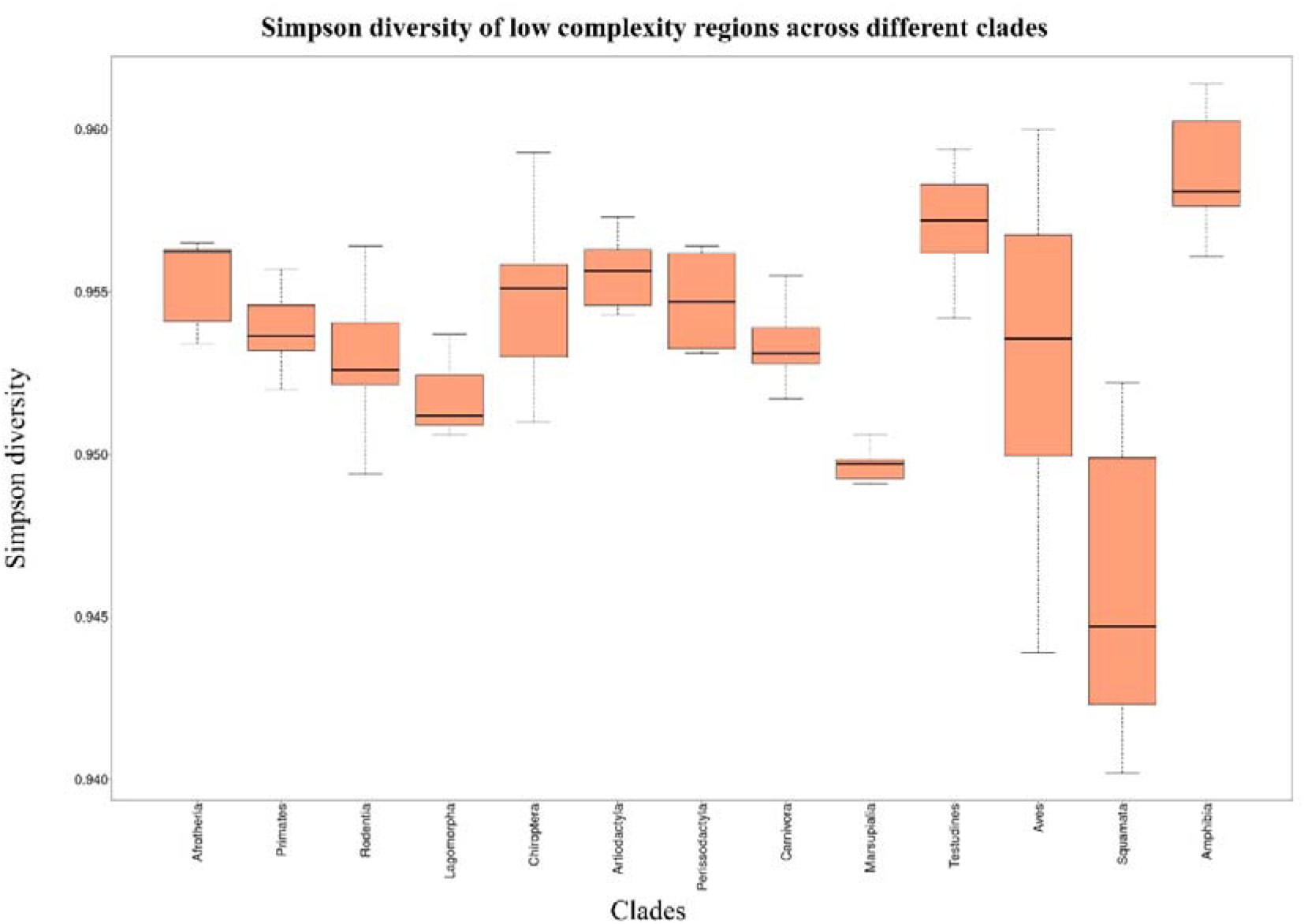
Diversity of LCRs across clades. The boxplot shows the amino acid LCRs diversity of each species in each clade. We calculate the diversity for each species using Simpson’s index of diversity. Simpson’s diversity index takes into account the type and abundance of LCRs. The homogeneous abundance of LCRs leads to a high diversity index, while an unequal abundance of different LCRs leads to a lower diversity index. The Amphibia clade shows an overall higher diversity, while the Squamata clade shows lower diversity of amino acid LCRs.

### Clades show variable repeat abundance

Factors like clade-specific codon usage, recombination slippage, or selection pressure can lead to a varying abundance of repeat types. A high proportion of a particular amino-acid repeat in a clade will imply bias towards that amino acid in mechanisms related to the origin or maintenance of repeats in that clade. We selected each clade’s twenty most abundant repeats and compared their proportion (**Fig. 3**). We observed that clades’ most abundant repeat types vary slightly (the top twenty repeats are not the same across all the clades). For instance, the proportion of the AG repeat is very low in the Amphibia clade but has a high proportion in other clades. However, the proportion for most repeats remains the same.

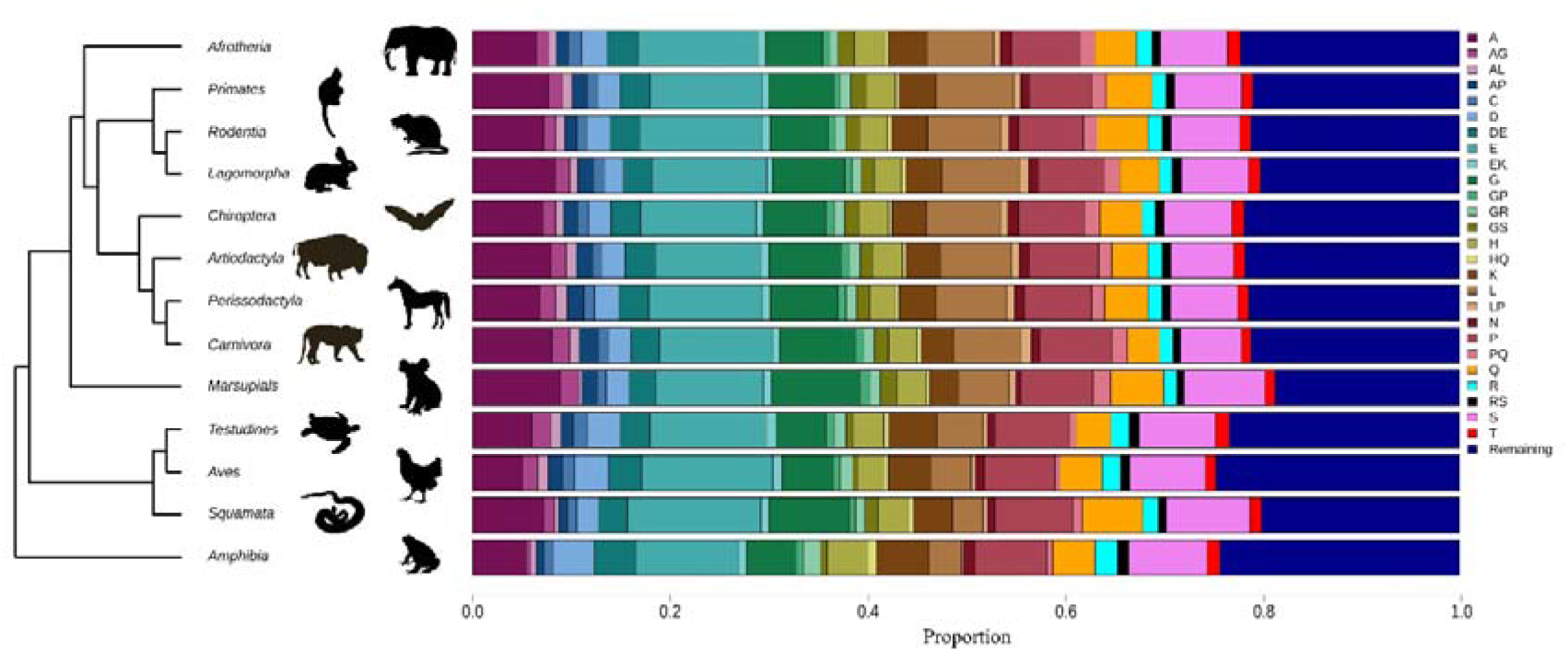
The proportion of the top twenty amino acids LCRs in each clade. The stacked barplot shows the proportion of the top twenty amino acid LCRs across all the clades along the phylogeny. Each color in the figure represents a different amino acid LCR. The deep blue color shows the summation of an abundance of all the remaining LCRs in the respective clade.

### LCRs exhibit amino acid-specific positional preference in a gene

Previous studies have emphasized a positional bias for amino acid repeats, where those containing poly-L, V, Q, H, N, C, A, and G prefer the N-terminal, while those with poly-S, K, I, and F favor the C-terminal (Albà and Guigó 2004; Siwach et al. 2006; Huntley and Clark 2007; Mier et al. 2017). These studies highlight an association between the type of amino acid and biological function but primarily focus on single amino acid repeats. To gain a more comprehensive understanding of the selection constraints acting on low-complexity regions within a biological context, we examined their positional preferences across a purity gradient ranging from 0% to 100%, with 10% intervals, as illustrated in **Figure S57-276**. Moreover, to investigate taxon-specific preferences, we extended our study in a clade-specific manner along the same purity gradient (**Figure S277-419**).

Interestingly, LCRs of amino acids A, G, L, and R exhibit a pronounced preference for the beginning of the gene, even for compositions of 0% and higher purity. Additionally, nine amino acid LCRs (A, V, R, Q, P, L, G, F, and C) exhibit an over-abundance at the N-terminus of genes, while two residues (D and E) show over-abundance at the C-terminus. Furthermore, three residues (Y, N, and M) show over-abundance at both termini at 70% purity. The distribution of leucine (L) amino acid shows an overabundance at the terminals of the gene sequence while avoiding the middle region (**Figure S163**). It is worth noting that none of the clades exhibit an overall preference for either terminal of the gene up to an LCR purity of 50%. However, for LCR compositions of 50% or higher, all clades prefer the N-terminal of the gene (**Figure S277-419**).

### Gene with LCRs face stronger purifying selection

Previous studies report the GC composition of genes containing repeats to be significantly higher than those without repeats (Albà and Guigó 2004; Albà et al. 2007; Teekas et al. 2022). These studies were either limited to specific organisms, specific gene sets, or amino acid tandem repeats. We compared the %GC of all the available protein-coding genes with and without LCRs (>70% purity) in the Tetrapoda clade. We find that the %GC of LCR-containing genes is significantly higher than those without LCRs (Wilcoxon test, p < 0.05 in all the comparisons; **Figure S420**).

Furthermore, we compared the distribution of ω (dN/dS: non-synonymous substitution rate/synonymous substitution rate) between genes with and without LCRs (>70% purity) after filtering out all the ω values greater than 2. Filtering out the values greater than two is necessary as sometimes PAML reports ω values as high as 999, potentially biasing the results. Our finding reveals that LCR-containing genes have significantly lower ω distribution than genes without LCRs across all the clades (Wilcoxon test, p < 0.05; **Fig. 5**). Most of the ω values are smaller than 1, implying purifying selection on the majority of the genes. The median and mean of ω in genes with LCRs are lower than those without LCRs. The result suggests that genes with LCRs face more substantial purifying selection than others. To rule out the possibility of sampling bias, we compared the distribution with all the ω values at different purity cutoffs of LCRs. The inference remains the same (**Figure S421-439**).

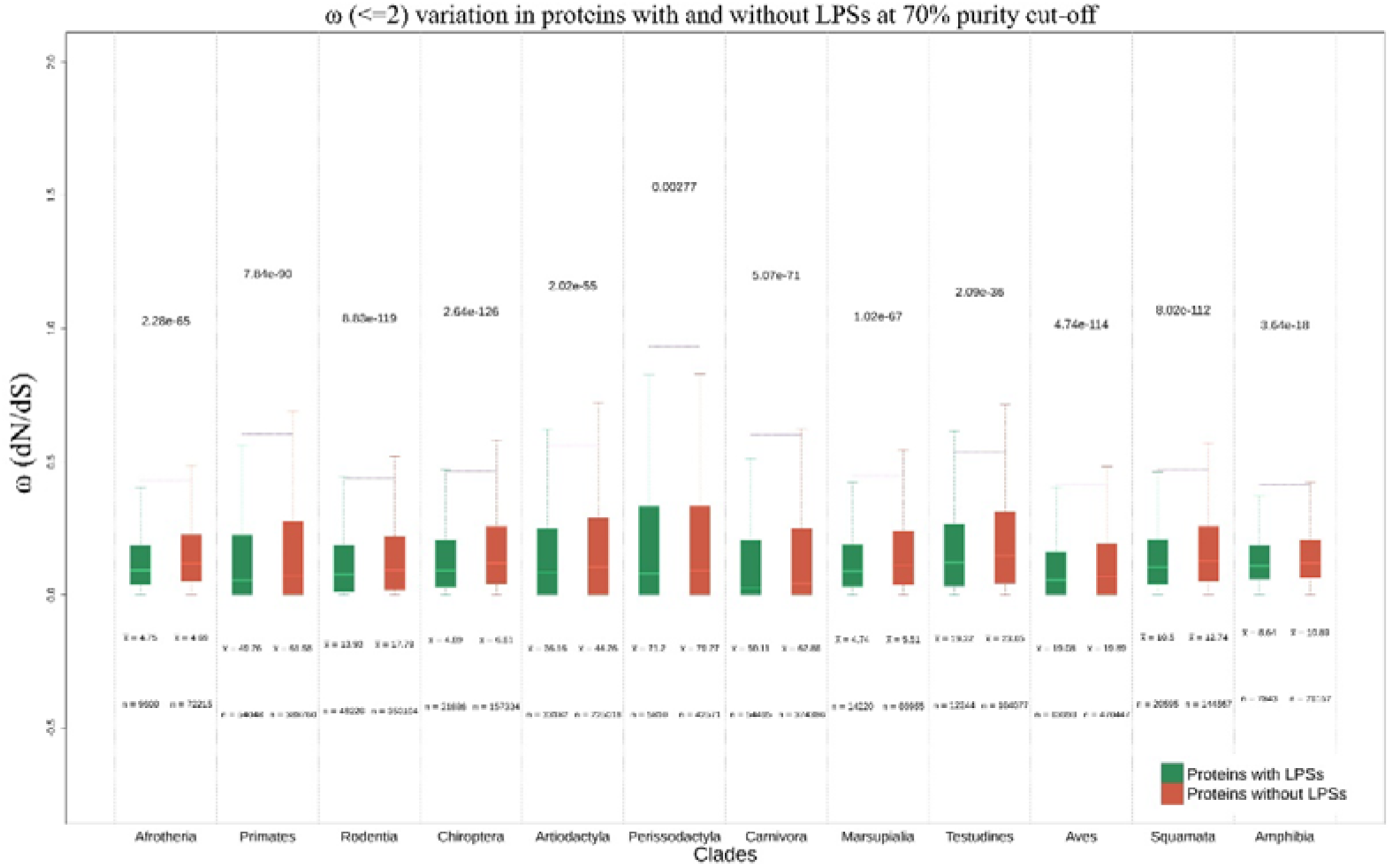
Distribution of ω (dN/dS) in proteins with LCRs and without LCRs. We calculated each lineage’s ω (dN/dS) using the free-ratio model of PAML and subsetted it according to the presence of LCR or absence. The boxplot in green and red color shows the ω distribution of protein sequences with LCRs and protein sequences without LCRs, respectively. The boxplots are visualized with ω <= 2 and without showing the outliers. The distributions of ω are compared using the Wilcoxon test in R in a clade-wise manner. The values above the boxplots represent the p-value of the distributions compared. The “x-bar” below the boxplot represents the mean ω for that distribution, and “n” represents the number of lineages used to plot the distribution.

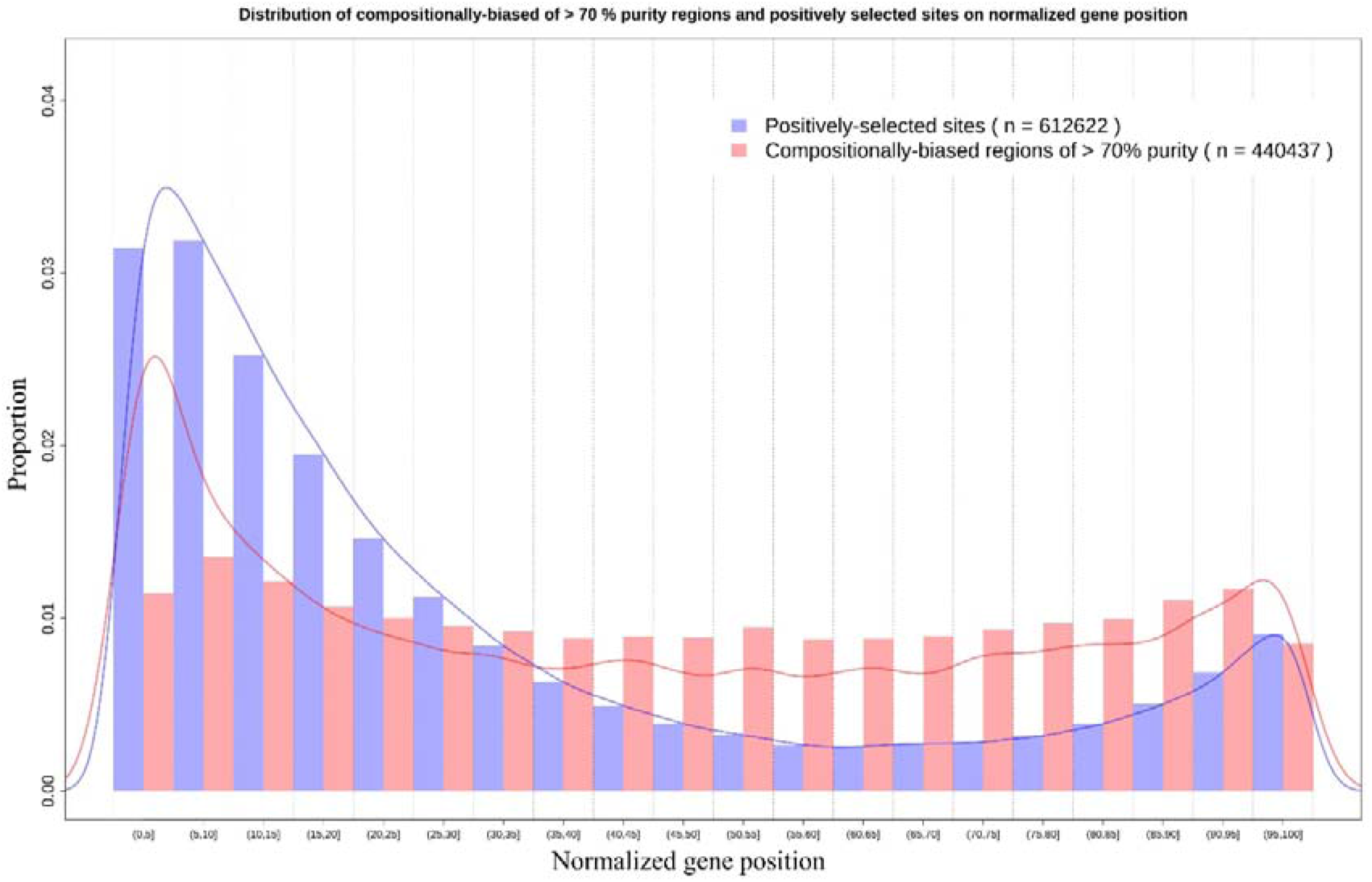
Positive selection and LCRs prefer similar regions in a gene. The overlapping histogram shows the abundance of positively selected sites, and LCRs mid-position along the normalized gene length. Selection and LCR show a preference towards the terminal regions of a gene.

### LCRs and selection prefer similar positions in a gene

We calculated the proportion of LCRs and positively selected sites in ten bins in a clade-wise manner in three purity classes of LCR composition, all LCRs, LCRS of > 70% purity, and LCRs of 100% purity, on a normalized gene length (**Fig. 5 and Figure S440-477**). Both LCRs and positively selected sites show a higher abundance at the terminals of the gene while keeping a low abundance in the remaining bins. Furthermore, we examined the distribution patterns of the three aforementioned LCR purity classes in each clade (**Figure S478-491**) to gain insight into changes in LCR distribution within genes. Our analysis indicates that while all LCRs show a relatively uniform distribution, those with >70% and 100% purity exhibit a preference for the terminal regions in all clades. To rule out the possibility of alignment coverage bias in the inference of results, we compared the distribution with alignment coverage (**Figure S492-517**). Alignment coverage shows a negative trend and correlation compared to positive sites and LCRs abundance distribution.

### Genes exhibit co-occurrence of LCRs and PSSs

Given the similarity in distribution patterns of LCRs (>70% purity) and PSSs within protein-coding genes, we explored the potential co-occurrence and overlap between the two. We analyzed the distribution and patterns of LCRs and PSSs within genes across three levels: gene-wise overlap, clade-wise overlap, and clade-wise co-occurrence. This comprehensive approach allowed us to understand better how these two features interact within genes. A total of 501 unique genes show a significant overlap (p < 0.05; Fisher’s exact test using bedtools fisher) of PSSs and LCRs in the Tetrapoda clade, while 363 unique genes exhibit an avoidance of overlap (**Supplementary Table 7**). Interestingly, eight of the twelve clades studied significantly favor PSSs in LCRs (p < 0.05; Fisher’s exact test using bedtools fisher; **Supplementary Table 8**). Three clades show non-significant results, while Afrotheria shows significant avoidance of PSSs in LCRs. Furthermore, our analysis of the co-occurrence of PSSs and LCRs showed significant (p < 0.05; Fisher’s exact test) co-occurrence in seven clades, except for Perissodactyla, where we observed mutual exclusion (**Figure S518-529** and **Supplementary Table 9**).

### PolyQ of MAML2 favors PSSs

We selected the MAML2 gene from the list of genes that significantly favor PSSs in LCRs, as it demonstrated a preference for PSSs in the polyQ LCR across five different clades (**Fig. 6** and **Figure S530-533**). MAML2 is a member of the mastermind-like protein family and functions as a coactivator of the canonical NOTCH signaling pathway, which is known to promote oncogenesis (Lin et al. 2002; Zhang et al. 2019). Members of the mastermind-like protein family are characterized by a high abundance of proline and glutamine residues, as well as a conserved basic domain located in their N-terminus that binds to the ankyrin repeat domain of the intracellular domain of Notch receptors (ICN1-4). In addition, these proteins possess a transcriptional activation domain in their C-terminus (Lin et al. 2002; Wu et al. 2002; Kovall and Hendrickson 2004; Zema et al. 2020). The polyglutamine region of MAML2 shows length polymorphism (Rozanska et al. 2007), which also contains positively selected sites.

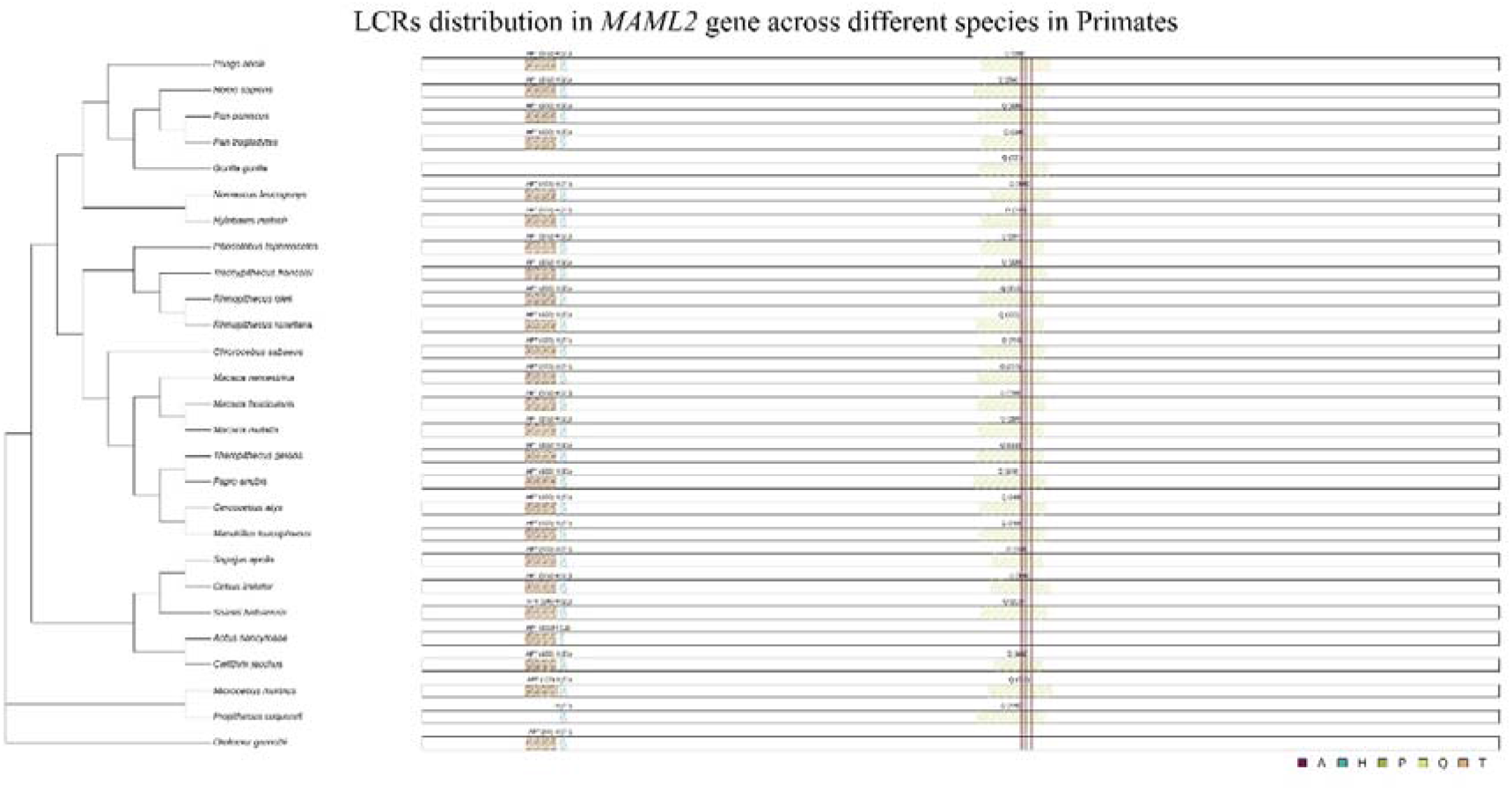
Evolutionary analysis of MAML2 in Primates reveals an overlap of positively selected sites with low complexity regions (LCRs) enriched in polyglutamine (polyQ) residues. The *MAML2* gene, which encodes a transcriptional coactivator, contains conserved LCRs with high purity (>70%) in Primates. The nucleotide lengths and base composition of the detected LCRs are represented above the gene. Additionally, positively selected sites (PSSs) detected using the site model of PAML are indicated by dull red vertical lines. This figure aims to illustrate the overlap of PSSs with the polyglutamine (polyQ) LCR in the *MAML2* gene along the Primates phylogeny.

## Discussion

Protein LCRs are unstable and introduce rapid, adaptive variations in a short evolutionary time, as opposed to longer-term evolutionary epistasis (Radó-Trilla and Albà 2012; Newton and Pask 2020). Moreover, LCRs also contribute to providing site-specific, frequent, and reversible mutations. These specific mutations works as a tuning knob for genes and genomes for efficient adaptation (King et al. 1997). Studies have proposed the Goldilocks length range within which the length variation has non-deleterious functional consequences. When the LCRs are very short, they have a negligible functional impact, while very long regions make protein aggregates and, thus, hampers function. The LCRs within this goldilocks range can provide functional diversity without being detrimental (Newton and Pask 2020). Examples like the CAG repeat of the *HTT* gene and the Poly-QA repeat in the *RUNX2* gene also support the detrimental effect of repeat above a critical length (Reiner et al. 2011; Newton and Pask 2020). The overwhelming evidence about the functional consequences of length variation suggests LCRs face stronger purifying selection above a critical length; an expansion may favor smaller LCRs, and the variation in length is nearly neutral within the critical limit (Hancock and Simon 2005). The LCR length instability provides a platform for selection to work. Since selection and LCR polymorphism work in tightly entangled manner, it is essential to understand their signatures across clades.

LCR-containing genes participate in pathways related to morphogenesis, neurogenesis, transcription regulation, and biosynthetic processes. Moreover, LCR-containing genes participate in a variety of binding-related processes like histone binding, chromatin binding, and DNA and RNA binding. LCRs are abundant in these specific processes and directly participate in many of them. For example, alanine repeats are known to have a role in mediating transcription repression (Janody et al. 2001; Maurer et al. 2003). The polyalanine domain of transcription factor HoxA11 works as a SIN3A/HDAC1Ldependent repressor domain by physically interacting with the PAH domain of the SIN3 protein (Lynch and Wagner 2021). Studies have highlighted that the expansion of polyalanine tracts may contribute to functional diversification in transcription factors (Lynch and Wagner 2021). Specific pathways prefer certain types of LCRs and amino acids. Transcription factors prefer regions of glutamine, aspartic acid, and asparagine (Hancock and Simon 2005). Similarly, leucine repeats are exploited both by hosts and pathogens in a host-pathogen interactive evolutionary arms race. The pathogens use leucine repeats for host cell attachment and entry, while host immunity utilizes leucine repeats to detect pathogen-specific molecules (Cohn 2002; KLdzierski et al. 2004). Enrichment of LCR-containing genes and direct participation of LCRs in various functions enhances the role of LCRs in rapid functional diversification.

Low-complexity regions contribute to rapid evolutionary novelty with length polymorphism. LCRs work in a length range for functional diversification (Newton and Pask 2020). Very distinct length distribution of orthologous LCRs across clades could imply functional diversification in that clade. We noticed the length distribution of LCRs across all the Tetrapoda clades is similar. Selective constraints regulate the length of most LCRs, with smaller regions being more common. Abnormal expansion of LCR length beyond a certain limit can potentially be detrimental. In a few exceptional cases of Primates, the LCR length is very large. We do not find a similar LCR length in other clade/species. We cannot rule out the possibility of annotation artifacts for such LCR-containing genes.

A varying number of species across clades can affect the unique LCRs detected from each clade. Moreover, poor genome assembly and annotation will lead to fewer detected genes overall. We use the concept of the species-area relationship curve to predict the slope of the log(number of unique repeats) to the log(number of sequences). The species-area relationship curve describes the number of species detected in a given area (Preston 1962). Here, we consider the number of gene sequences equivalent to the area and the number of LCRs detected equivalent to the number of species detected. This method compares the detection intensity of LCRs across different clades. In most clades, the distribution is between 5.4 to 5.6, implying that in all the clades, we can detect an almost equal number of unique LCRS if the number of gene sequences detected is the same. This result also establishes the essentiality of LCRs across different tetrapod clades. Since clades have varying levels of annotations and number of species (which can lead to a variable number of gene sequences detected), we calculated Simpson’s and Shannon-Weiner’s diversity index to compare LCR community structure across the clades (Simpson 1949; Spellerberg and Fedor 2003). Both diversity indices take richness (unique LCRs) and evenness (uniform proportion of LCRs) into account. The Simpson’s diversity index (D) ranges from 0 to 1, where 0 represents no diversity and 1 represents maximum diversity. Similarly, Shannon’s diversity index (H) ranges from 0 to n (n is any positive number), where 0 is no diversity. A similar diversity index distribution of LCRs across all the clades will imply similar LCR composition irrespective of the number of genes considered for each species. We observe very similar indices (D: 0.94 - 0.96 and H: 3.6 - 3.8) for all the species across all the clades. This similar diversity index implies the LCRs have an equally important role across the diverse species of the tetrapod clade. An indispensable role of LCRs in diverse functions helps in the maintenance of repeat diversity across different clades.

We also compared the proportions of the twenty most abundant LCRs in each clade. Of all the amino acid LCR types, alanine, glycine, and proline are the most frequent across all the clades. Specific amino acid LCRS have functional implications in specific processes (Hancock and Simon 2005). The profound presence of particular types of LCRs in particular processes is proposed to contribute to the evolution of that specific pathway (Hancock and Simon 2005). In mammalian proteins, alanine repeat-containing genes are over-represented in DNA-and RNA-binding (Albà and Guigó 2004). Alanine repeats play a prominent role in neurogenesis, signaling, and development (Cocquet et al. 2003; Lavoie et al. 2003; Caburet et al. 2004; Lynch and Wagner 2021). Hence, it is unsurprising to observe a high abundance of alanine repeats across the Tetrapoda clades. Interestingly, alanine, proline, and glycine LCRs have more evolutionary effects than other repeats. PolyP, polyA, and polyG repeats are known to modulate protein-protein interactions and regulate transcription (Mitchell and Tjian 1989; Gerber et al. 1994; Emili et al. 1994; Wilkins and Lis 1999). Proline rich regions prohibit aggregate formation by the PolyQ region when the PolyP is on the C-terminal side of the glutamine repeat, but the effect is abolished if PolyP is moved to the N-terminal side of the PolyQ (Bhattacharyya et al. 2006). The essentiality of these specific LCRs and their functional impact in regulating other LCRs can explain their higher abundance in the Tetrapoda clade.

Apart from the amino acid LCR-specific functions, specific LCRs have a positional preference and position-dependent roles (Coletta et al. 2010). Leucine repeats prefer a gene’s beginning and end terminals while mainly avoiding the mid-portions. Most repeats prefer the ’gene’s beginning to any other portion, except for D, E, K, and Y repeats. Aspartate, glutamate, lysine, and tyrosine repeats prefer the end terminal more than any other position. Interestingly, serine repeats do not show any positional preference. Moreover, C, F, I, M, N, V, W, and Y repeats show very low abundance across all the clades, with W repeats showing the least abundance. One previous study about the abundance and distribution of amino acid repeats failed to detect any tryptophan (W) repeats (Mier et al. 2017); we noticed a very low abundance. Other amino acid repeats’ proportion and position preferences remain consistent with their study (Mier et al. 2017). Most leucine repeats at the beginning have signal peptide roles and are secreted (Mier et al. 2017). These findings indicate position-dependent functions of amino-acid LCRs.

In an interesting finding of overall LCR abundance across the genes, we noticed that the terminal regions are more preferred than the central region. The N-terminal region contains more LCRs than the C-terminal region, indicating that the ’5’ end of the gene is more dynamic in generating LCRs (Albà and Guigó 2004). Moreover, avoiding the middle region of a gene by the LCRs might be an evolutionary consequence of avoiding the globular regions and domain misfolding (Albà and Guigó 2004; Mier et al. 2017). These findings highlight that selective constraints are region-dependent on a gene, and some regions can tolerate more diversity than others. The profound occurrence of LCRs in a gene’s malleable regions might facilitate functional diversification. Interestingly, we observe a similar trend for positively selected sites along the genes across the Tetrapoda clade. LCRs and positively selected sites both prefer the initial region of a gene the most, with the least proportion in the middle region.

The PSS genes are enriched for neurogenesis, transcription regulation, and morphogenesis. PSS genes are also enriched in molecular functions like tyrosine kinase, transporter activity, and ATP binding. LCR-containing and PSS genes are enriched for similar biological processes, suggesting an equally important role of positive selection and LCRs in maintaining these processes and their diversity. Interestingly, mid-PSS-and terminal-PSS-containing genes are enriched for distinct molecular functions. Terminal PSS genes are mainly enriched for ATP and nucleotide binding, while central PSS genes show enrichment for glycosaminoglycan binding. Of the two categories of enriched functions, only seven genes are common to both, with the majority being unique to each function. An over-abundance of unique genes in both implies a position-dependent role of selection based on function.

A previous study observed a relationship between the distance from the LCR, the average substitution rate per site, and the proportion of genes with gaps in the alignment at each site. Additionally, the study analyzed the dN/dS ratios and found a negative relationship between the proportion of sites undergoing negative selection and the distance from the LCR (Lenz et al. 2014). Another interesting study found that sites with evidence of positive selection tend to be located near the edges of LCRs, with more sites clustered on the N-terminal side of the LCR boundary than on the C-terminal side. Specifically, half of these sites were found within the first 26 residues closest to the LCR boundary, while 95% were within 201 residues. The same trend was observed even when only focusing on the most strongly selected sites. The skewness might be related to the direction of transcription rather than the structure of the protein (Huntley and Clark 2007). We suspected the role of poor alignment coverage in the terminal regions might influence the detection of positively selected sites. The program that detects the PSS, PAML, treats poorly aligned regions as missing data (Huntley and Golding 2006). Regions with variable lengths can have poor alignment and may be excluded from the analysis. We detected a negative trend and correlation between alignment coverage with both PSS and LCR regions. The result shows that LCR regions have poor alignment due to variable length. However, low alignment coverage does not lead to the detection of a lower number of positively selected sites in the region.

The comparison of the GC content revealed that LCR-containing genes have a significantly higher %GC than genes without LCRs across the whole Tetrapoda clade (Wilcoxon test; p < 0.05). The result is consistent with previously published studies on specific gene sets (Albà and Guigó 2004; Teekas et al. 2022). A high %GC in LCR-containing genes suggests that GC-rich regions are prone to LCR formation. GC-rich codons primarily encode LCRs, increasing overall %GC (Albà and Guigó 2004). A high GC content increases mutation and recombination rates and affects length by DNA polymerase slippage (Kiktev et al. 2018, 2021). Replication slippage is one of the central mechanisms for the emergence and maintenance of protein LCRs (Mar Albà et al. 1999; Kashi and King 2006). The above statements suggest a high GC content promotes a predisposition of LCRs, increases mutation and recombination rates, and leads to higher DNA polymerase slippage. The emergence of LCRs, in turn, increases the overall GC content of the gene, as GC-rich codons mainly encode LCRs. So, protein LCRs and GC content work in positive feedback dynamics where the existence of one can promote the emergence and maintenance of the other if no other biological constraint exists.

Natural selection plays a crucial role in the emergence and maintenance of repeats (Mularoni et al. 2010; Persi et al. 2016). A newly emerged LCR faces relaxed and positive selection, which promotes rapid diversification. But, a functionally crucial fixed repeat is under purifying selection (Persi et al. 2016). Moreover, purifying selection keeps the repeat’s composition pure, while relaxed selection leads to compositional amalgamation (Fondon and Garner 2004). We observe that LCR-containing genes have significantly lower ω (dN/dS: non-synonymous substitution rate/synonymous substitution rate) than genes without LCRs across all the clades of Tetrapoda. This finding is consistent with a previous finding on a smaller dataset with the comparison of dN (non-synonymous substitution rate) only (Hancock and Simon 2005). They observed a lower median dN for proteins with orthologous LCRs versus proteins with non-orthologous LCRs. An overall ω distribution close to 0 implies that most genes are under purifying selection, with LCR-containing genes under more intense purifying selection. Protein LCRs, under intensified purifying selection but still a significant source of functional diversification and evolutionary novelty, imply that a tight interplay of selection and mutation governs protein LCR functionality. The mutation-selection dynamics act on length polymorphism, with selection being nearly neutral within the normal length range, and purifying above a critical length (Hancock and Simon 2005).

Of the 7590 genes containing both PSS and LCR in a clade-specific manner, 607 genes significantly favored PSSs within LCRs. We observed that the polyQ tract of MAML2 gene significantly favors PSSs in five clades. Mastermind-like transcriptional co-activator 2 (MAML2) is a member of the mastermind-like protein family and functions as a co-activator for the oncogenic NOTCH signaling pathway (Lin et al. 2002; Wu et al. 2002). The activation of NOTCH signaling has been implicated in carcinogenesis, where it plays a crucial role in cell transformation (Zhang et al. 2019). There are three orthologs of the Mastermind-like protein (MAML) found in vertebrates, all of which have an N-terminal domain capable of creating a complex with CSL transcription factors and intracellular Notch (ICN) ankyrin repeats to form a trimer (Kitagawa 2015). These homologs have similar protein architecture that involves two acidic domains and regions with polyglutamine (polyQ) that help Mam/MAMLs activate the transcription of genes targeted by Notch (McElhinny et al. 2008; Kitagawa 2015; Sinha et al. 2021). Interestingly, polyQ is known to play a role in protein-protein interaction and transcription activation (Gemayel et al. 2015; Newton and Pask 2020). The presence of PSSs and the direct functional implication of polyQ in MAML2 further establish the importance of LCRs in an evolutionary context.

## Conclusions

Protein LCRs are a source of rapid functional diversity and evolutionary novelty with composition, position, and length-dependent biological functions. A close interplay of mutation and selection governs the length and composition of LCRs. Furthermore, the origin and maintenance mechanisms of pure amino acid repeats and regions of compositional bias are different. Despite this, studies focusing on the positional preference of selection and LCRs with a change in composition are very scarce. Our findings indicate that site-specific positive selection and LCRs prefer the same positions in a gene and cooccur in most of the clades in Tetrapoda. Moreover, the polyQ of MAML2 gene favors PSSs in LCRs, further establishing the evolutionary role of low-complexity regions. LCRs are enriched in specific pathways and have similar diversity across different clades of Tetrapoda. LCRs occur in functionally important genes, are under intense purifying selection, and have a higher %GC than genes without LCRs. PSS and LCRs both prefer terminal regions of a gene over the central region. Moreover, central-and terminal-PSS genes and central-and terminal-LCR genes have distinct biological processes. Our study also highlights that the terminal regions of a gene are more flexible to accommodate variations and harbor volatile LCRs and positively selected sites. We believe our study contributes to understanding the nature and dynamics of selection and mutation for the emergence and maintenance of protein LCRs.

### Limitations and future directions

In this study, we included thirteen clades comprising more than 300 species with more than 18000 genes to rule out any sampling bias. But, this does not take care of all the confounding factors. Species have different genome assembly and annotation levels. Furthermore, the number of species between clades varied because of the unavailability of proper annotation in some clades. A clade comprising only a few species with good annotation will reflect the results of only those species and not the whole clade. Clade-specific results can potentially be affected by the number and combination of species selected for the study.

We used fLPS 2.0 for LCR sequence identification and filtered out sequences with less than 70% LCR composition, less than four amino acid lengths, and more than four amino acid combinations. The results can vary if the filtering criteria or program to identify LCRs are changed. Moreover, we used the MUSCLE aligner with 100 bootstraps for sequence alignment. Using a different aligner can potentially affect the alignment and, thus, the results associated with the aligned files. Most positively selected sites are in low-alignment coverage regions. This result needs further analysis to rule out the effect of alignment.

Despite the potential confounding factors, the results provided open a new avenue for further exploration of the evergrowing field of protein LCRs evolution. Our results explore only the Tetrapoda clade. Analyzing other clades can help us understand the mechanism and evolution of protein LCRs in the genomes of those clades. Furthermore, the orthologous LCRs vary in length between species of the same clade. The impact of LCR length variation on protein structure can allow us to understand their role in diseases in greater detail.

## Availability of data

All data associated with this study are available in the Supplementary Materials and GitHub link https://github.com/ceglablokdeep/LCRs_global_patterns

We have uploaded the entire dataset and the necessary scripts to Mendeley datasets, as some files exceed the 25MB limit for uploading on GitHub.

## Contributions

LT and SS wrote the manuscript with NV. LT and SS analyzed the data and generated the results. All authors reviewed the manuscript.

## Acknowledgment

We thank the Ministry of Human Resource Development for fellowship to LT and SS. Computational analyses were done on the Har Gobind Khorana Computational Biology cluster established and maintained by combining funds from IISER Bhopal under Grant # INST/BIO/2017/019, IYBA 2018 from the Department of Biotechnology (Grant no. BT/11/IYBA/2018/03), and ECRA from Science and Engineering Research Board (Grant no. ECR/2017/001430).

