## Supplementary Figures for "Terminal regions of a protein are a hotspot for low complexity regions (LCRs) and selection"

**Supplementary figure 1**

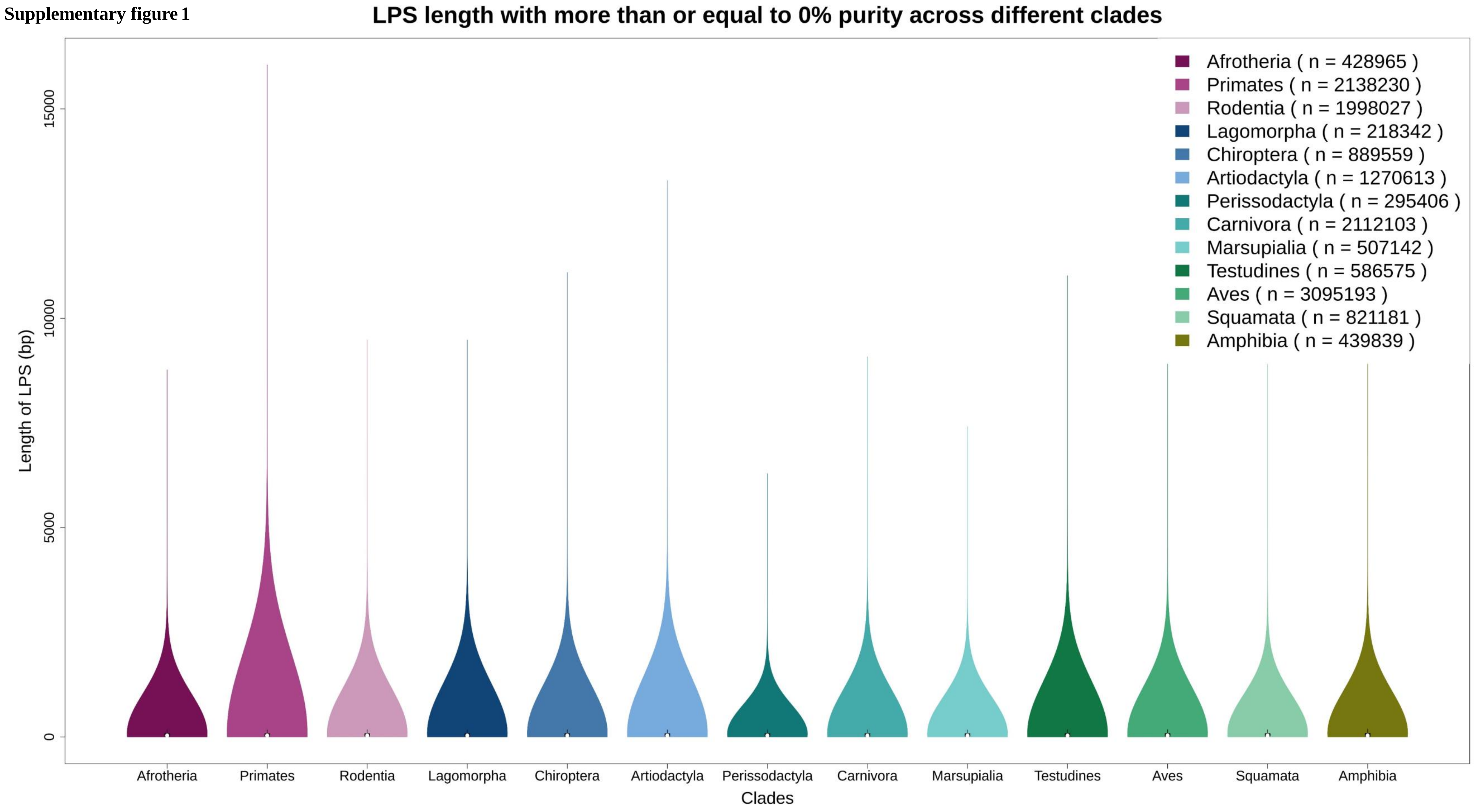

**Supplementary figure 2**

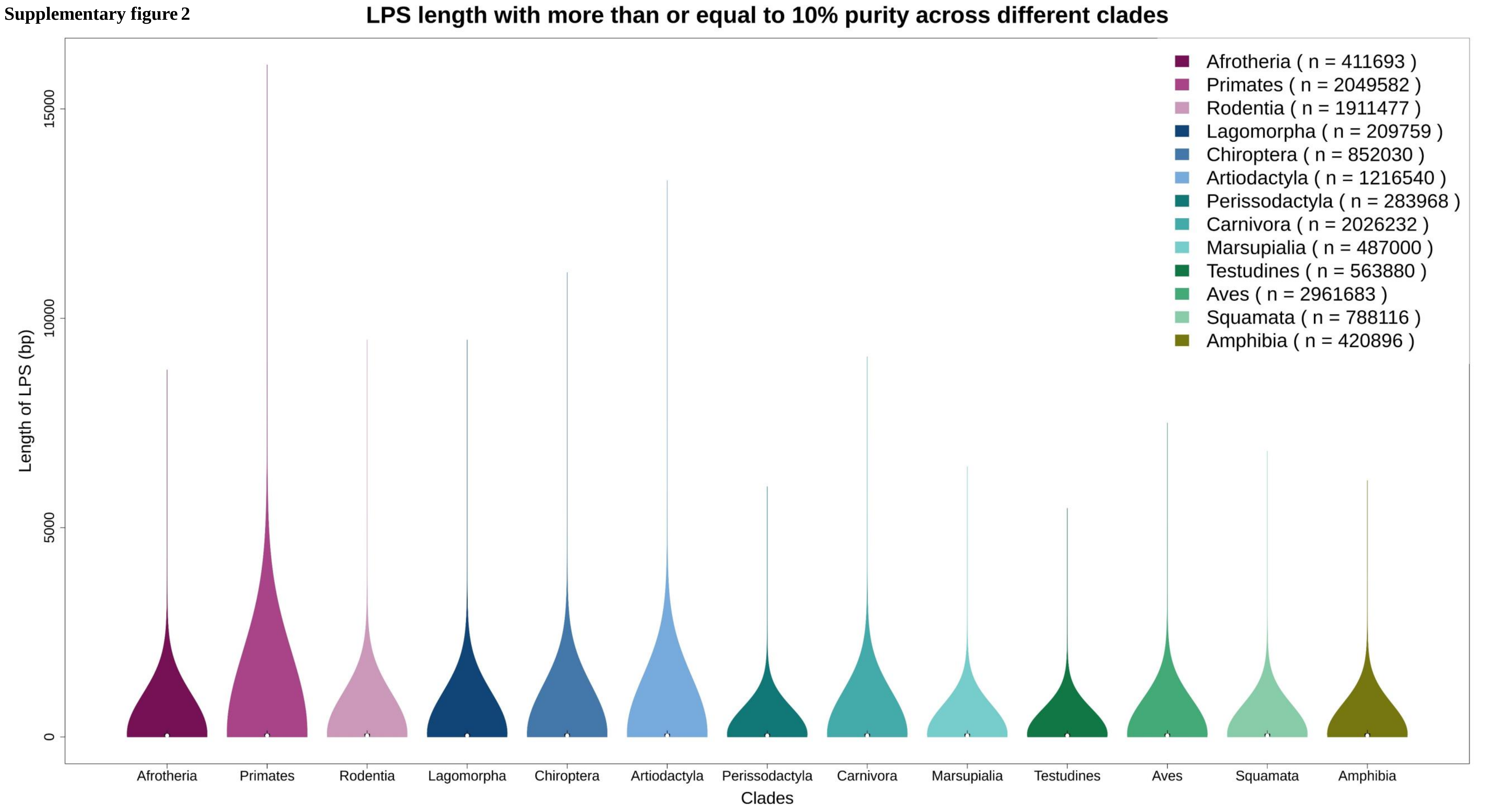

**Supplementary figure 3**

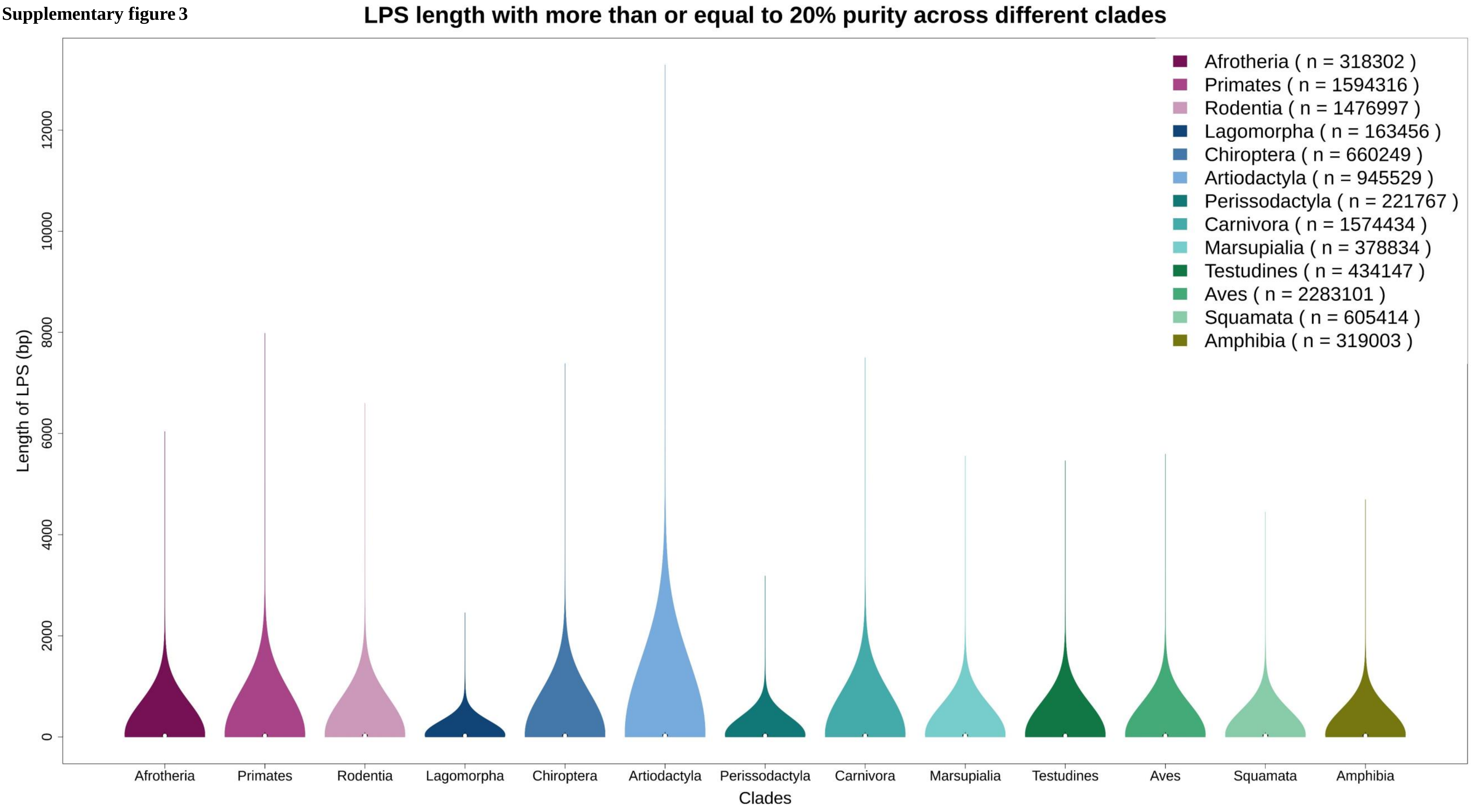

Supplementary figure 4

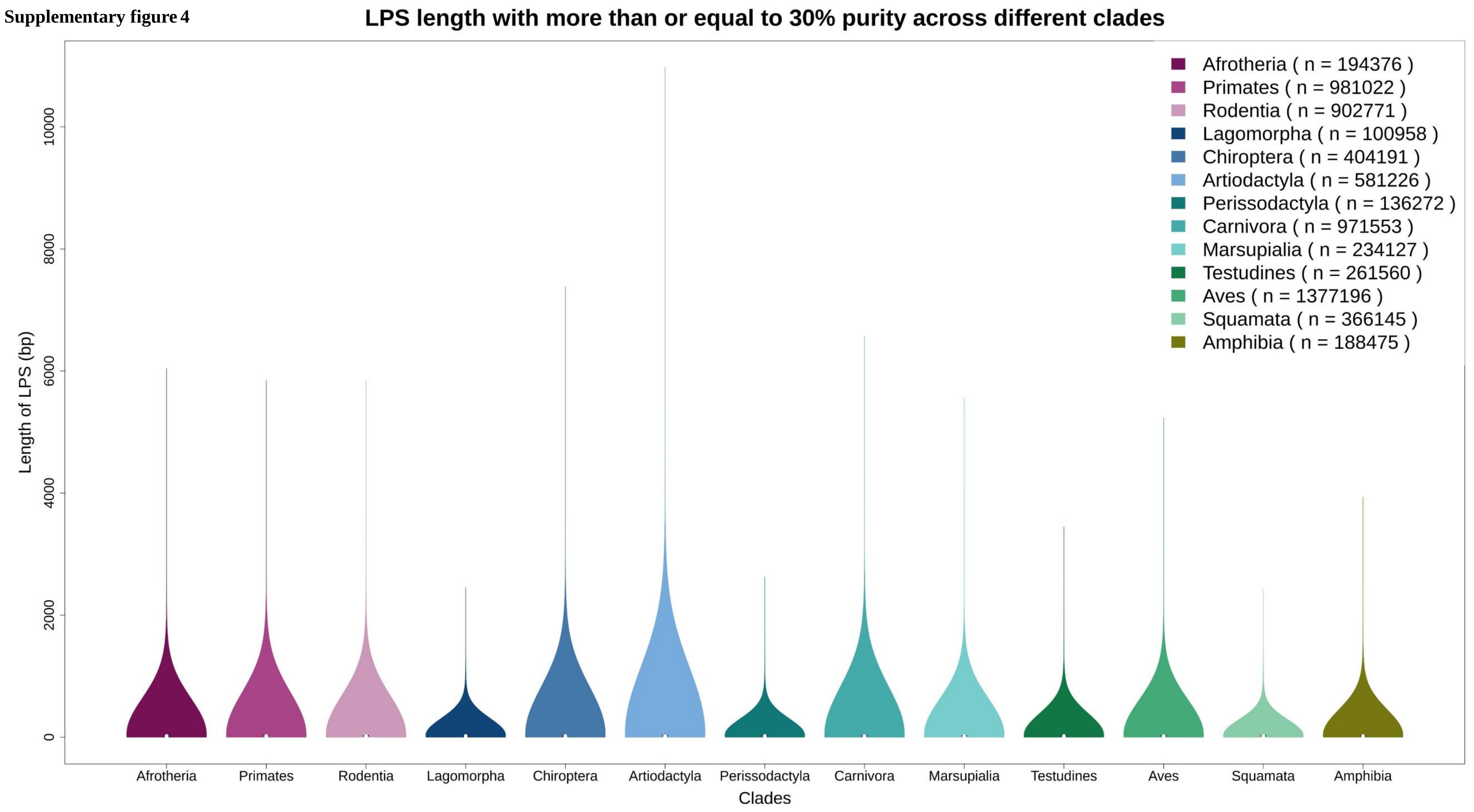

**Supplementary figure 5**

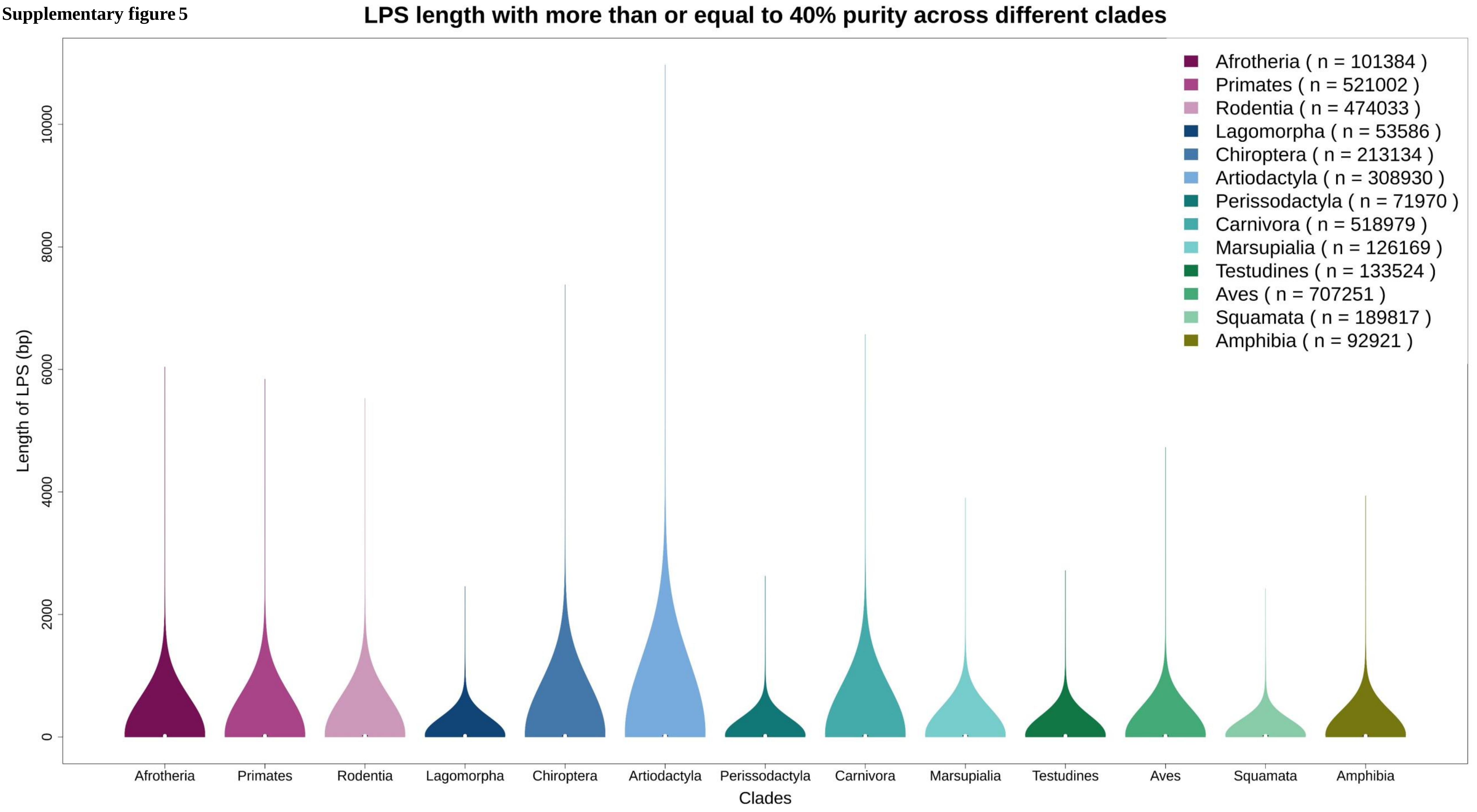

**Supplementary figure 6**

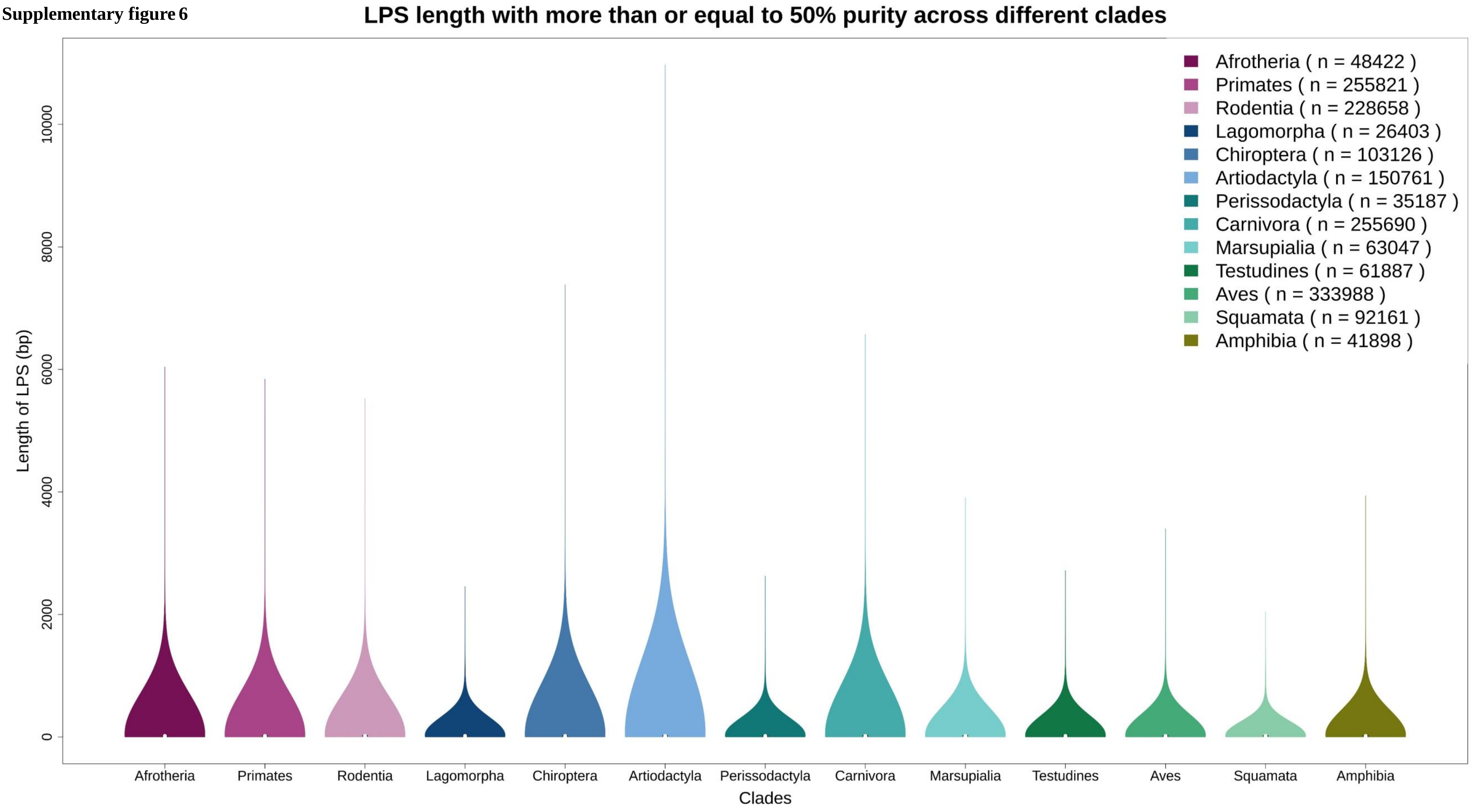

**Supplementary figure 7**

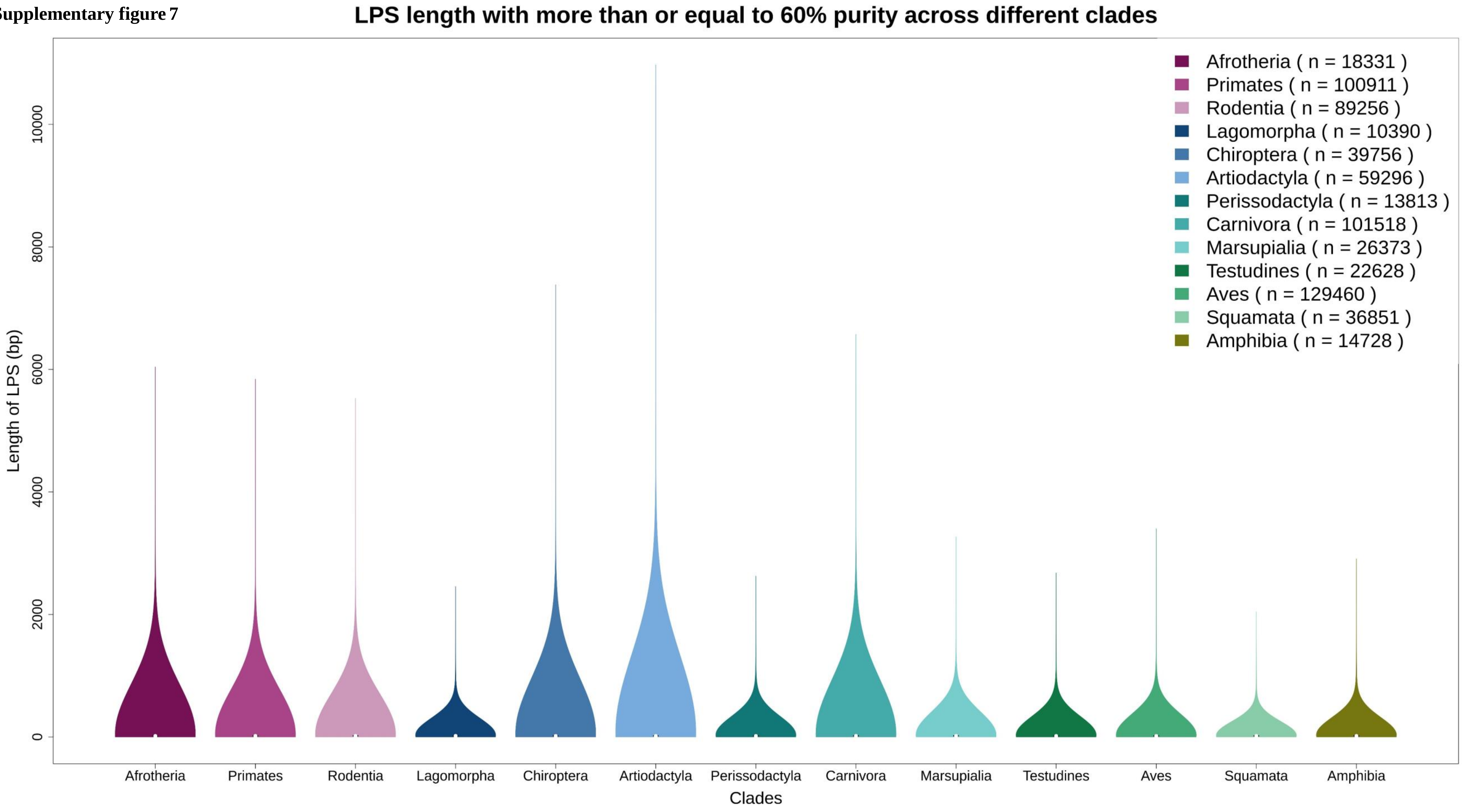

**Supplementary figure 8**

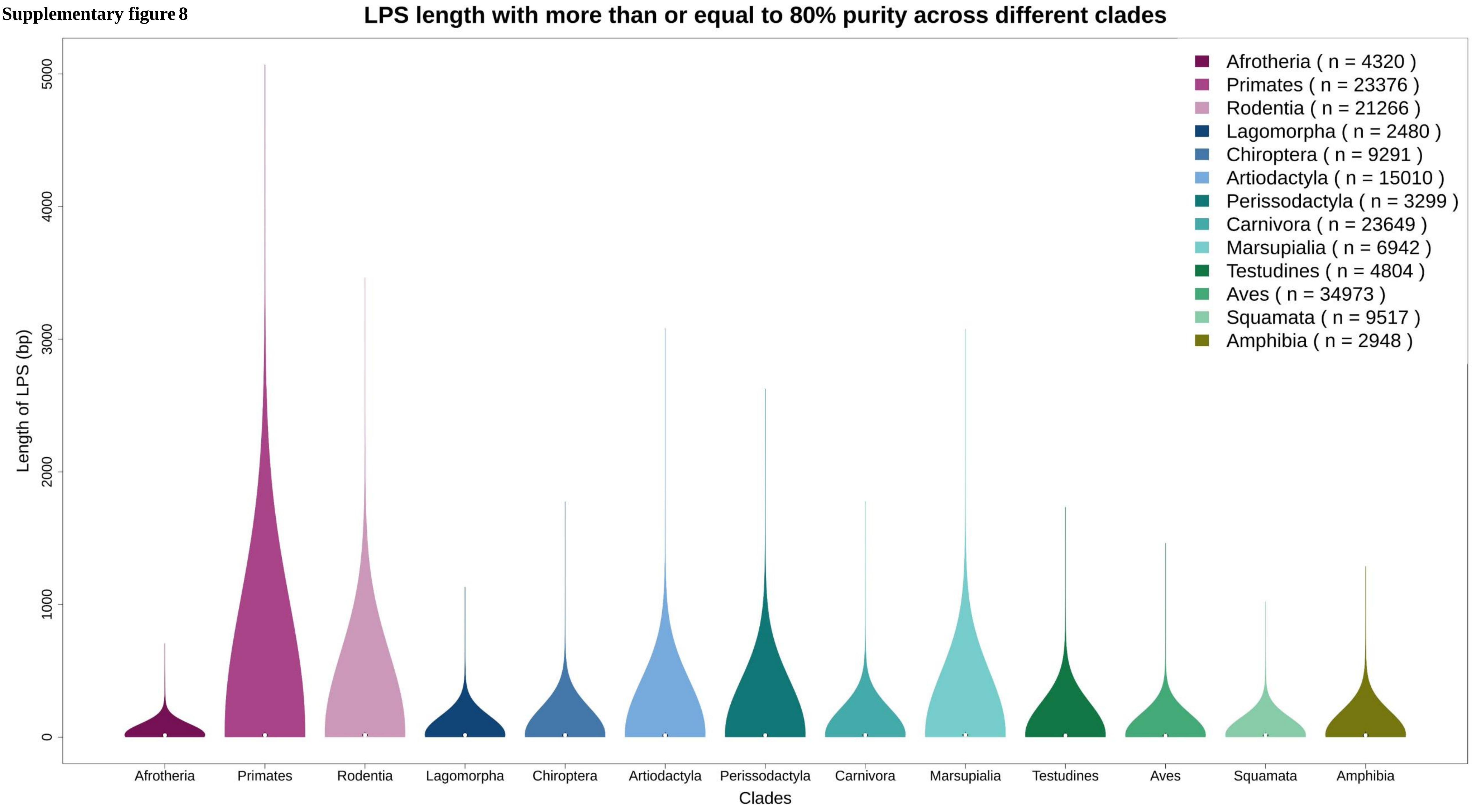

**Supplementary figure 9**

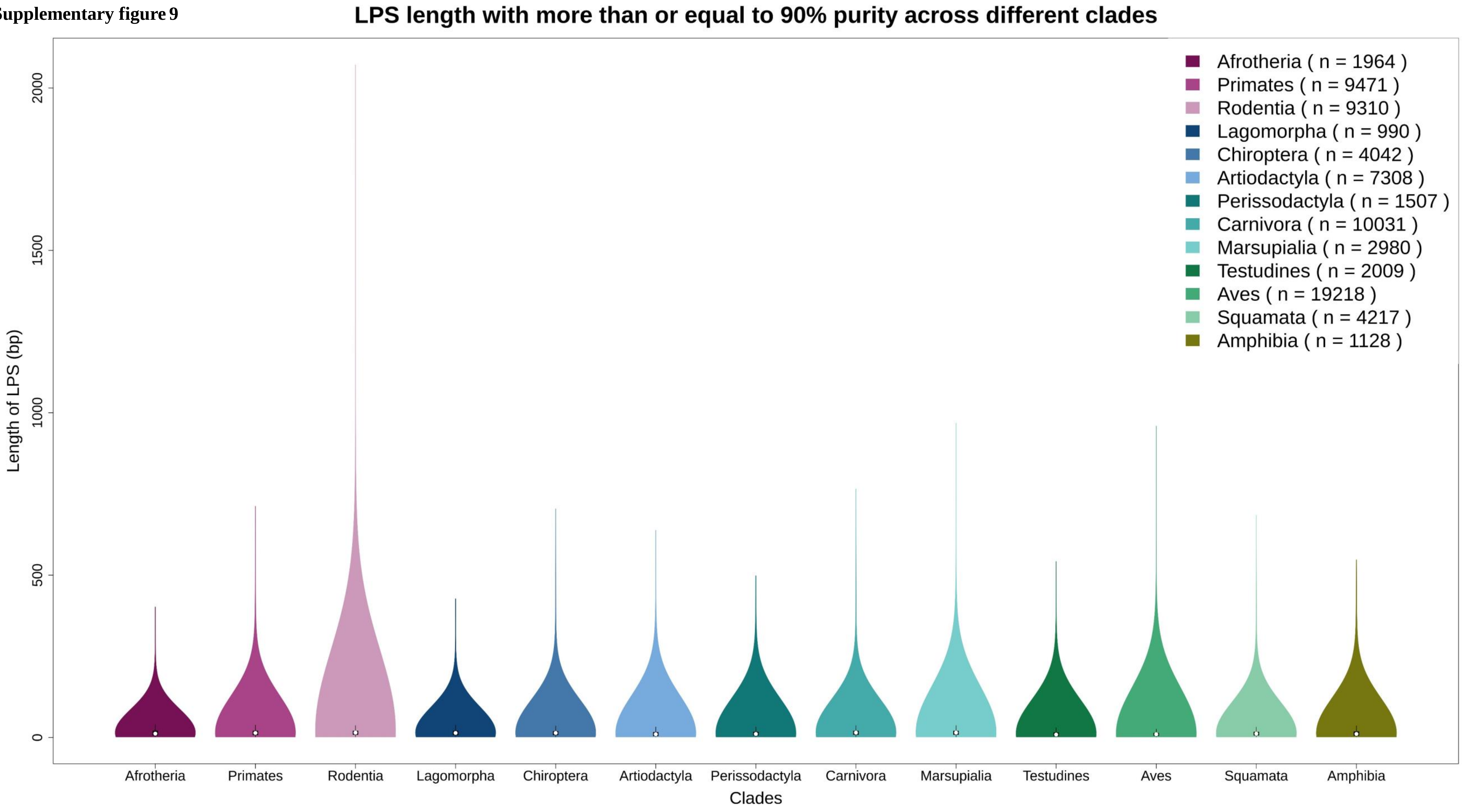

**Supplementary figure 10**

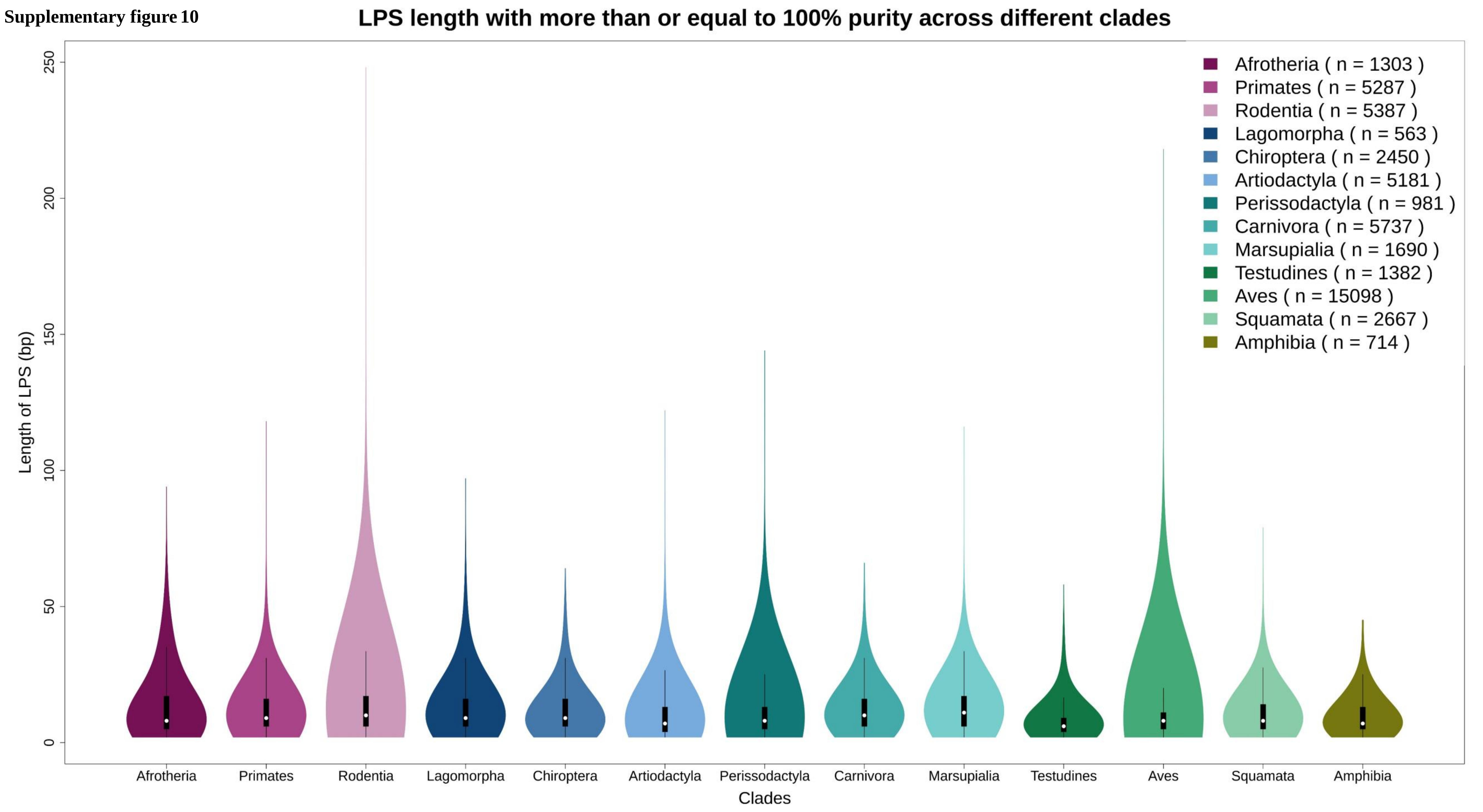

**Supplementary figure 11**

Proportion of low-complexity regions and purity in different clades

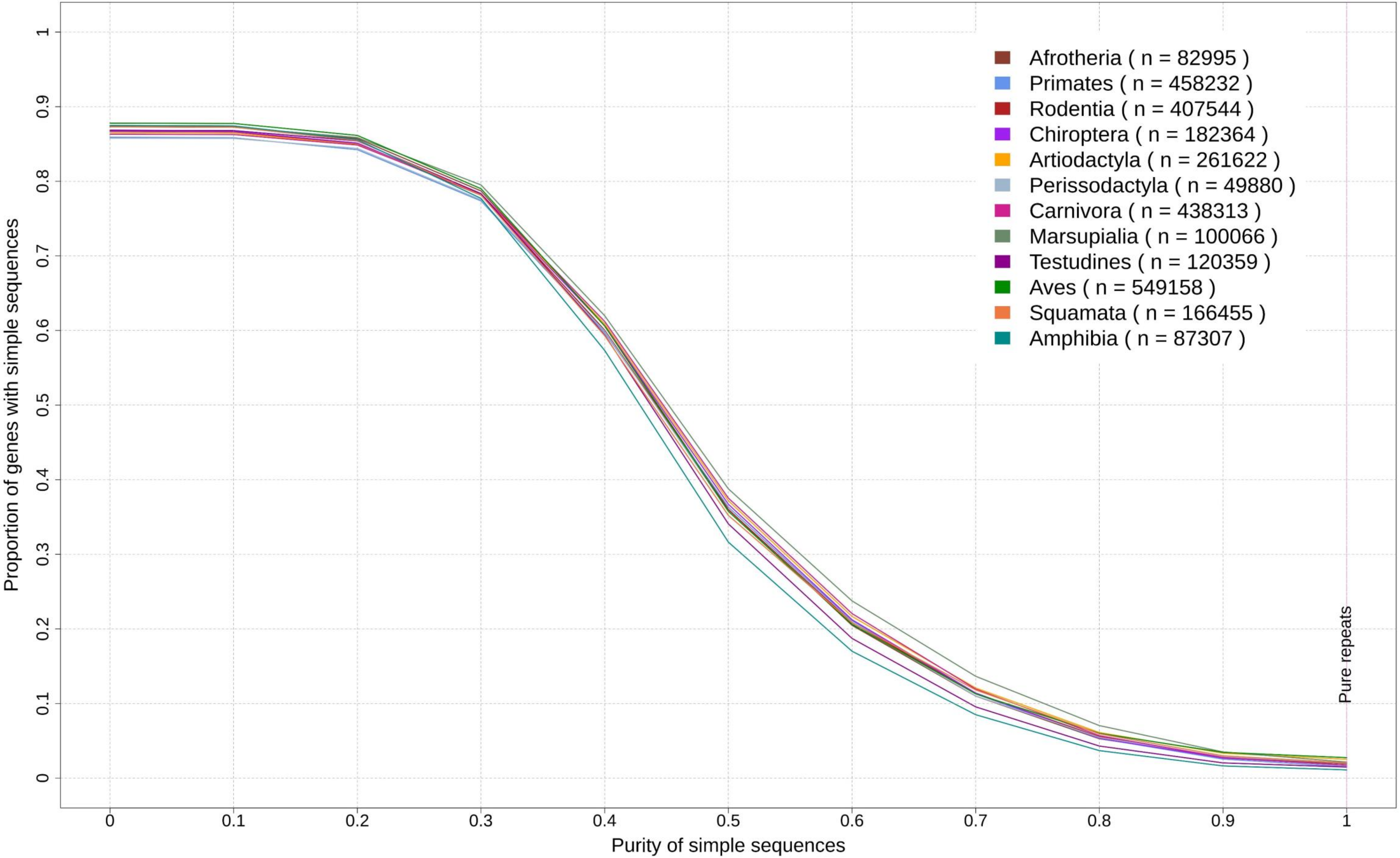

**Supplementary figure 12**

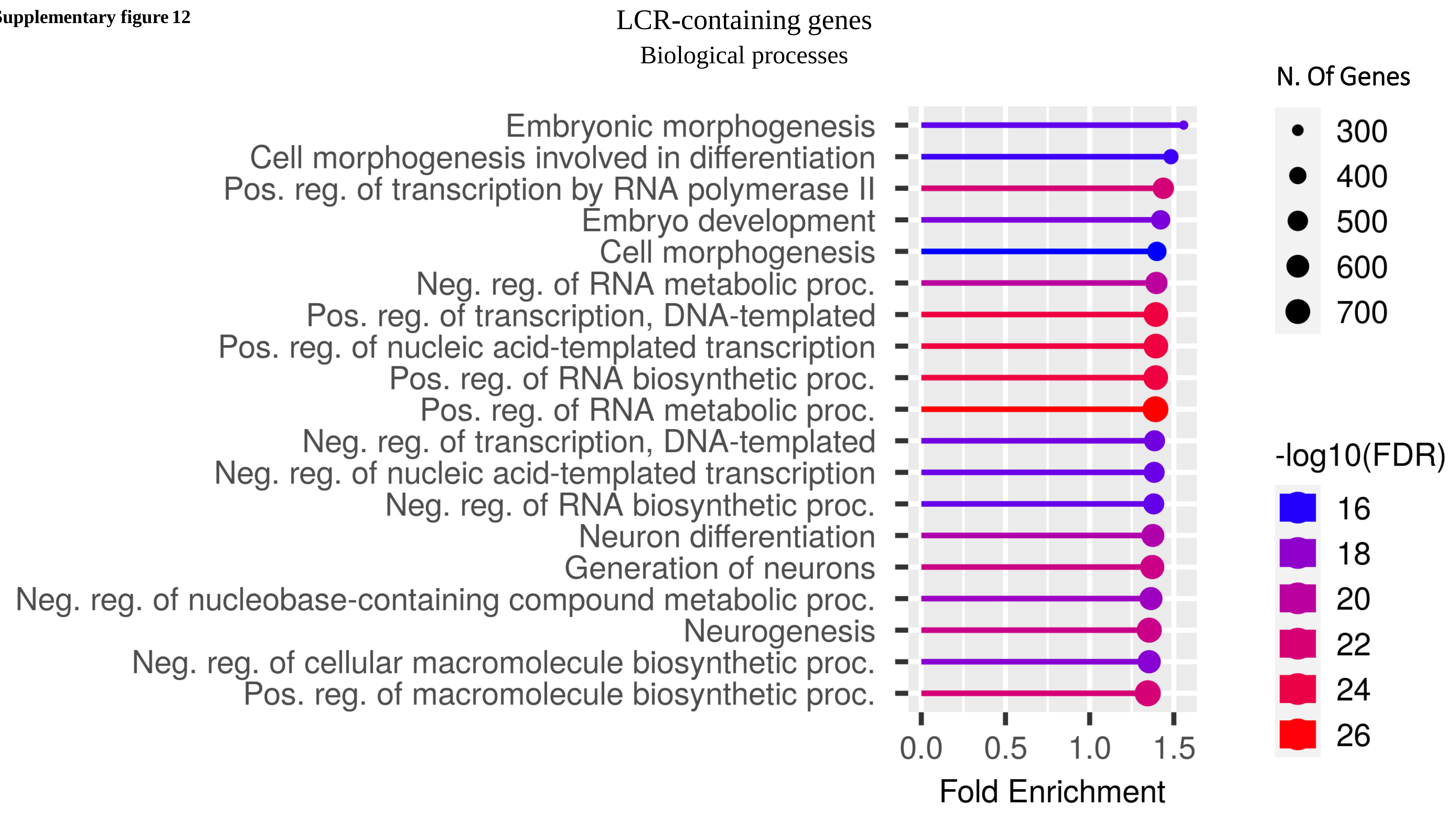

**Supplementary figure 13**

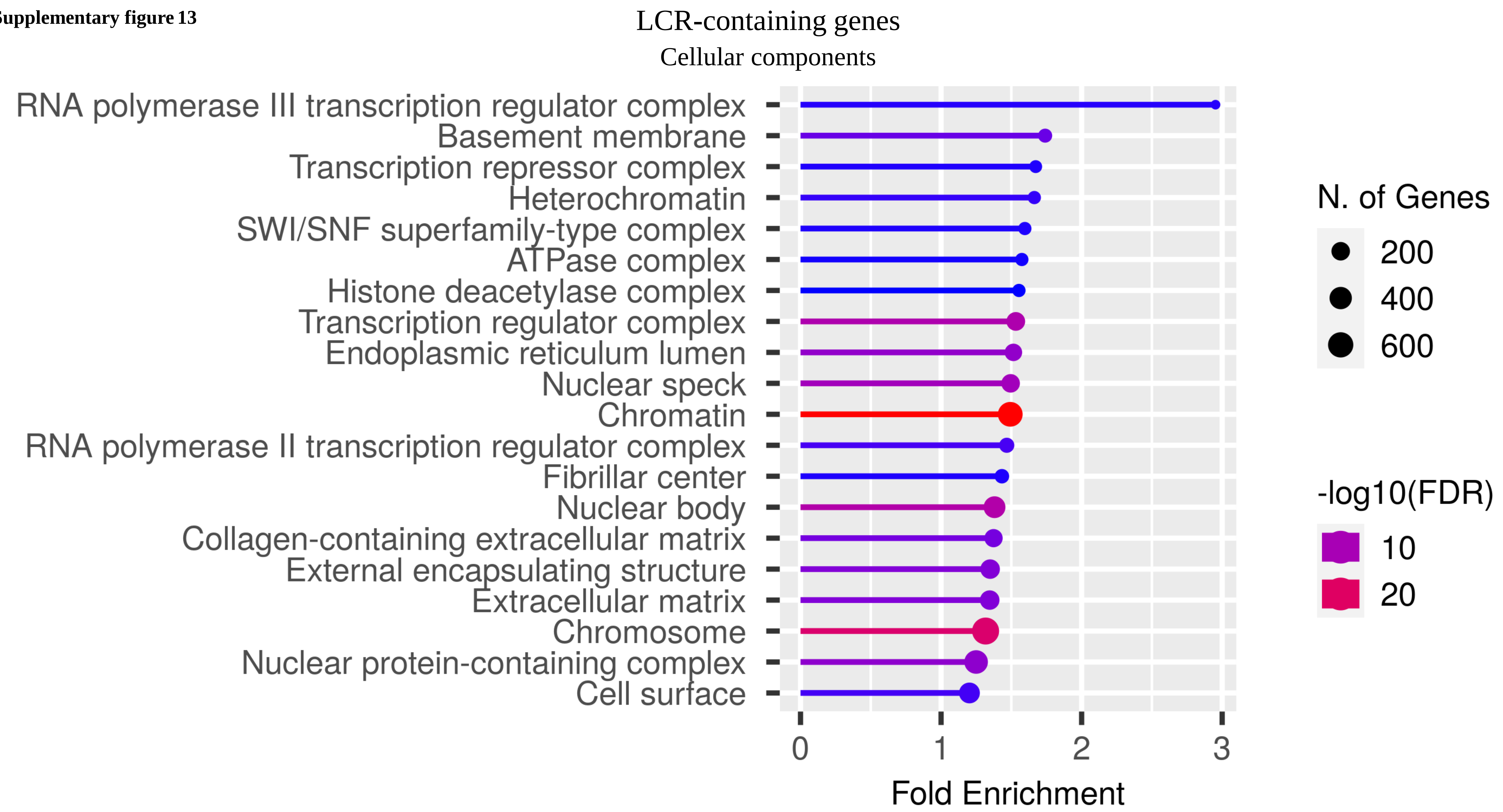

**Supplementary figure 14**

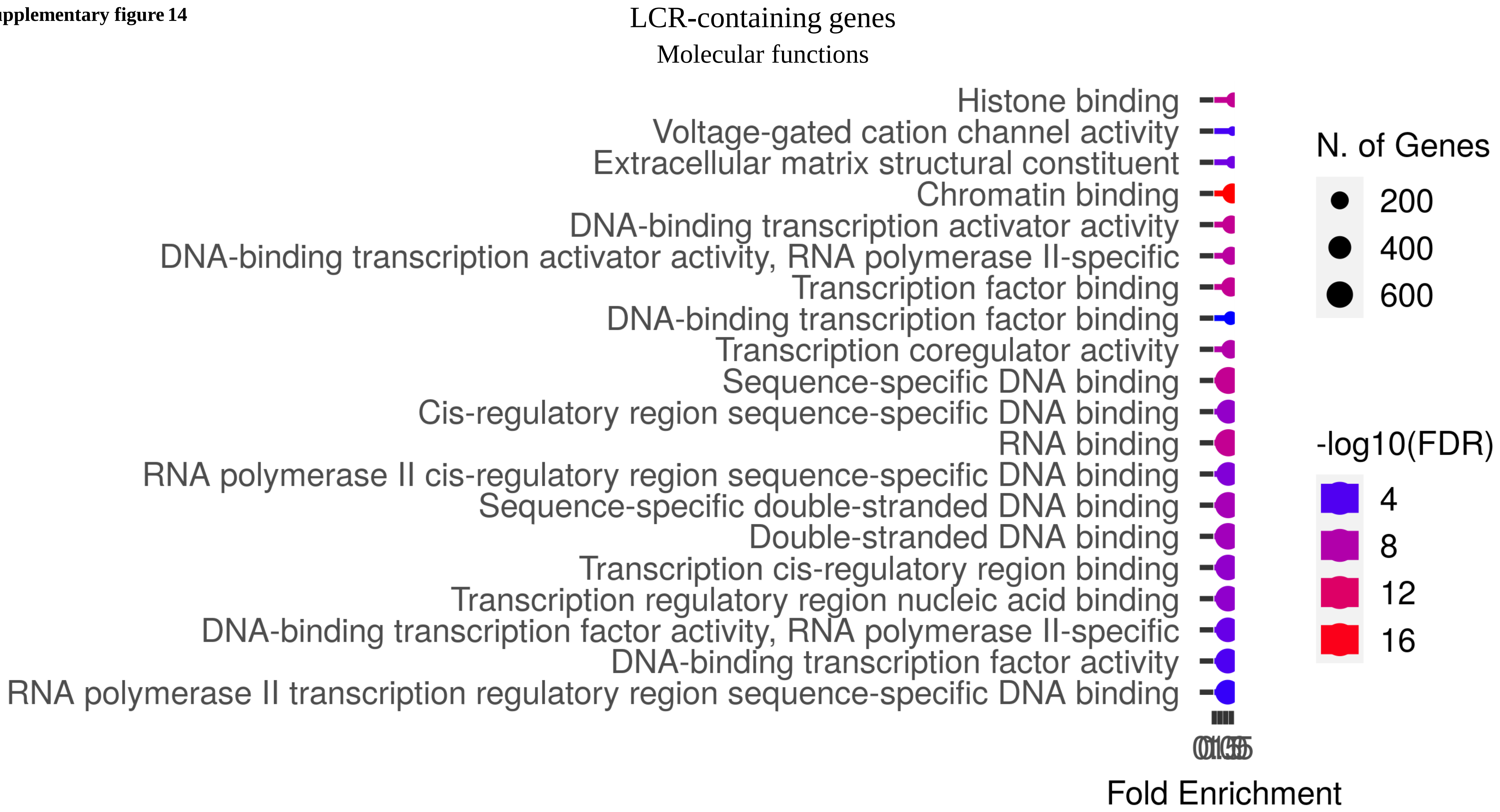

**Supplementary figure 15**

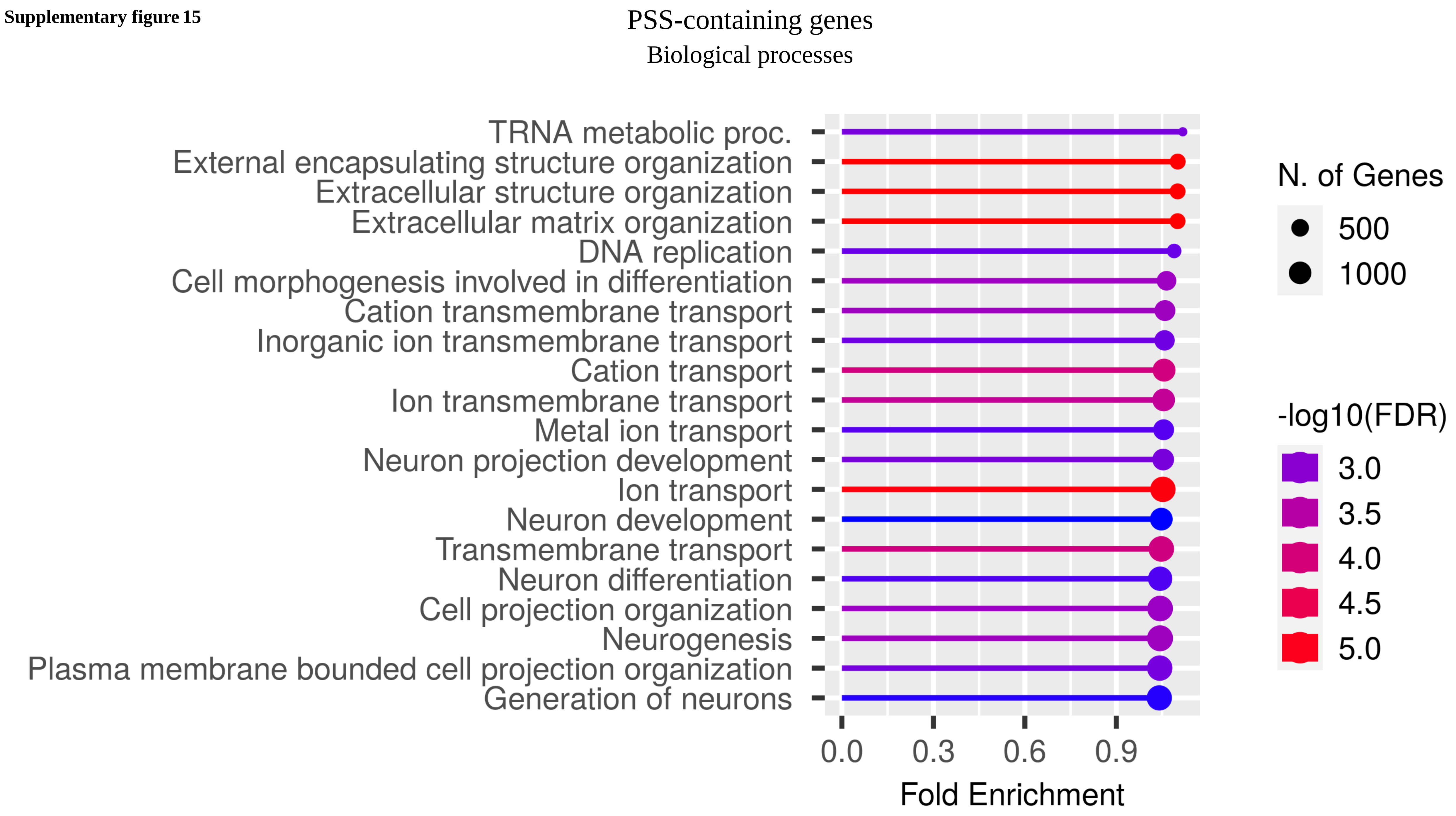

**Supplementary figure 16**

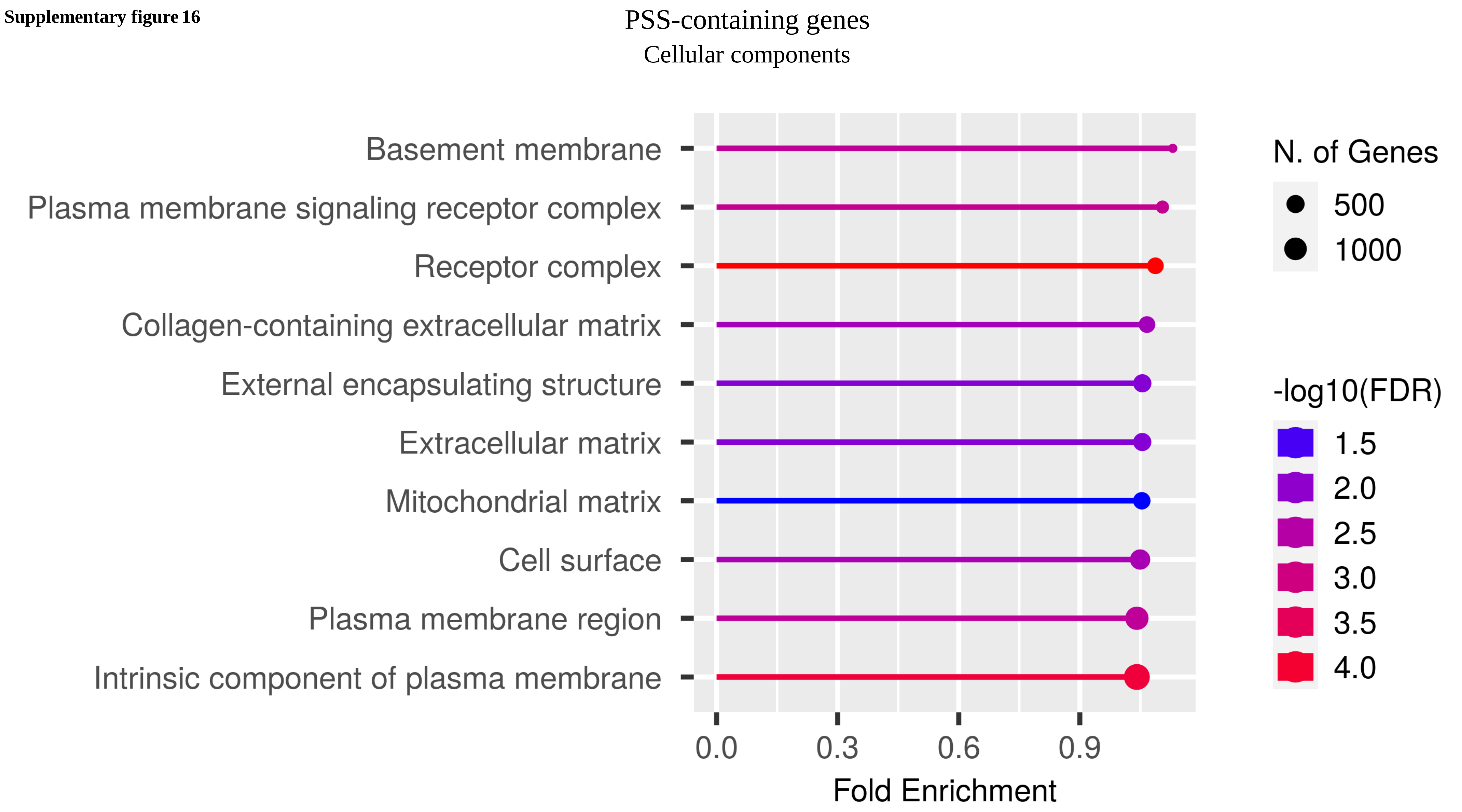

Supplementary figure 17

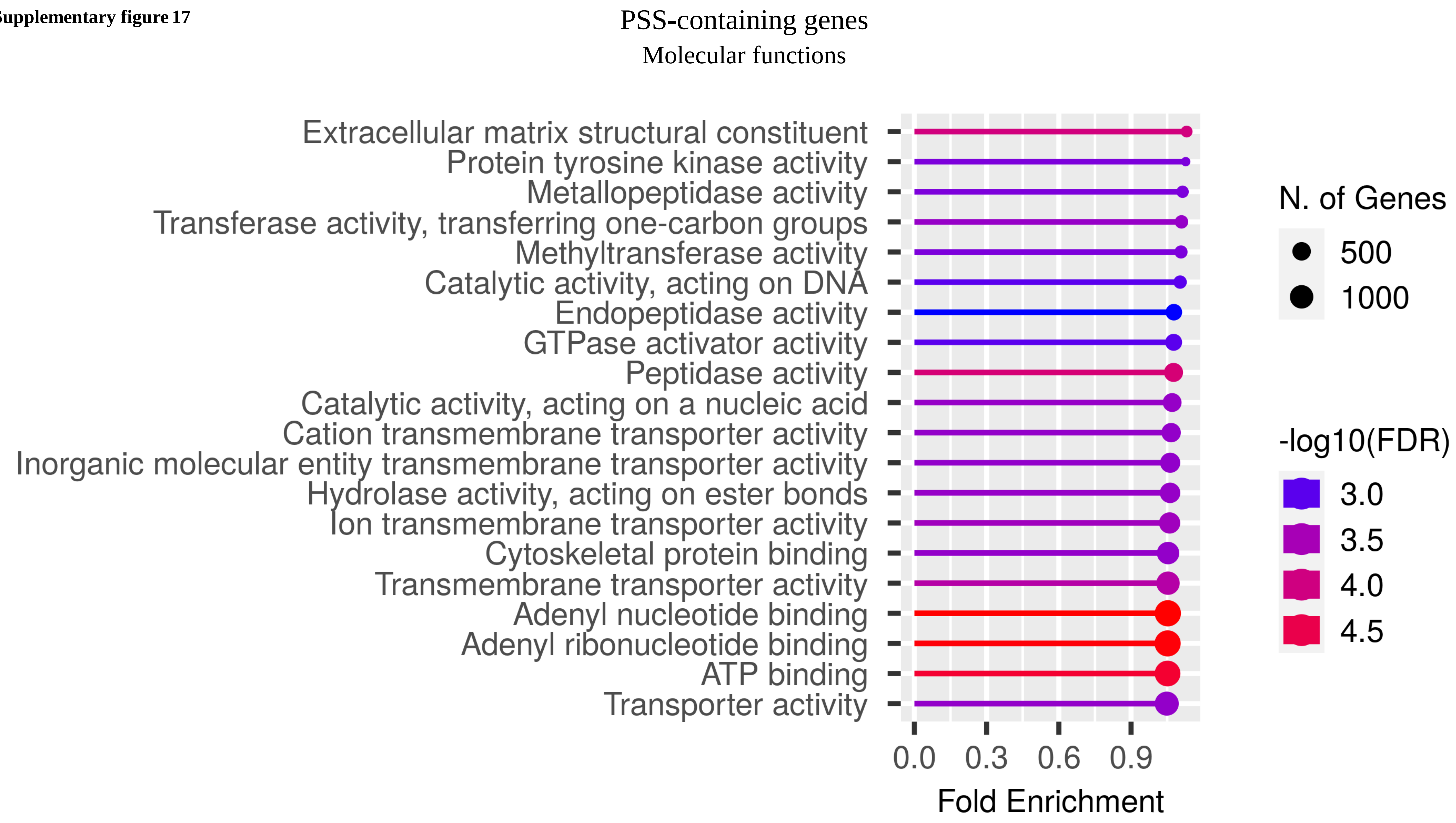

**Supplementary figure 18**

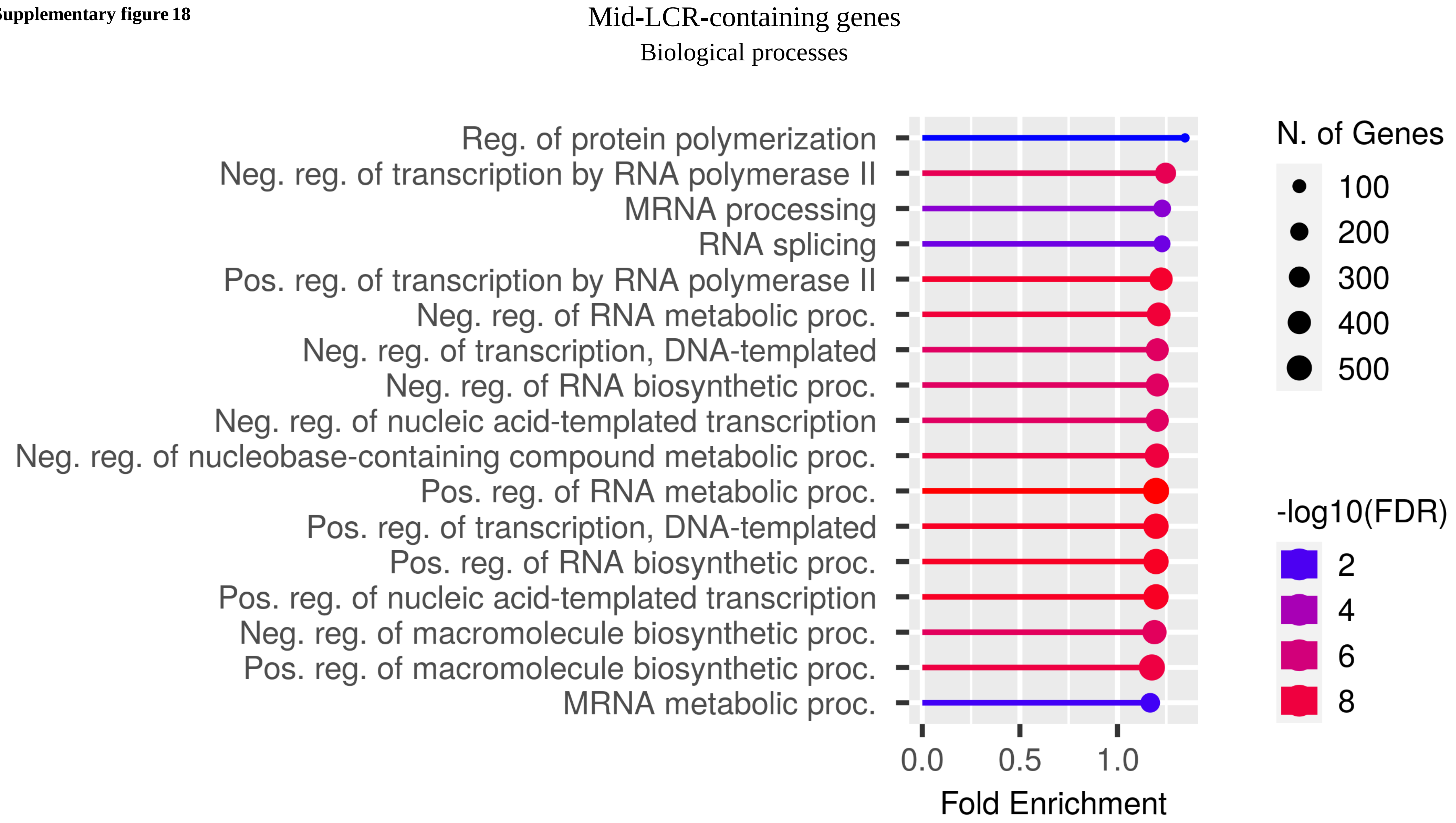

**Supplementary figure 19**

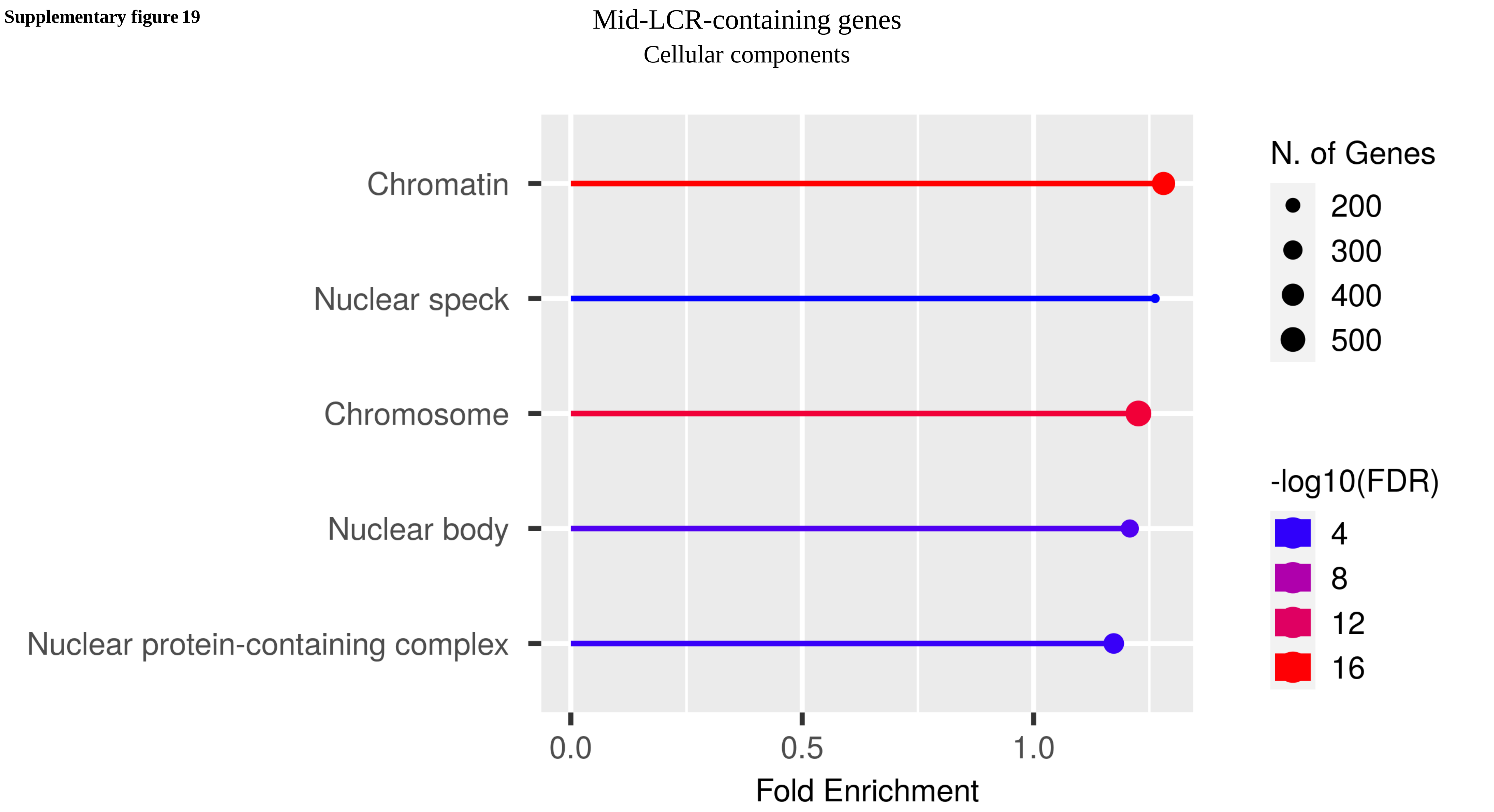

**Supplementary figure 20**

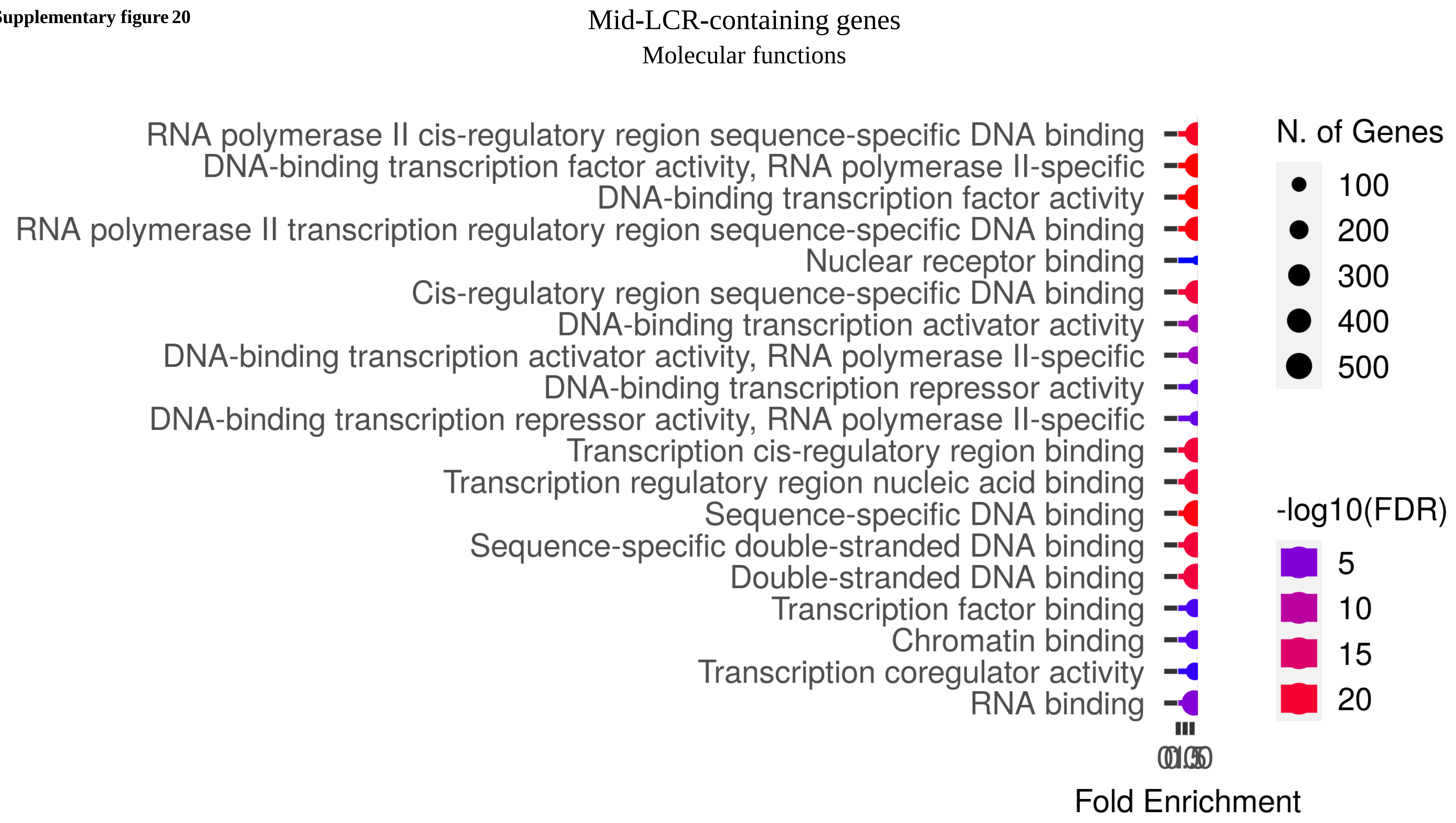

**Supplementary figure 21**

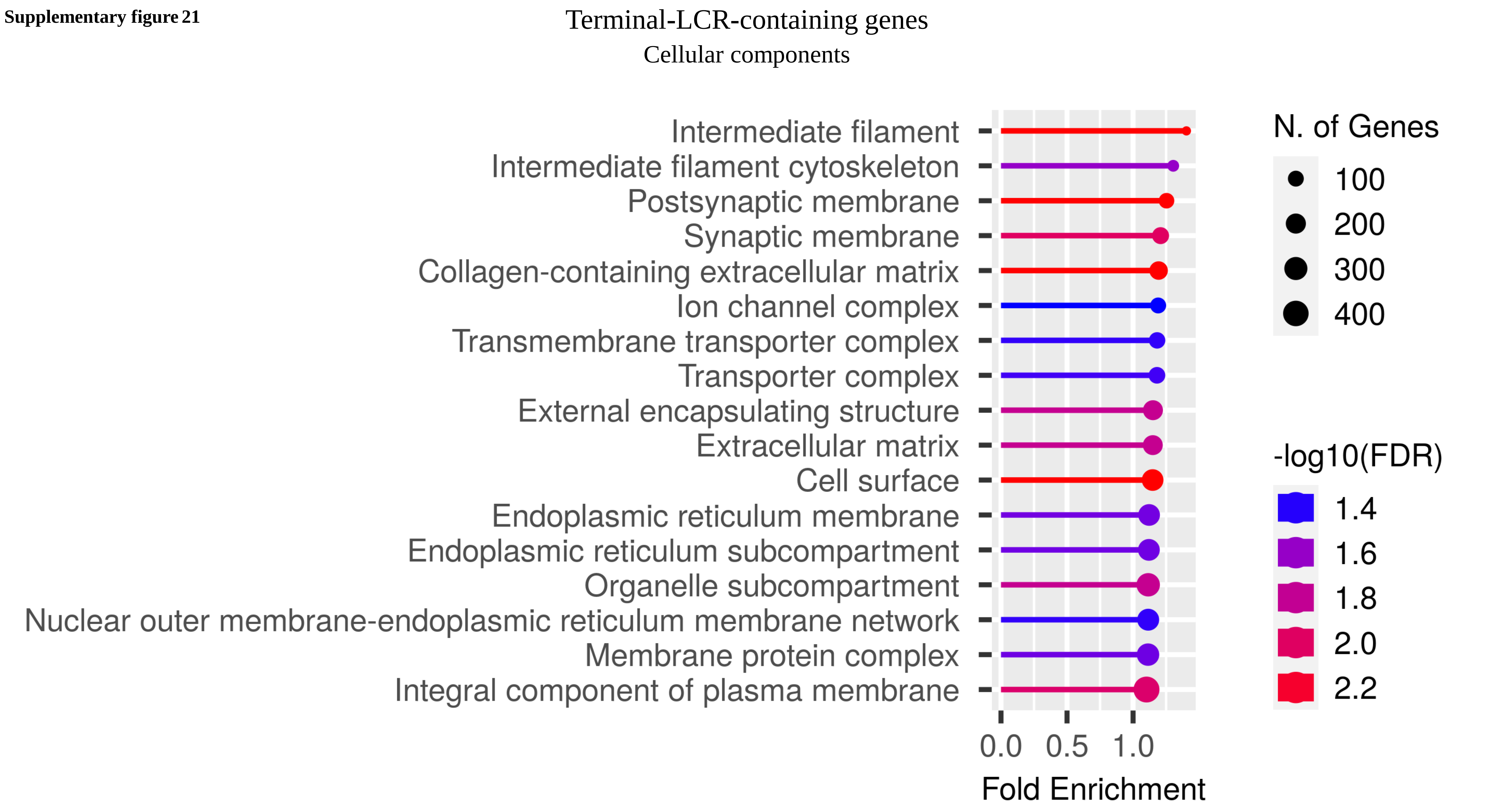

**Supplementary figure 22**

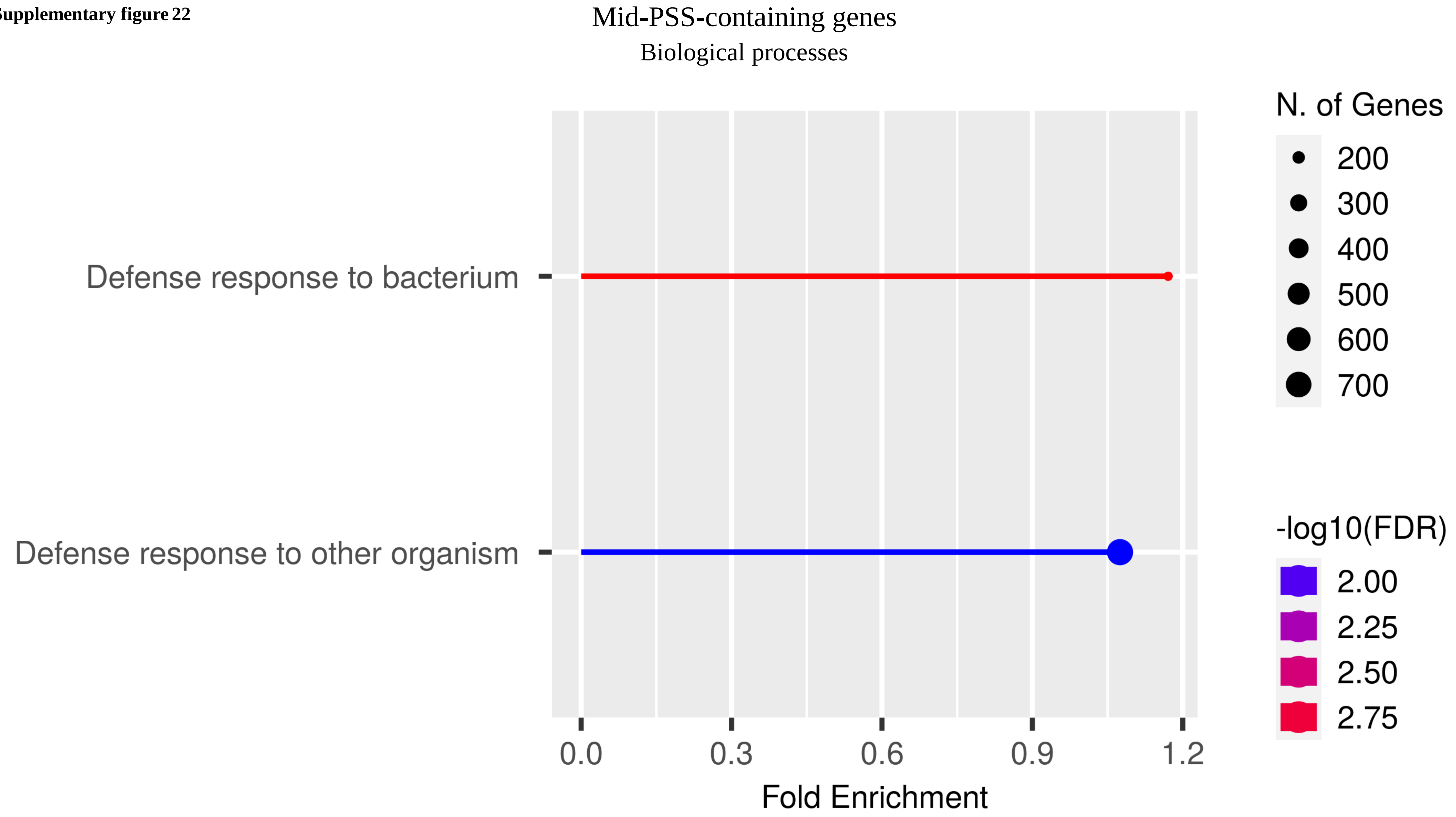

Supplementary figure 23

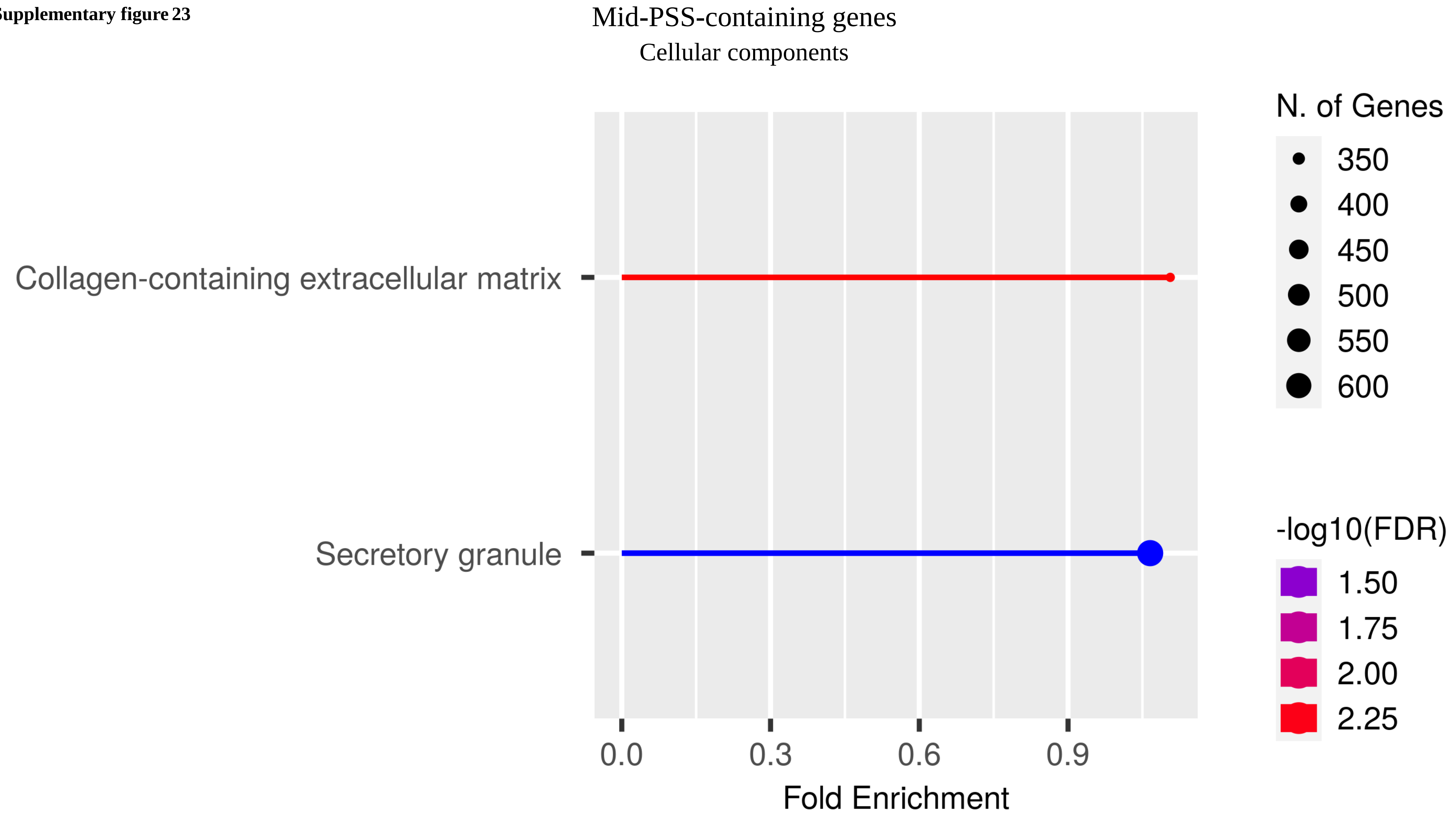

**Supplementary figure 24**

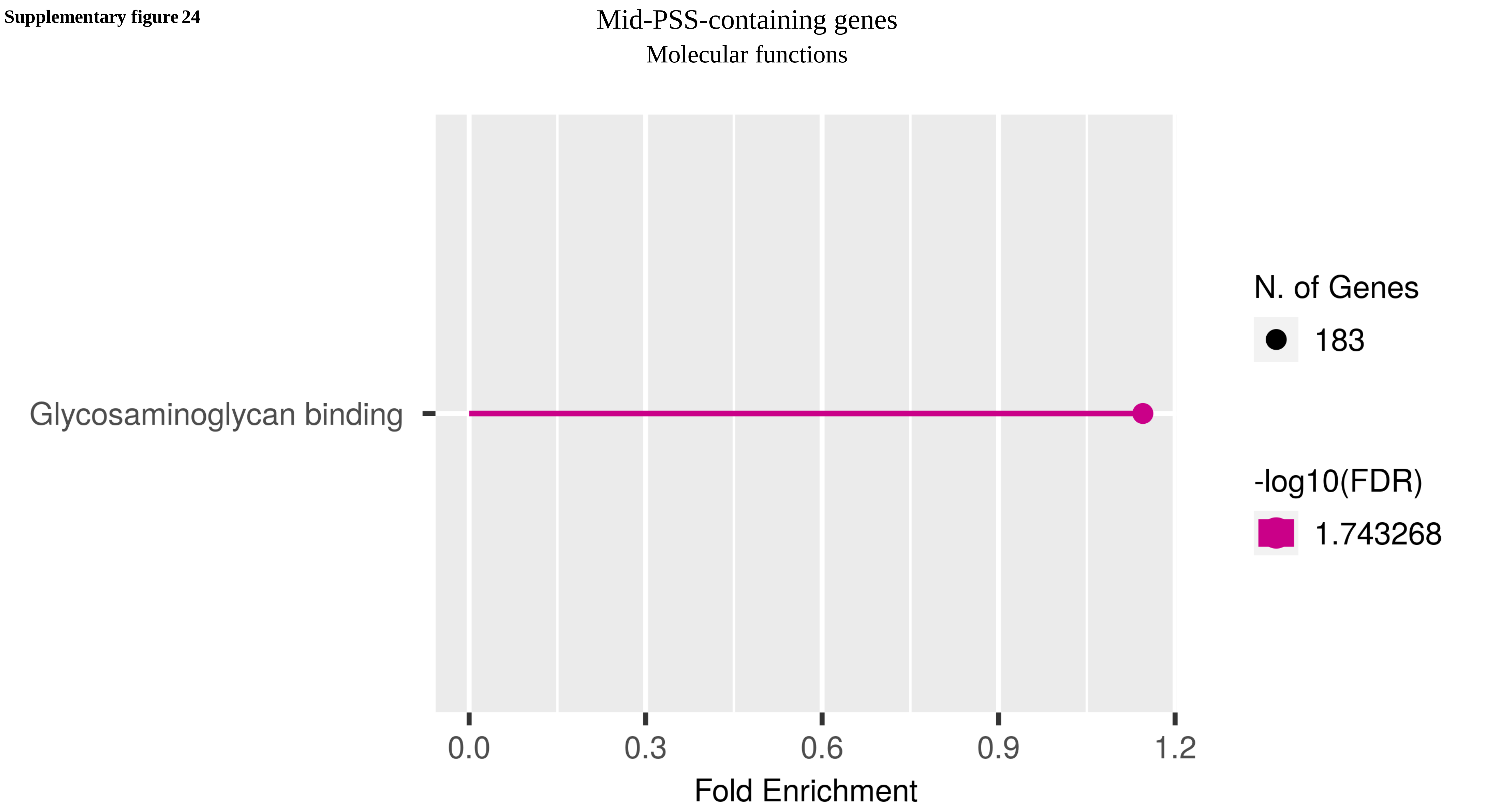

**Supplementary figure 25**

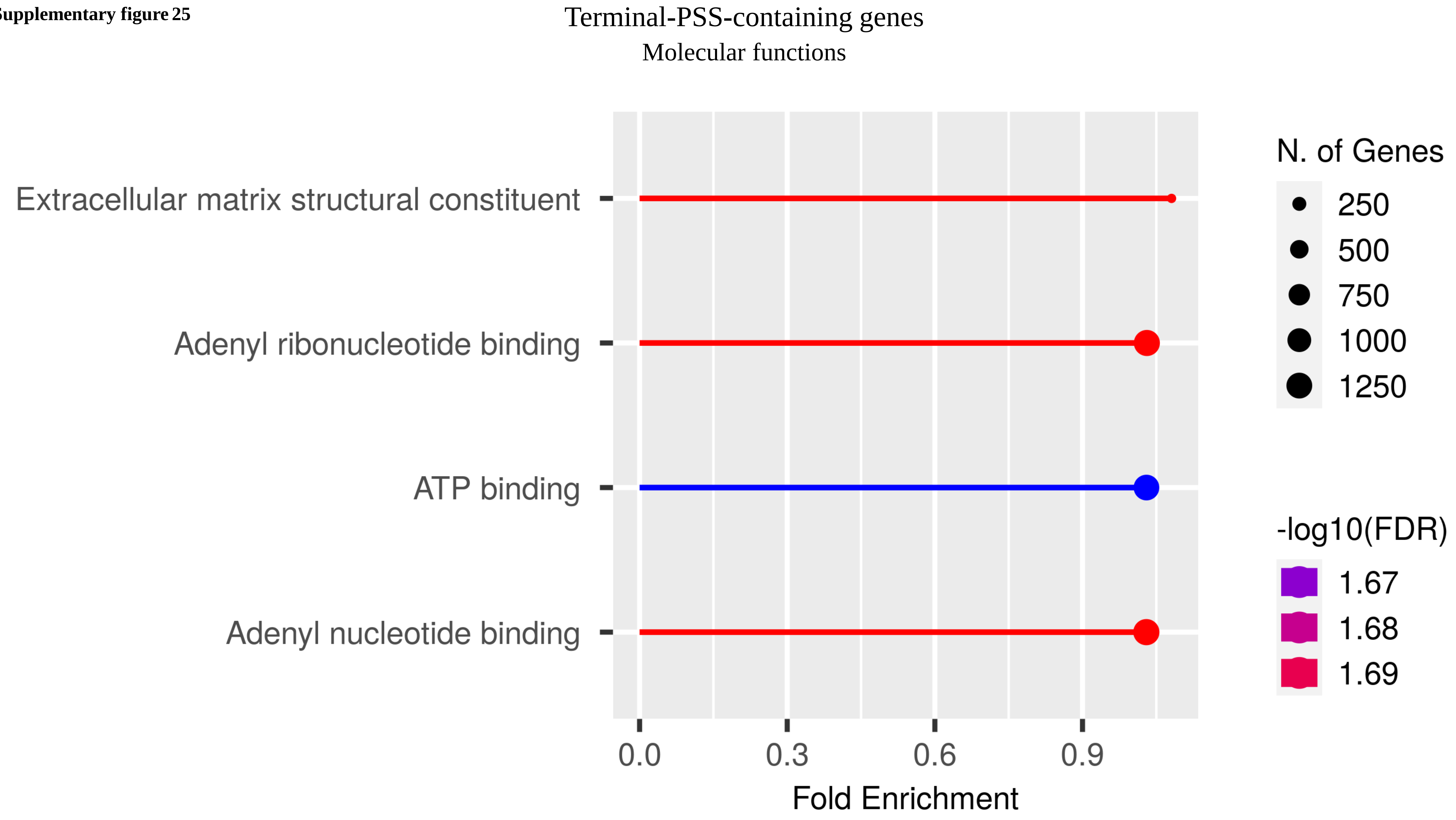

**Supplementary figure 26**

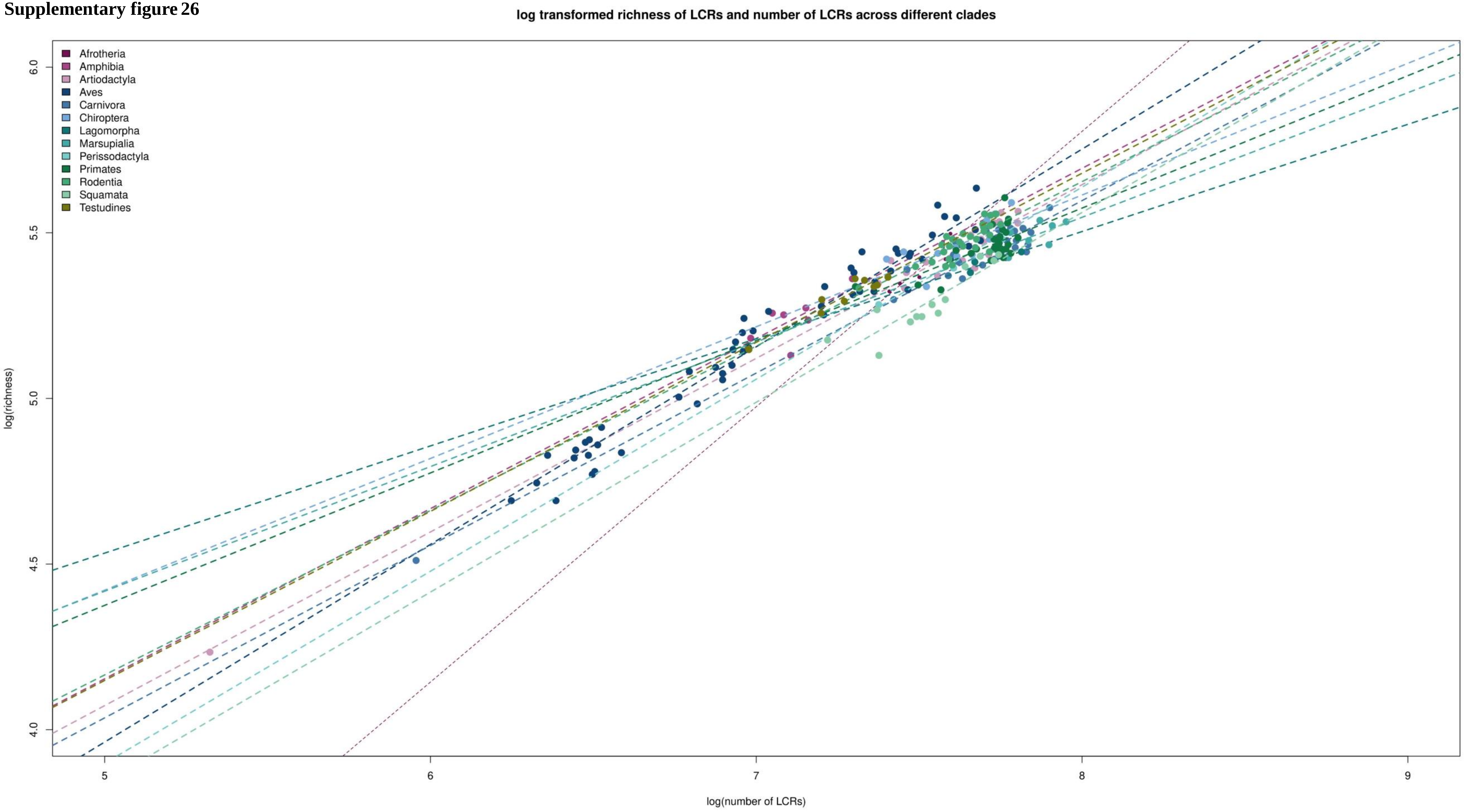

**Supplementary figure 27**

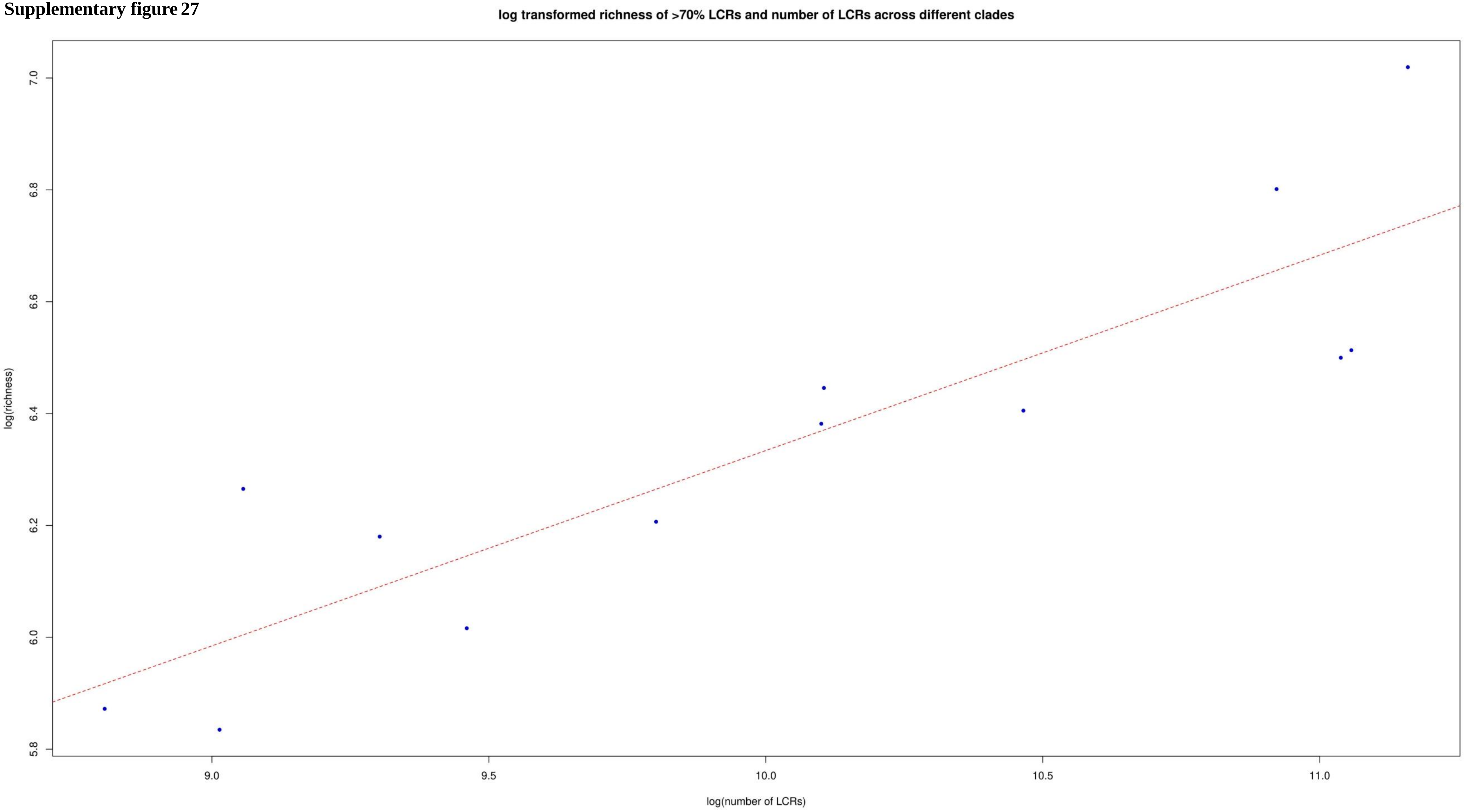

**Supplementary figure 28**

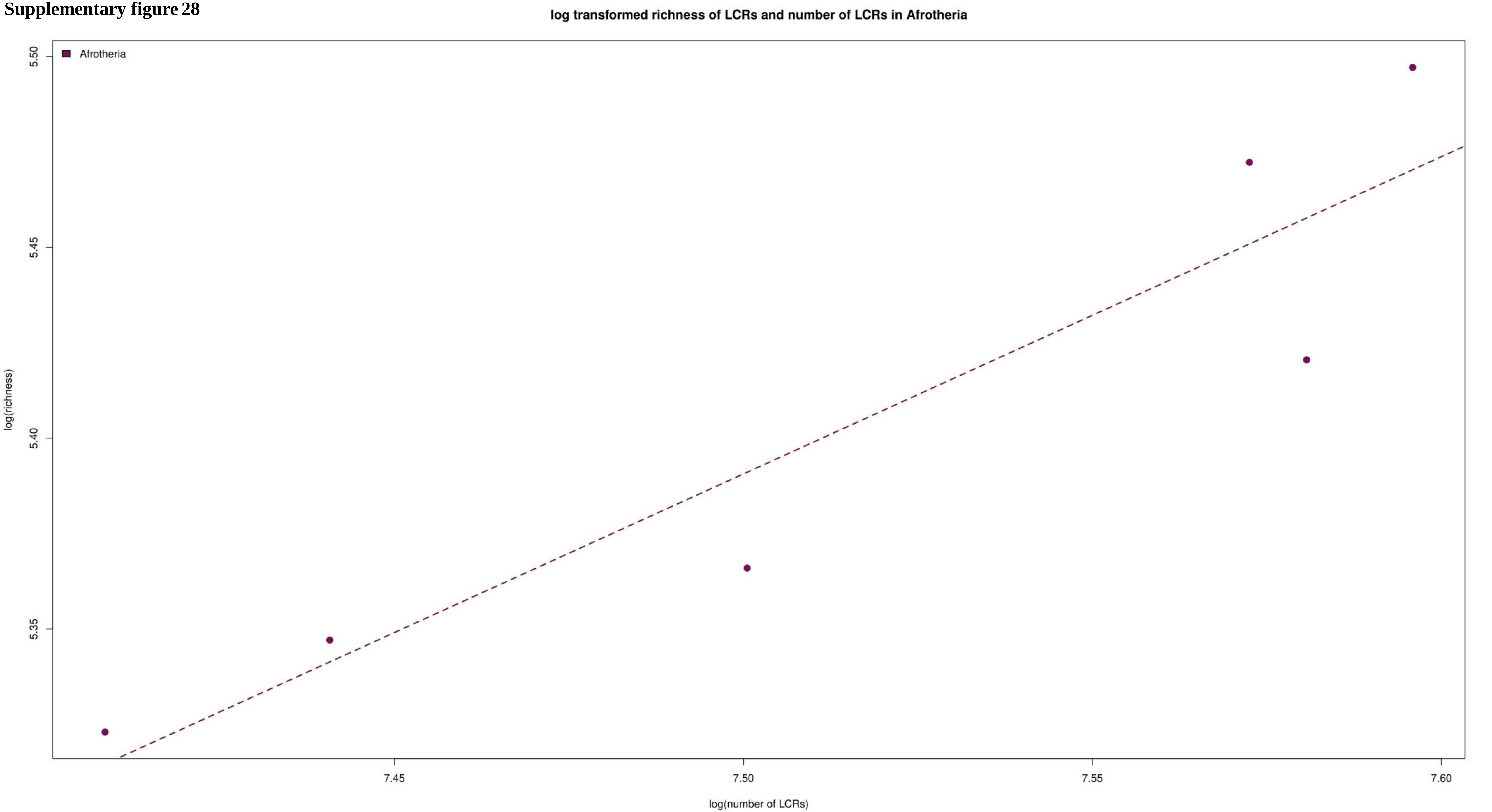

**Supplementary figure 29**

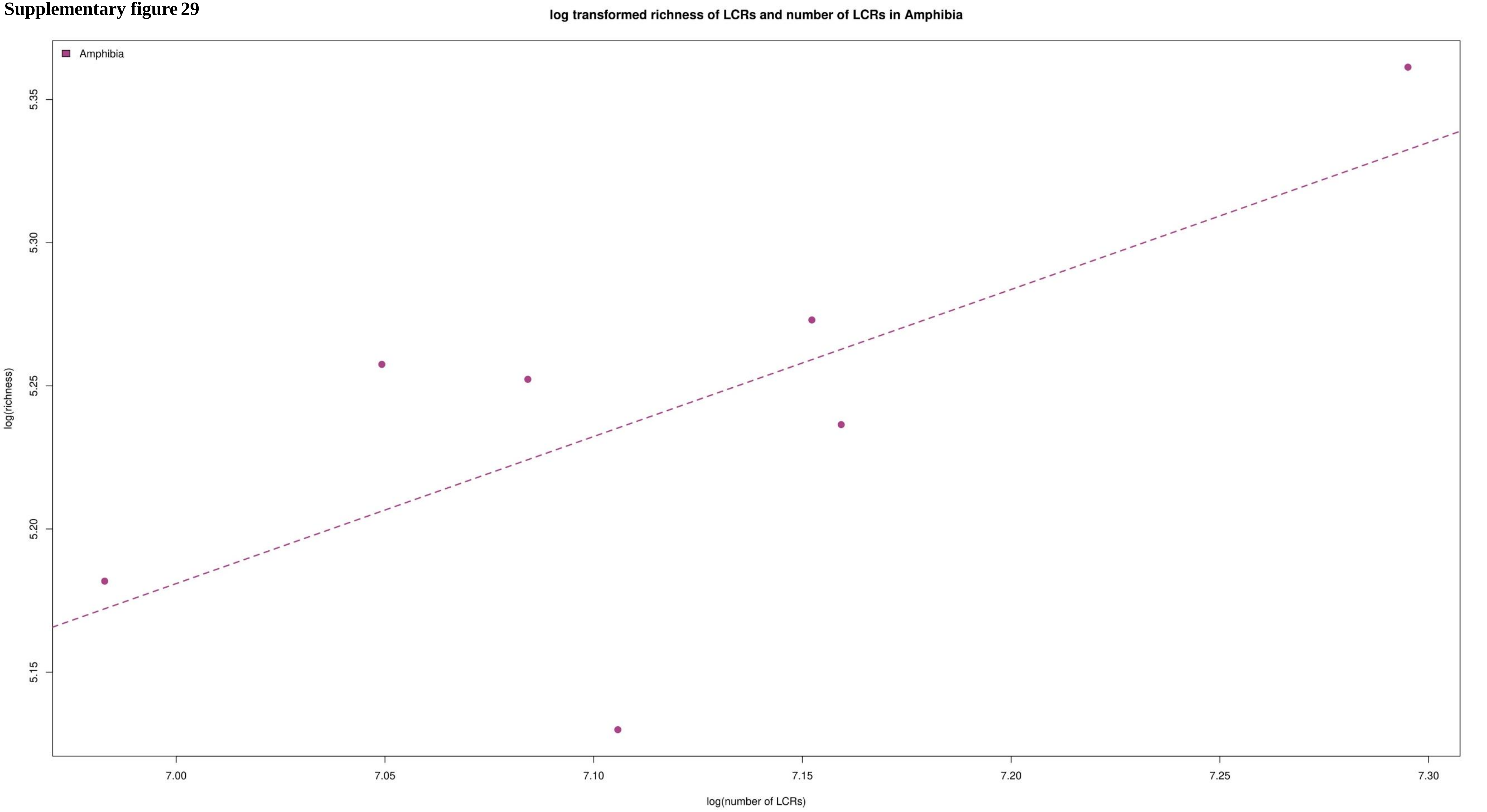

**Supplementary figure 30**

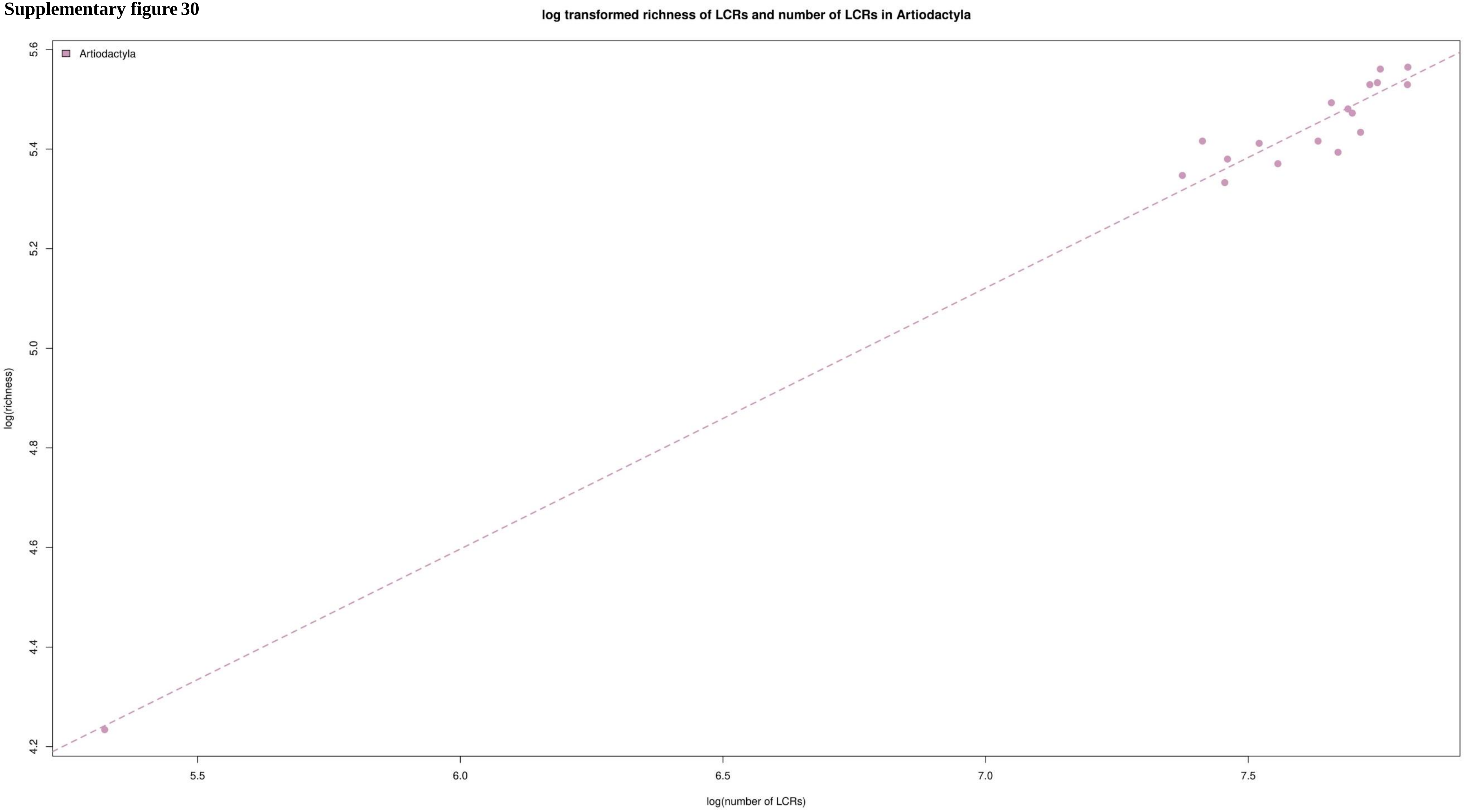

**Supplementary figure 31**

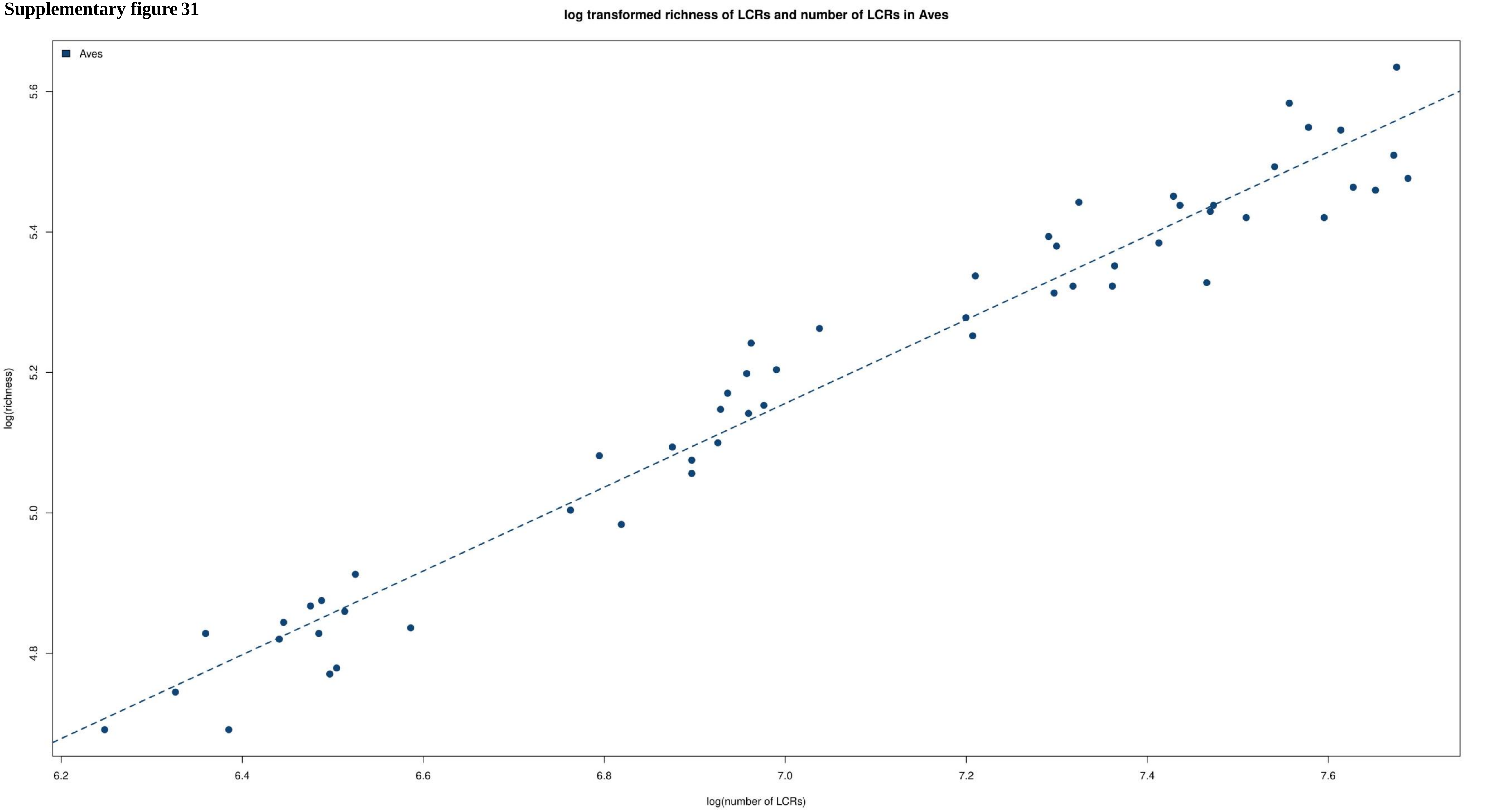

**Supplementary figure 32**

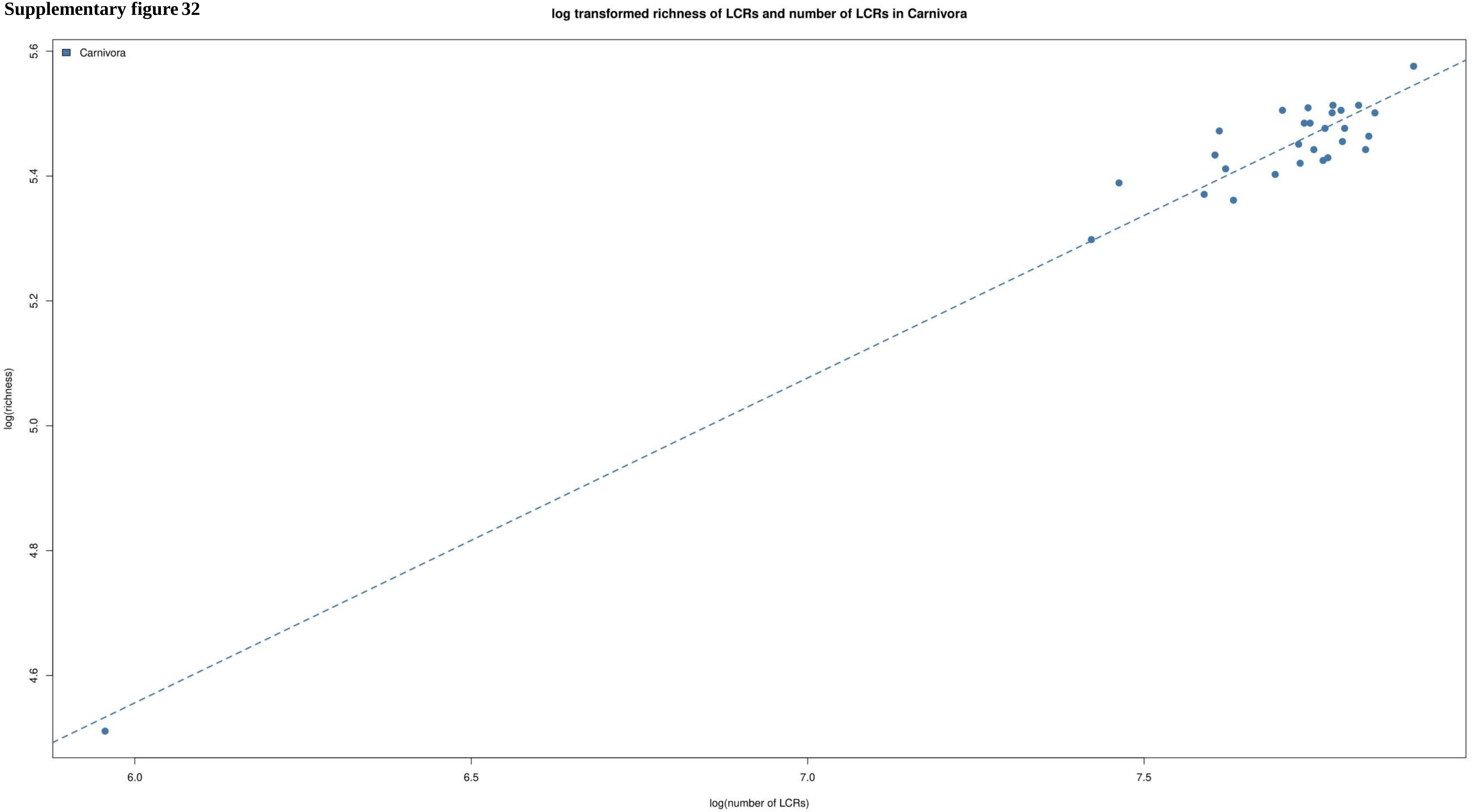

**Supplementary figure 33**

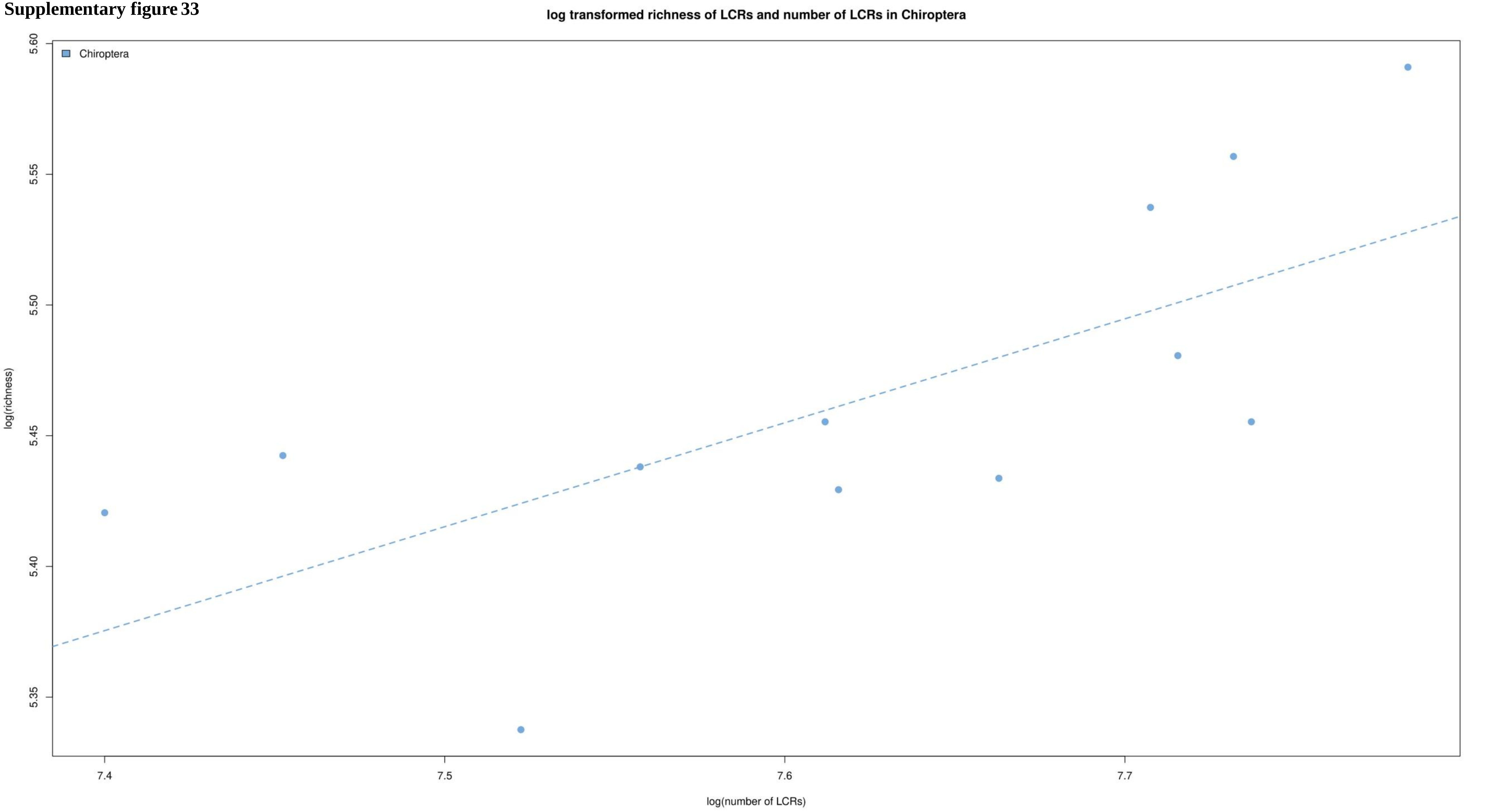

**Supplementary figure 34**

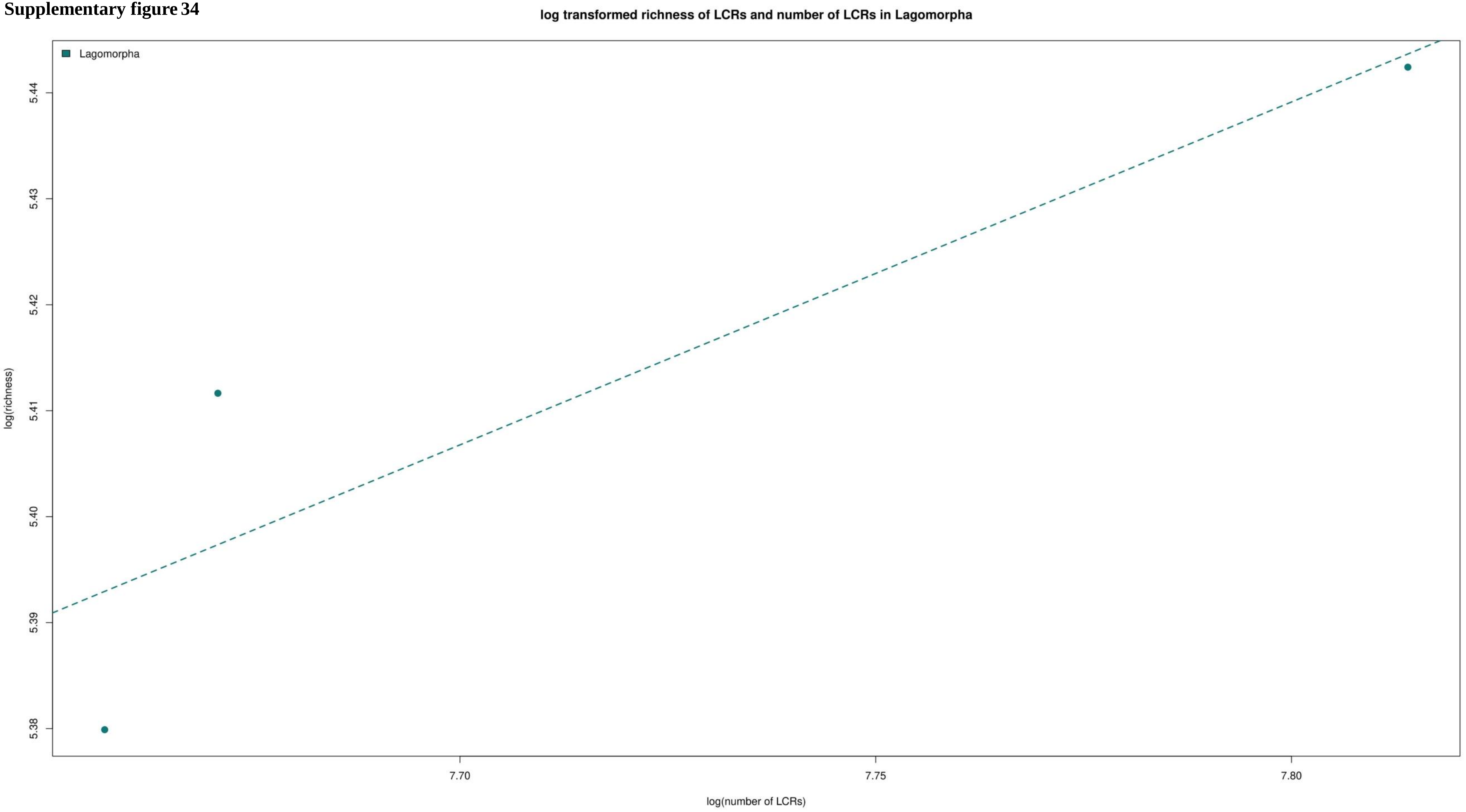

**Supplementary figure 35**

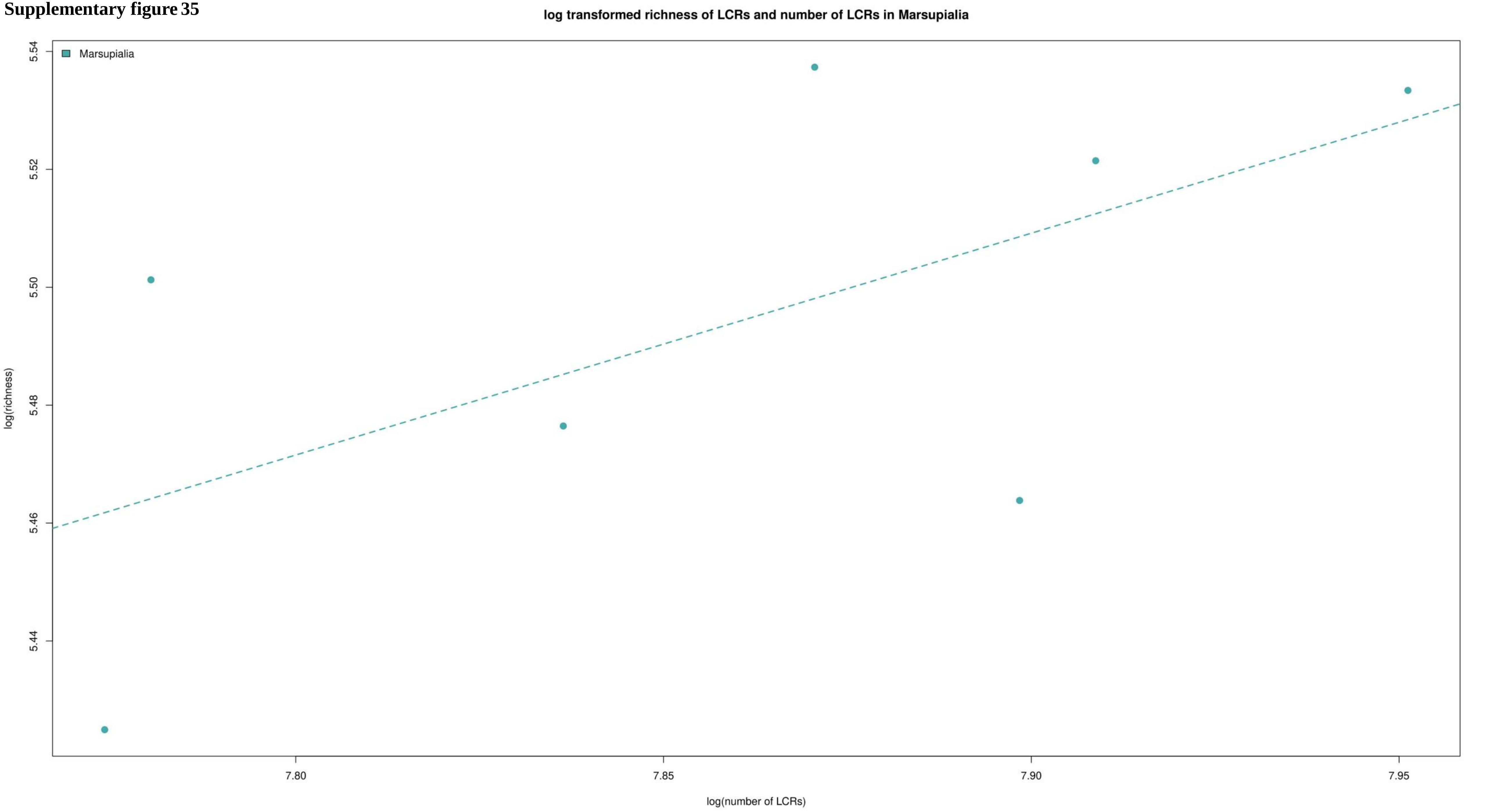

**Supplementary figure 36**

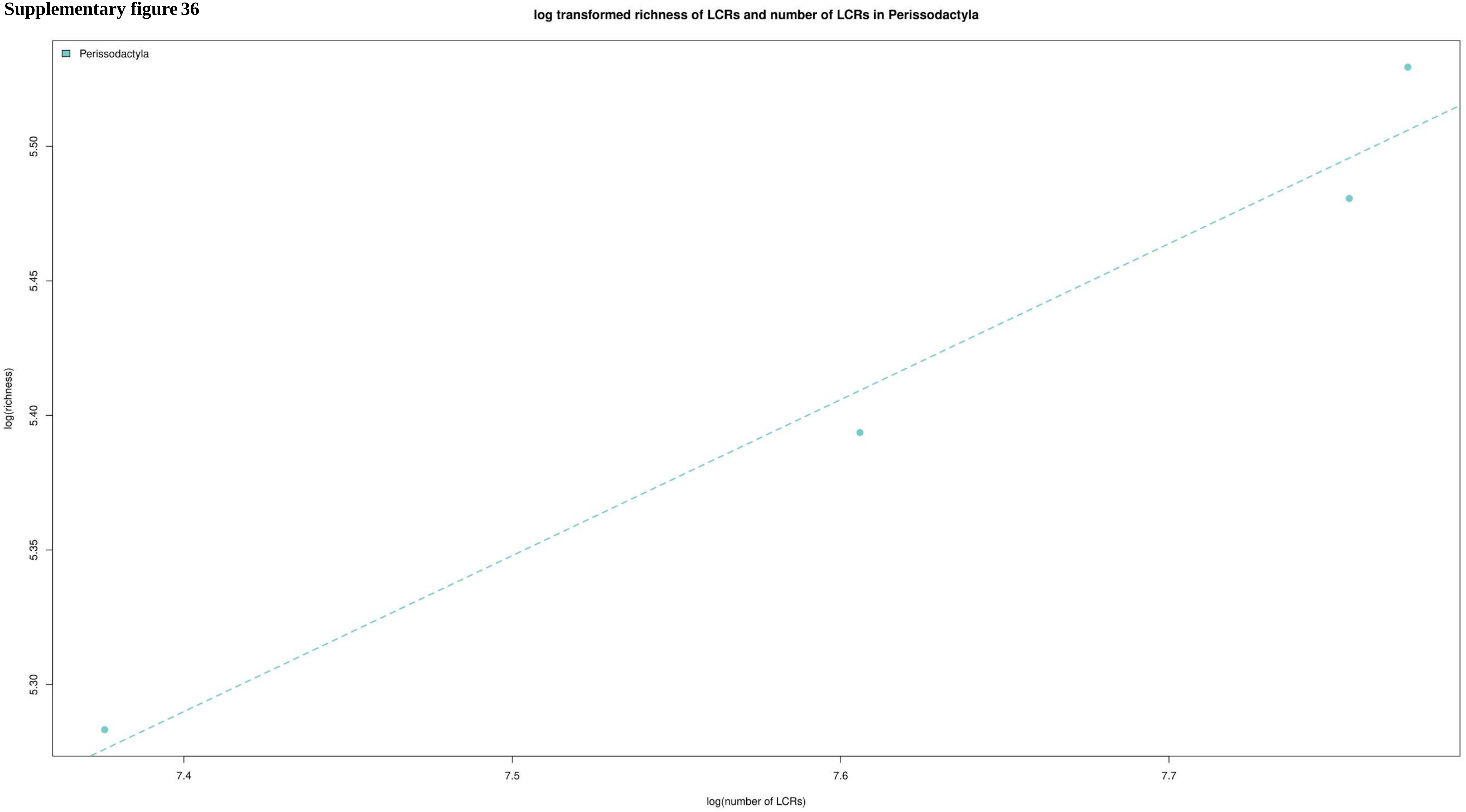

**Supplementary figure 37**

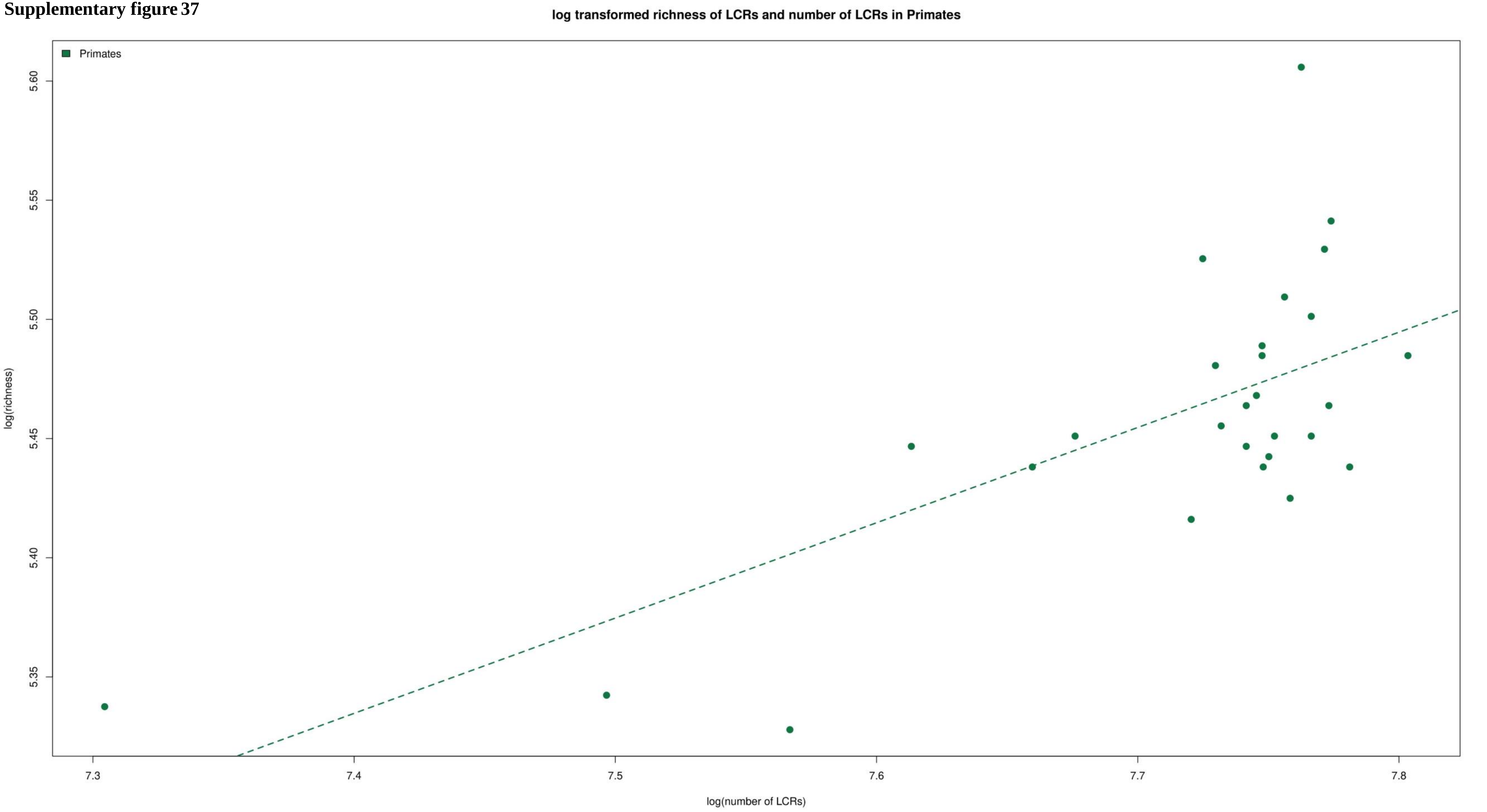

**Supplementary figure 38**

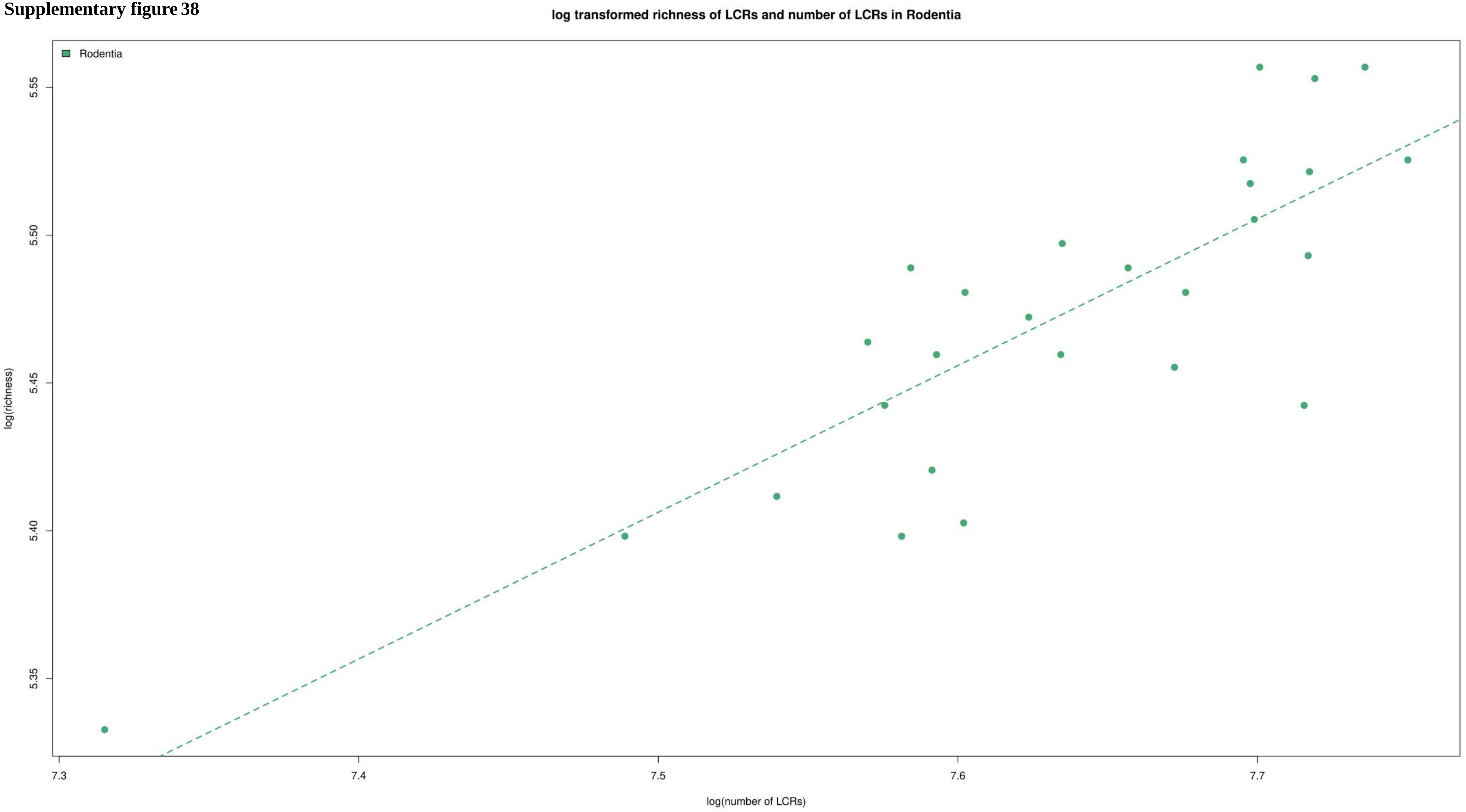

**Supplementary figure 39**

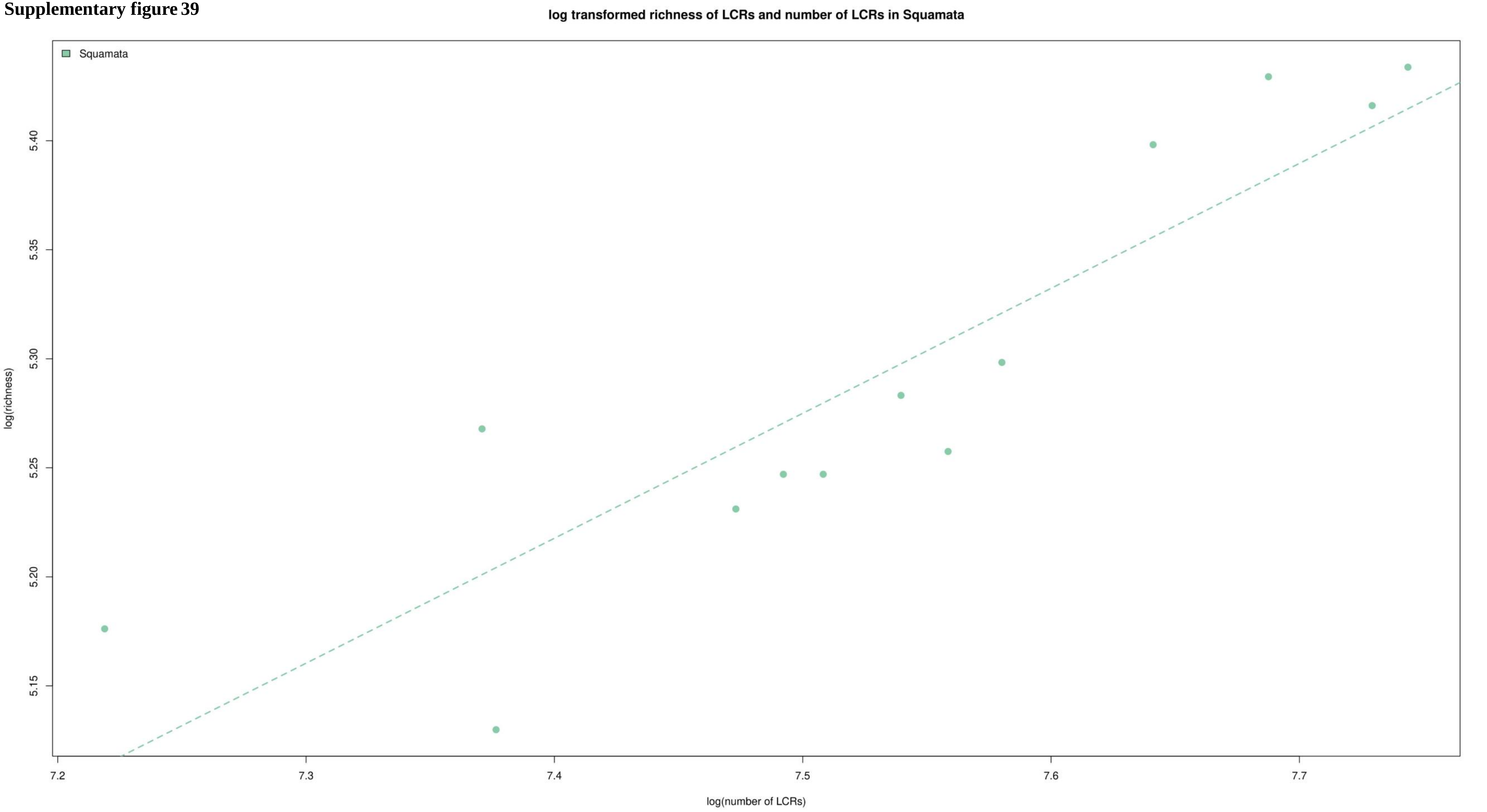

**Supplementary figure 40**

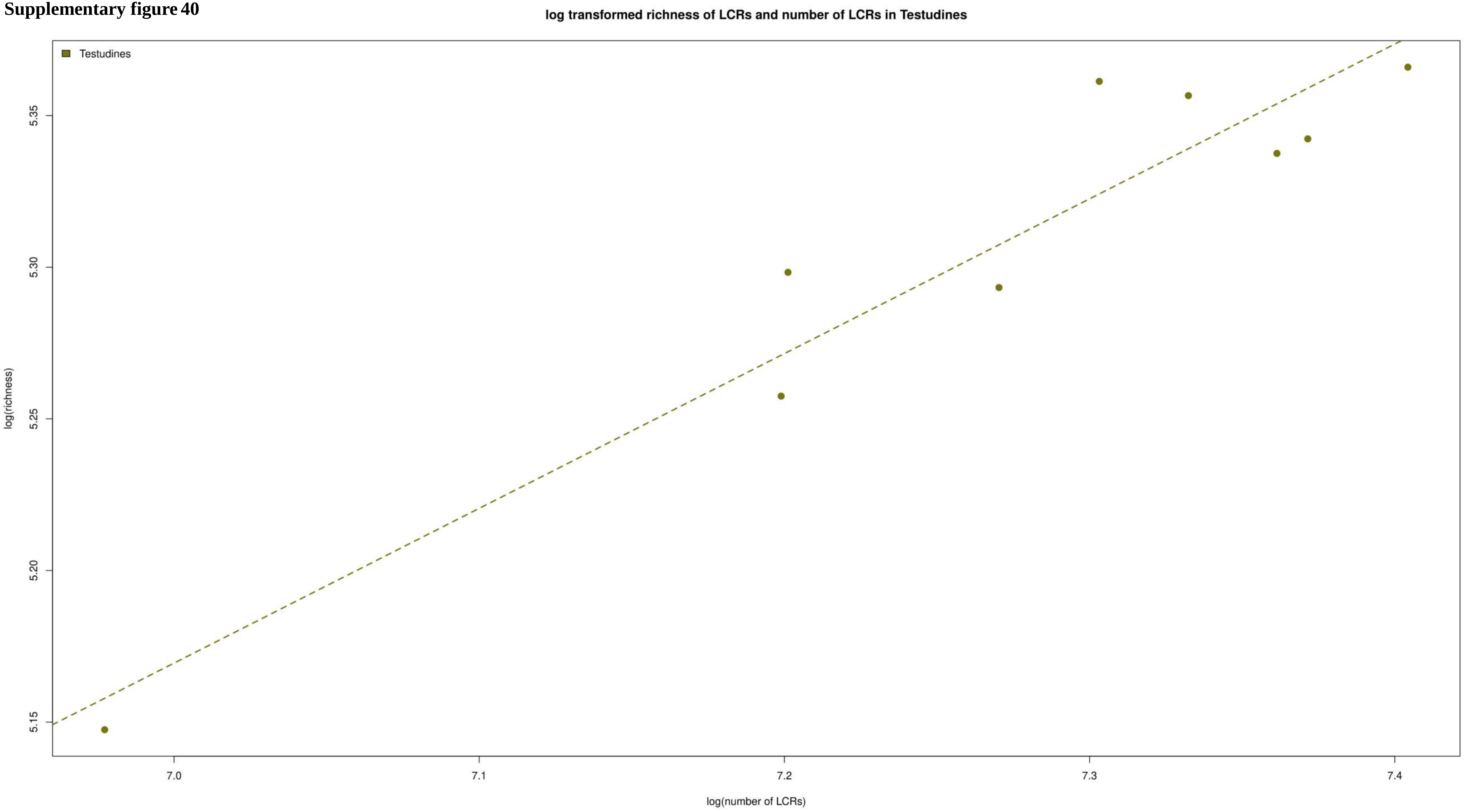

**Supplementary figure 41**

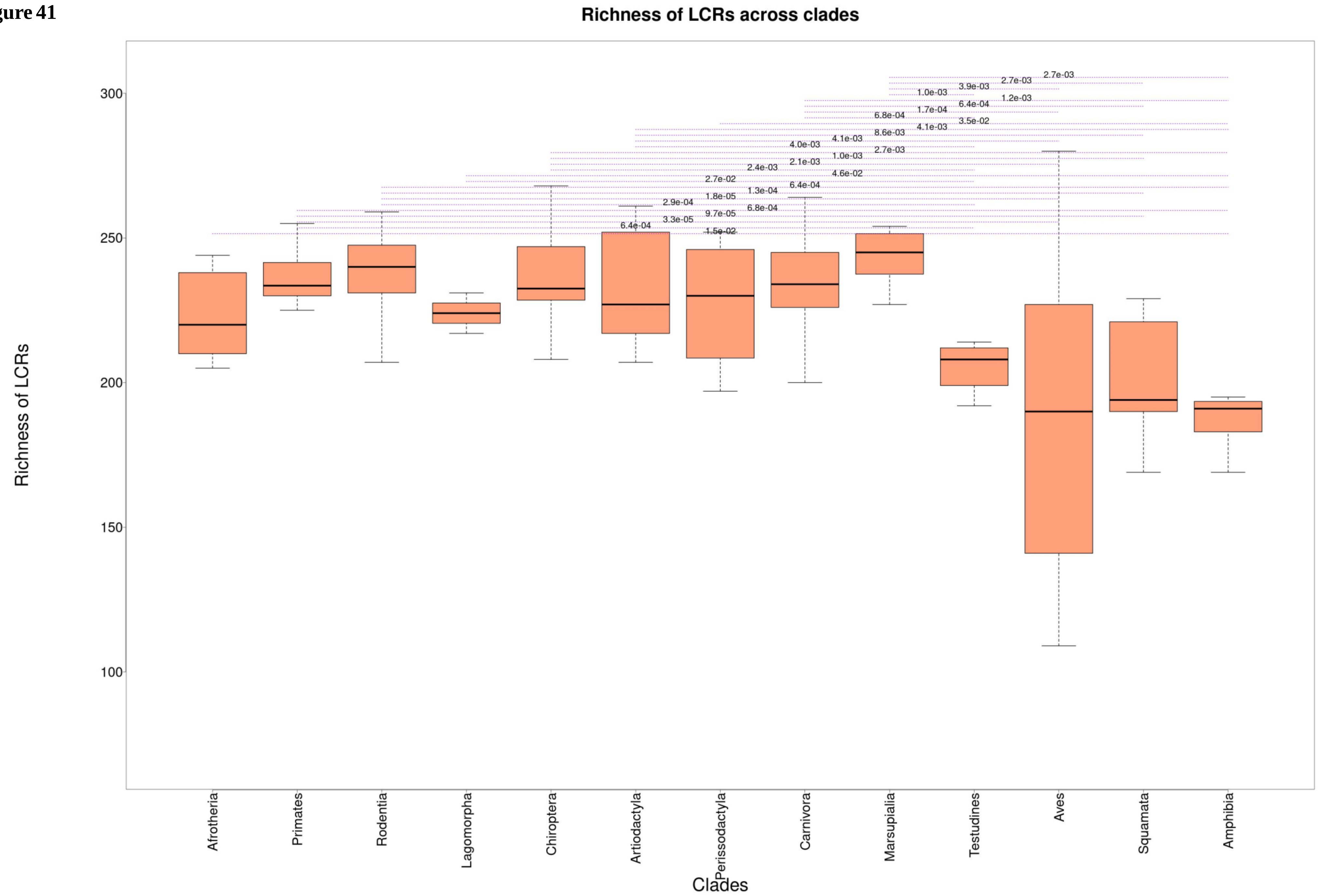

**Supplementary figure 42**

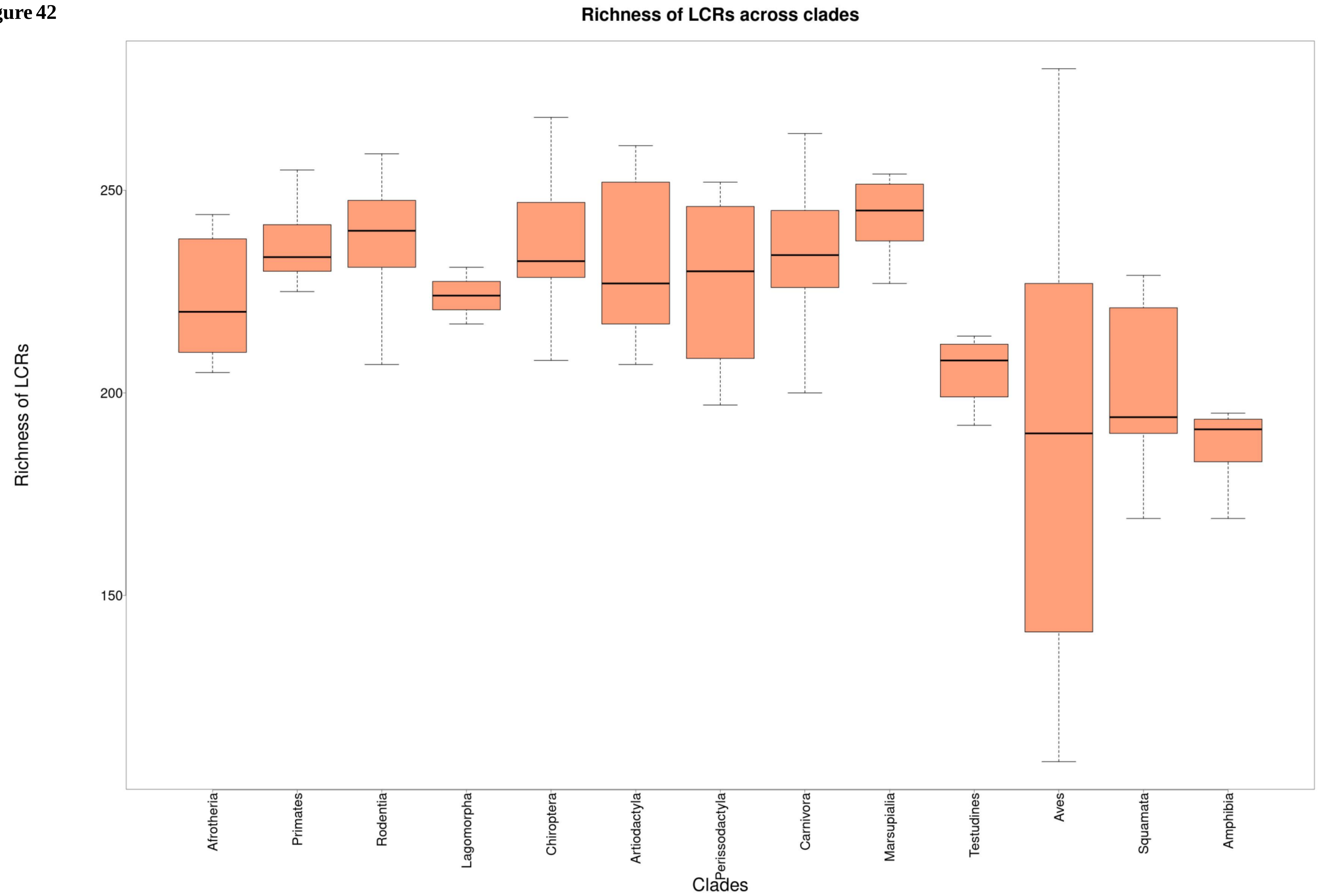

**Supplementary figure 43**

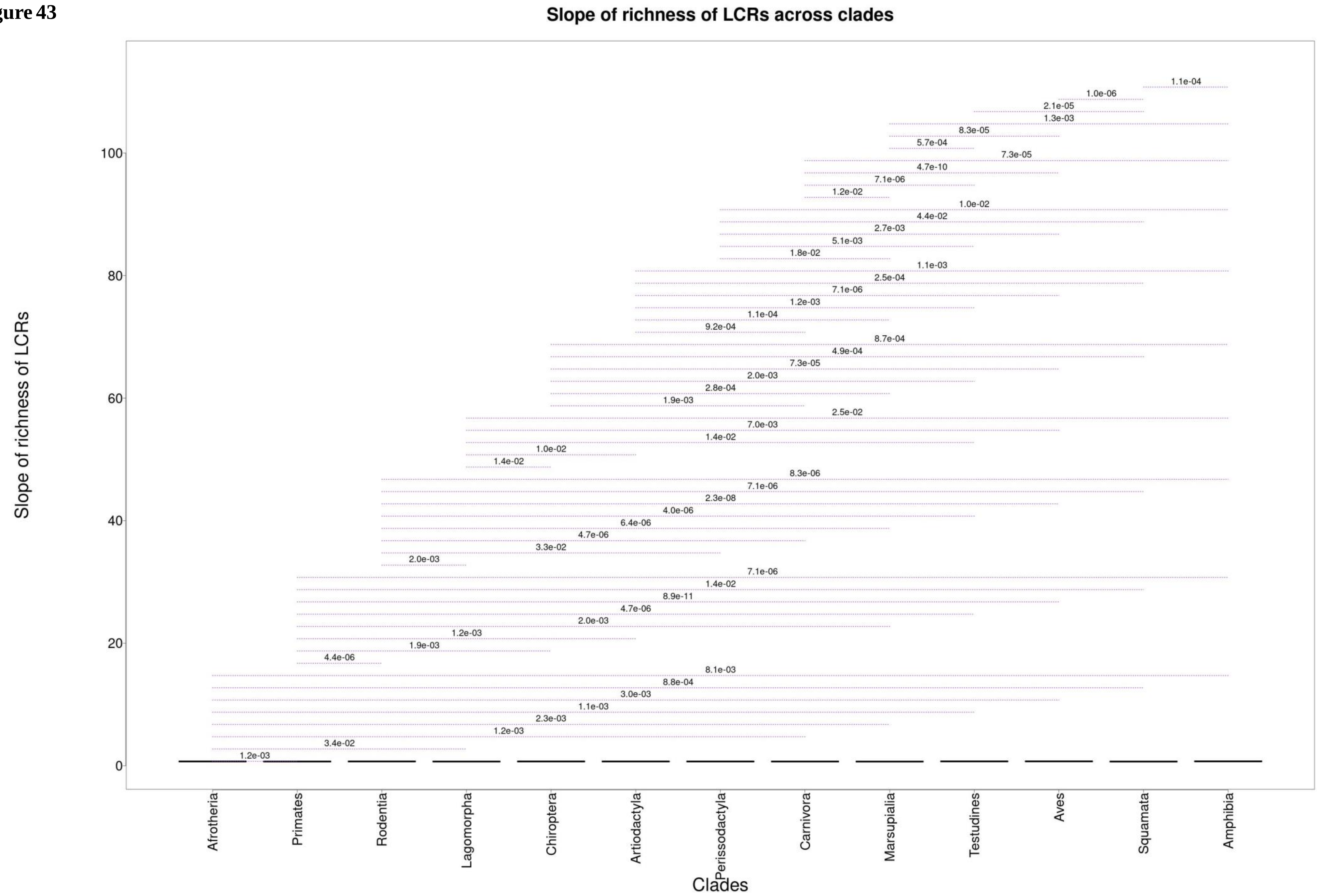

**Supplementary figure 44**

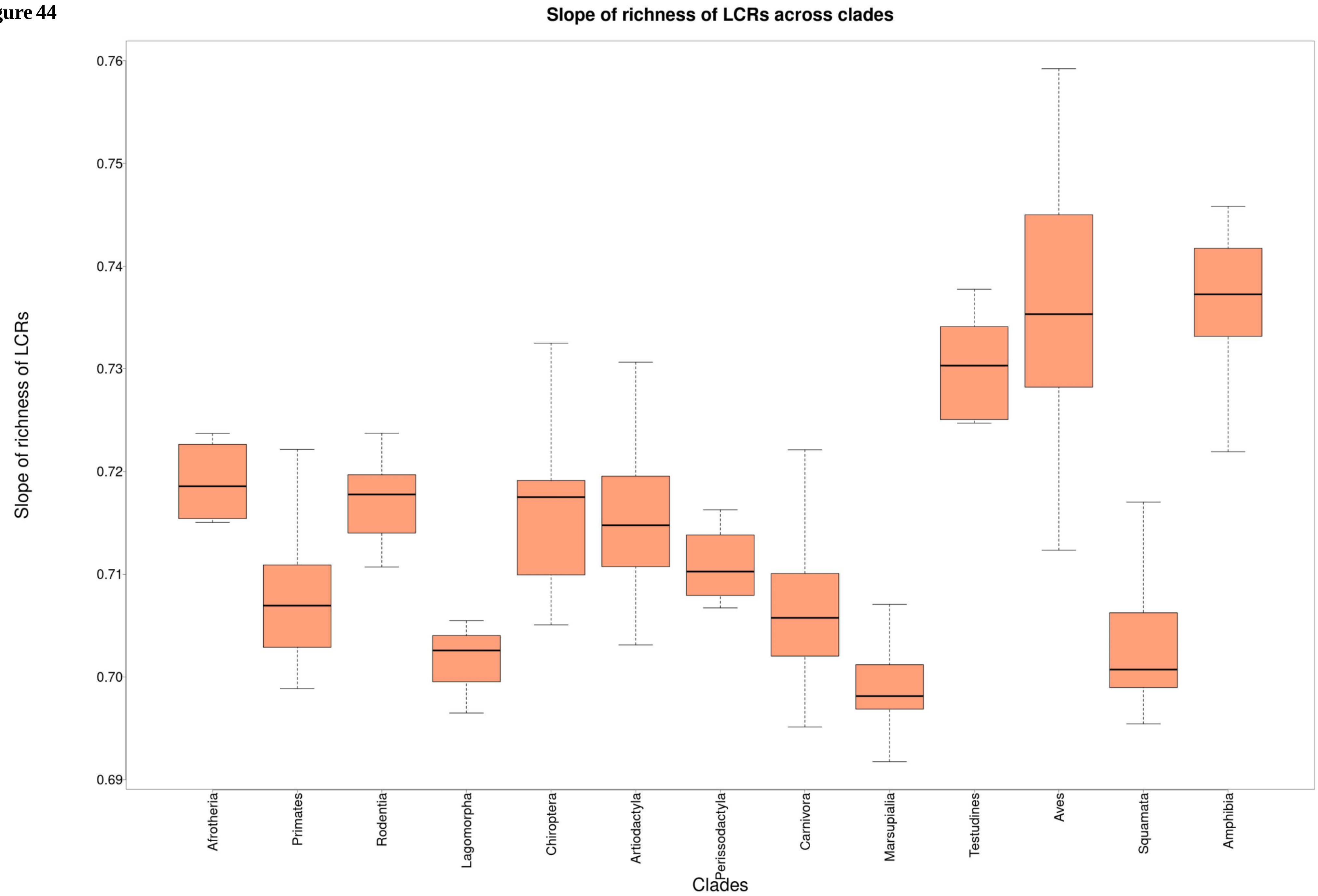

**Supplementary figure 45**

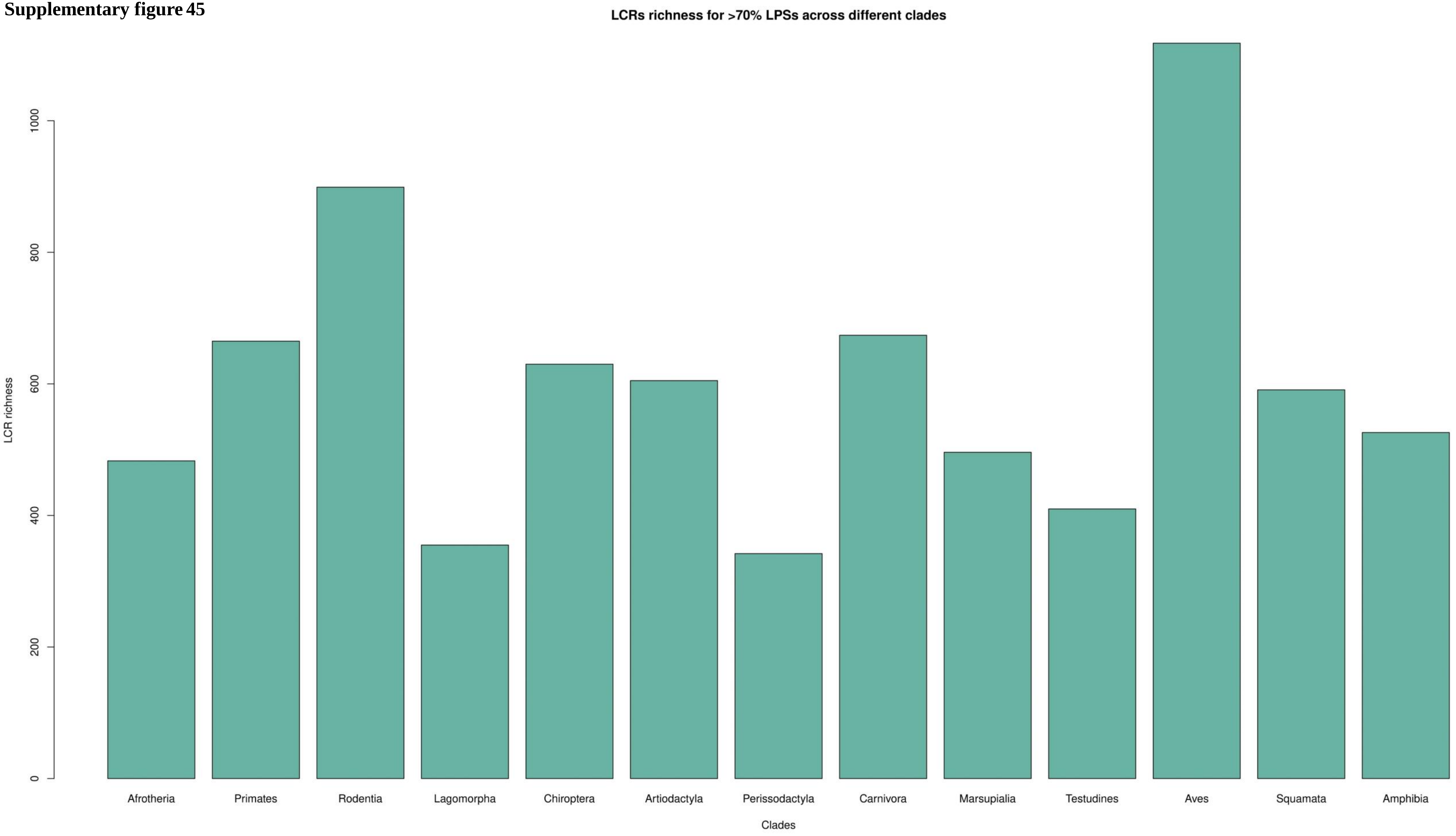

**Supplementary figure 46**

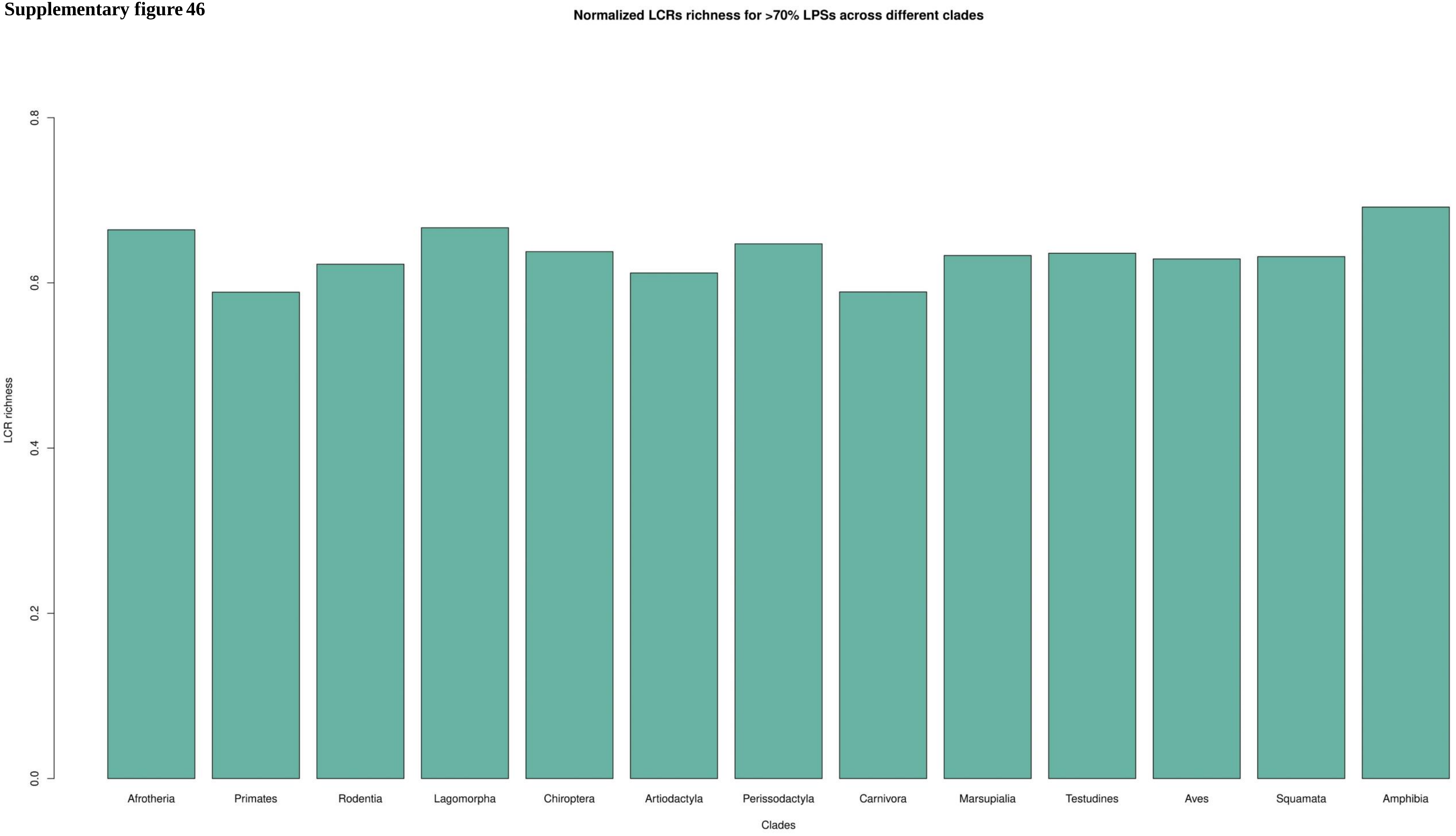

**Supplementary figure 47**

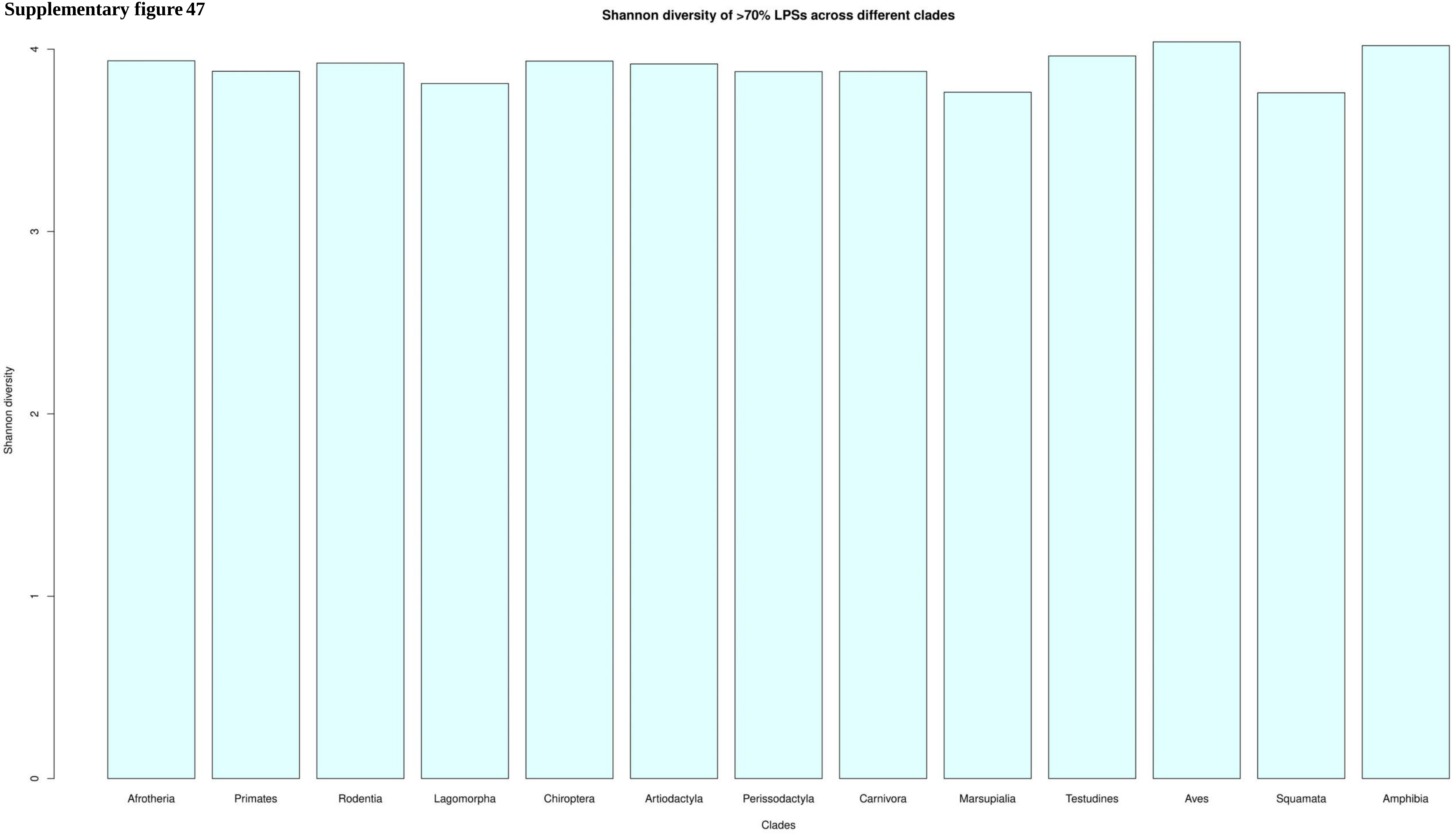

**Supplementary figure 48**

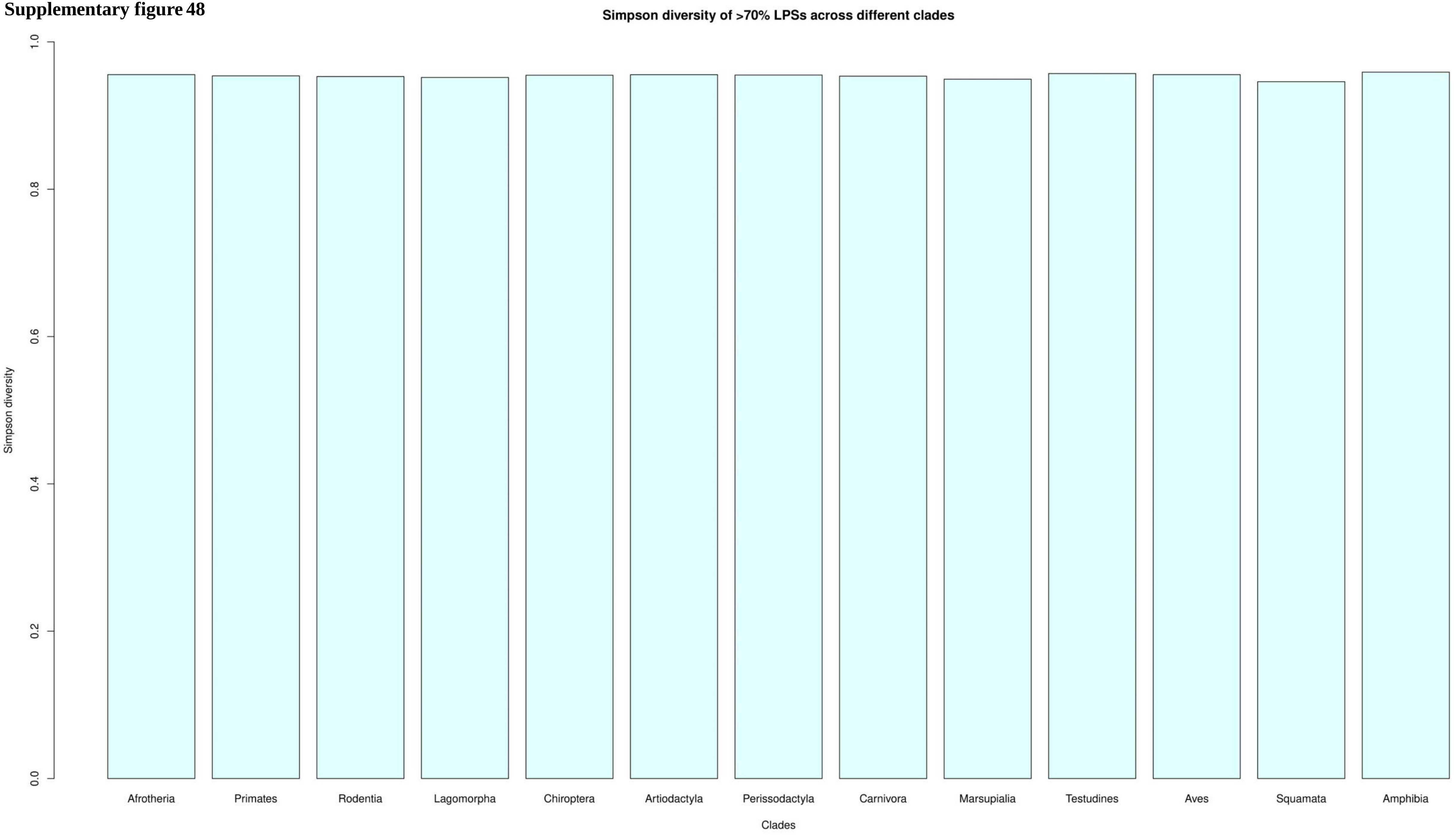

**Supplementary figure 49**

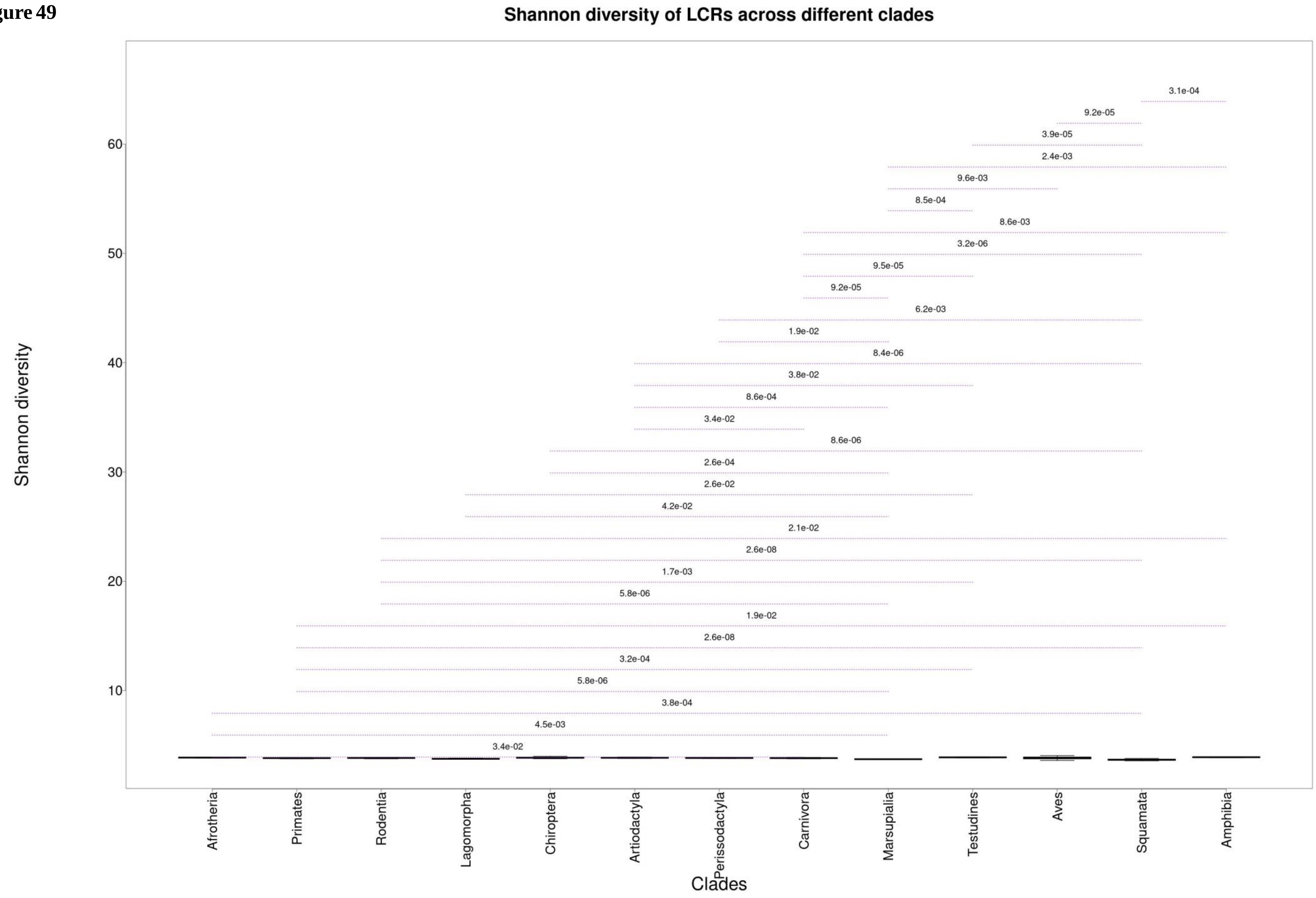

**Supplementary figure 50**

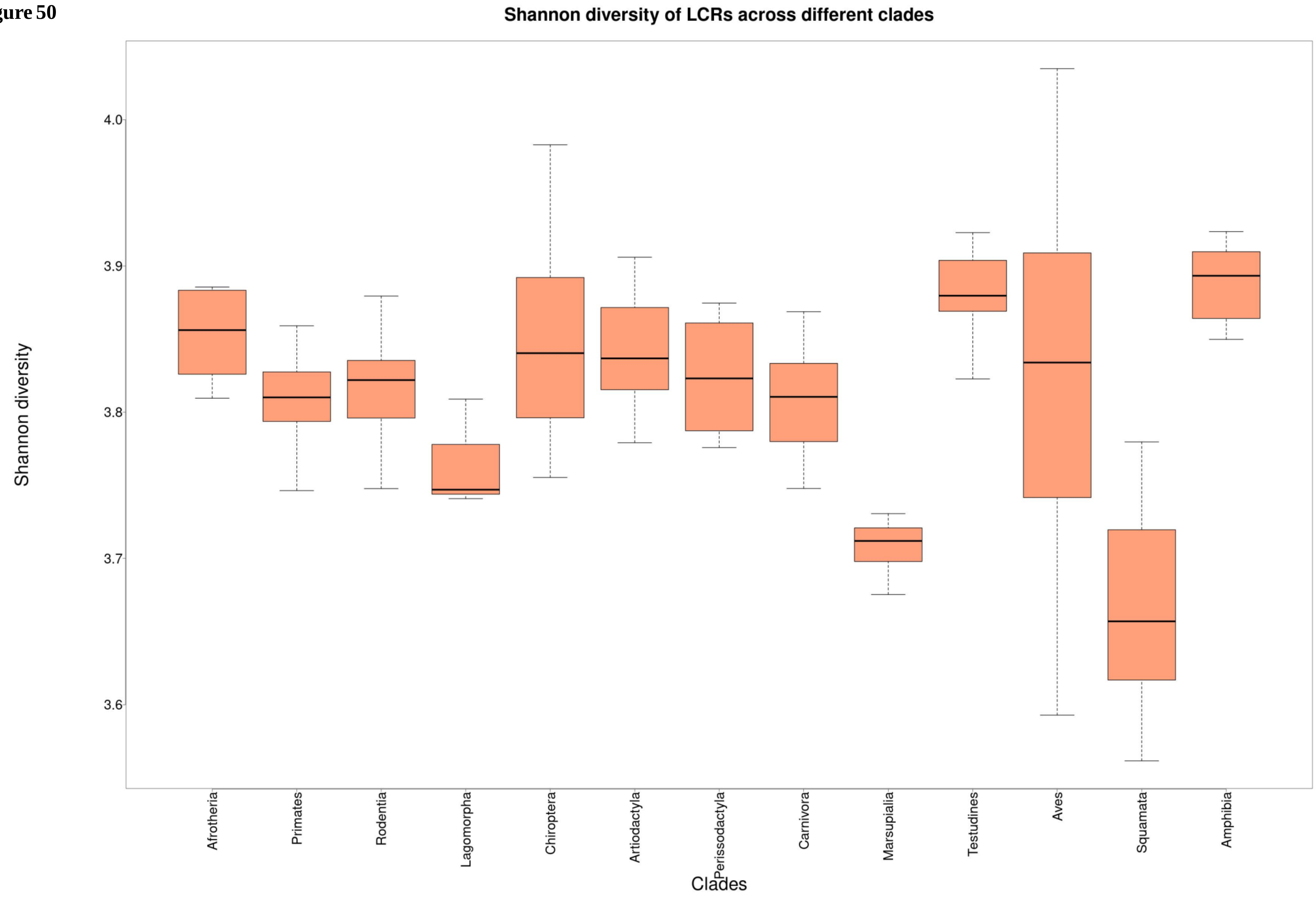

**Supplementary figure 51**

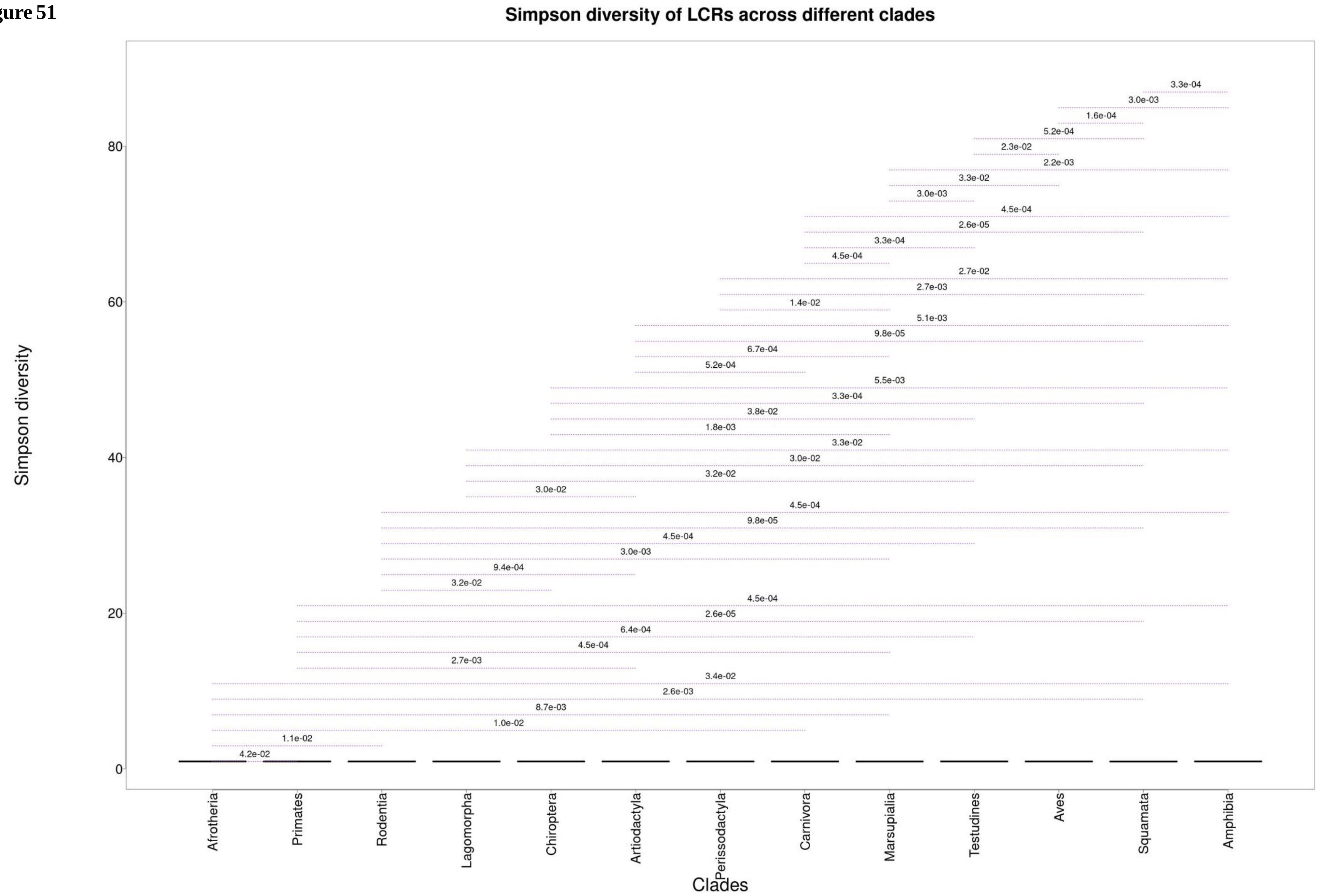

**Supplementary figure 52**

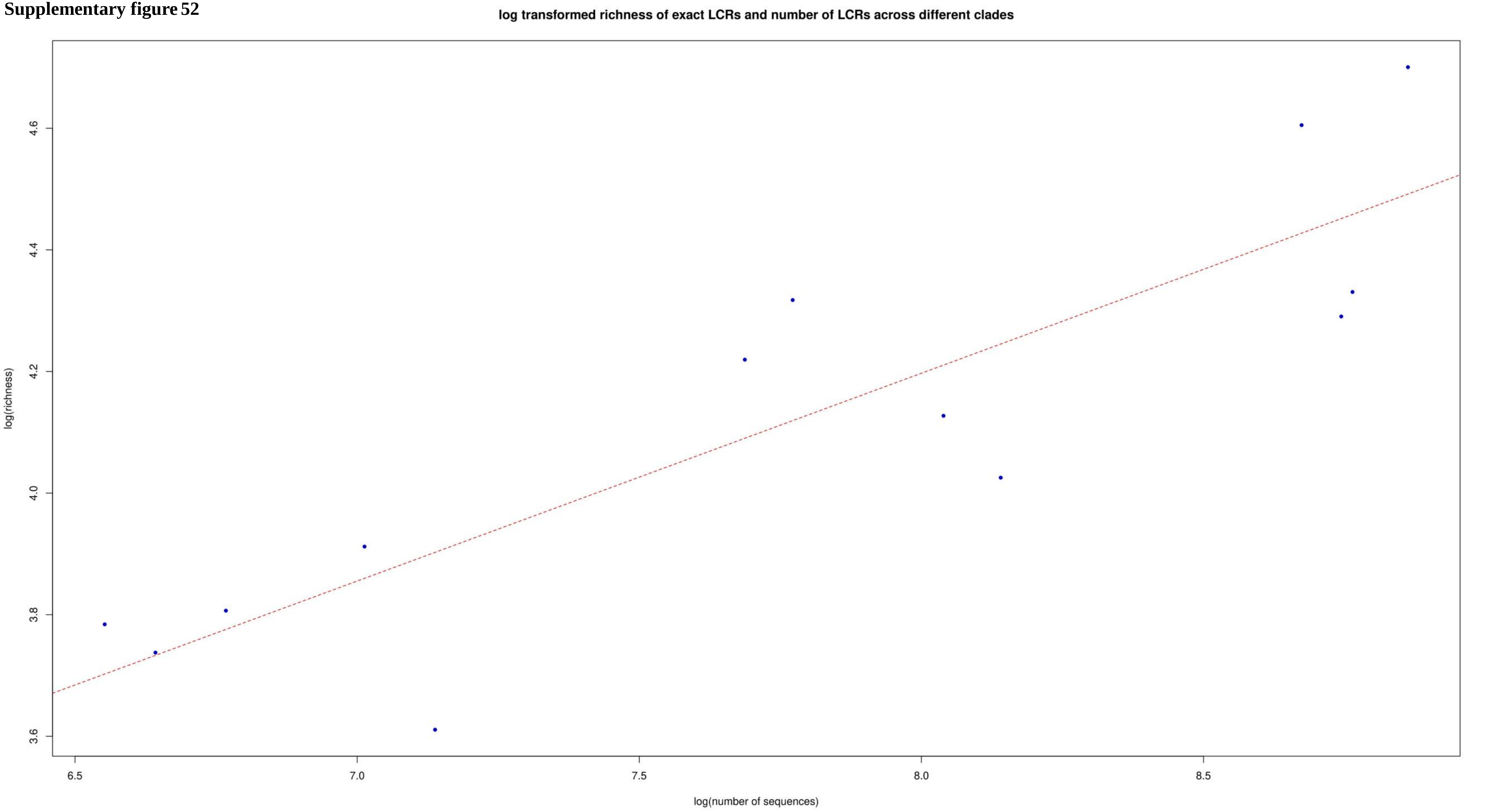

**Supplementary figure 53**

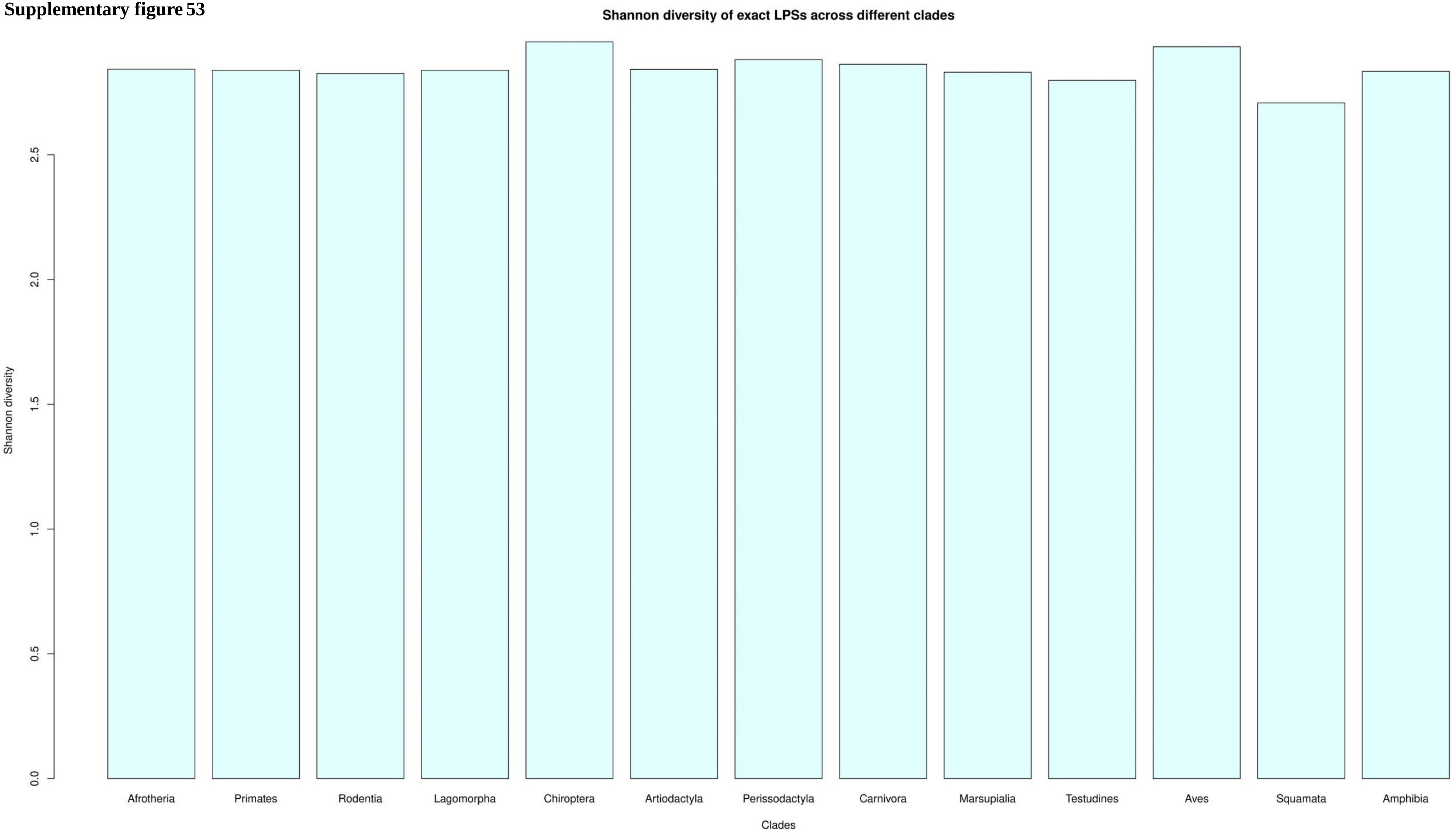

**Supplementary figure 54**

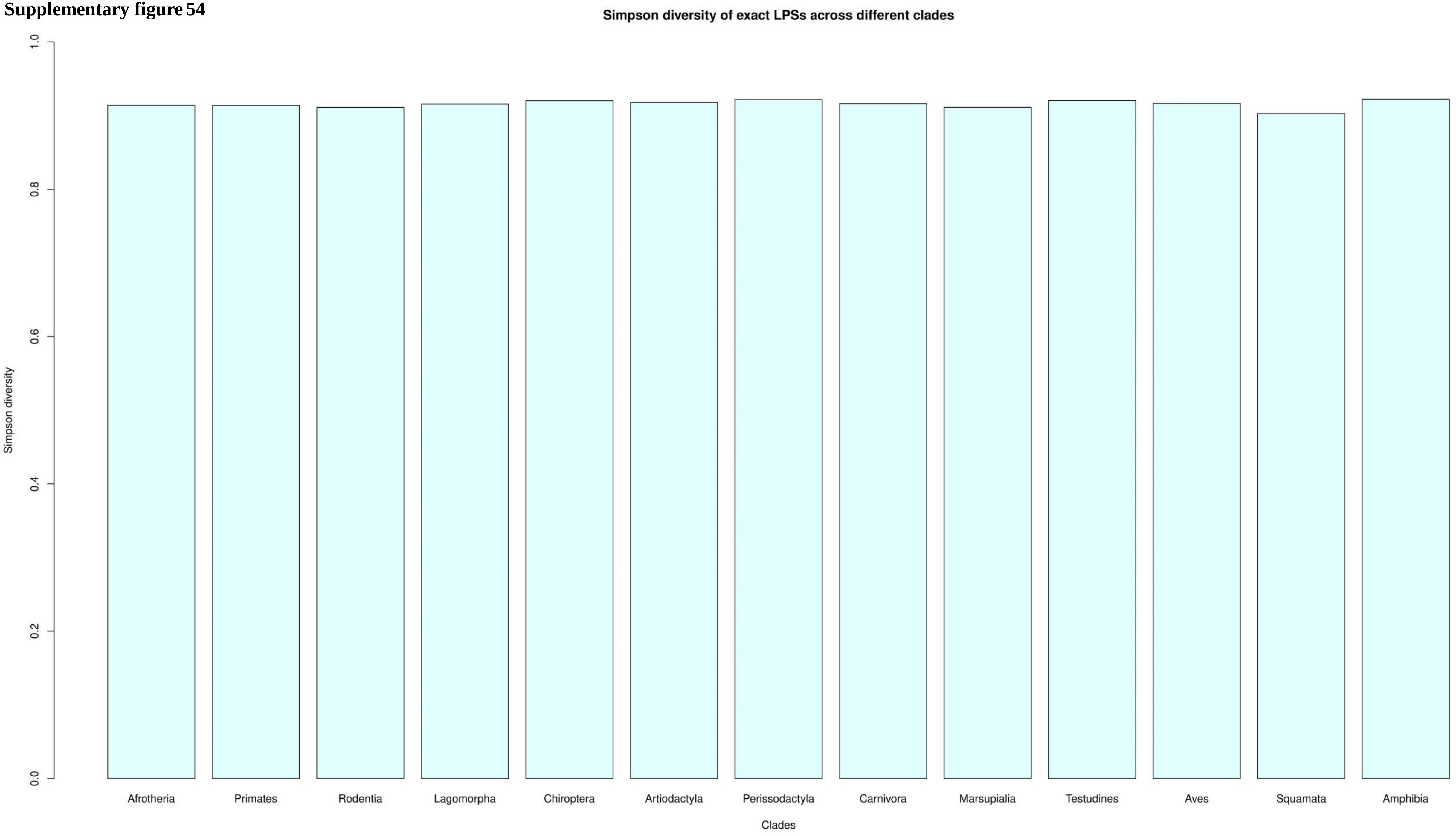

**Supplementary figure 55**

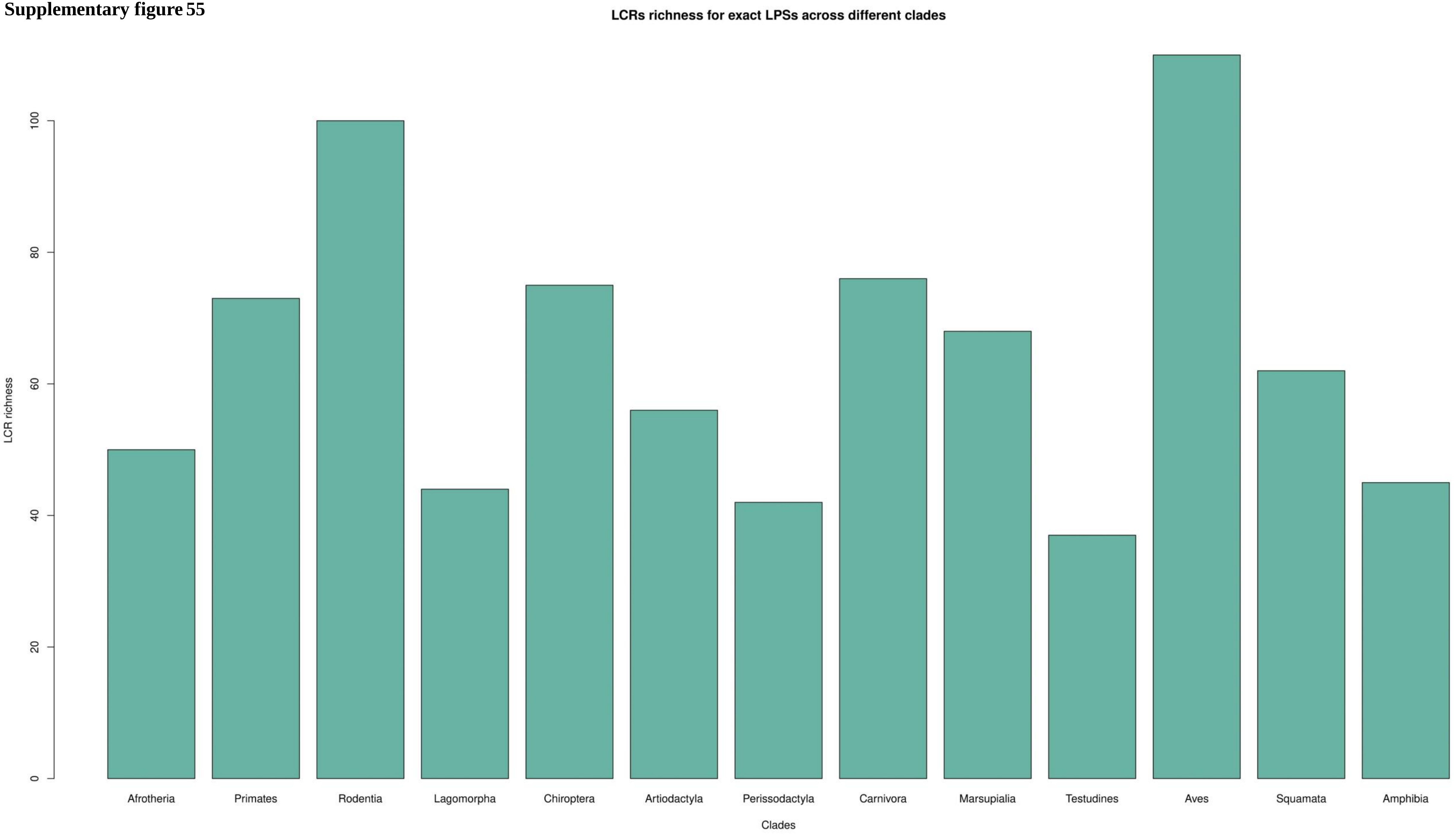

**Supplementary figure 56**

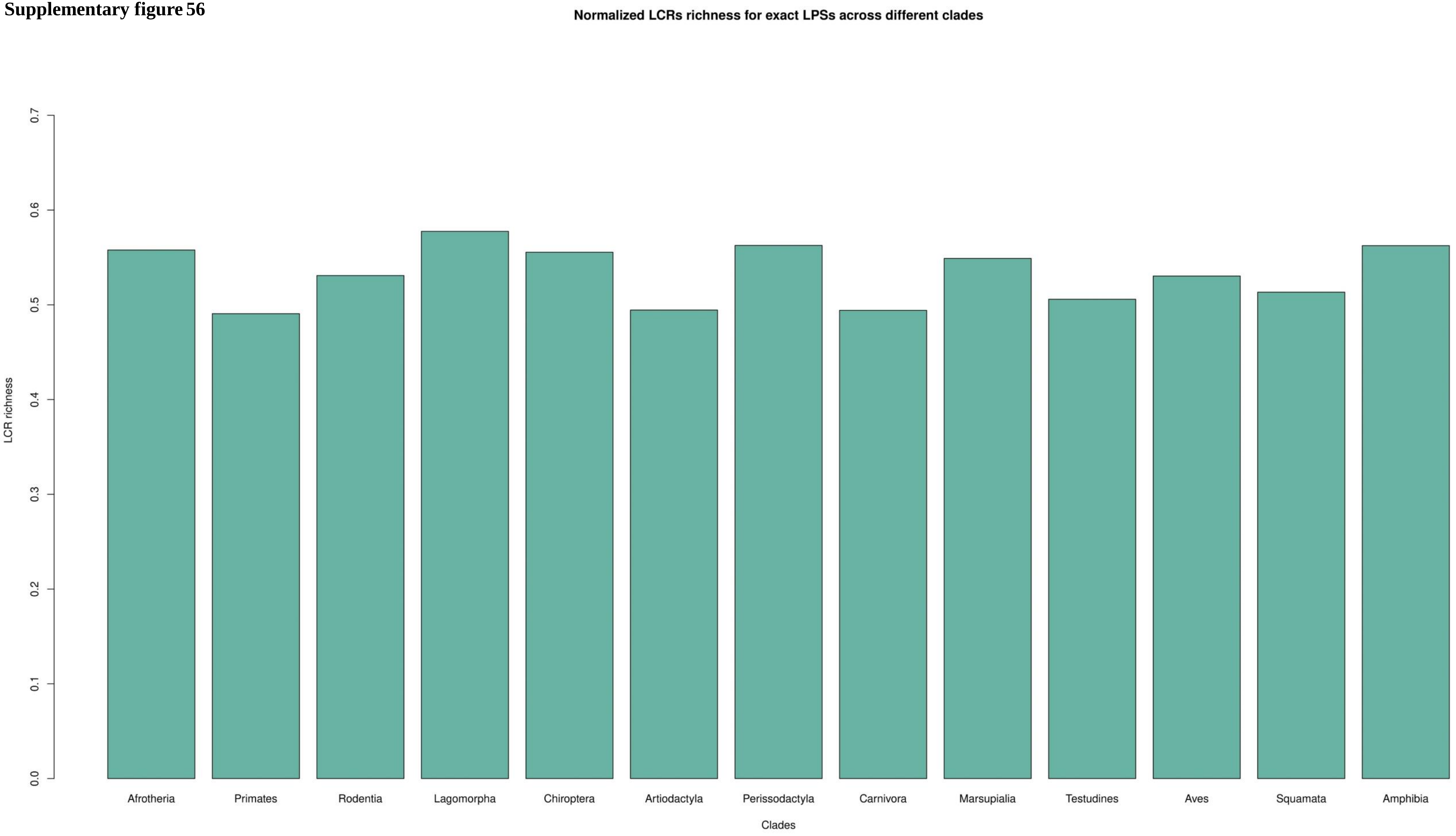

**Supplementary figure 57**

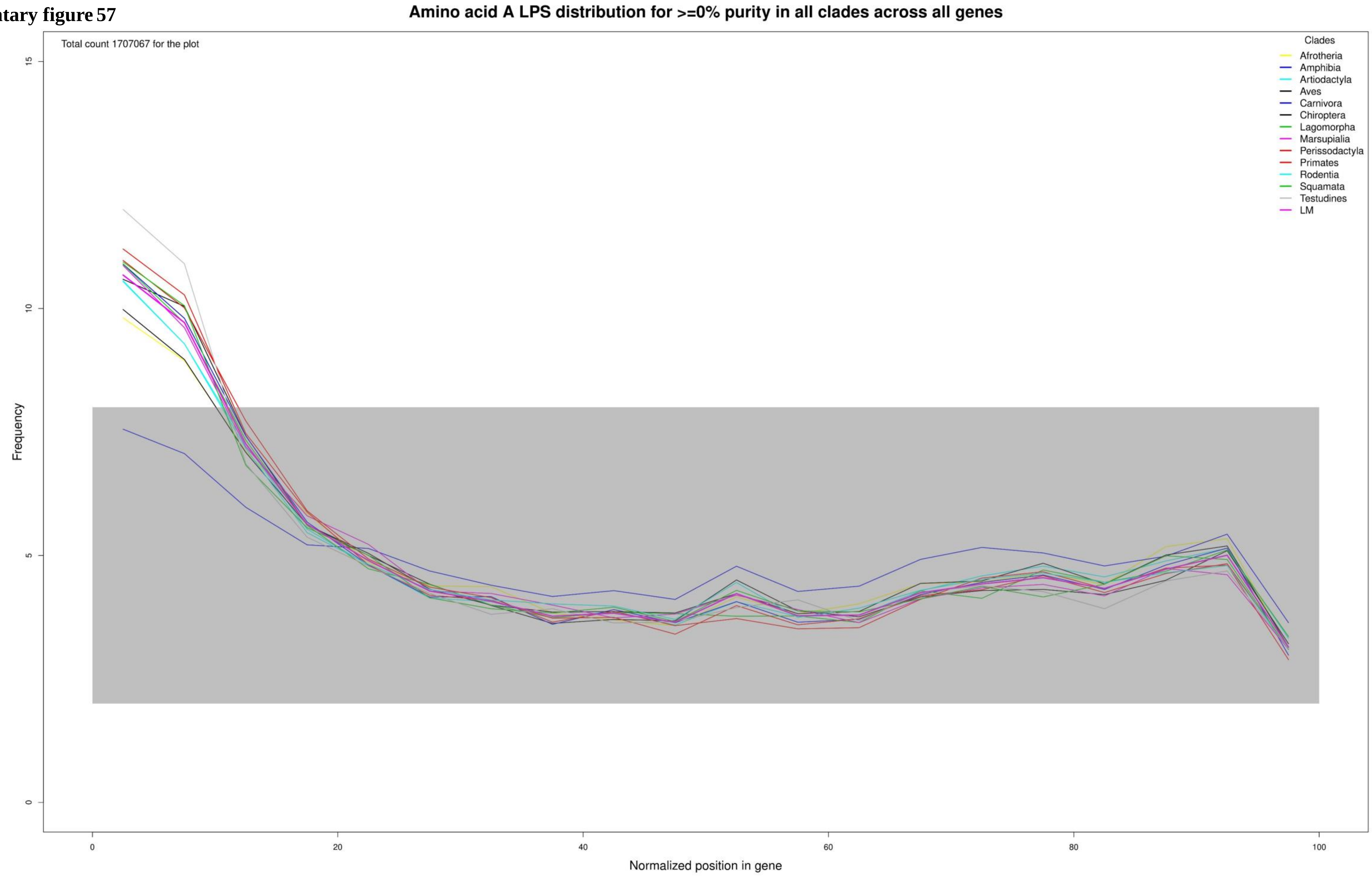

**Supplementary figure 58**

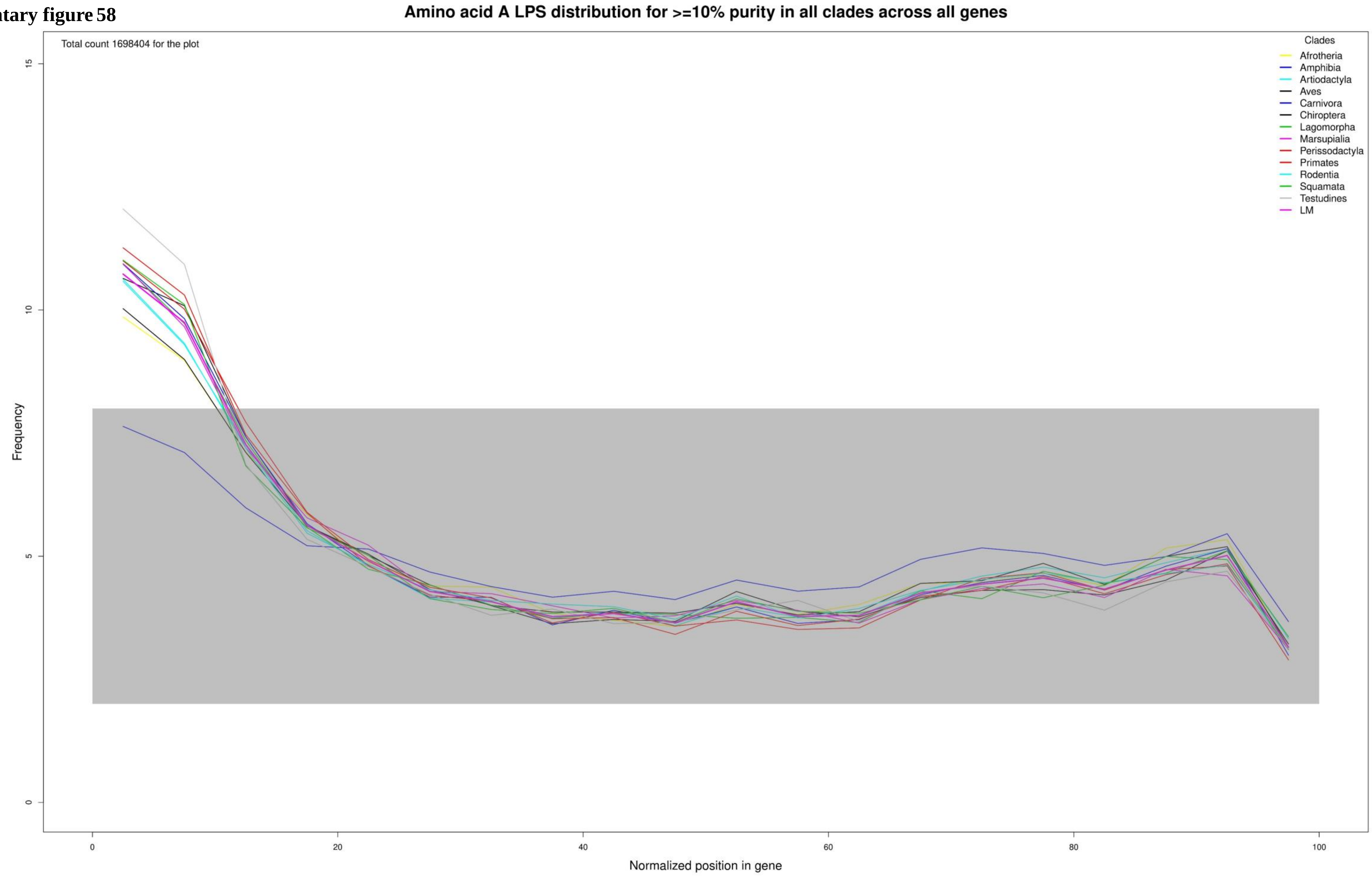

Supplementary figure 59

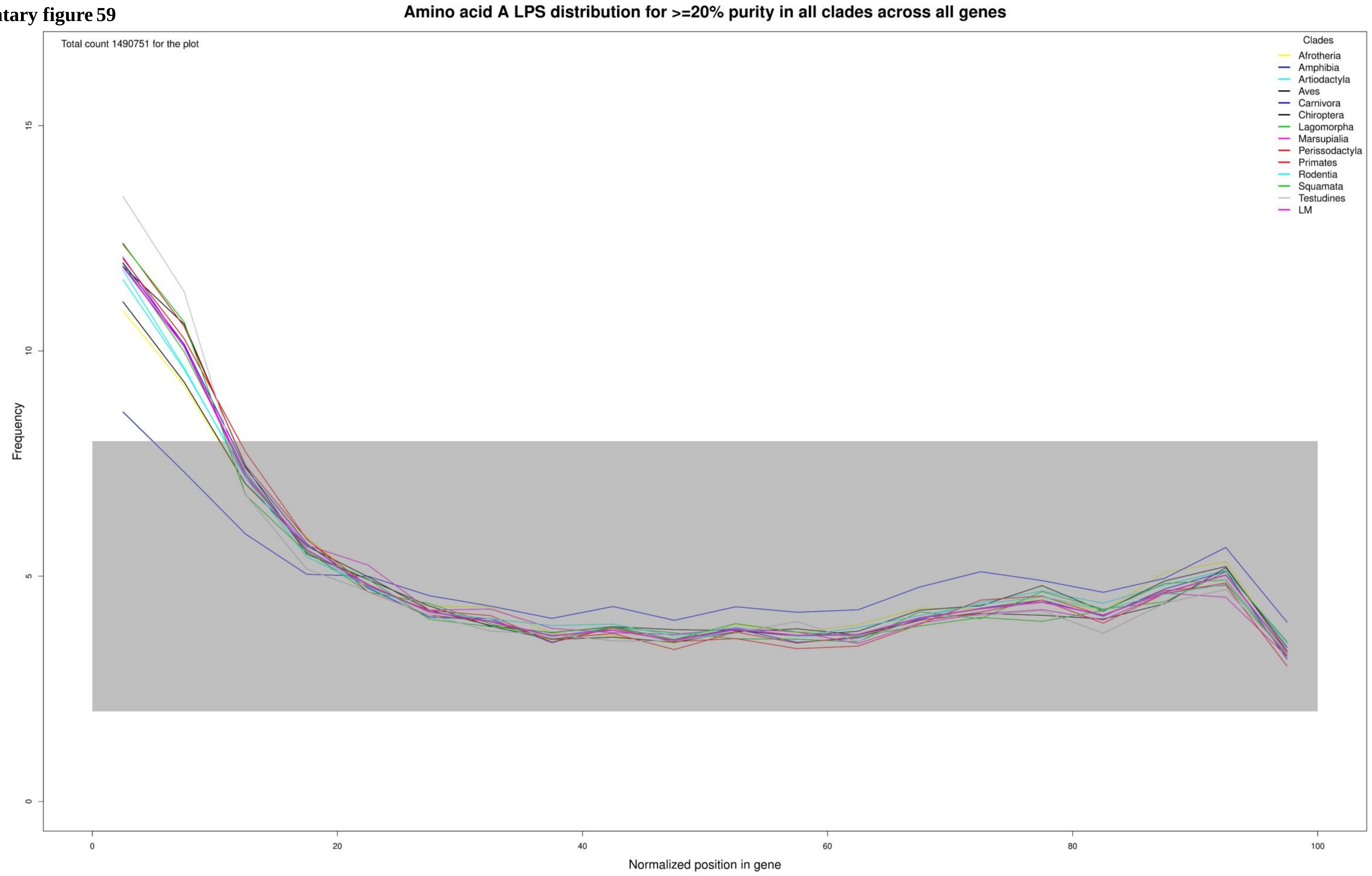

**Supplementary figure 60**

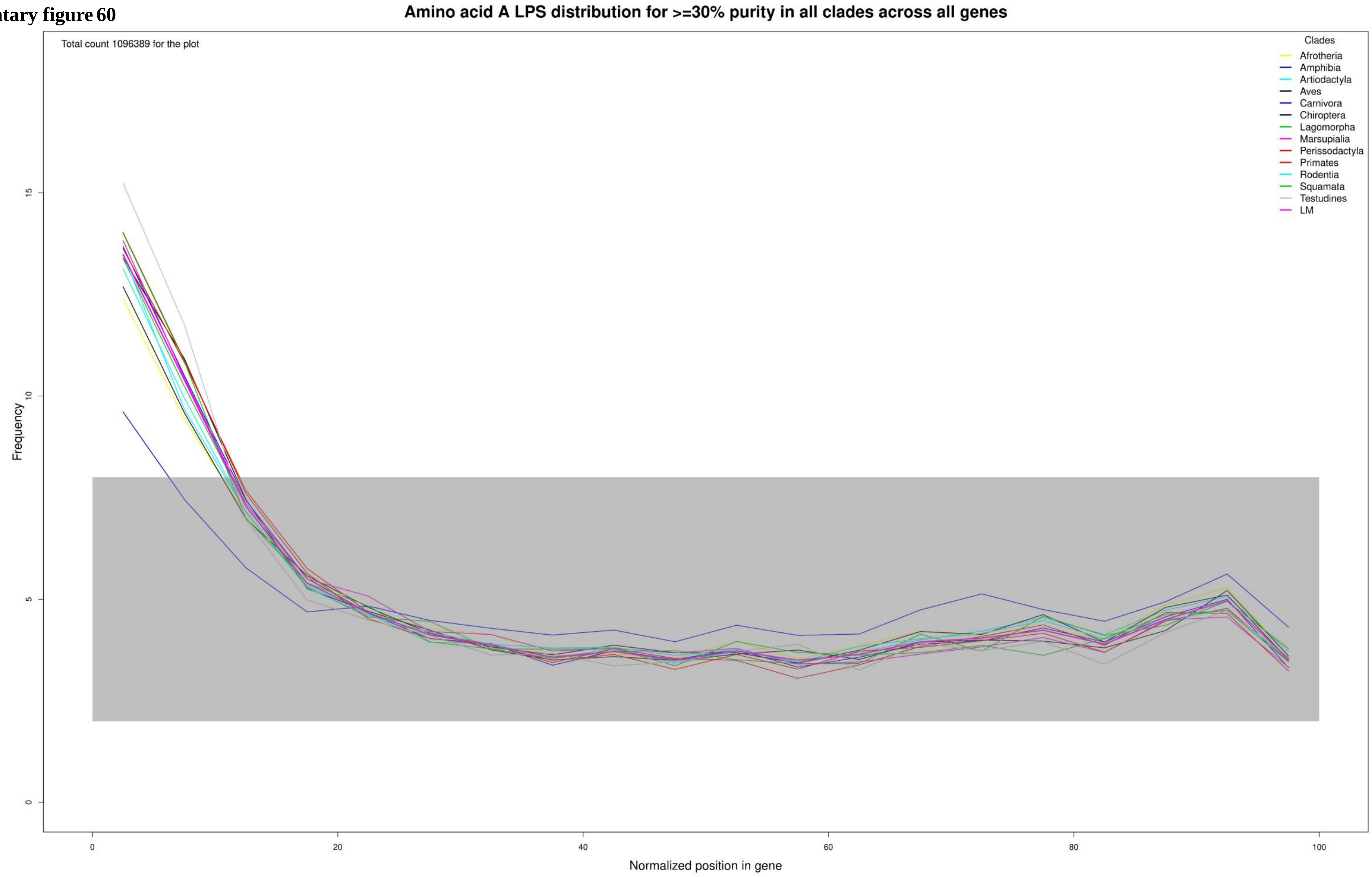

Supplementary figure 61

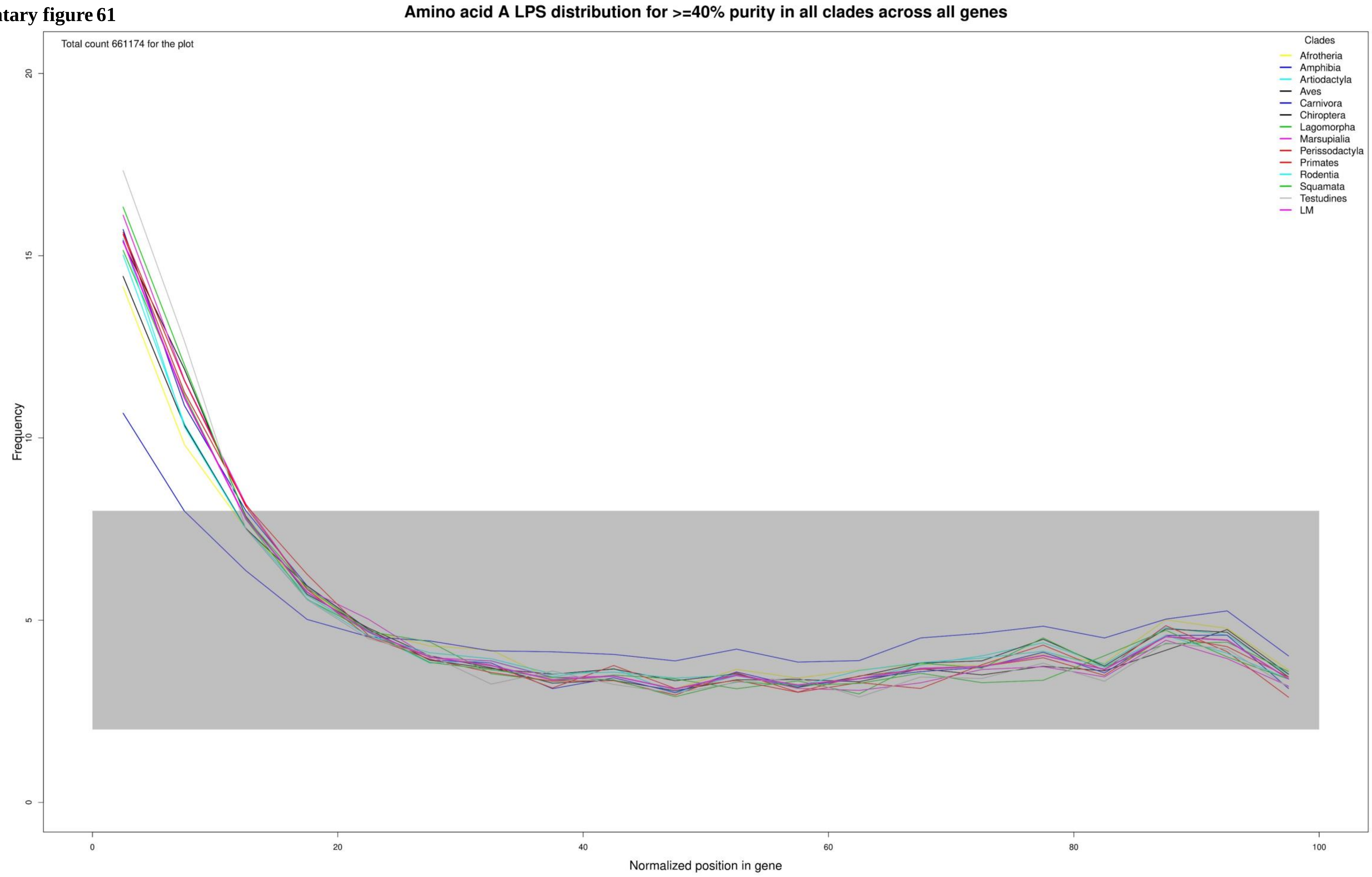

**Supplementary figure 62**

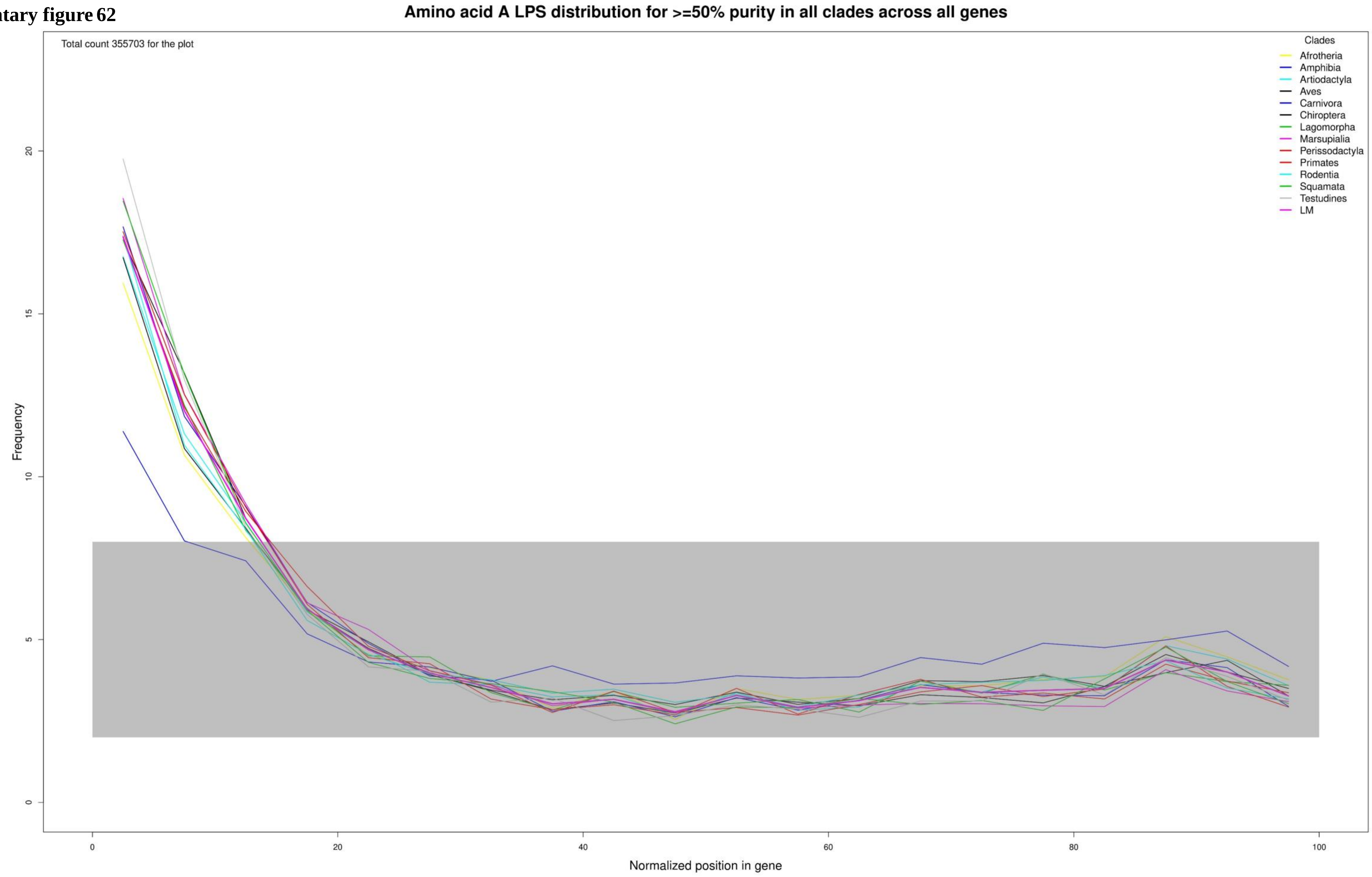

**Supplementary figure 63**

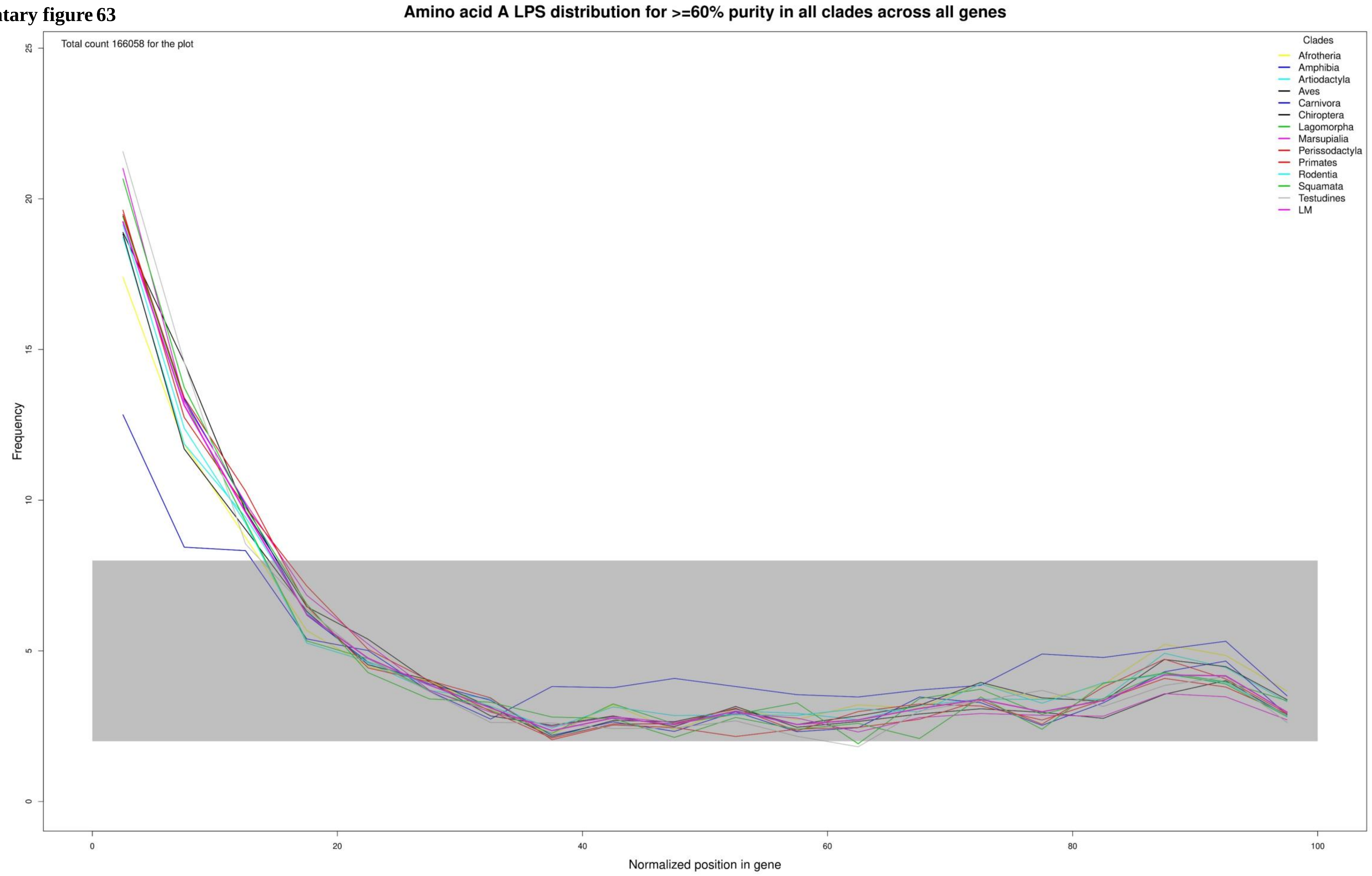

**Supplementary figure 64**

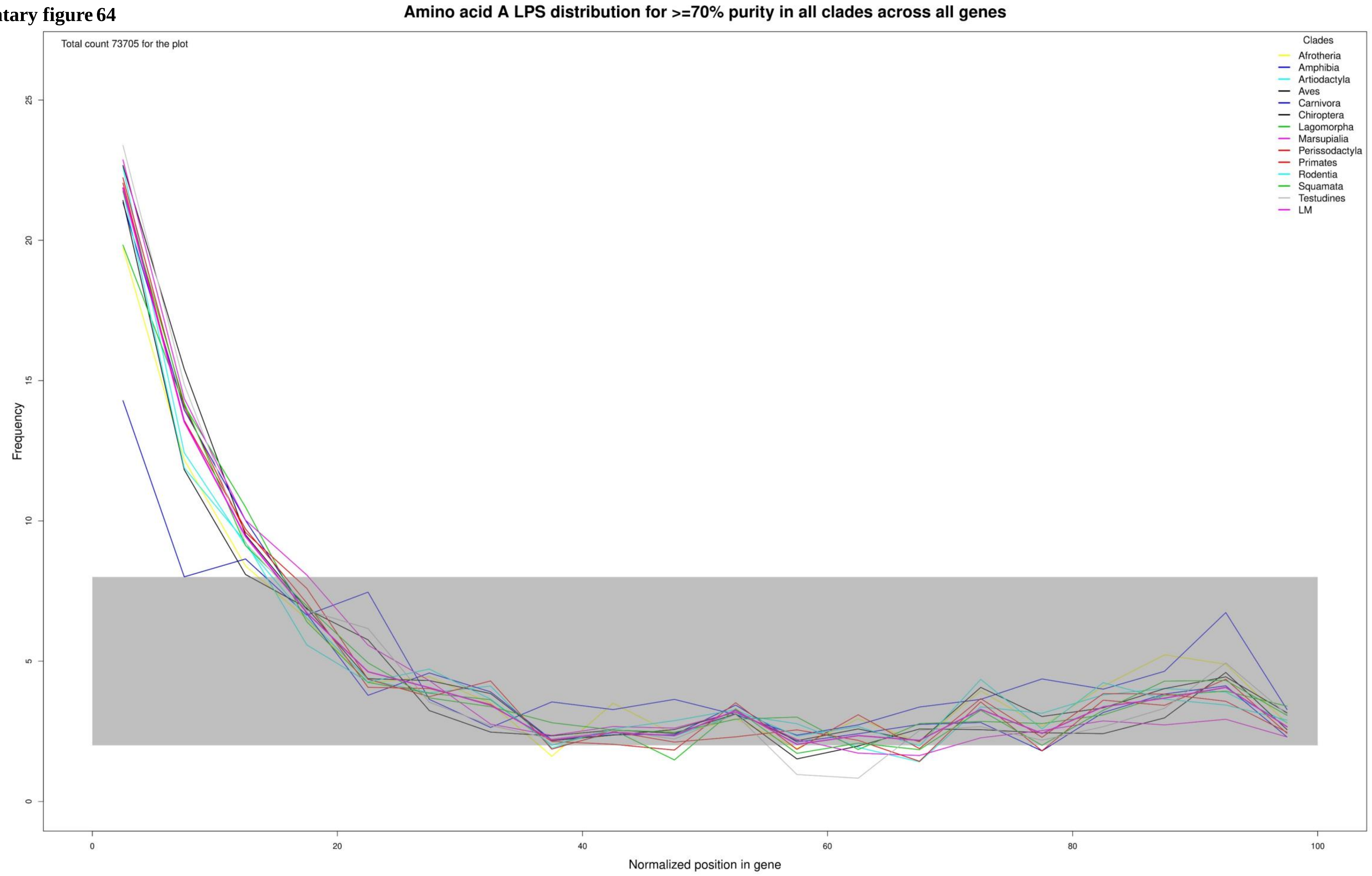

Supplementary figure 65

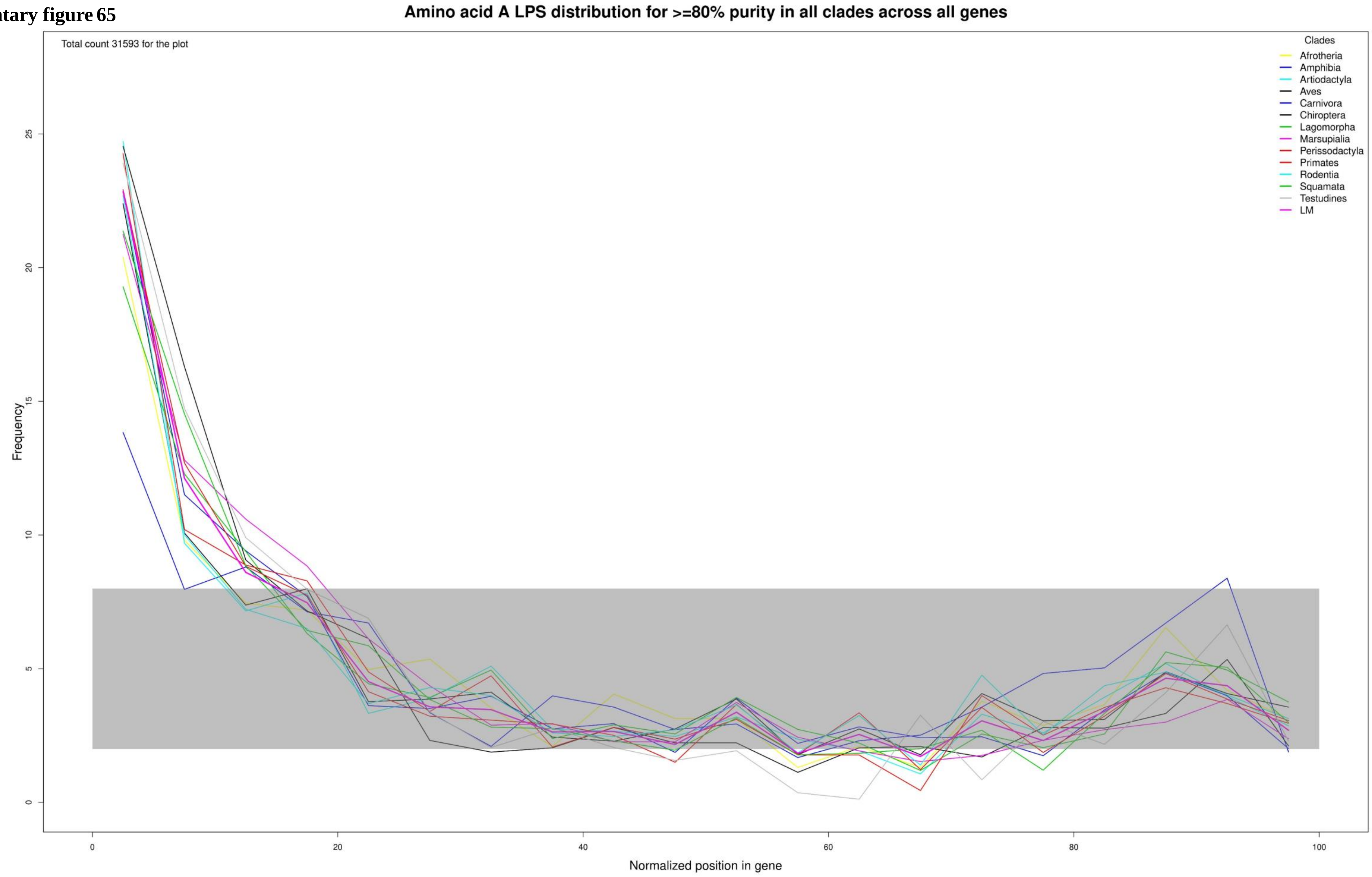

**Supplementary figure 66**

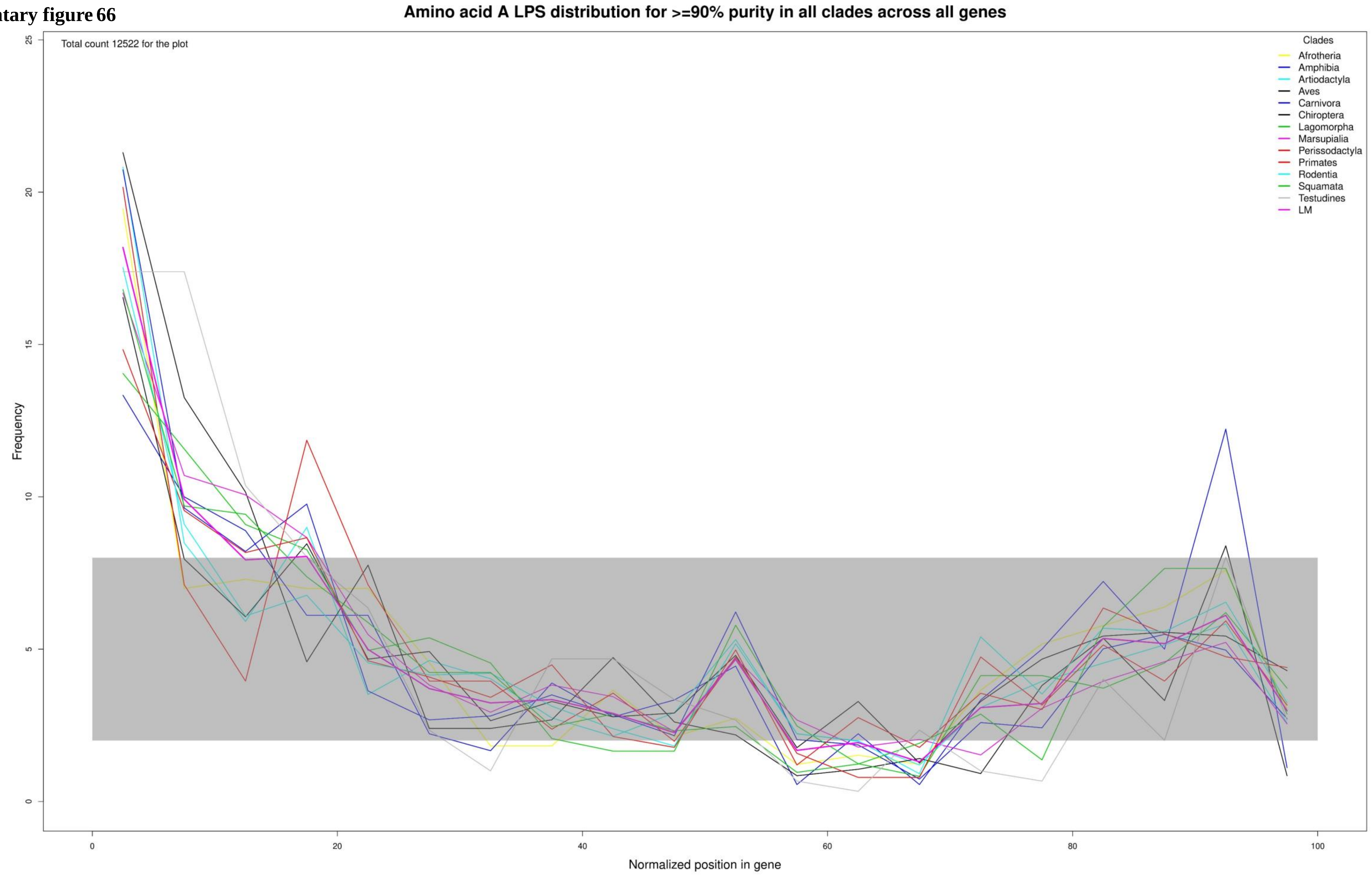

Supplementary figure 67

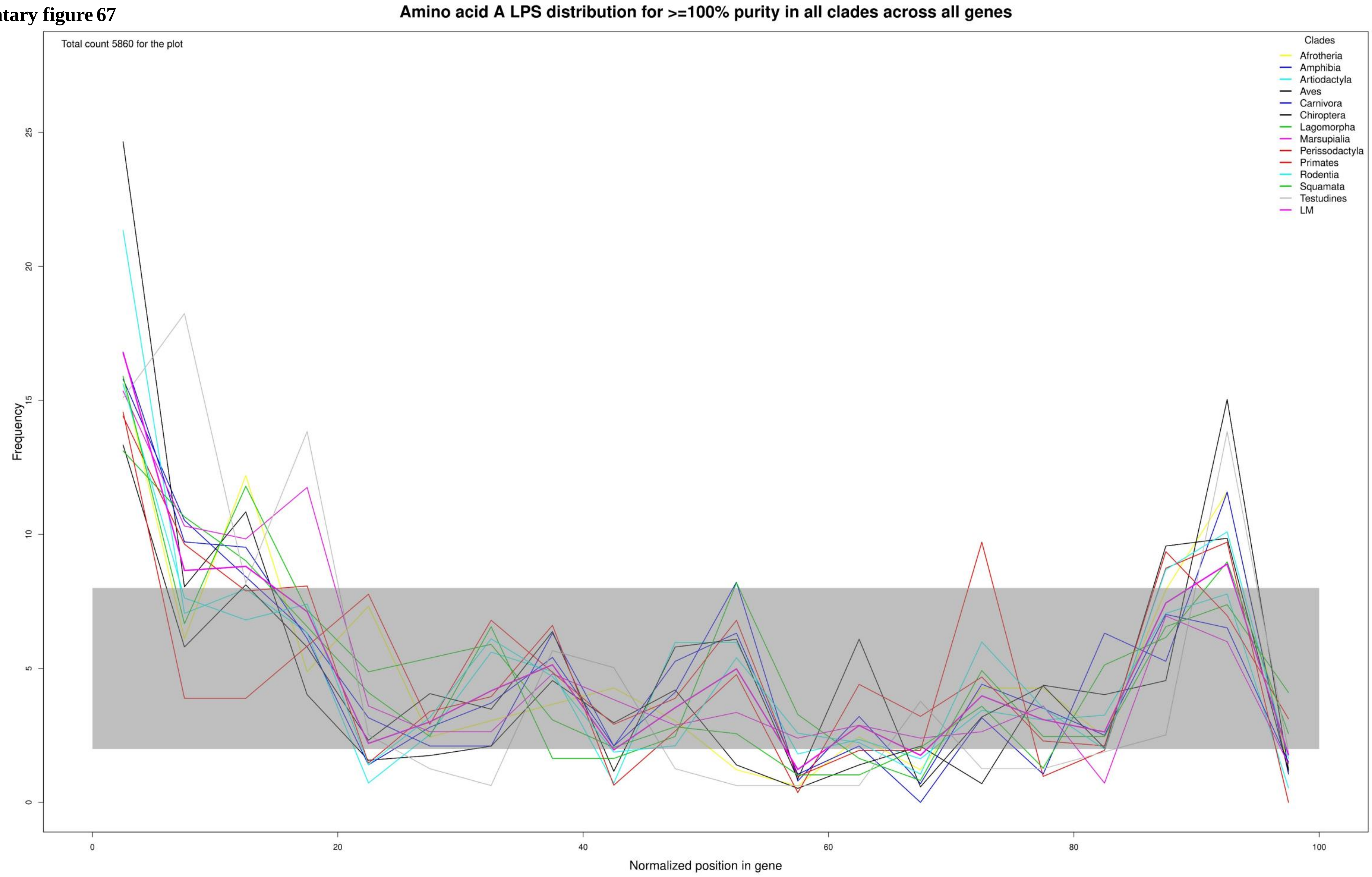

**Supplementary figure 68**

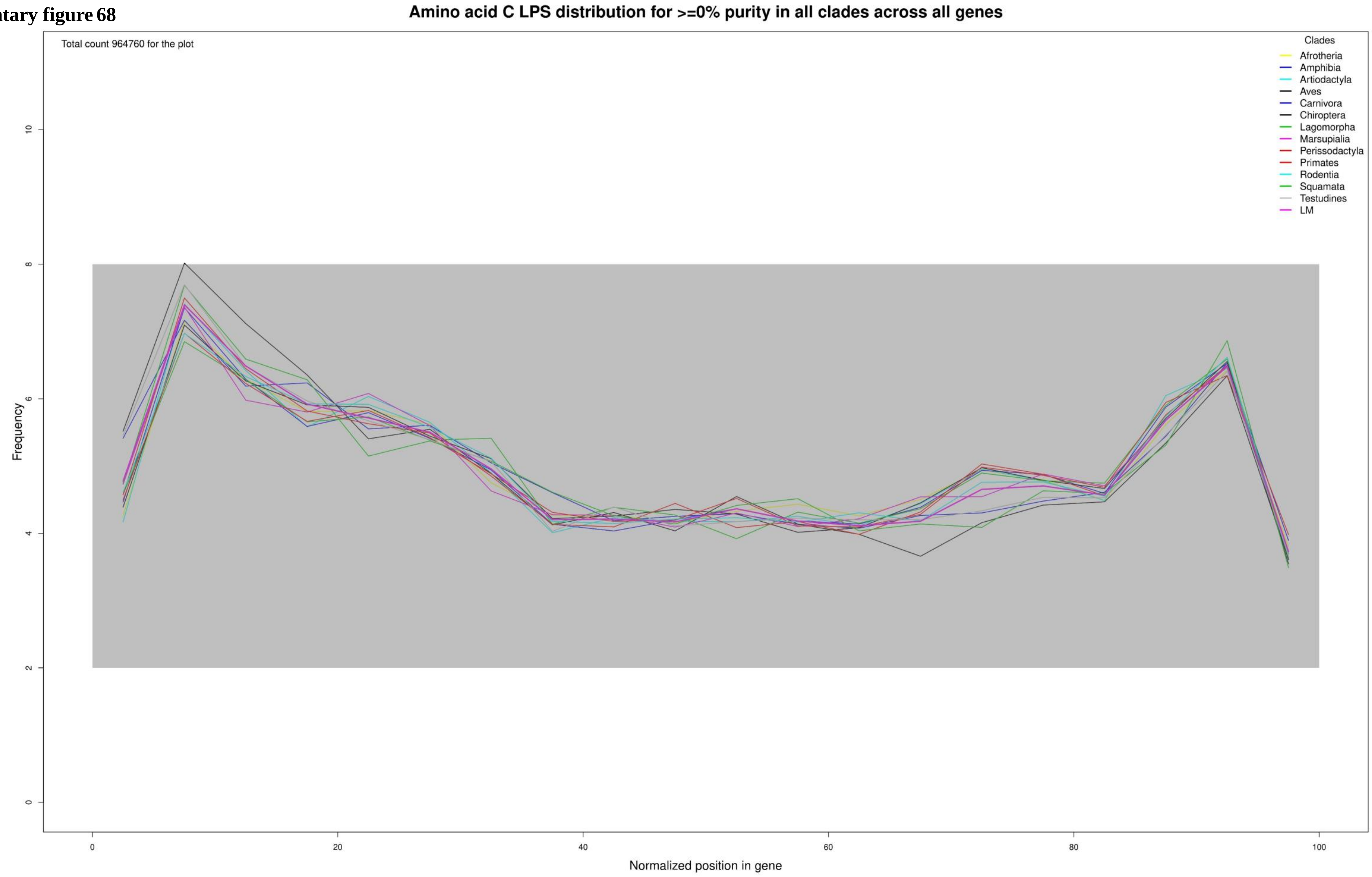

**Supplementary figure 69**

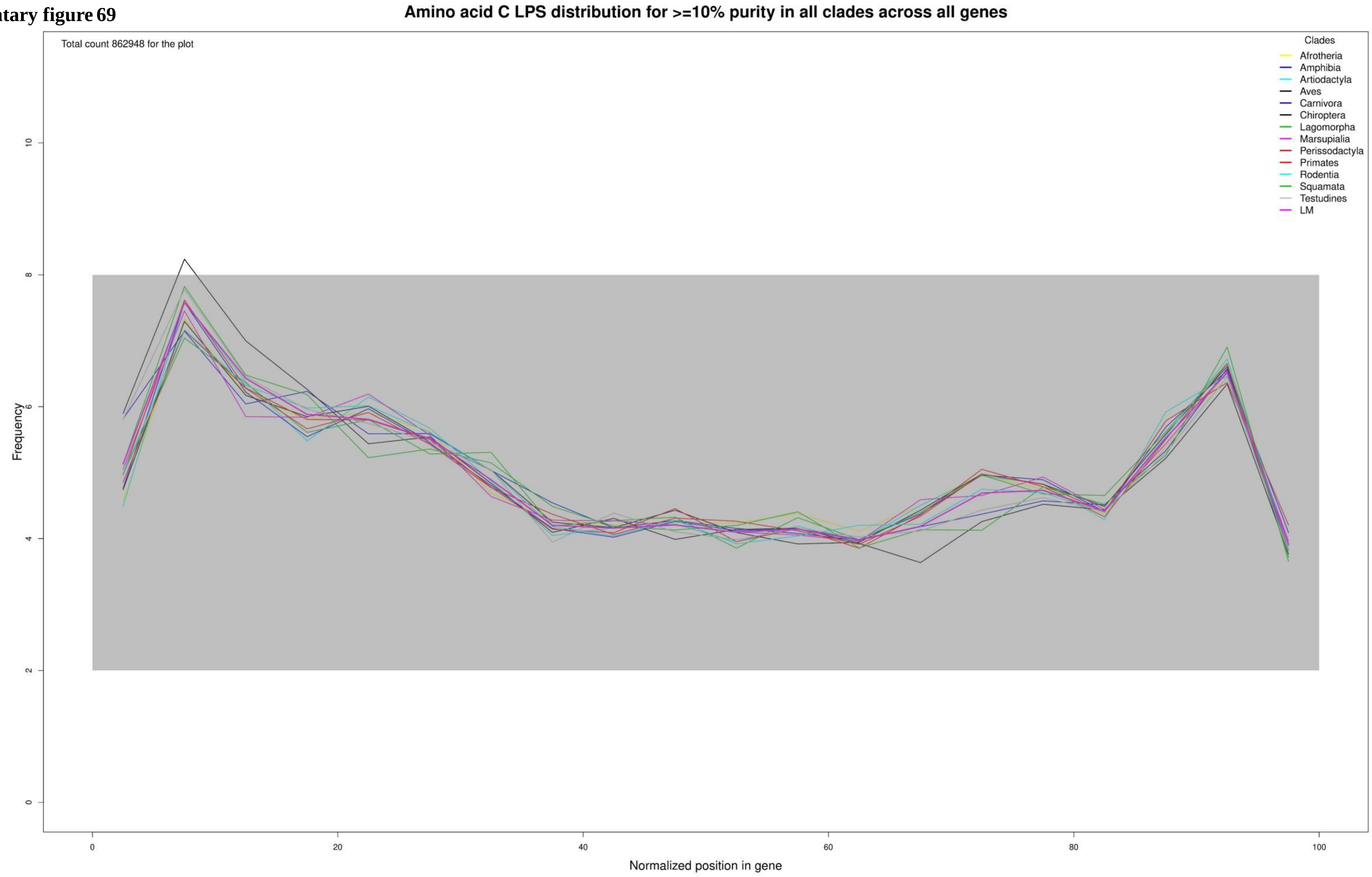

**Supplementary figure 70**

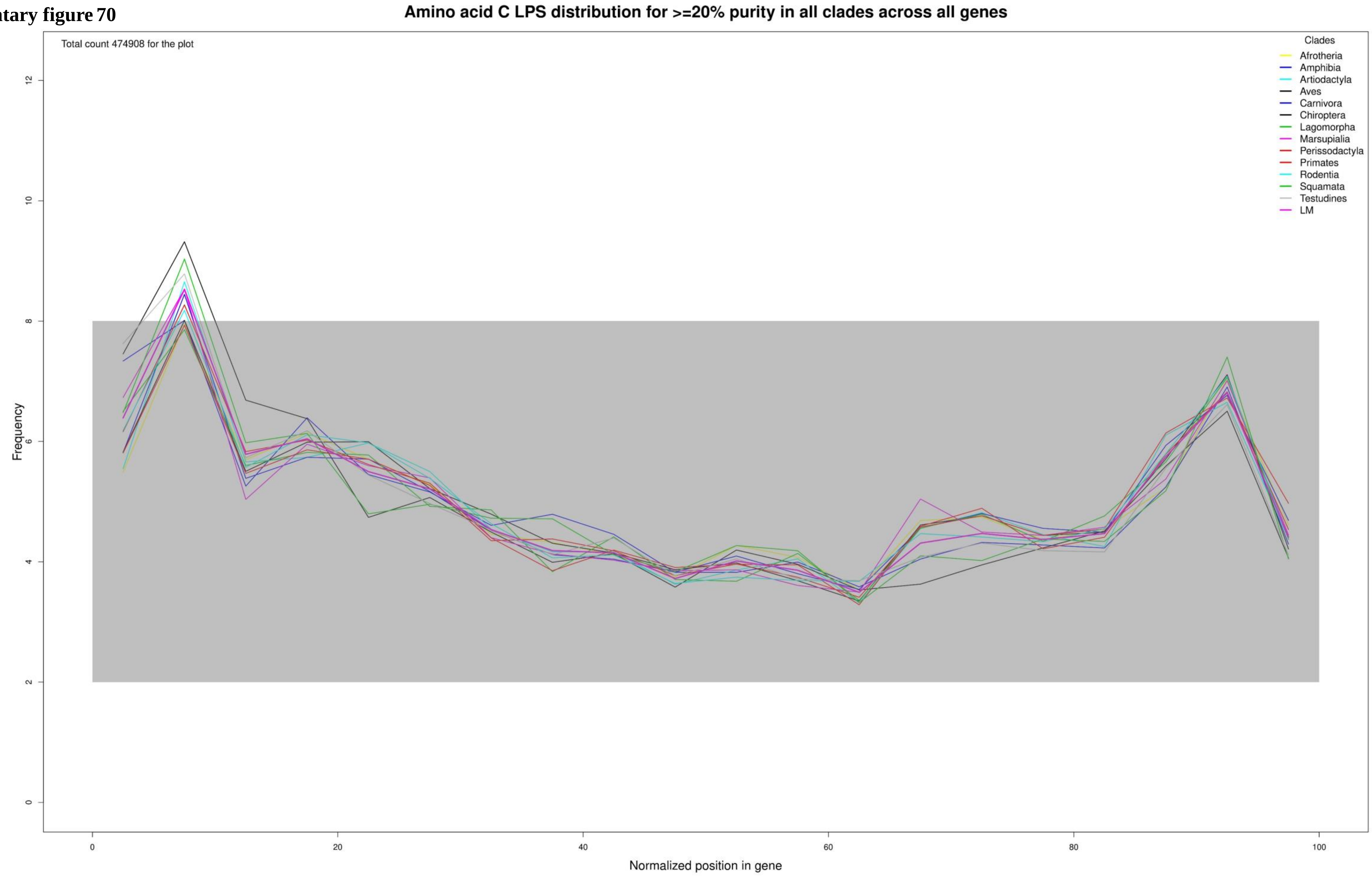

**Supplementary figure 71**

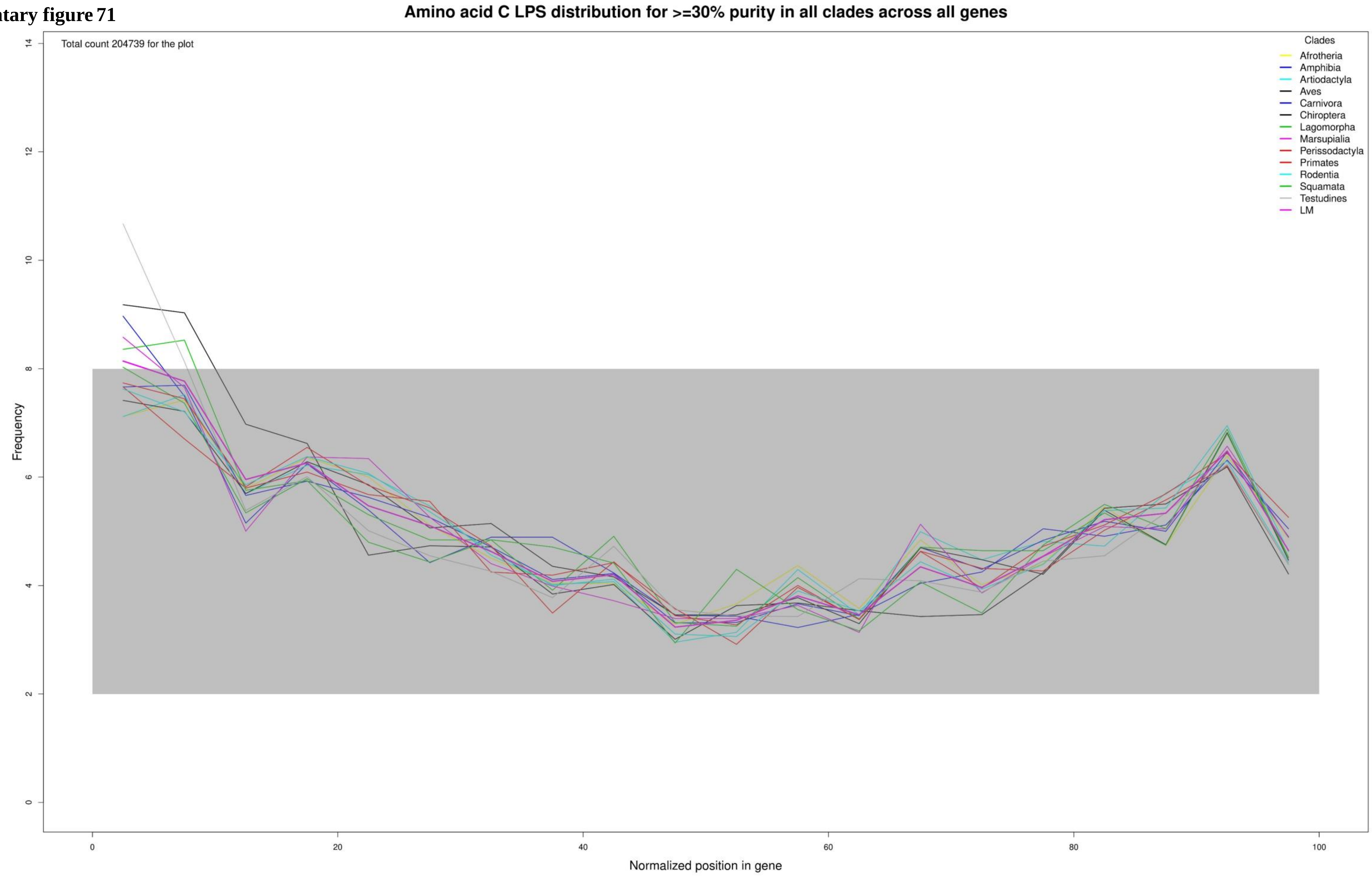

**Supplementary figure 72**

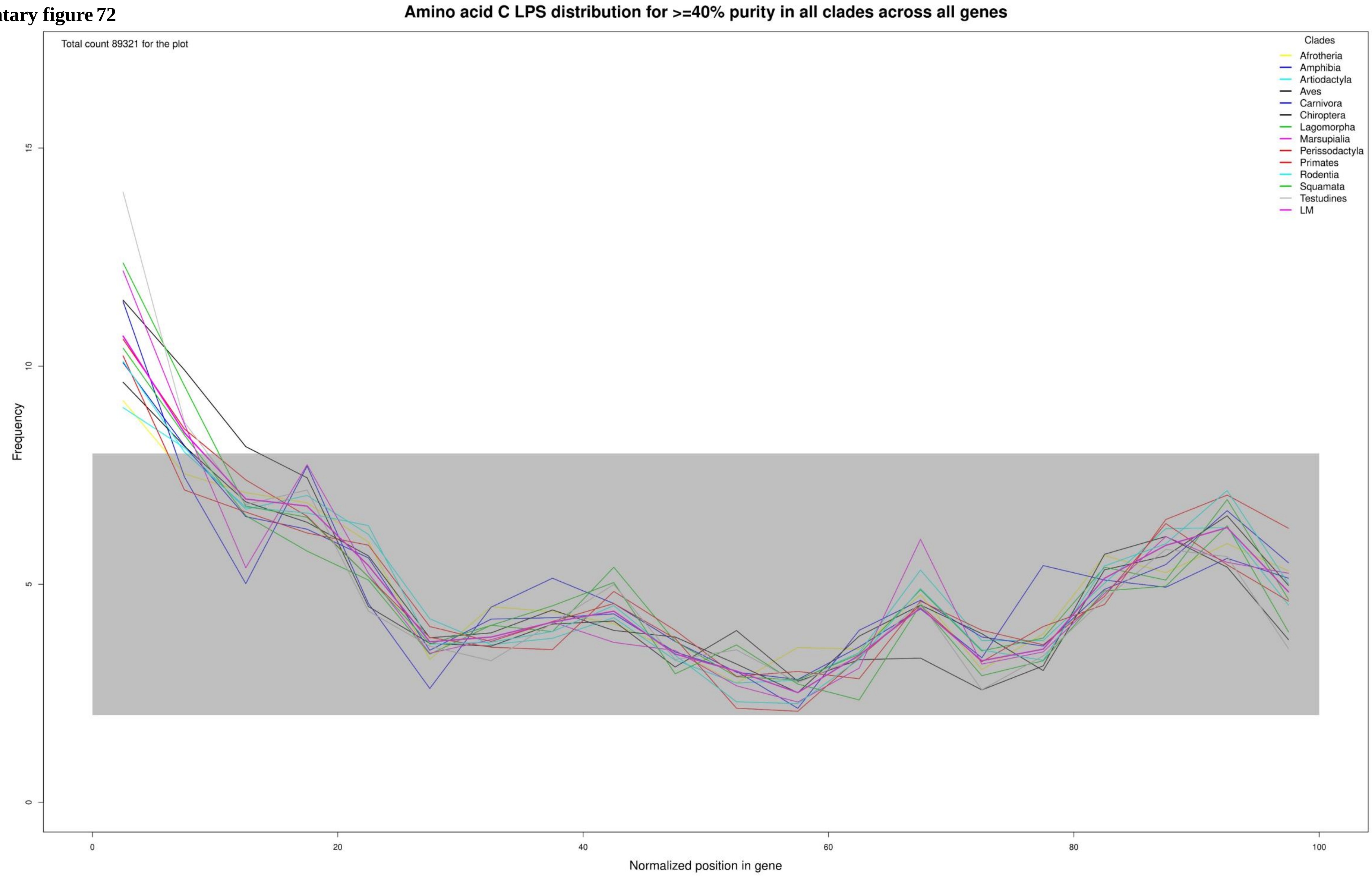

**Supplementary figure 73**

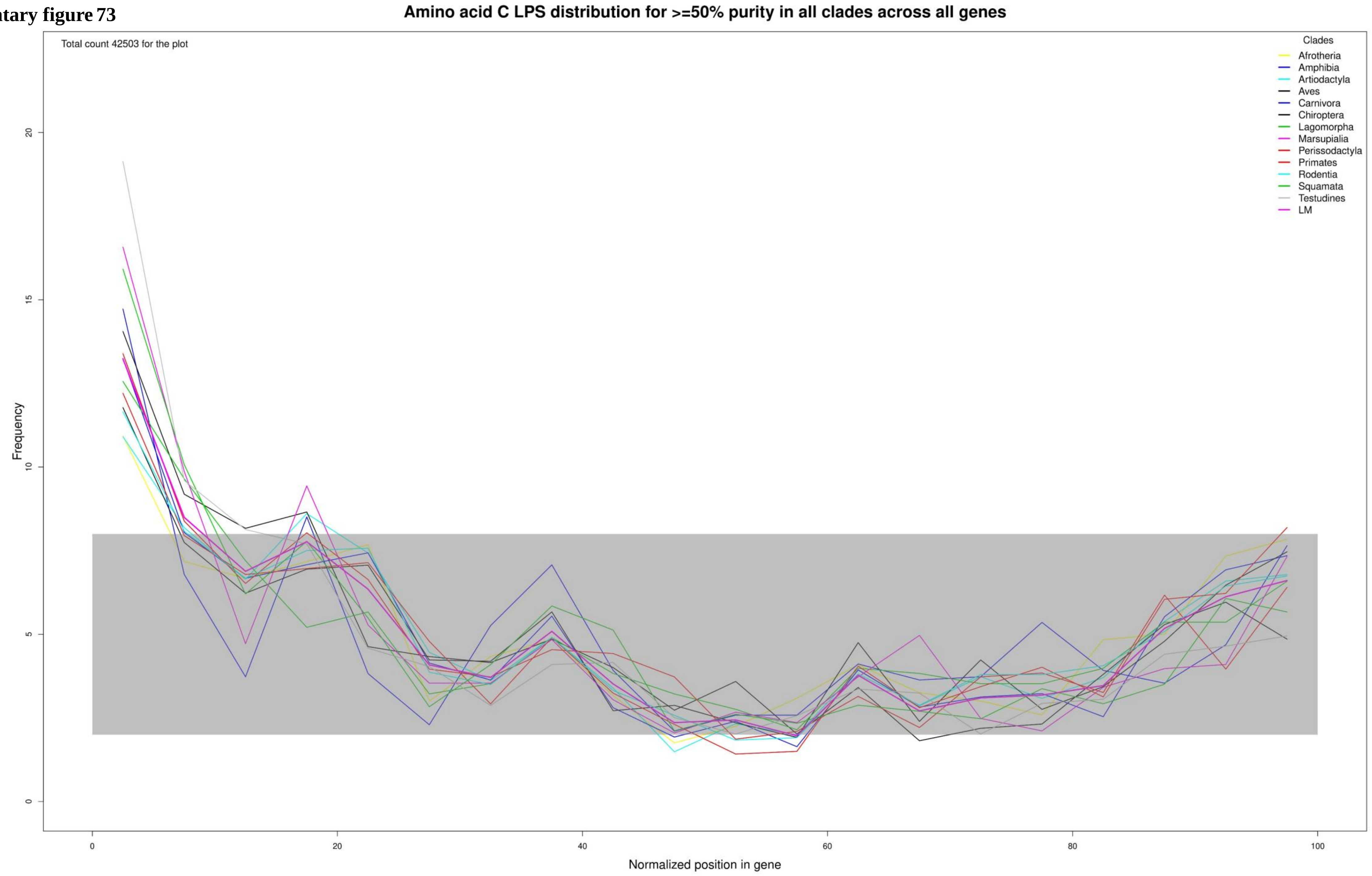

**Supplementary figure 74**

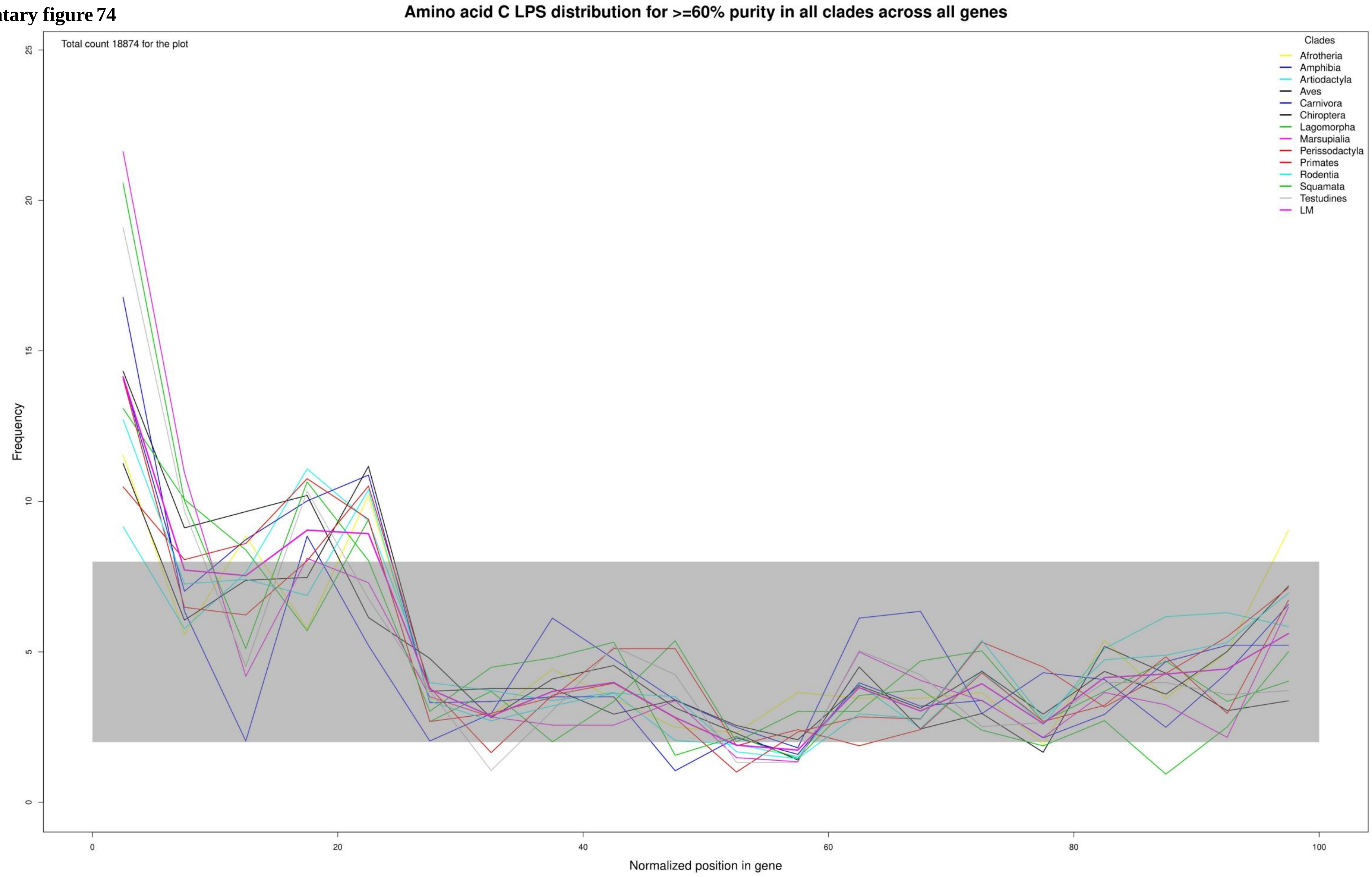

**Supplementary figure 75**

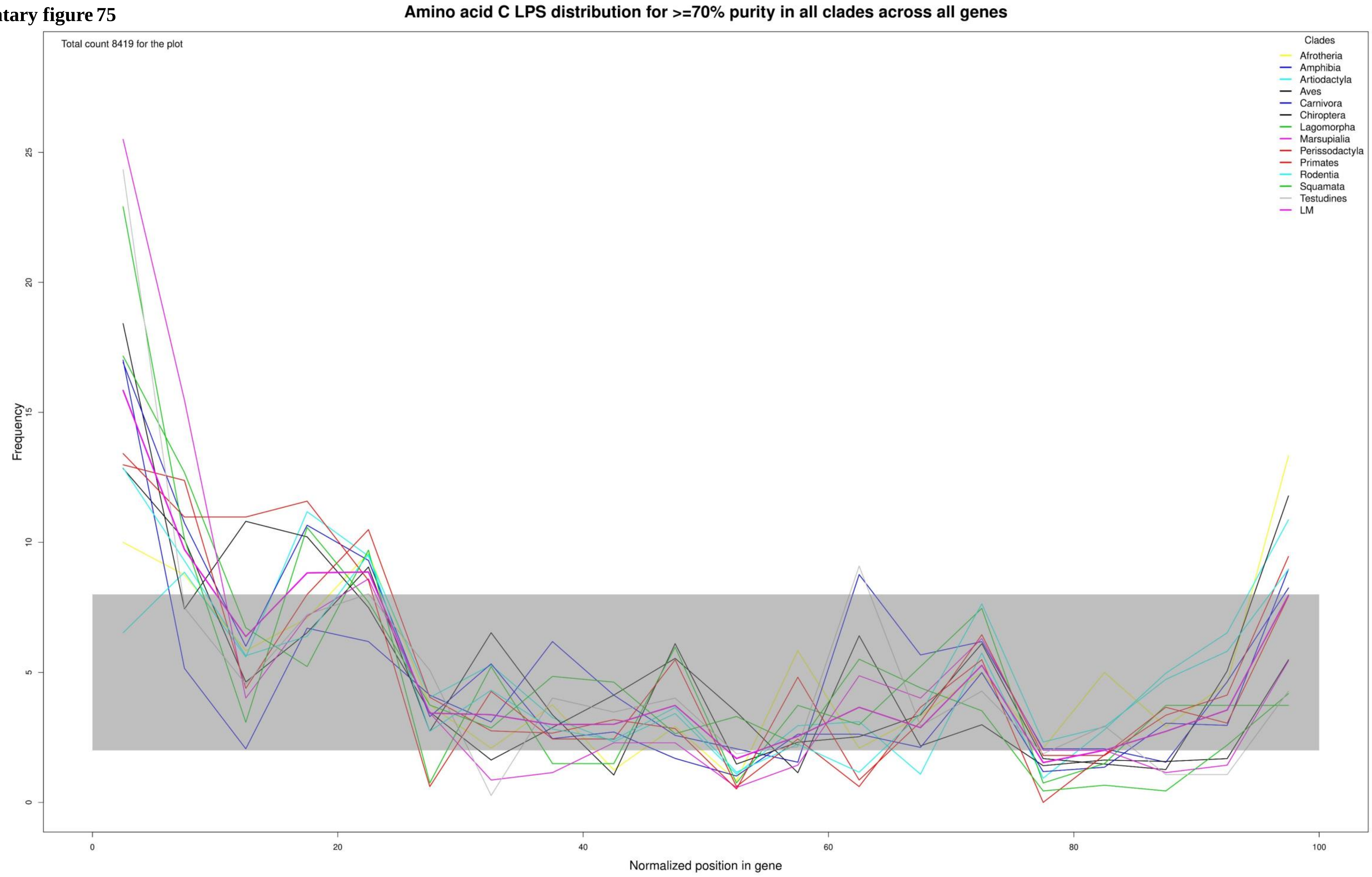

**Supplementary figure 76**

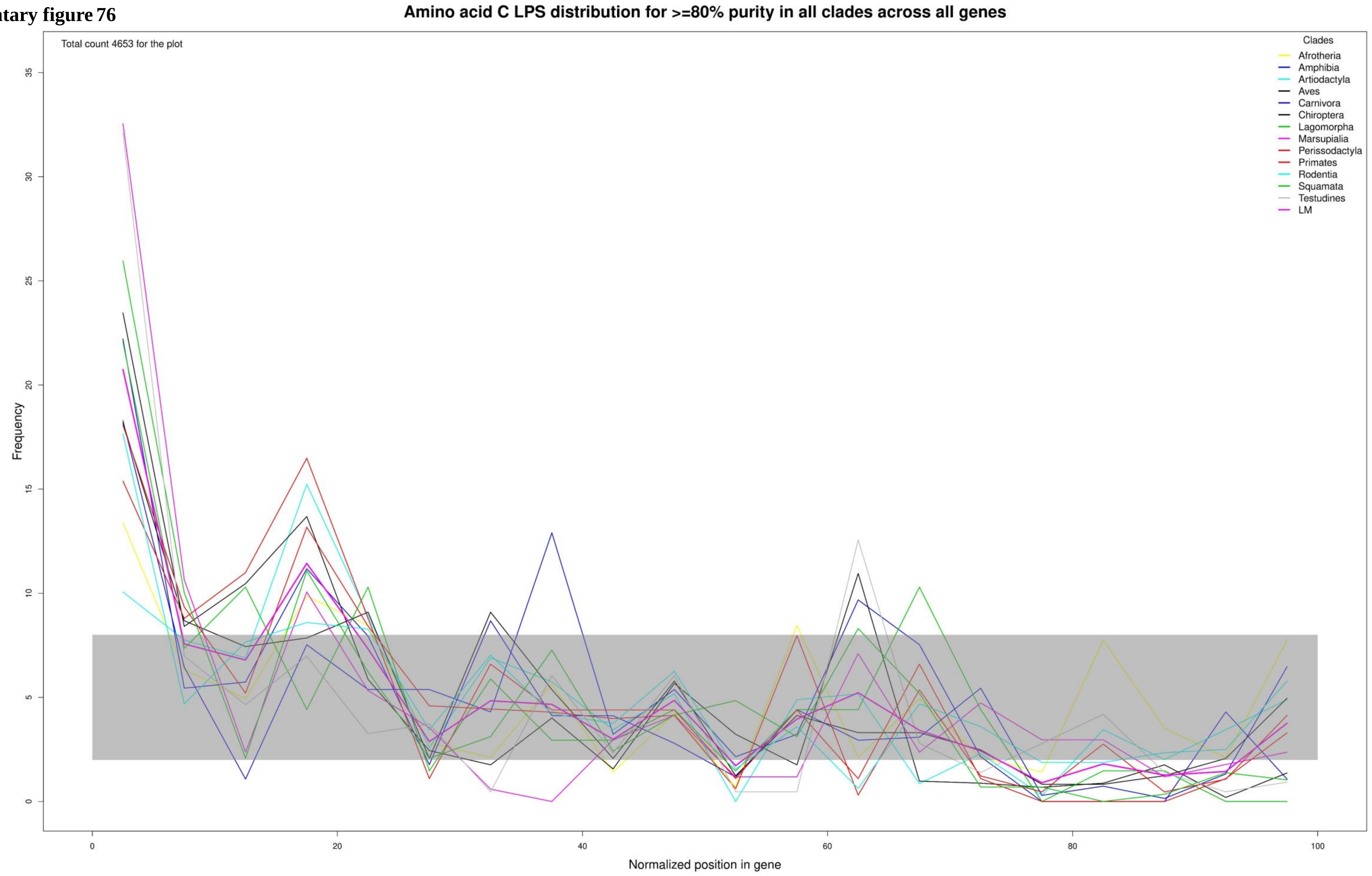

**Supplementary figure 77**

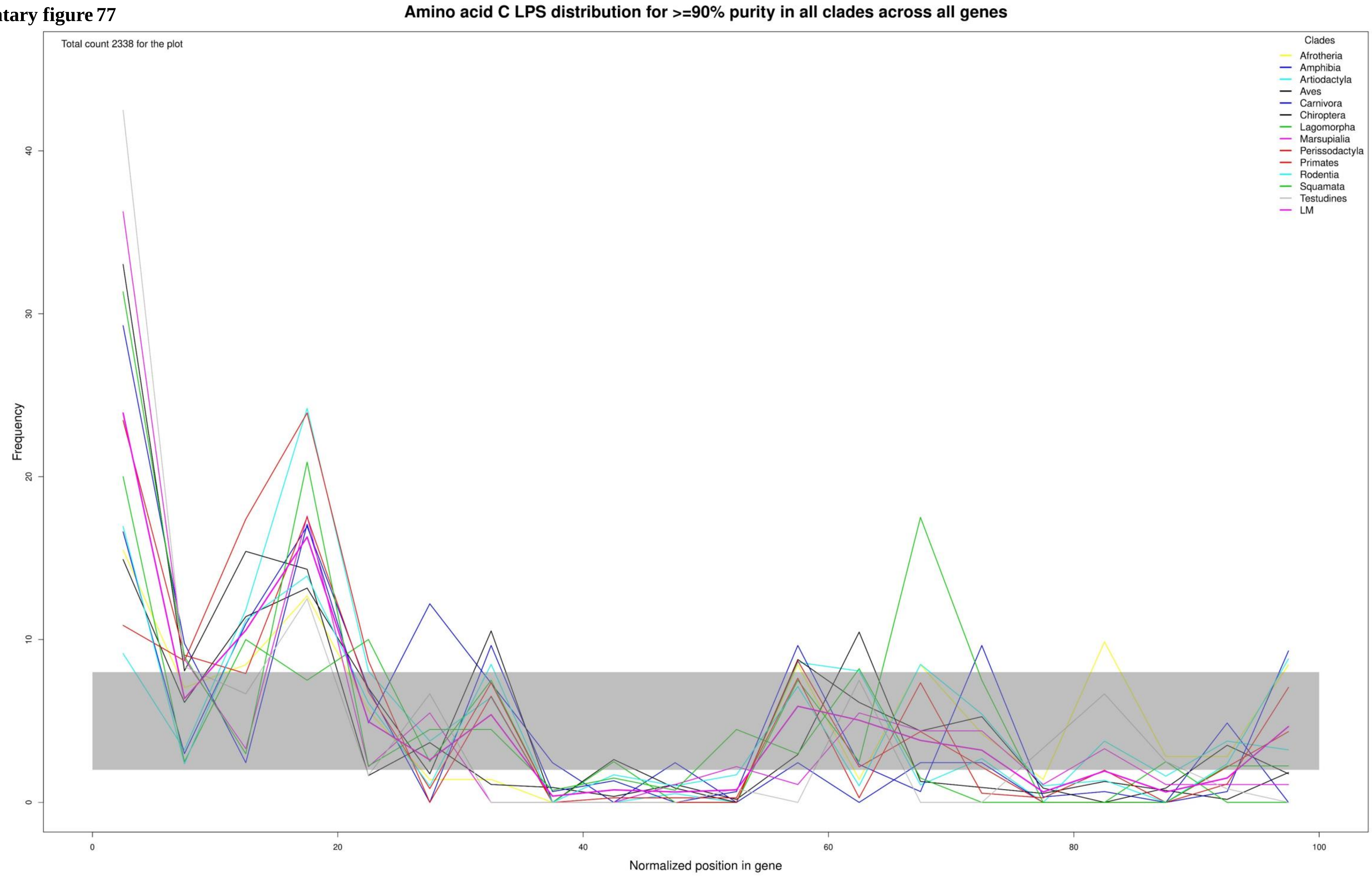

**Supplementary figure 78**

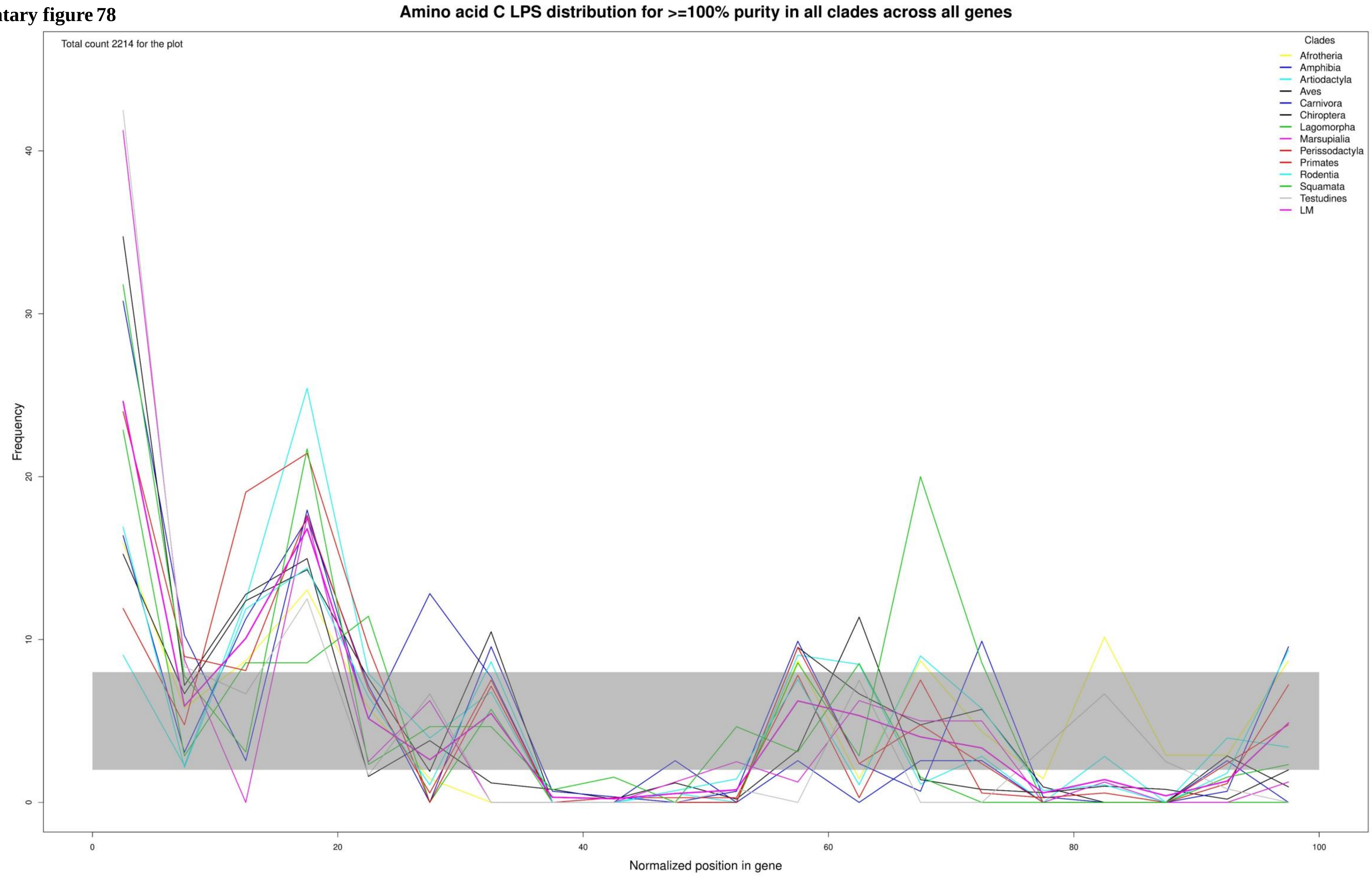

**Supplementary figure 79**

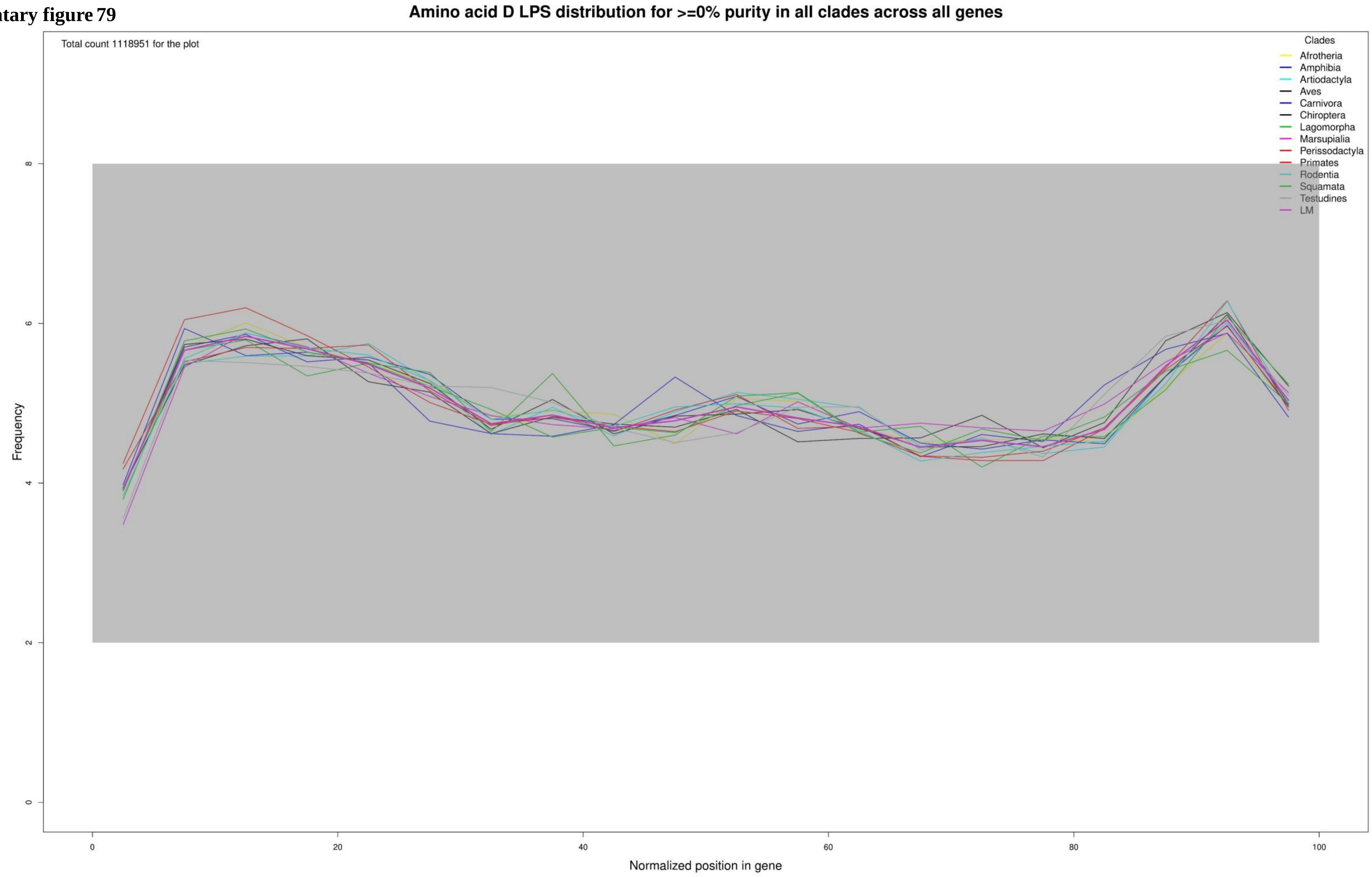

**Supplementary figure 80**

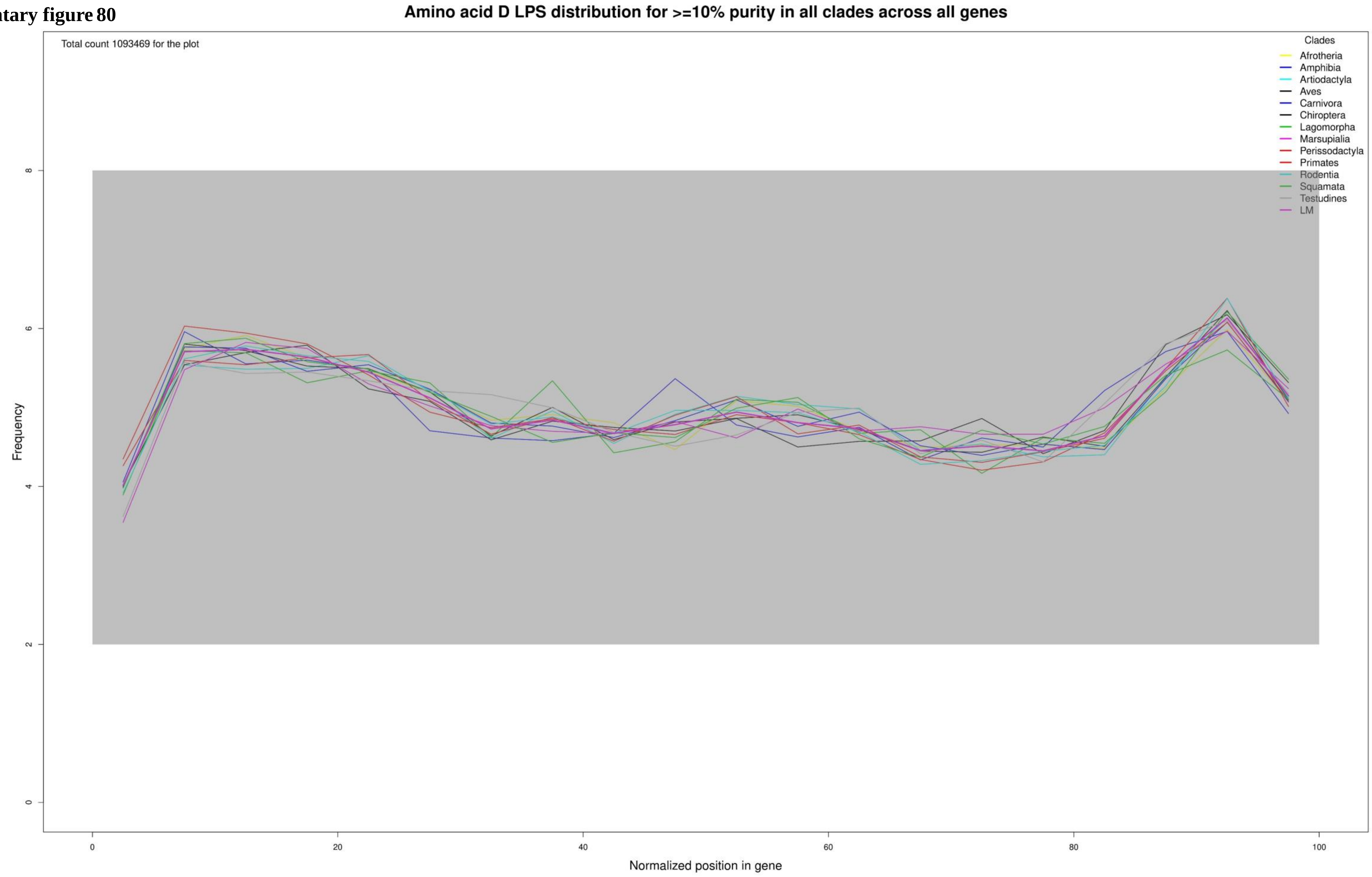

**Supplementary figure 81**

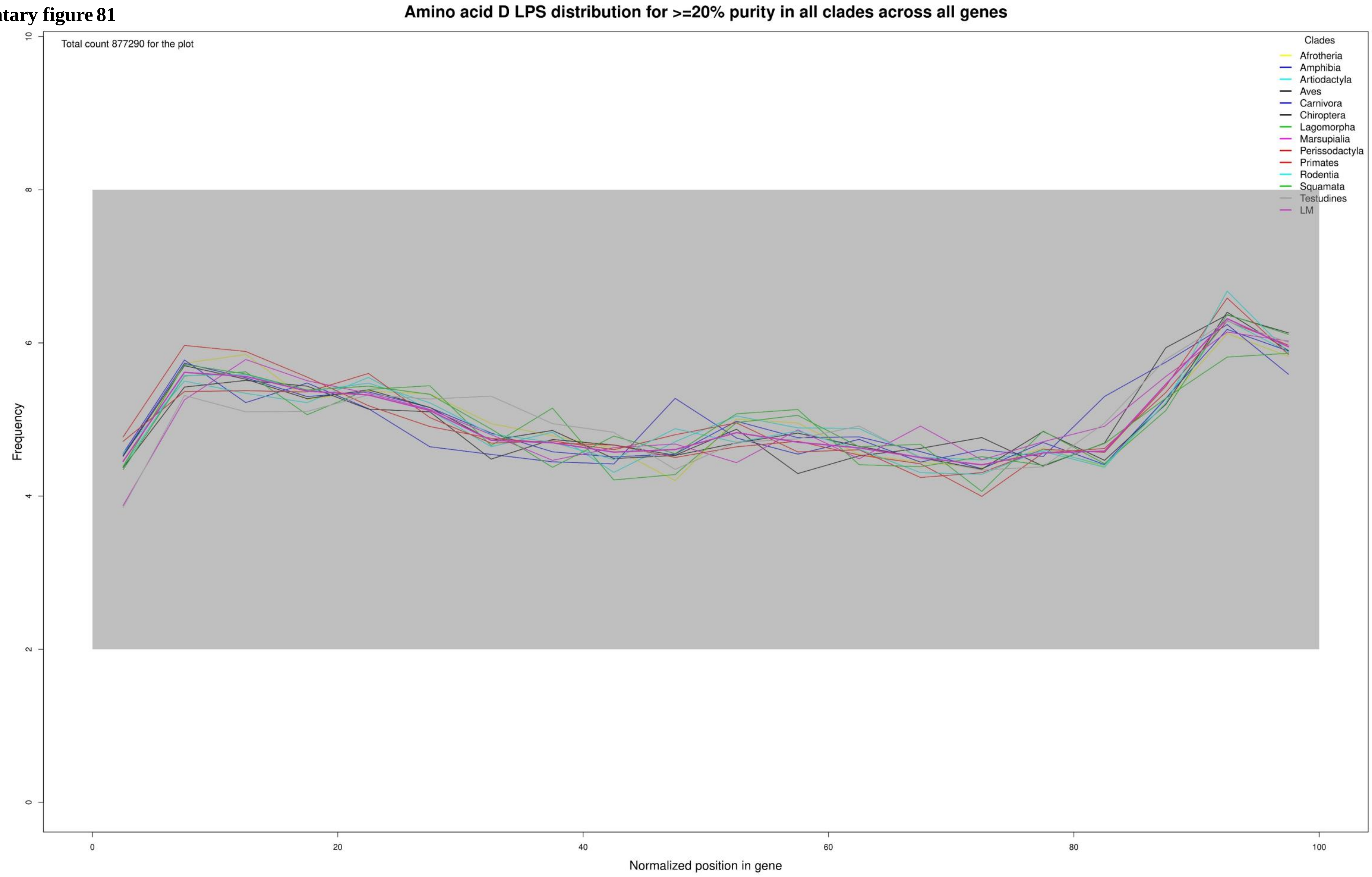

**Supplementary figure 82**

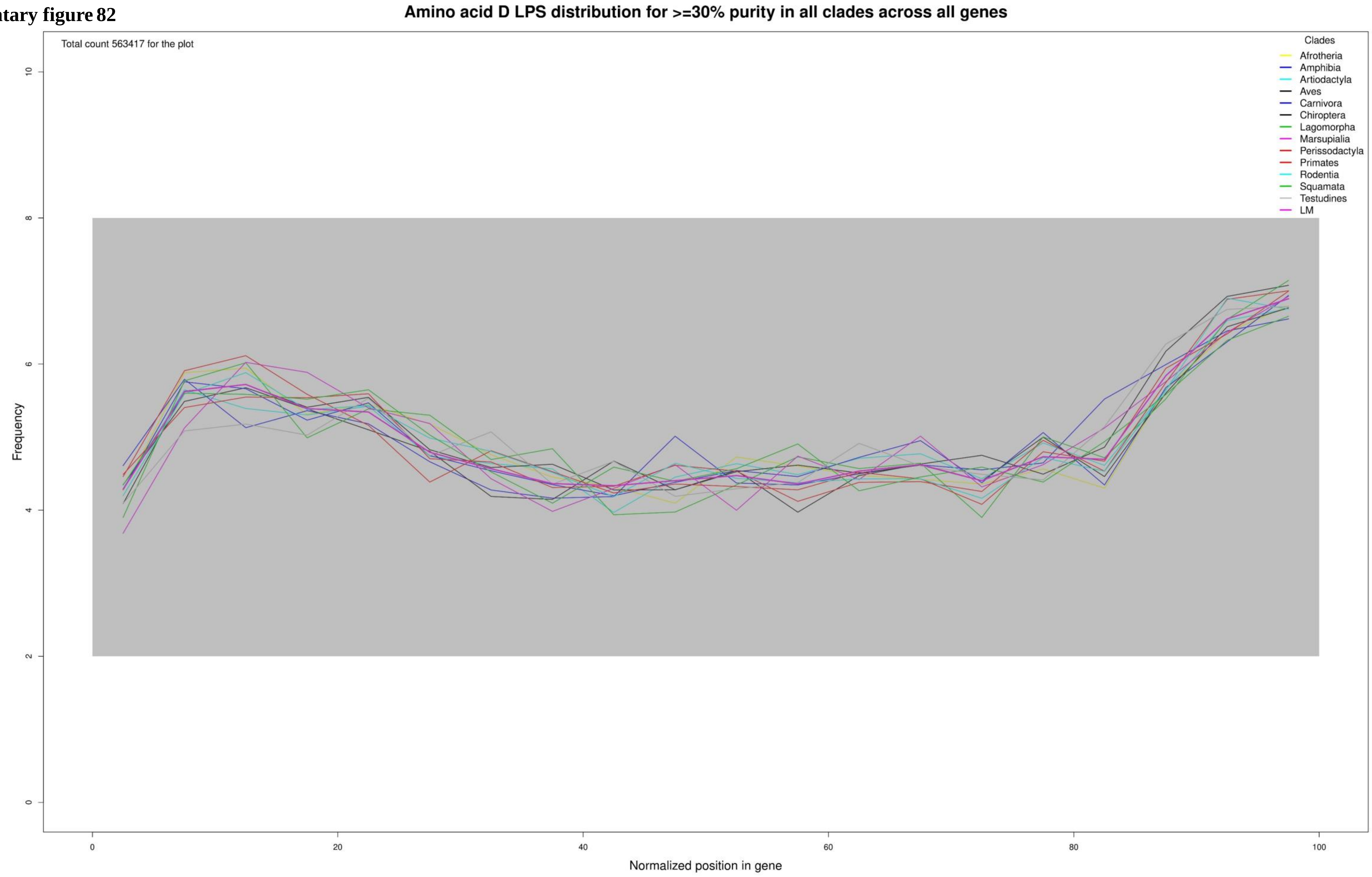

**Supplementary figure 83**

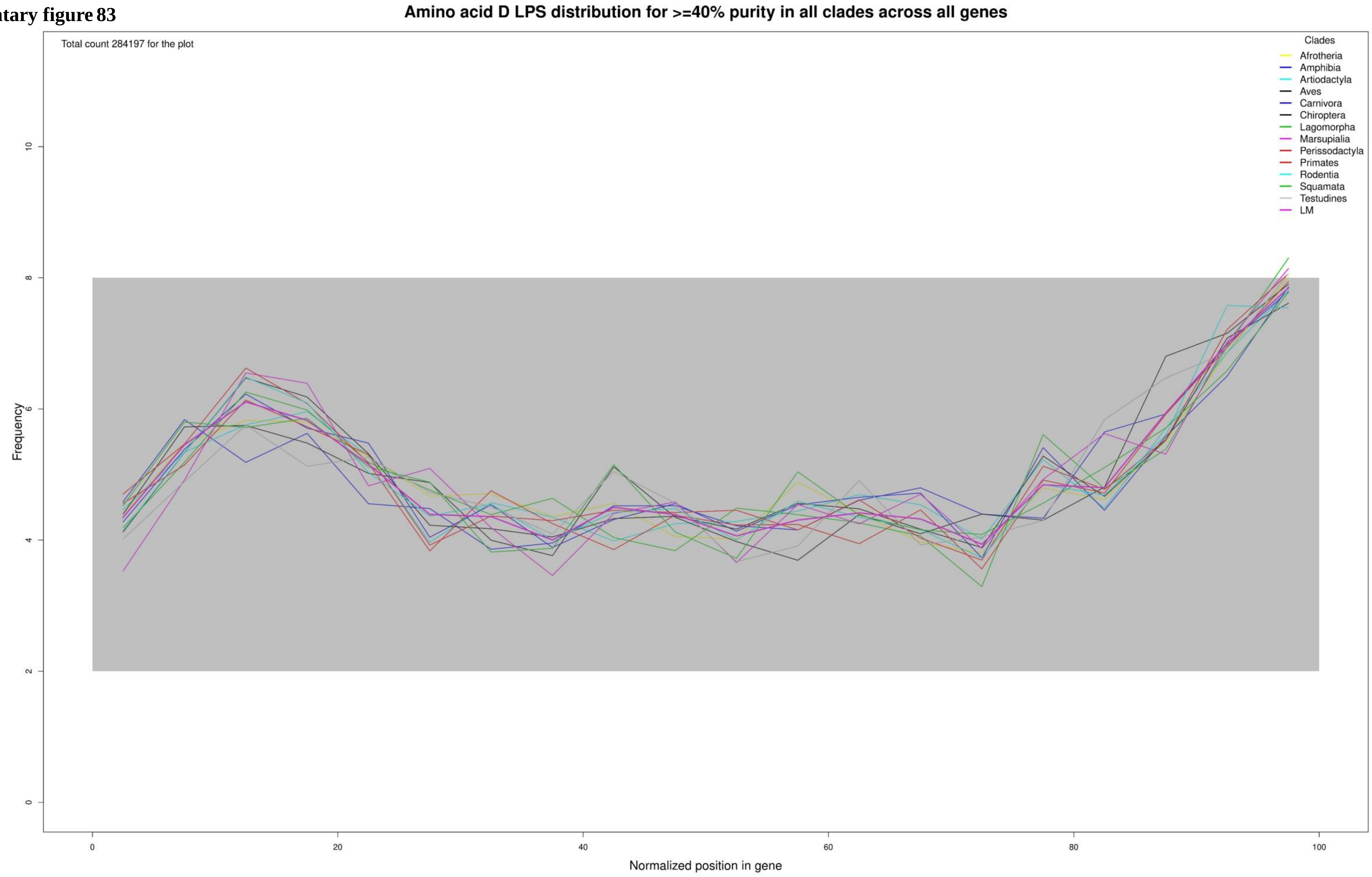

**Supplementary figure 84**

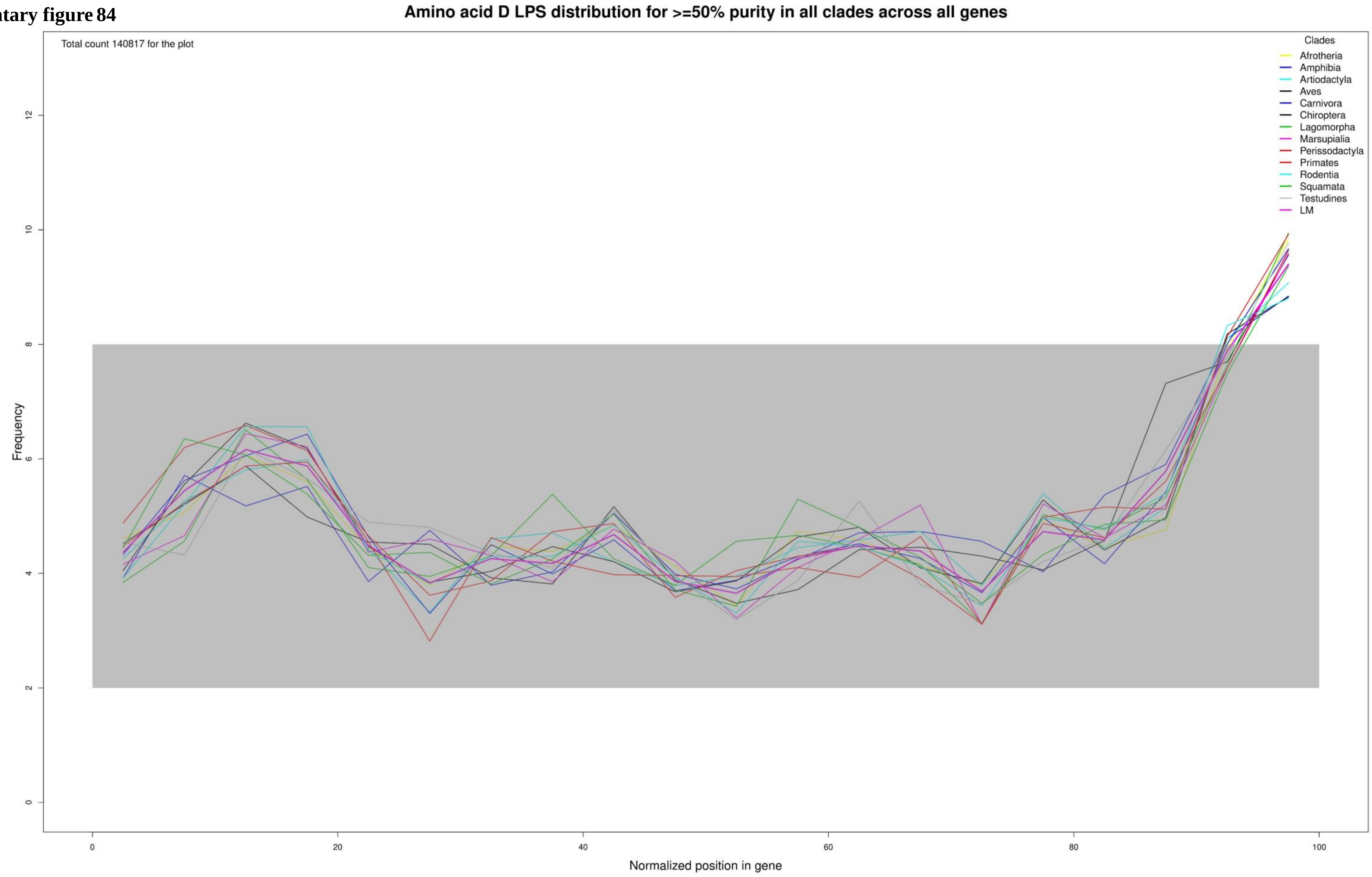

**Supplementary figure 85**

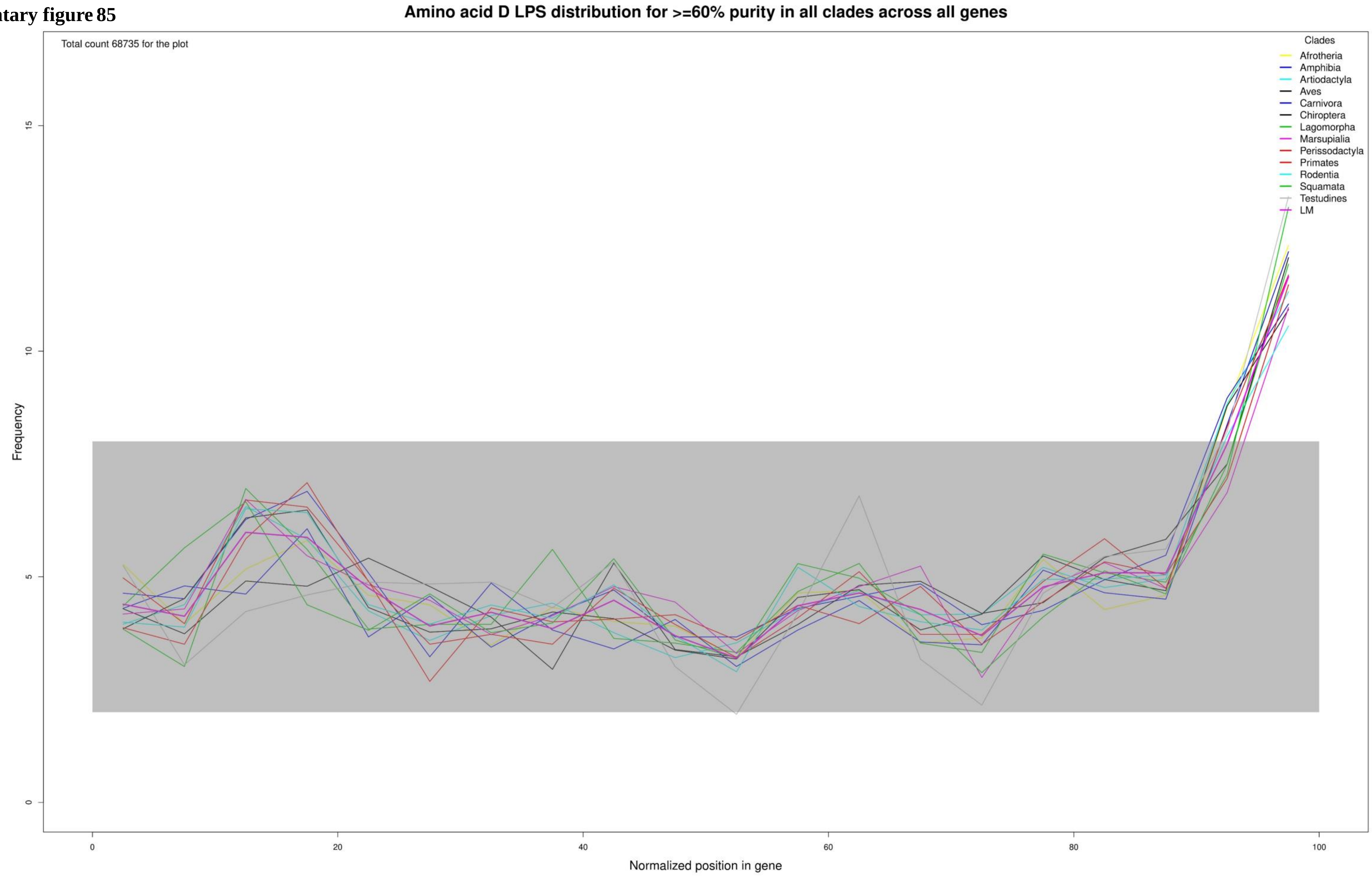

**Supplementary figure 86**

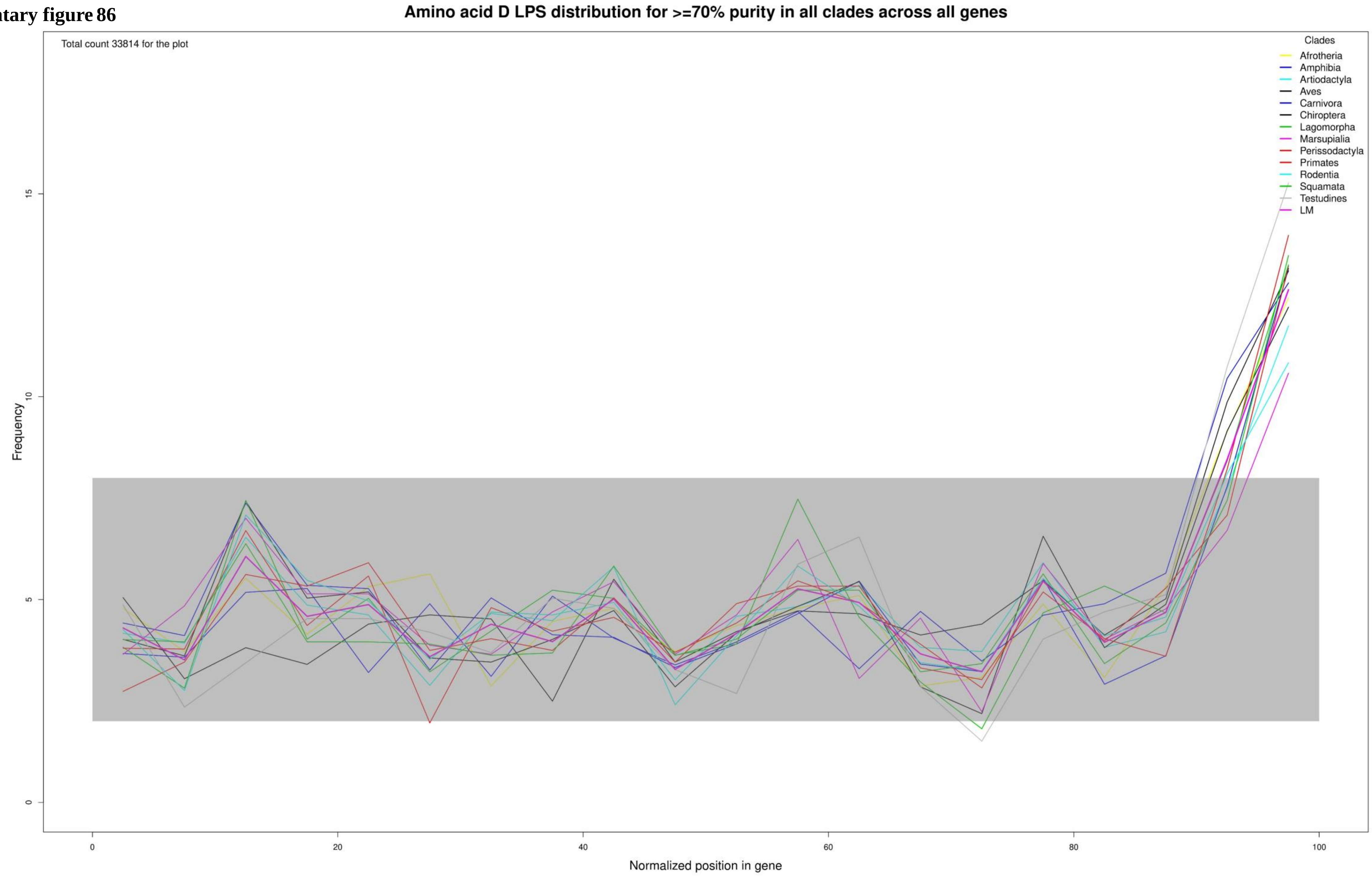

**Supplementary figure 87**

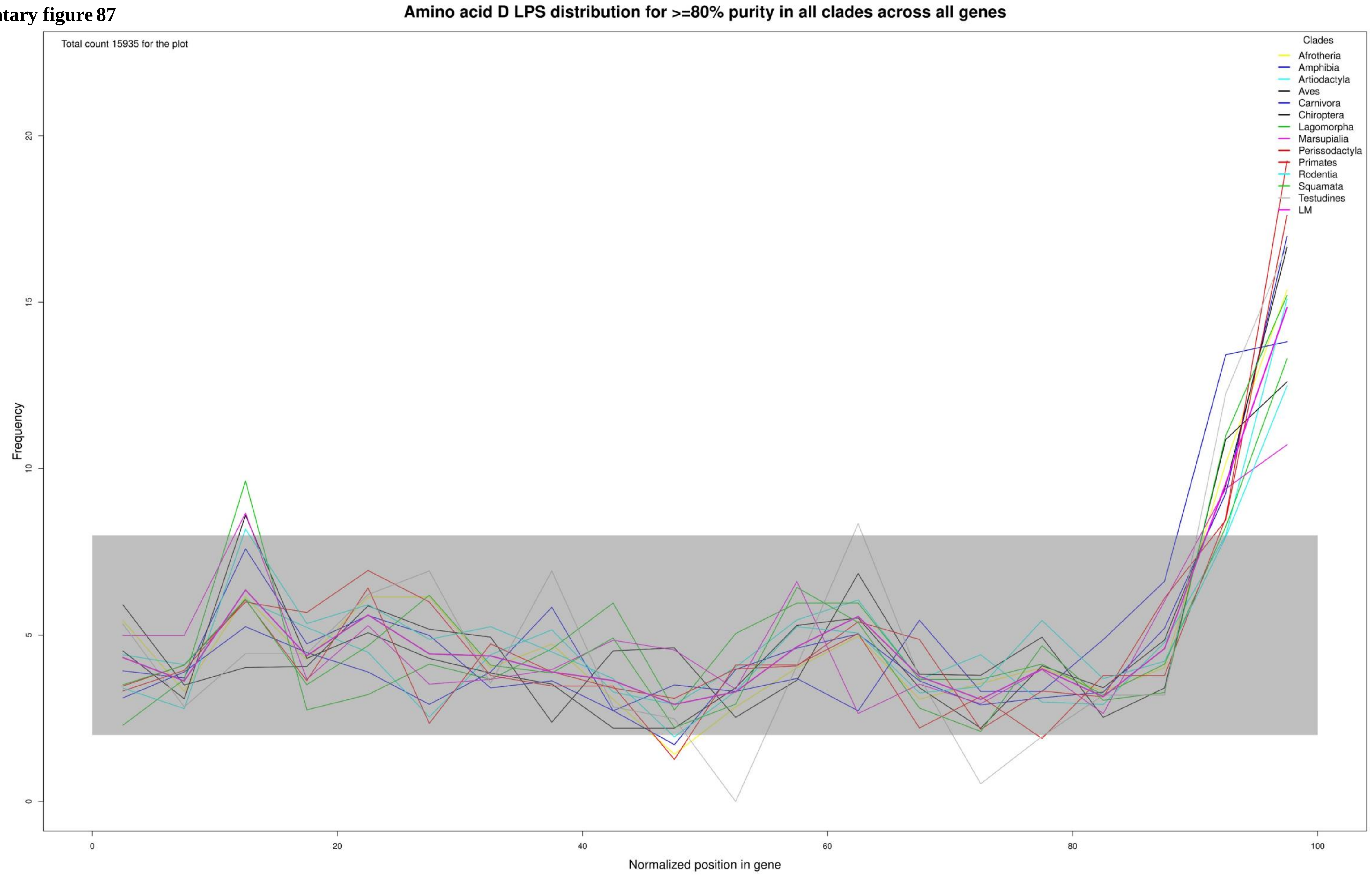

**Supplementary figure 88**

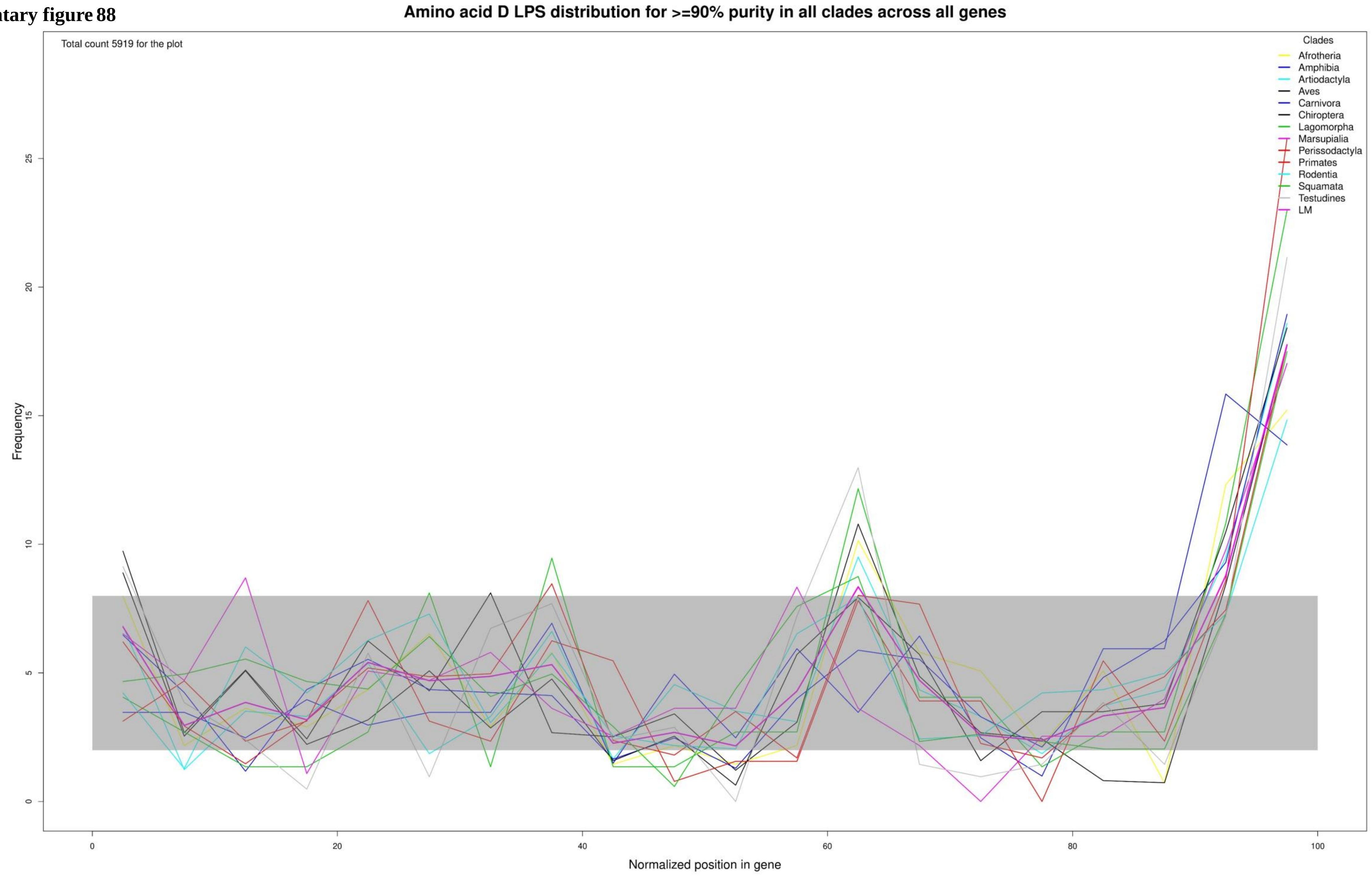

**Supplementary figure 89**

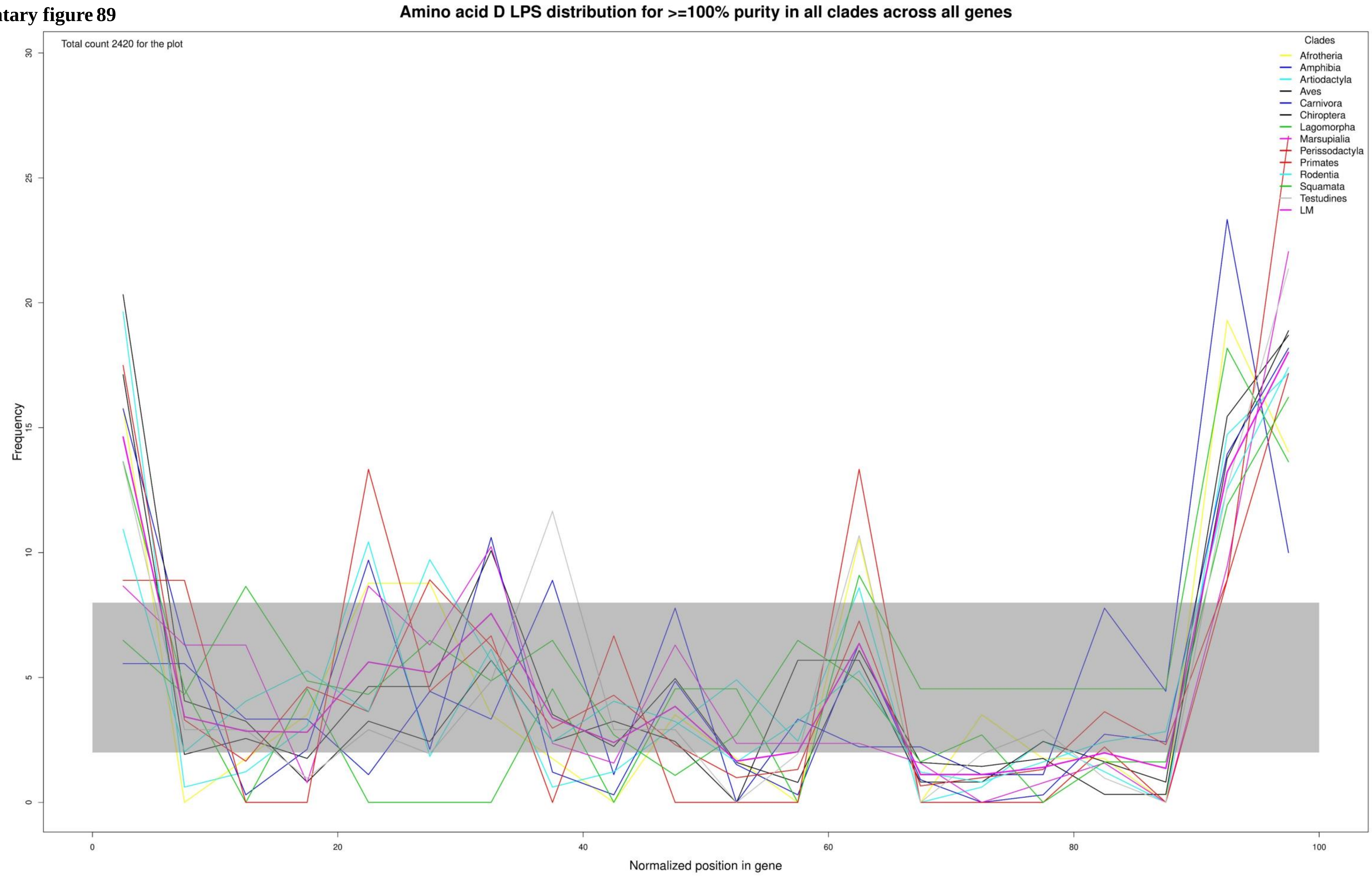

**Supplementary figure 90**

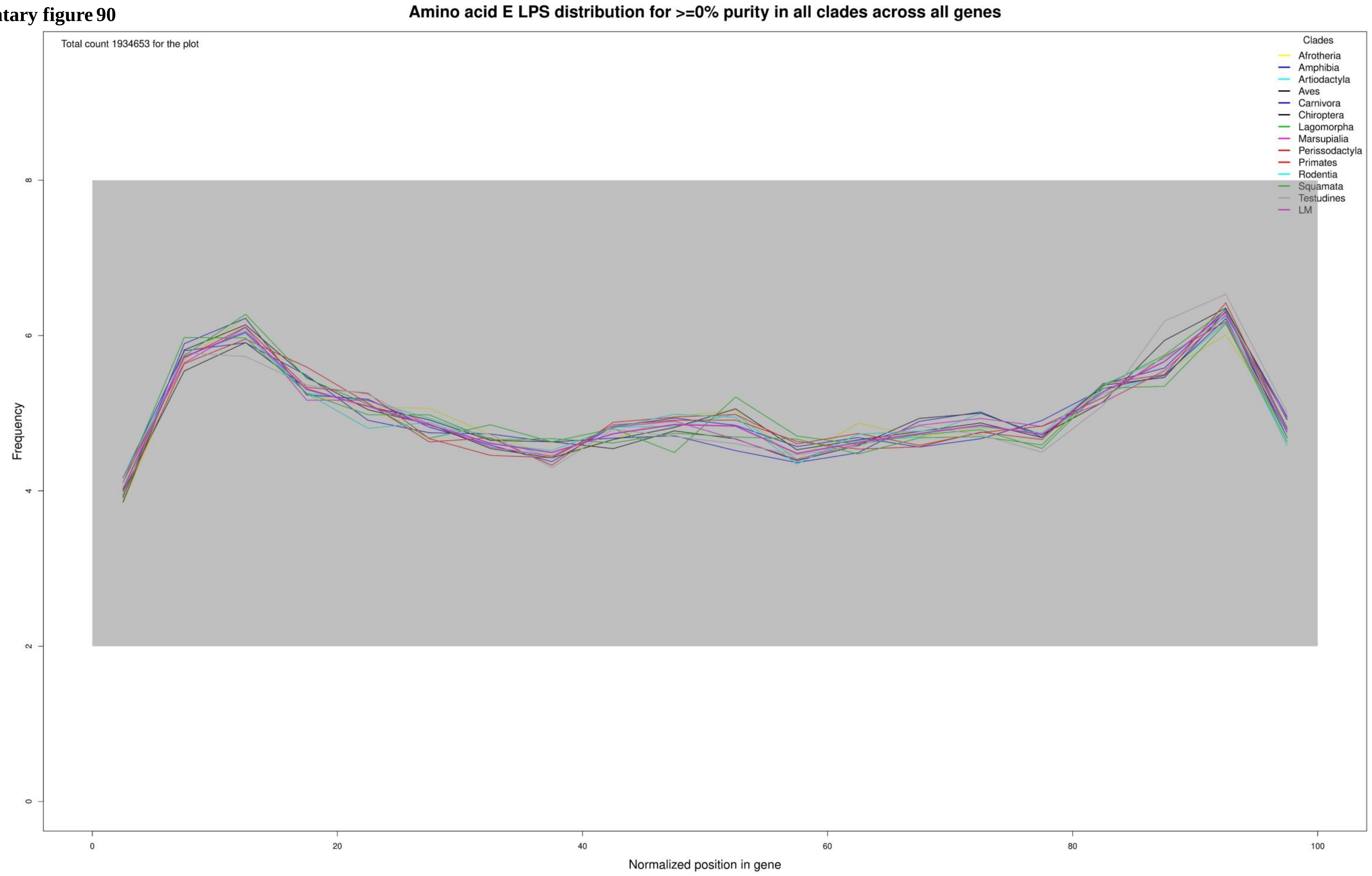

**Supplementary figure 91**

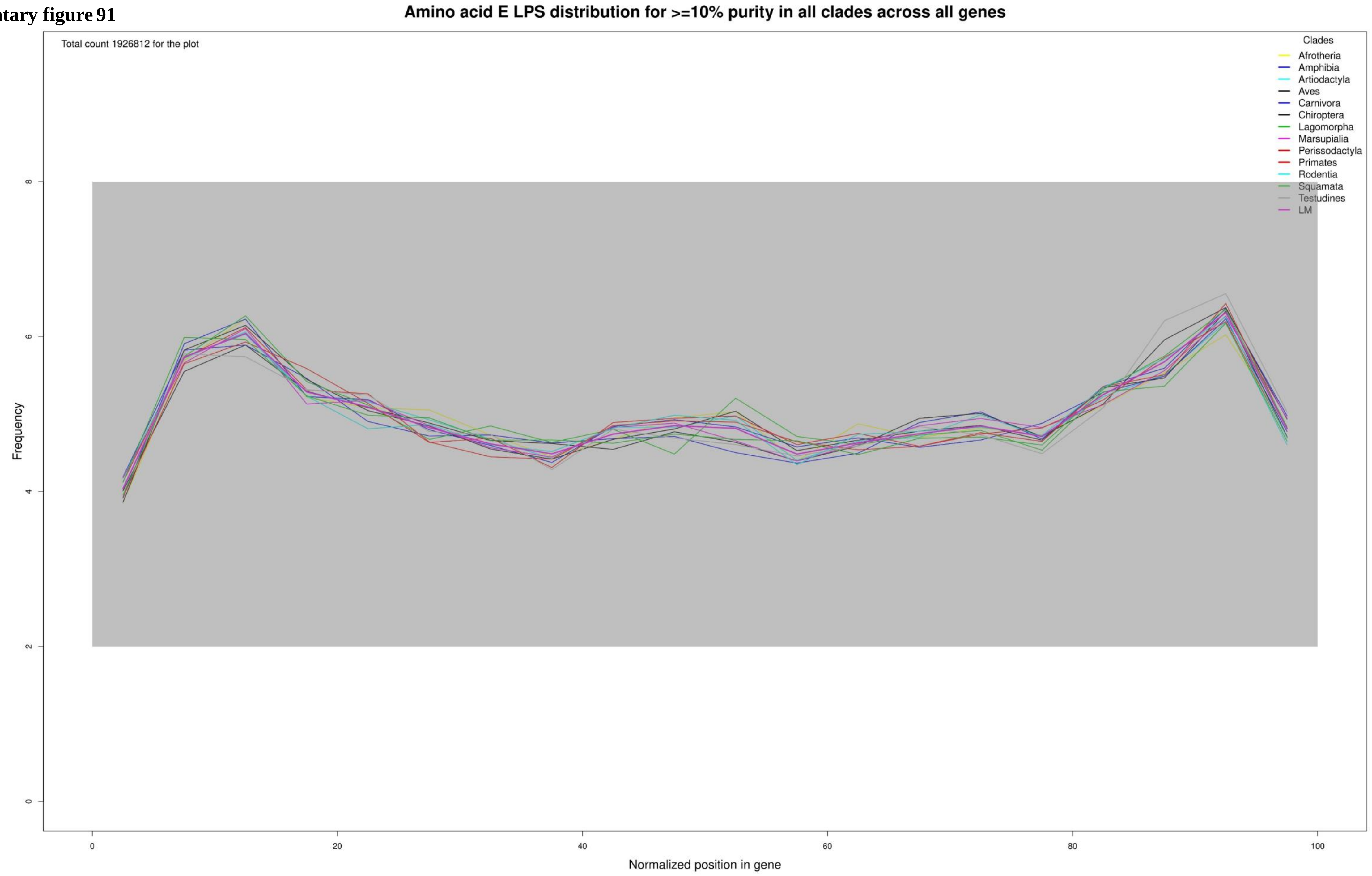

**Supplementary figure 92**

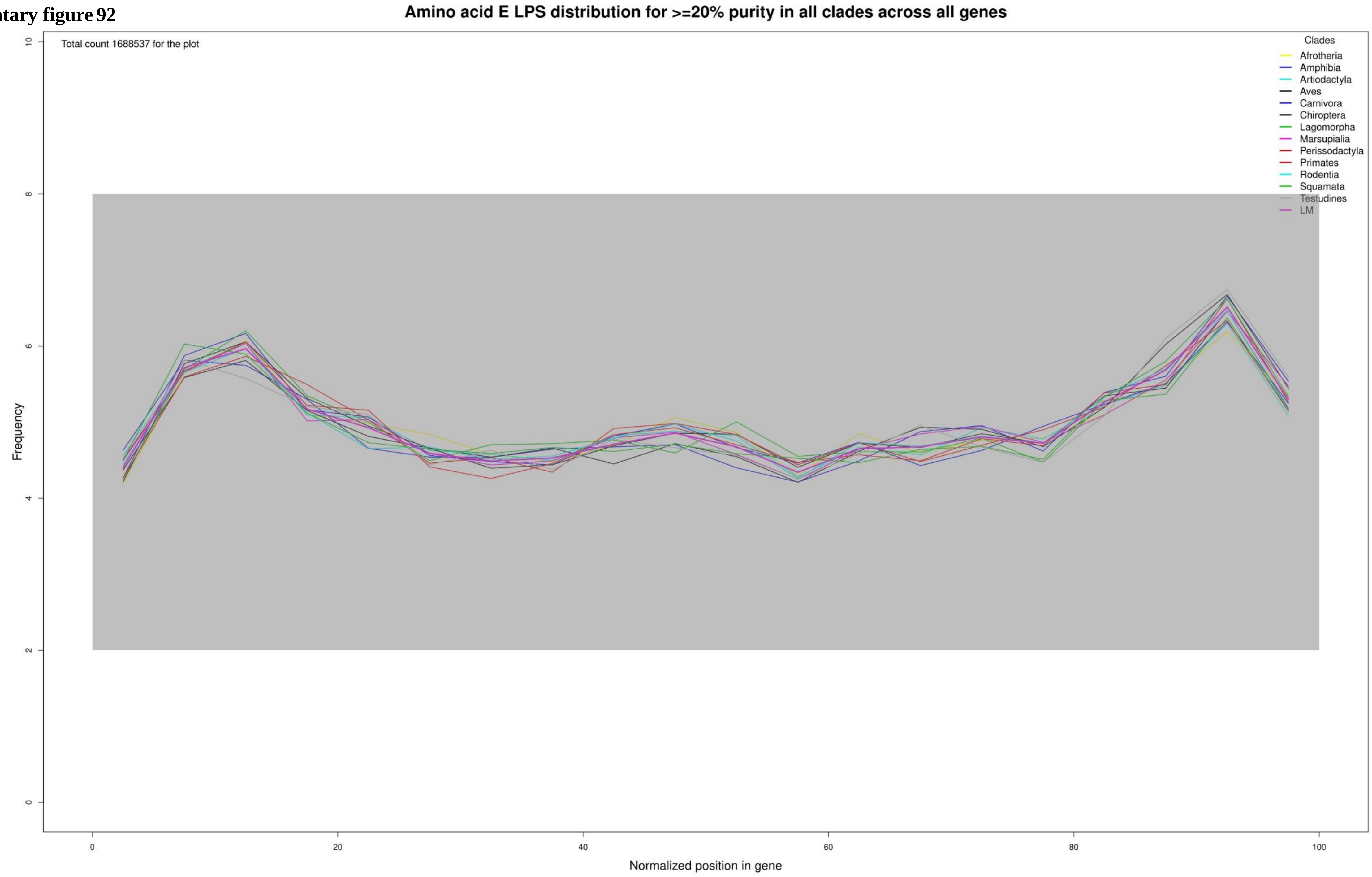

**Supplementary figure 93**

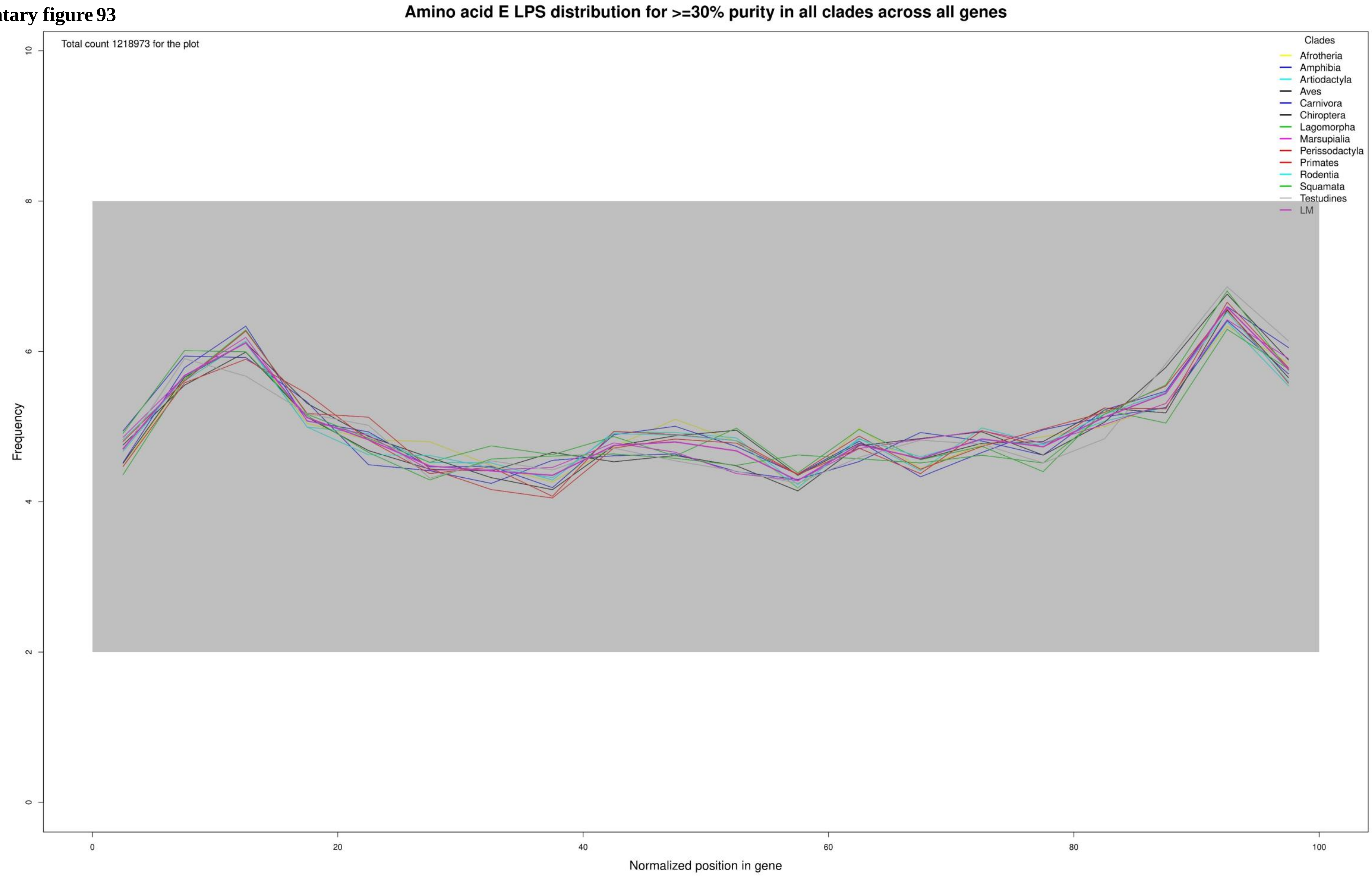

**Supplementary figure 94**

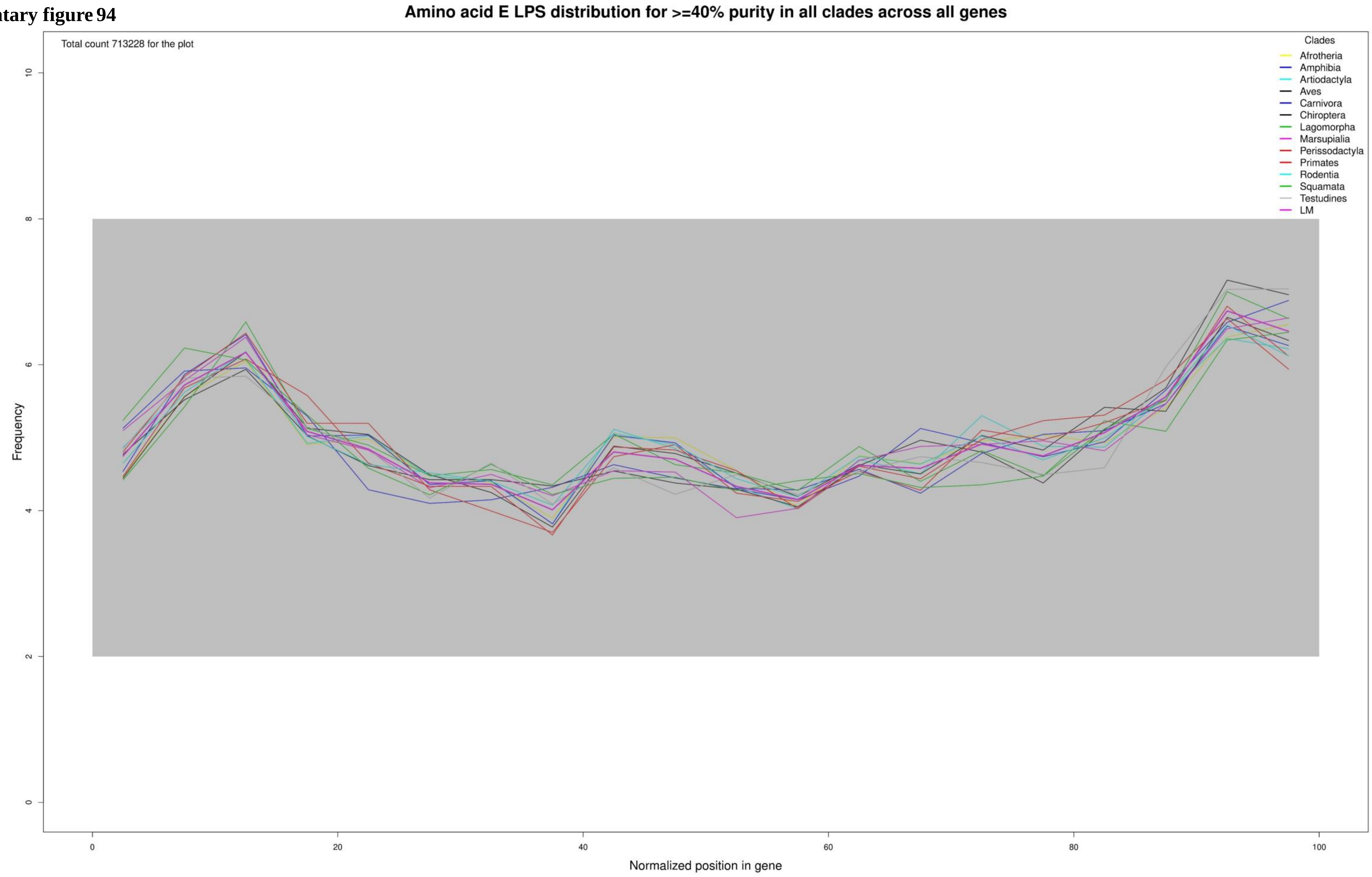

**Supplementary figure 95**

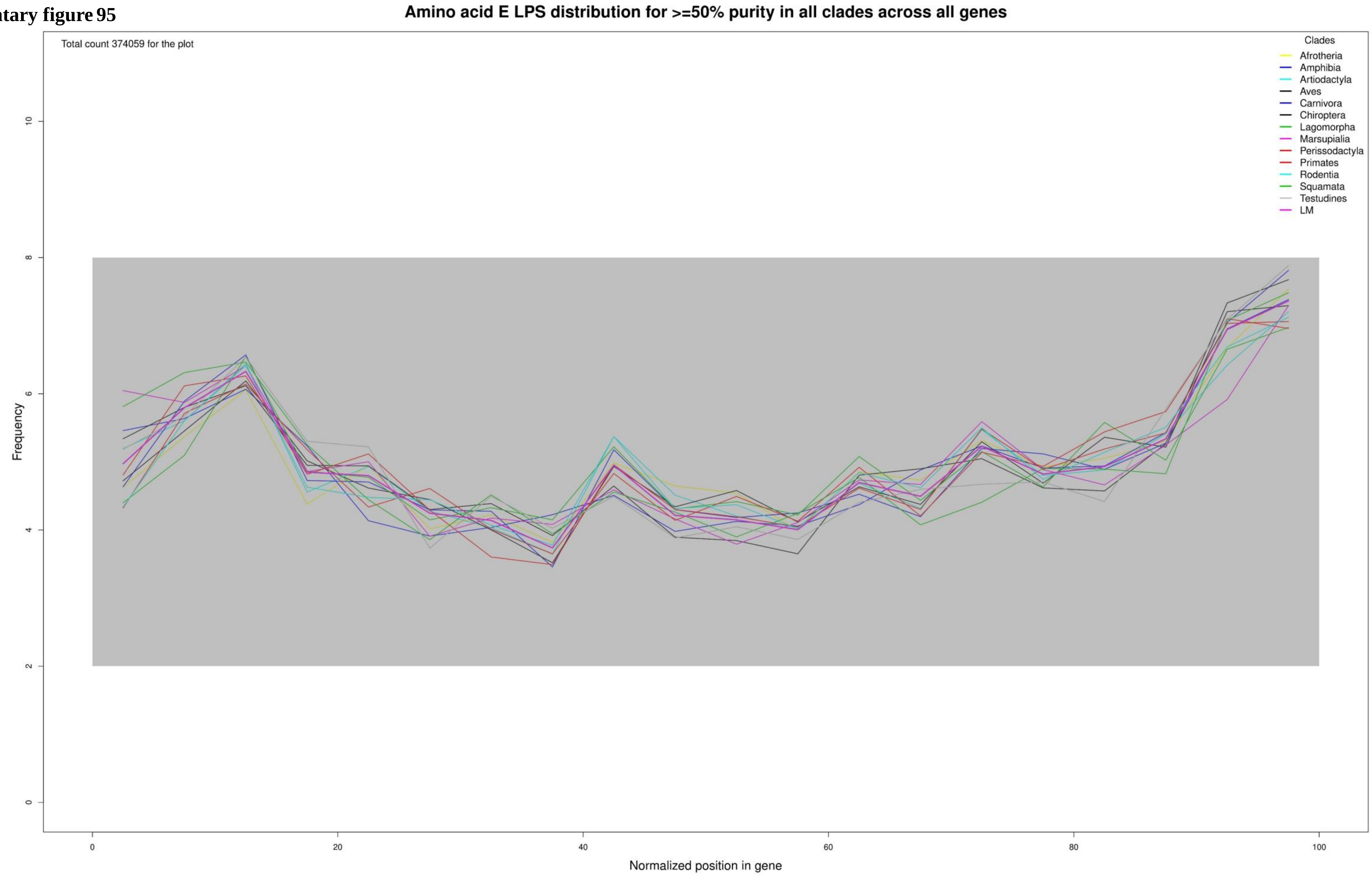

**Supplementary figure 96**

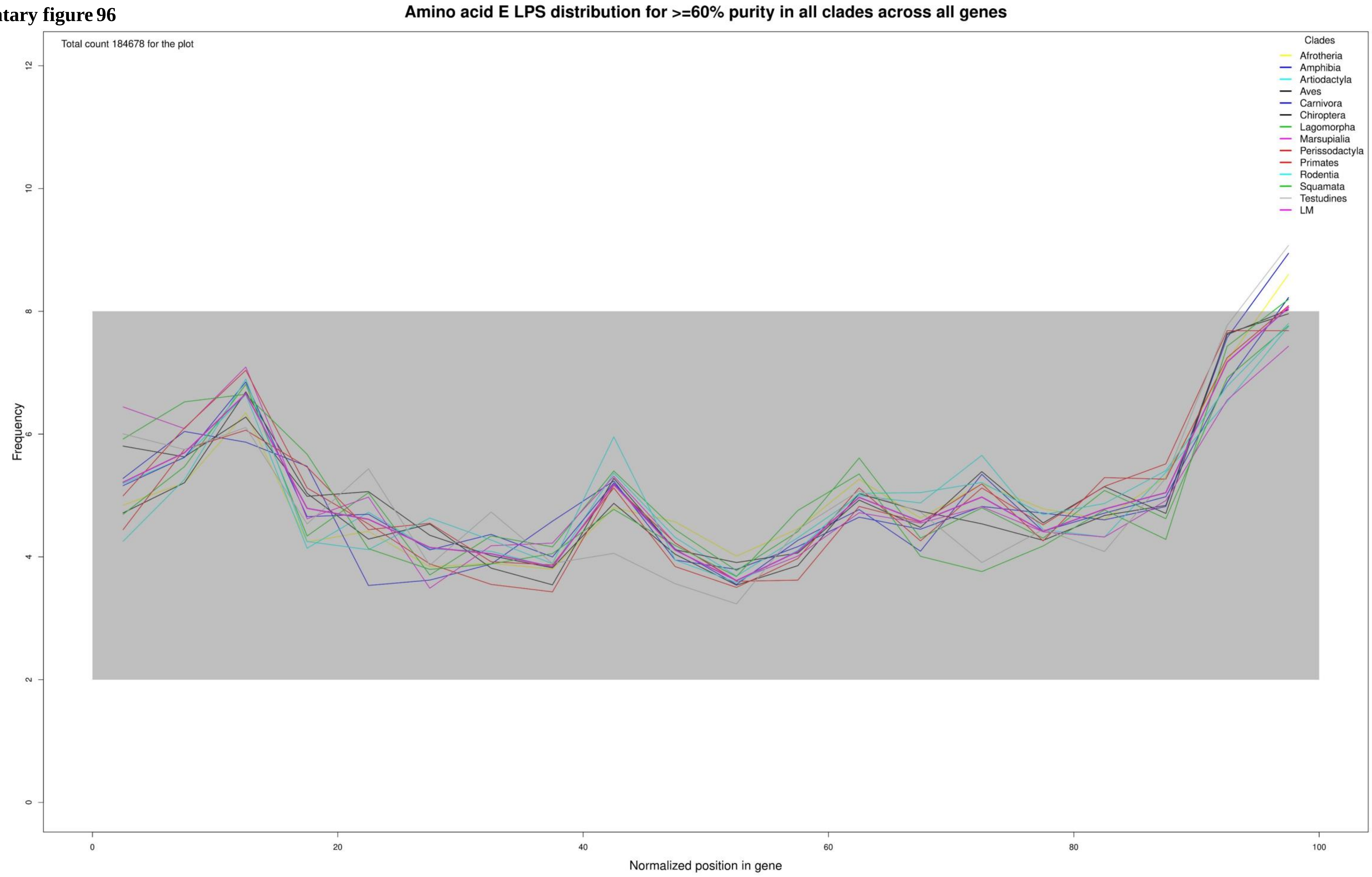

**Supplementary figure 97**

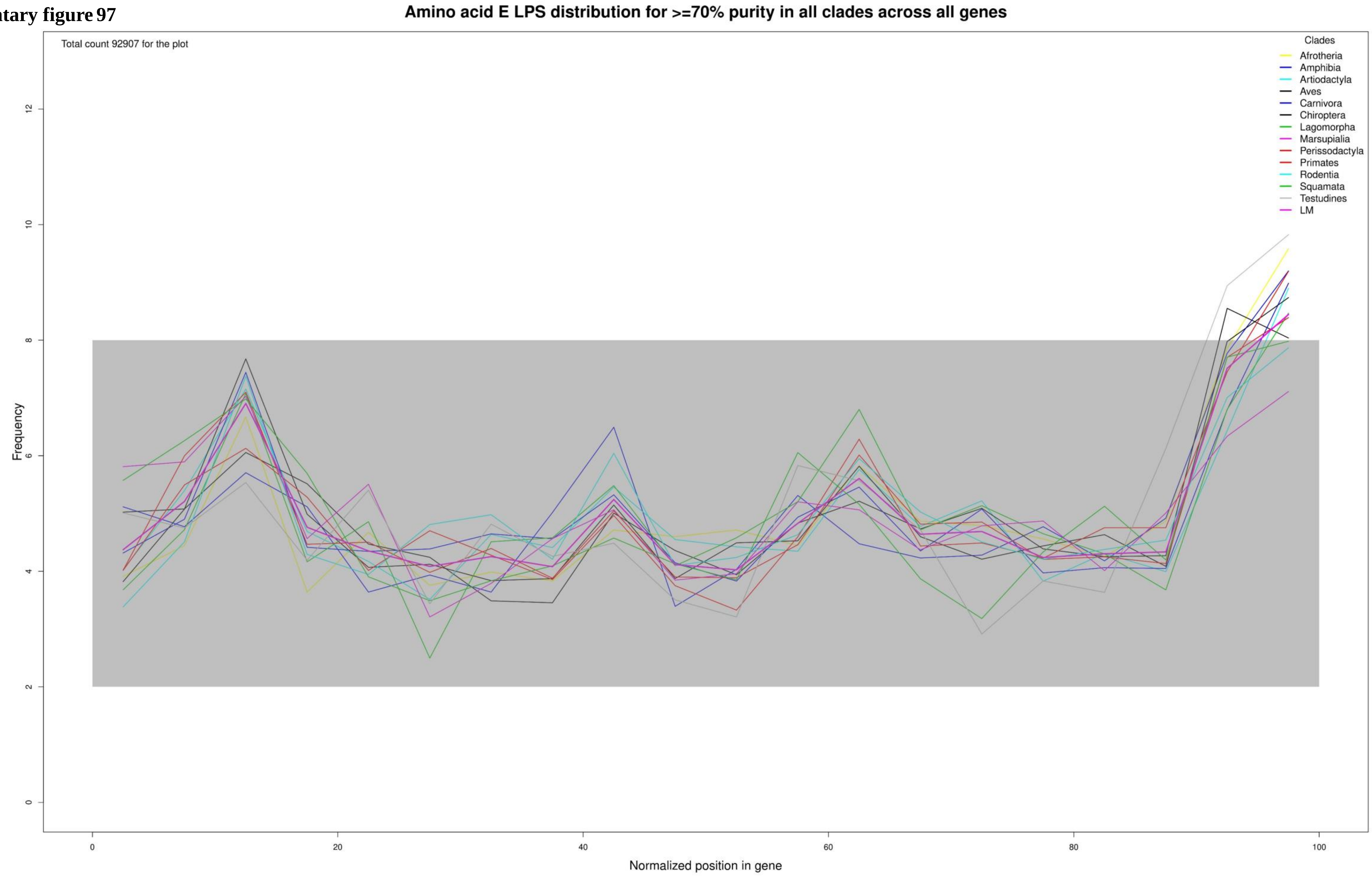

**Supplementary figure 98**

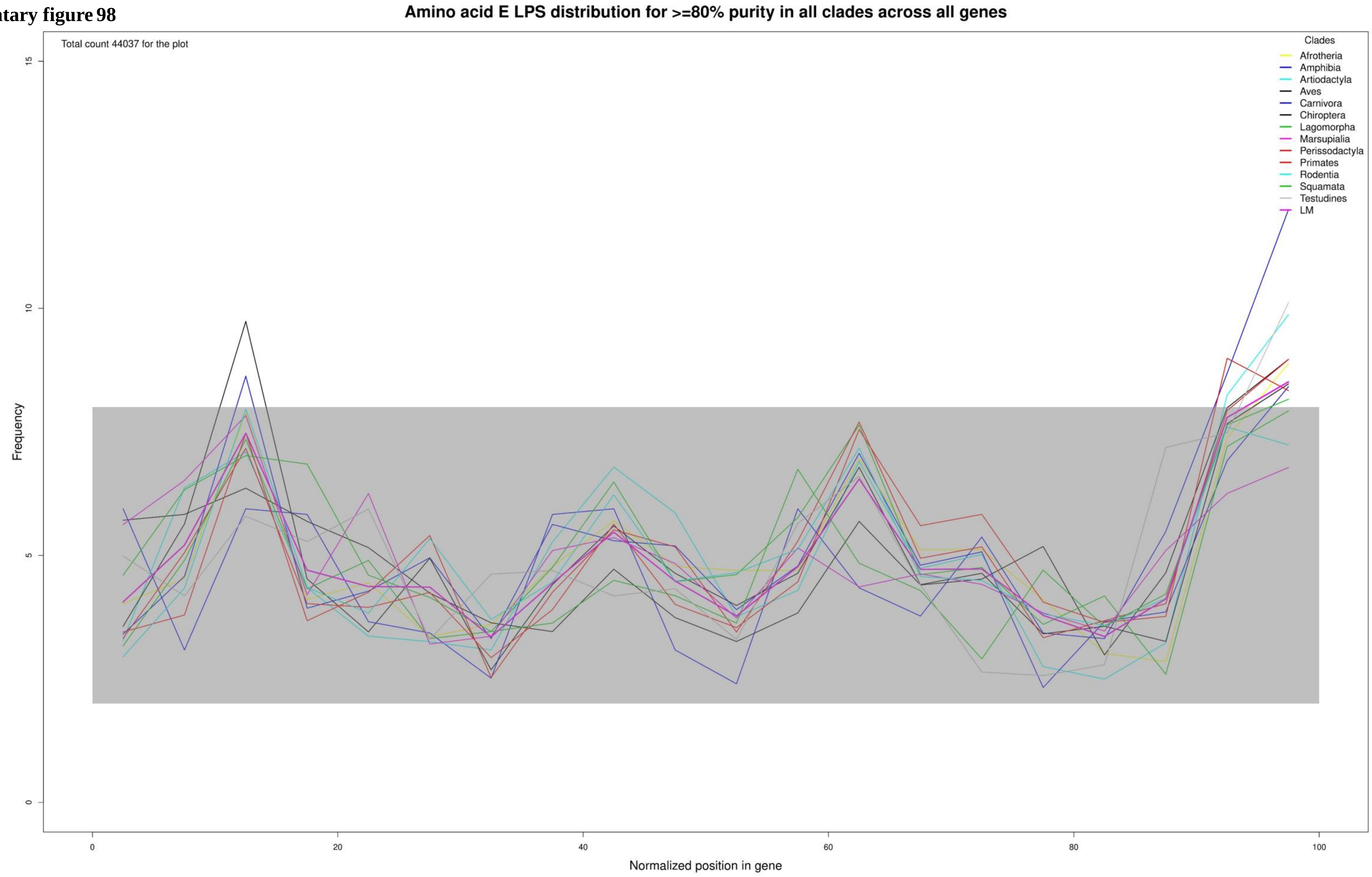

**Supplementary figure 99**

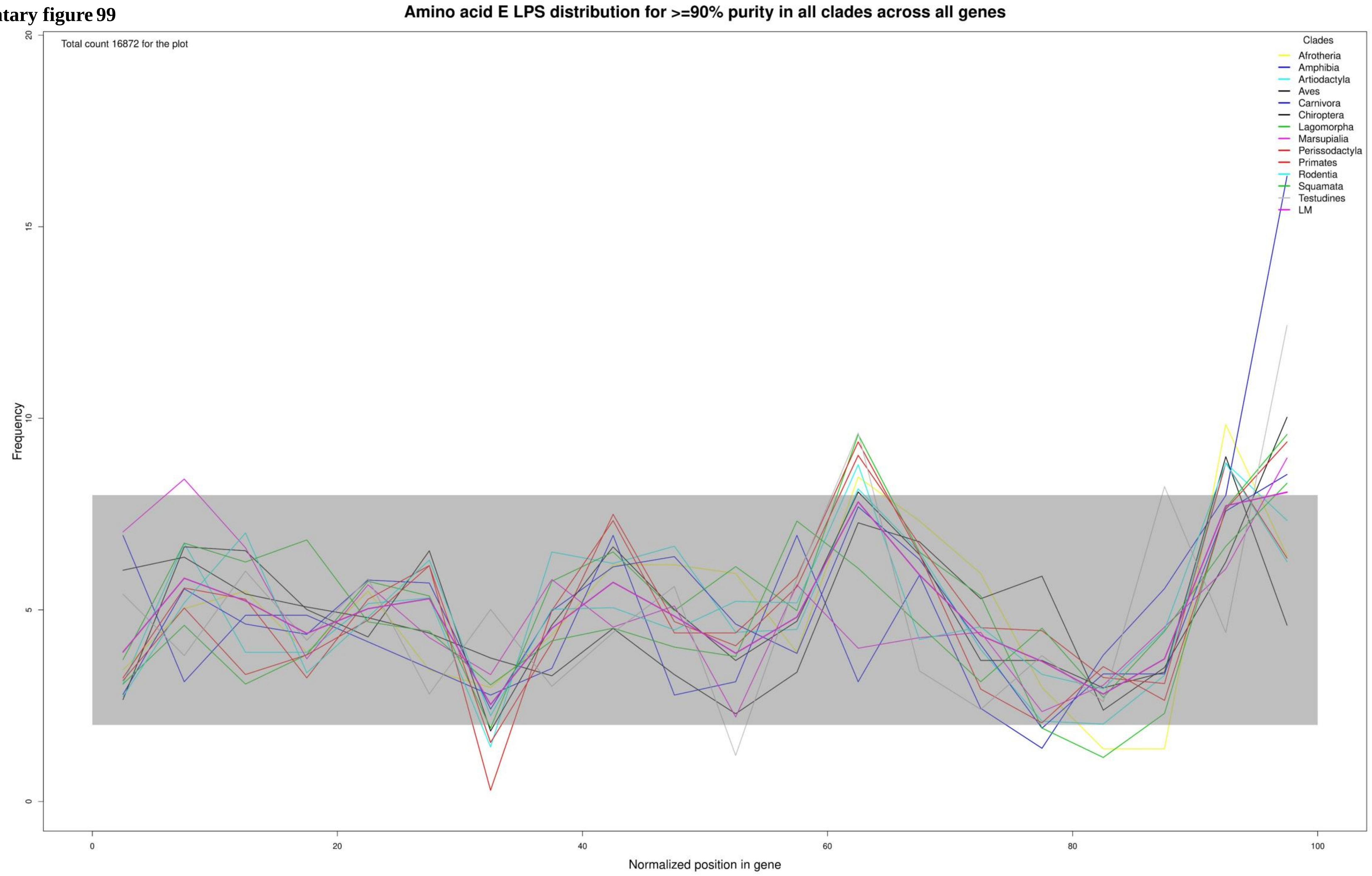

**Supplementary figure 100**

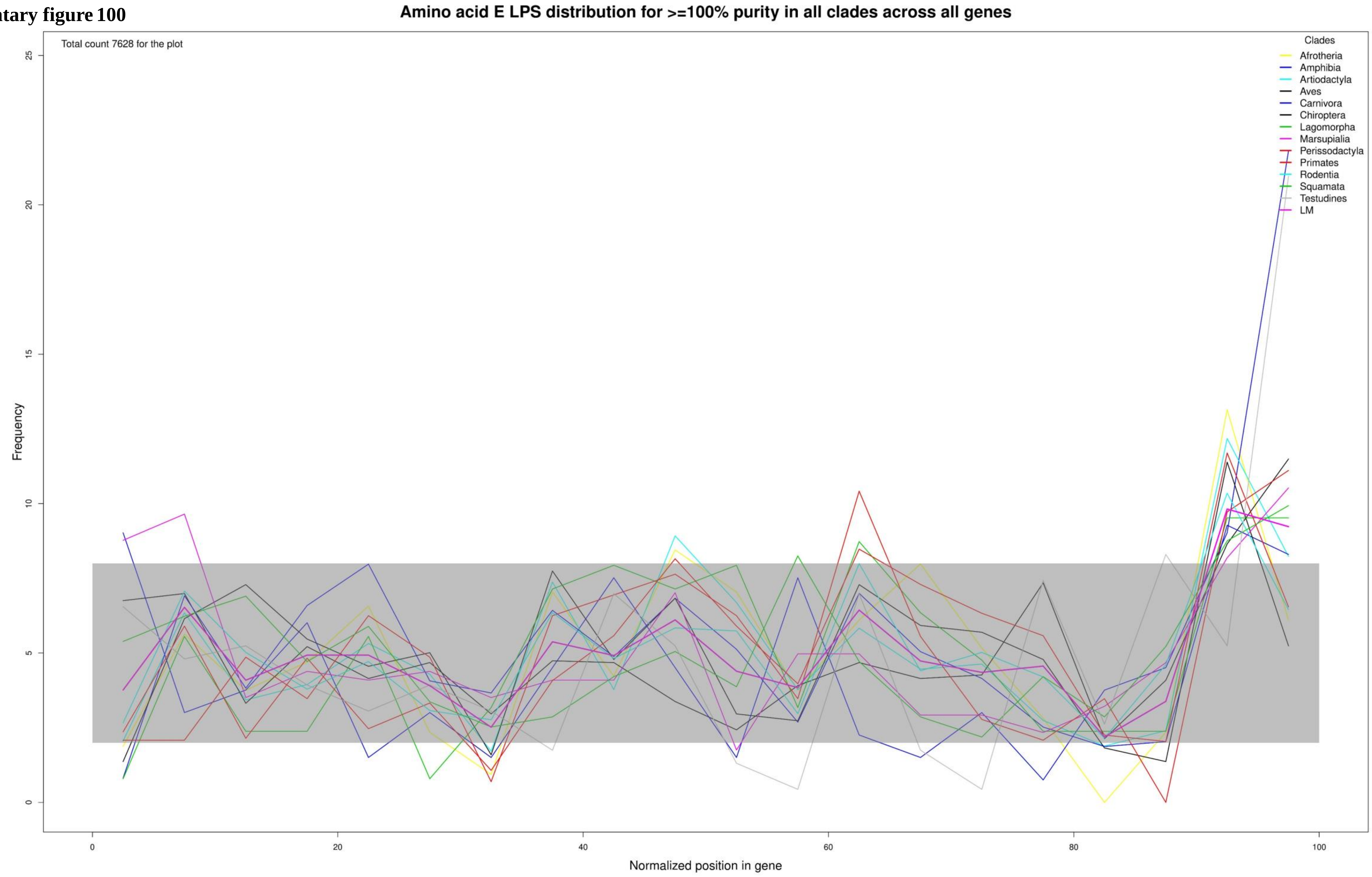

**Supplementary figure 101**

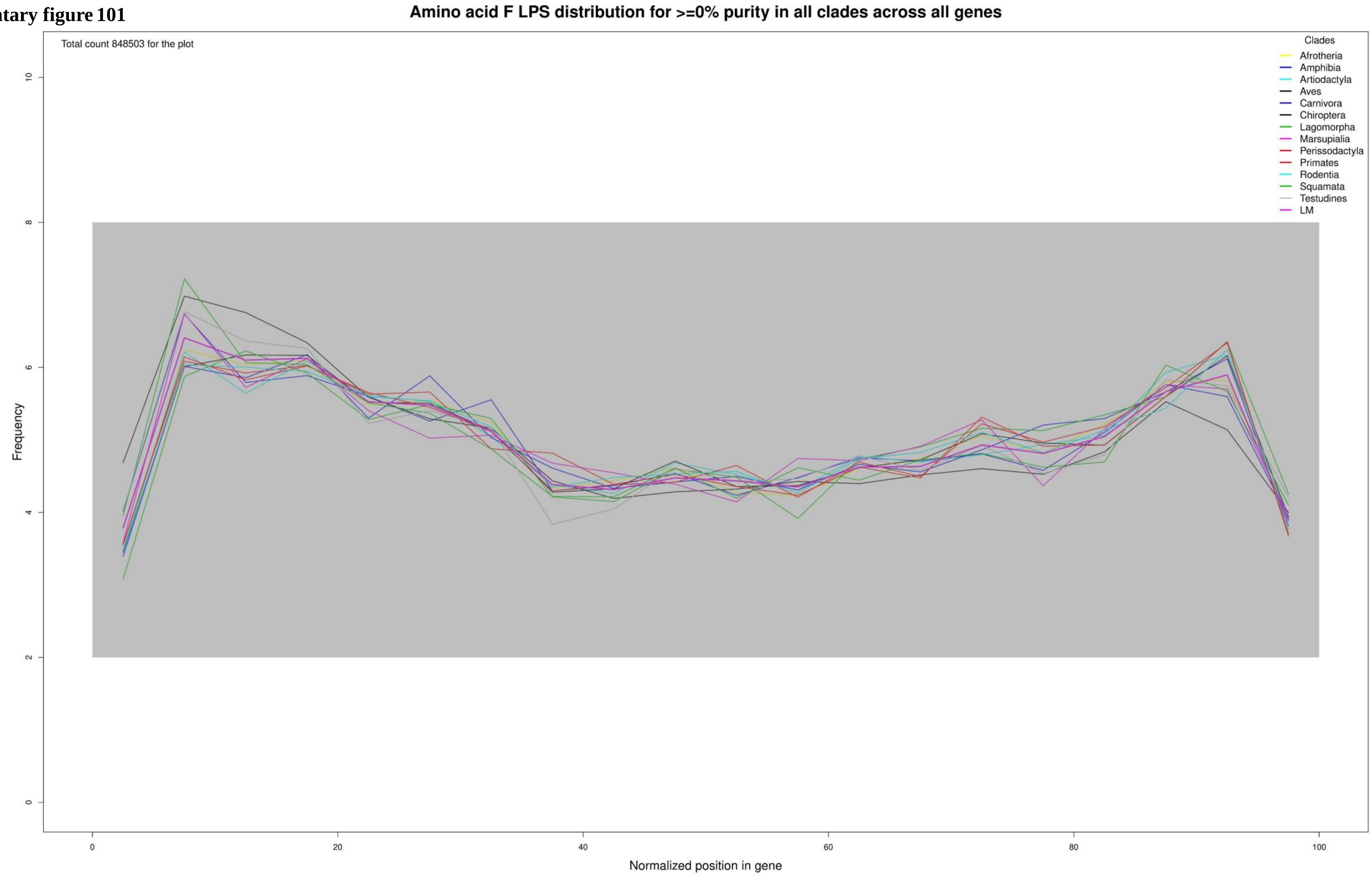

**Supplementary figure 102**

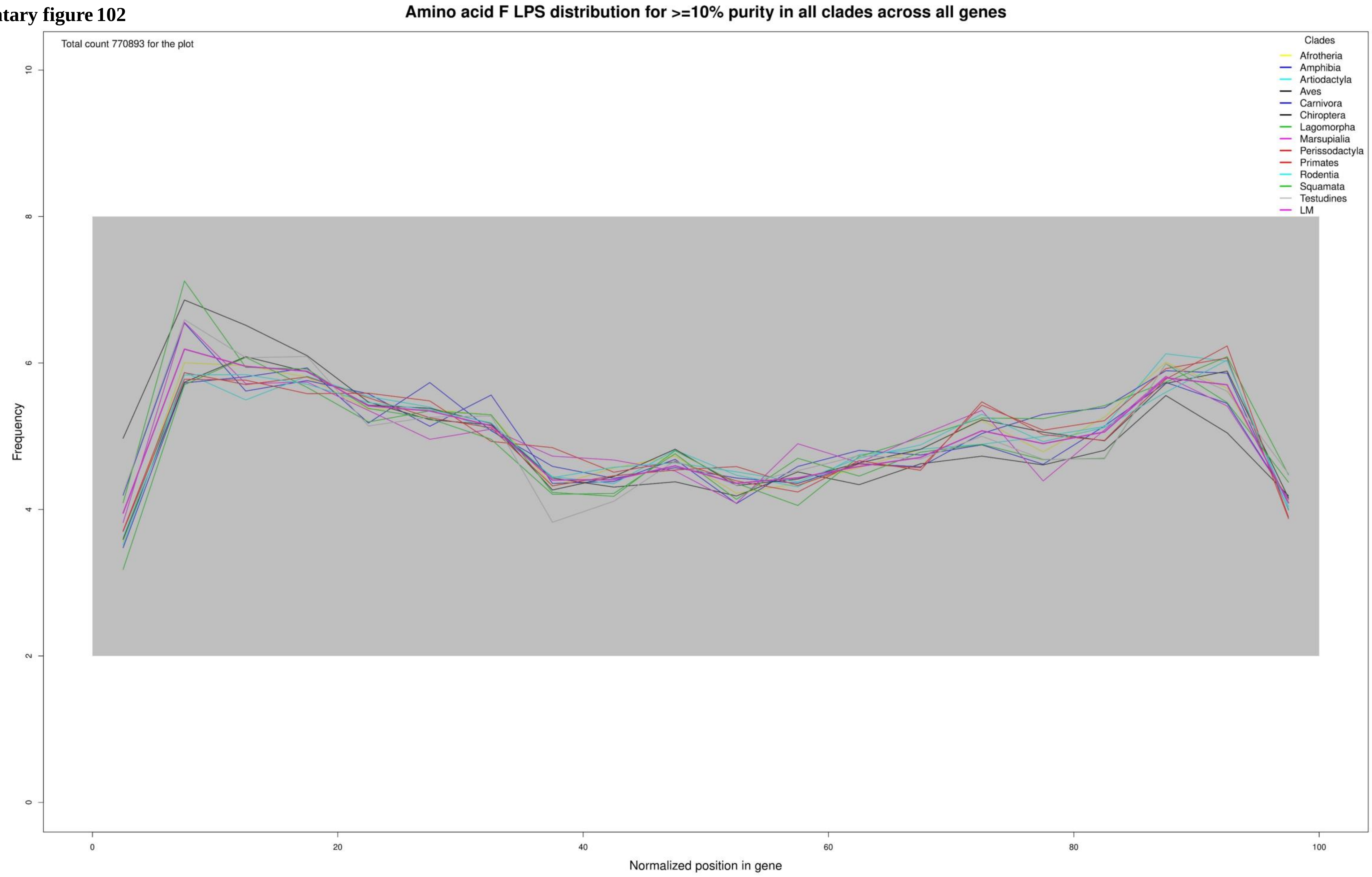

**Supplementary figure 103**

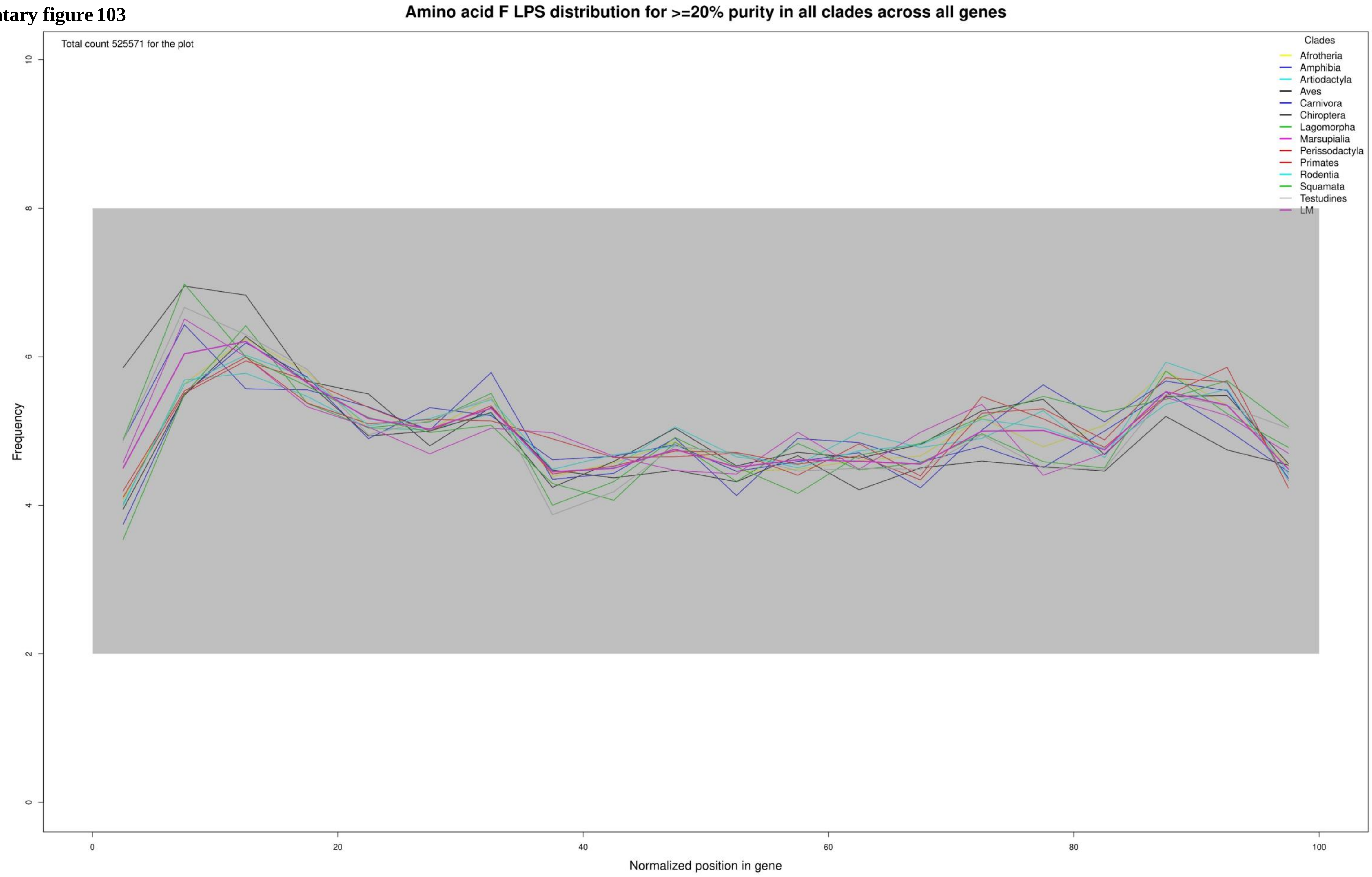

**Supplementary figure 104**

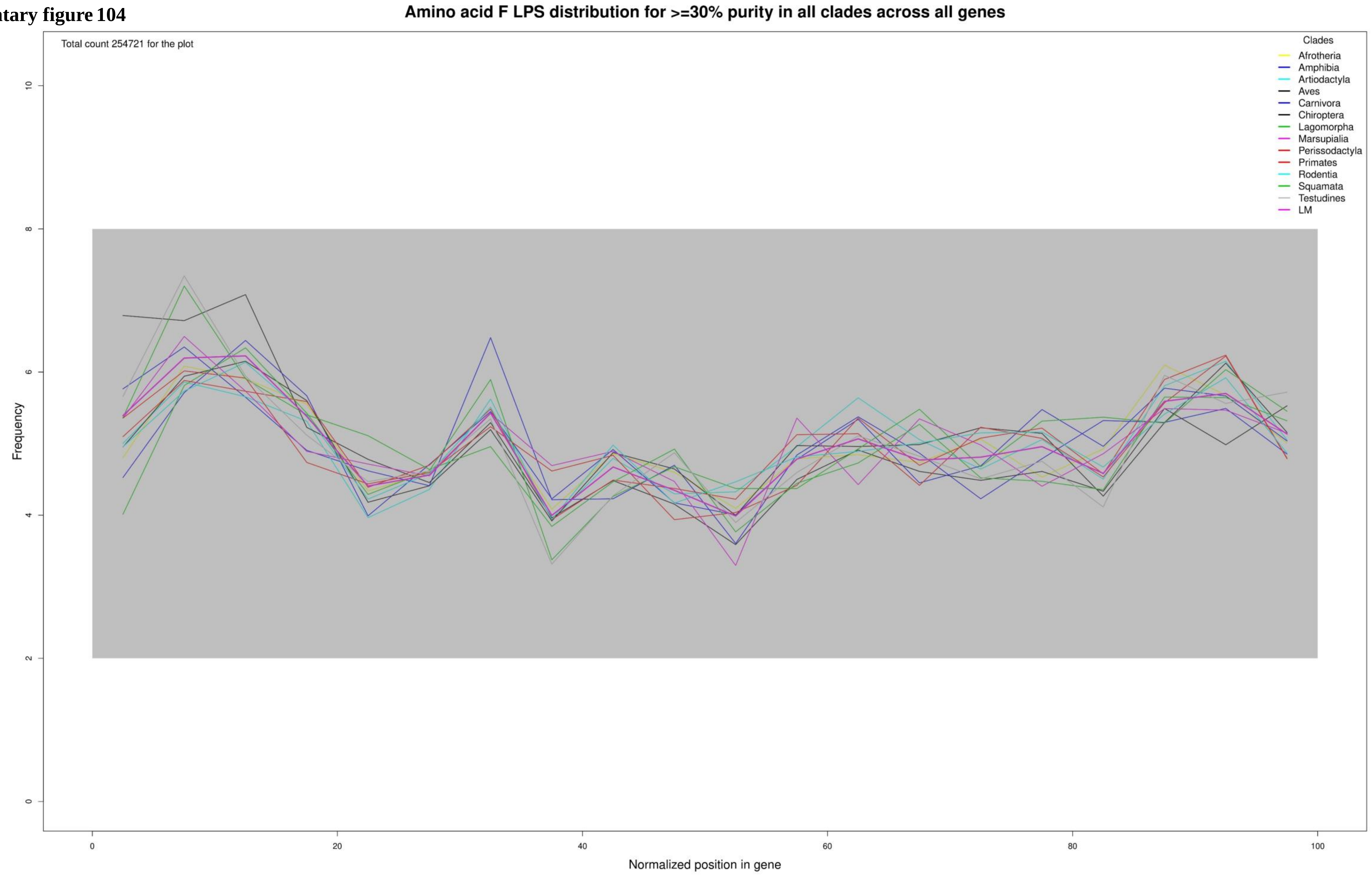

**Supplementary figure 105**

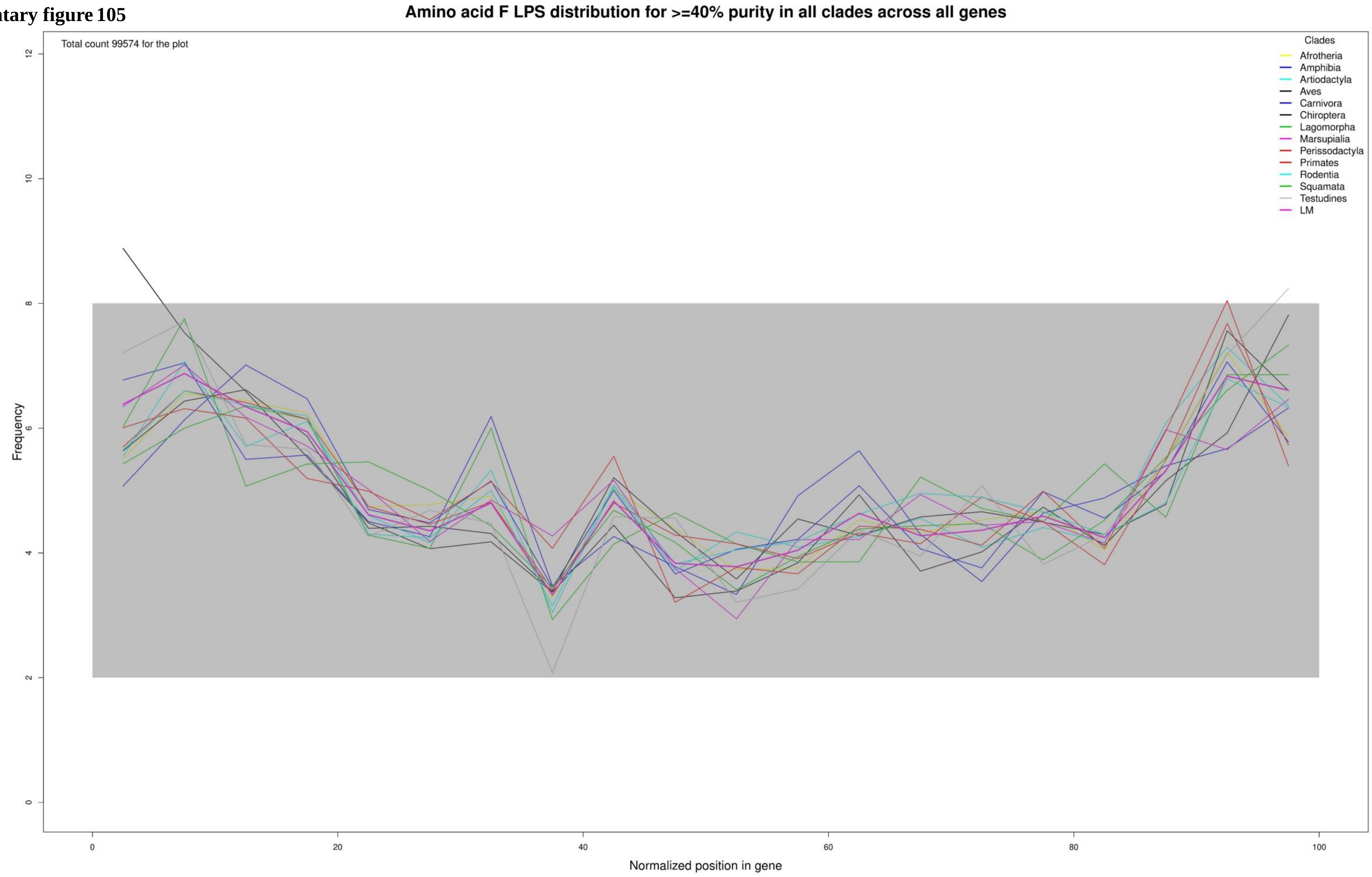

**Supplementary figure 106**

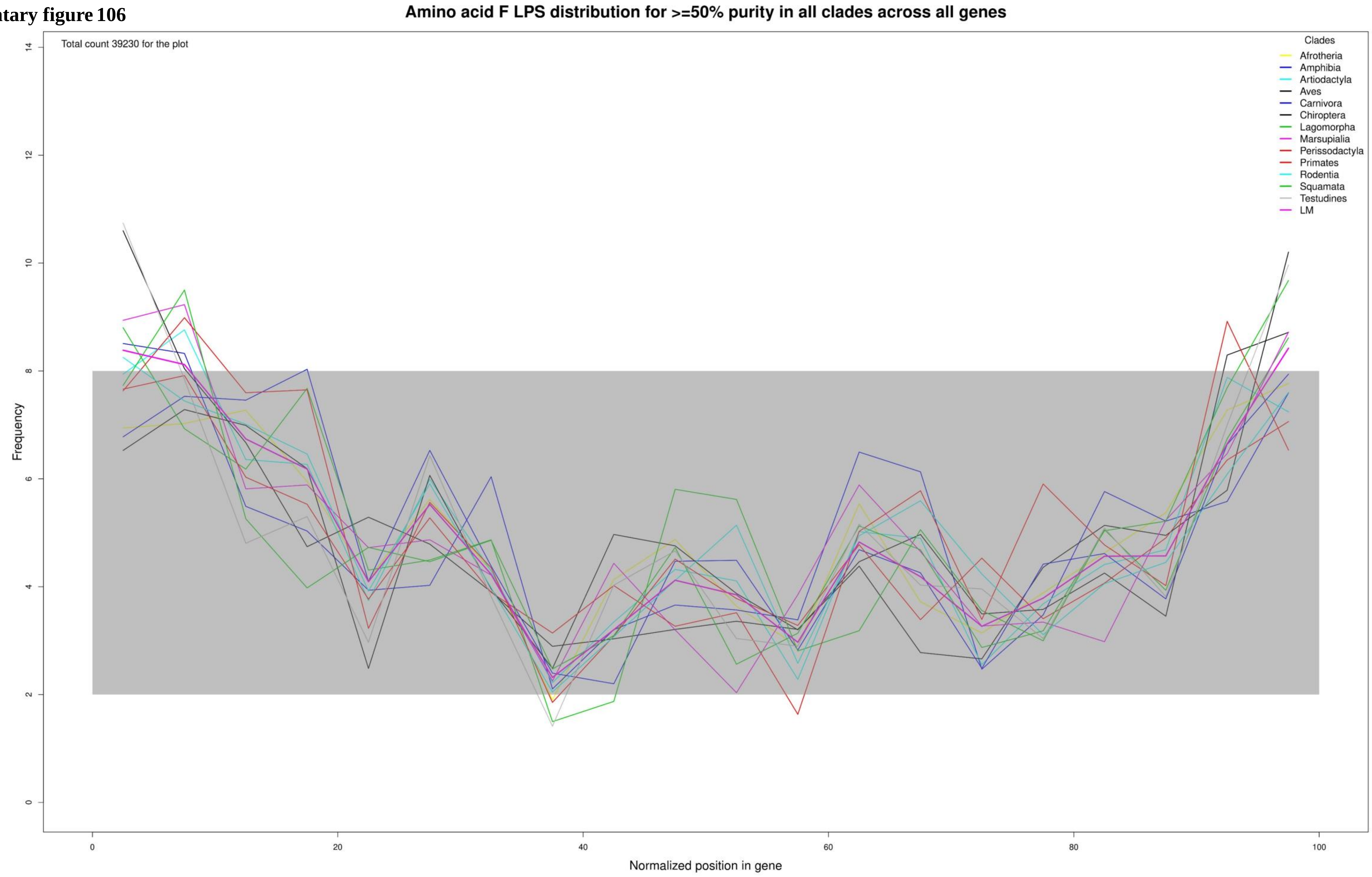

Supplementary figure 107

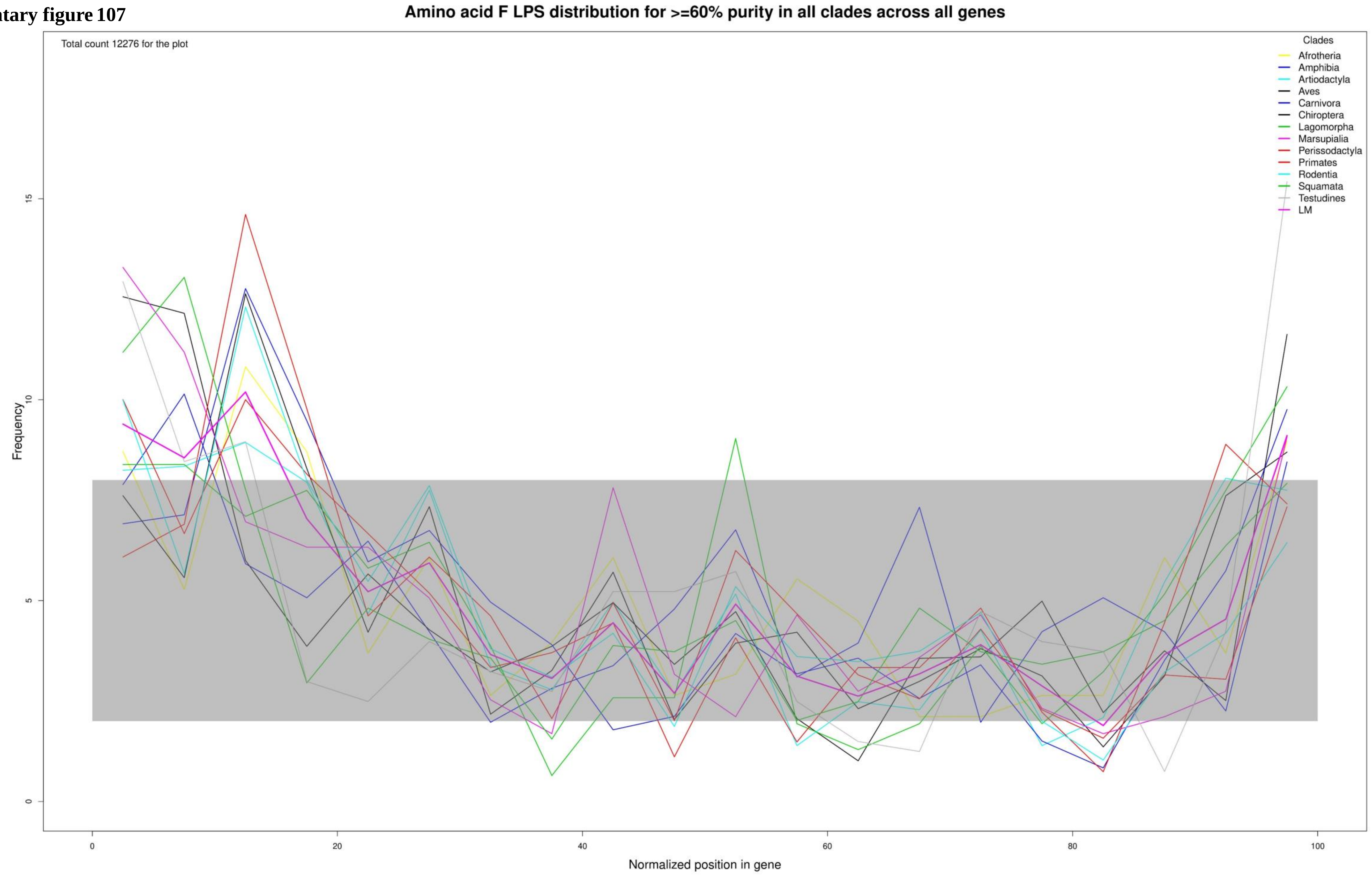

**Supplementary figure 108**

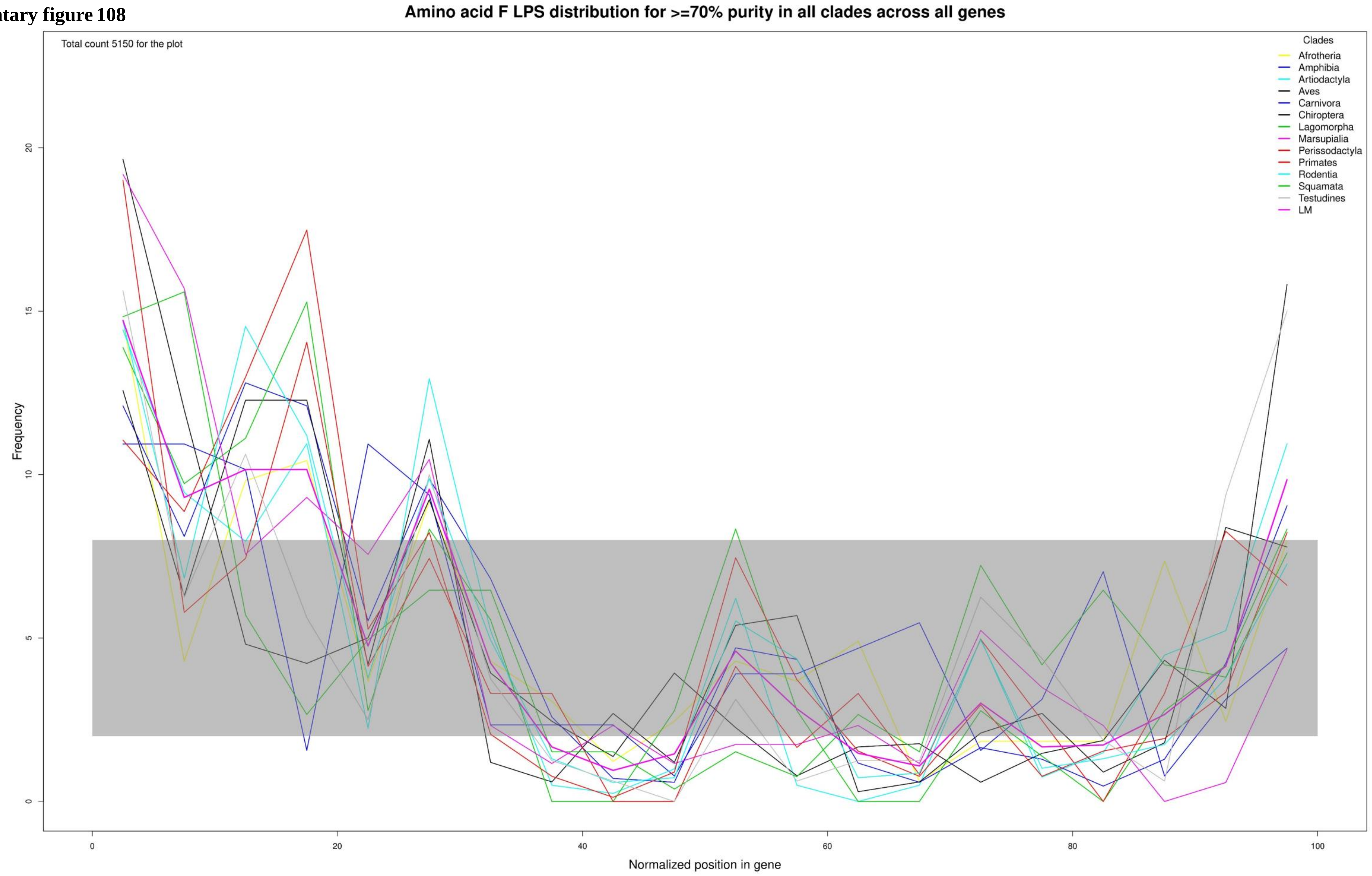

**Supplementary figure 109**

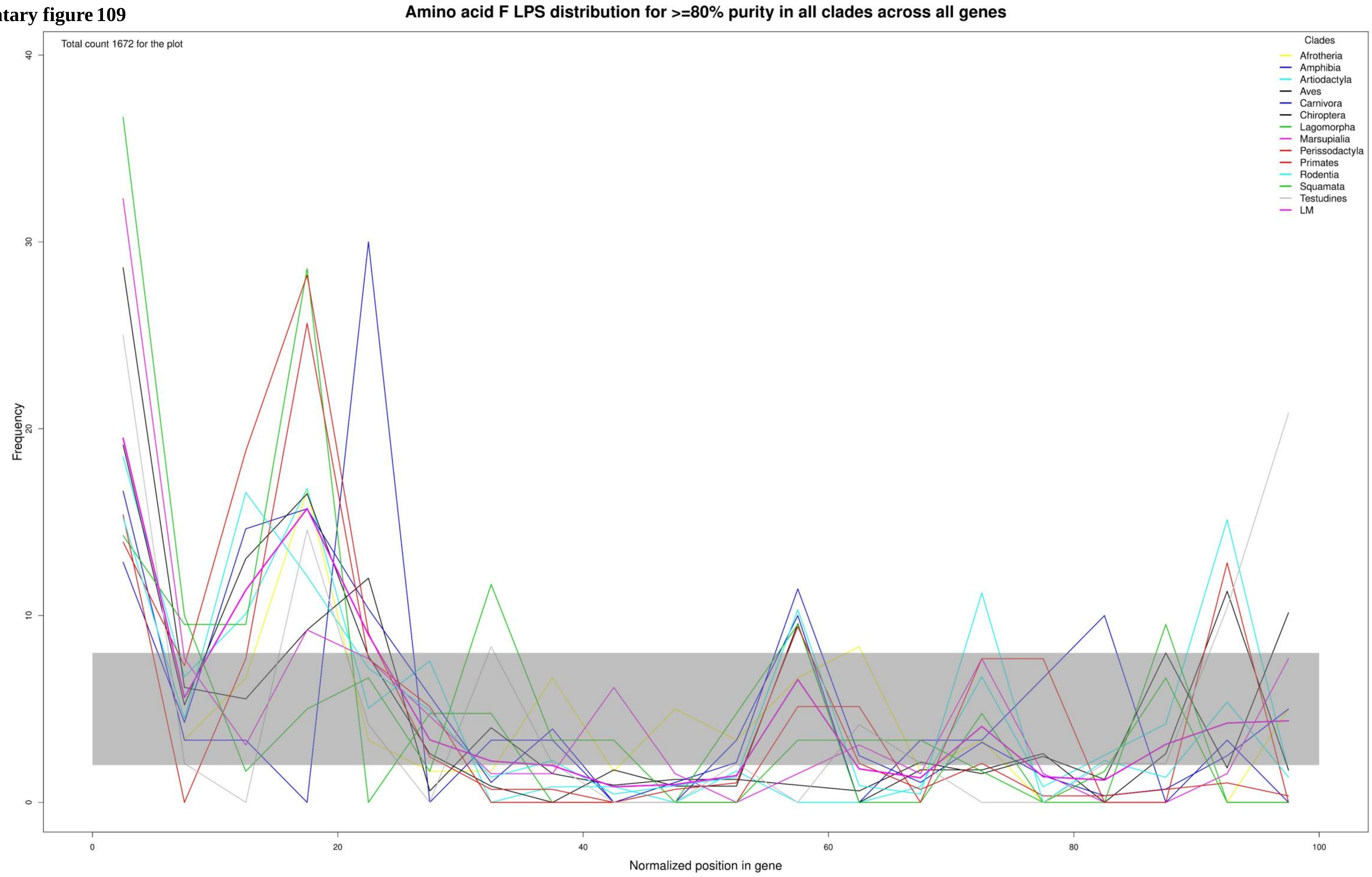

**Supplementary figure 110**

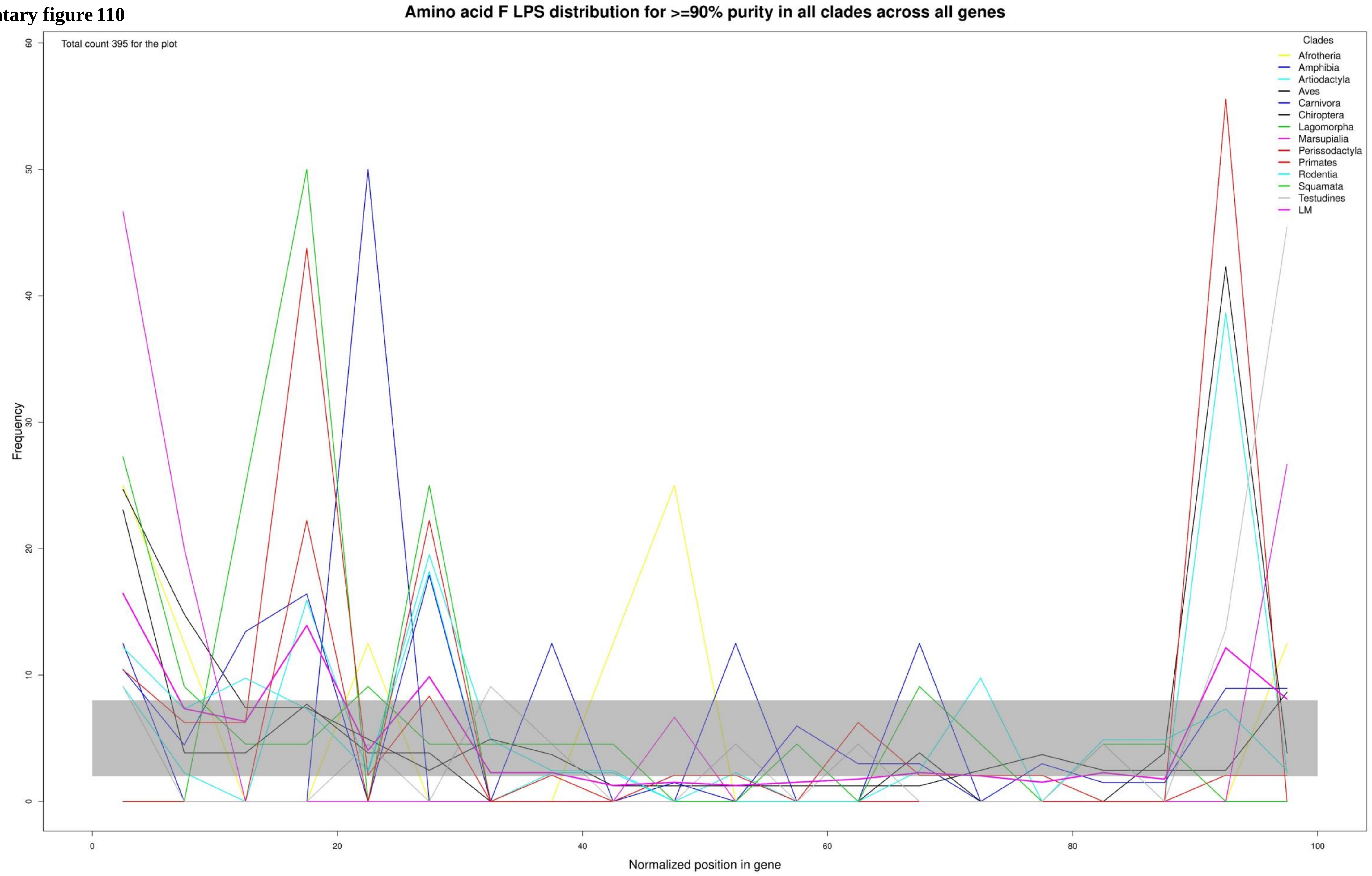

**Supplementary figure 111**

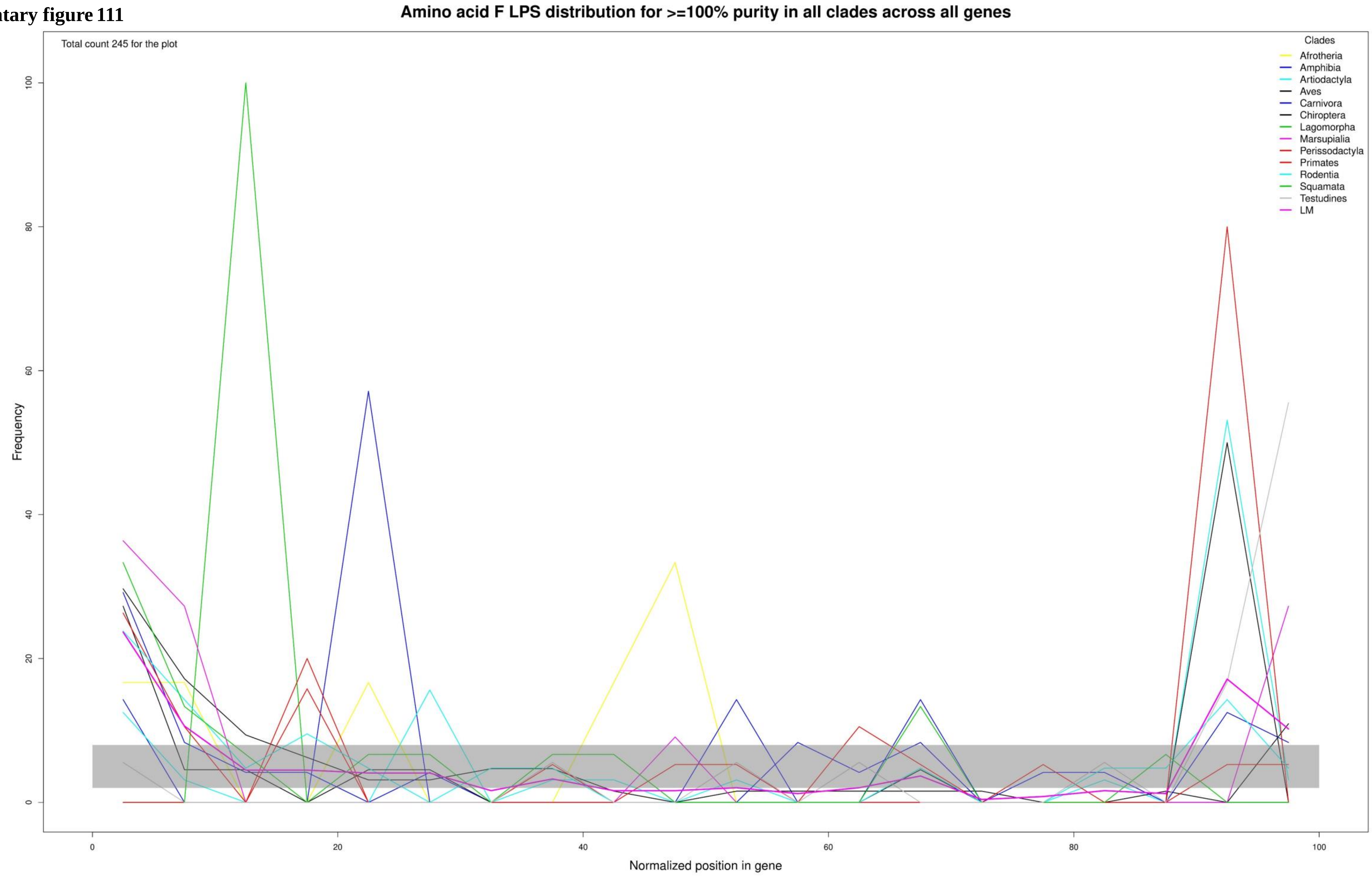

Supplementary figure 112

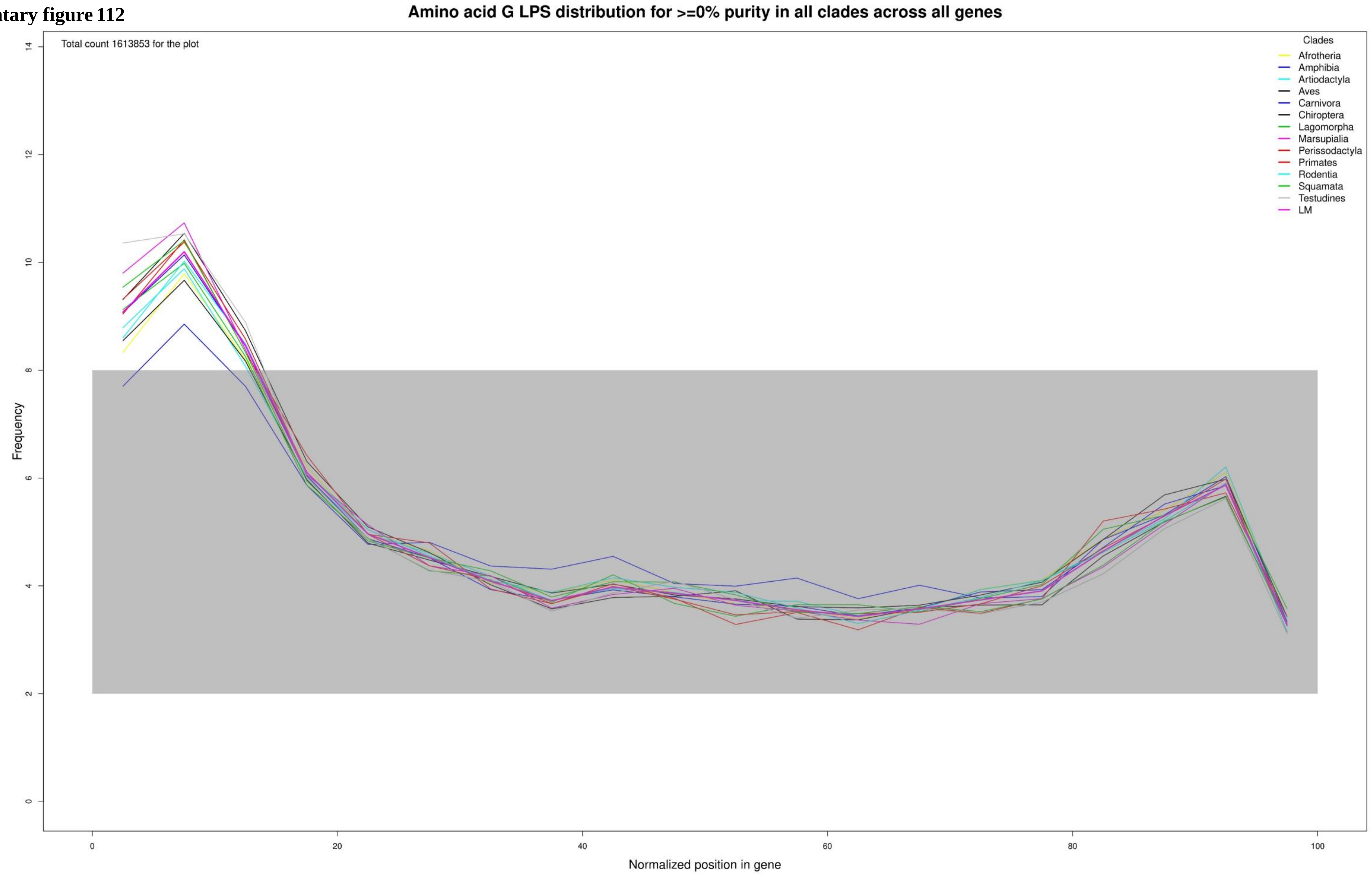

**Supplementary figure 113**

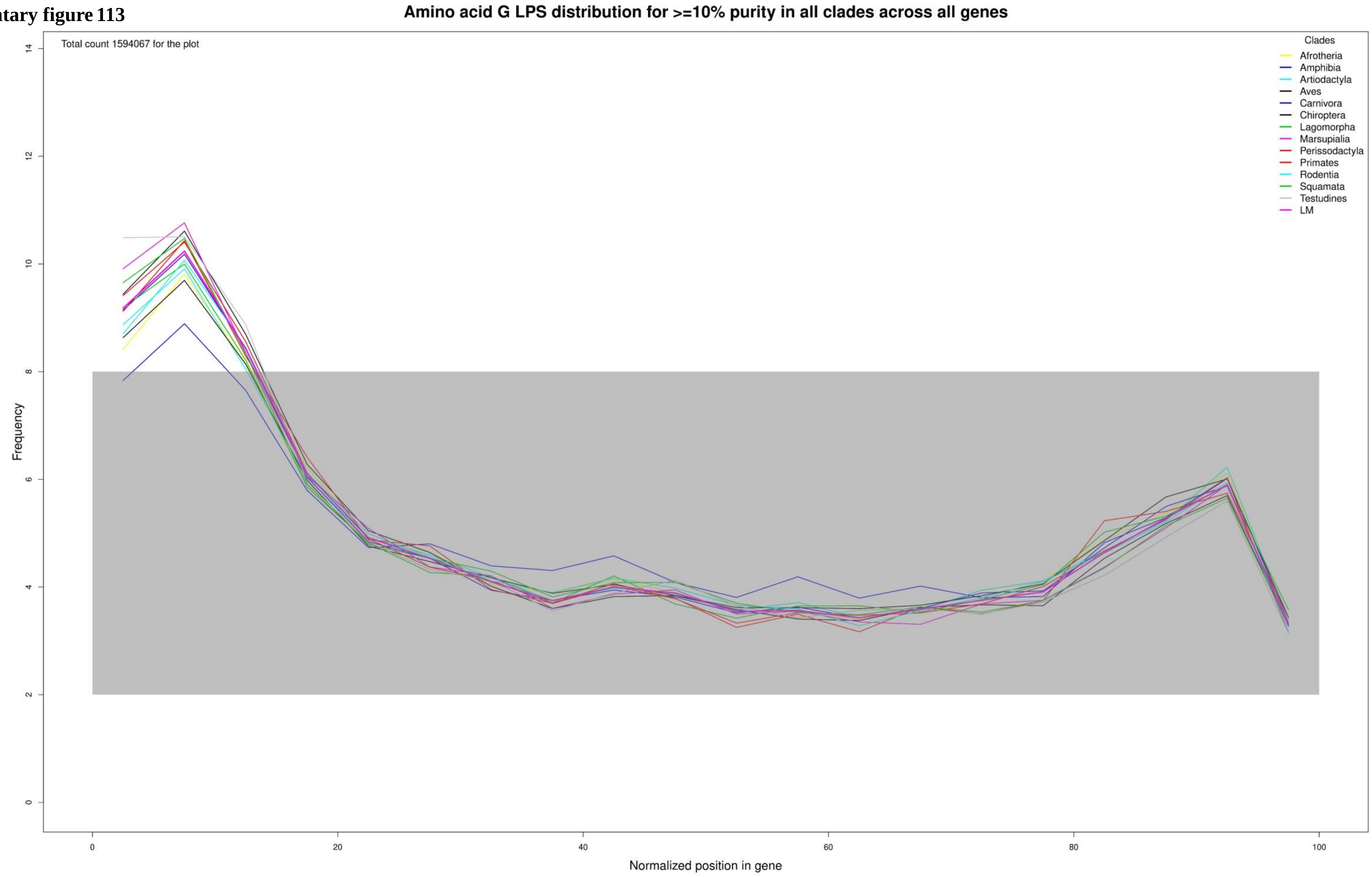

**Supplementary figure 114**

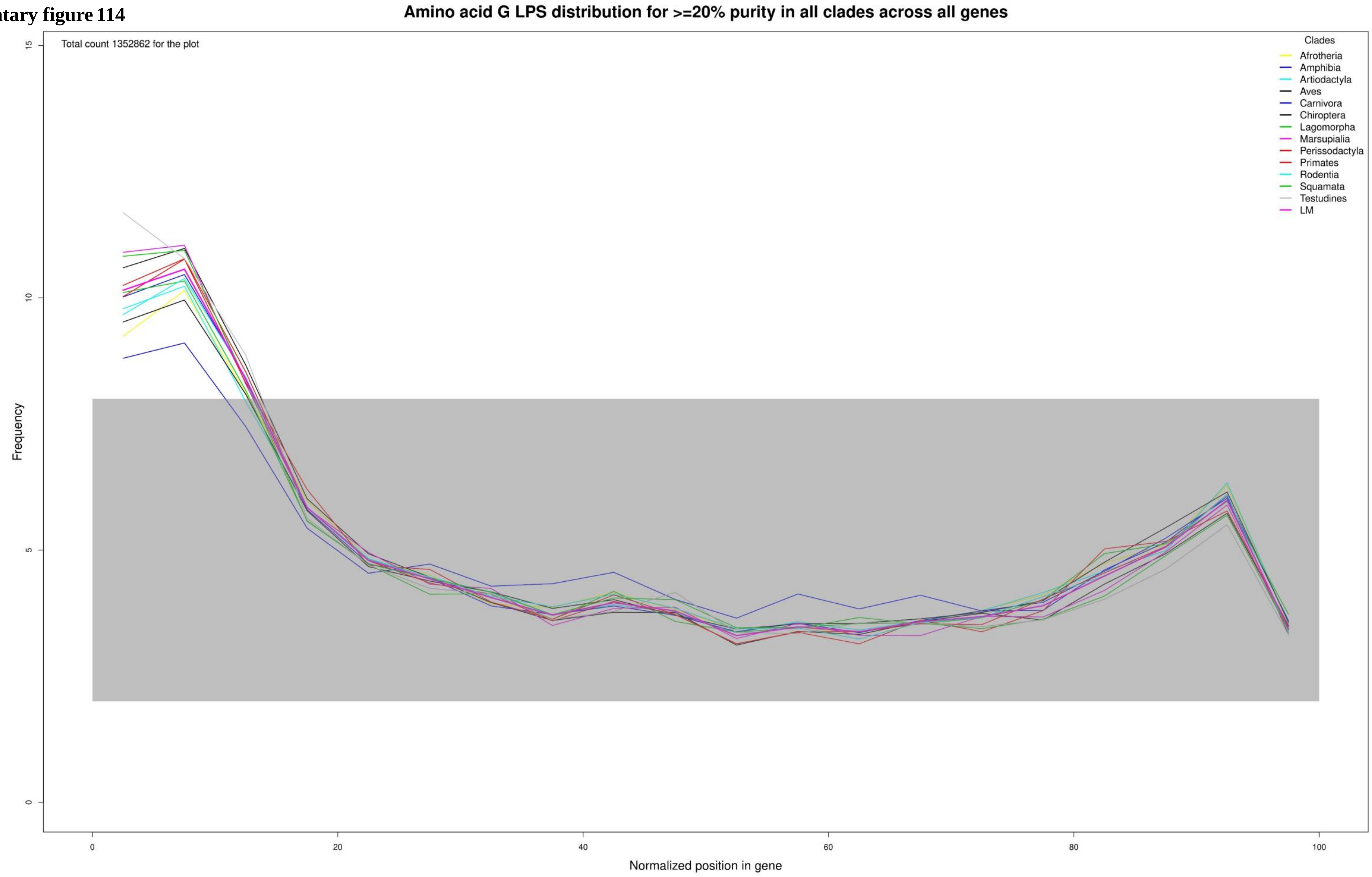

**Supplementary figure 115**

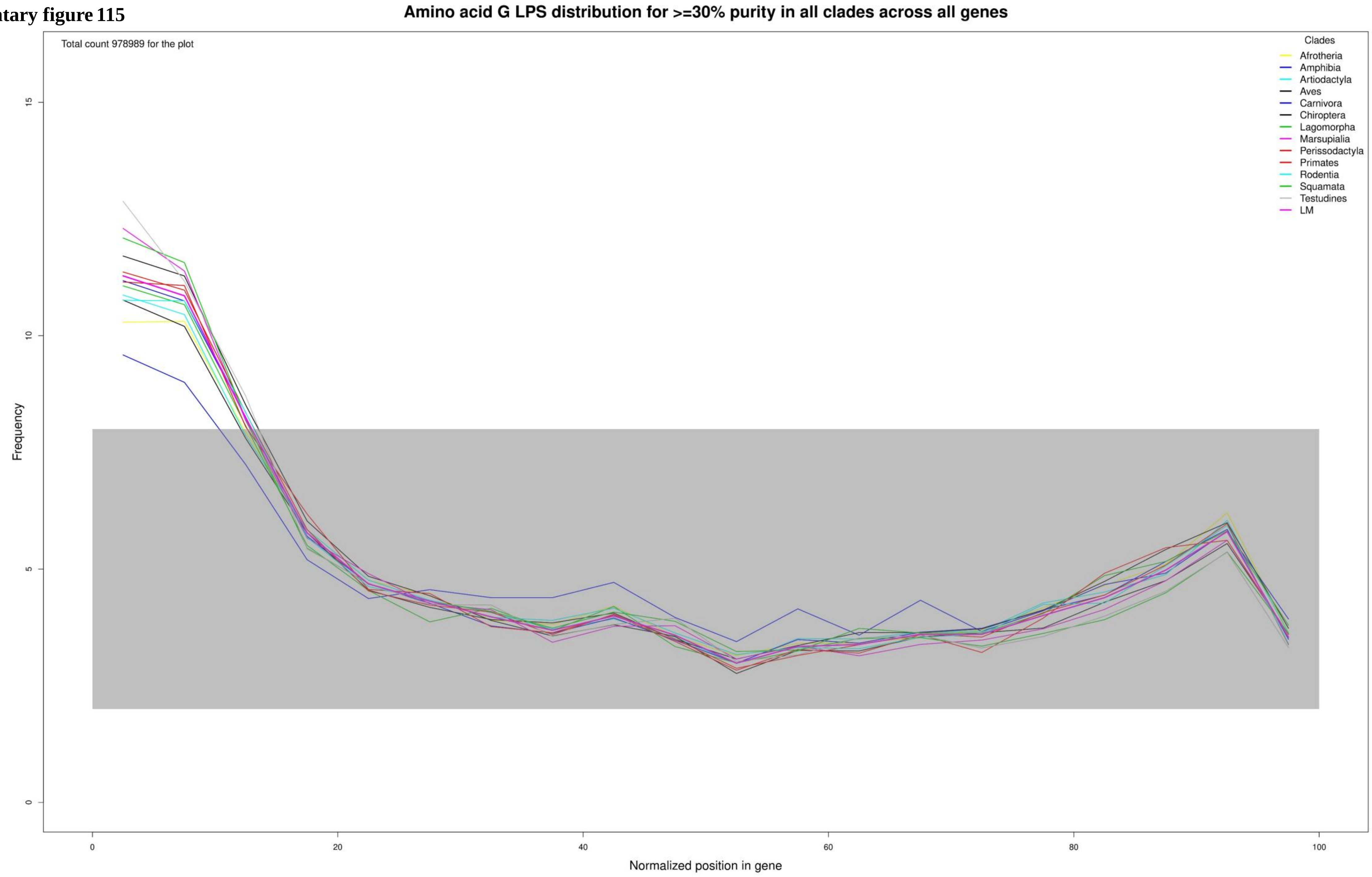

**Supplementary figure 116**

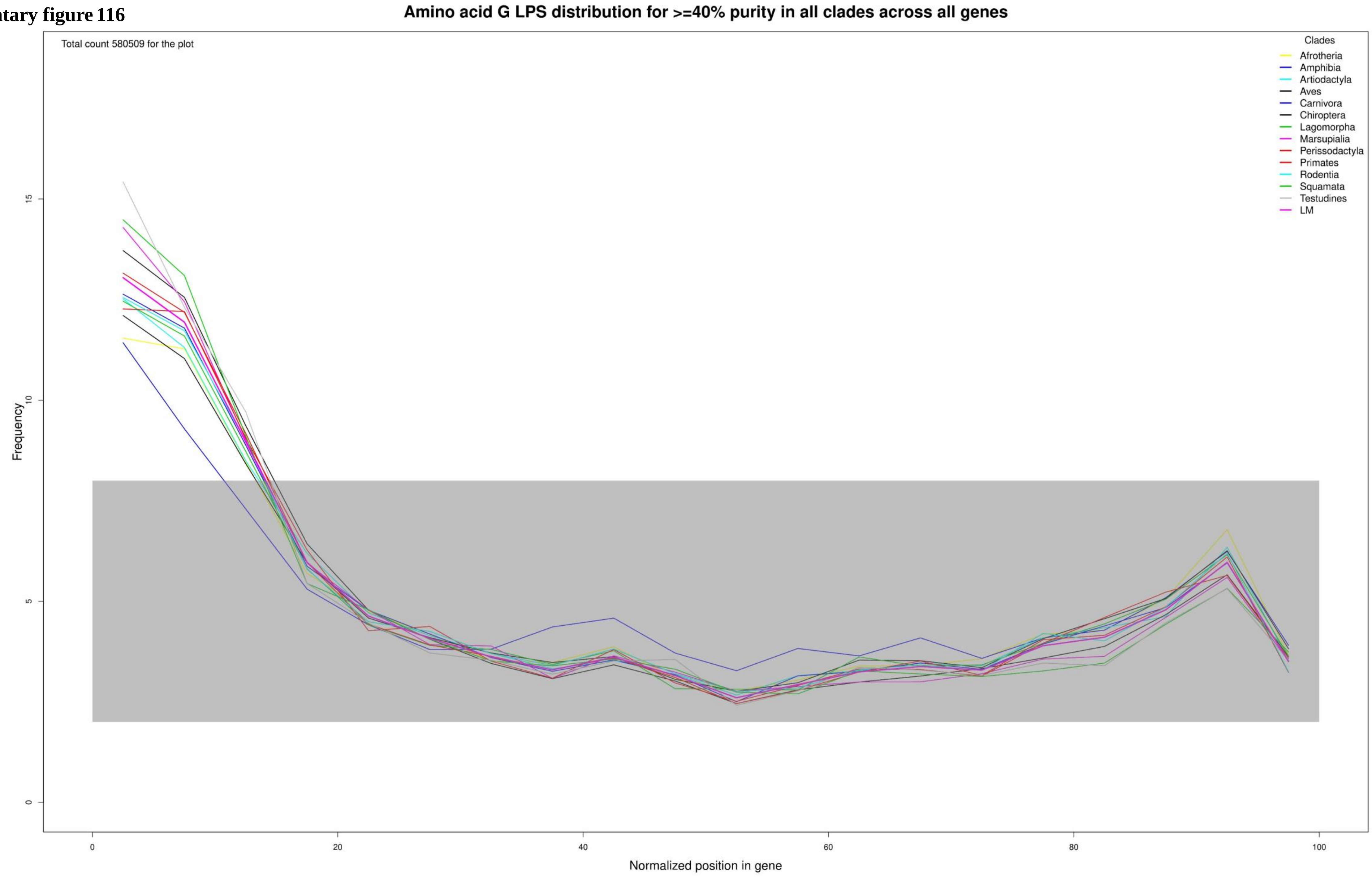

**Supplementary figure 117**

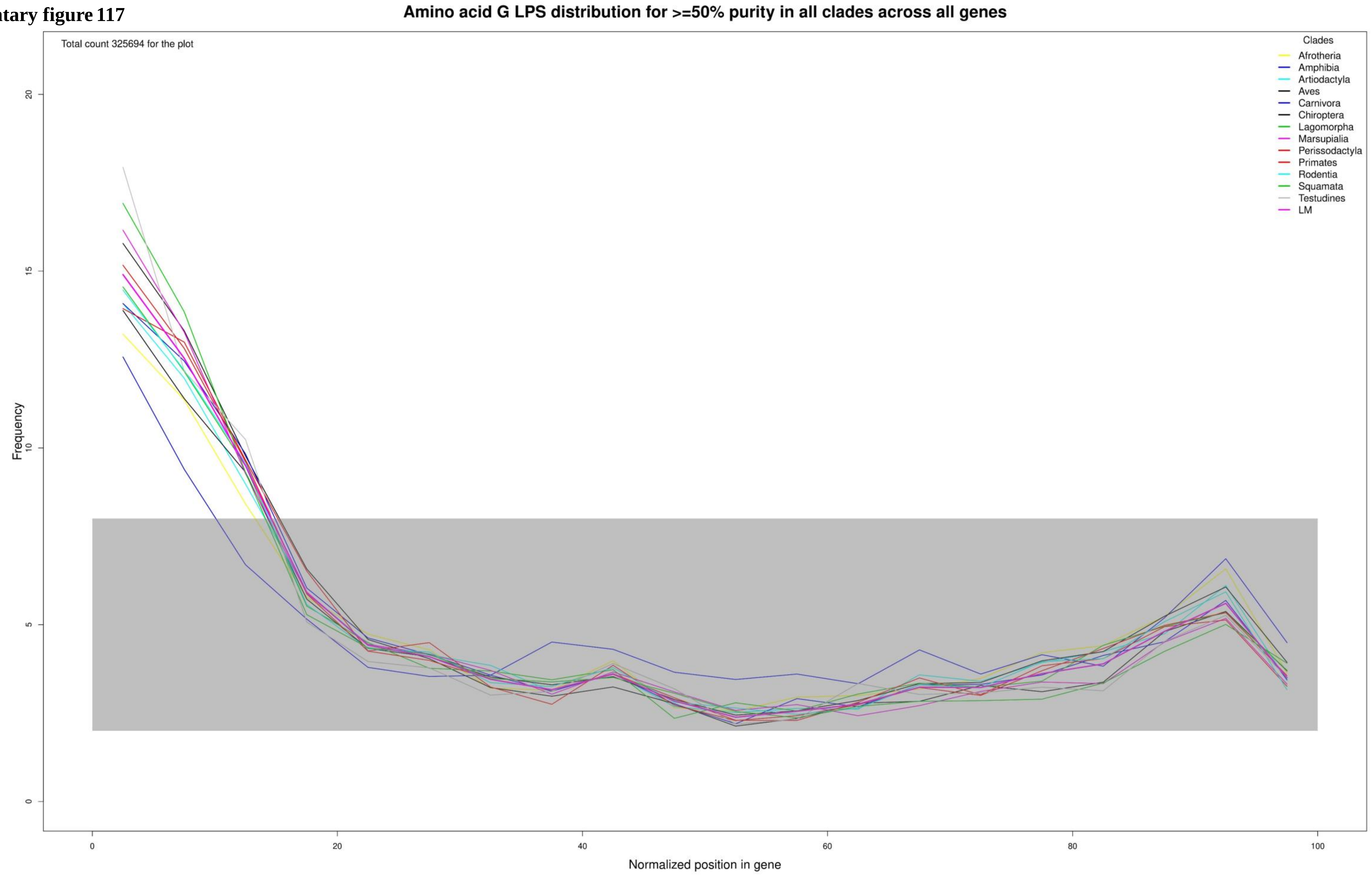

**Supplementary figure 118**

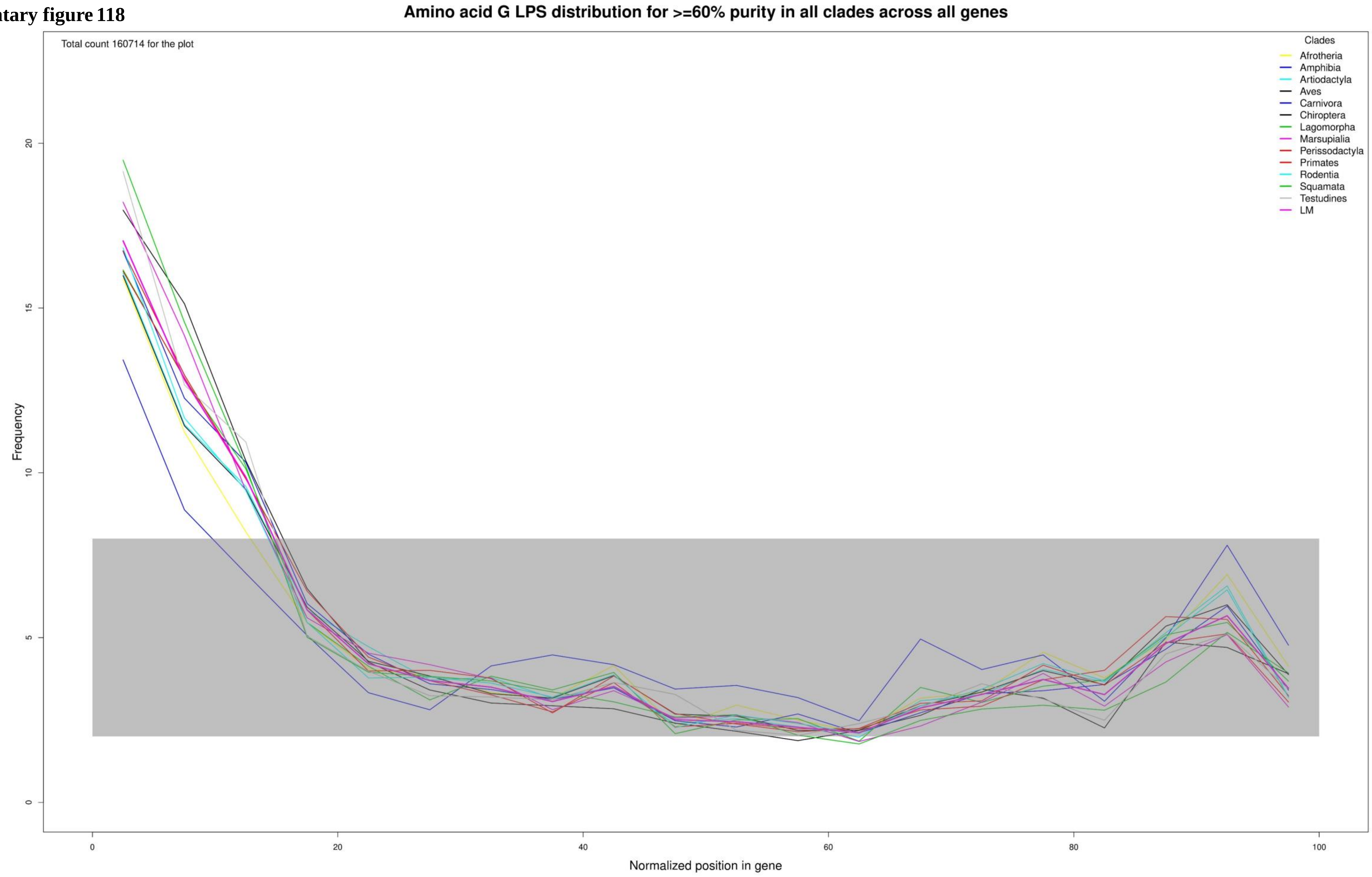

**Supplementary figure 119**

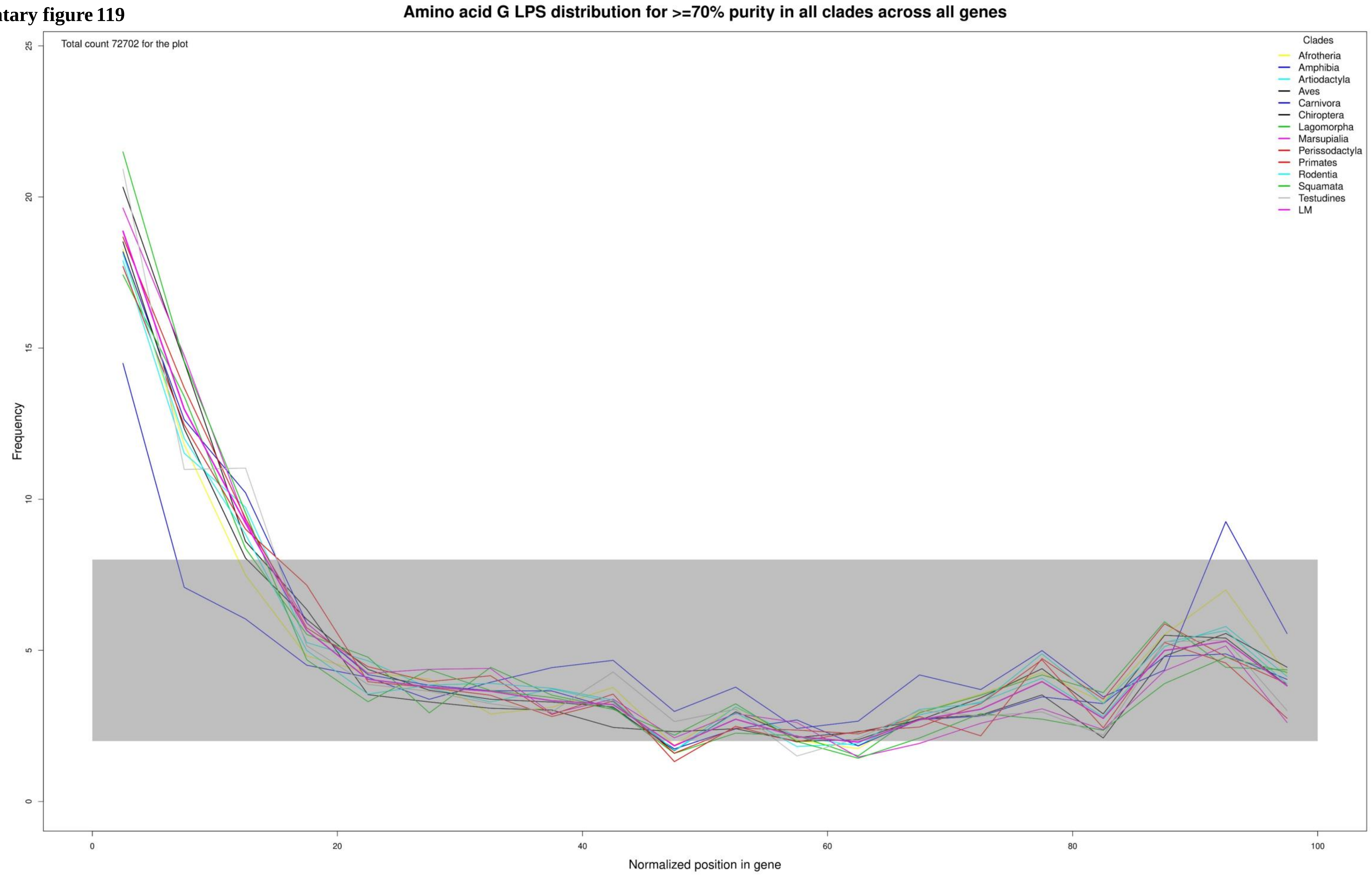

Supplementary figure 120

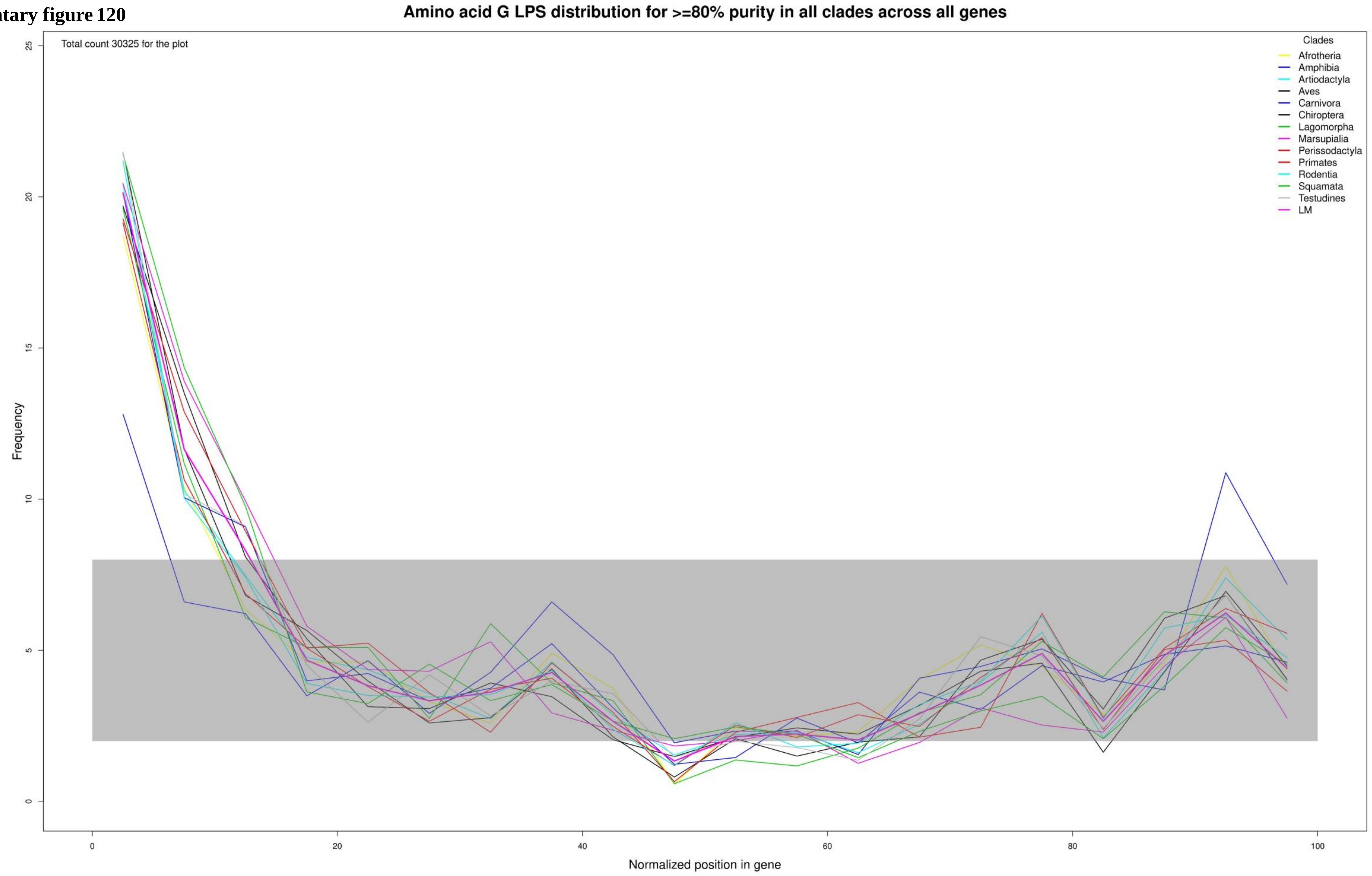

**Supplementary figure 121**

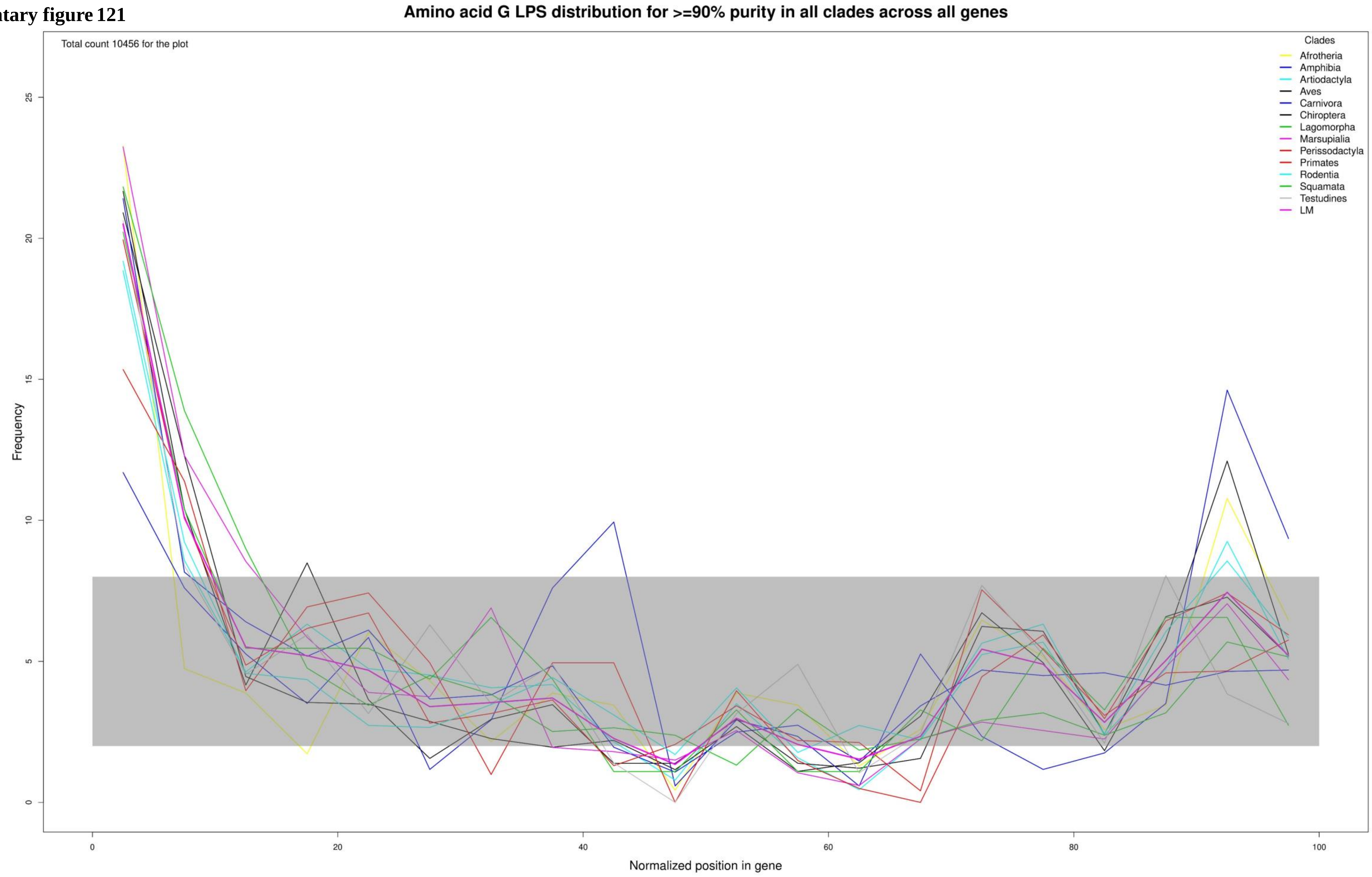

**Supplementary figure 122**

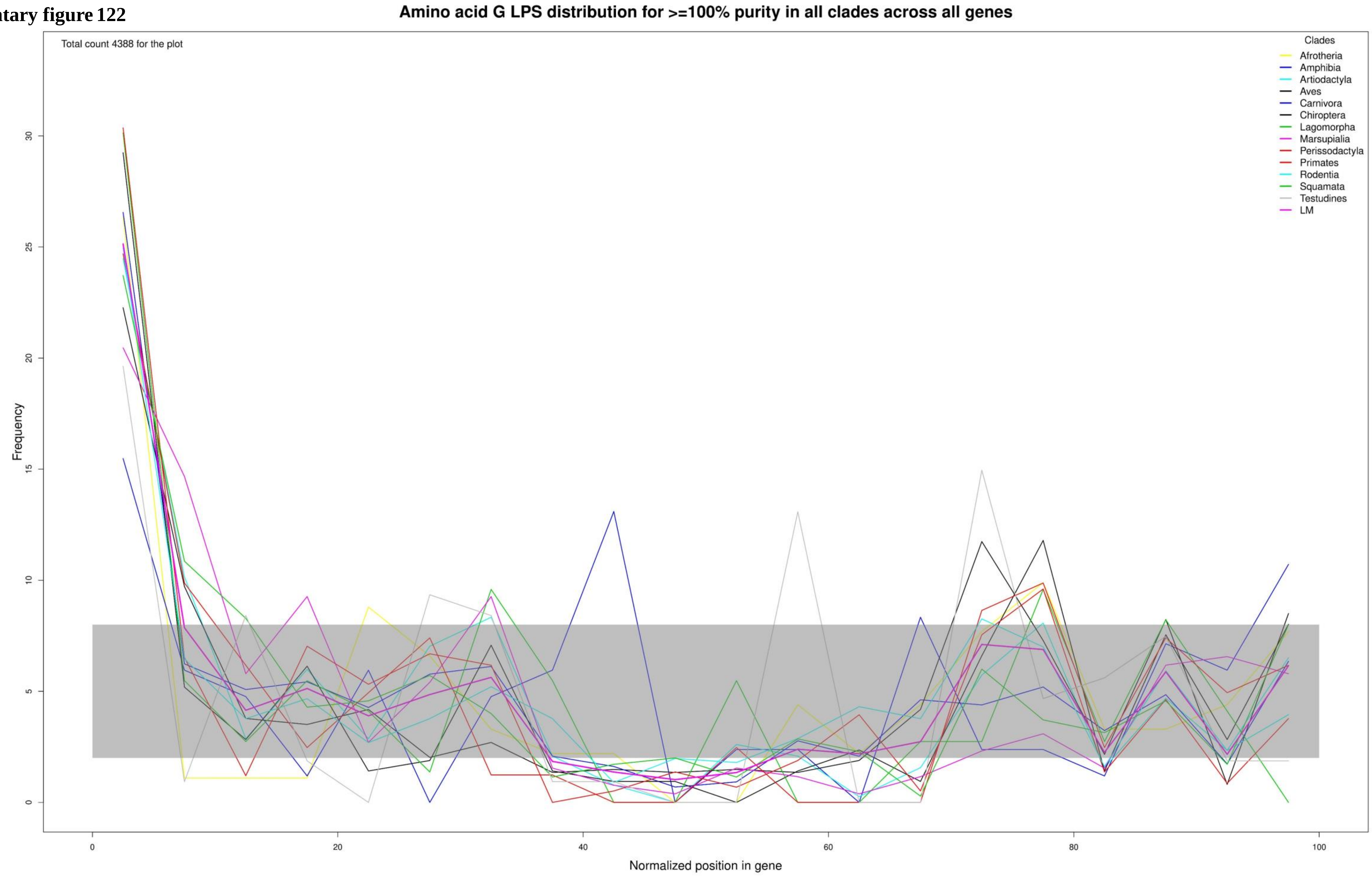

**Supplementary figure 123**

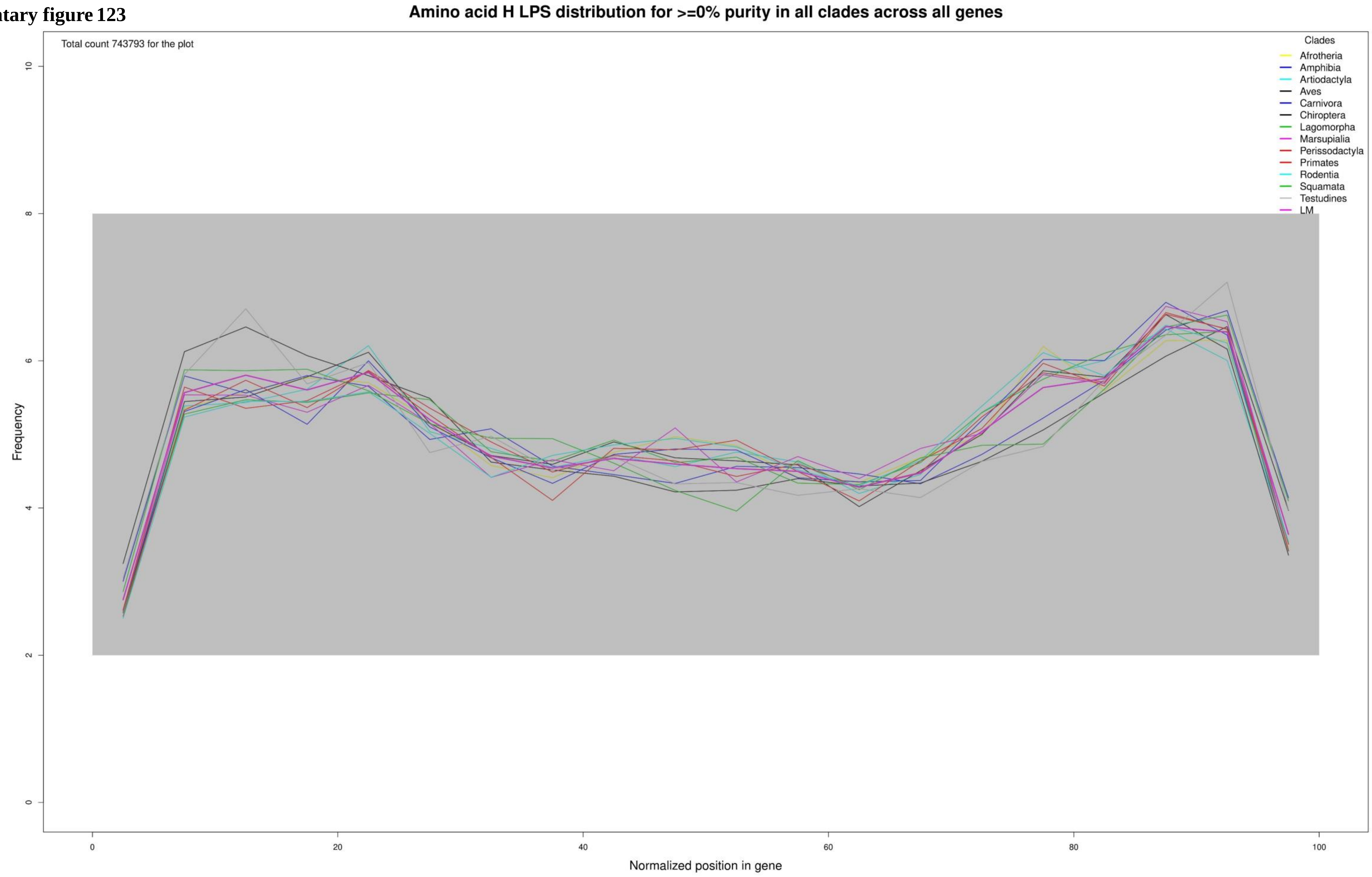

**Supplementary figure 124**

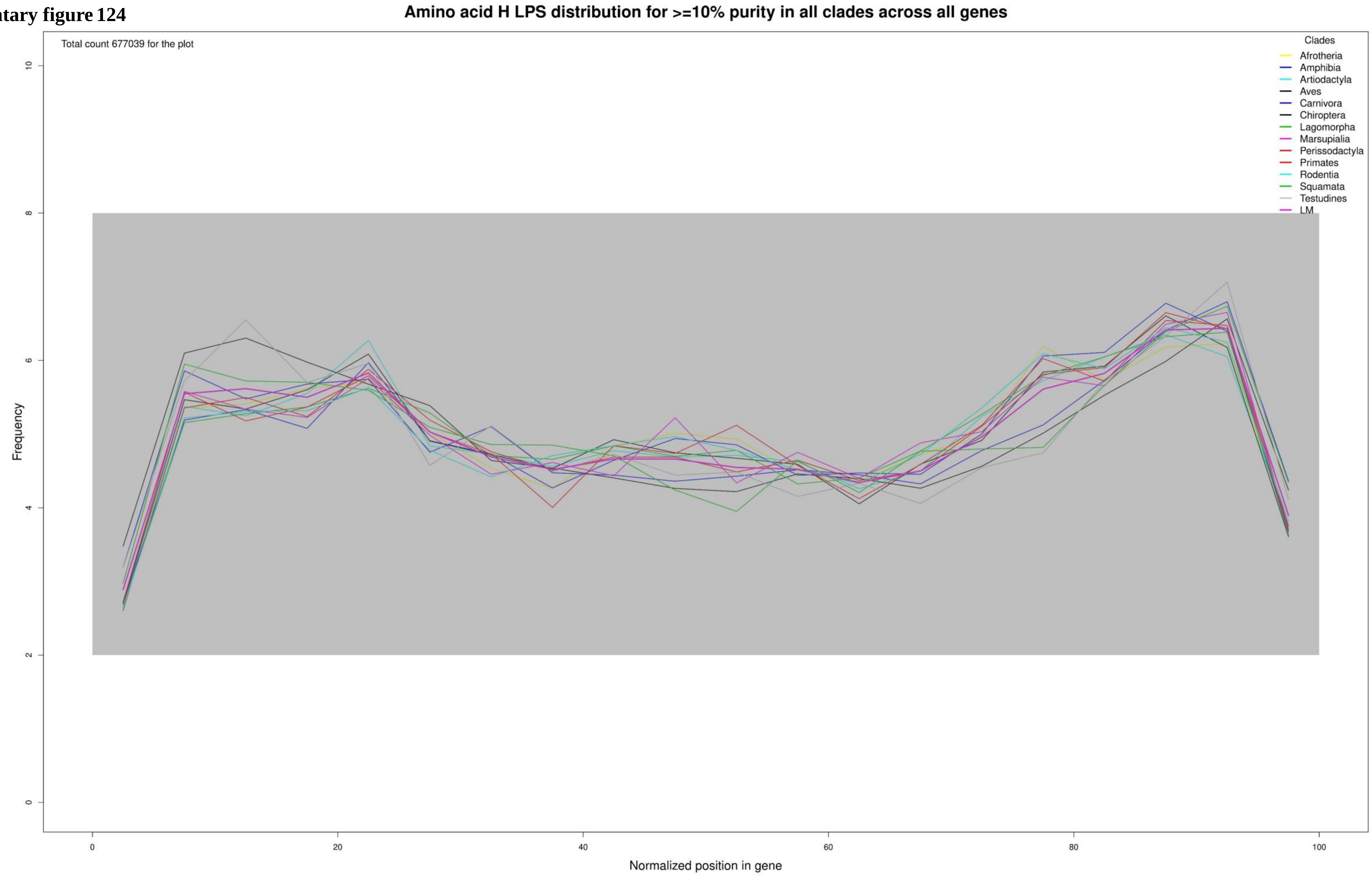

**Supplementary figure 125**

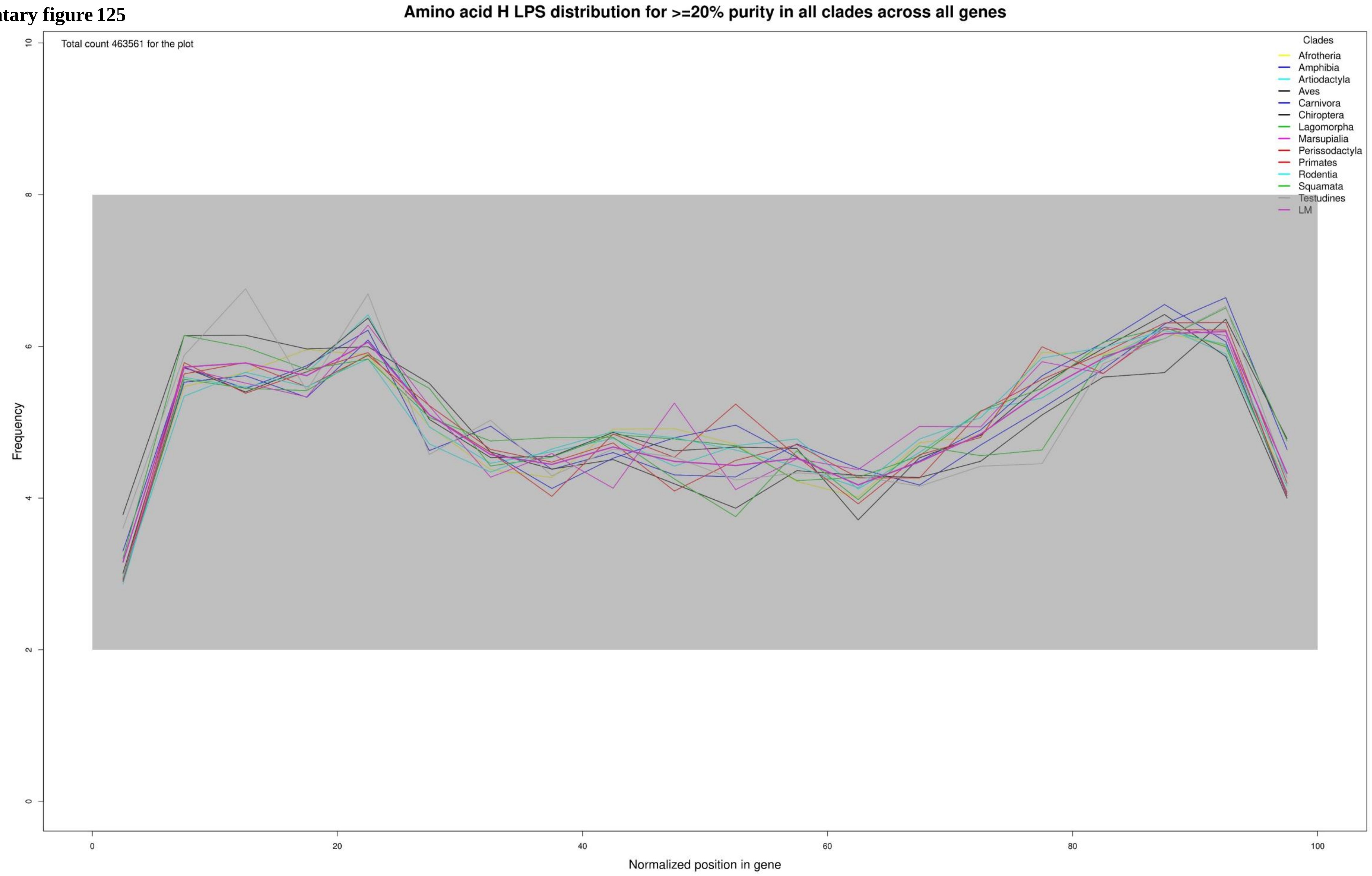

**Supplementary figure 126**

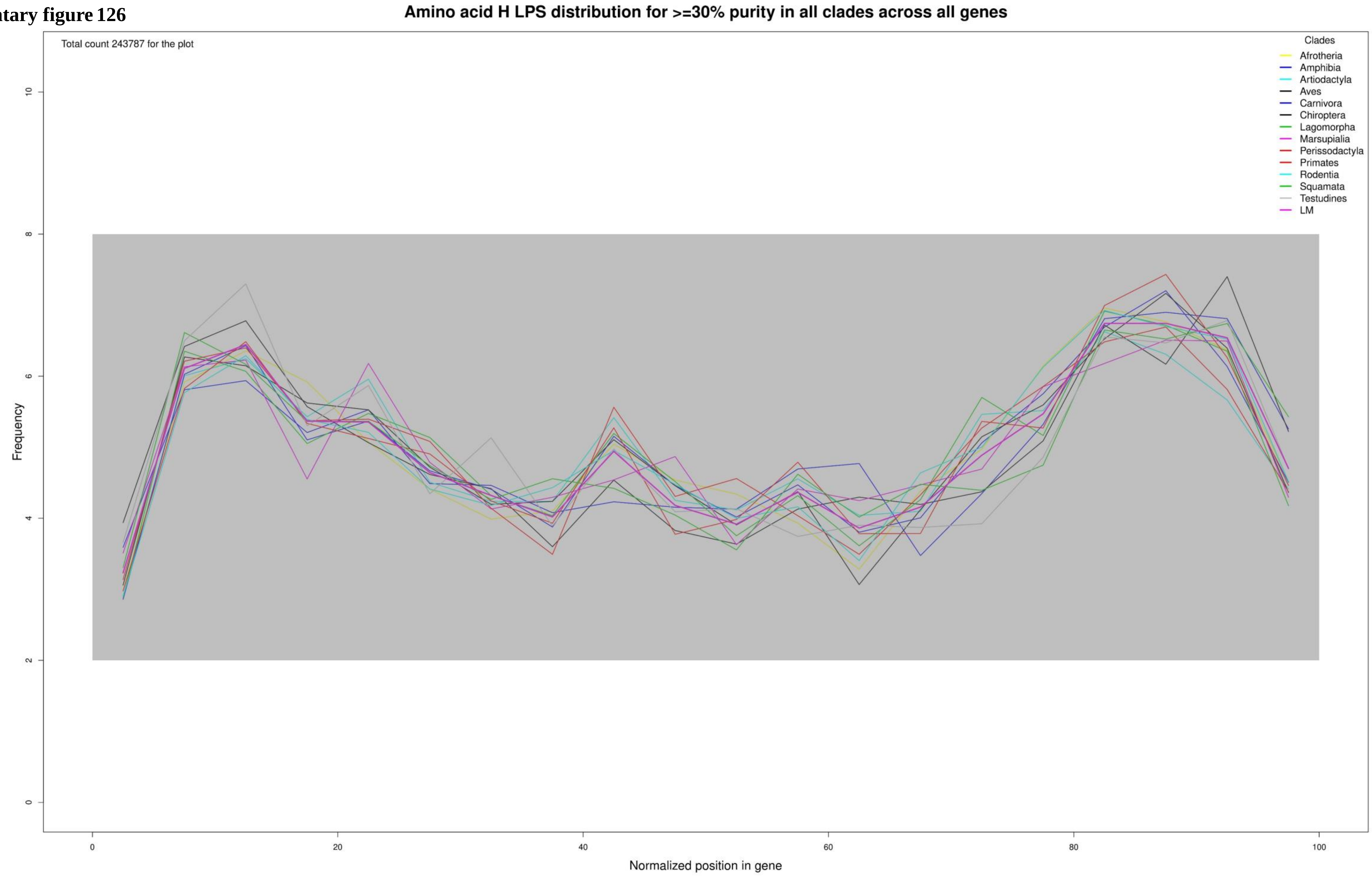

**Supplementary figure 127**

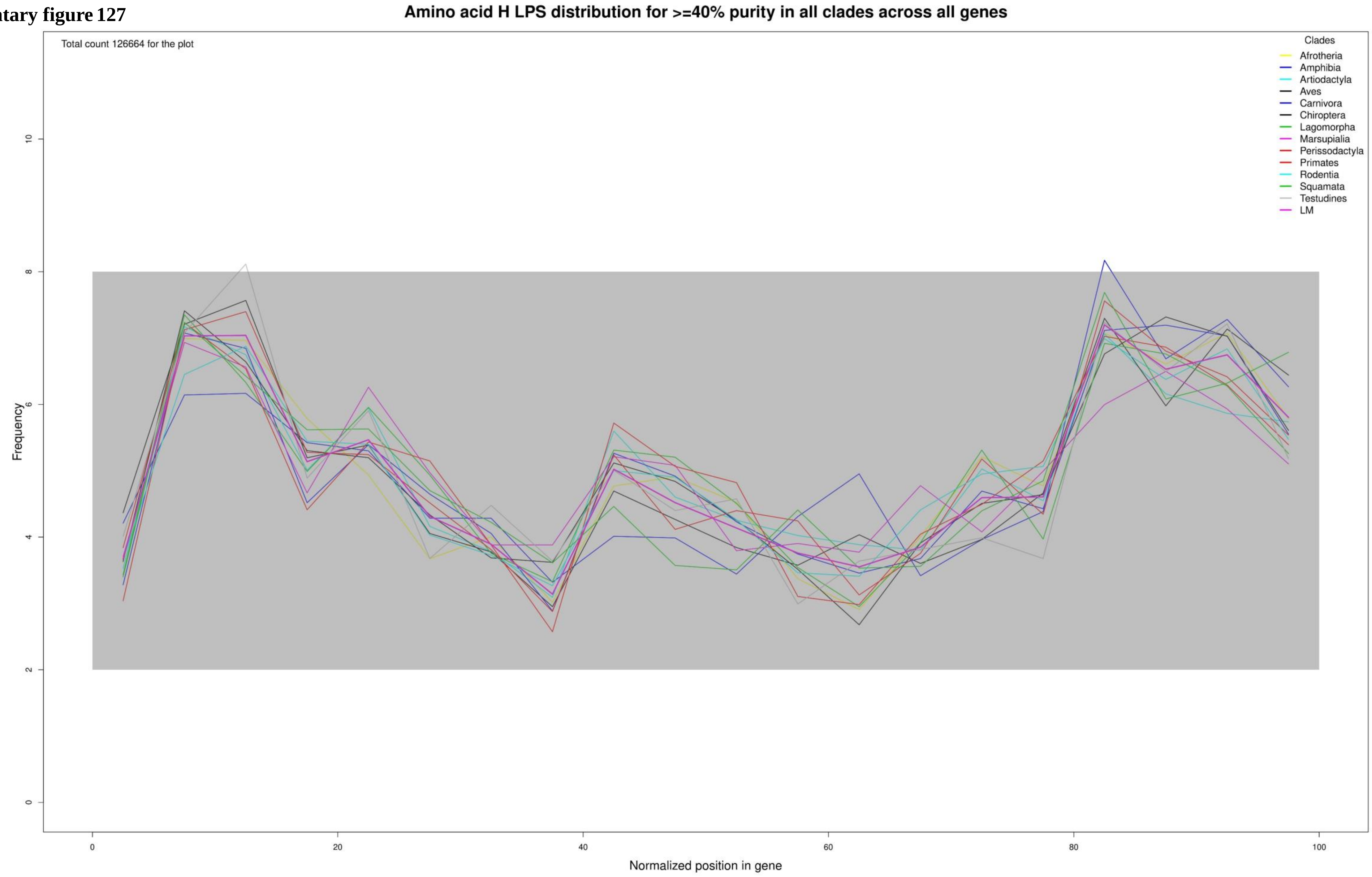

**Supplementary figure 128**

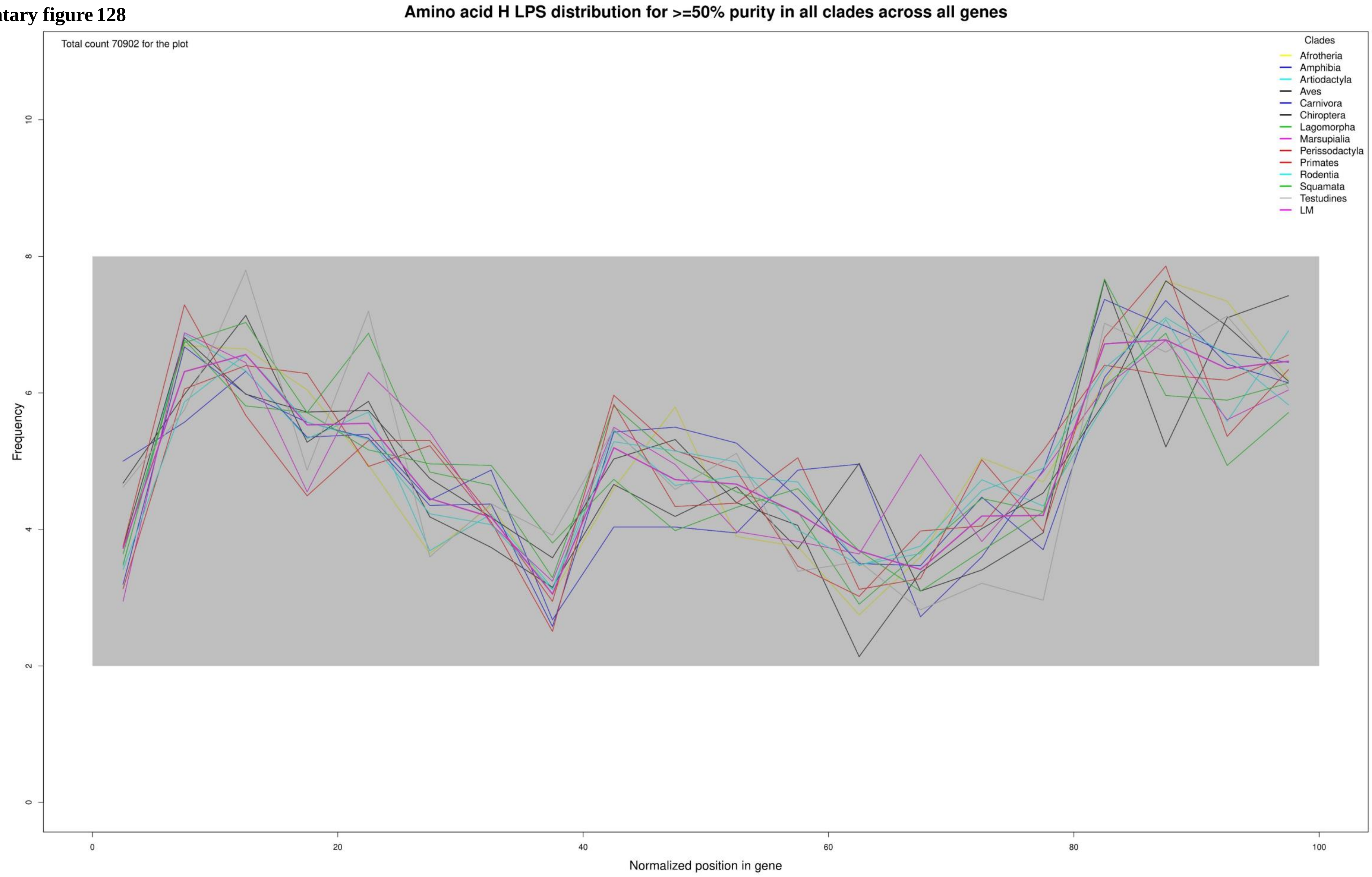

**Supplementary figure 129**

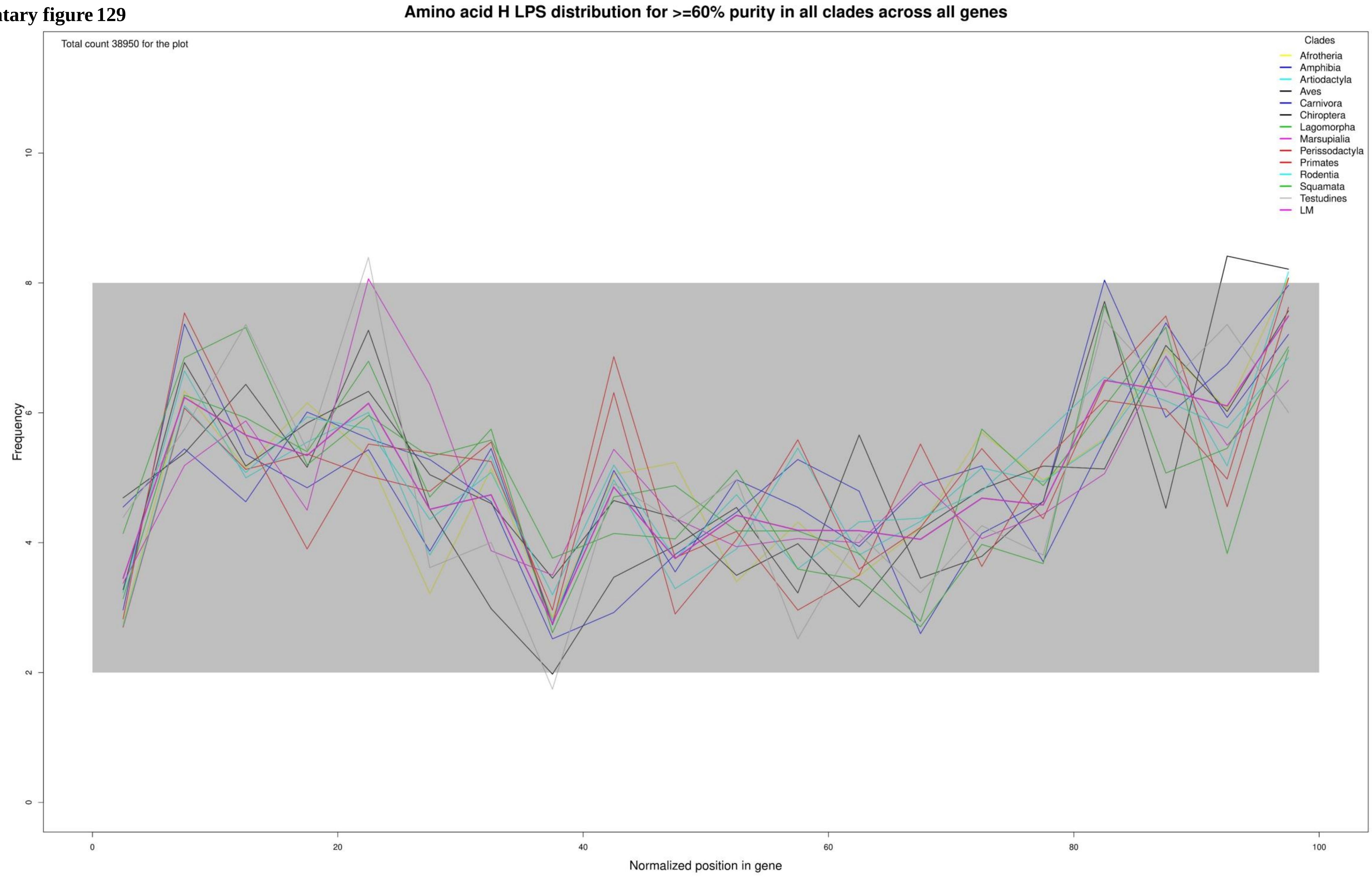

**Supplementary figure 130**

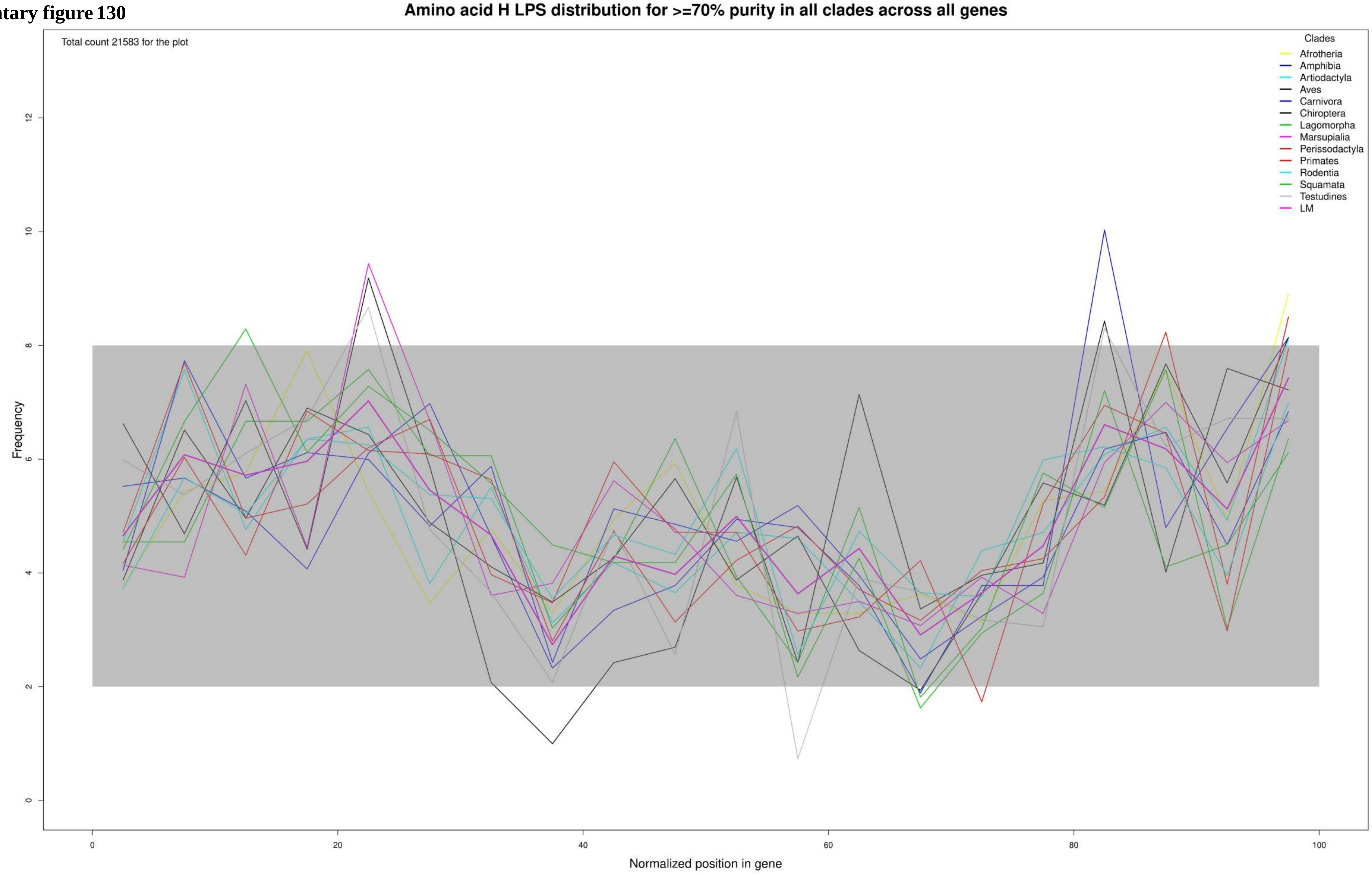

**Supplementary figure 131**

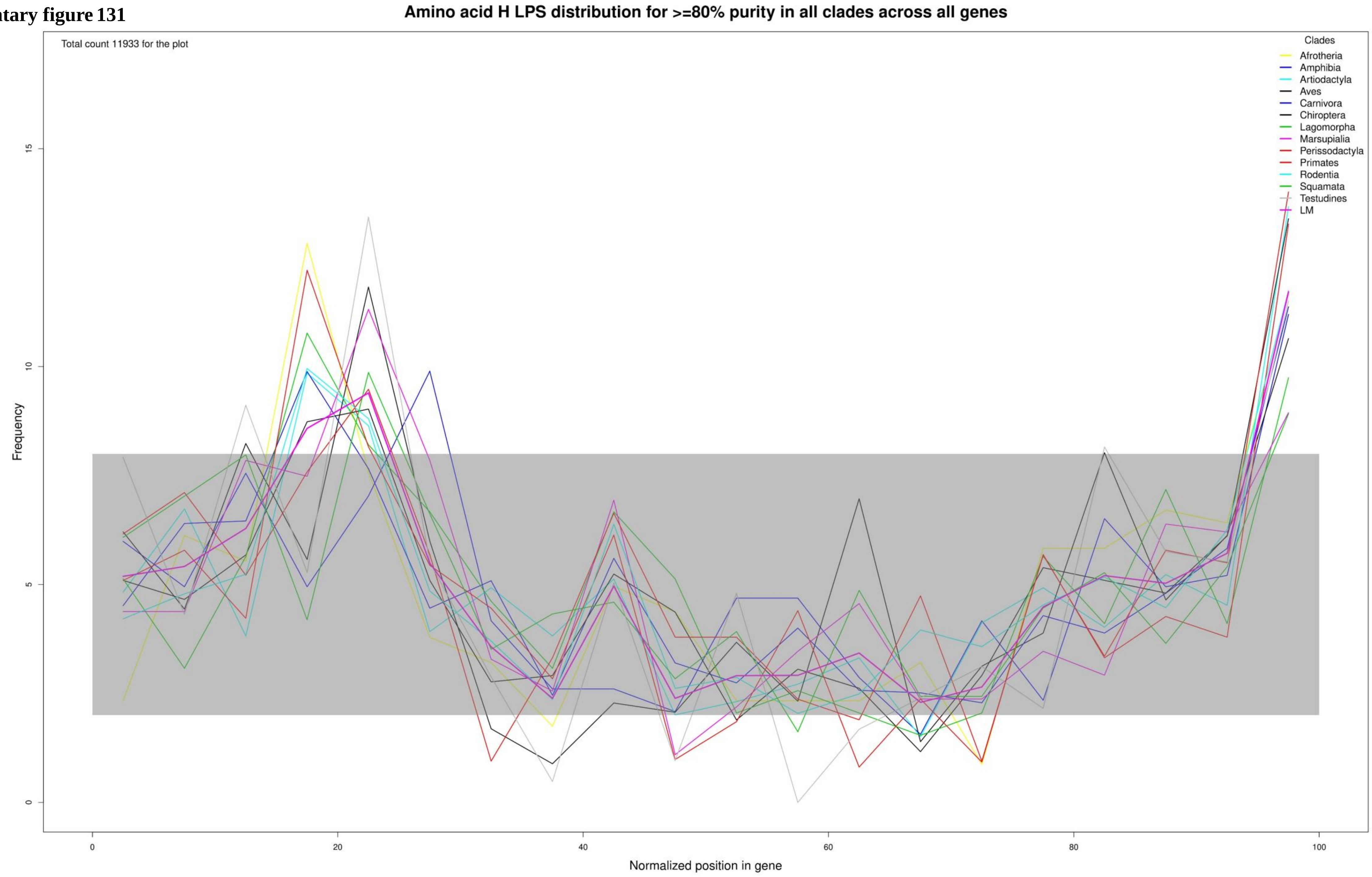

**Supplementary figure 132**

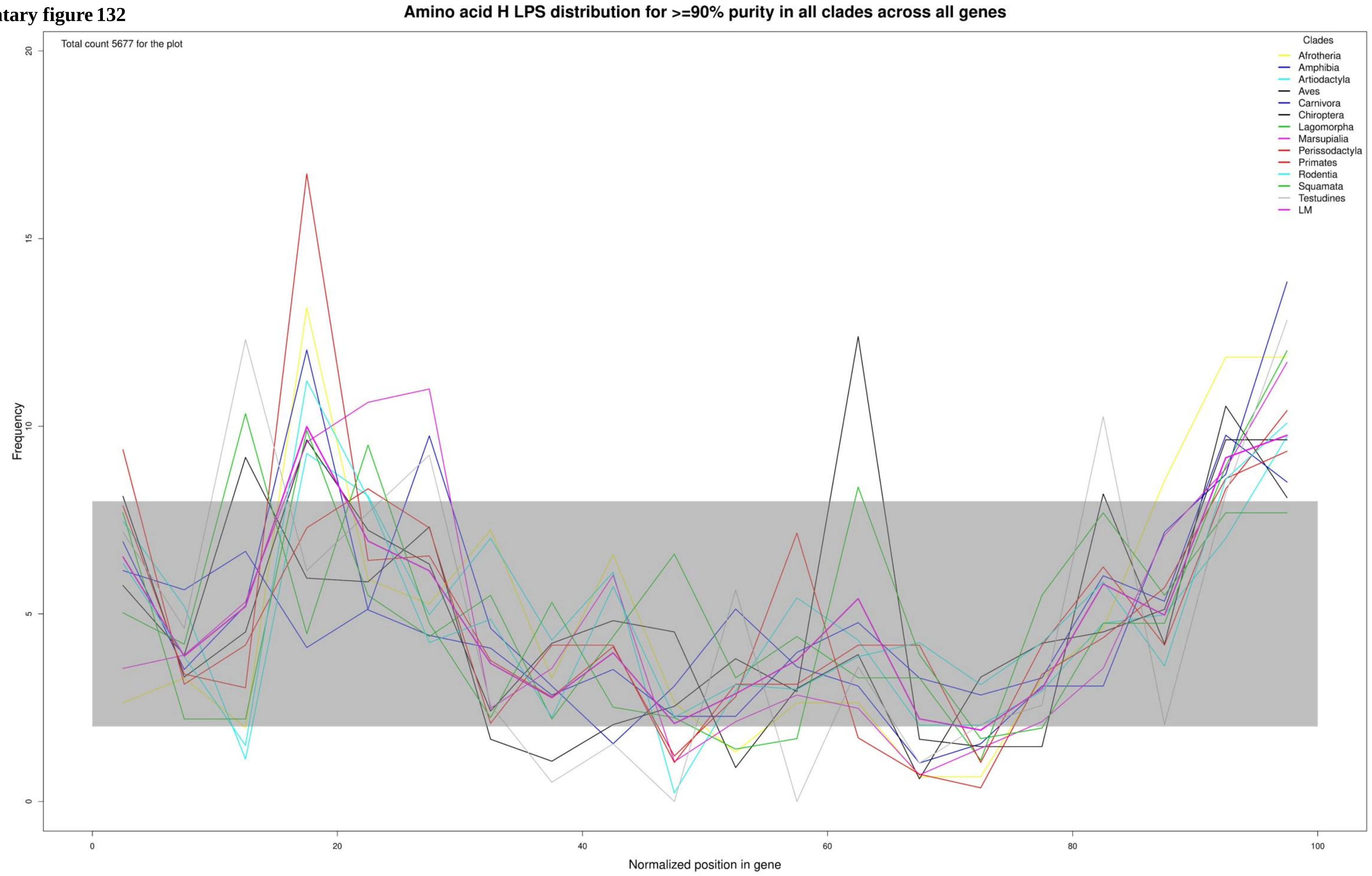

**Supplementary figure 133**

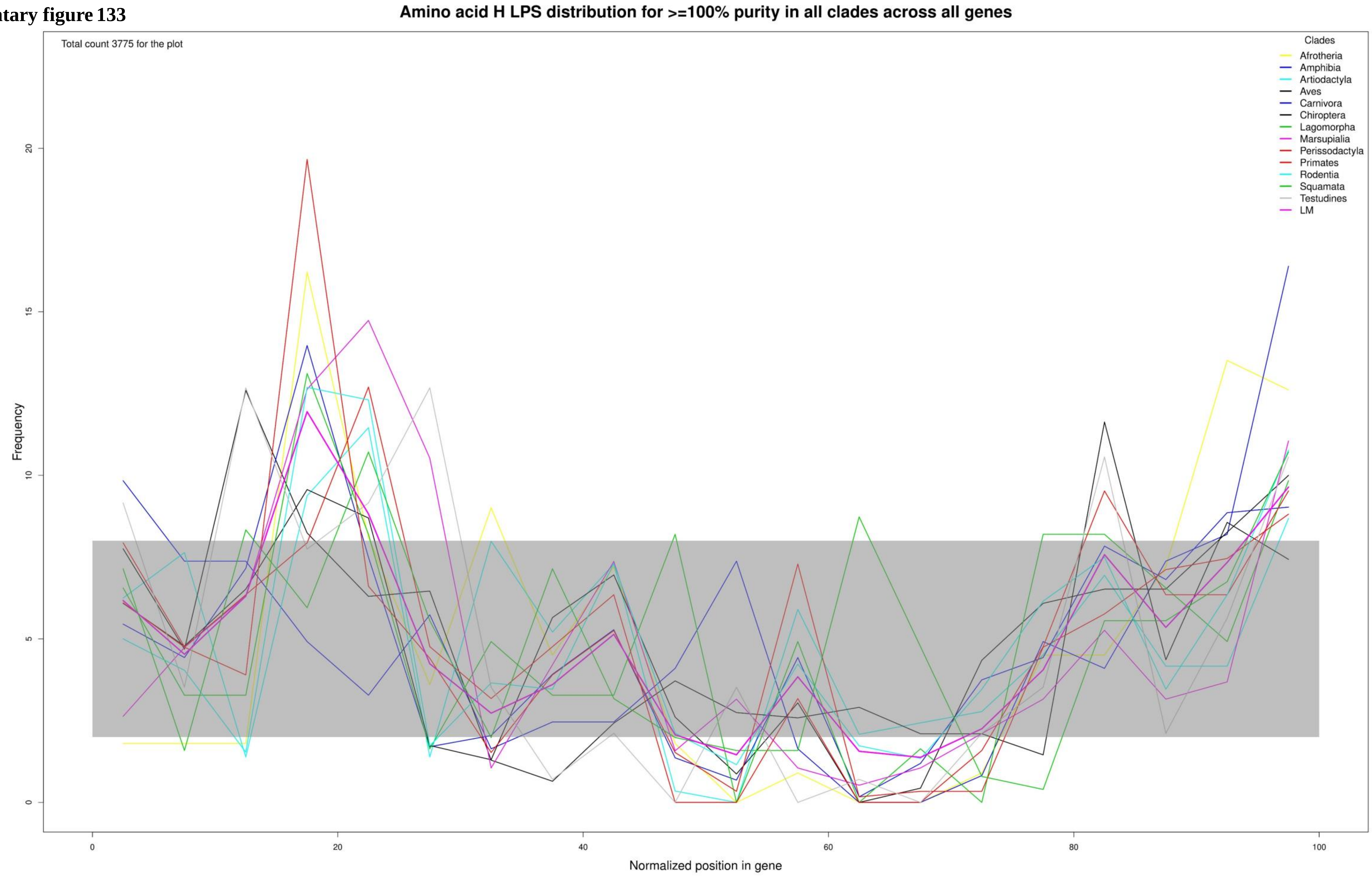

**Supplementary figure 134**

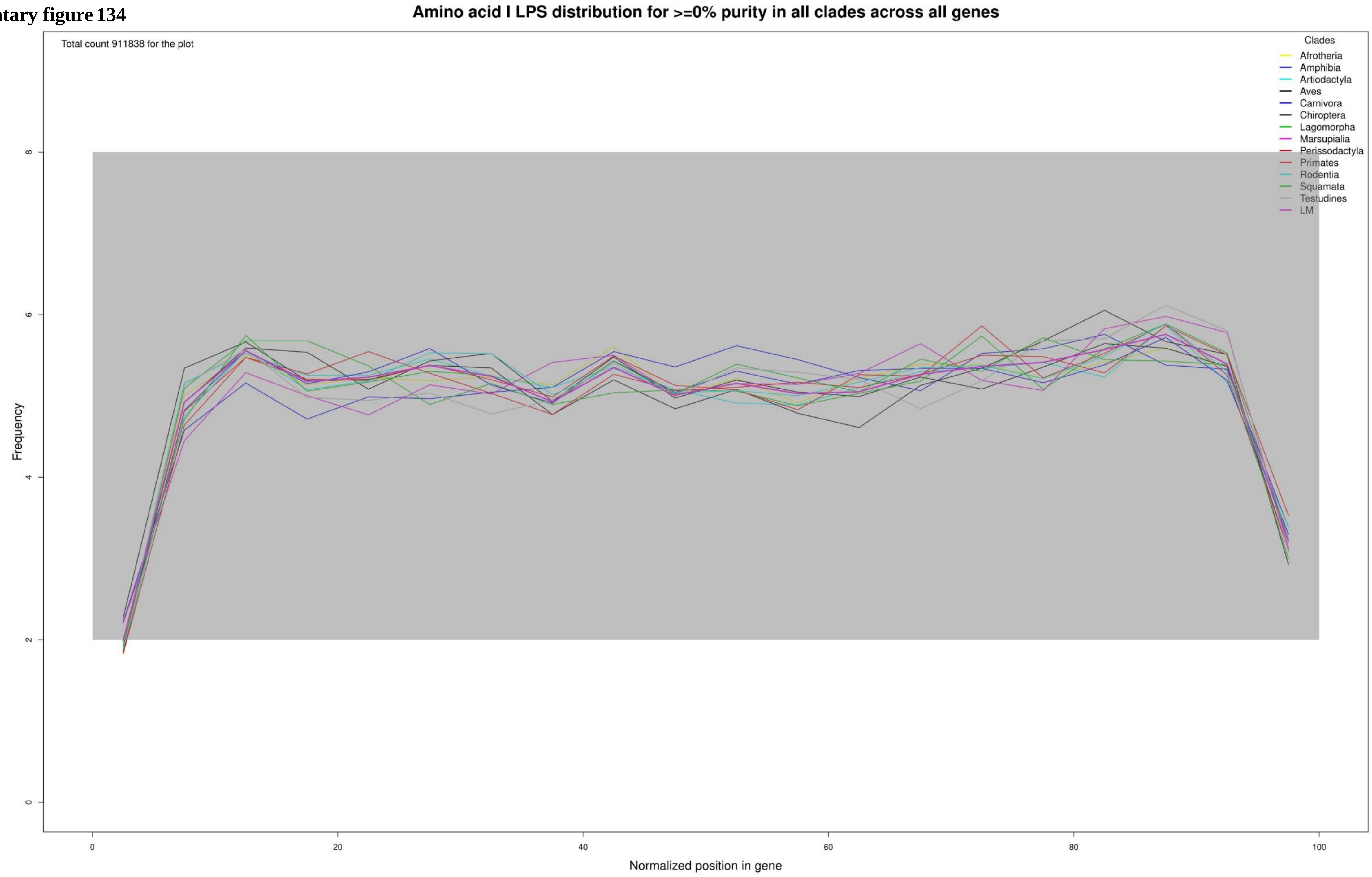

Supplementary figure 135

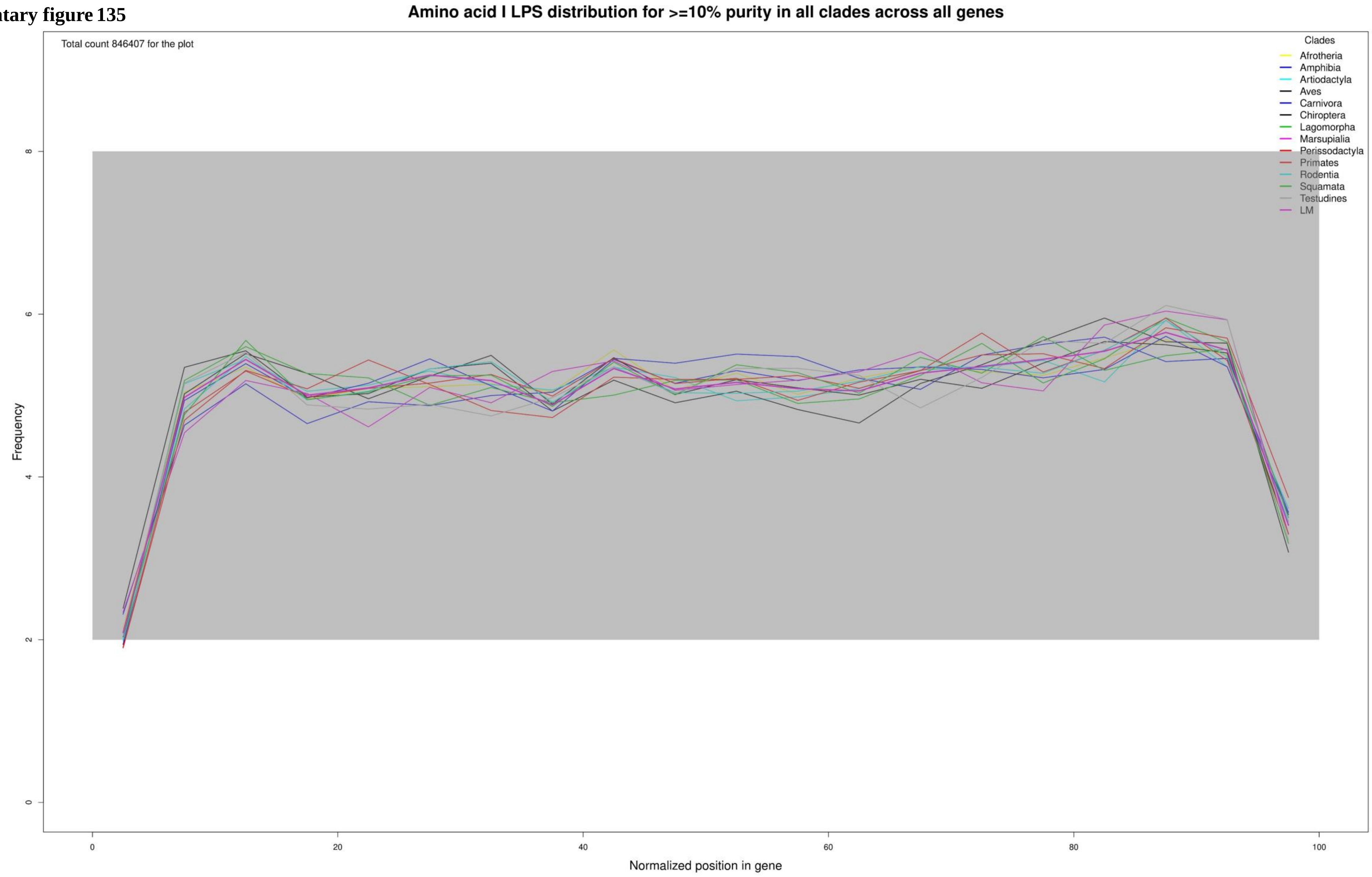

**Supplementary figure 136**

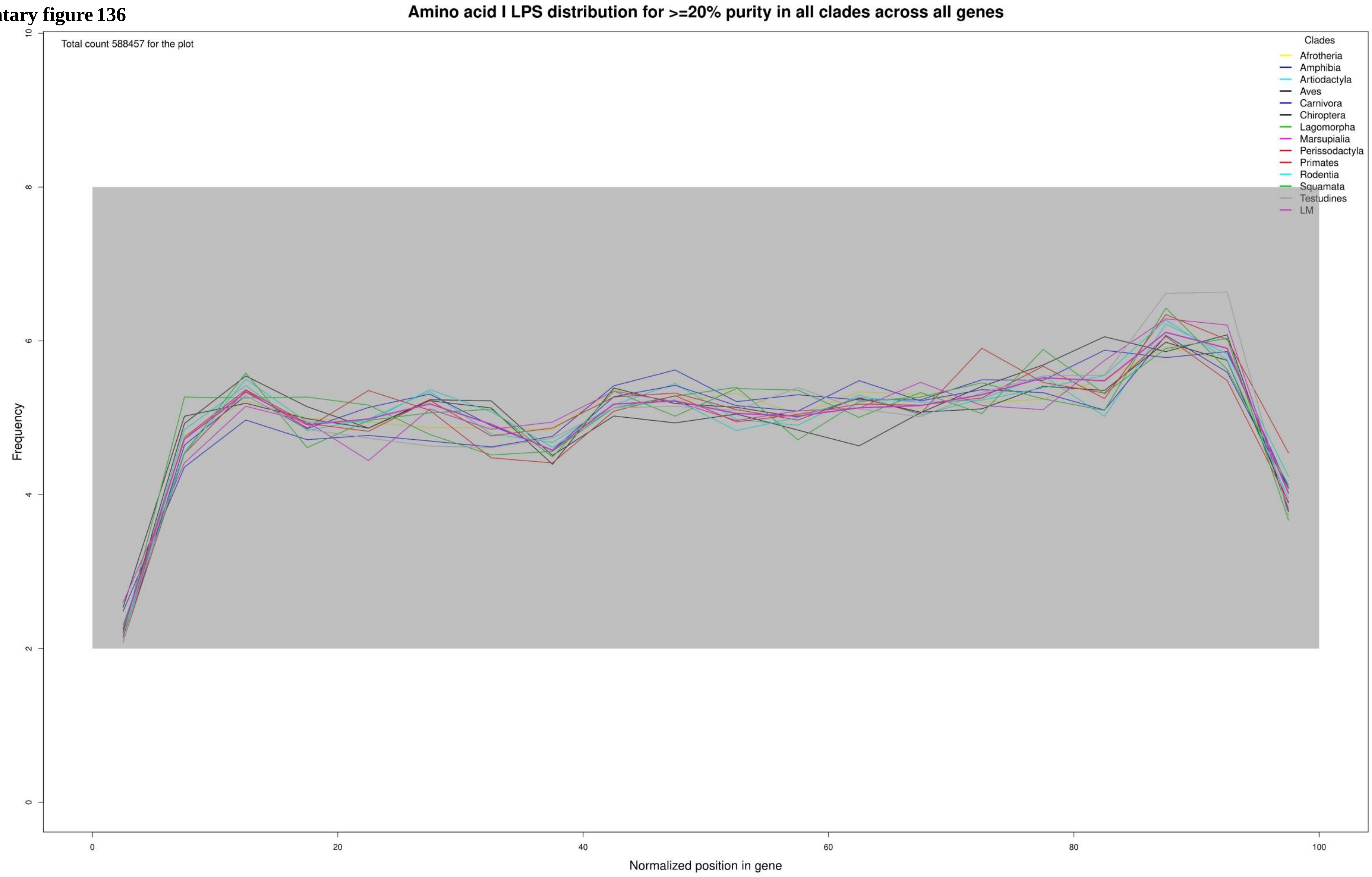

**Supplementary figure 137**

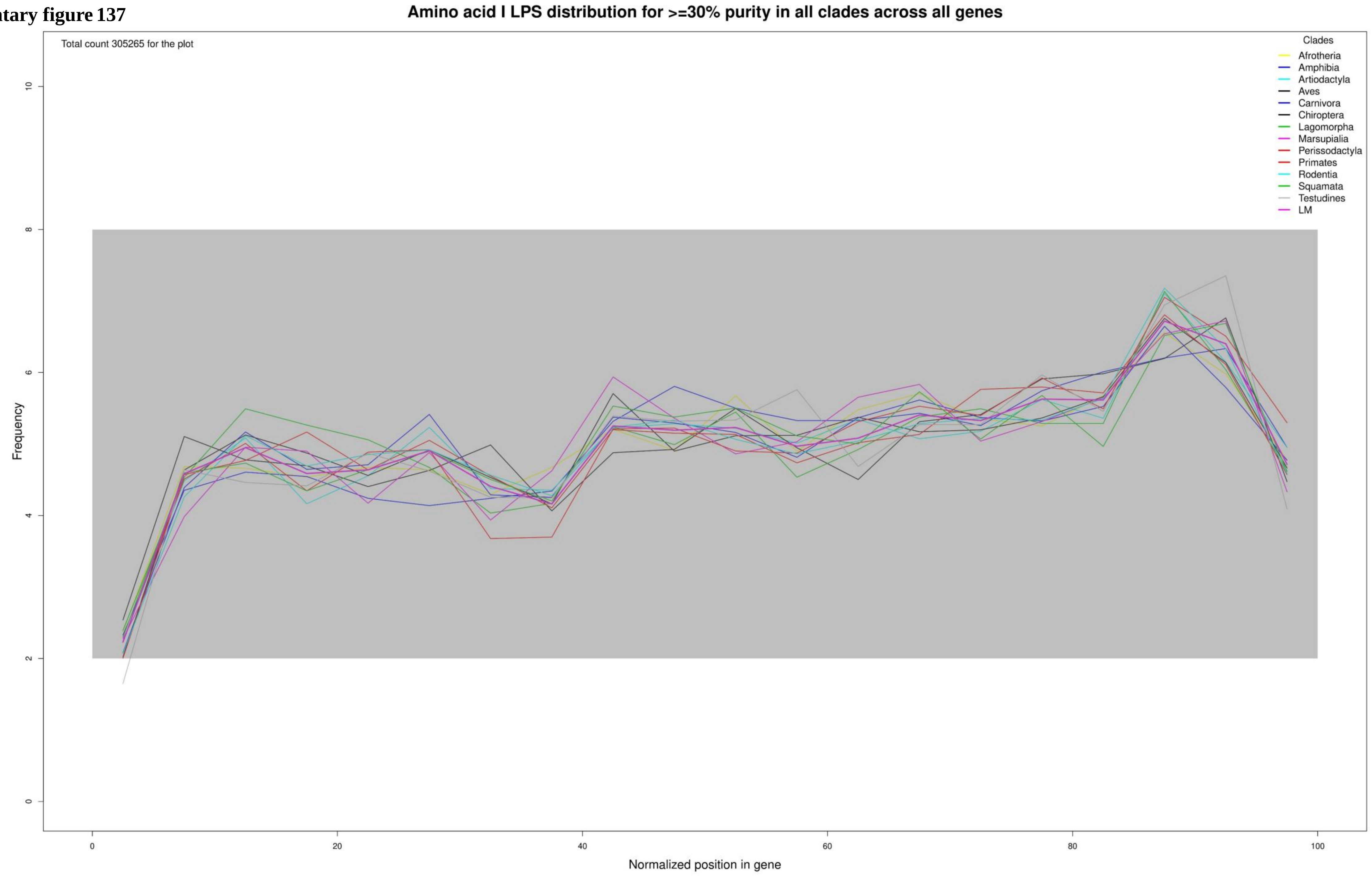

**Supplementary figure 138**

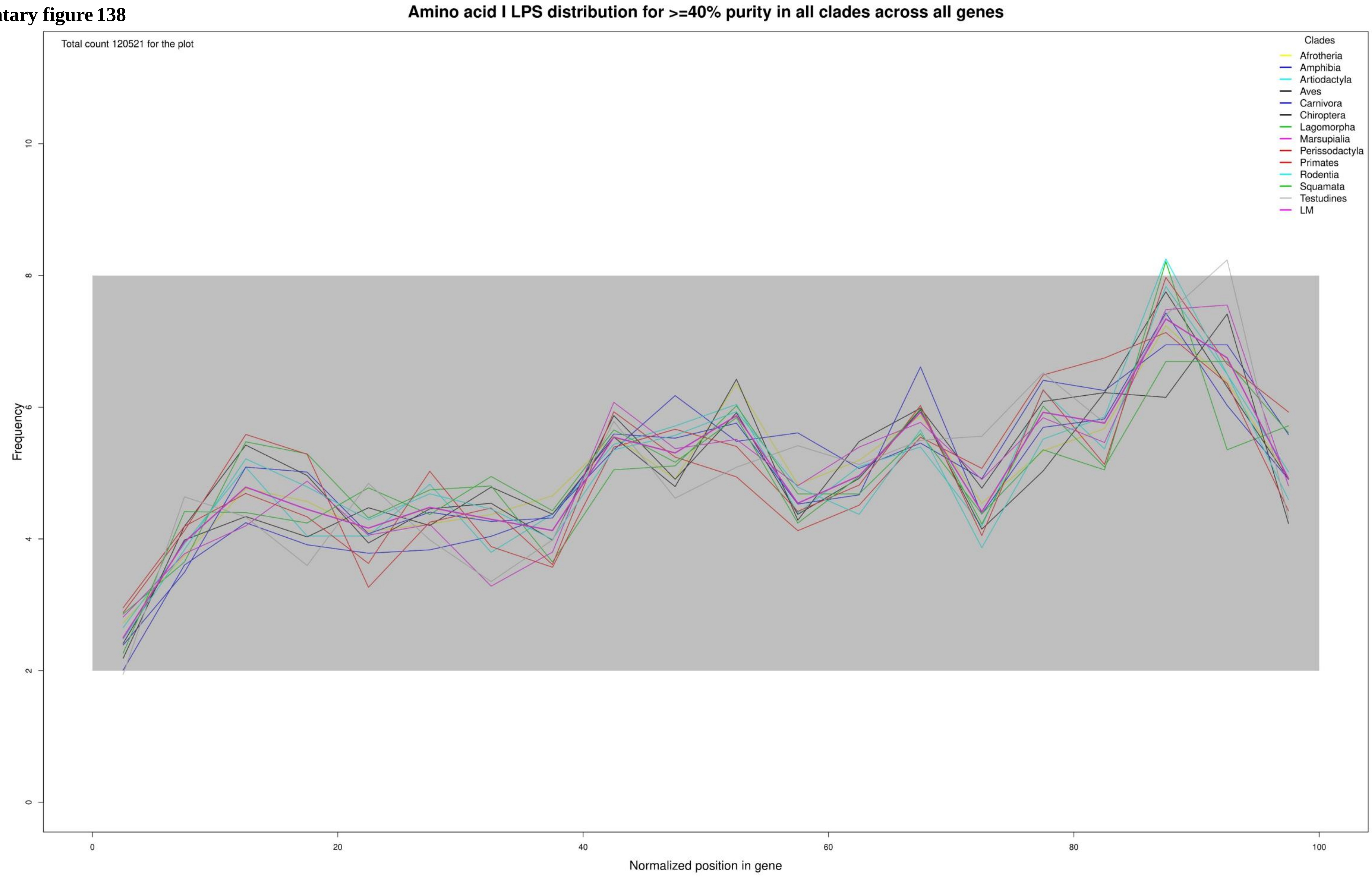

**Supplementary figure 139**

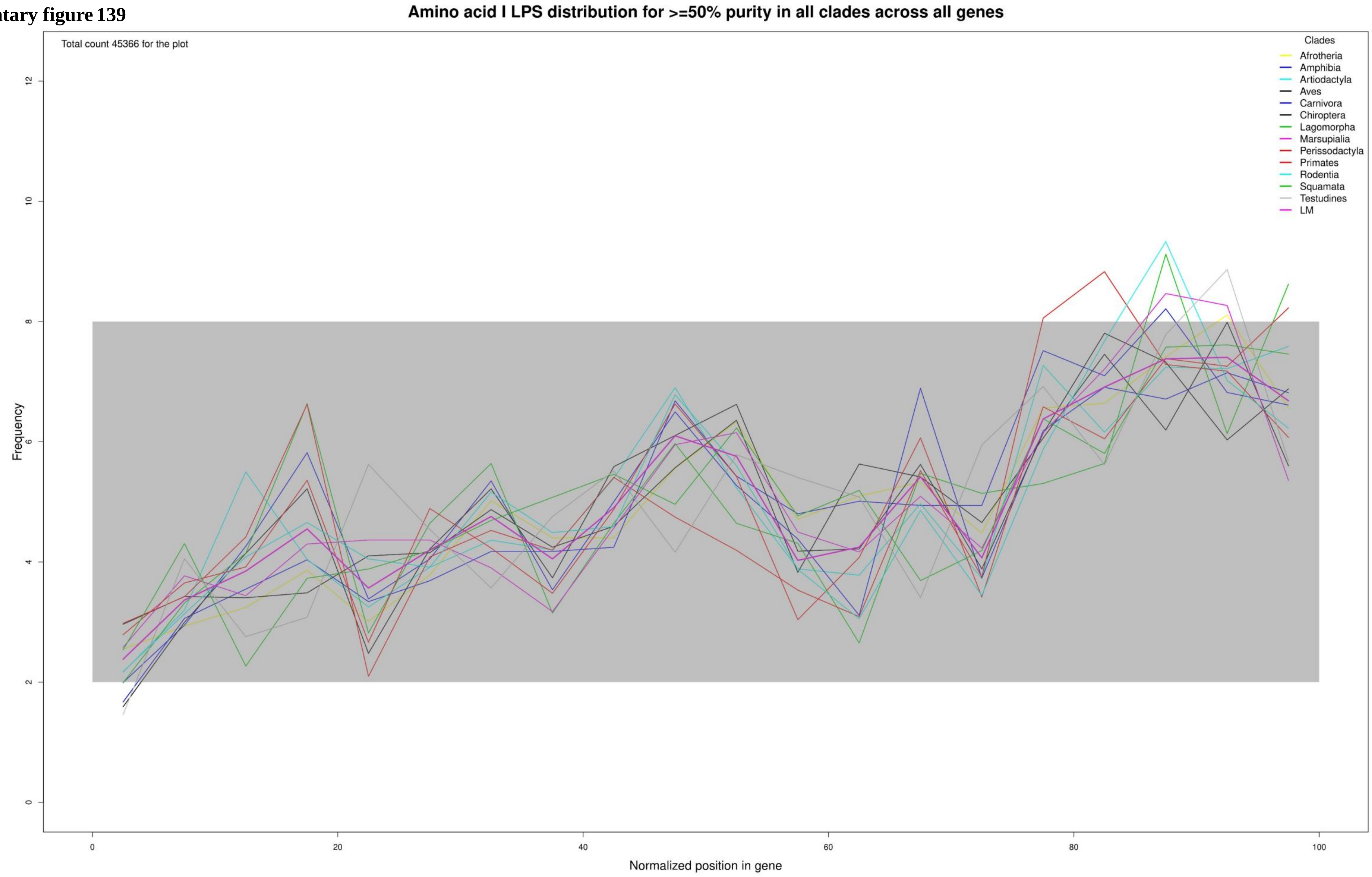

**Supplementary figure 140**

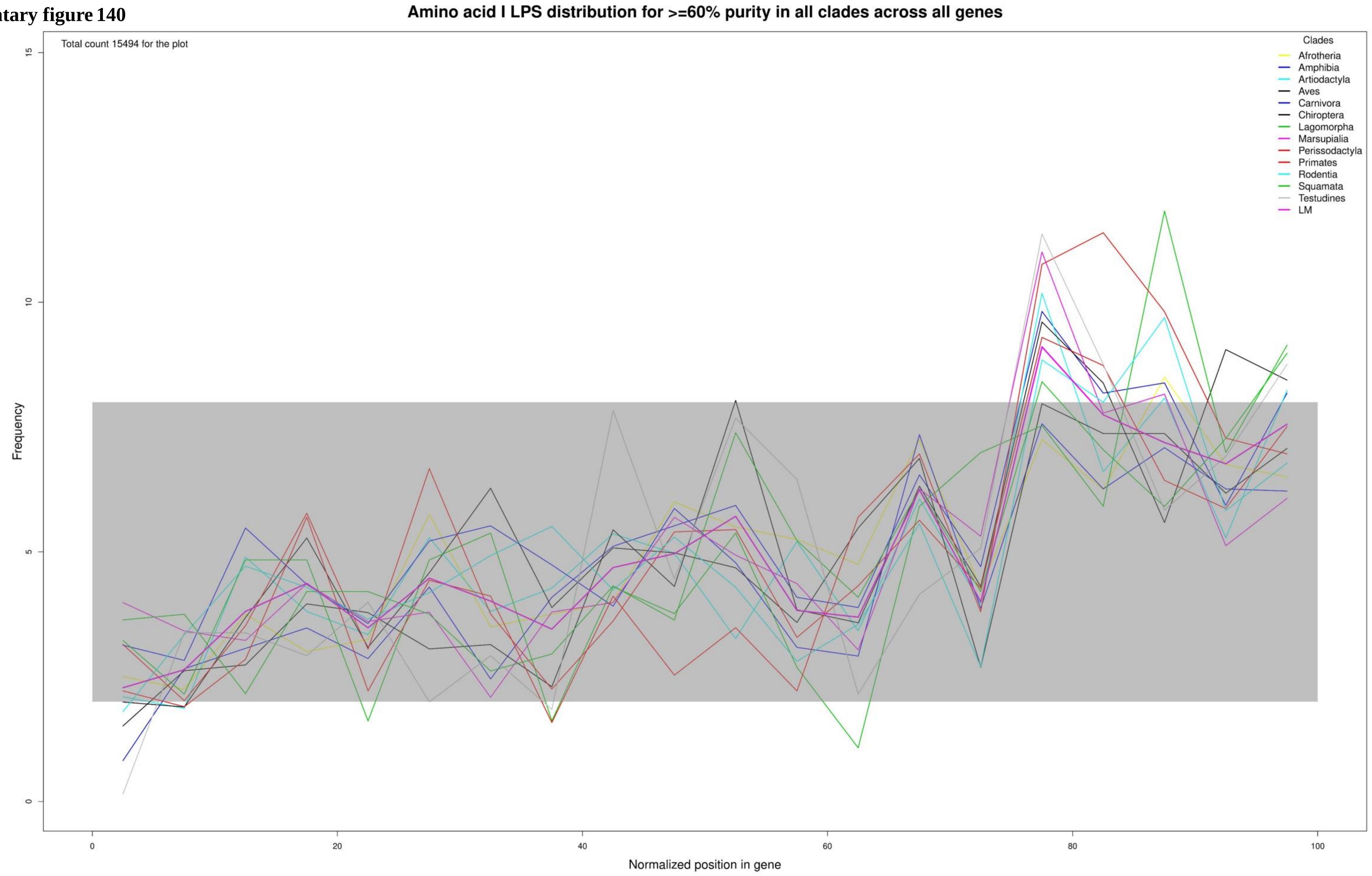

**Supplementary figure 141**

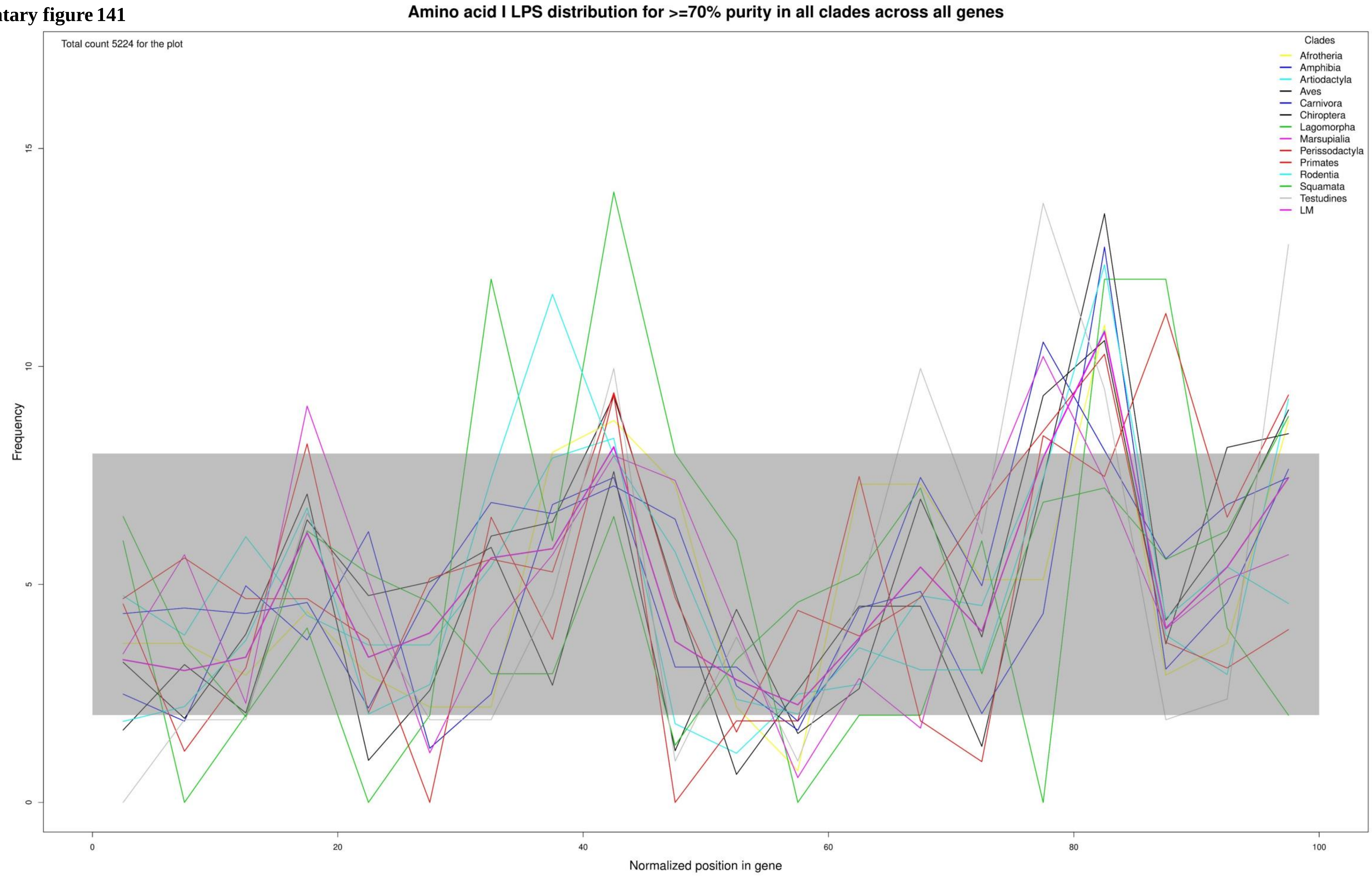

**Supplementary figure 142**

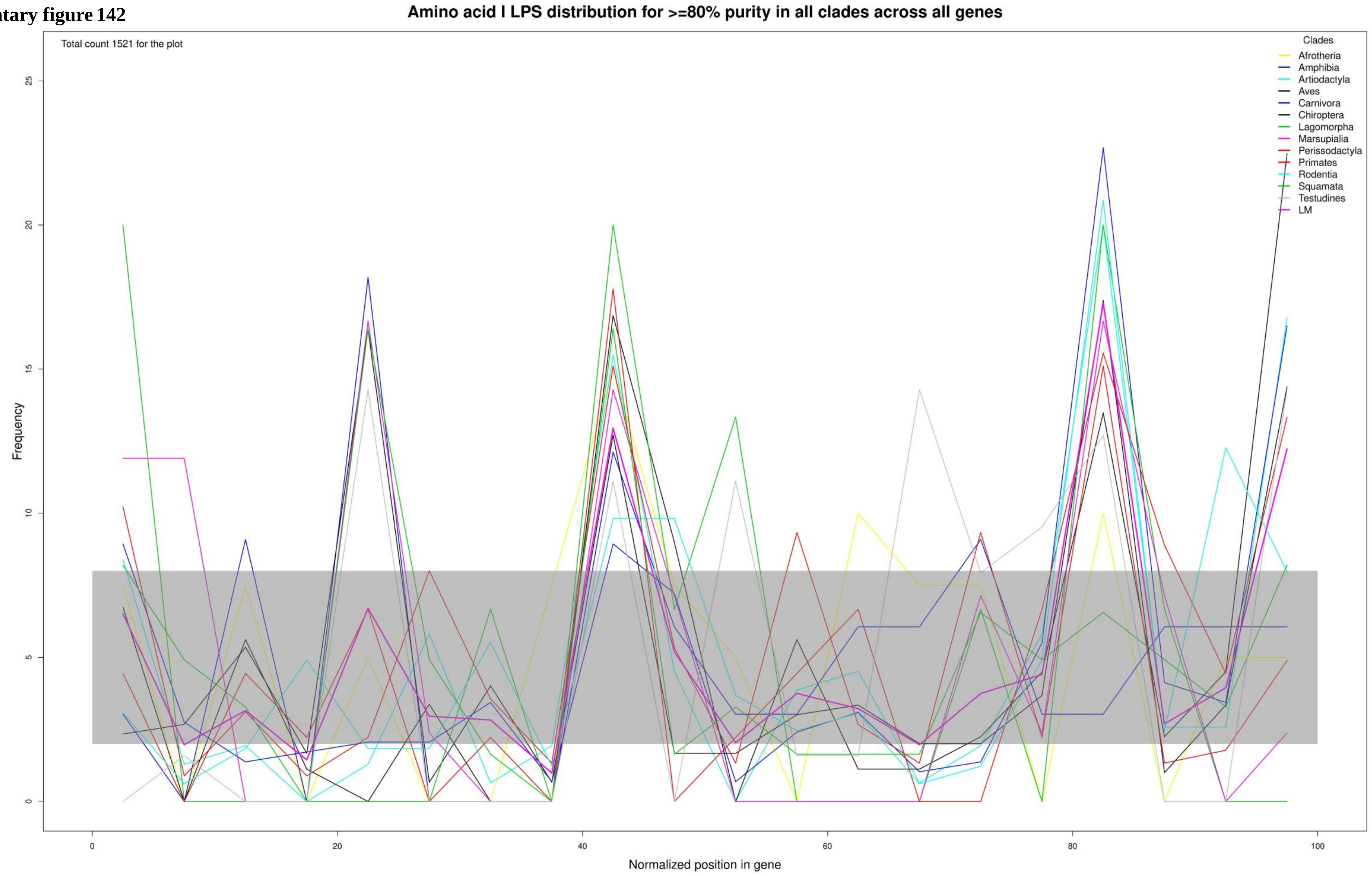

Supplementary figure 143

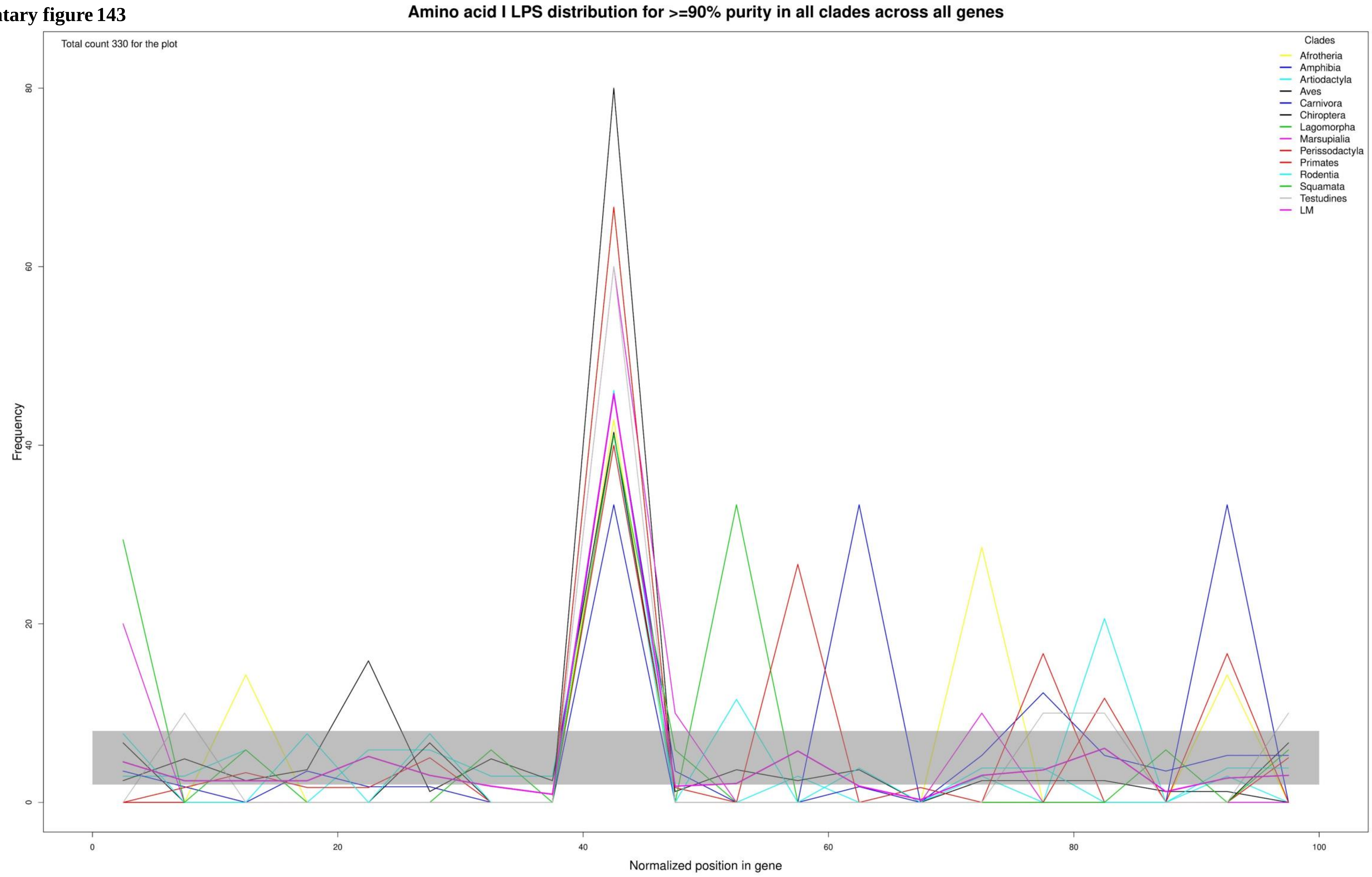

**Supplementary figure 144**

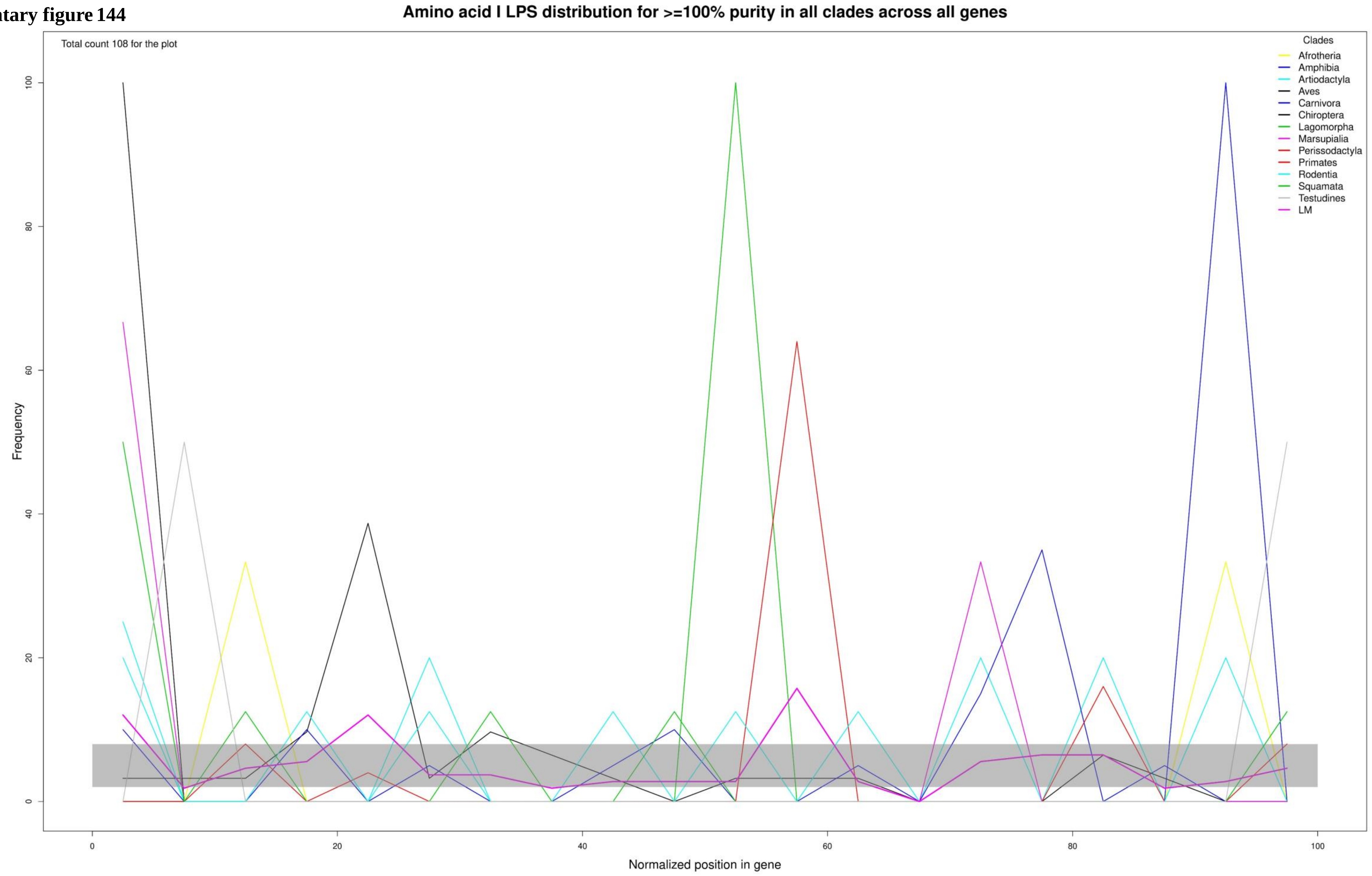

**Supplementary figure 145**

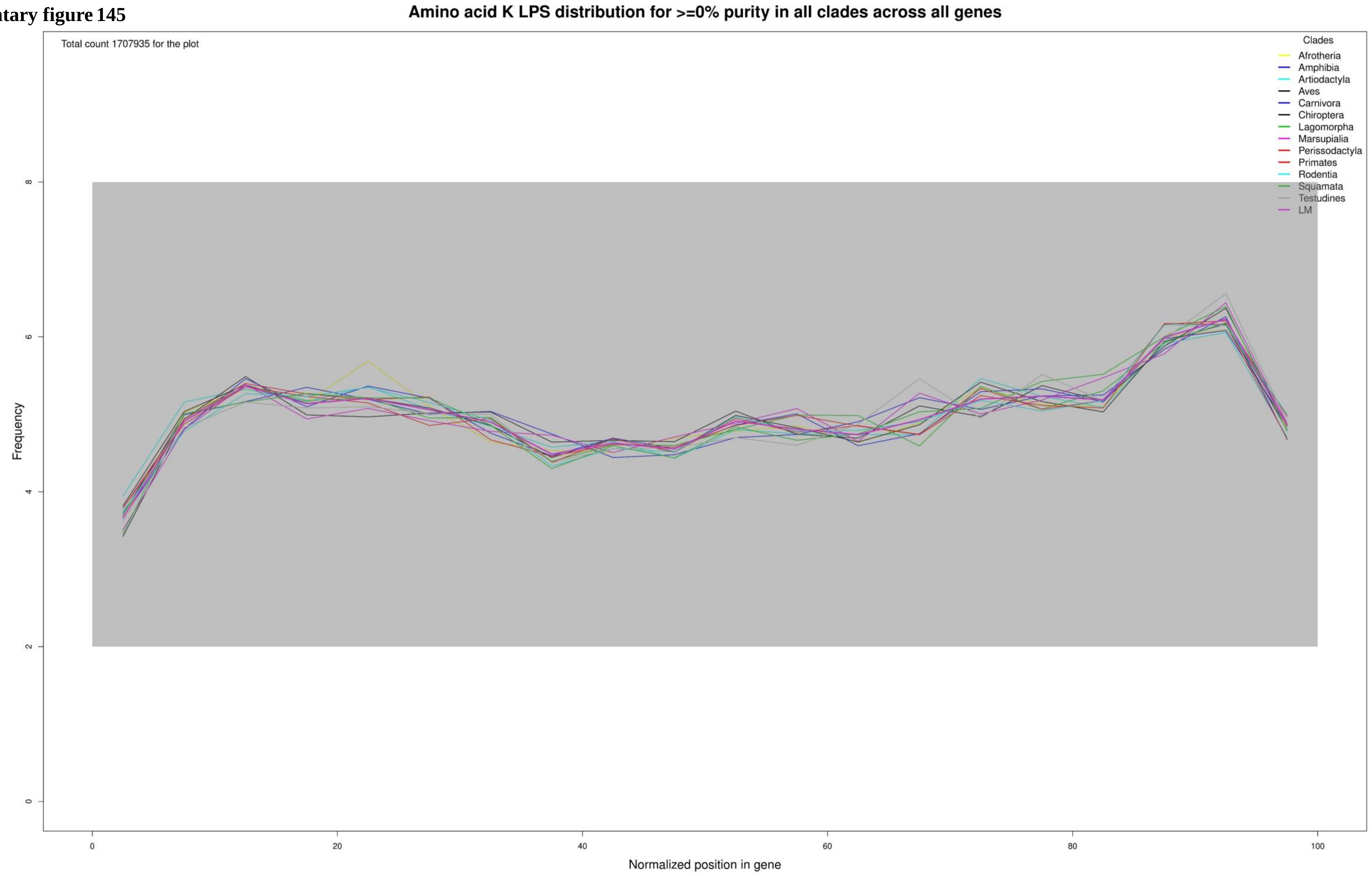

**Supplementary figure 146**

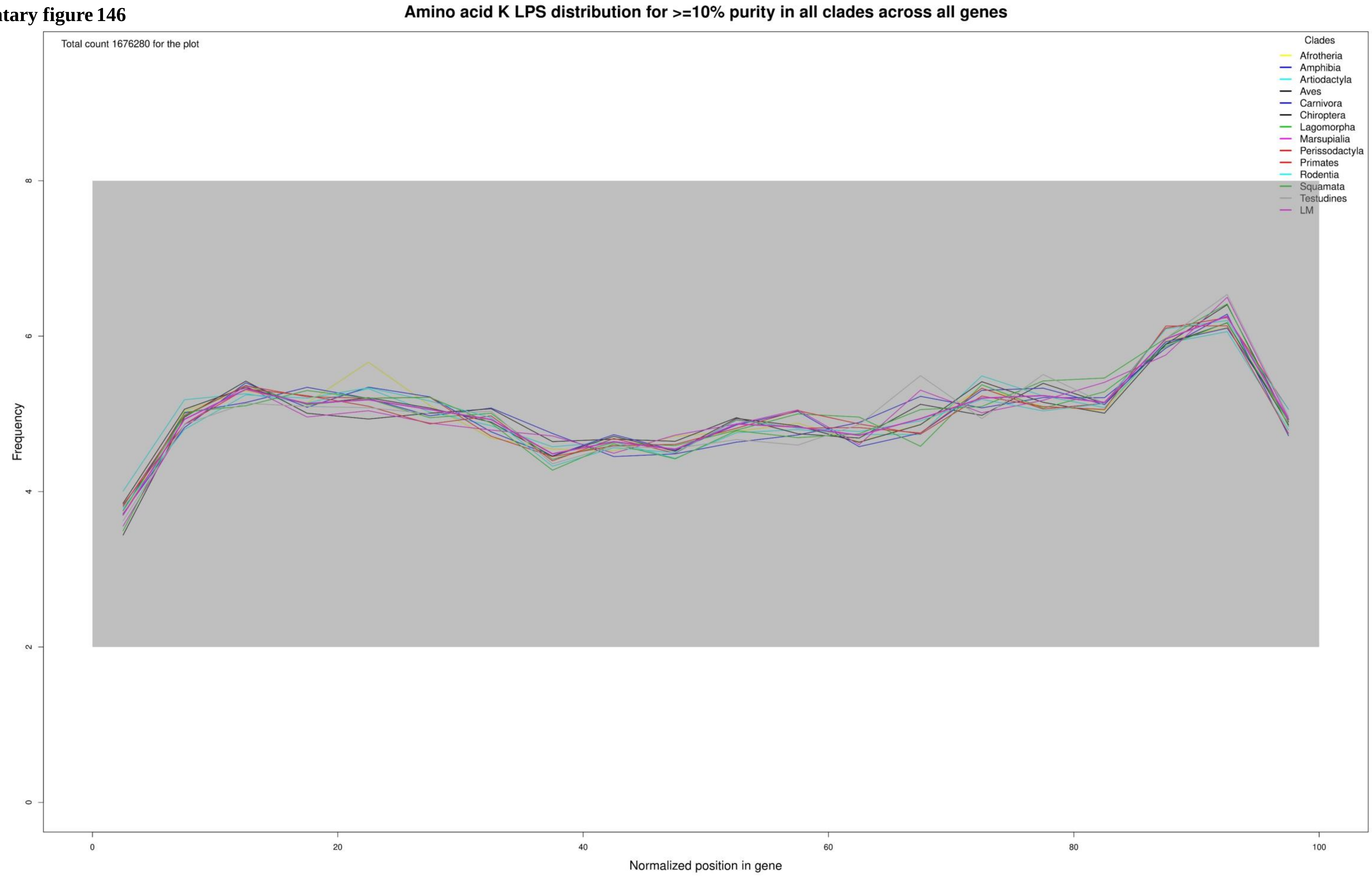

**Supplementary figure 147**

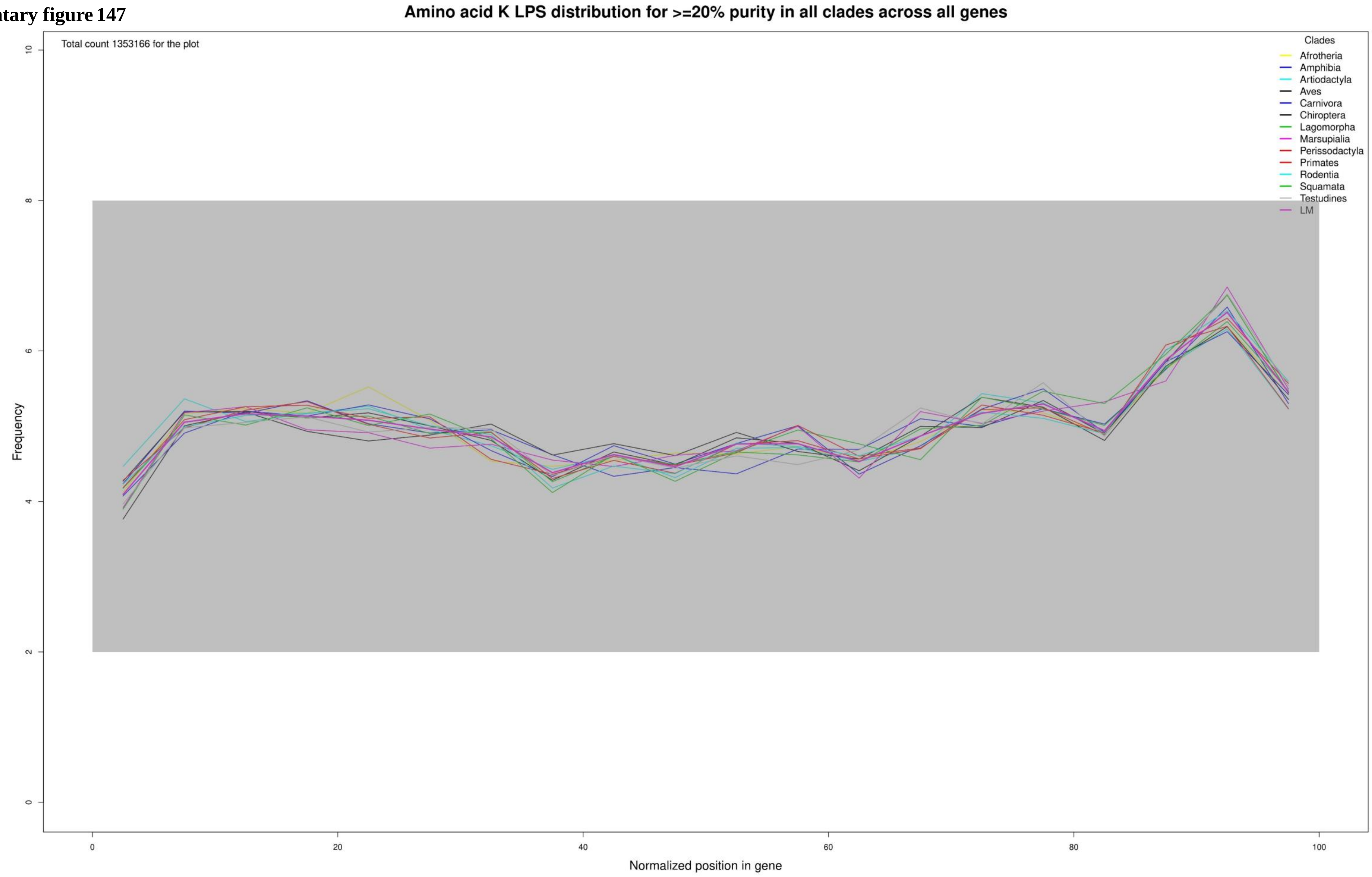

**Supplementary figure 148**

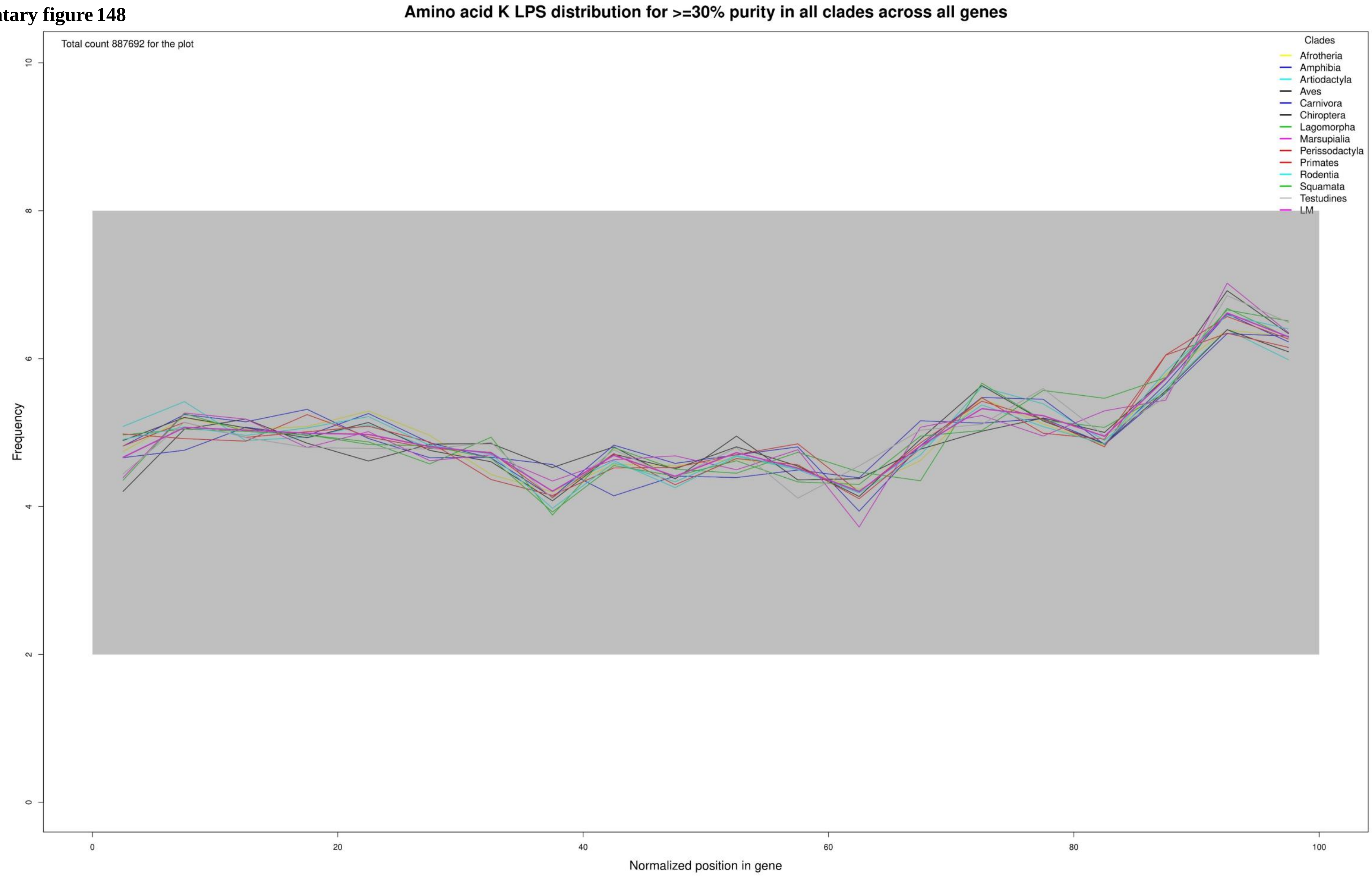

**Supplementary figure 149**

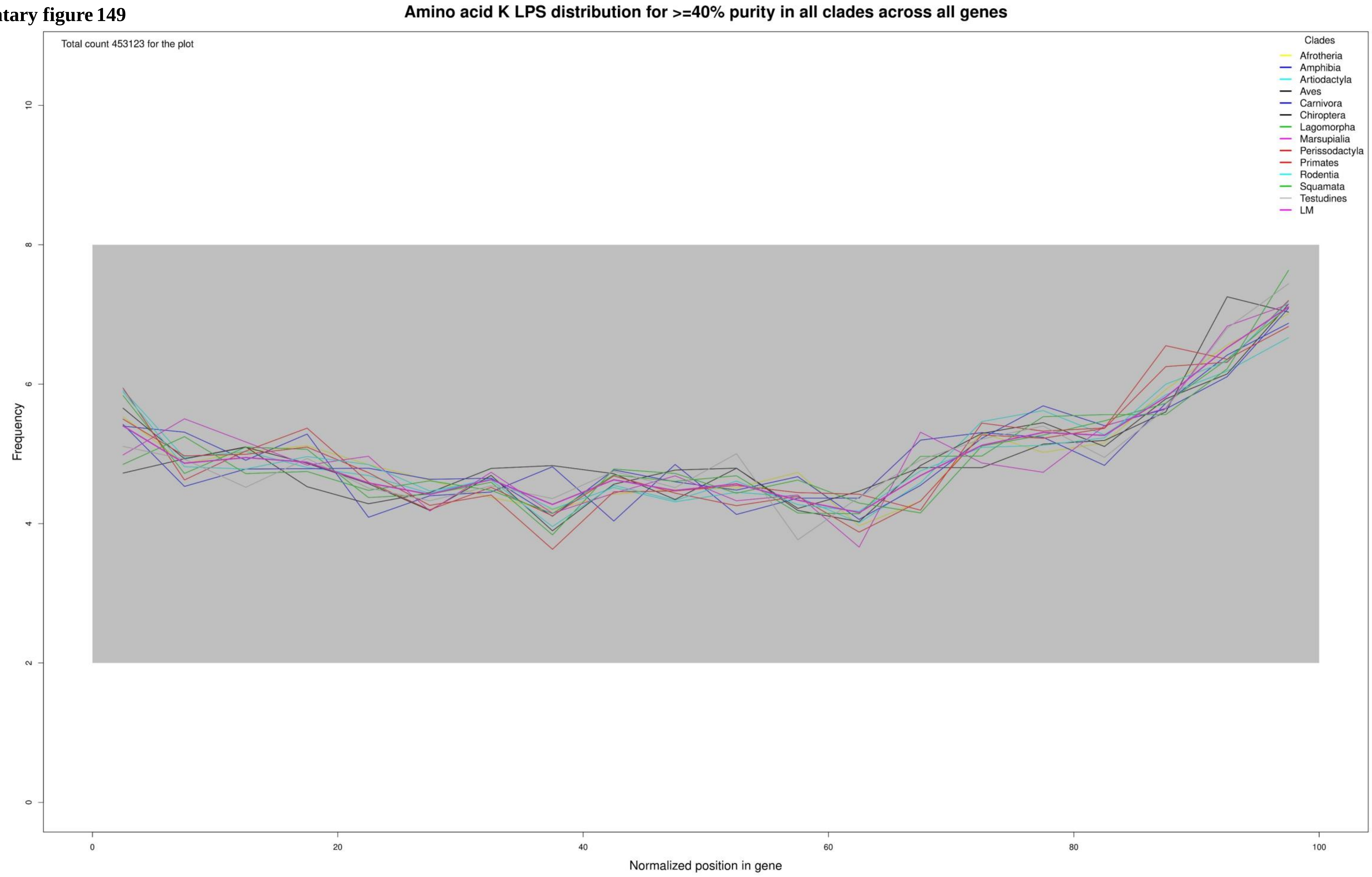

**Supplementary figure 150**

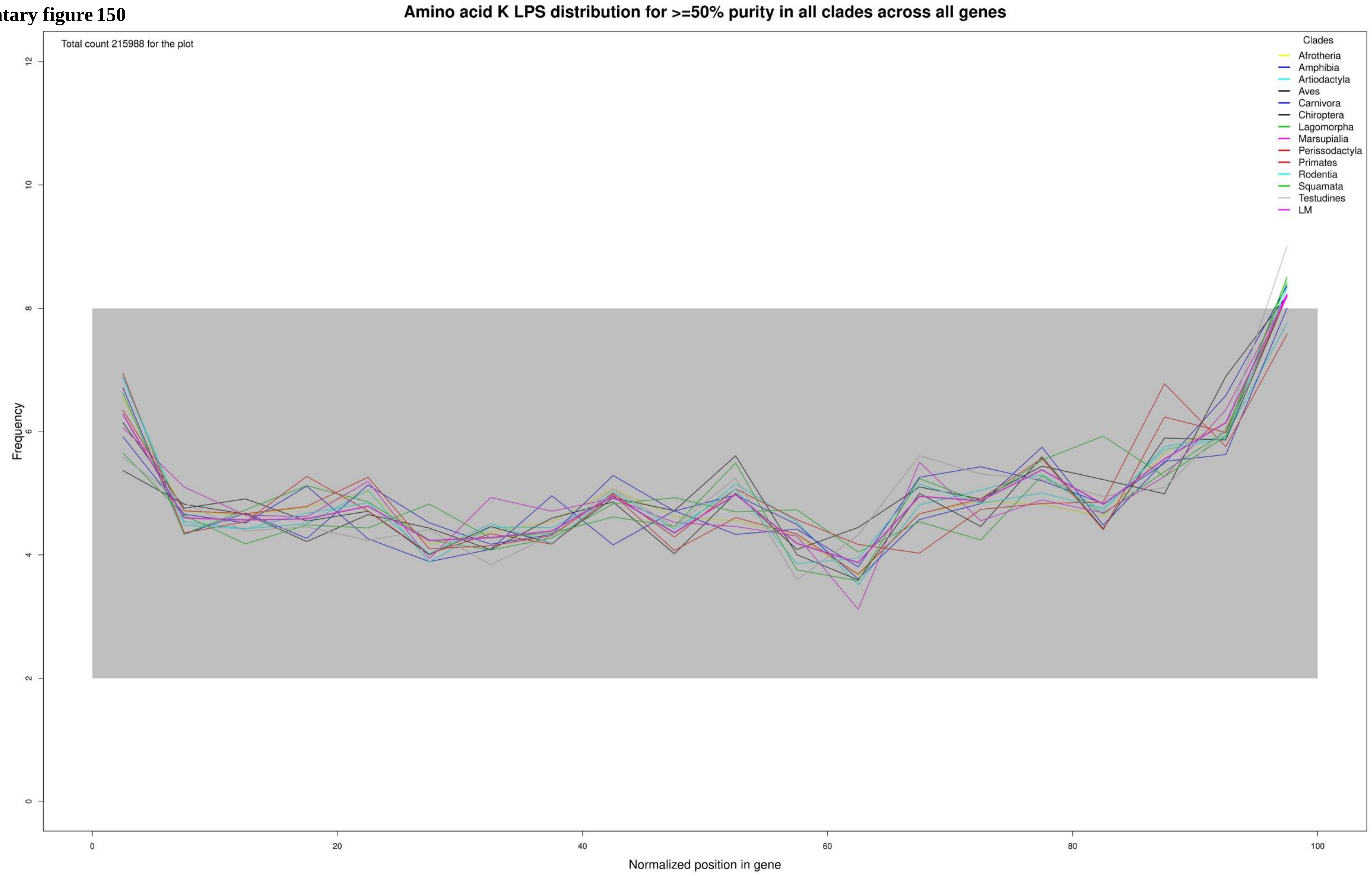

**Supplementary figure 151**

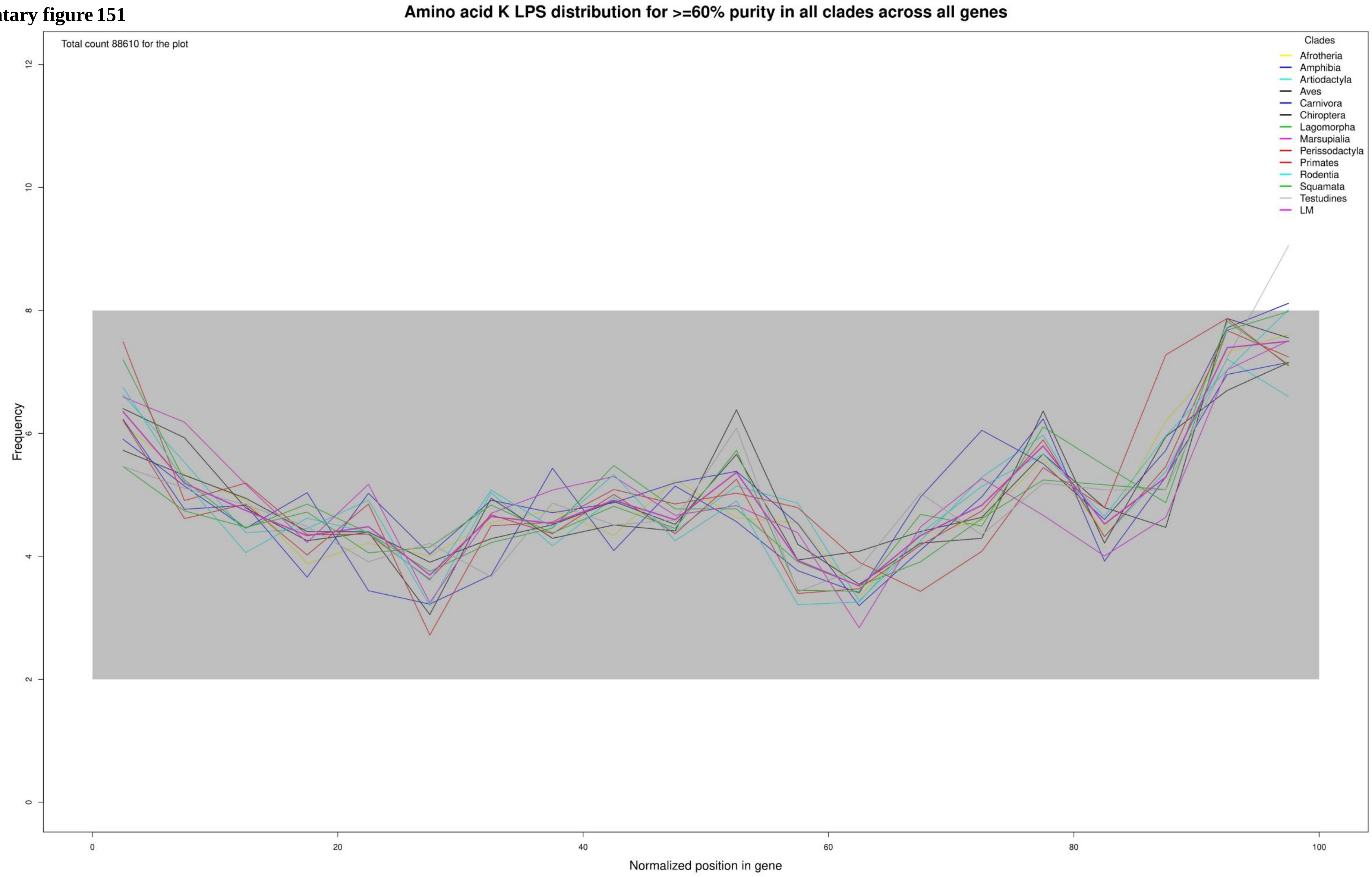

**Supplementary figure 152**

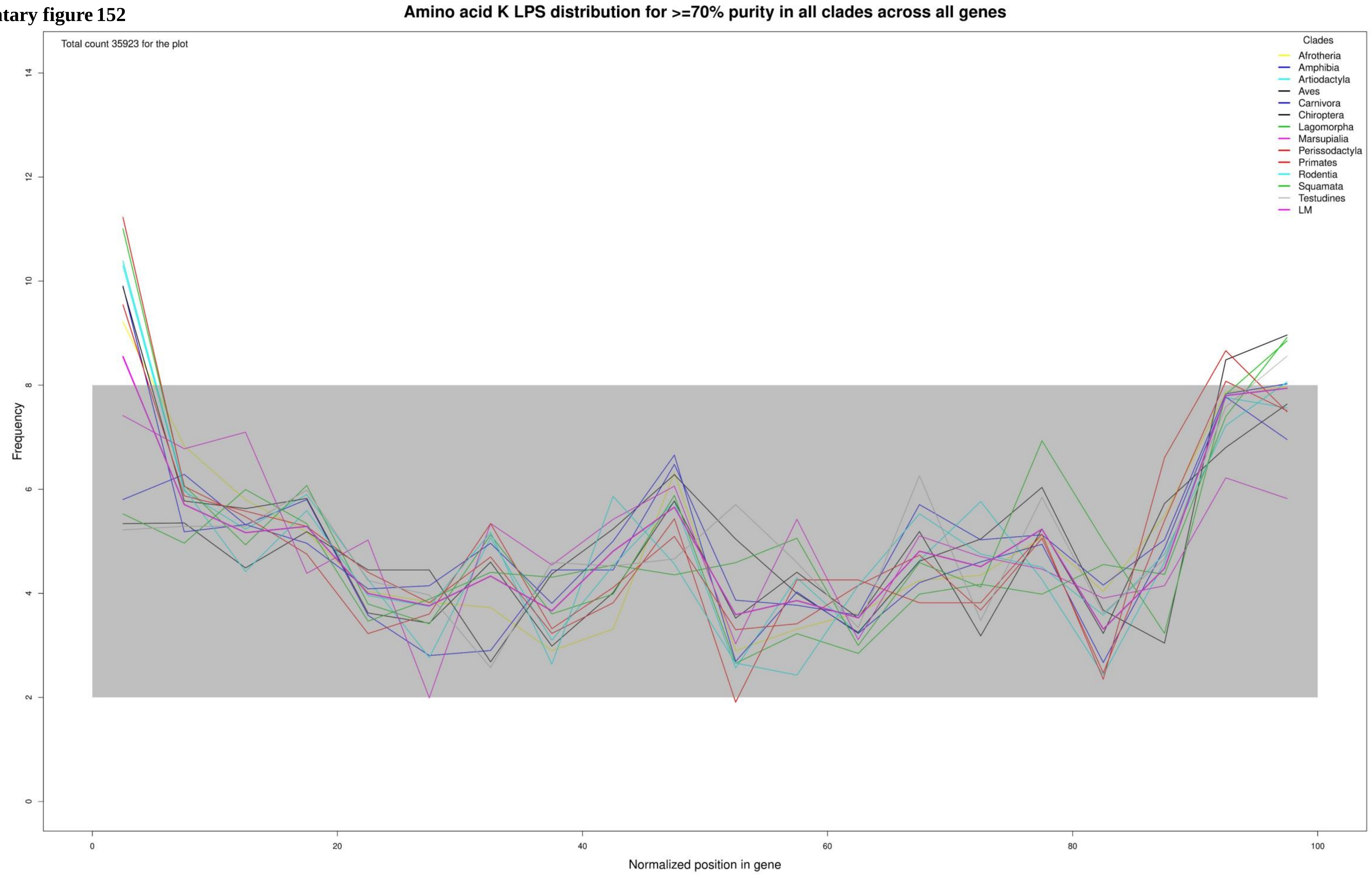

**Supplementary figure 153**

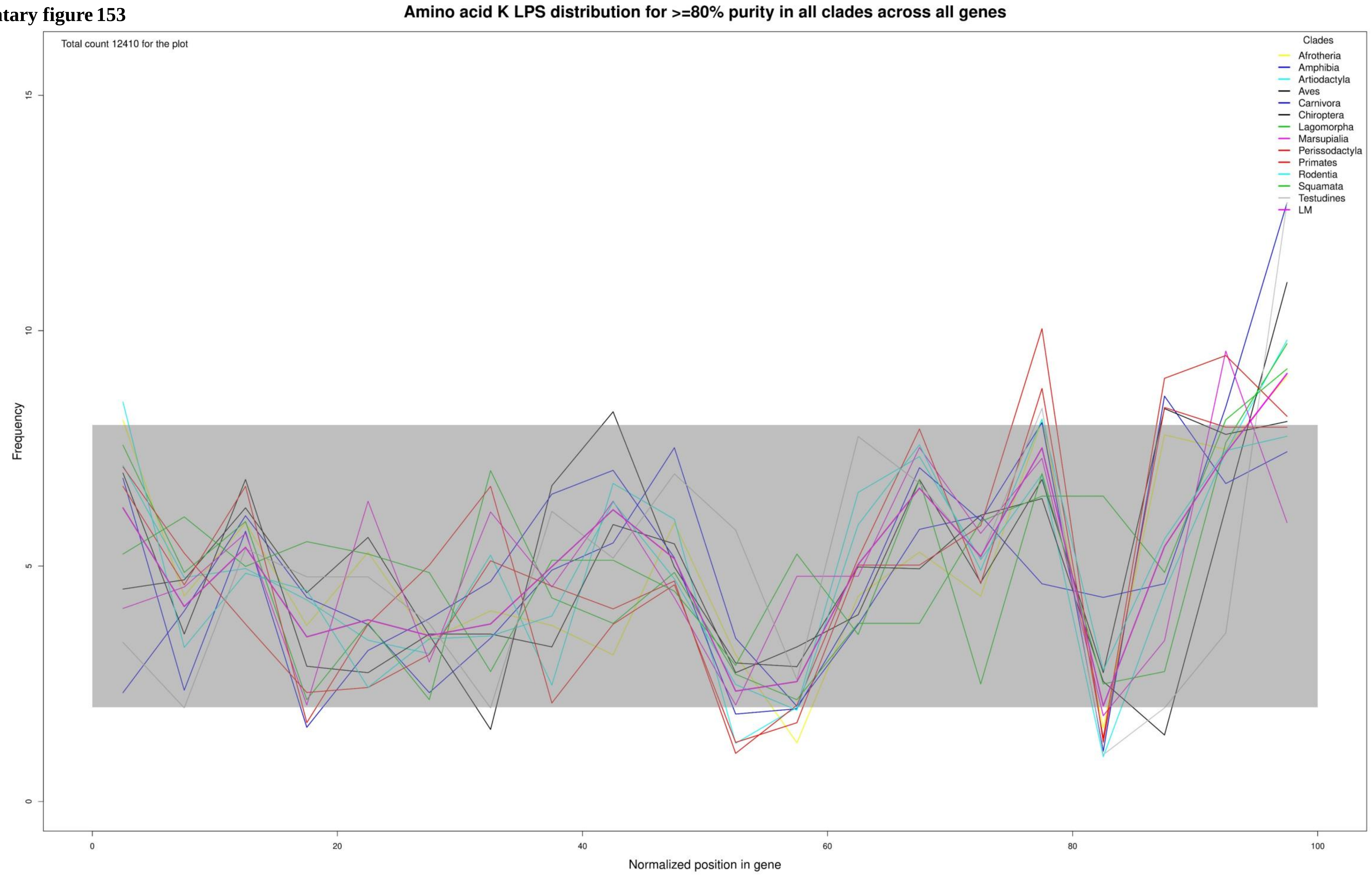

**Supplementary figure 154**

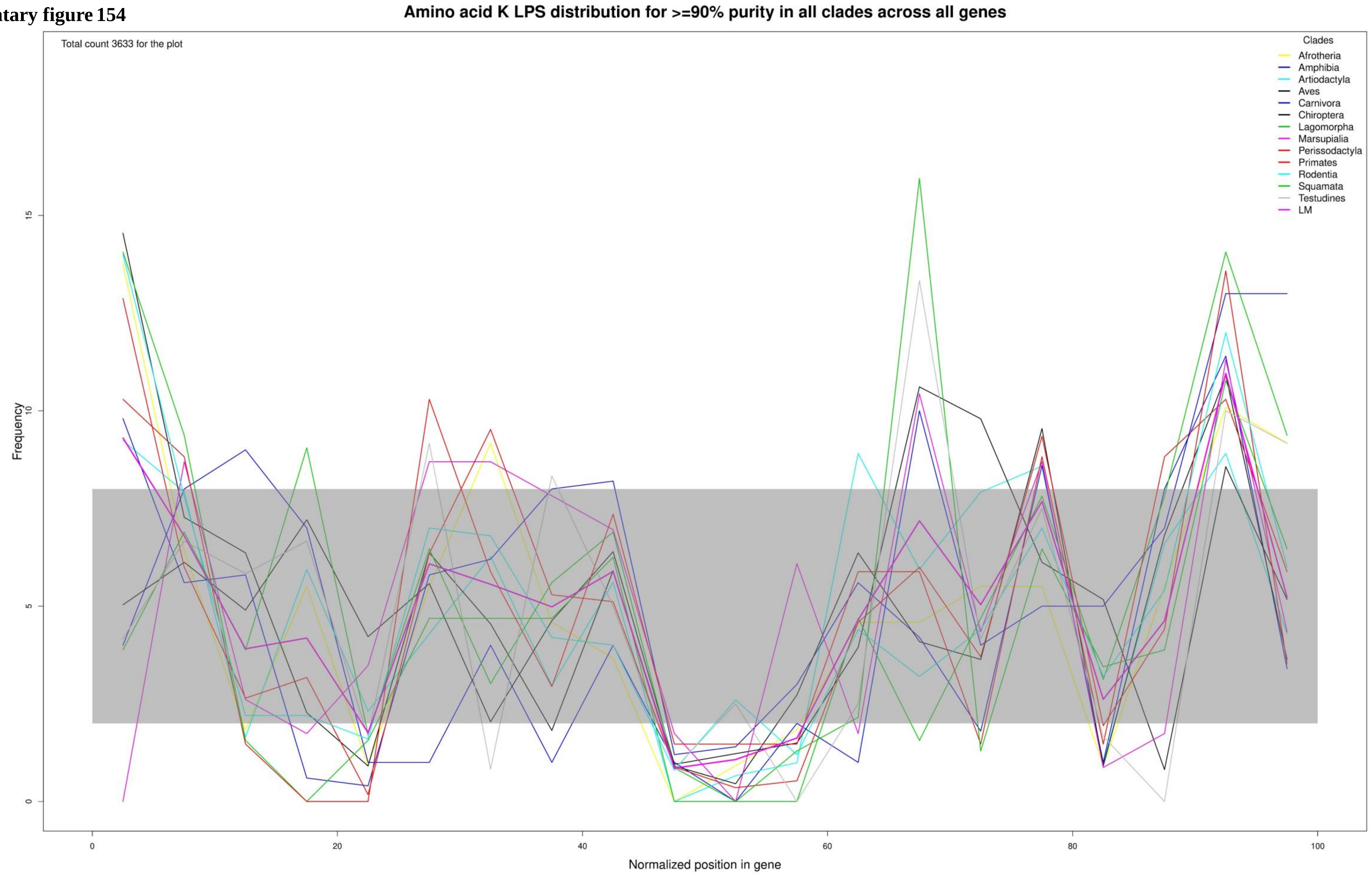

**Supplementary figure 155**

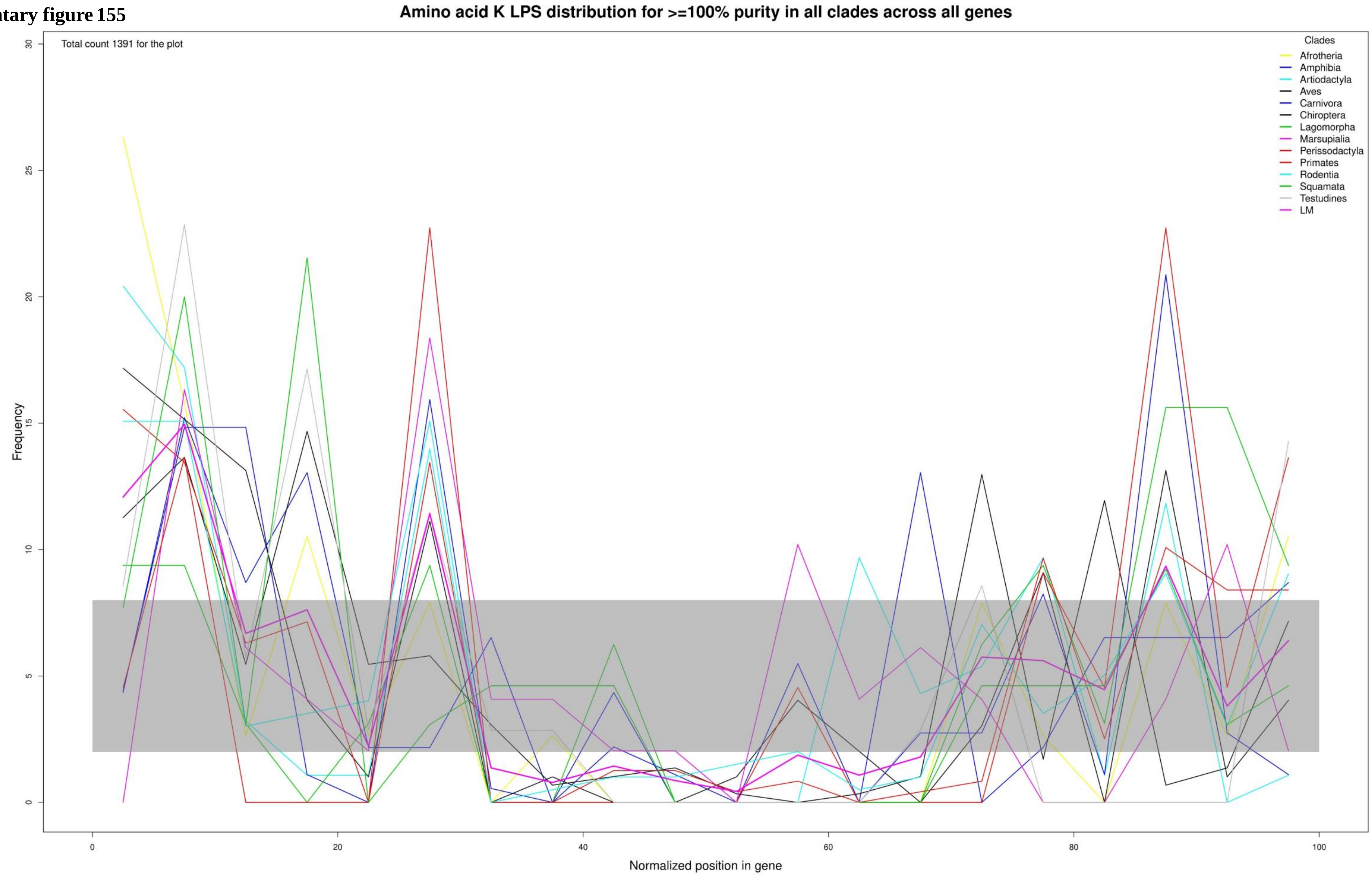

**Supplementary figure 156**

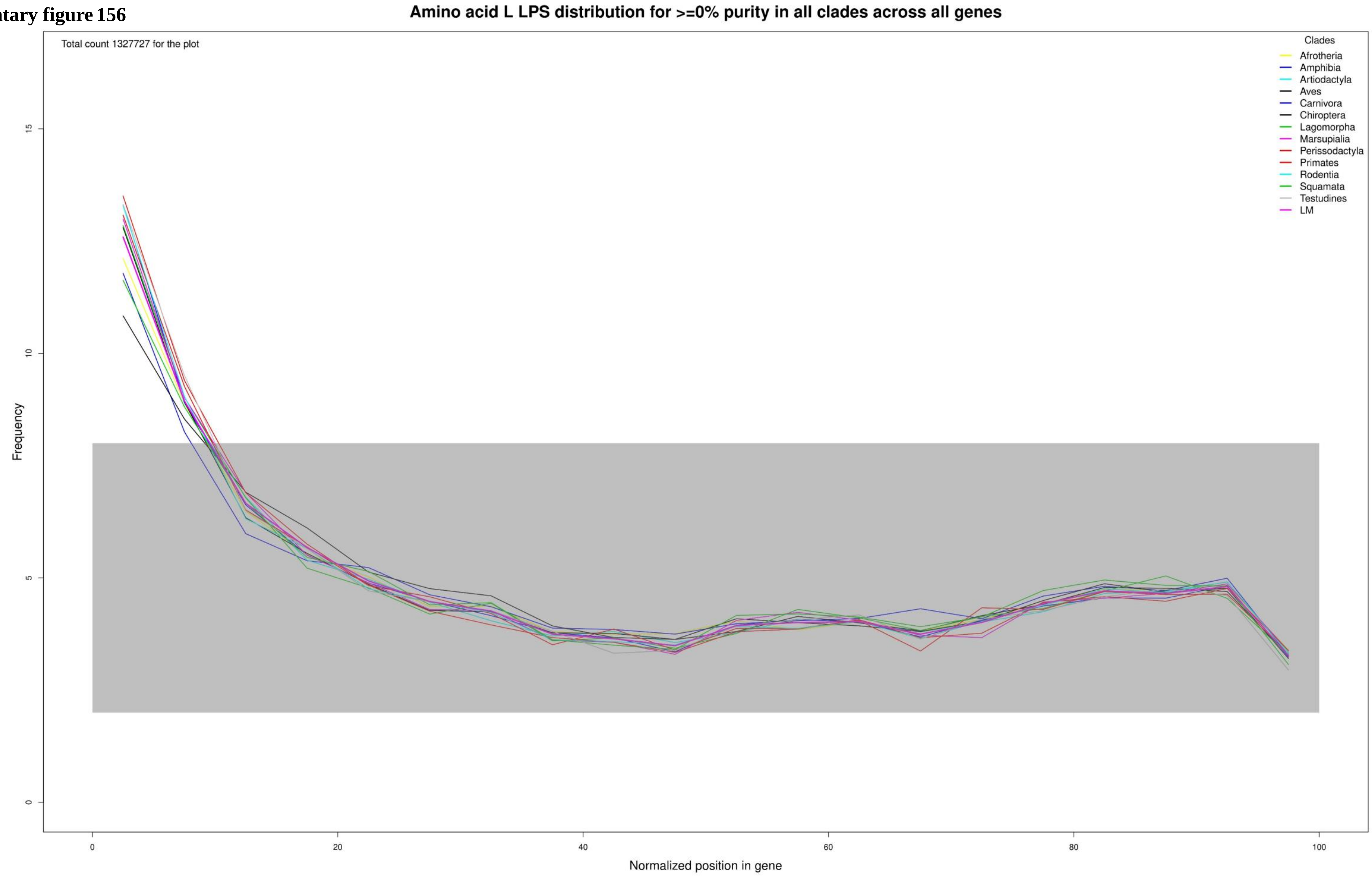

Supplementary figure 157

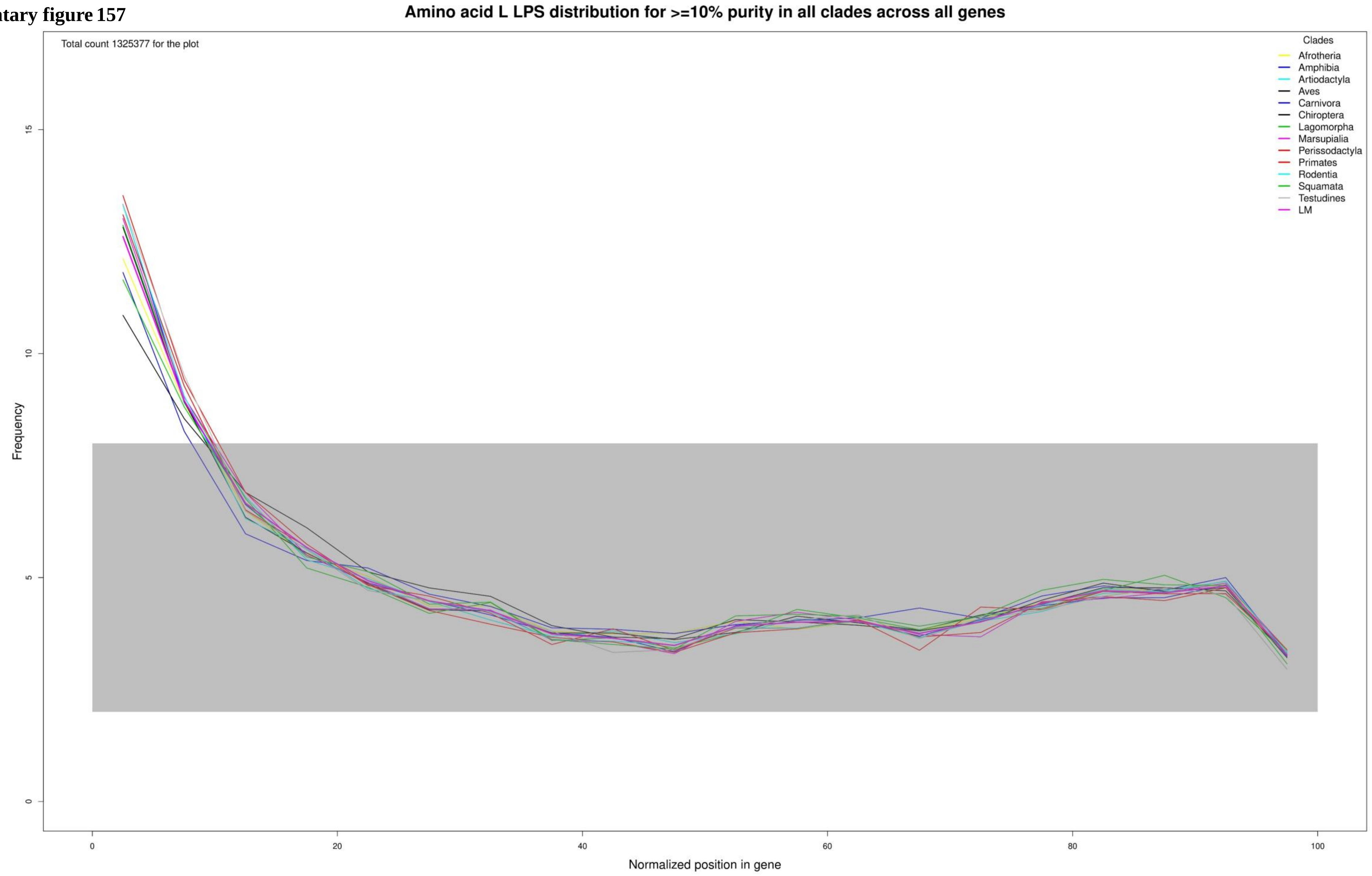

Supplementary figure 158

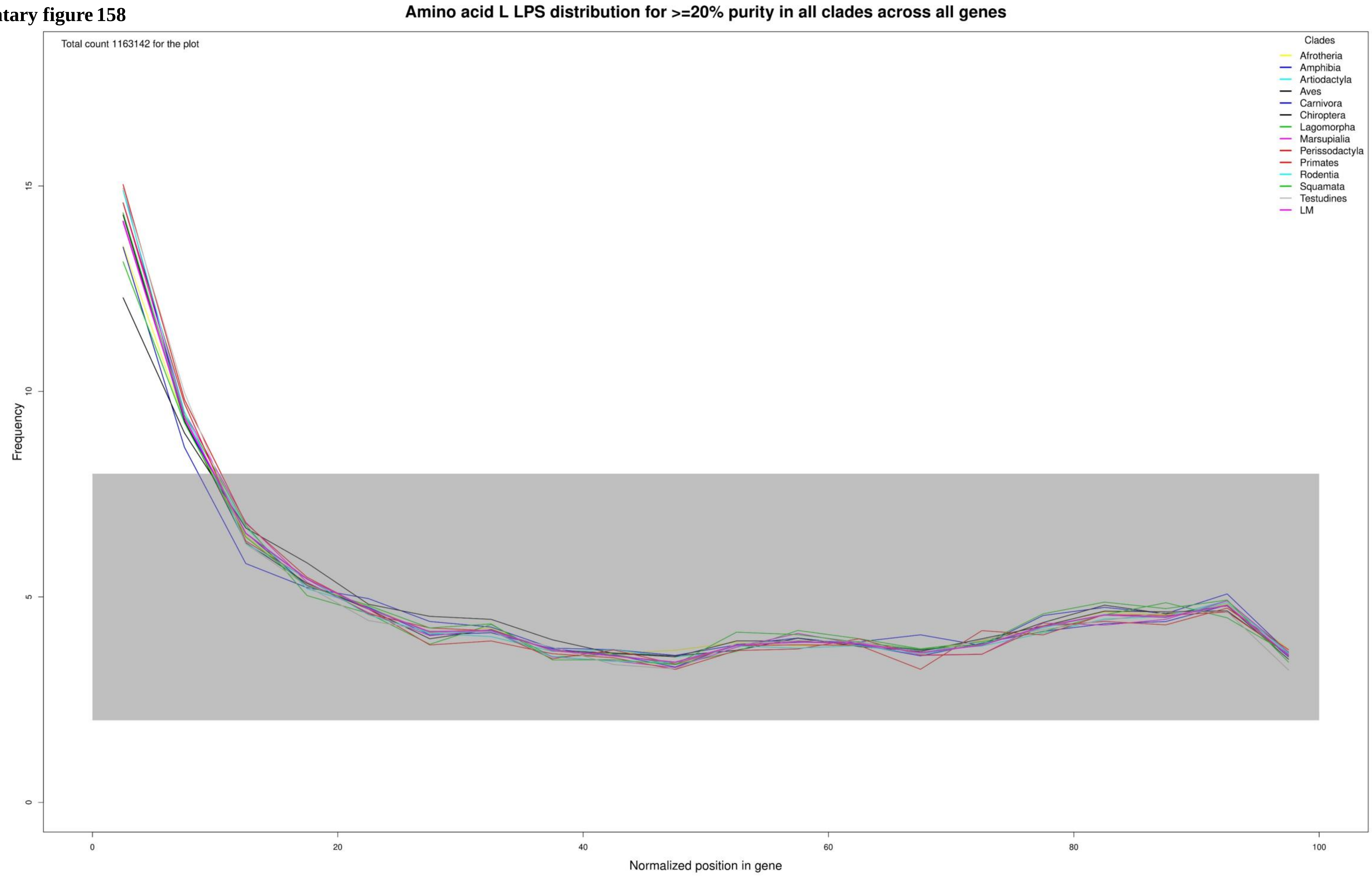

**Supplementary figure 159**

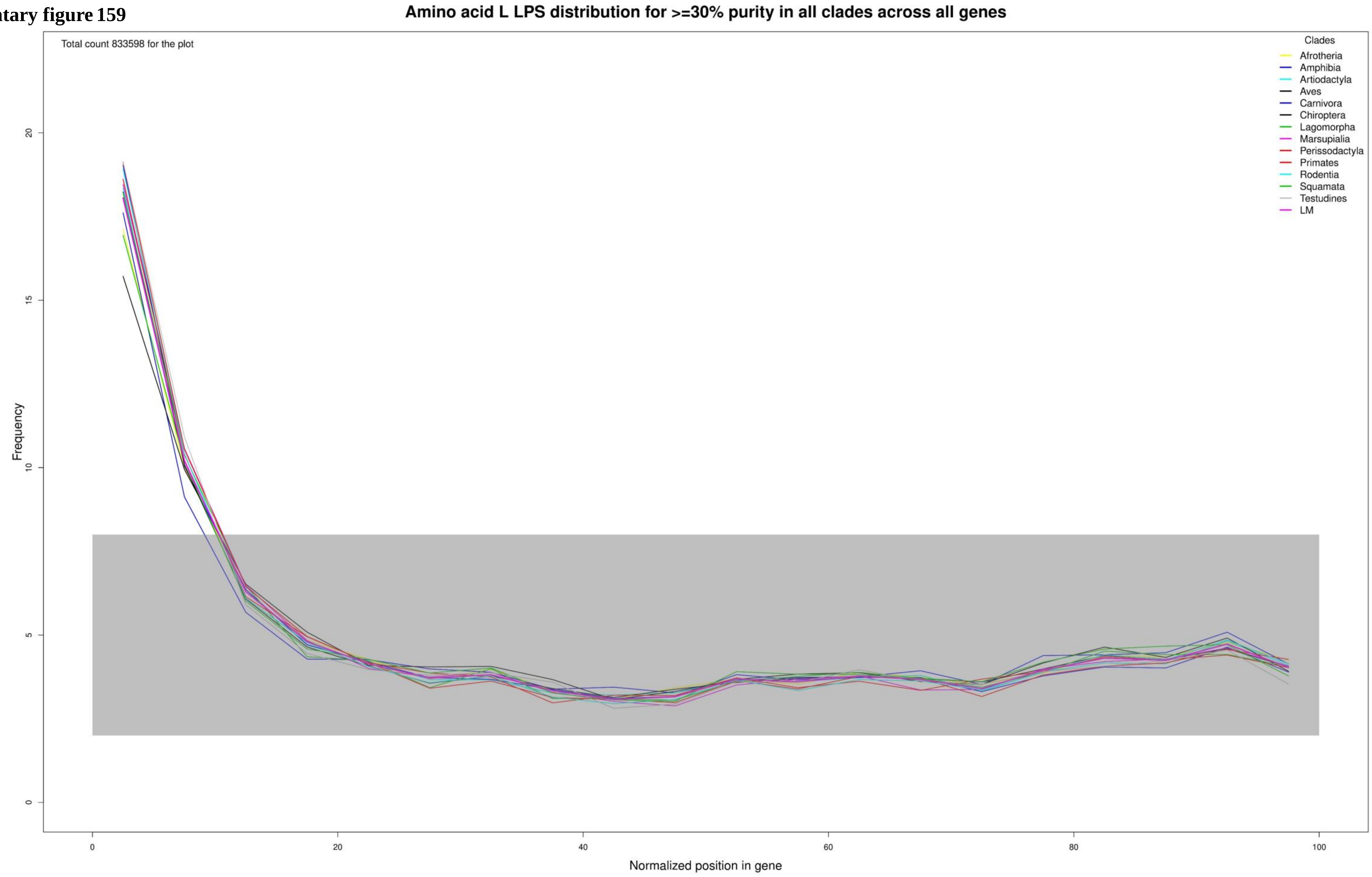

Supplementary figure 160

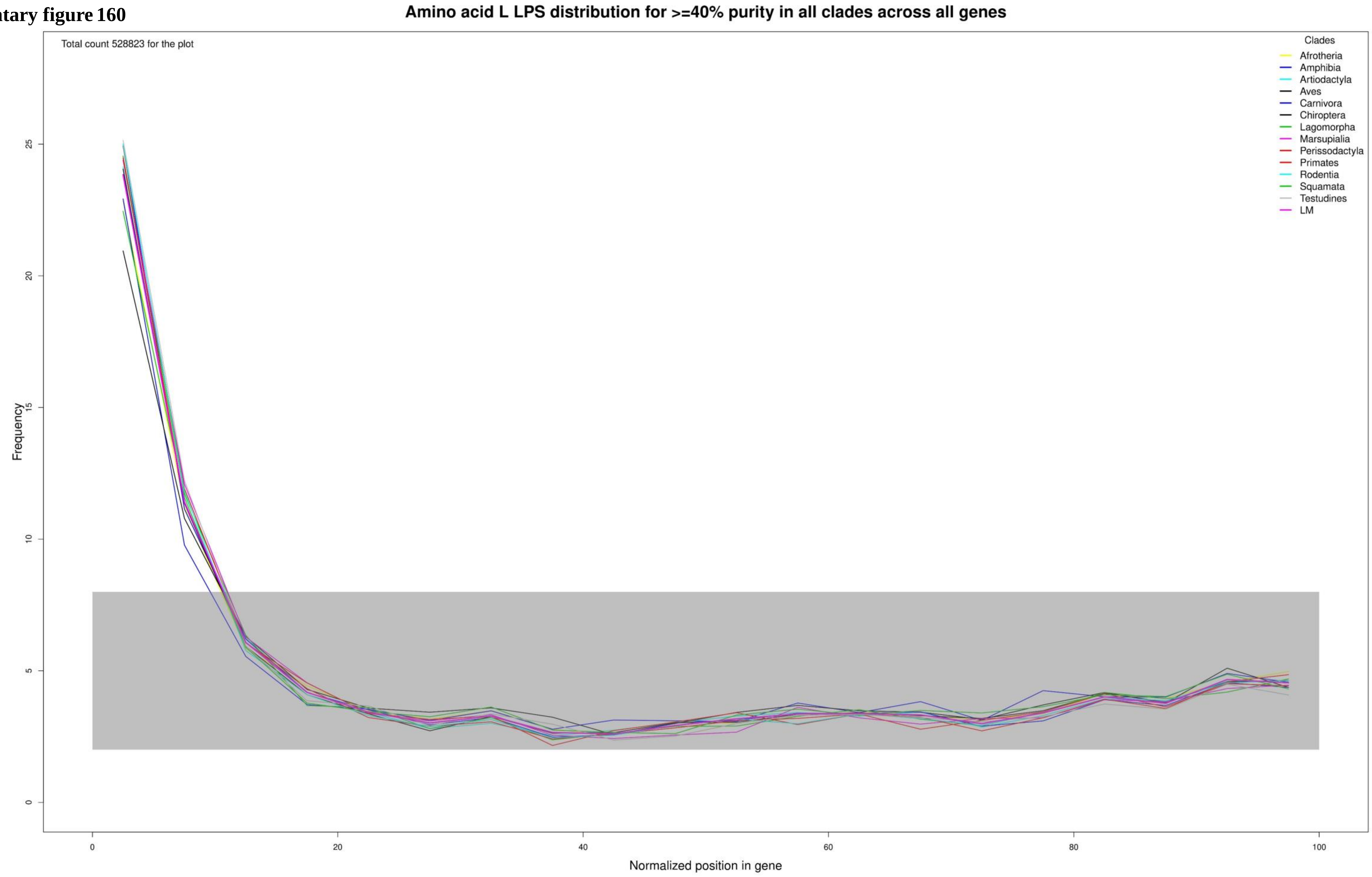

**Supplementary figure 161**

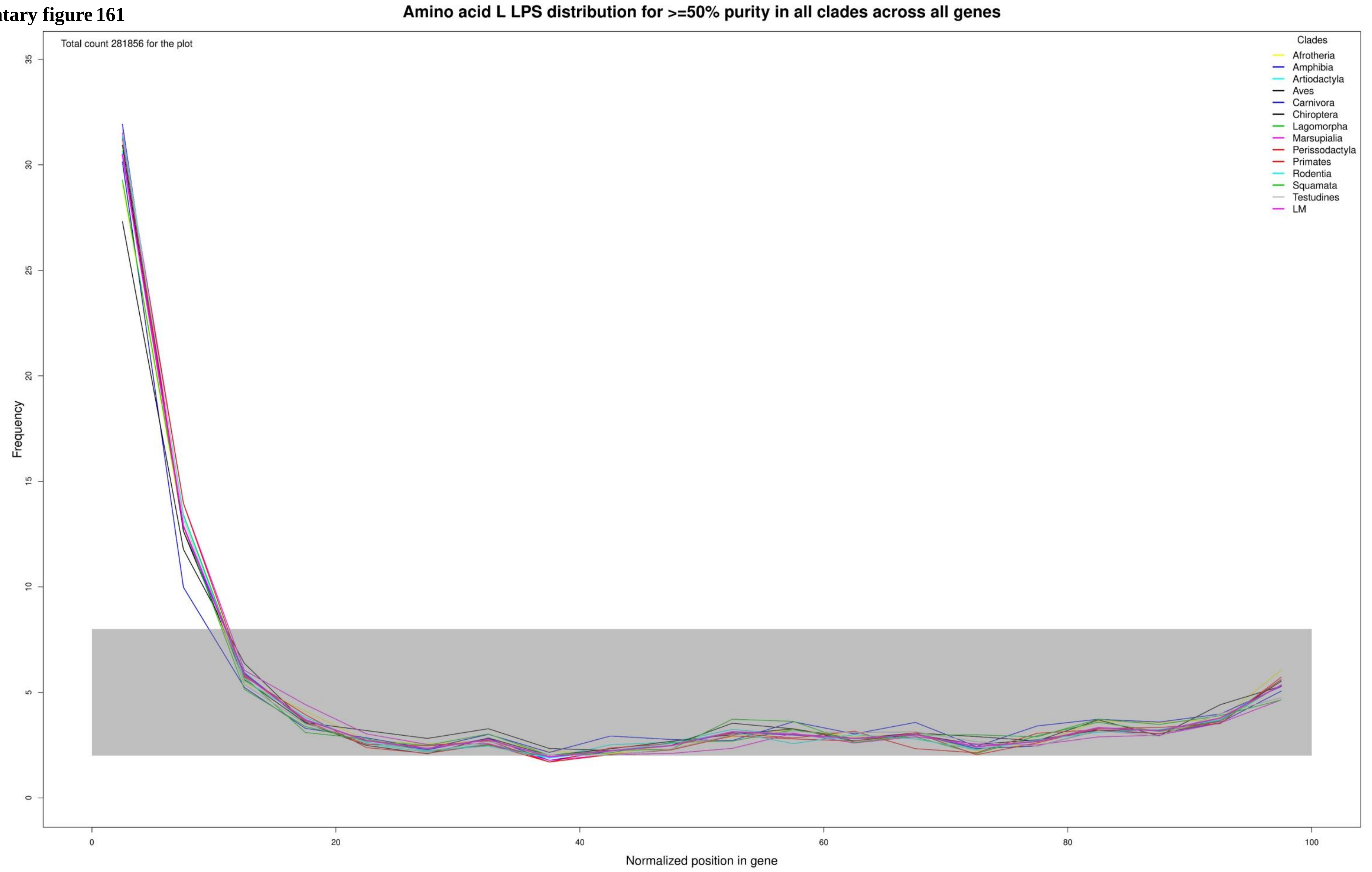

**Supplementary figure 162**

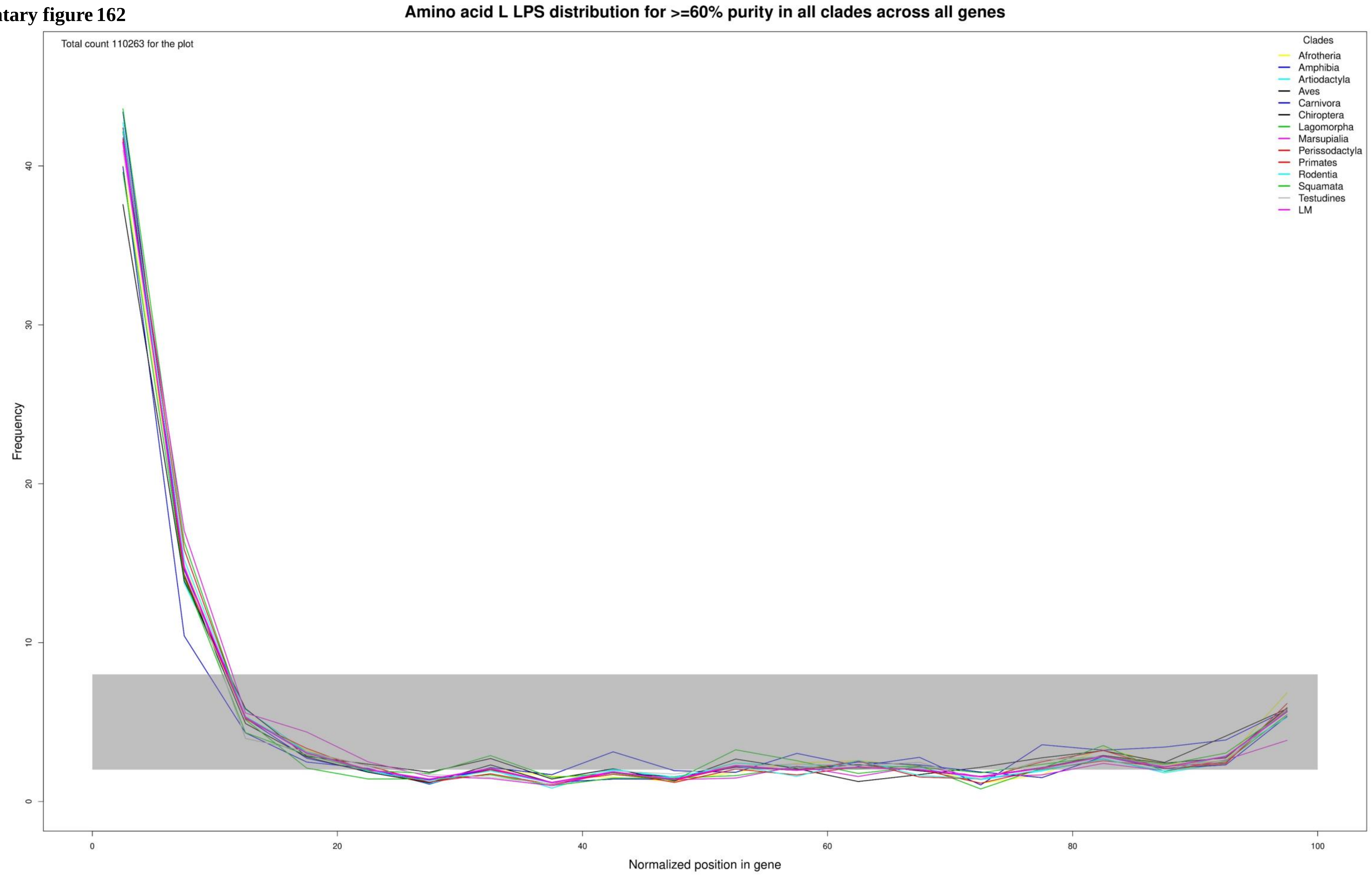

**Supplementary figure 163**

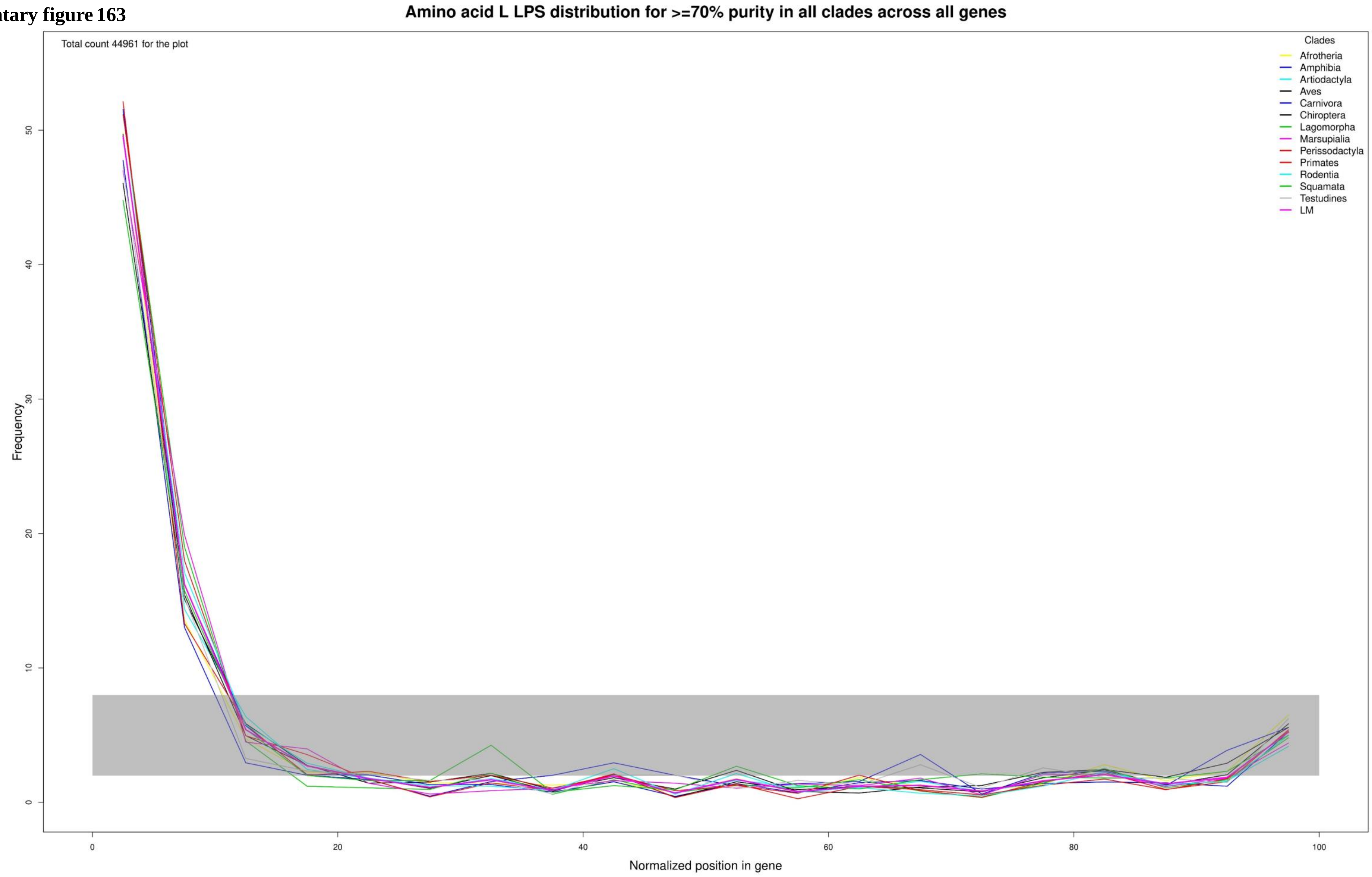

**Supplementary figure 164**

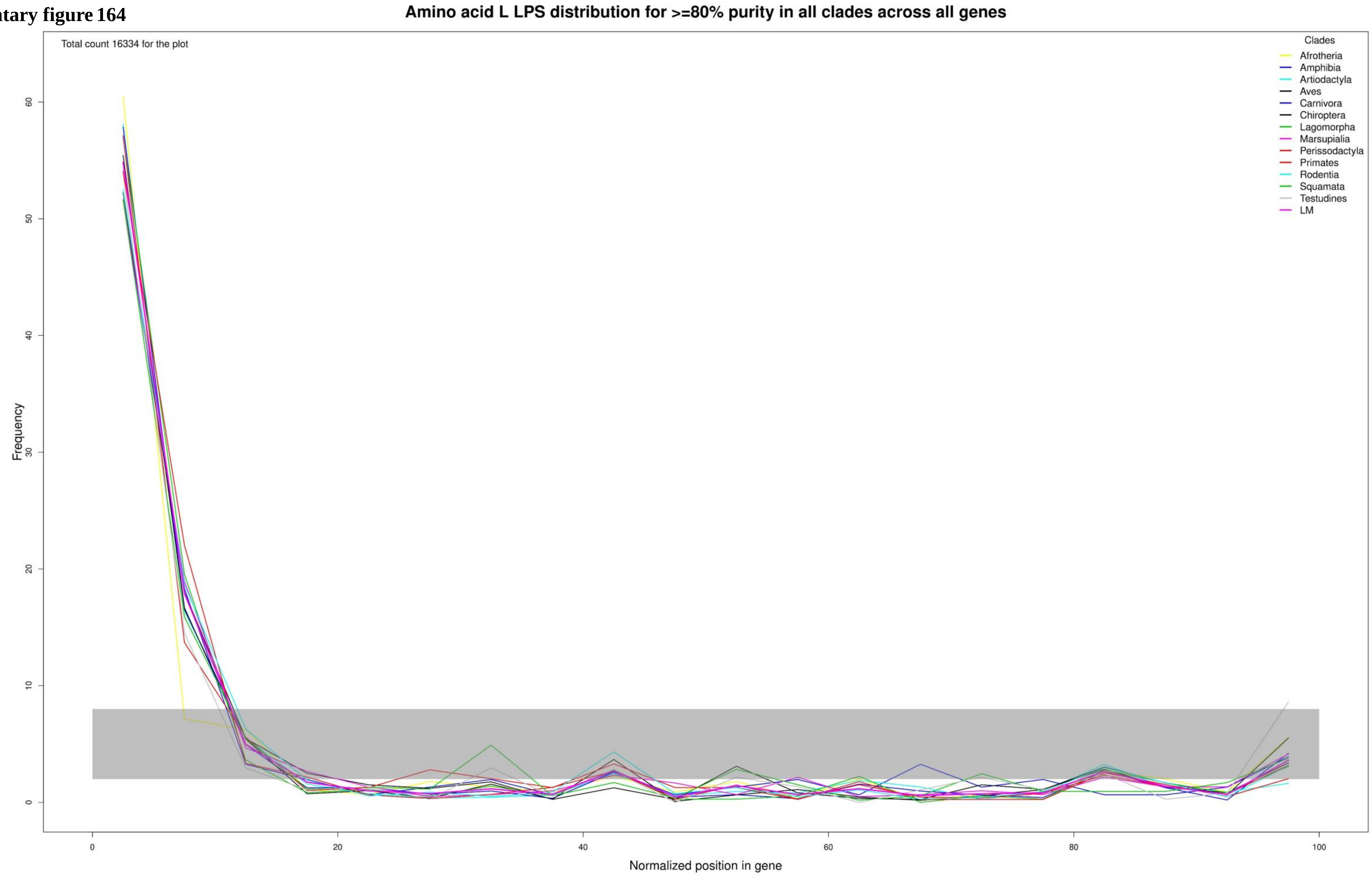

**Supplementary figure 165**

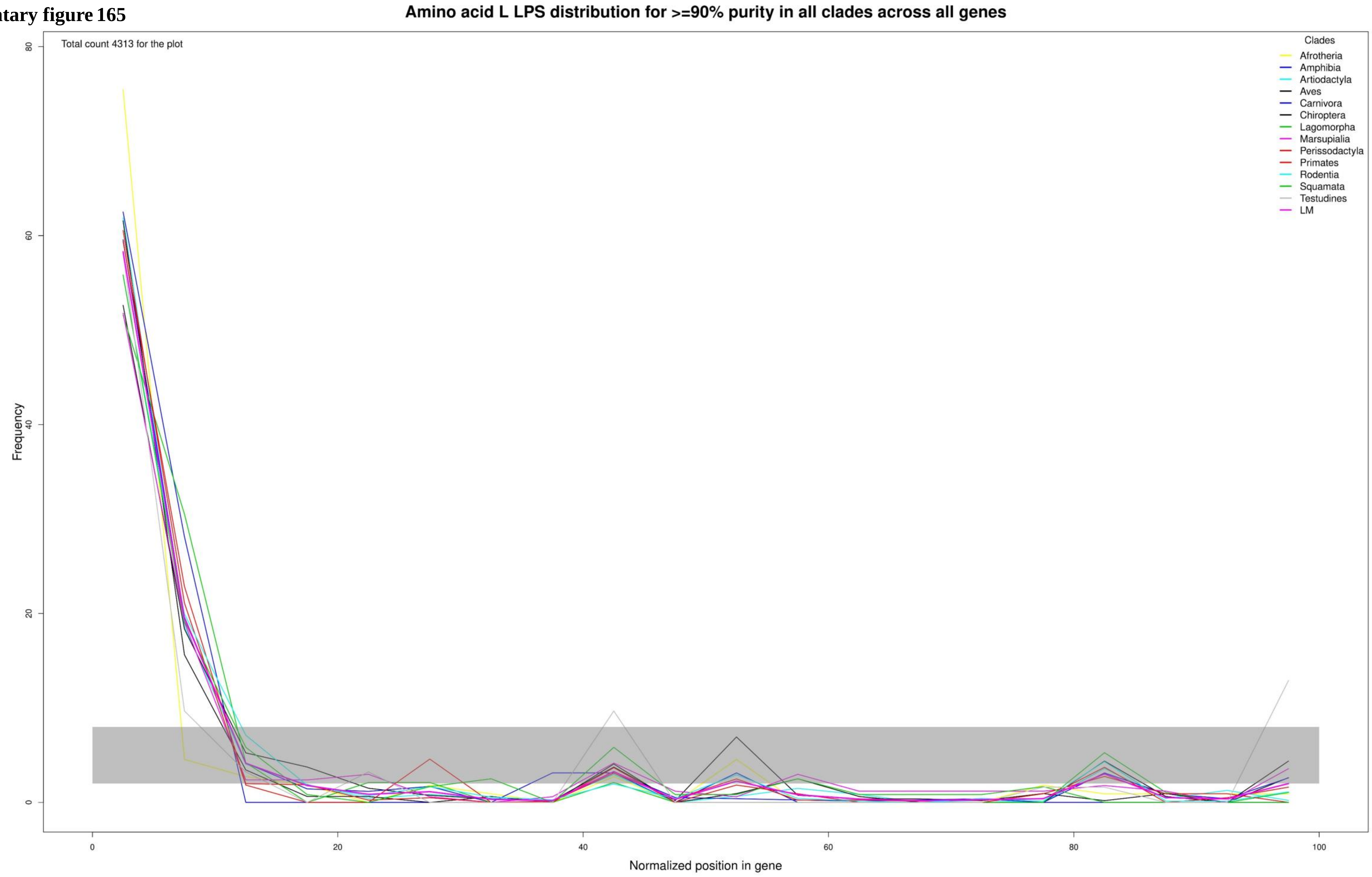

**Supplementary figure 166**

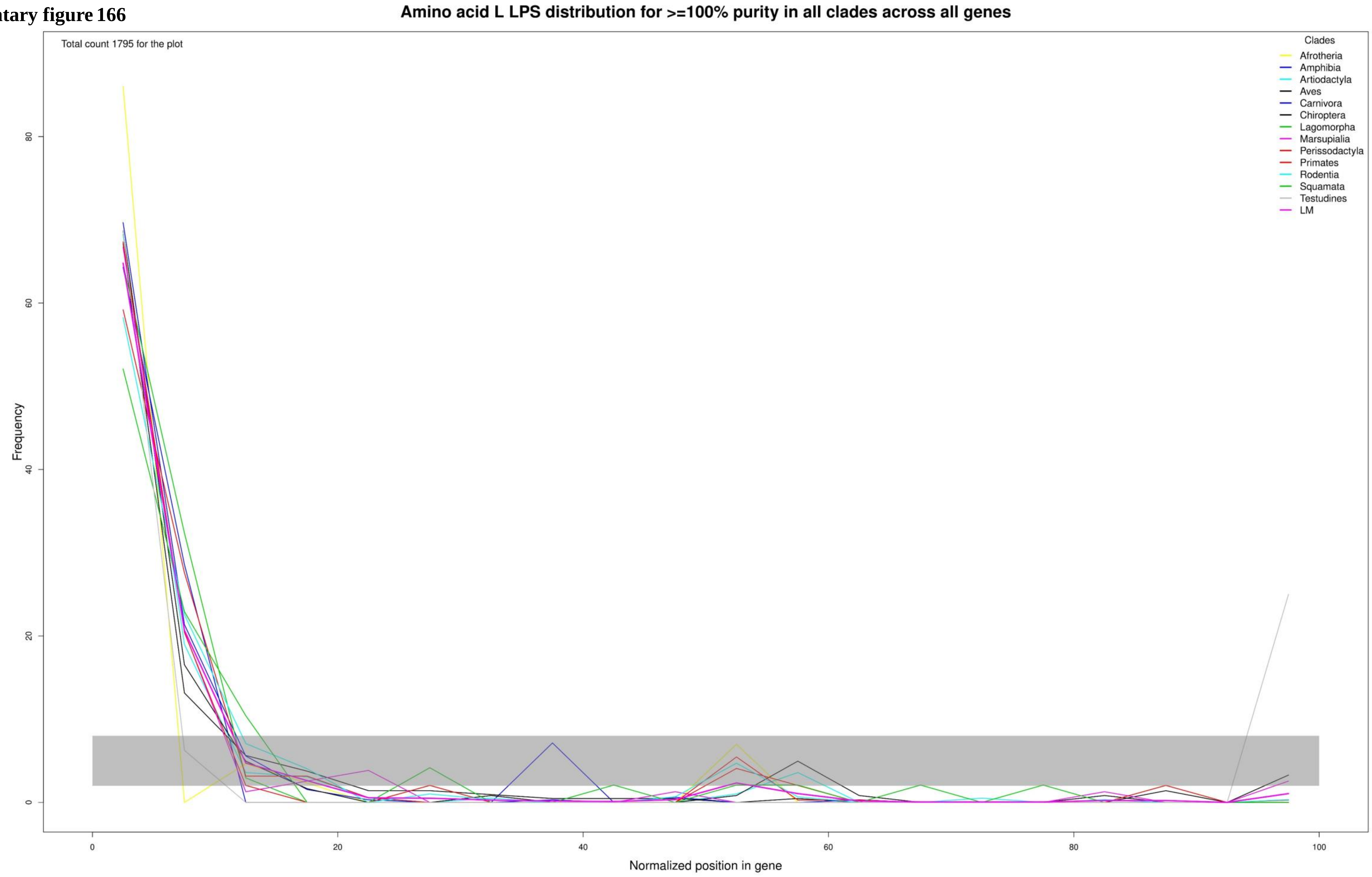

**Supplementary figure 167**

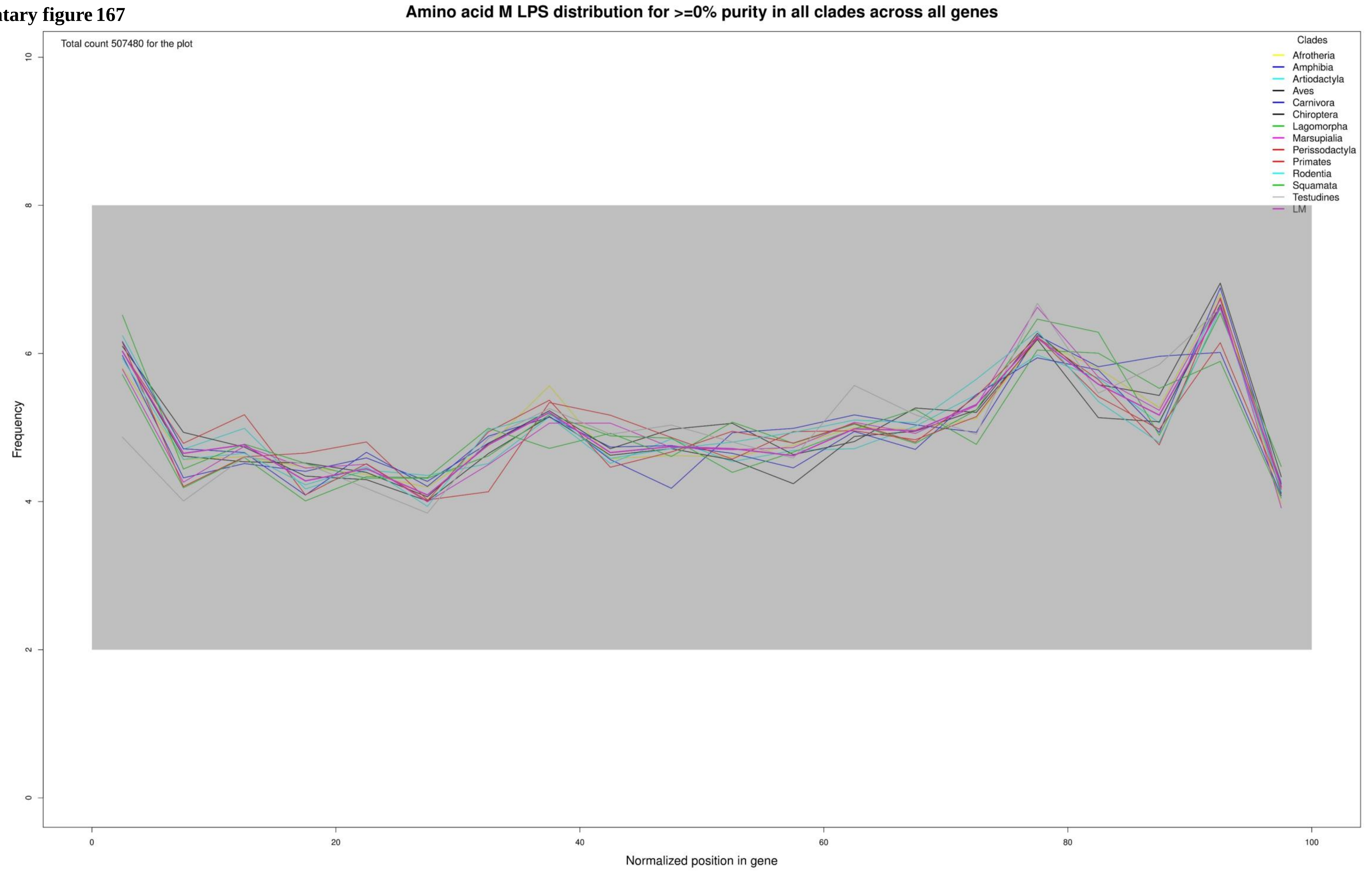

**Supplementary figure 168**

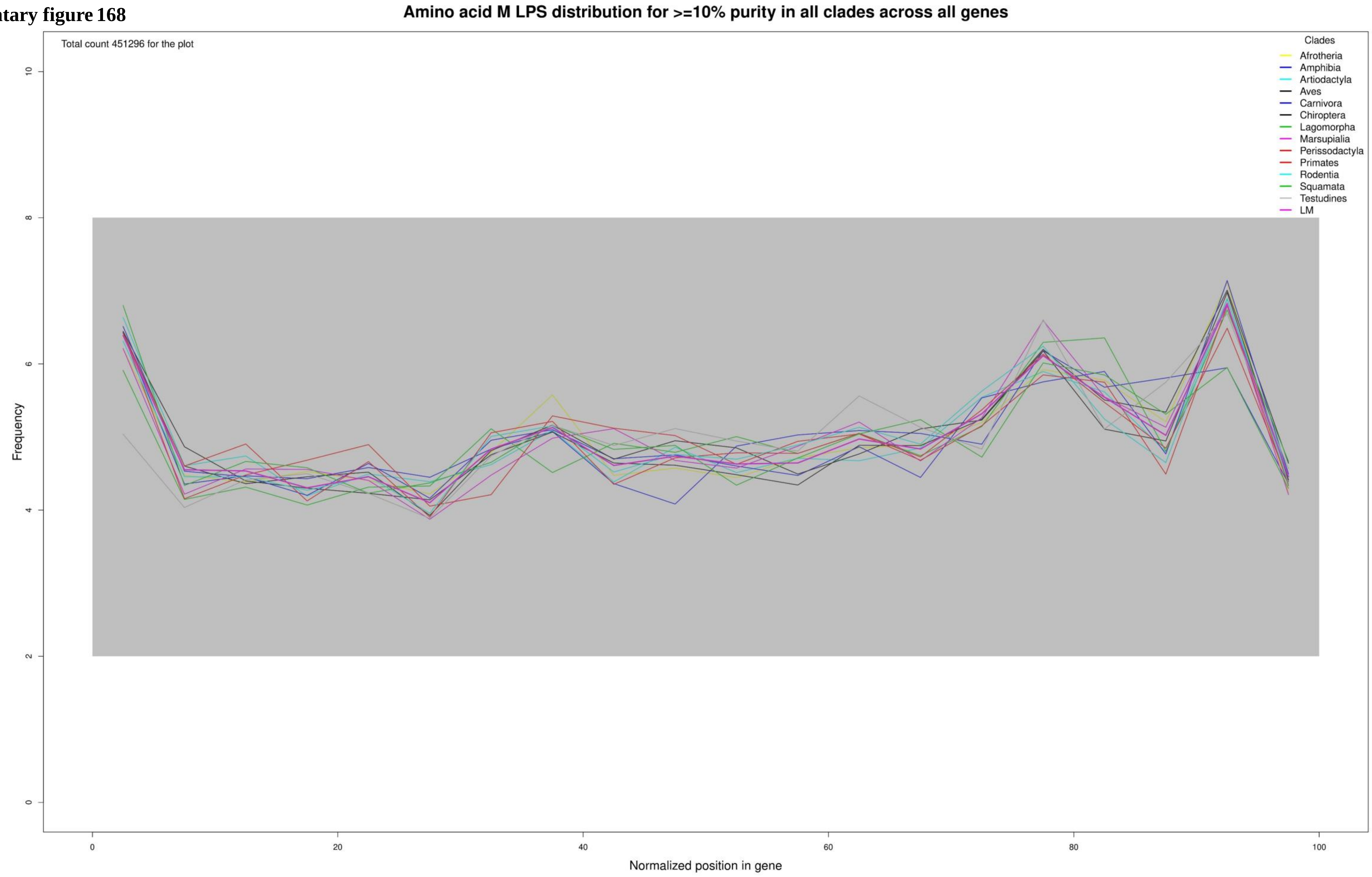

**Supplementary figure 169**

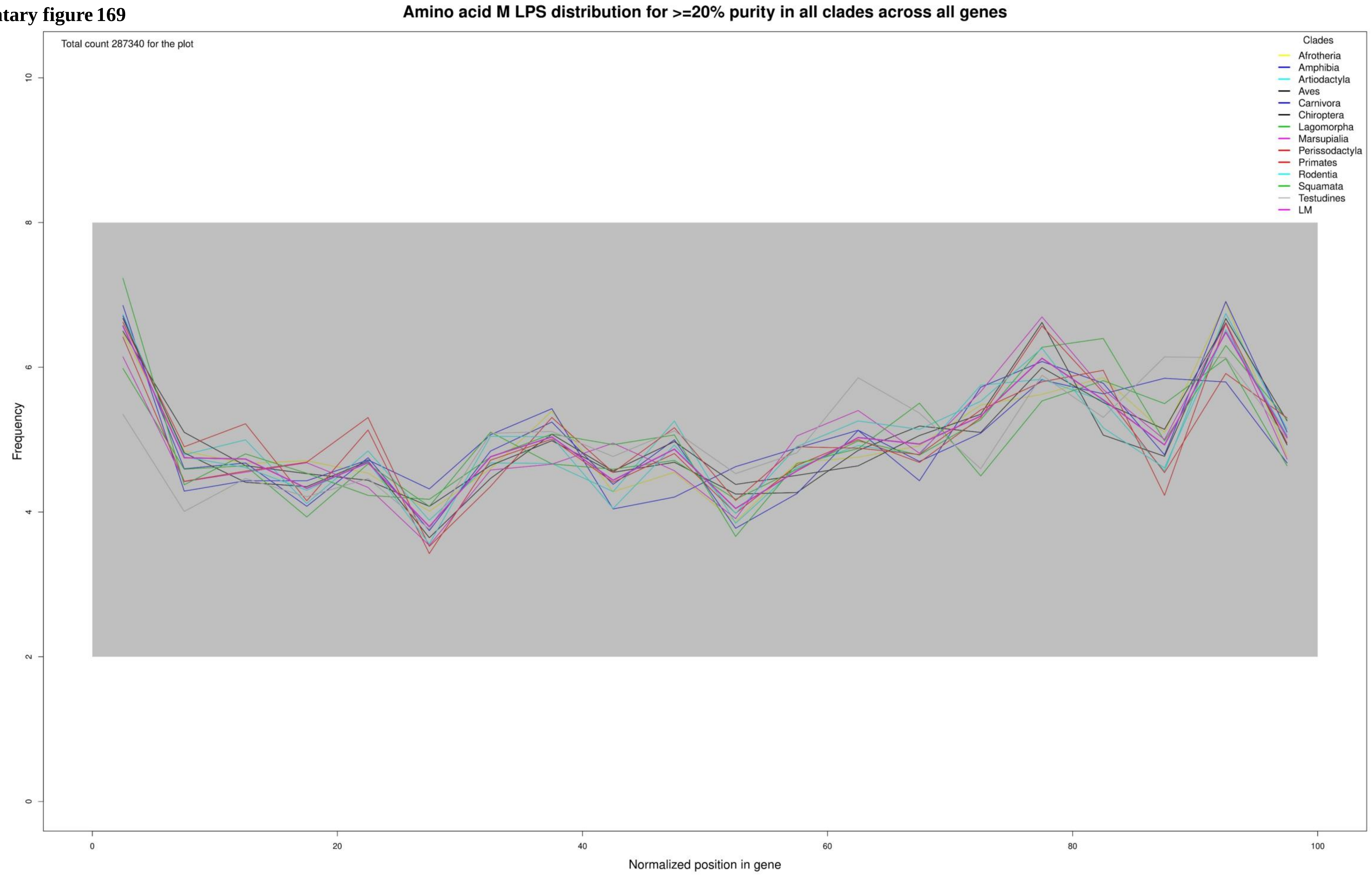

**Supplementary figure 170**

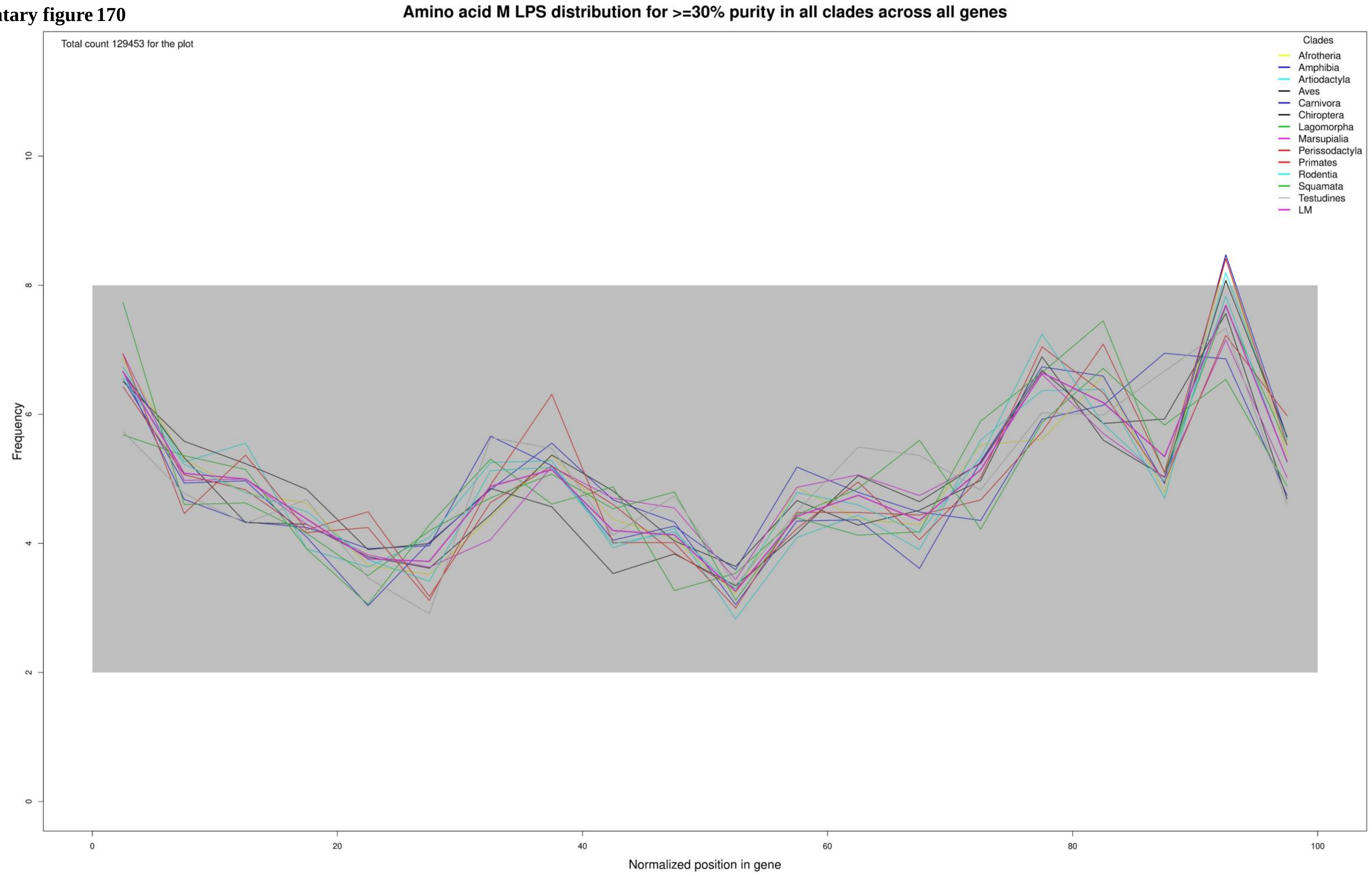

**Supplementary figure 171**

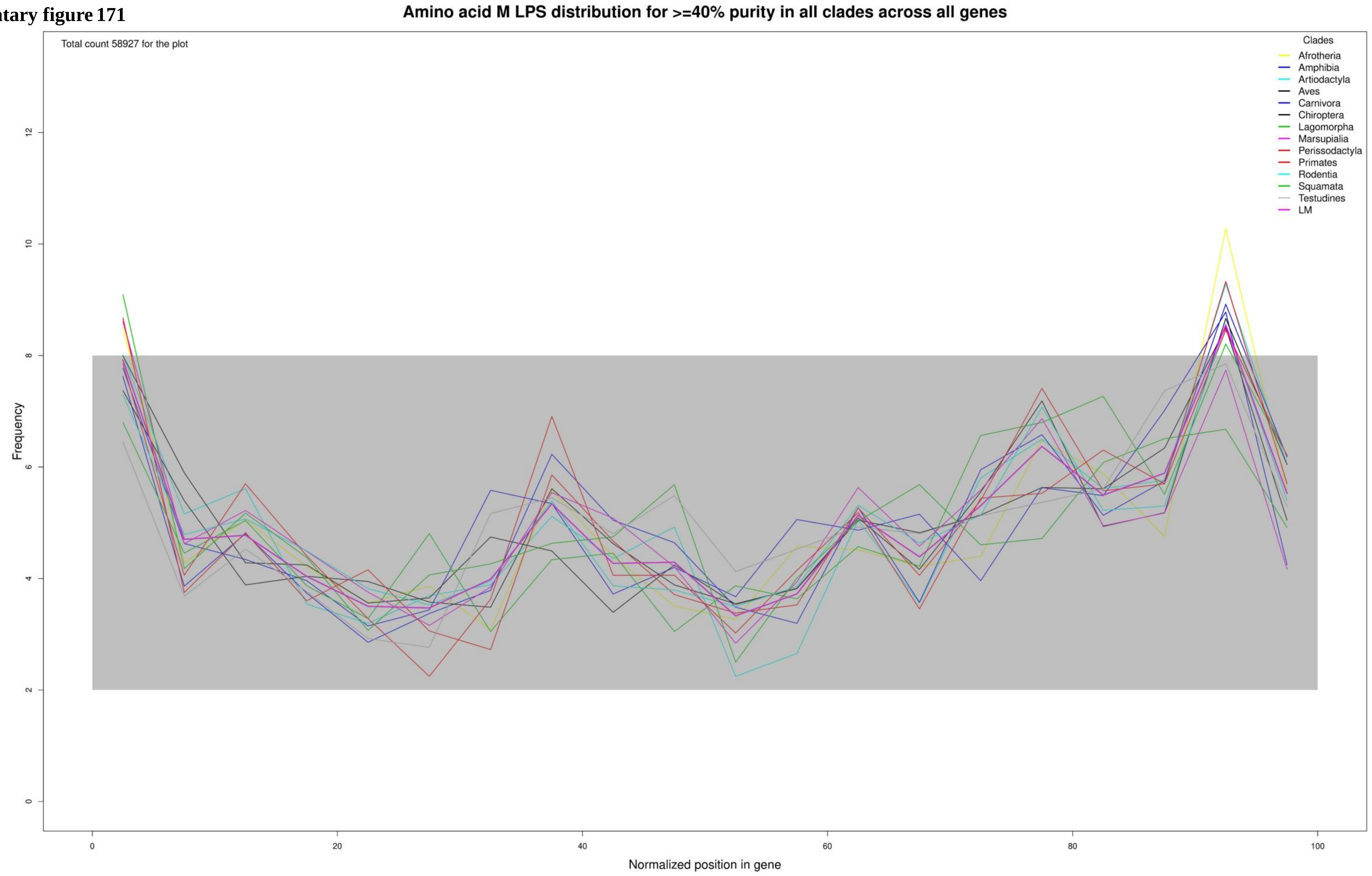

**Supplementary figure 172**

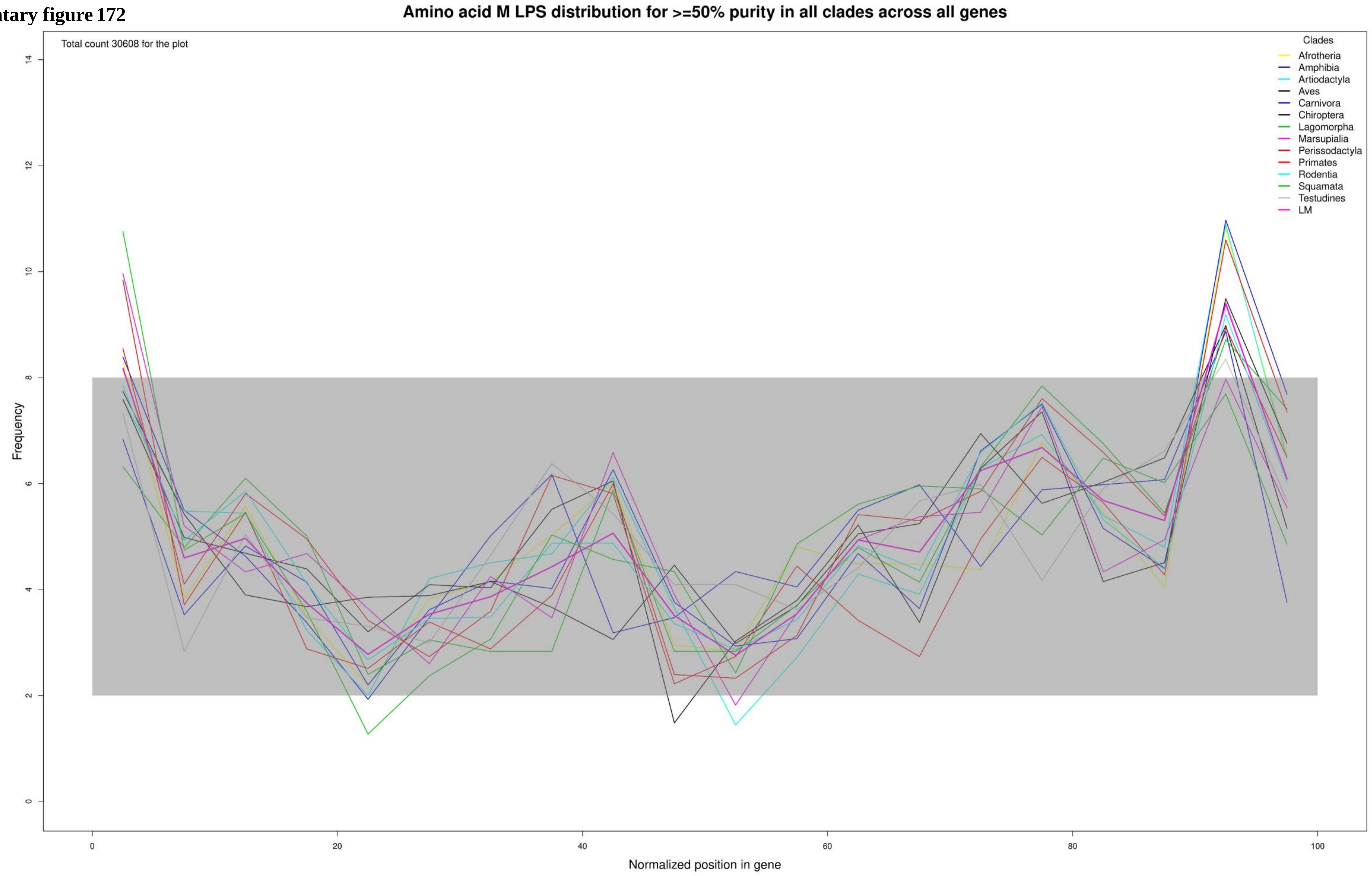

**Supplementary figure 173**

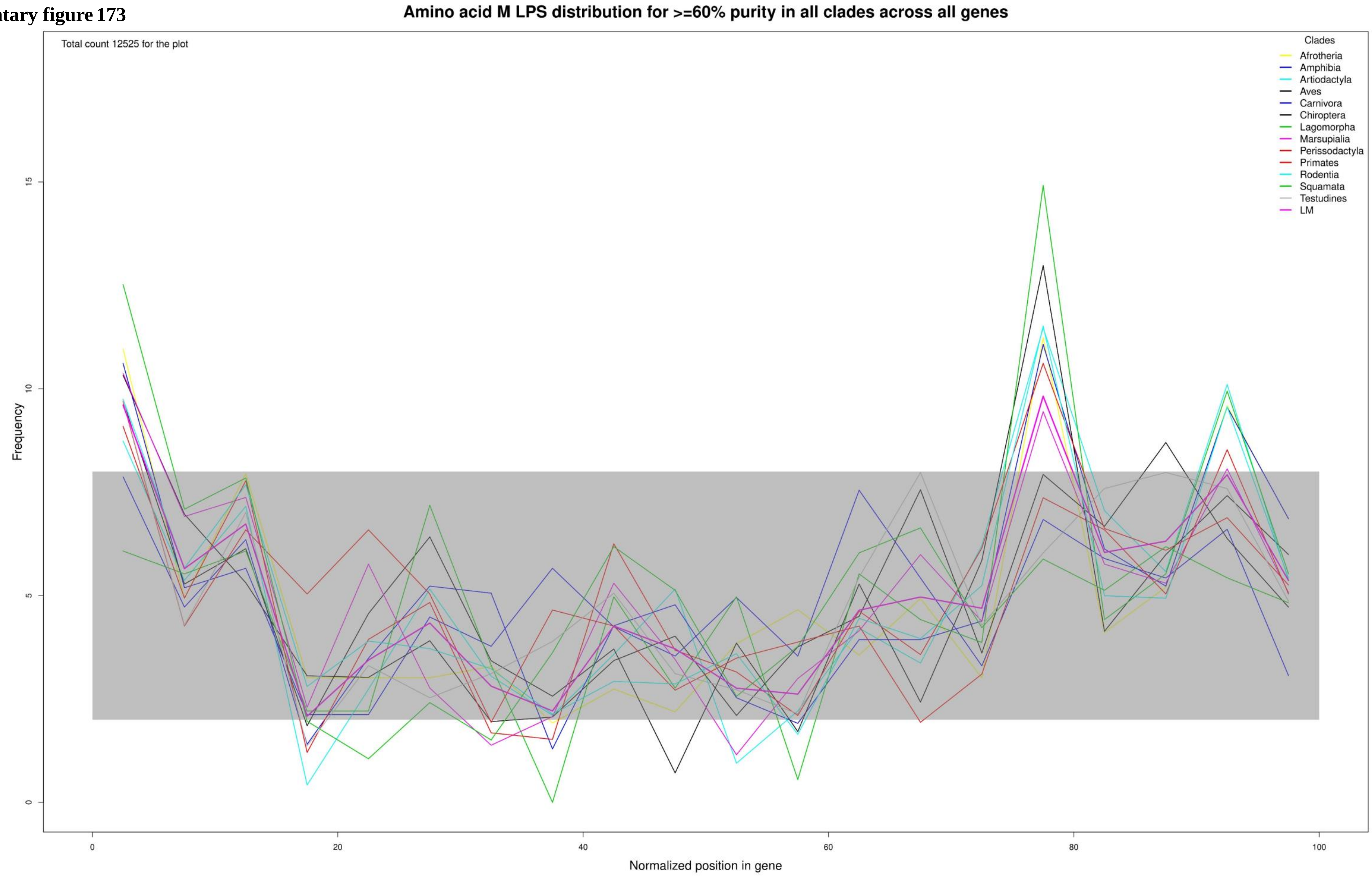

Supplementary figure 174

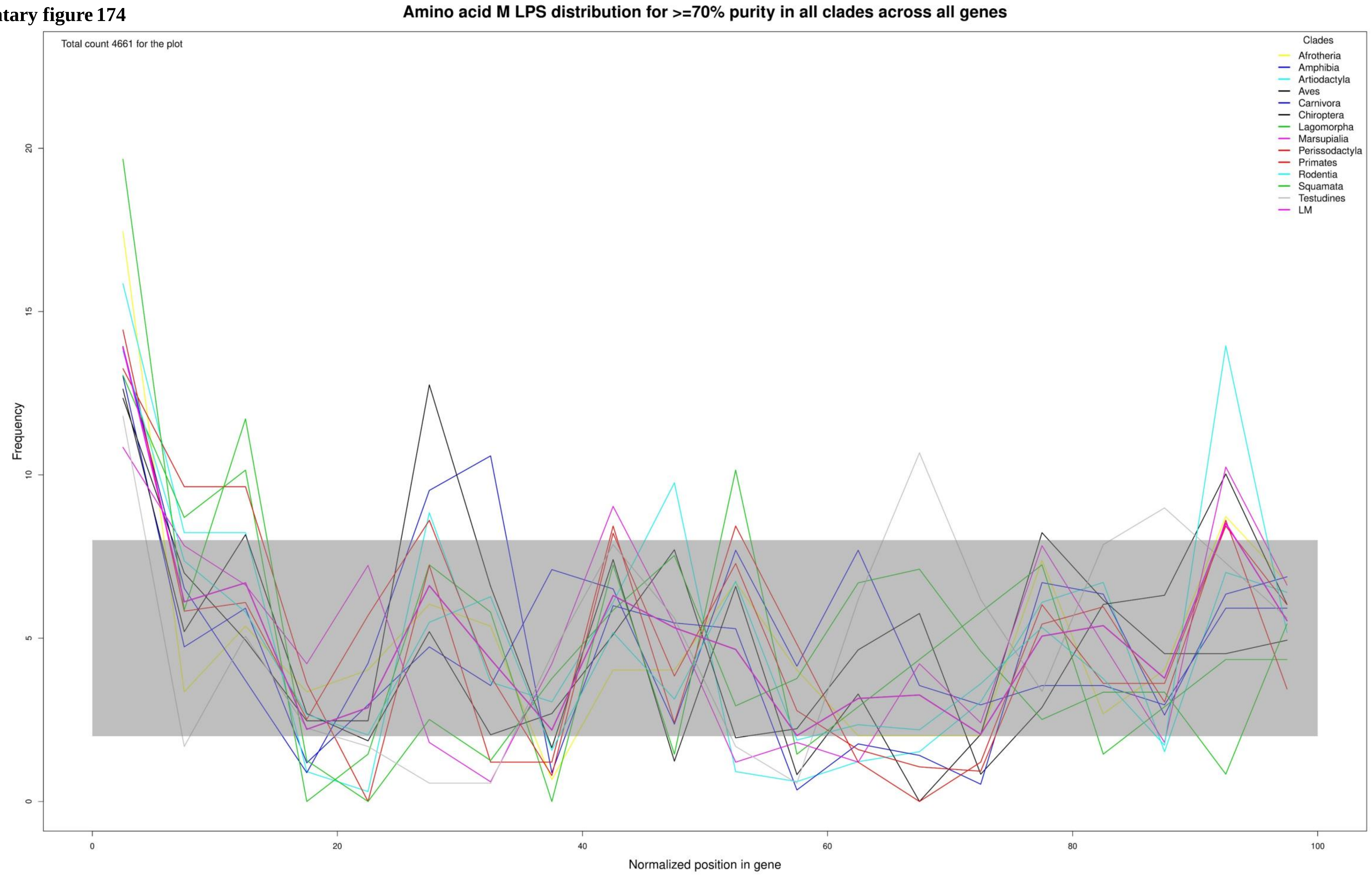

**Supplementary figure 175**

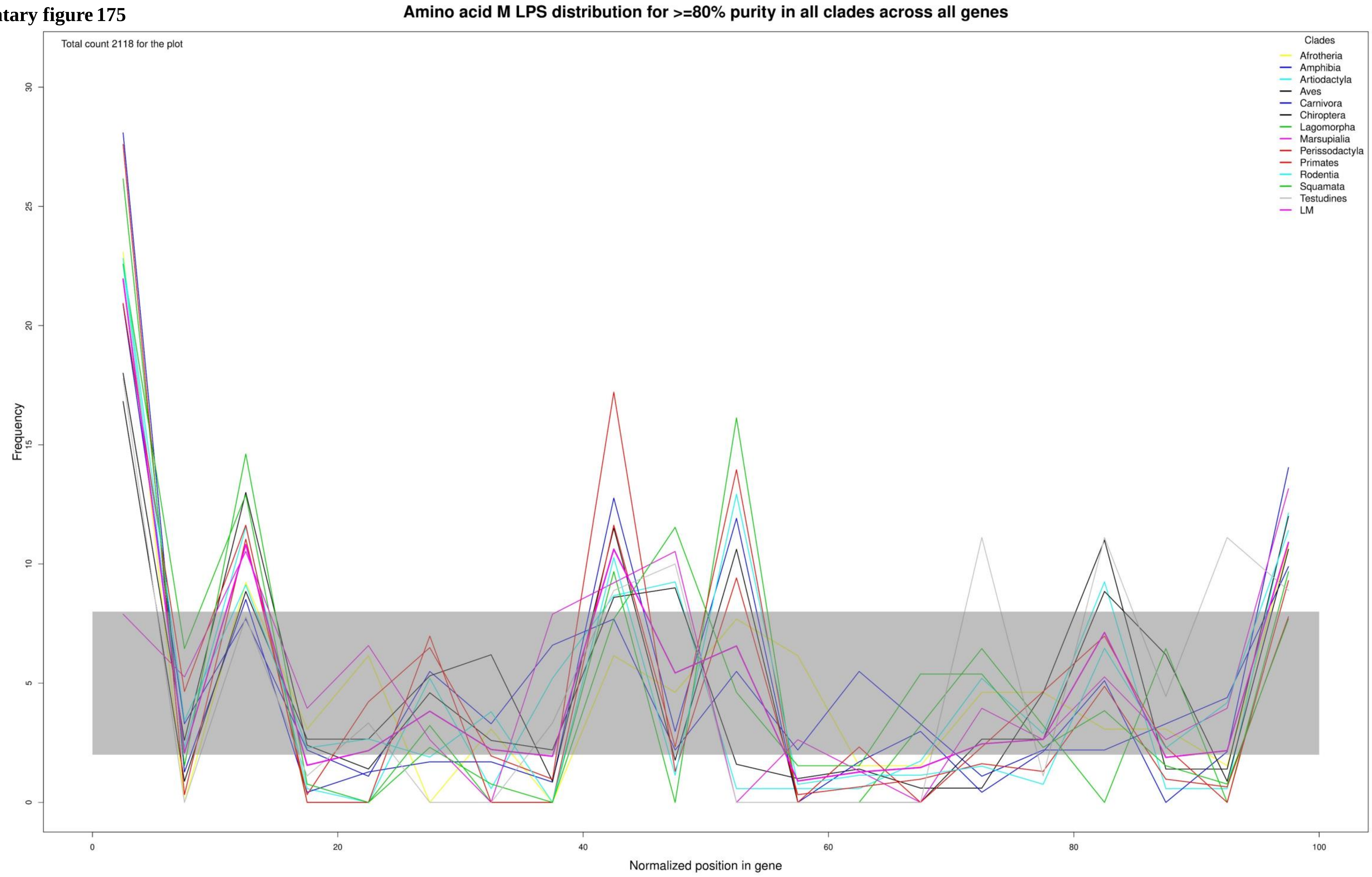

**Supplementary figure 176**

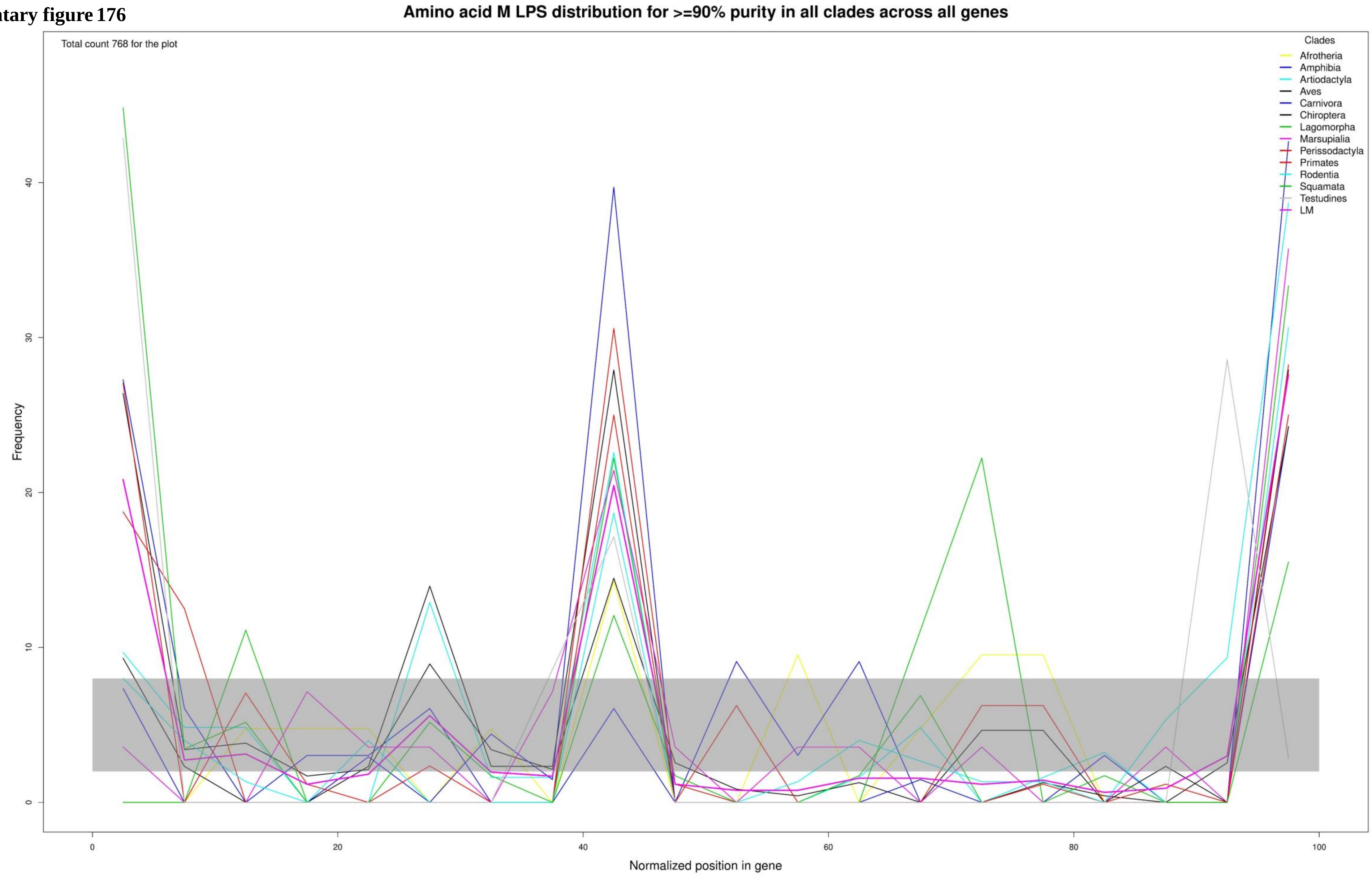

**Supplementary figure 177**

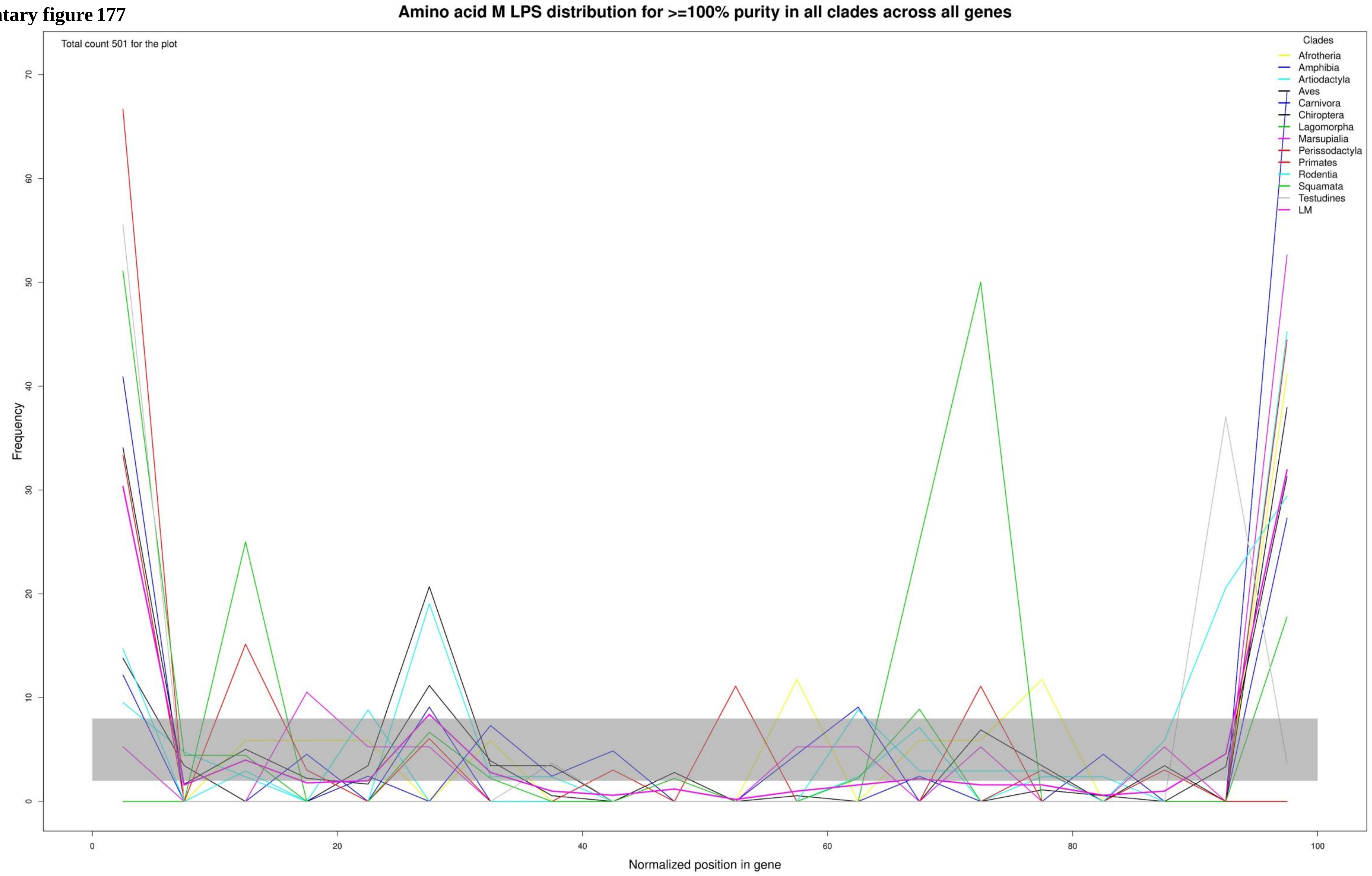

Supplementary figure 178

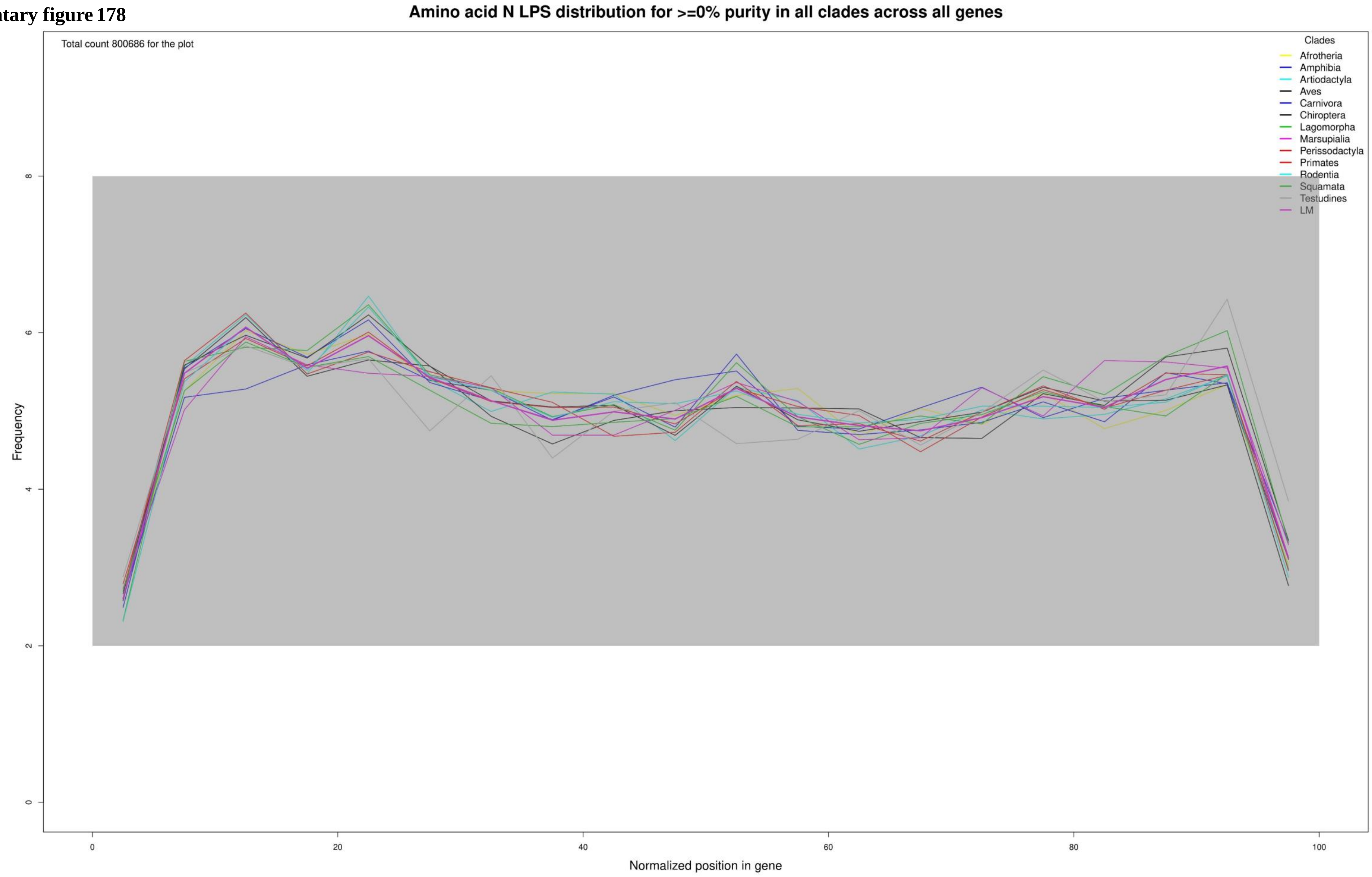

**Supplementary figure 179**

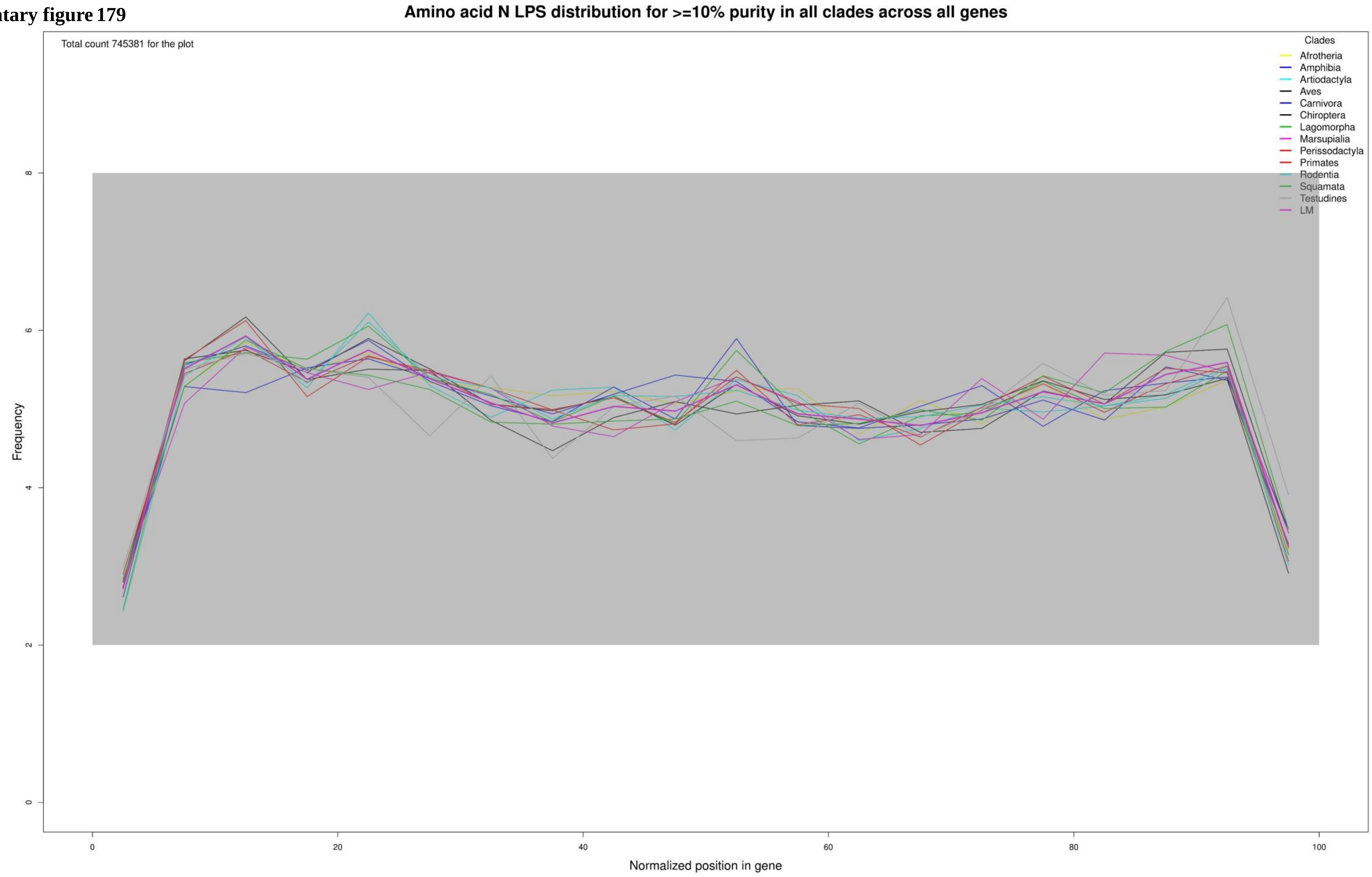

**Supplementary figure 180**

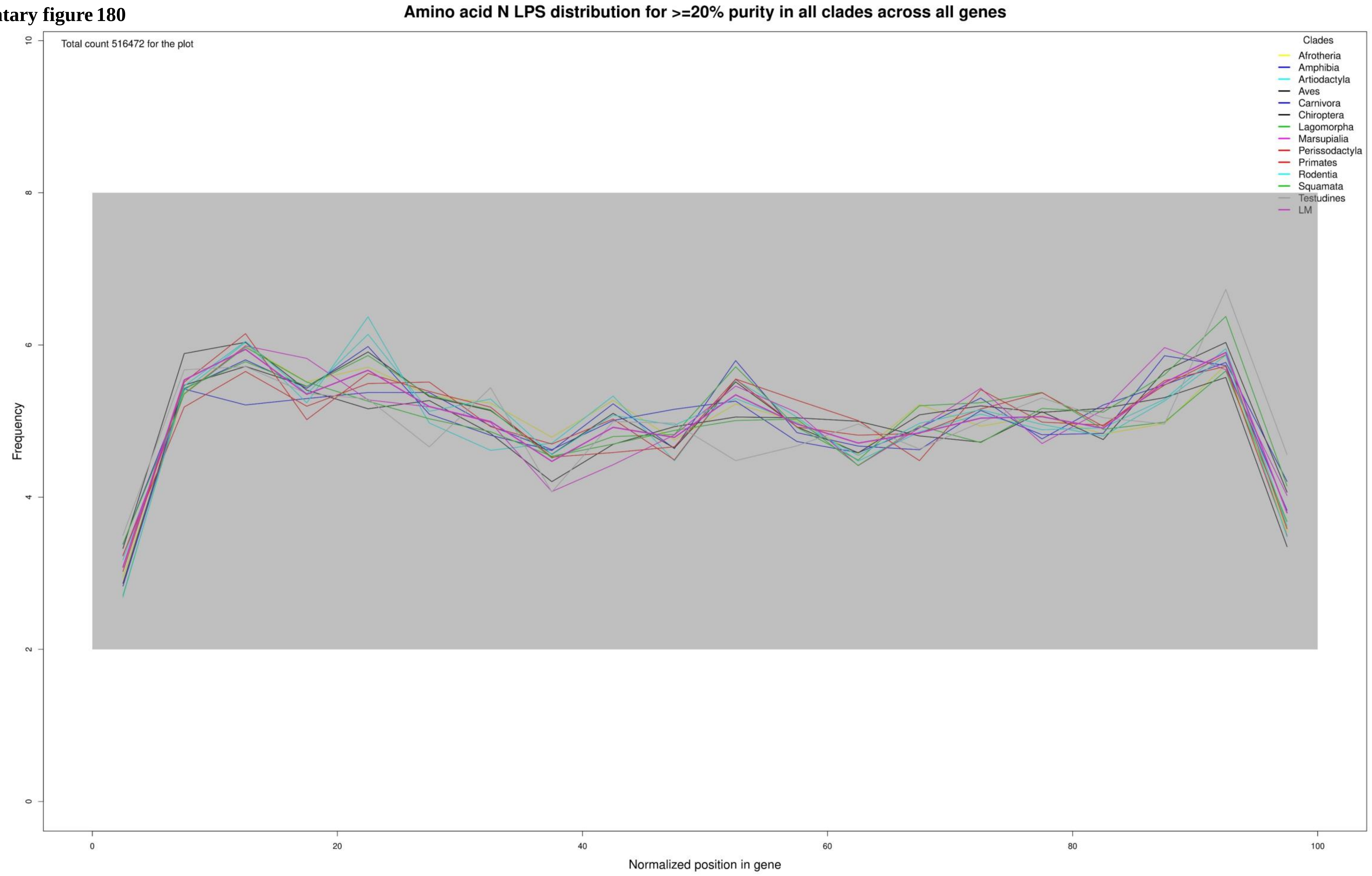

**Supplementary figure 181**

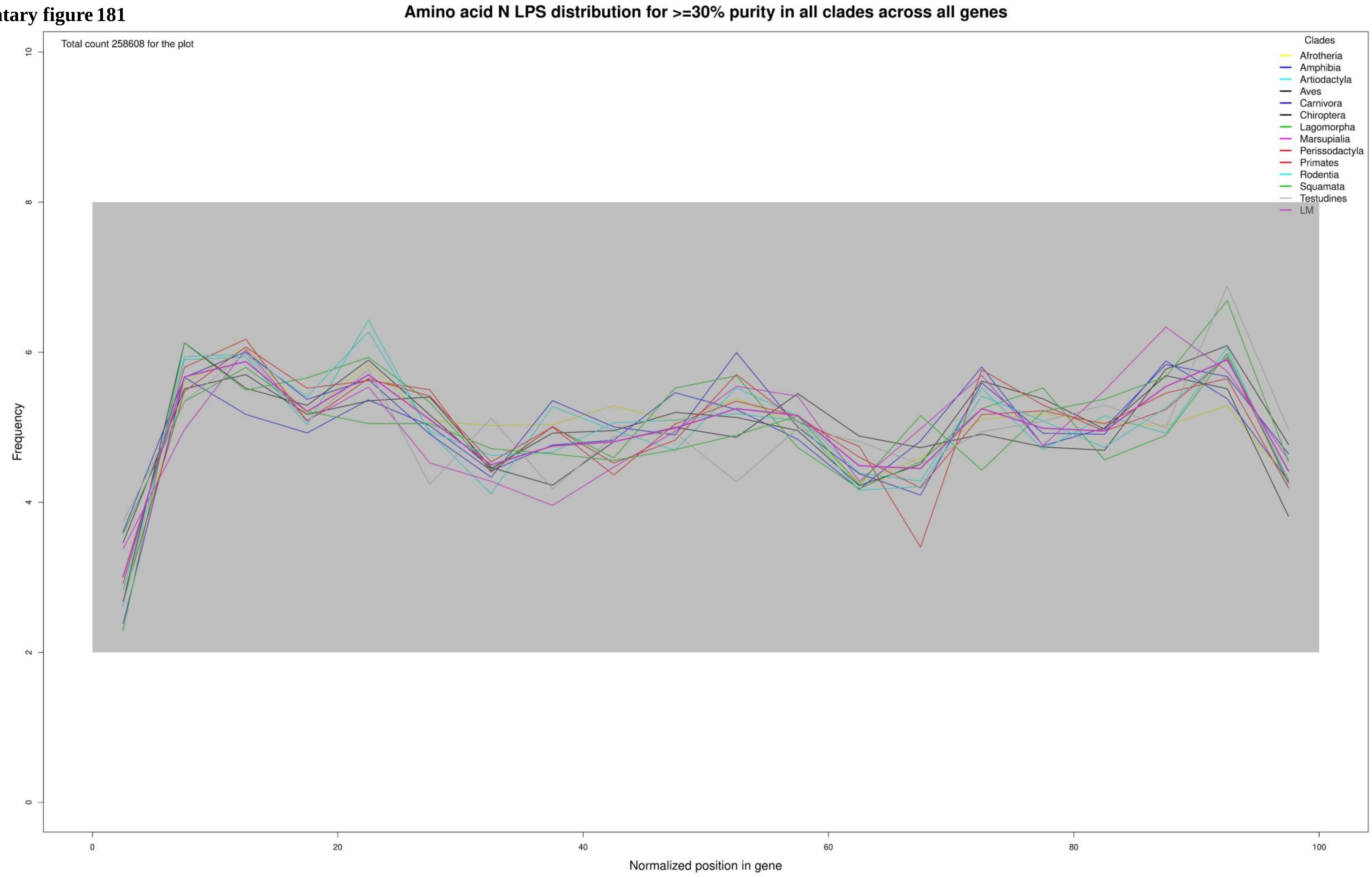

**Supplementary figure 182**

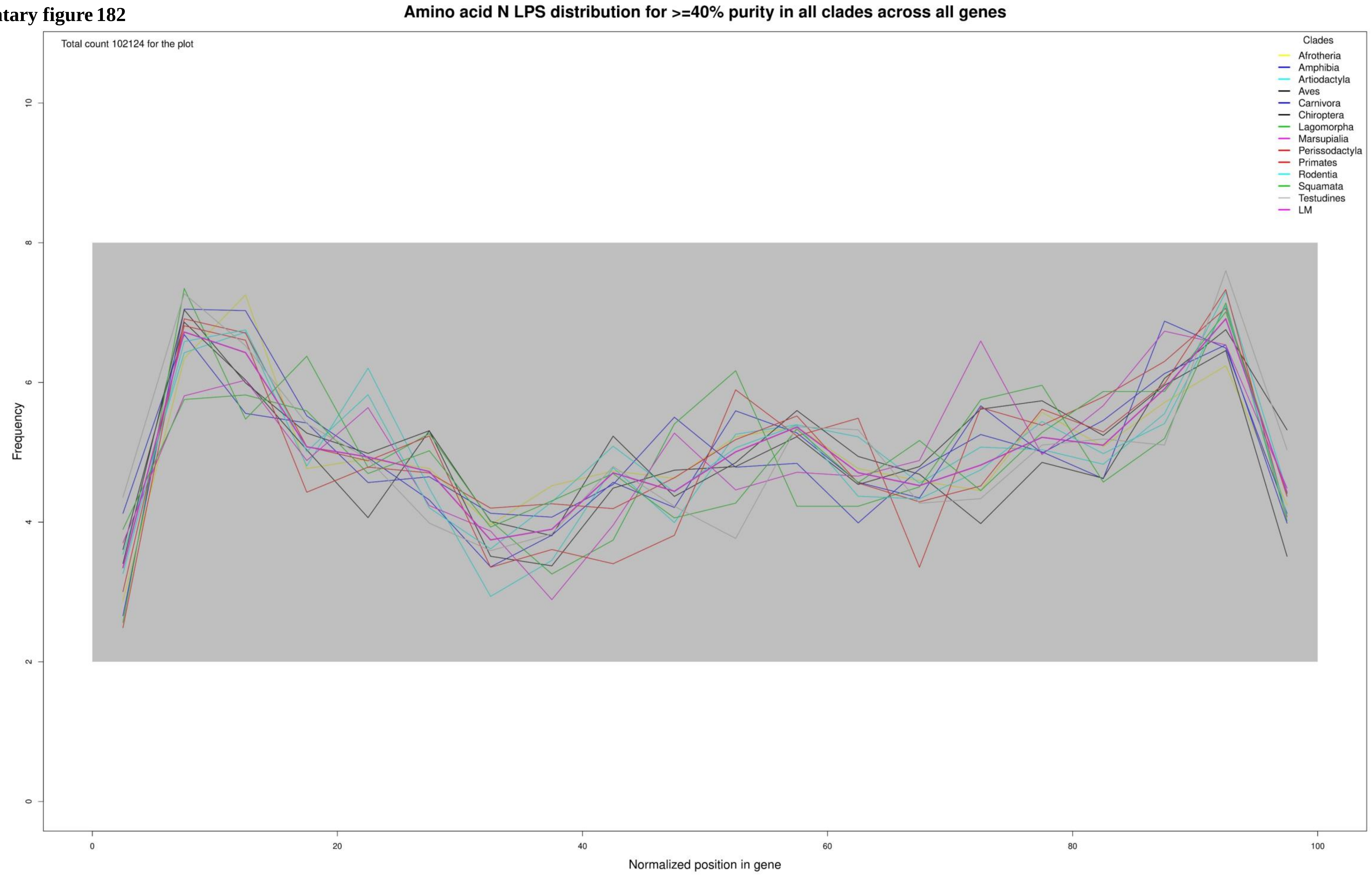

**Supplementary figure 183**

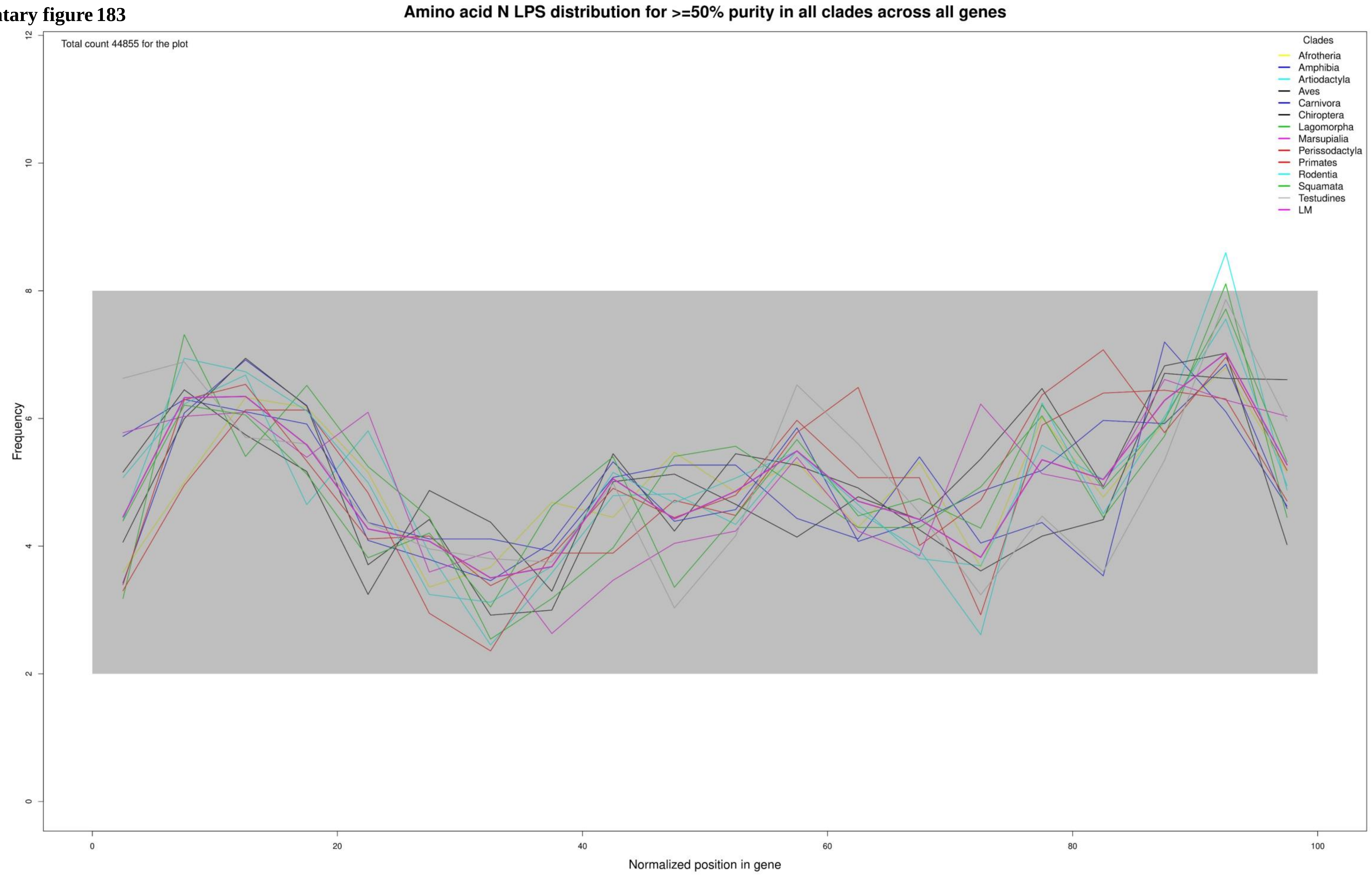

**Supplementary figure 184**

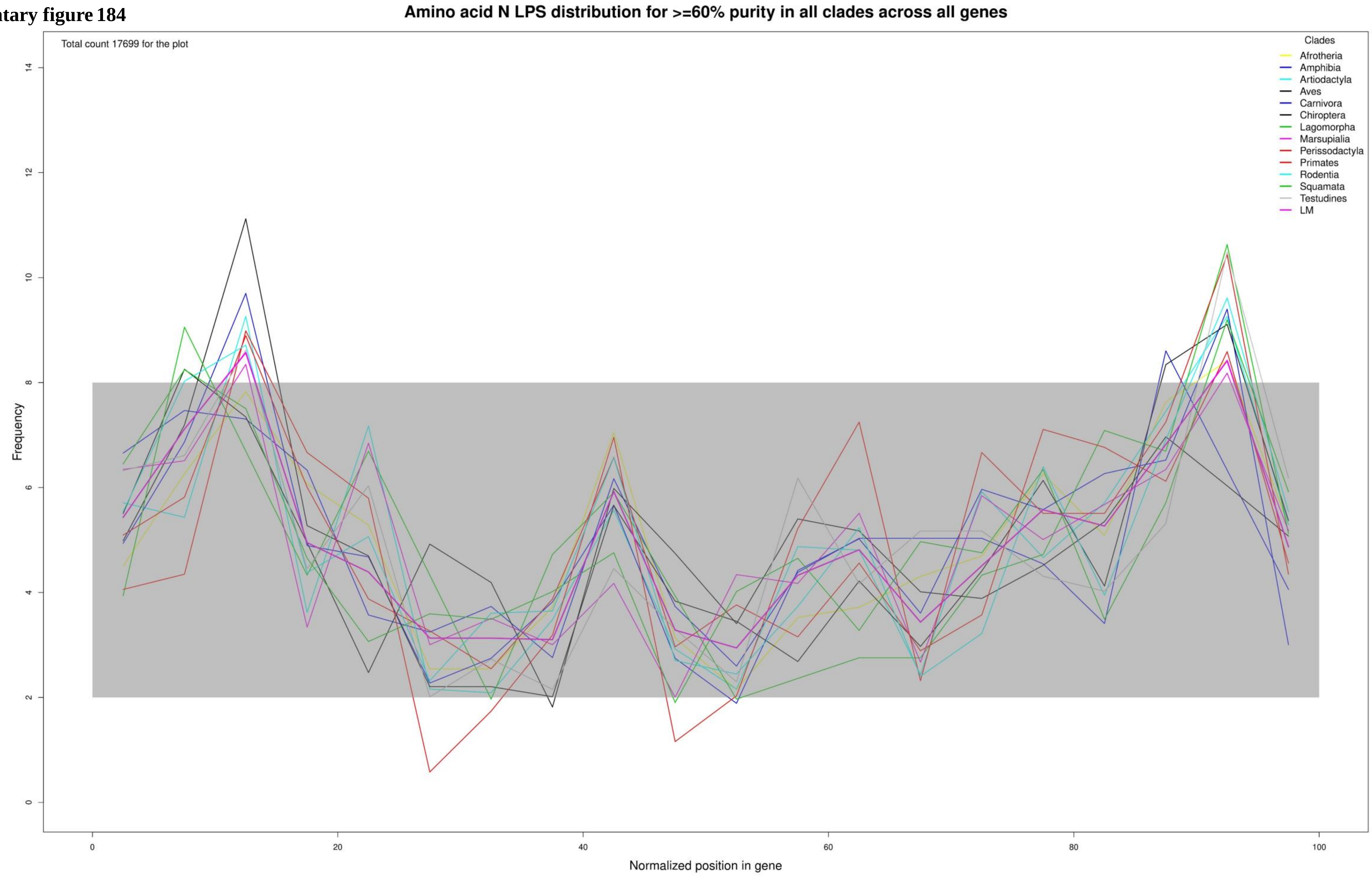

**Supplementary figure 185**

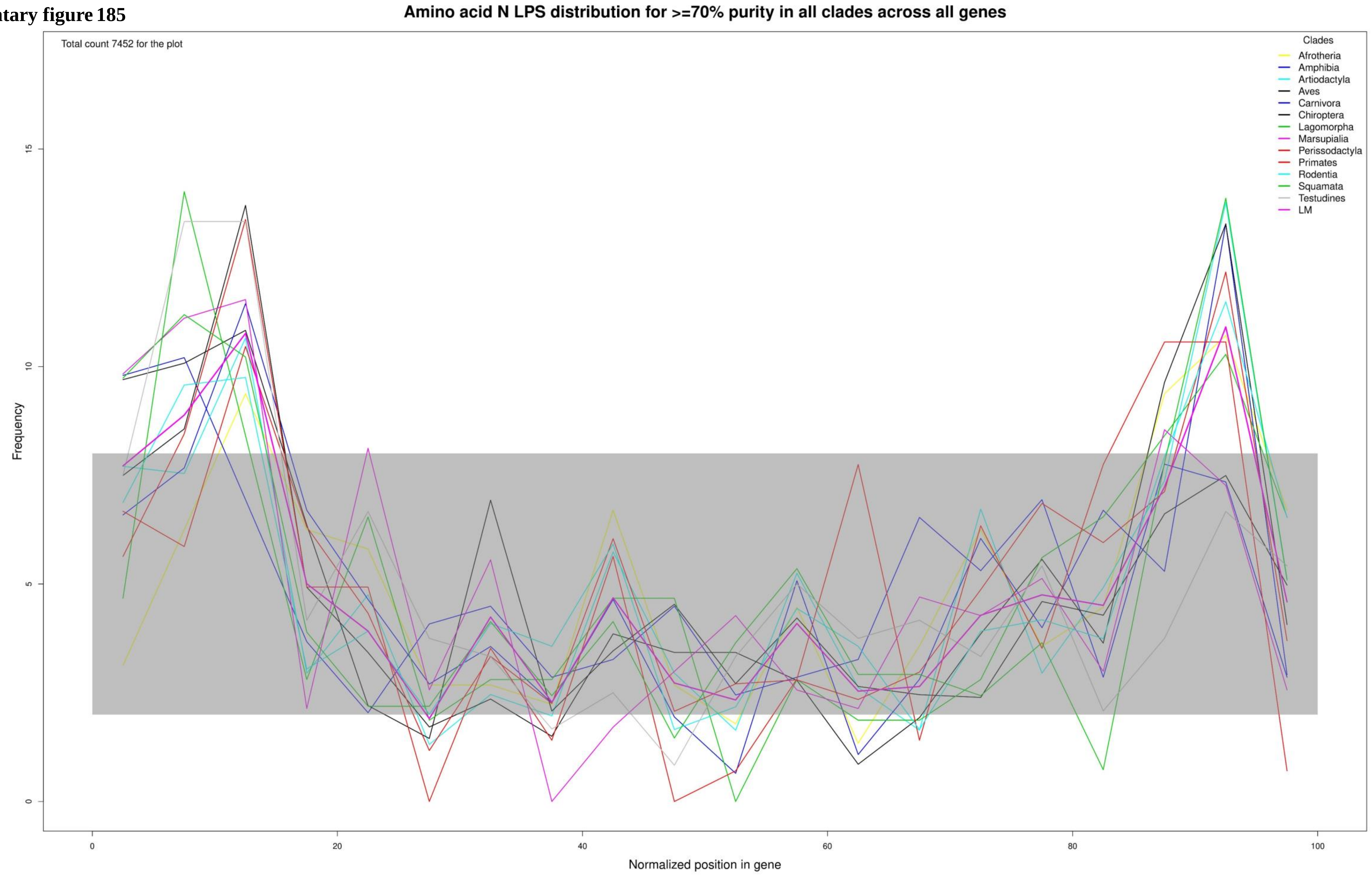

**Supplementary figure 186**

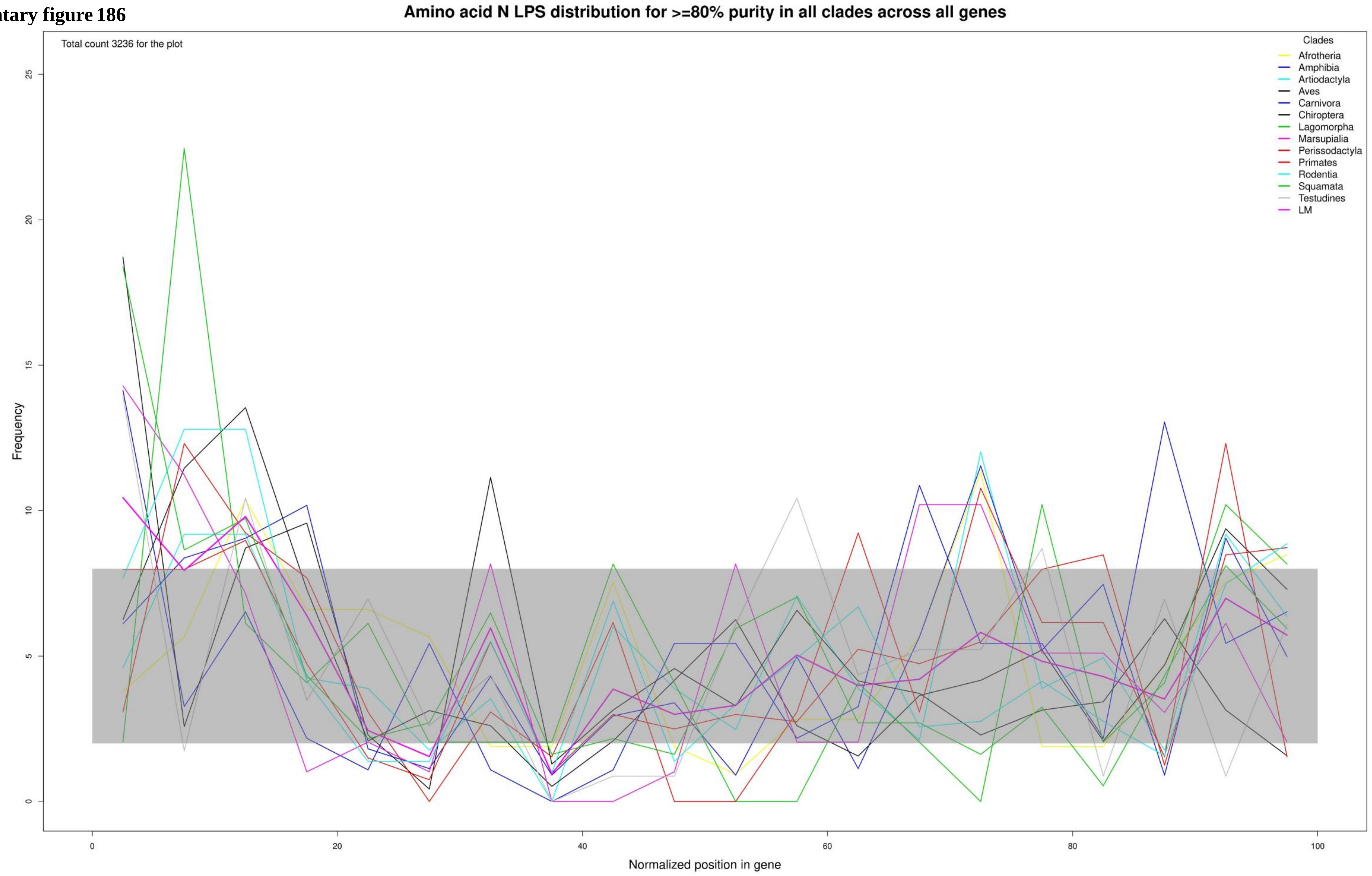

**Supplementary figure 187**

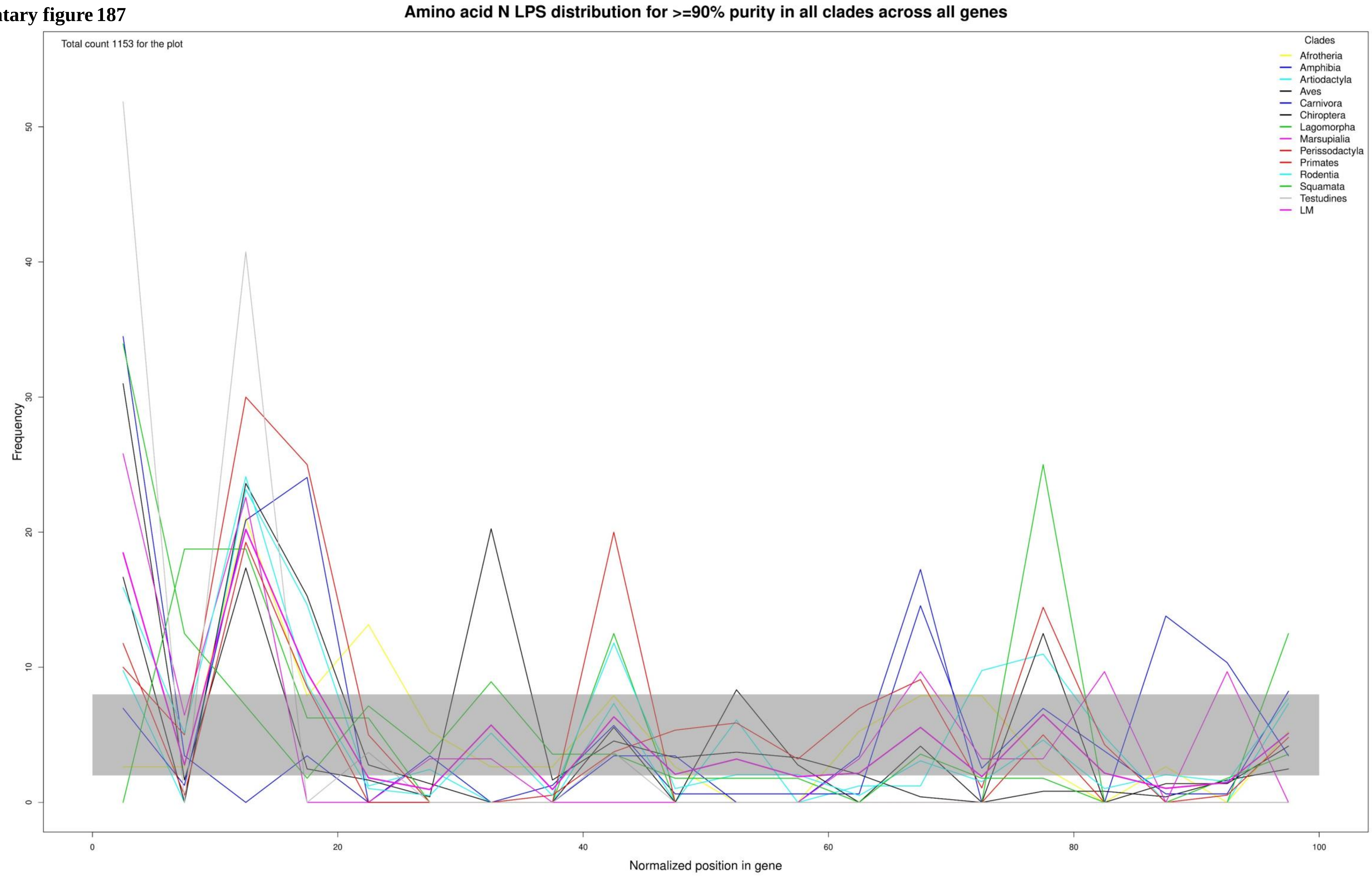

**Supplementary figure 188**

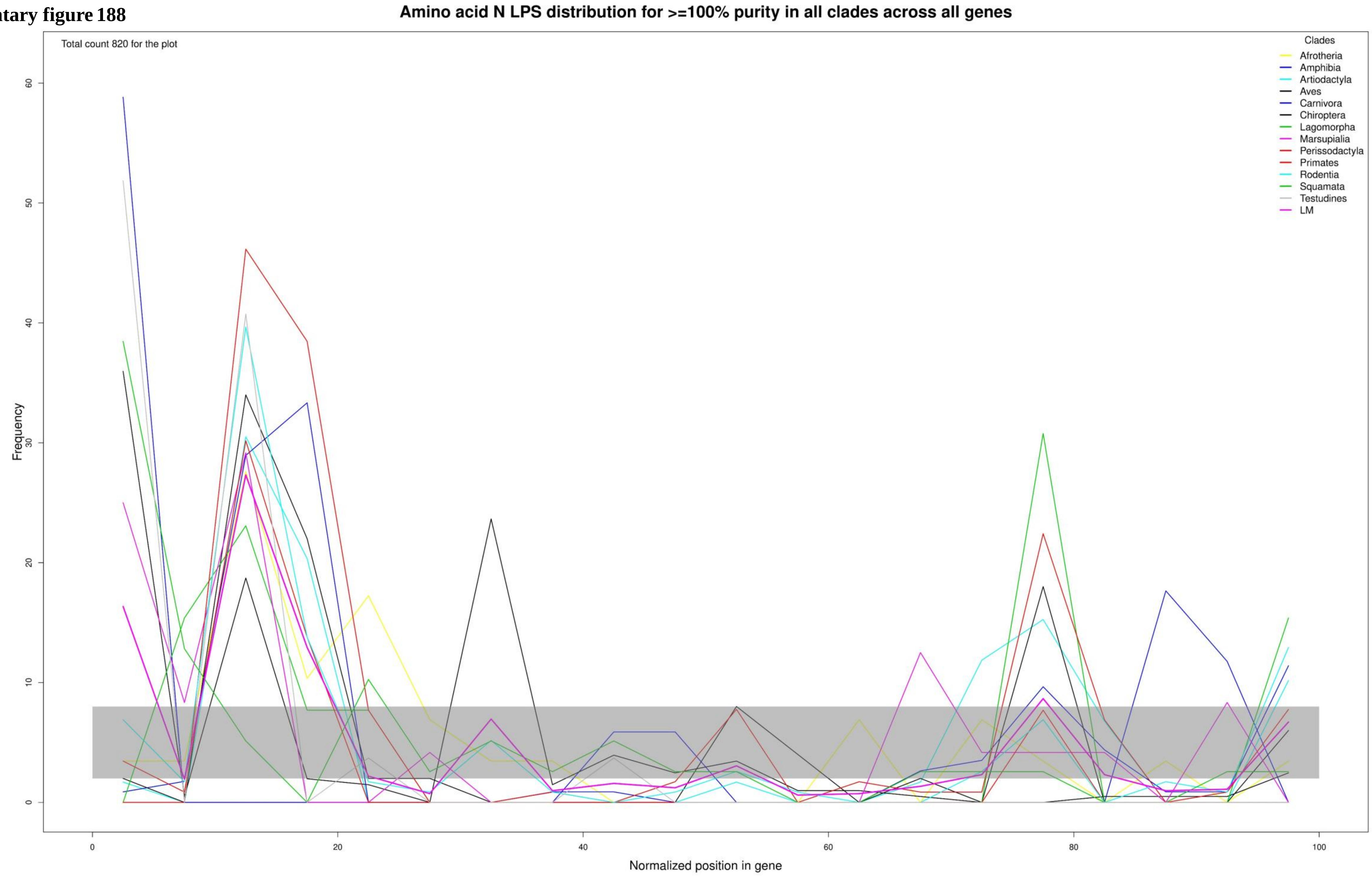

**Supplementary figure 189**

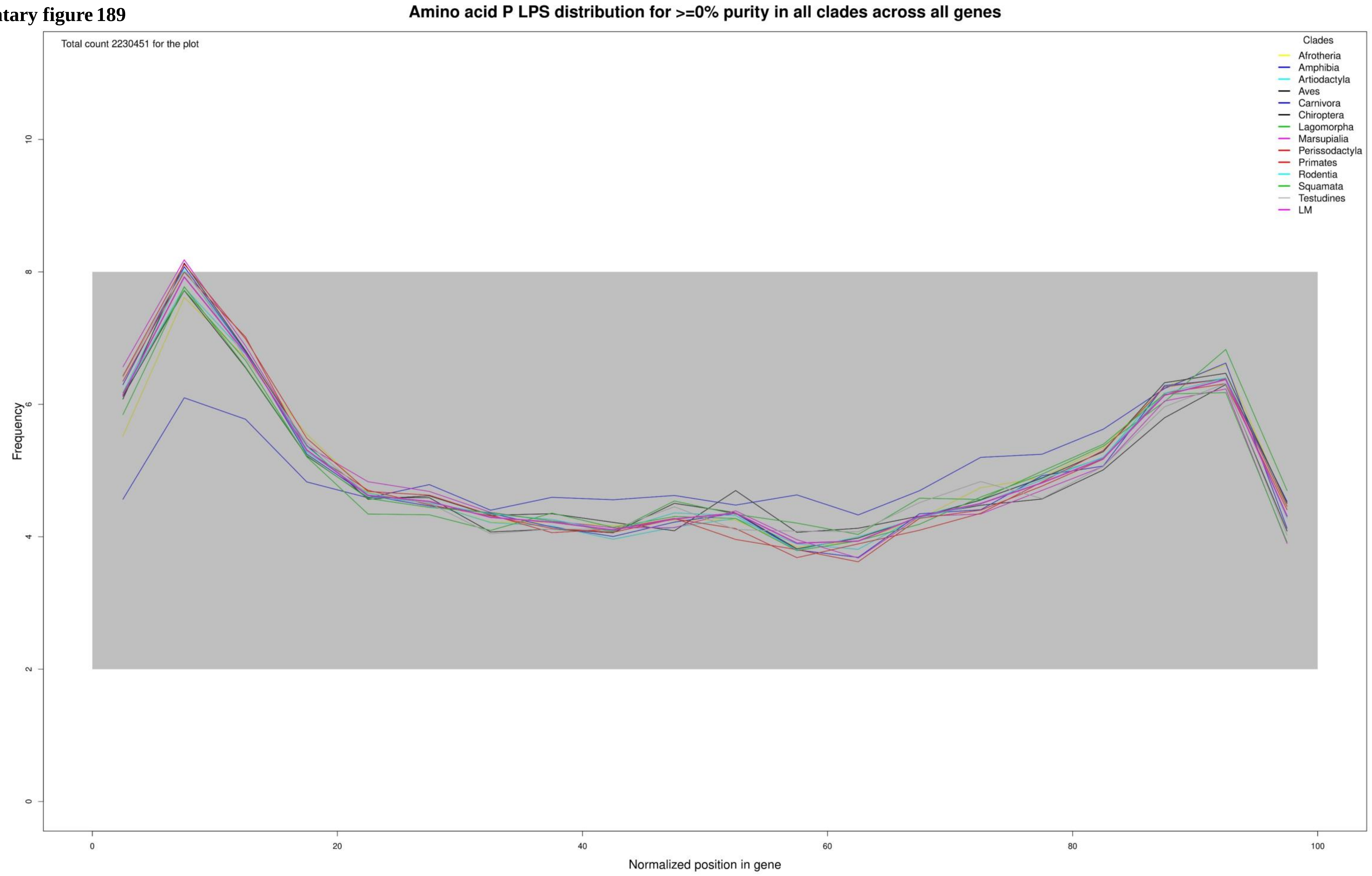

**Supplementary figure 190**

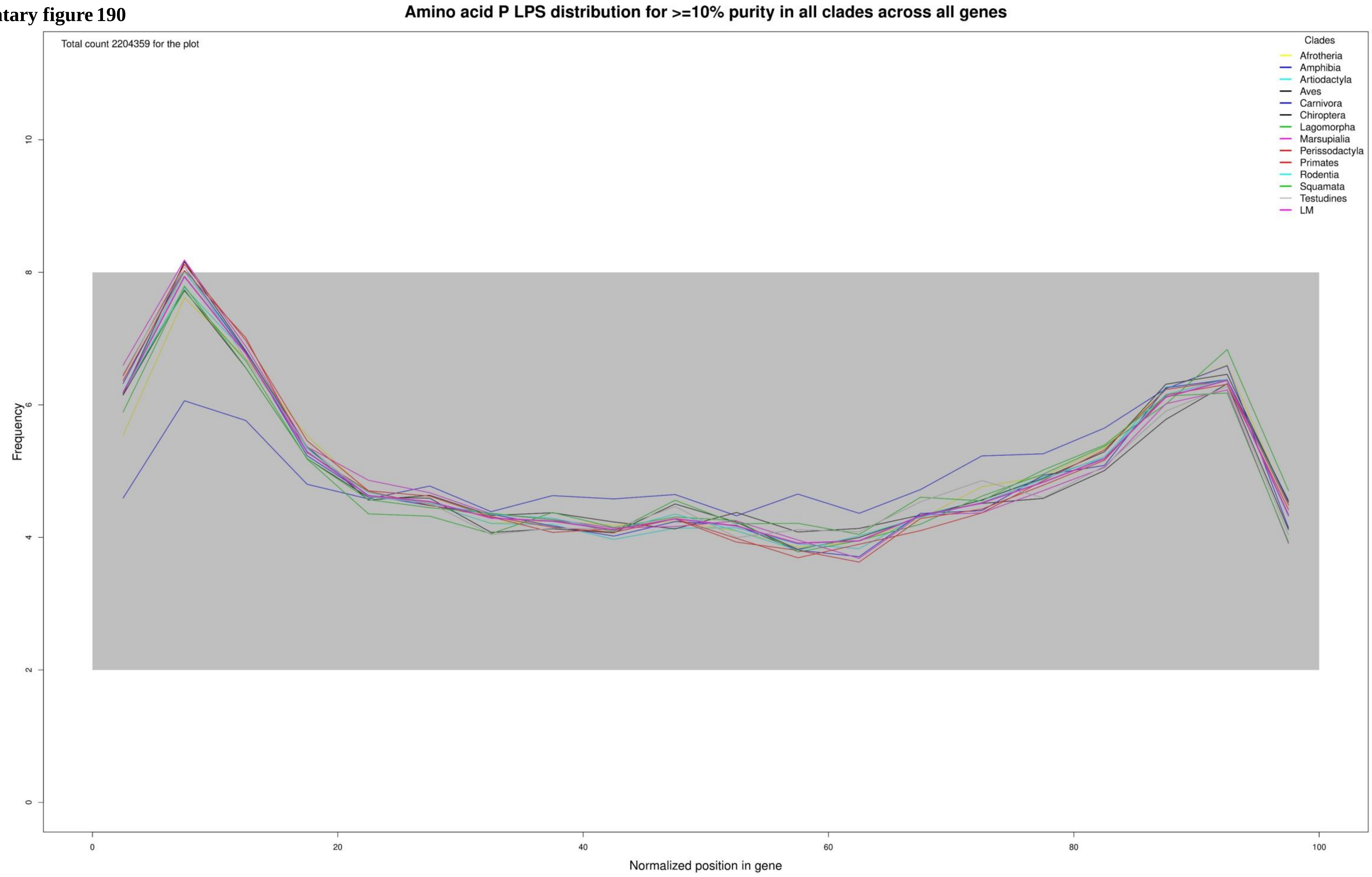

**Supplementary figure 191**

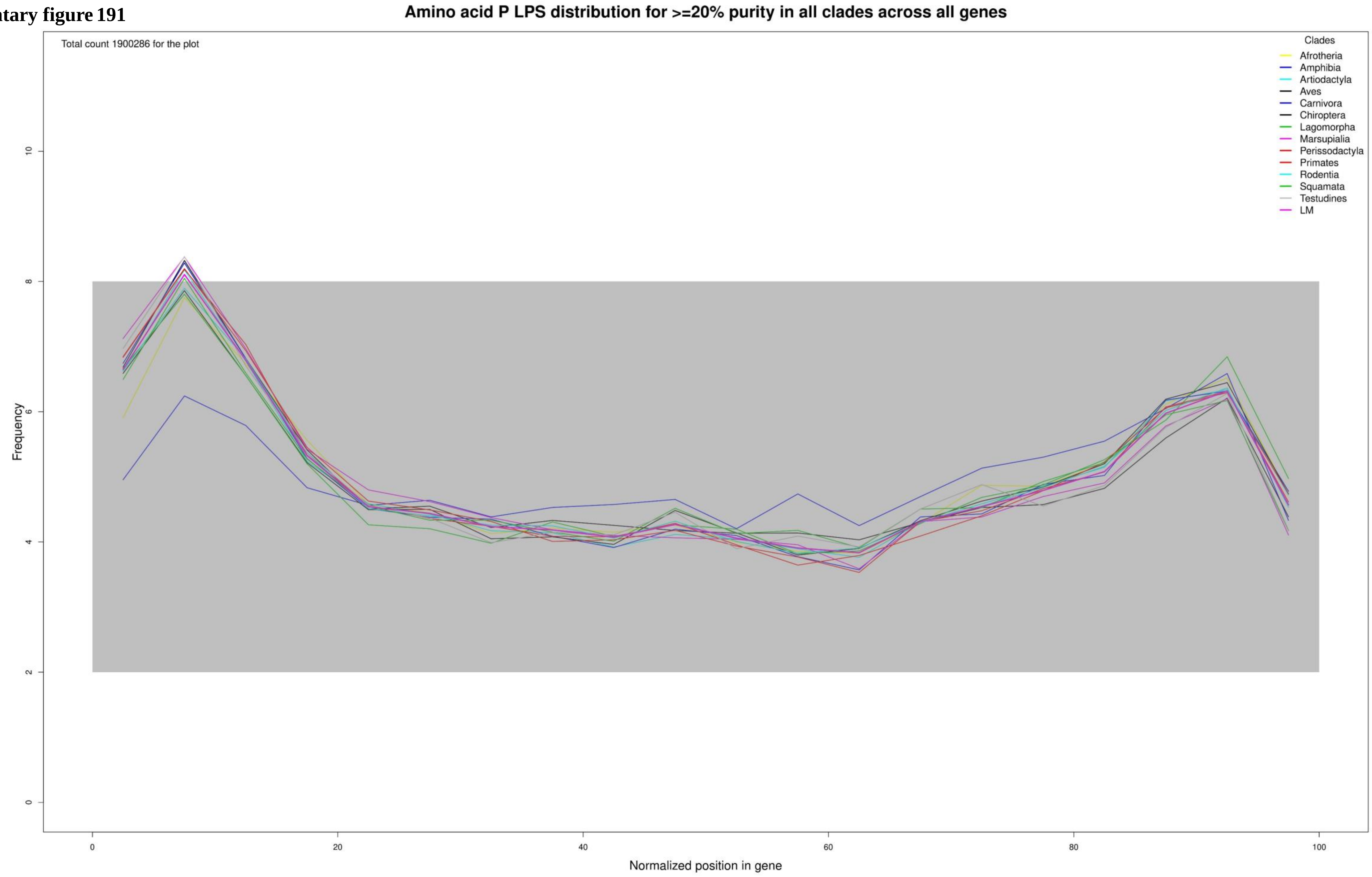

**Supplementary figure 192**

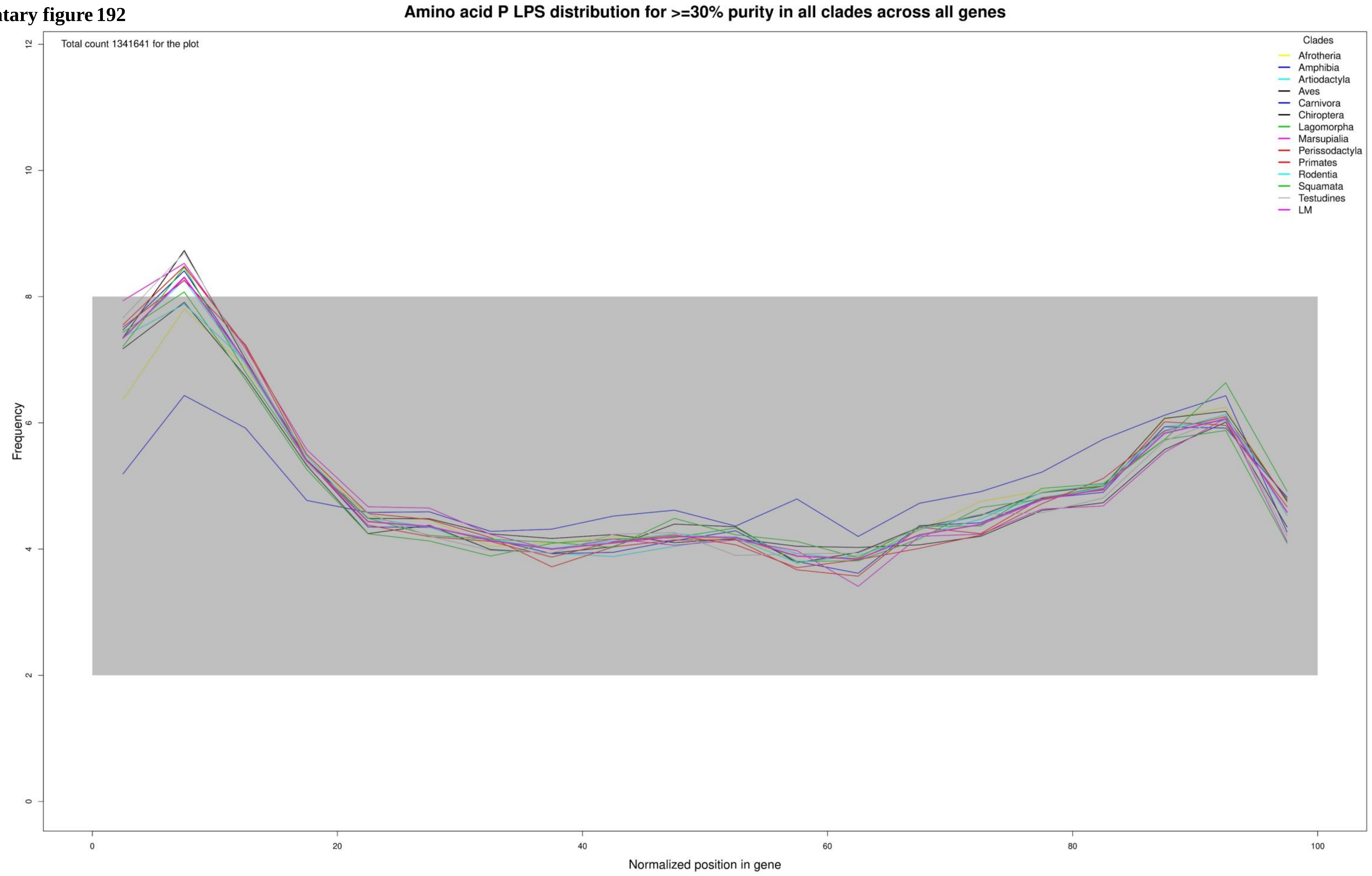

**Supplementary figure 193**

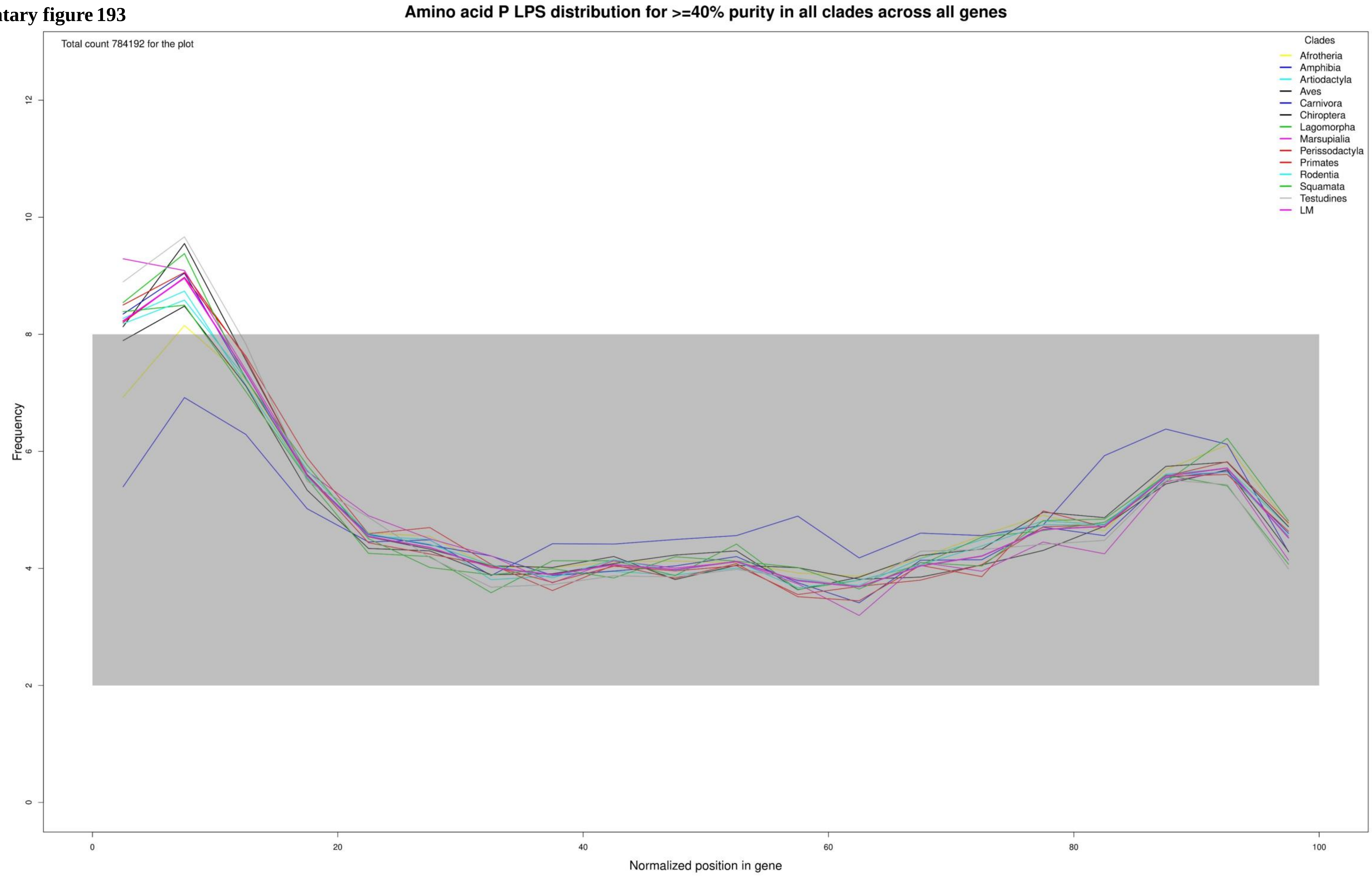

**Supplementary figure 194**

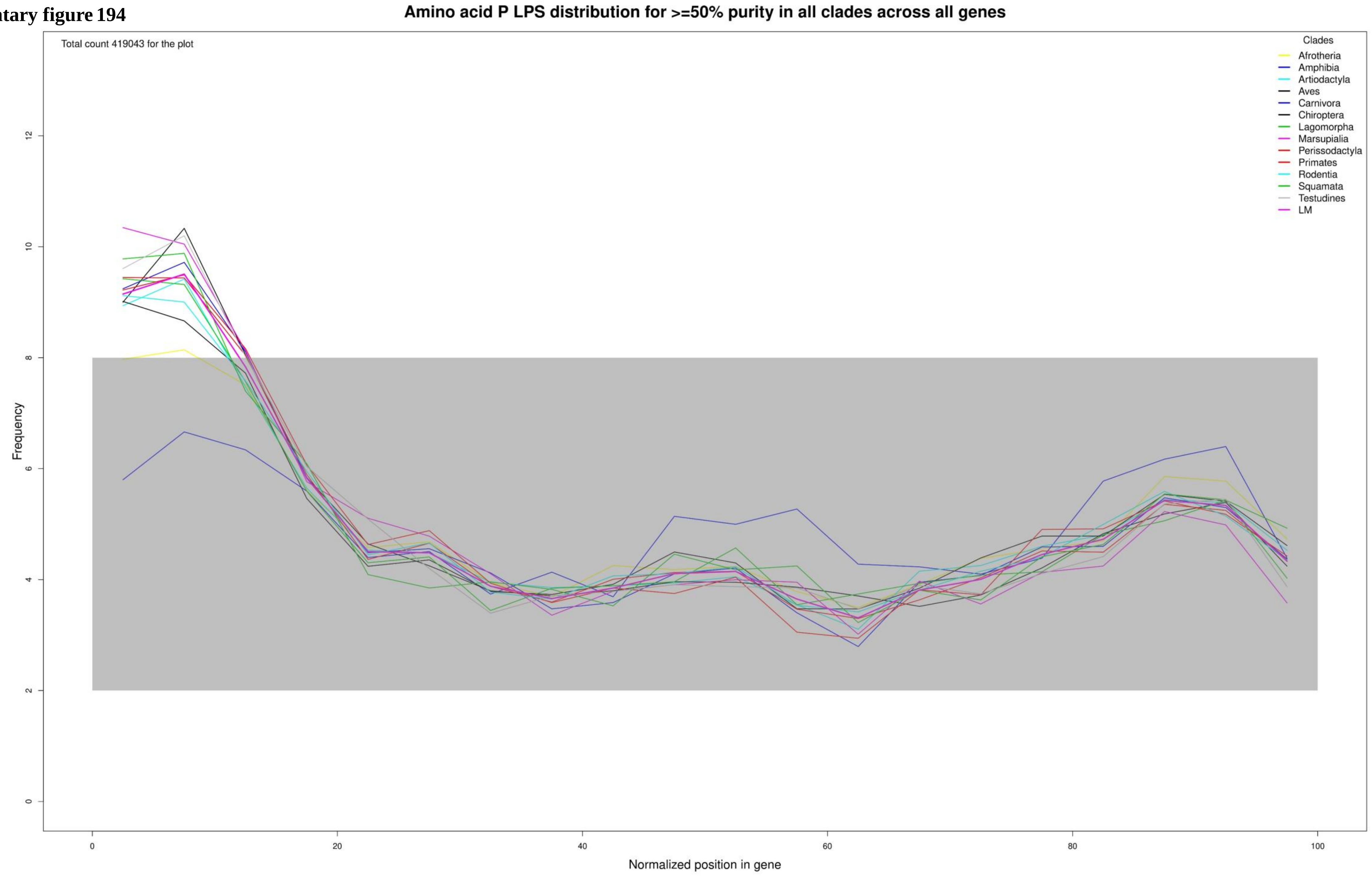

**Supplementary figure 195**

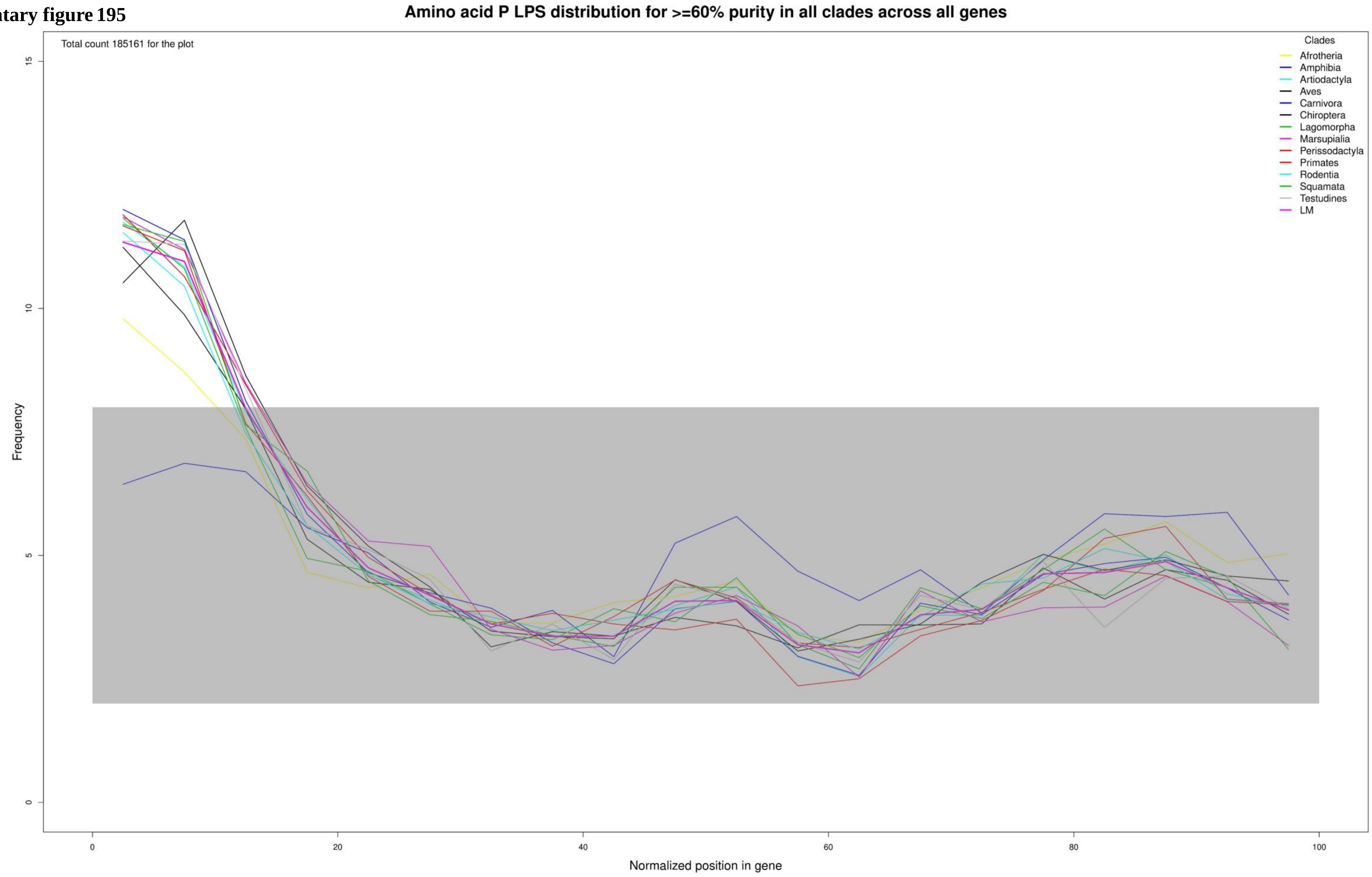

**Supplementary figure 196**

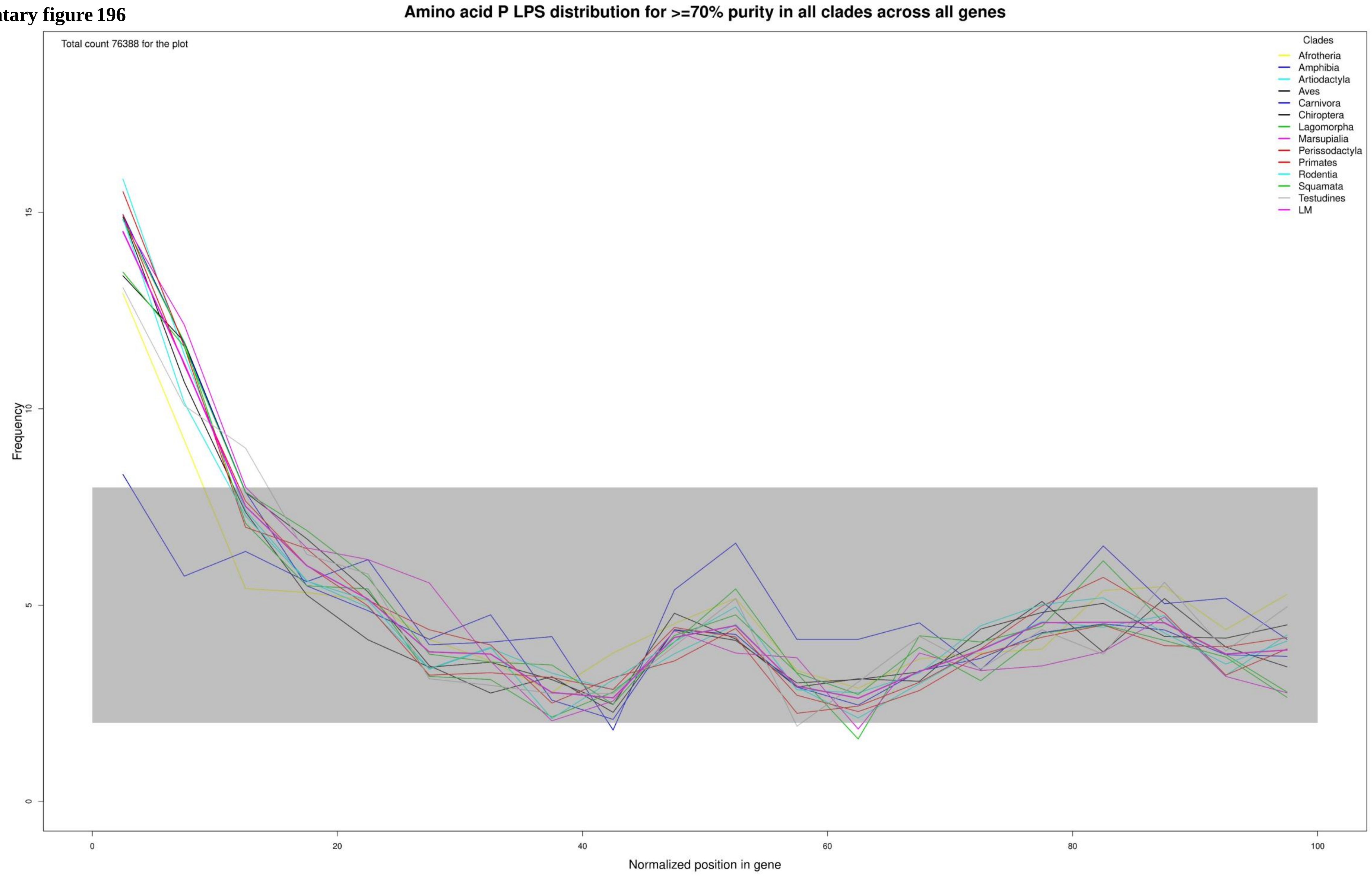

**Supplementary figure 197**

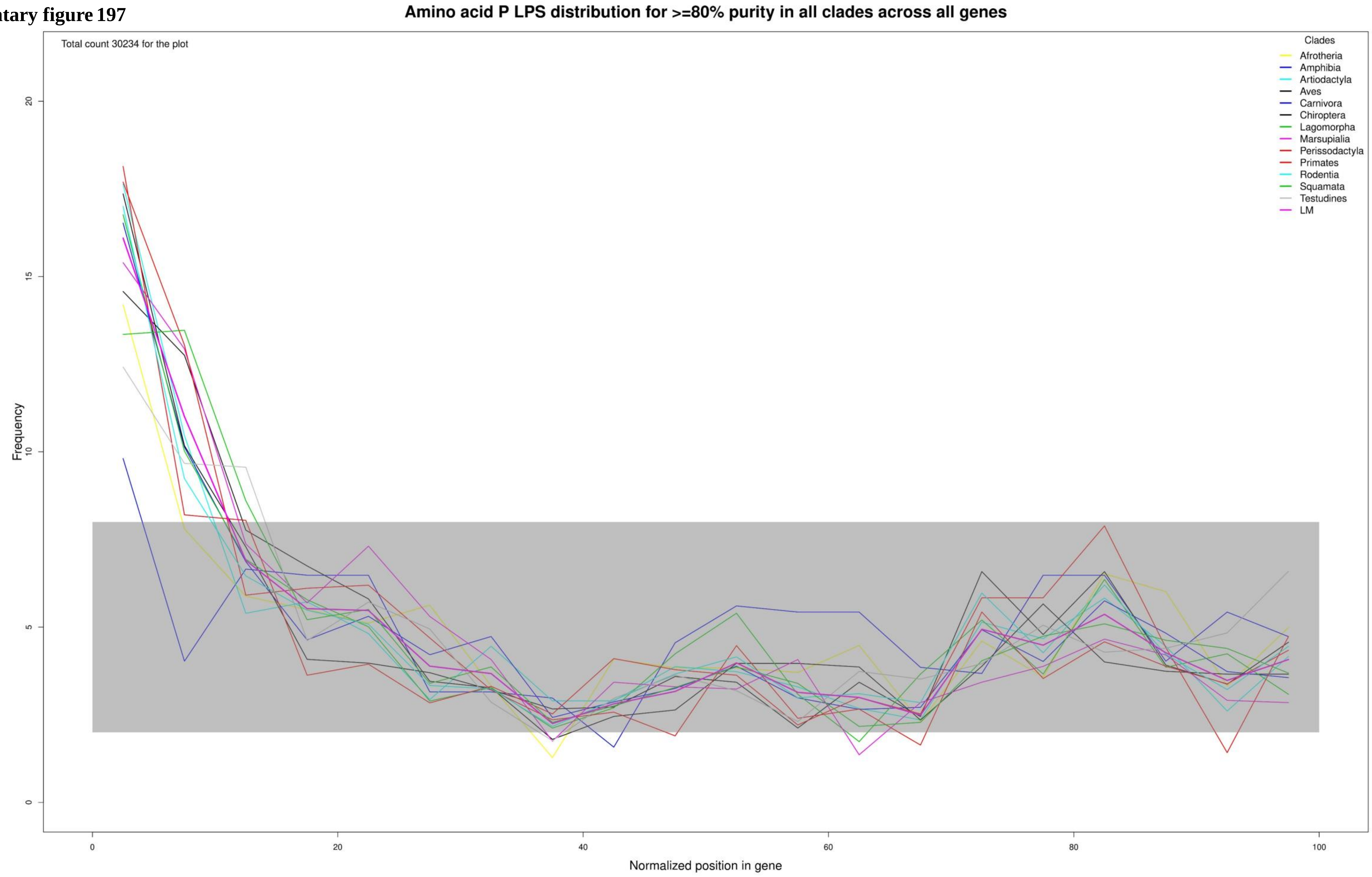

**Supplementary figure 198**

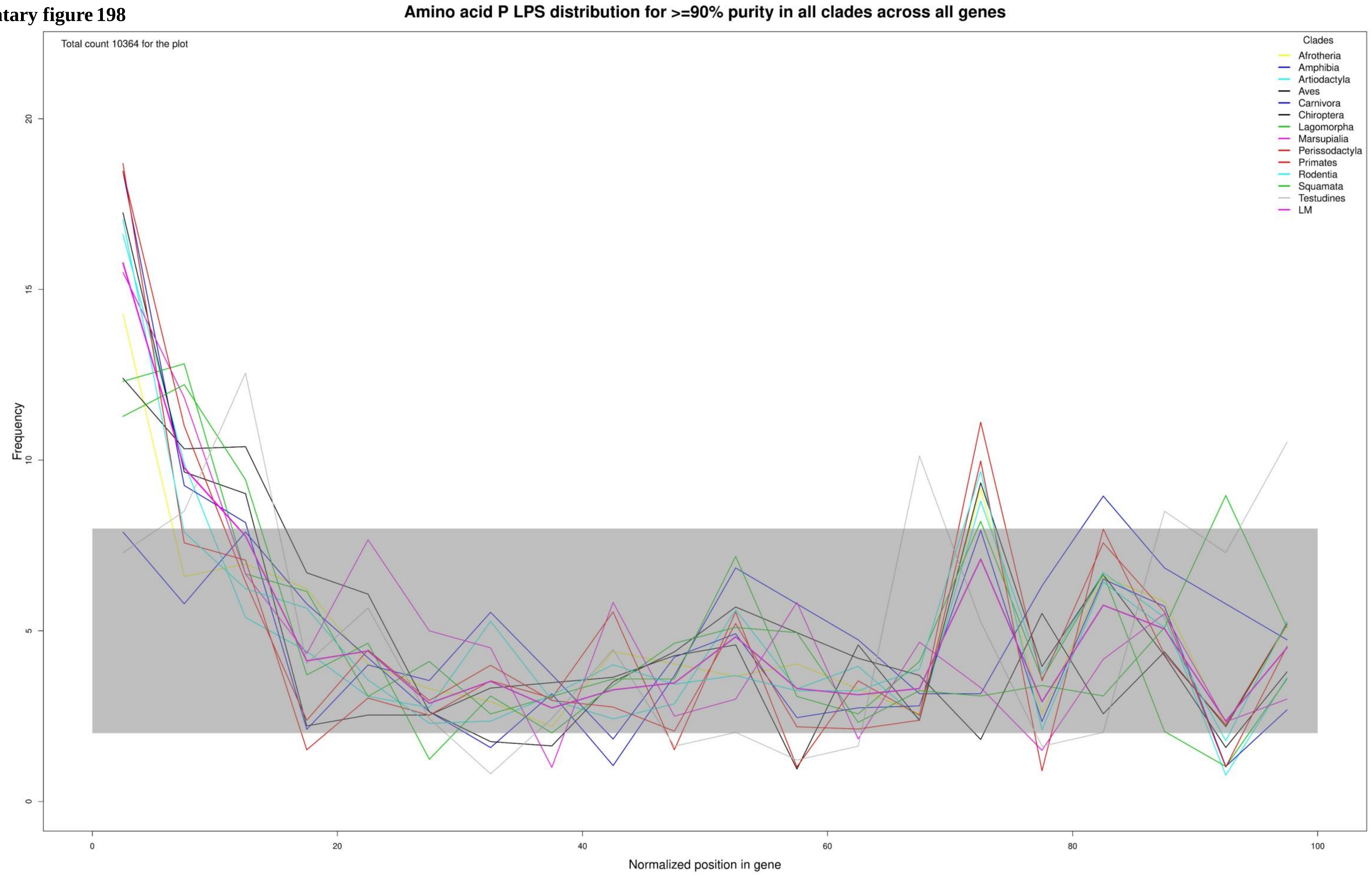

**Supplementary figure 199**

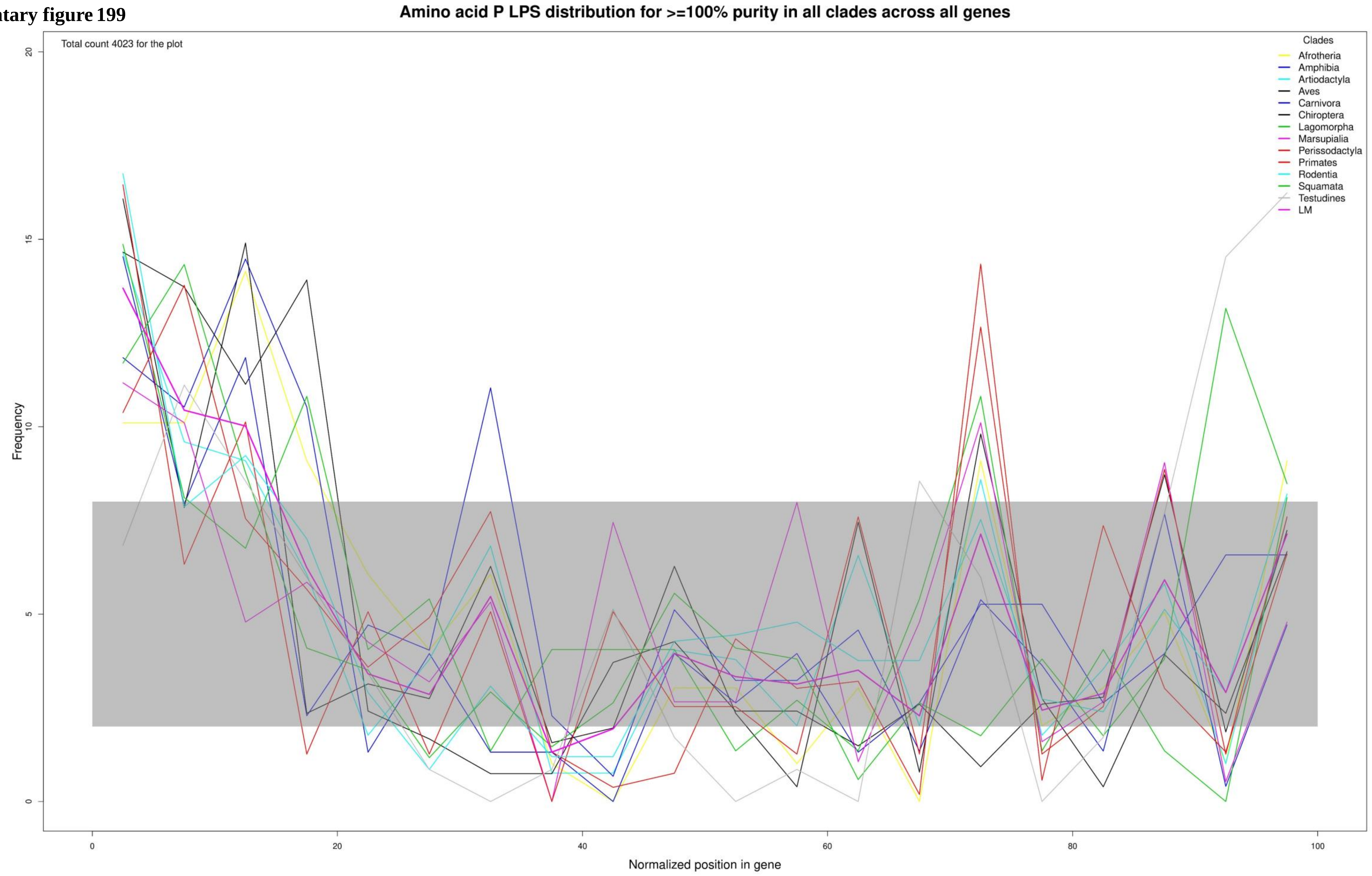

**Supplementary figure 200**

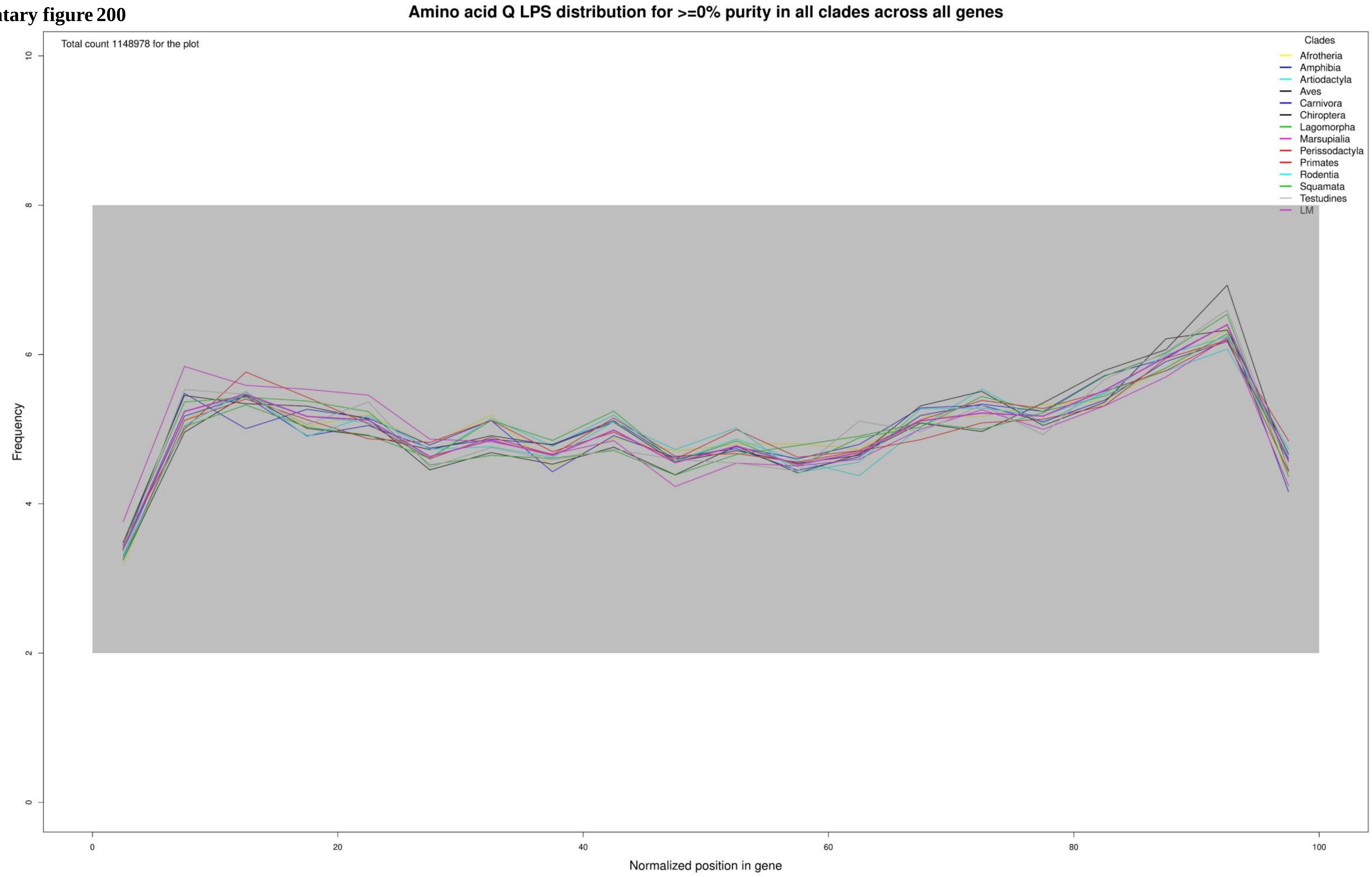

**Supplementary figure 201**

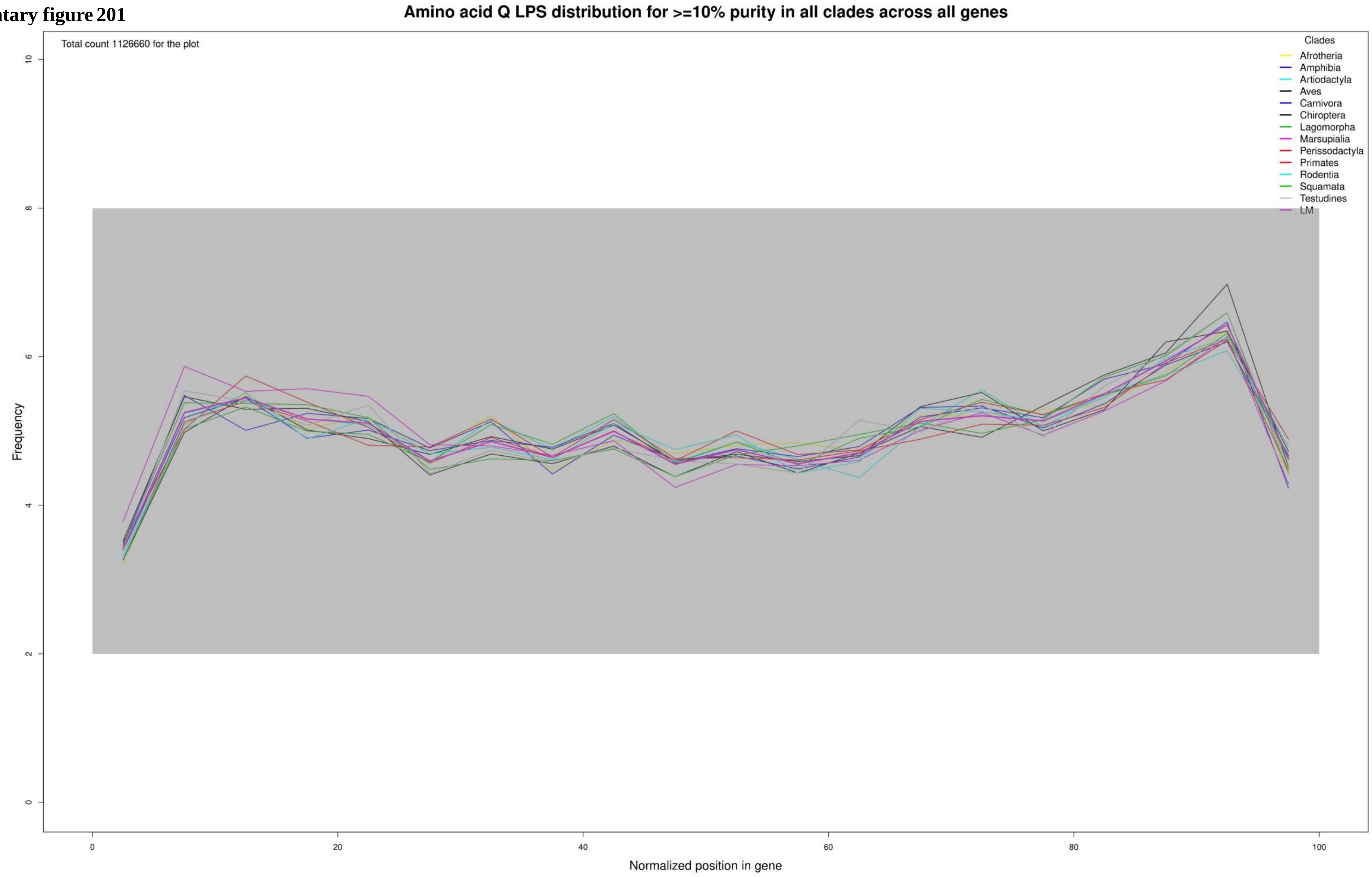

**Supplementary figure 202**

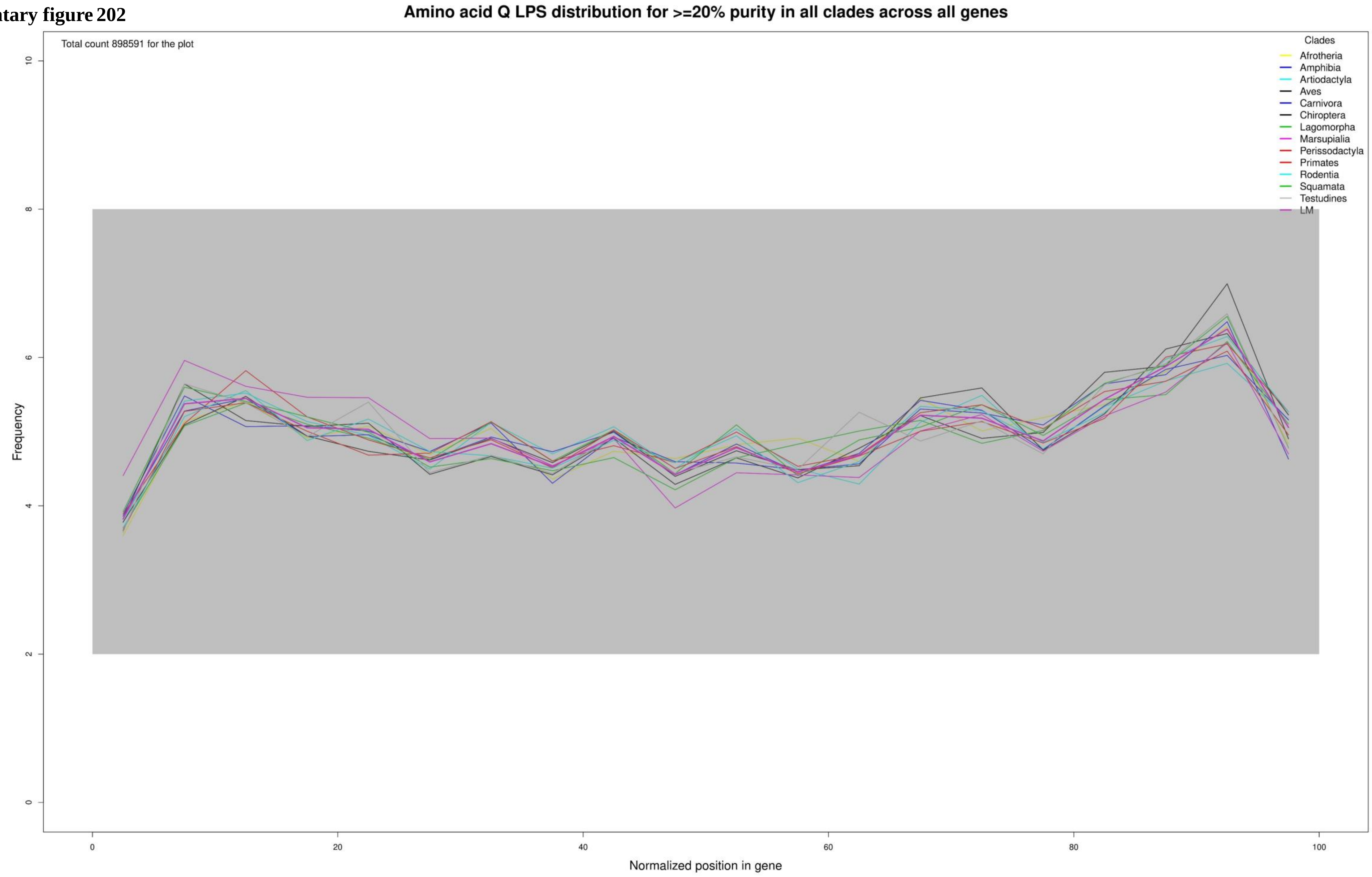

**Supplementary figure 203**

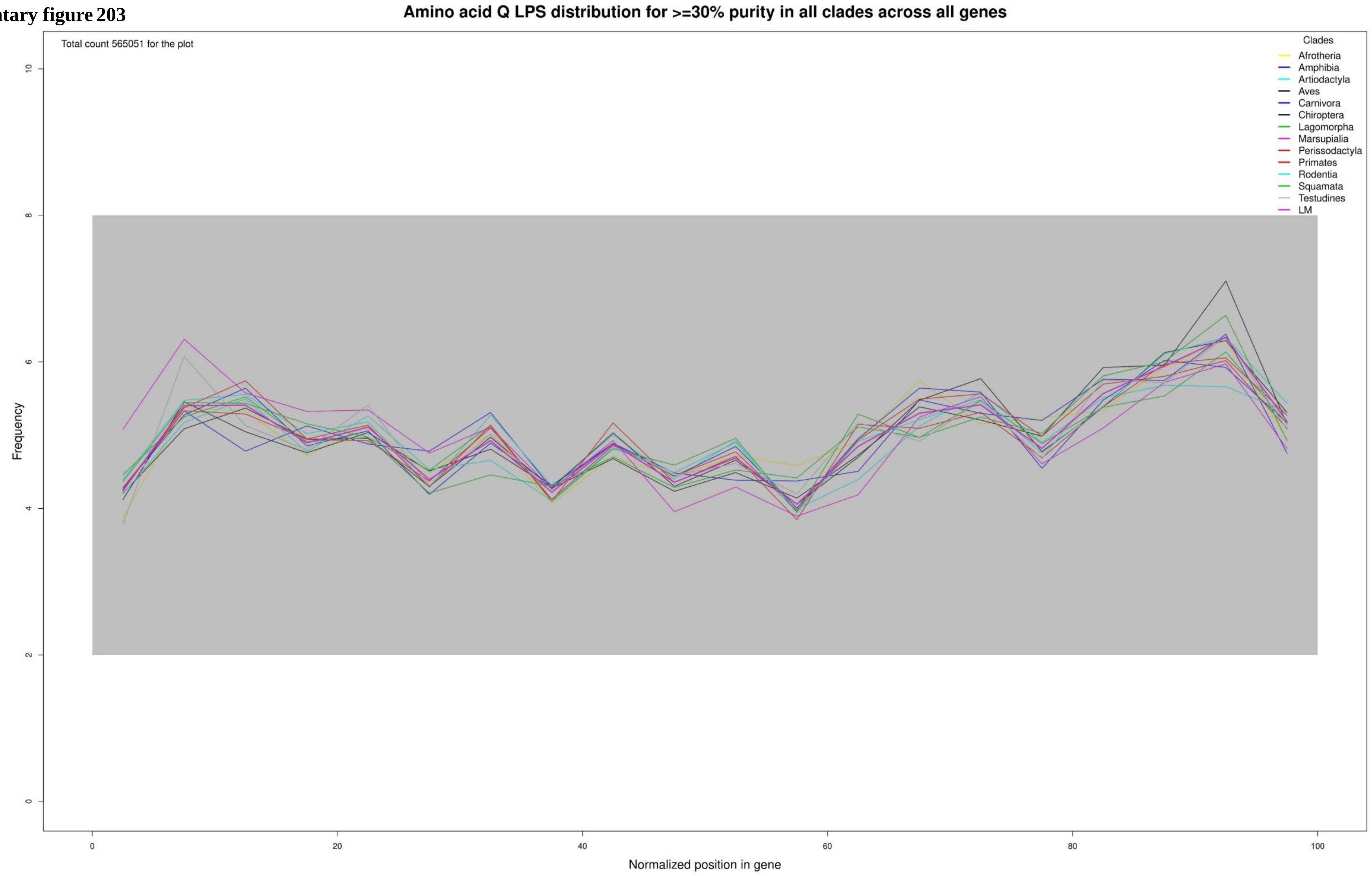

**Supplementary figure 204**

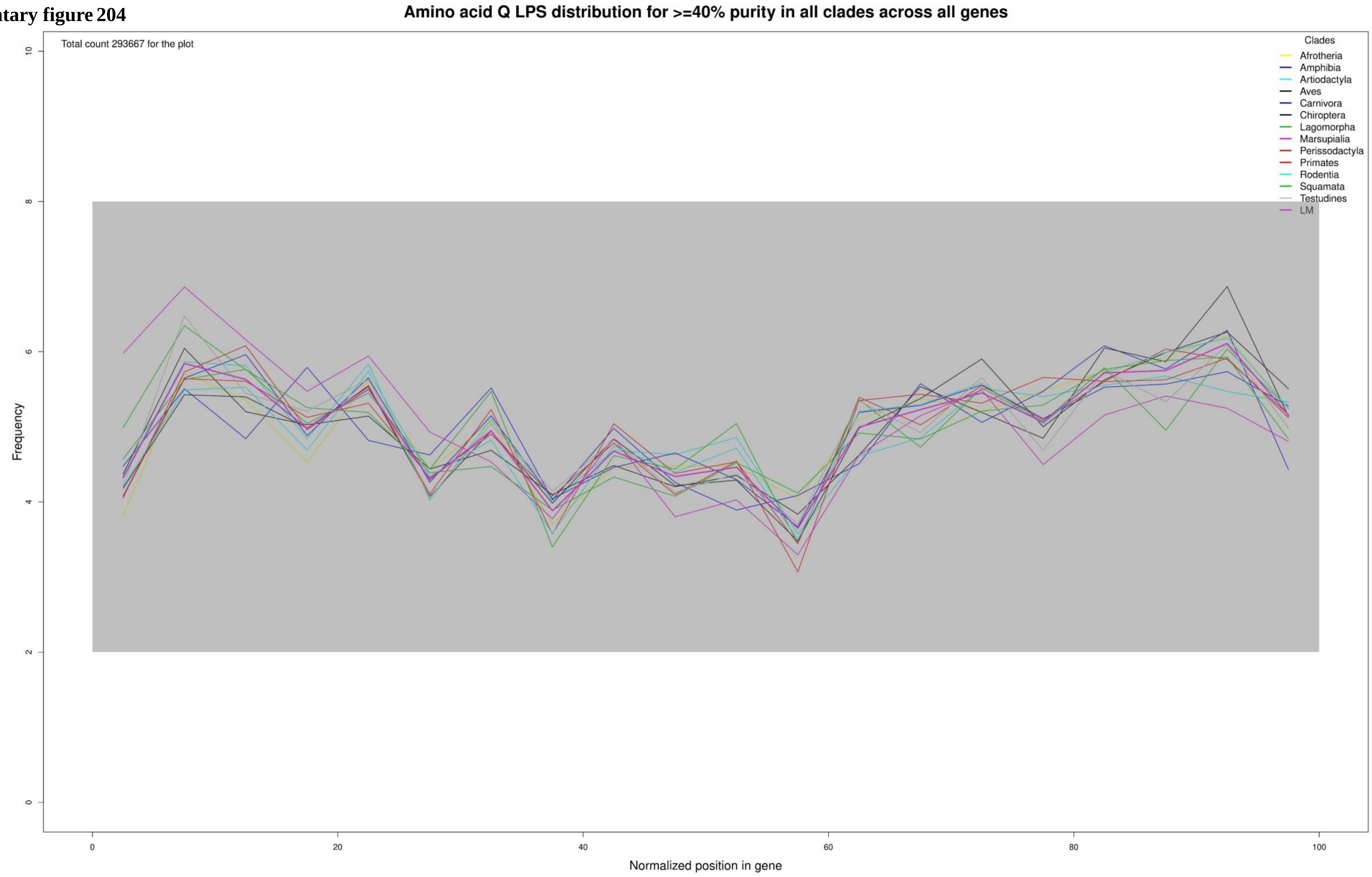

**Supplementary figure 205**

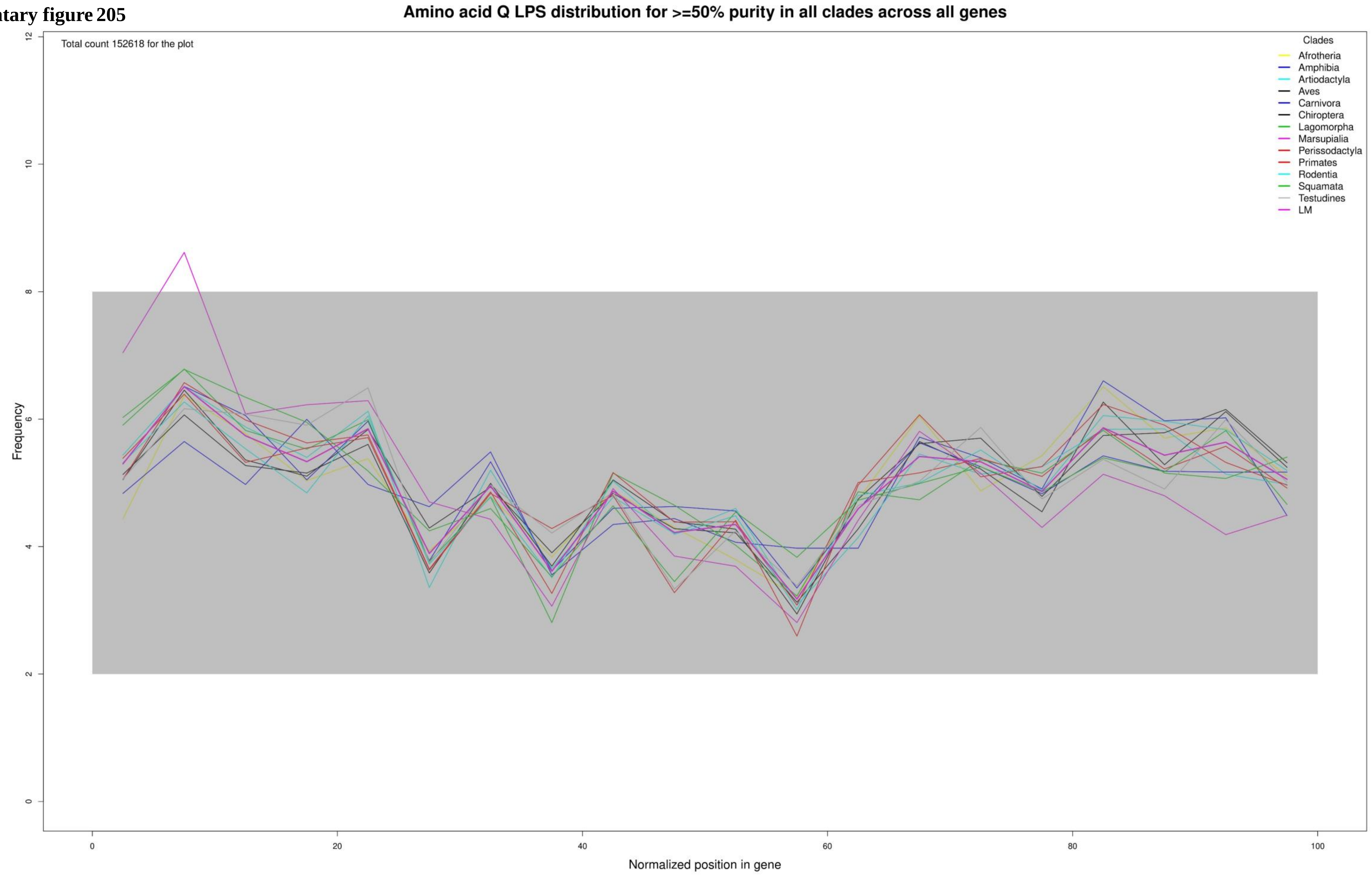

**Supplementary figure 206**

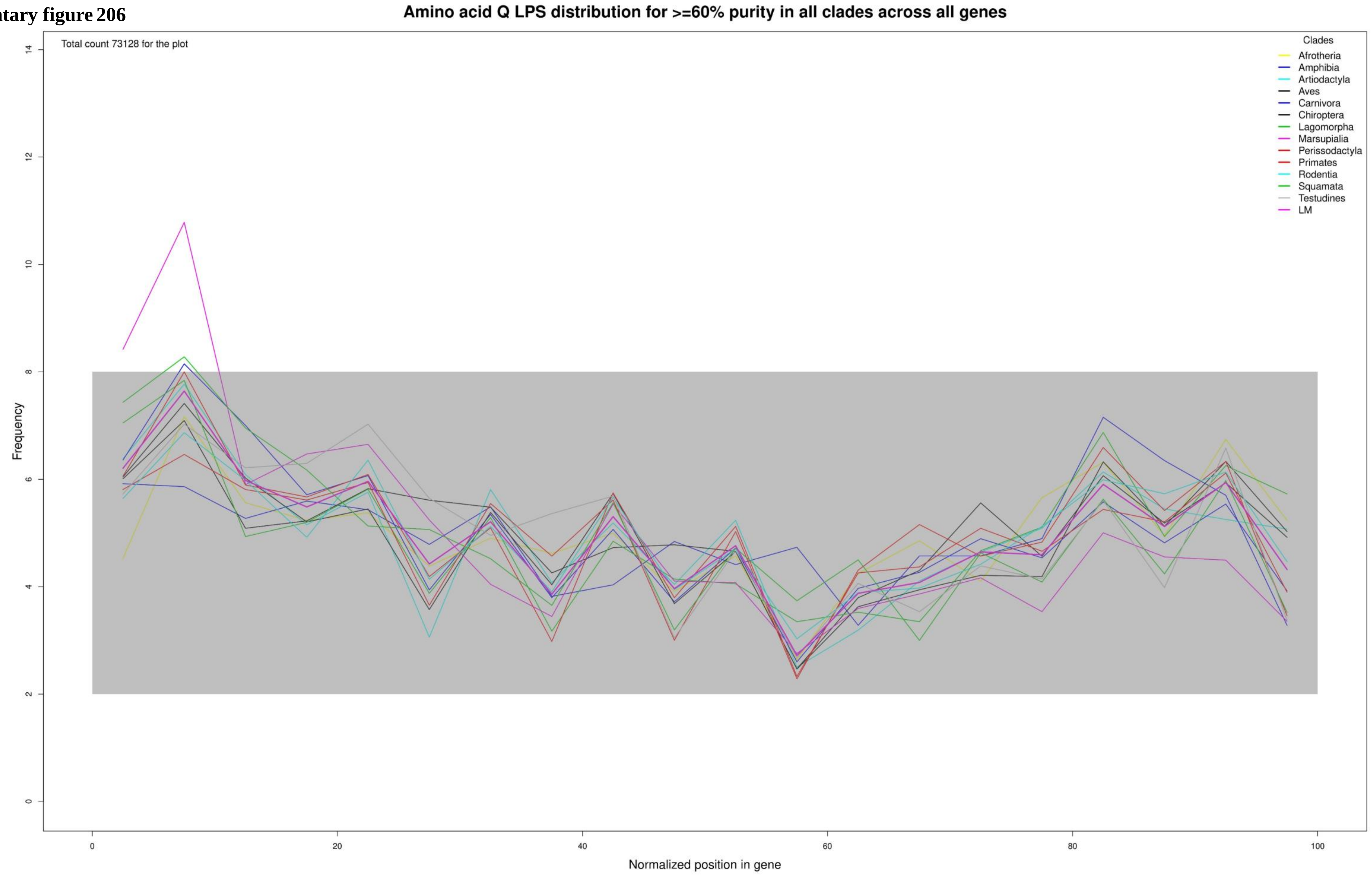

**Supplementary figure 207**

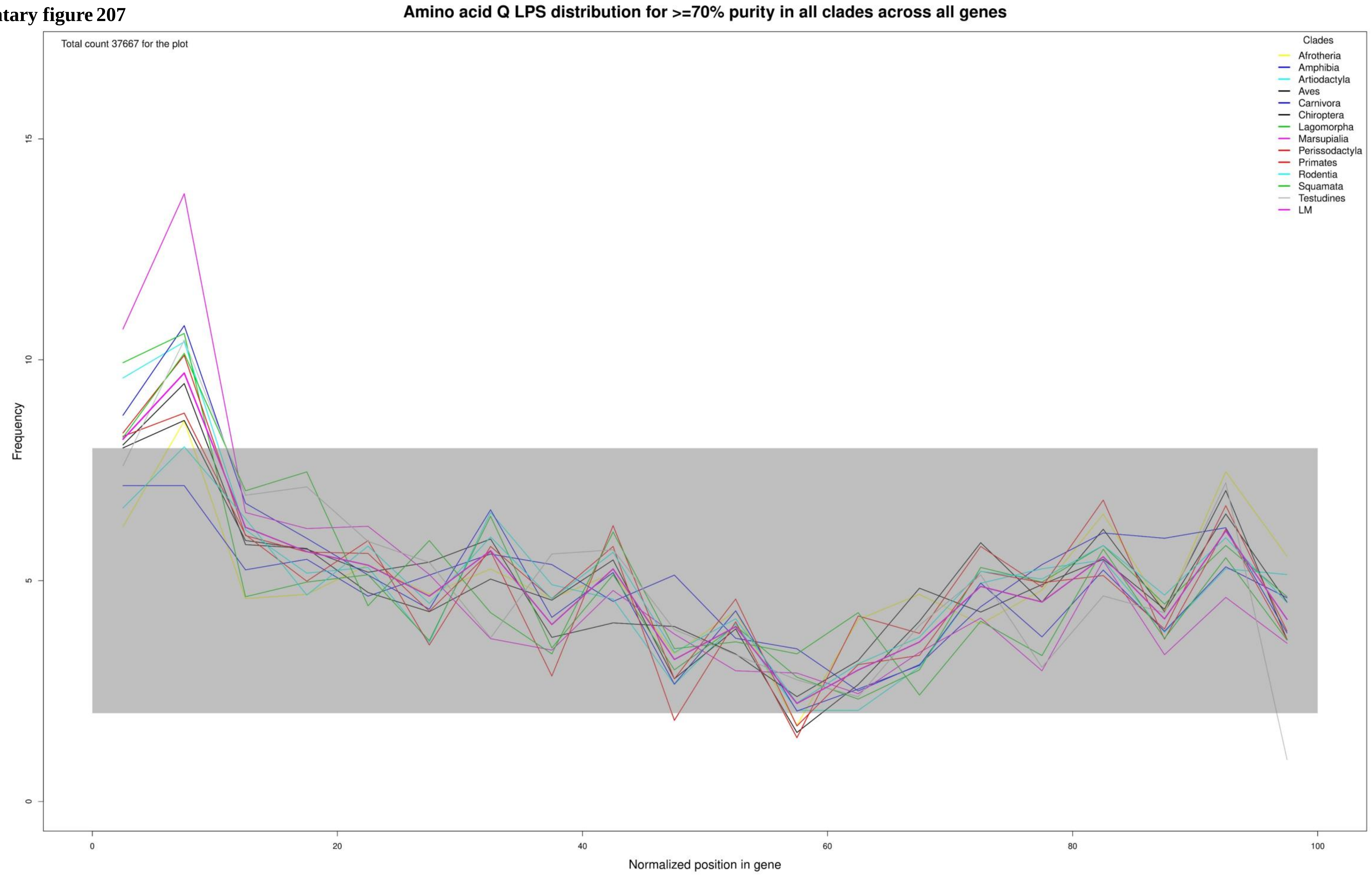

**Supplementary figure 208**

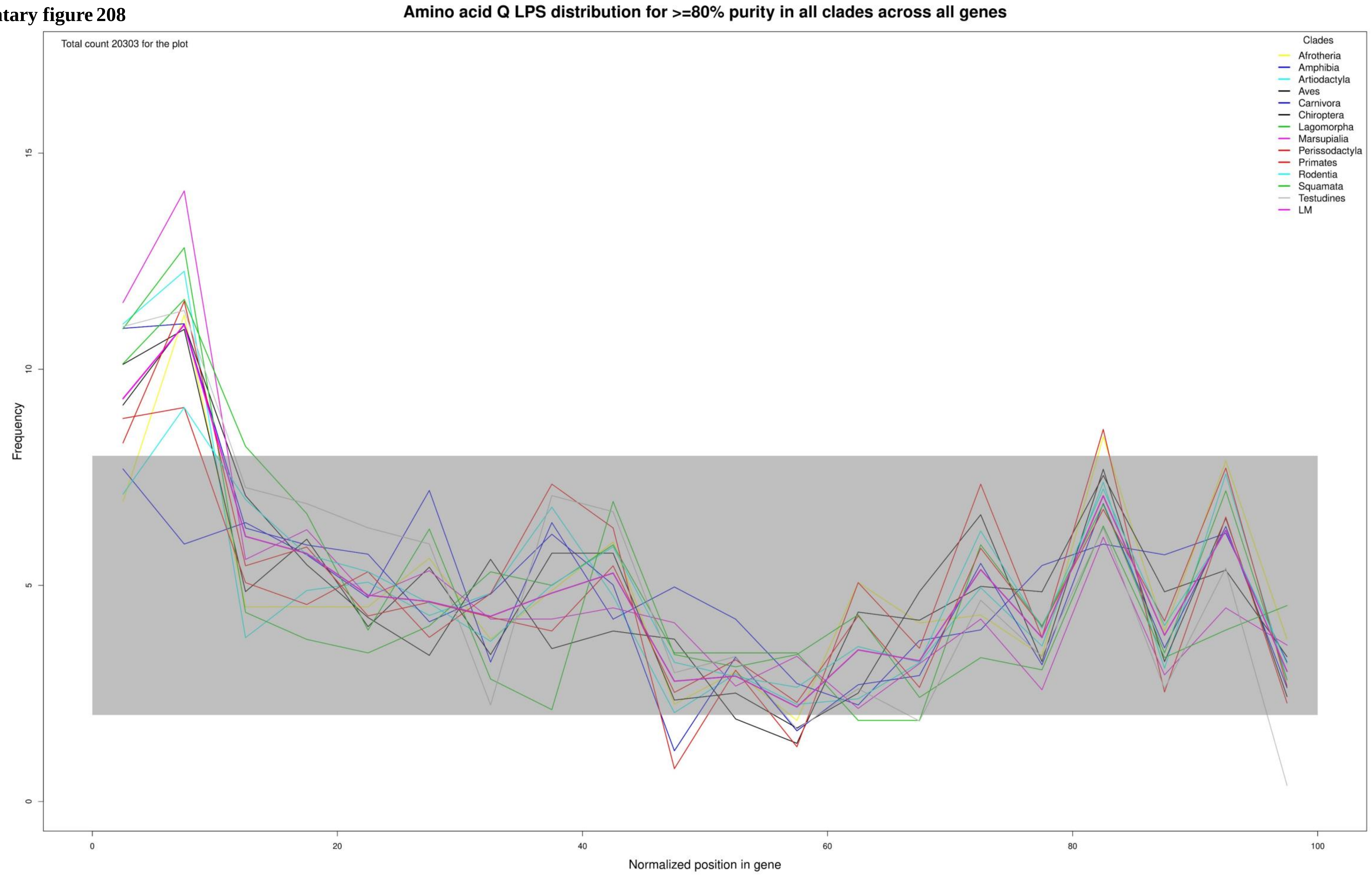

**Supplementary figure 209**

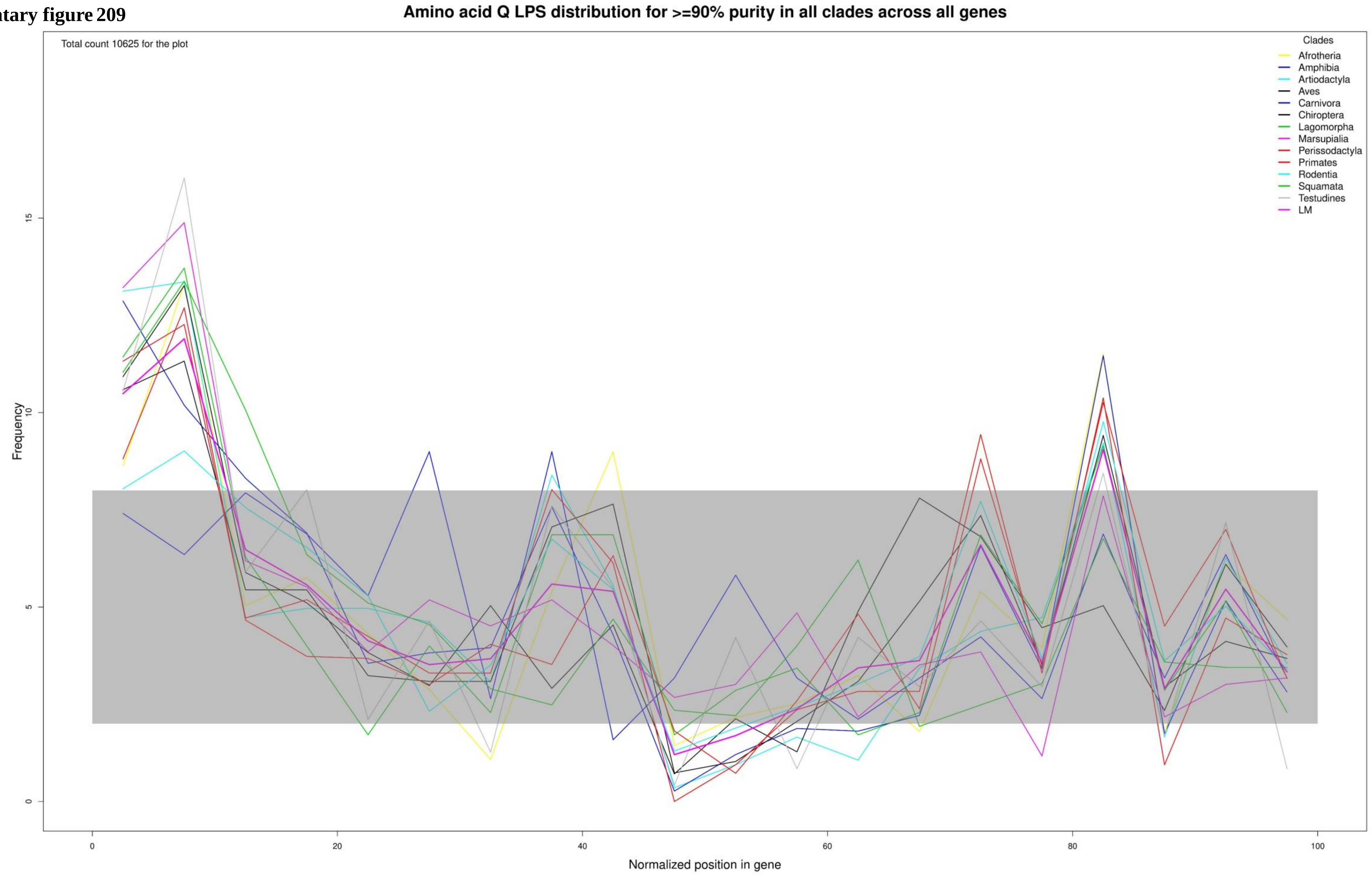

Supplementary figure 210

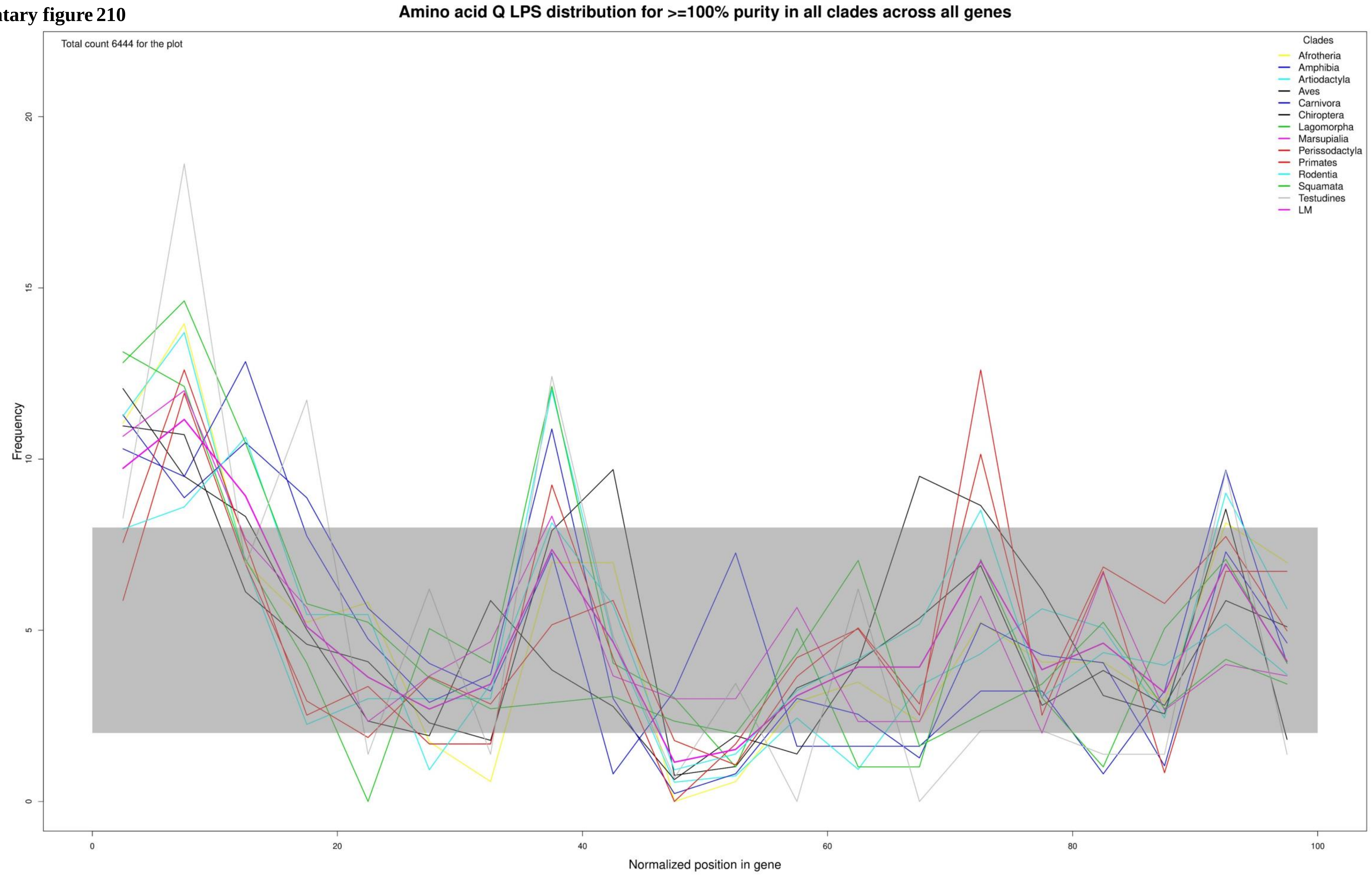

**Supplementary figure 211**

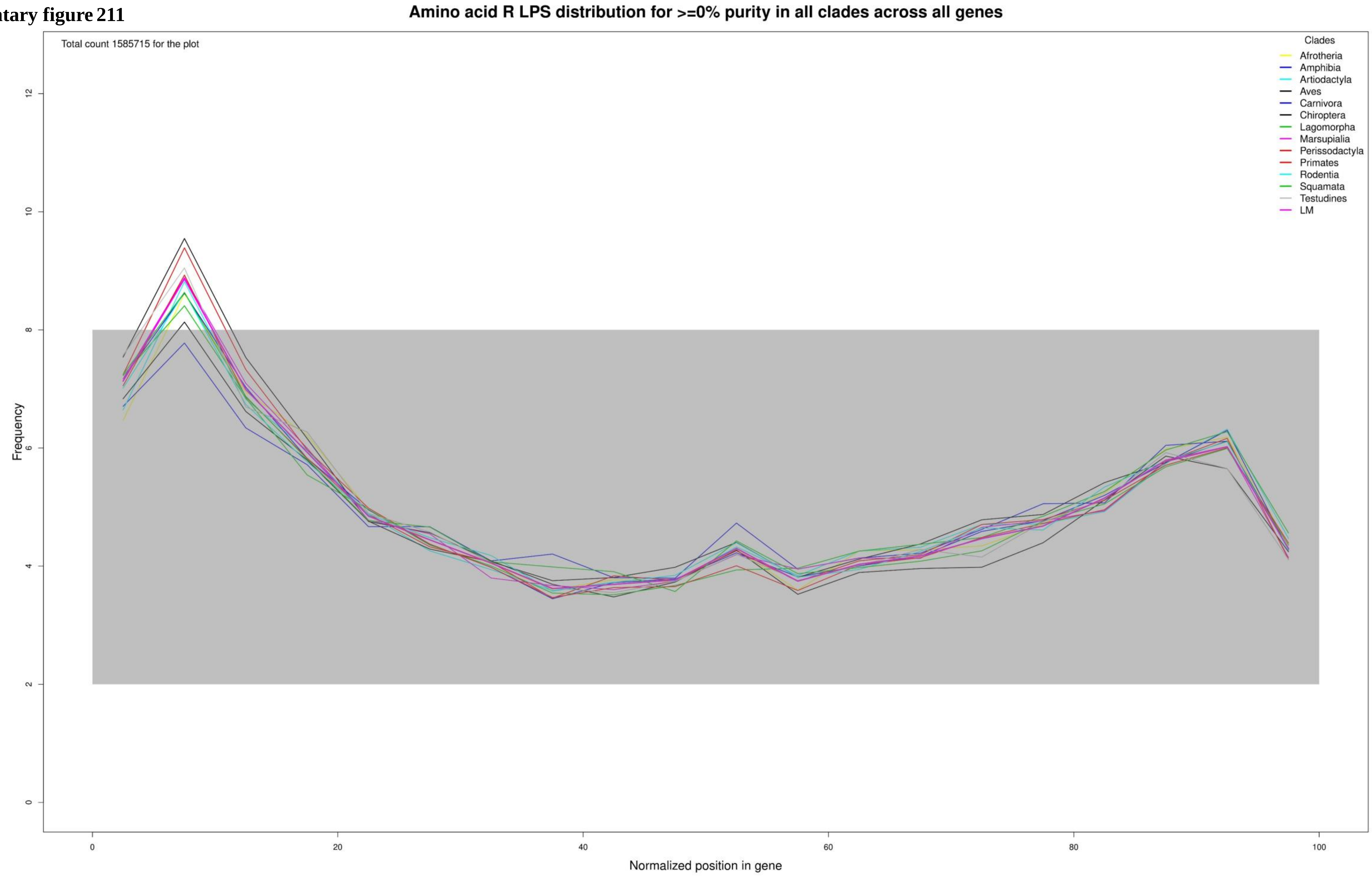

**Supplementary figure 212**

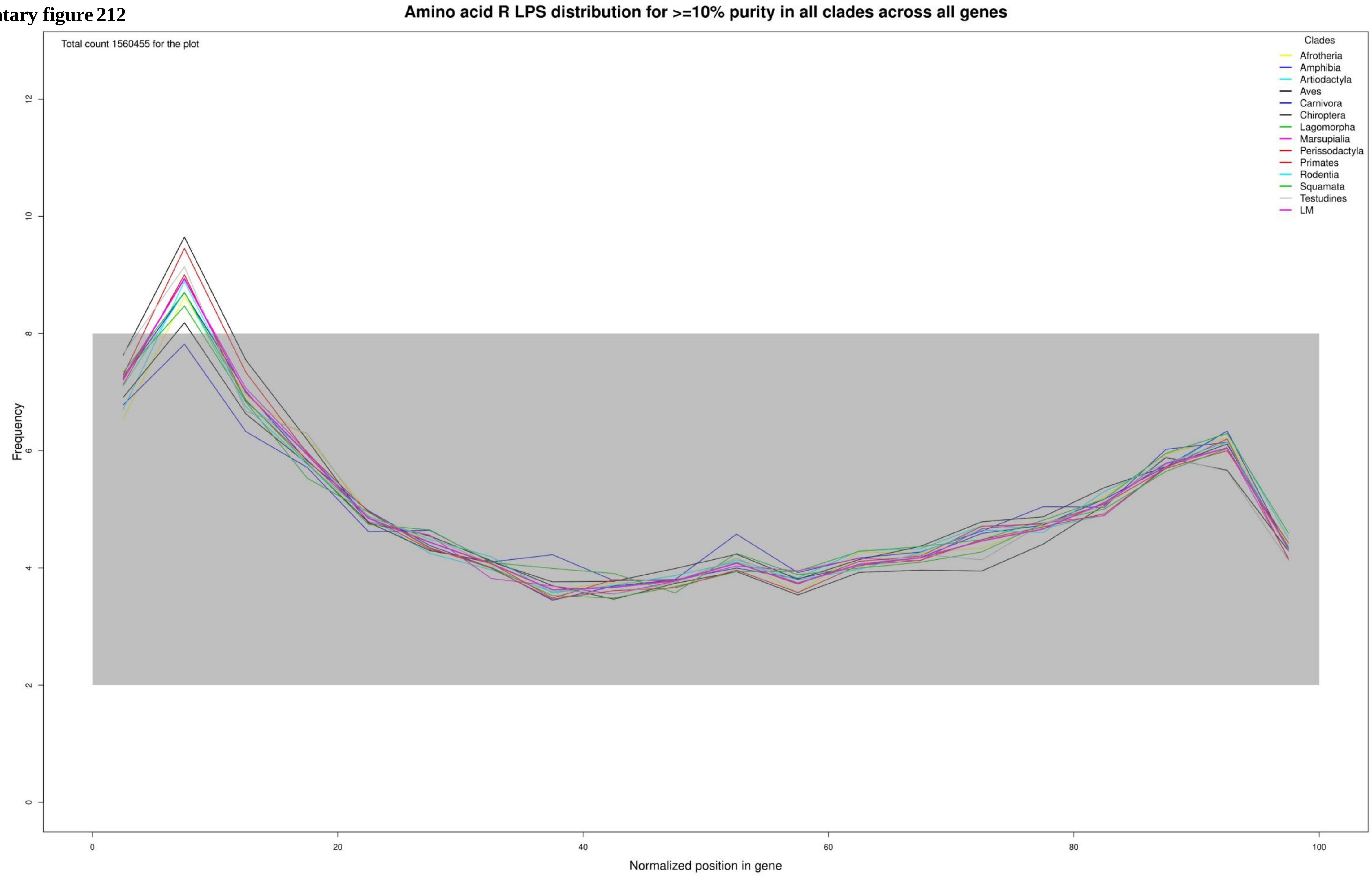

**Supplementary figure 213**

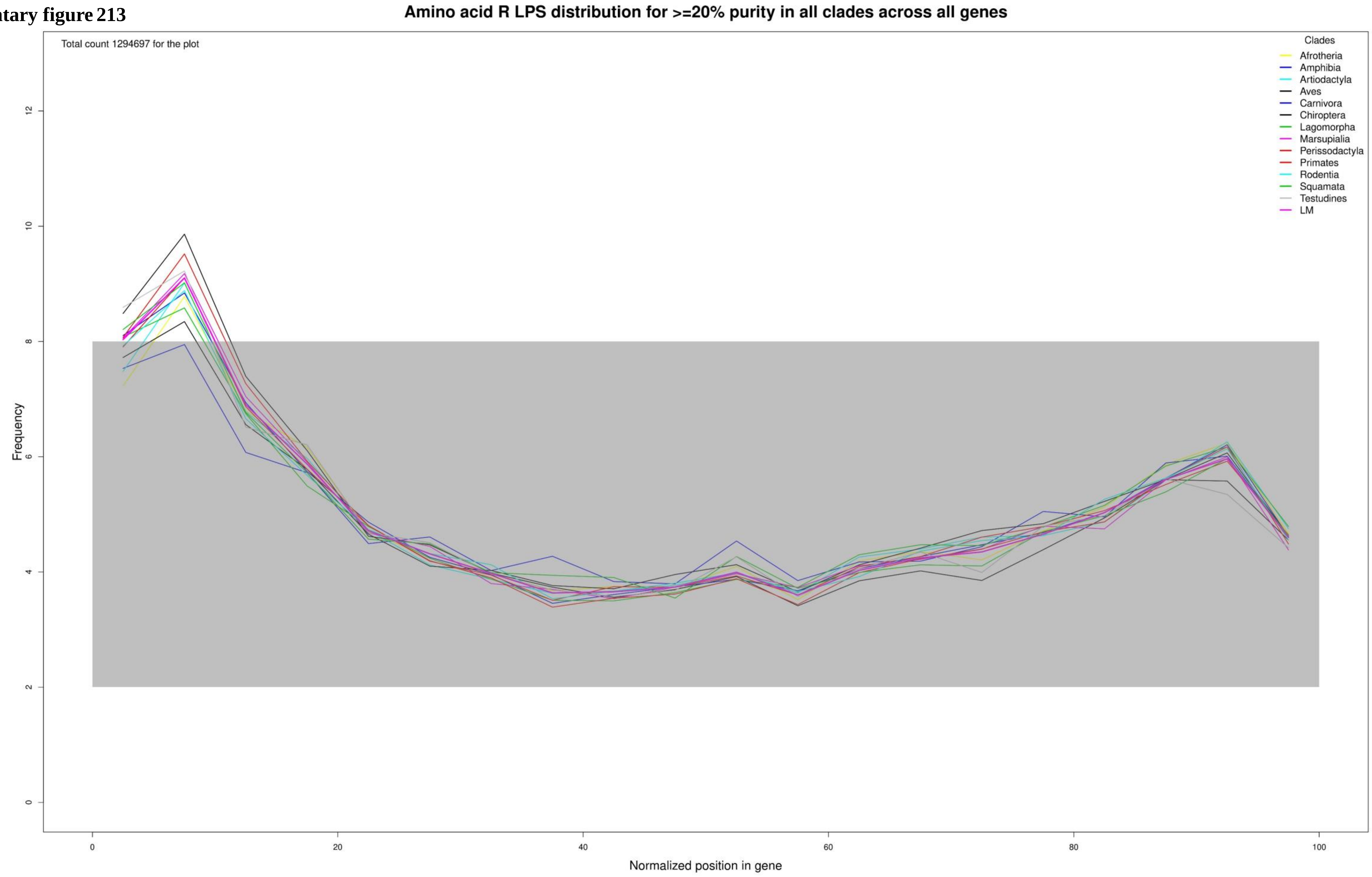

**Supplementary figure 214**

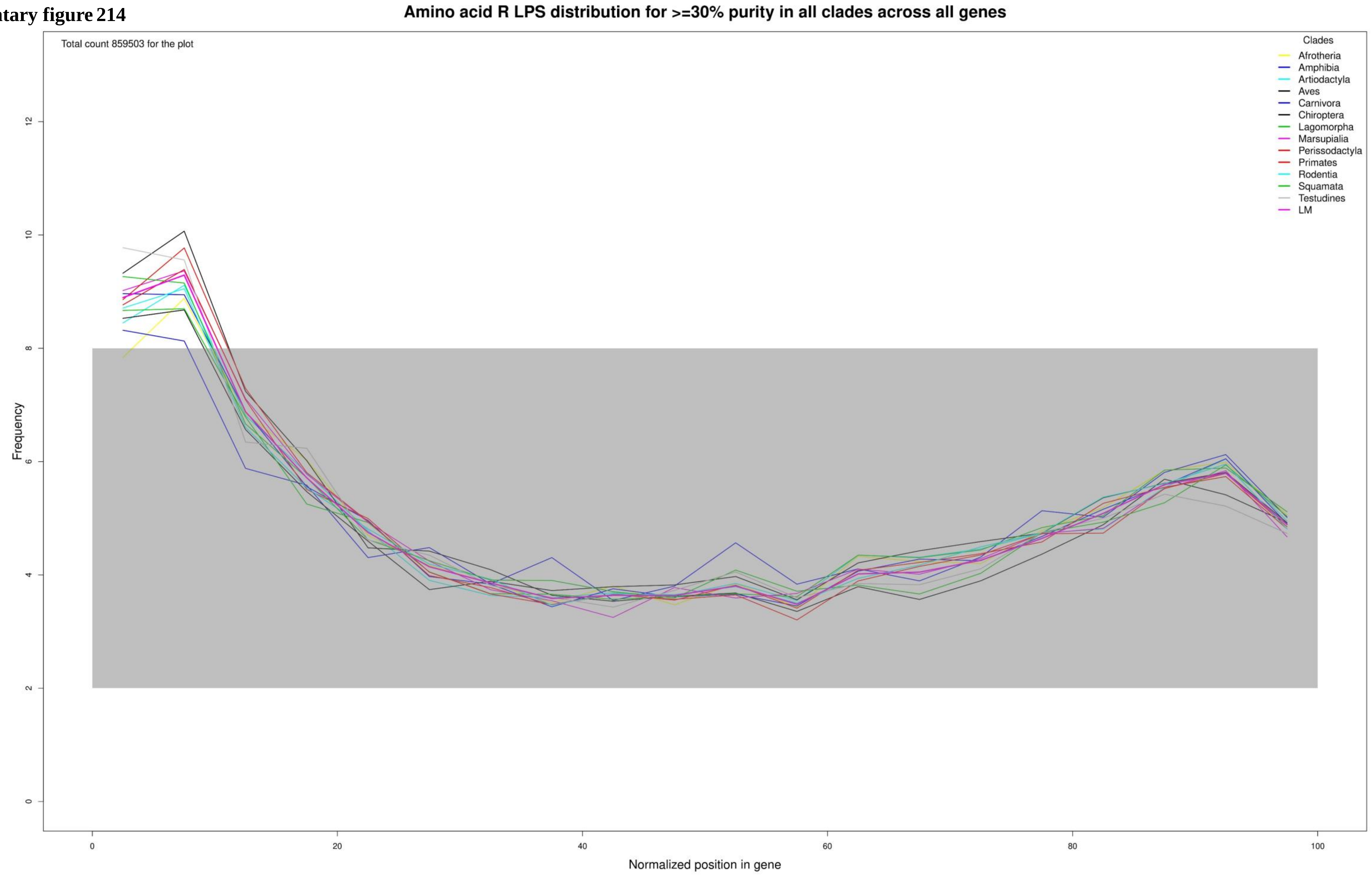

**Supplementary figure 215**

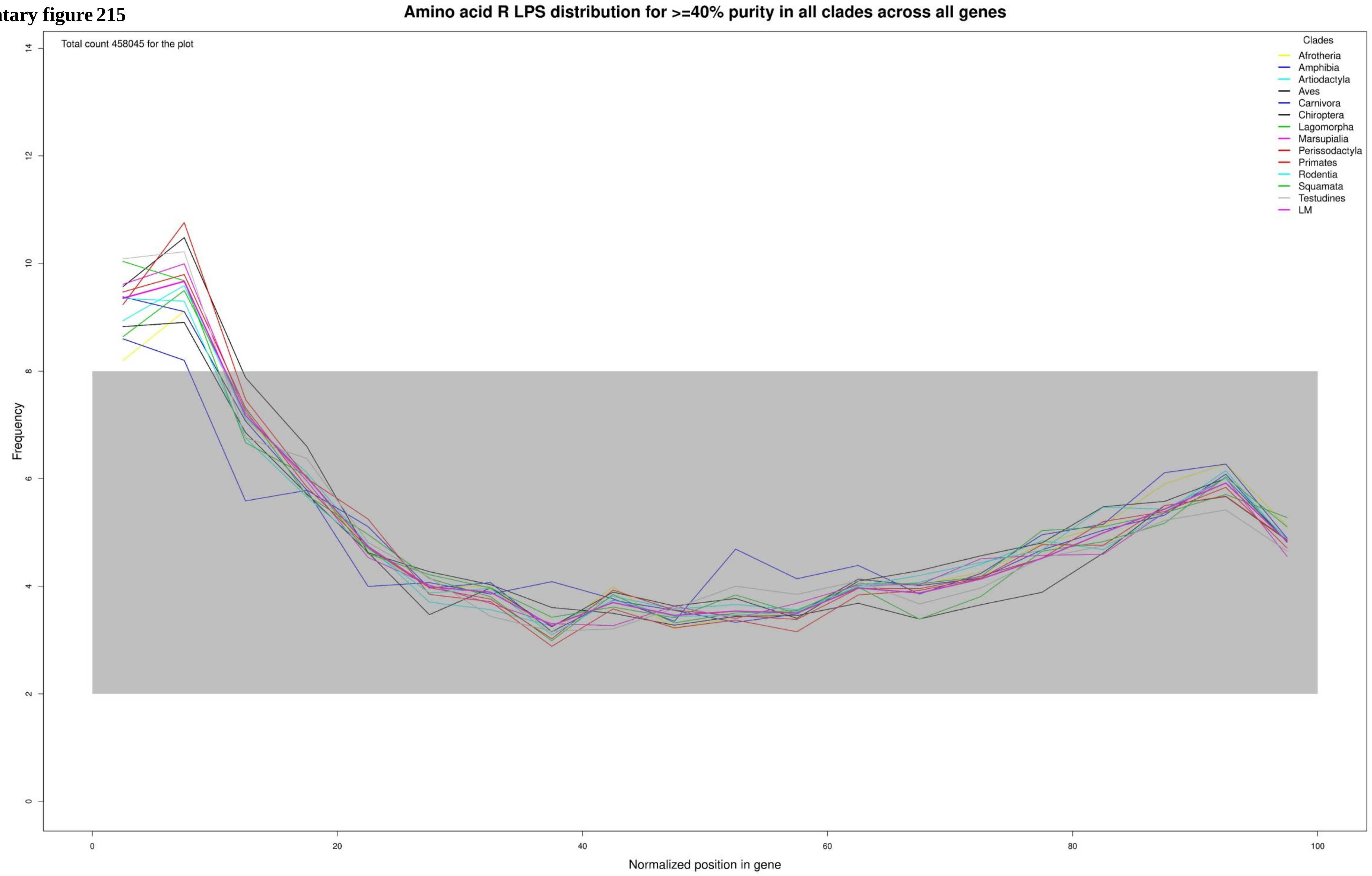

**Supplementary figure 216**

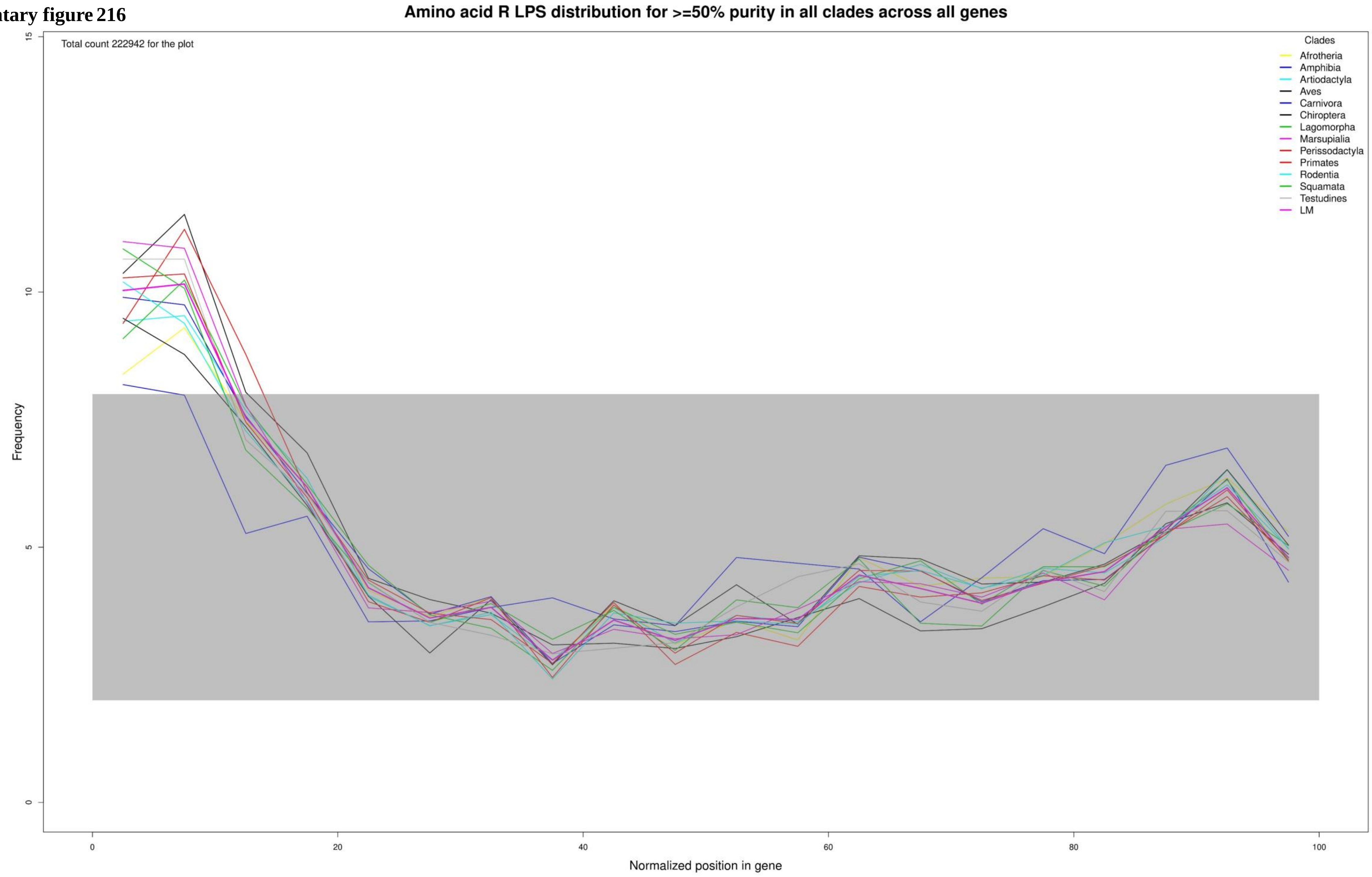

**Supplementary figure 217**

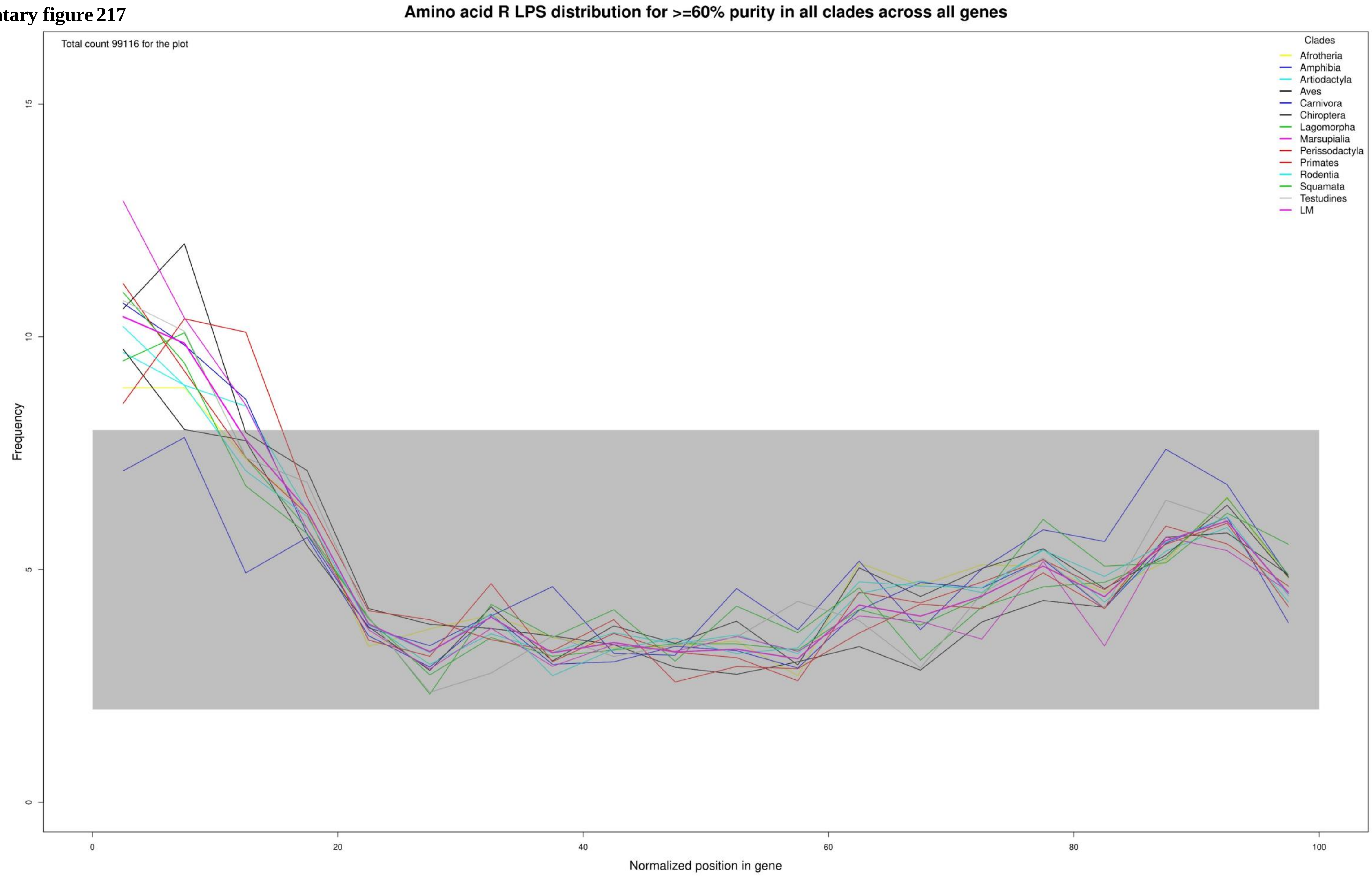

**Supplementary figure 218**

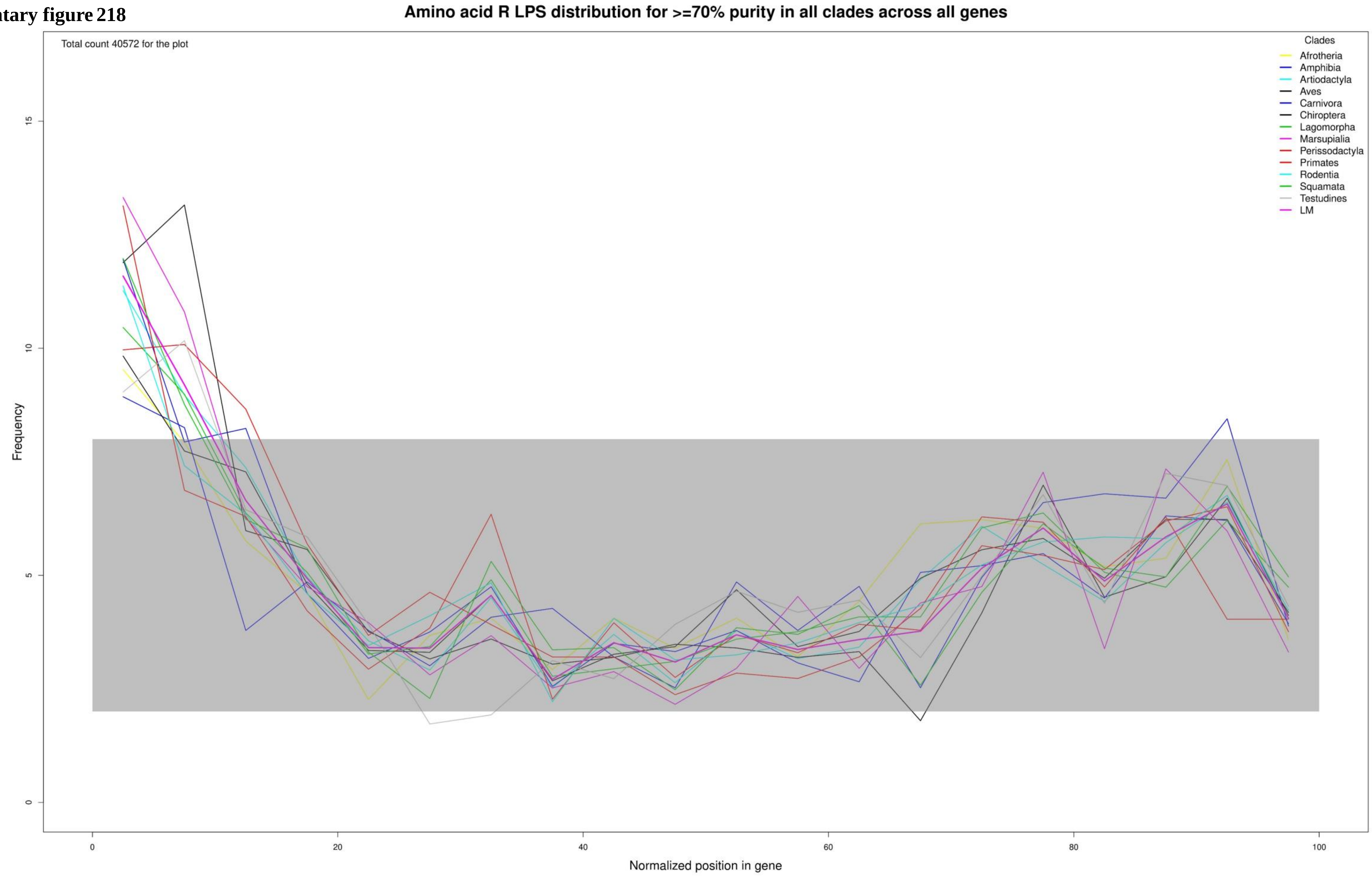

**Supplementary figure 219**

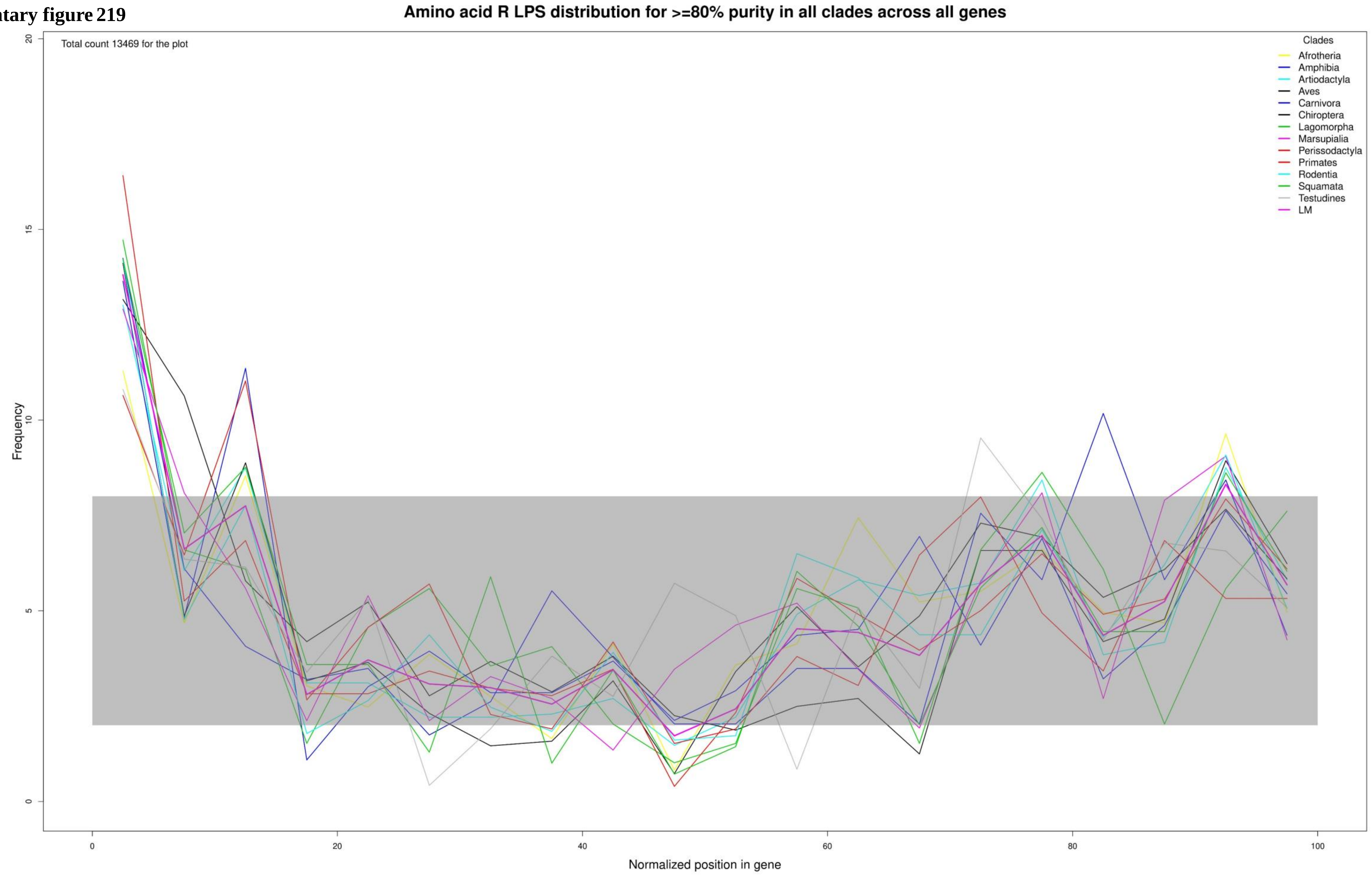

**Supplementary figure 220**

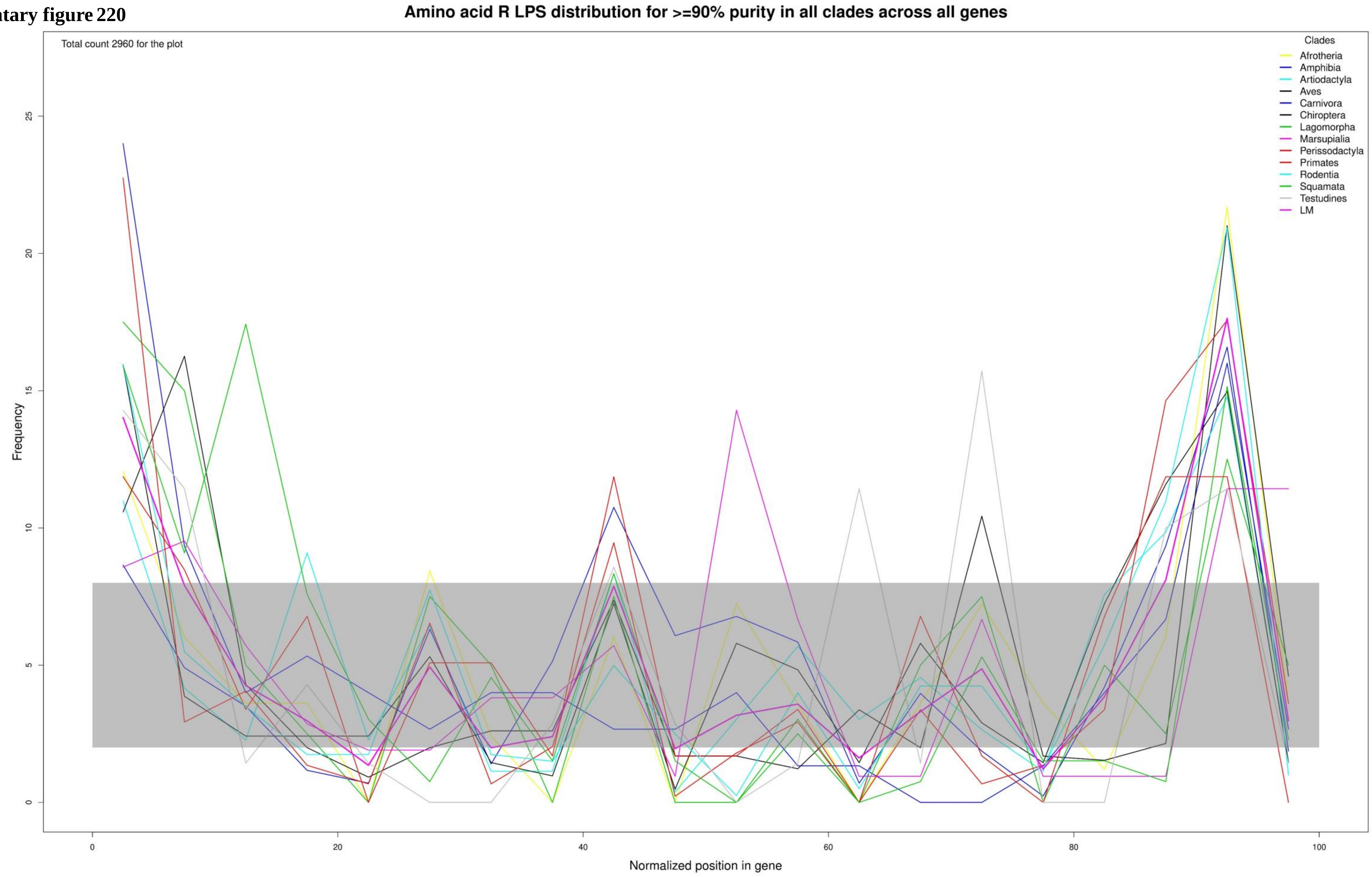

Supplementary figure 221

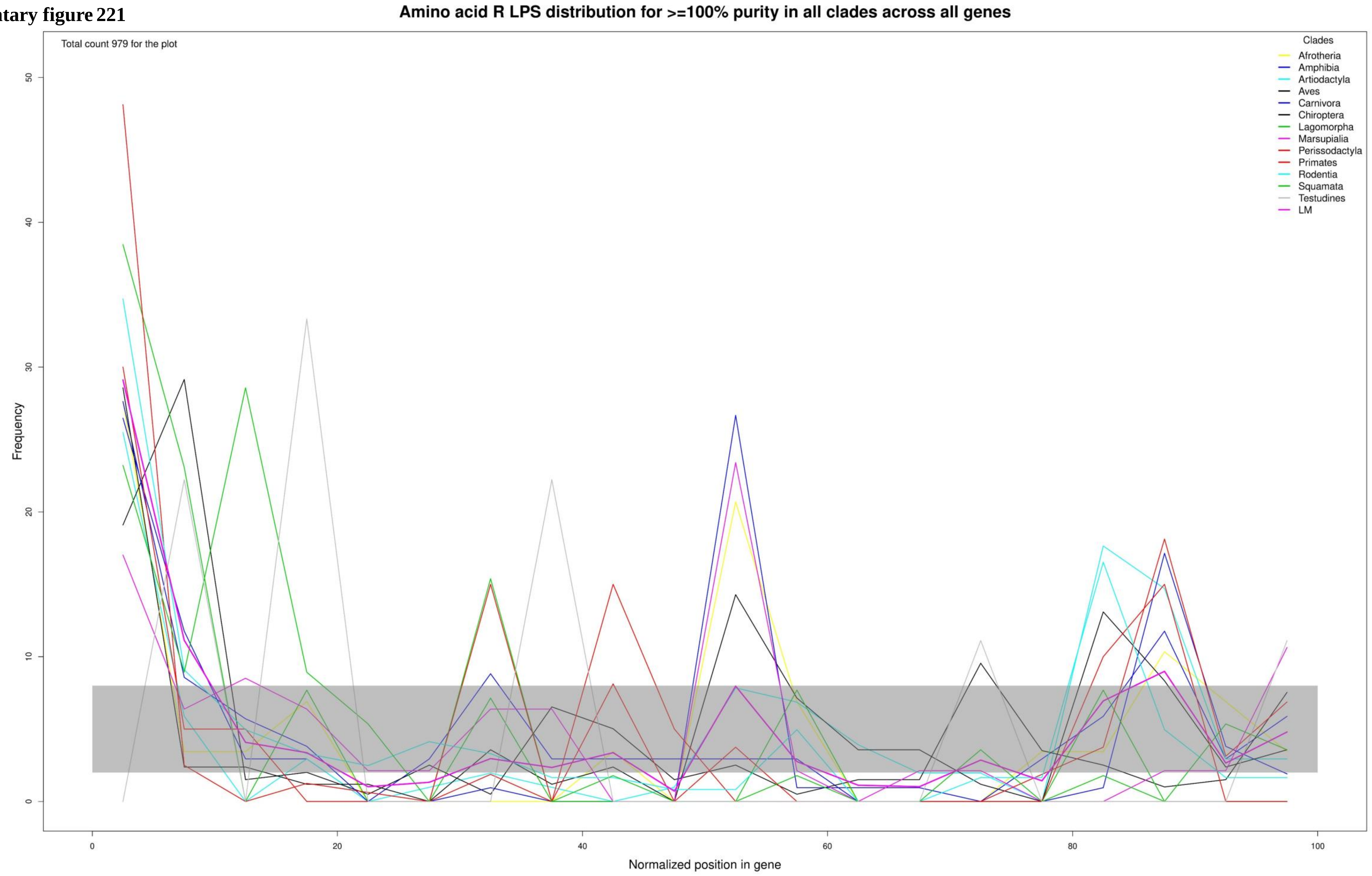

**Supplementary figure 222**

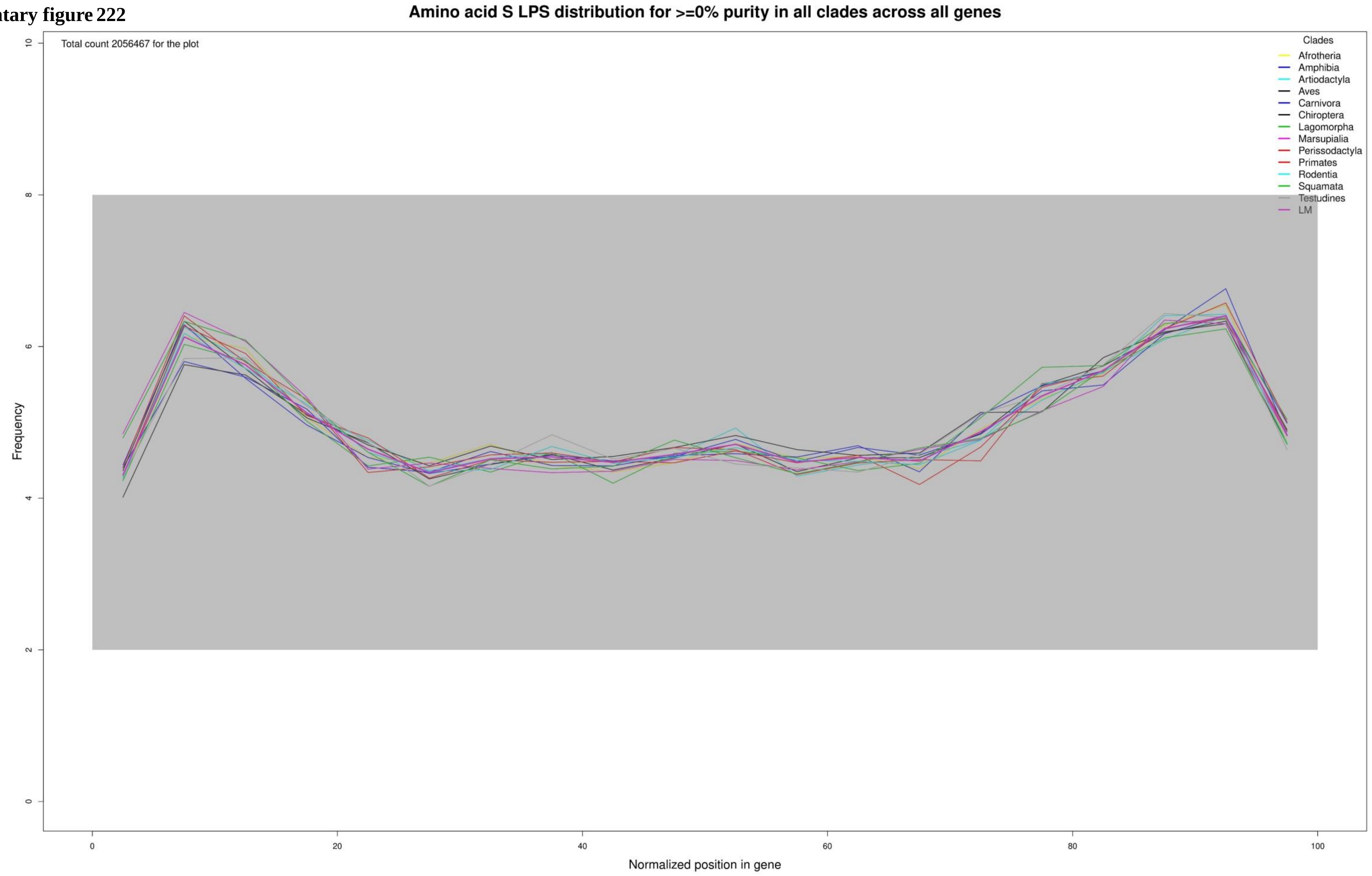

**Supplementary figure 223**

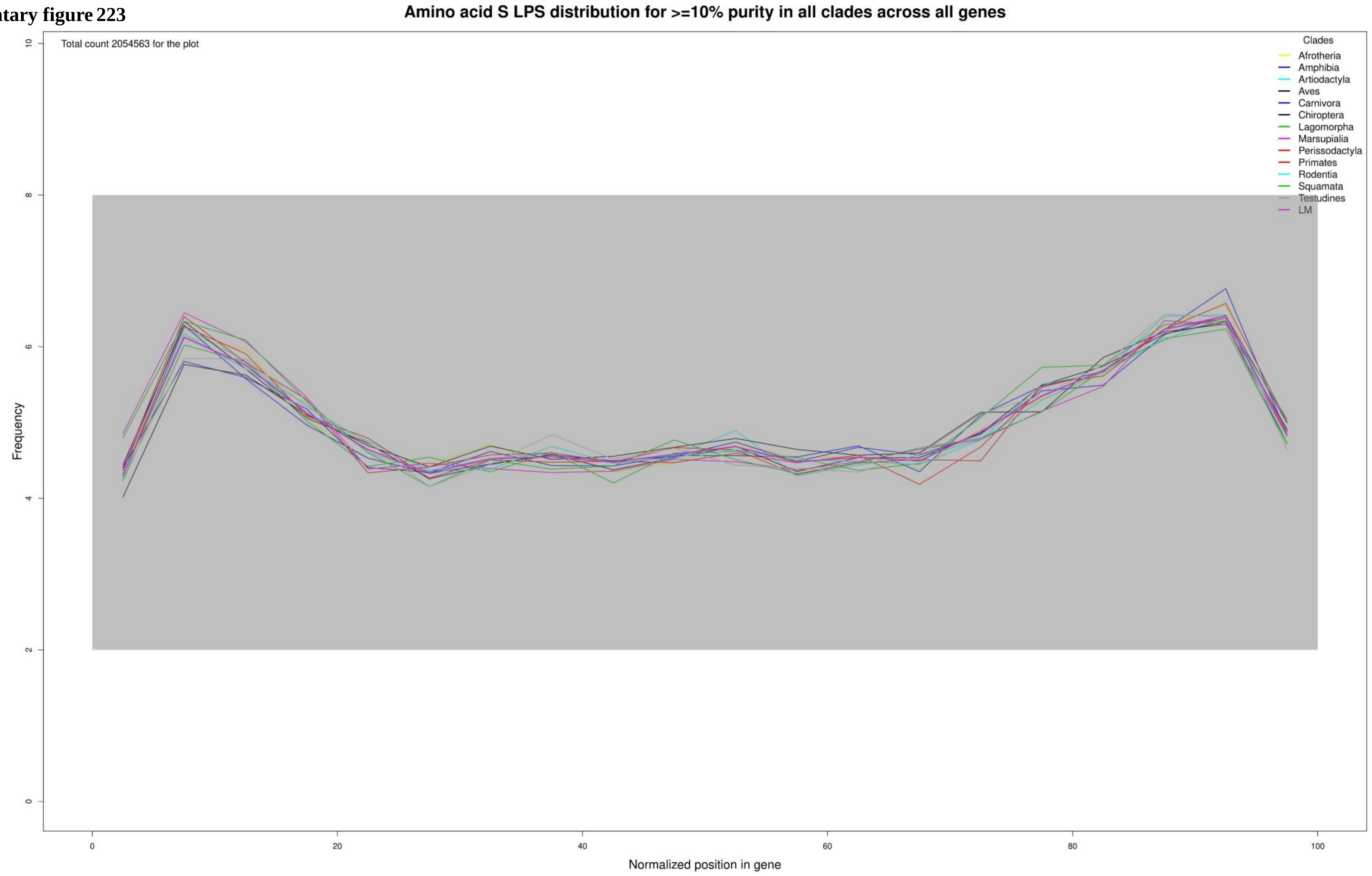

**Supplementary figure 224**

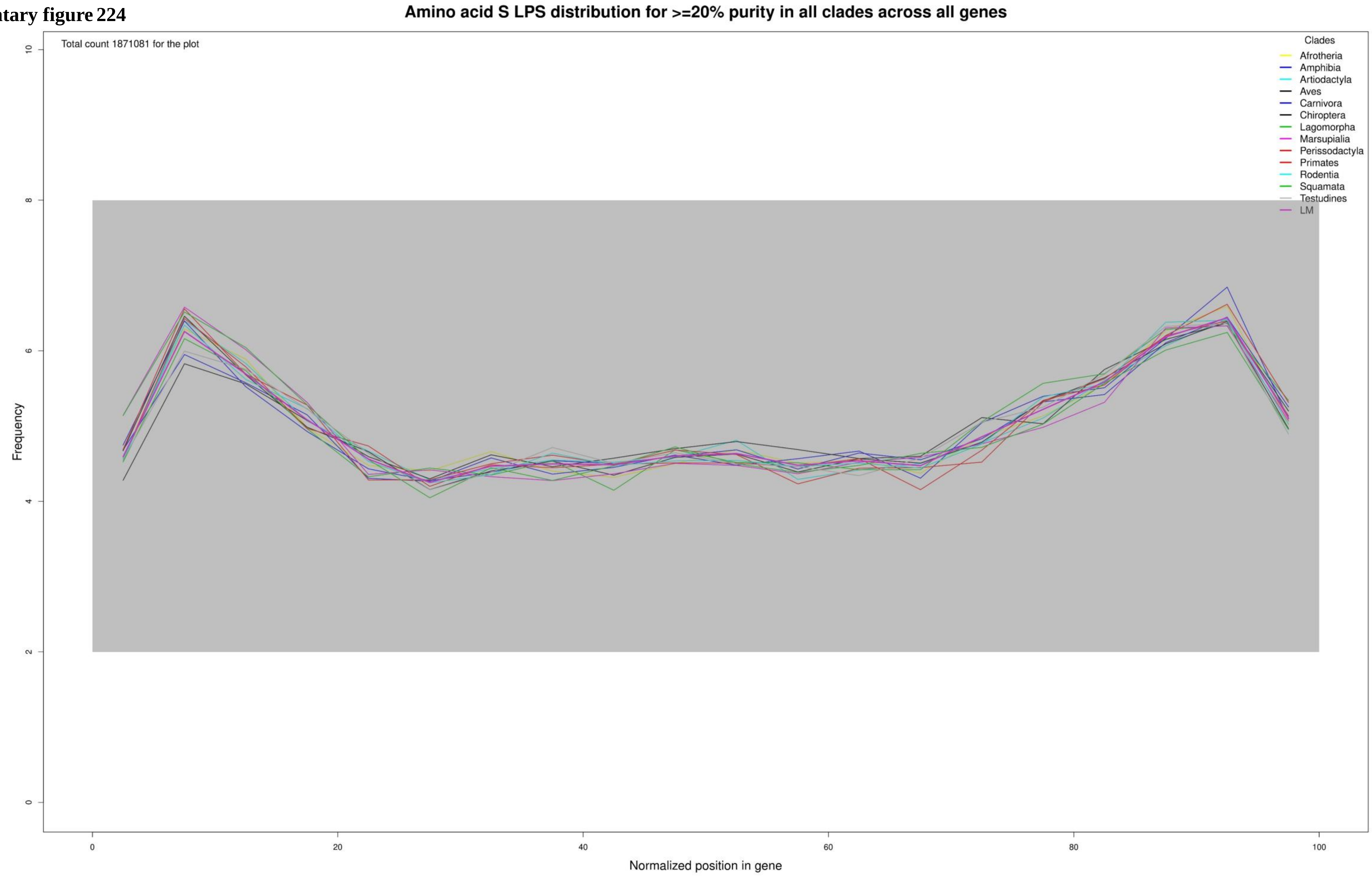

**Supplementary figure 225**

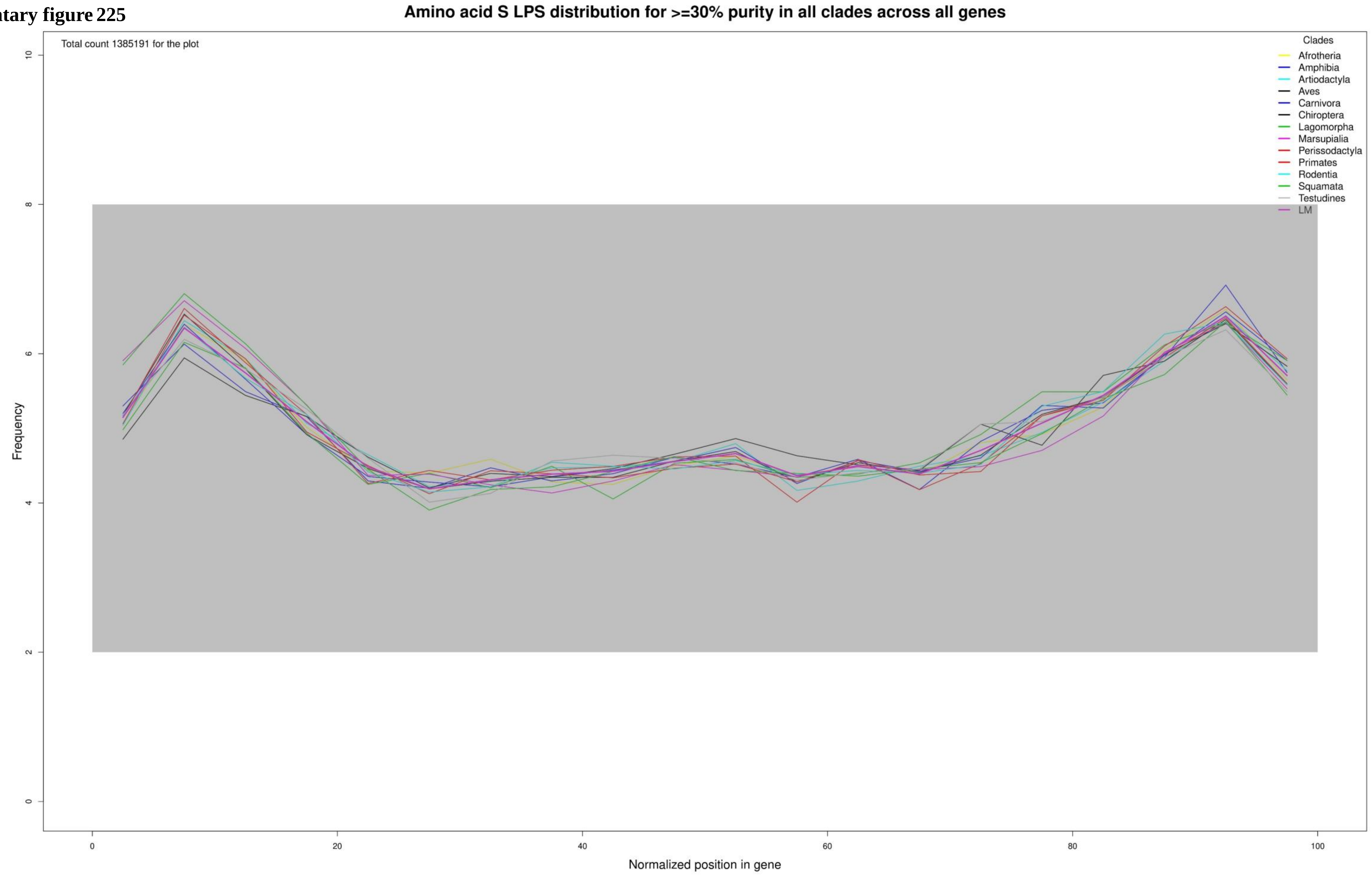

**Supplementary figure 226**

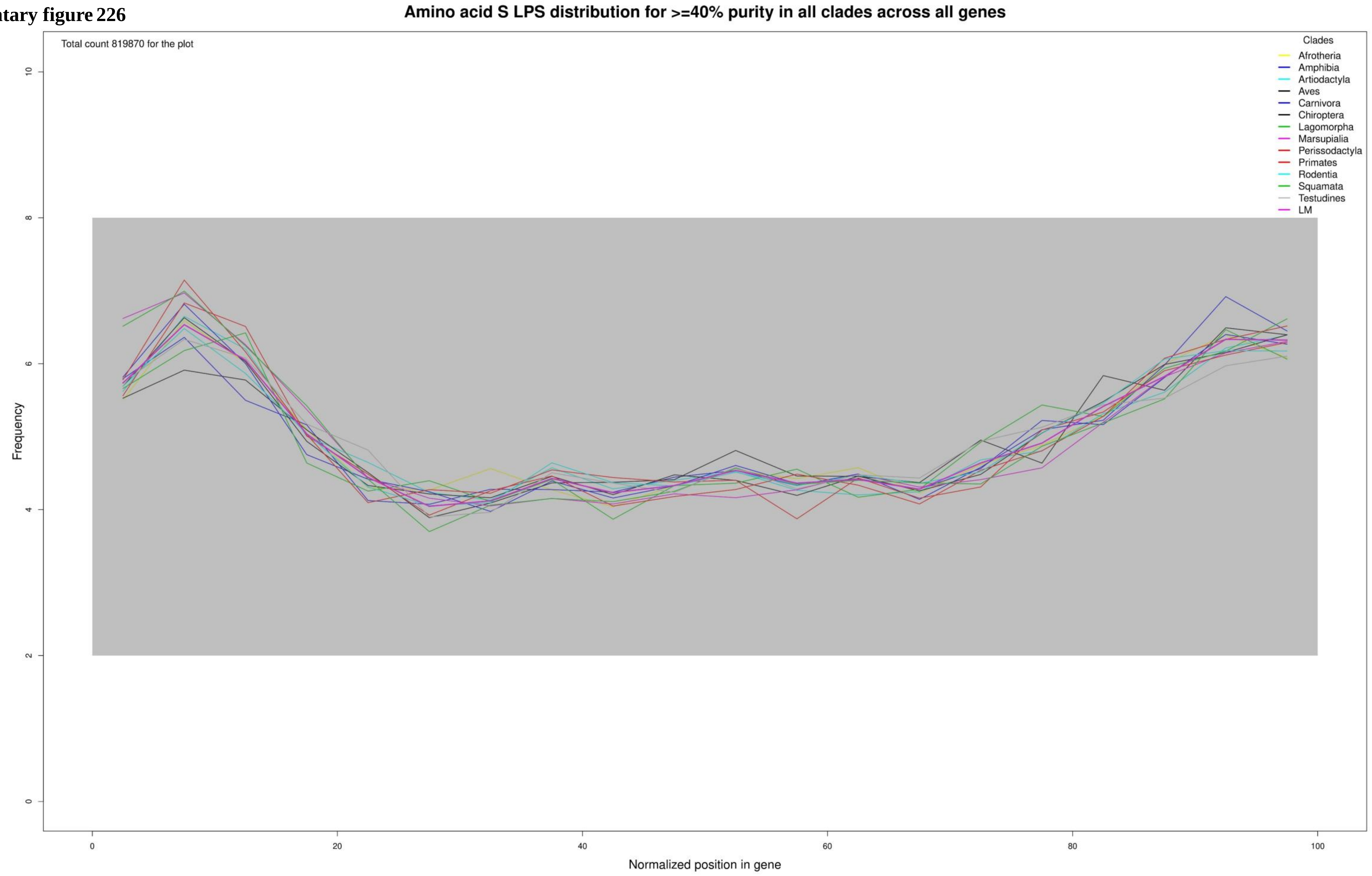

**Supplementary figure 227**

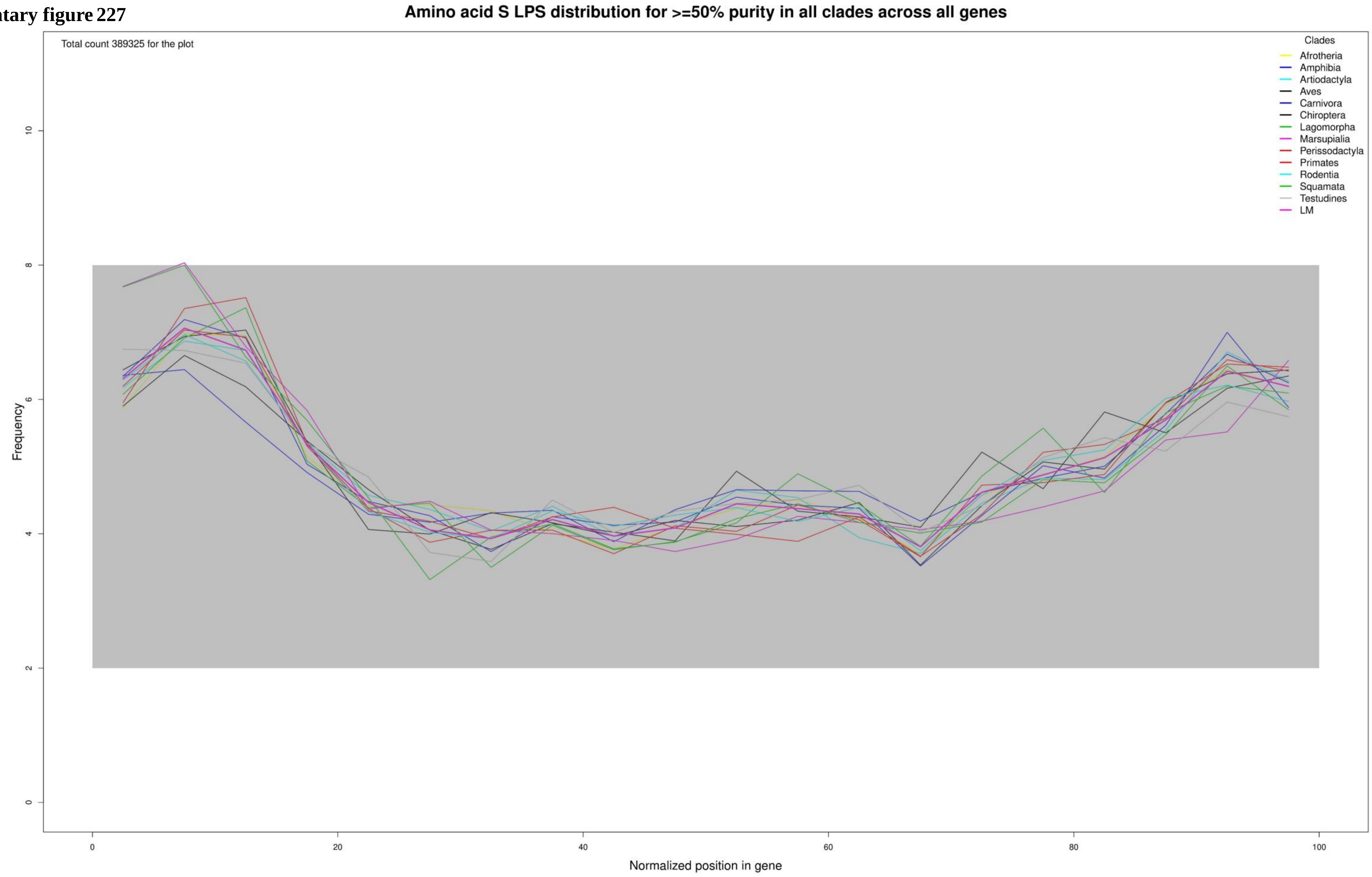

**Supplementary figure 228**

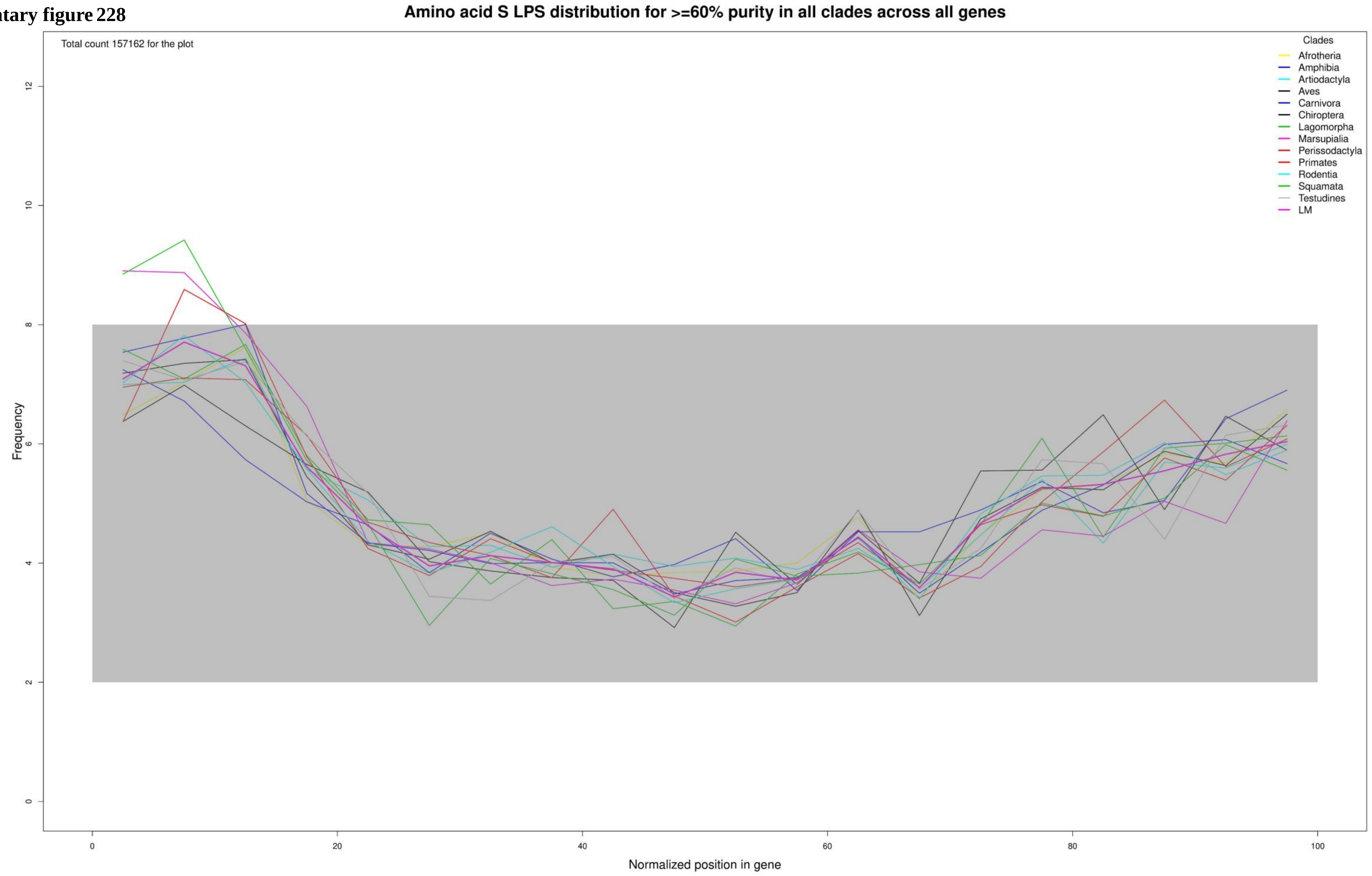

**Supplementary figure 229**

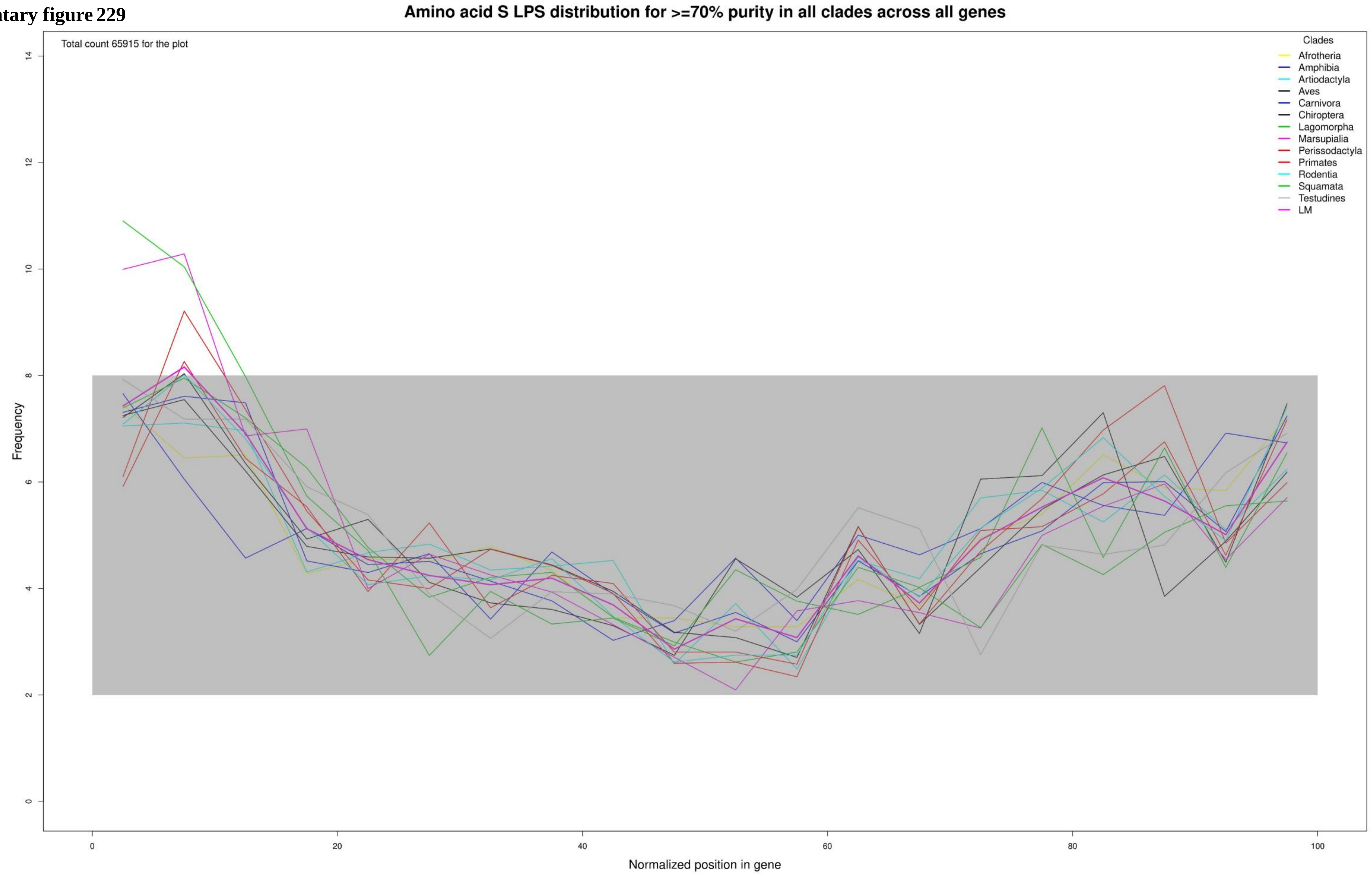

**Supplementary figure 230**

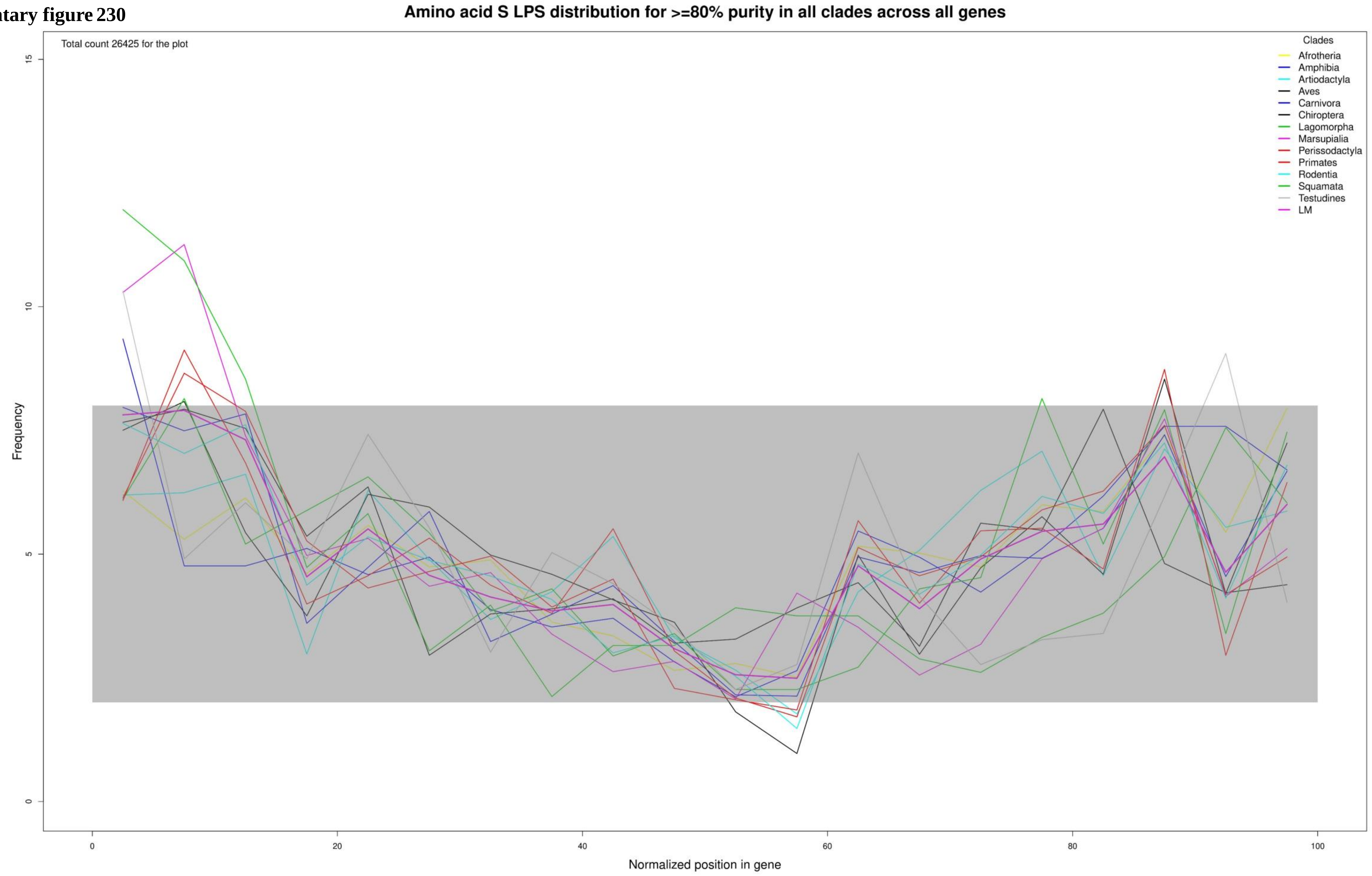

**Supplementary figure 231**

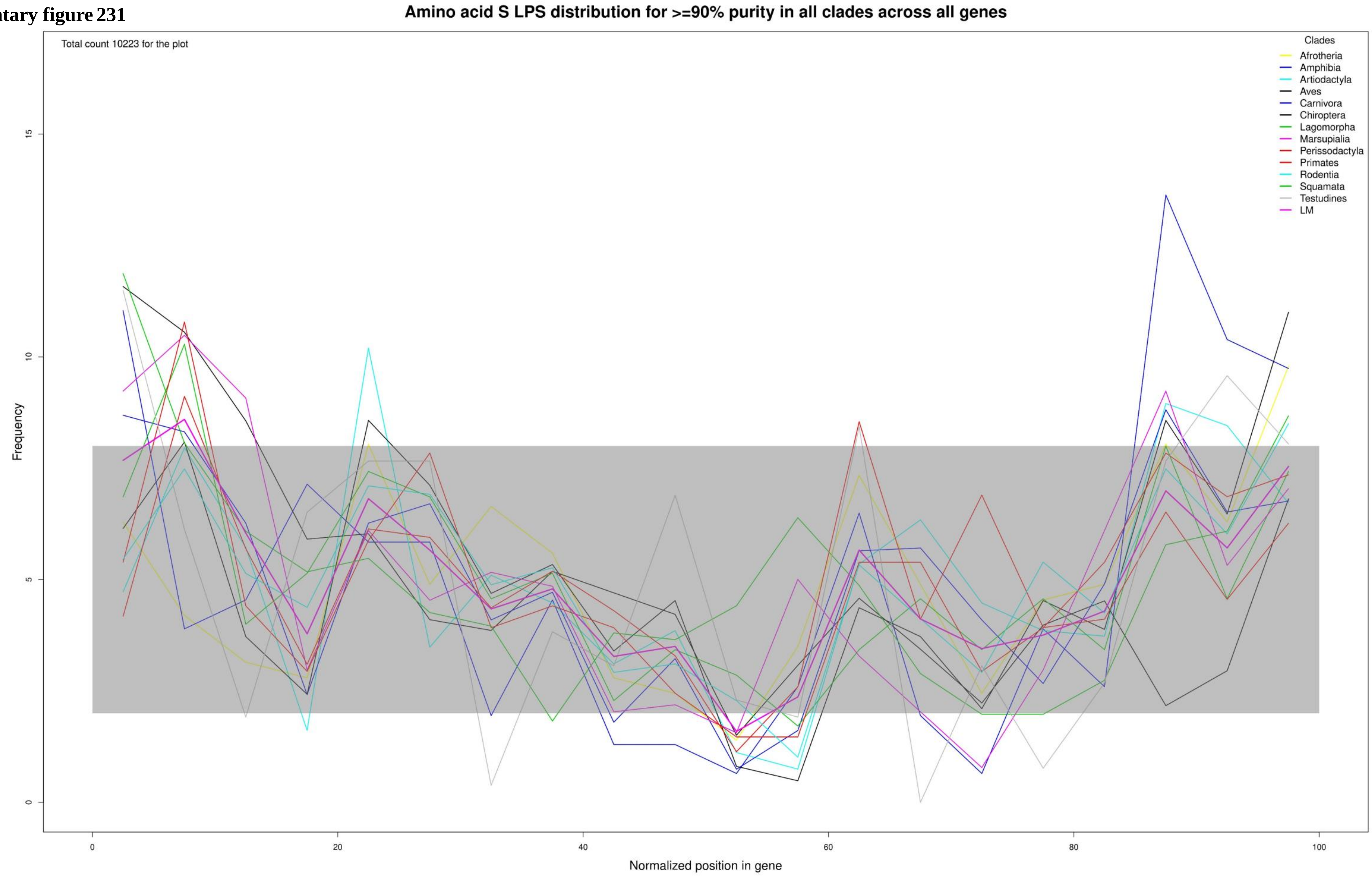

**Supplementary figure 232**

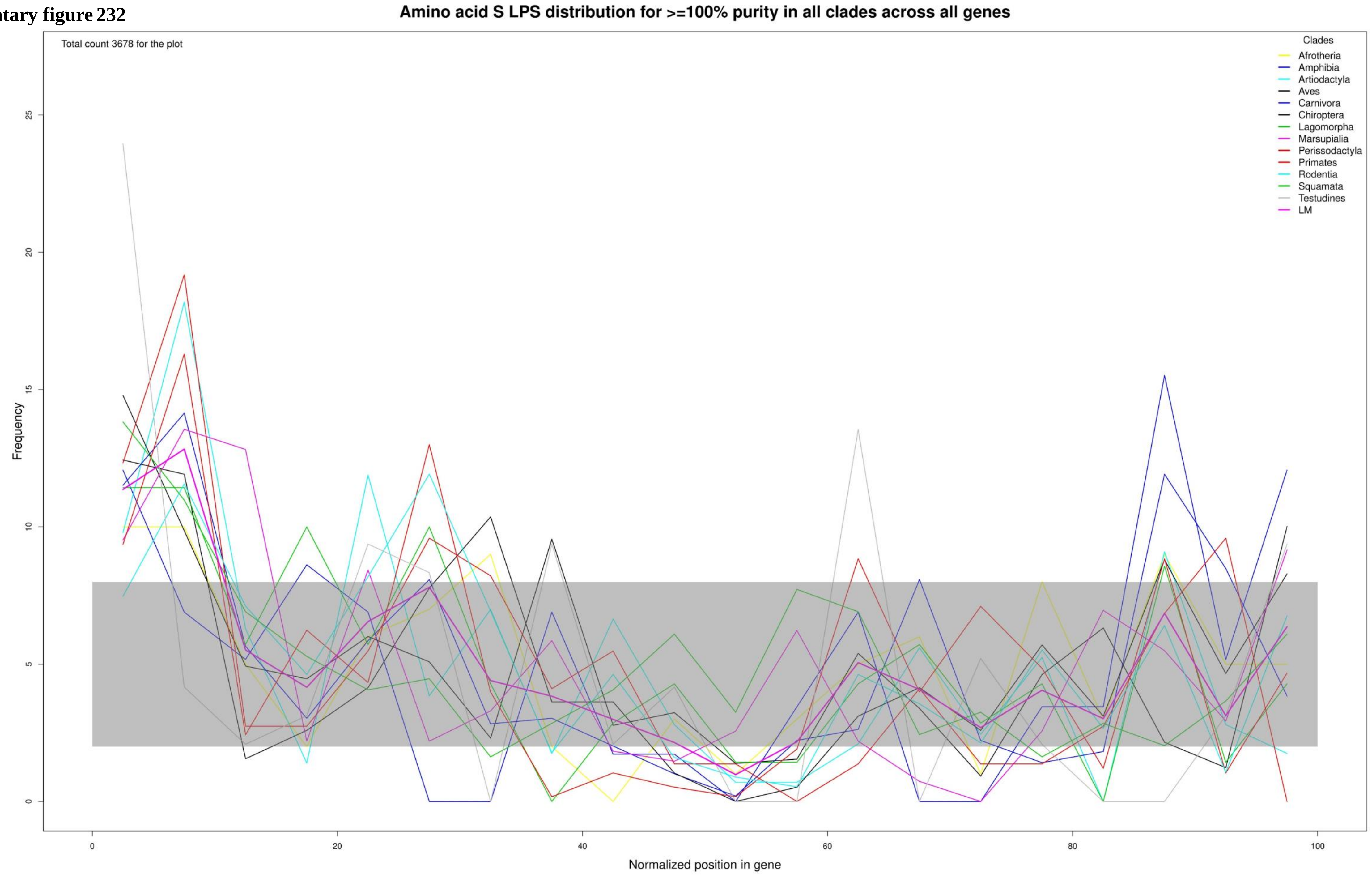

Supplementary figure 233

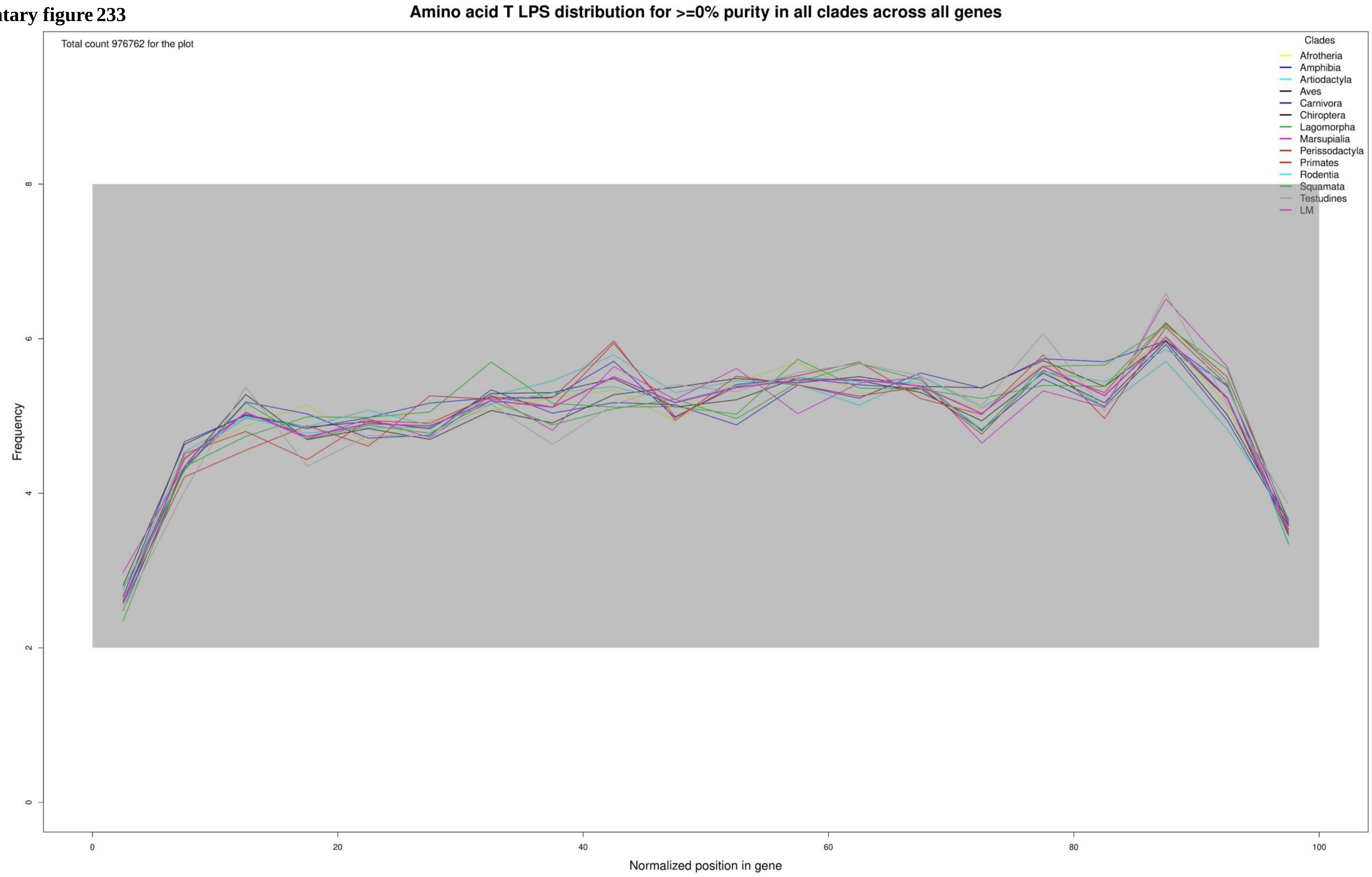

**Supplementary figure 234**

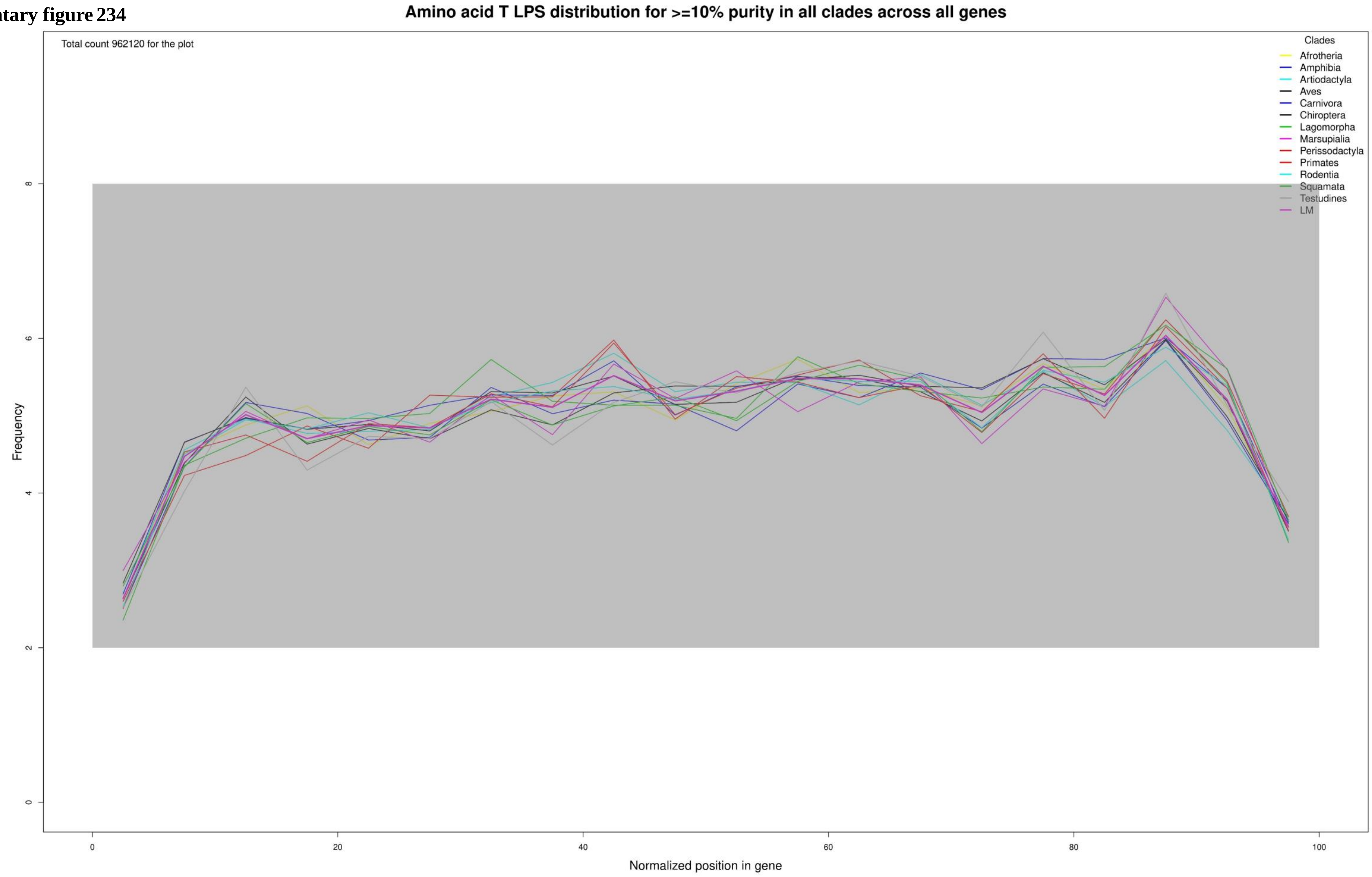

**Supplementary figure 235**

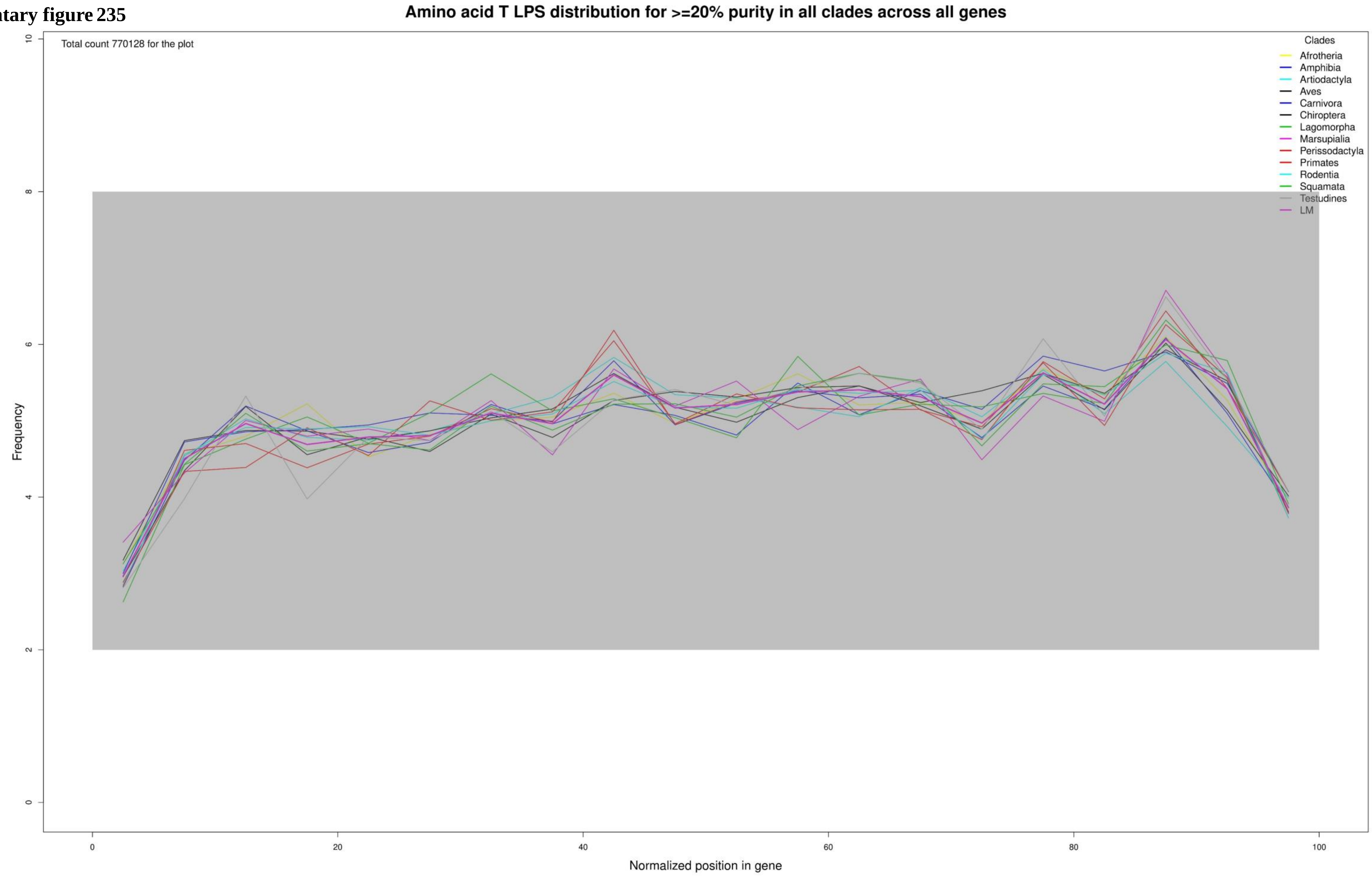

**Supplementary figure 236**

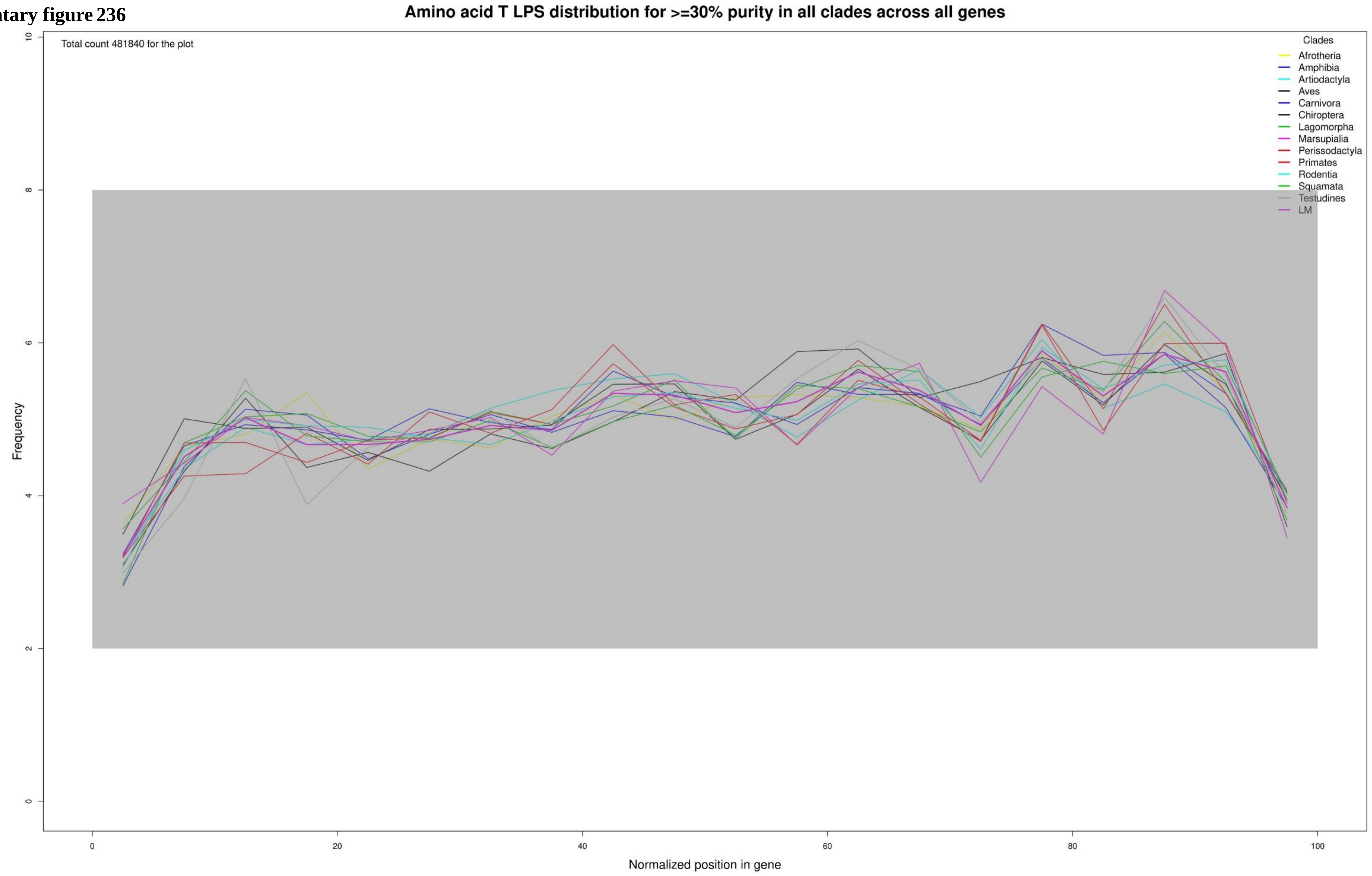

**Supplementary figure 237**

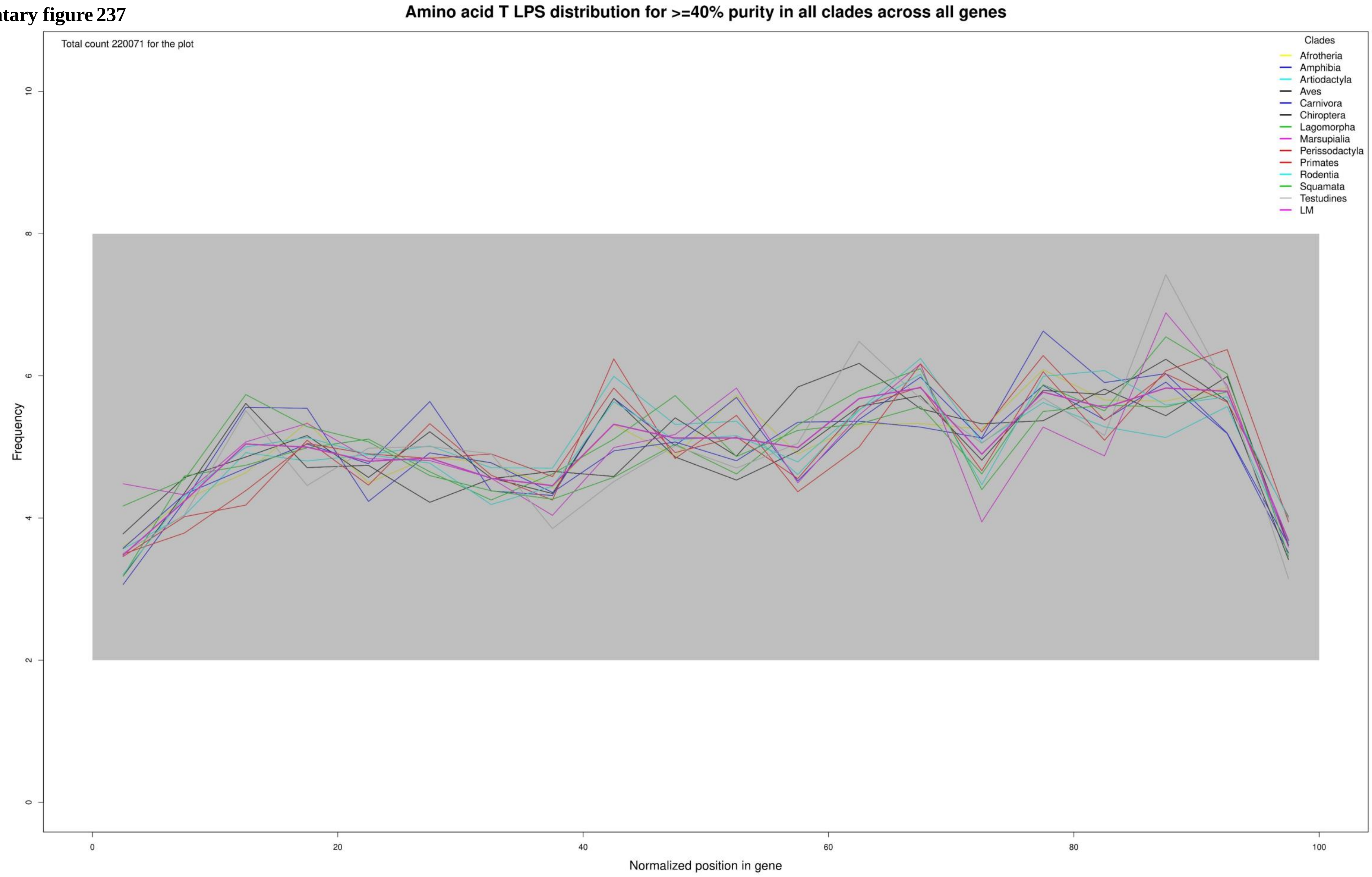

**Supplementary figure 238**

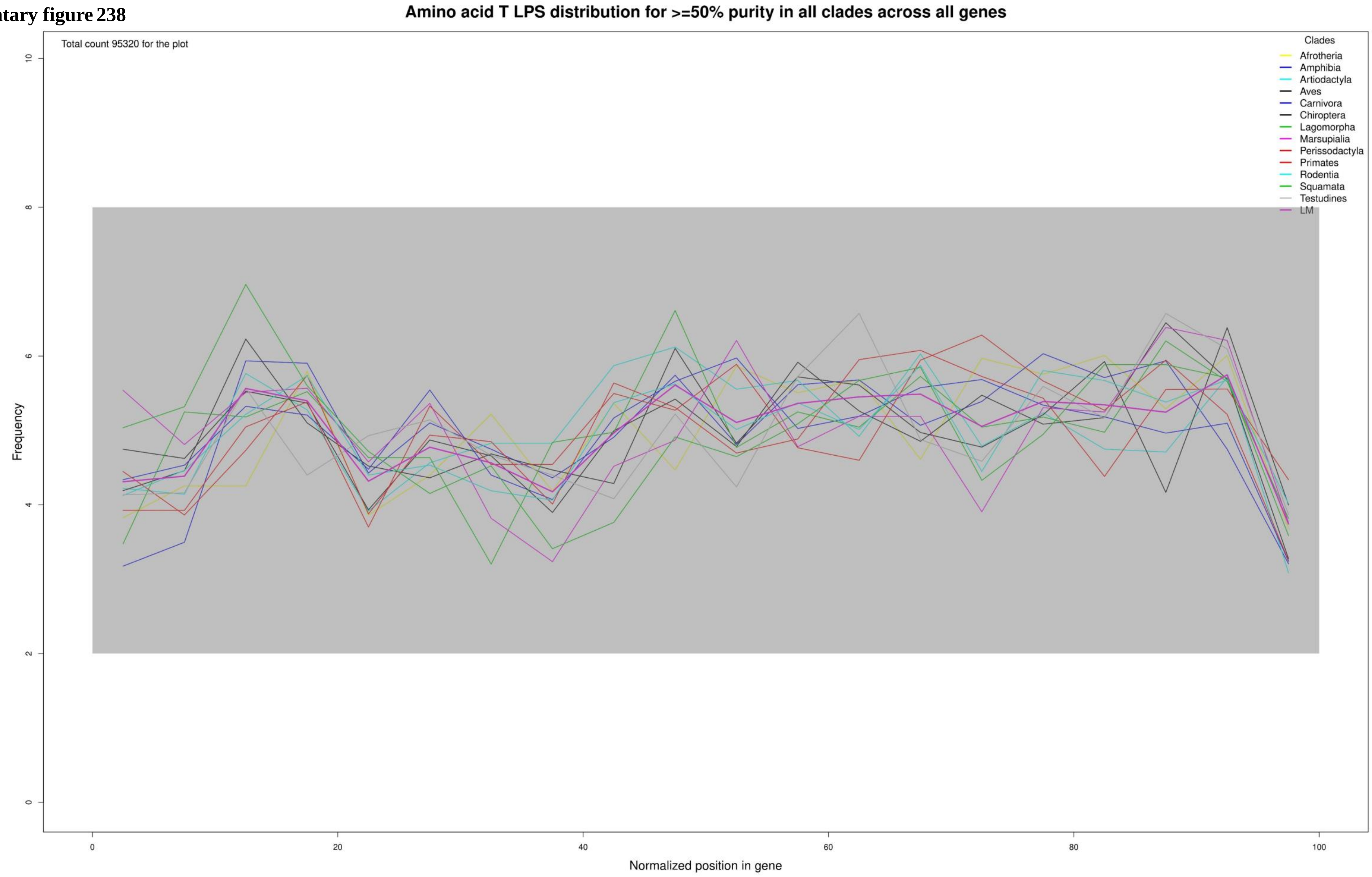

**Supplementary figure 239**

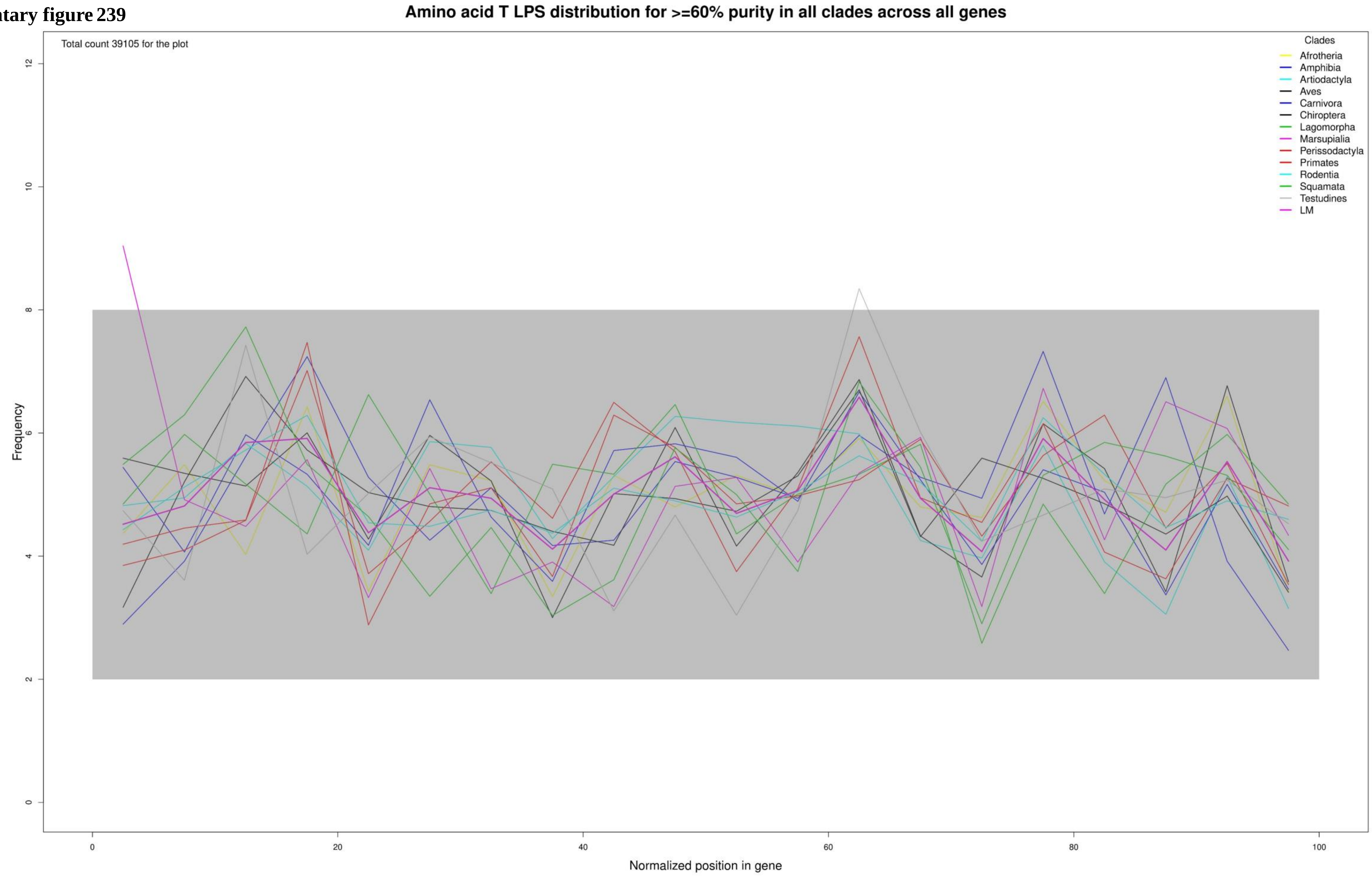

**Supplementary figure 240**

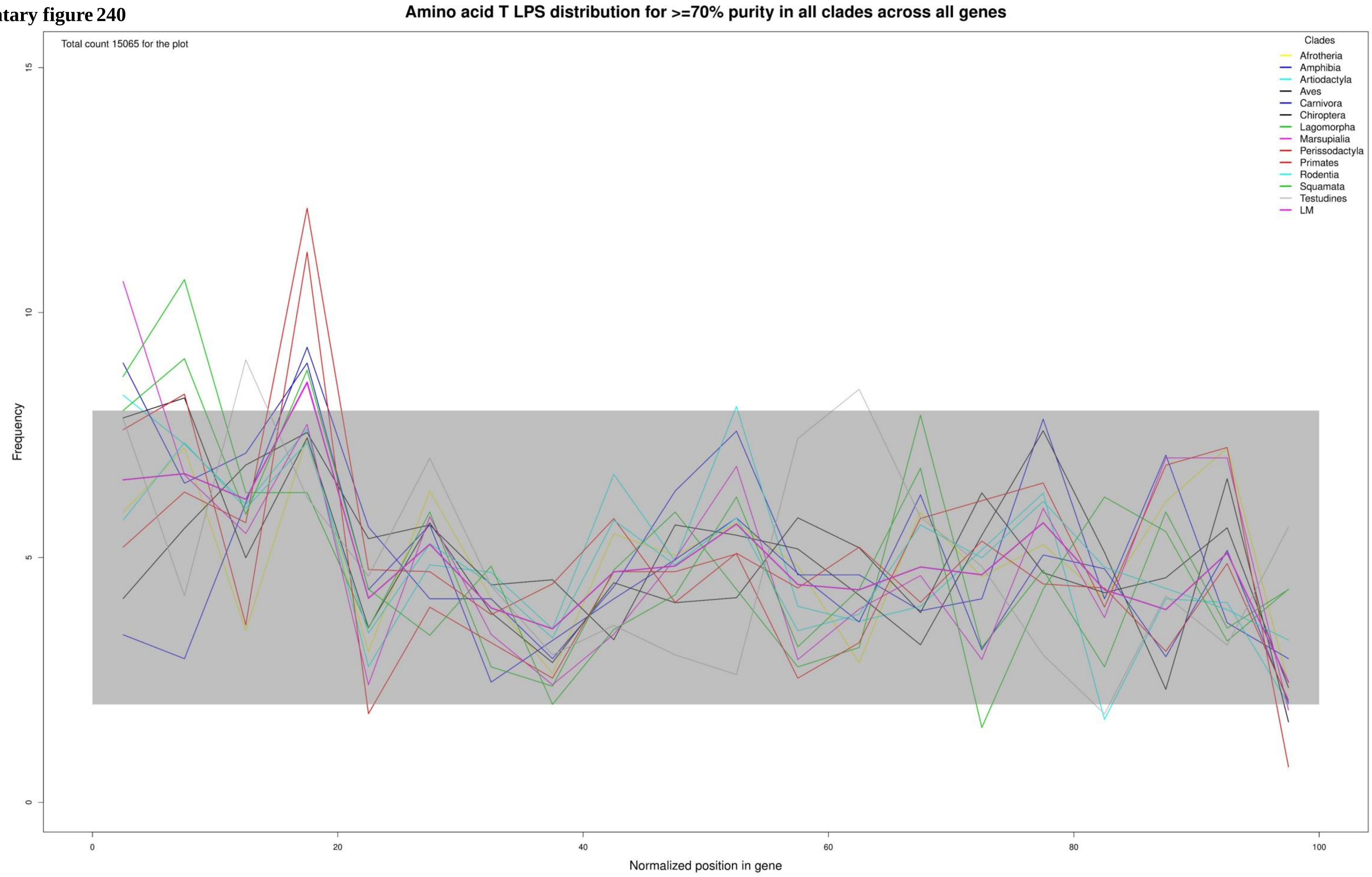

**Supplementary figure 241**

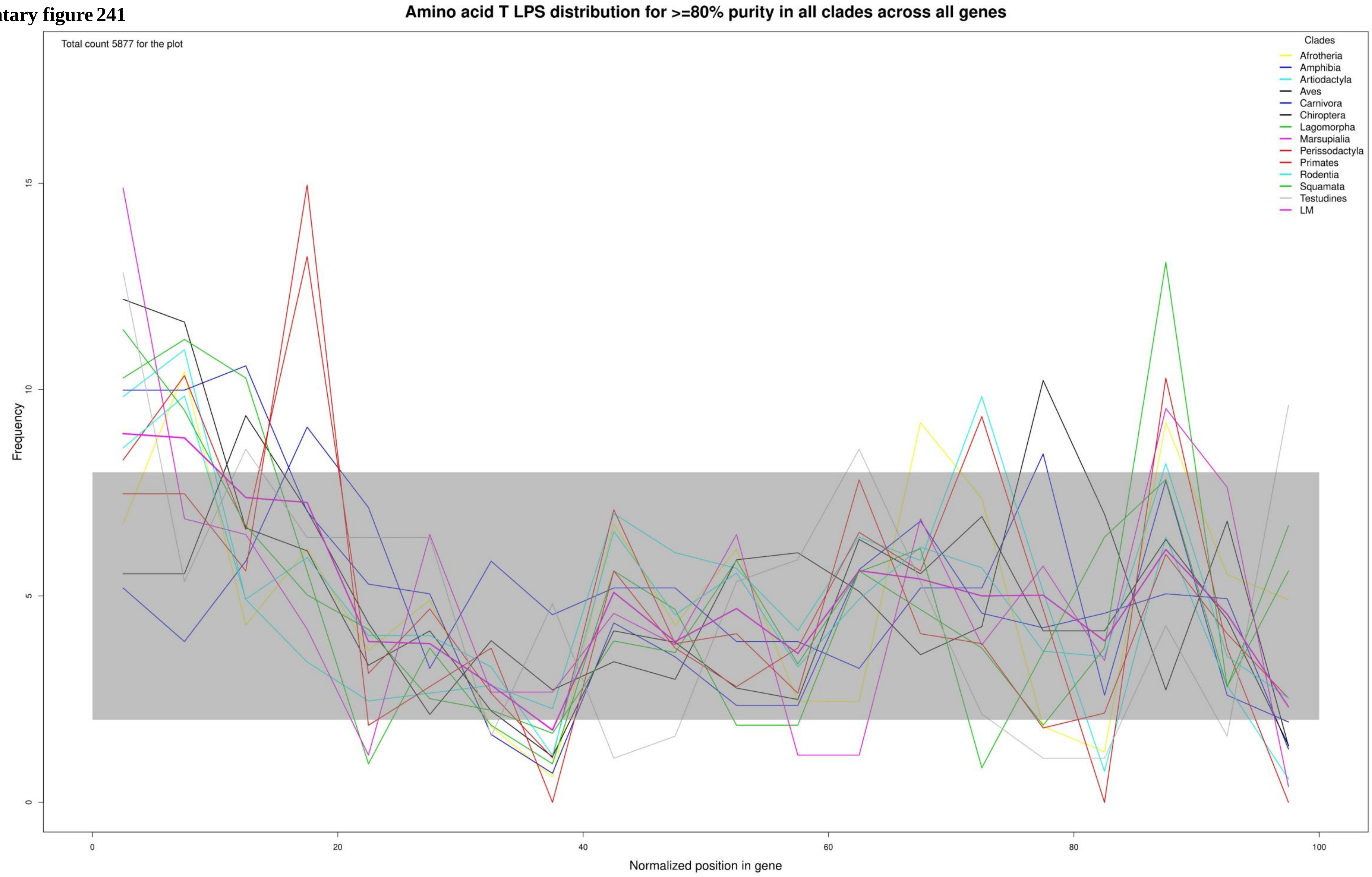

**Supplementary figure 242**

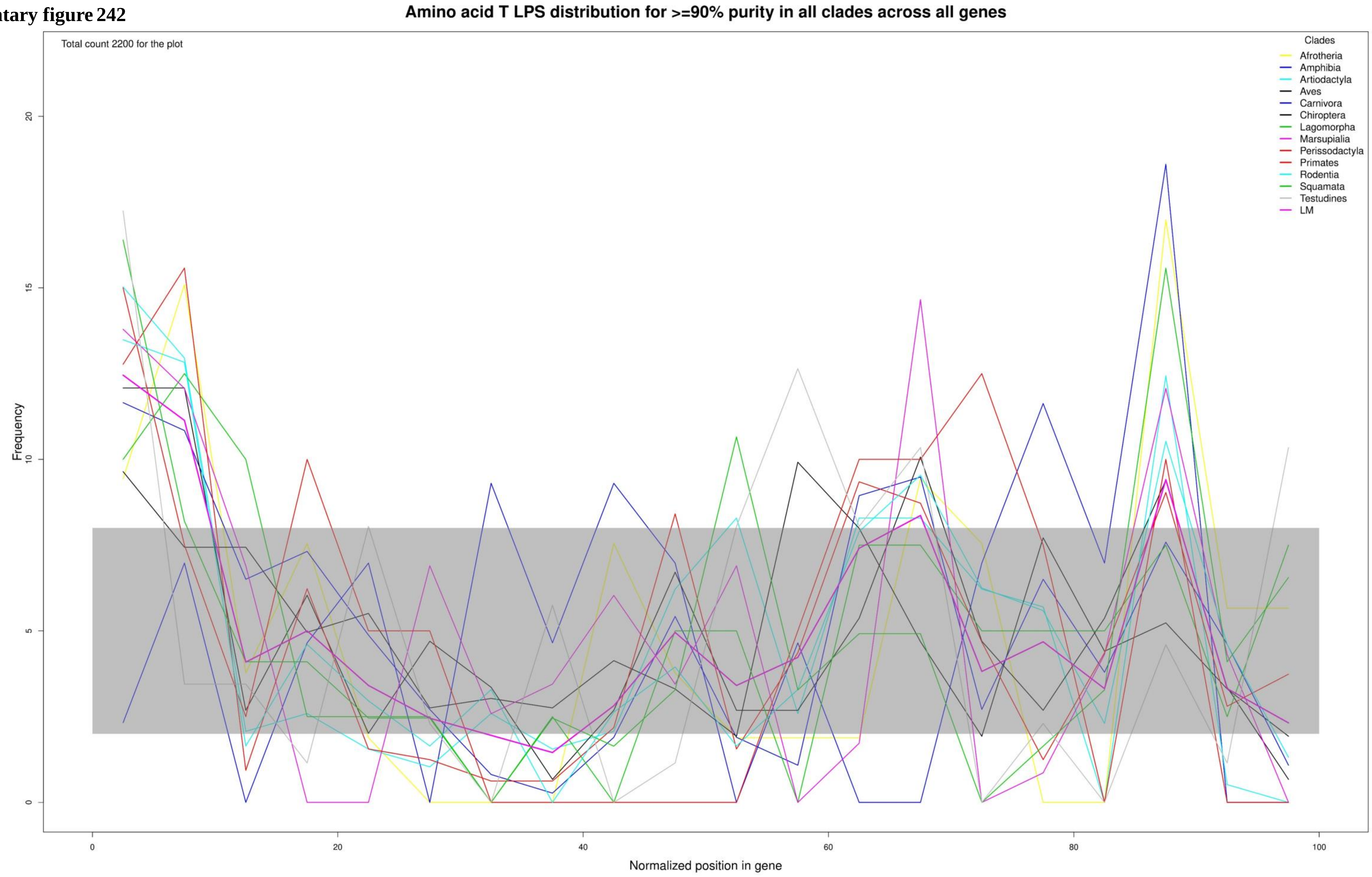

**Supplementary figure 243**

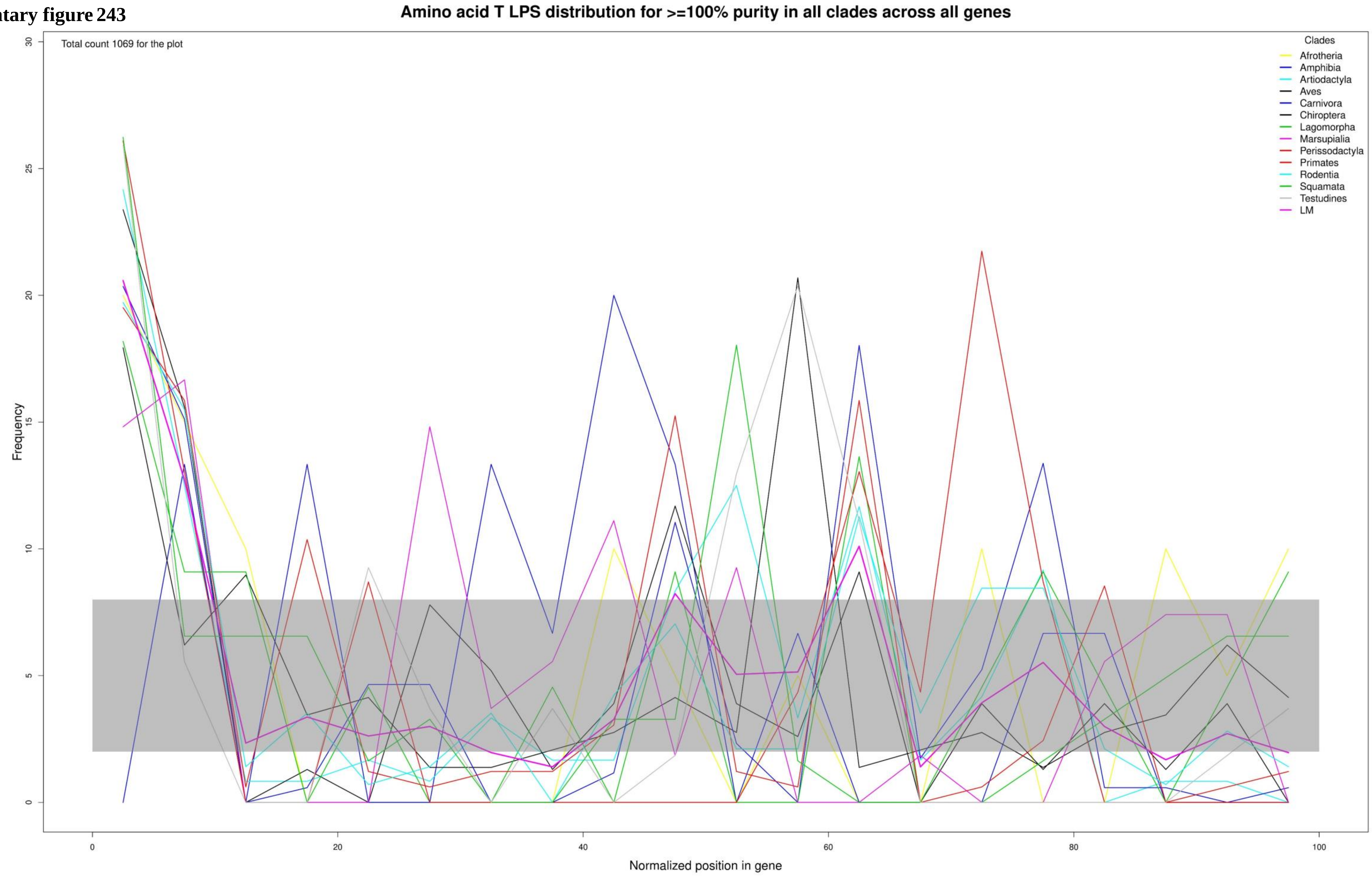

Supplementary figure 244

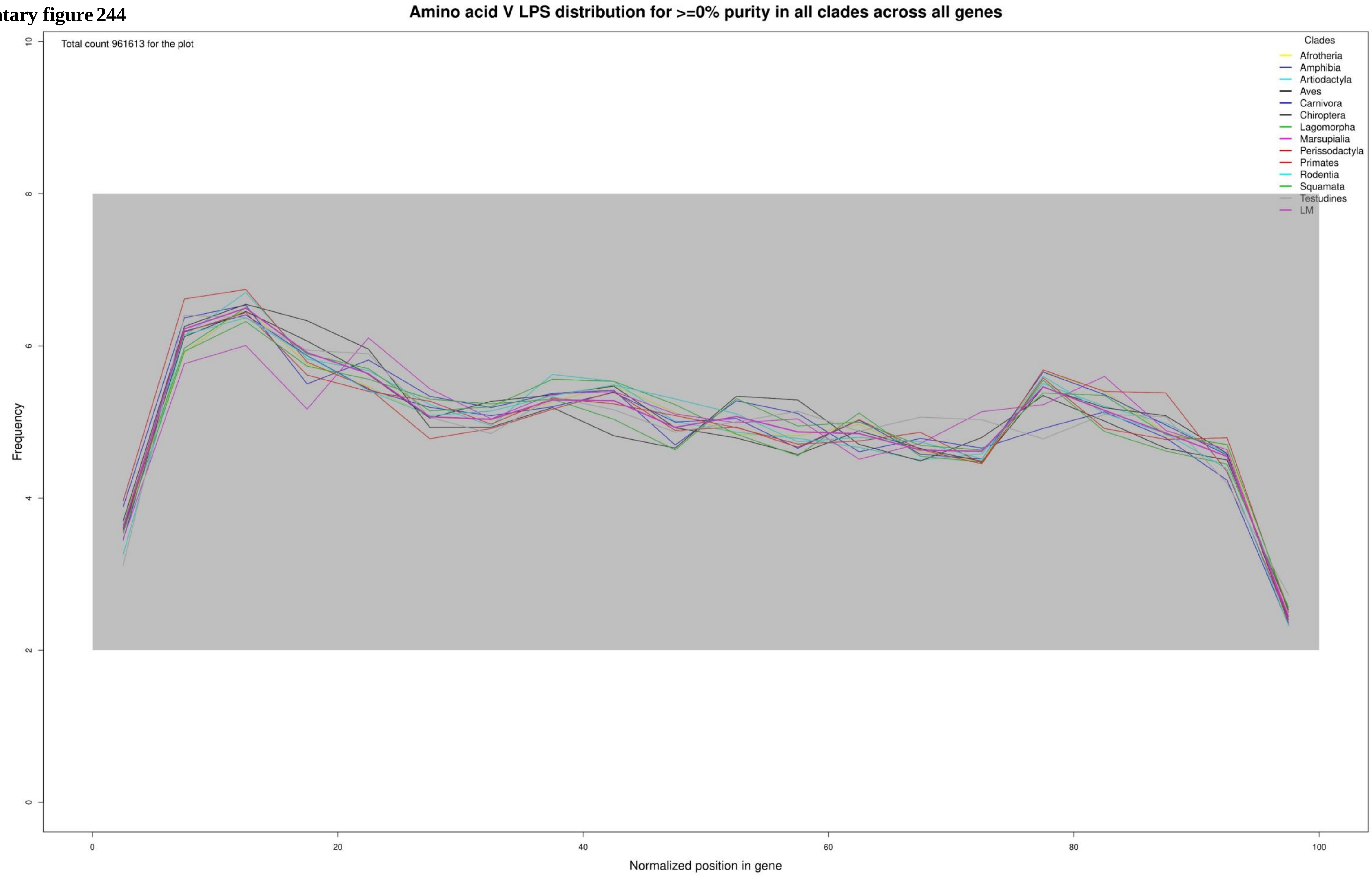

**Supplementary figure 245**

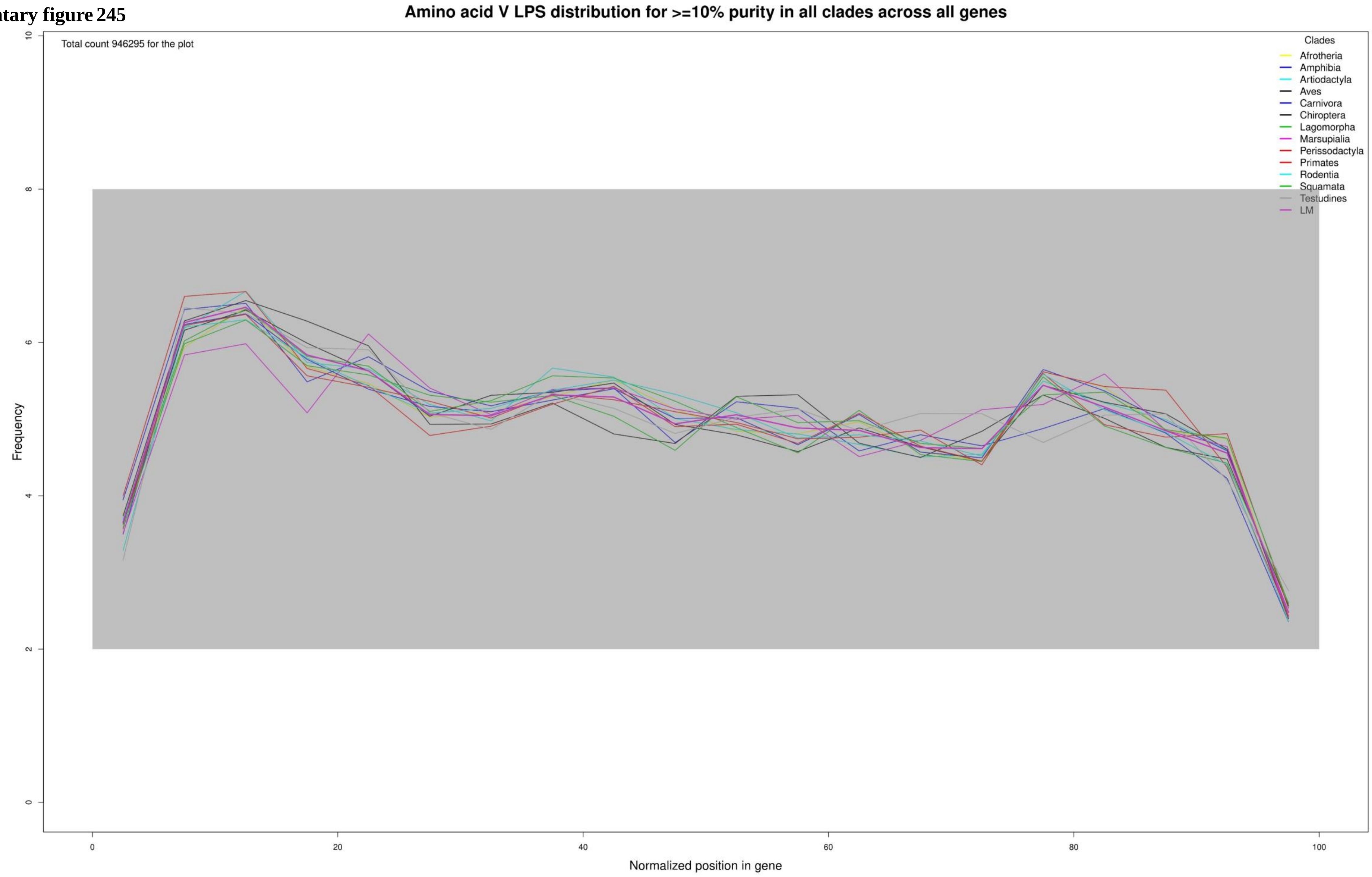

**Supplementary figure 246**

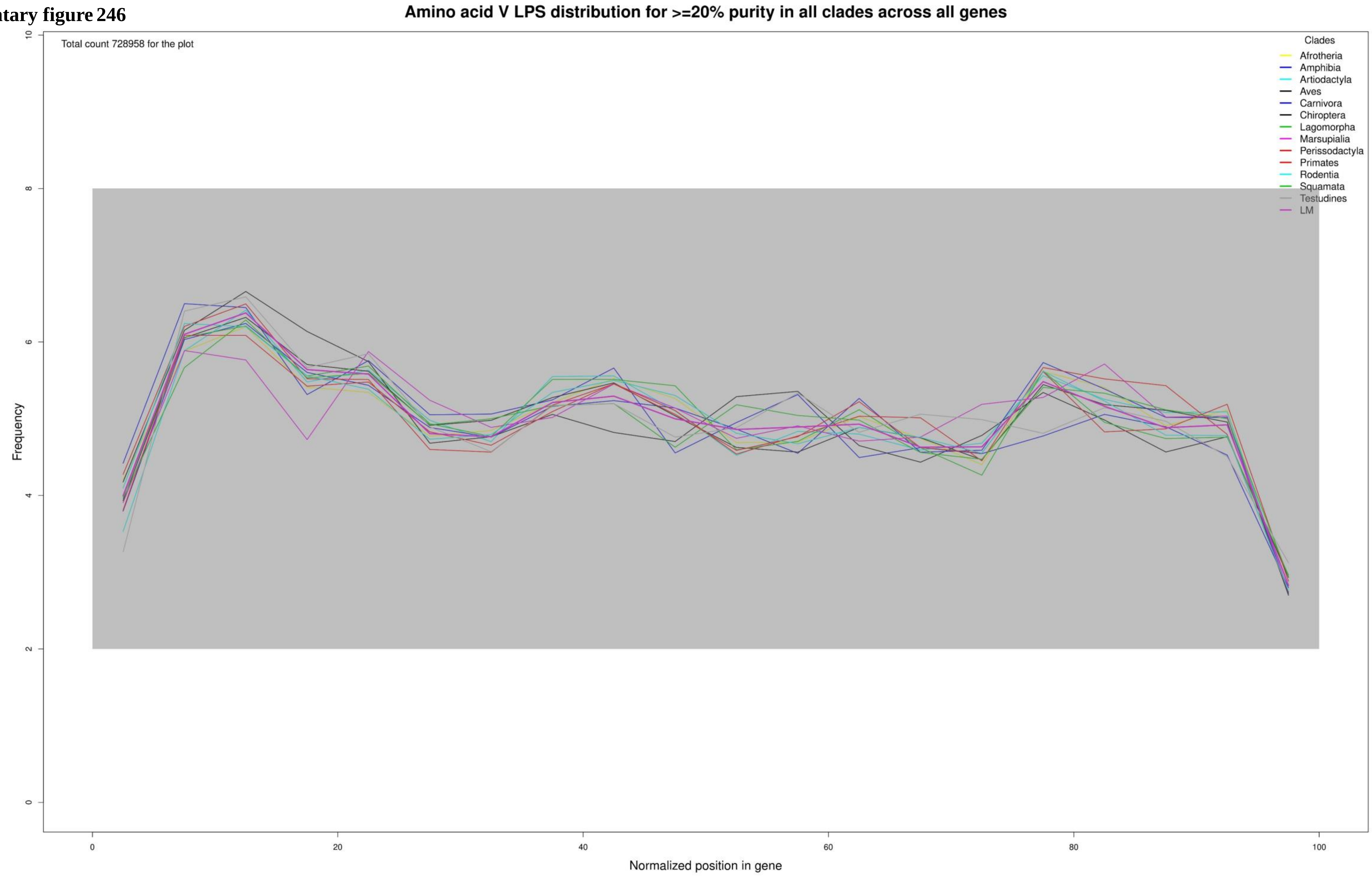

**Supplementary figure 247**

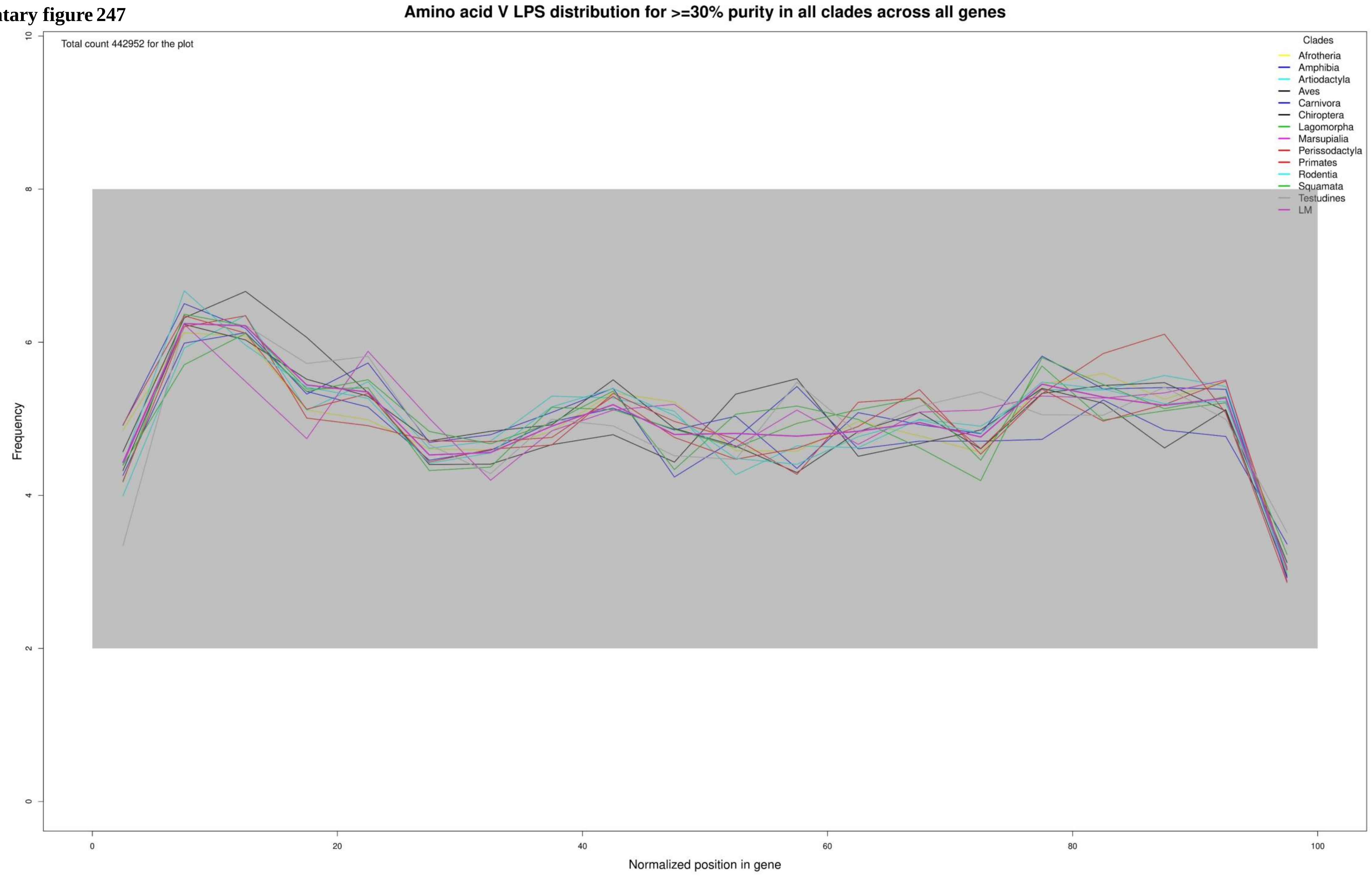

**Supplementary figure 248**

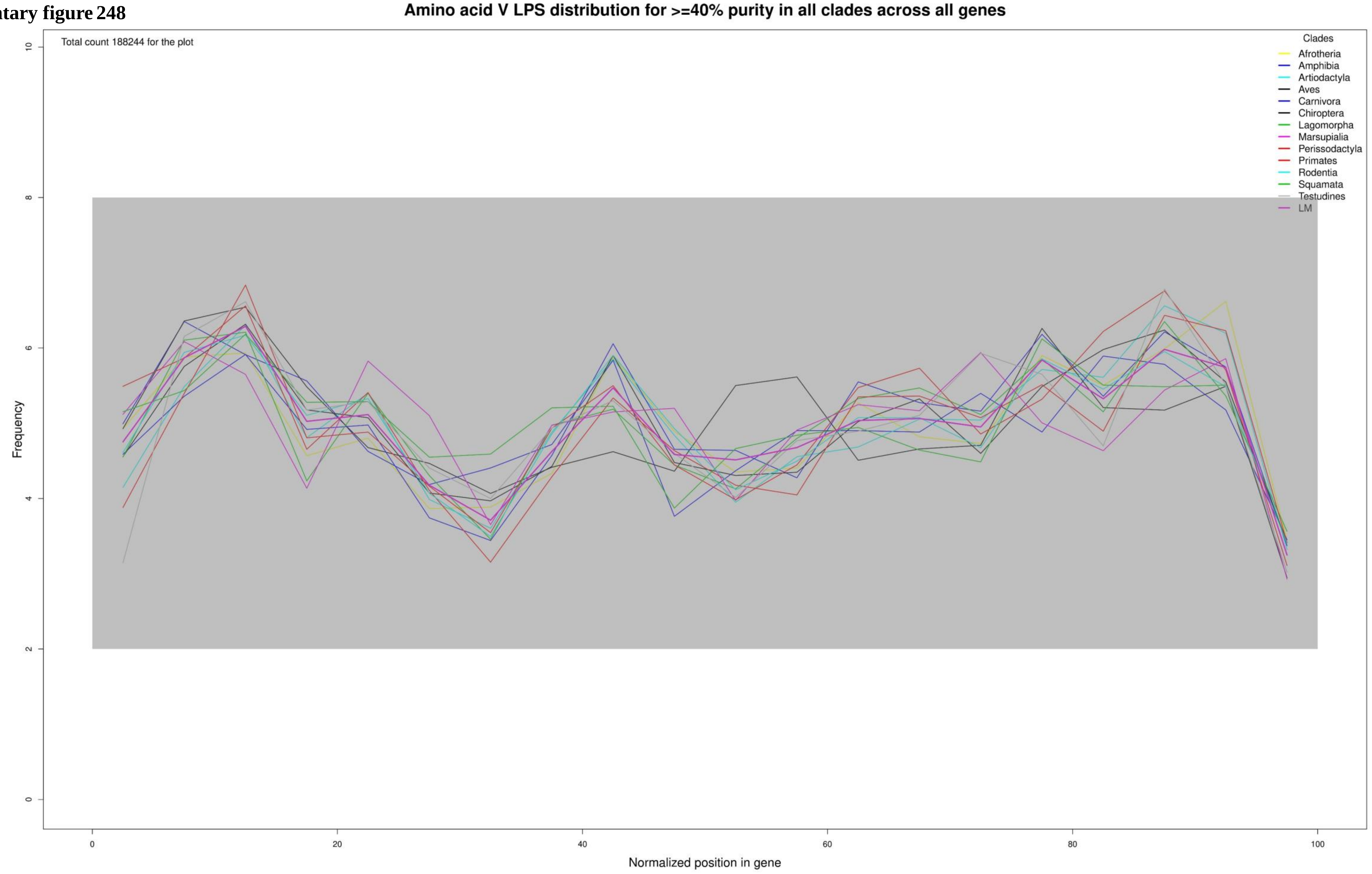

**Supplementary figure 249**

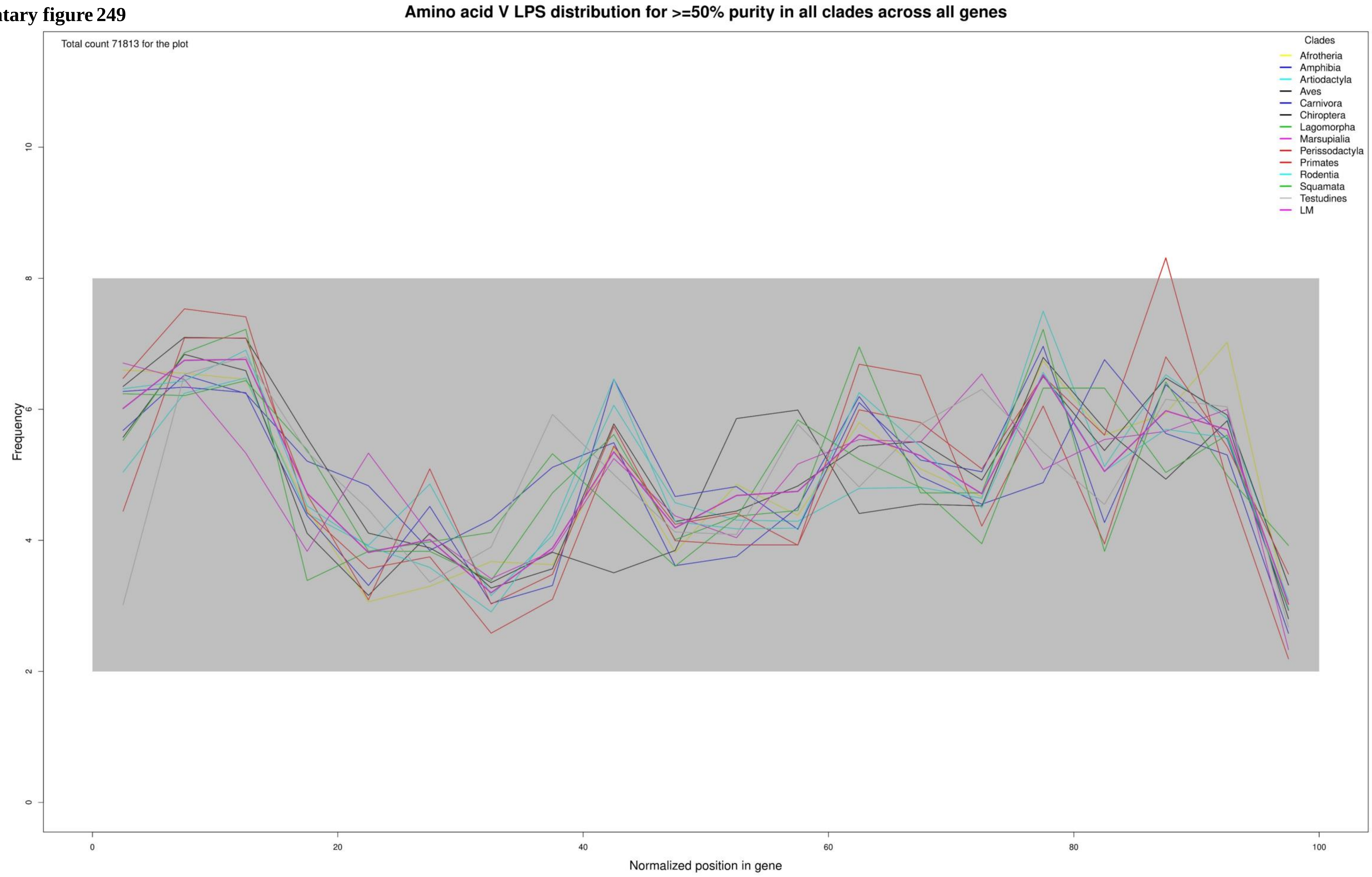

**Supplementary figure 250**

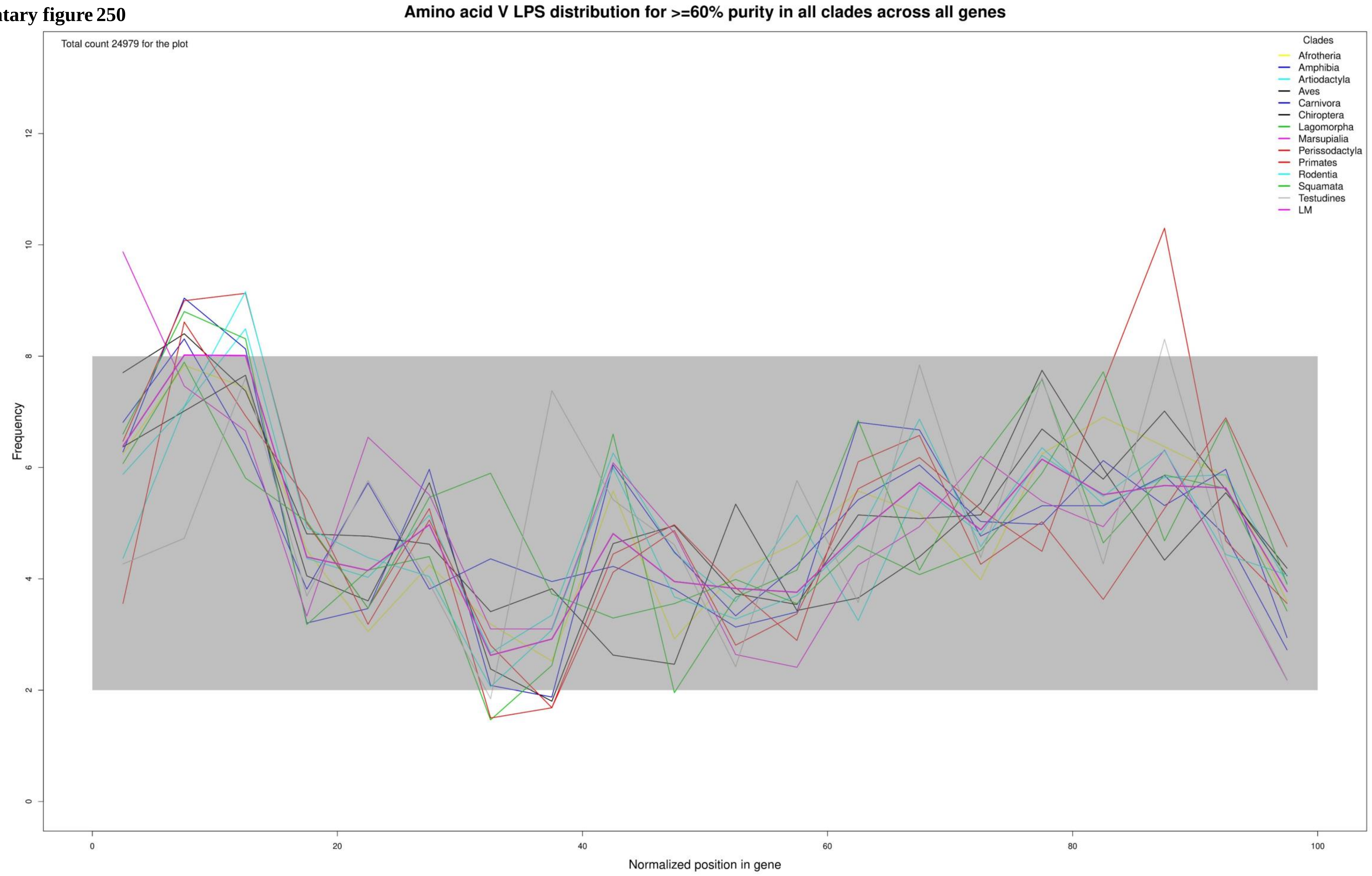

**Supplementary figure 251**

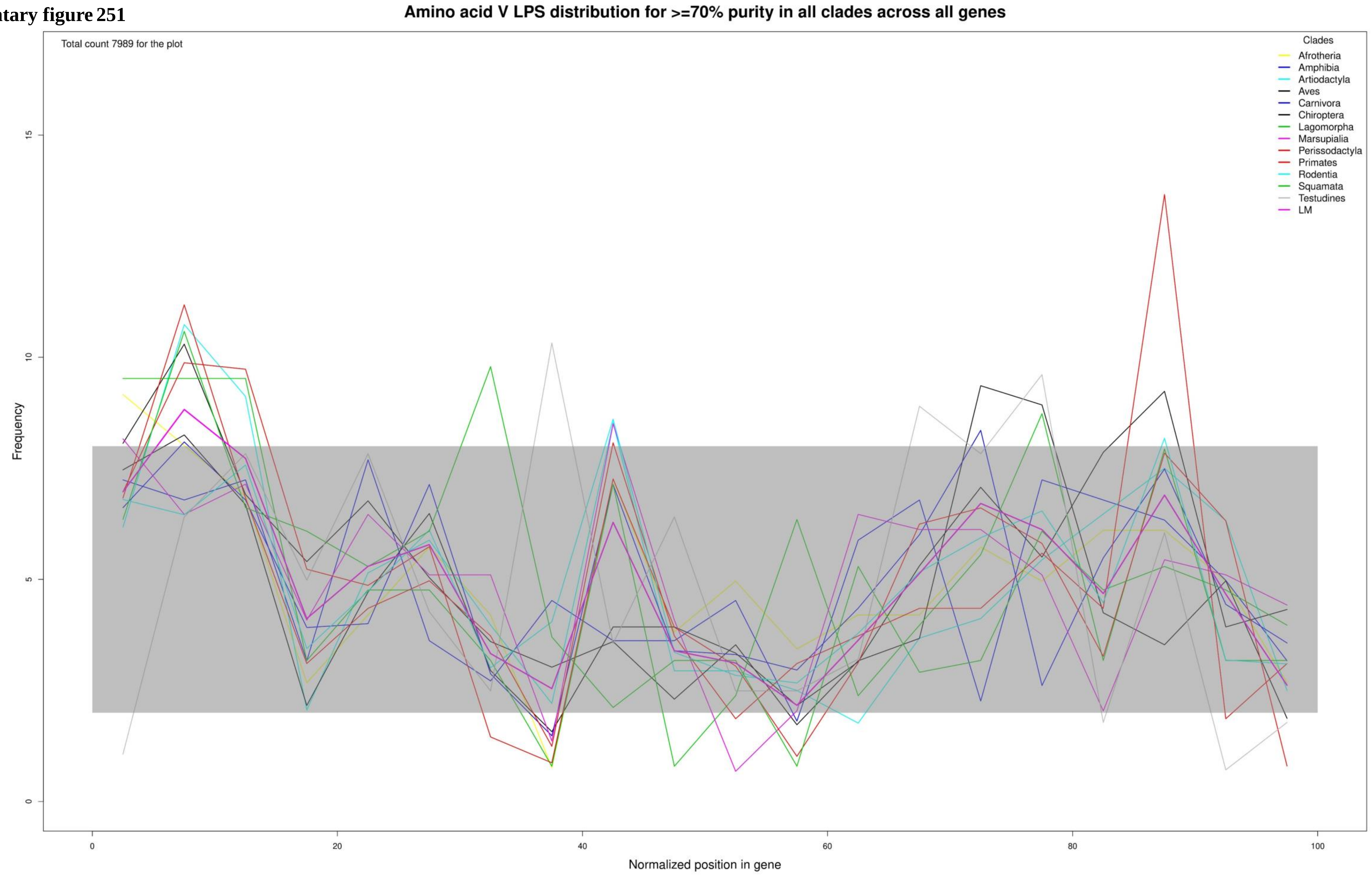

Supplementary figure 252

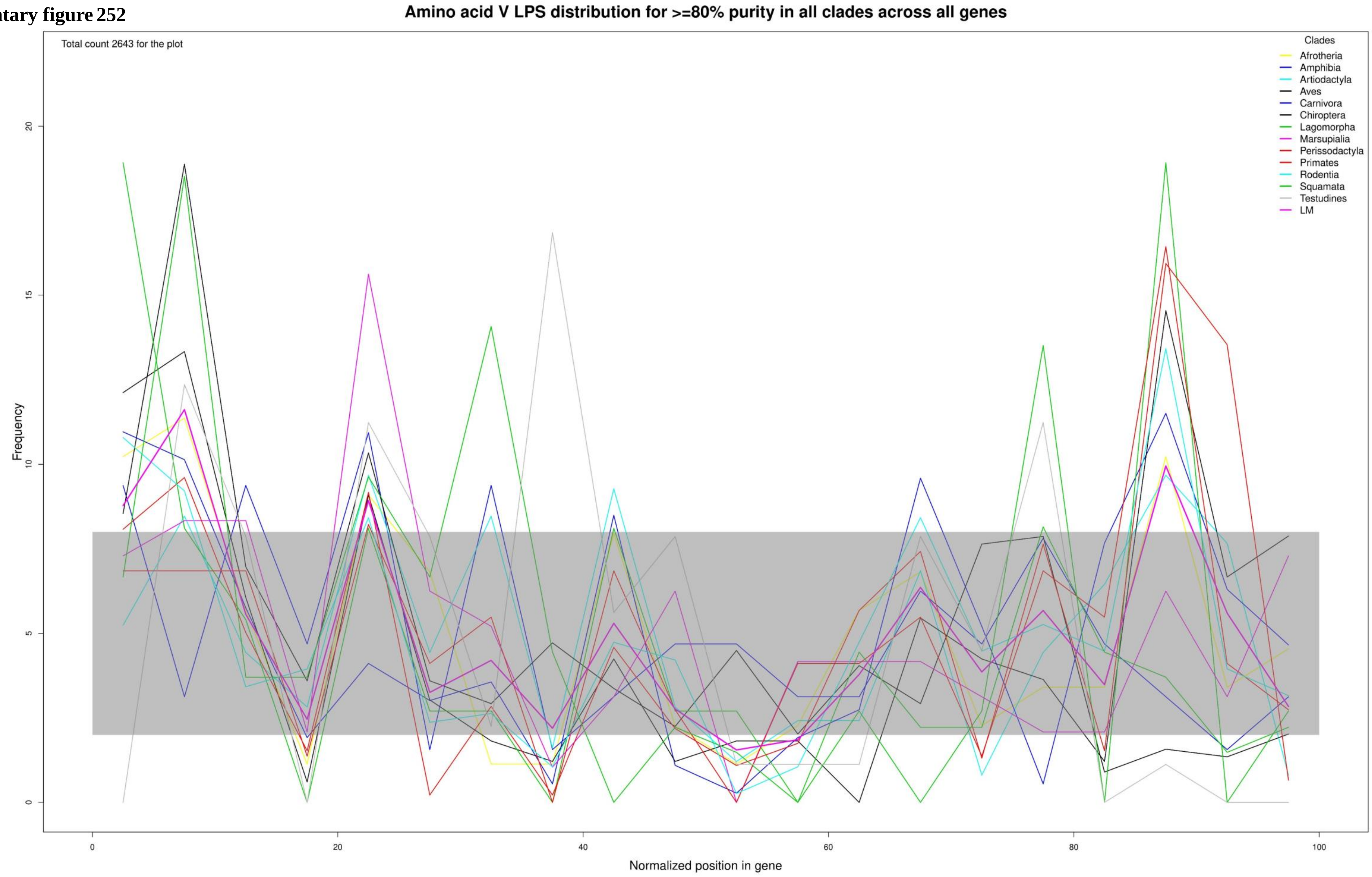

**Supplementary figure 253**

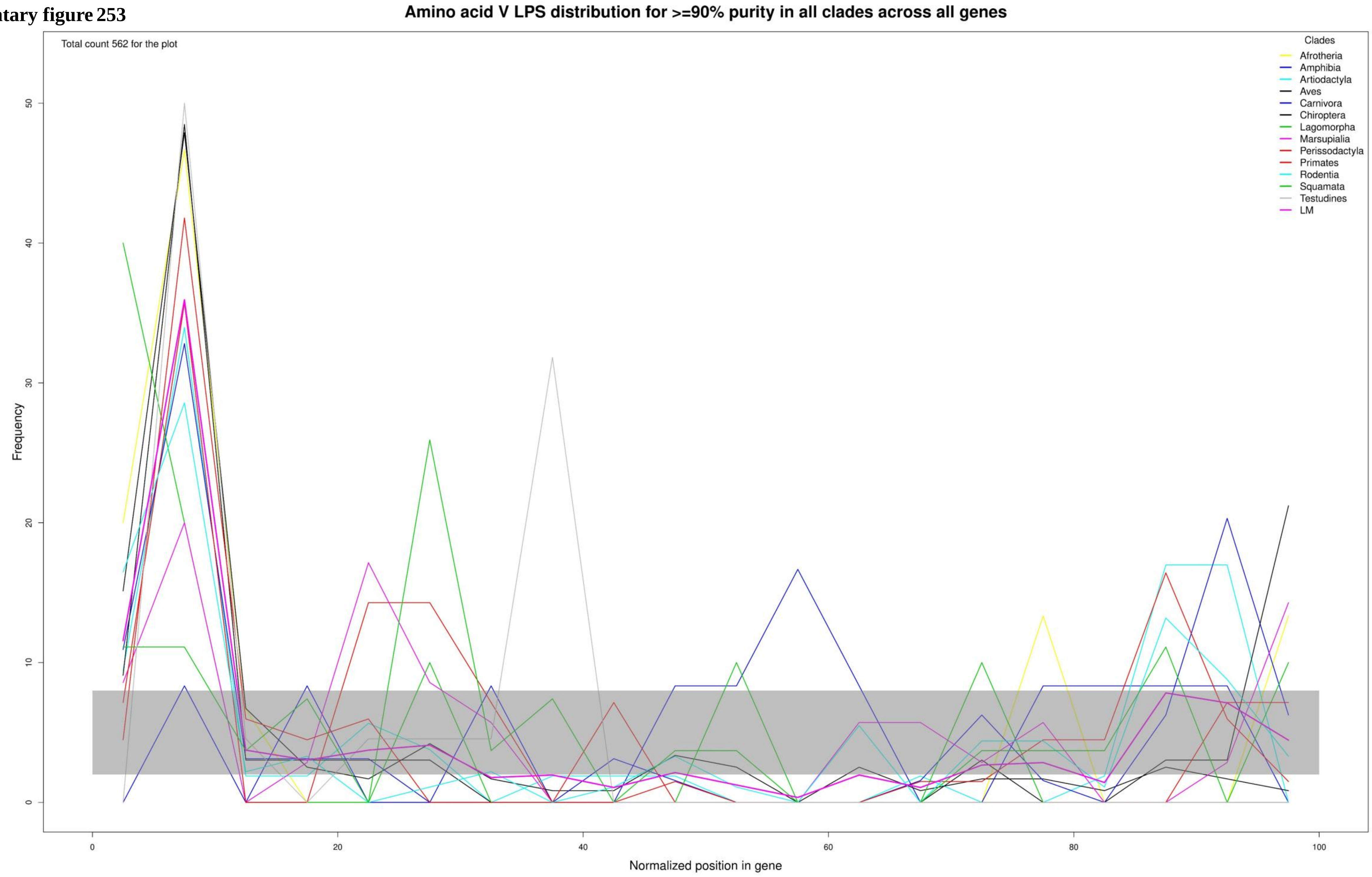

Supplementary figure 254

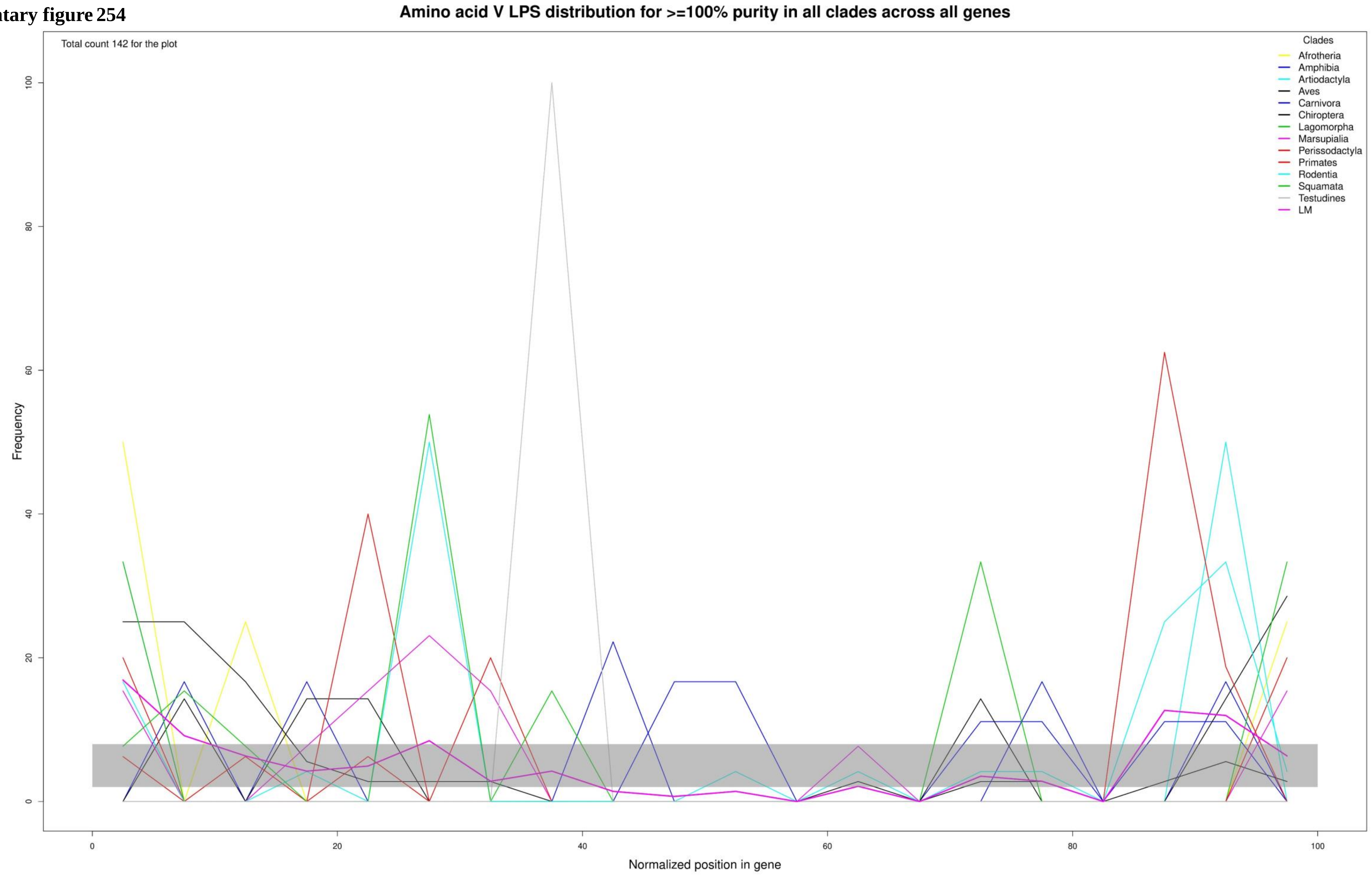

**Supplementary figure 255**

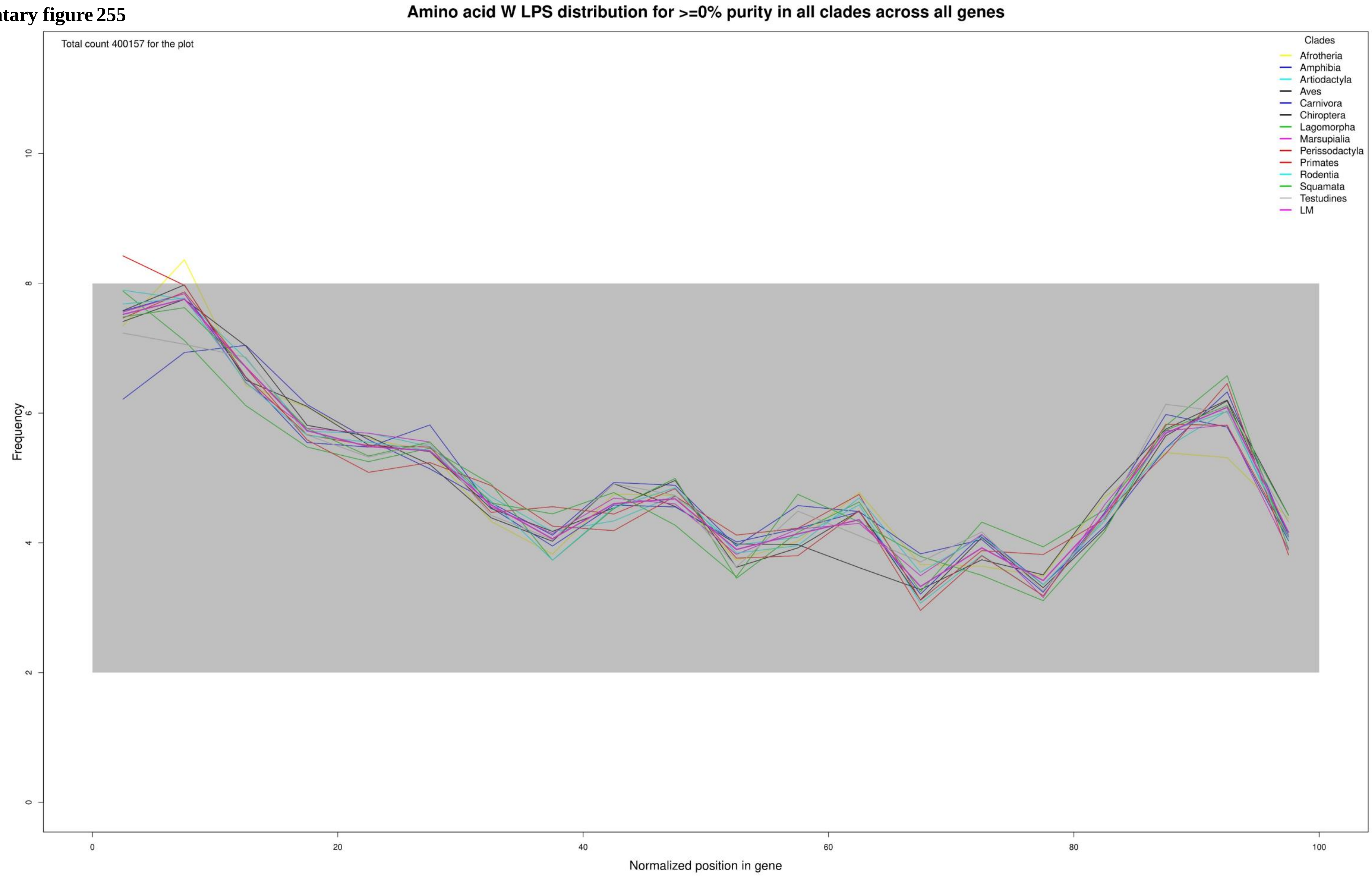

**Supplementary figure 256**

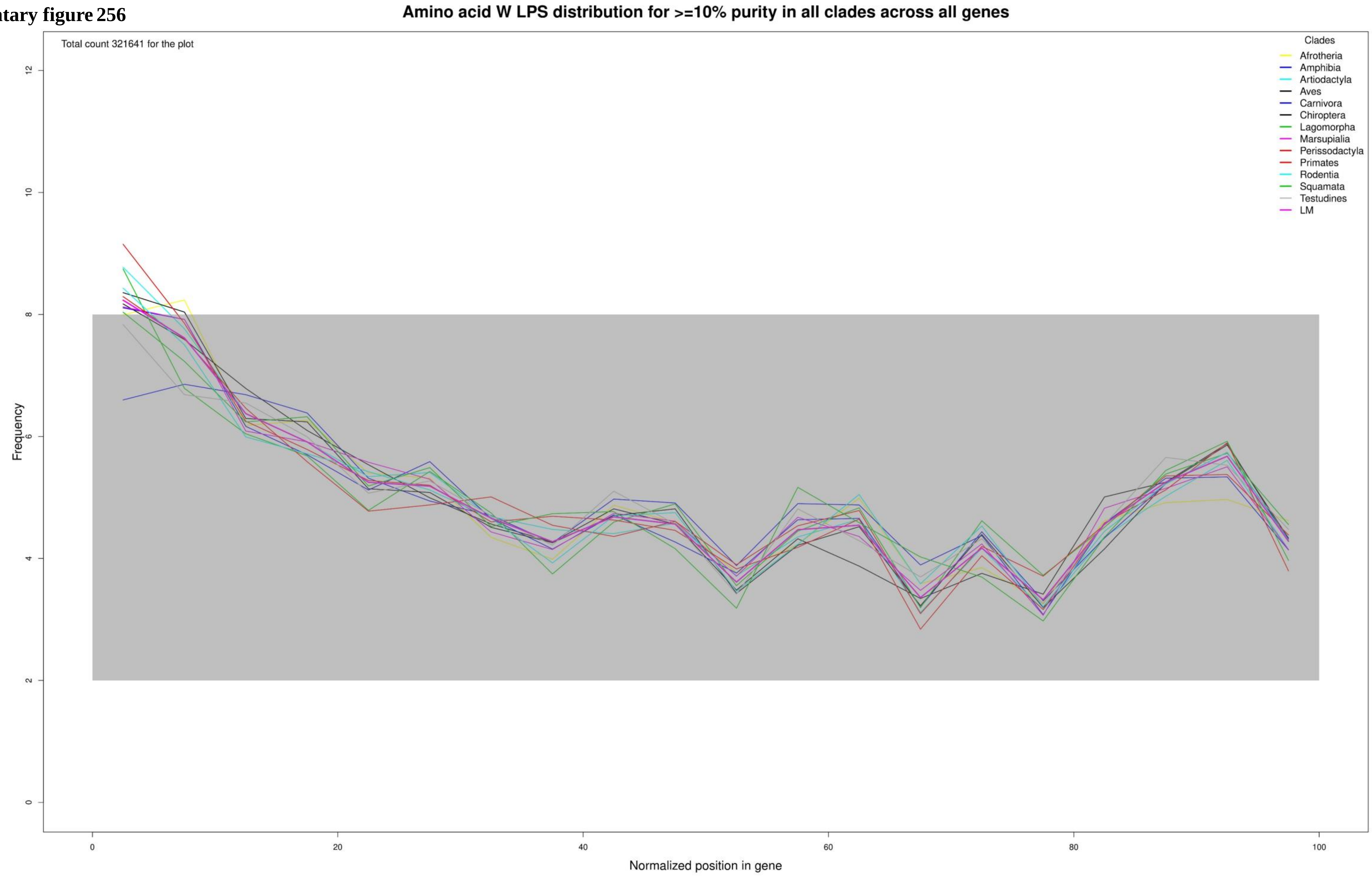

**Supplementary figure 257**

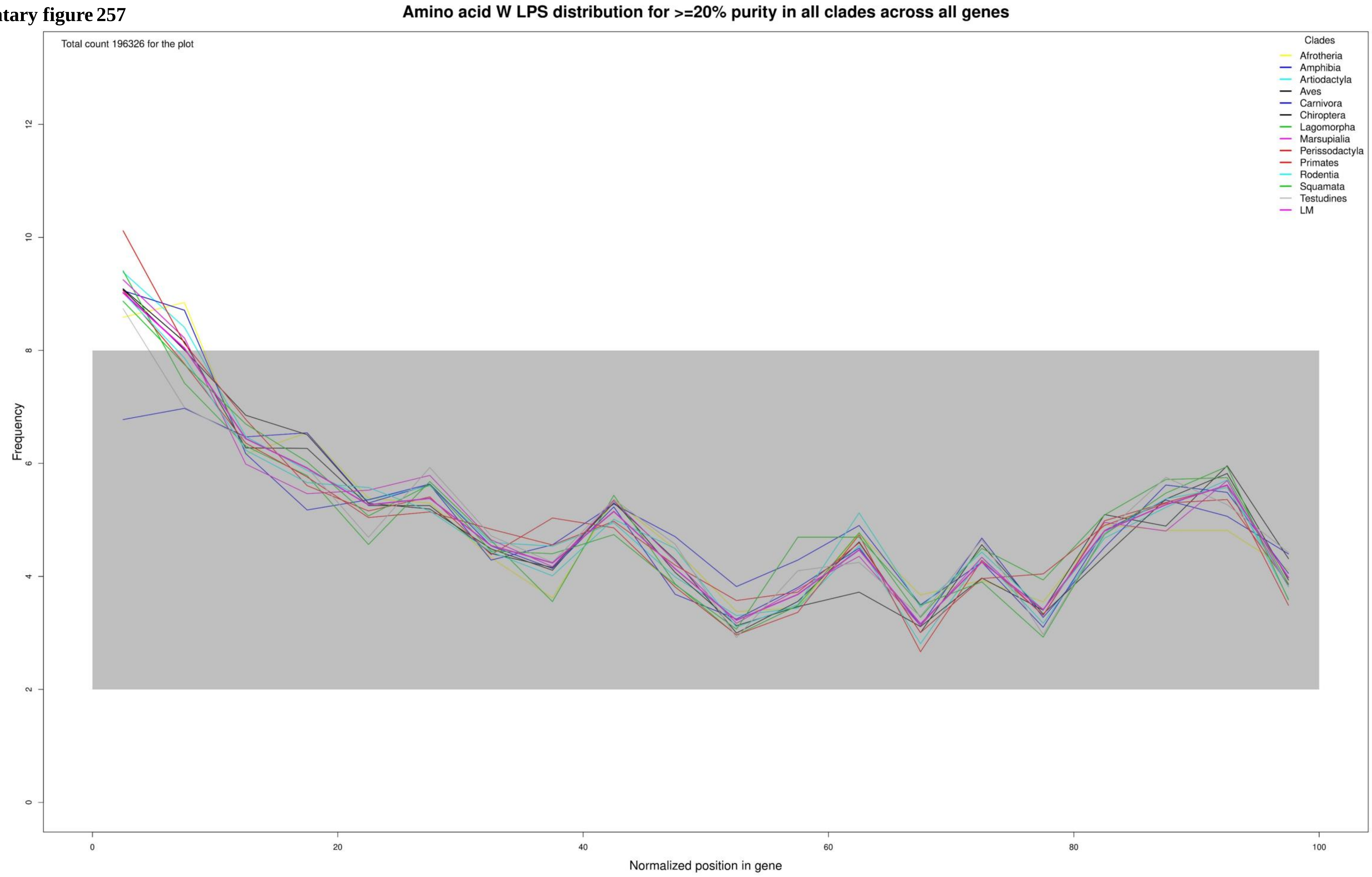

**Supplementary figure 258**

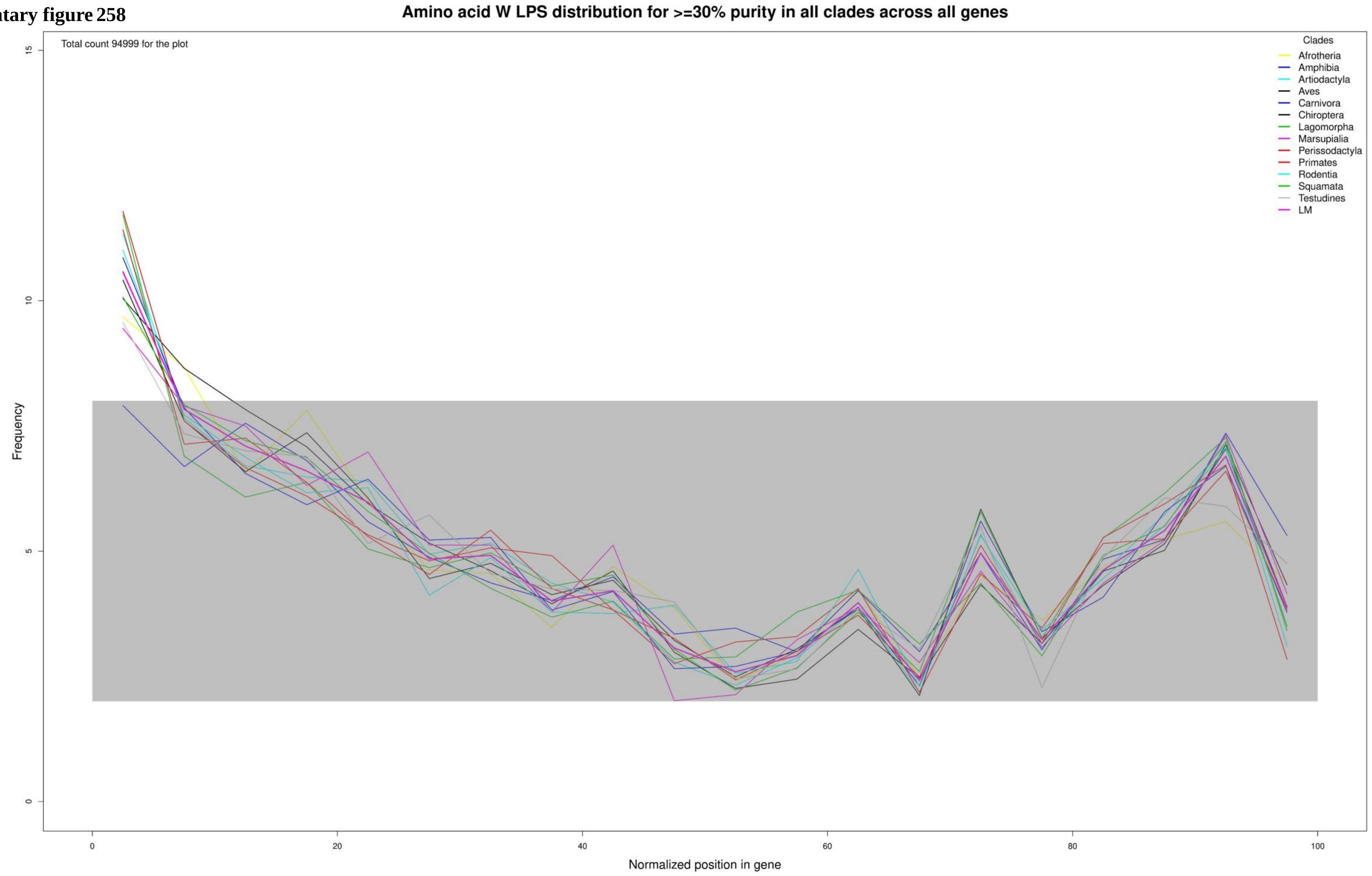

**Supplementary figure 259**

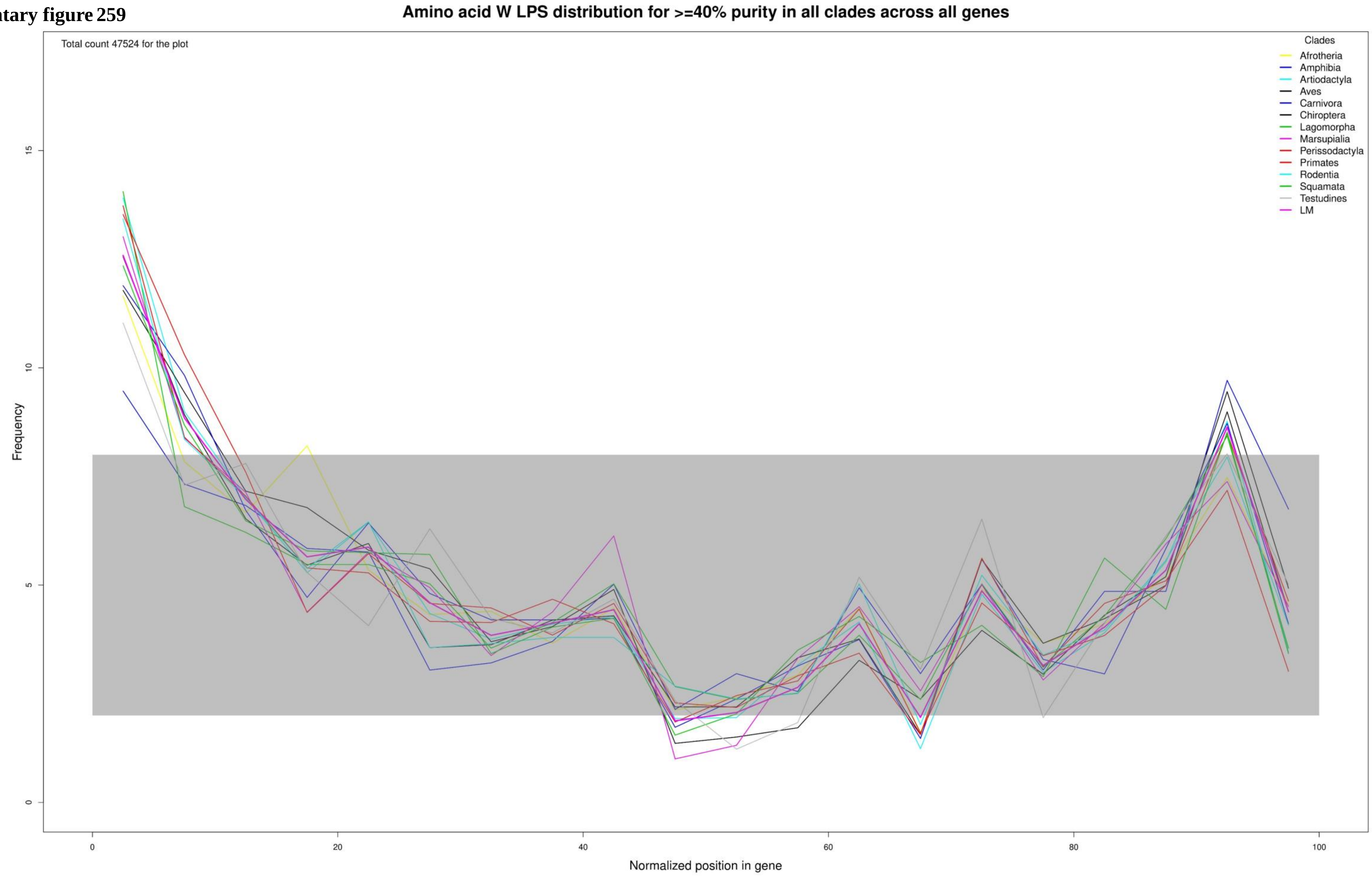

**Supplementary figure 260**

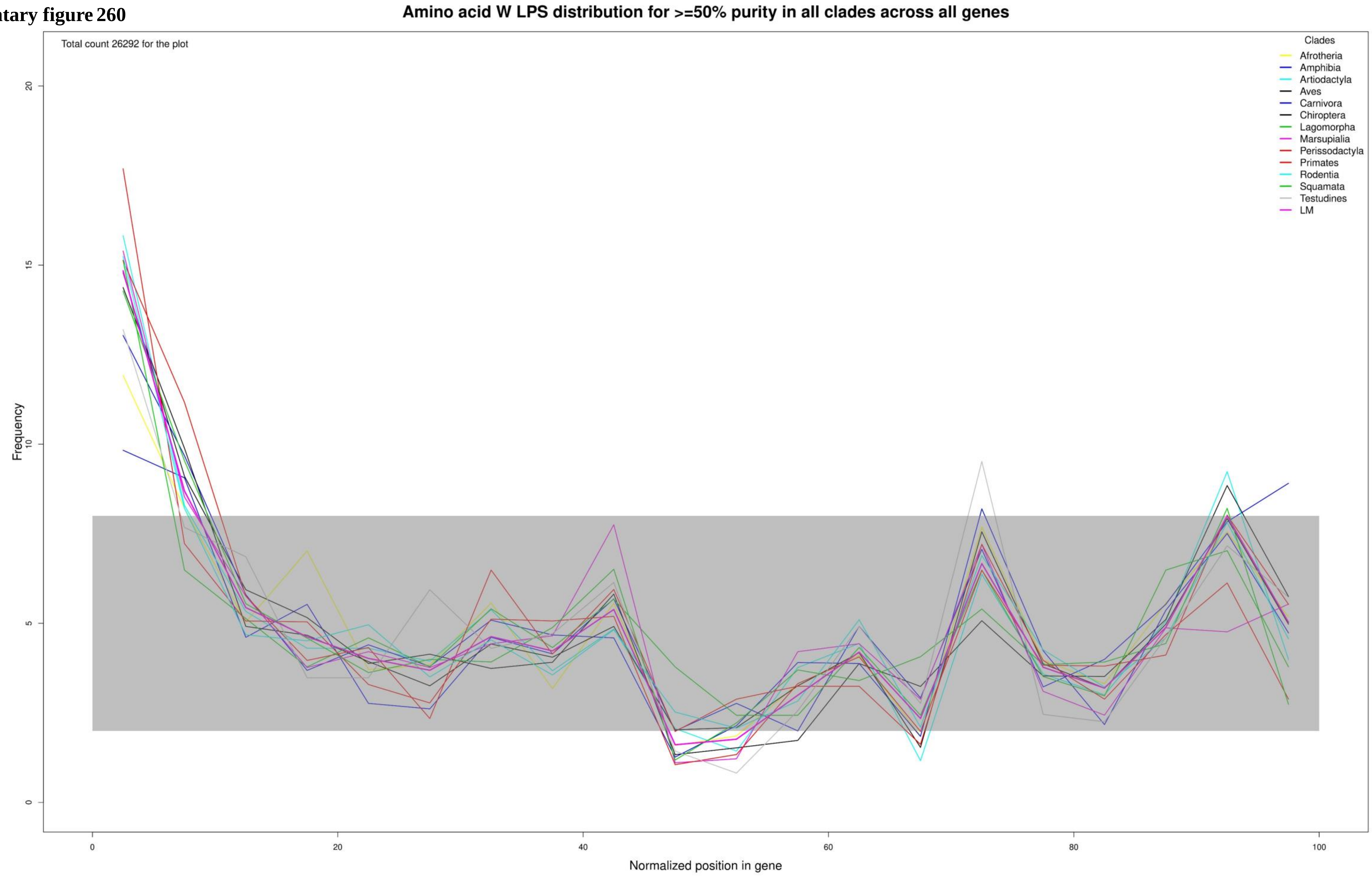

Supplementary figure 261

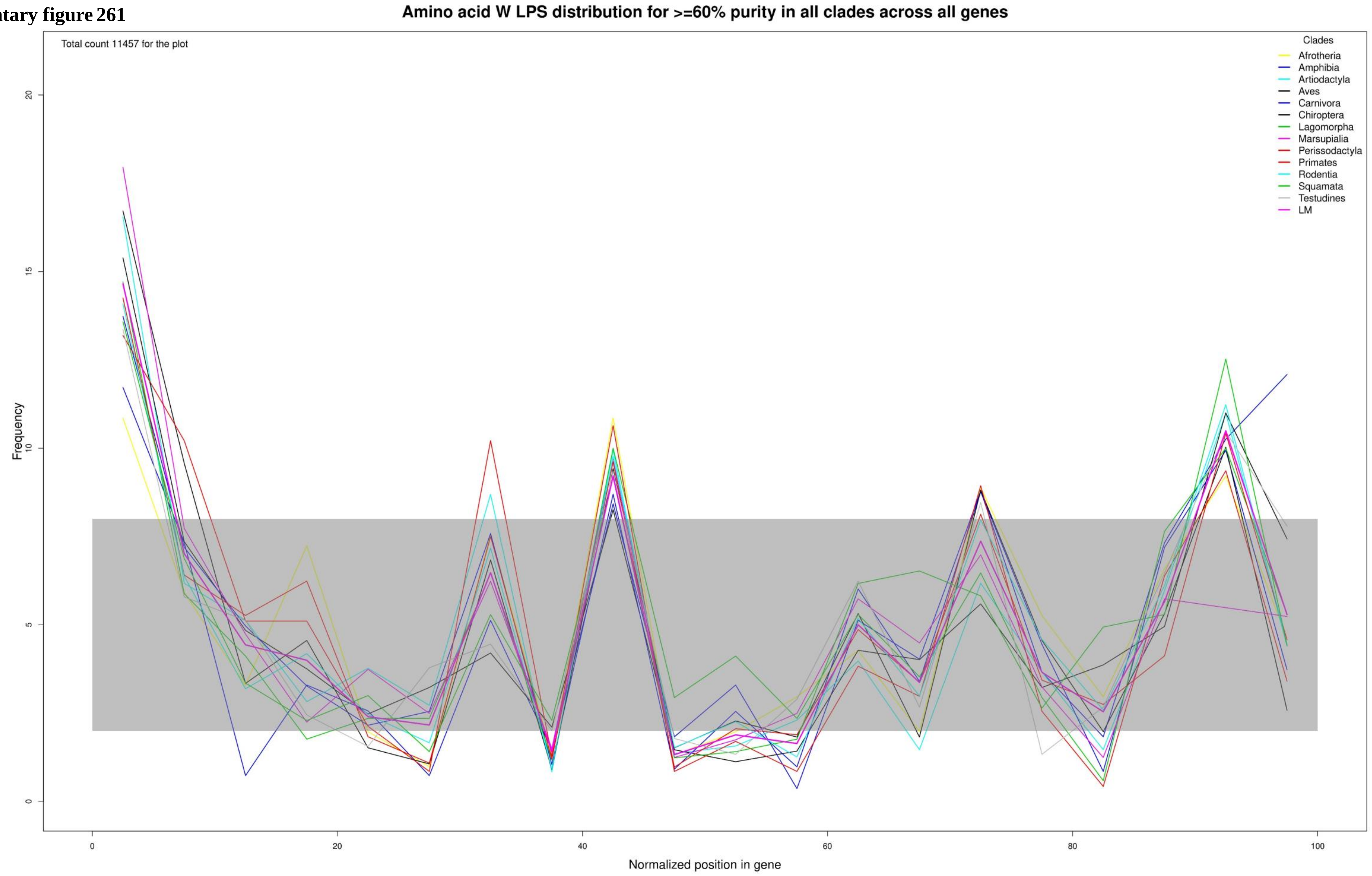

Supplementary figure 262

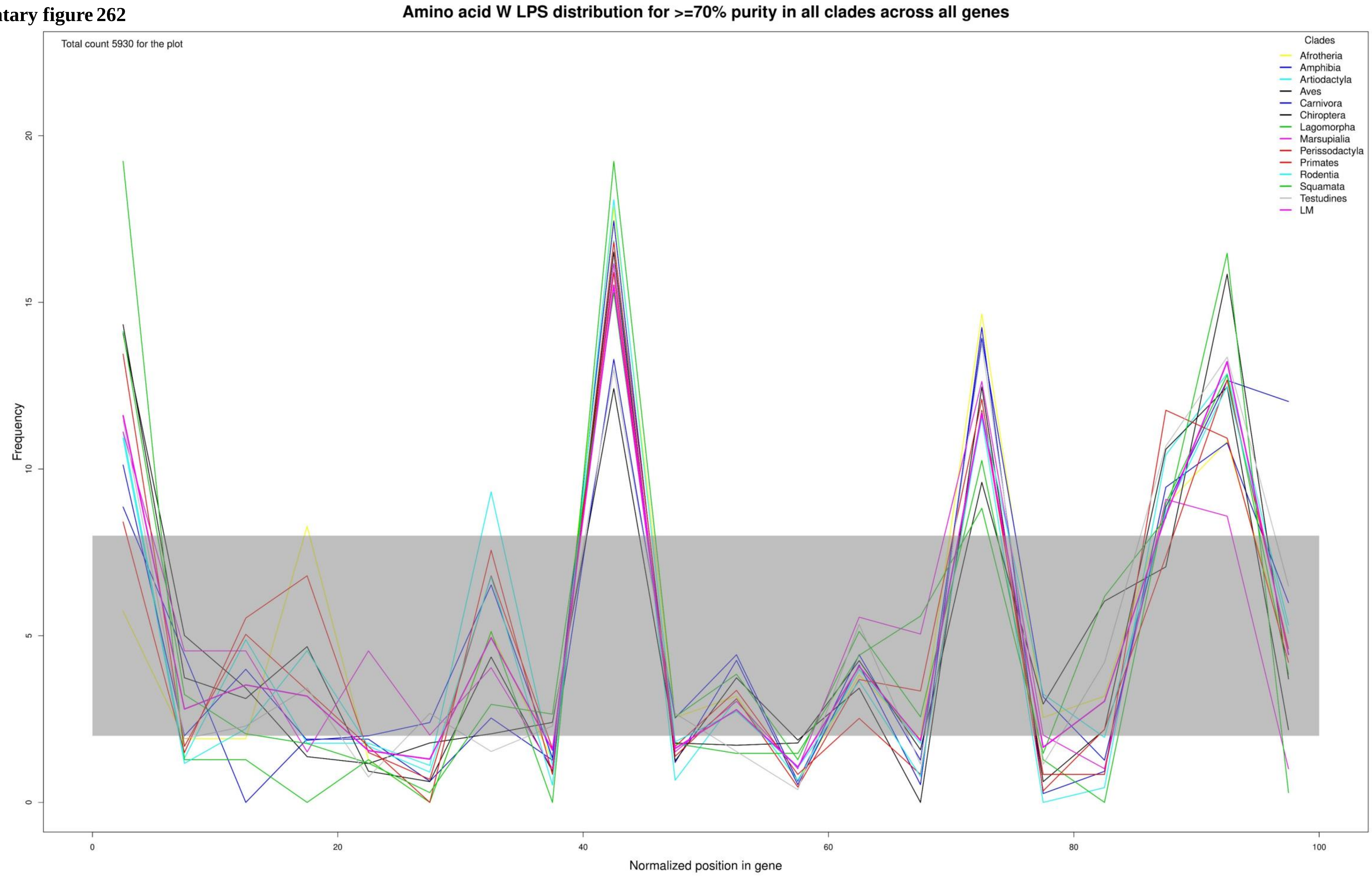

**Supplementary figure 263**

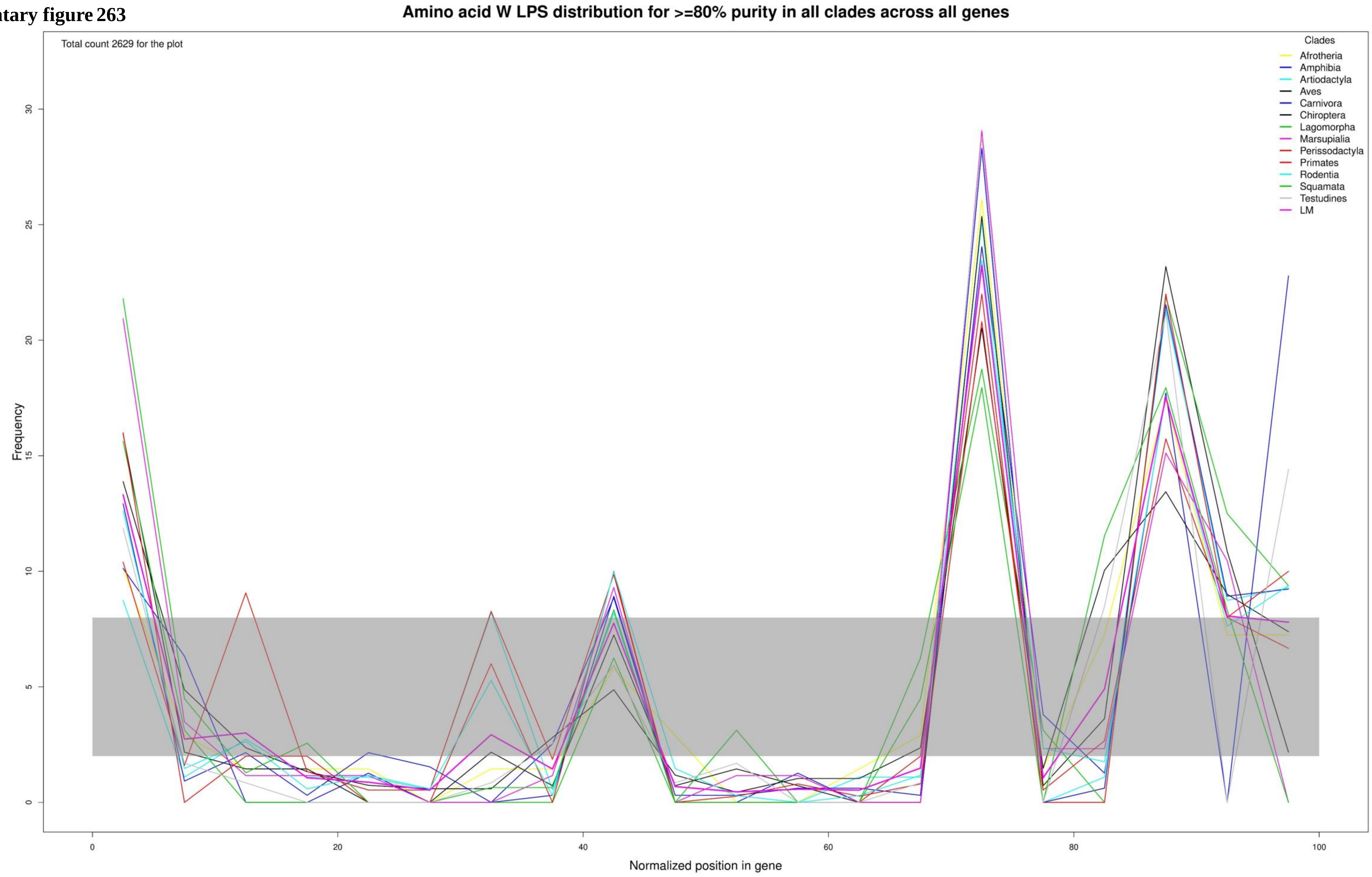

Supplementary figure 264

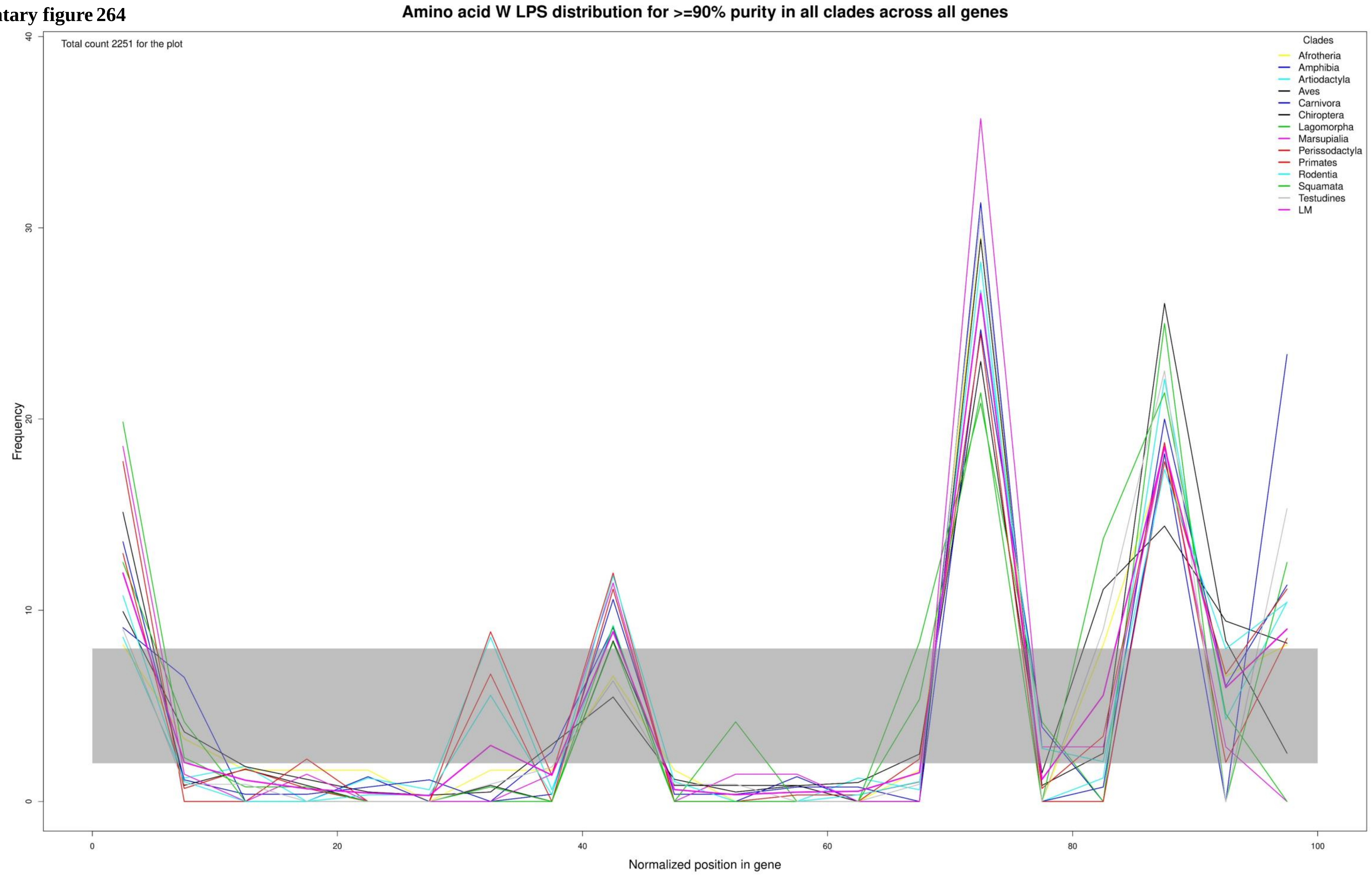

**Supplementary figure 265**

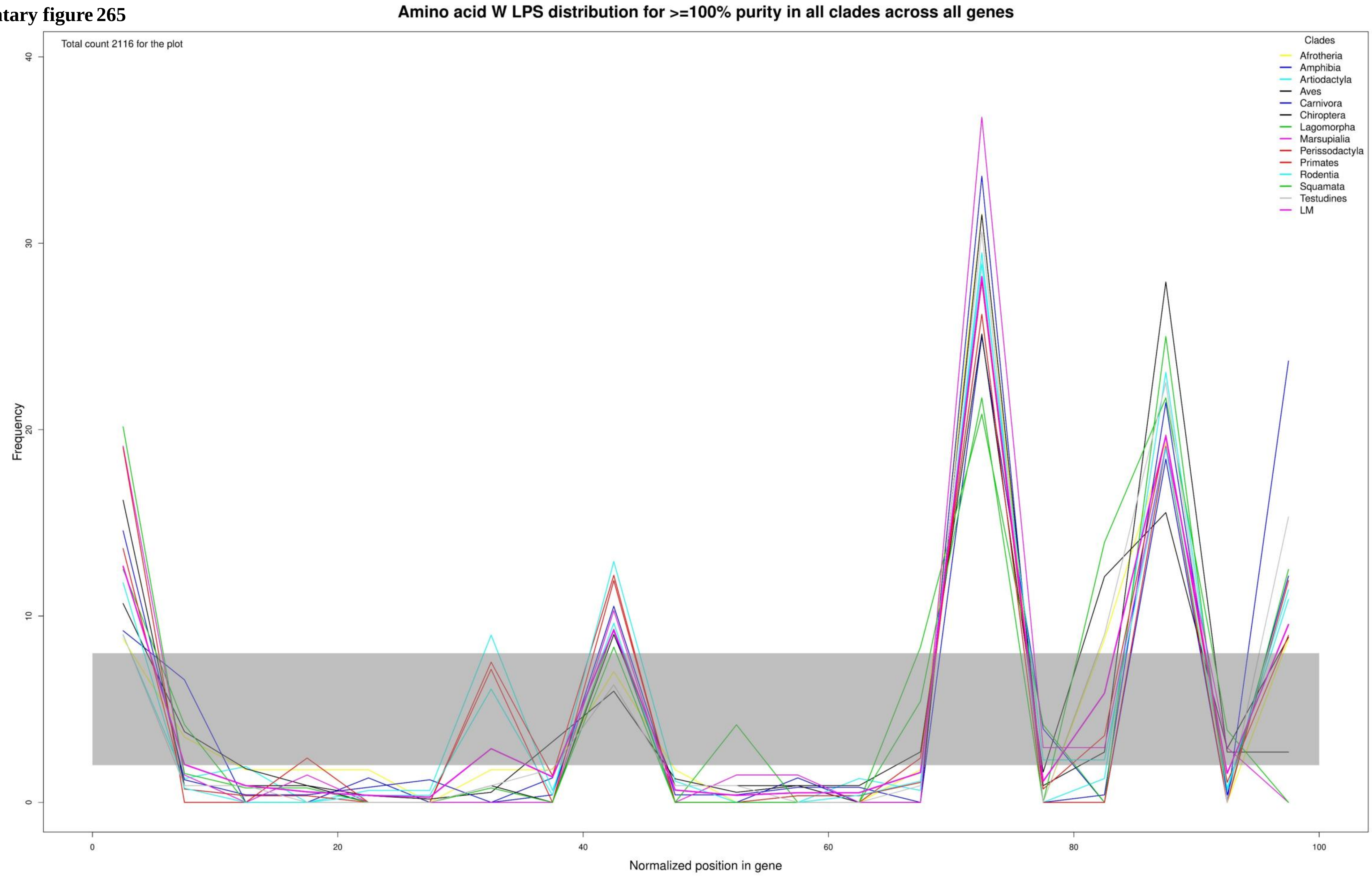

**Supplementary figure 266**

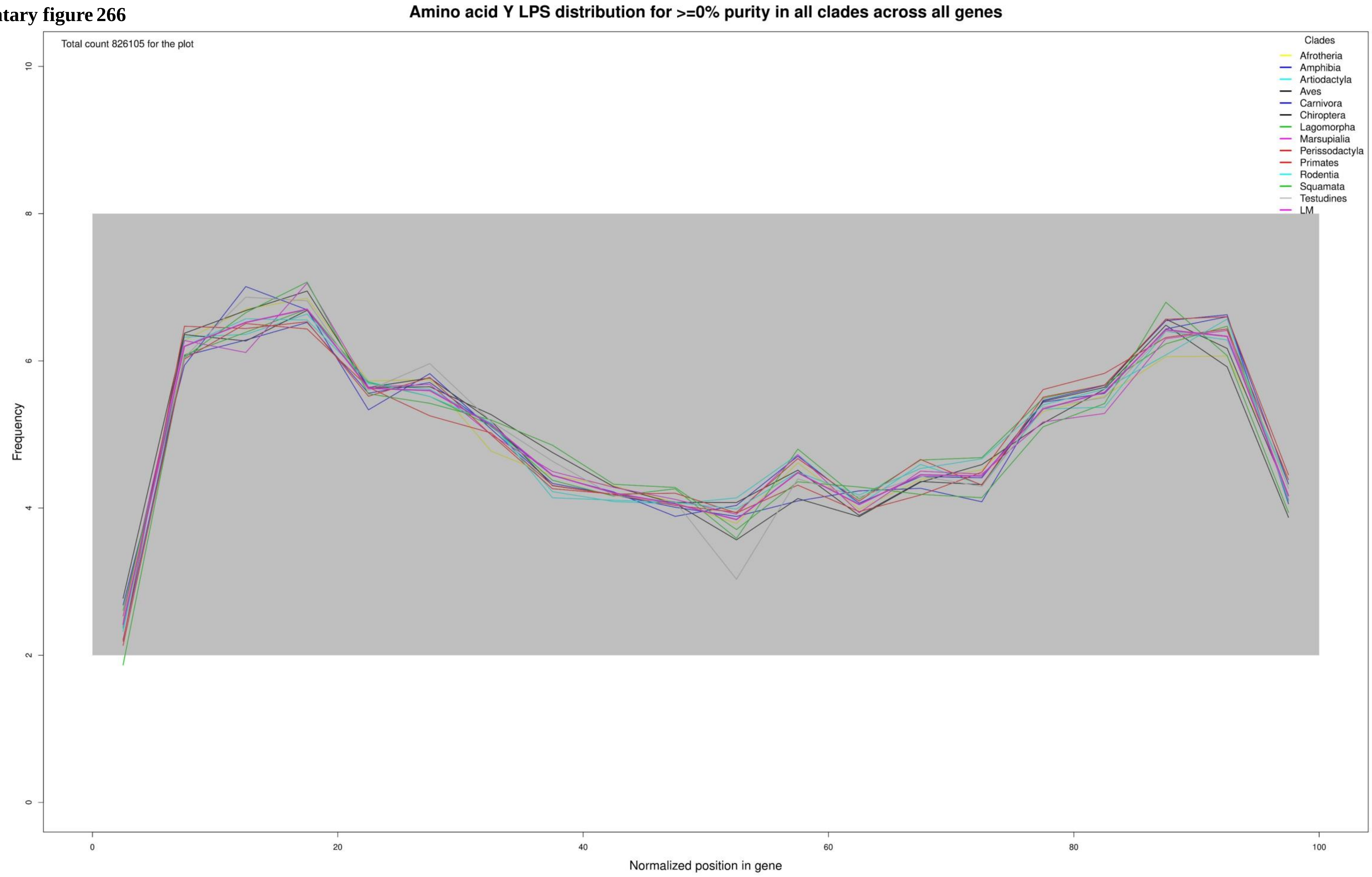

**Supplementary figure 267**

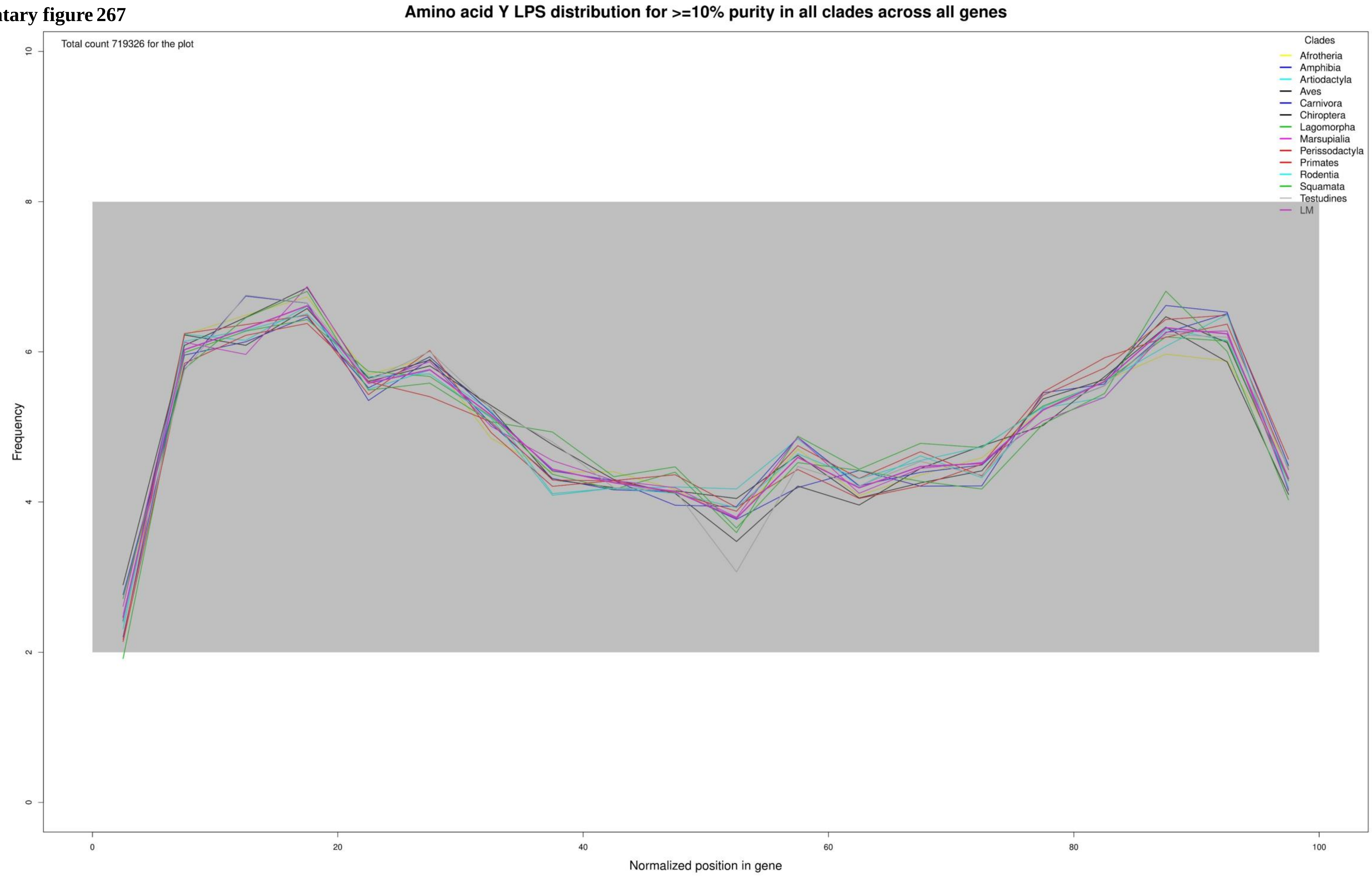

**Supplementary figure 268**

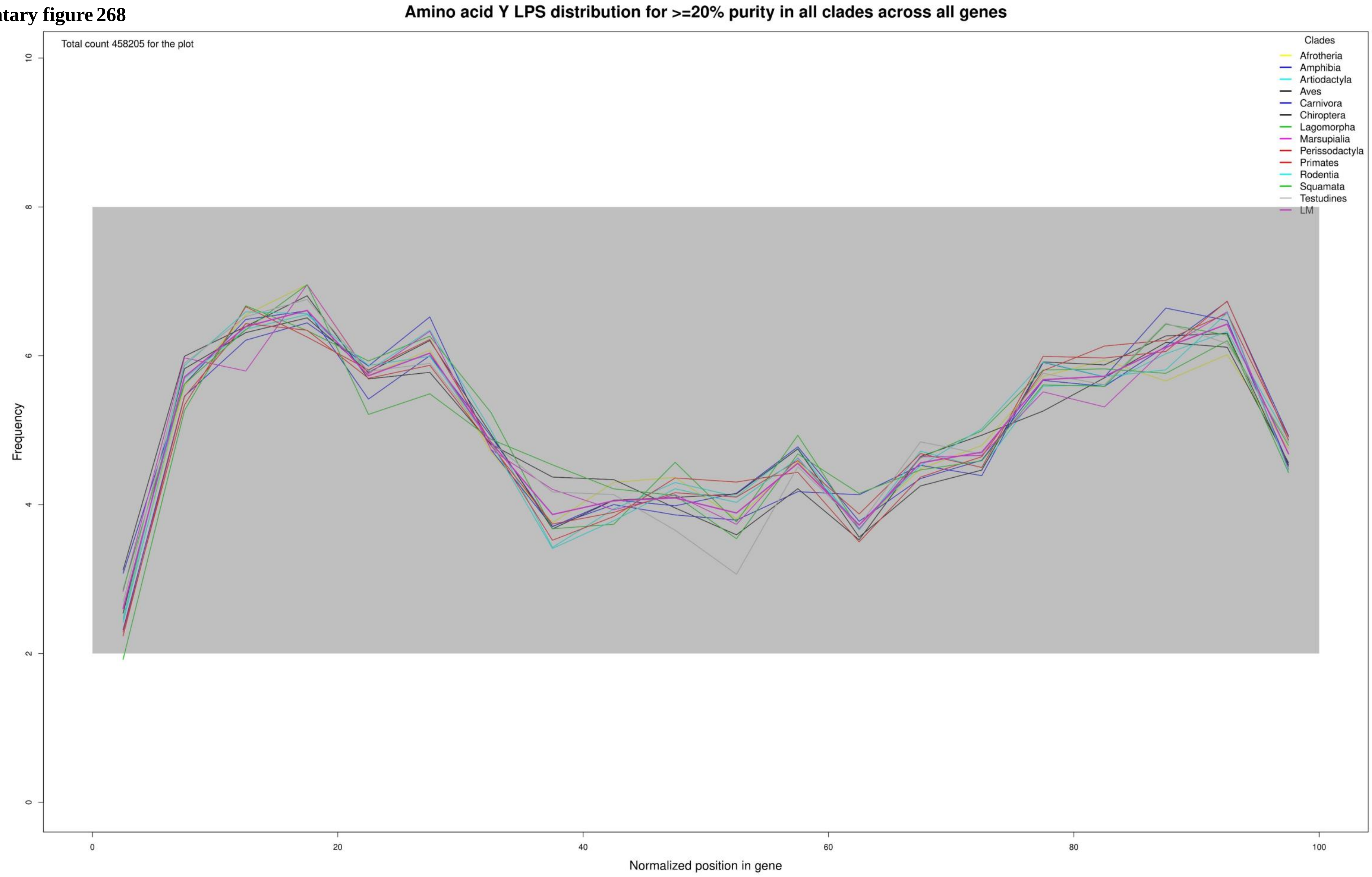

Supplementary figure 269

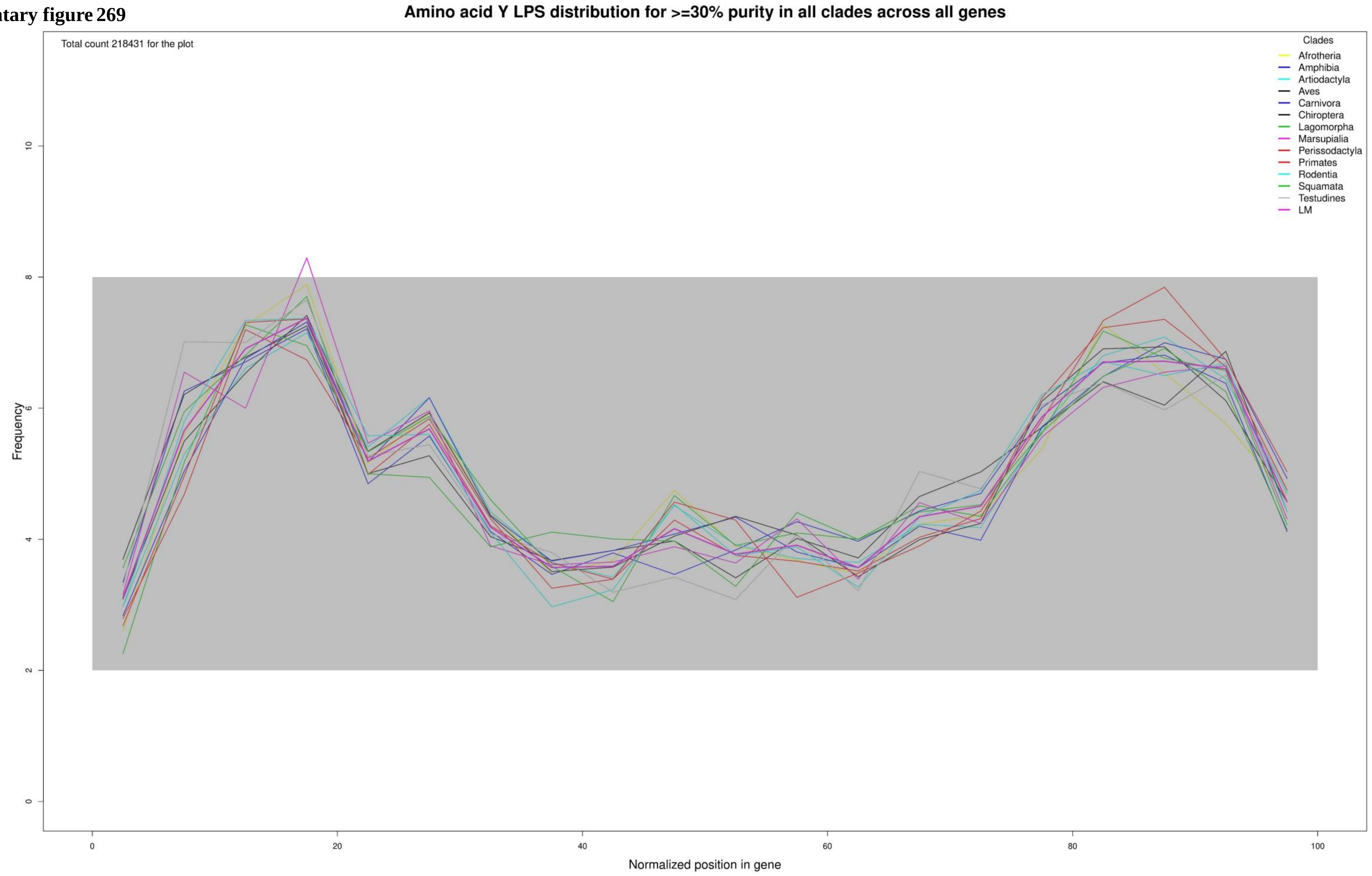

**Supplementary figure 270**

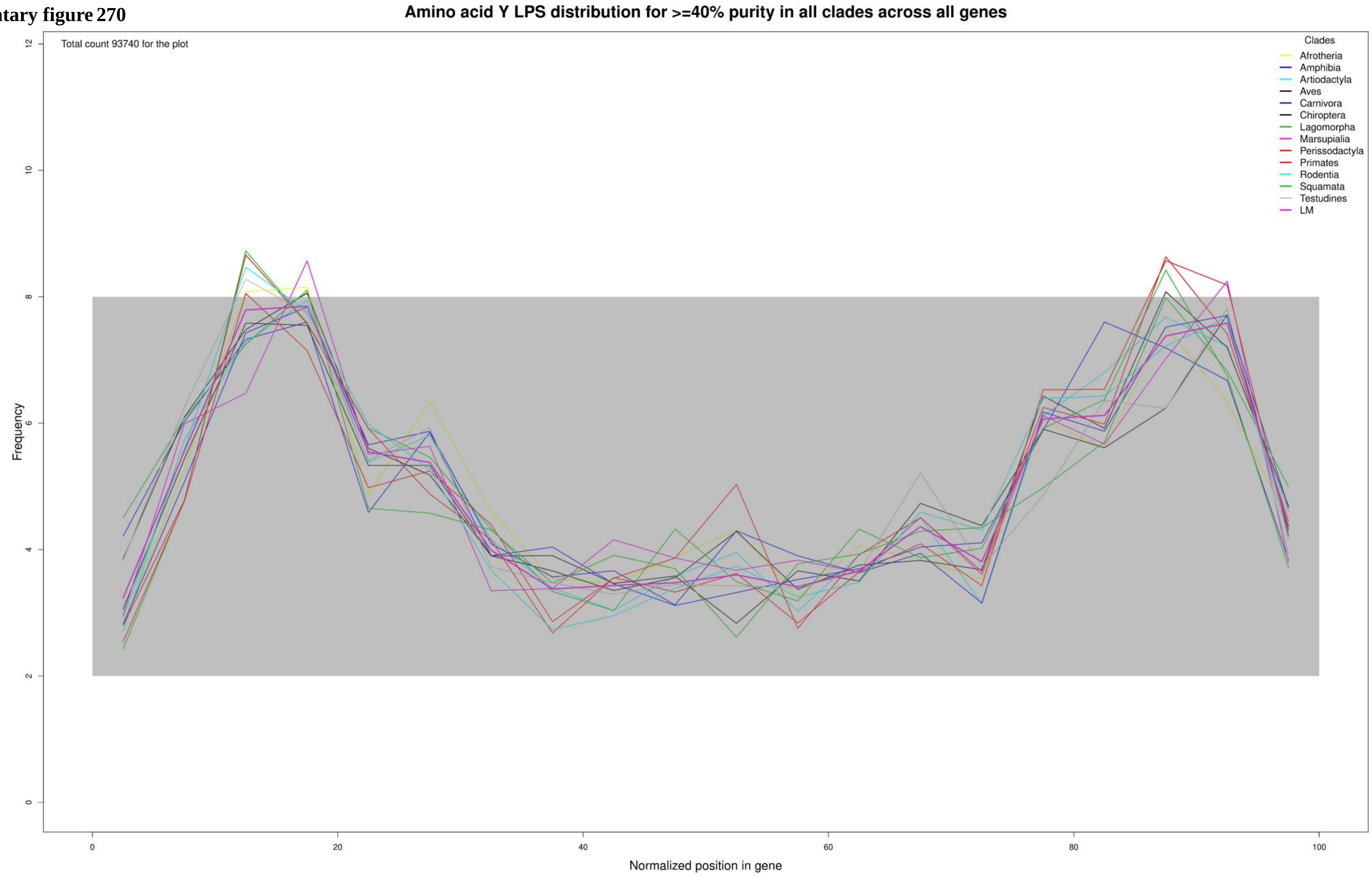

**Supplementary figure 271**

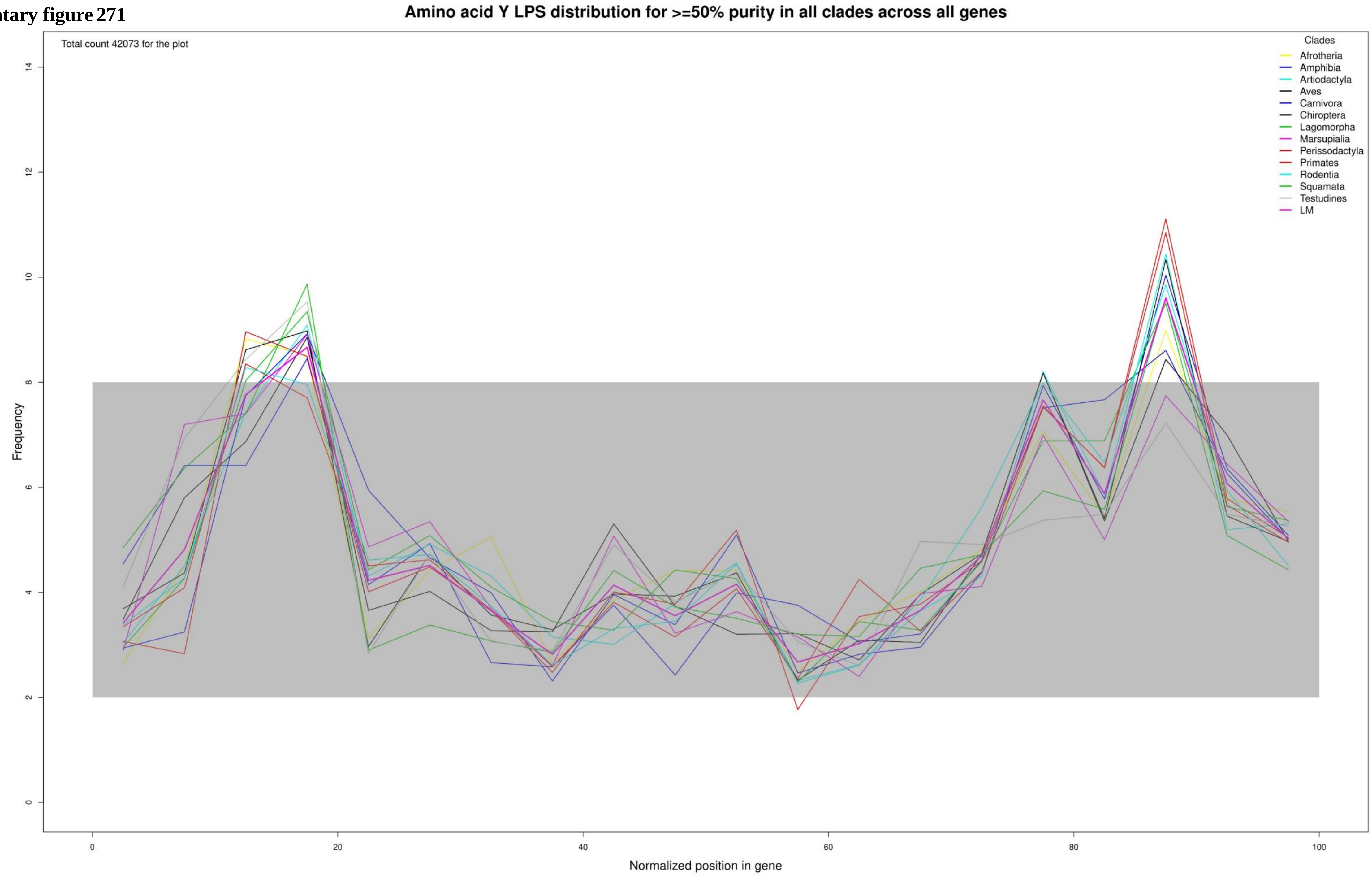

**Supplementary figure 272**

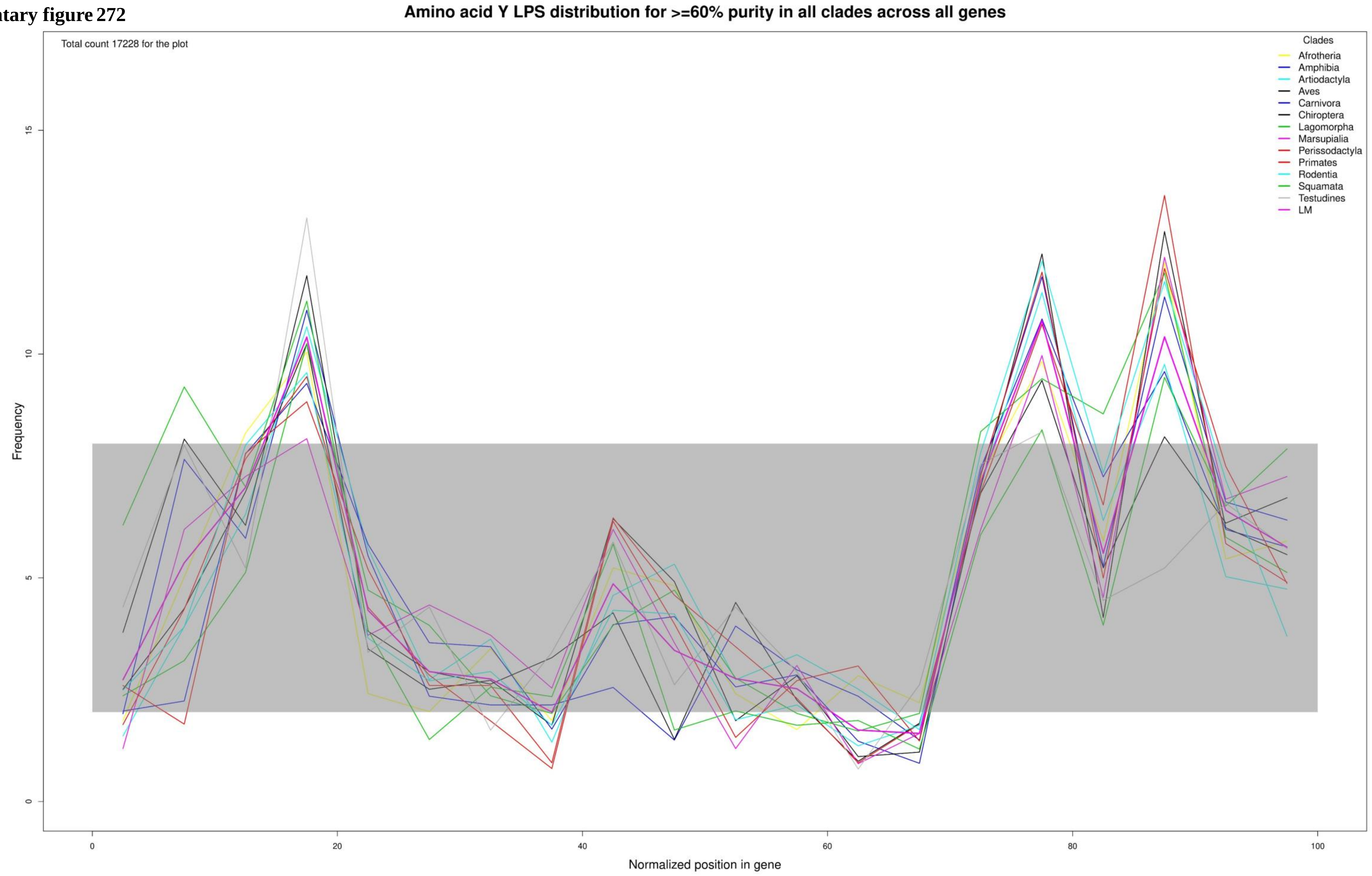

**Supplementary figure 273**

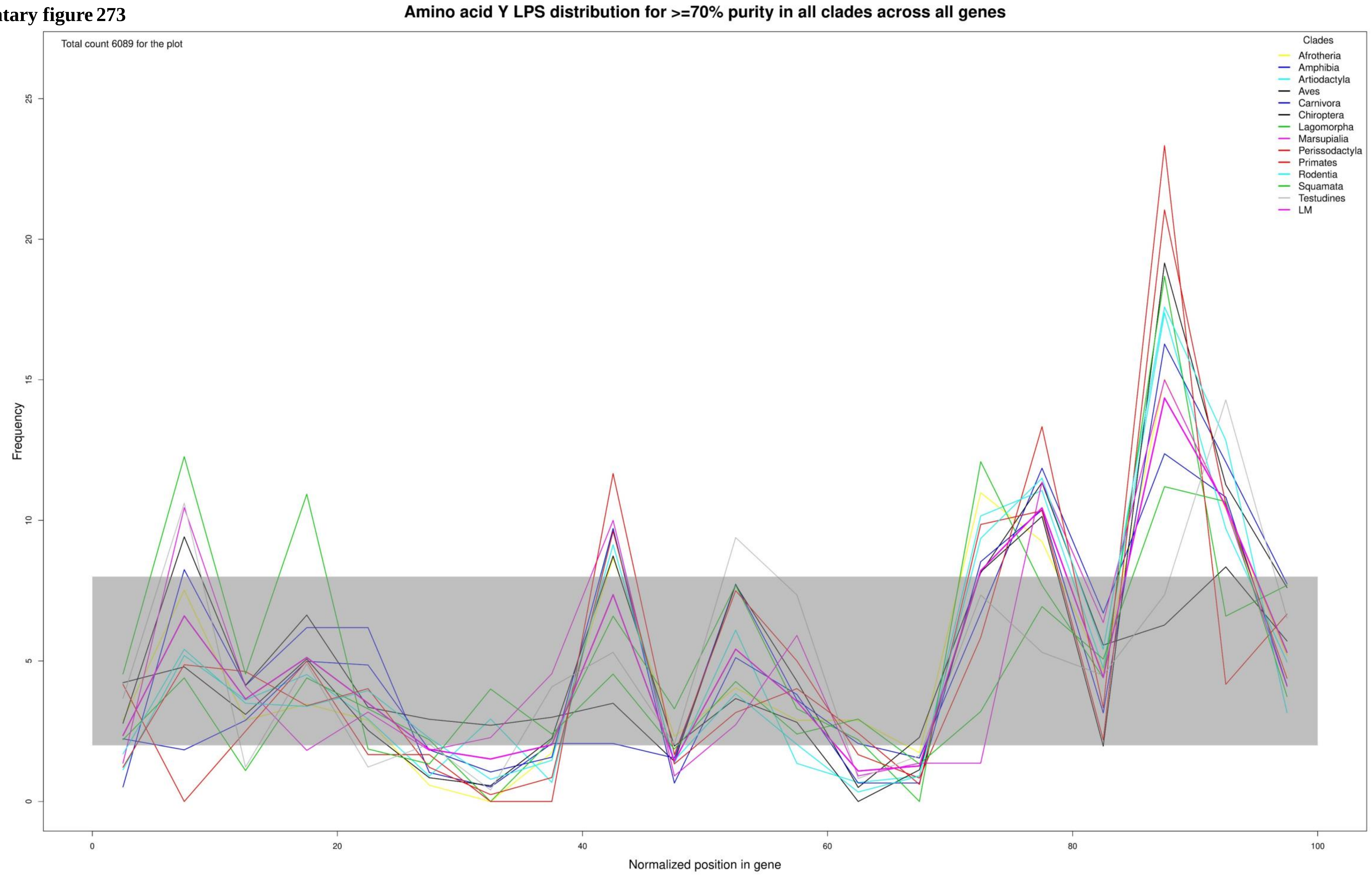

**Supplementary figure 274**

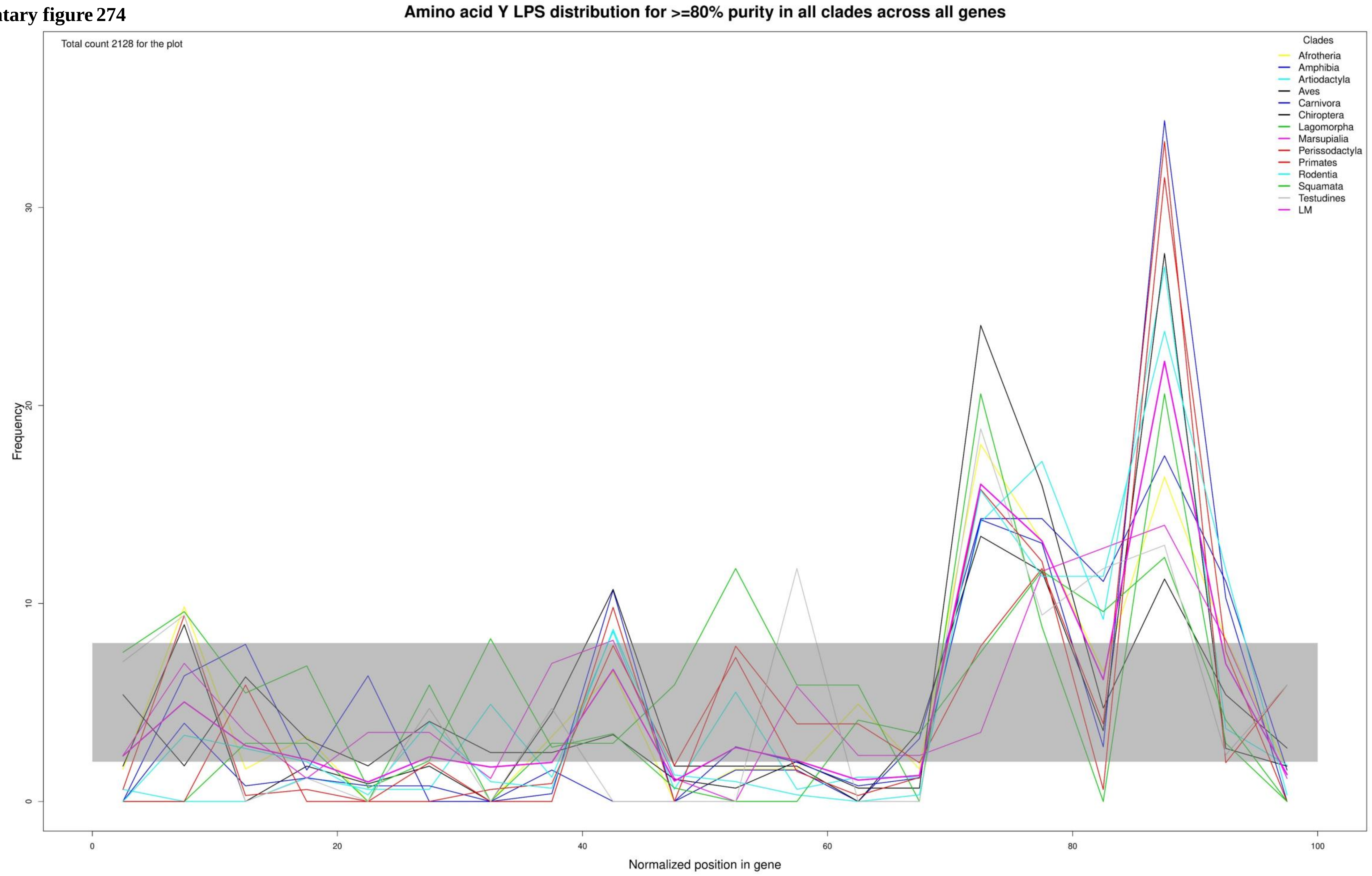

**Supplementary figure 275**

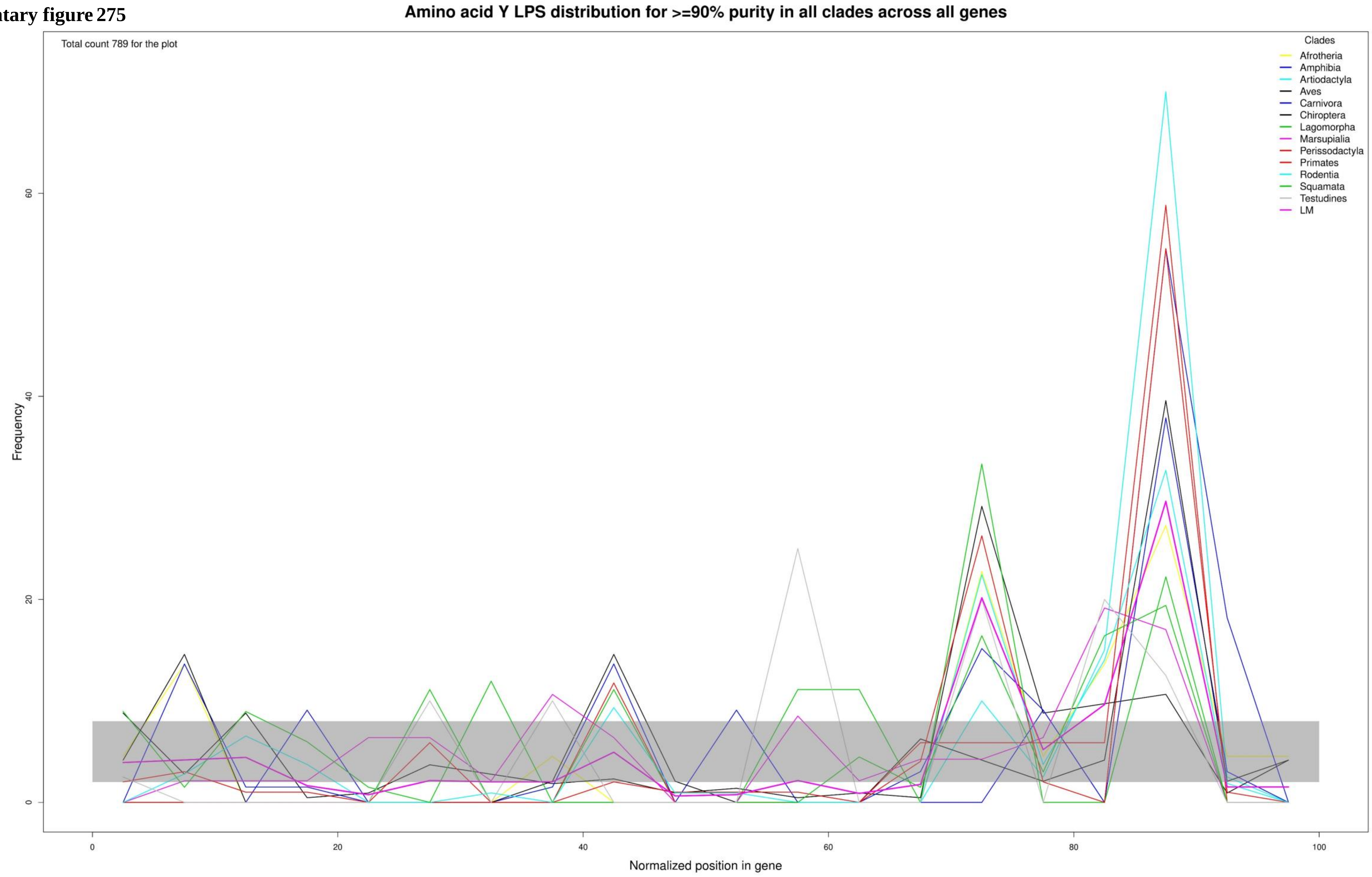

**Supplementary figure 276**

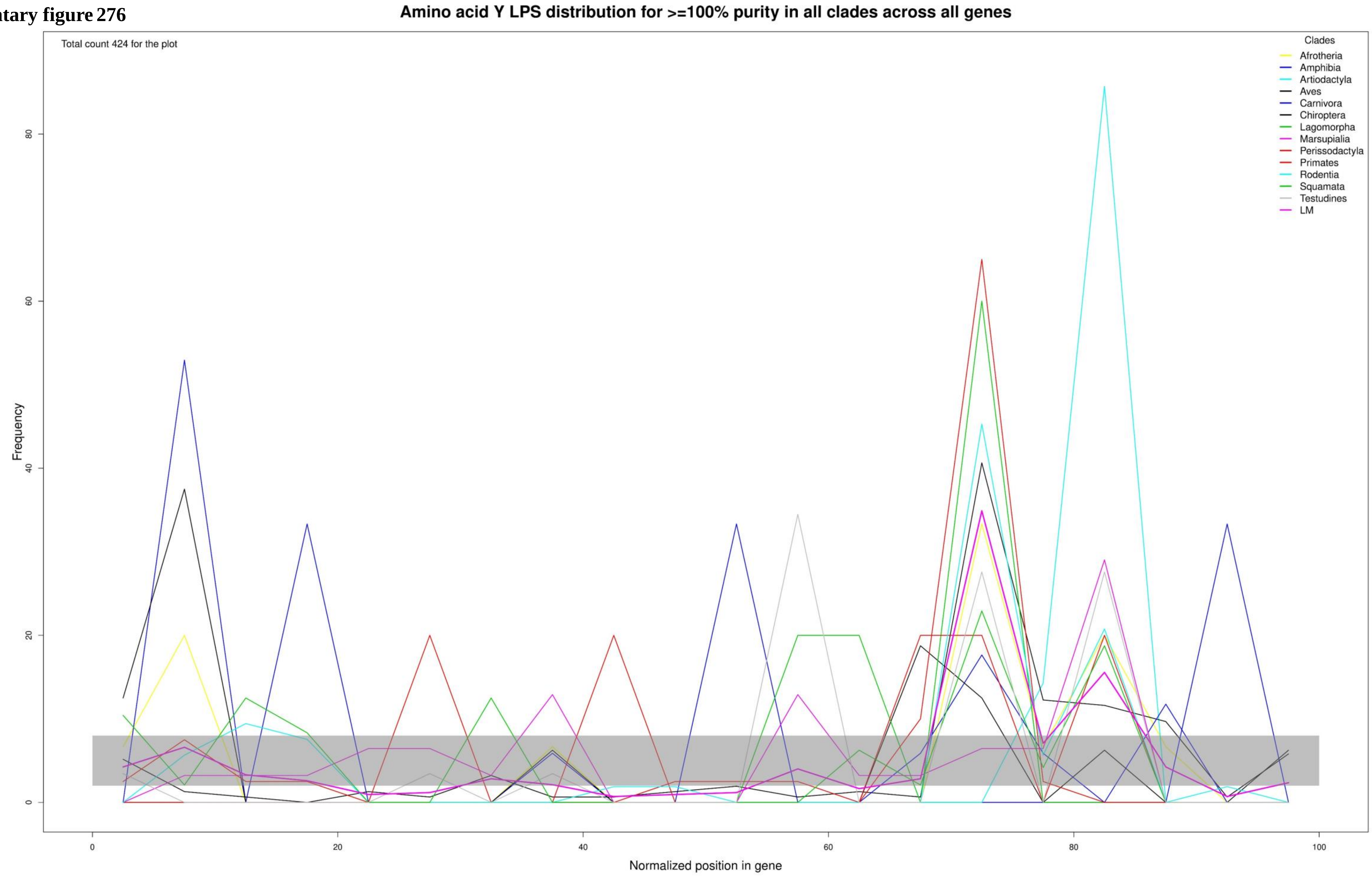

Supplementary figure 277

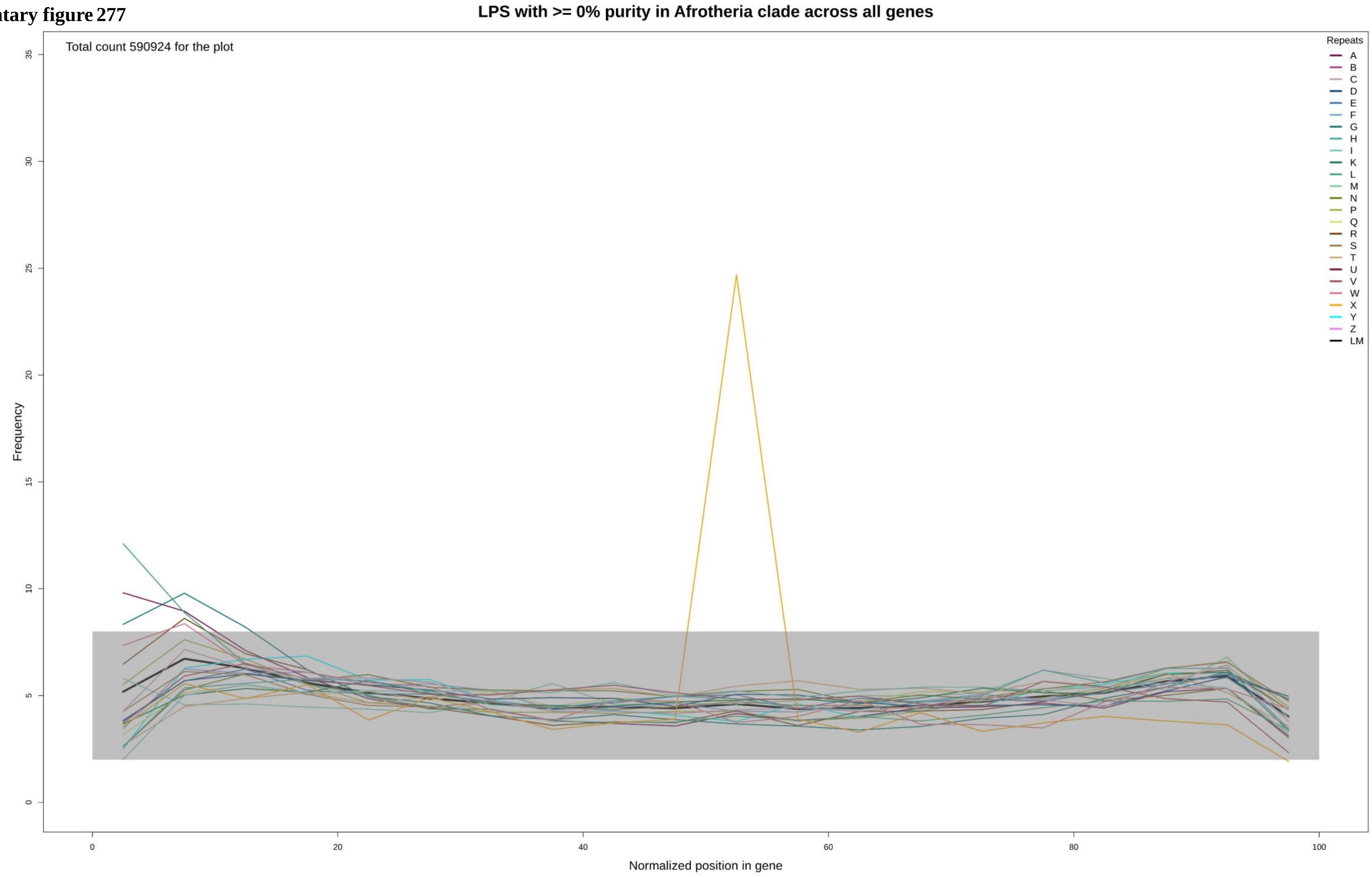

Supplementary figure 278

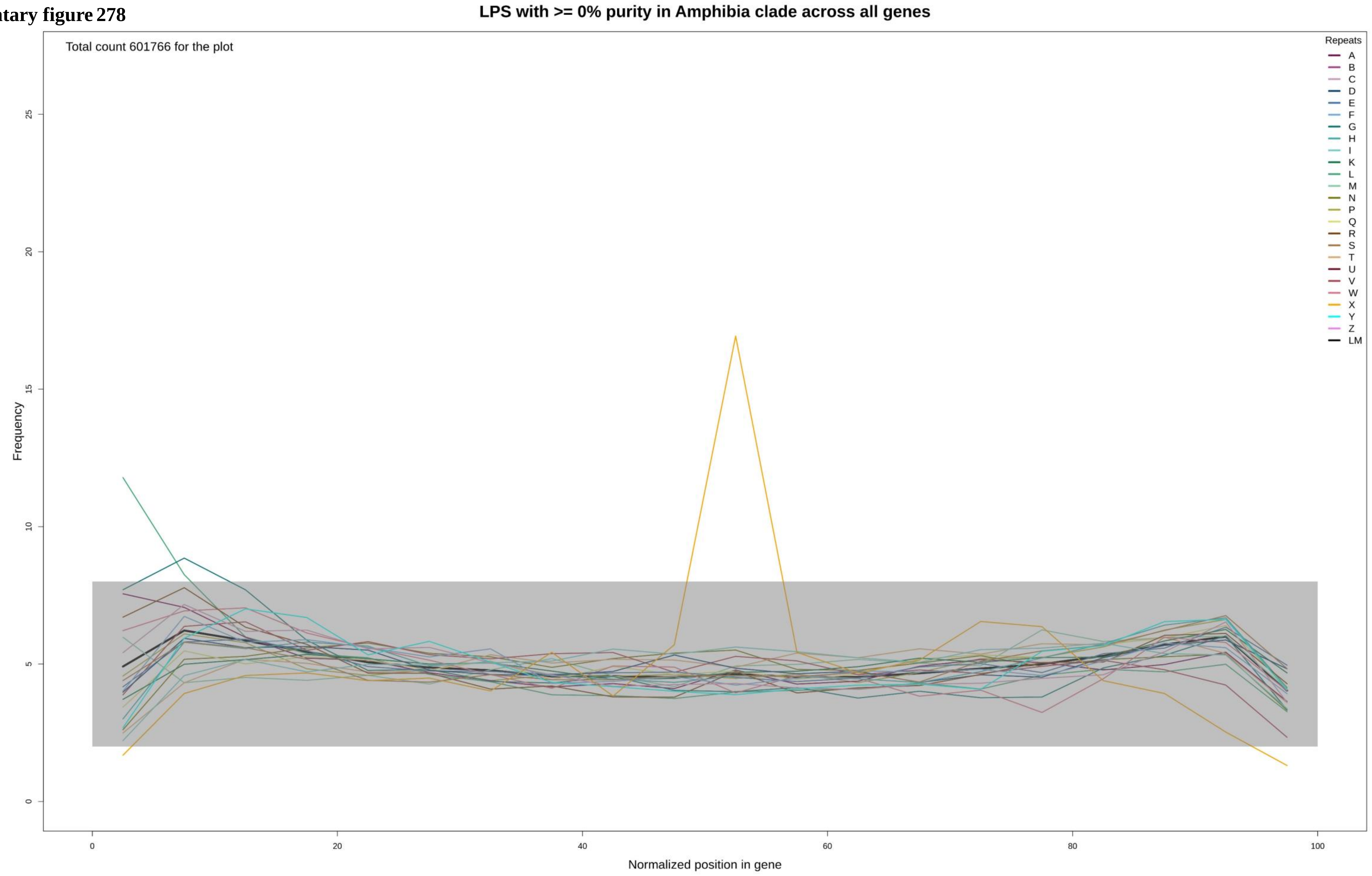

**Supplementary figure 279**

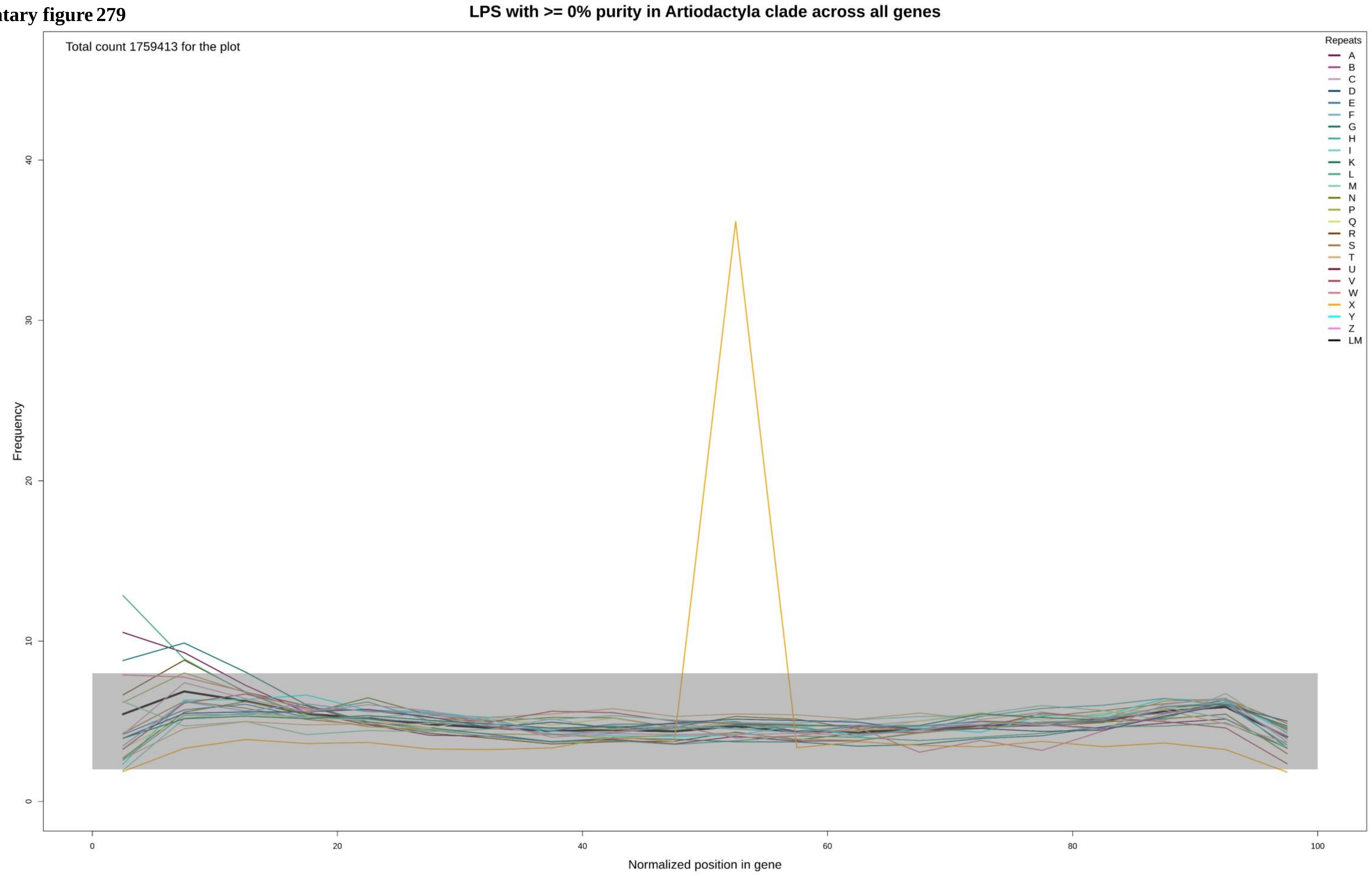

Supplementary figure 280

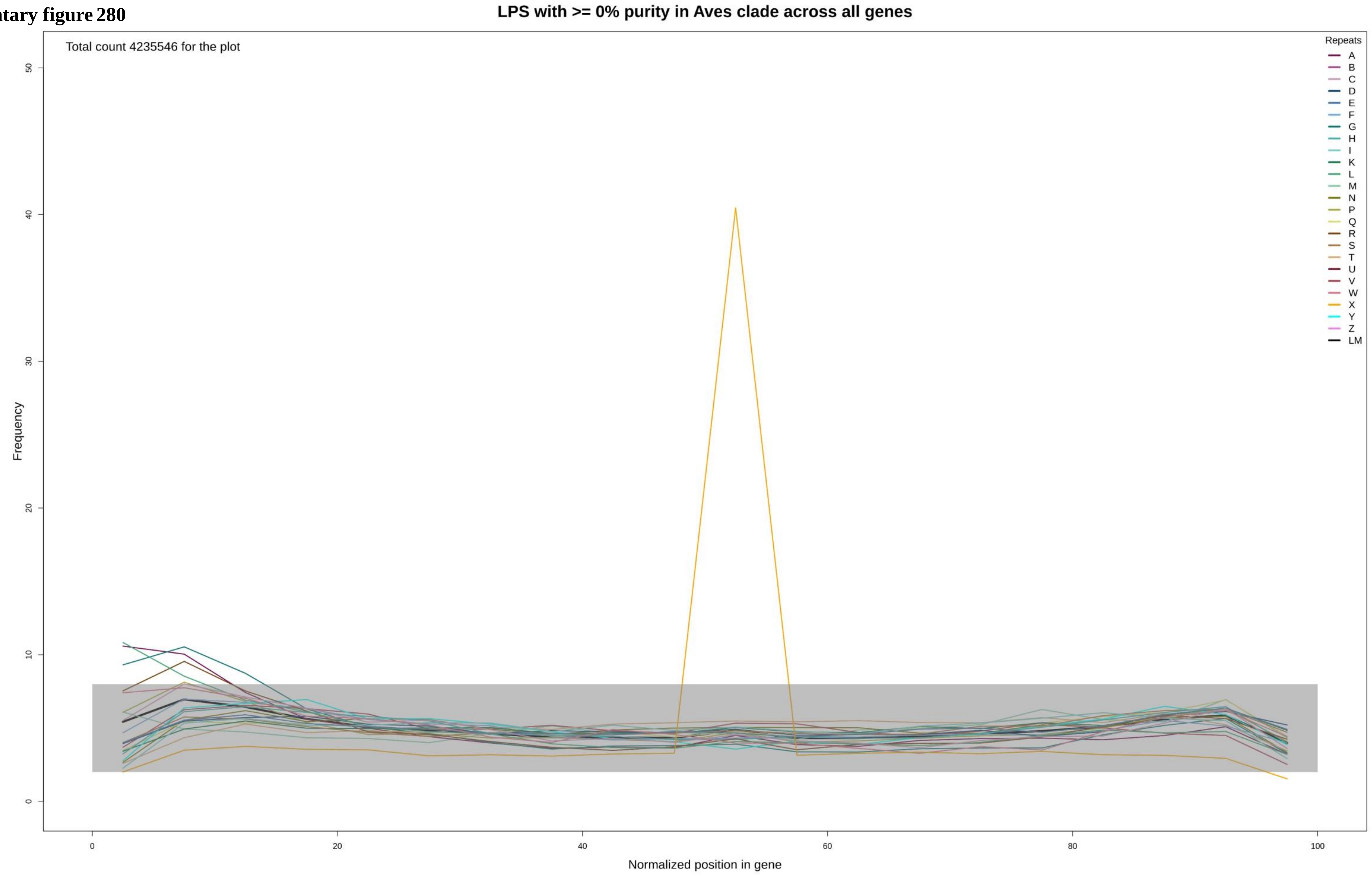

**Supplementary figure 281**

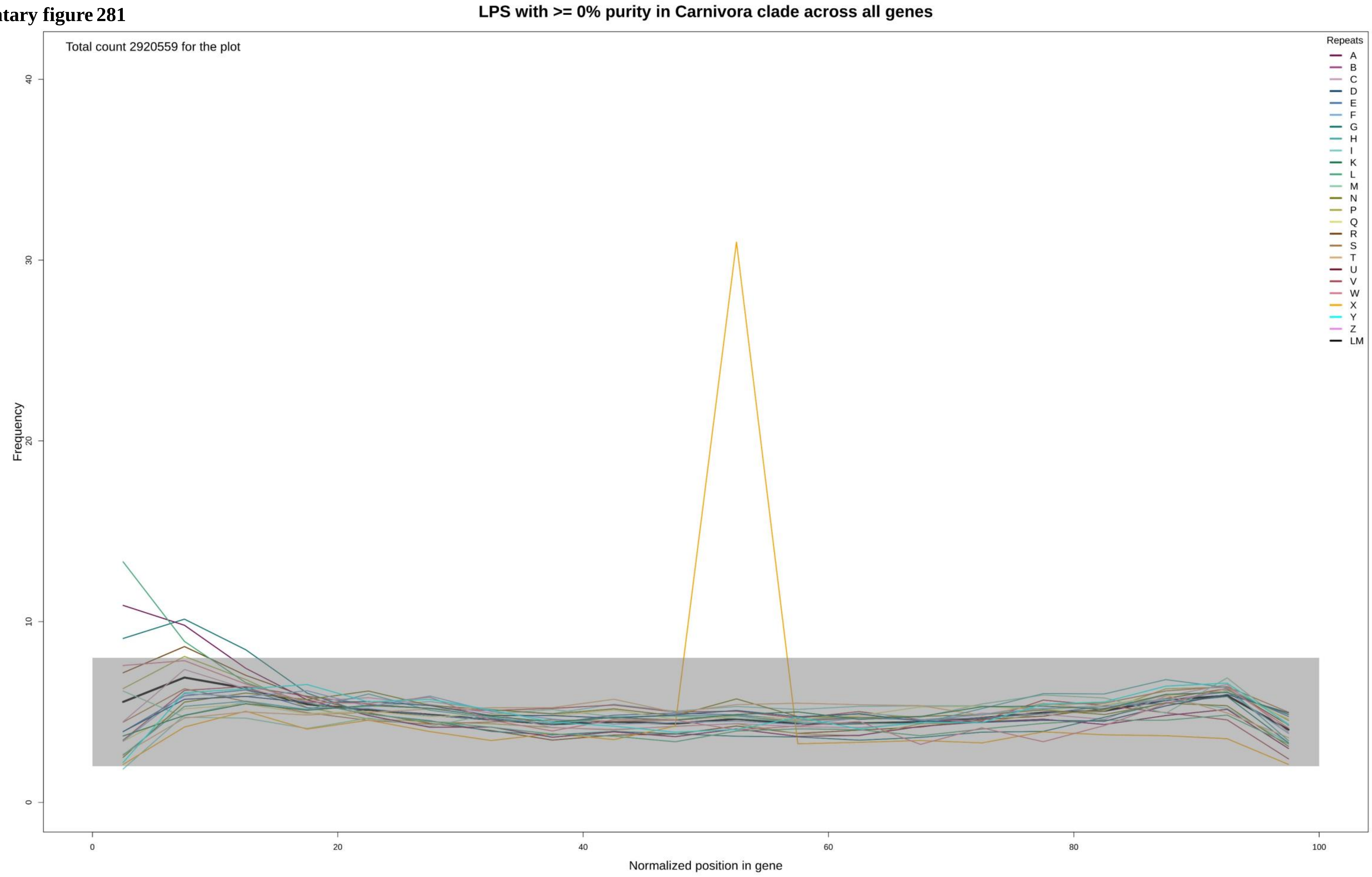

**Supplementary figure 282**

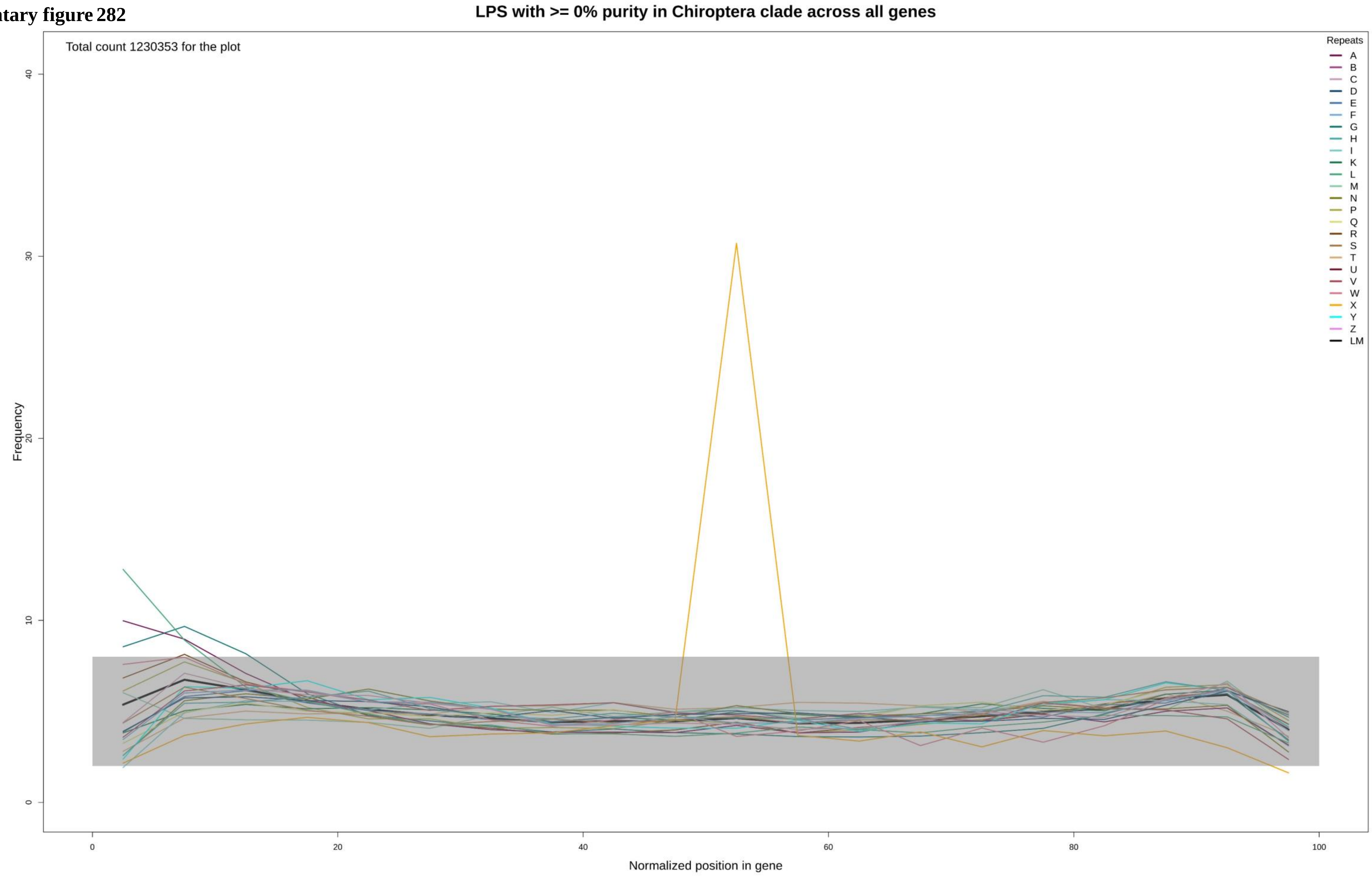

**Supplementary figure 283**

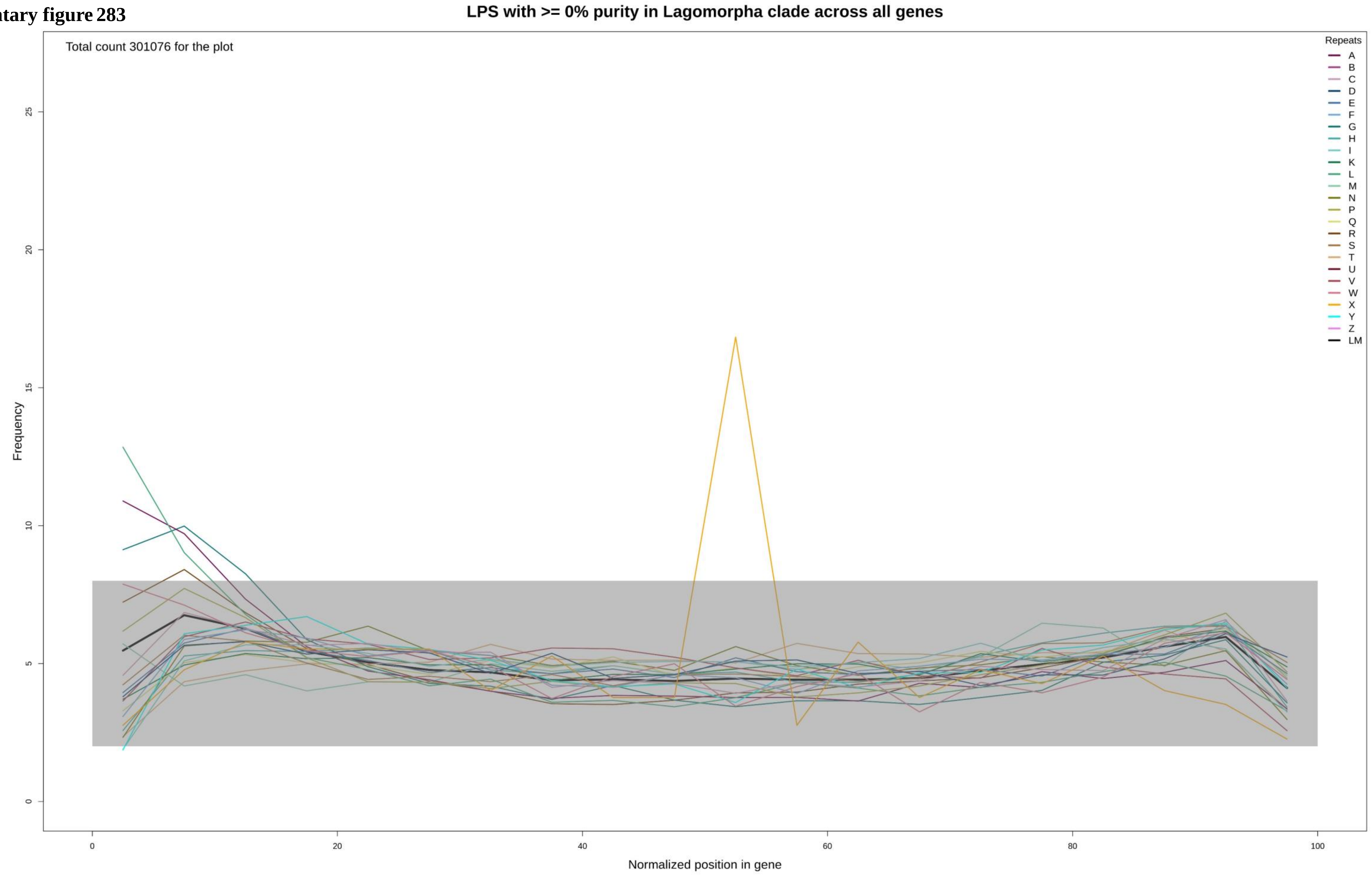

Supplementary figure 284

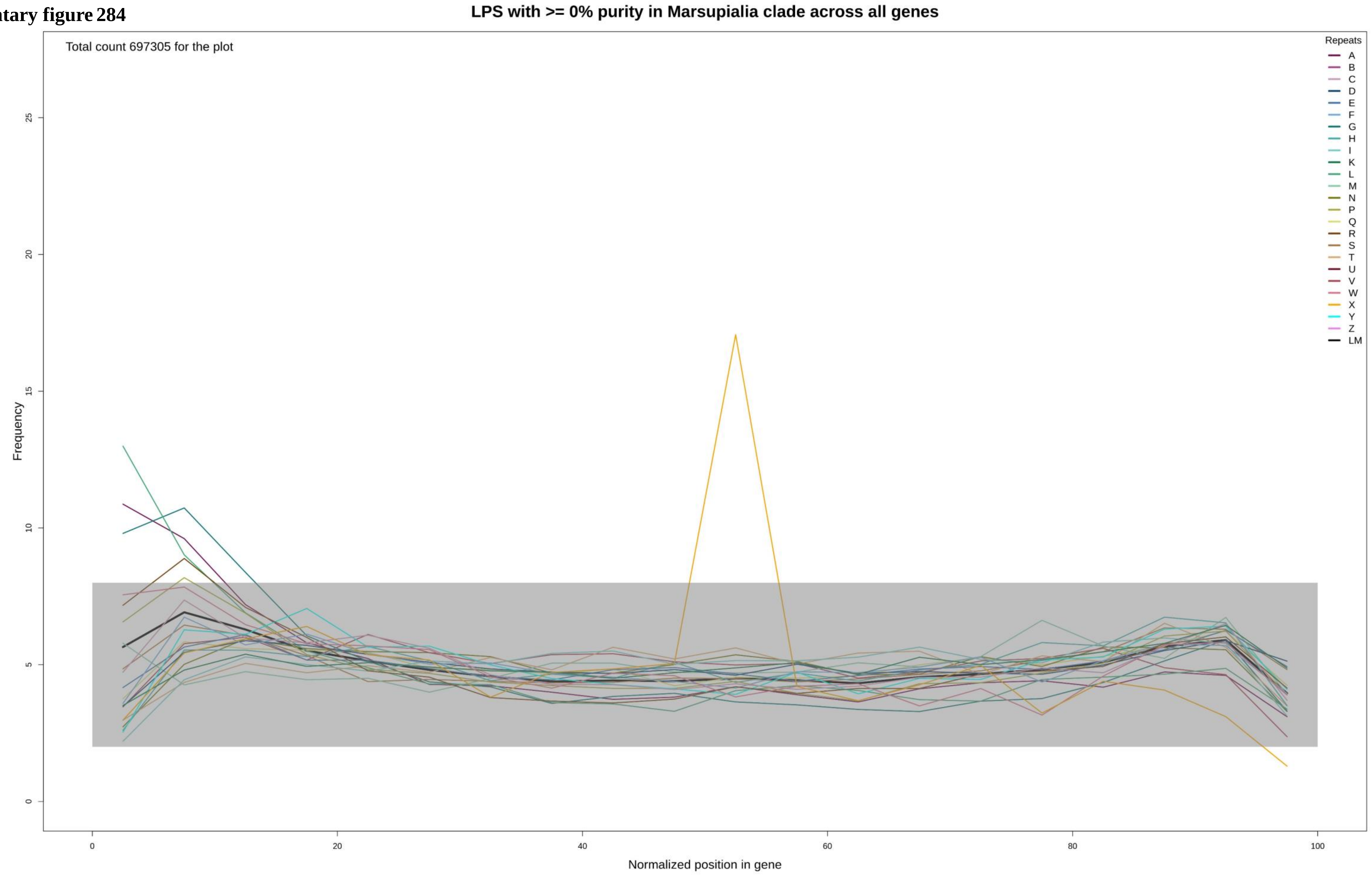

**Supplementary figure 285**

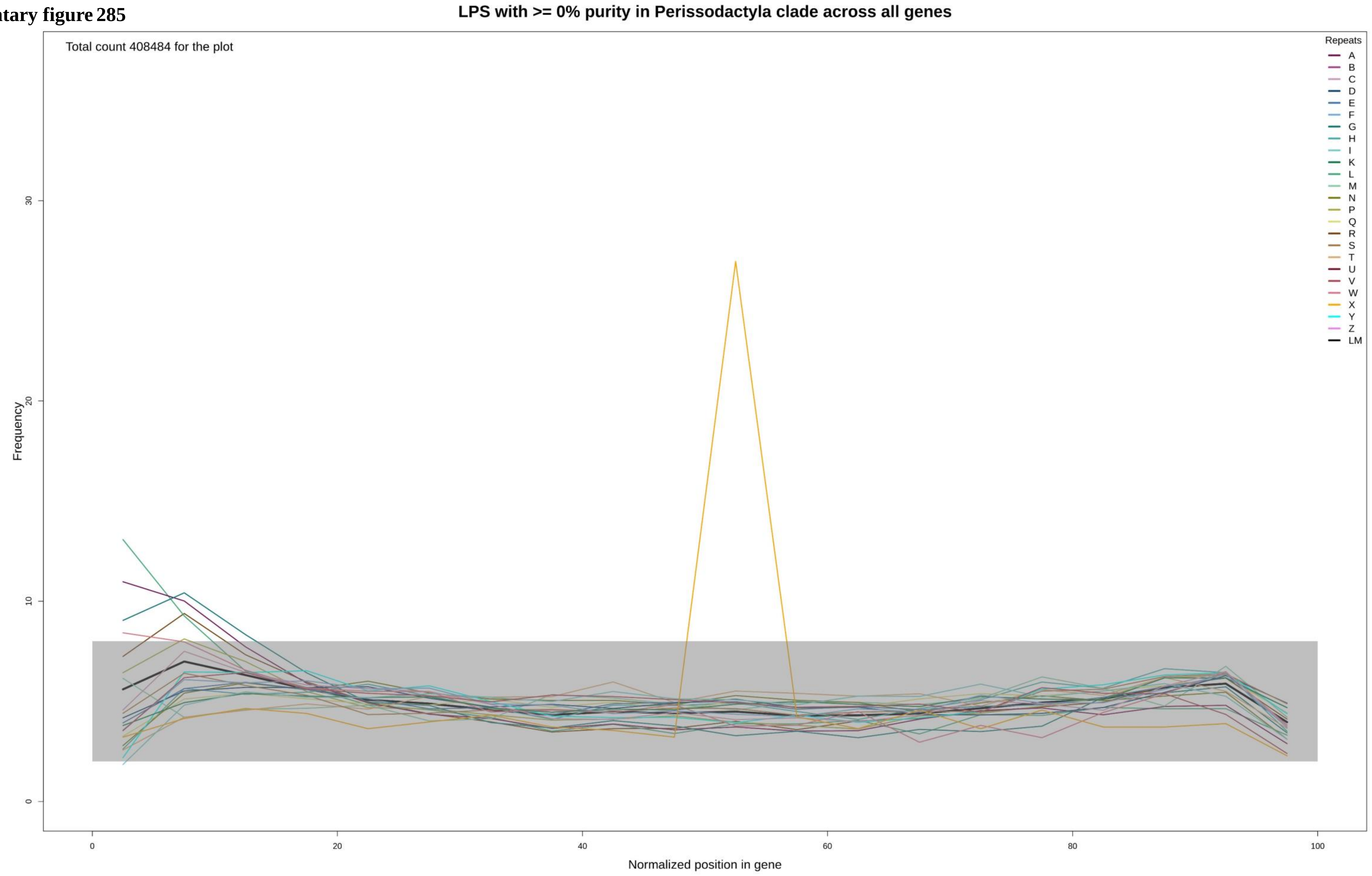

**Supplementary figure 286**

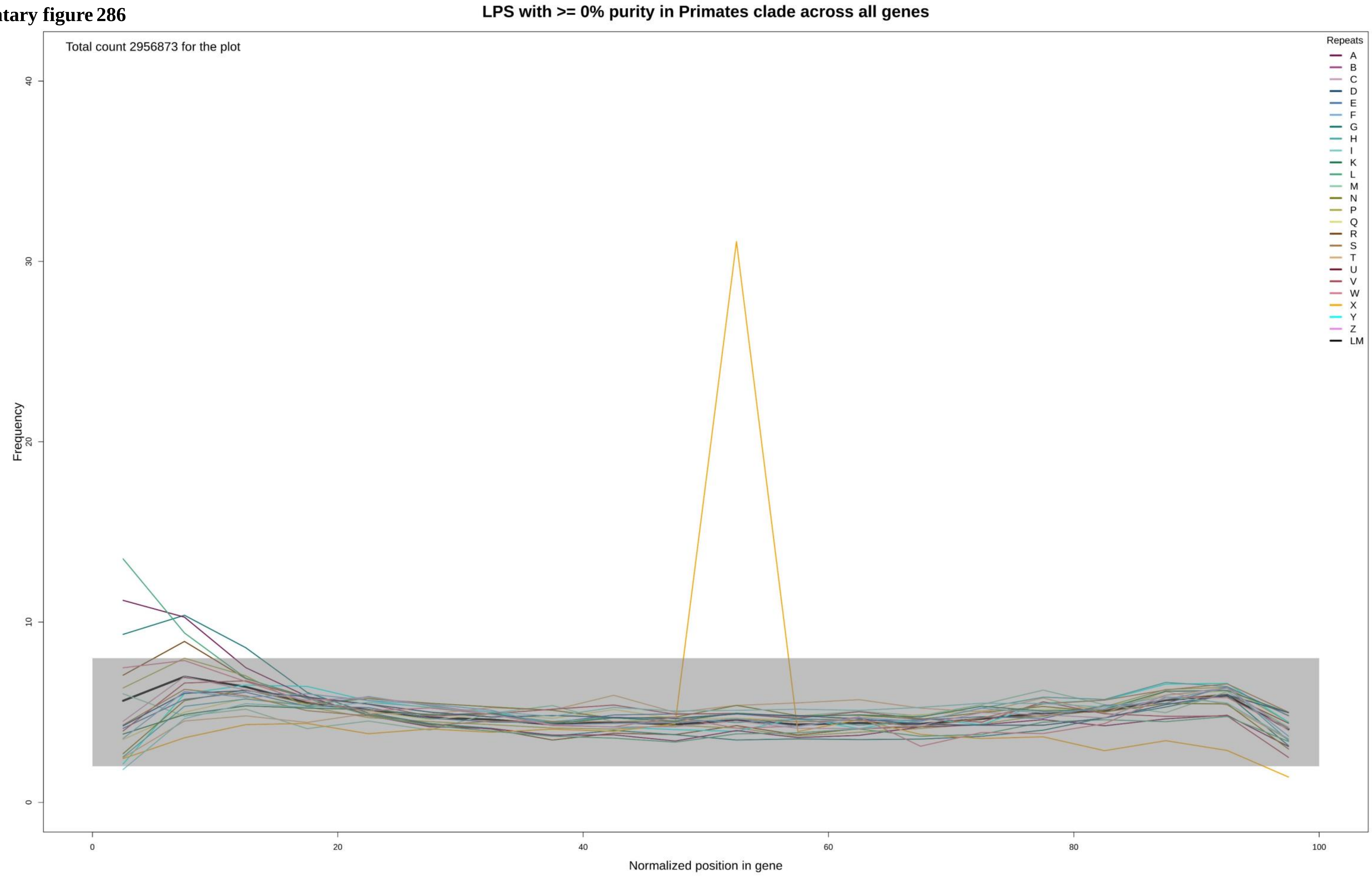

Supplementary figure 287

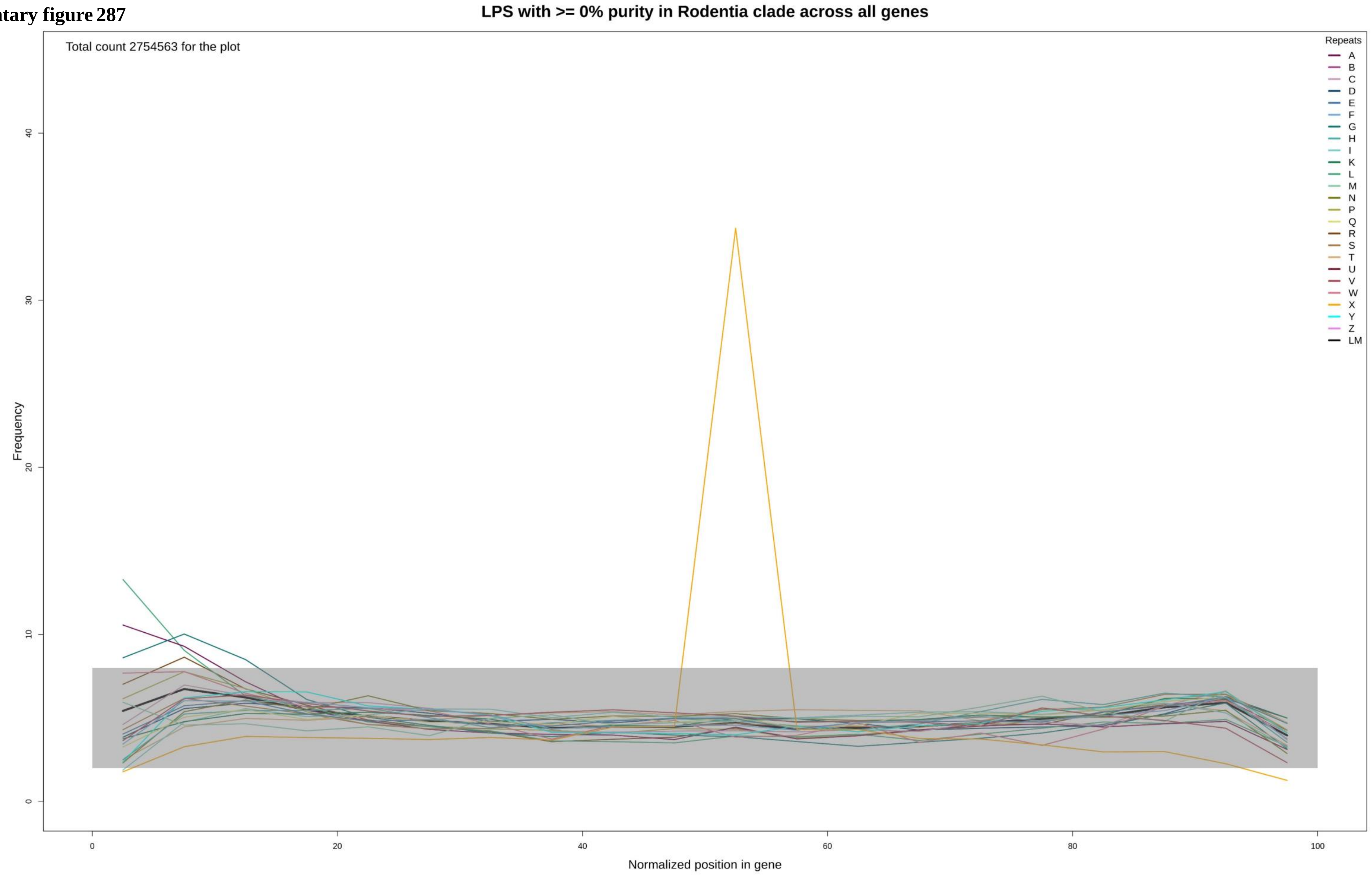

Supplementary figure 288

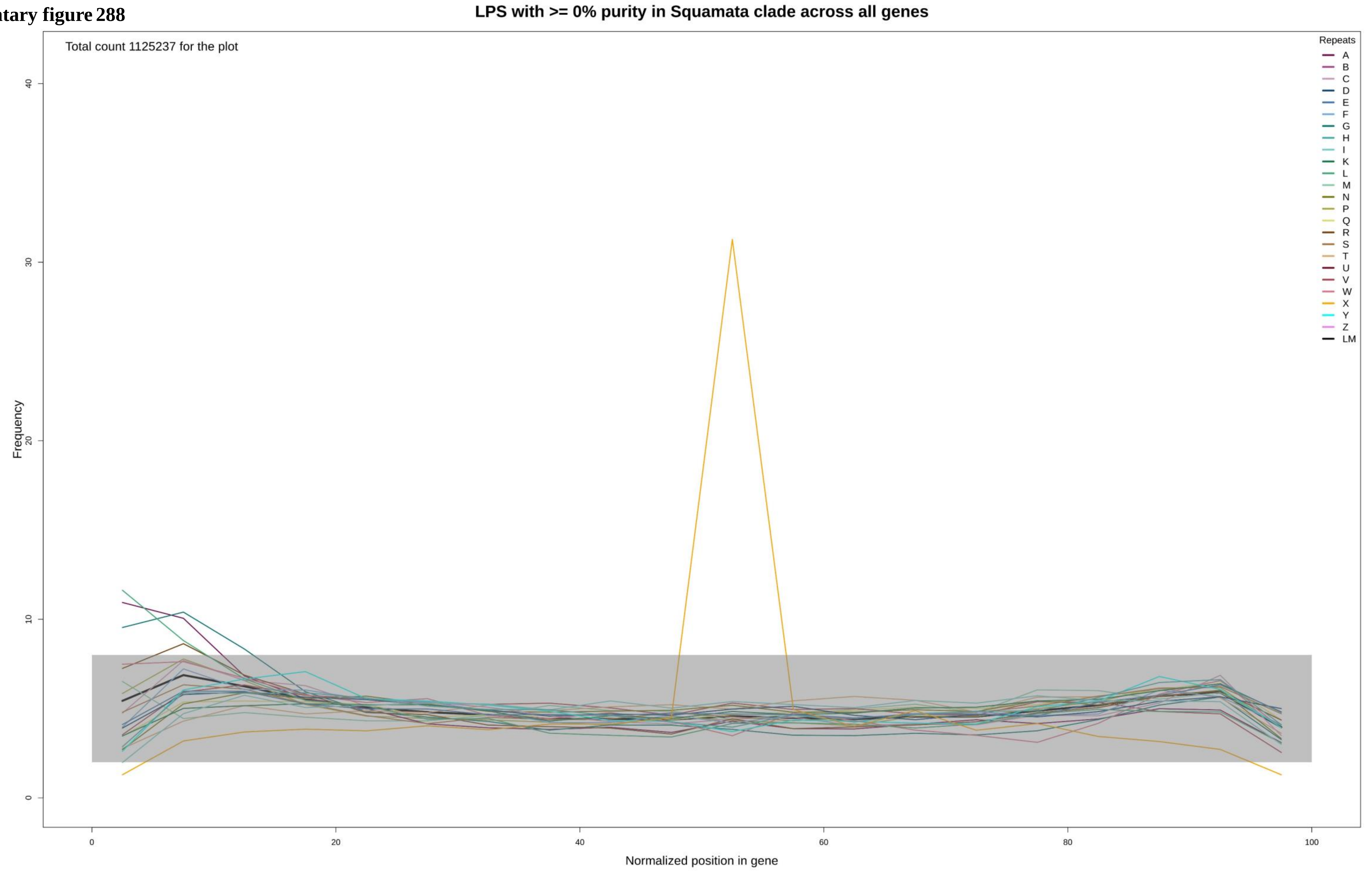

Supplementary figure 289

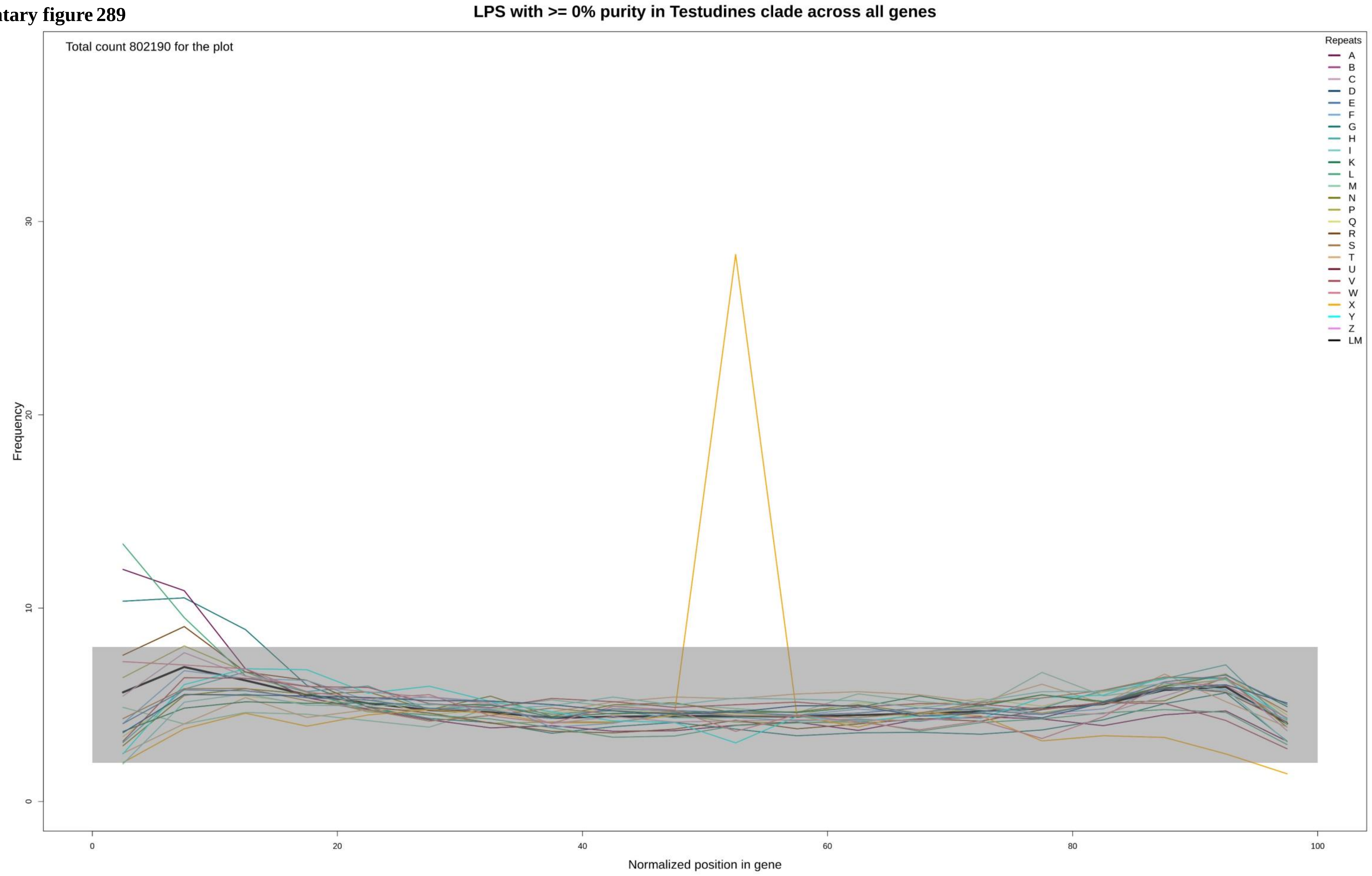

**Supplementary figure 290**

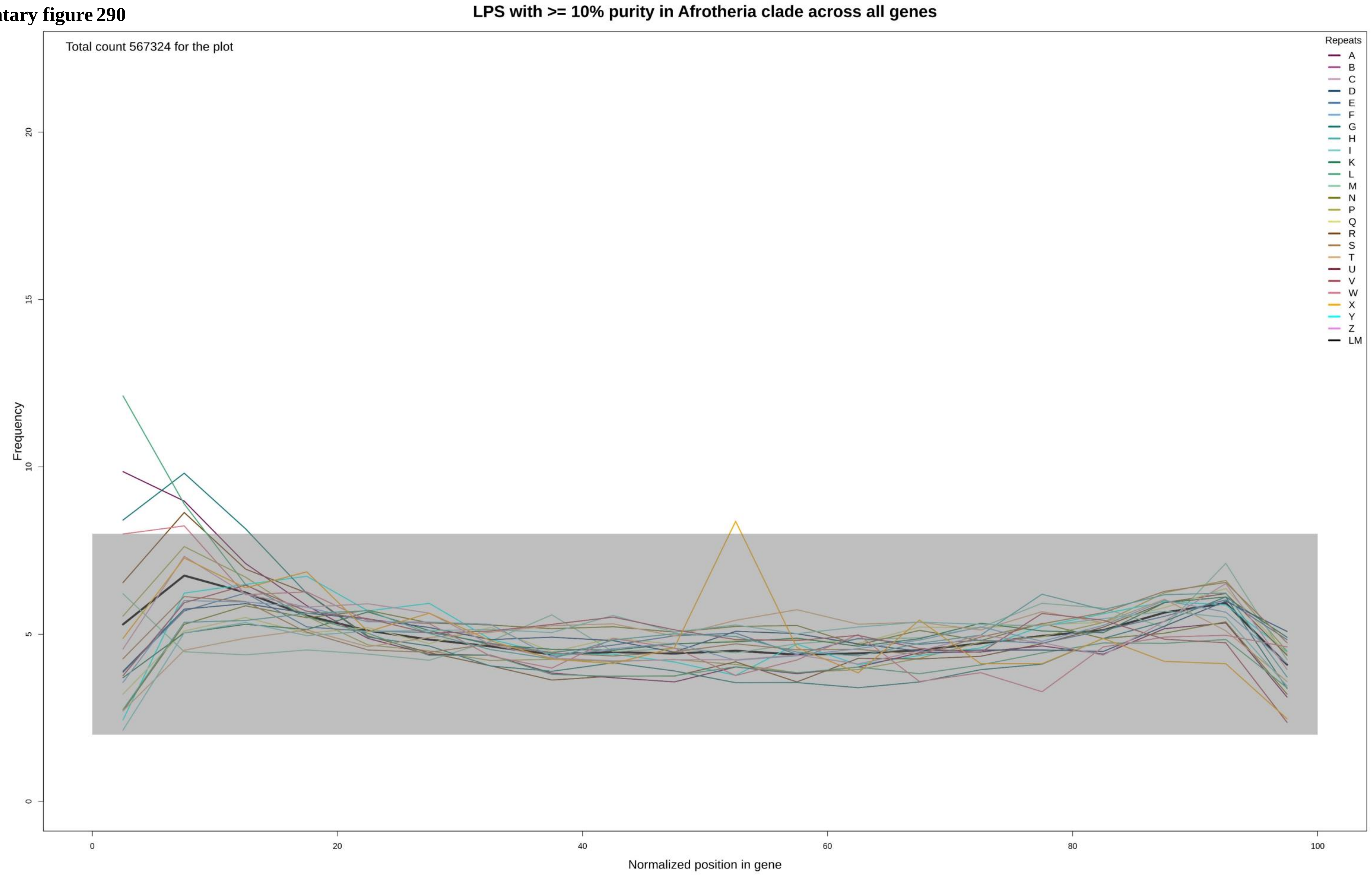

**Supplementary figure 291**

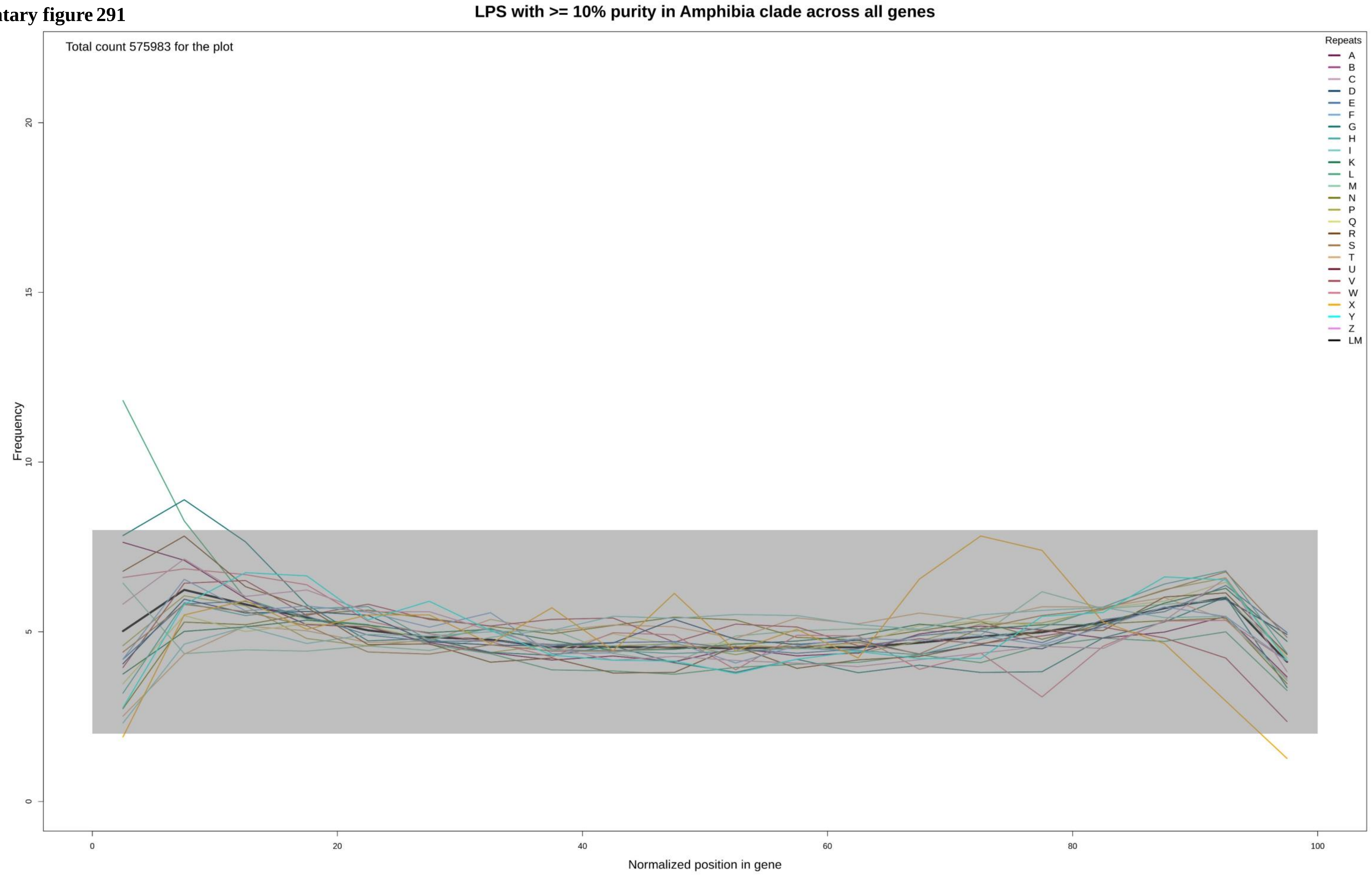

**Supplementary figure 292**

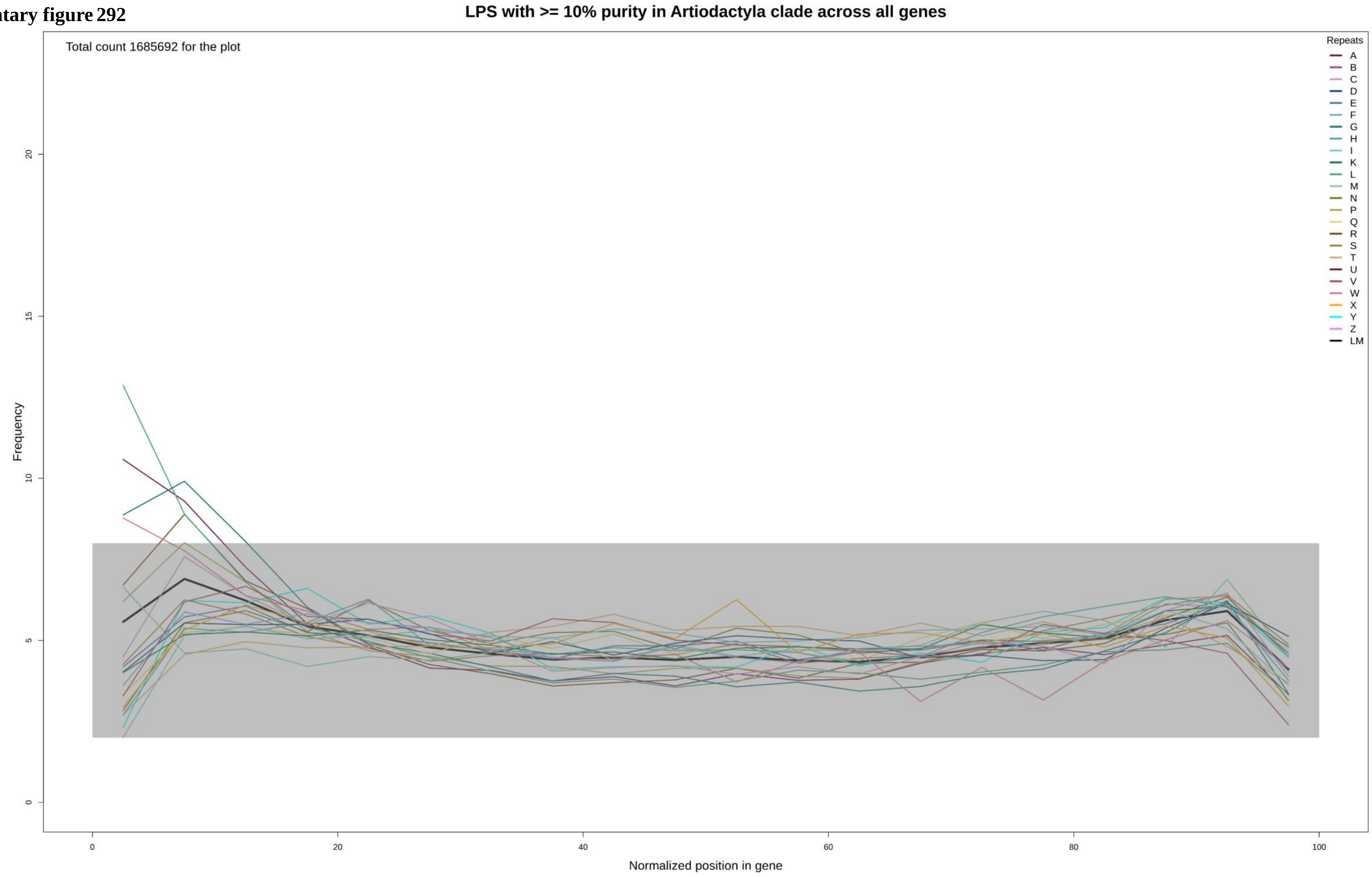

**Supplementary figure 293**

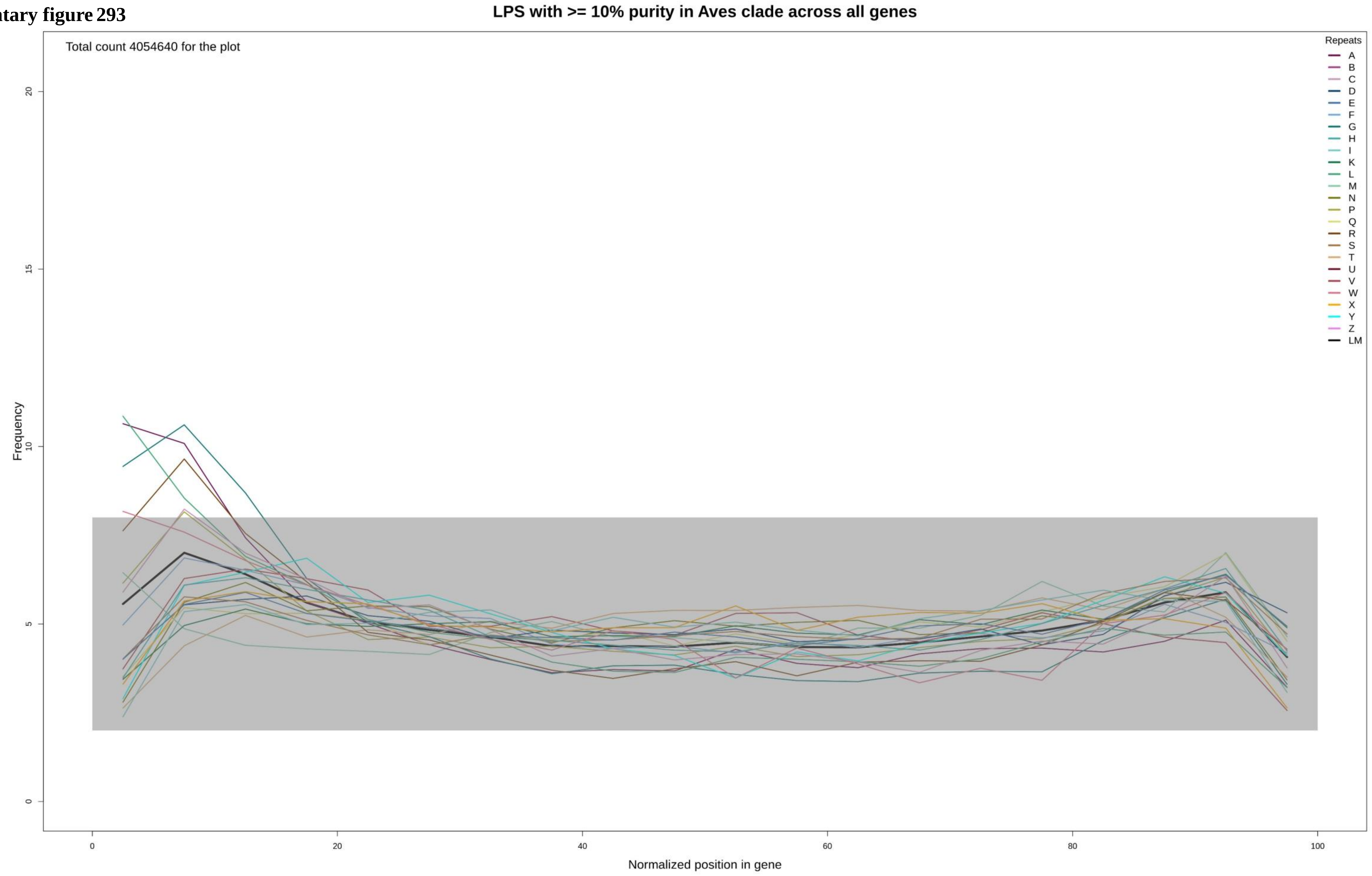

**Supplementary figure 294**

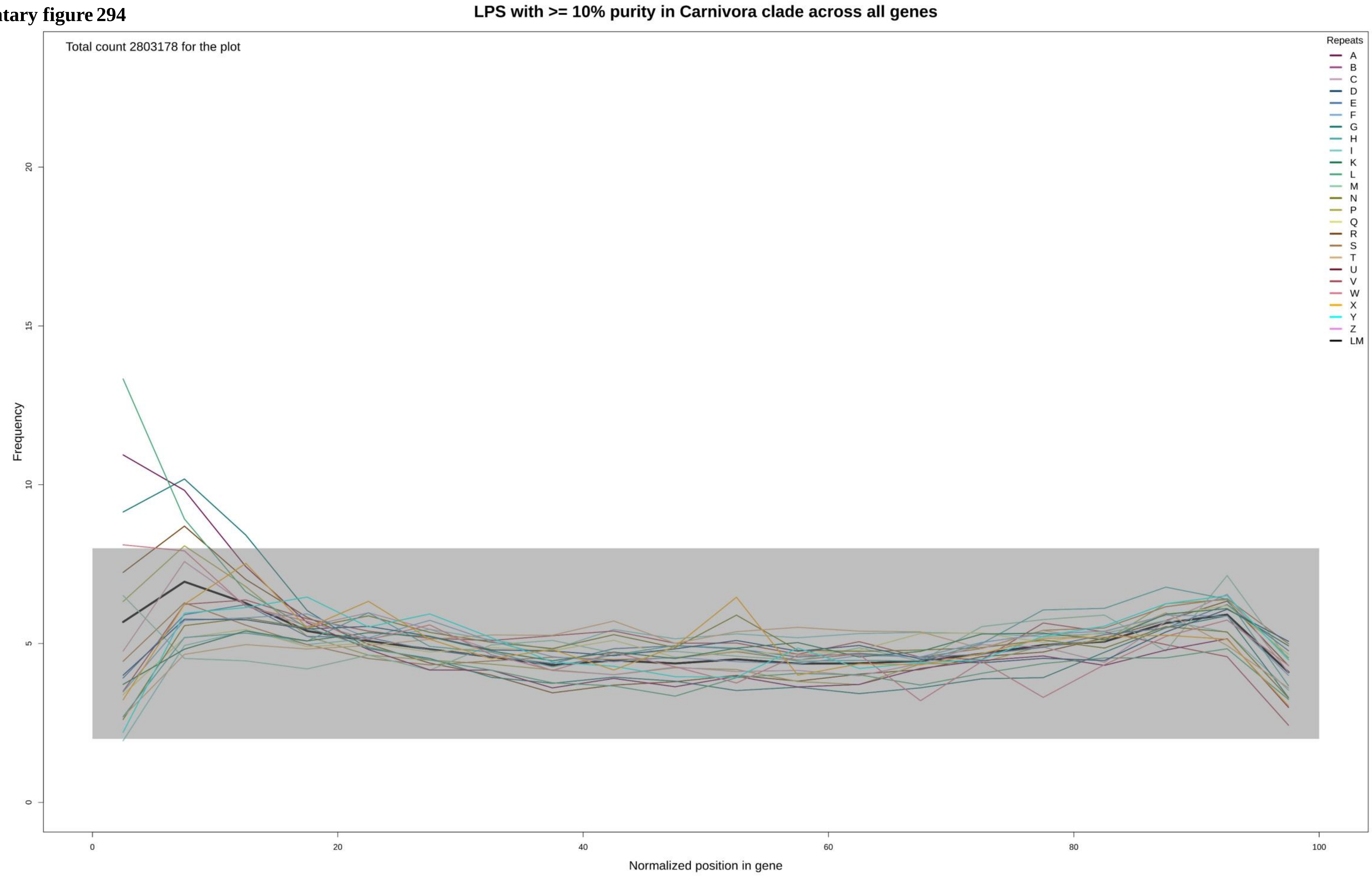

**Supplementary figure 295**

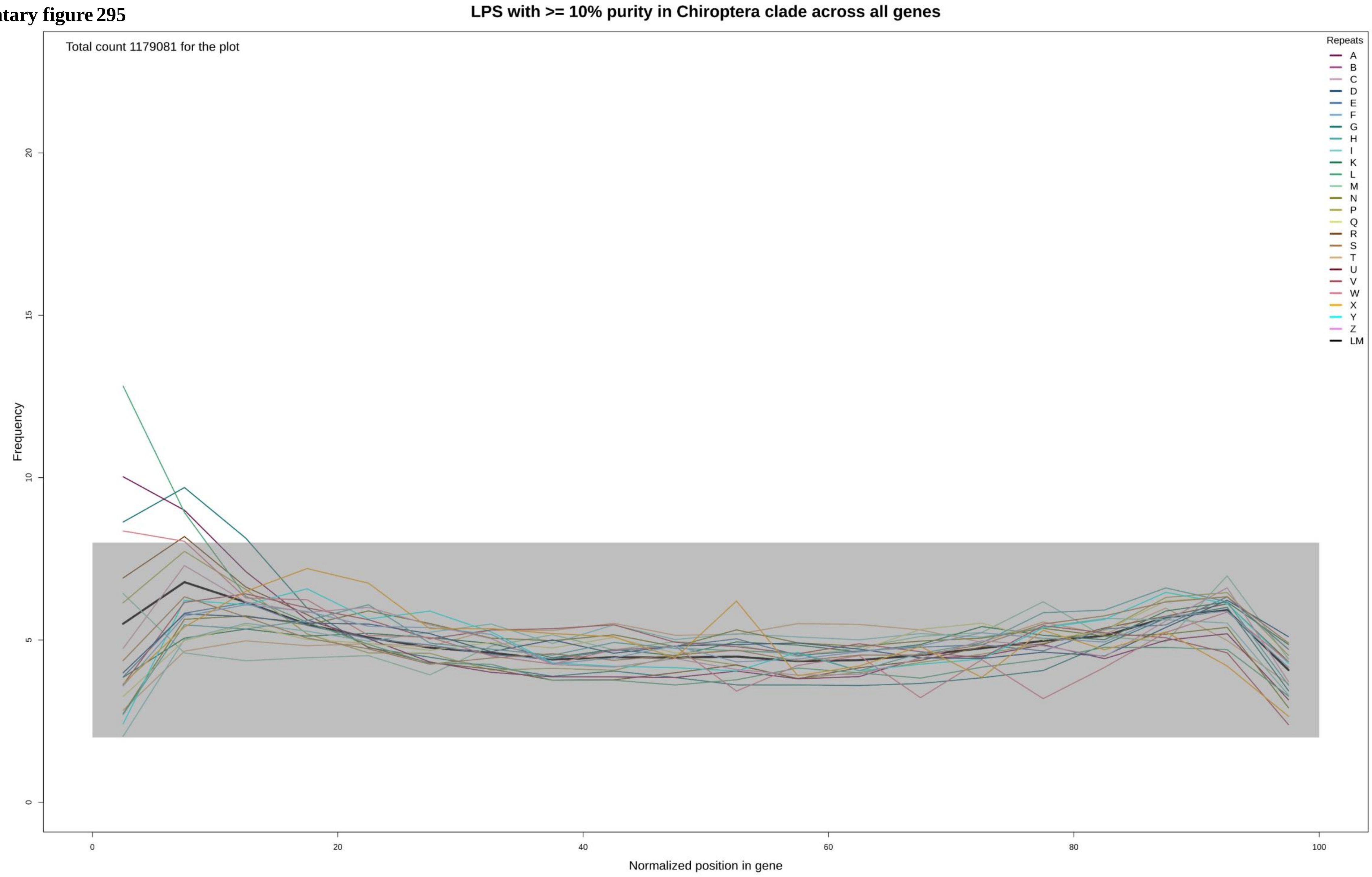

**Supplementary figure 296**

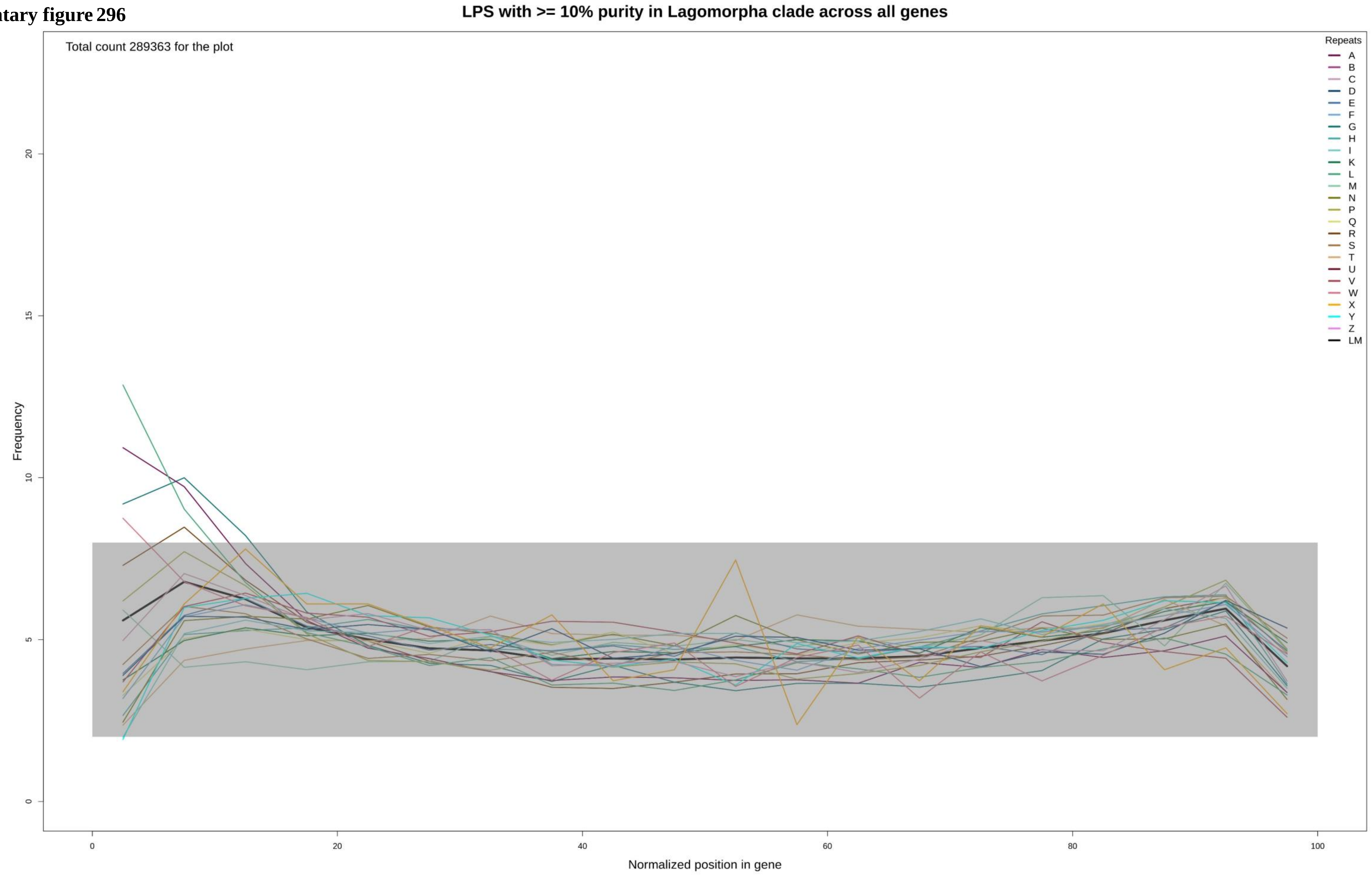

**Supplementary figure 297**

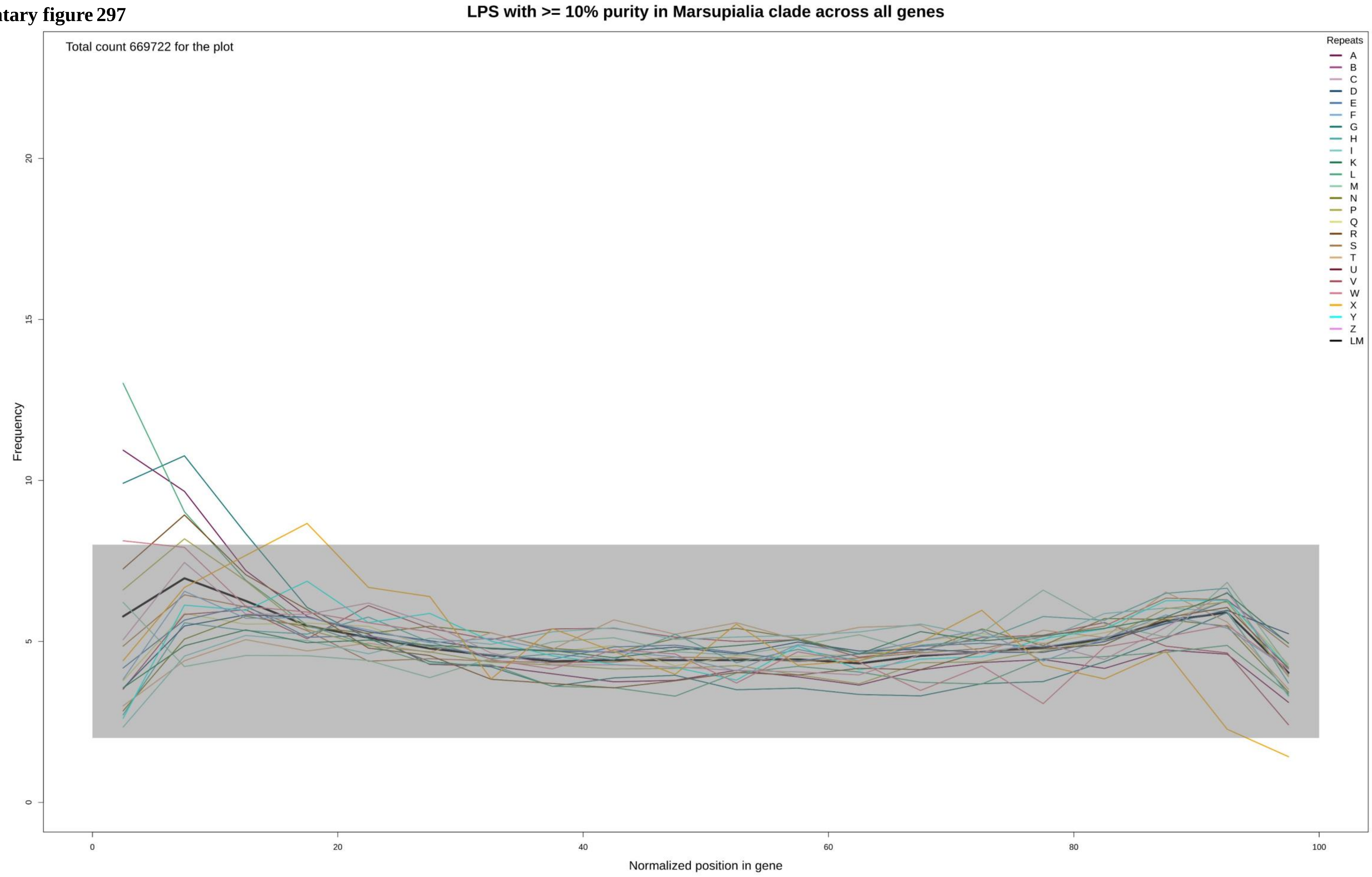

**Supplementary figure 298**

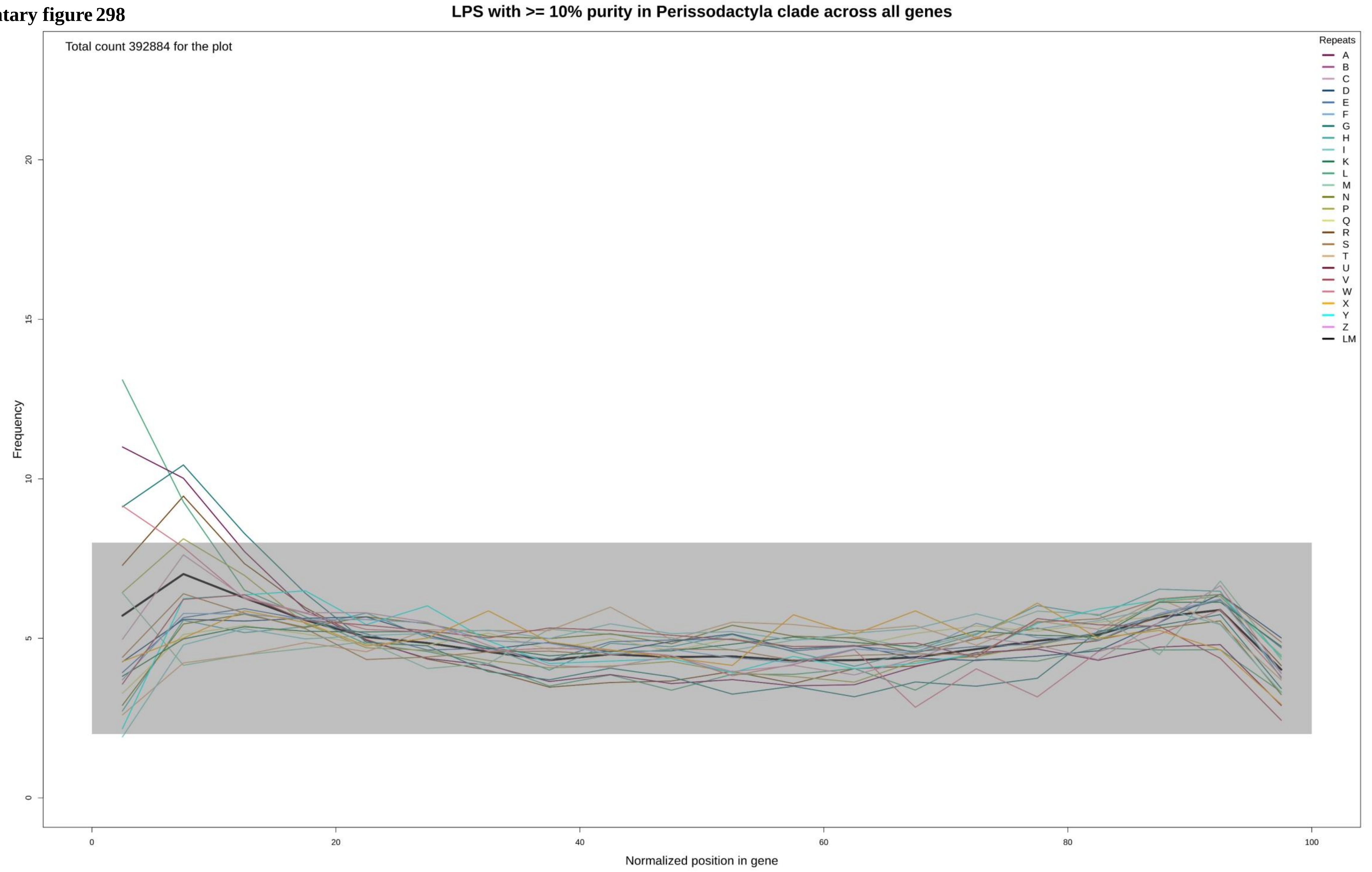

Supplementary figure 299

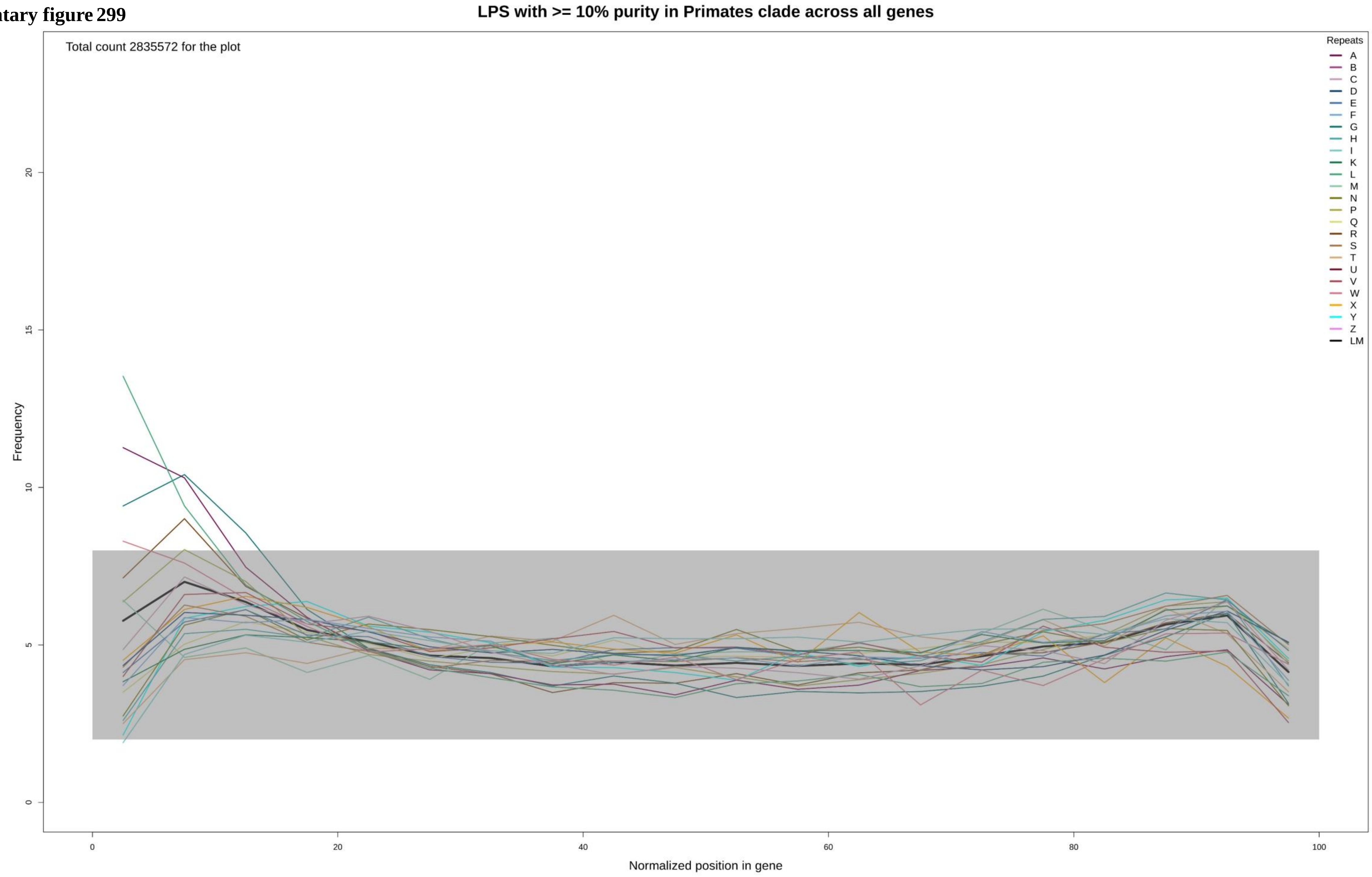

**Supplementary figure 300**

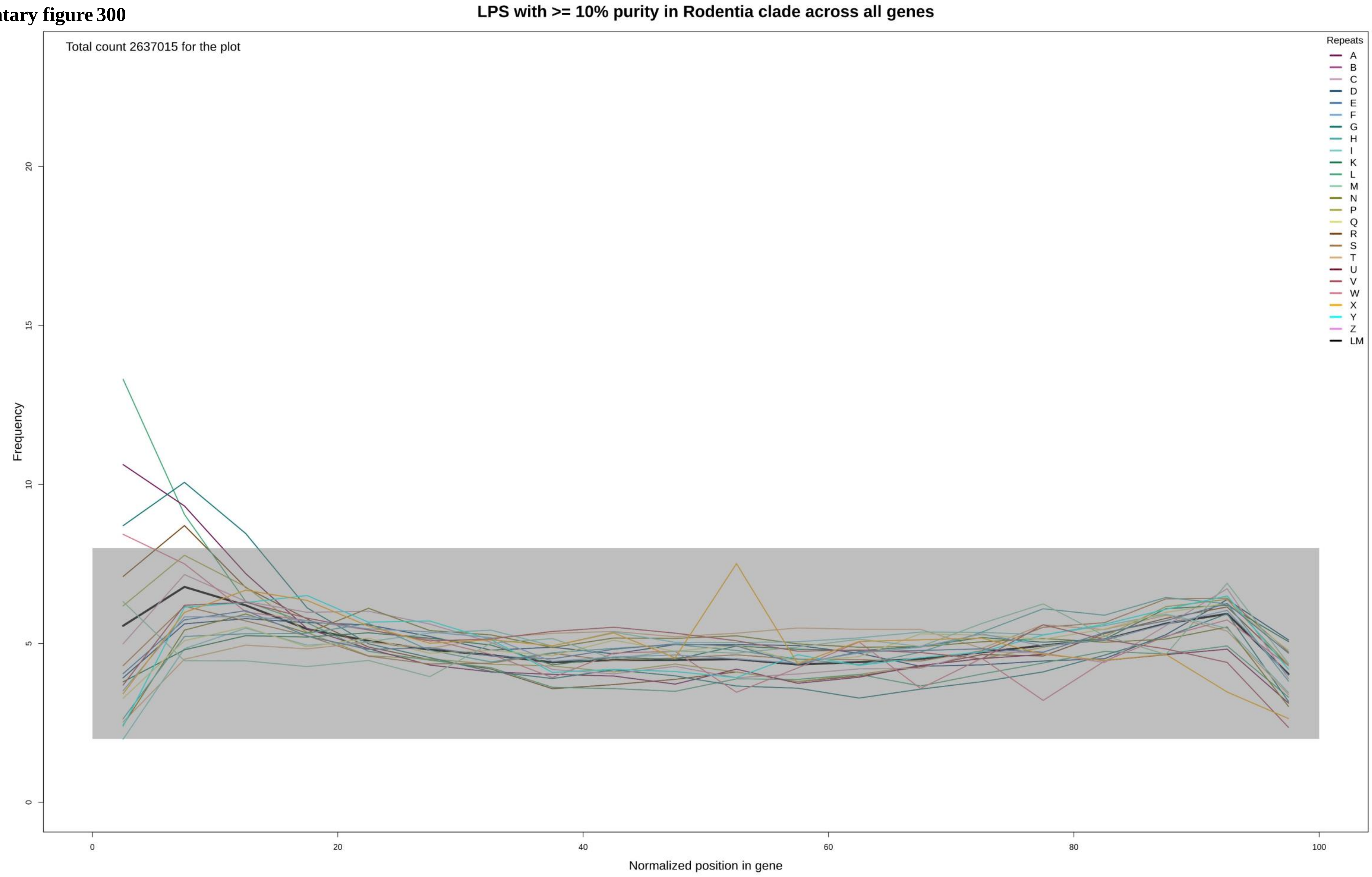

Supplementary figure 301

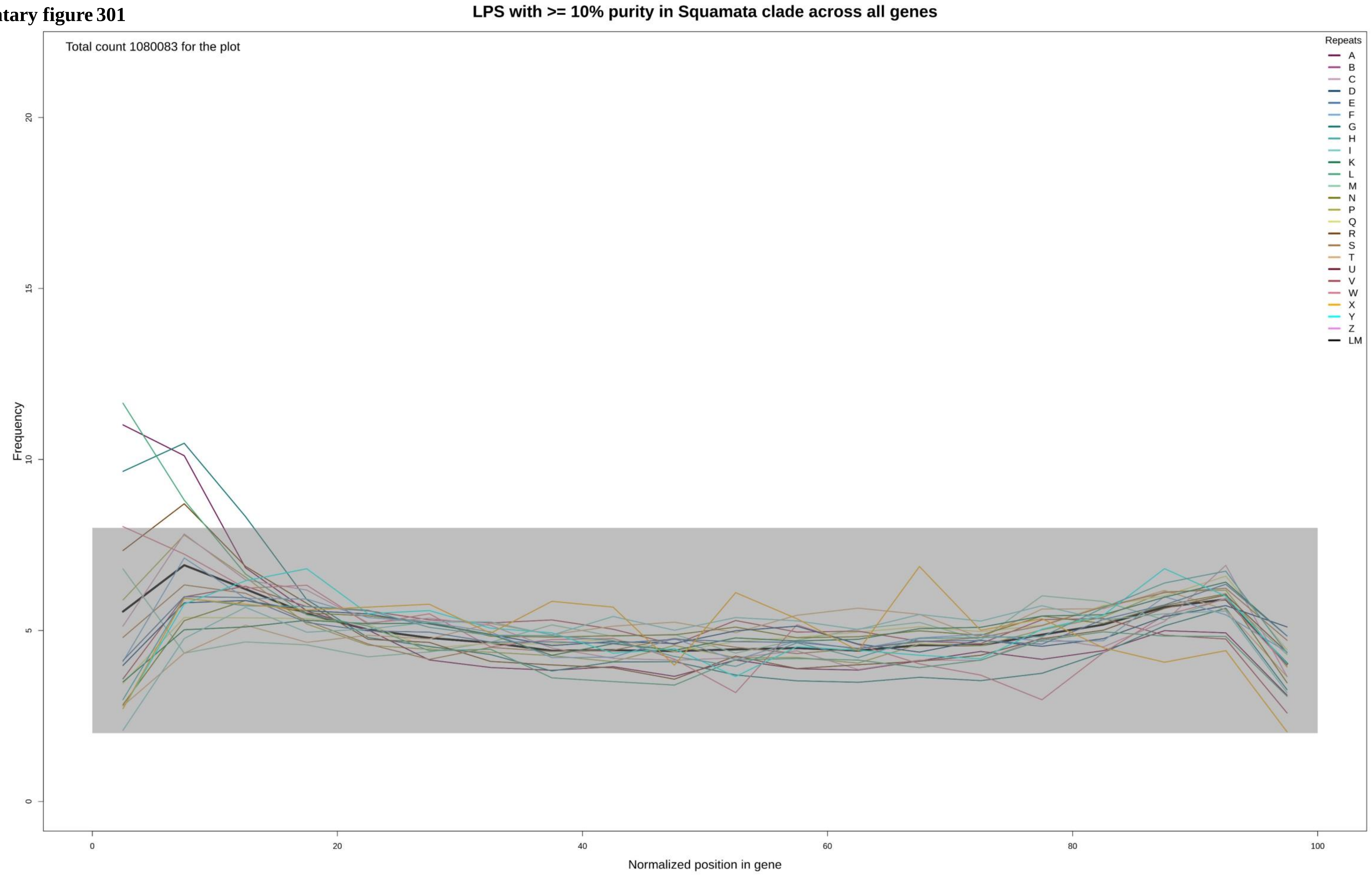

**Supplementary figure 302**

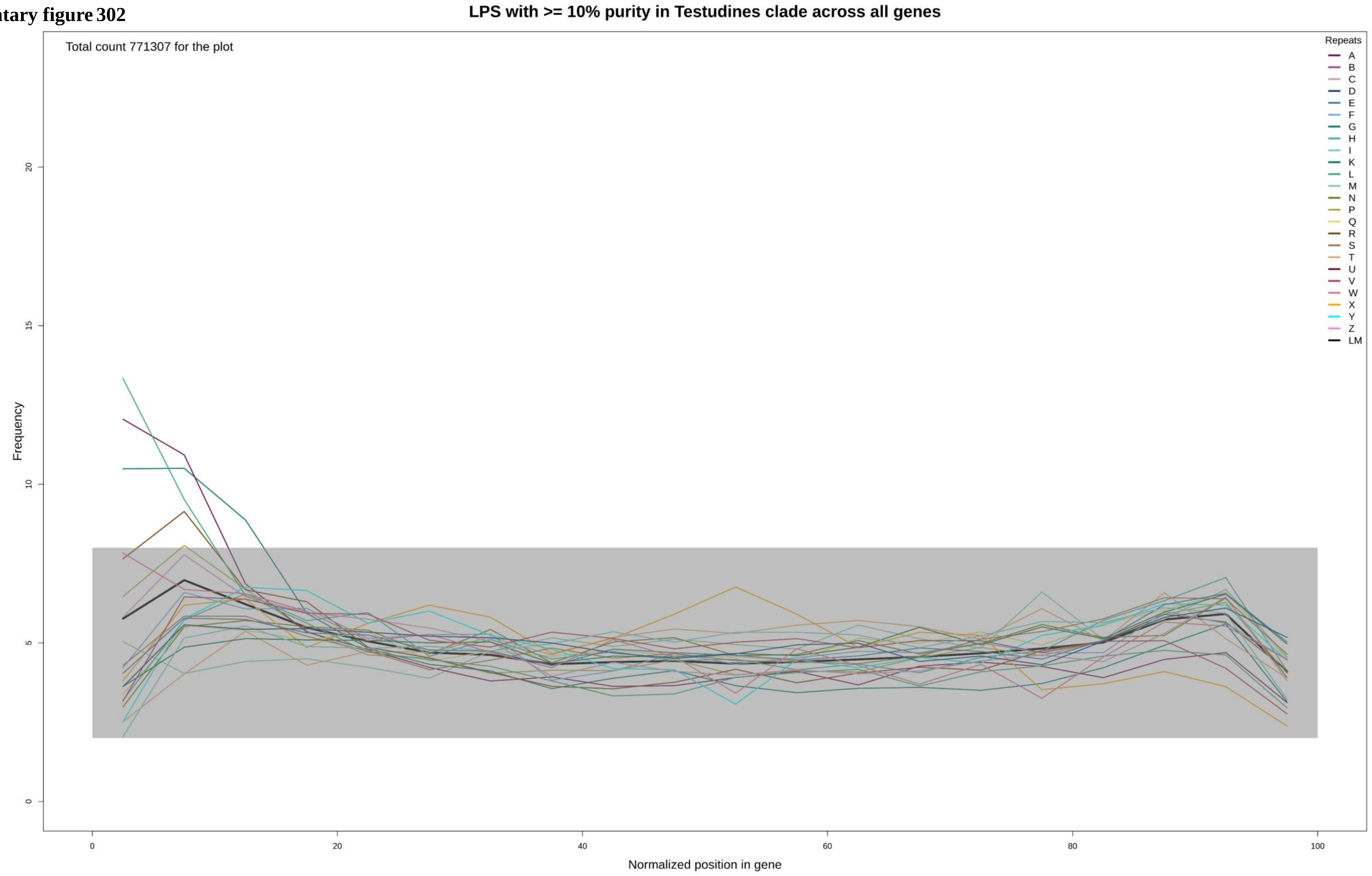

**Supplementary figure 303**

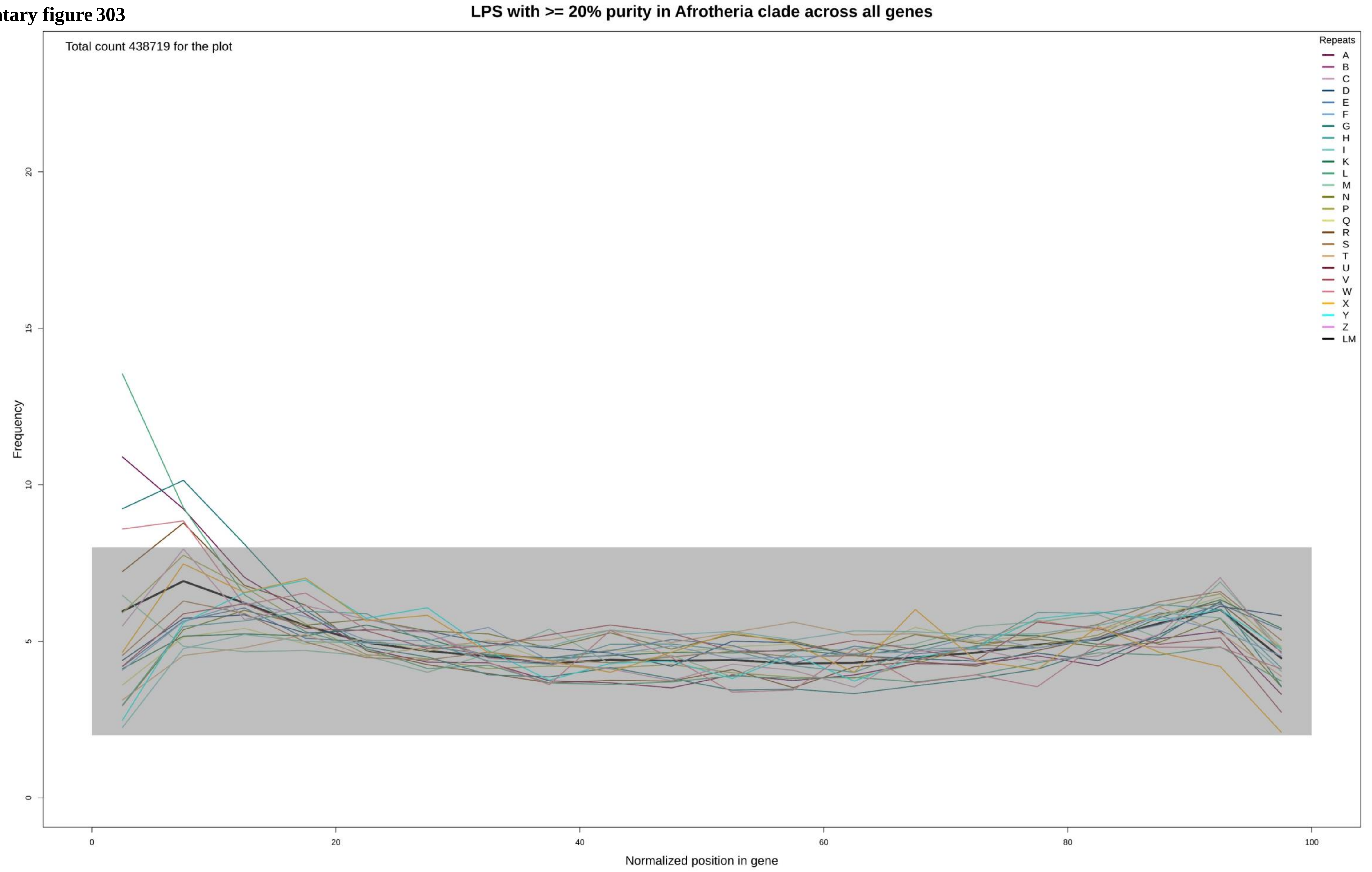

**Supplementary figure 304**

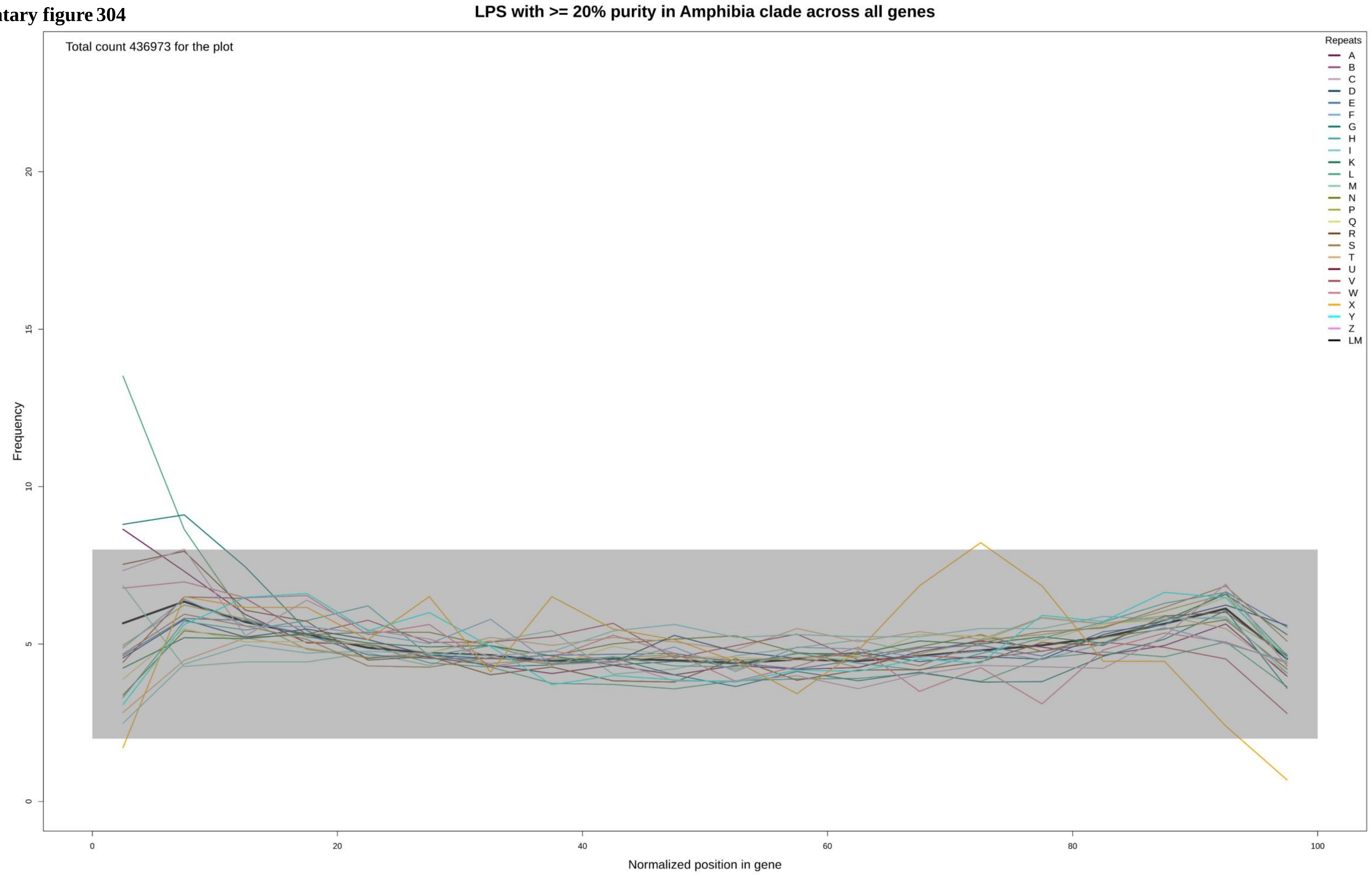

Supplementary figure 305

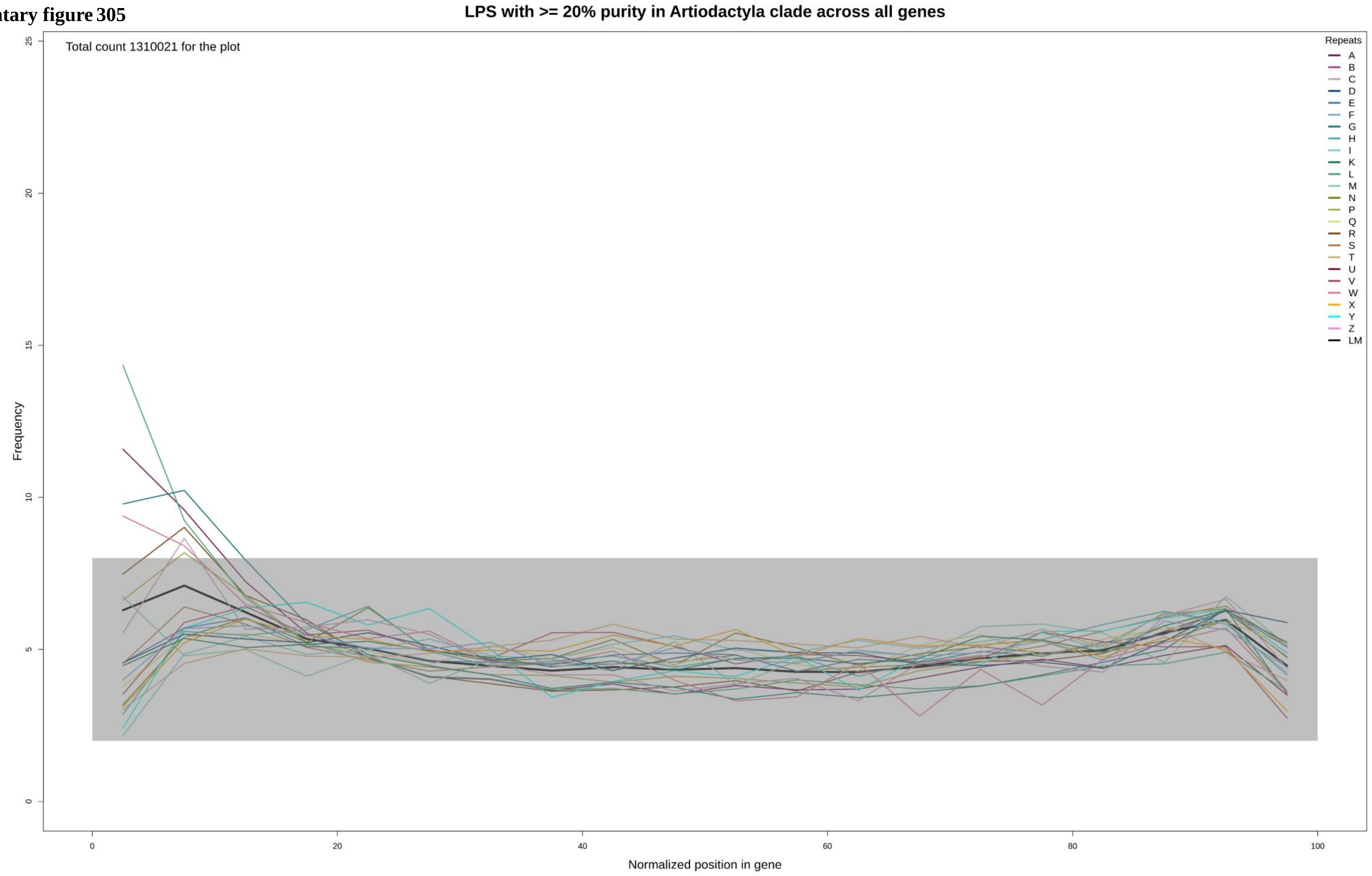

Supplementary figure 306

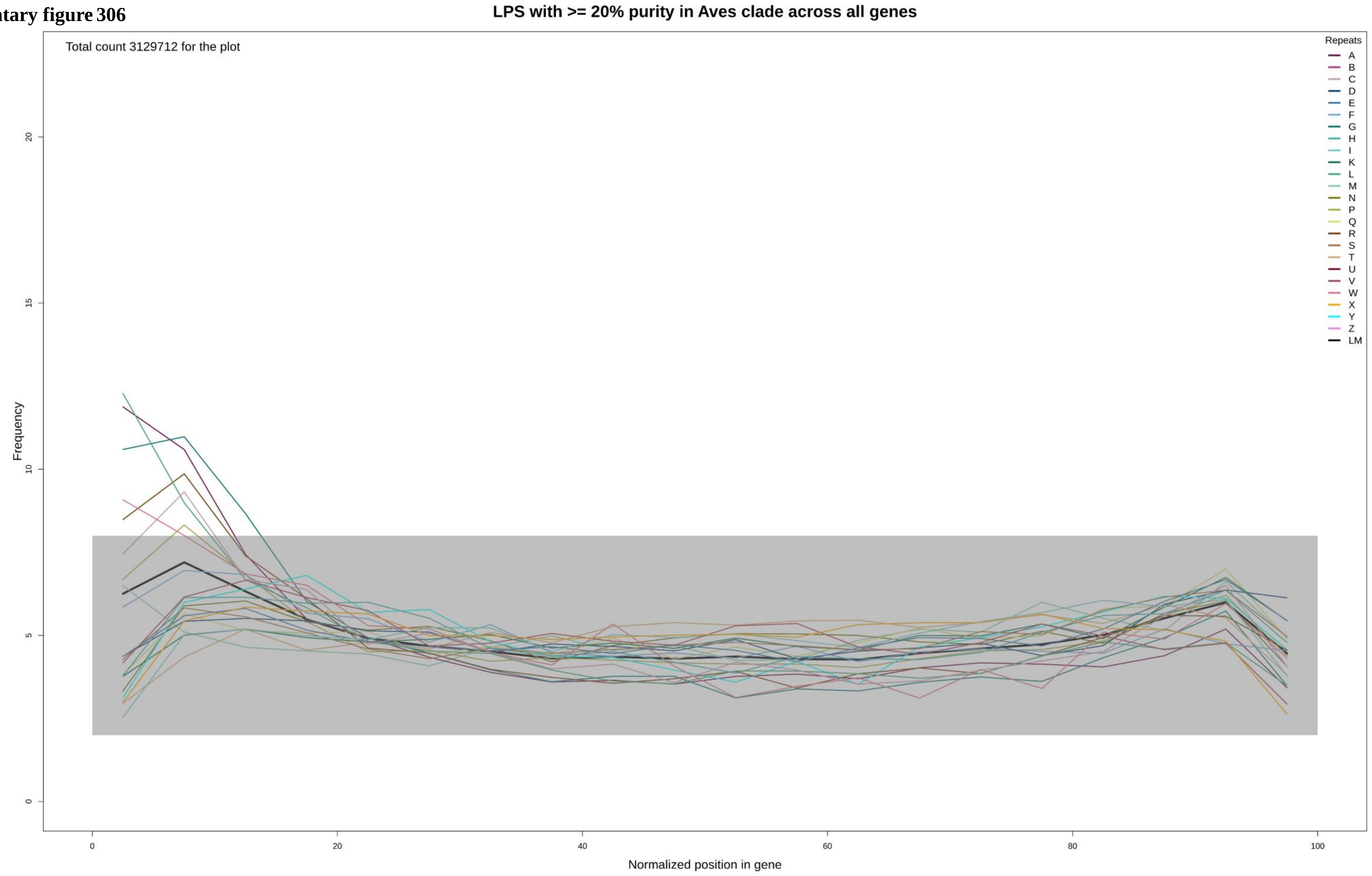

**Supplementary figure 307**

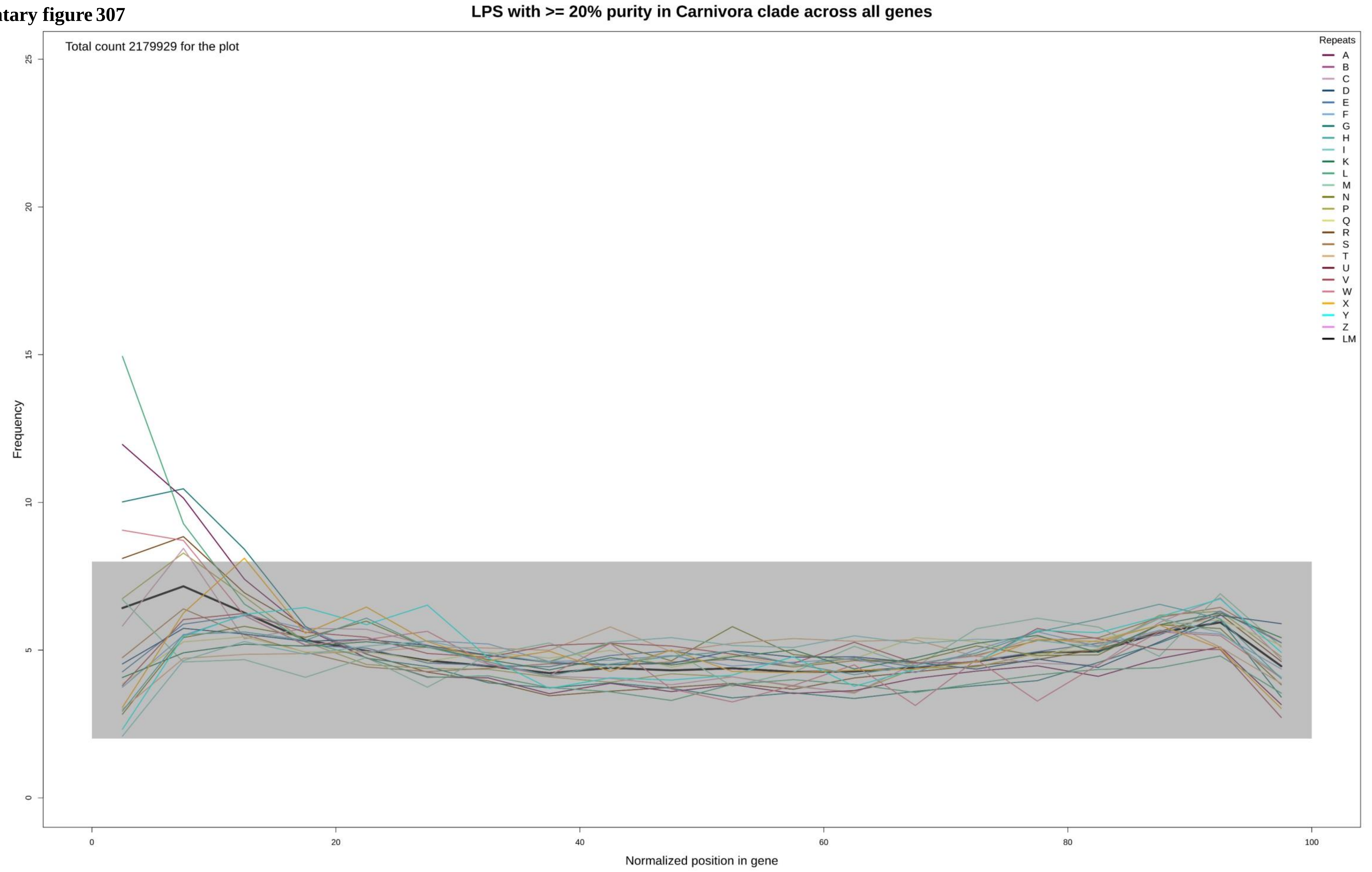

**Supplementary figure 308**

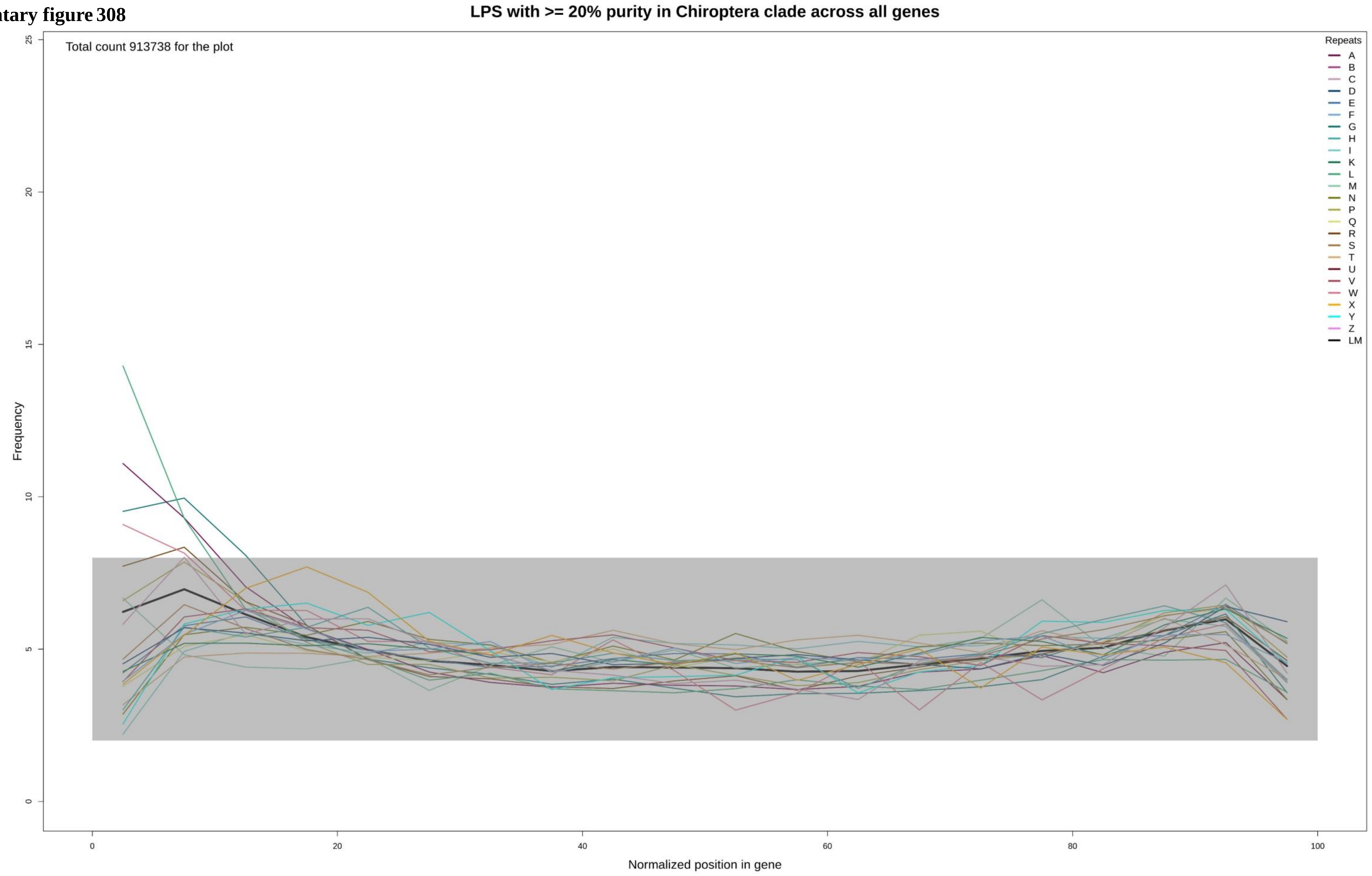

**Supplementary figure 309**

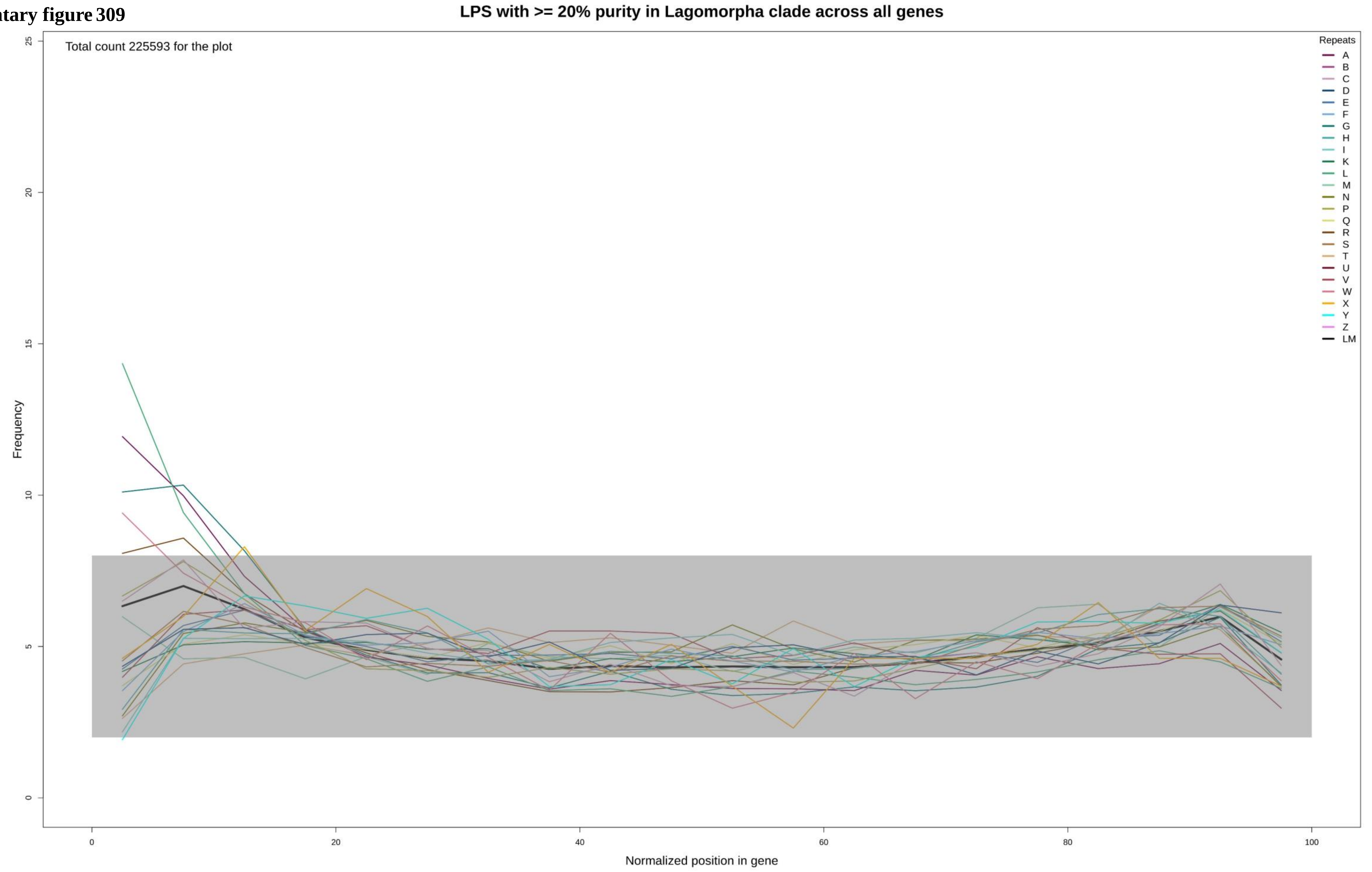

**Supplementary figure 310**

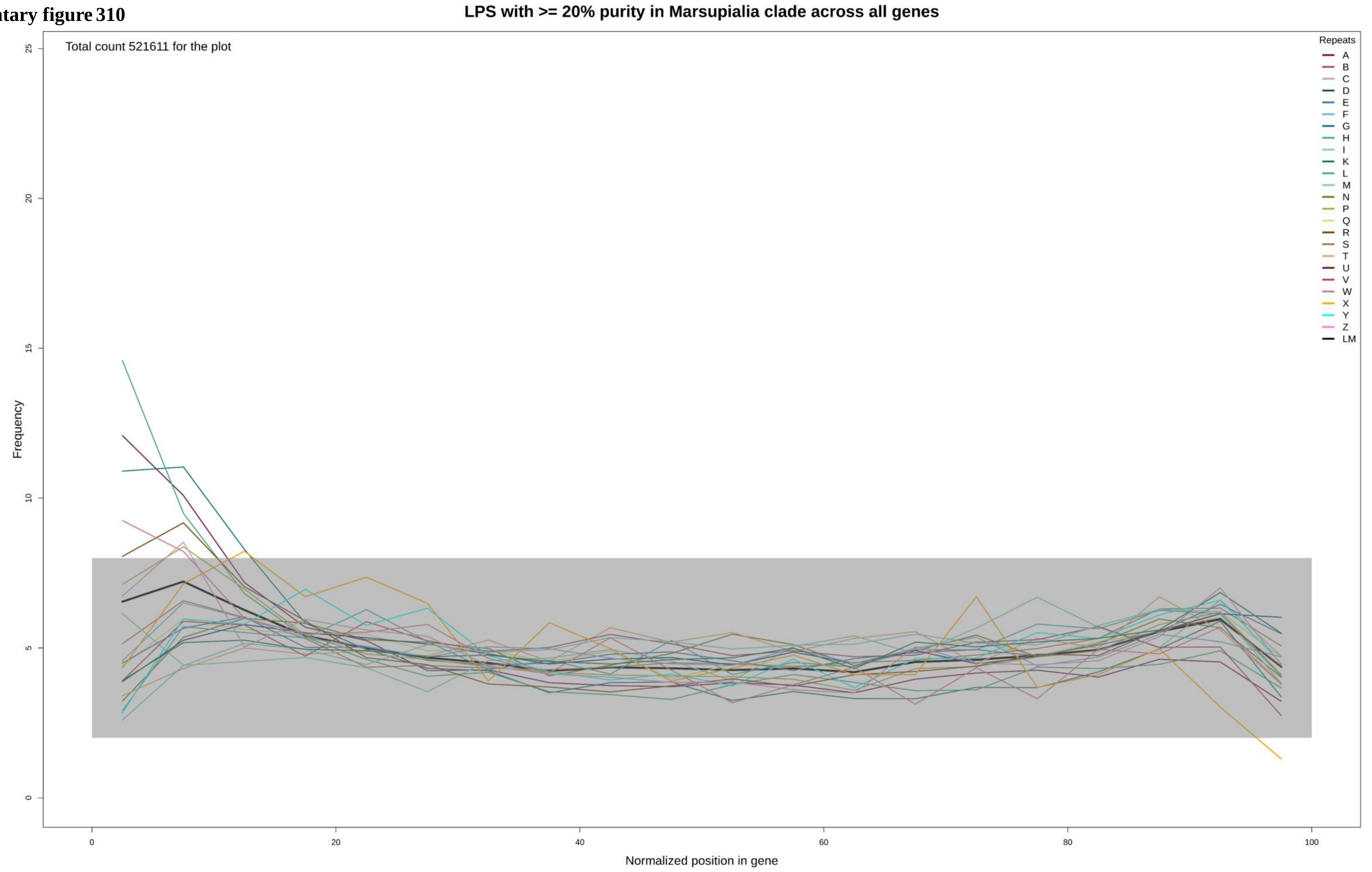

**Supplementary figure 311**

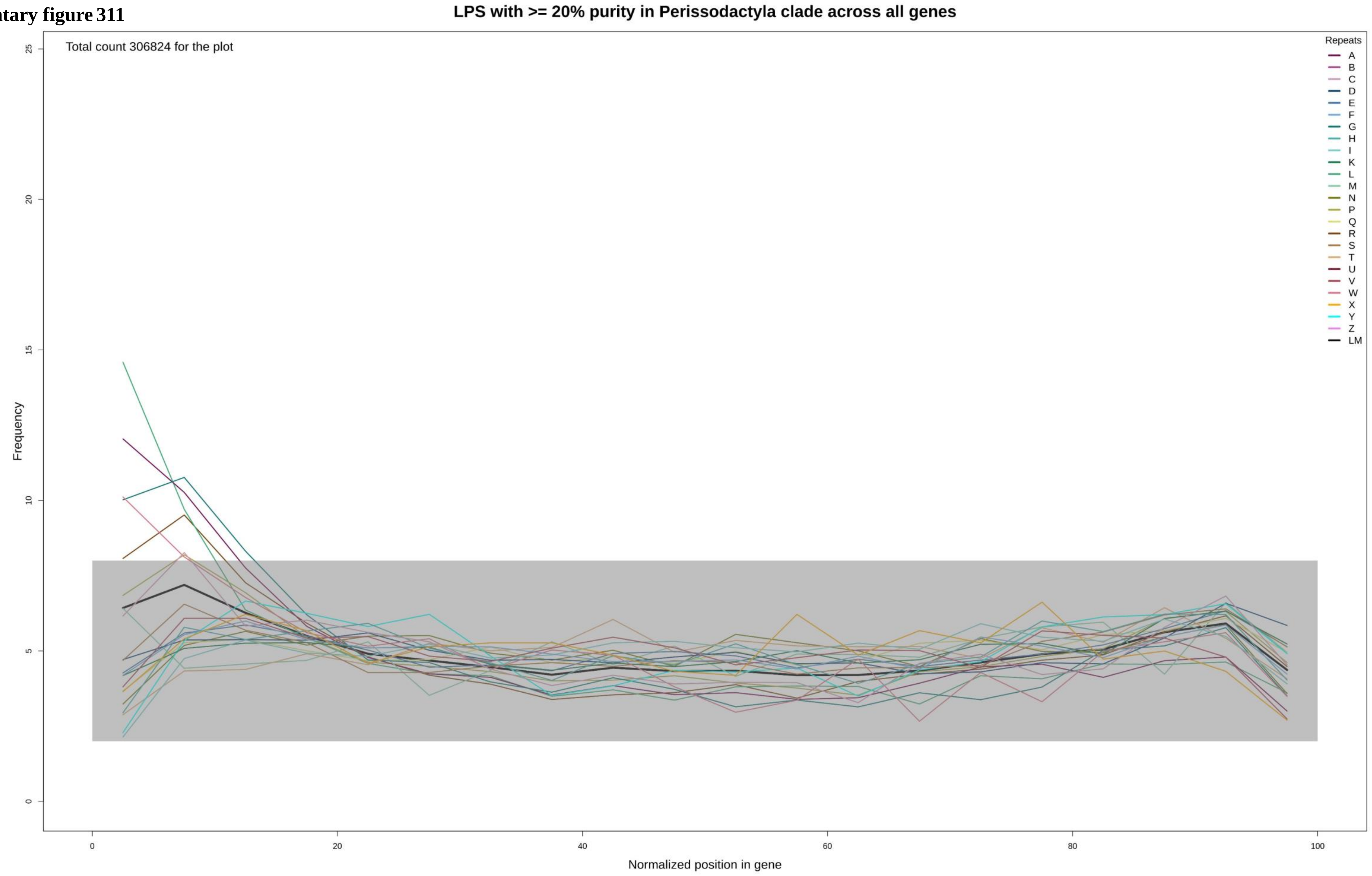

**Supplementary figure 312**

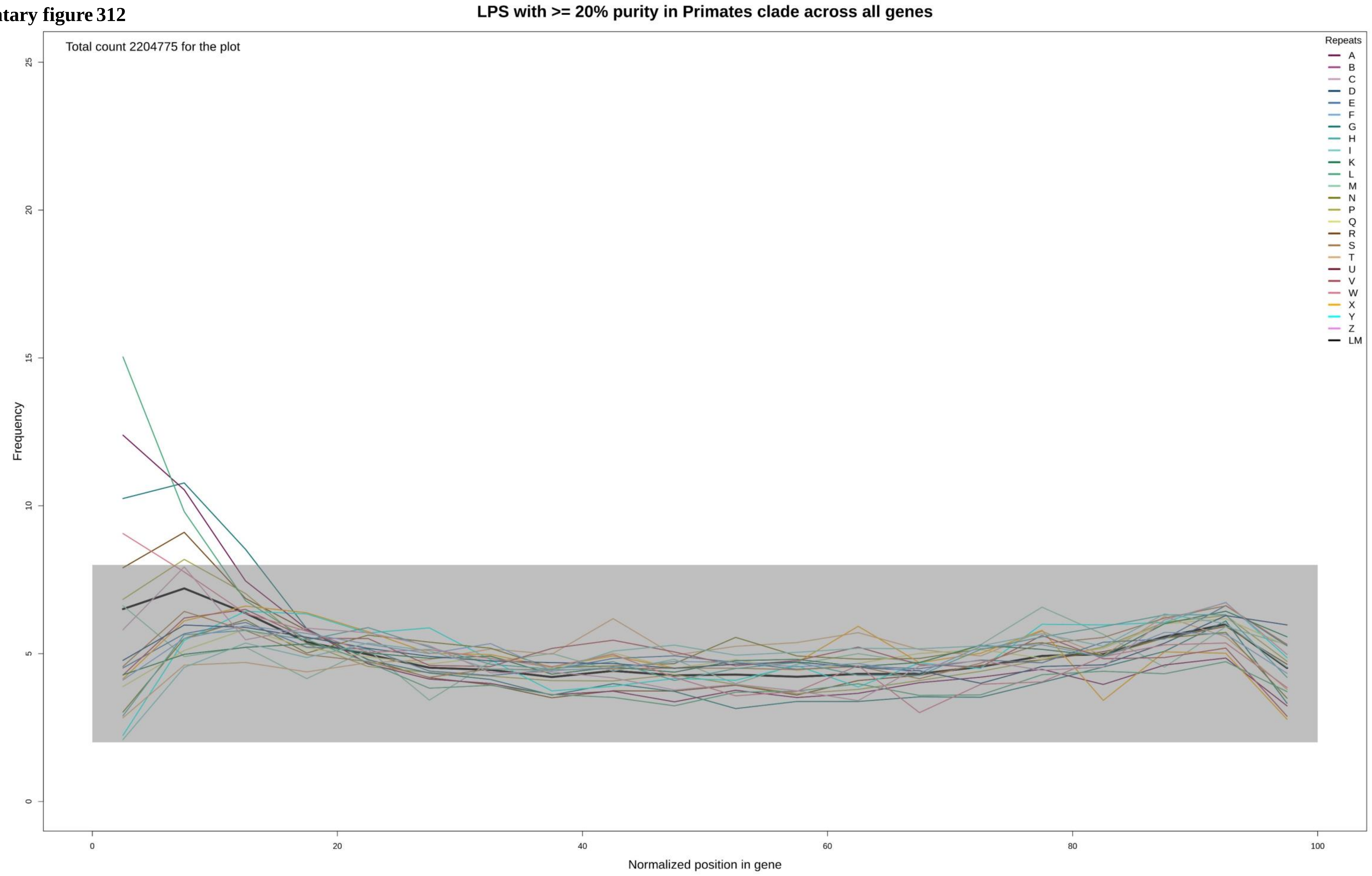

**Supplementary figure 313**

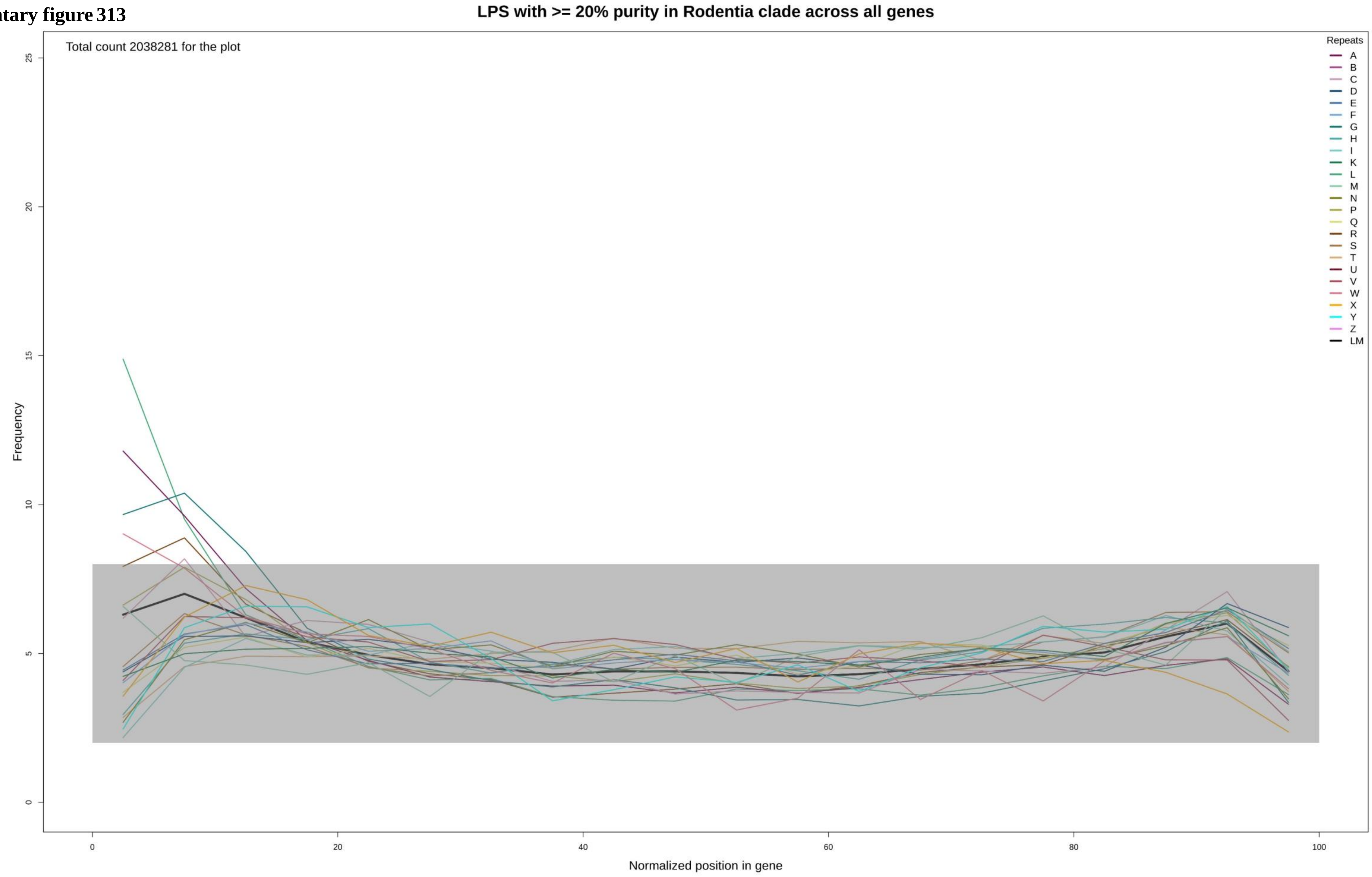

**Supplementary figure 314**

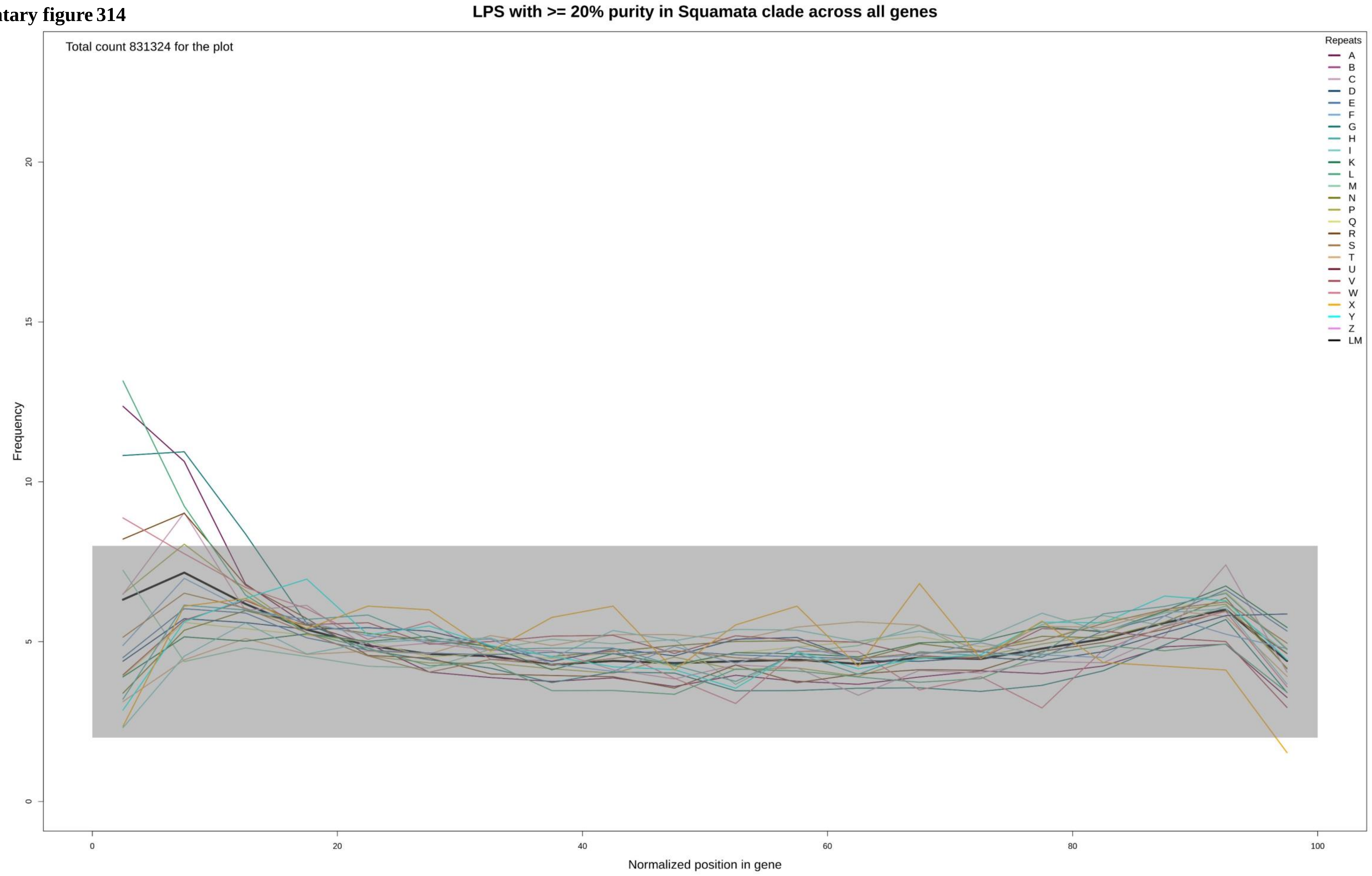

Supplementary figure 315

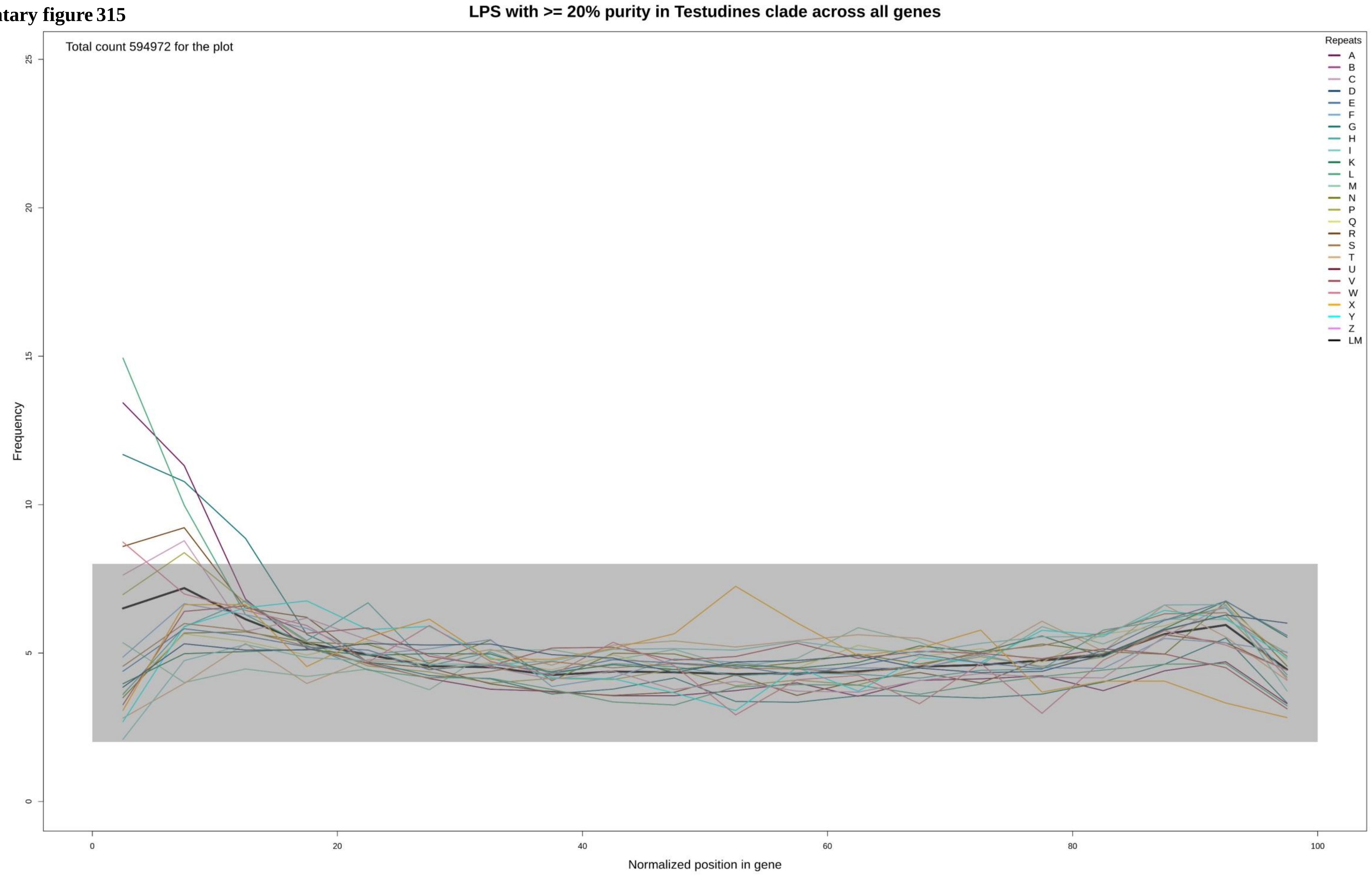

**Supplementary figure 316**

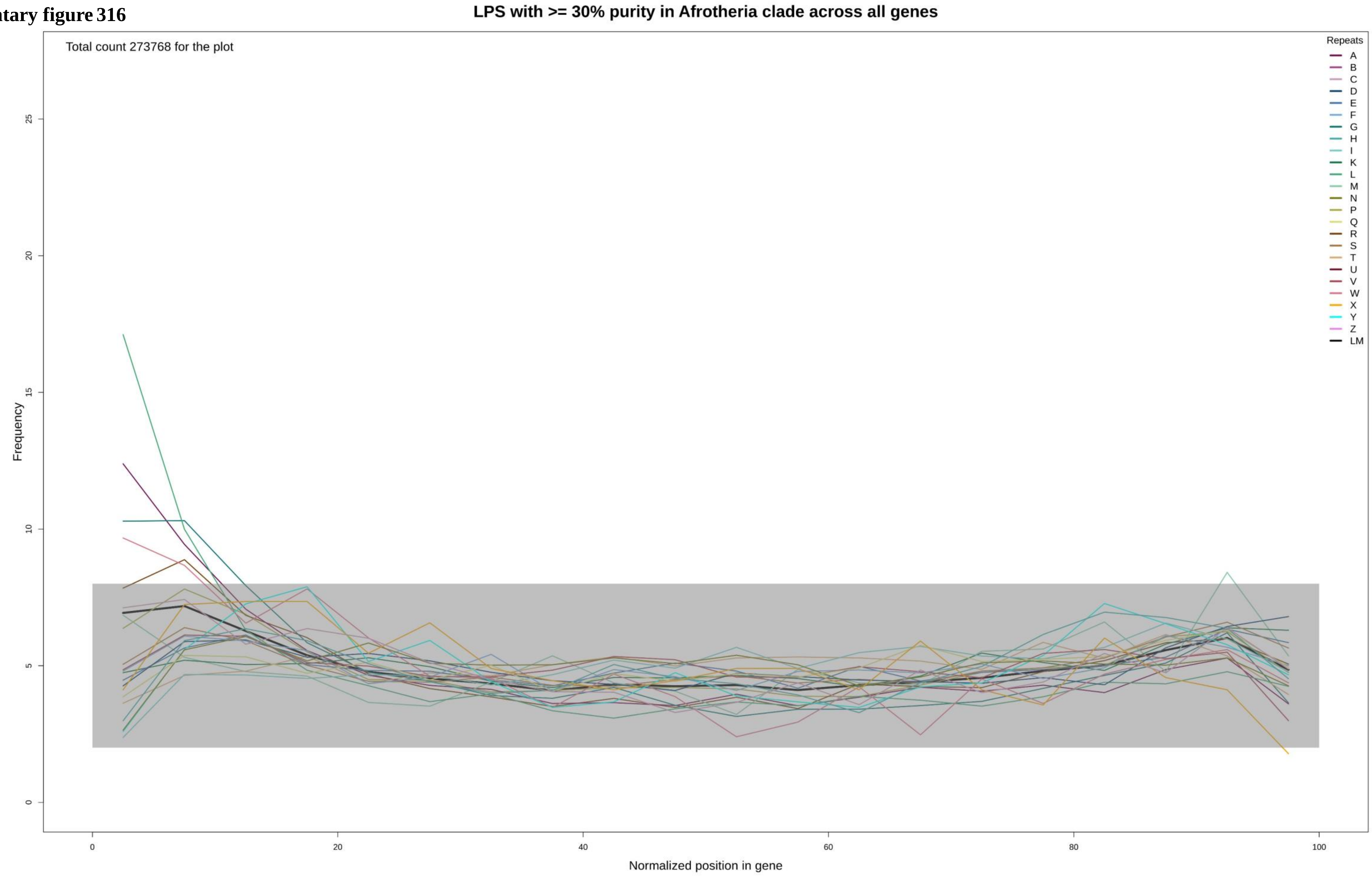

Supplementary figure 317

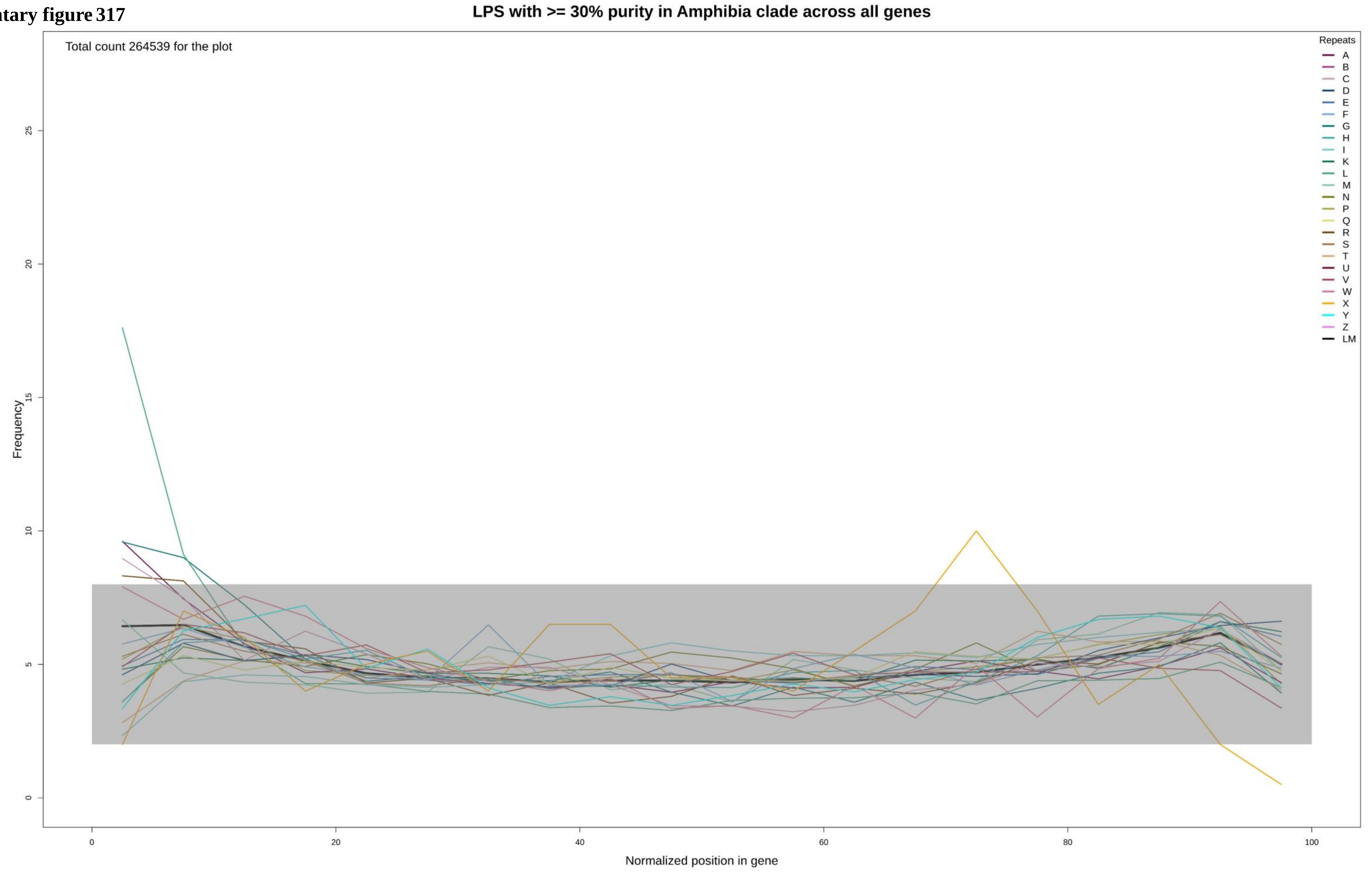

Supplementary figure 318

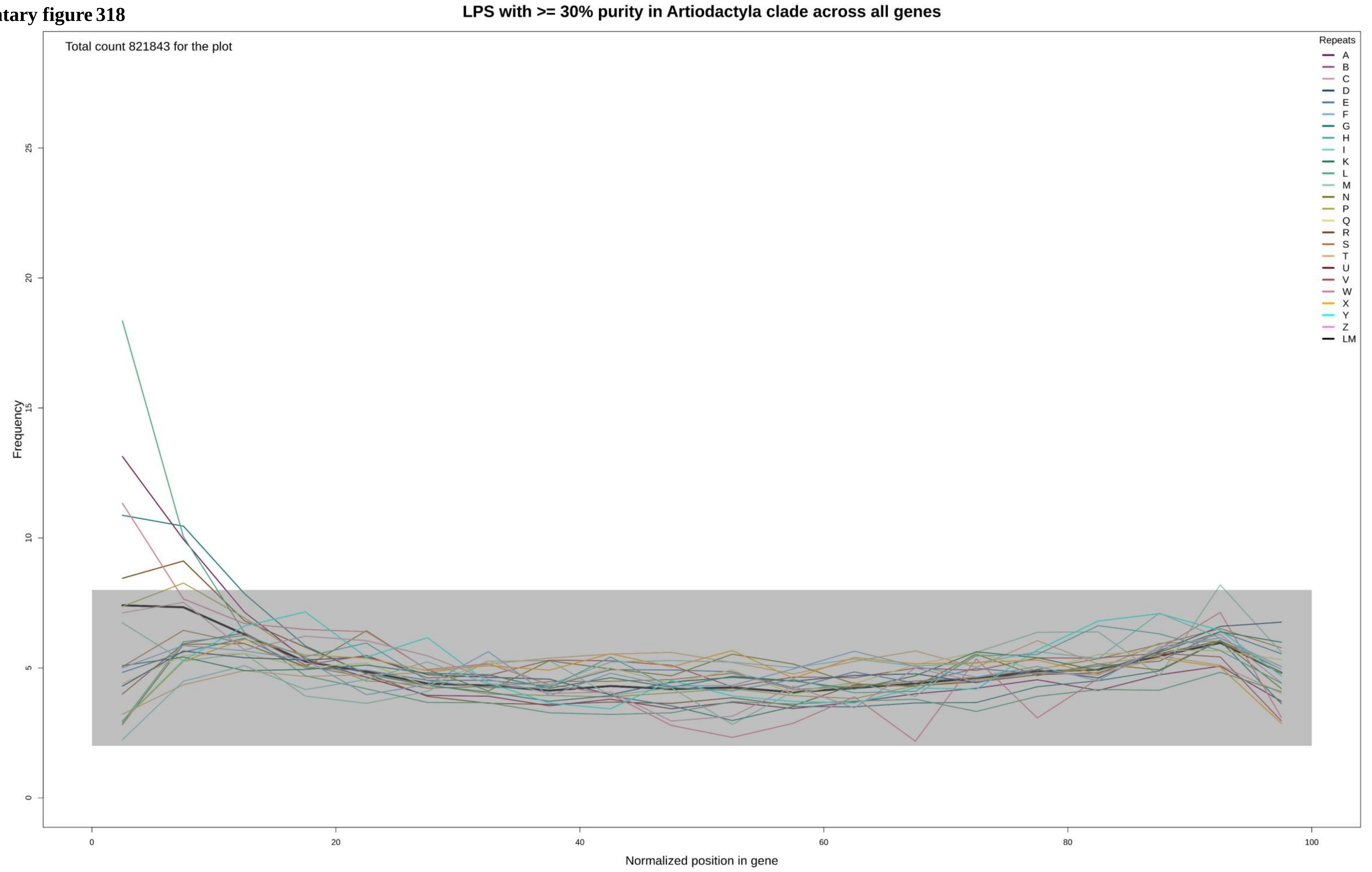

**Supplementary figure 319**

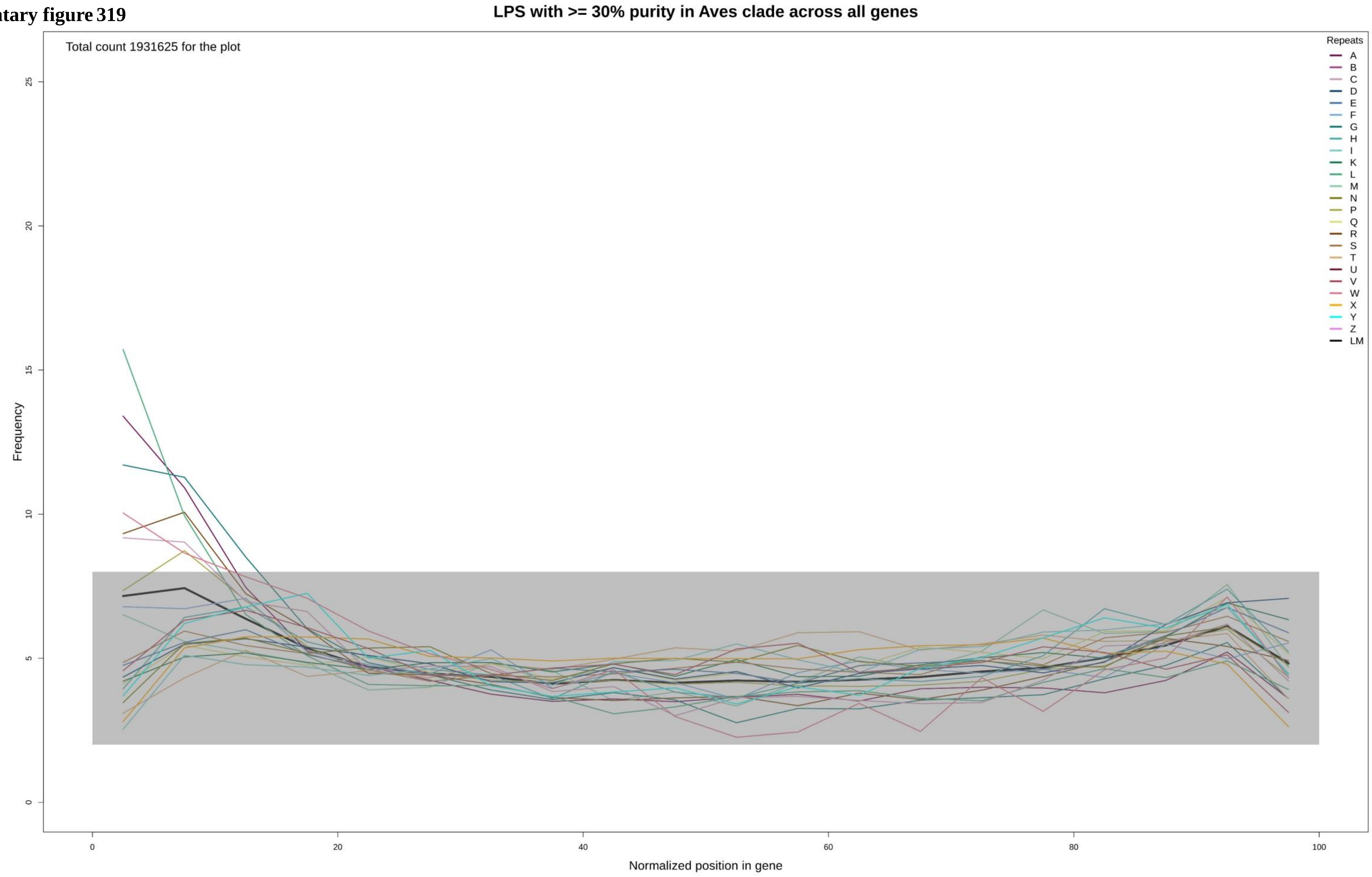

Supplementary figure 320

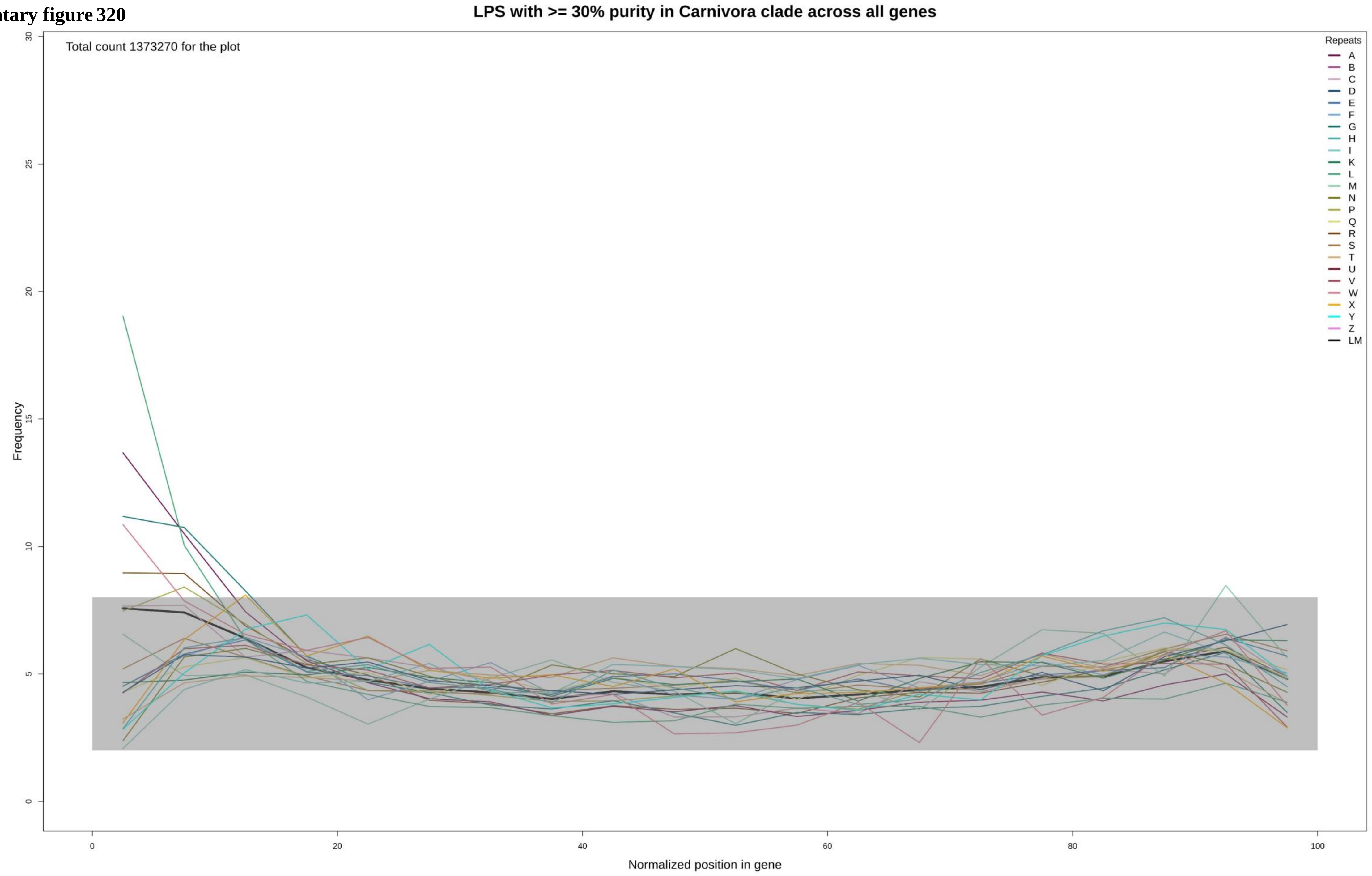

**Supplementary figure 321**

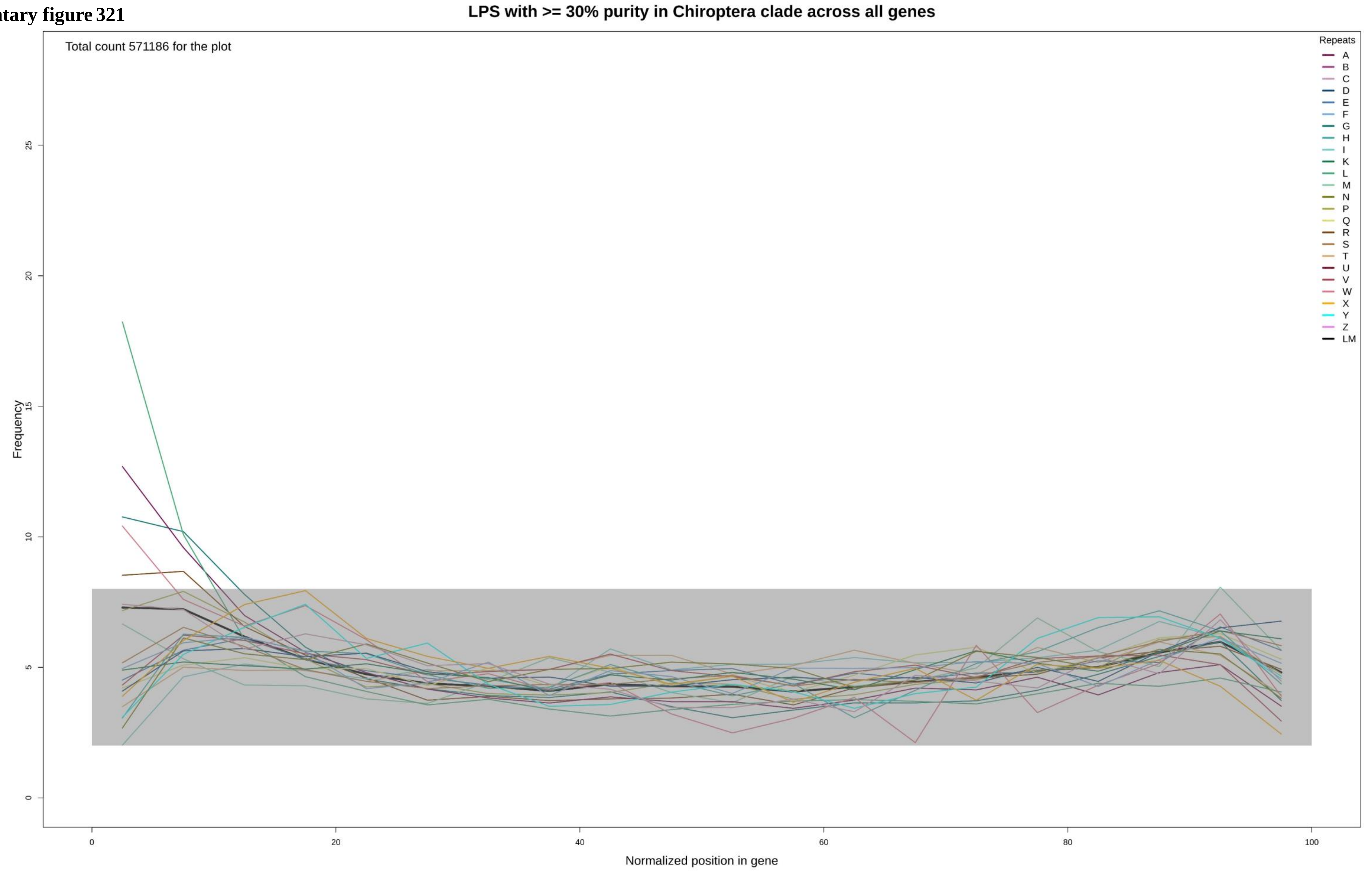

Supplementary figure 322

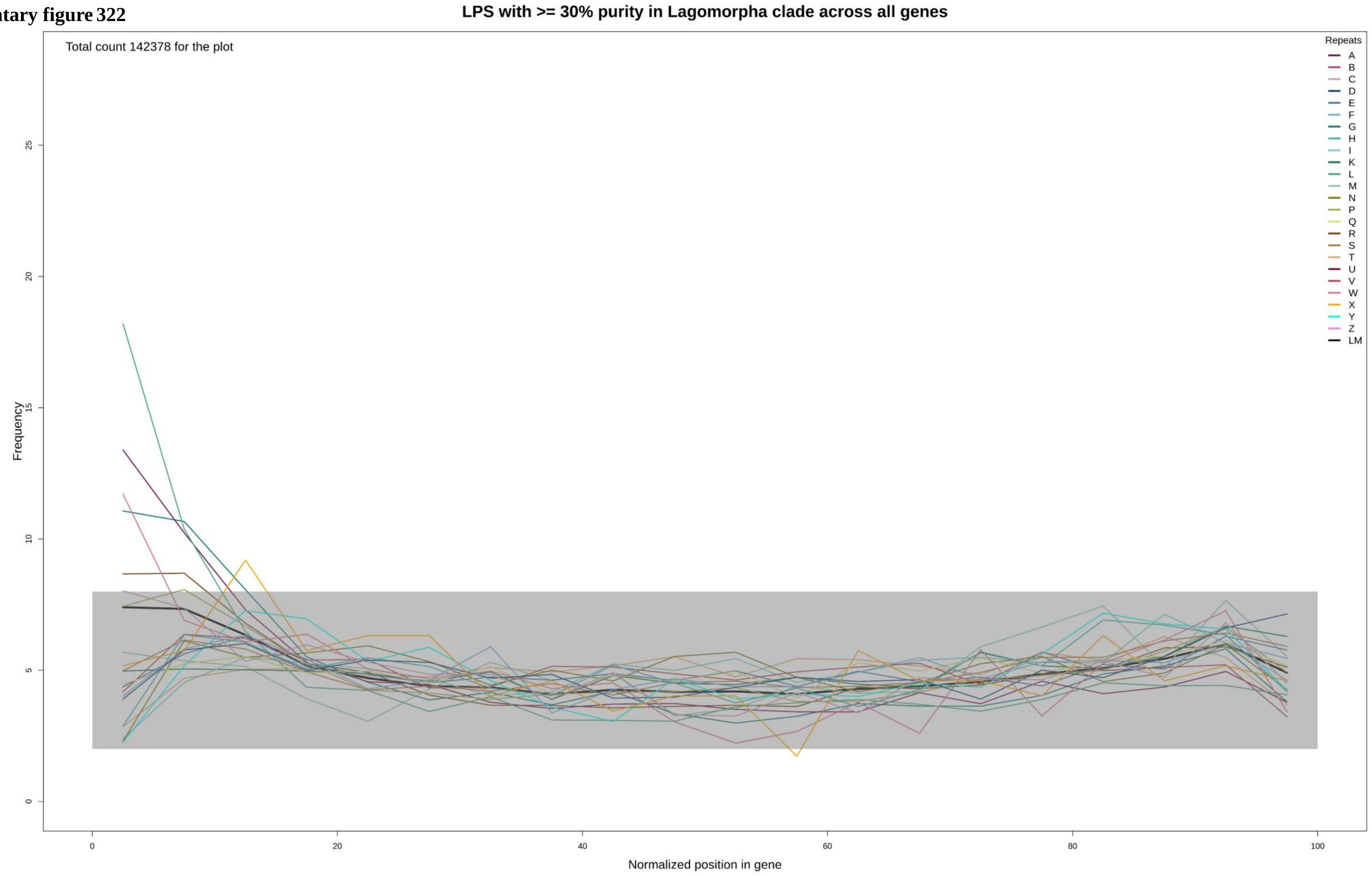

**Supplementary figure 323**

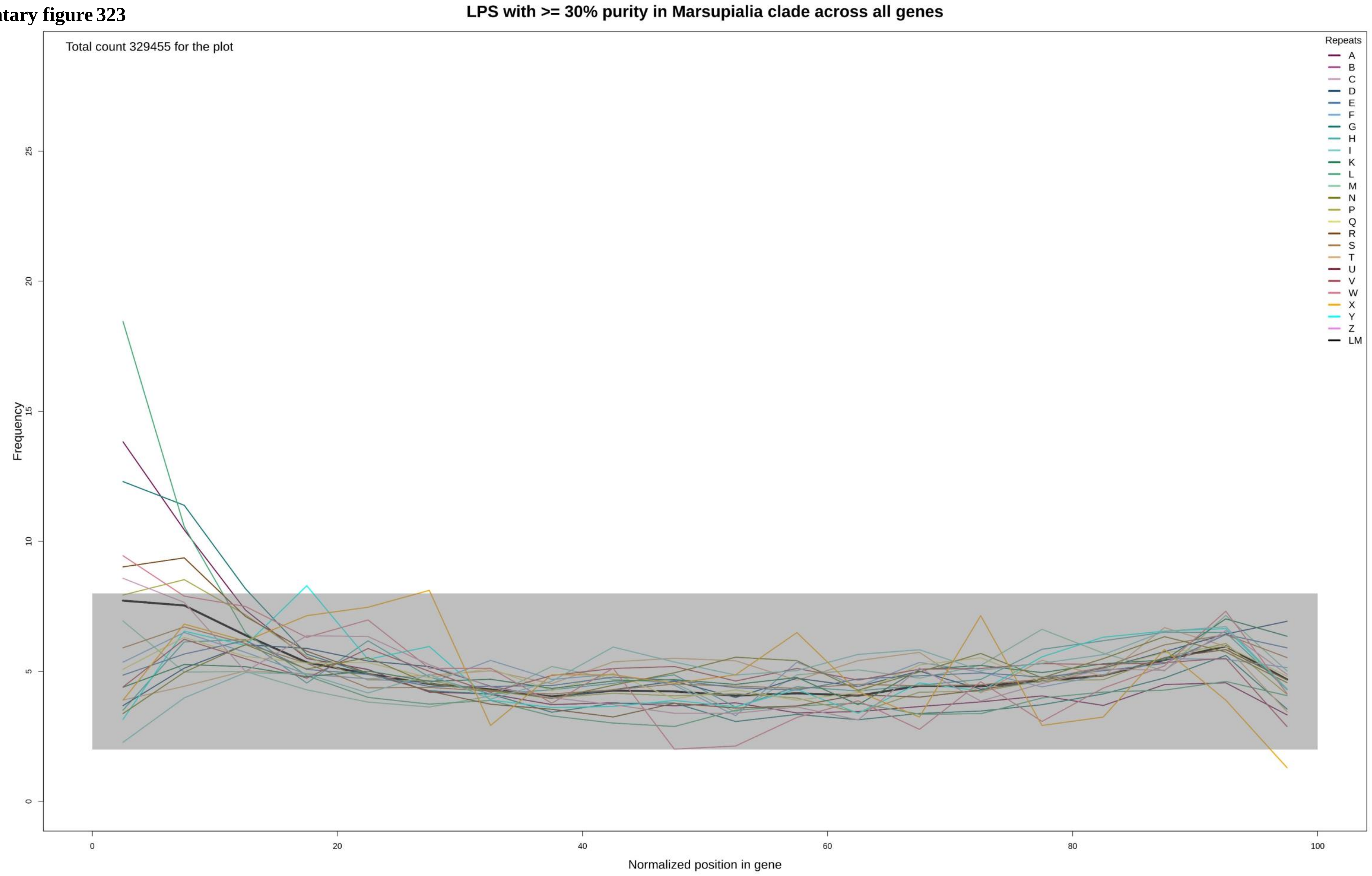

Supplementary figure 324

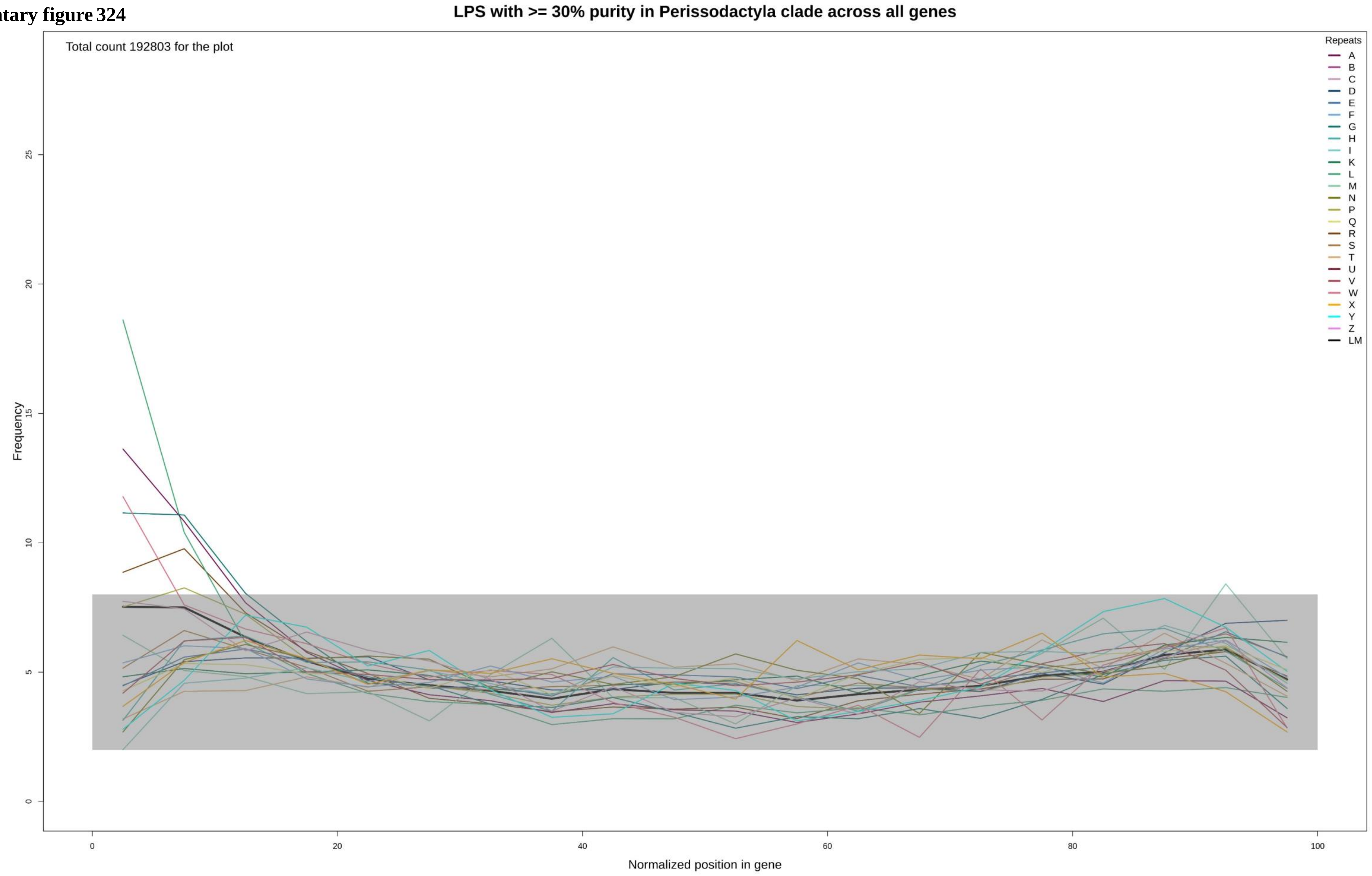

**Supplementary figure 325**

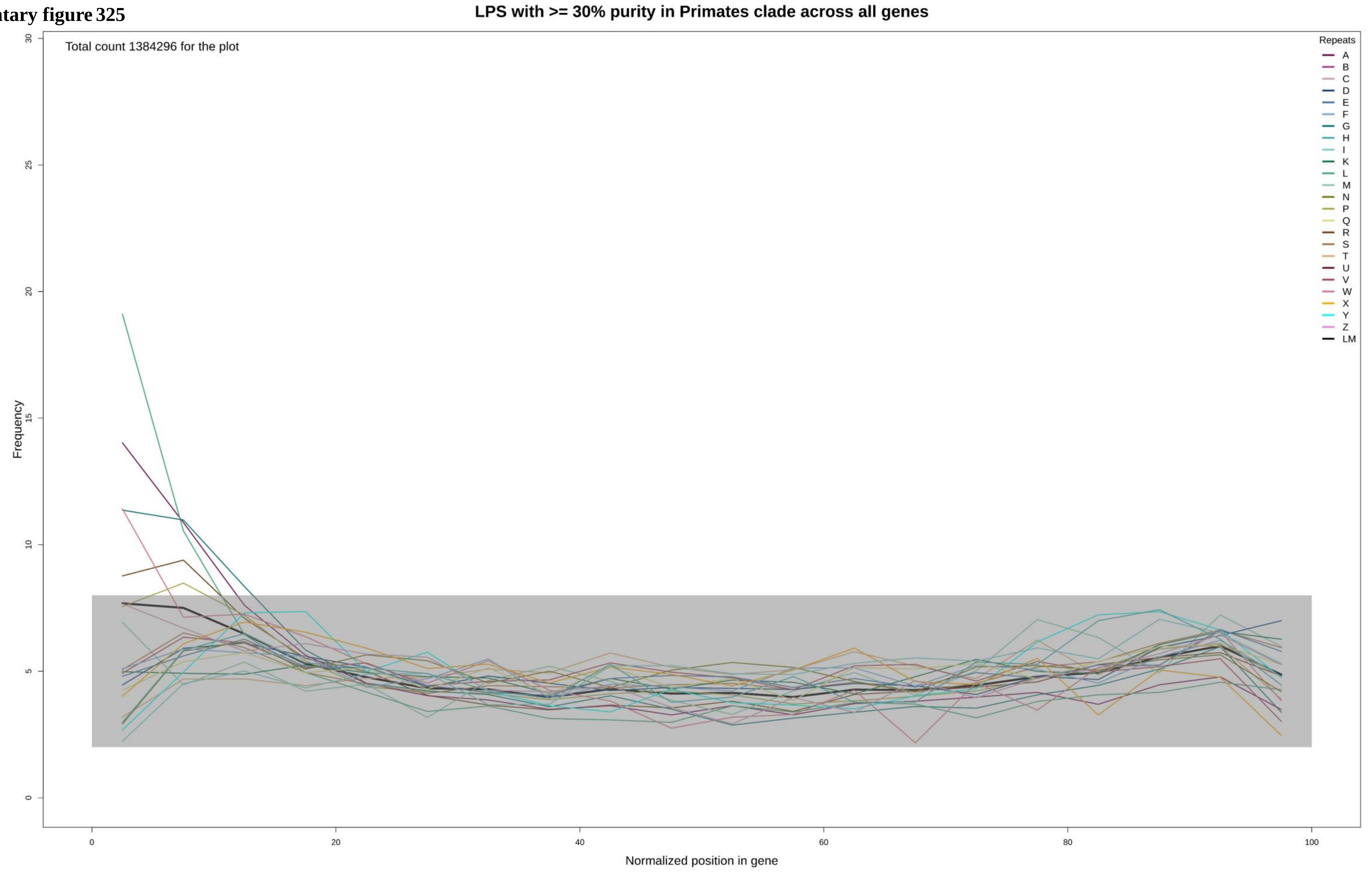

**Supplementary figure 326**

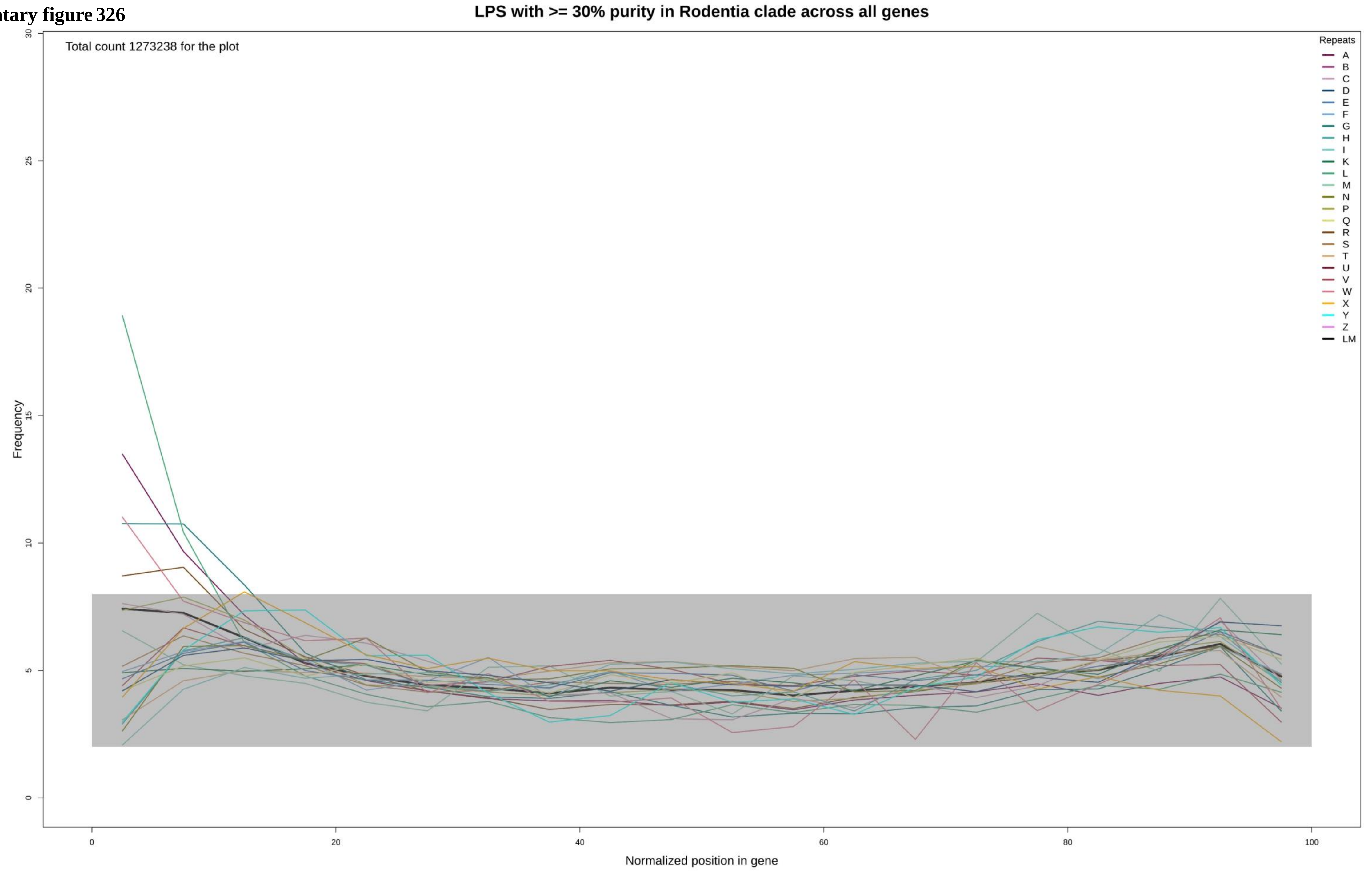

Supplementary figure 327

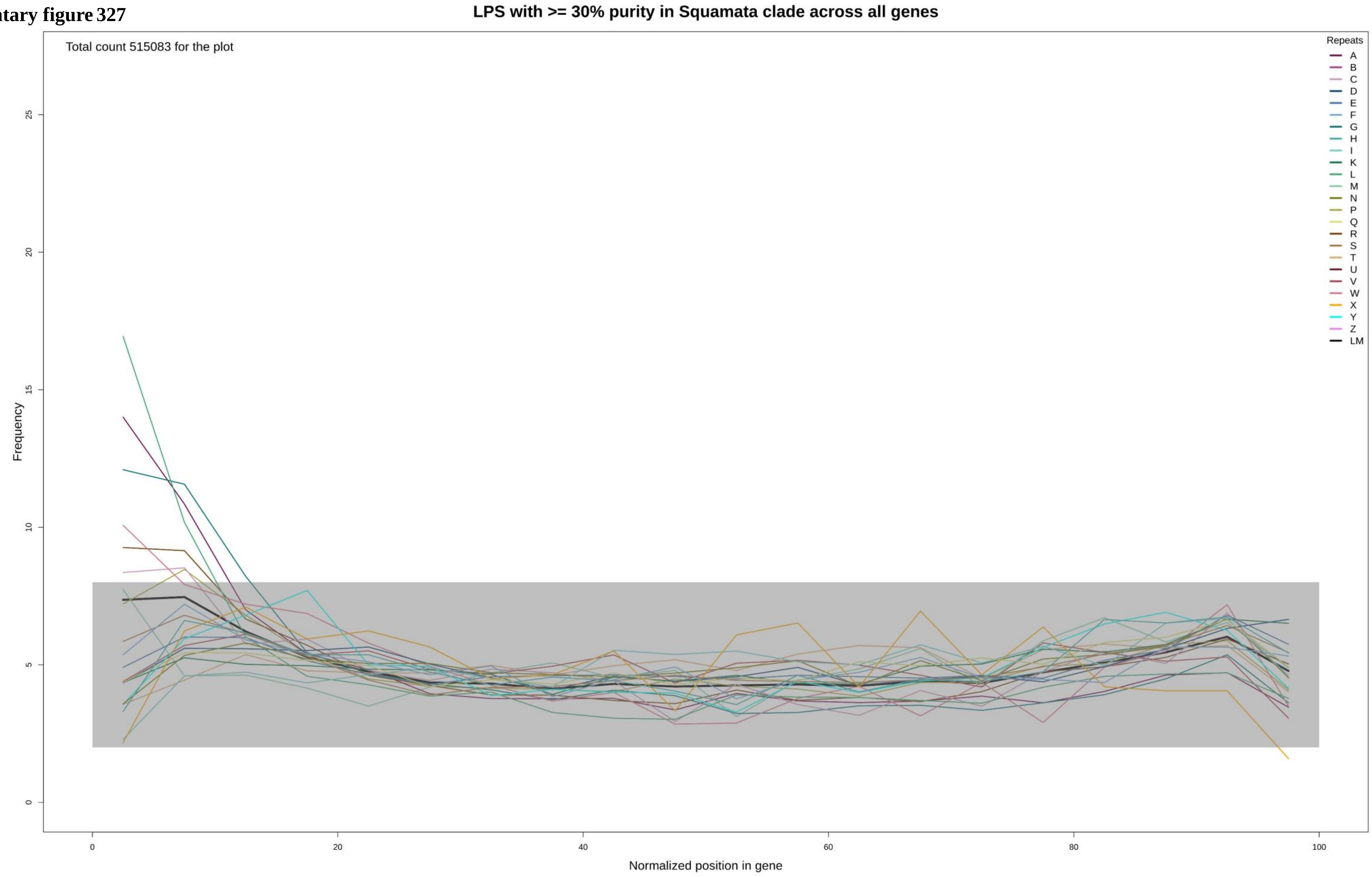

Supplementary figure 328

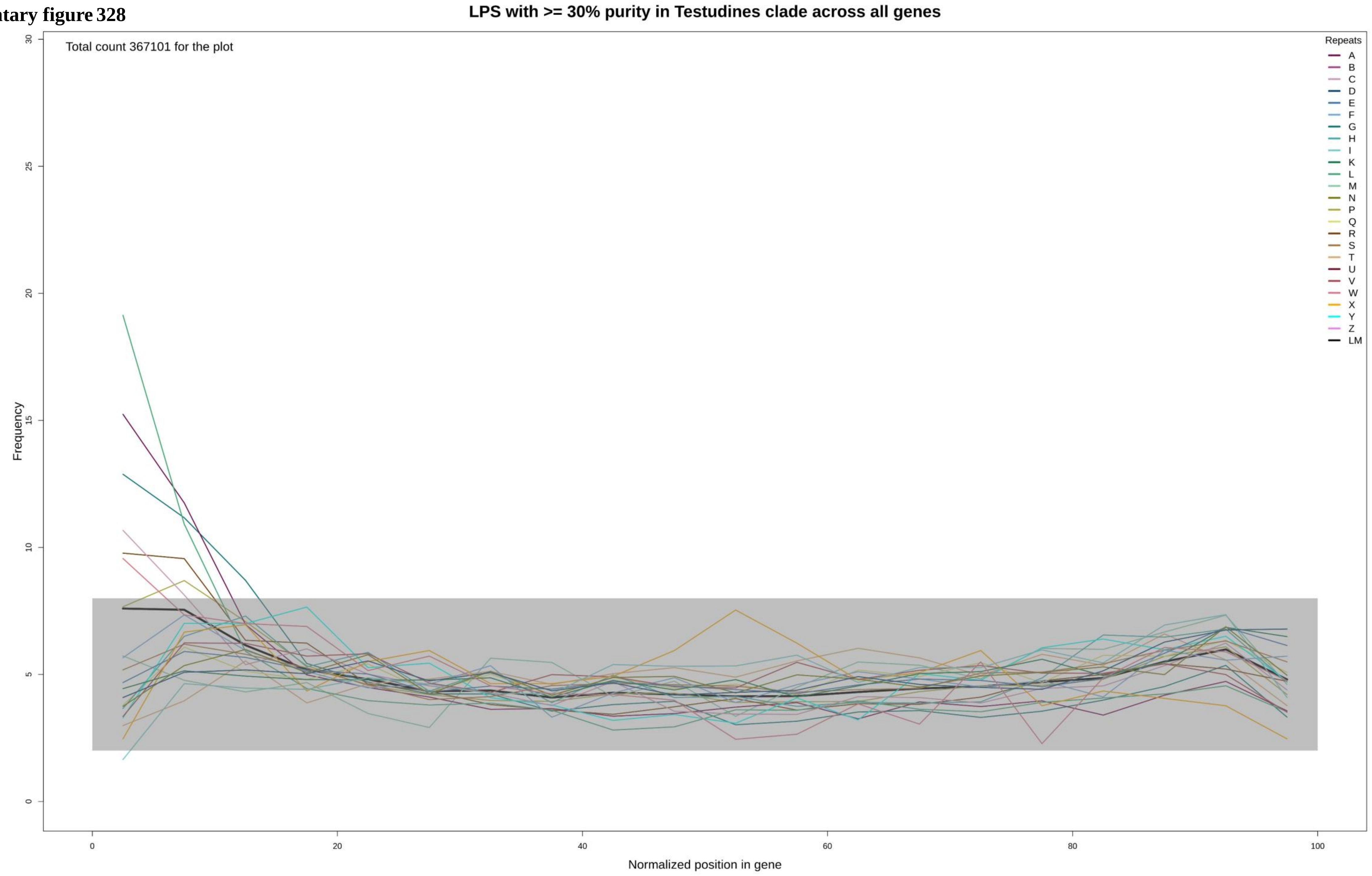

**Supplementary figure 329**

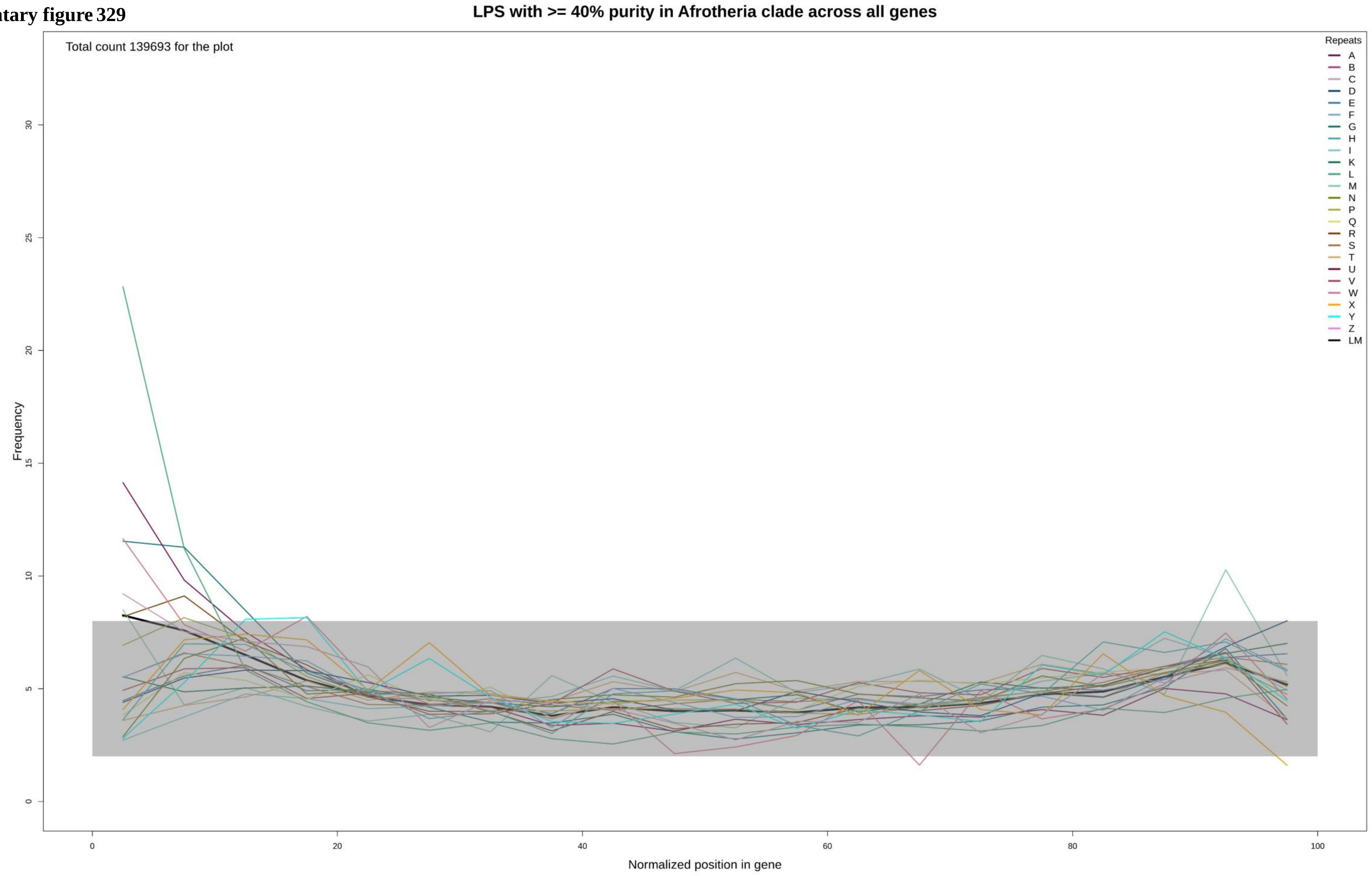

Supplementary figure 330

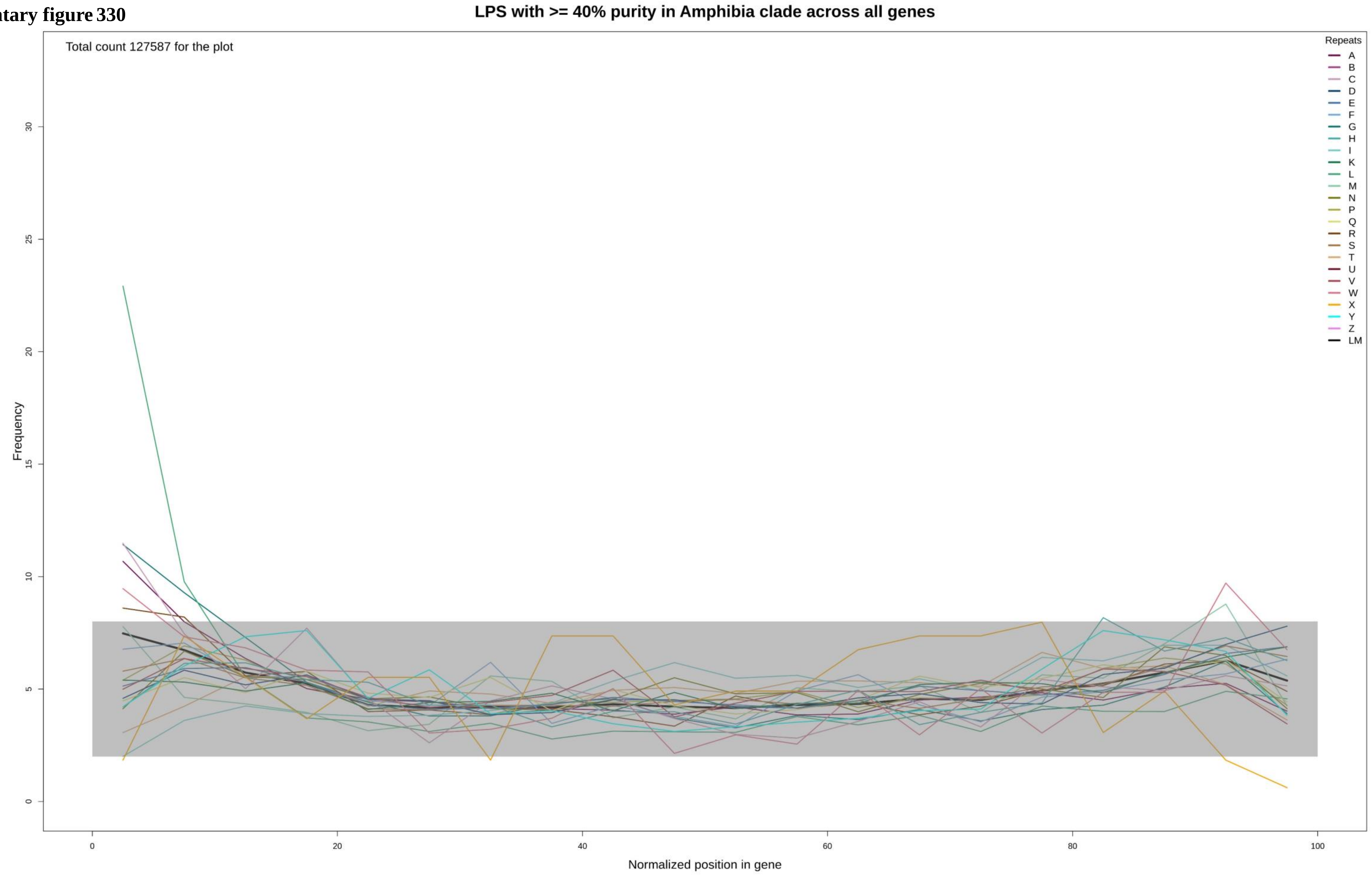

**Supplementary figure 331**

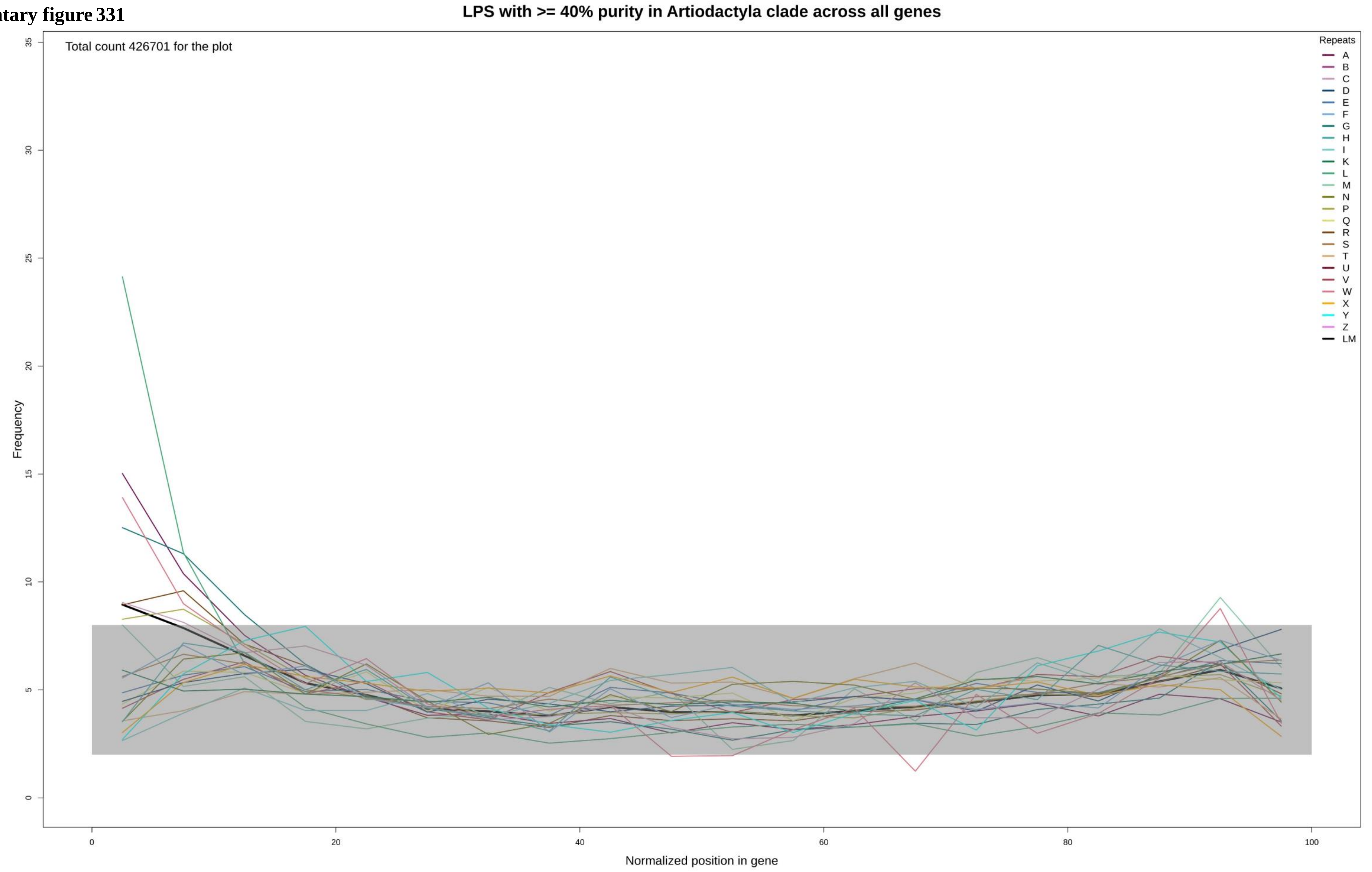

**Supplementary figure 332**

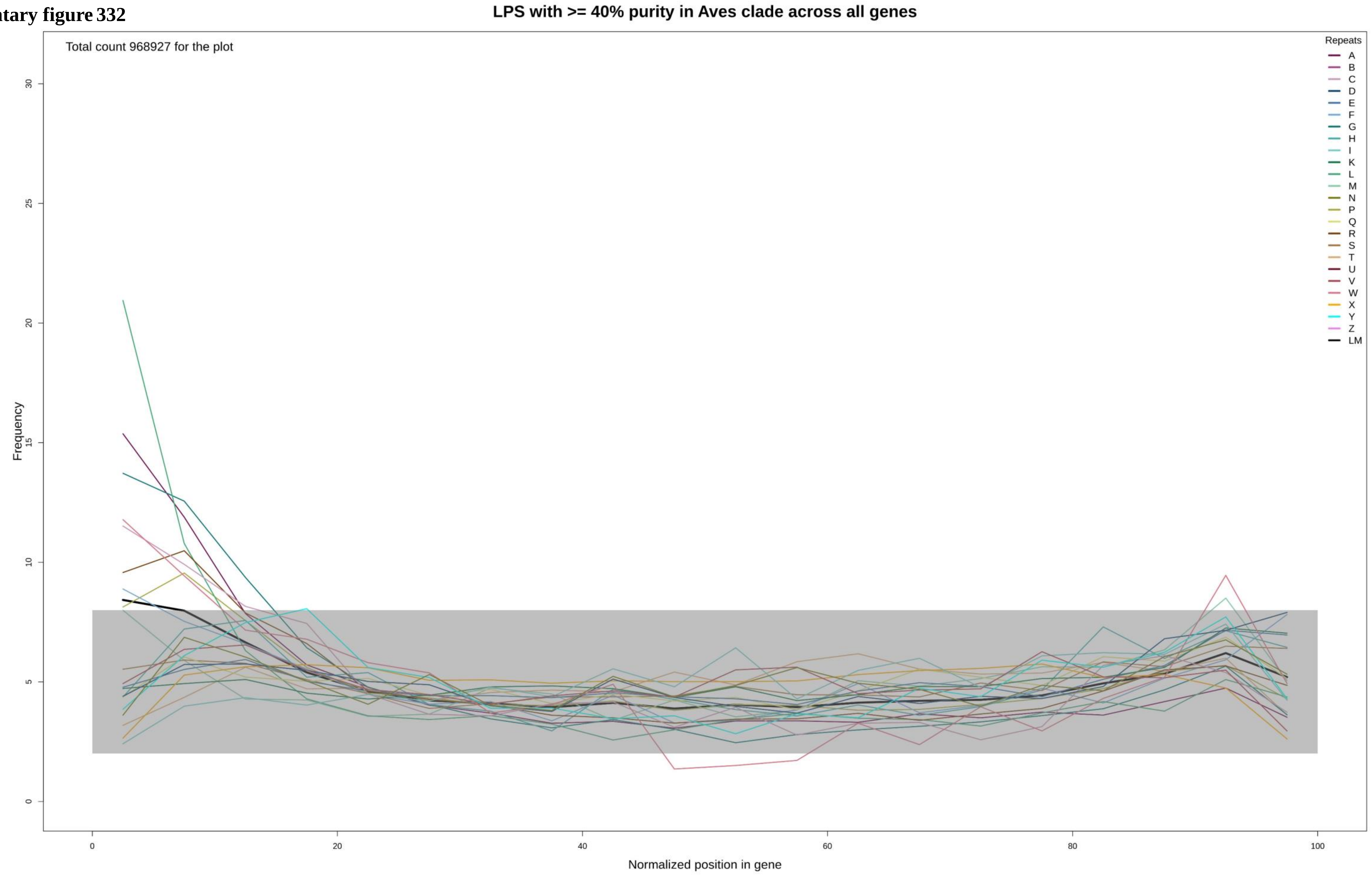

**Supplementary figure 333**

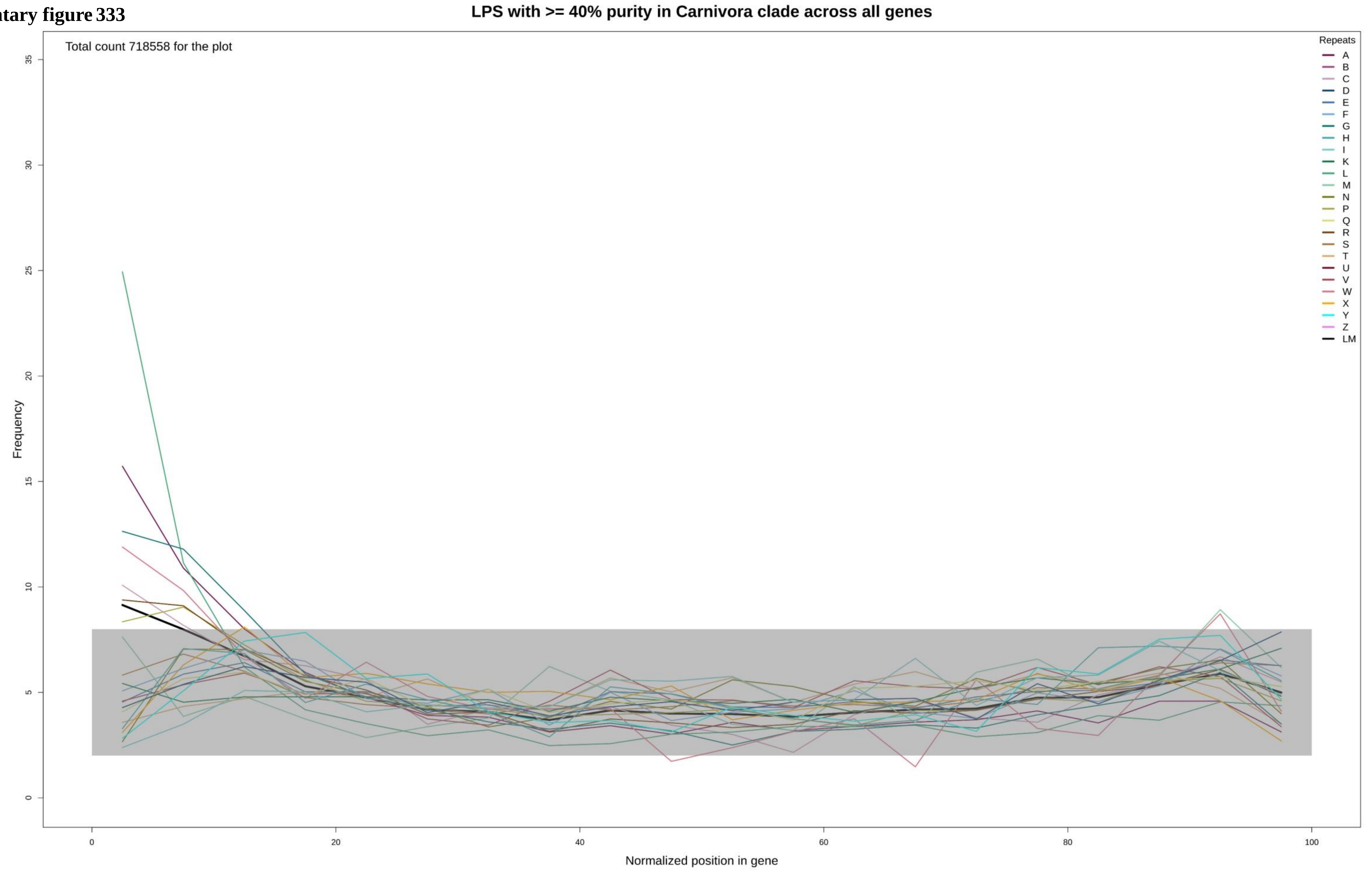

Supplementary figure 334

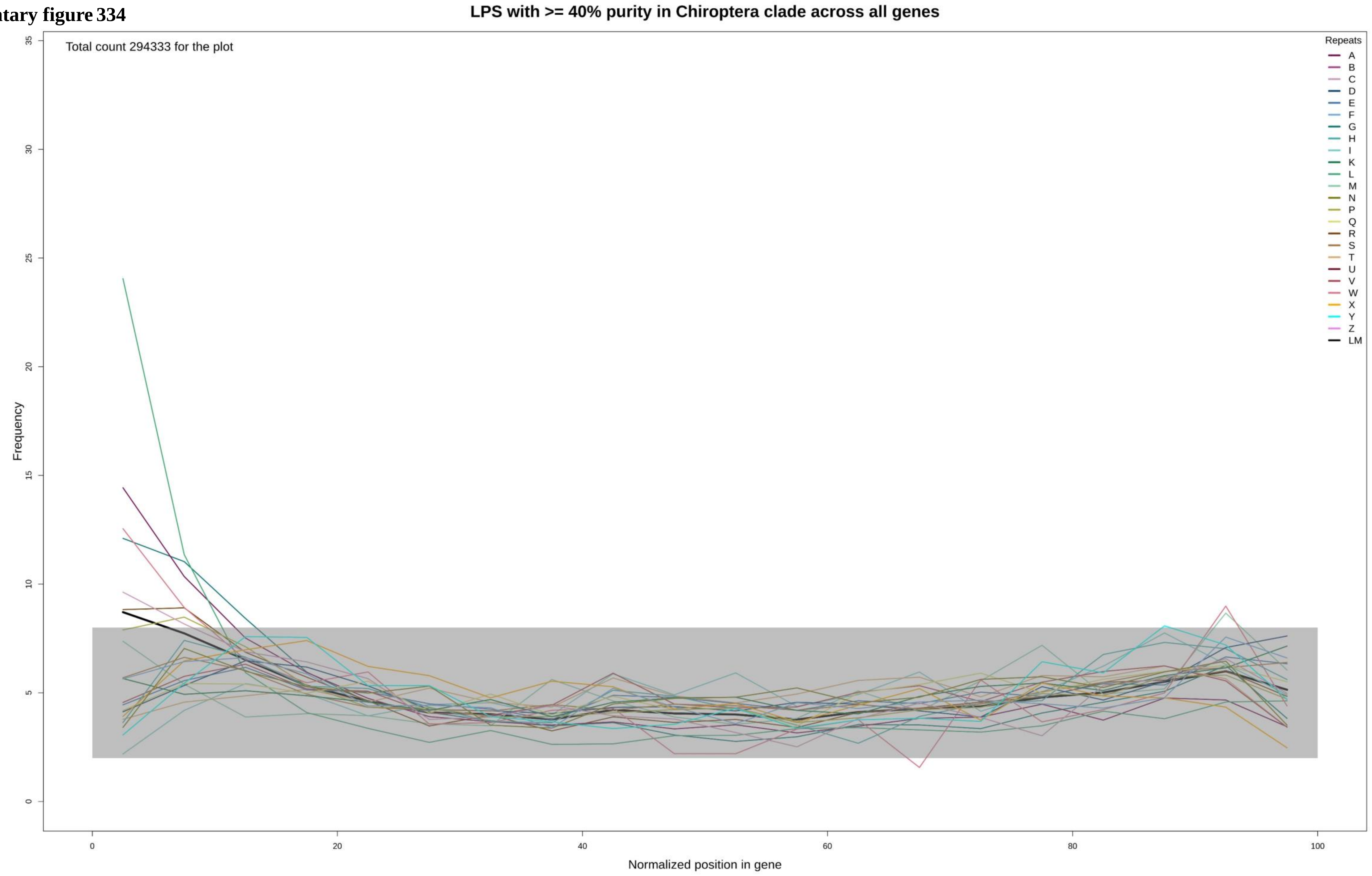

Supplementary figure 335

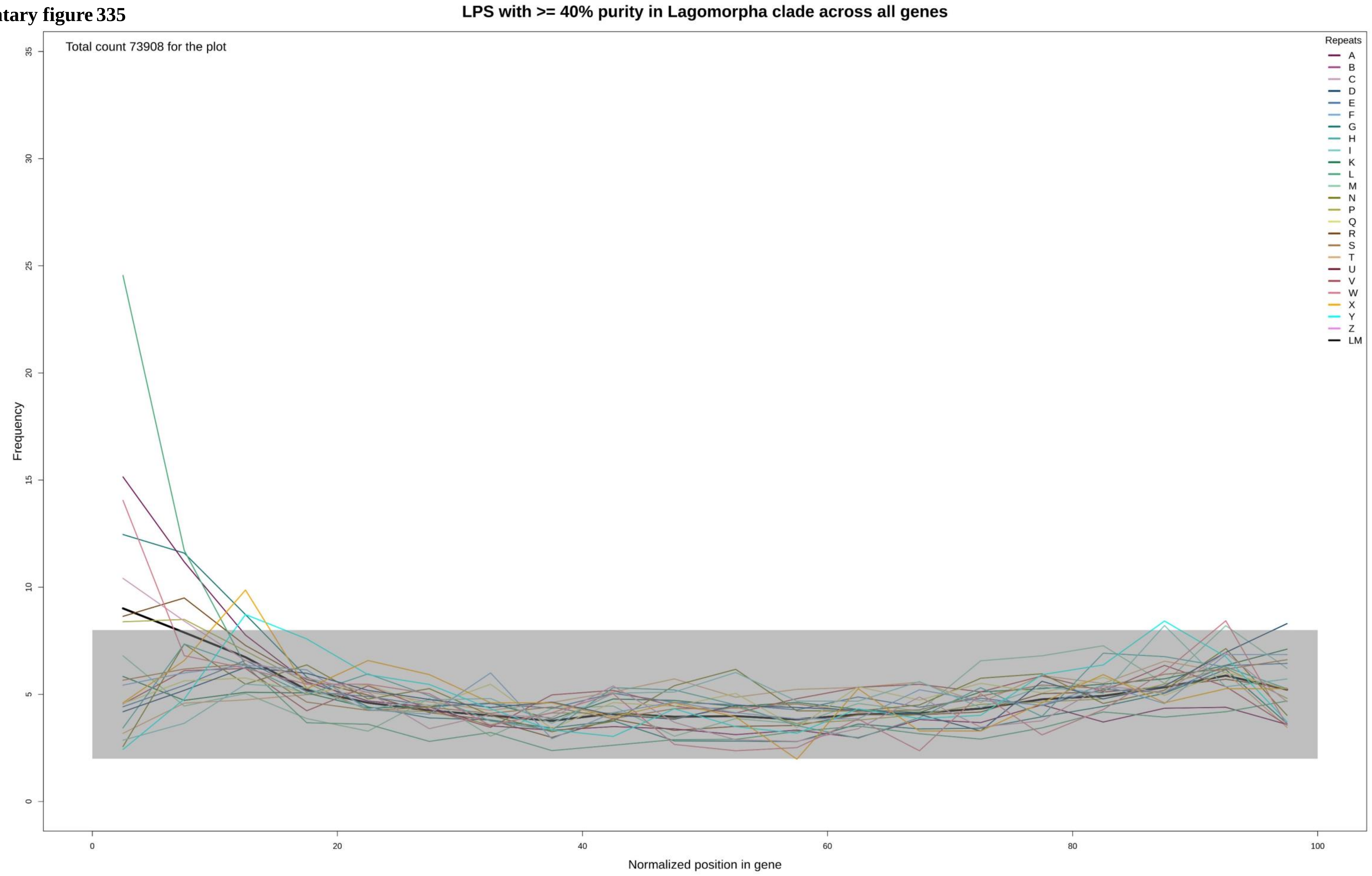

Supplementary figure 336

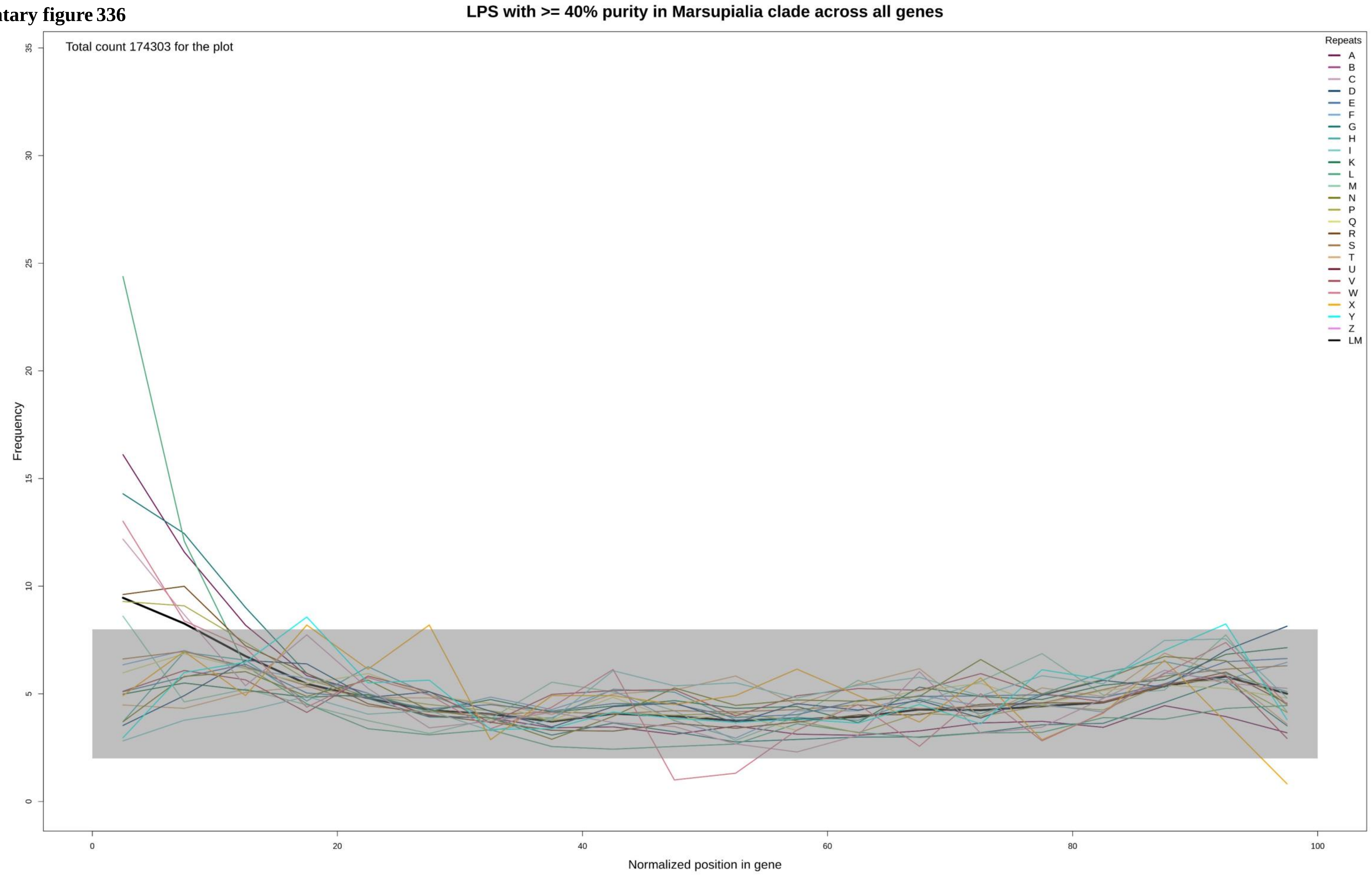

**Supplementary figure 337**

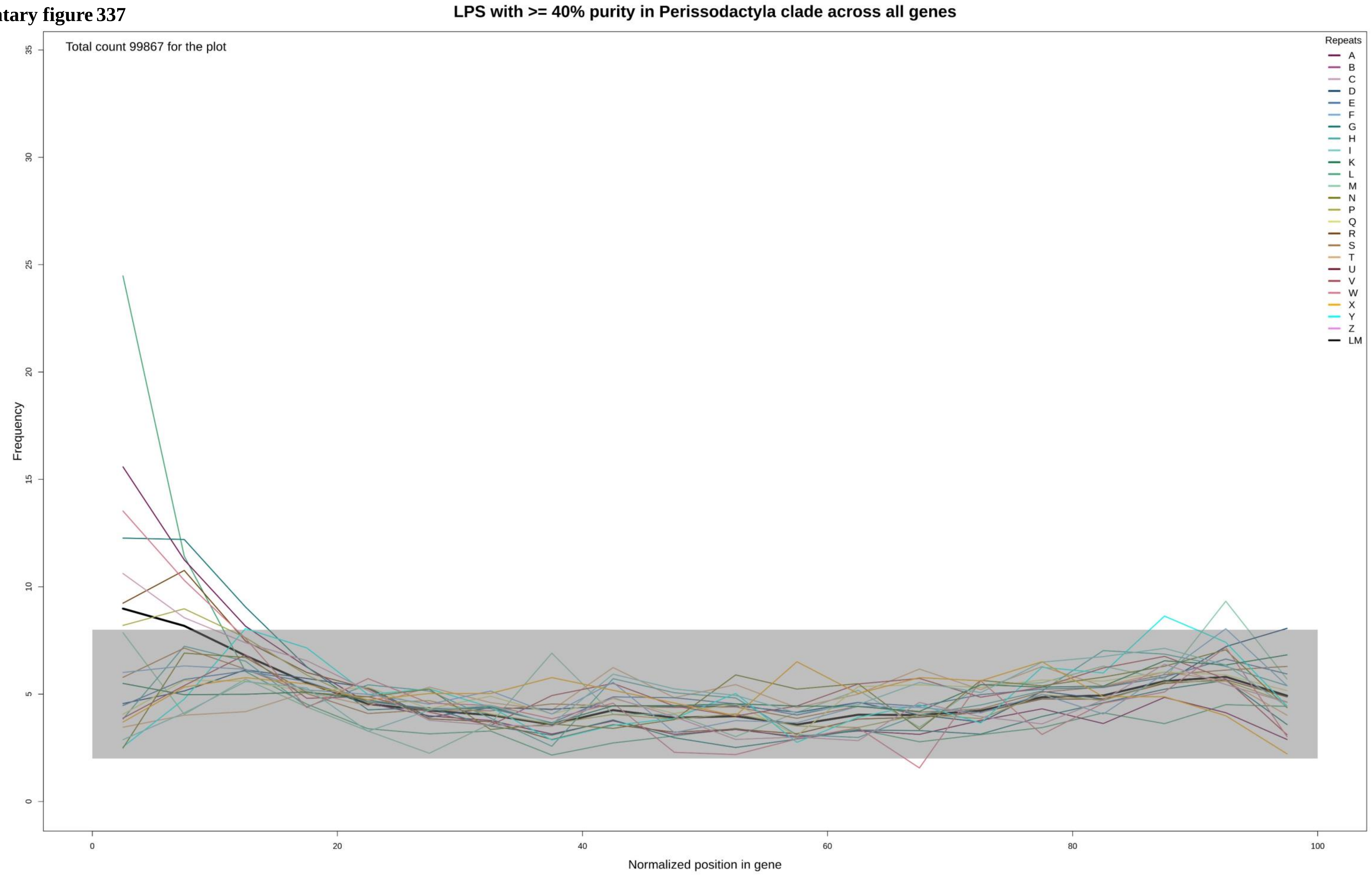

**Supplementary figure 338**

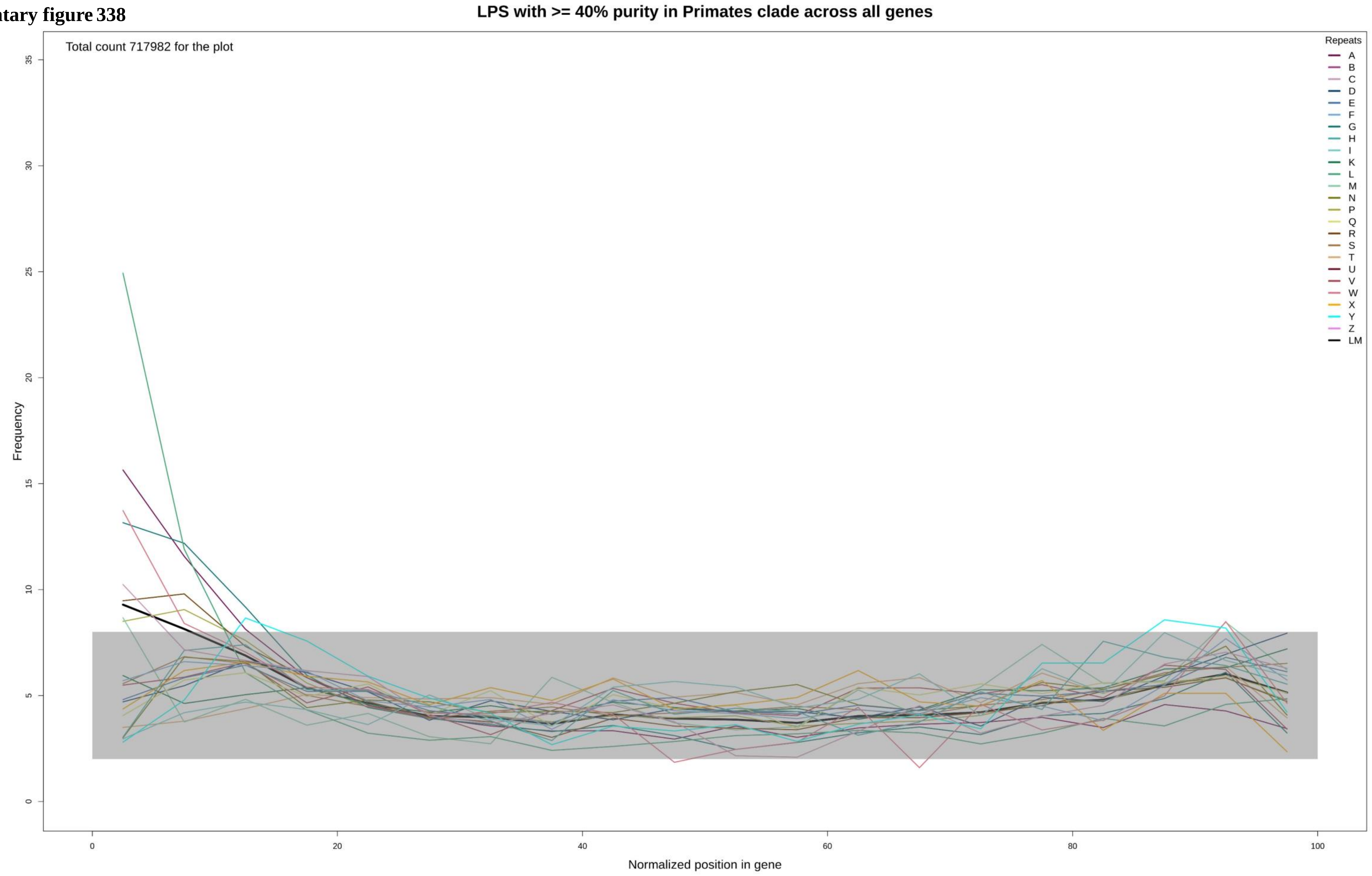

**Supplementary figure 339**

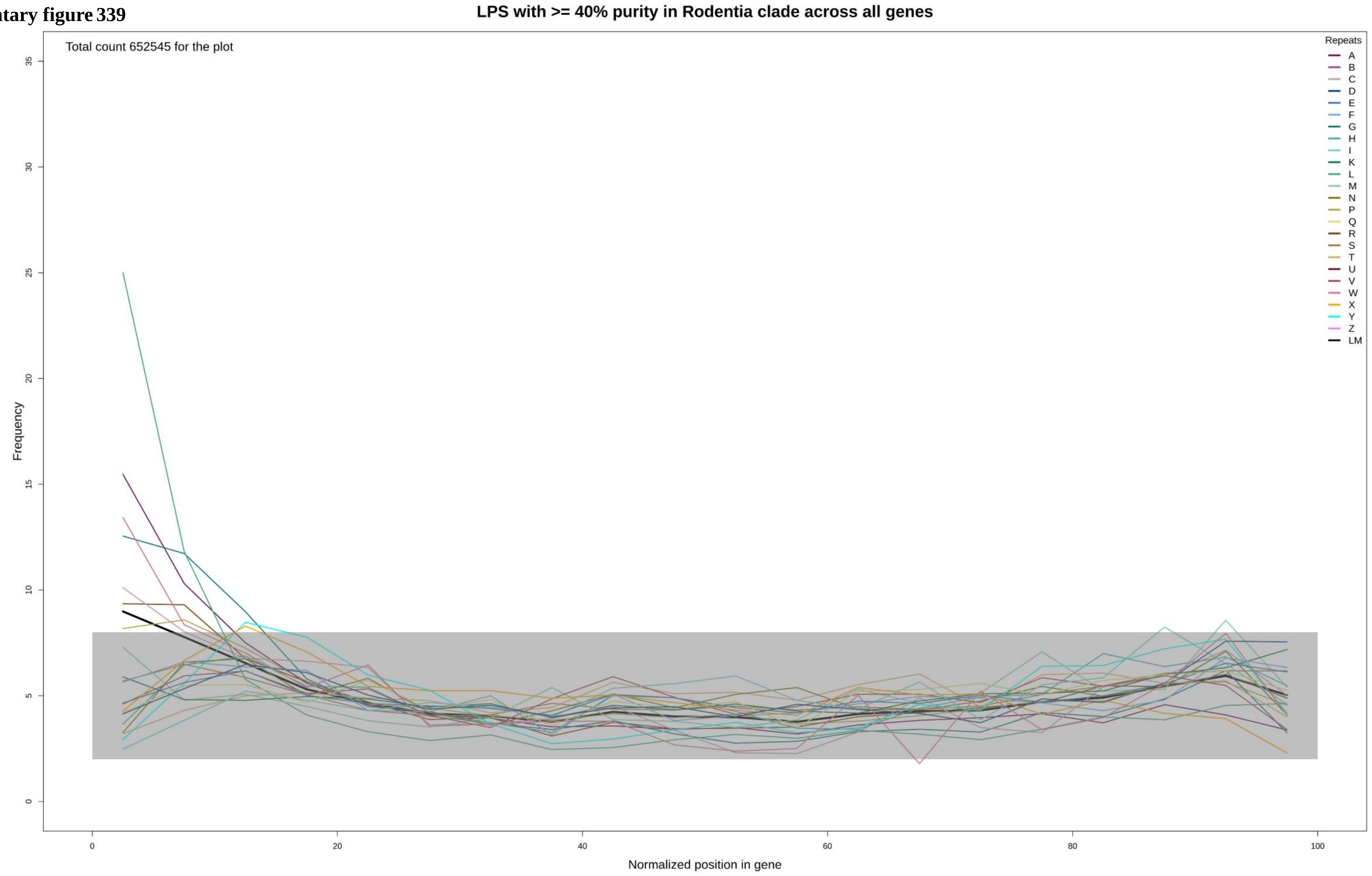

**Supplementary figure 340**

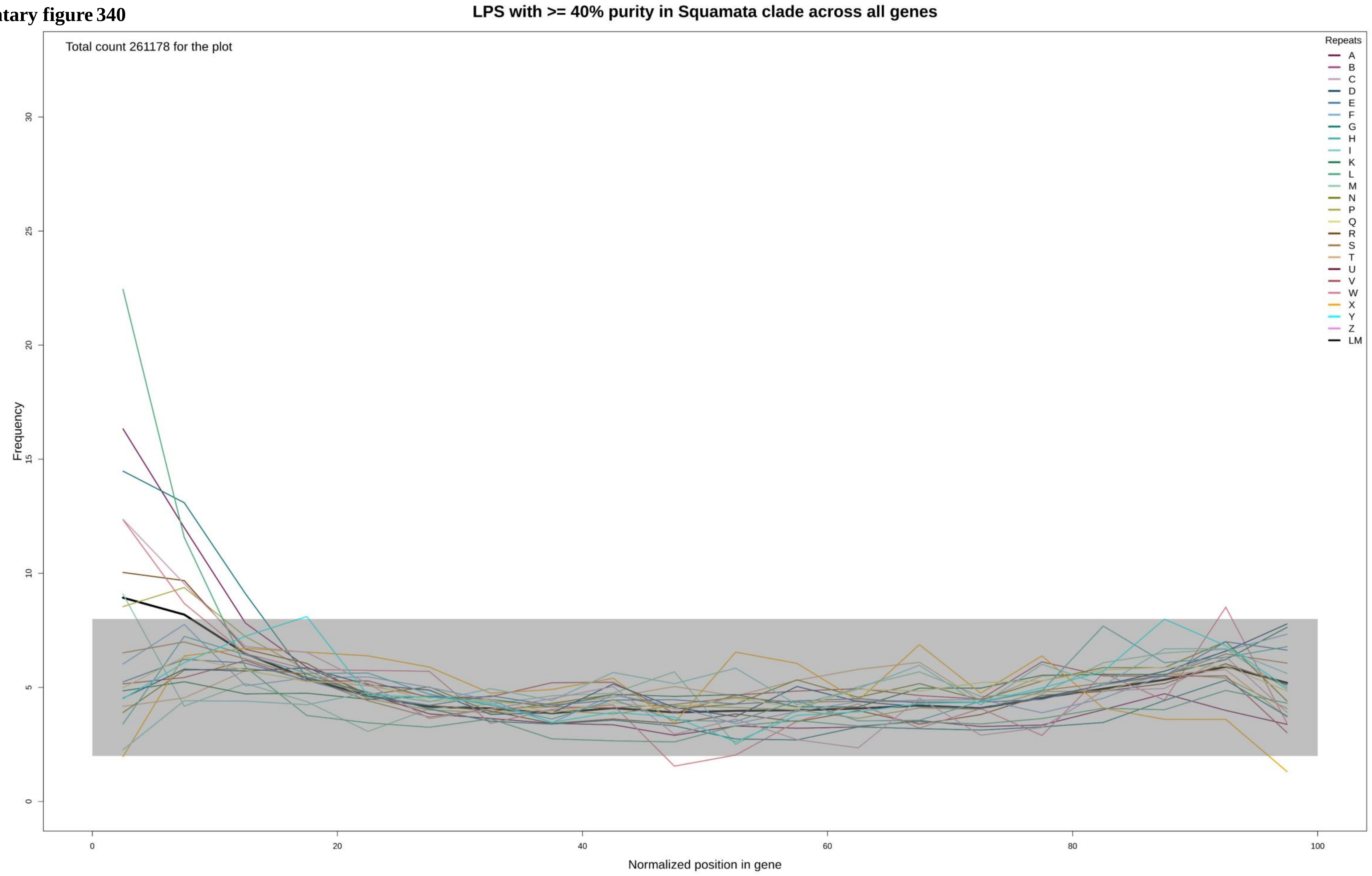

Supplementary figure 341

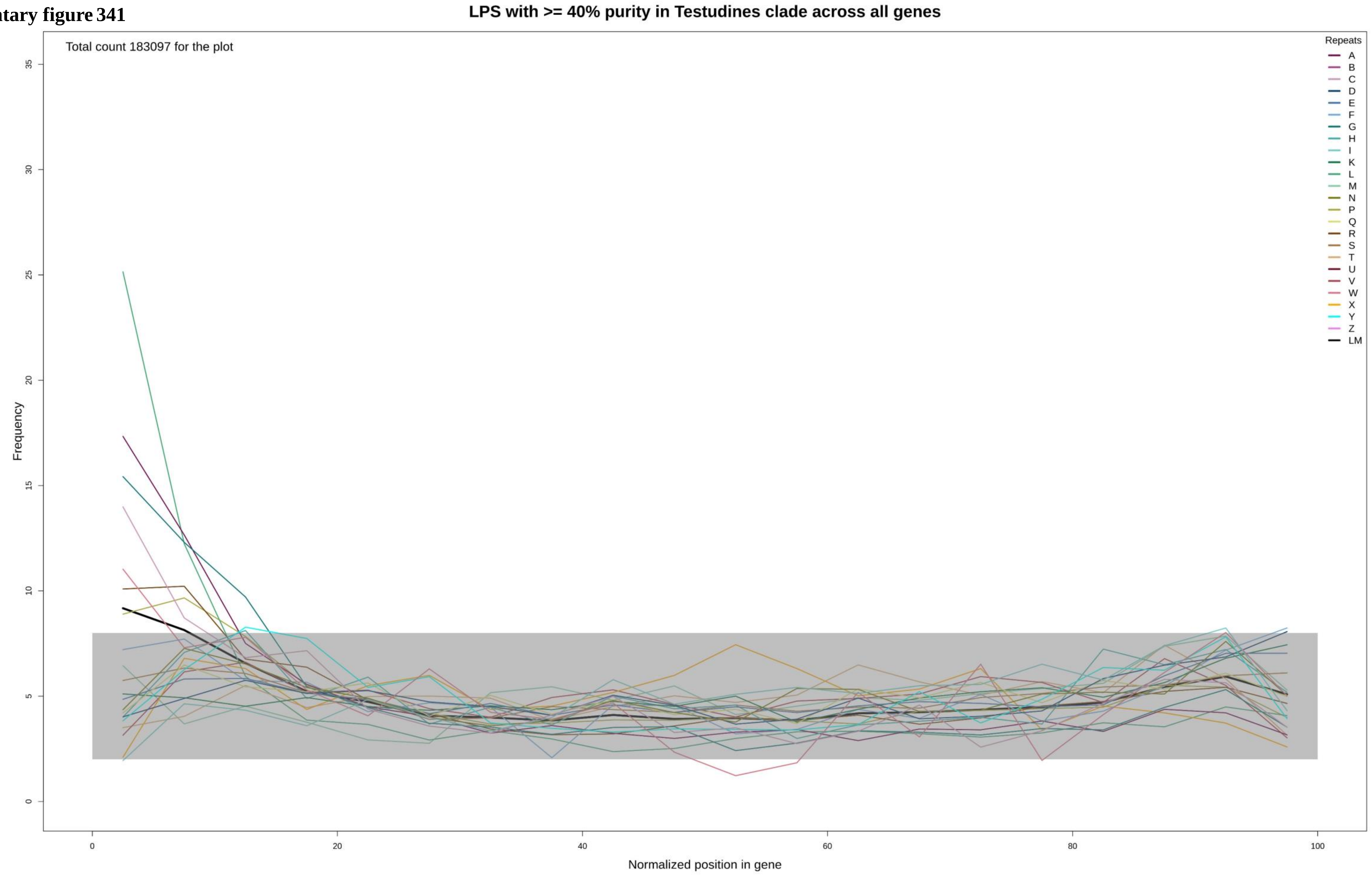

**Supplementary figure 342**

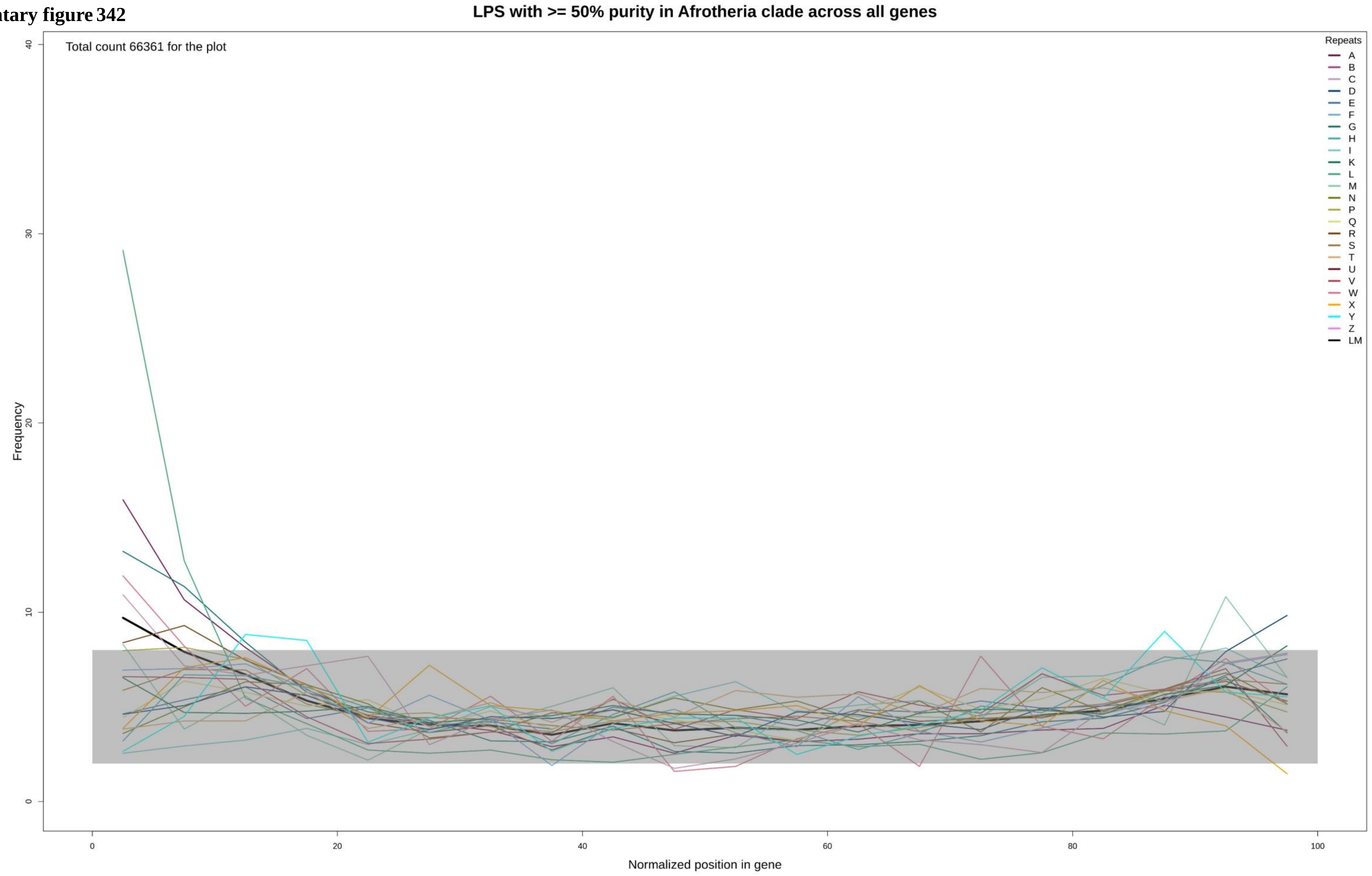

**Supplementary figure 343**

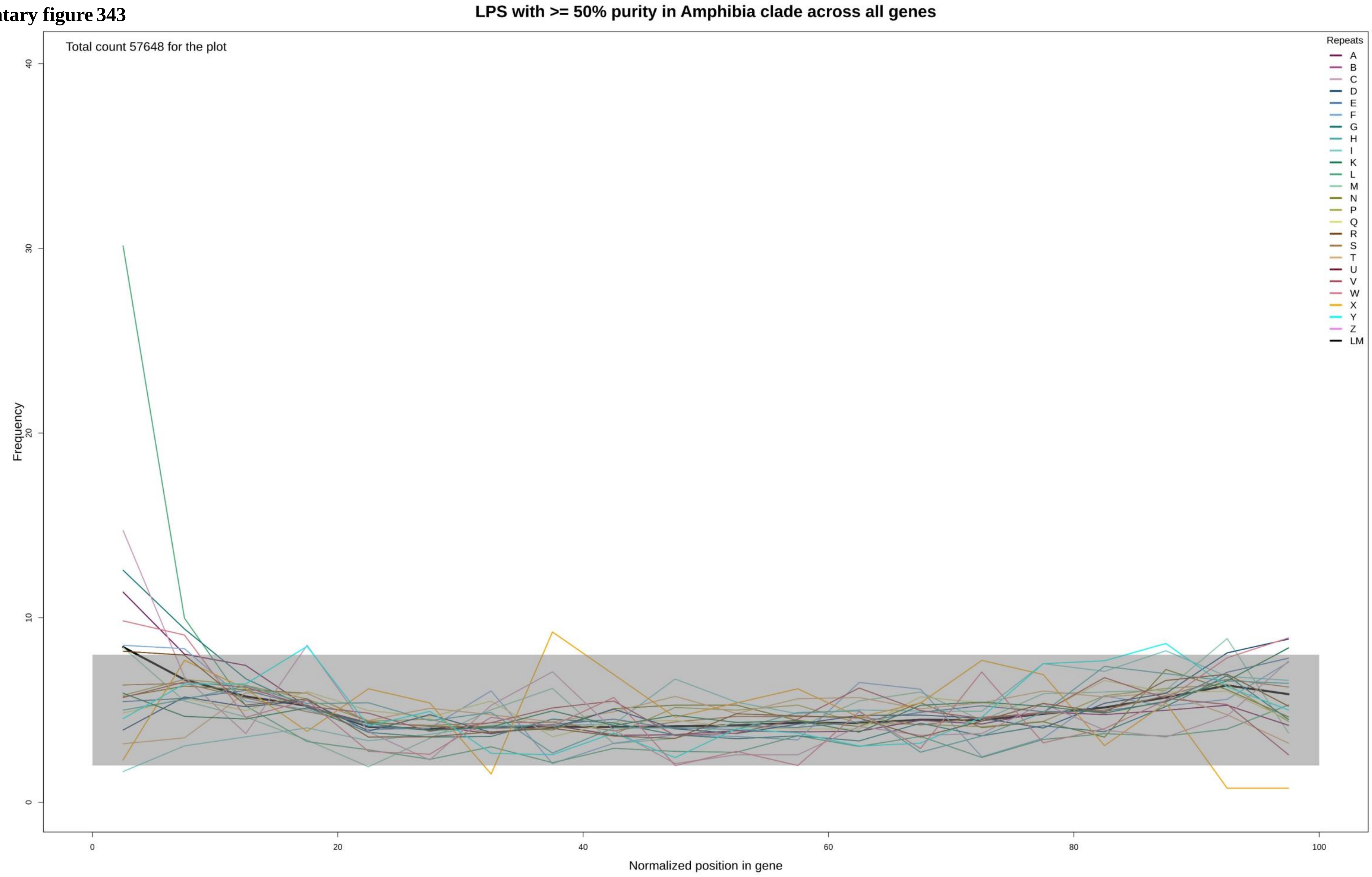

**Supplementary figure 344**

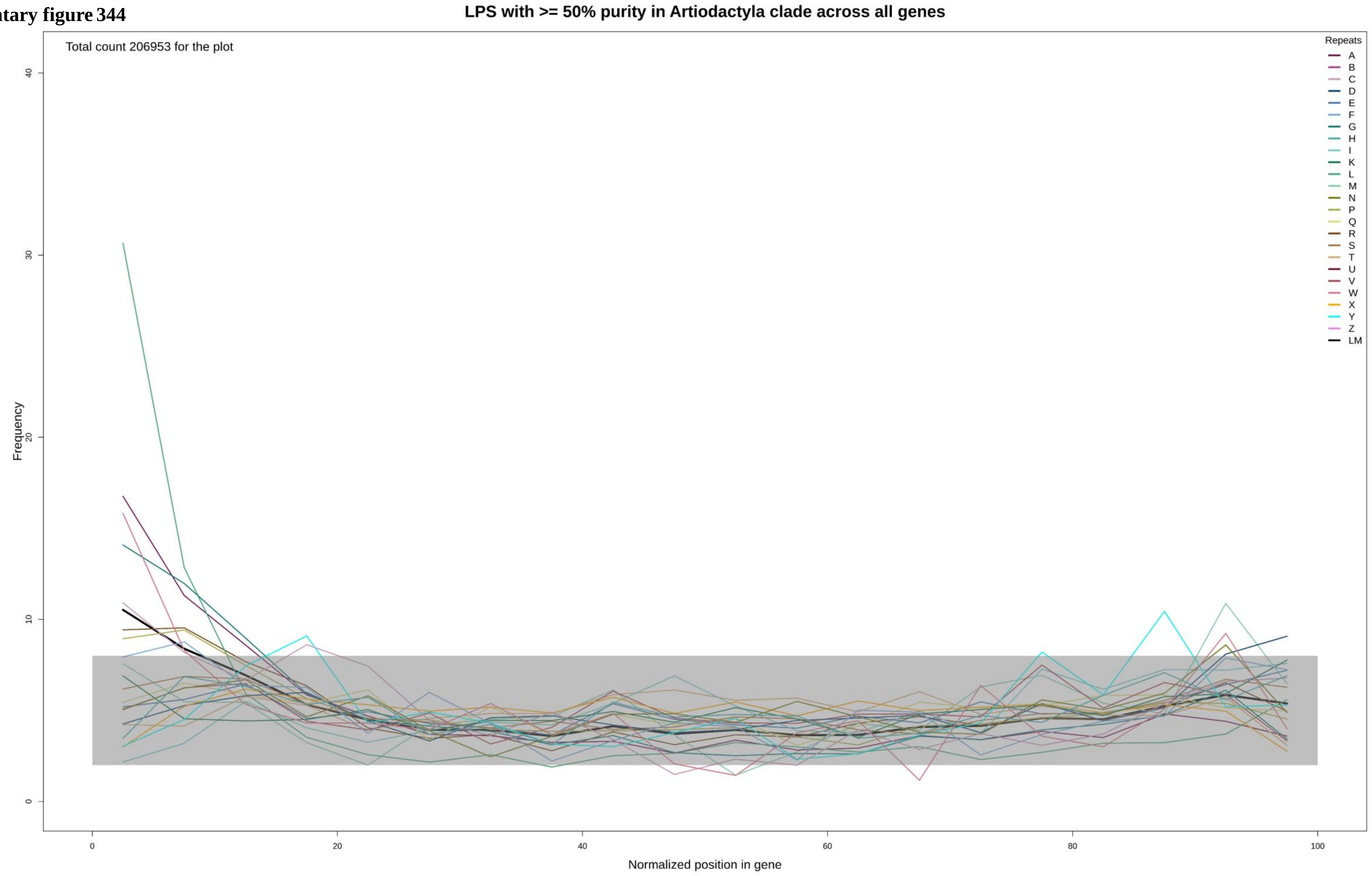

**Supplementary figure 345**

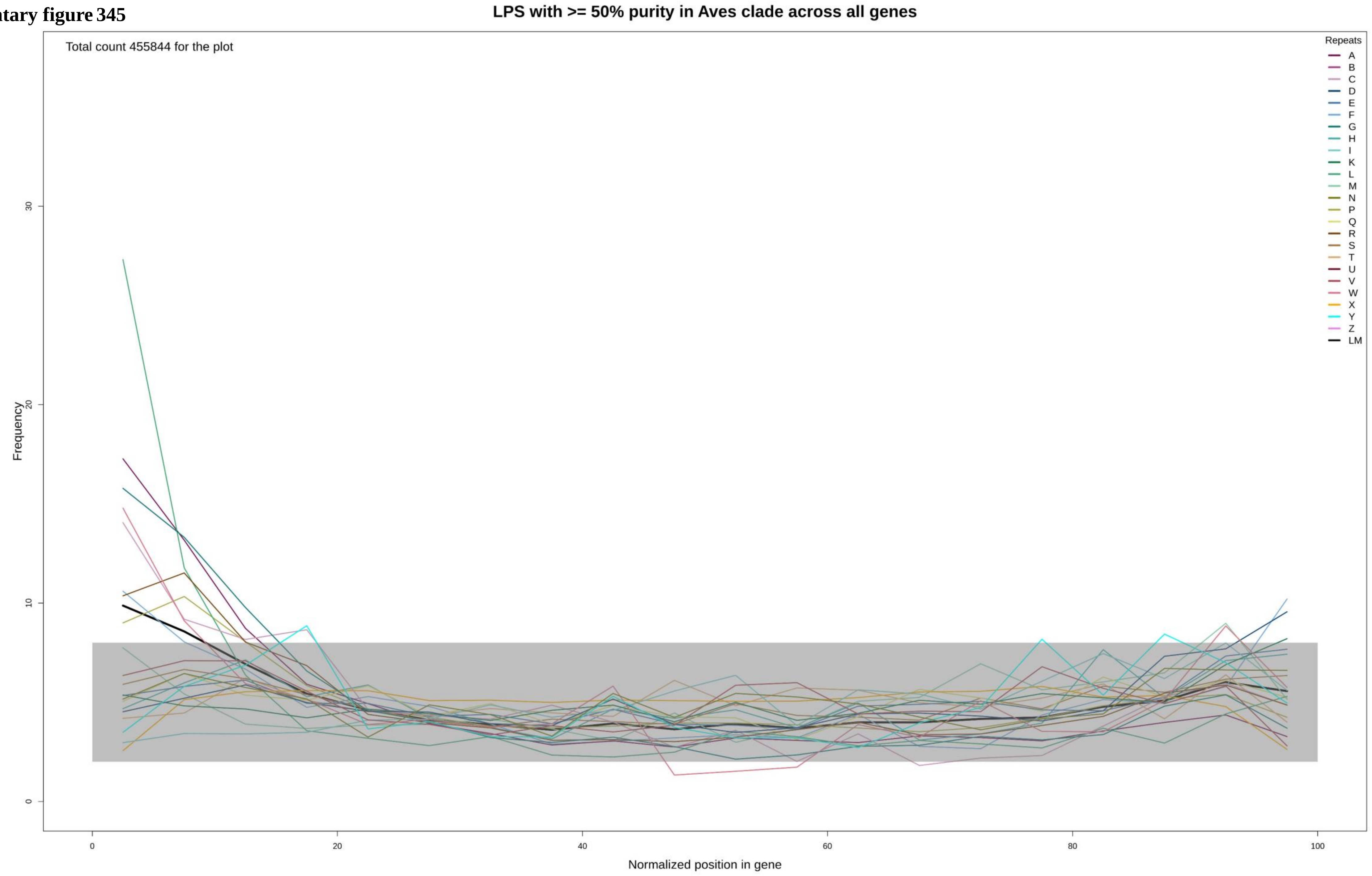

**Supplementary figure 346**

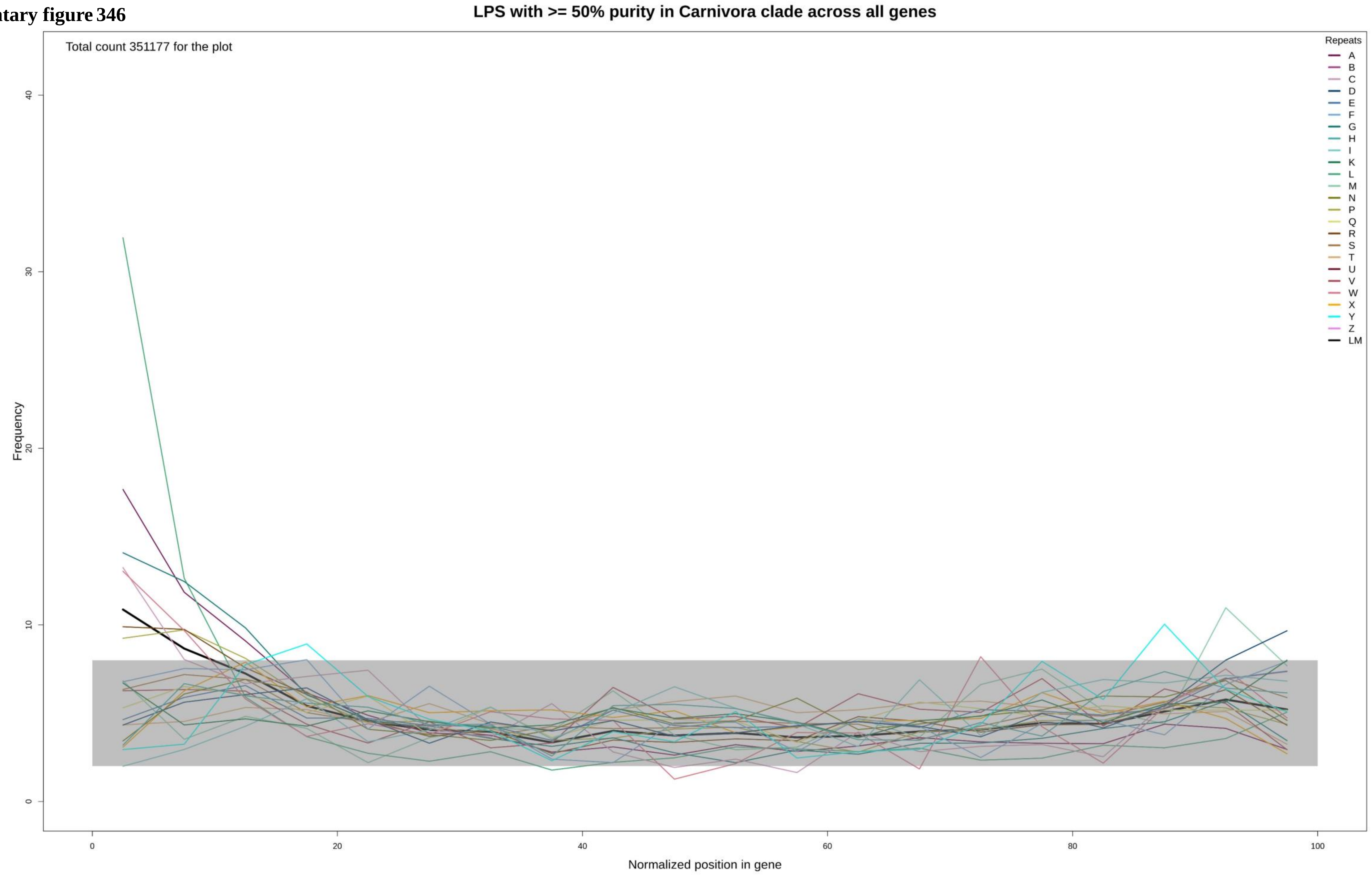

**Supplementary figure 347**

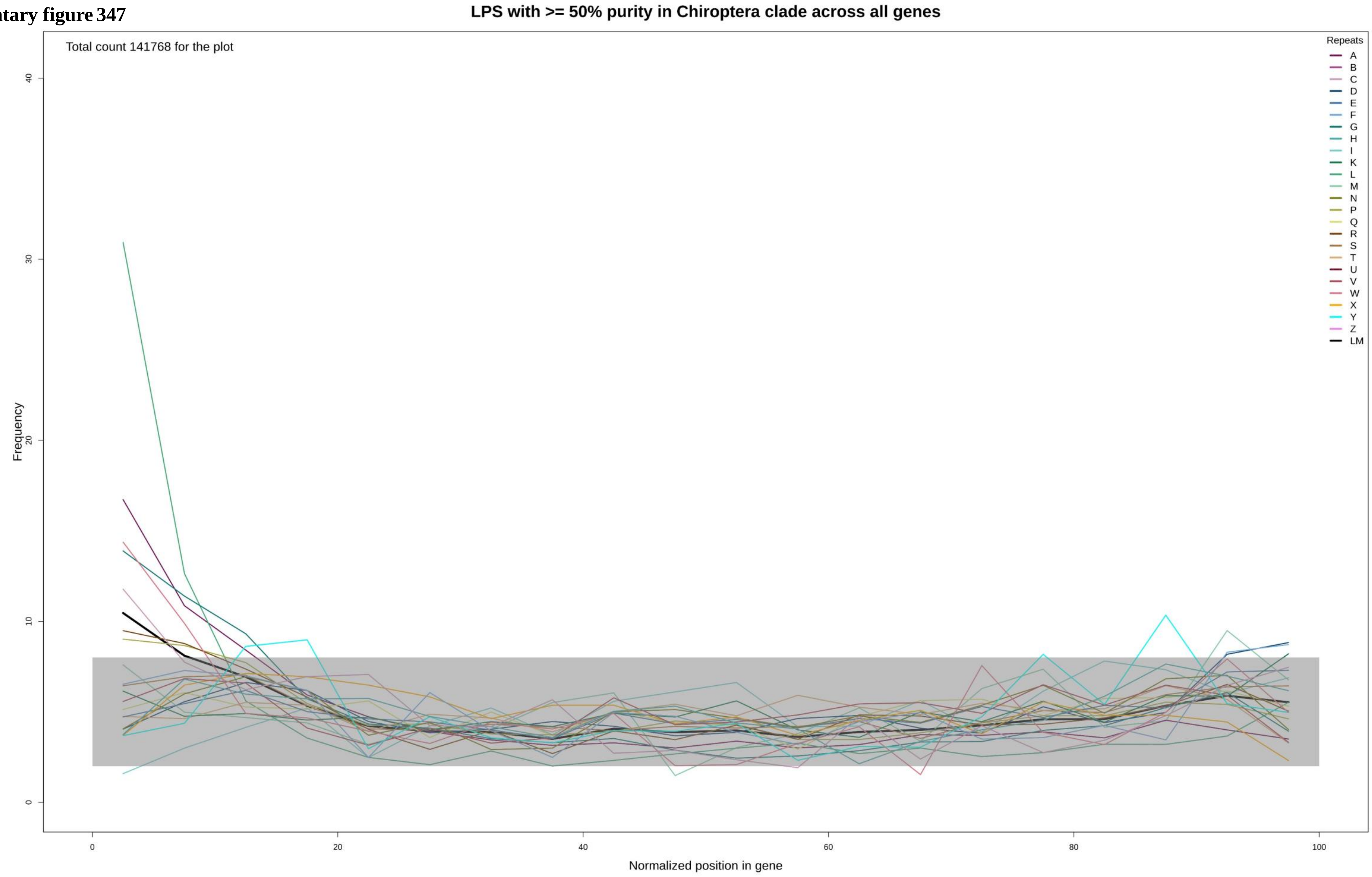

**Supplementary figure 348**

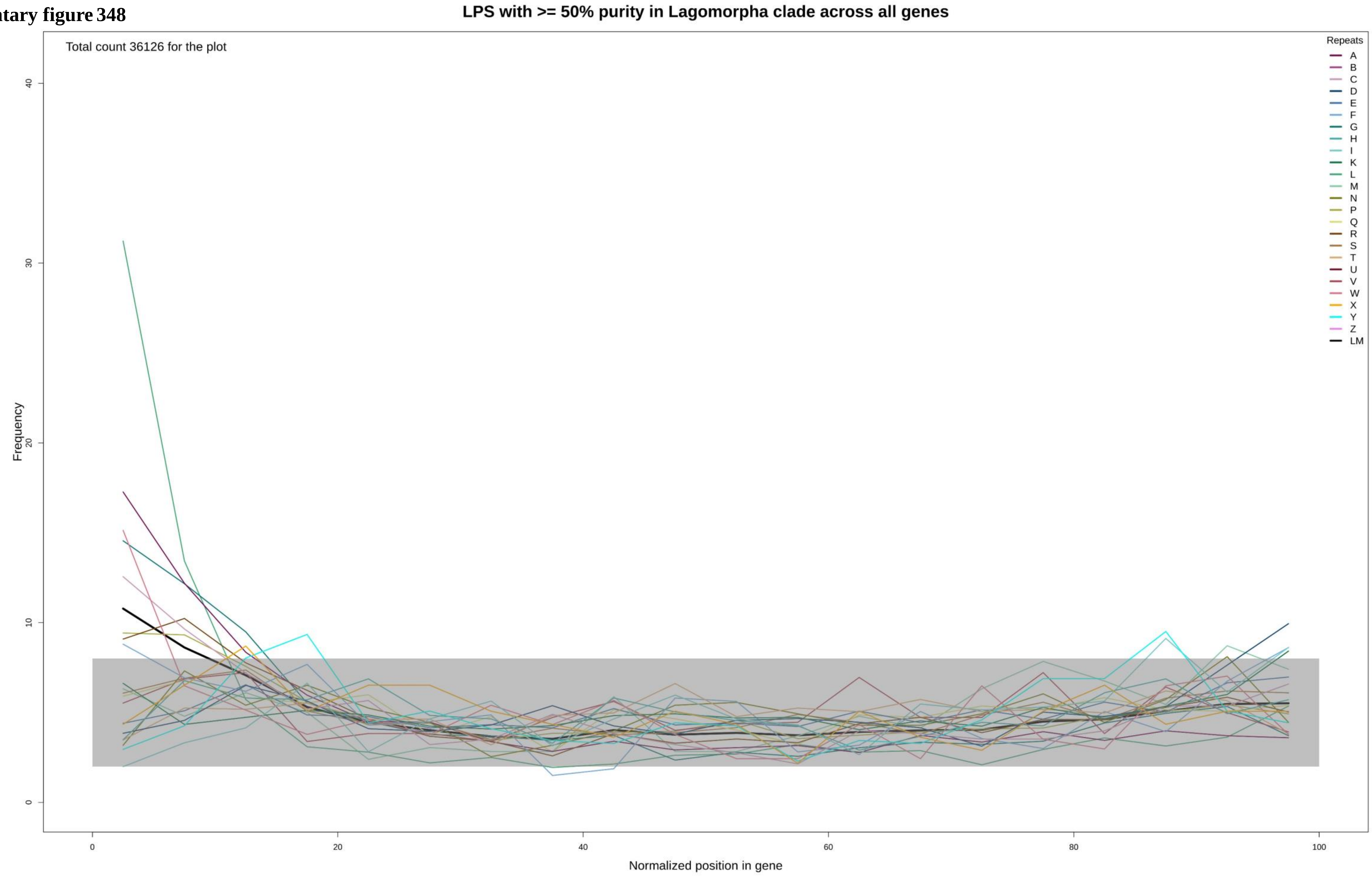

Supplementary figure 349

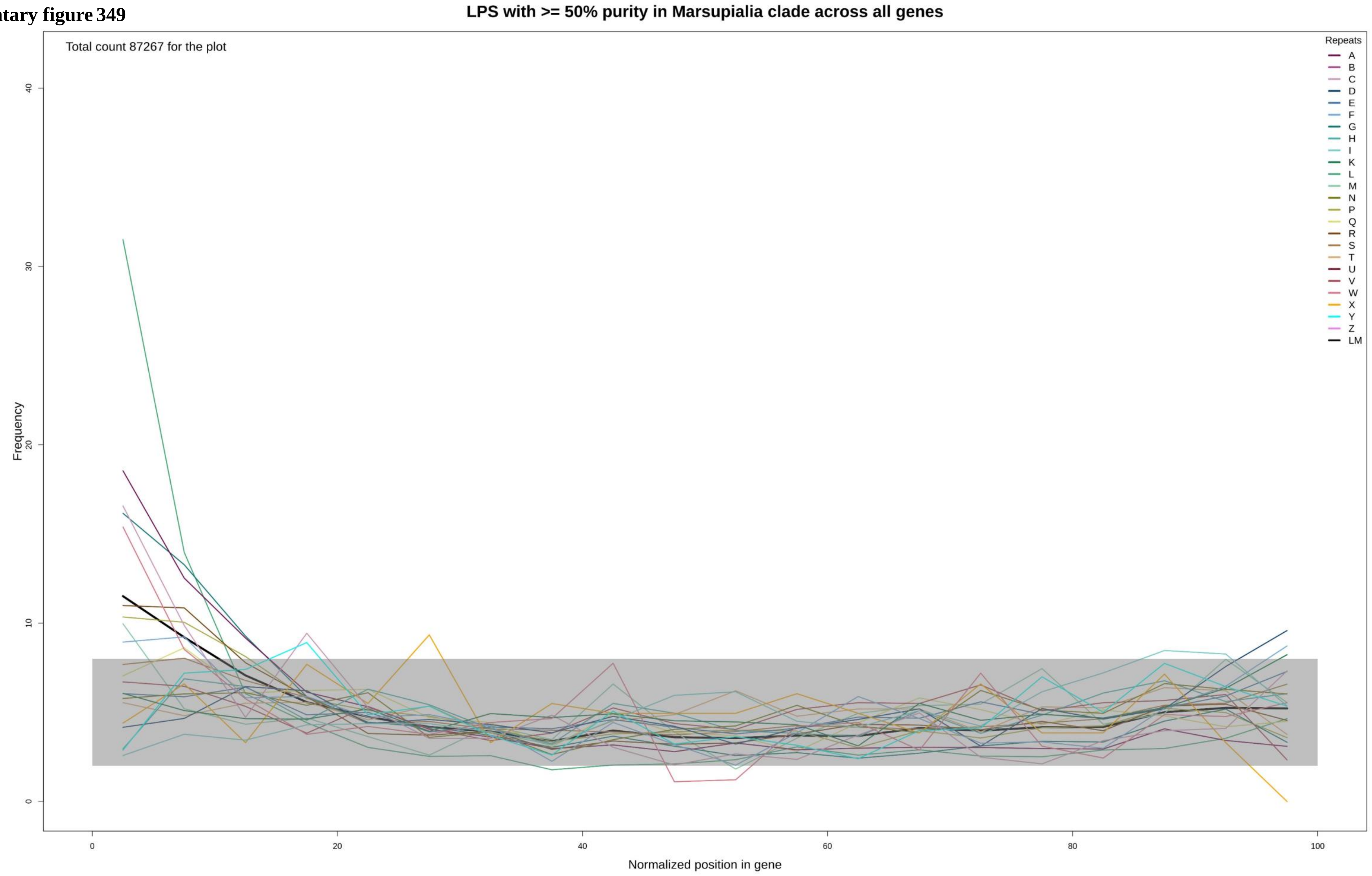

Supplementary figure 350

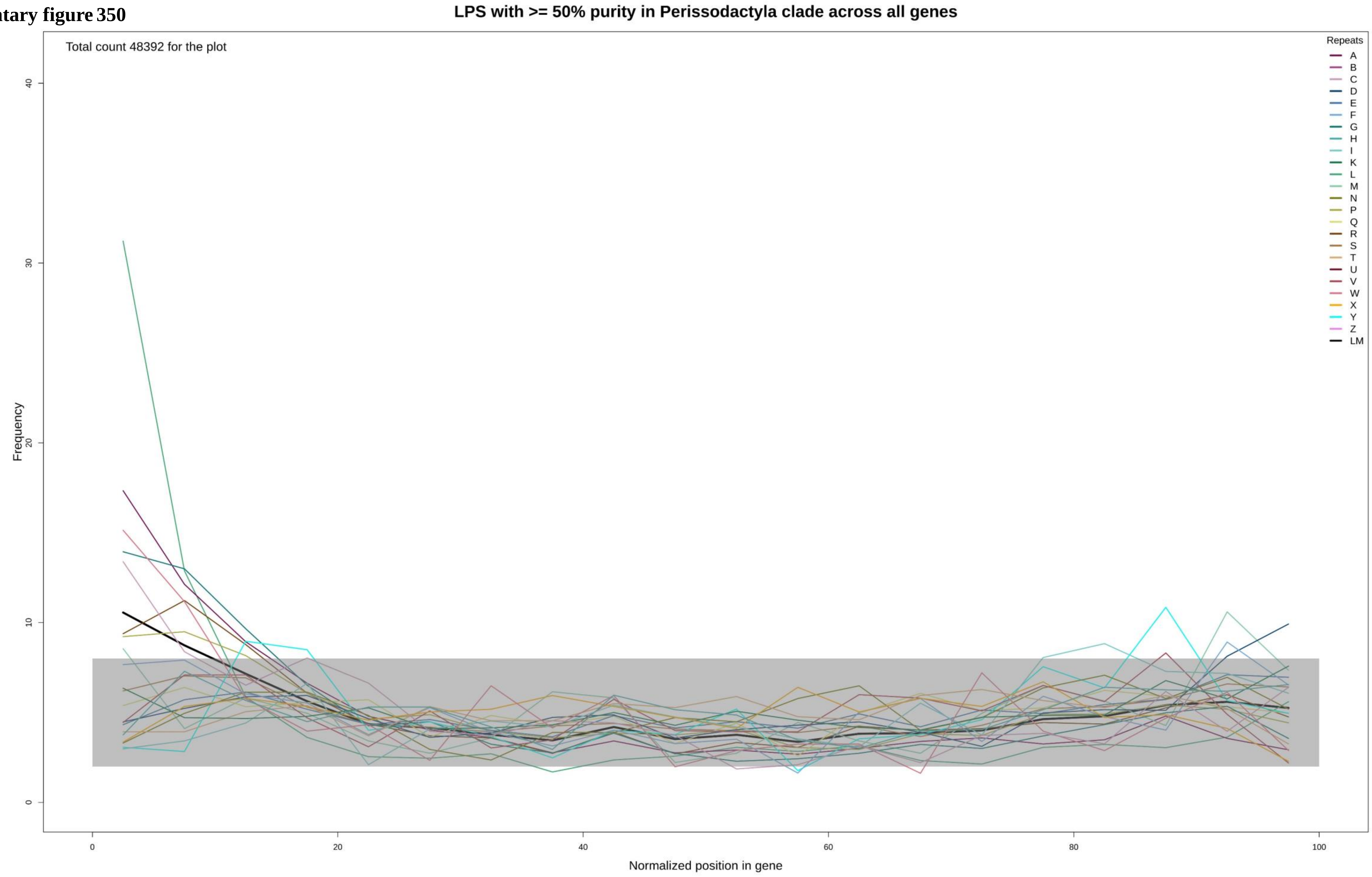

**Supplementary figure 351**

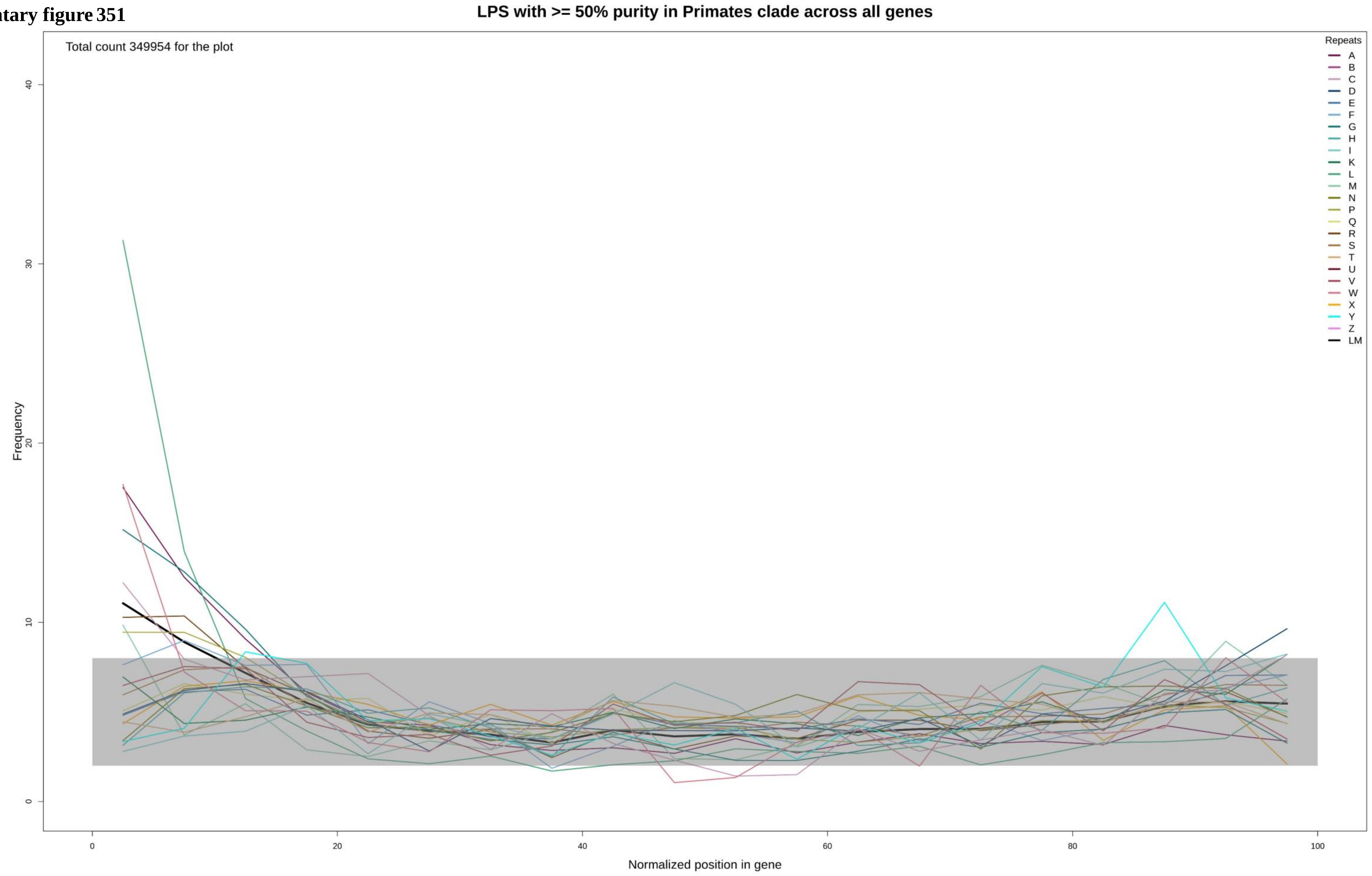

Supplementary figure 352

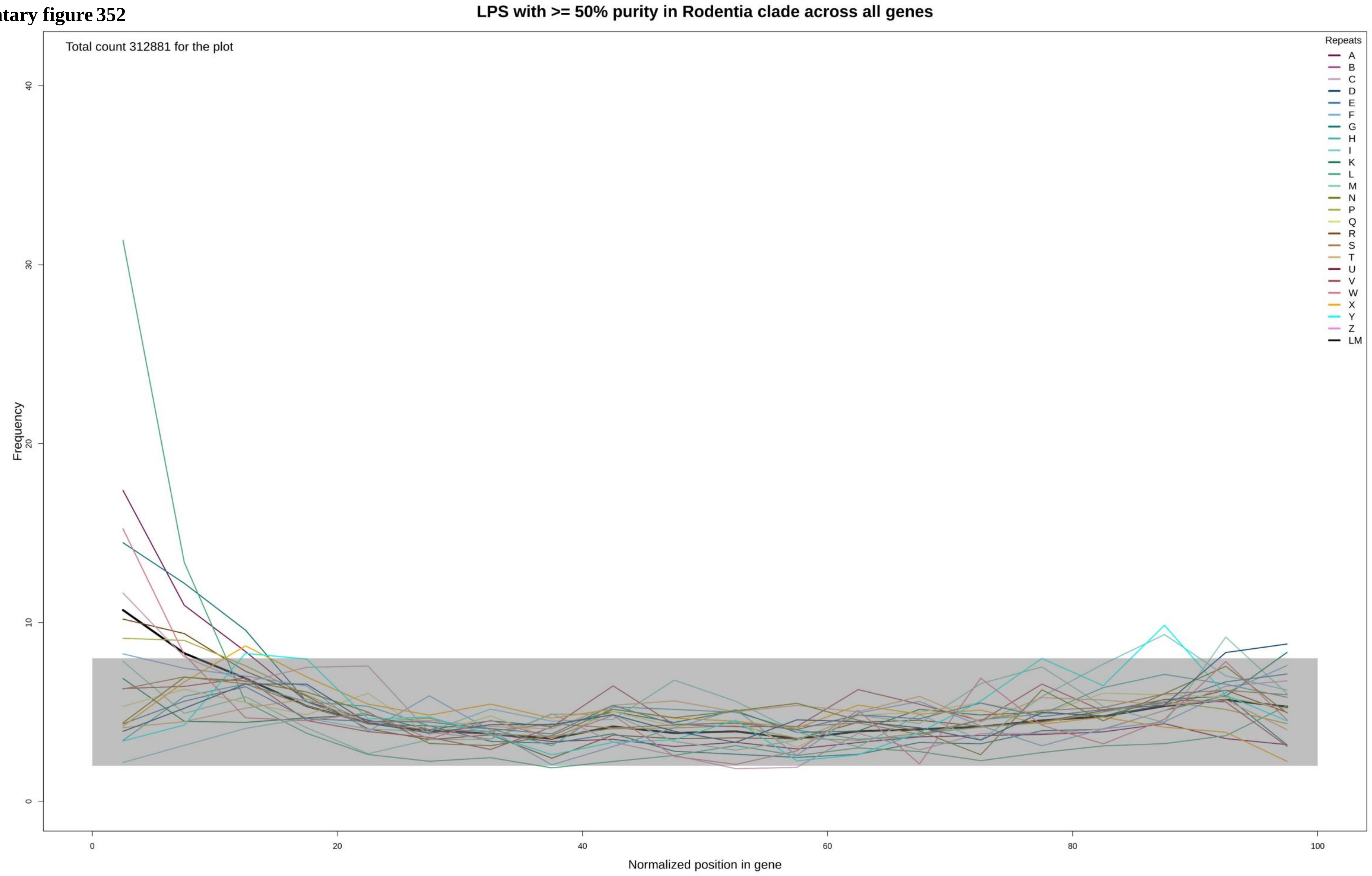

**Supplementary figure 353**

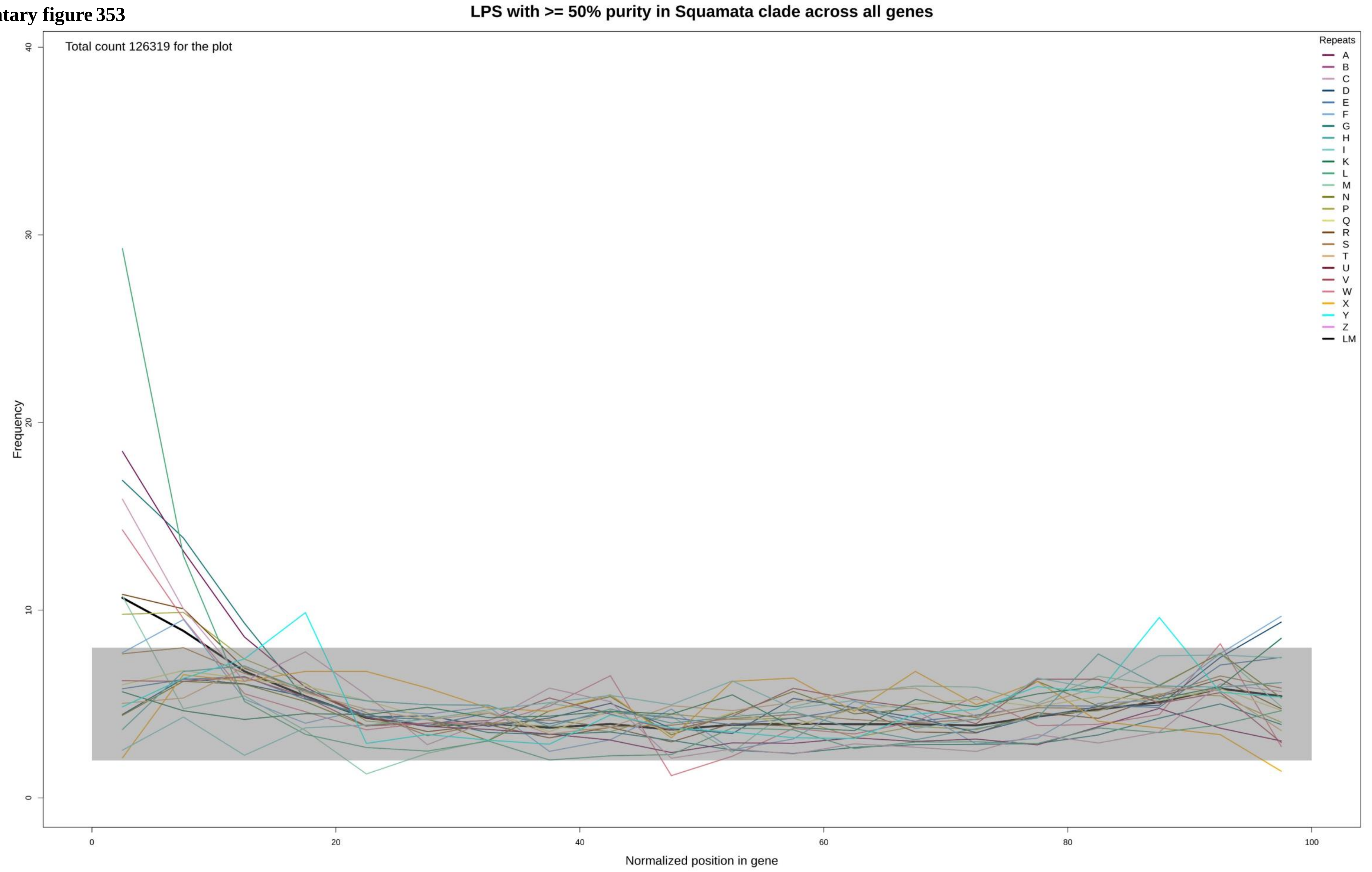

Supplementary figure 354

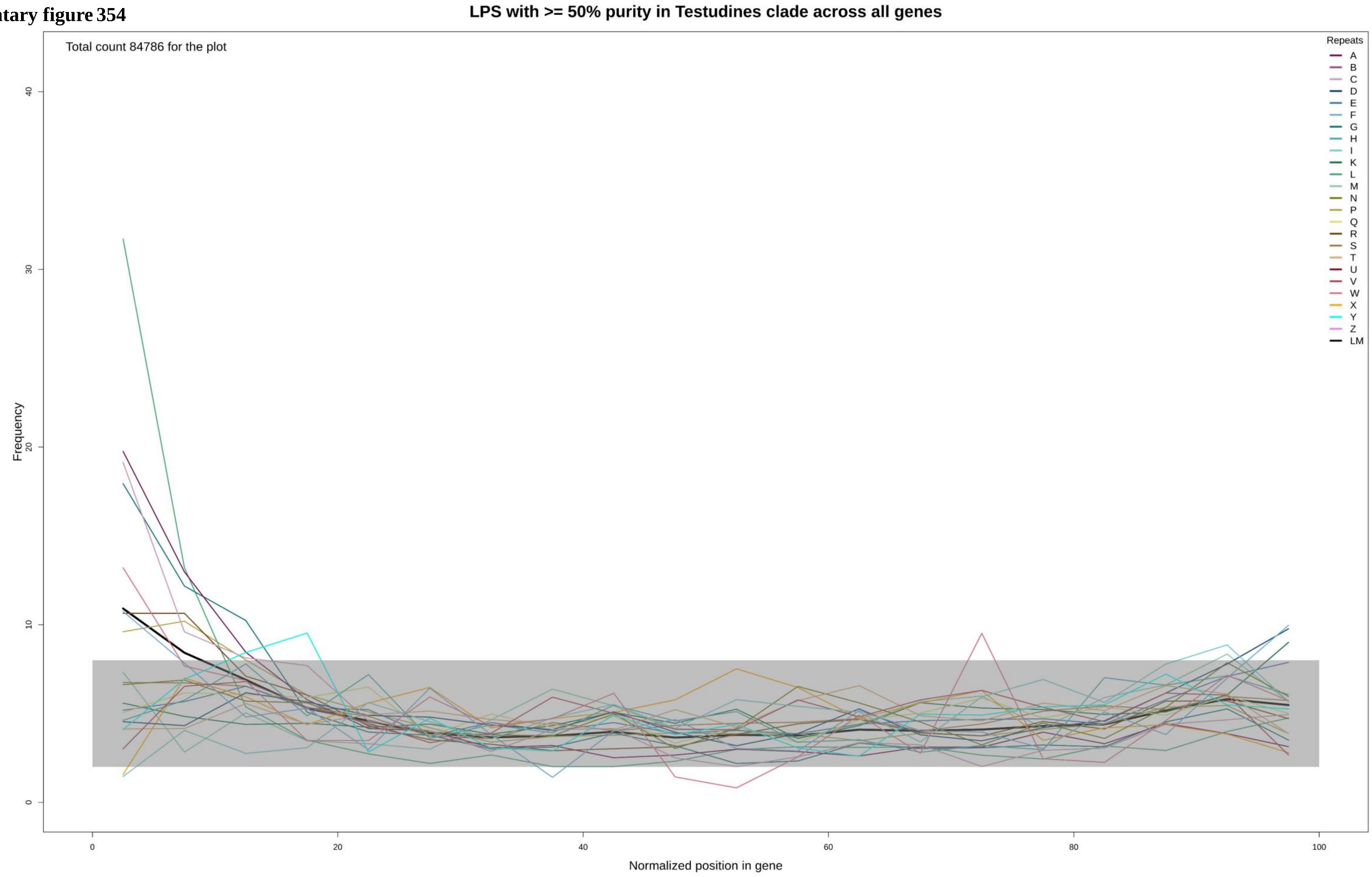

**Supplementary figure 355**

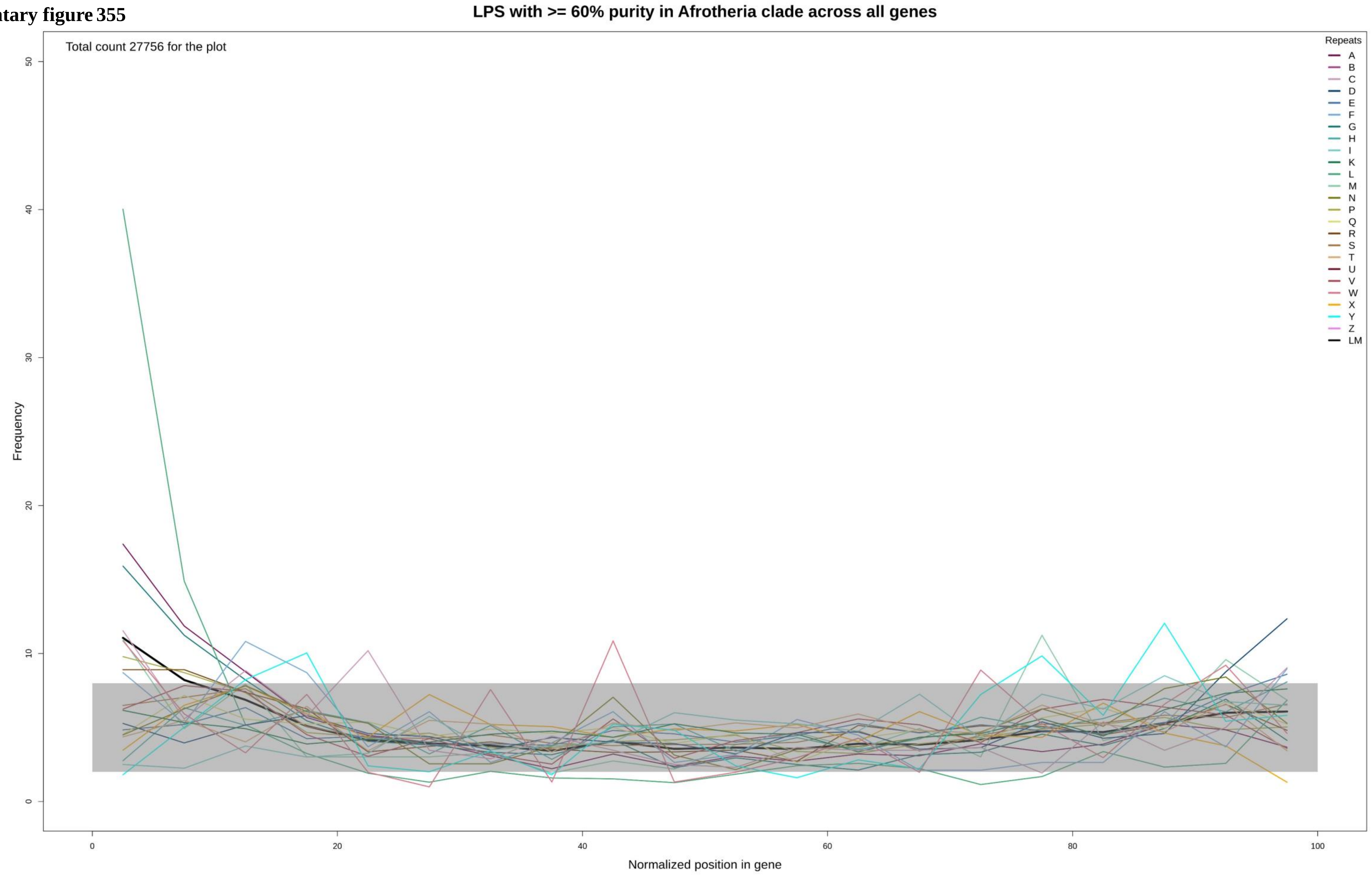

Supplementary figure 356

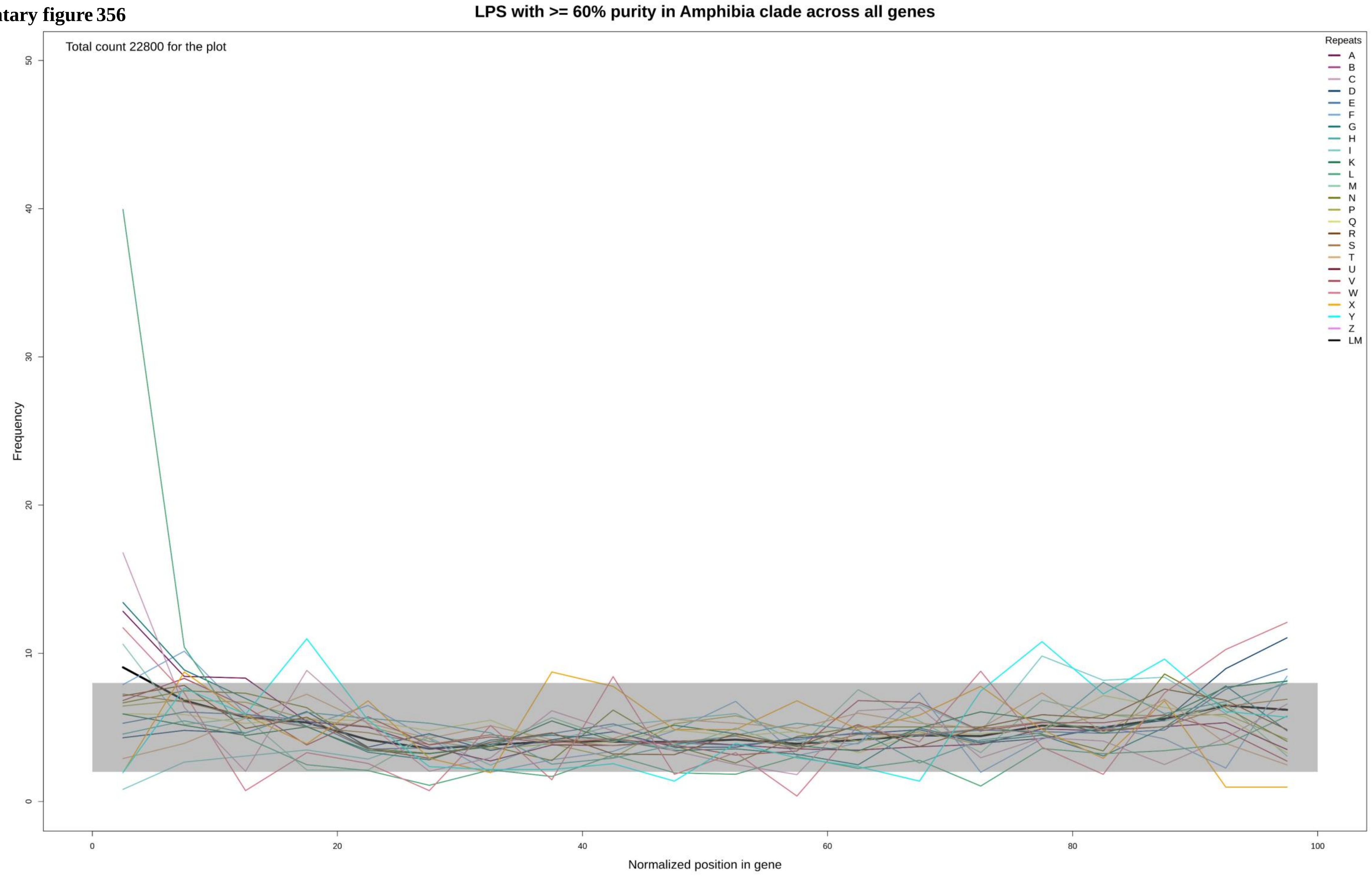

Supplementary figure 357

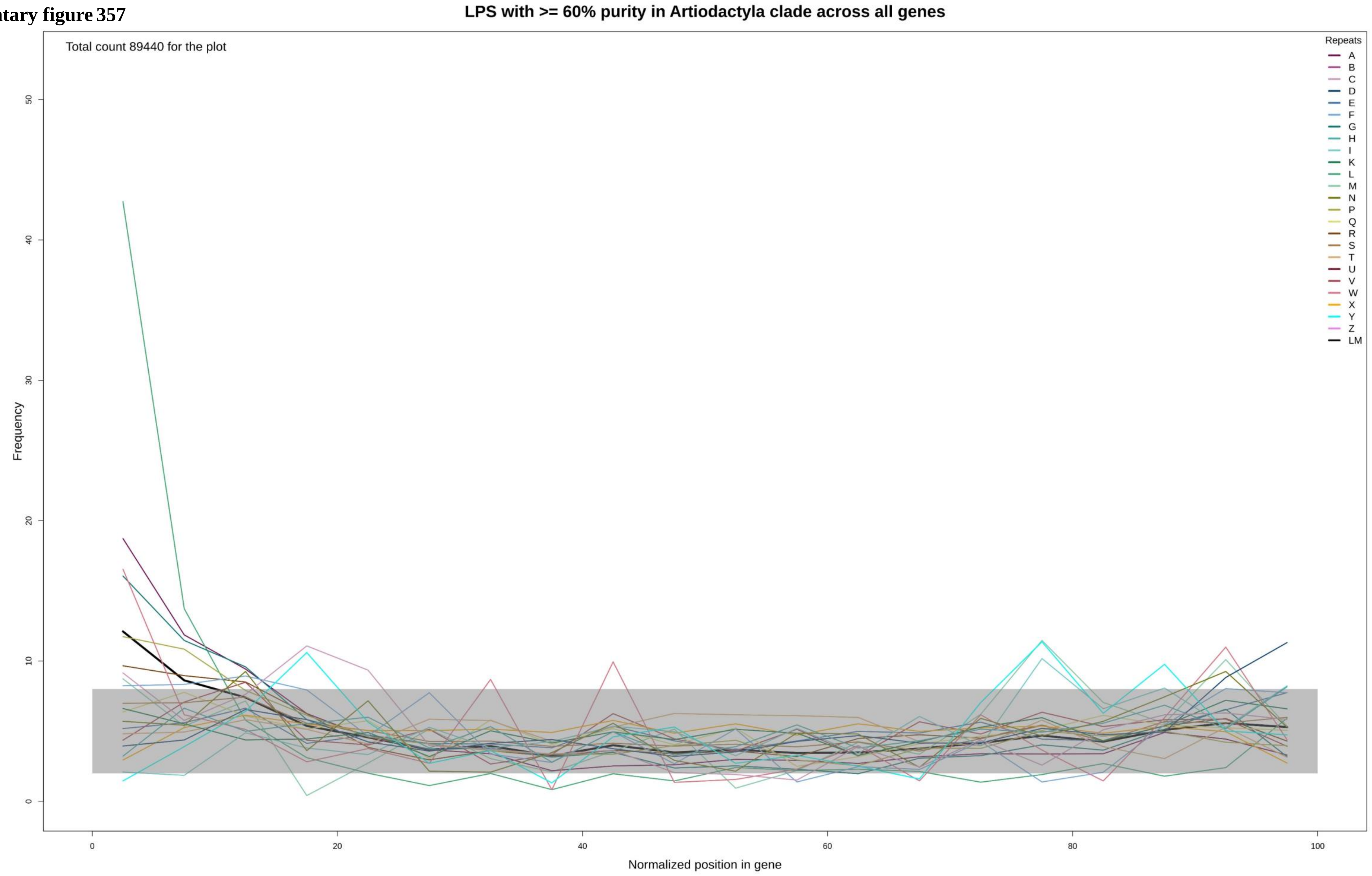

**Supplementary figure 358**

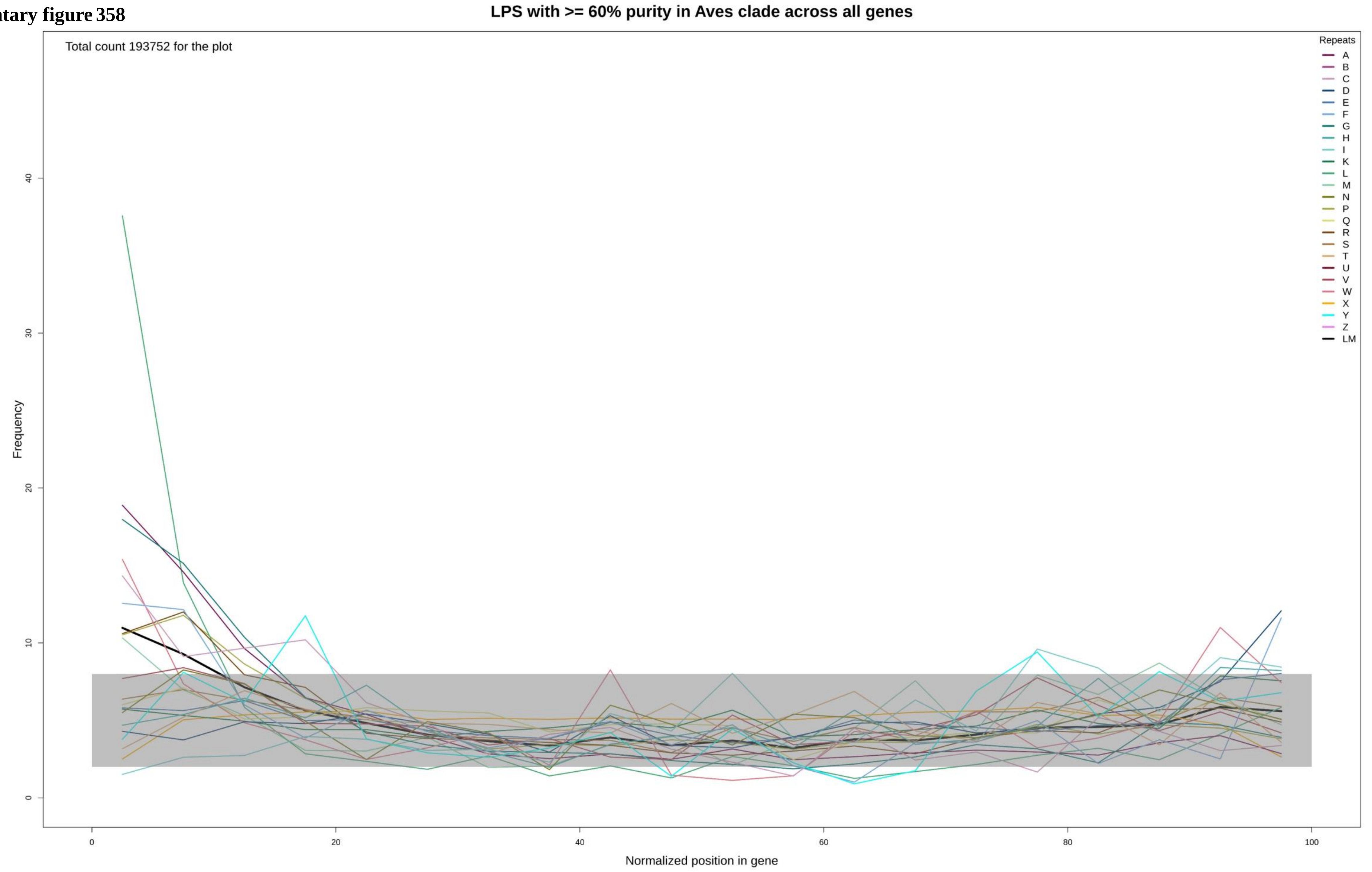

**Supplementary figure 359**

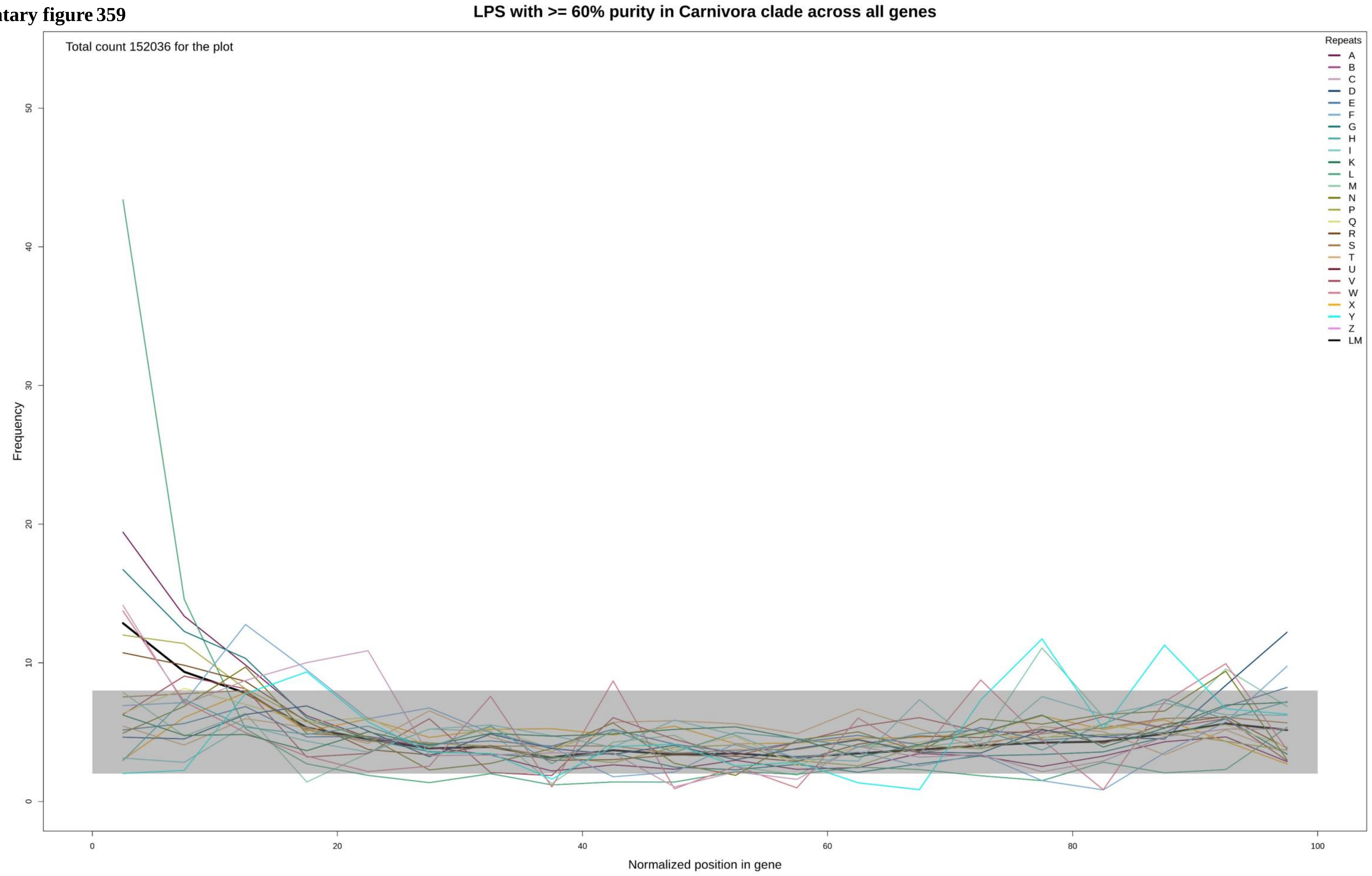

**Supplementary figure 360**

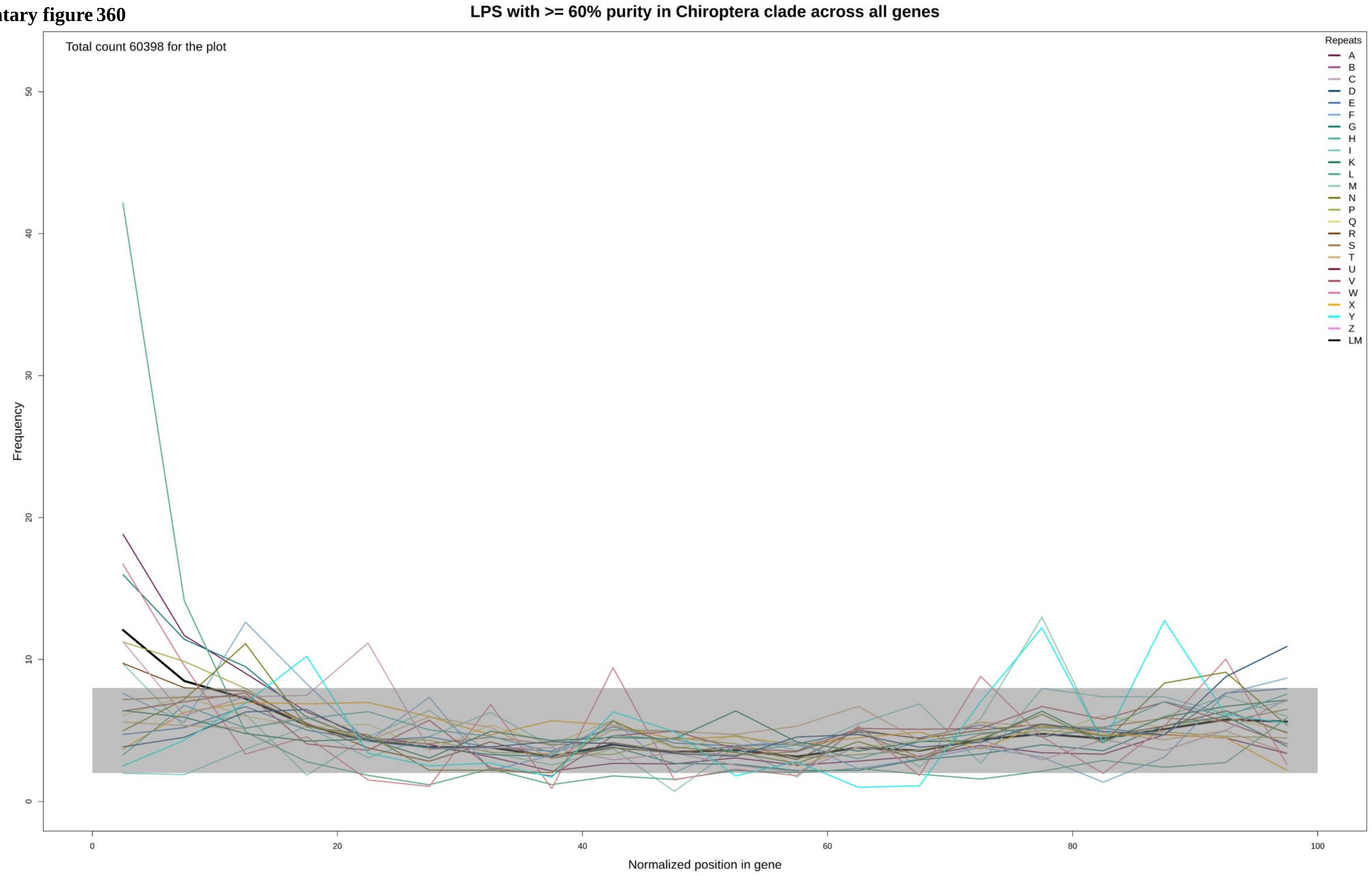

**Supplementary figure 361**

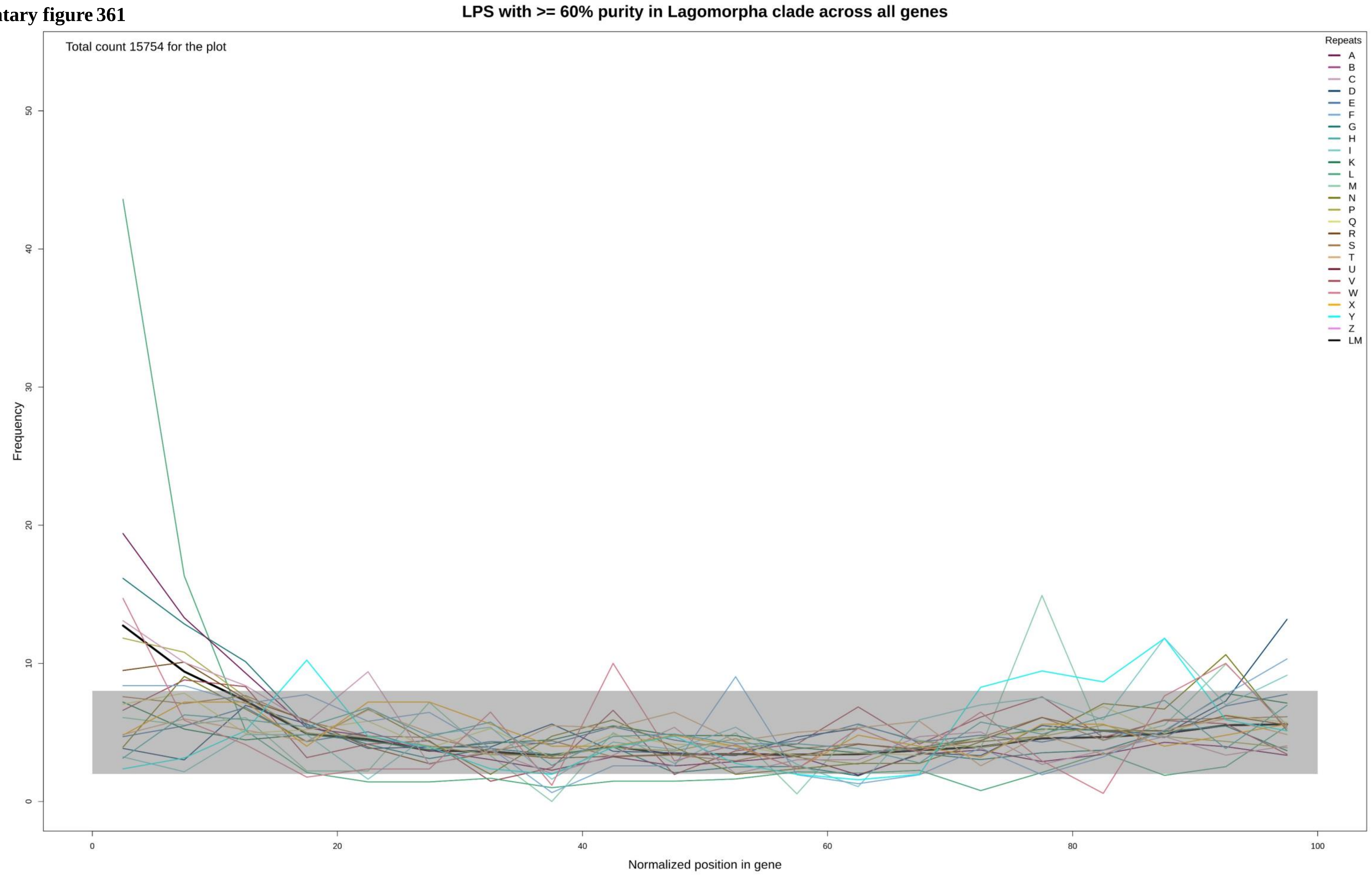

Supplementary figure 362

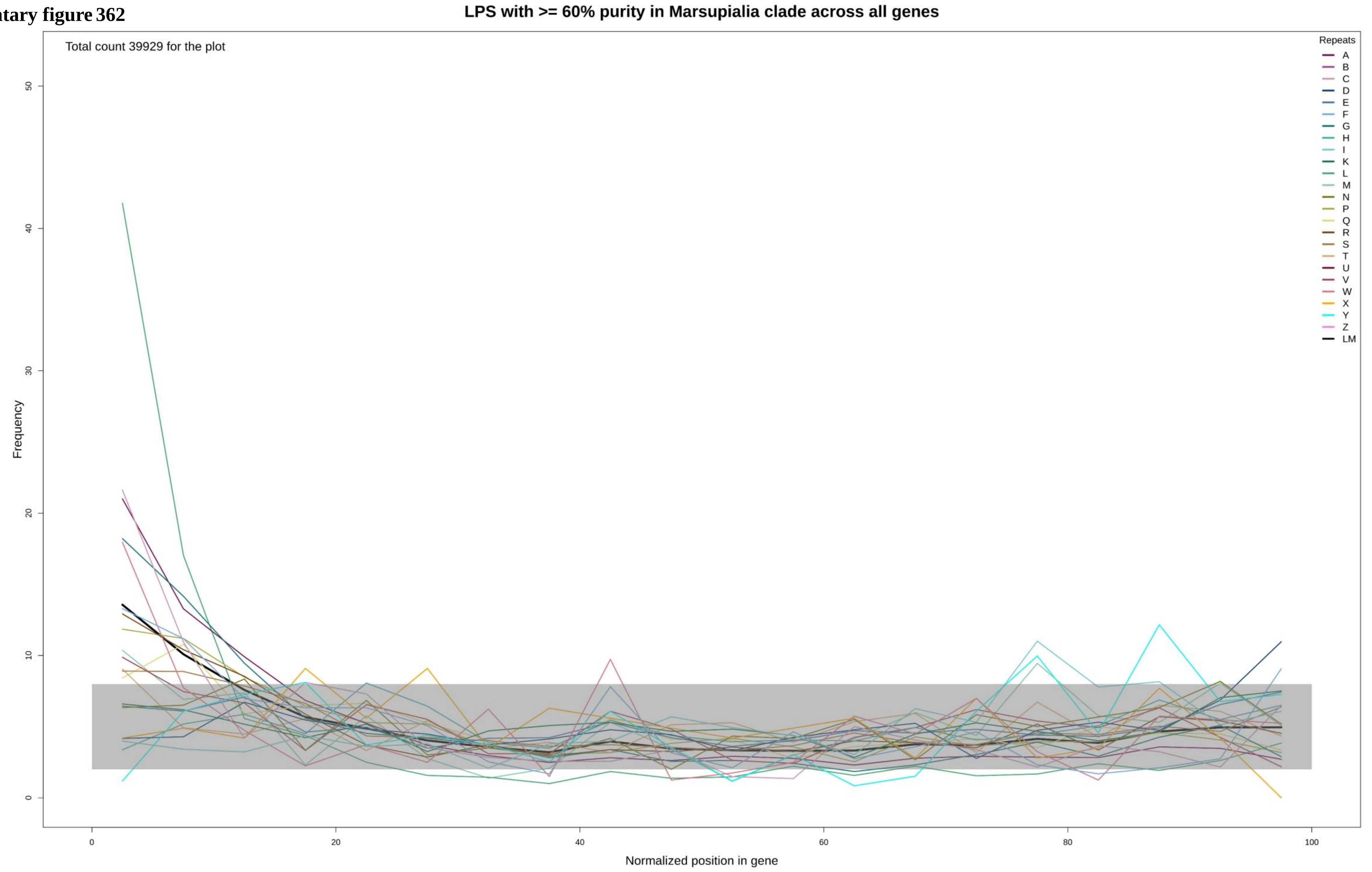

Supplementary figure 363

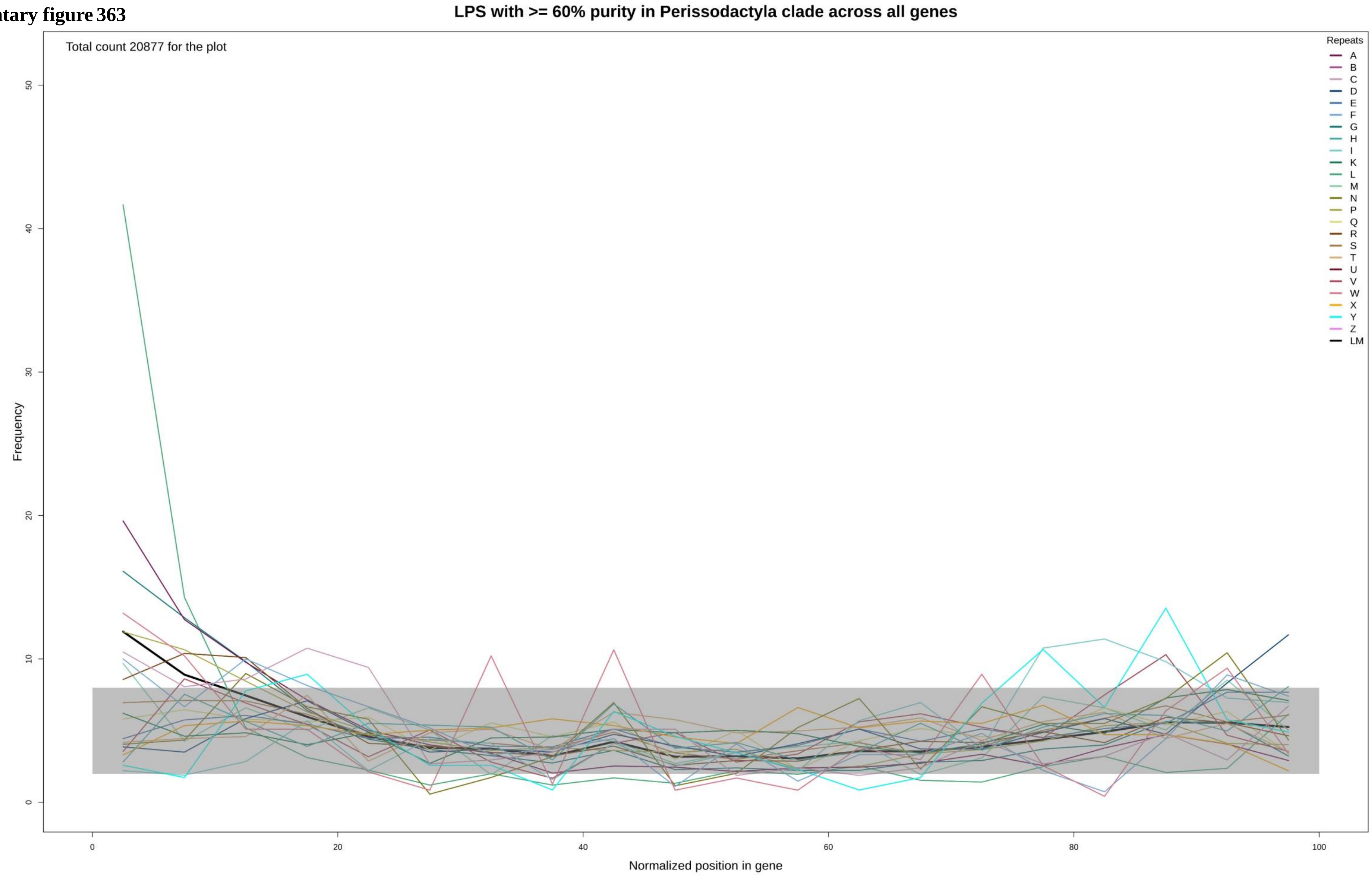

**Supplementary figure 364**

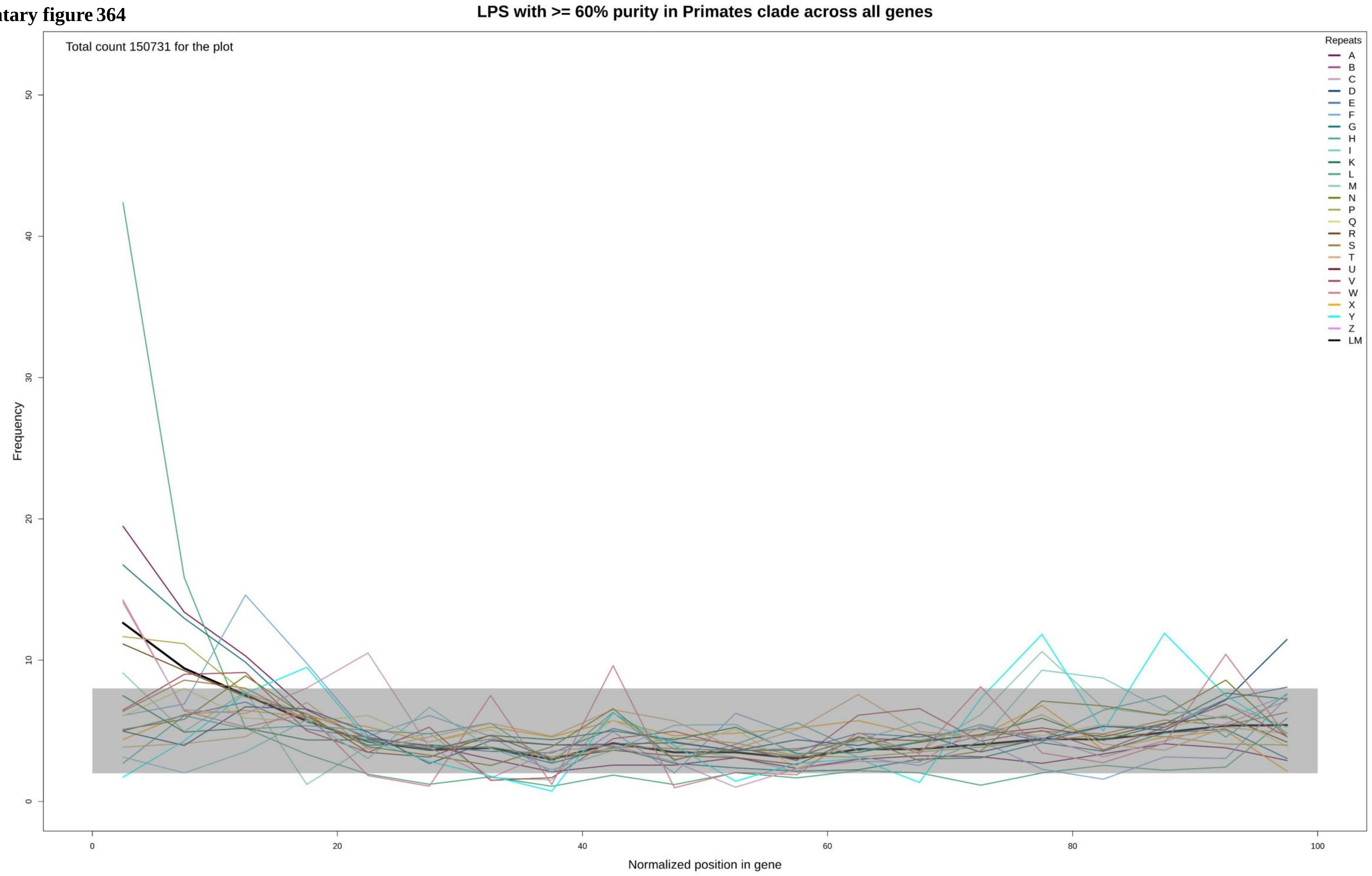

**Supplementary figure 365**

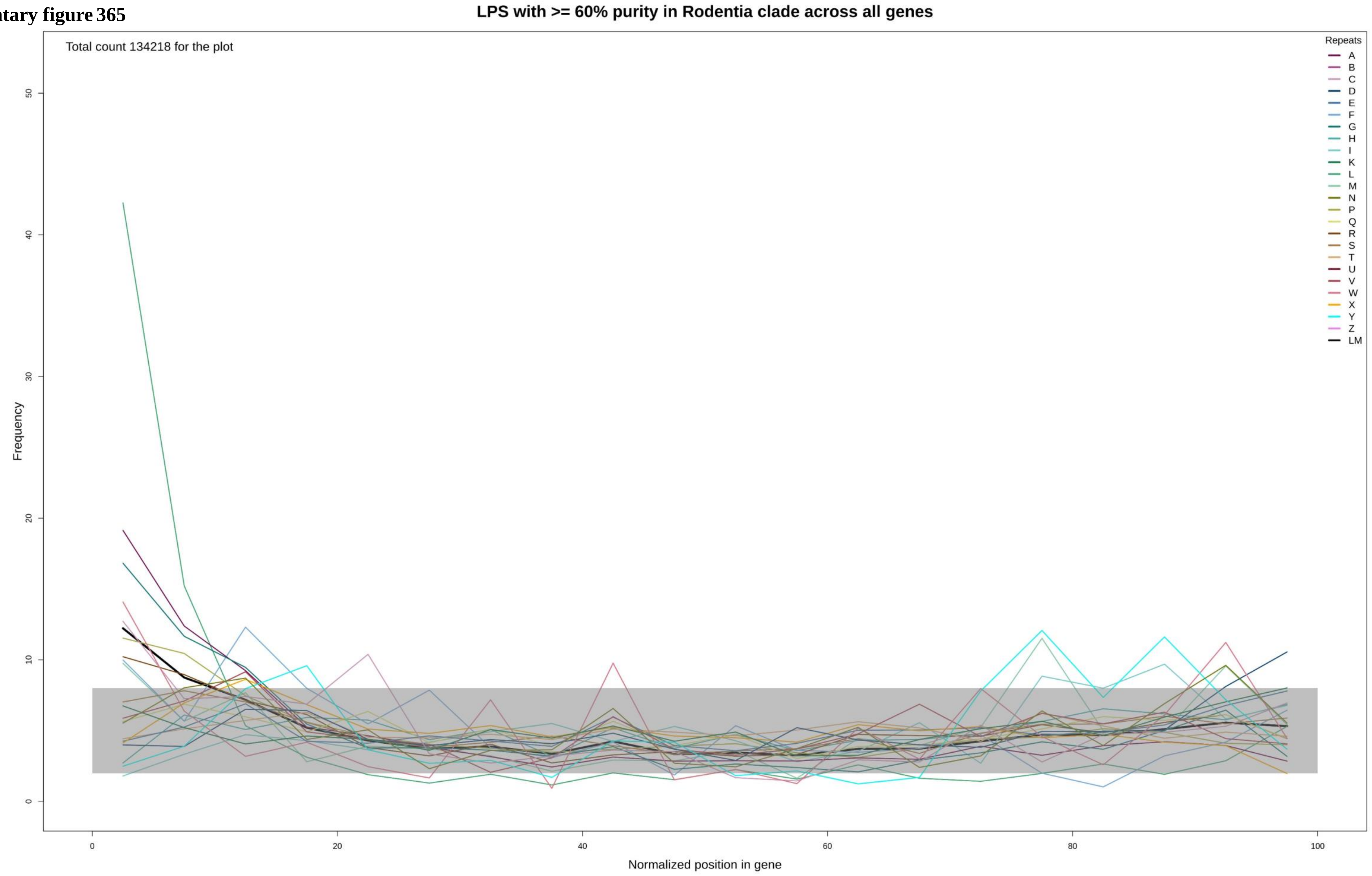

**Supplementary figure 366**

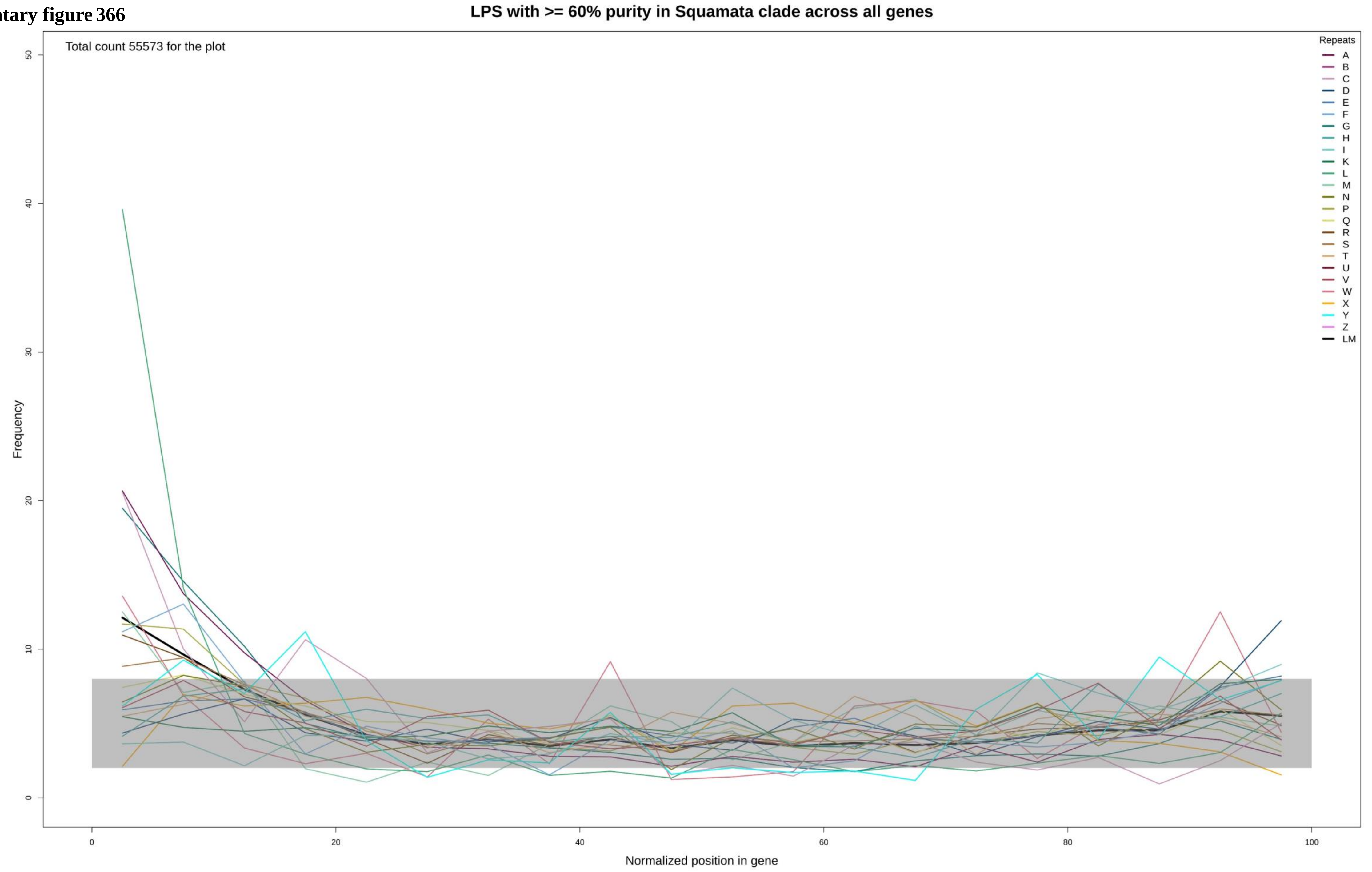

**Supplementary figure 367**

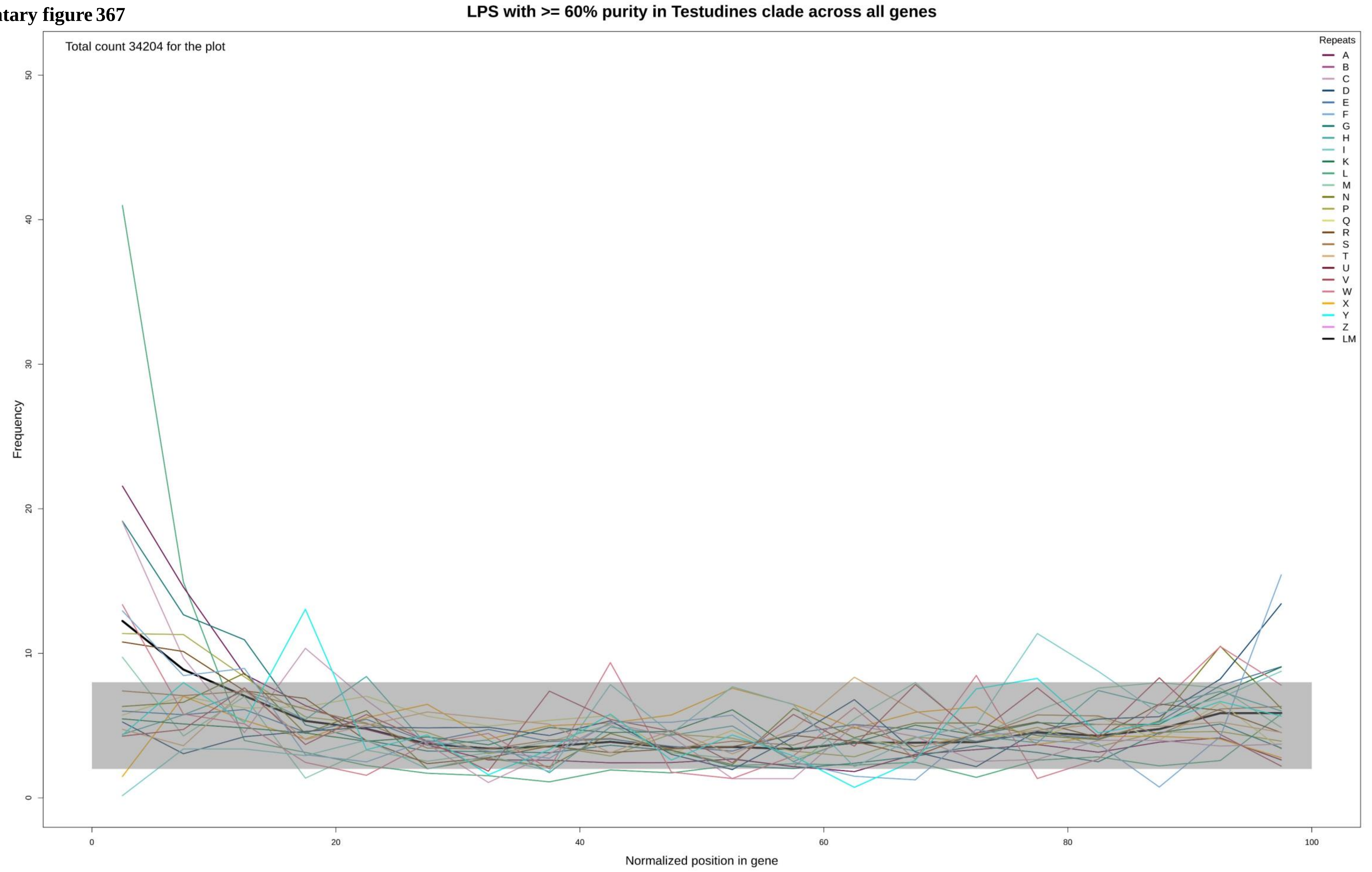

**Supplementary figure 368**

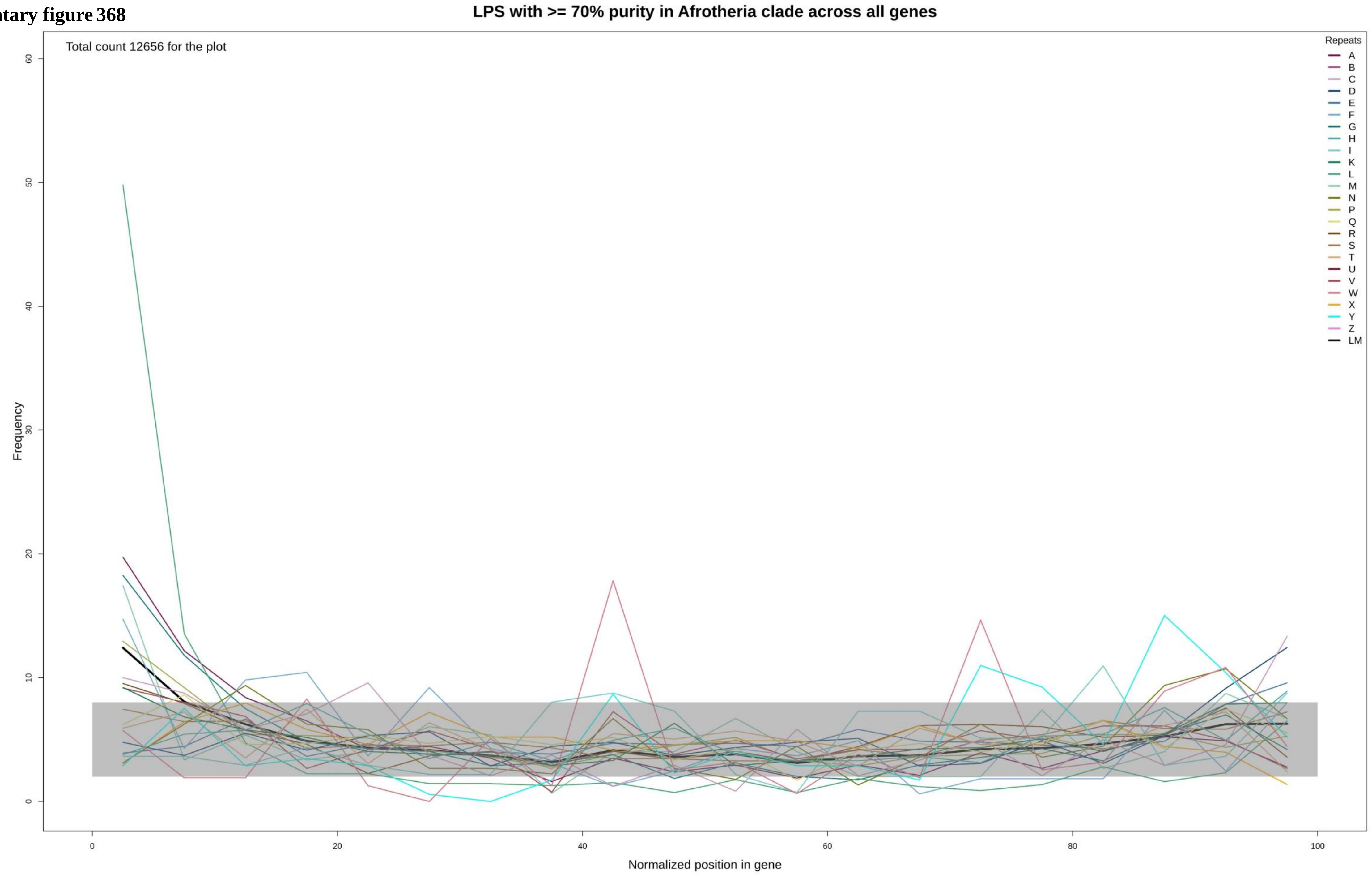

Supplementary figure 369

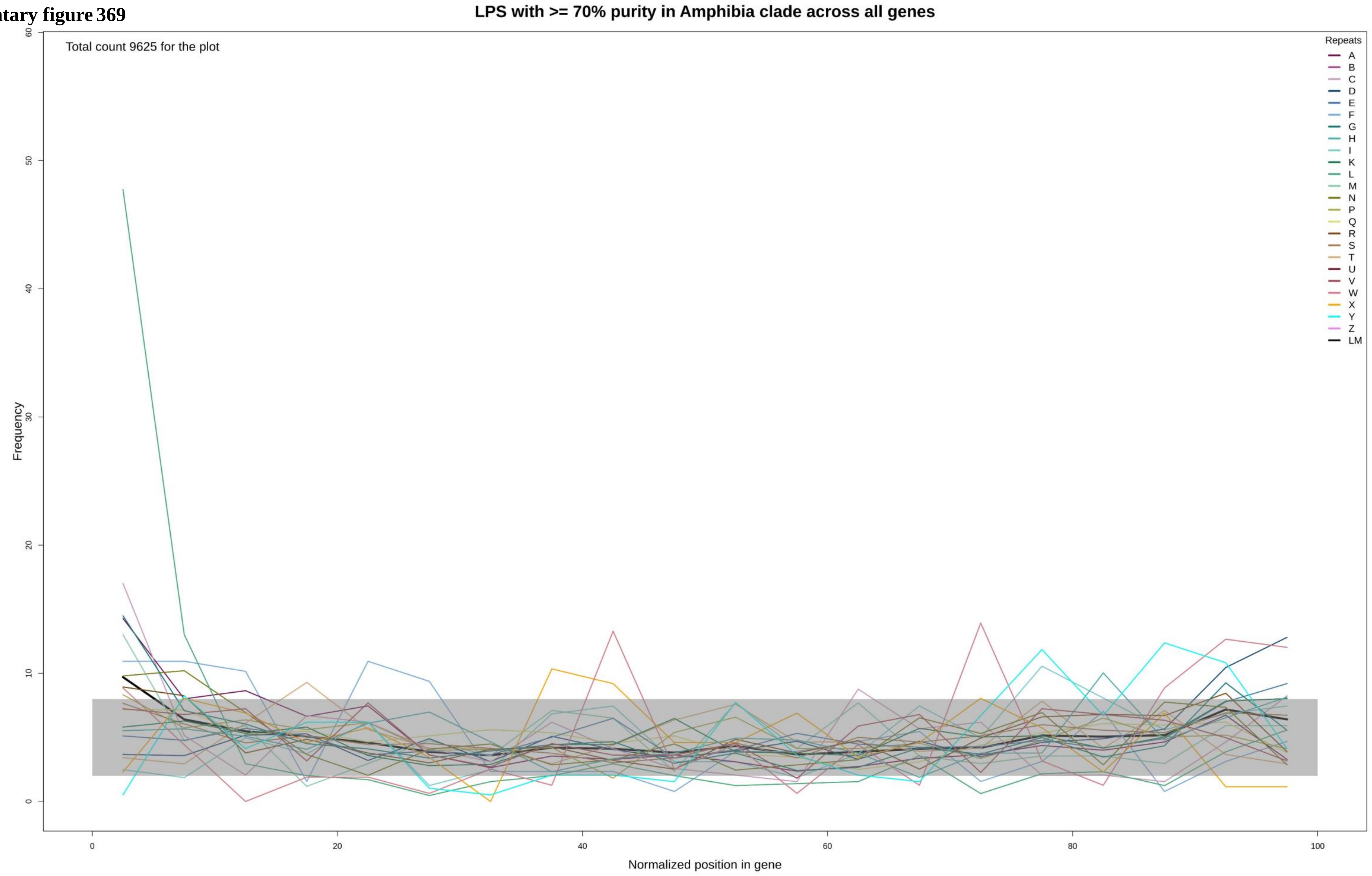

**Supplementary figure 370**

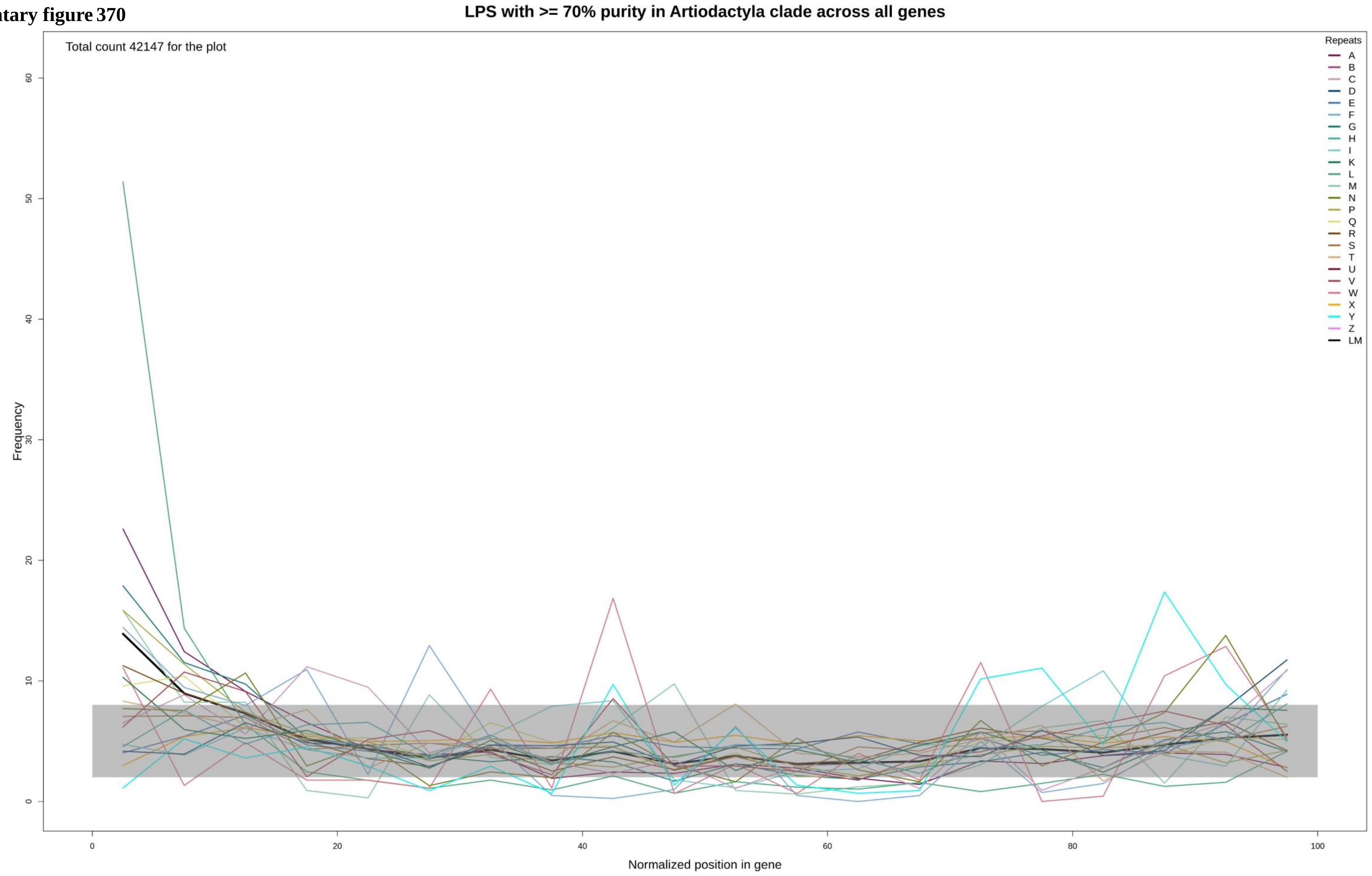

Supplementary figure 371

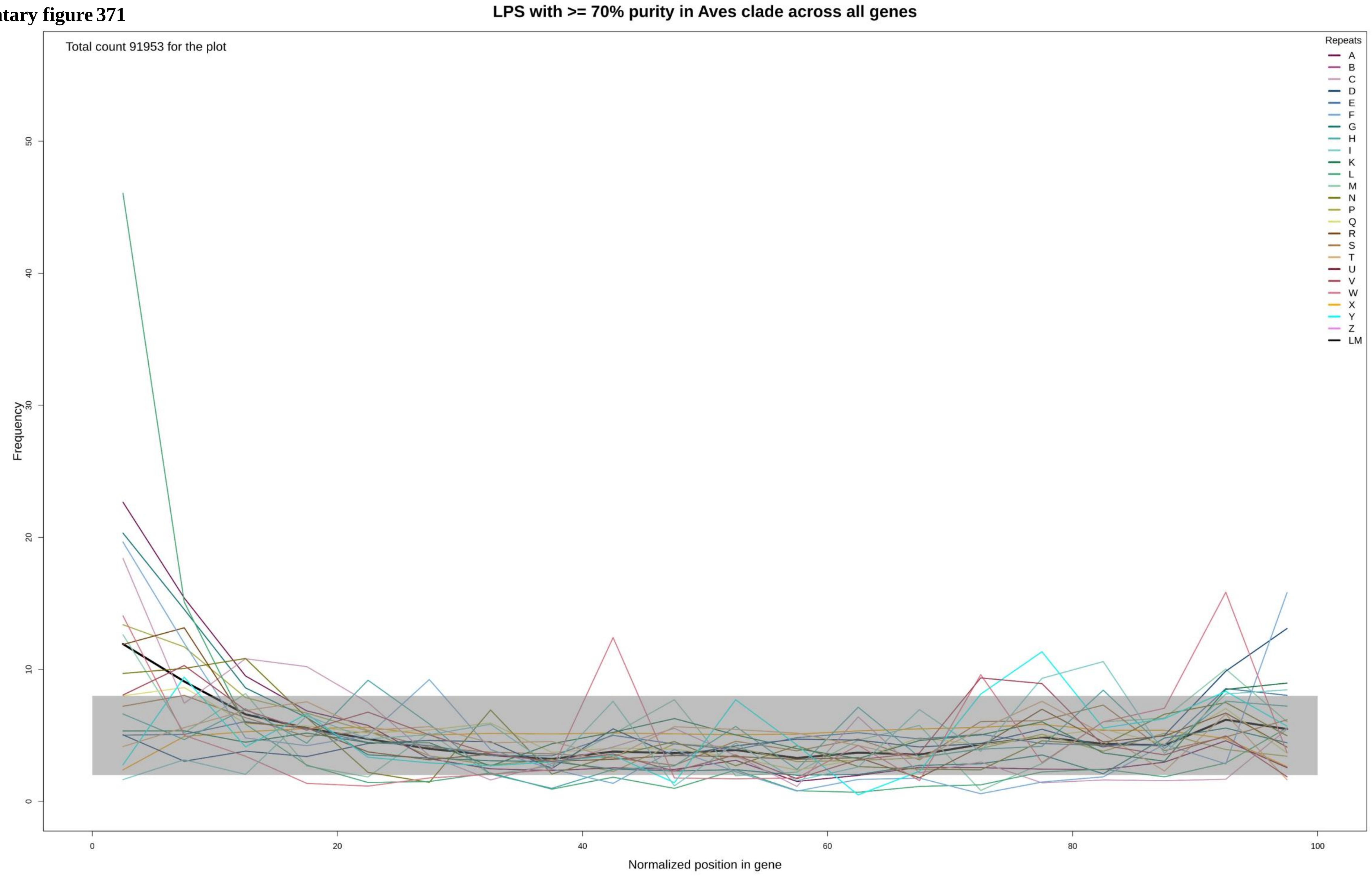

**Supplementary figure 372**

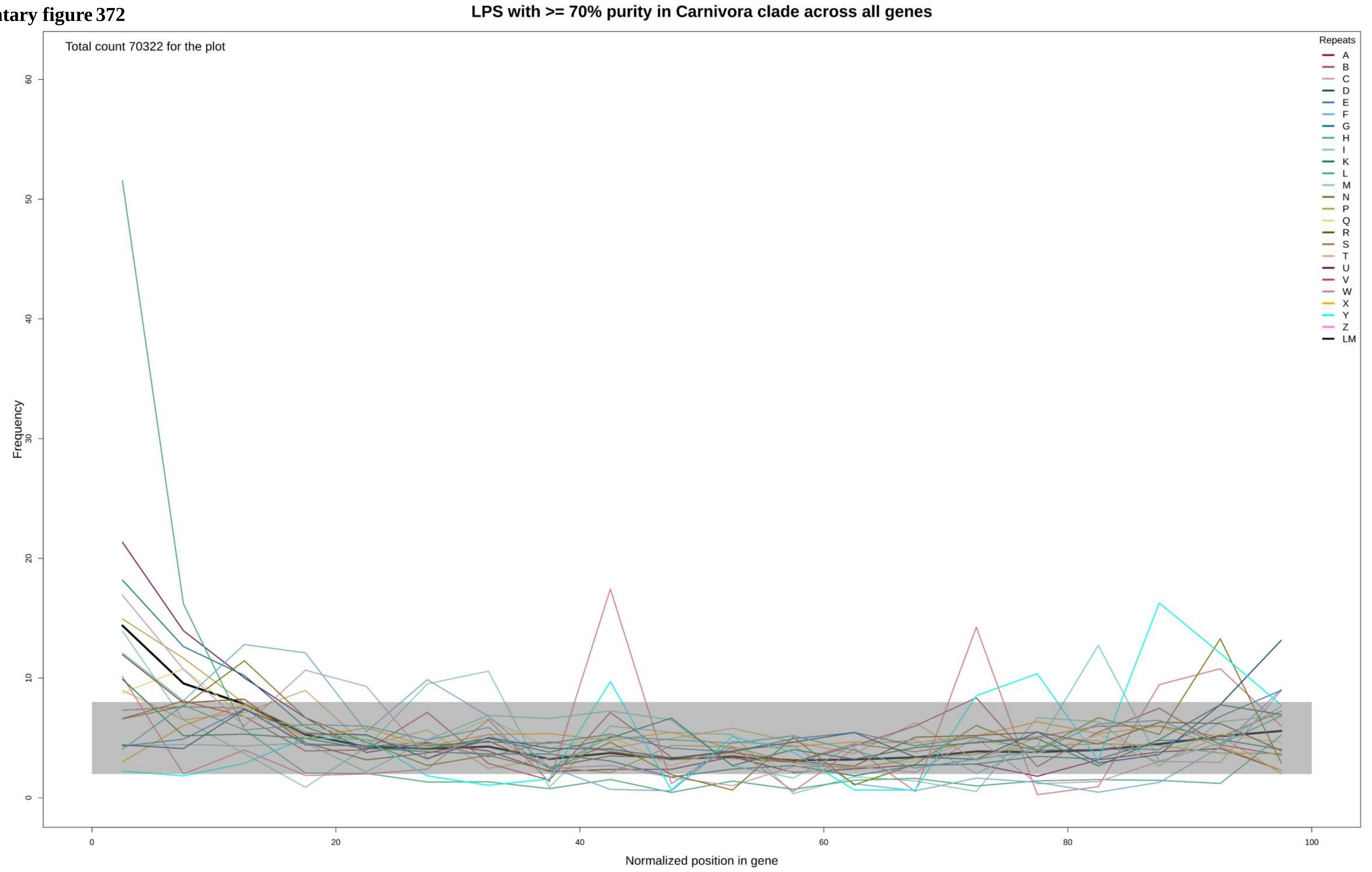

**Supplementary figure 373**

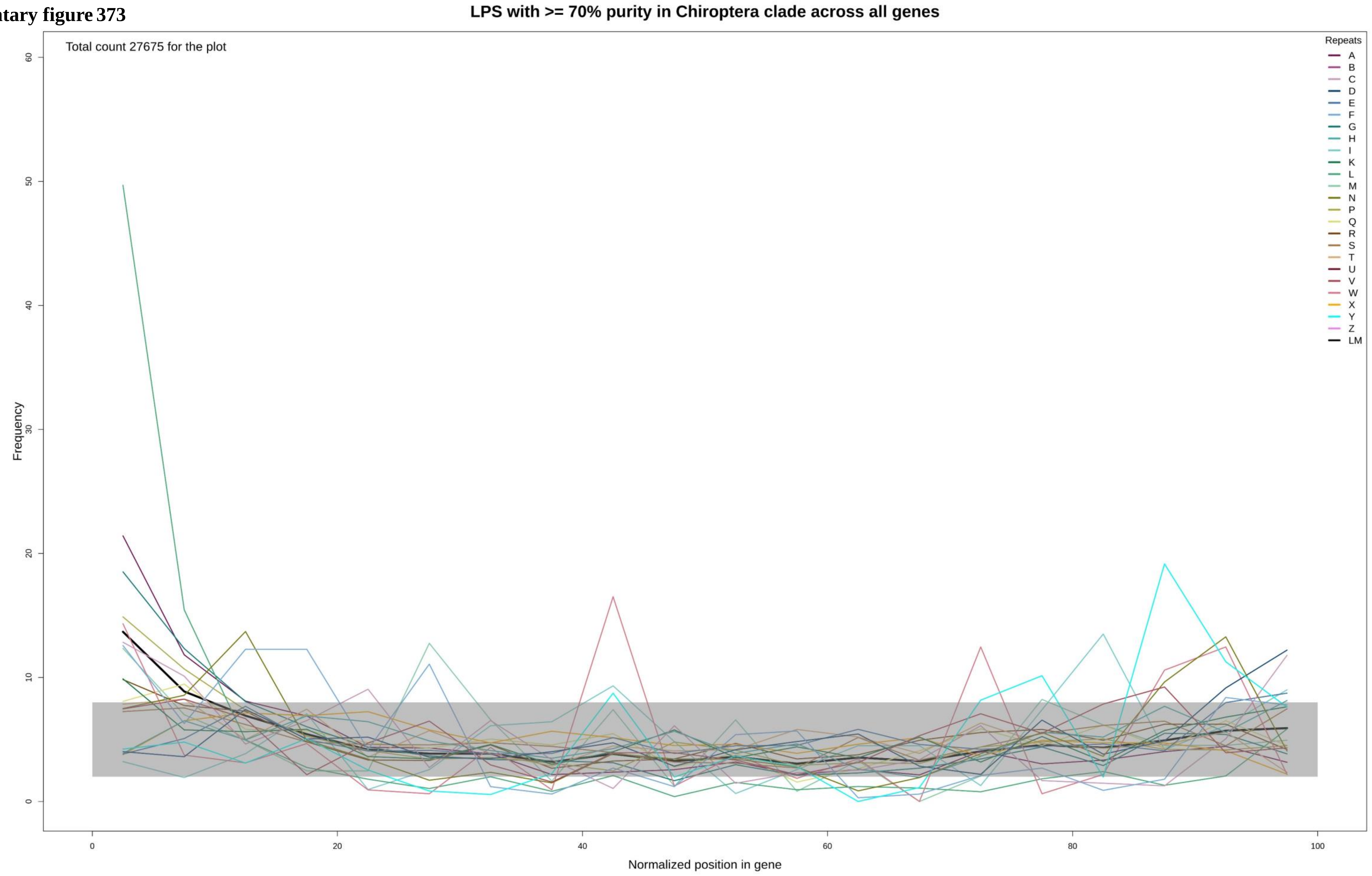

**Supplementary figure 374**

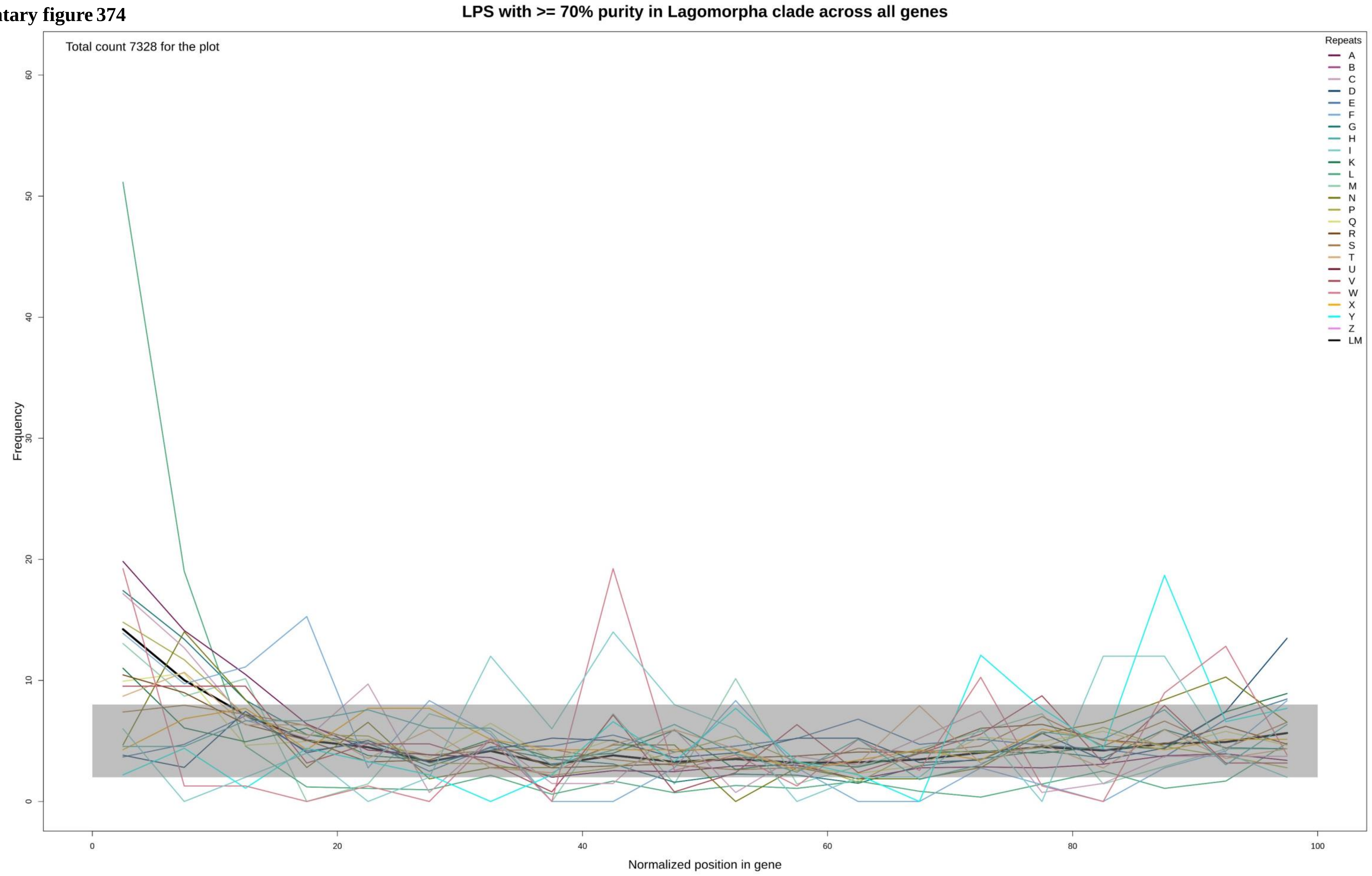

**Supplementary figure 375**

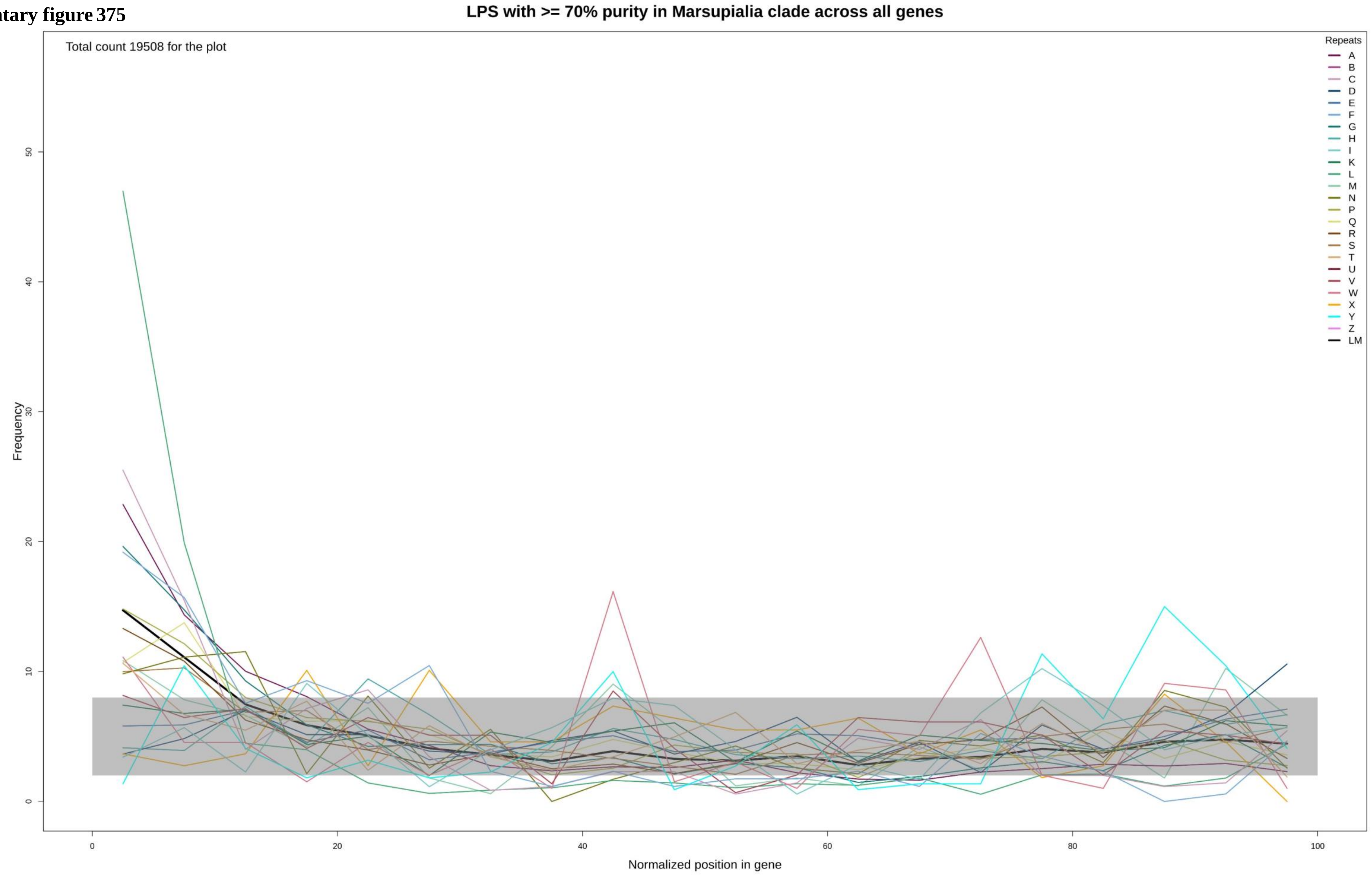

**Supplementary figure 376**

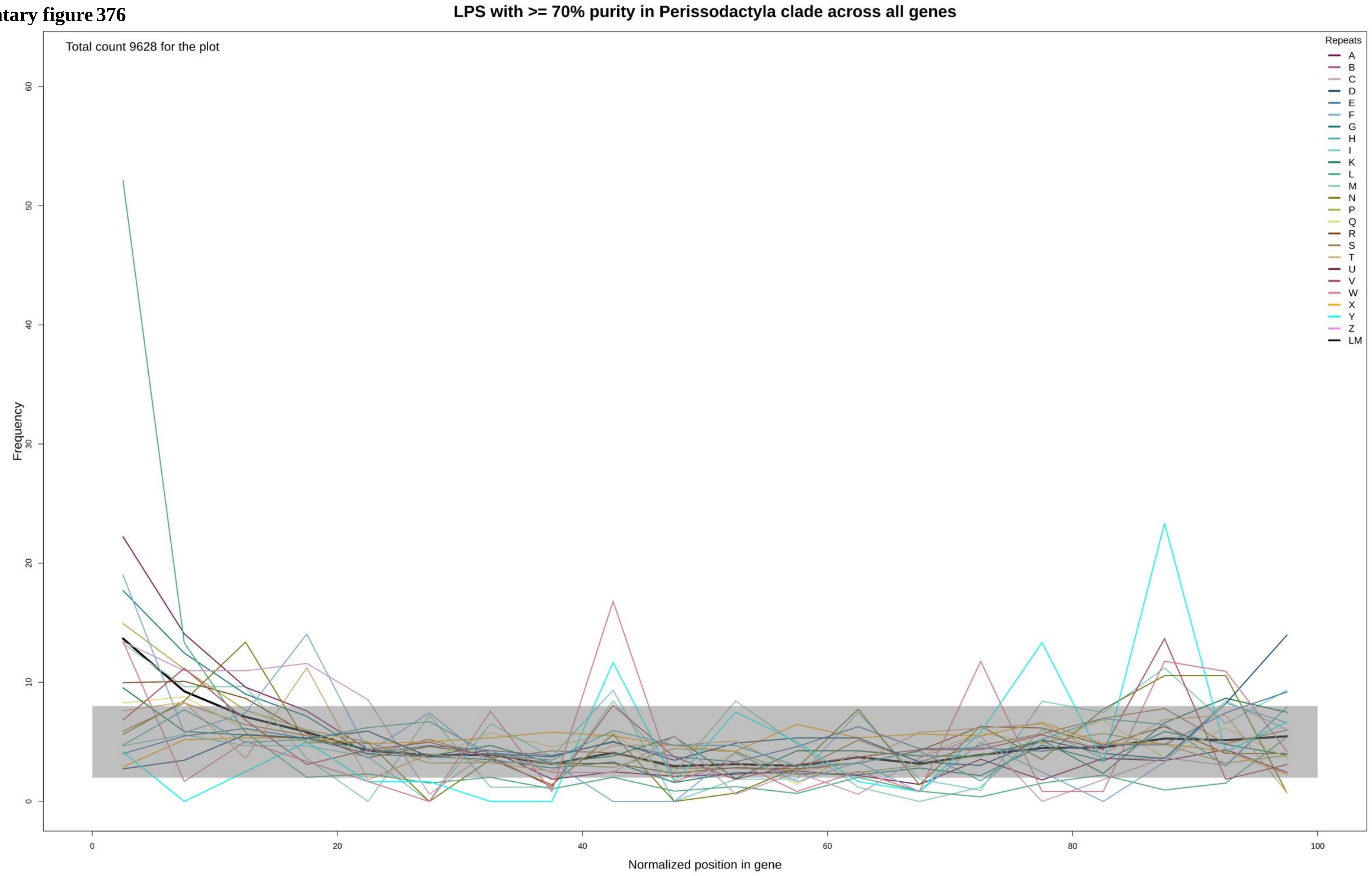

**Supplementary figure 377**

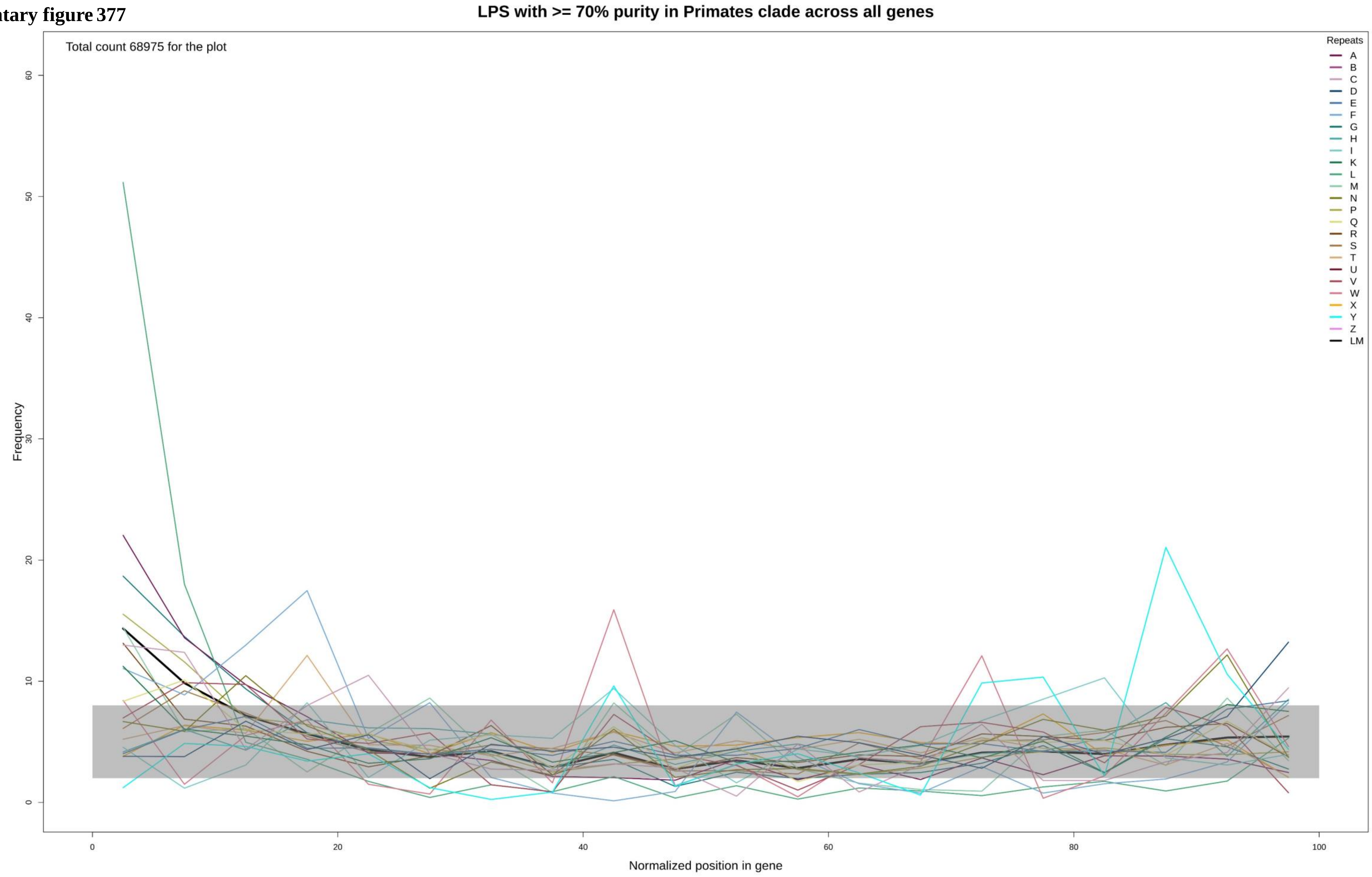

Supplementary figure 378

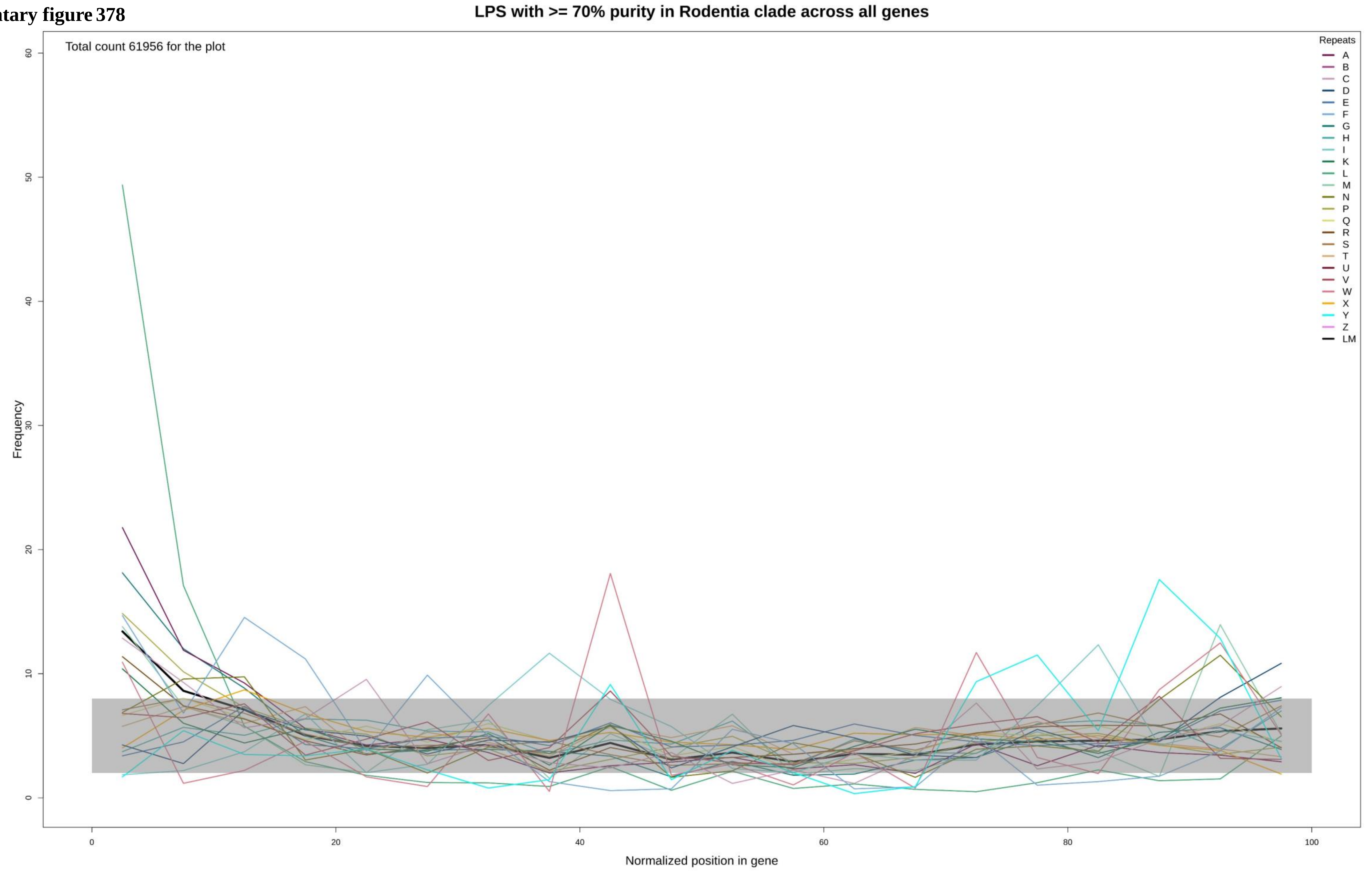

**Supplementary figure 379**

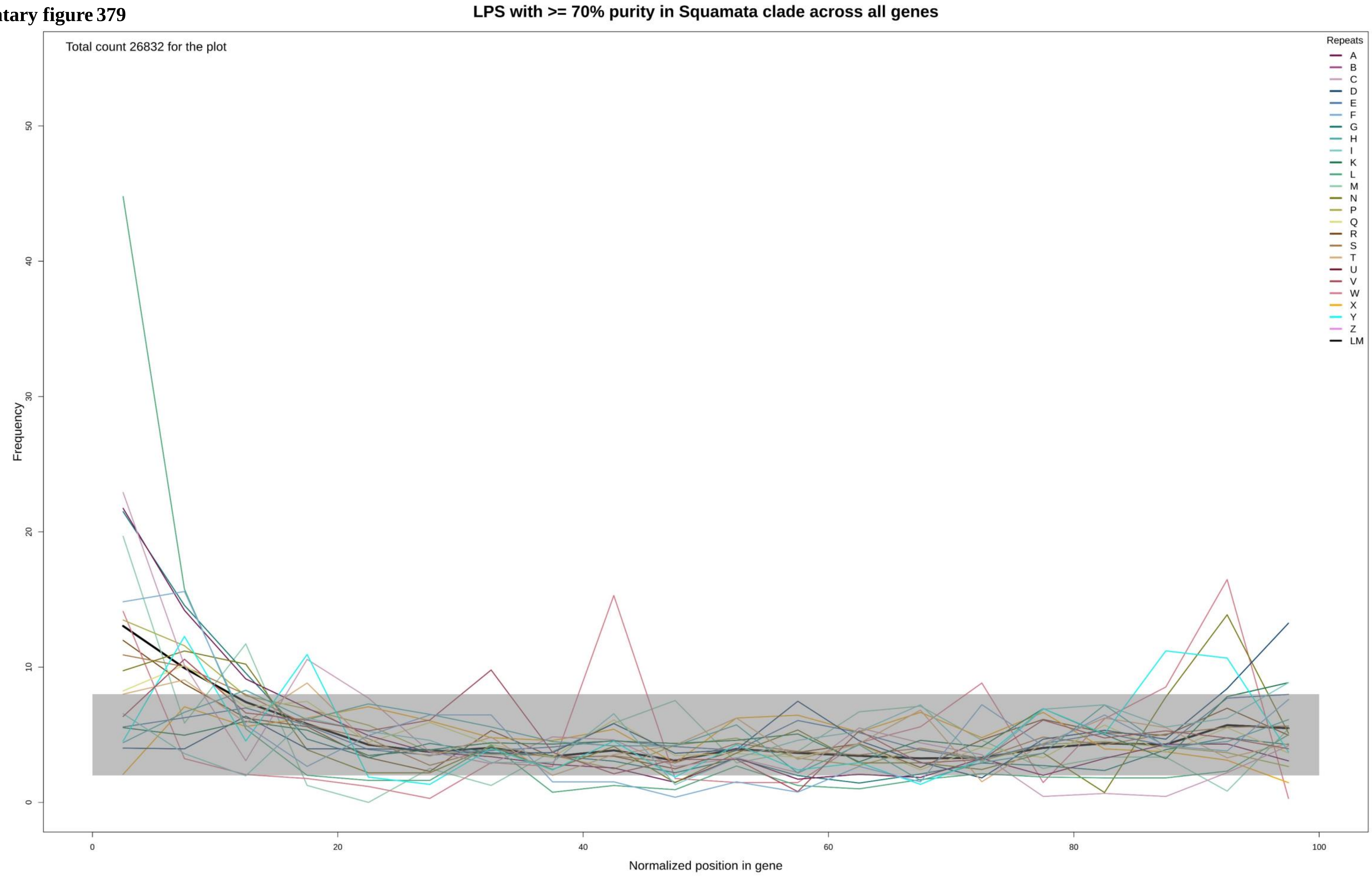

**Supplementary figure 380**

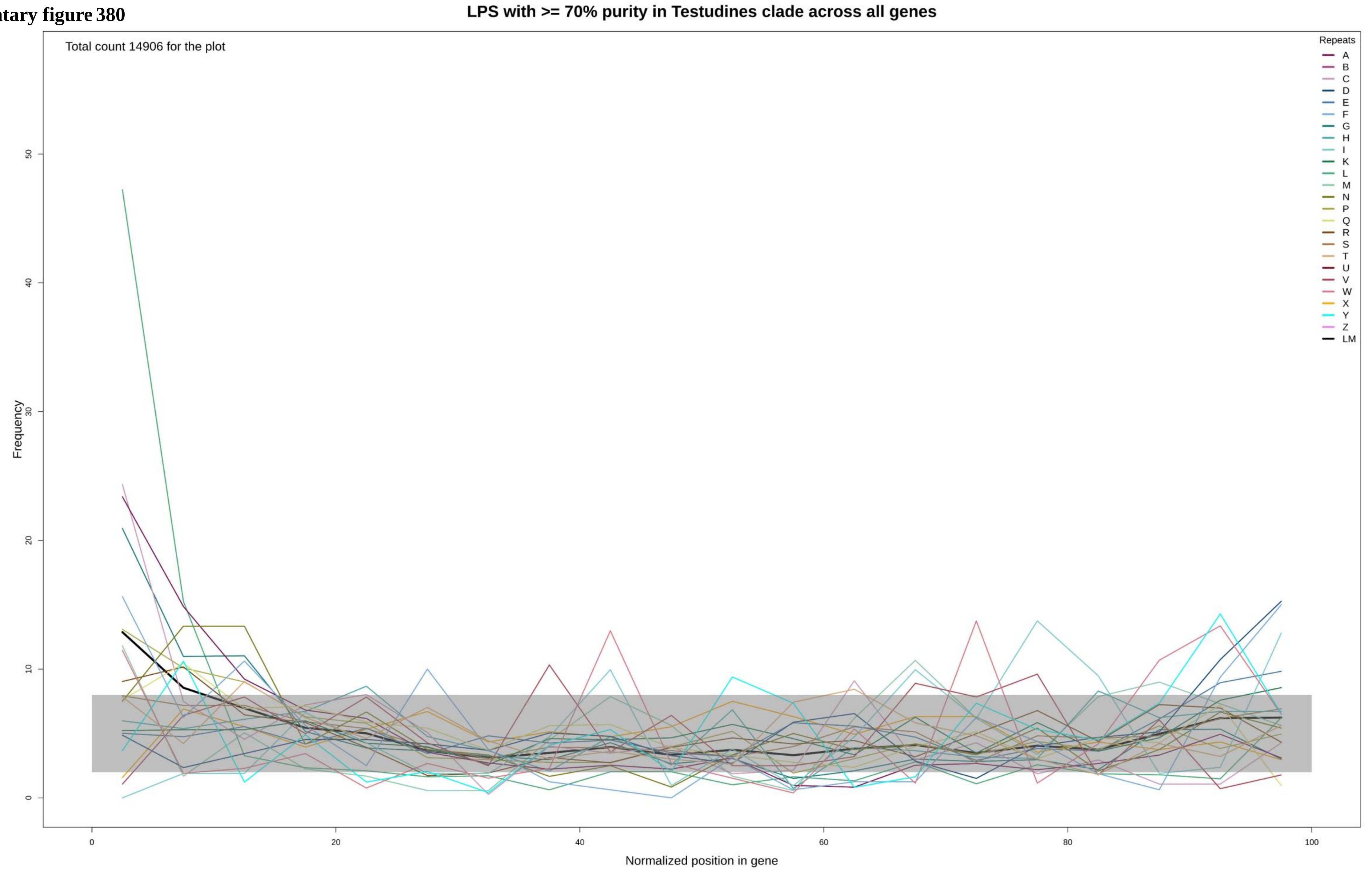

Supplementary figure 381

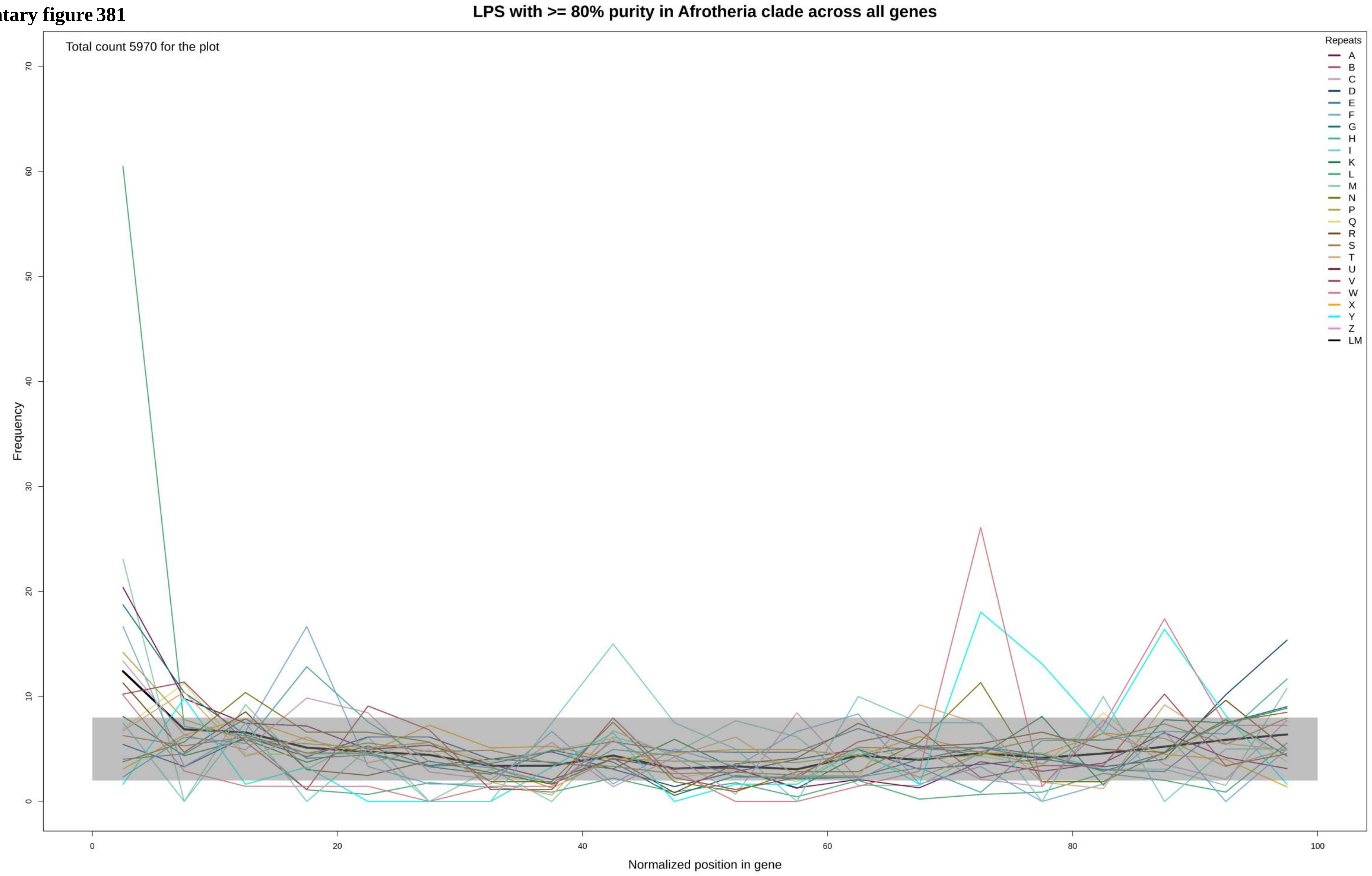

Supplementary figure 382

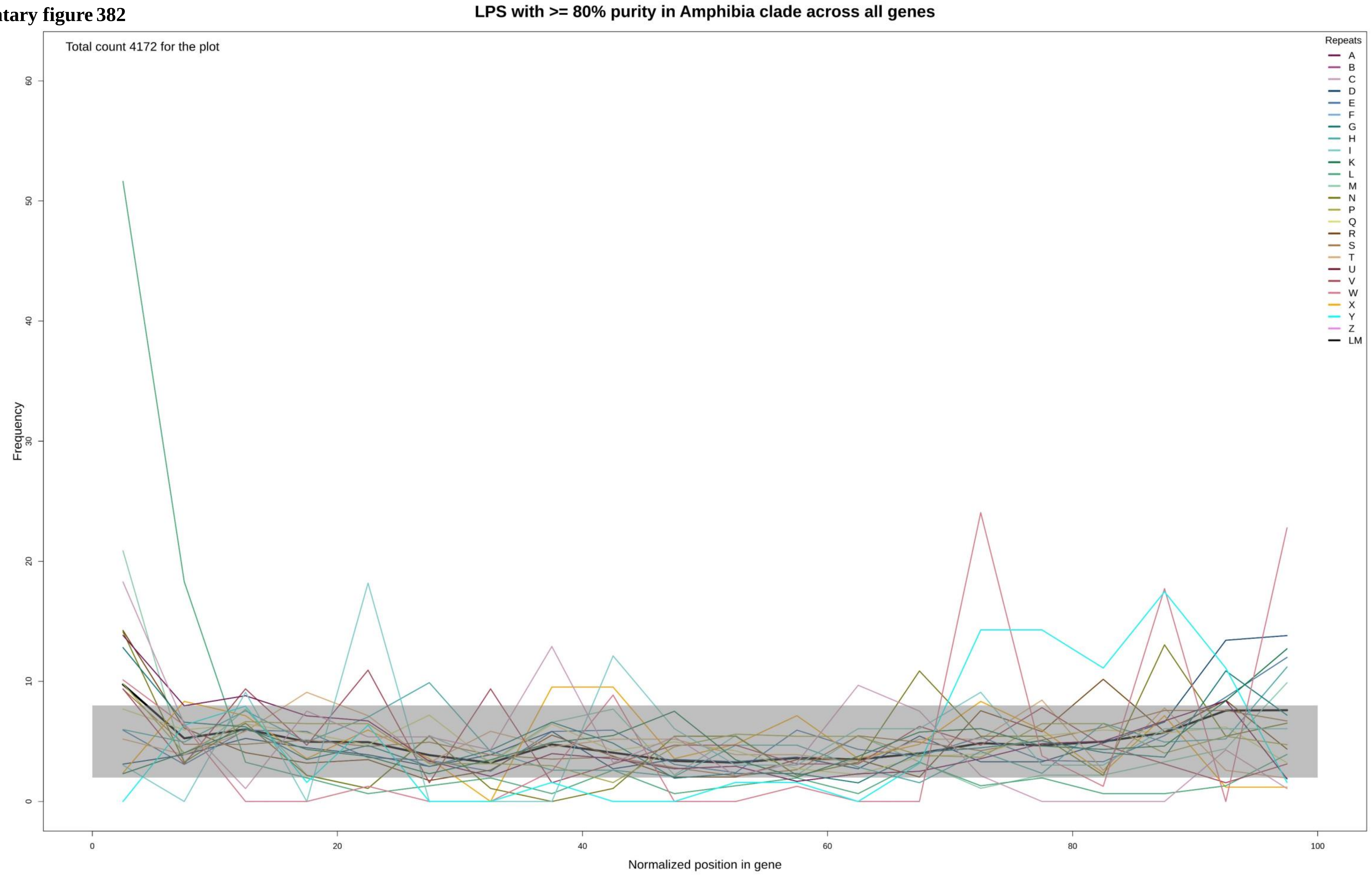

Supplementary figure 383

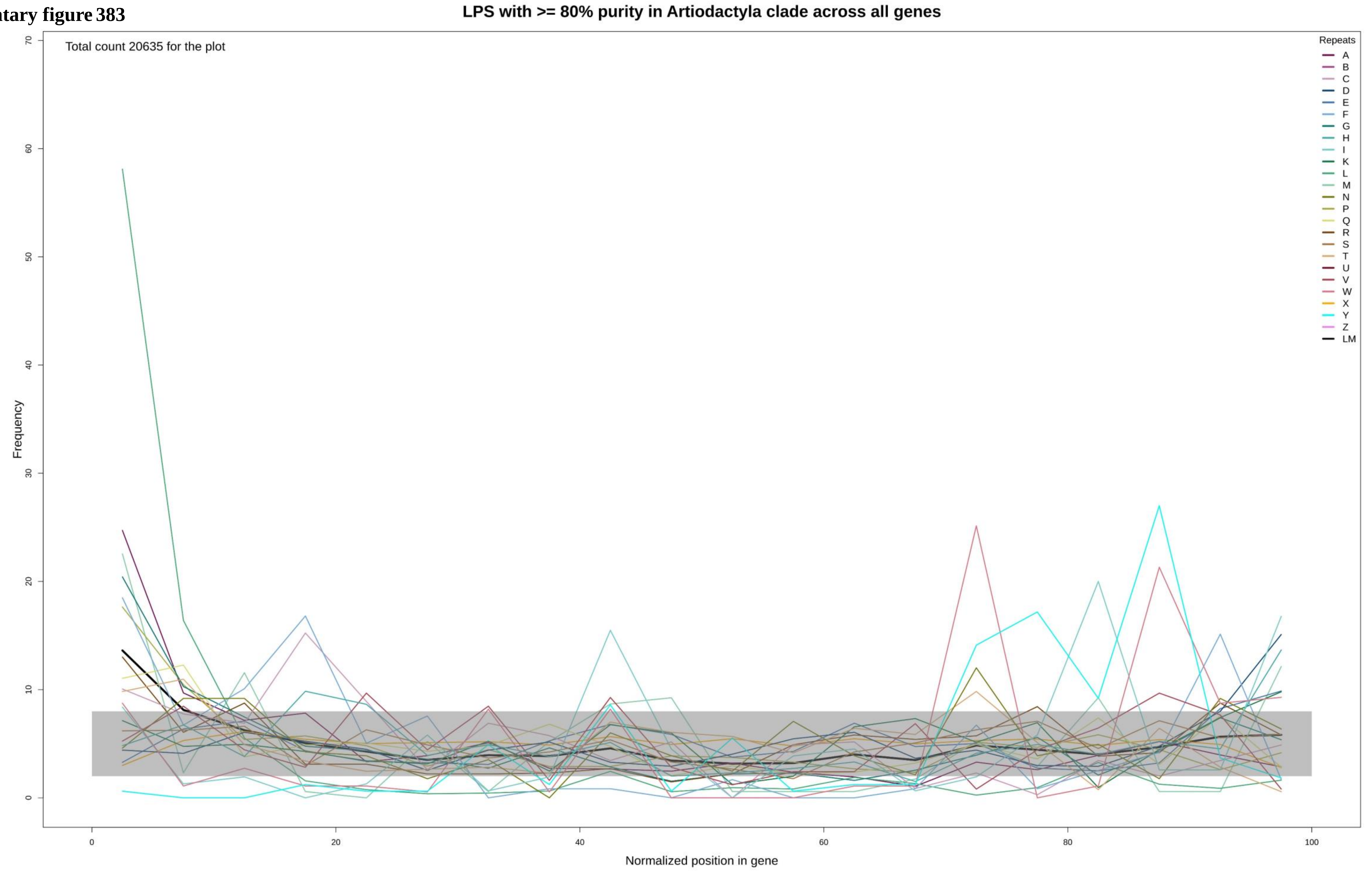

**Supplementary figure 384**

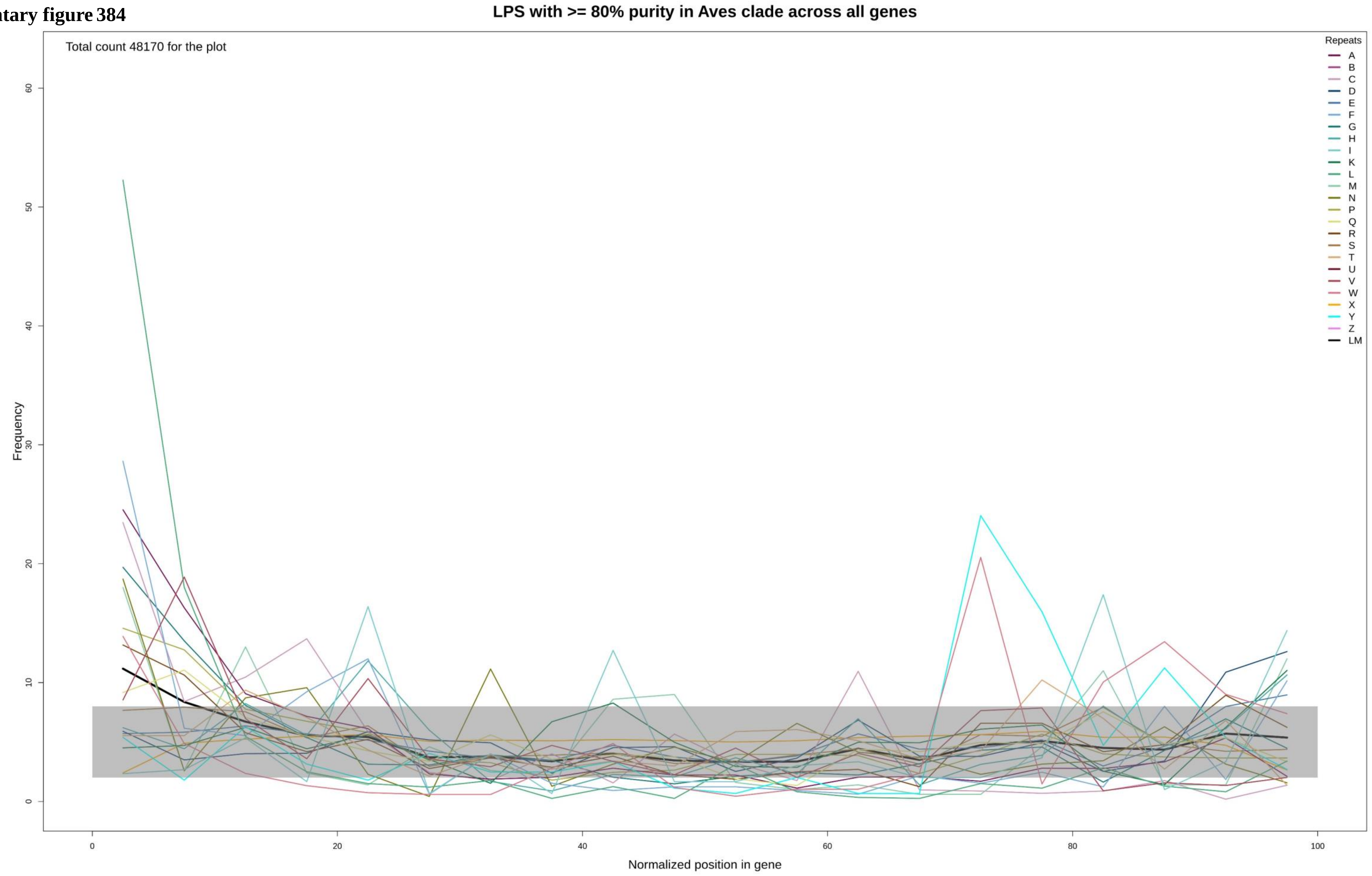

Supplementary figure 385

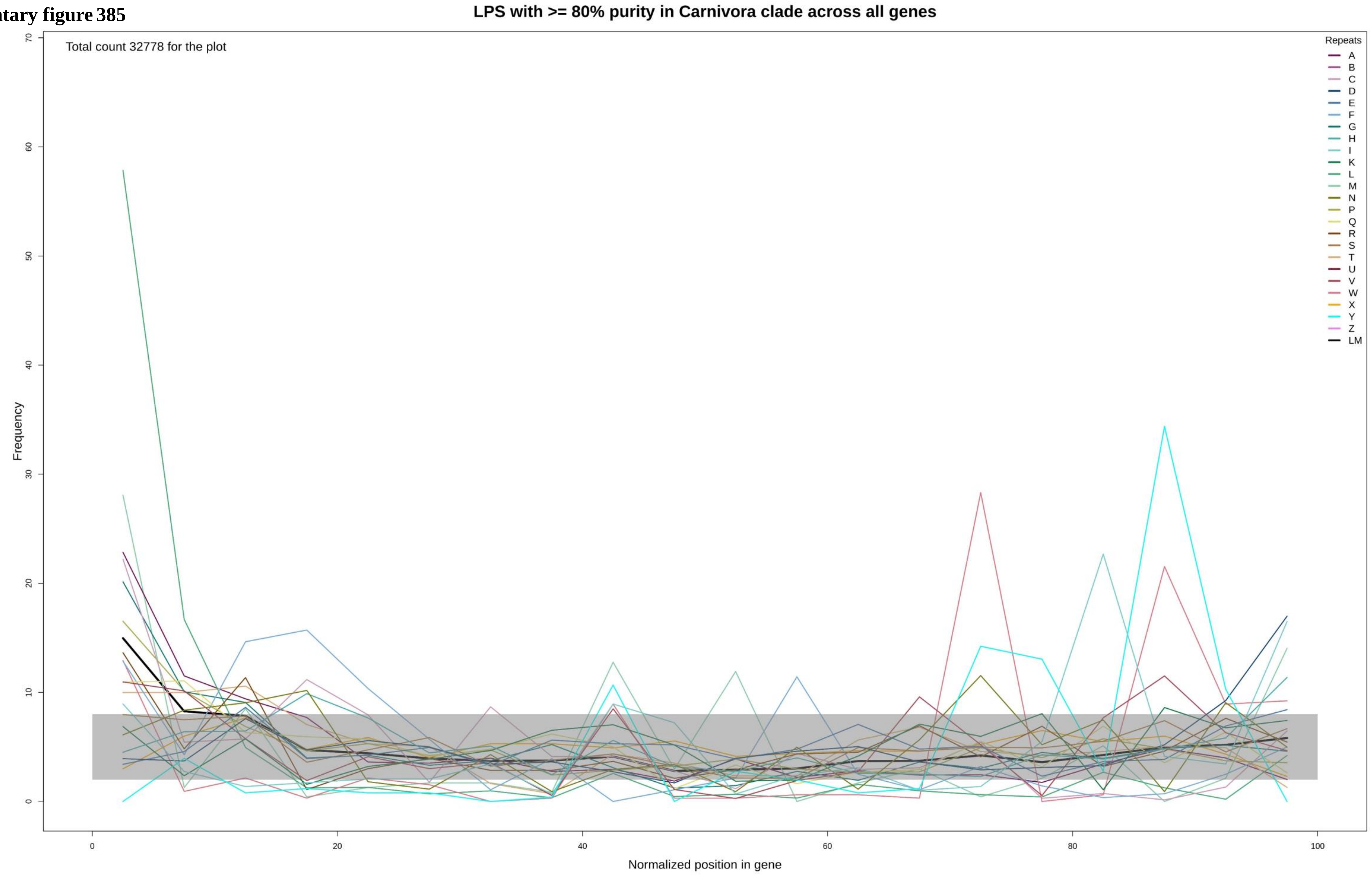

Supplementary figure 386

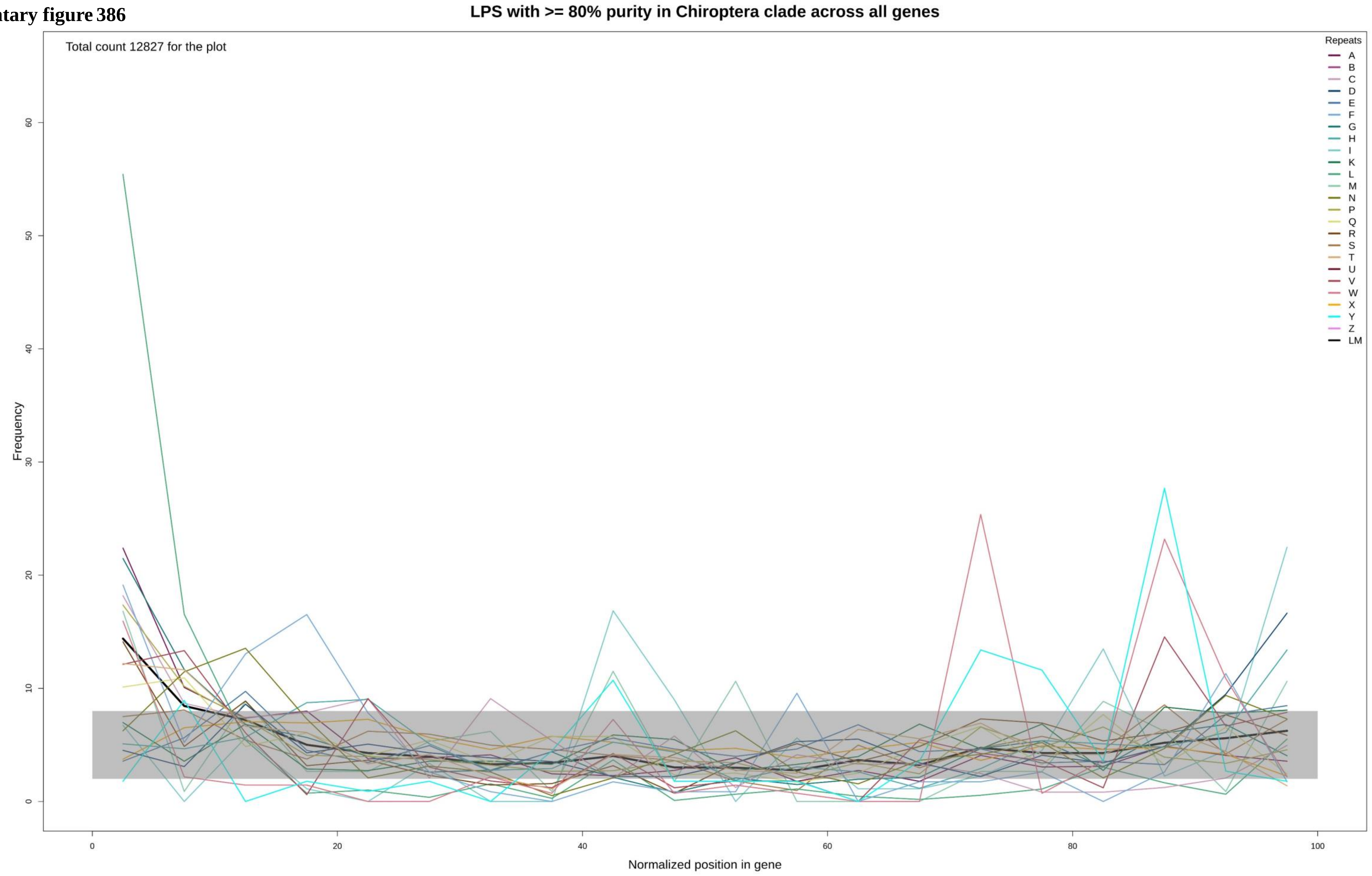

Supplementary figure 387

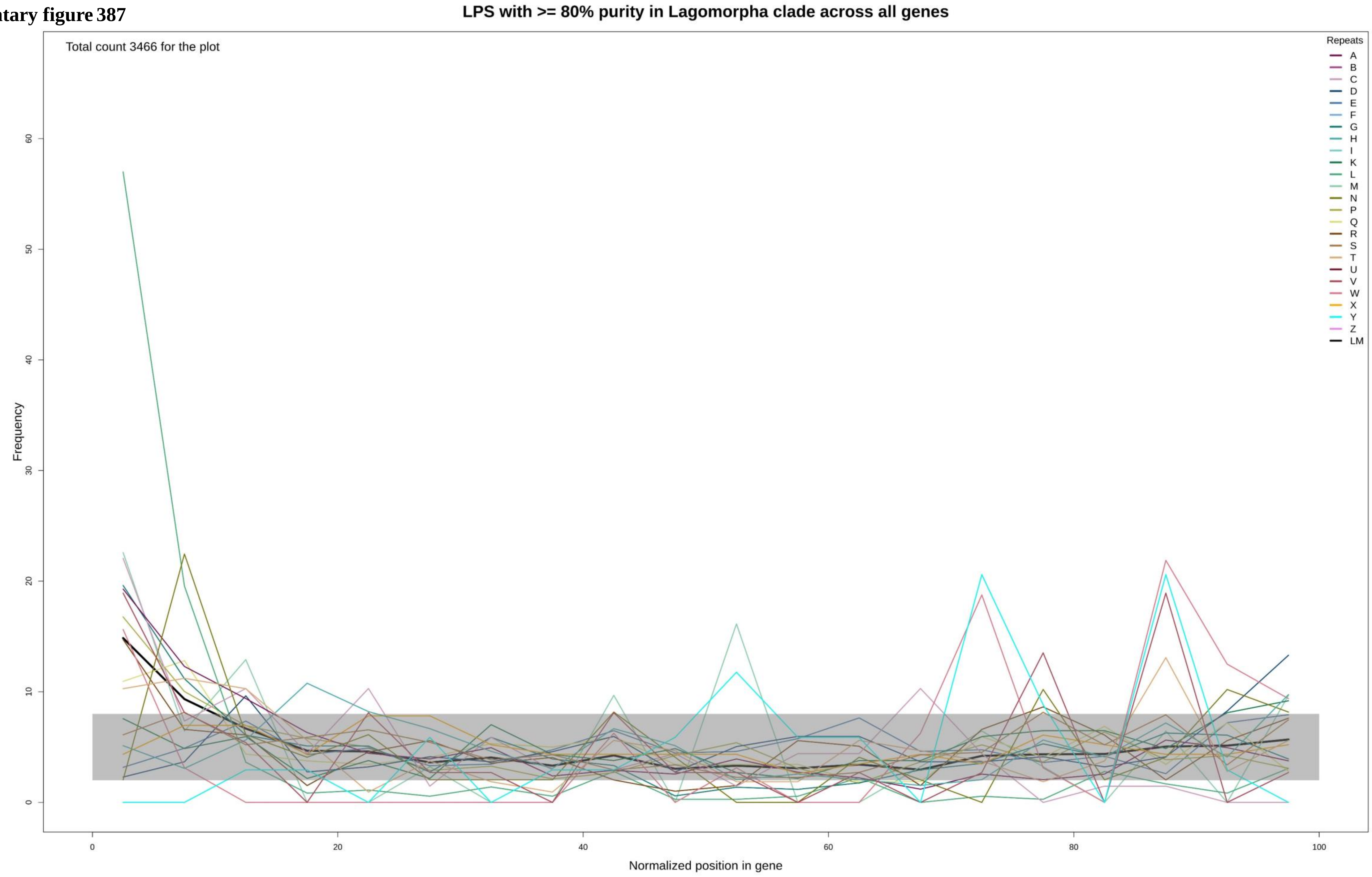

**Supplementary figure 388**

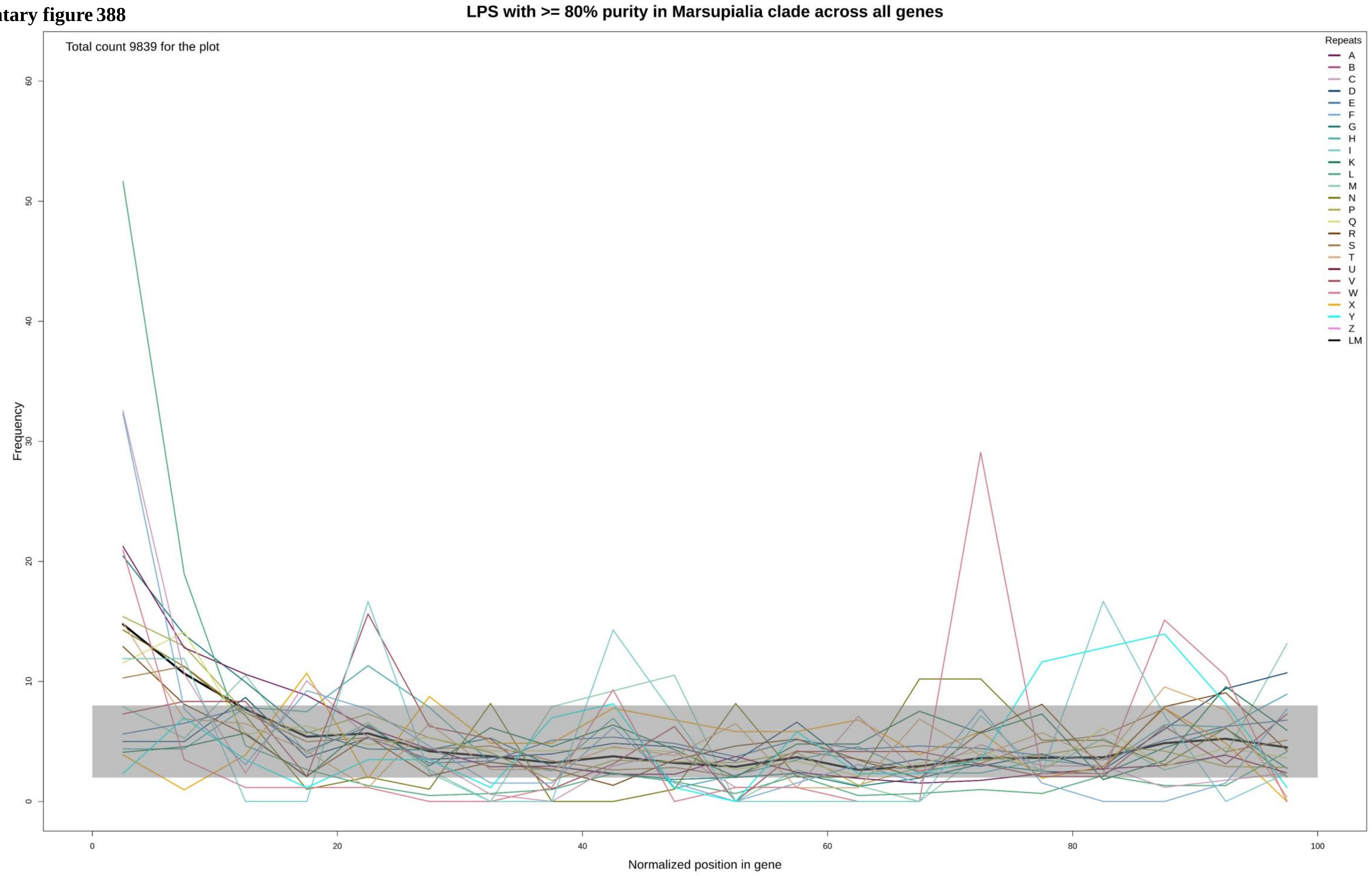

Supplementary figure 389

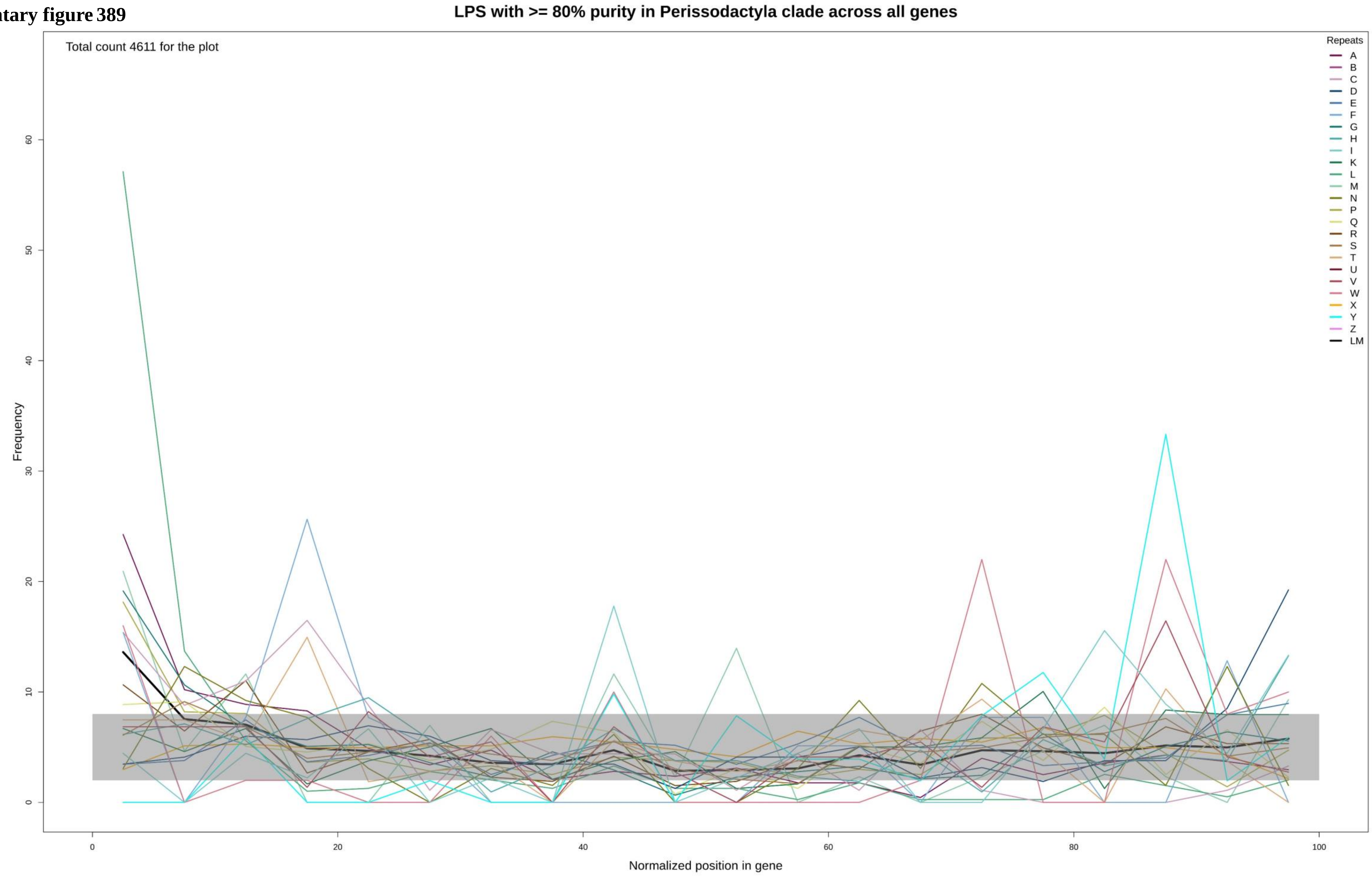

Supplementary figure 390

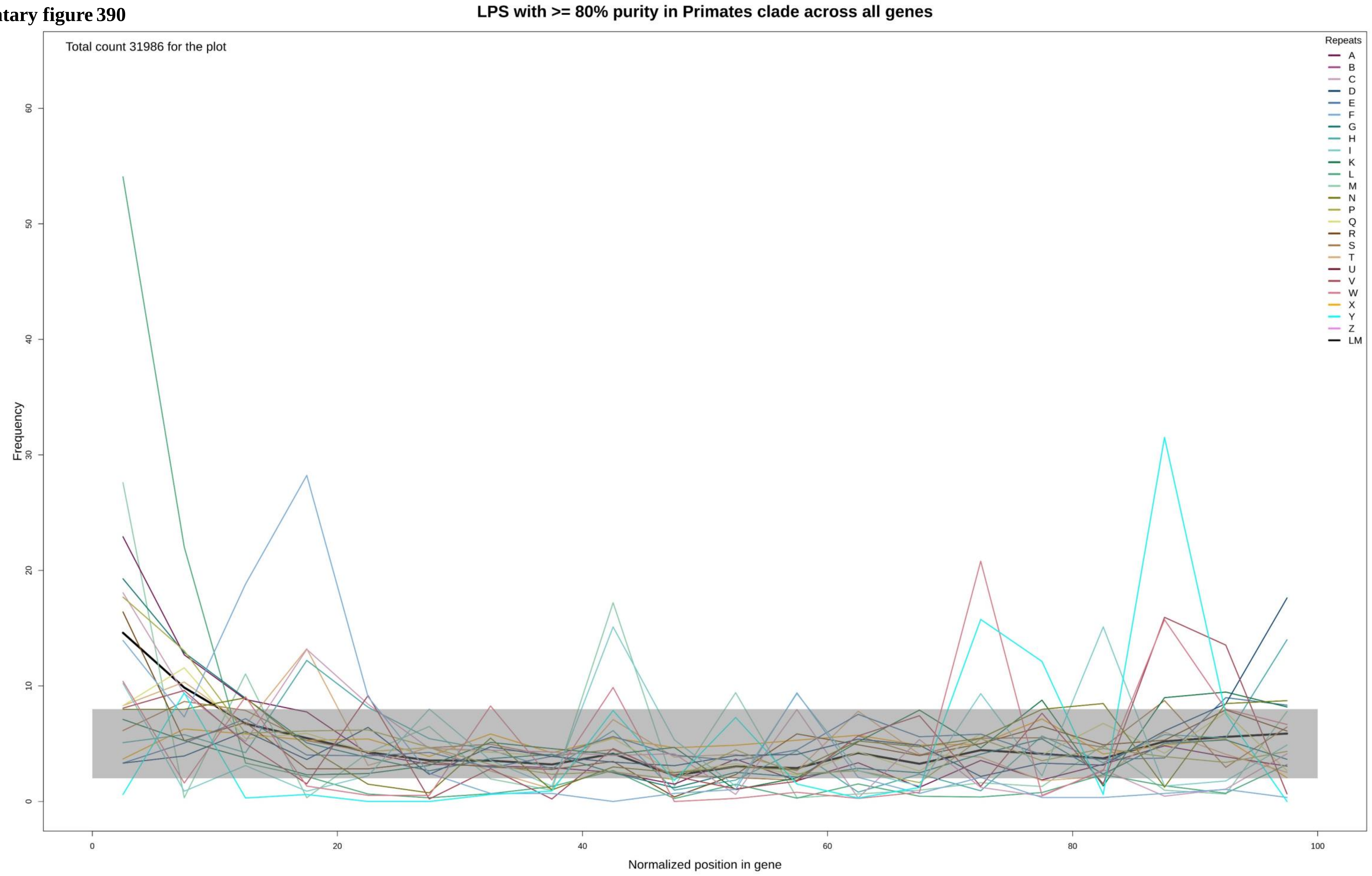

**Supplementary figure 391**

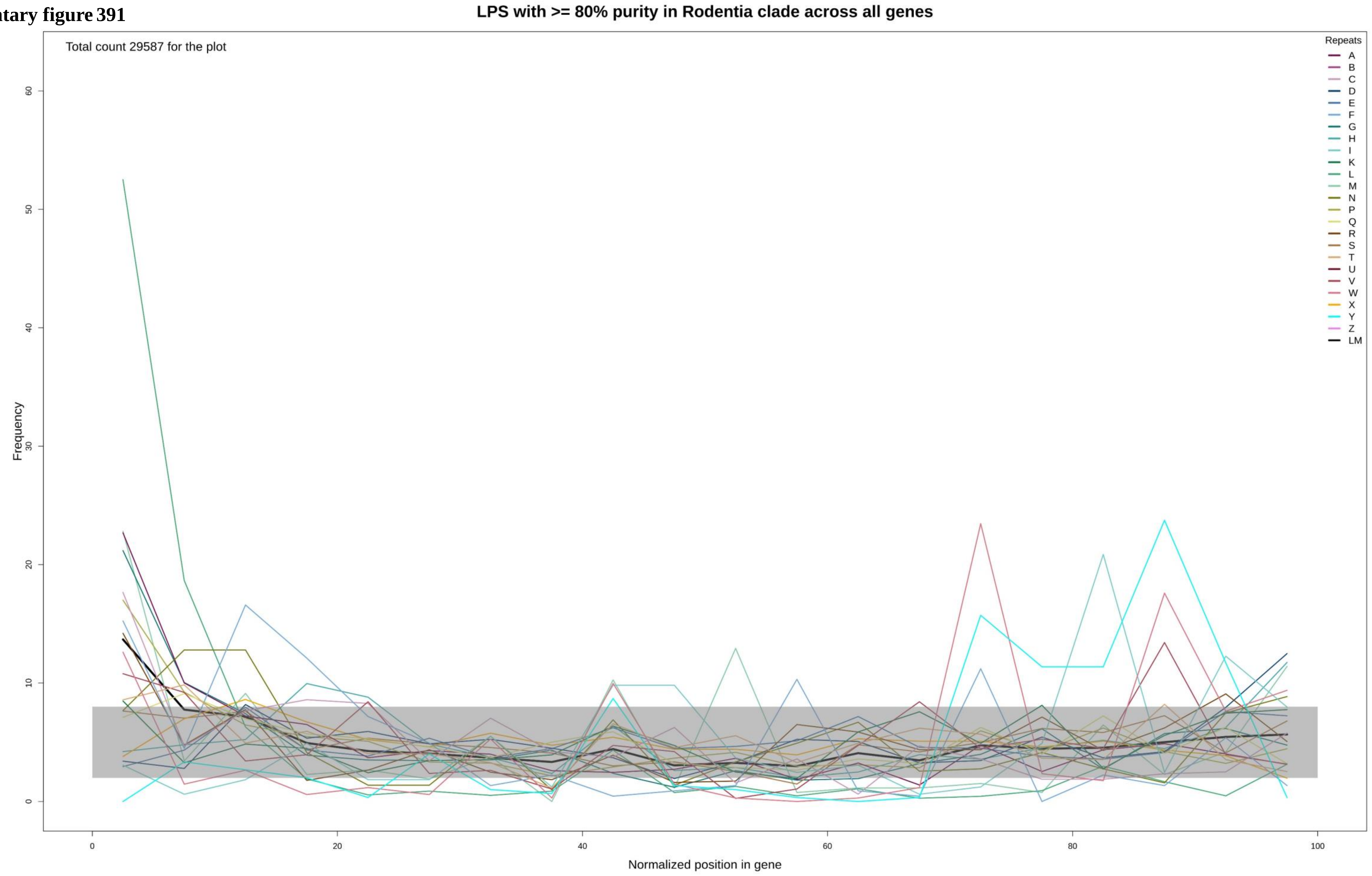

Supplementary figure 392

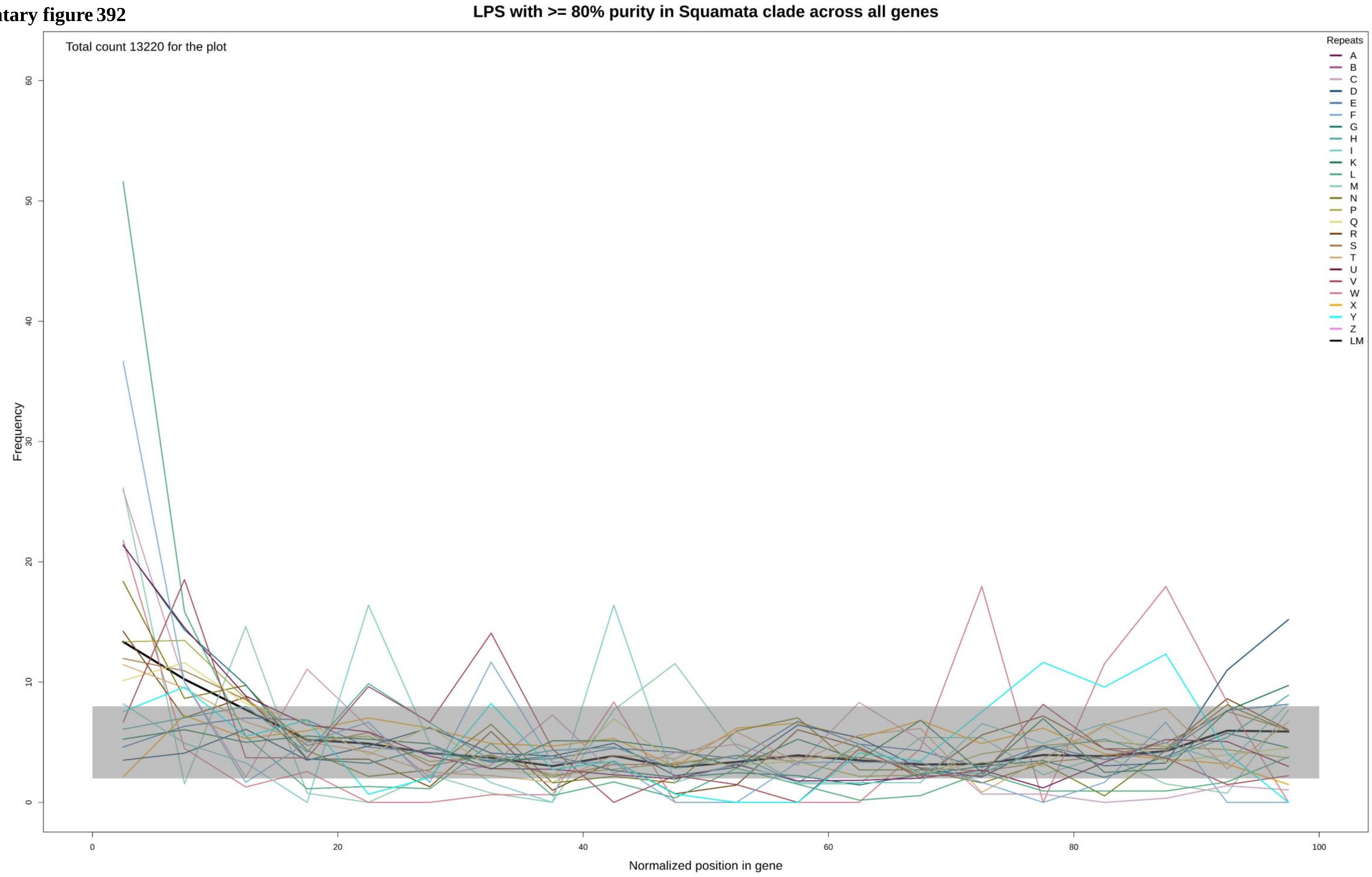

**Supplementary figure 393**

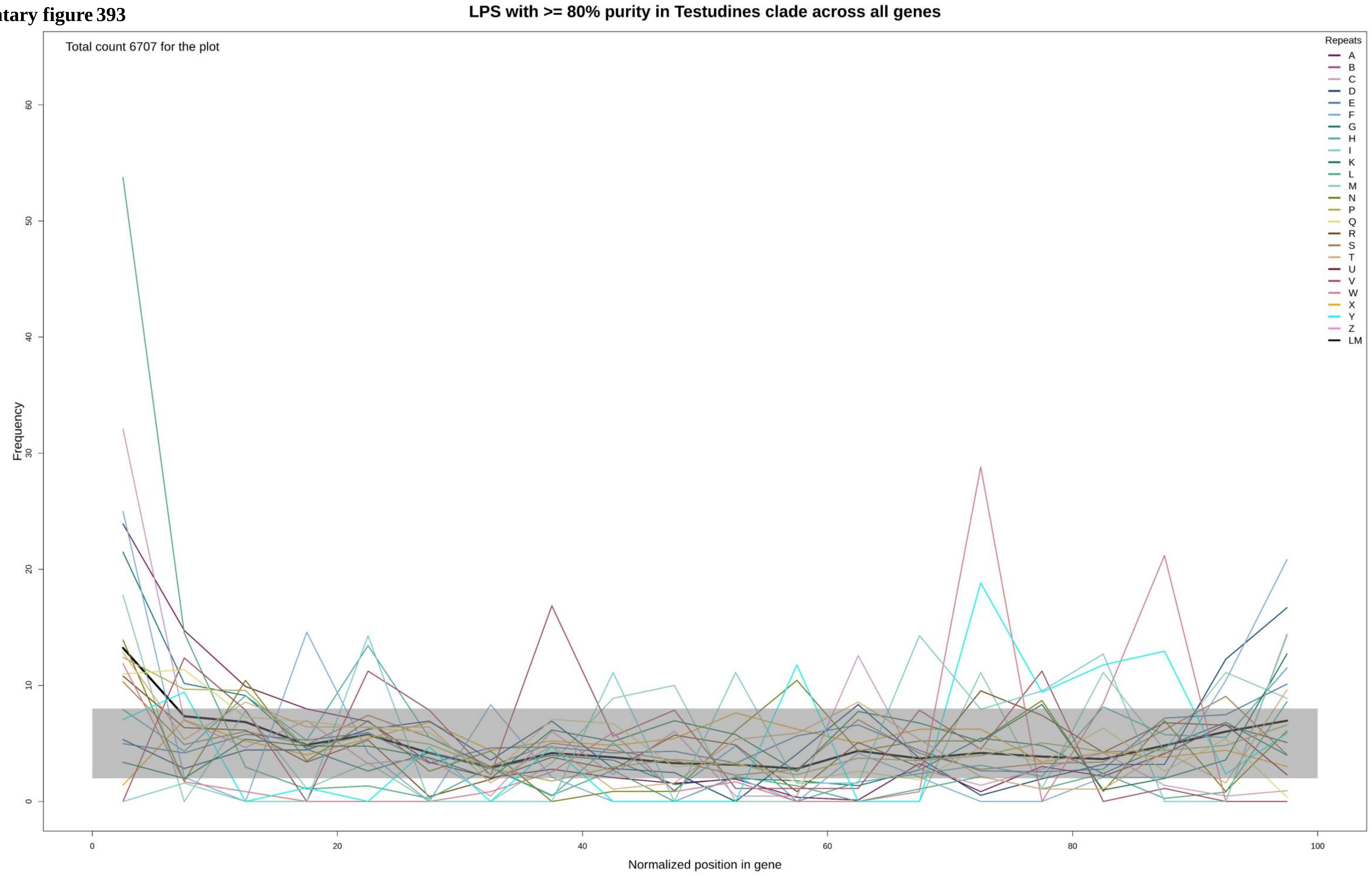

Supplementary figure 394

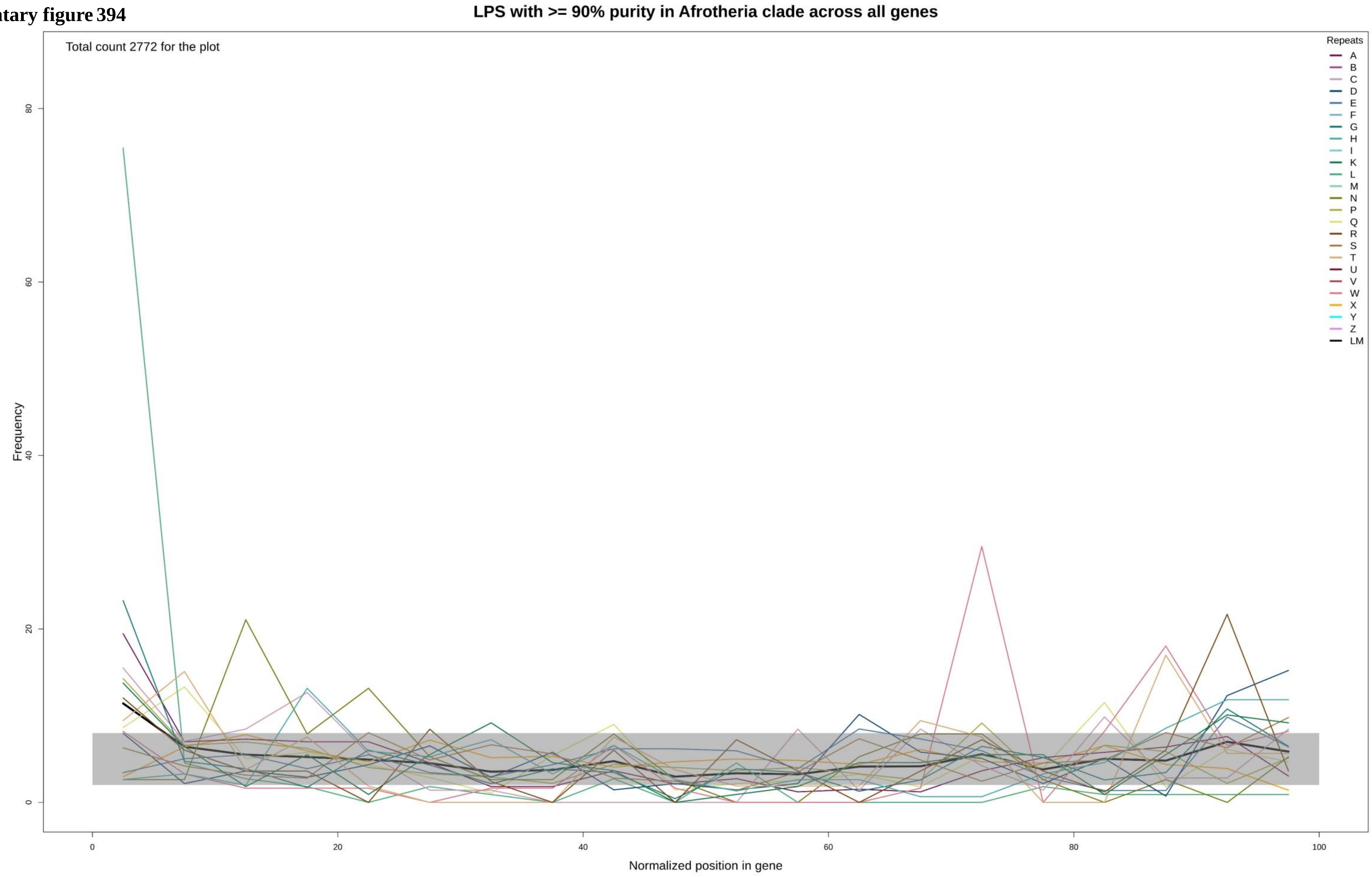

Supplementary figure 395

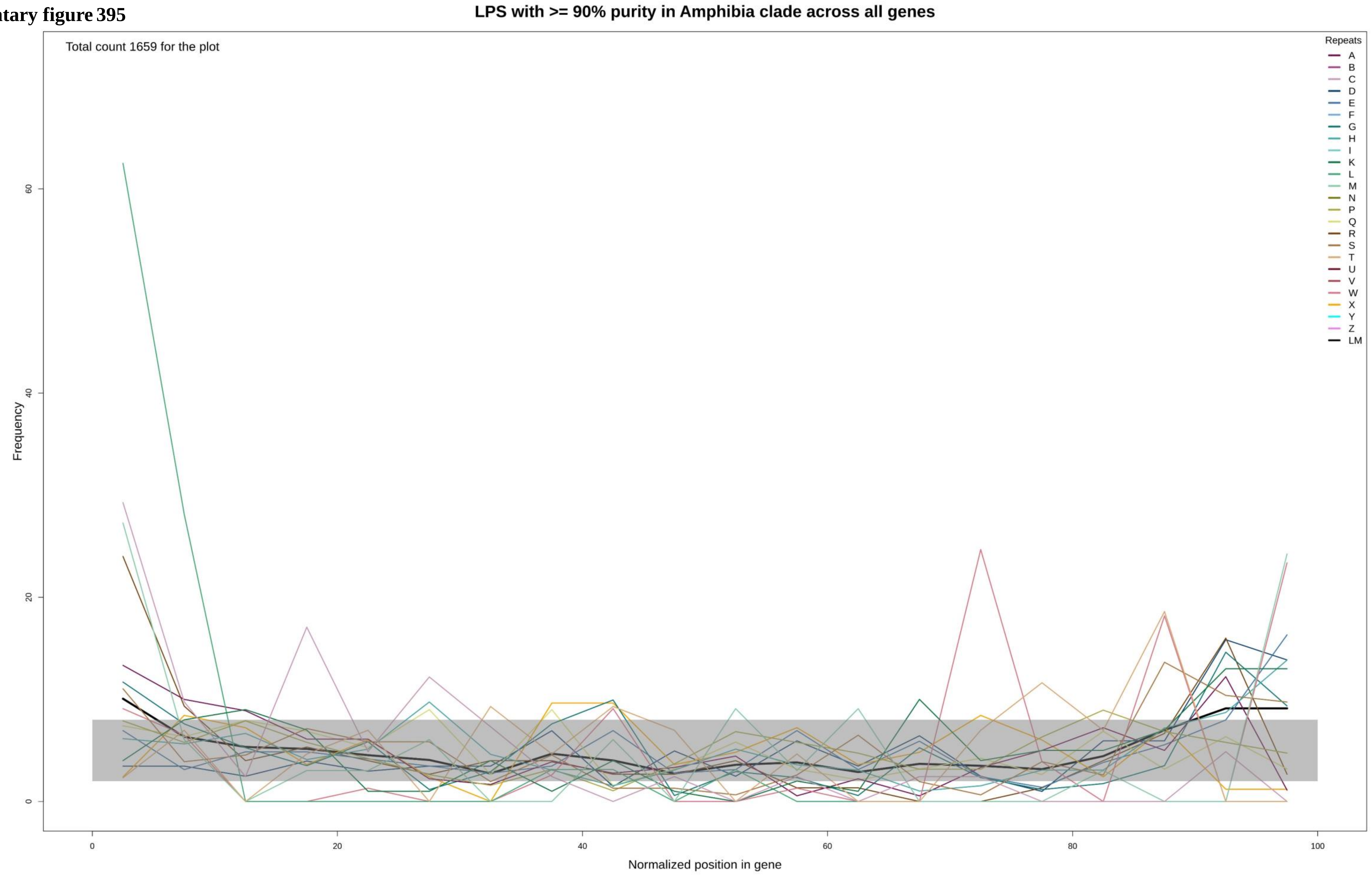

Supplementary figure 396

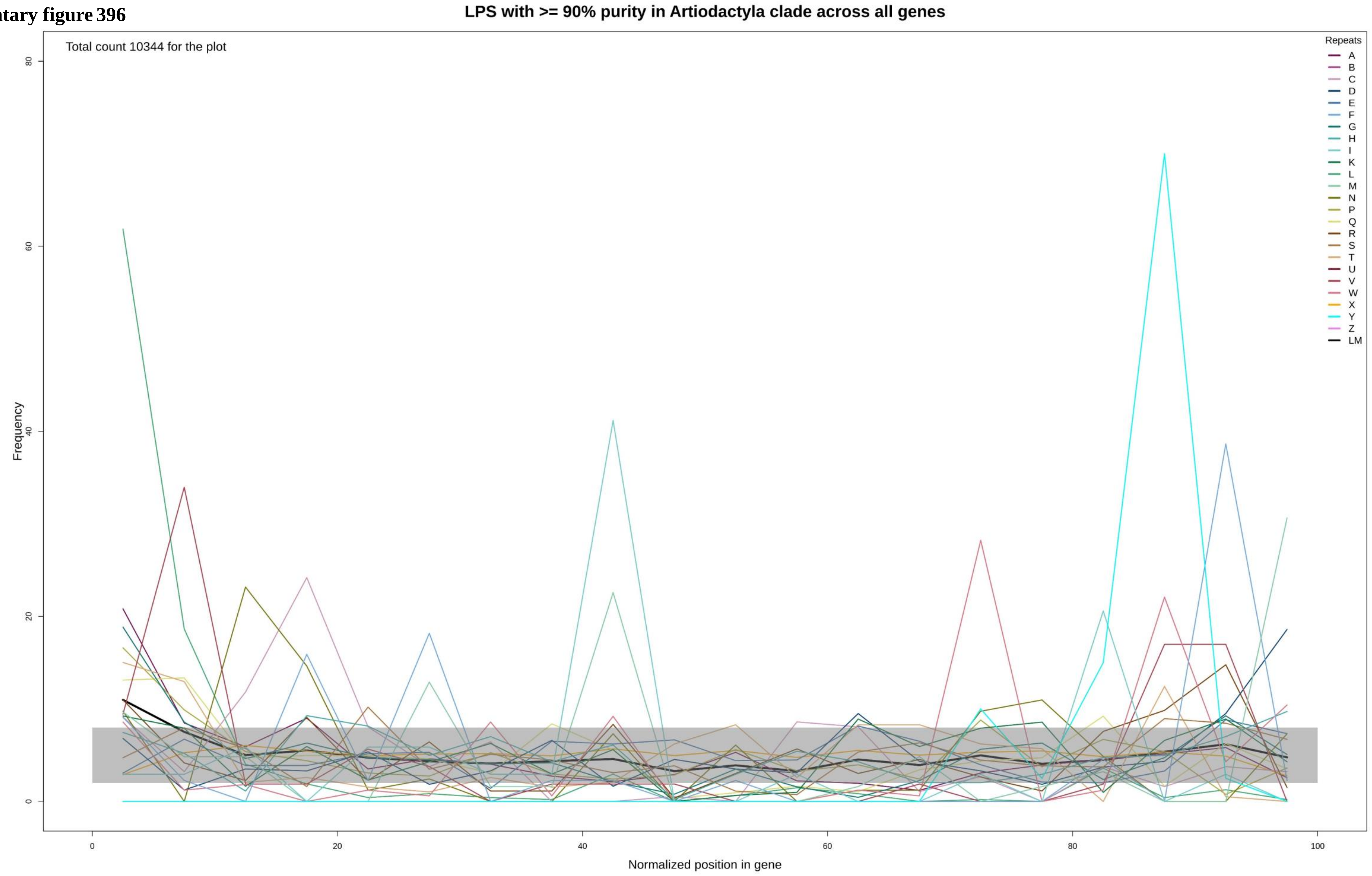

Supplementary figure 397

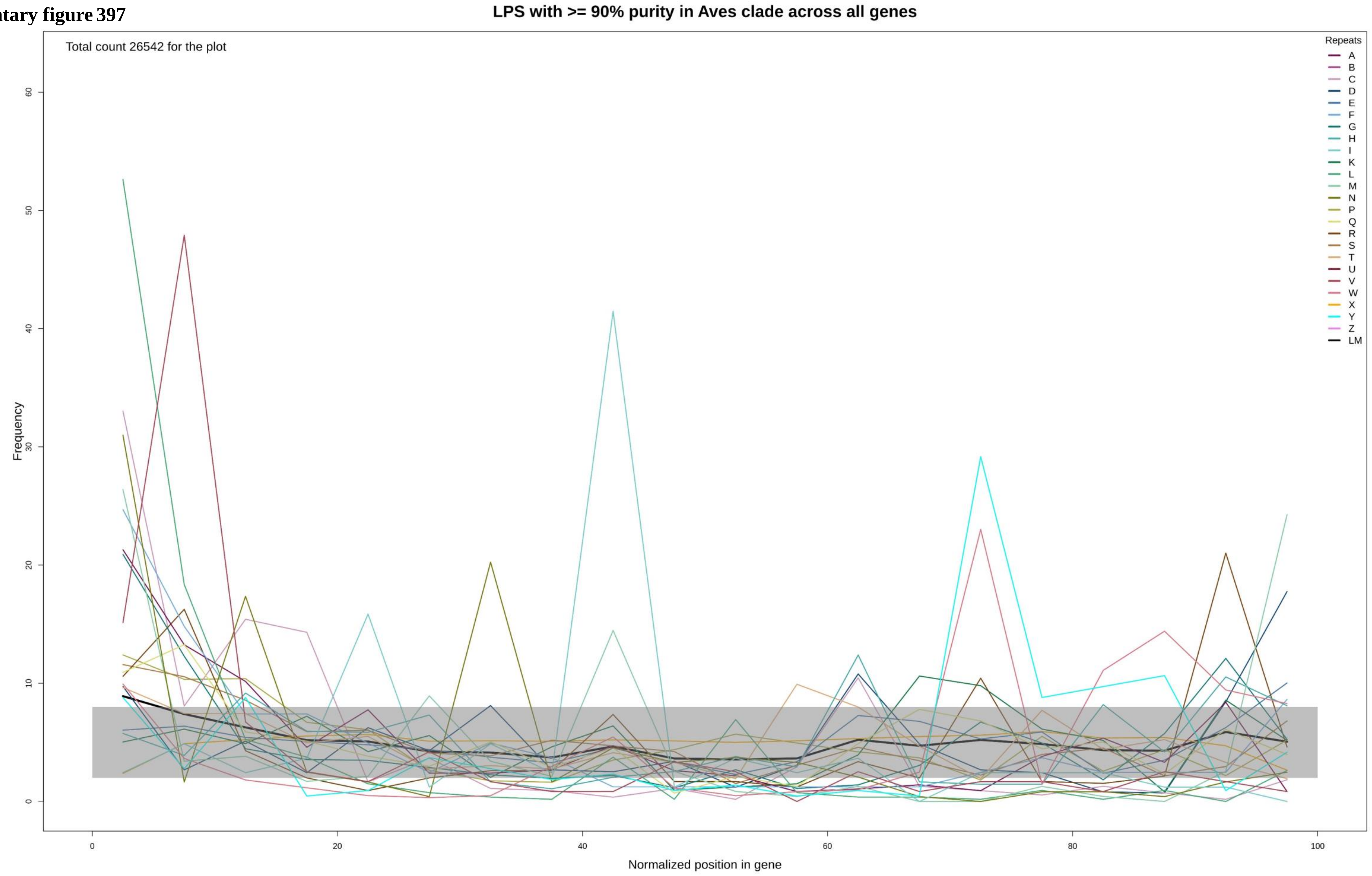

**Supplementary figure 398**

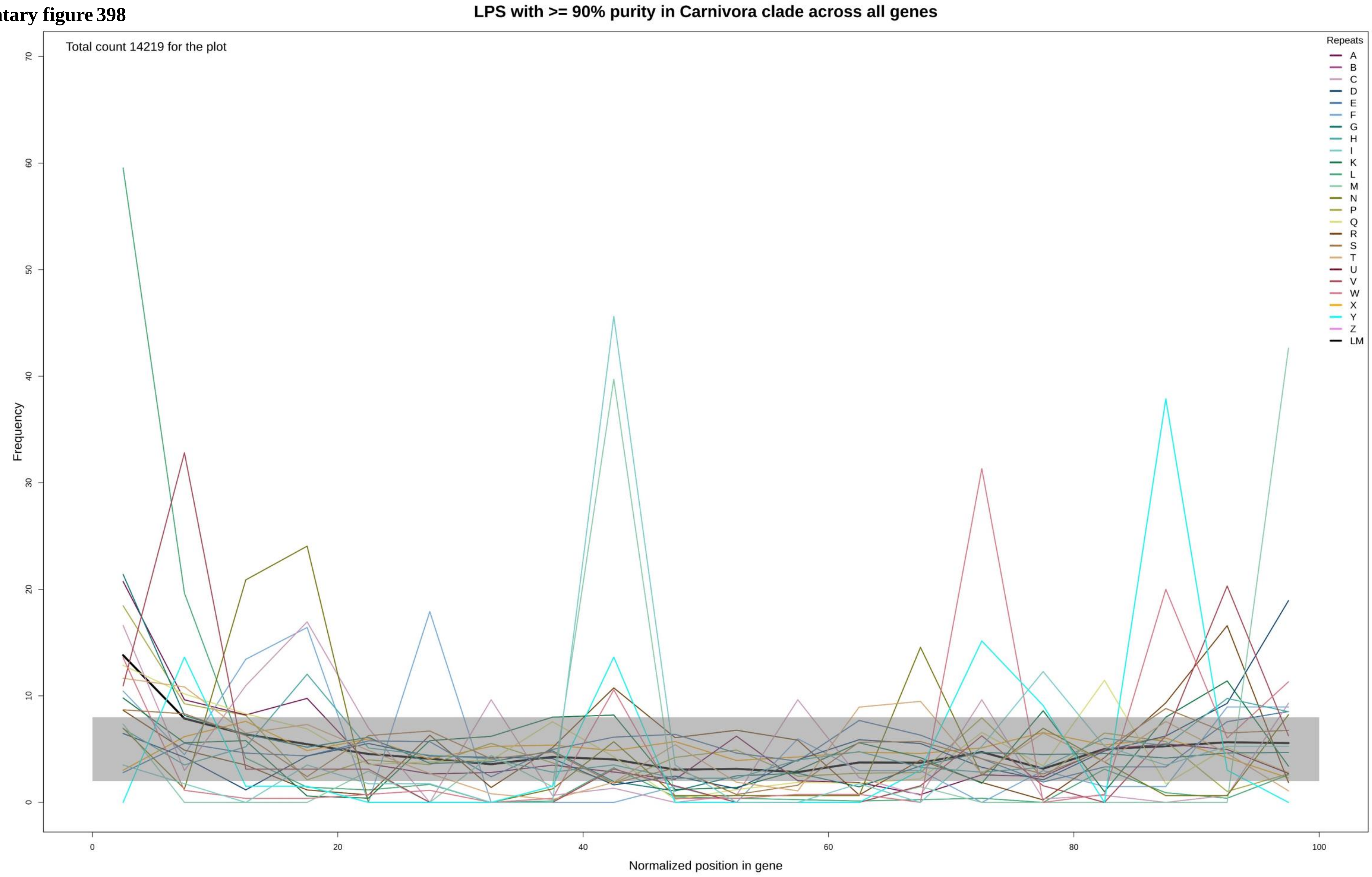

**Supplementary figure 399**

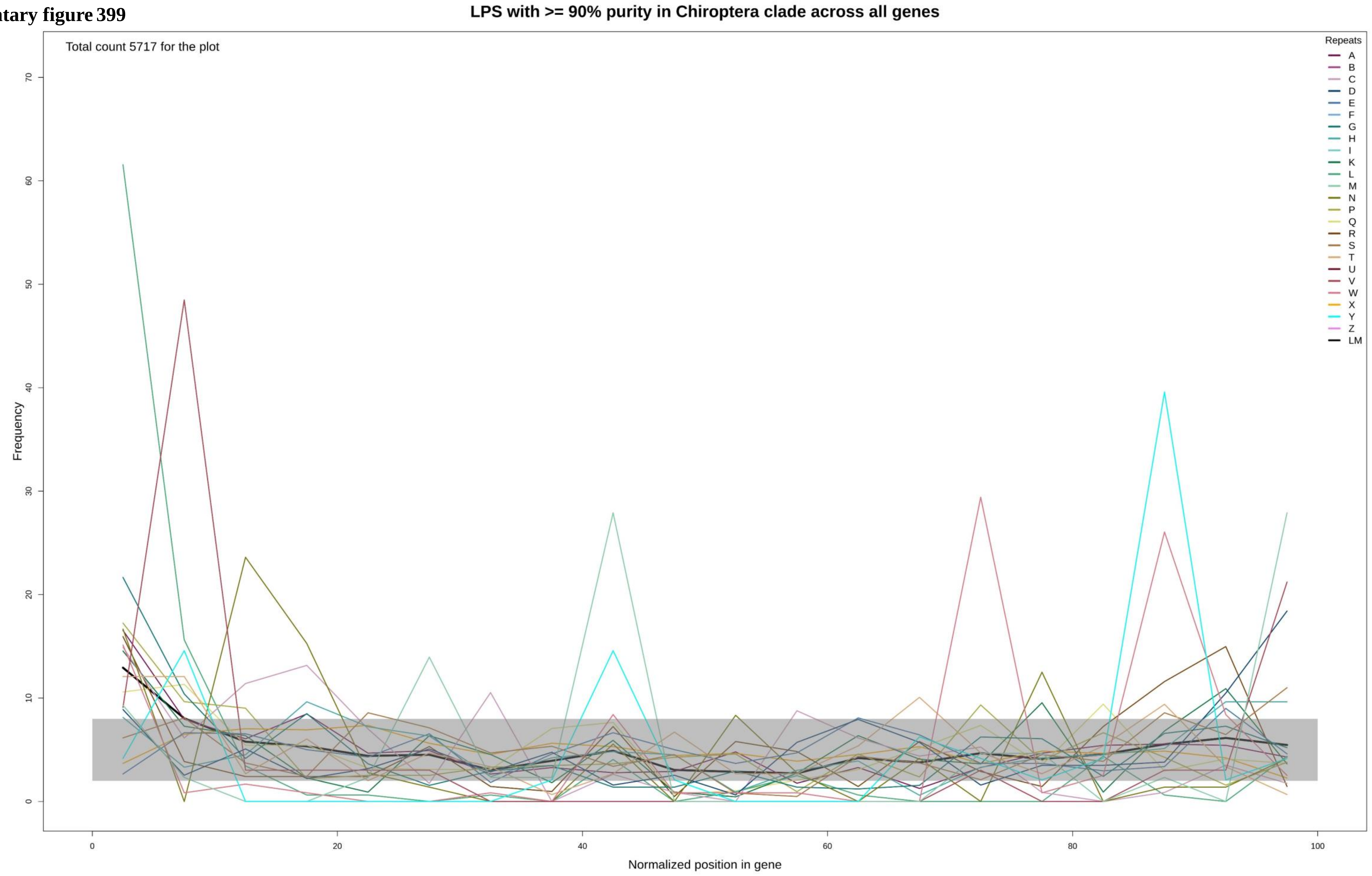

Supplementary figure 400

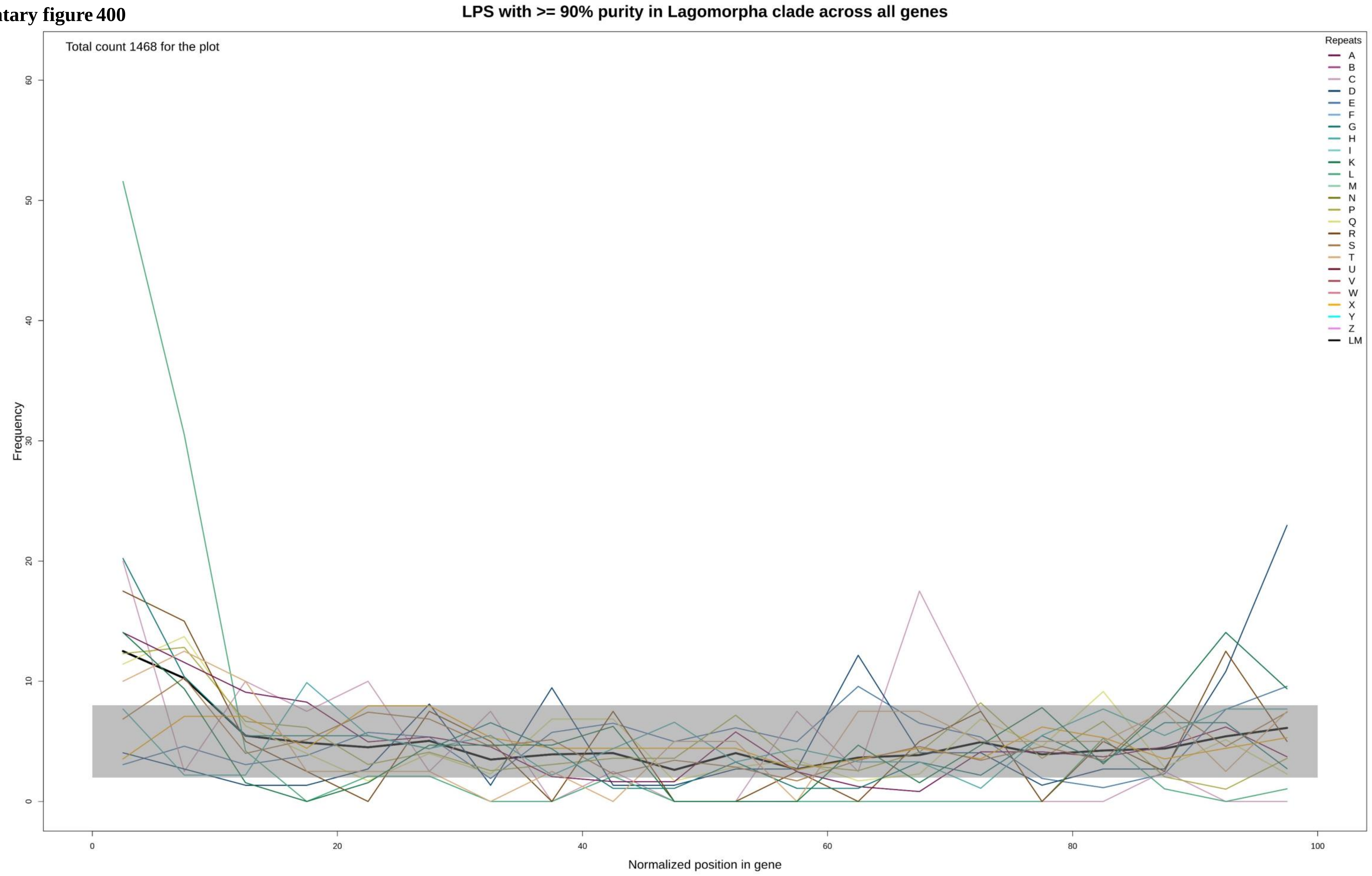

**Supplementary figure 401**

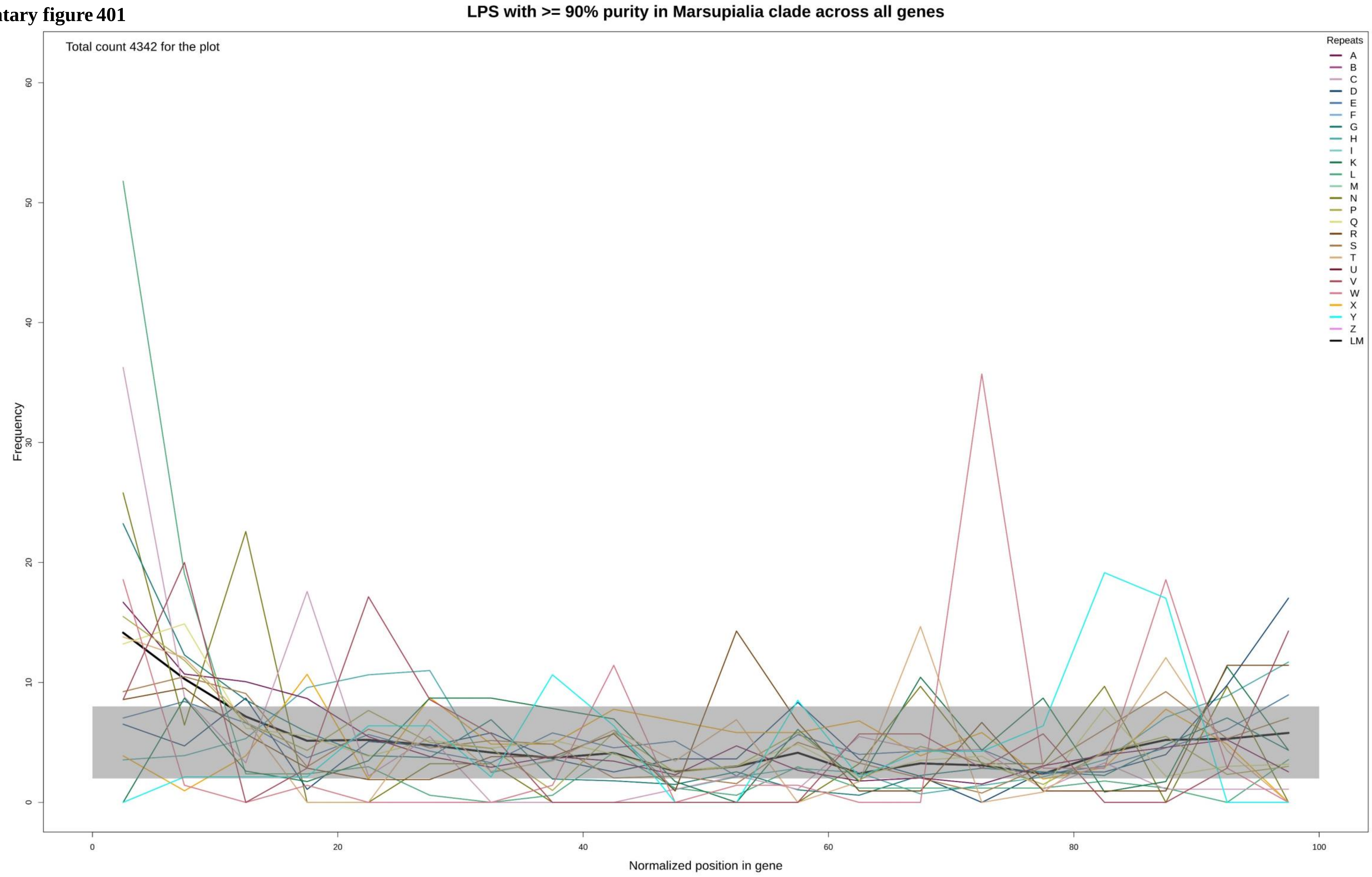

Supplementary figure 402

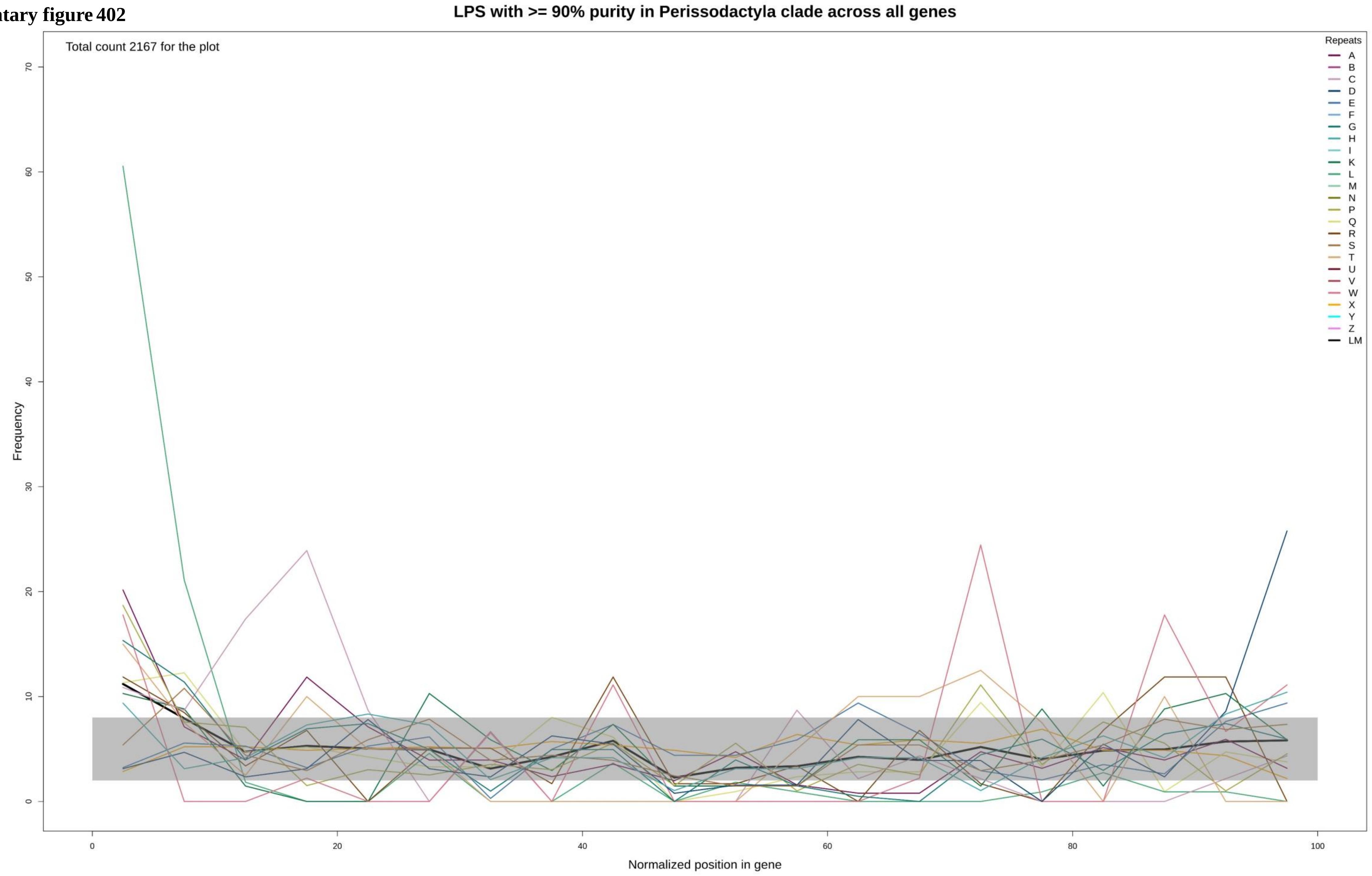

**Supplementary figure 403**

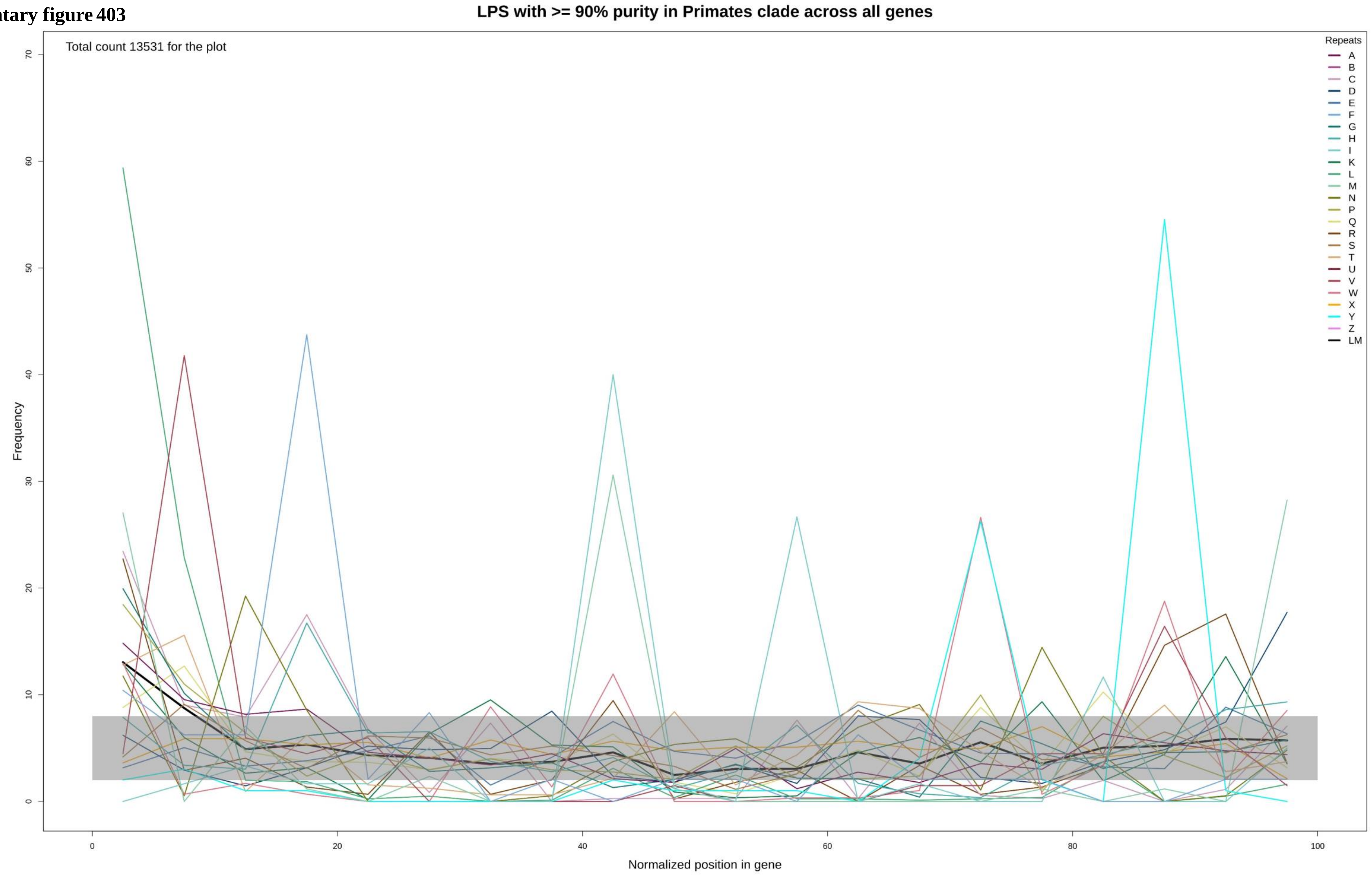

**Supplementary figure 404**

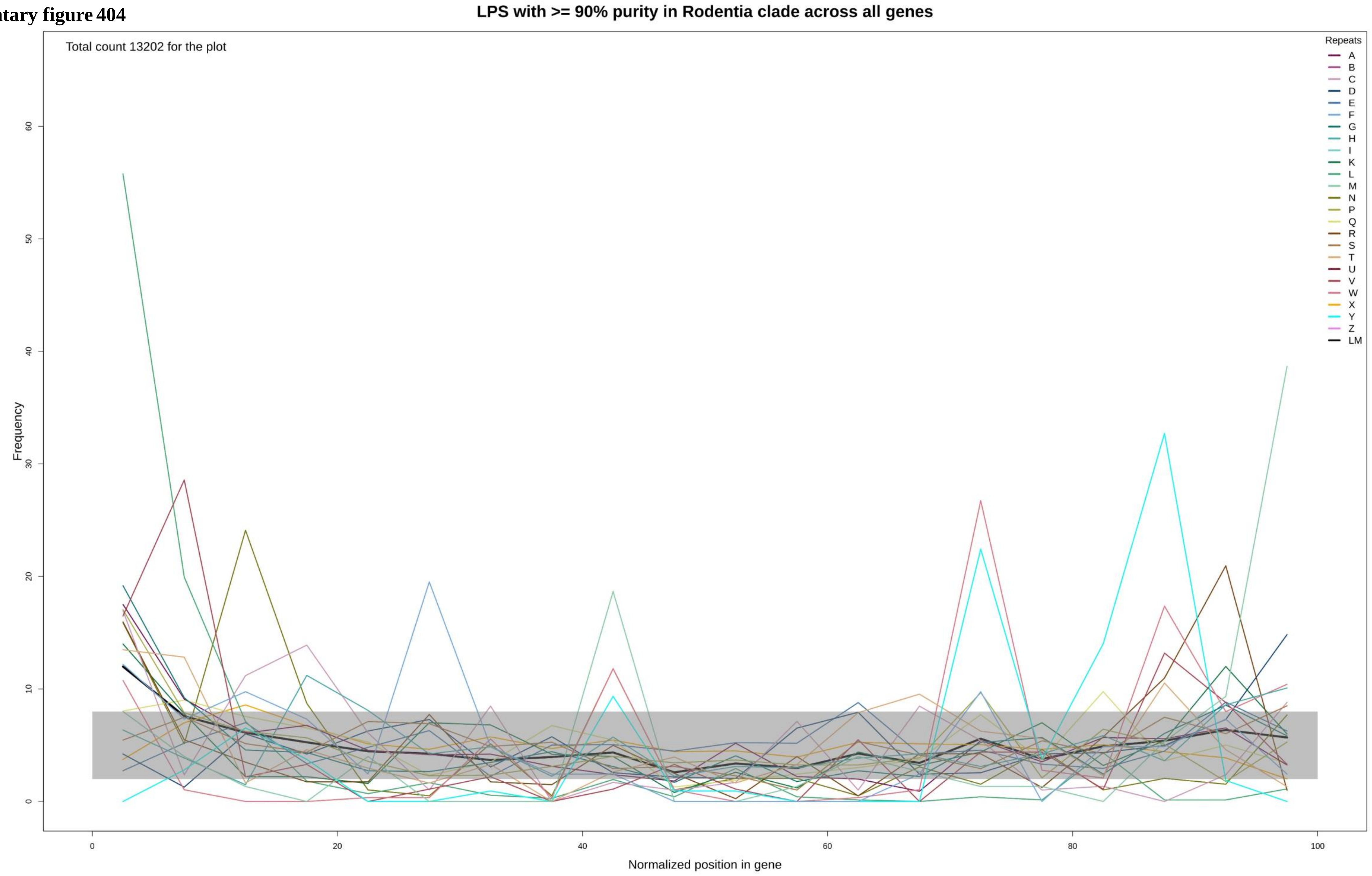

Supplementary figure 405

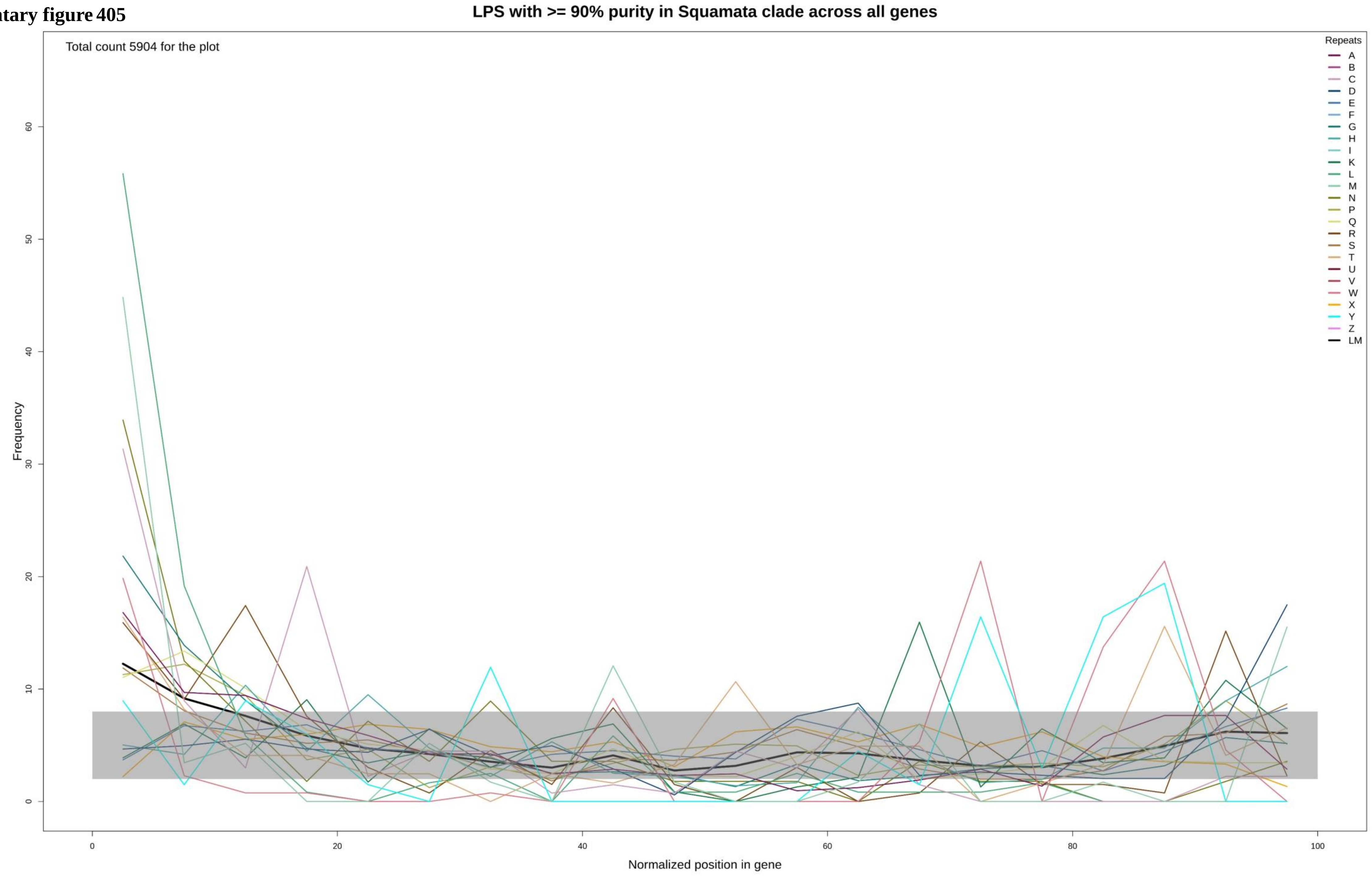

**Supplementary figure 406**

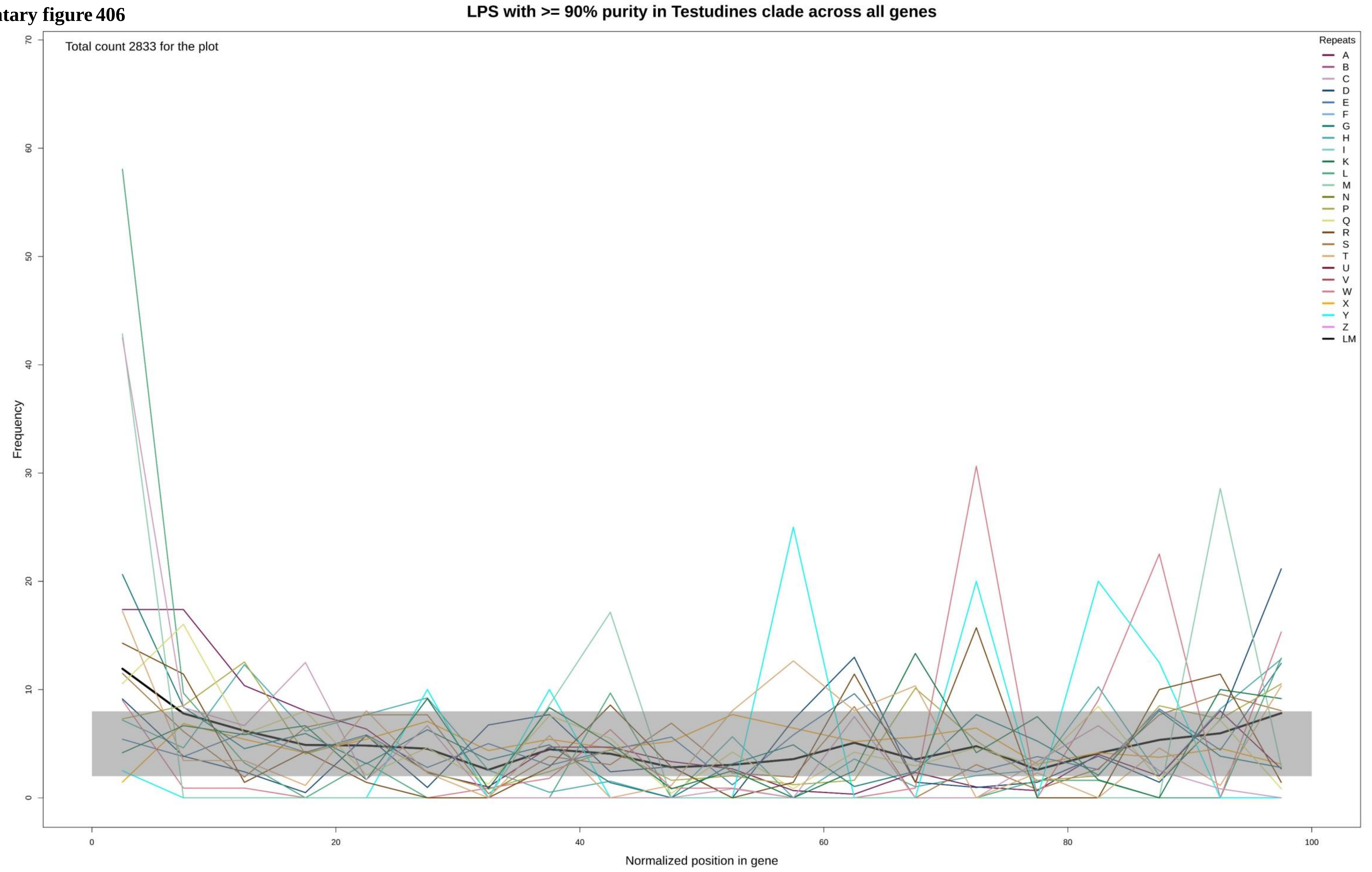

**Supplementary figure 407**

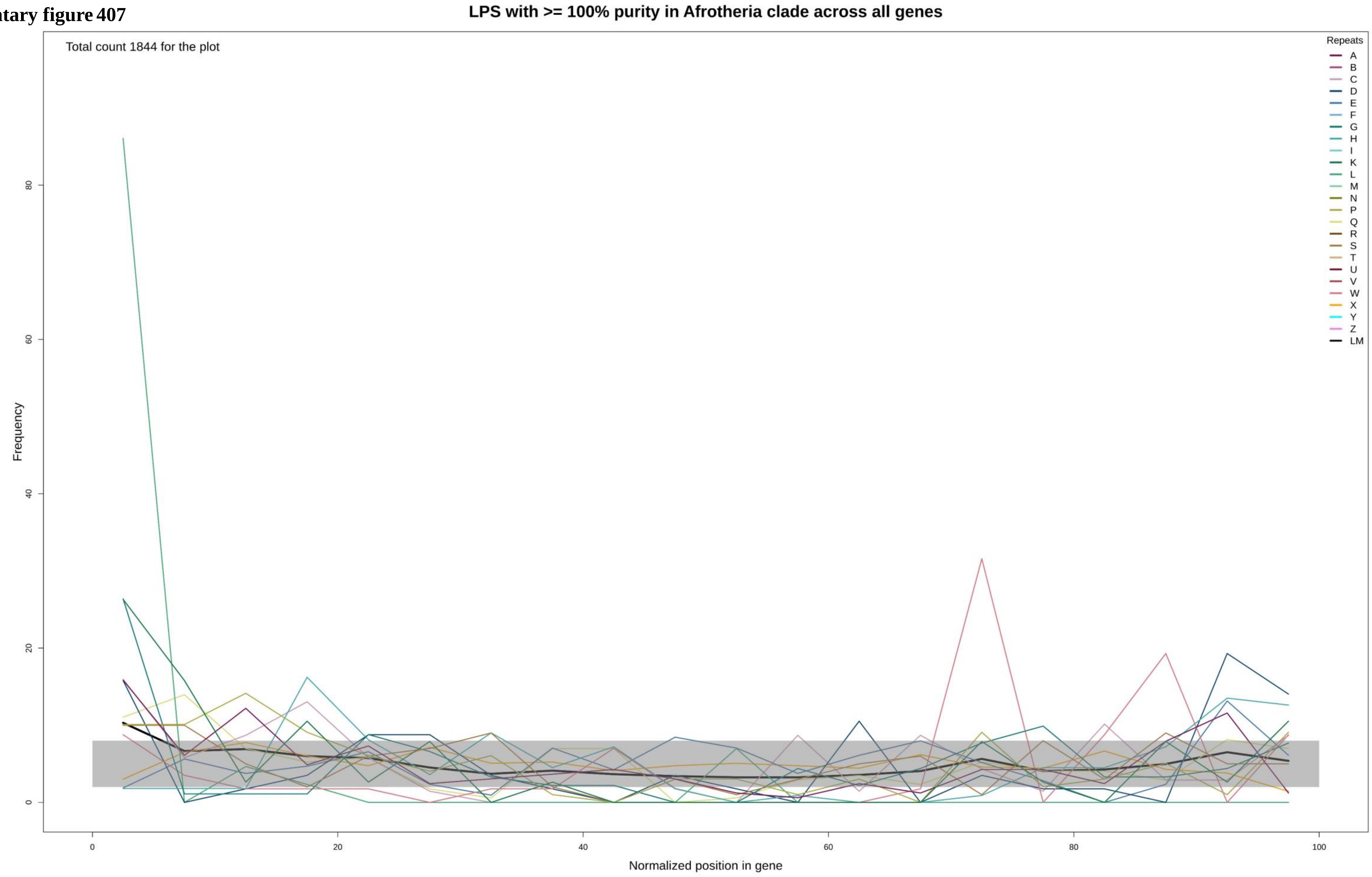

Supplementary figure 408

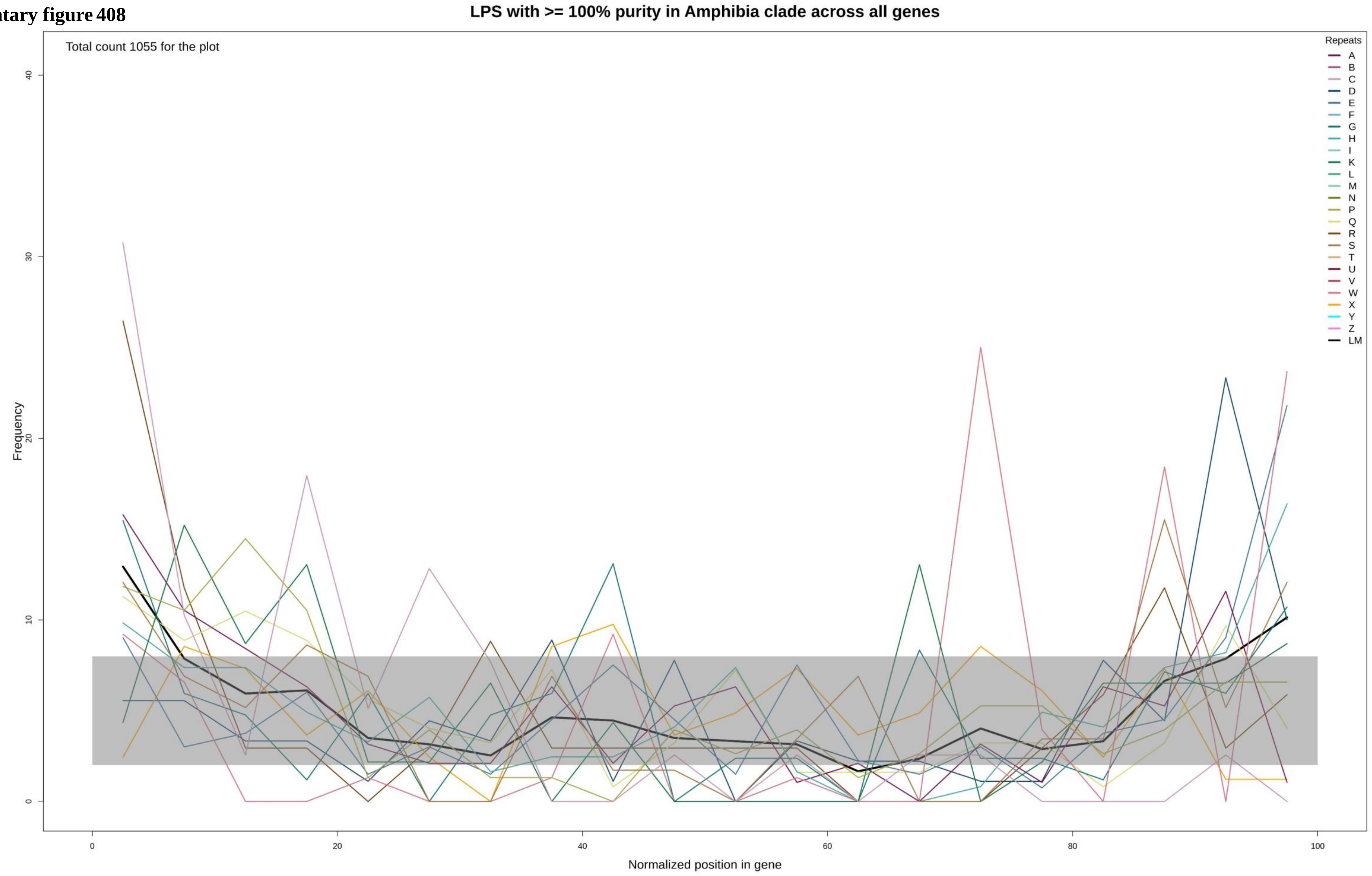

**Supplementary figure 409**

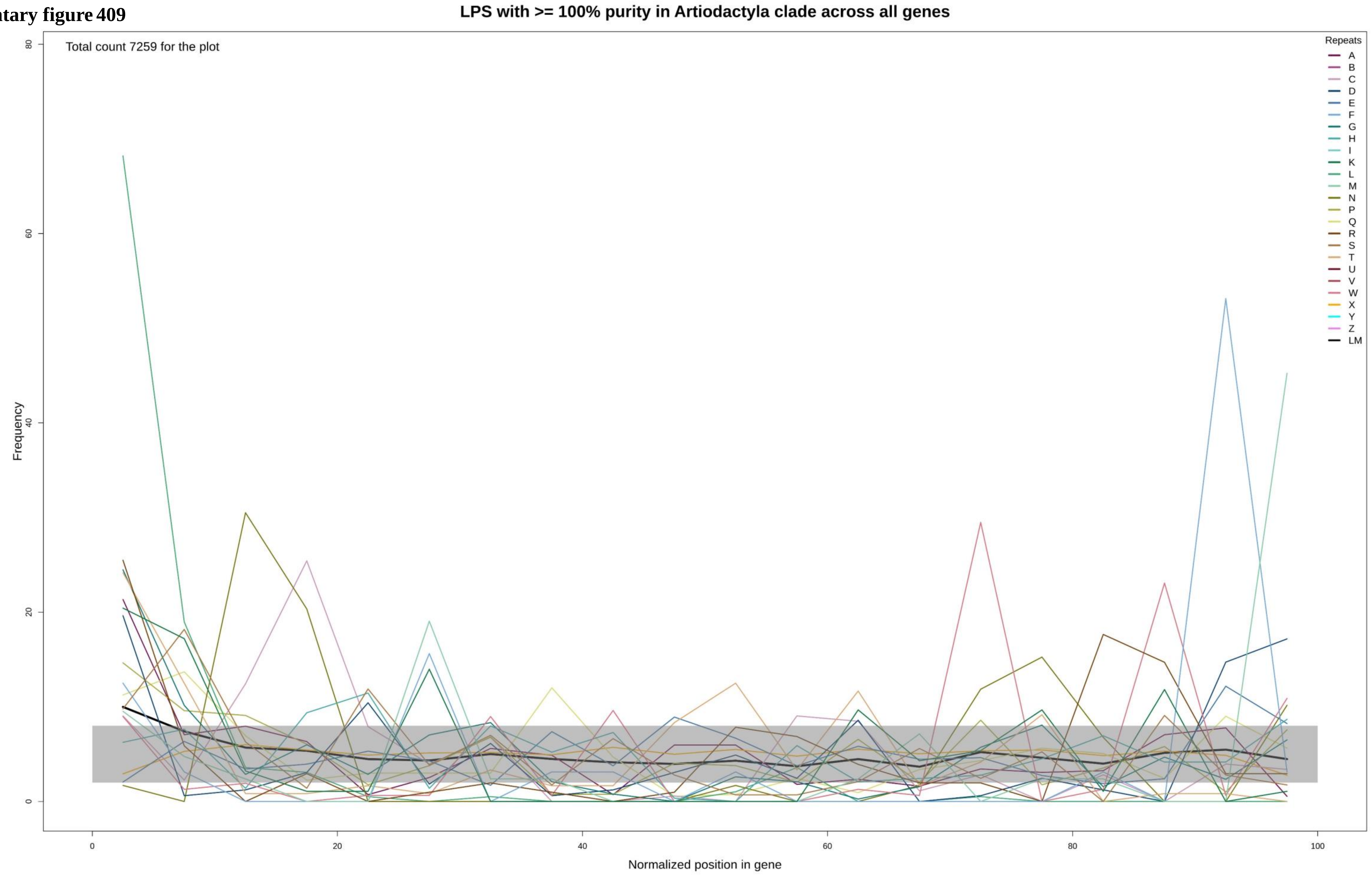

**Supplementary figure 410**

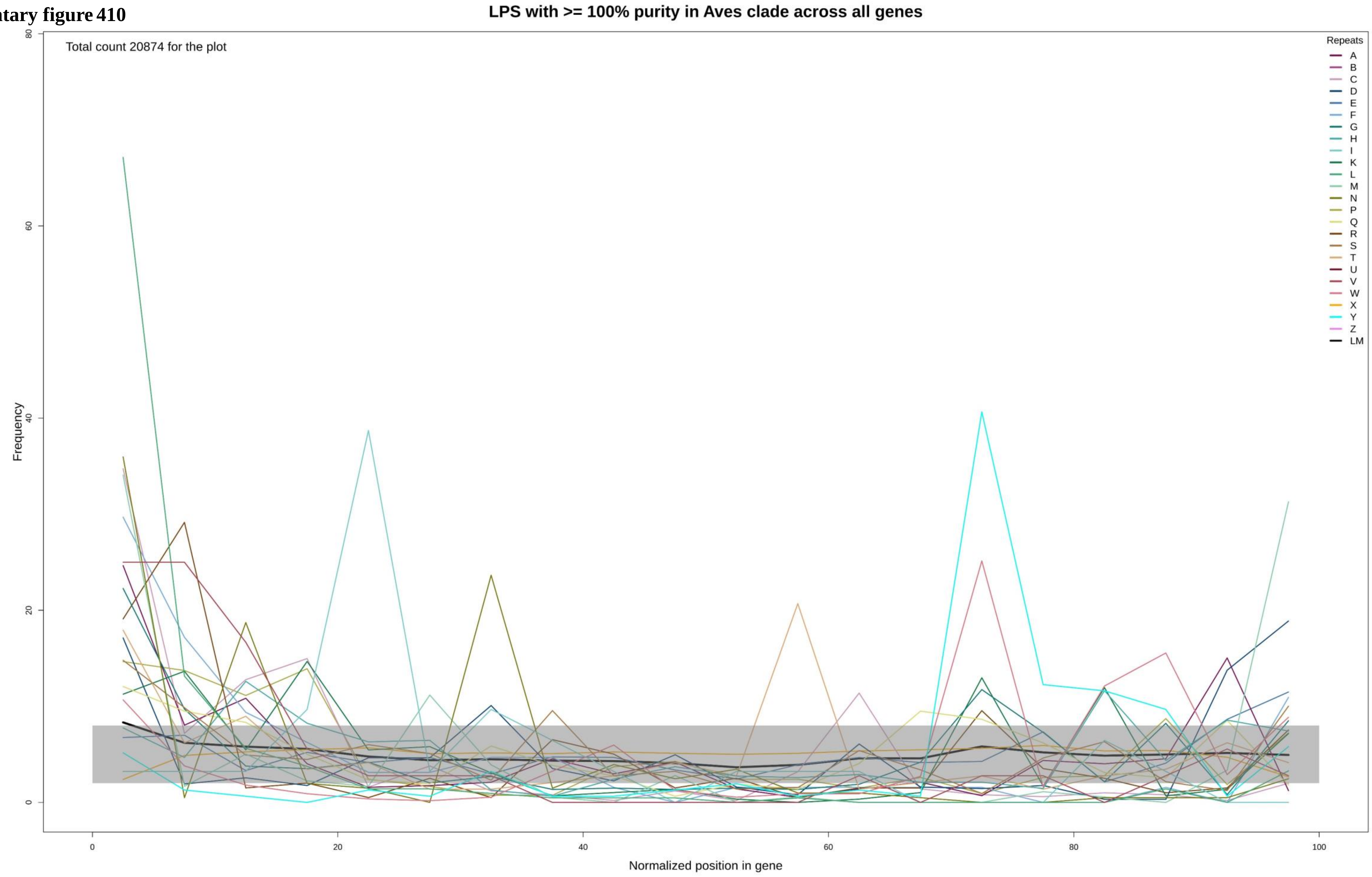

**Supplementary figure 411**

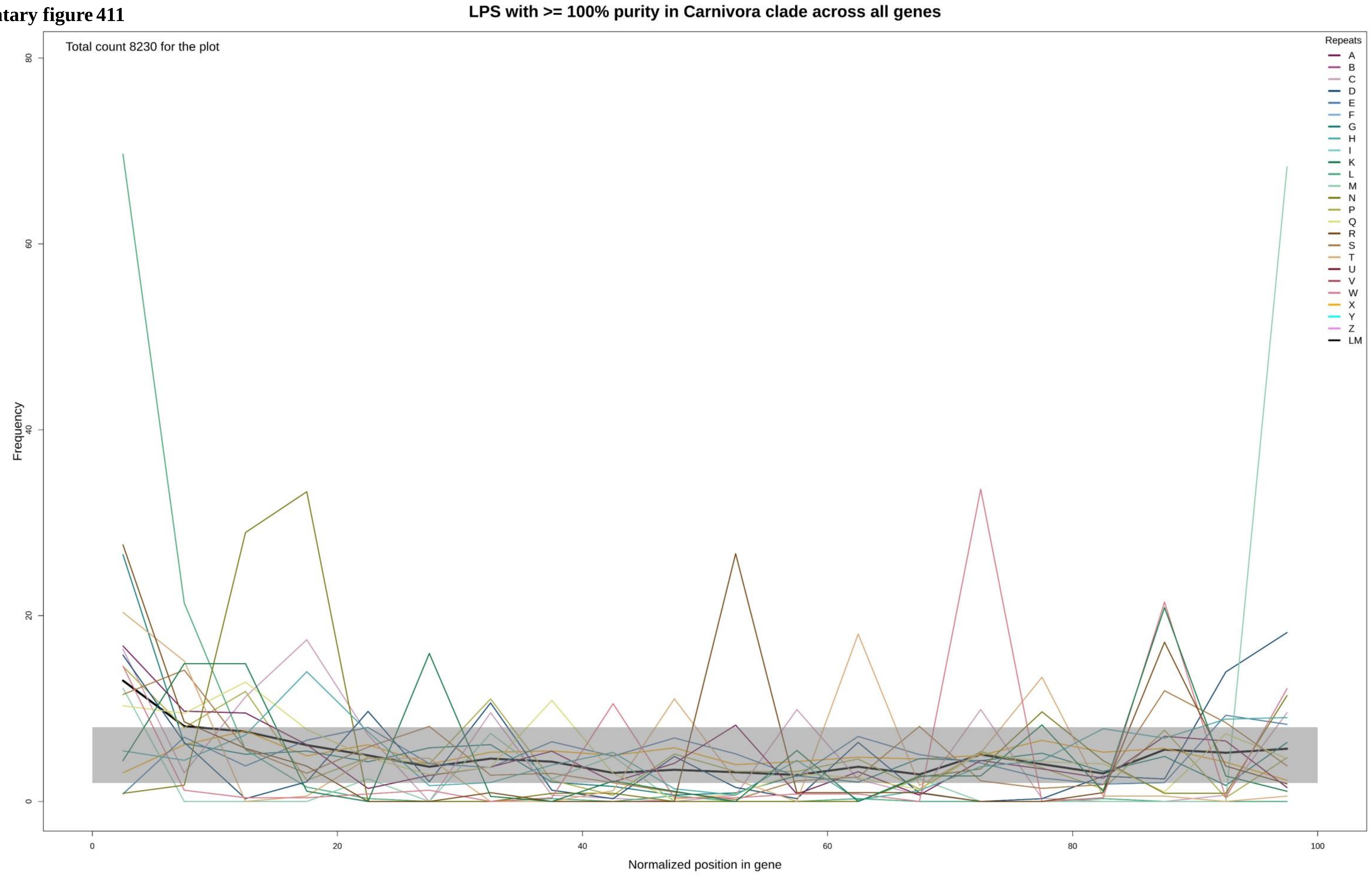

**Supplementary figure 412**

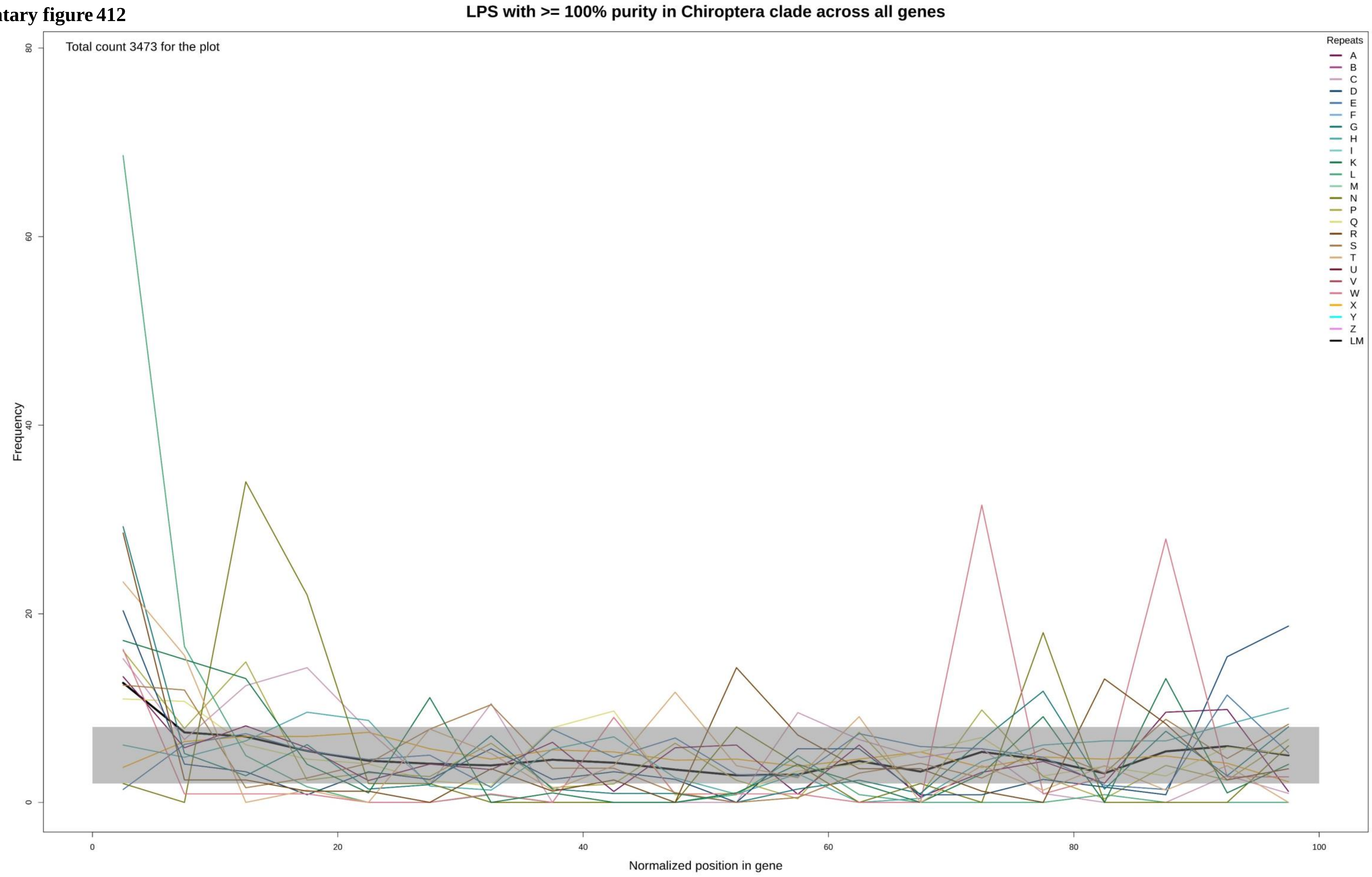

**Supplementary figure 413**

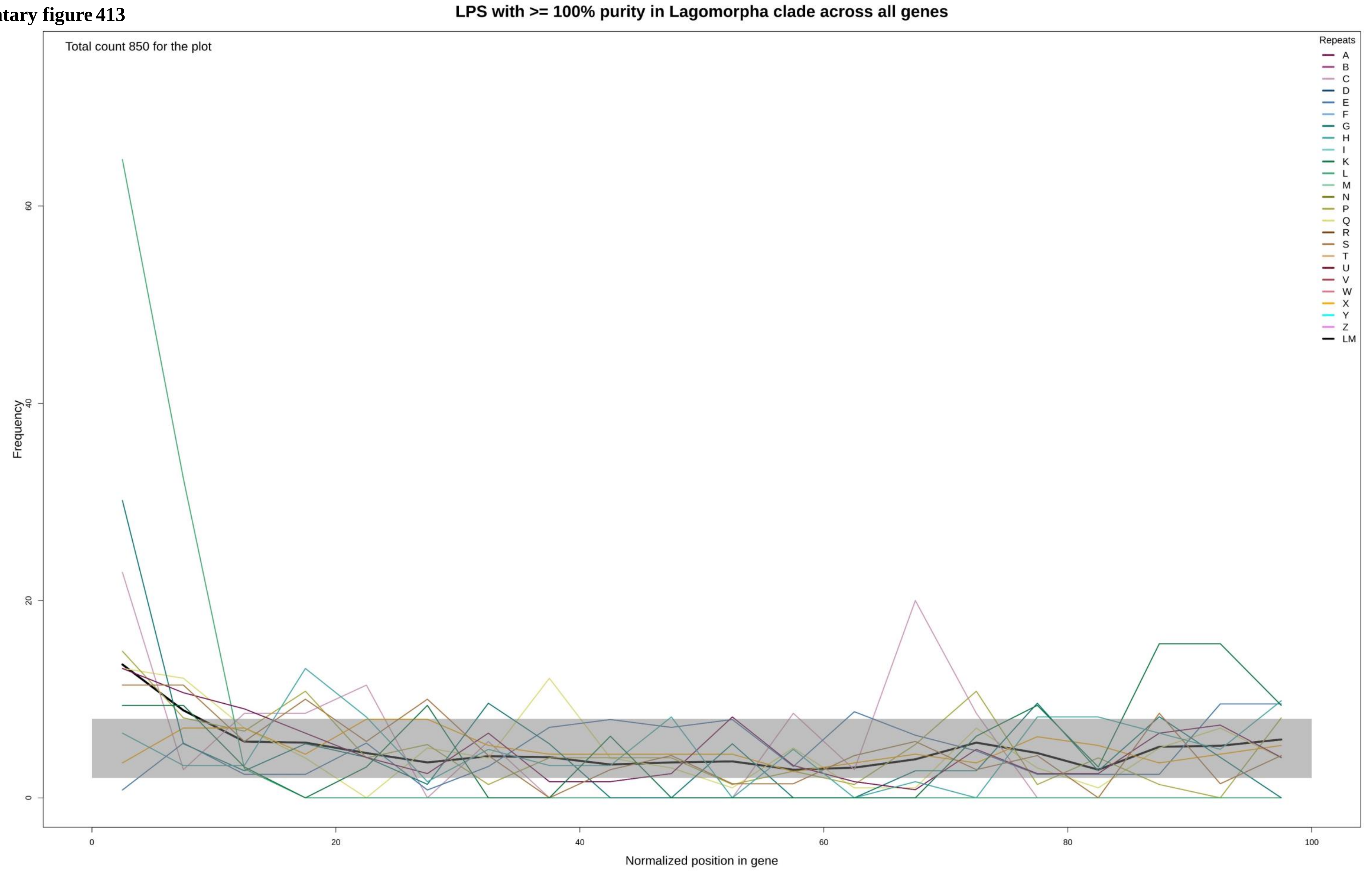

**Supplementary figure 414**

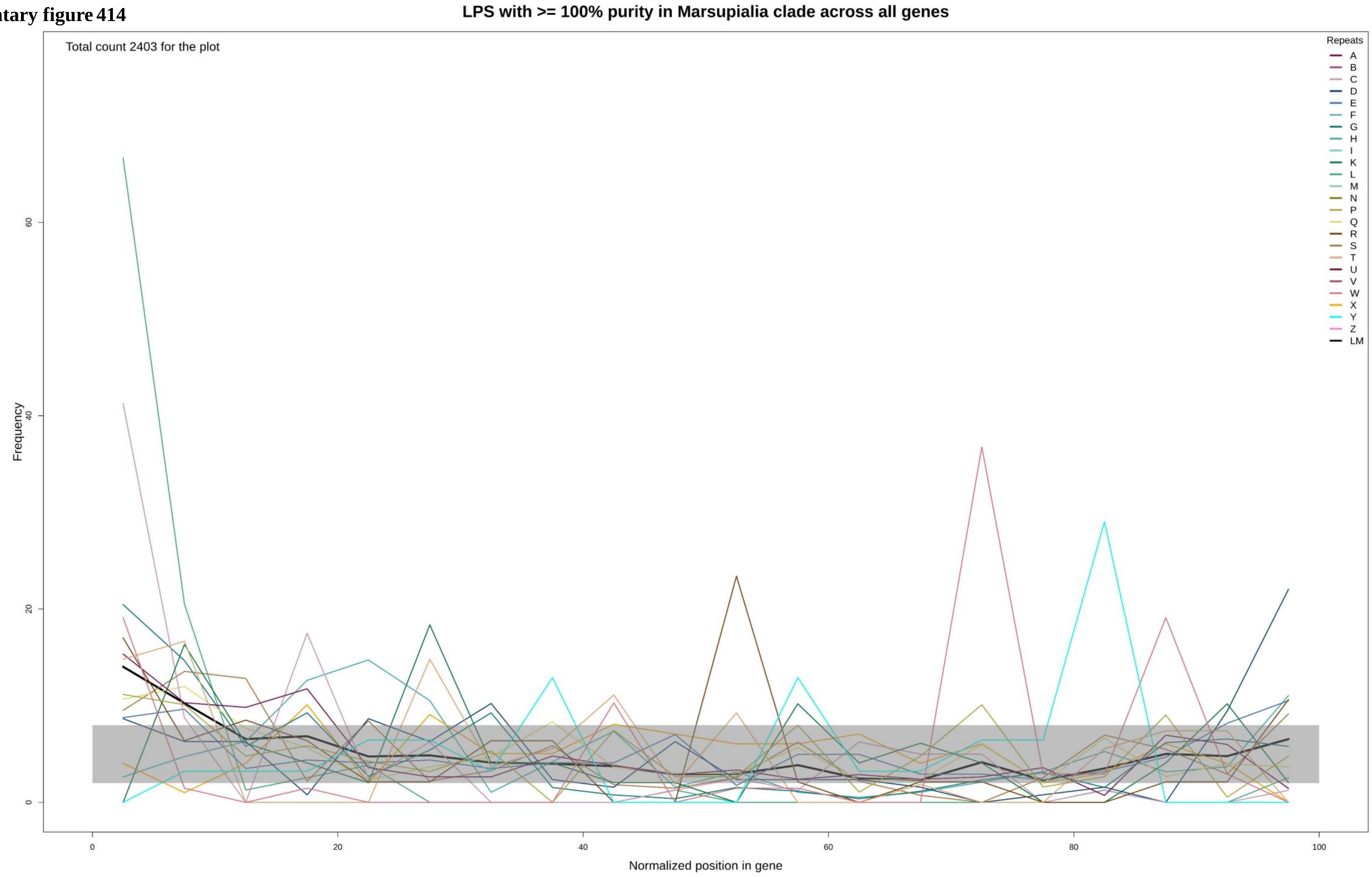

**Supplementary figure 415**

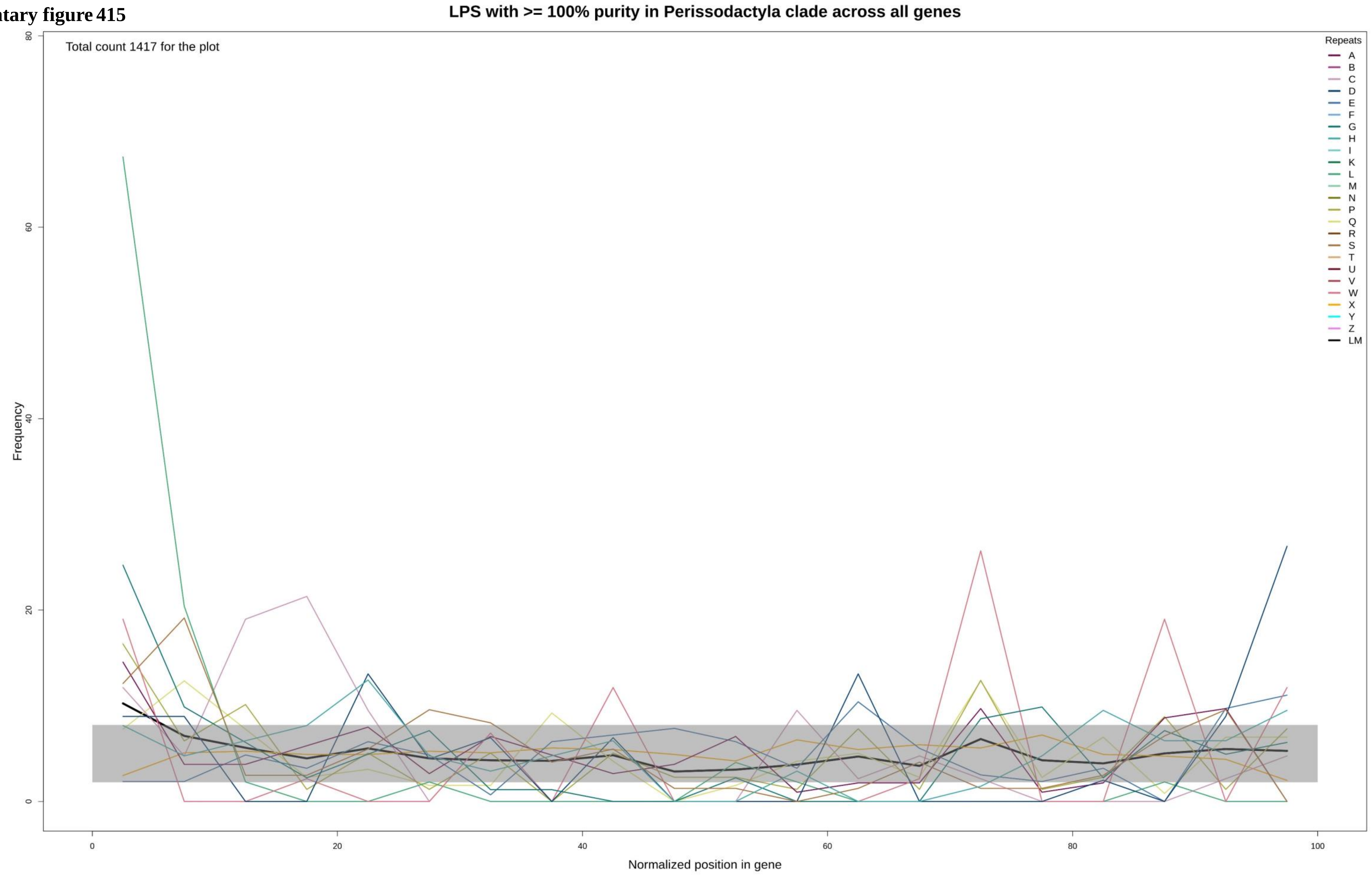

**Supplementary figure 416**

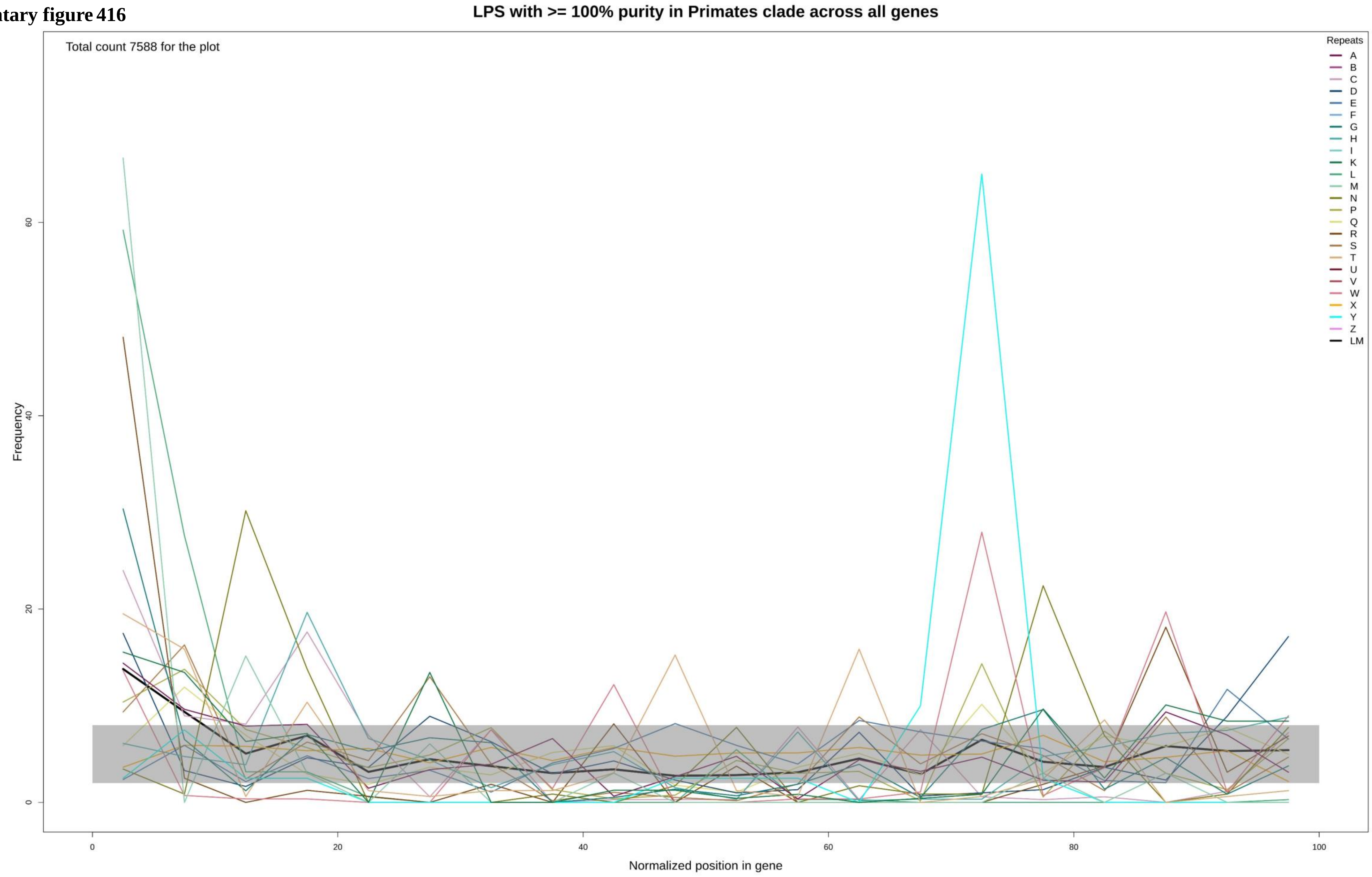

**Supplementary figure 417**

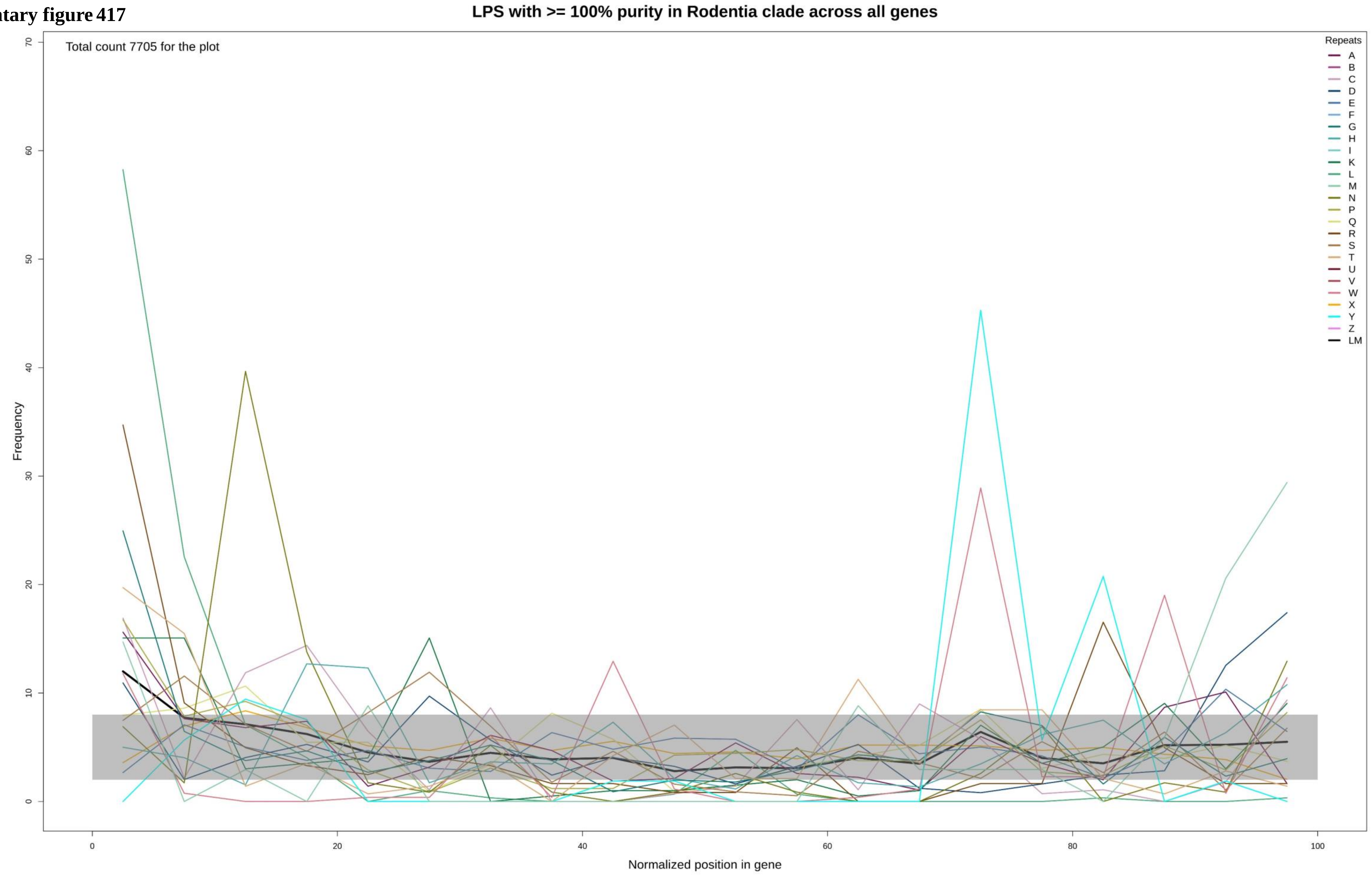

**Supplementary figure 418**

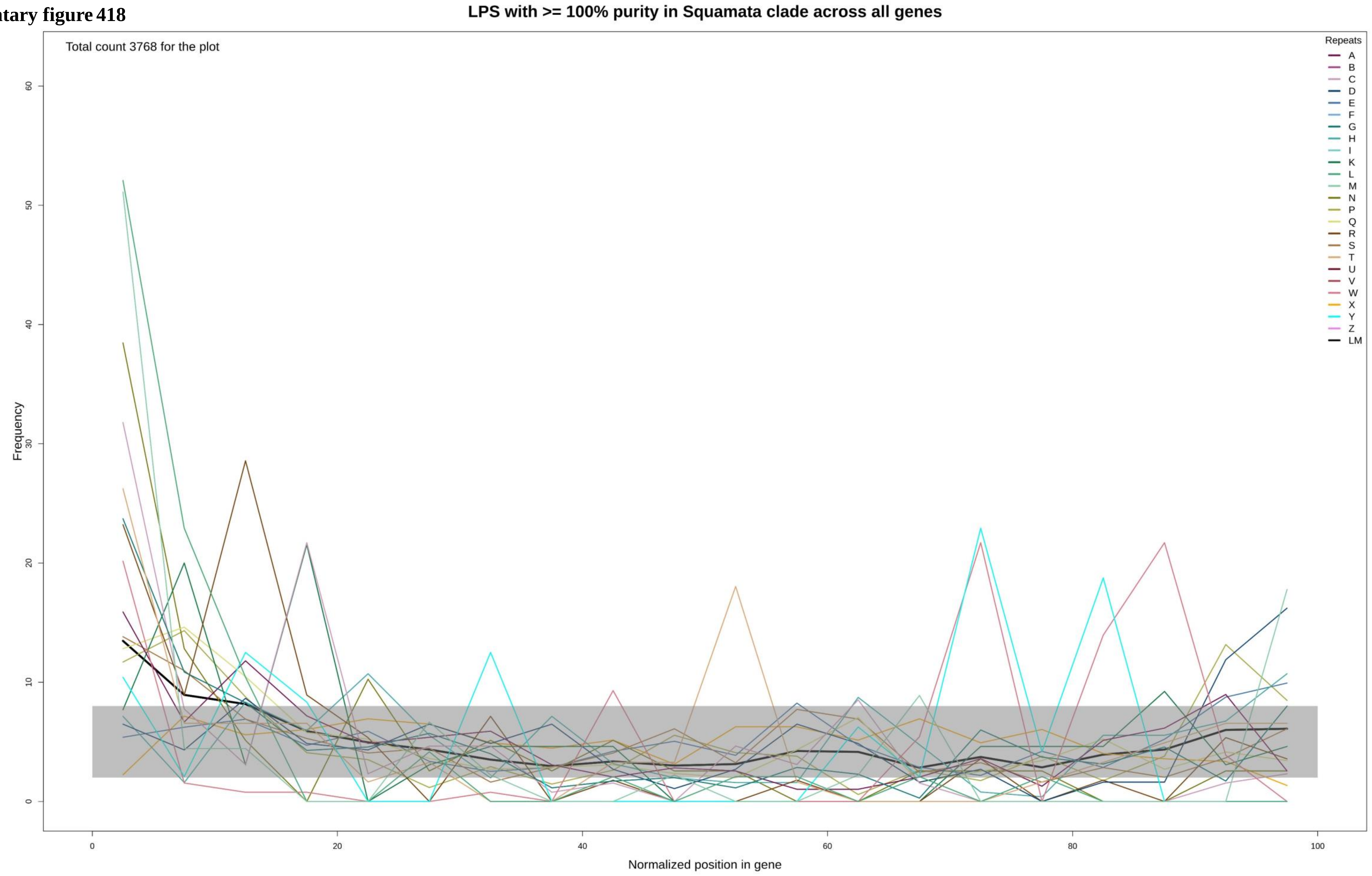

**Supplementary figure 419**

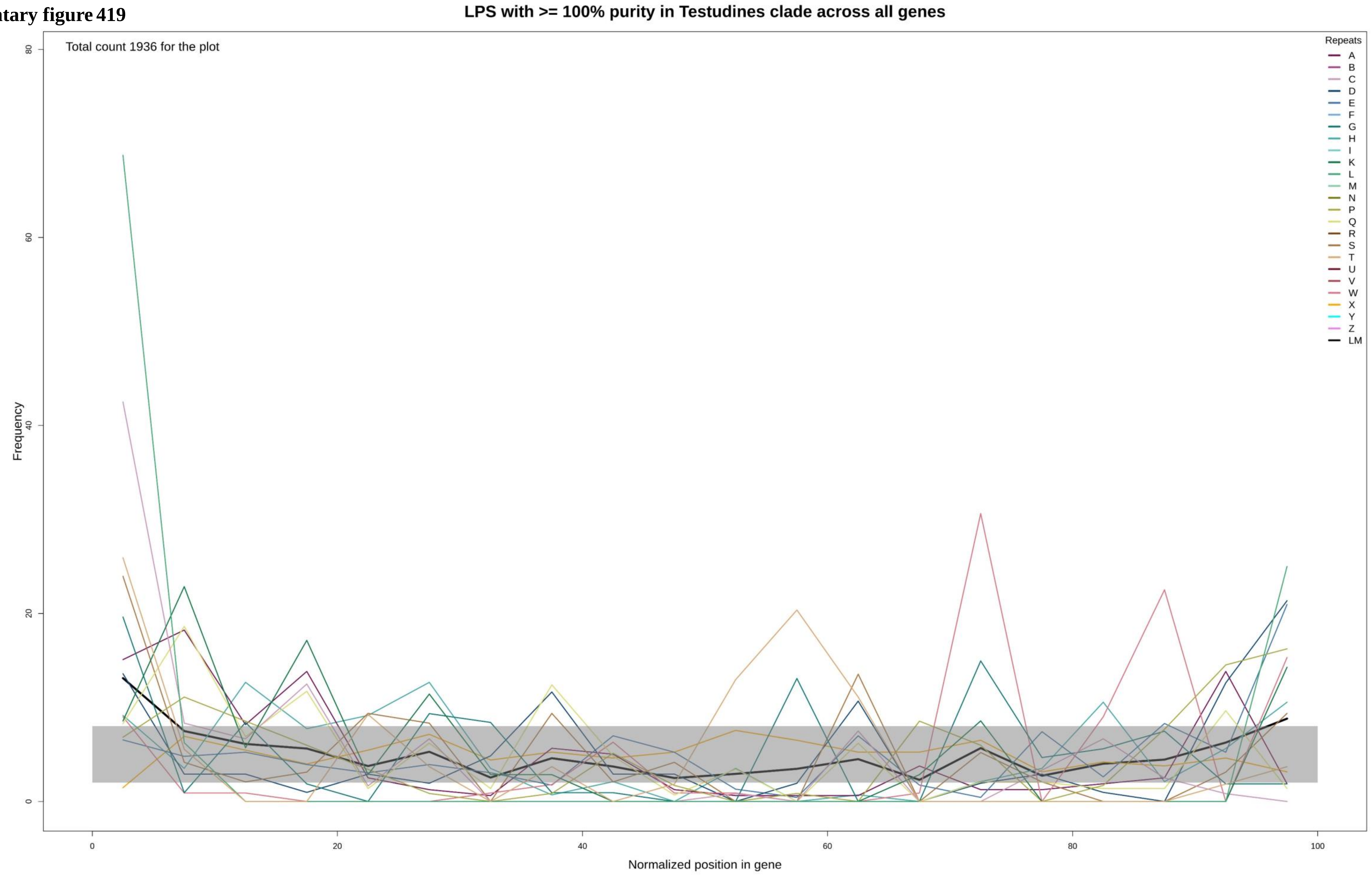

**Supplementary figure 420**

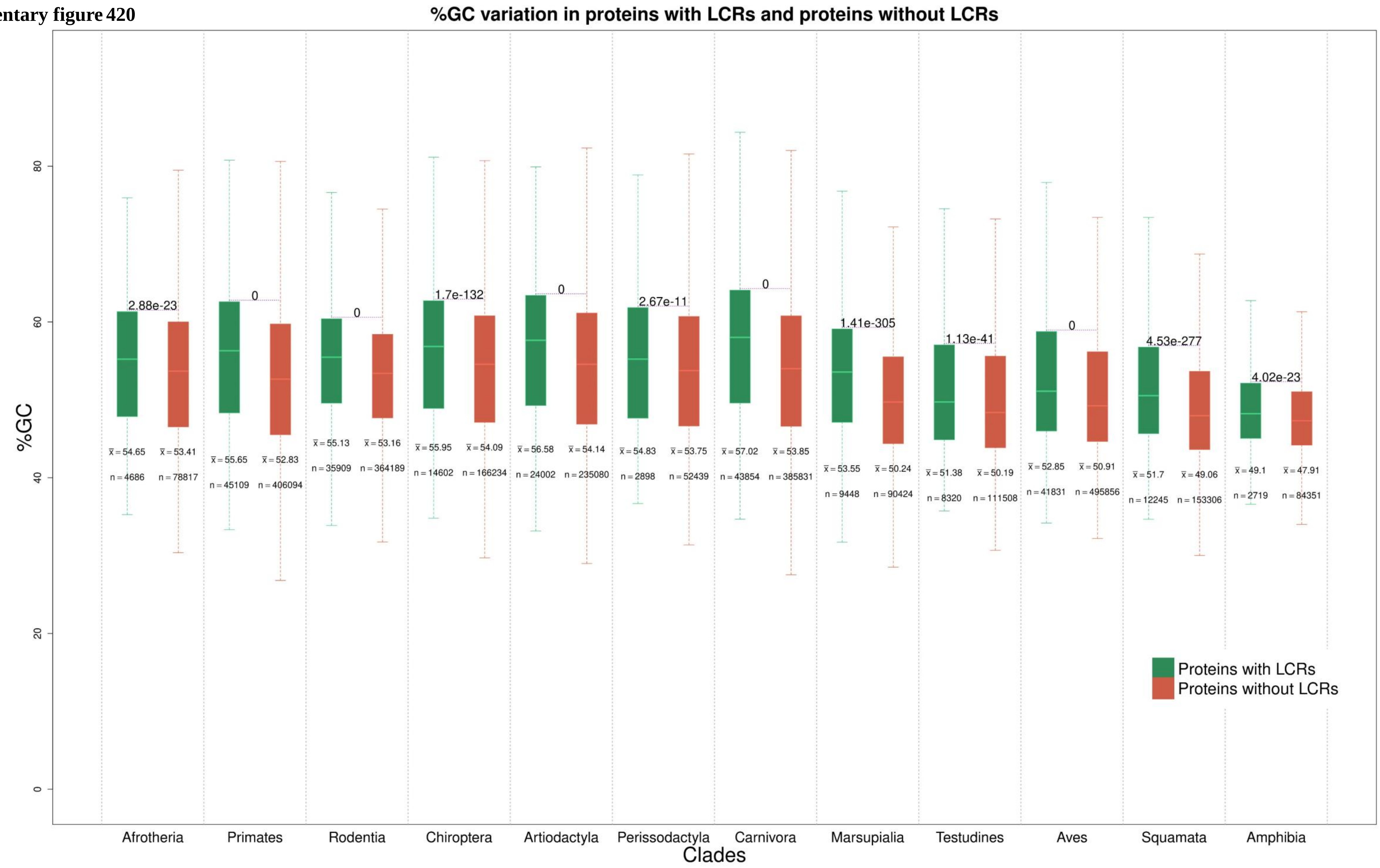

**Supplementary figure 421**

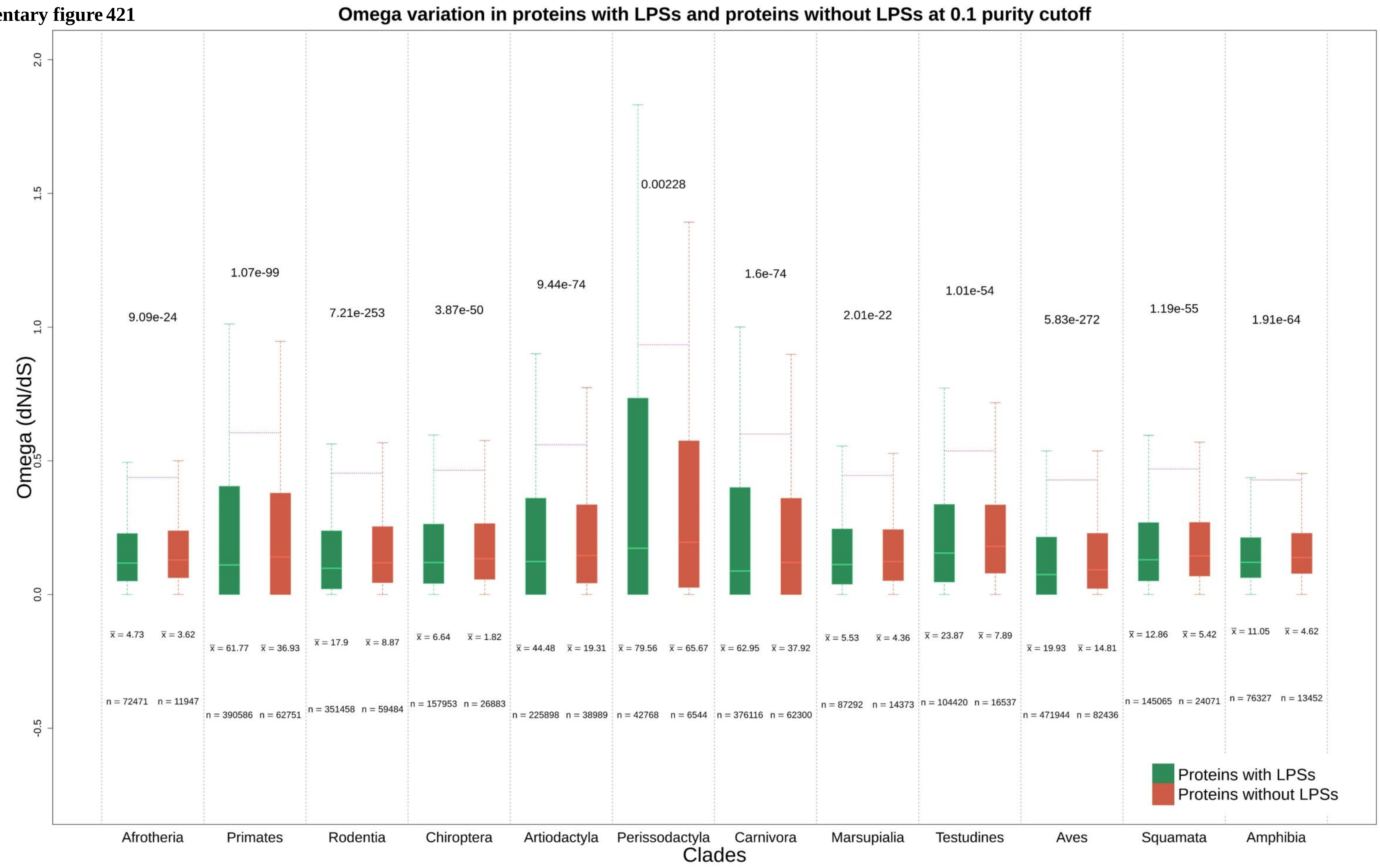

Supplementary figure 422

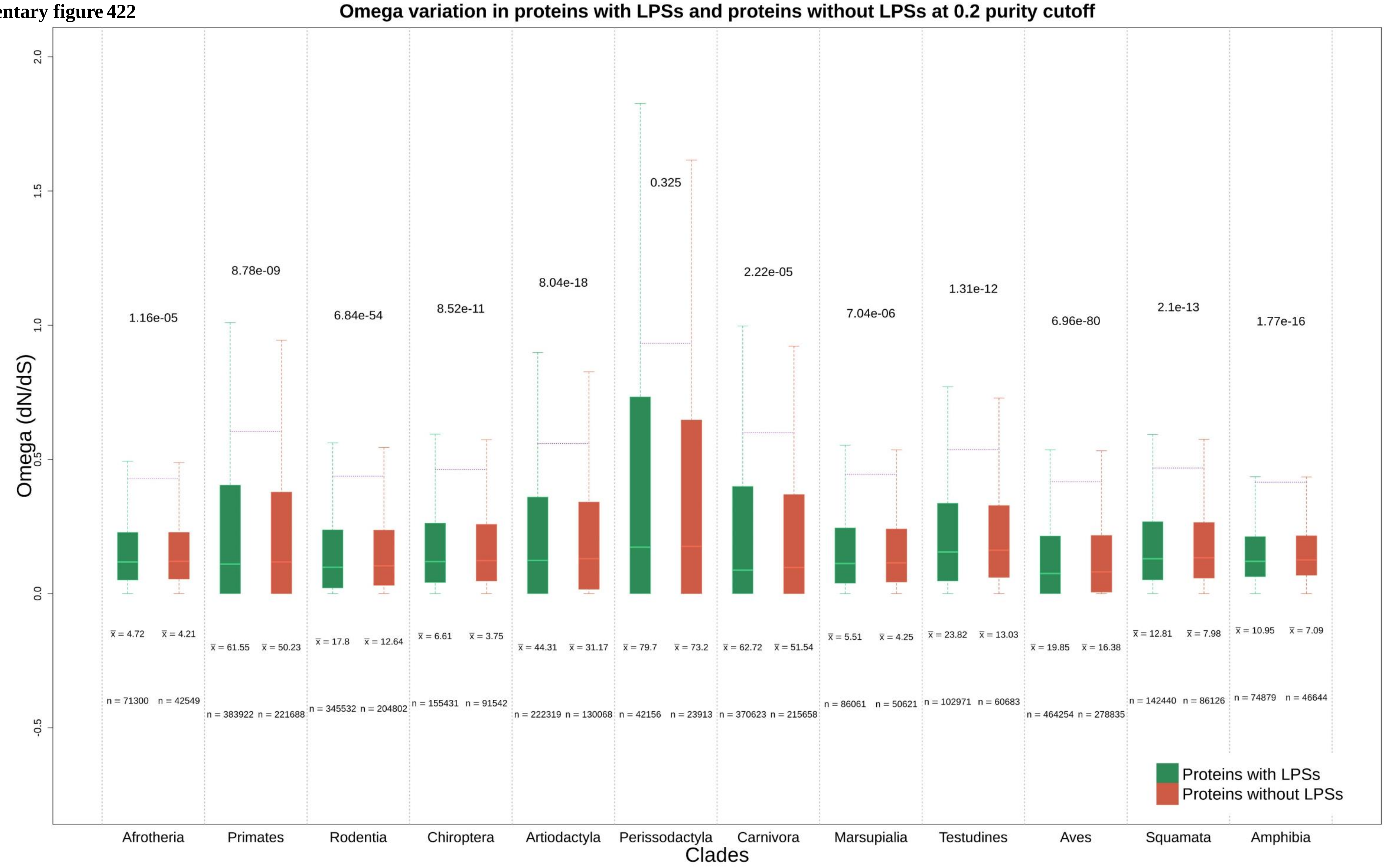

**Supplementary figure 423**

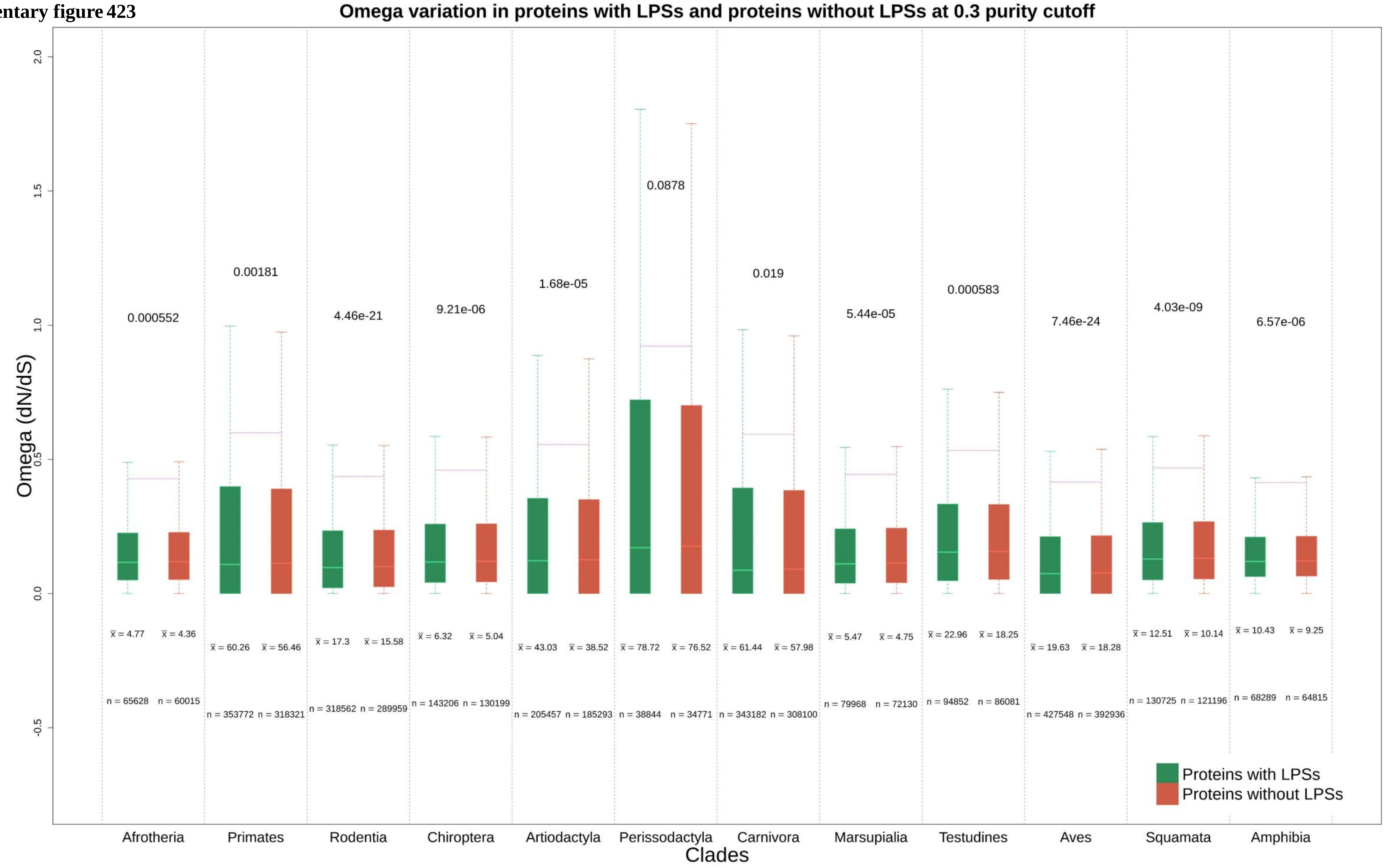

**Supplementary figure 424**

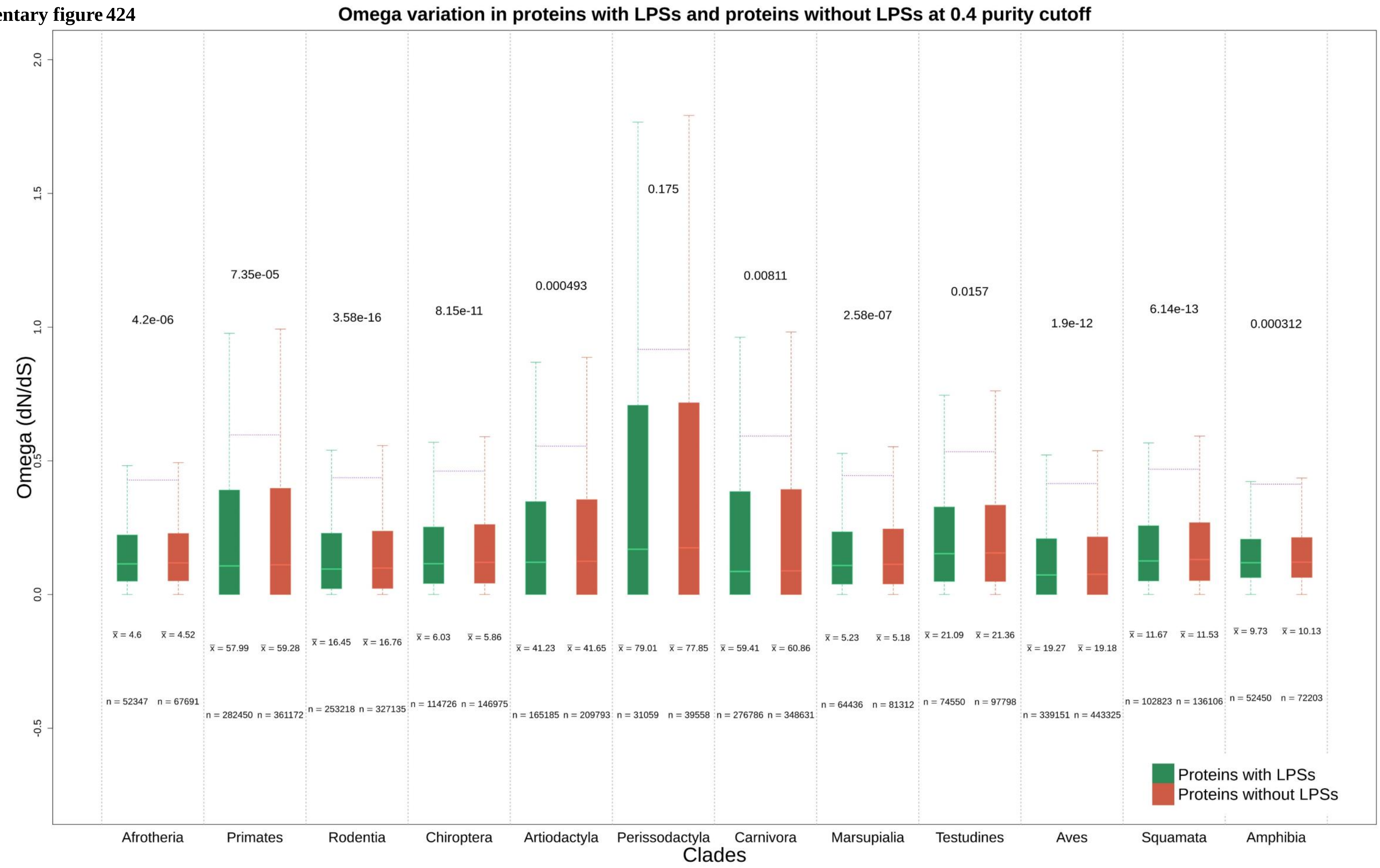

Supplementary figure 425

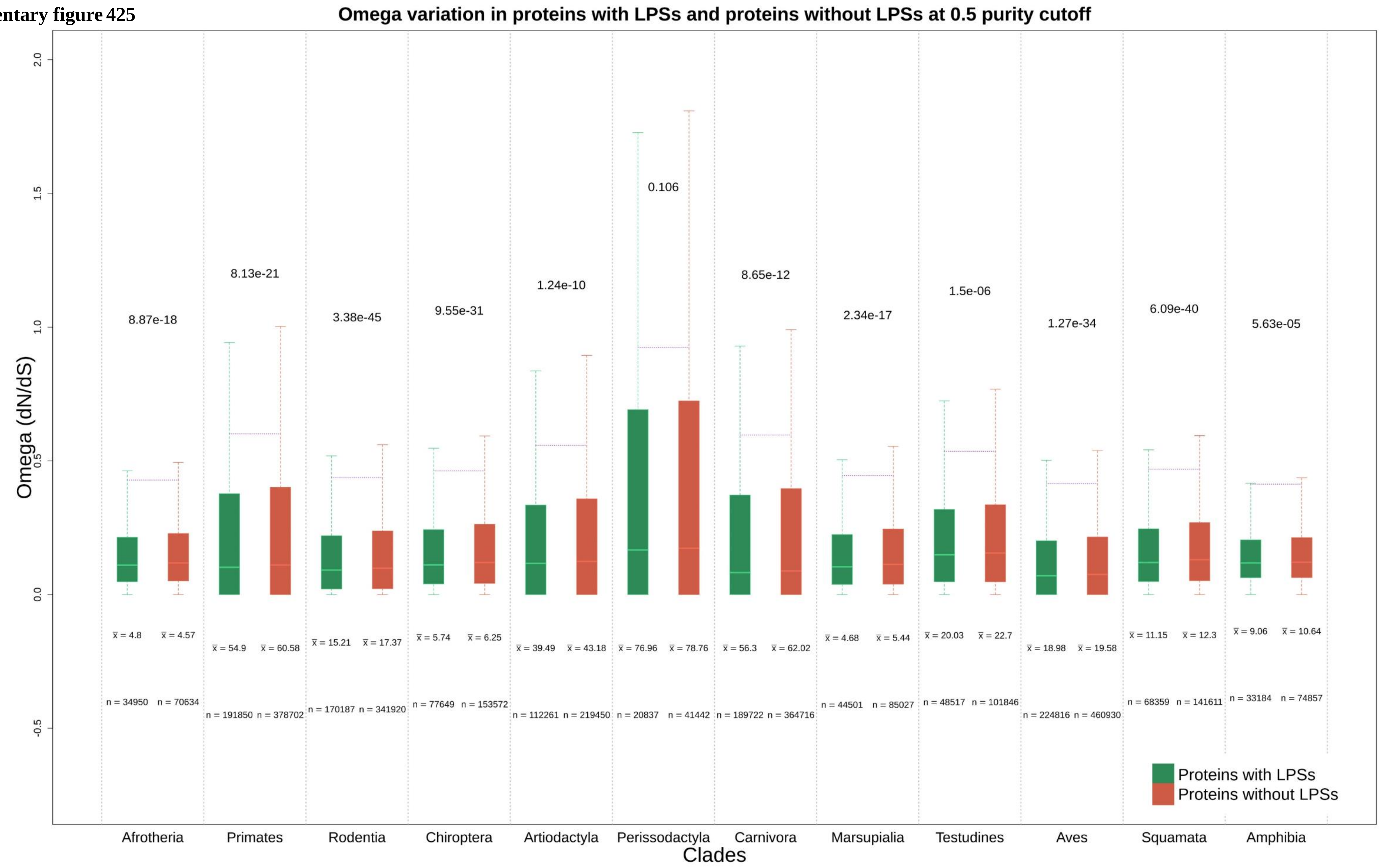

Supplementary figure 426

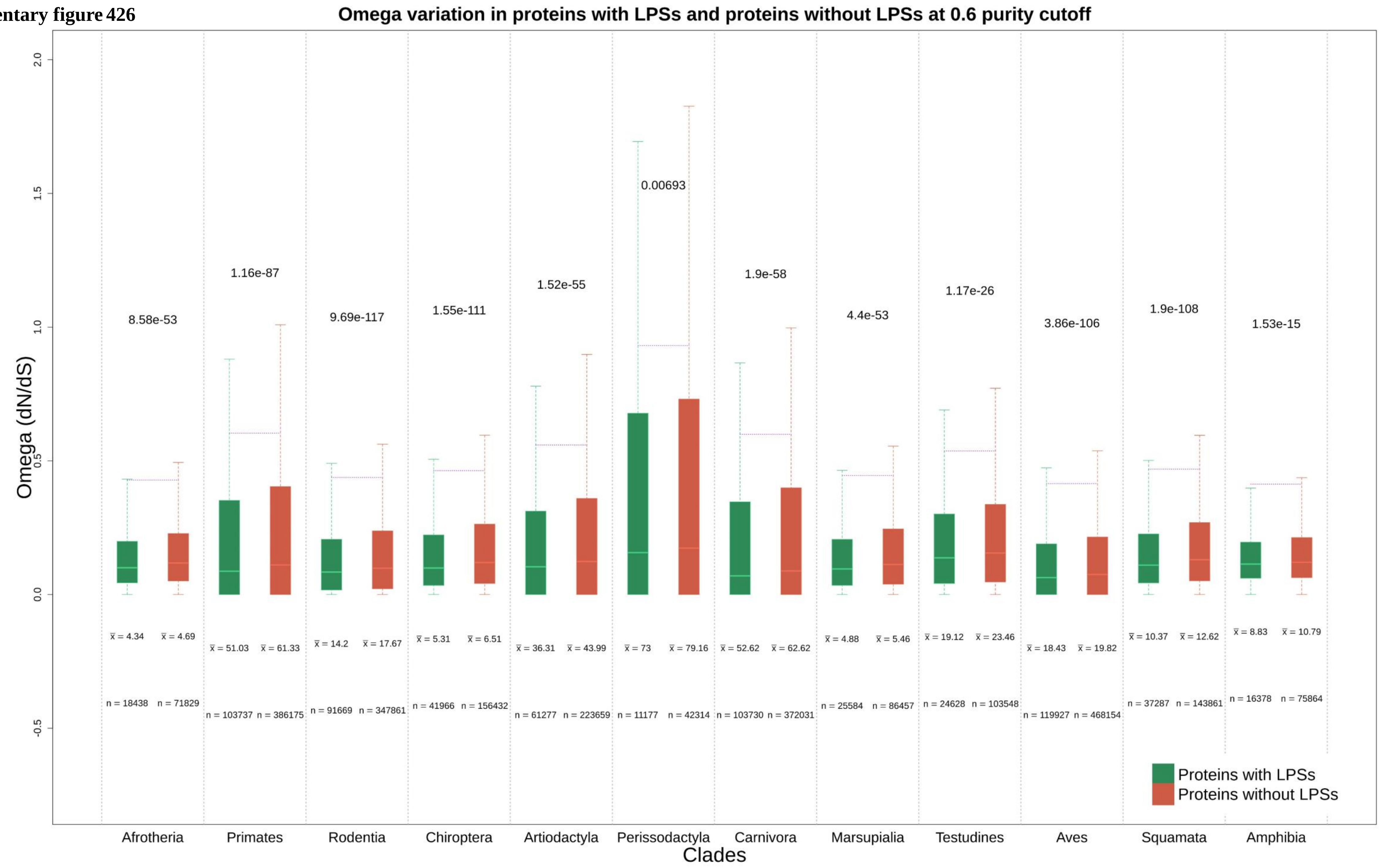

**Supplementary figure 427**

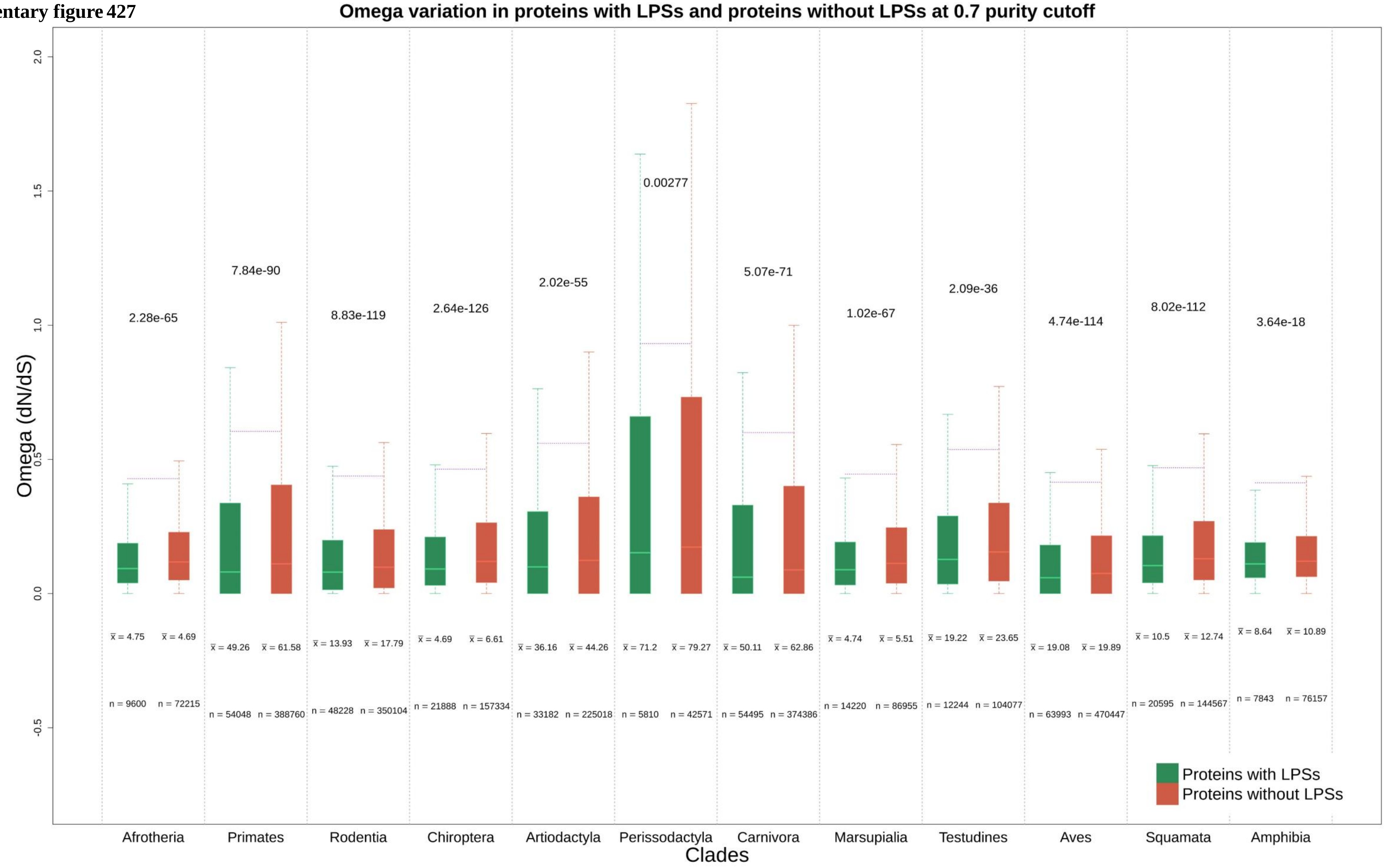

**Supplementary figure 428**

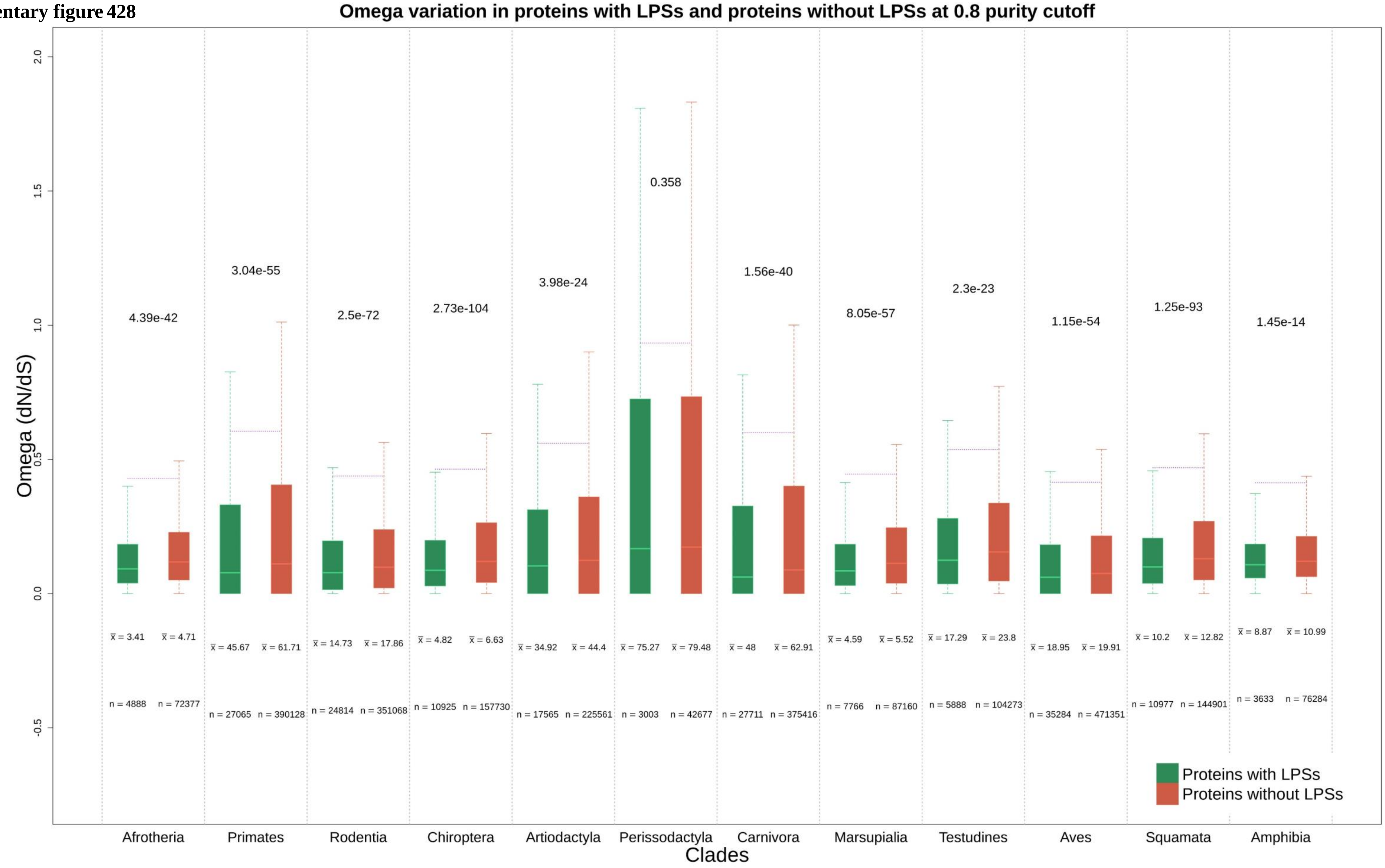

Supplementary figure 429

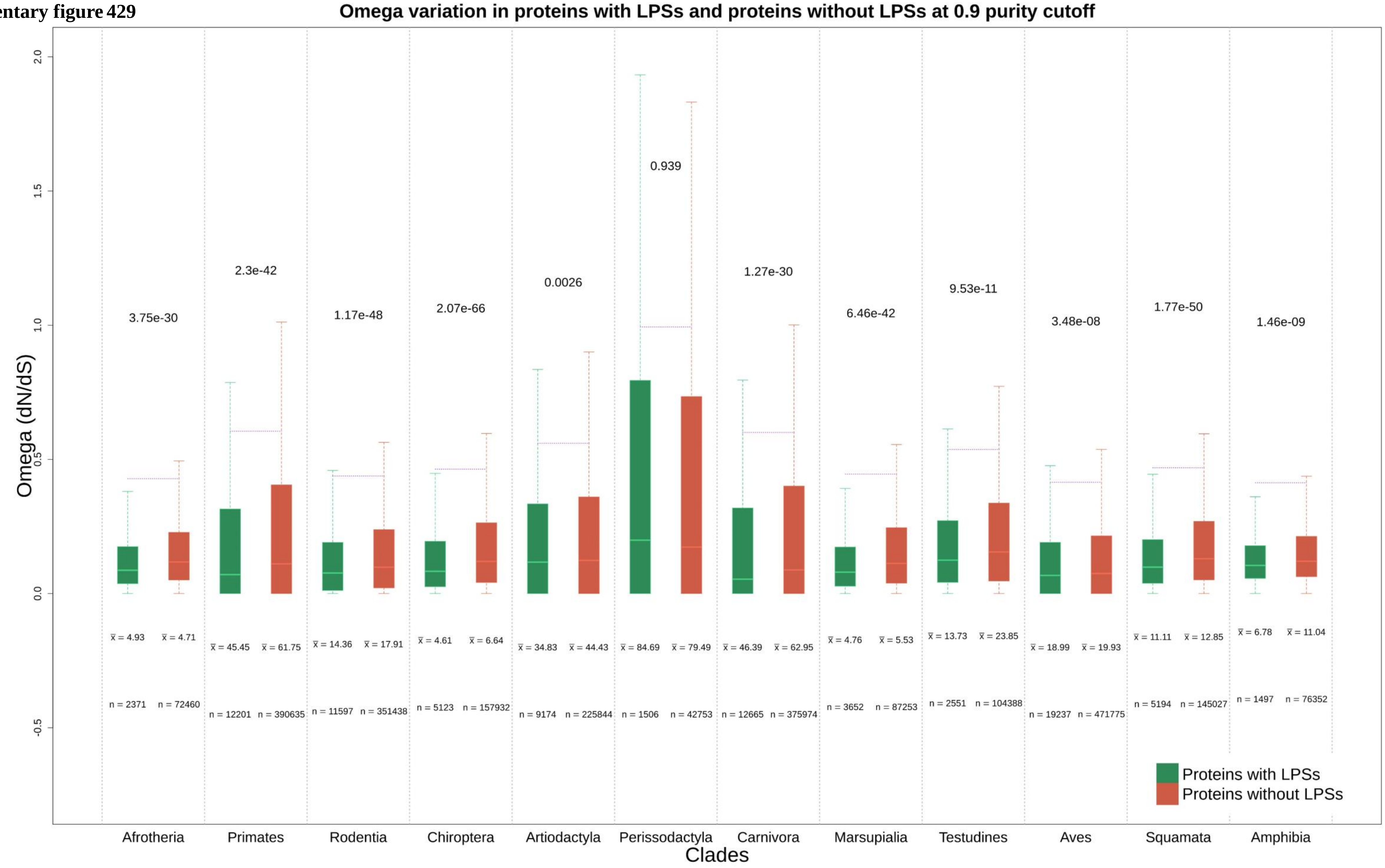

Supplementary figure 430

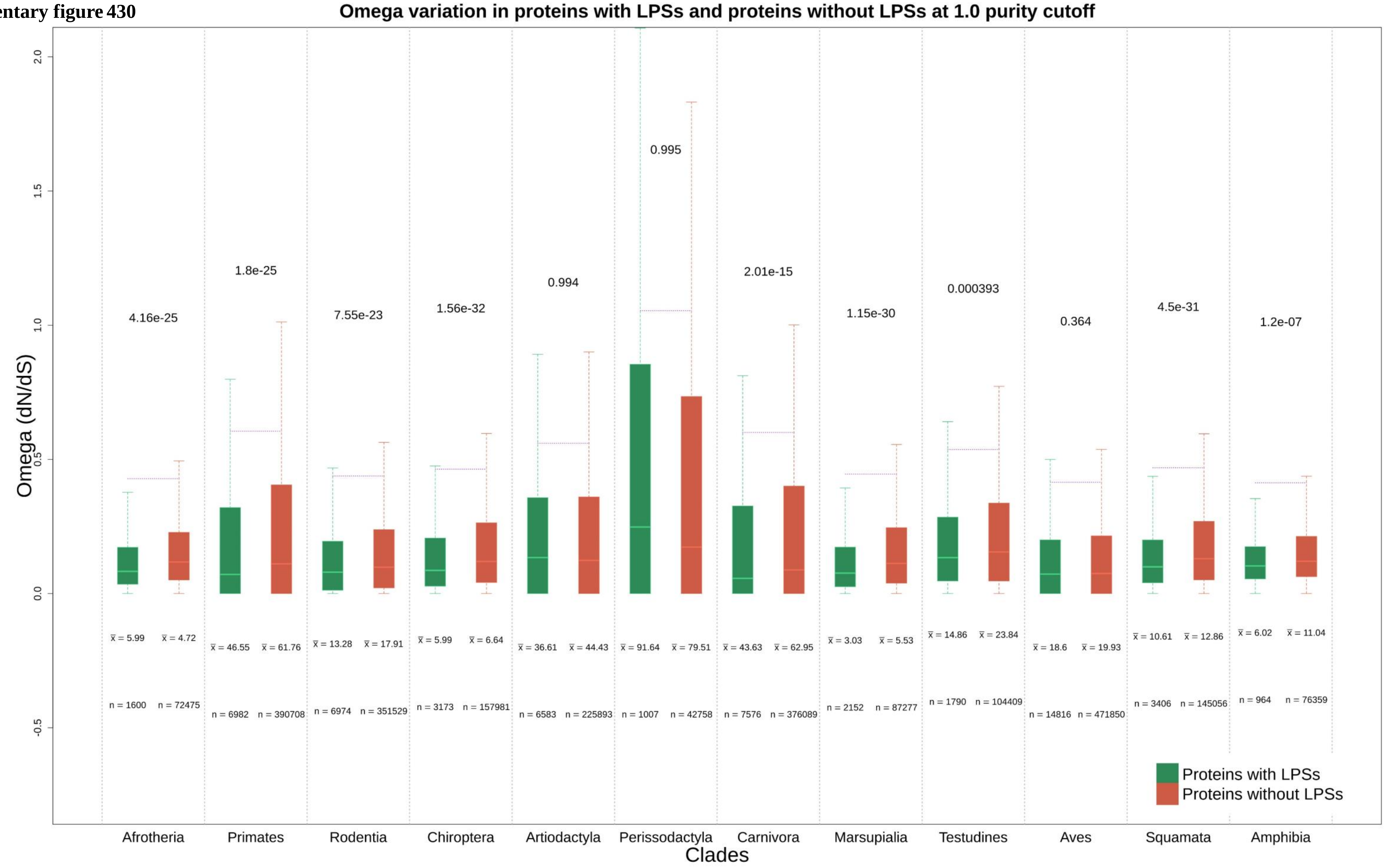

**Supplementary figure 431**

**Omega (<=2) variation in proteins with LPSs and proteins without LPSs at 0.1 purity cutoff**

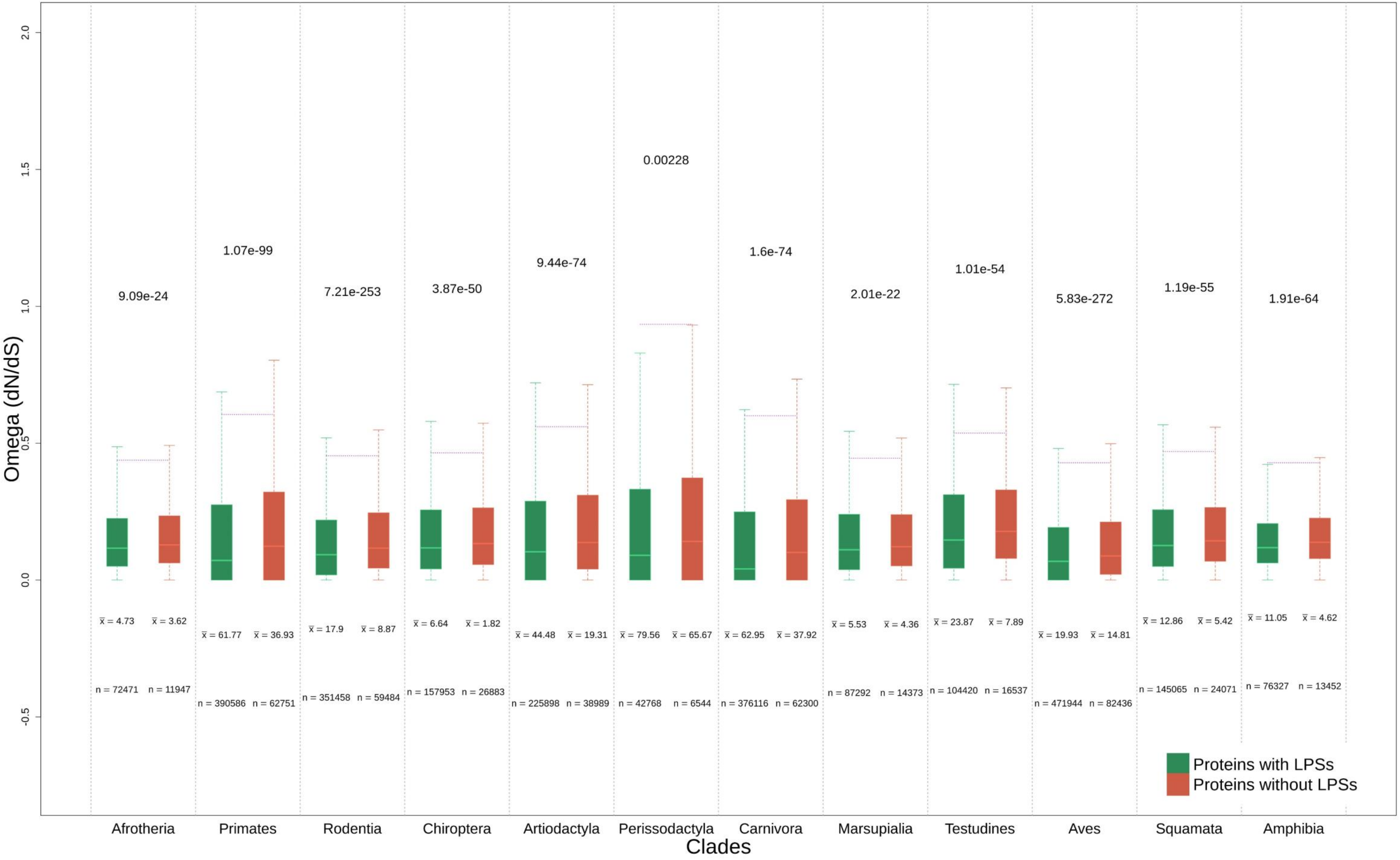

Supplementary figure 432

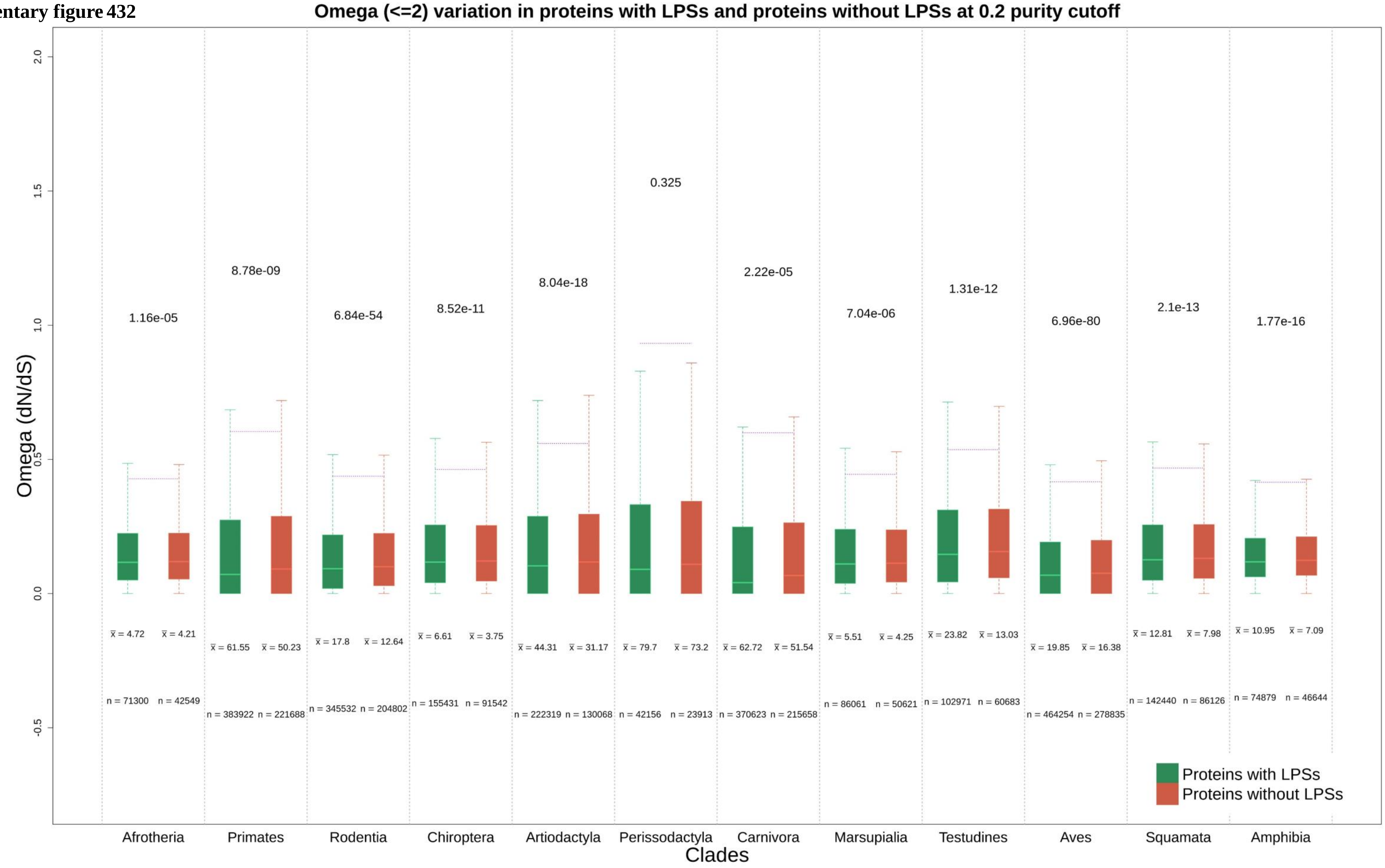

Supplementary figure 433

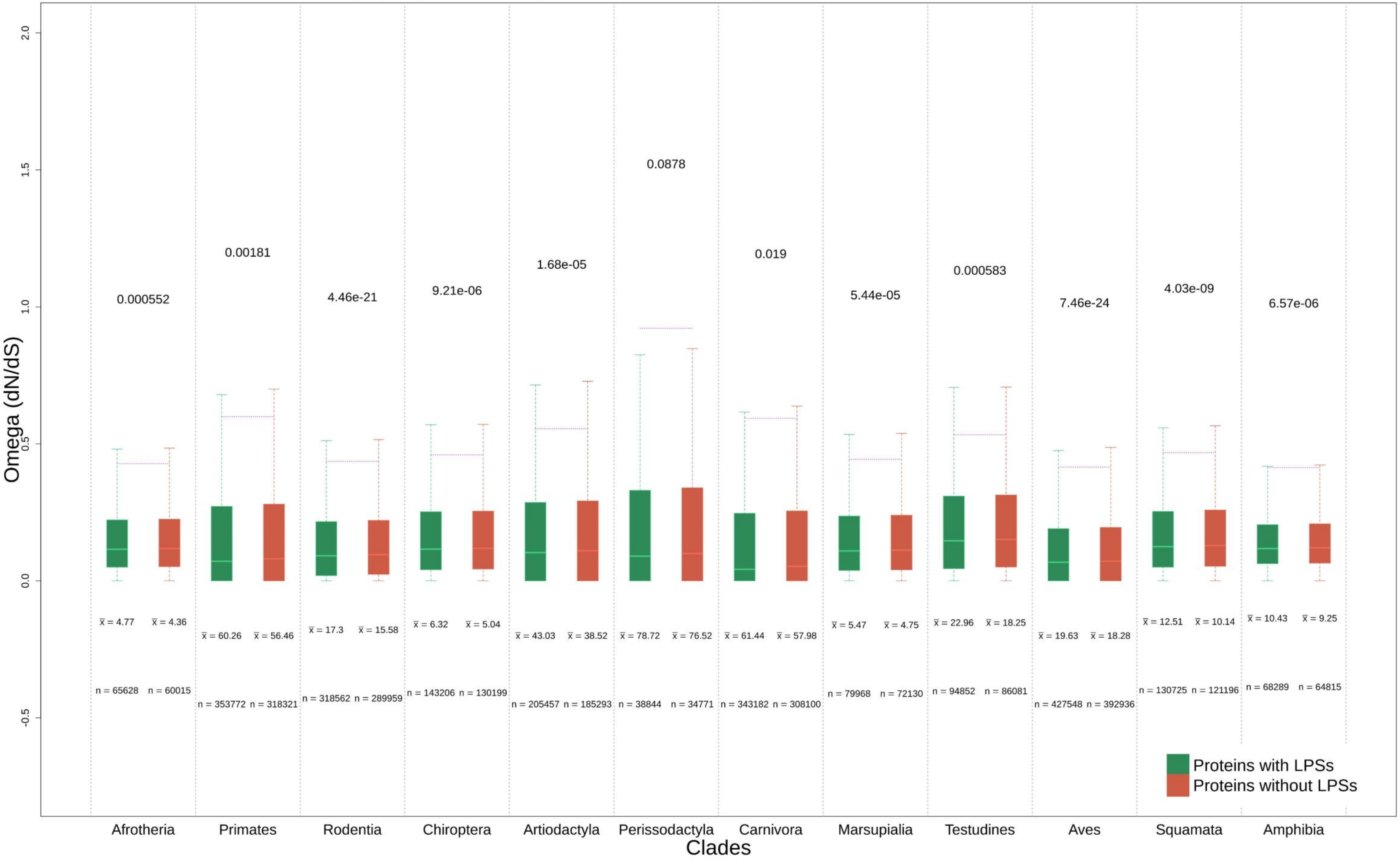

**Supplementary figure 434**

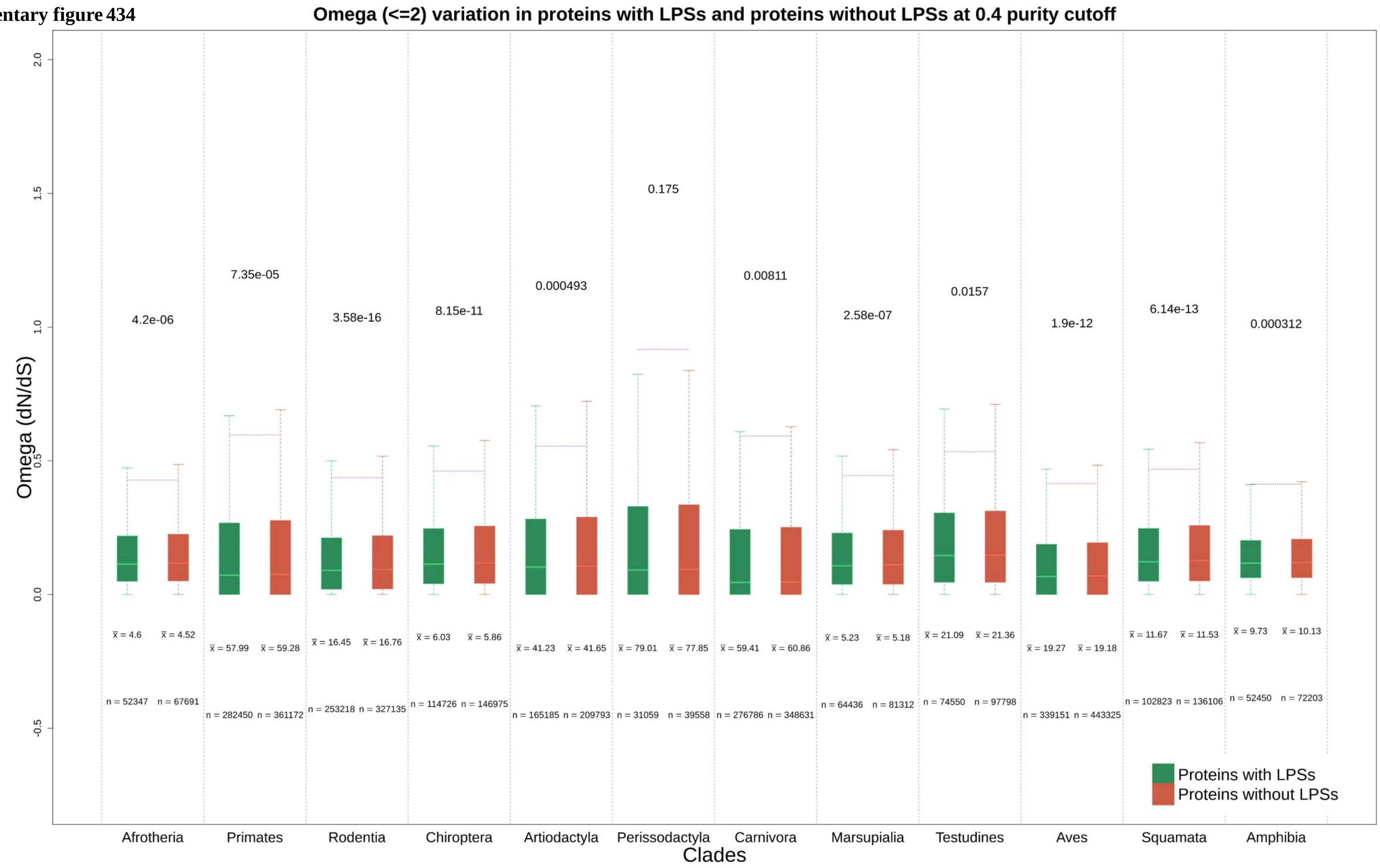

**Supplementary figure 435**

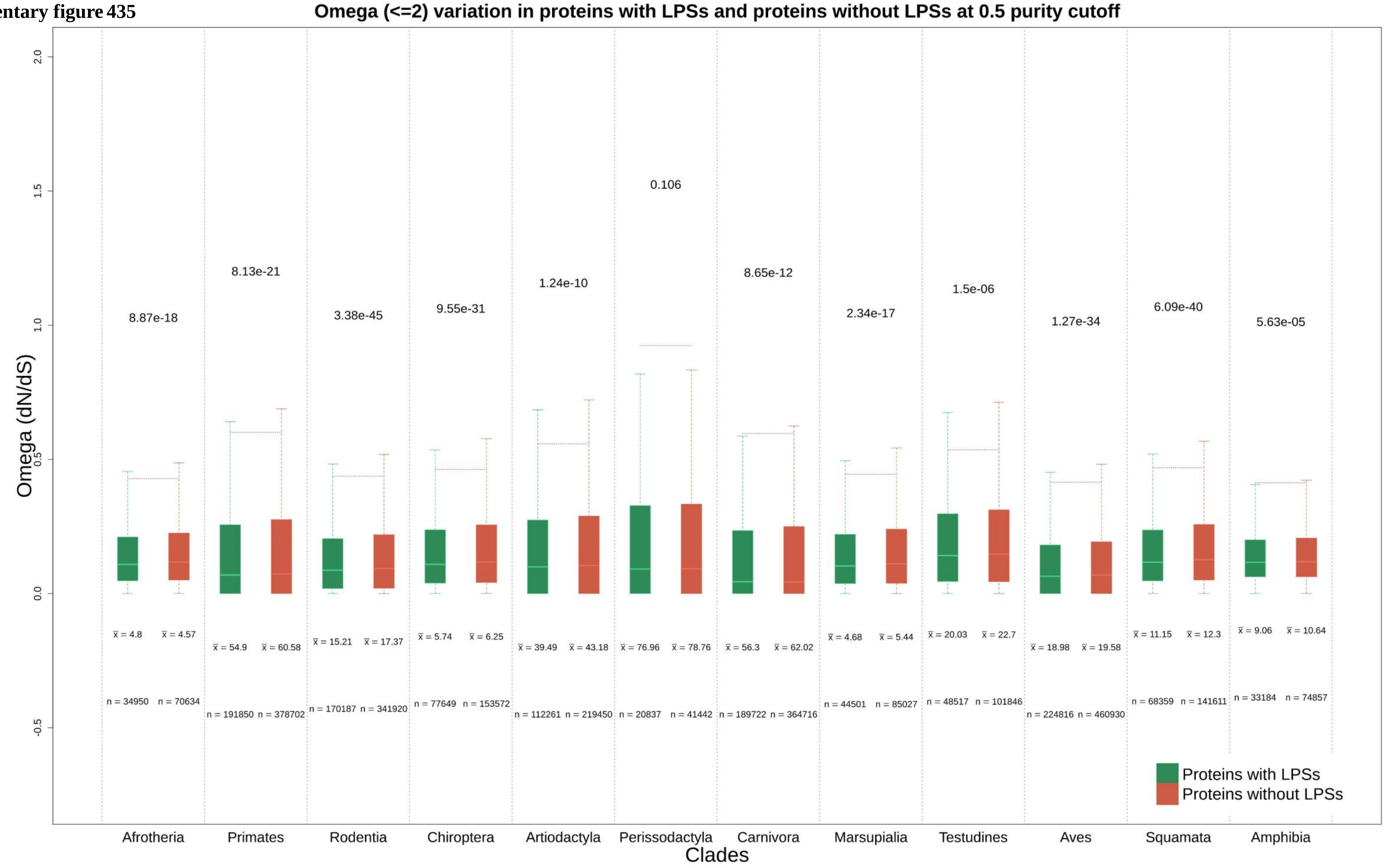

Supplementary figure 436

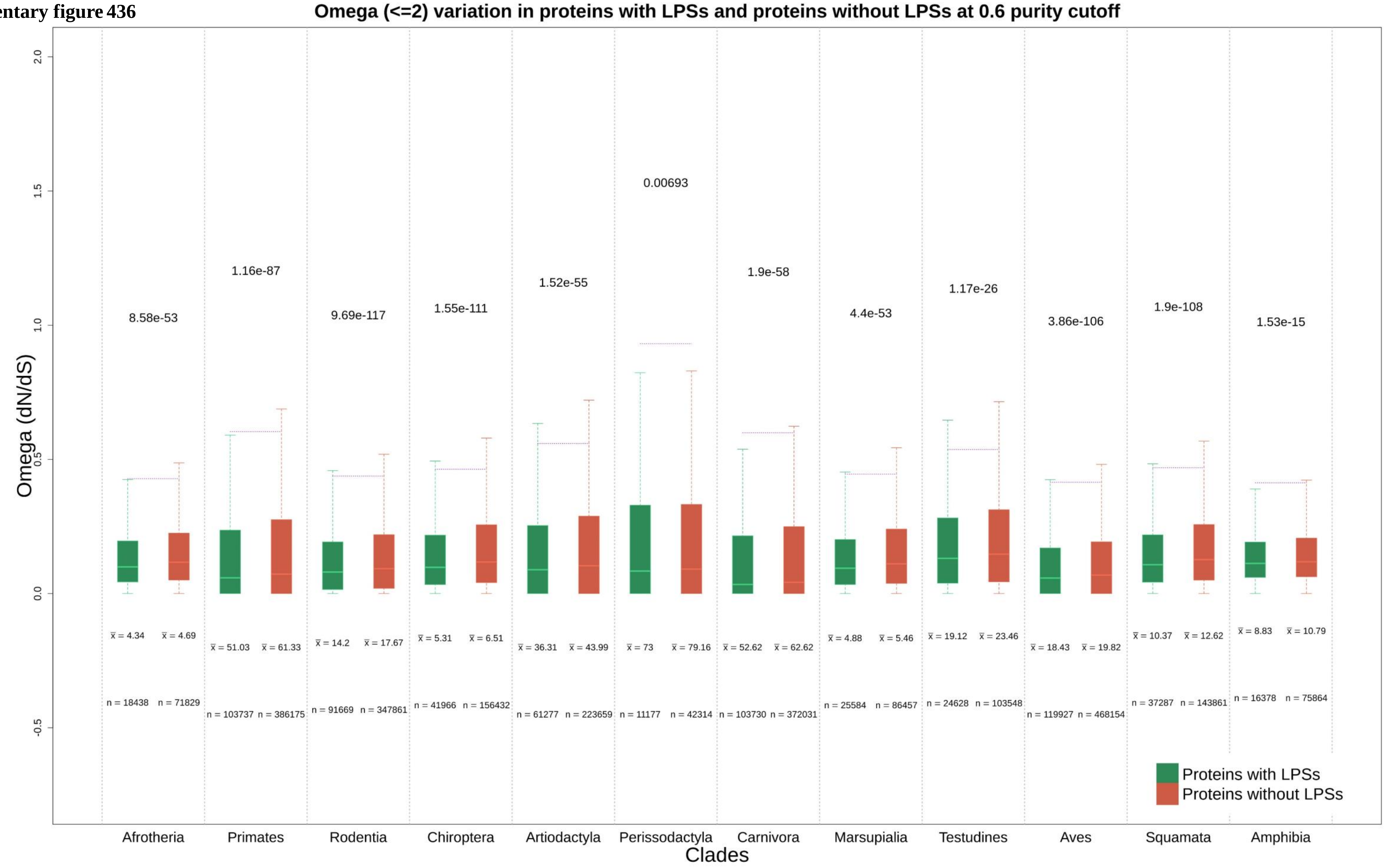

Supplementary figure 437

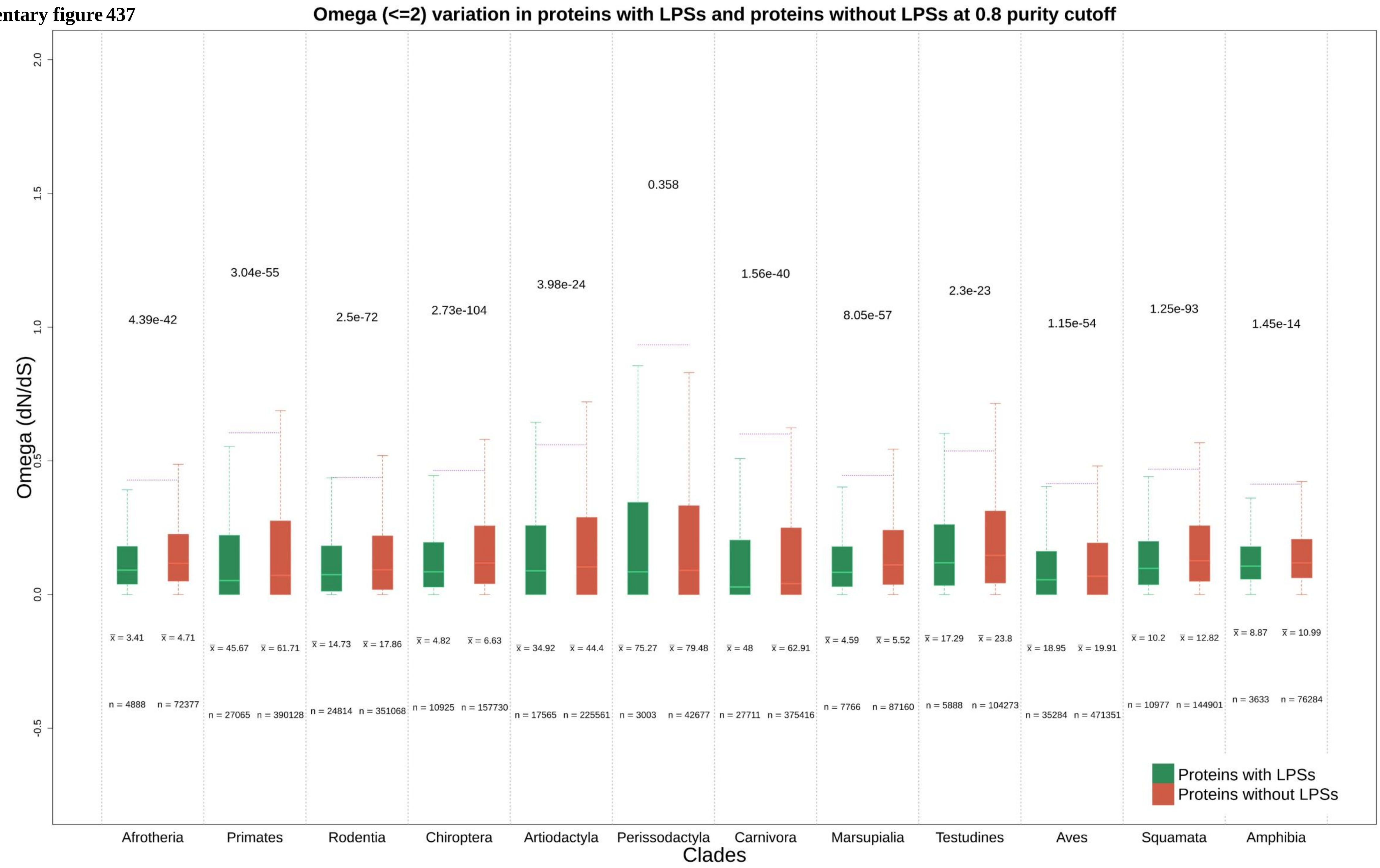

**Supplementary figure 438**

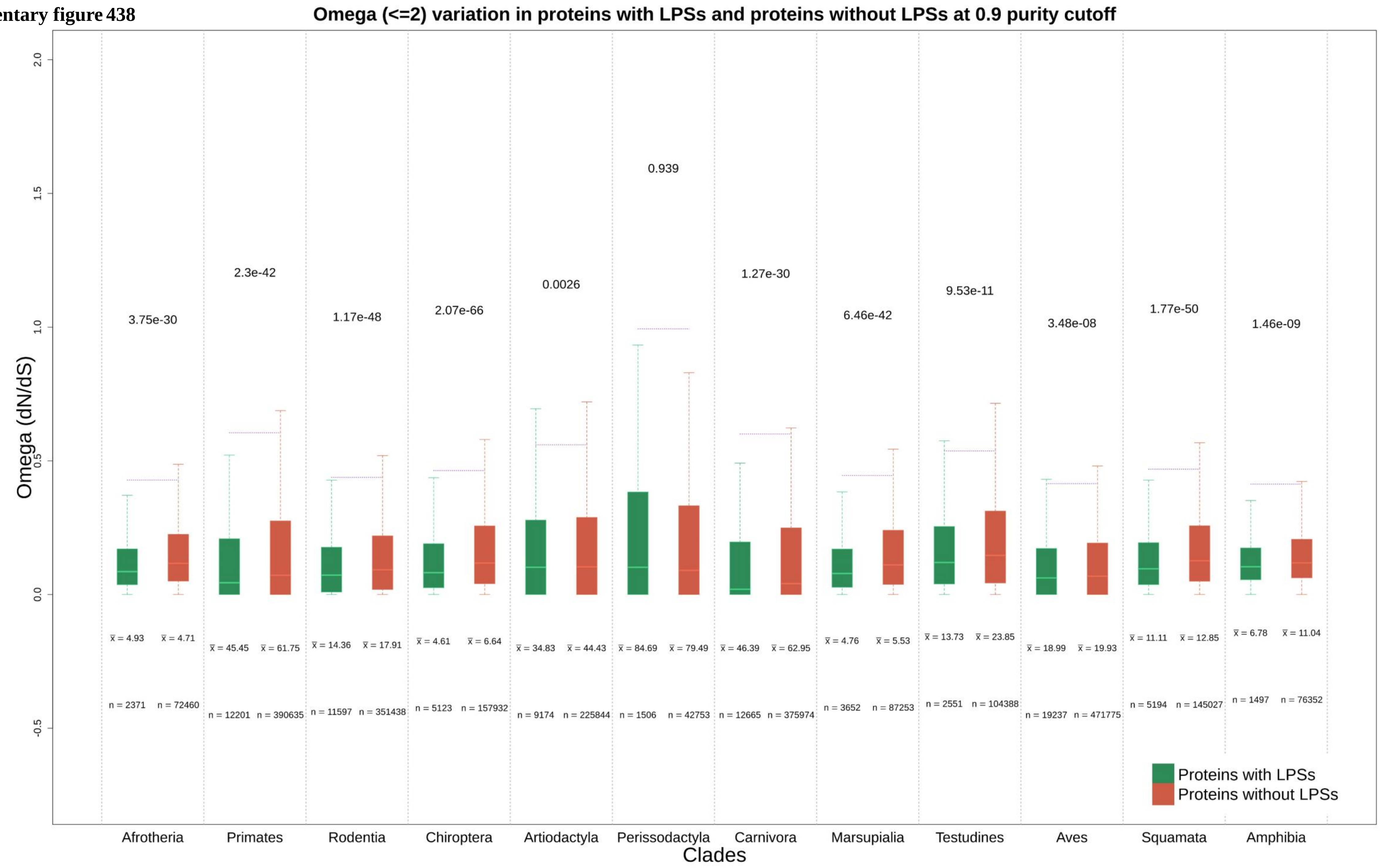

Supplementary figure 439

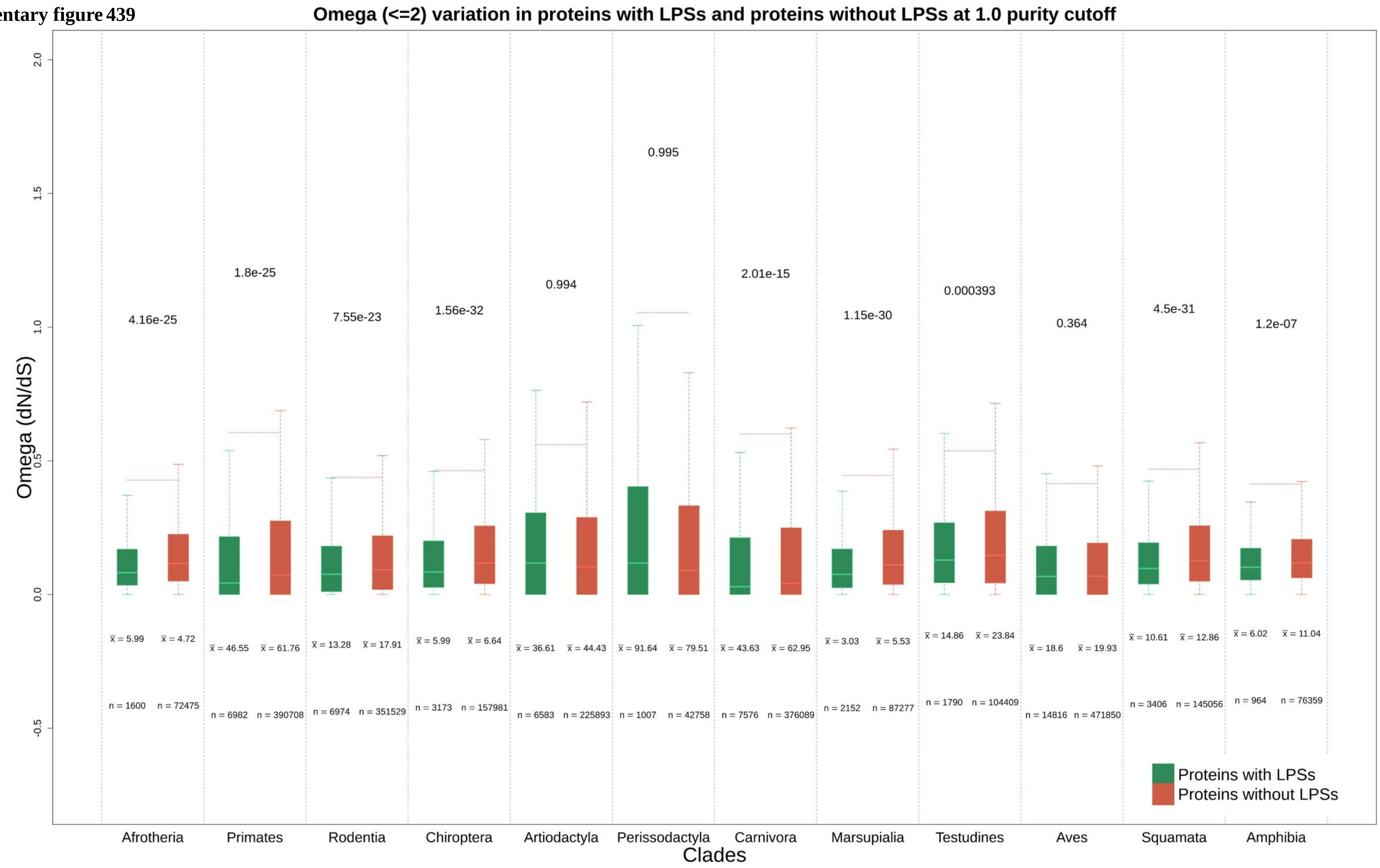

**Supplementary figure 440**

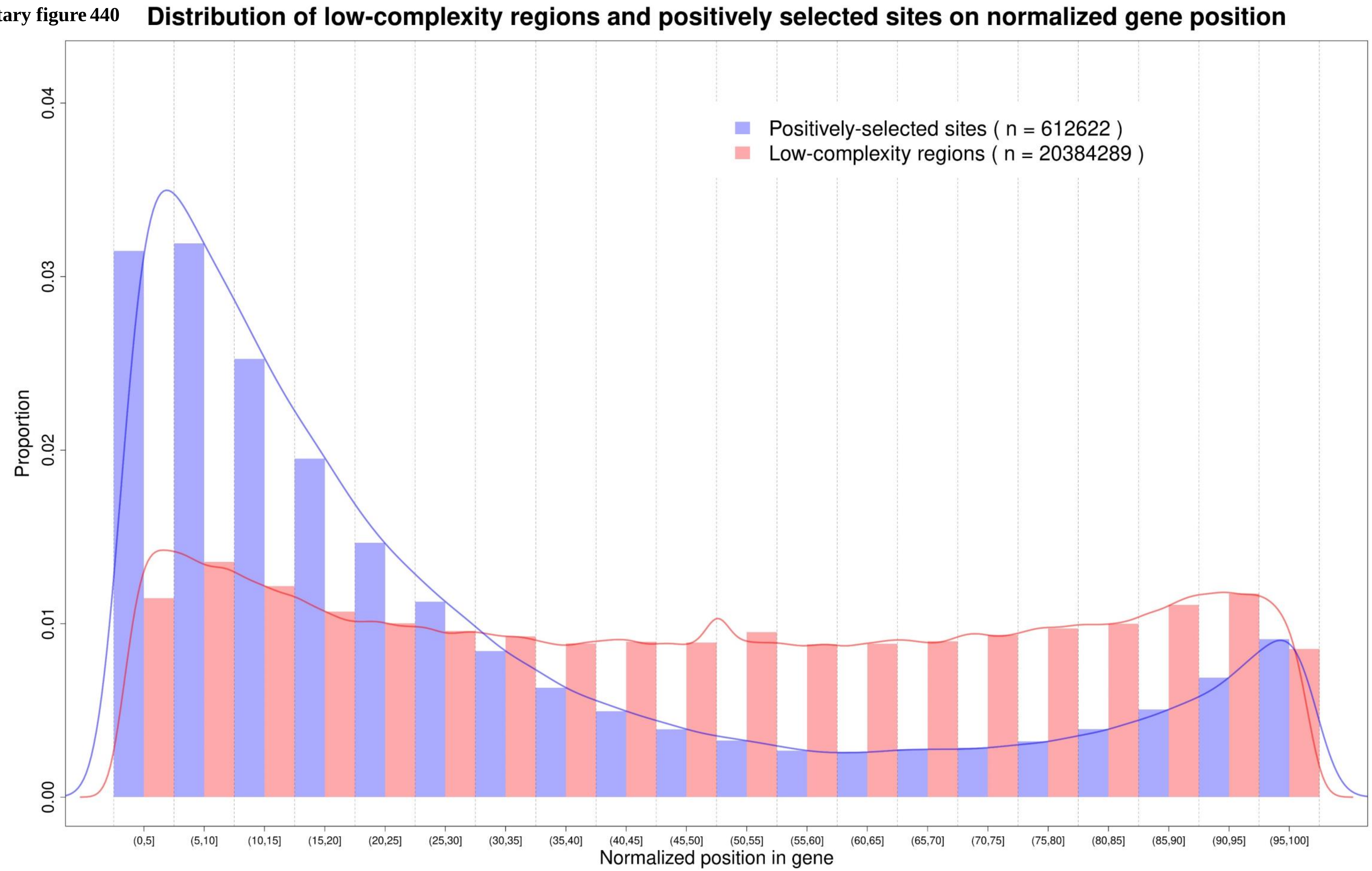

Supplementary figure 441

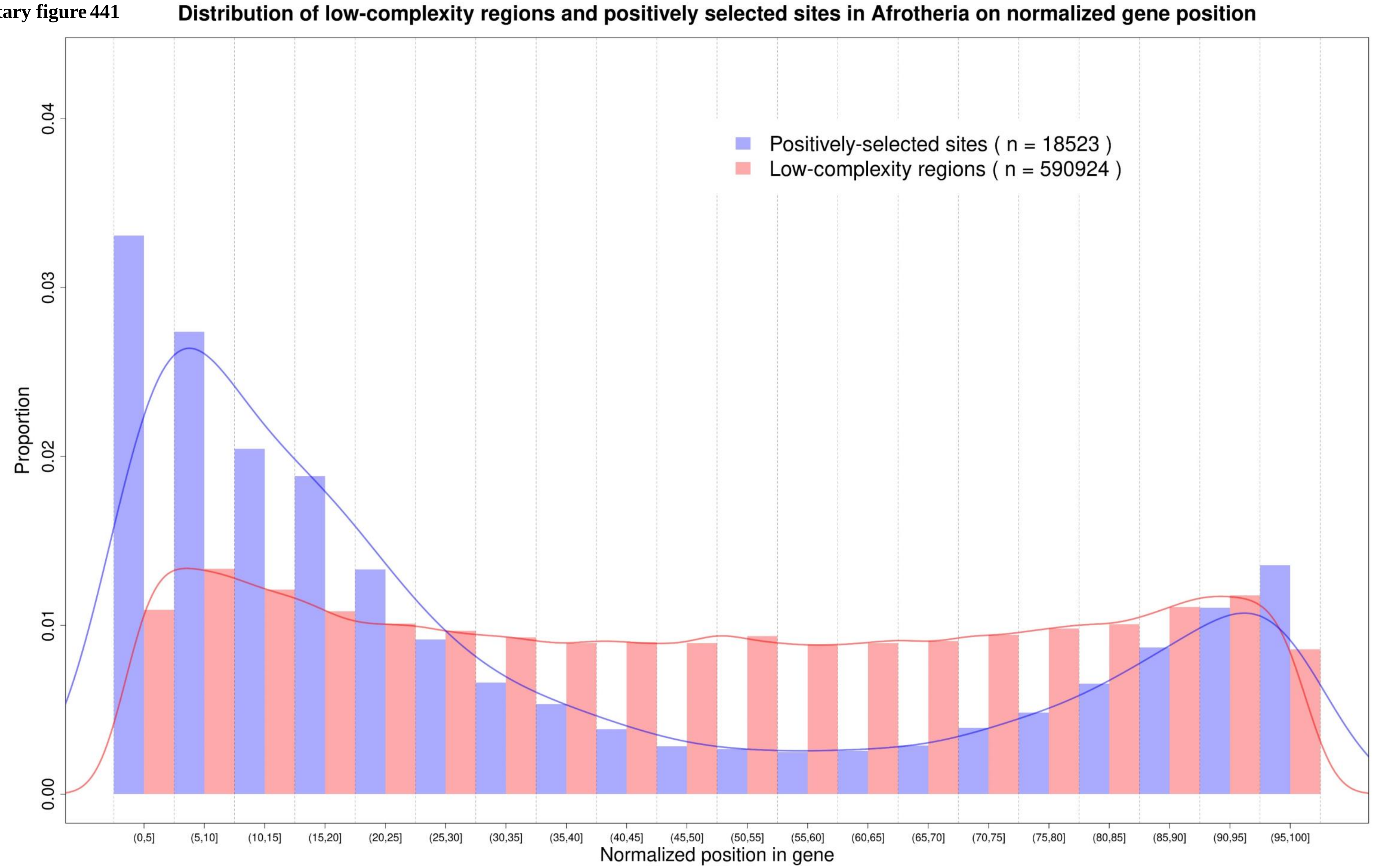

**Supplementary figure 442**

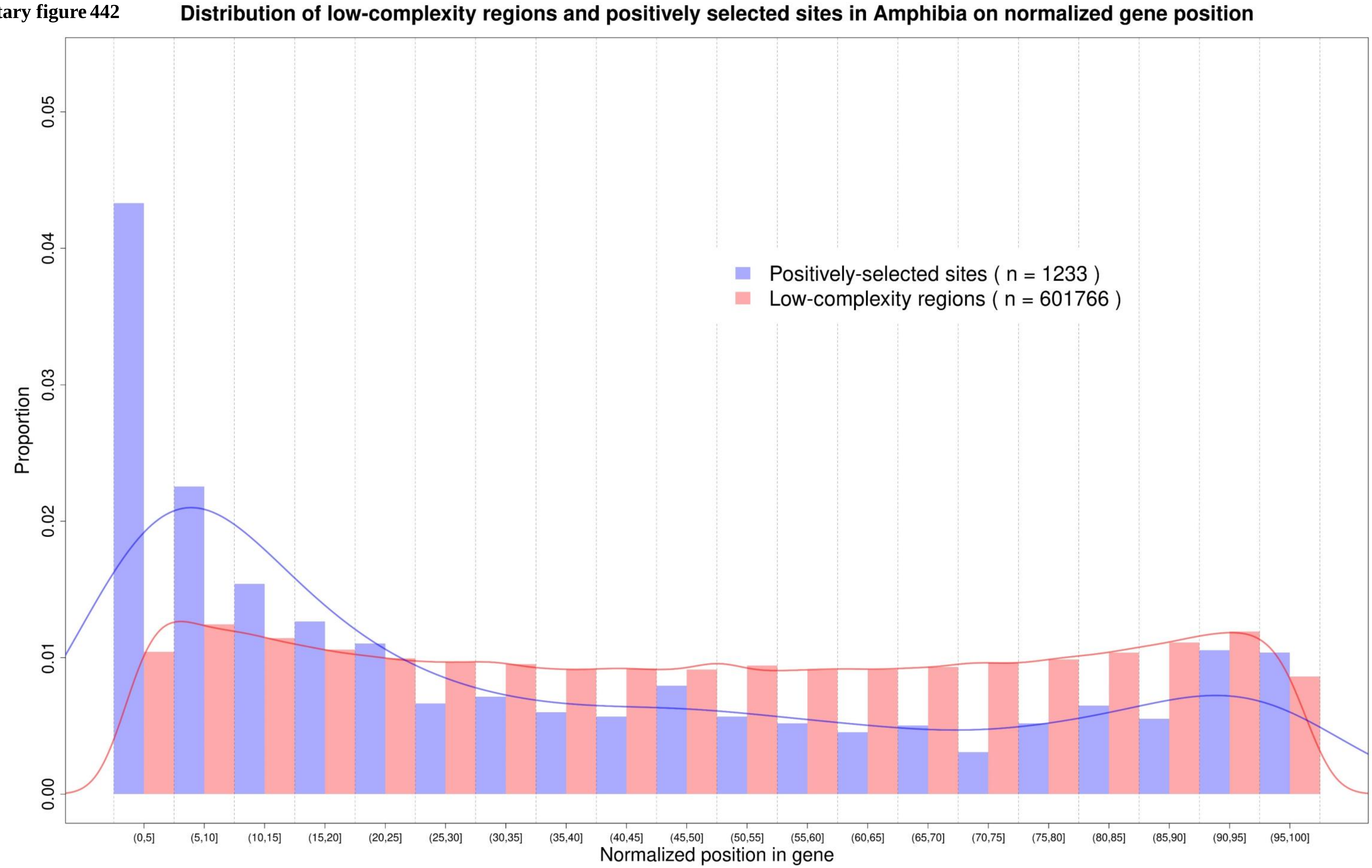

**Supplementary figure 443** **Distribution of low-complexity regions and positively selected sites in Artiodactyla on normalized gene position**

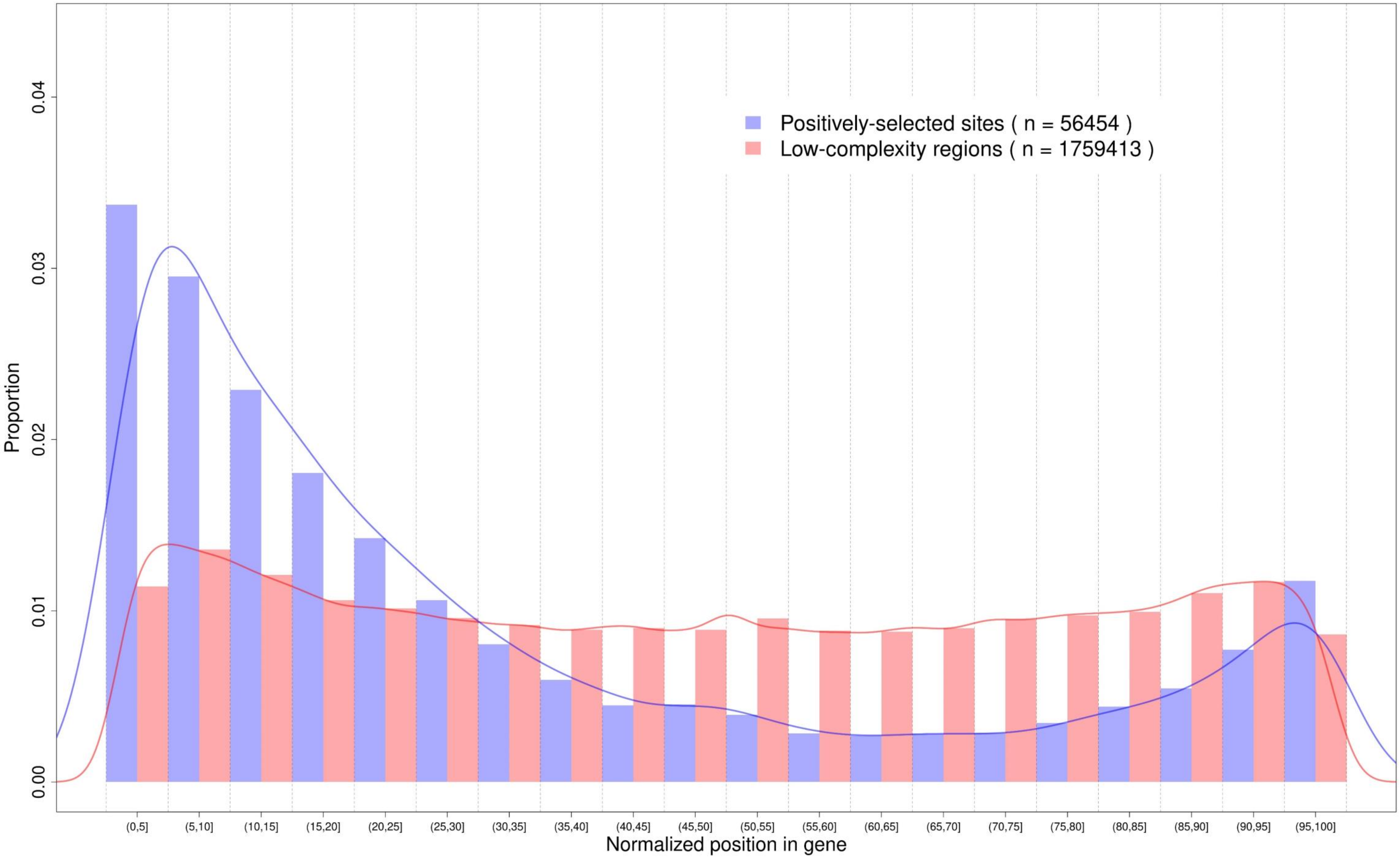

**Supplementary figure 444**

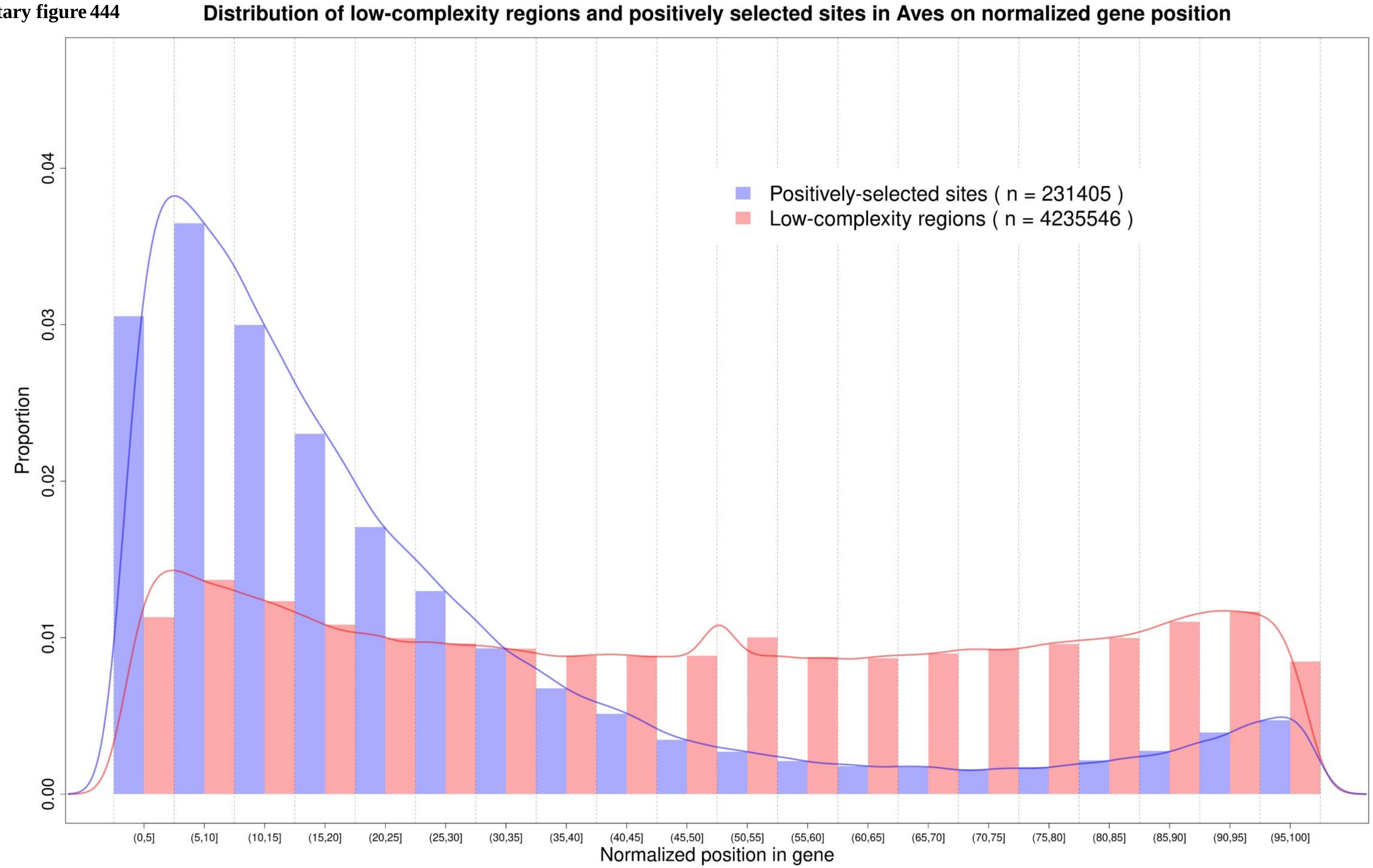

Supplementary figure 445

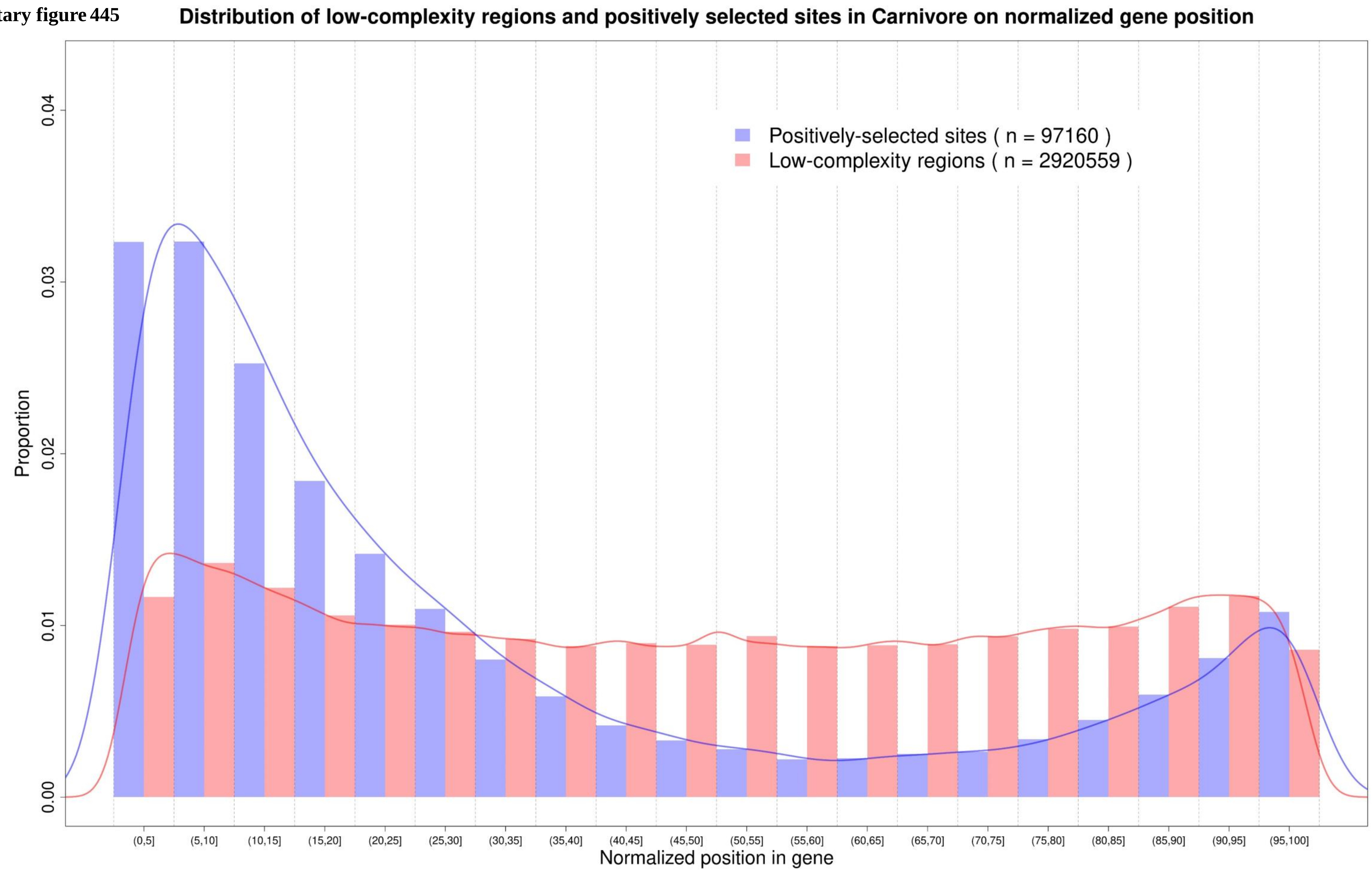

**Supplementary figure 446**

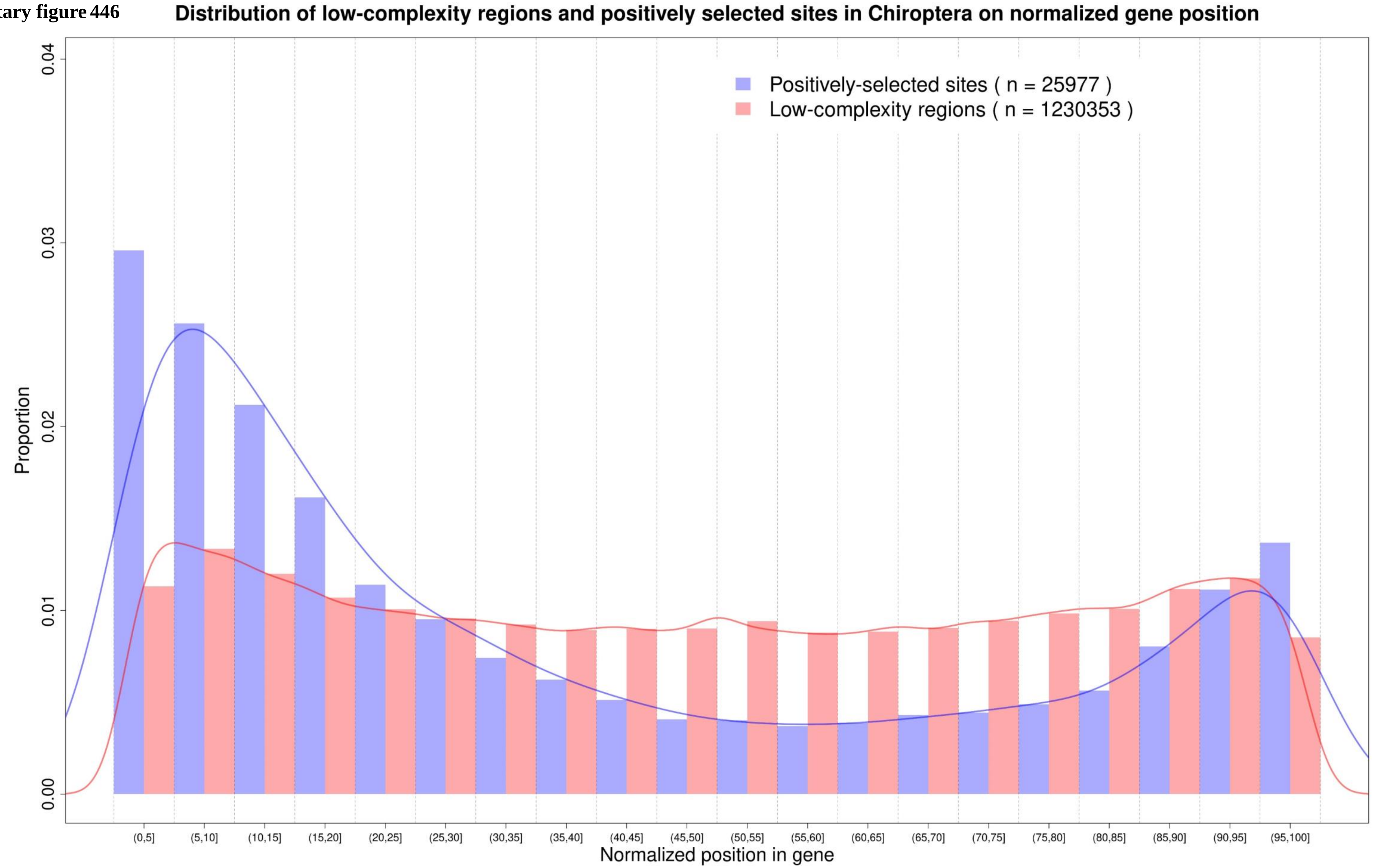

**Supplementary figure 447**

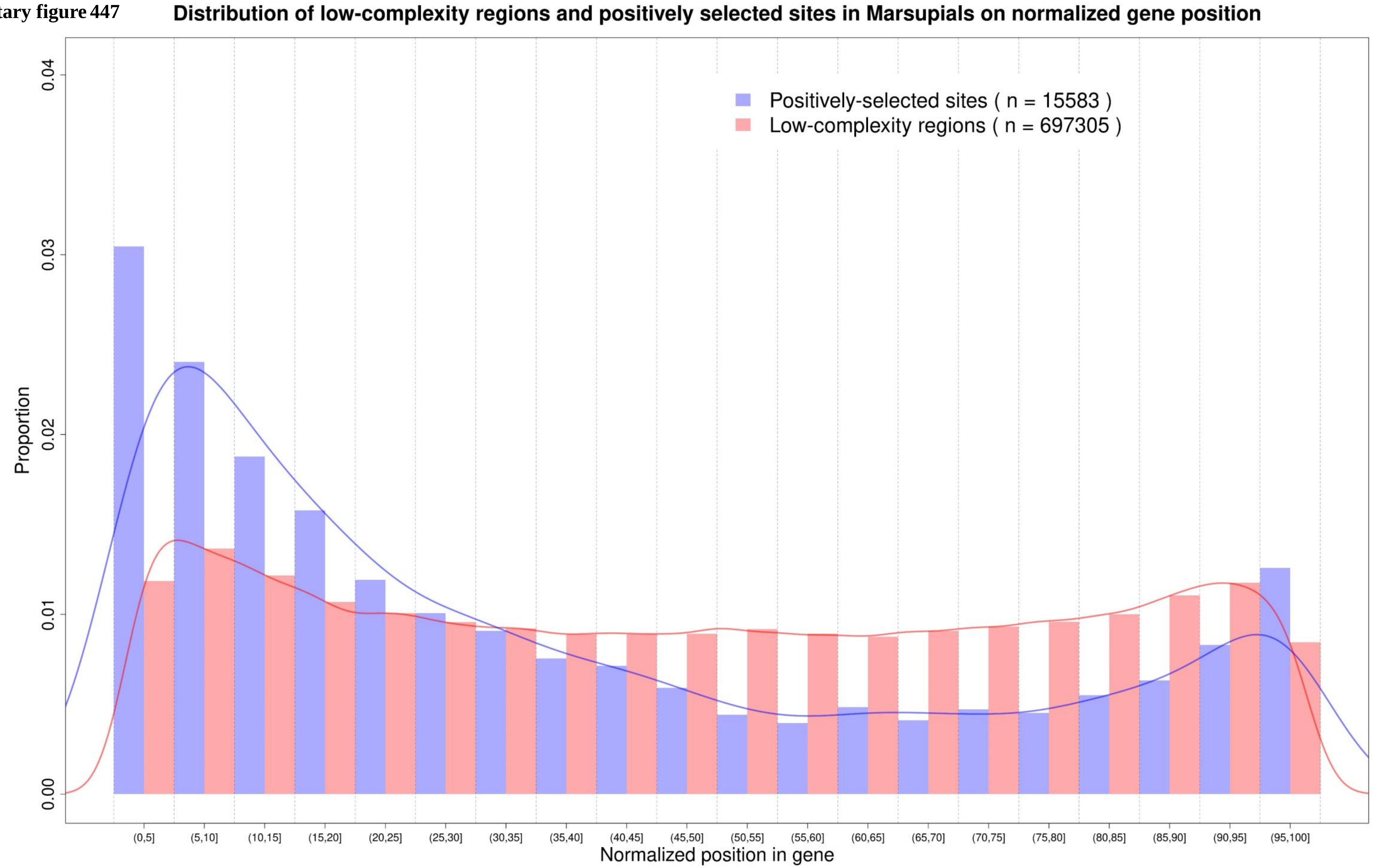

Supplementary figure 448

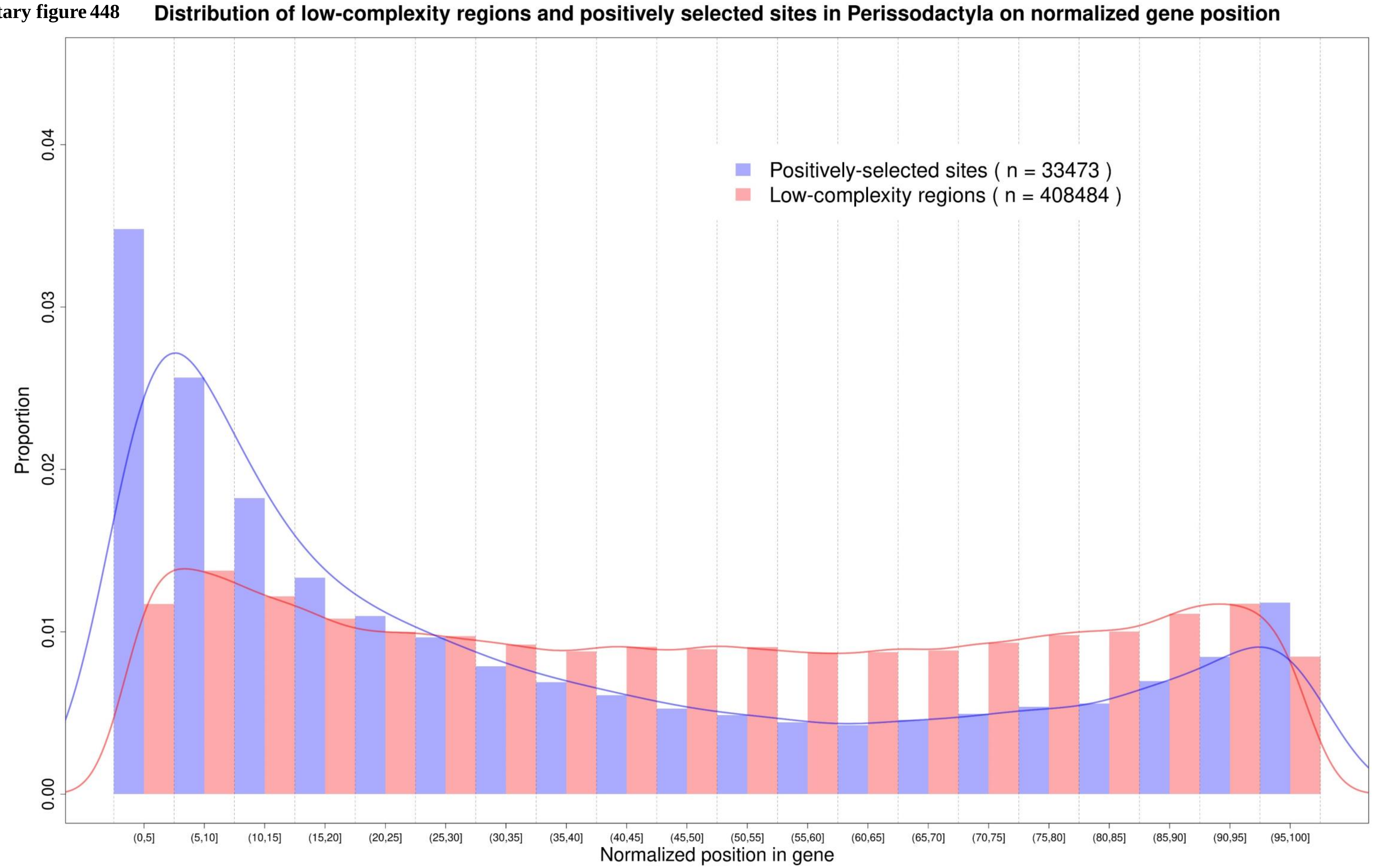

Supplementary figure 449

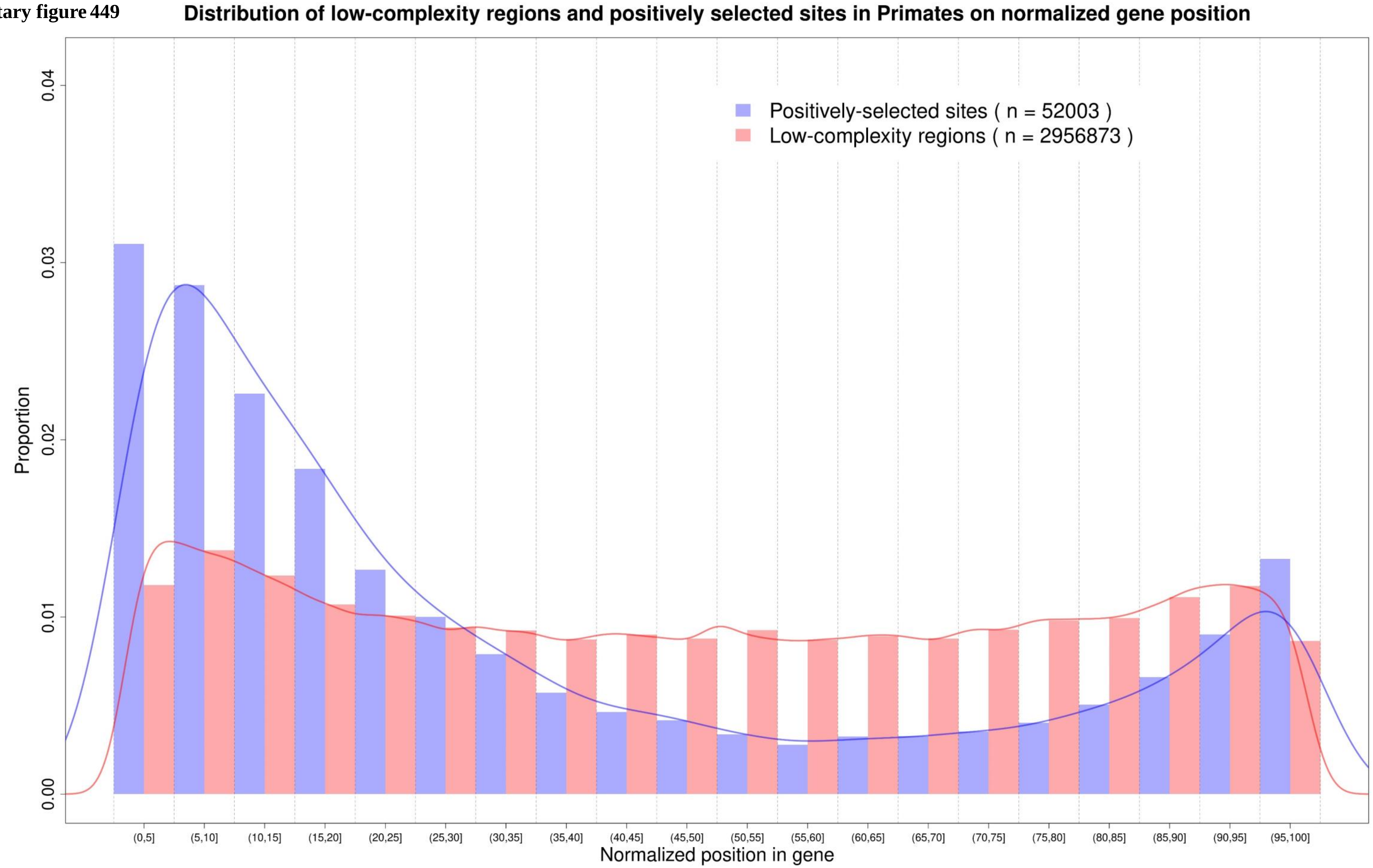

**Supplementary figure 450**

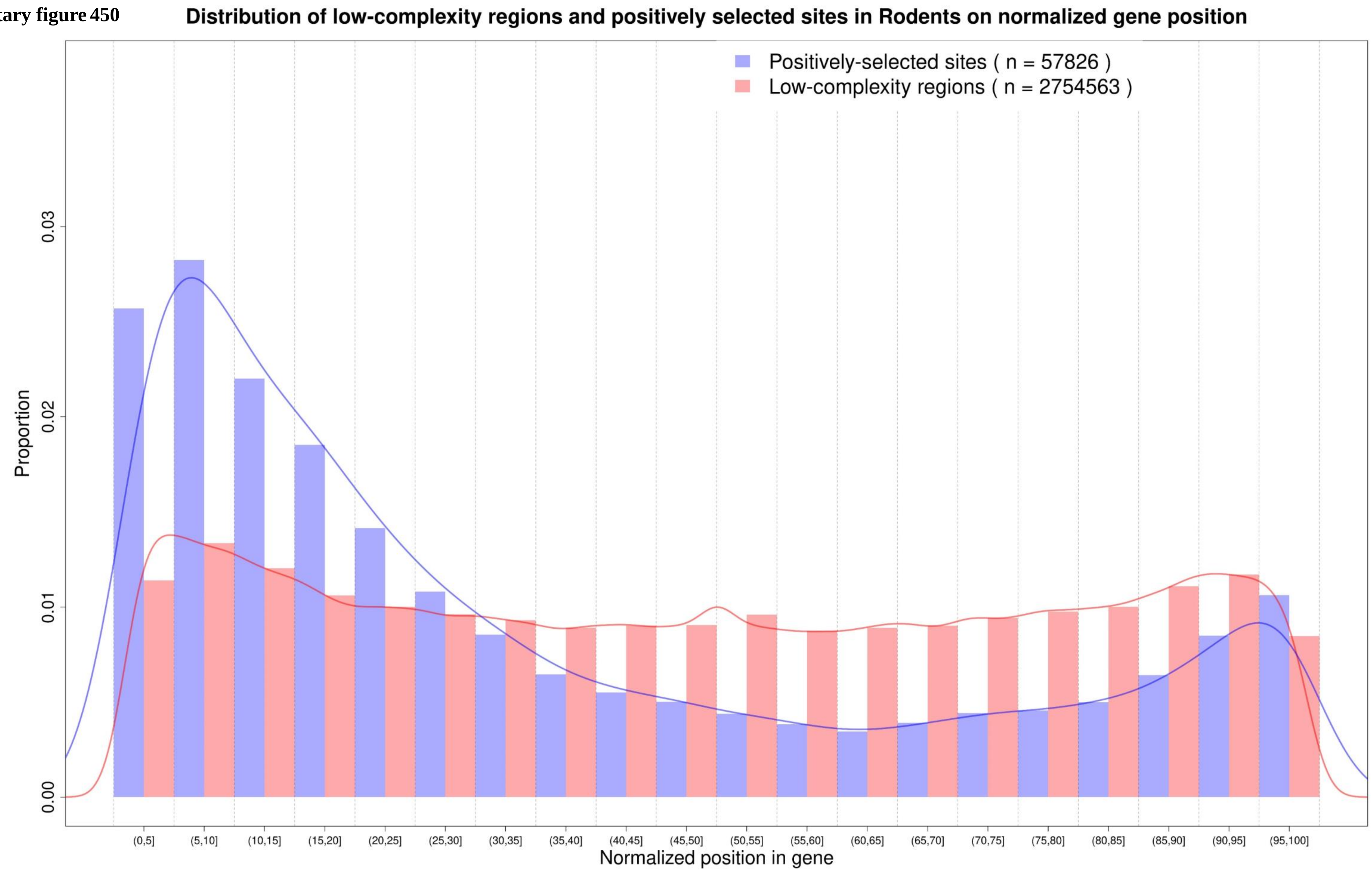

**Supplementary figure 451**

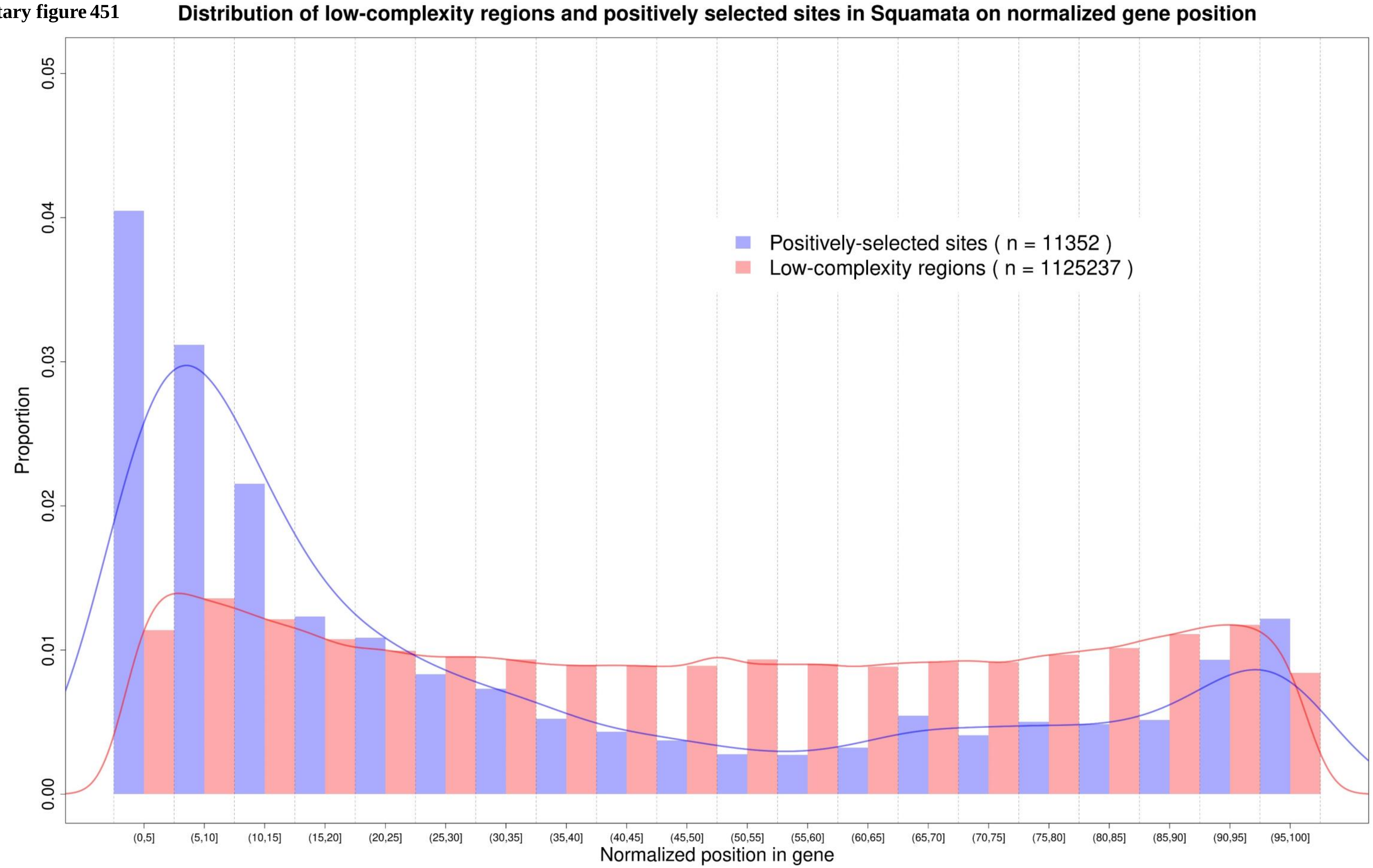

Supplementary figure 452

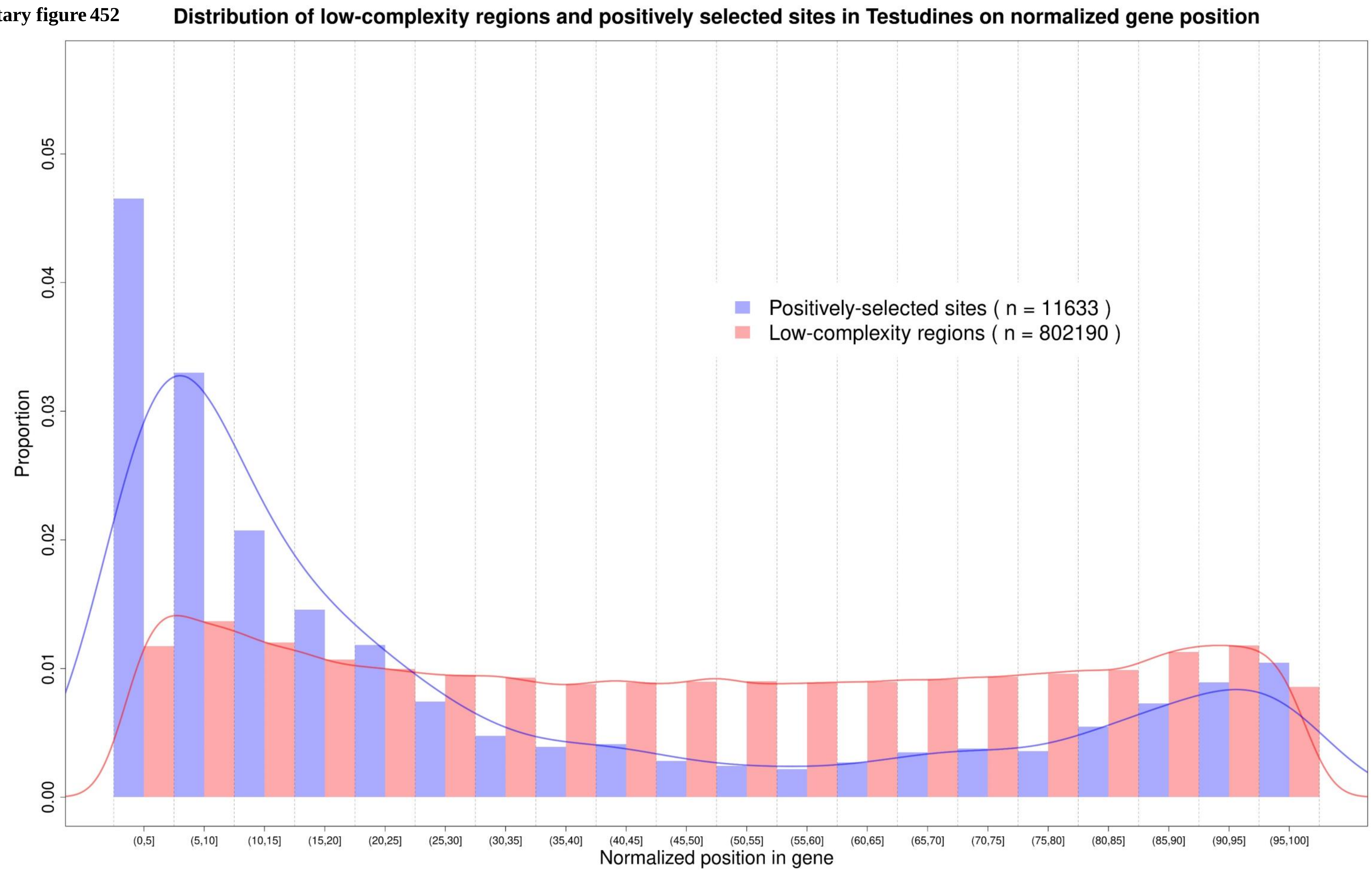

Supplementary figure 453

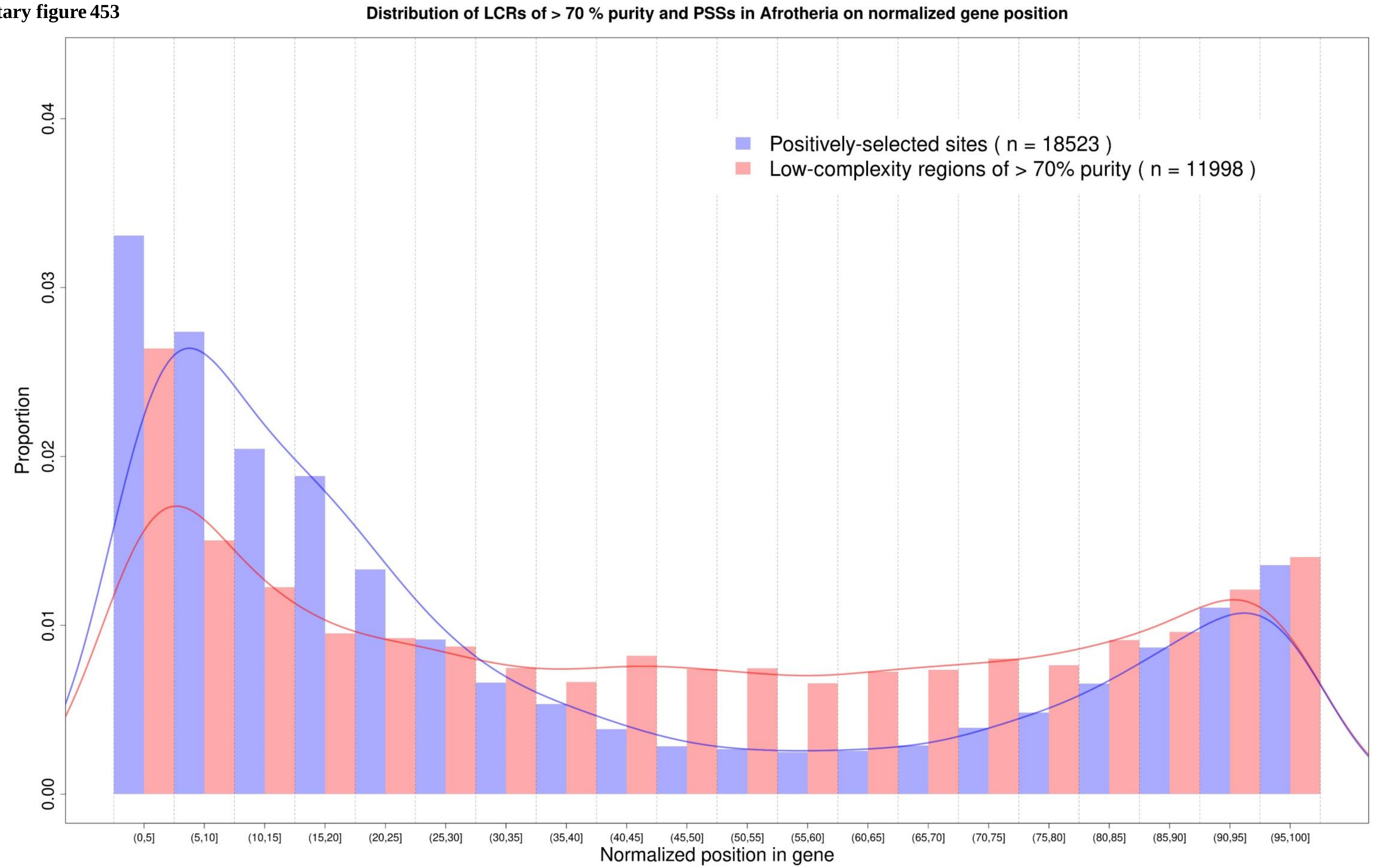

**Supplementary figure 454**

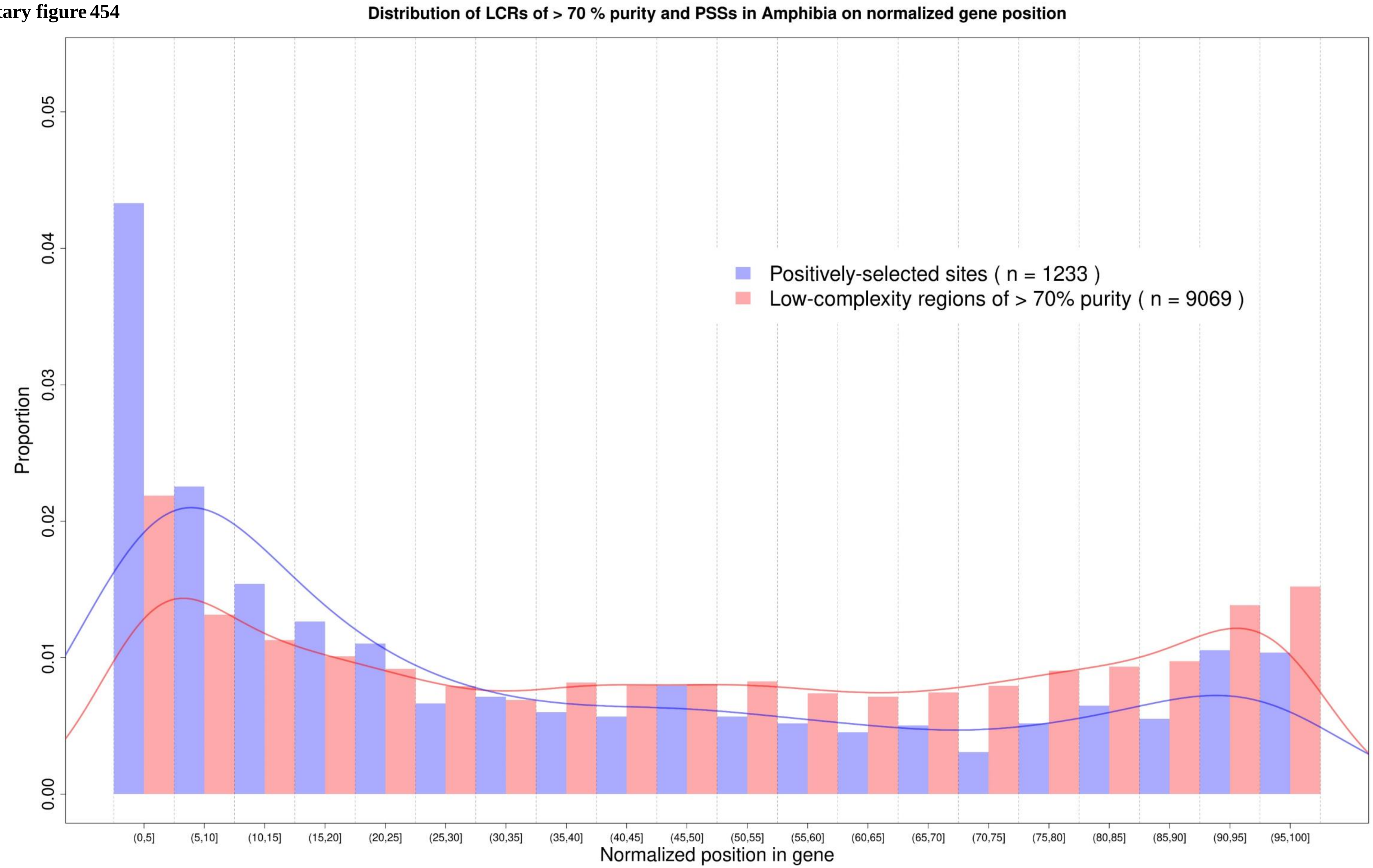

**Supplementary figure 455**

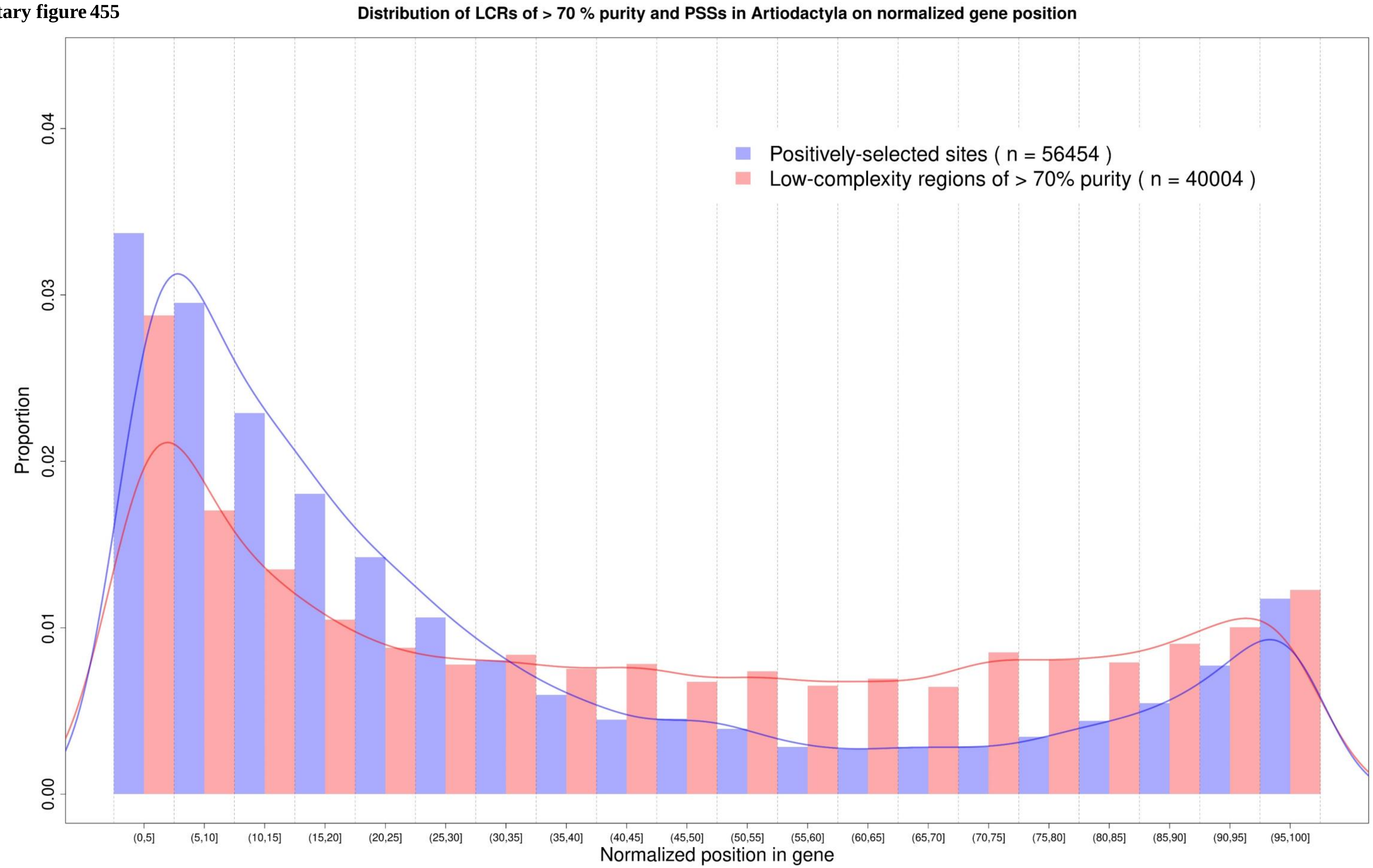

Supplementary figure 456

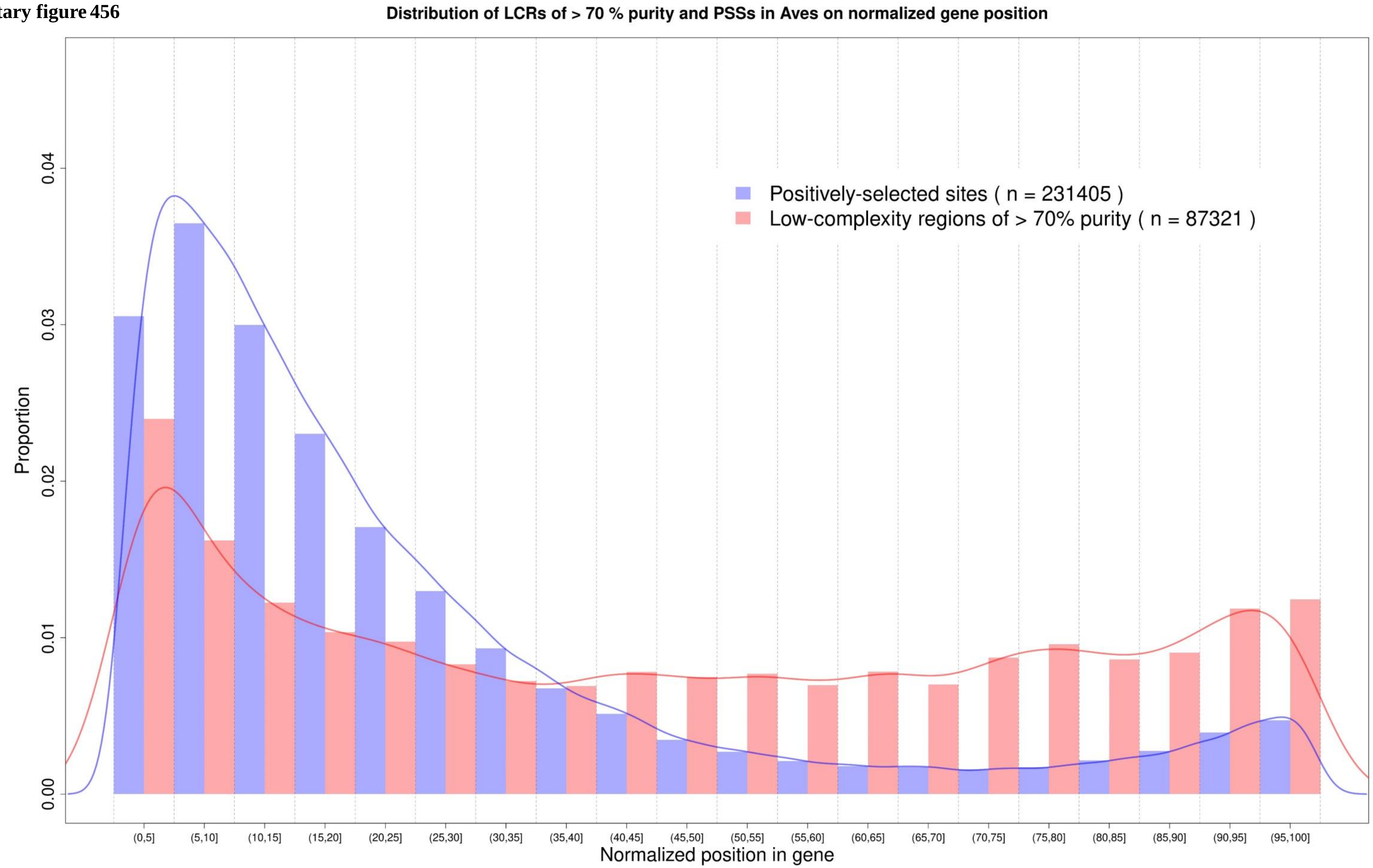

**Supplementary figure 457**

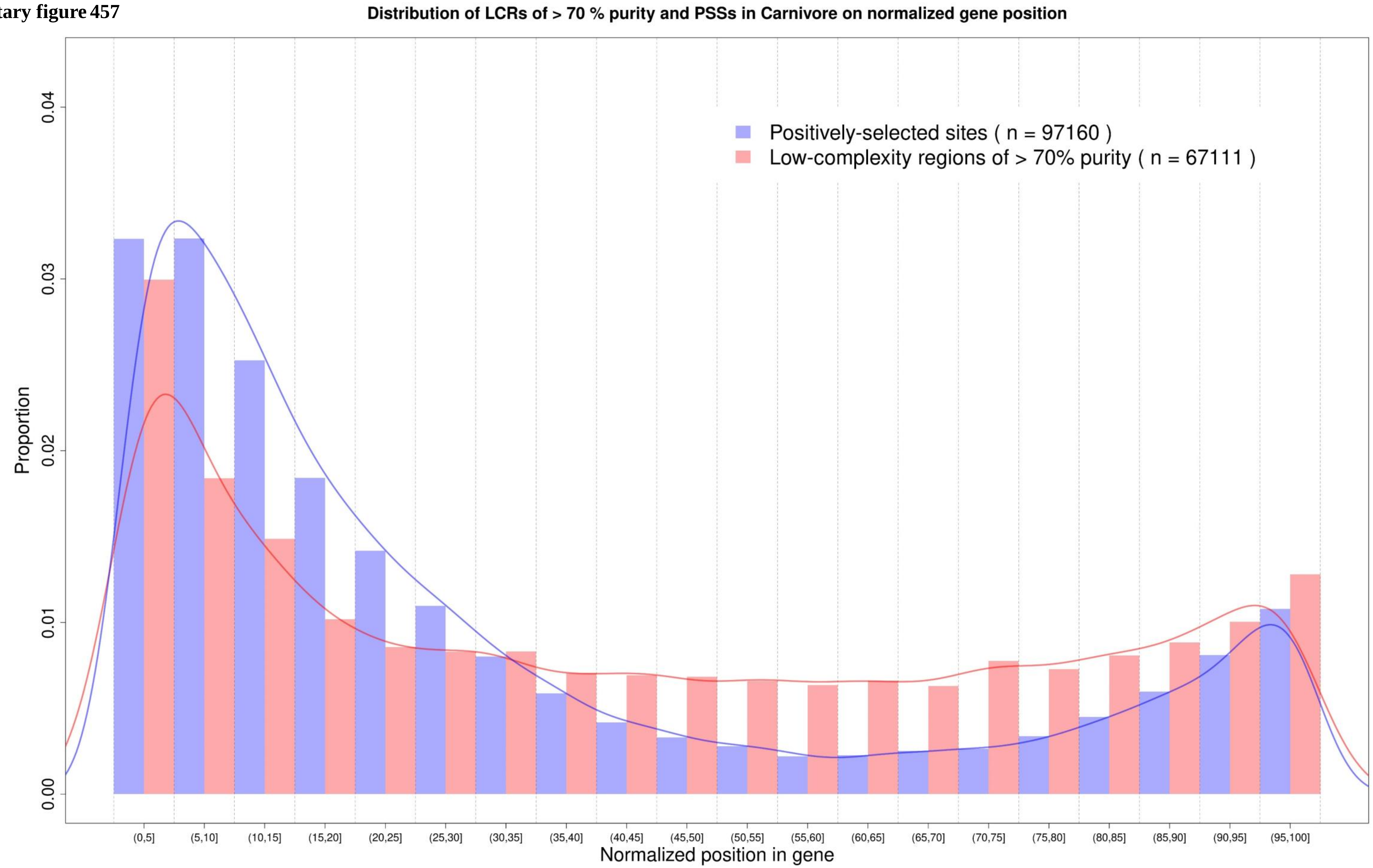

**Supplementary figure 458**

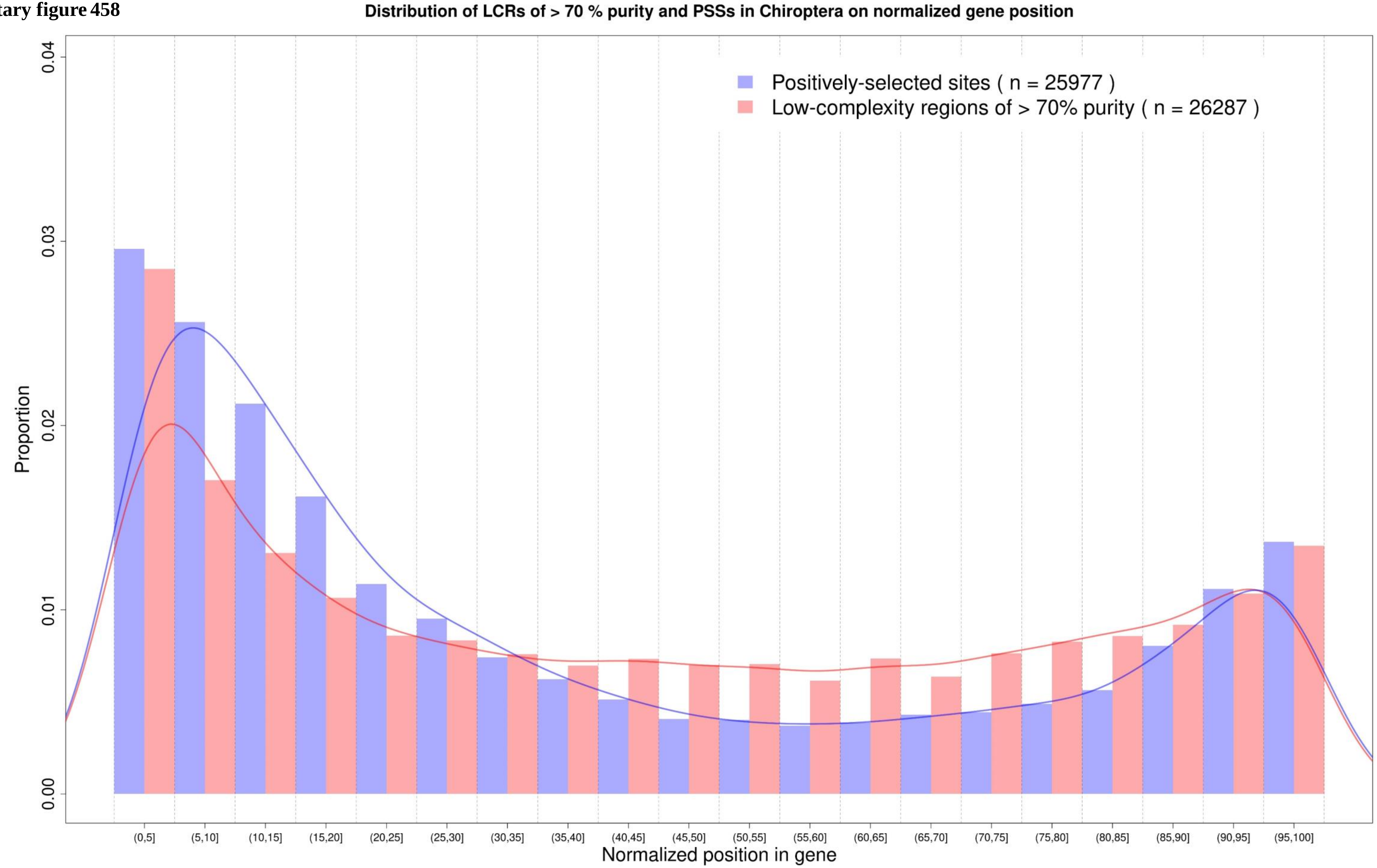

**Supplementary figure 459**

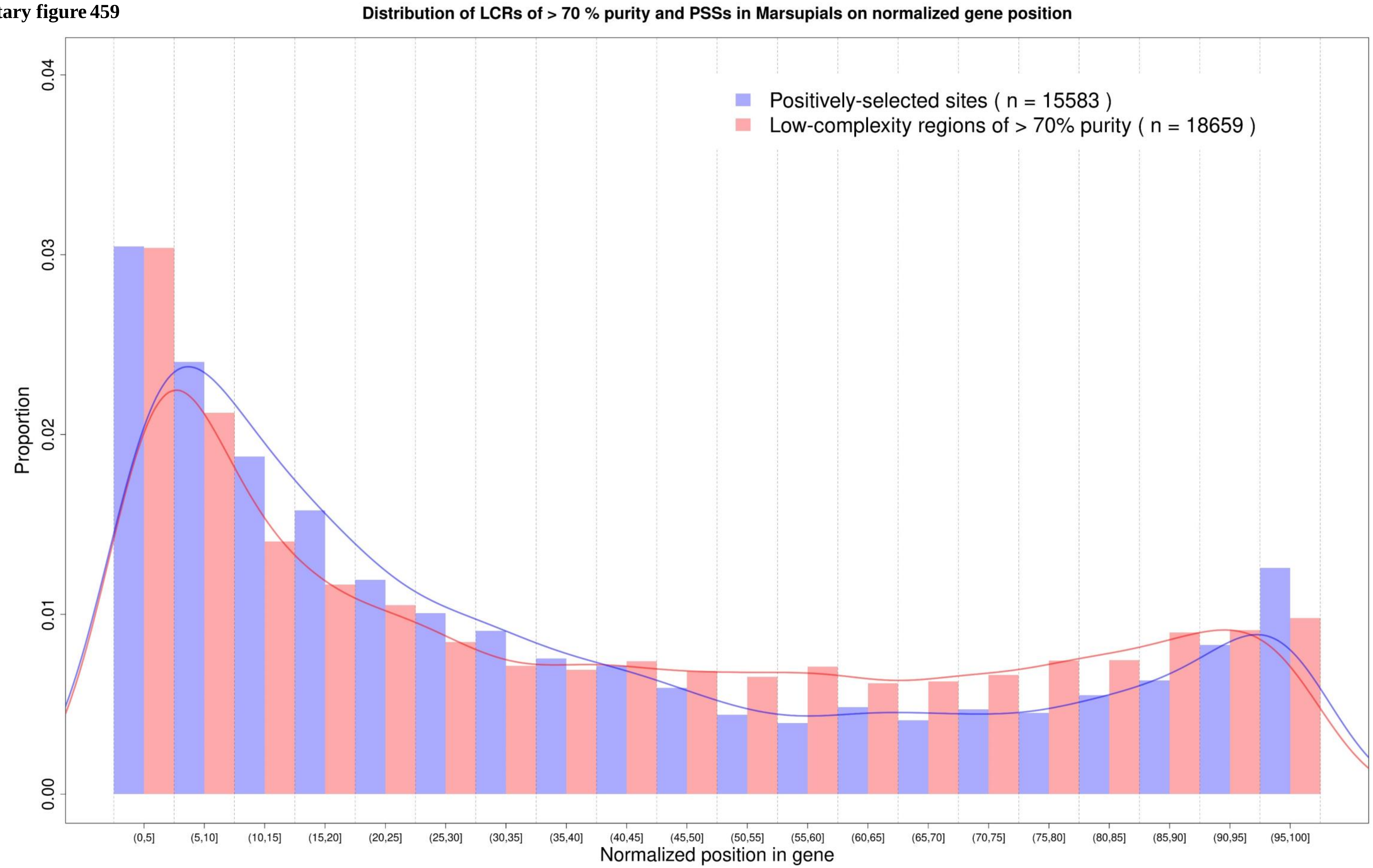

**Supplementary figure 460**

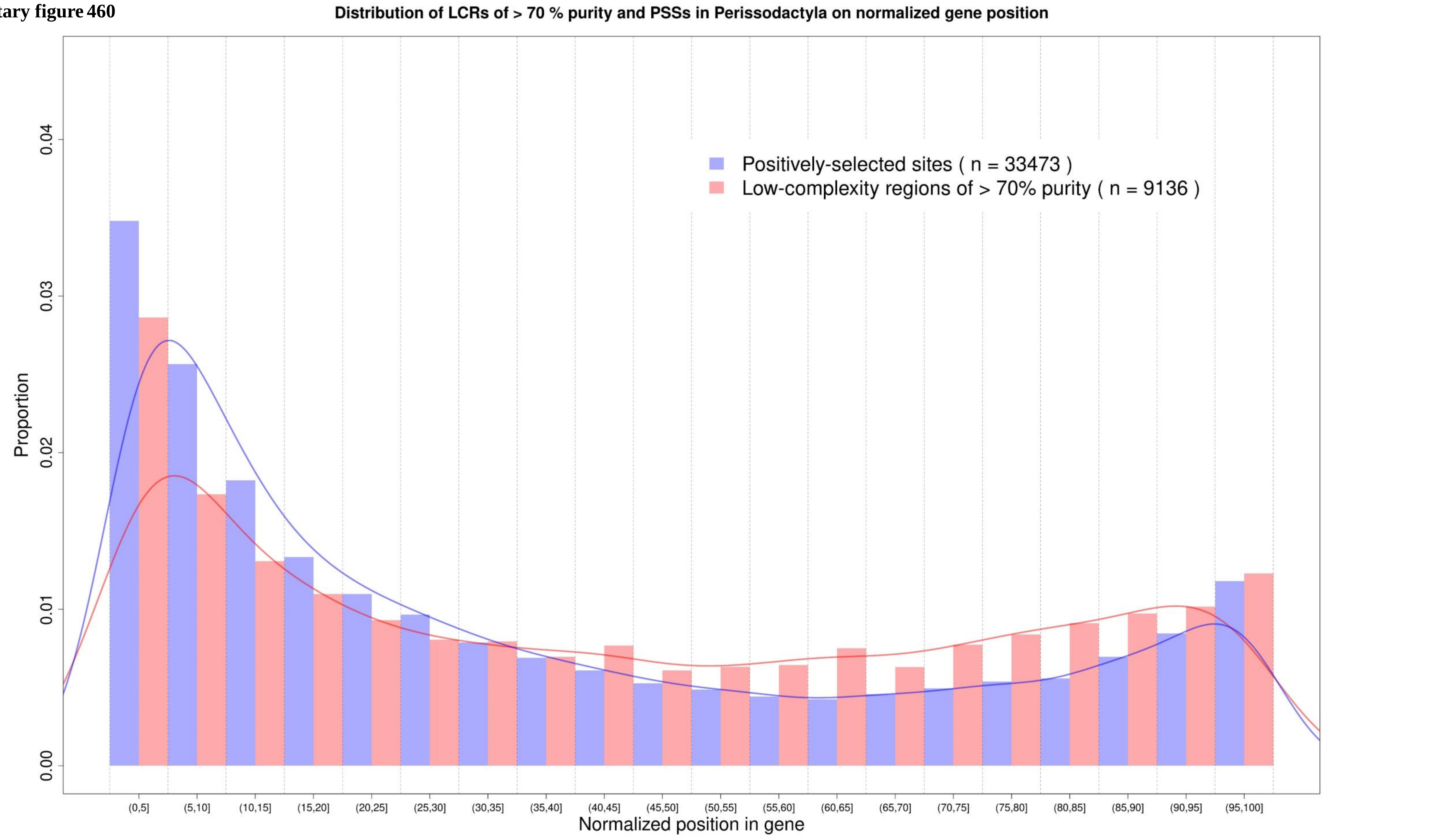

**Supplementary figure 461**

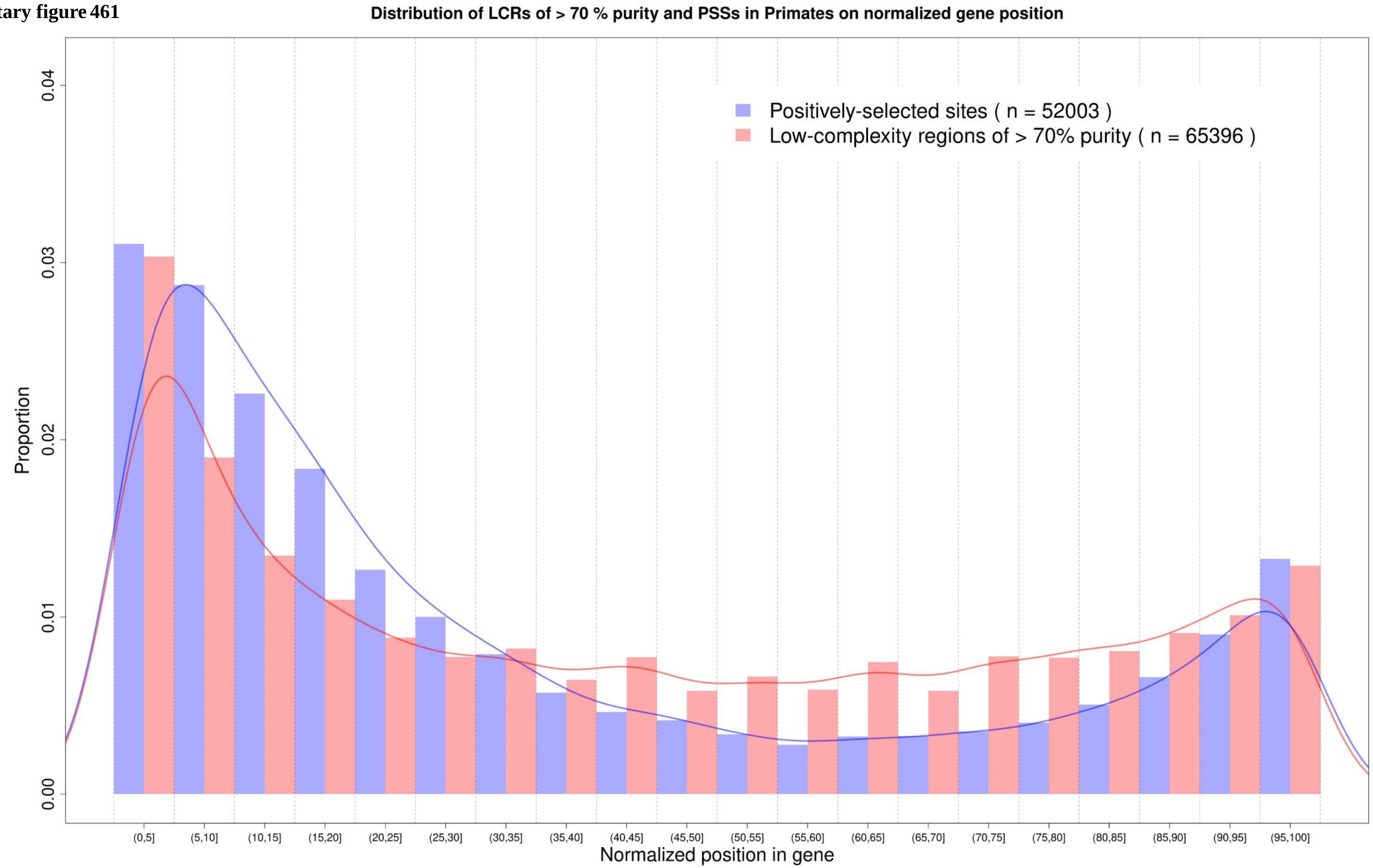

**Supplementary figure 462**

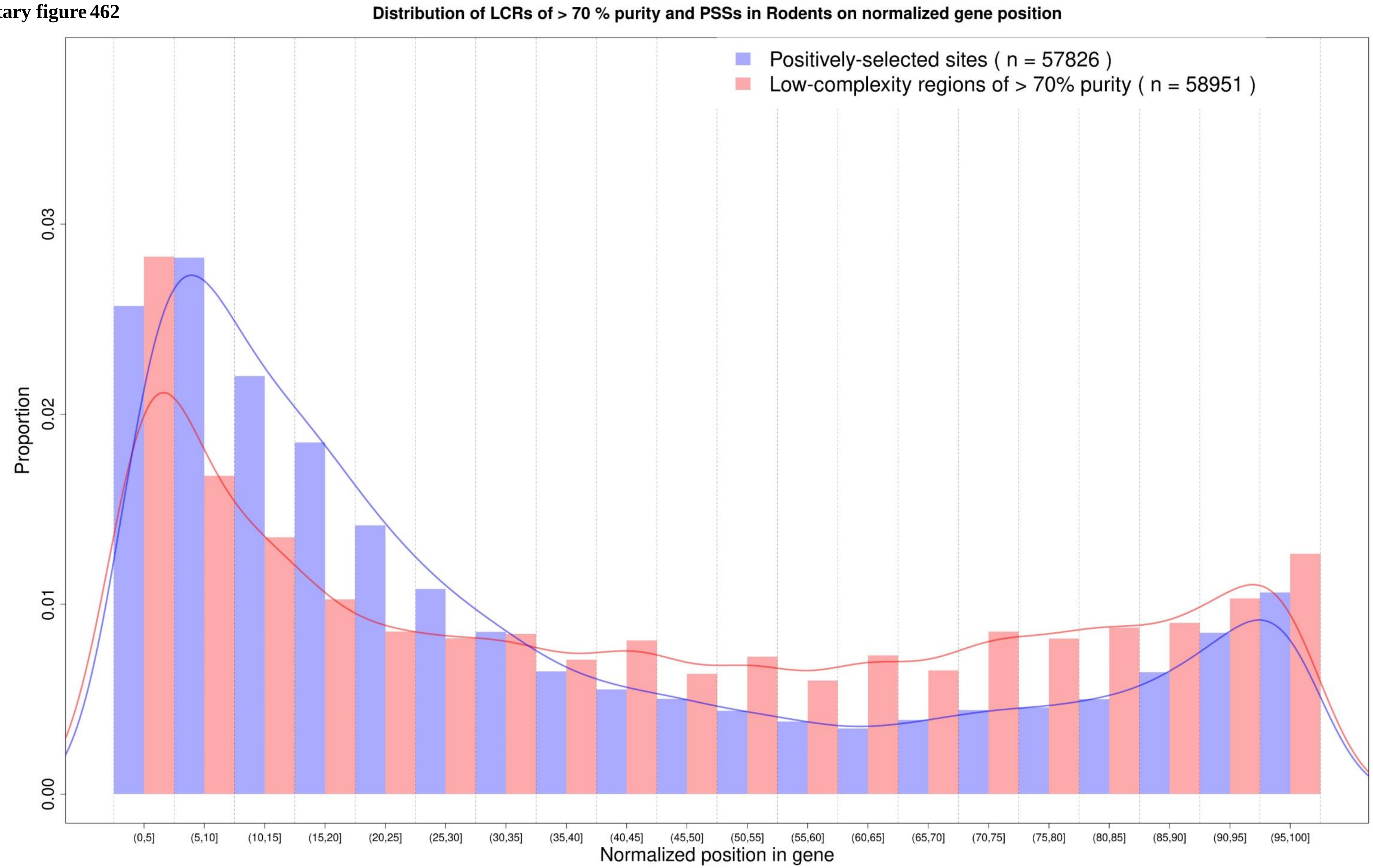

**Supplementary figure 463**

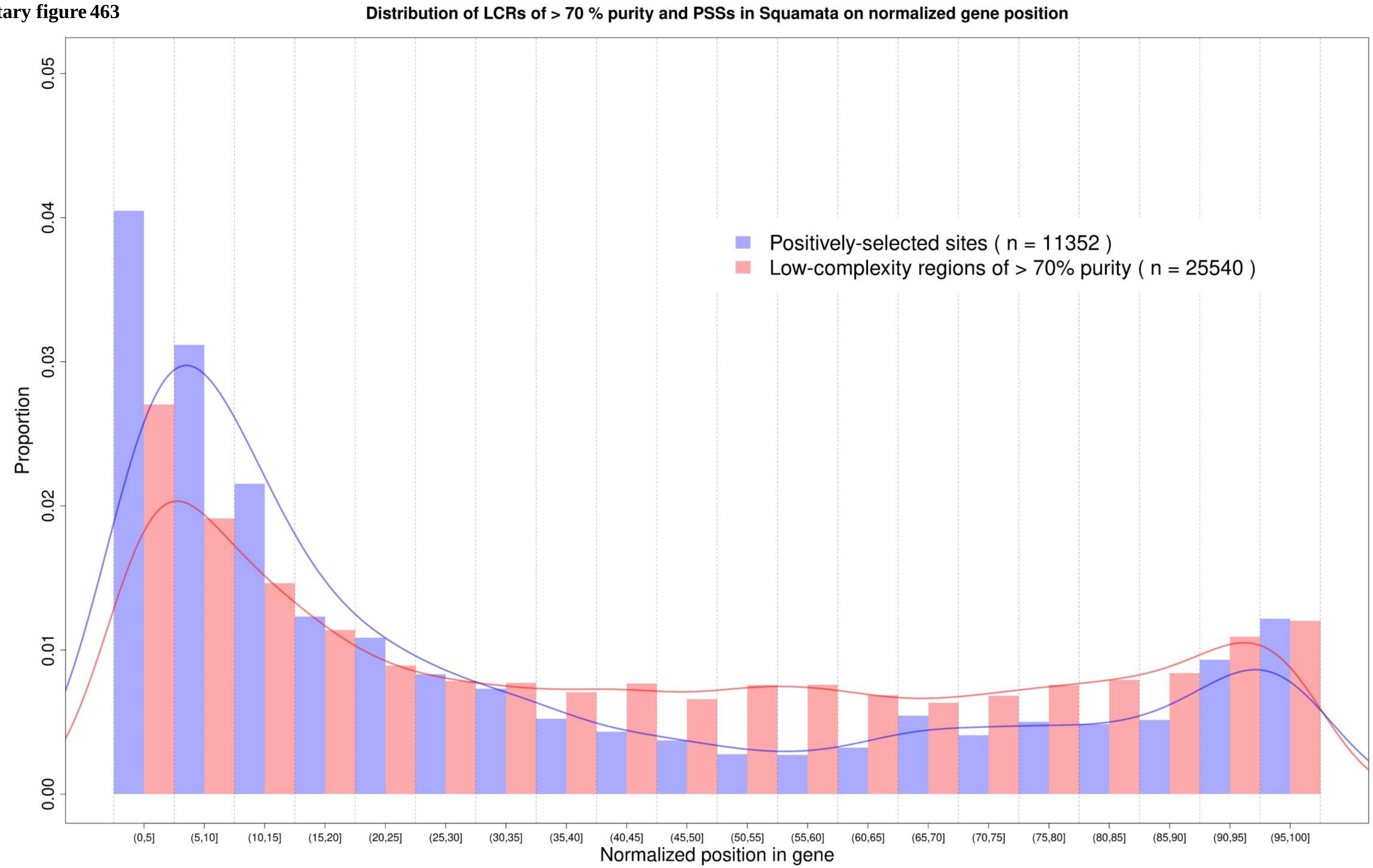

**Supplementary figure 464**

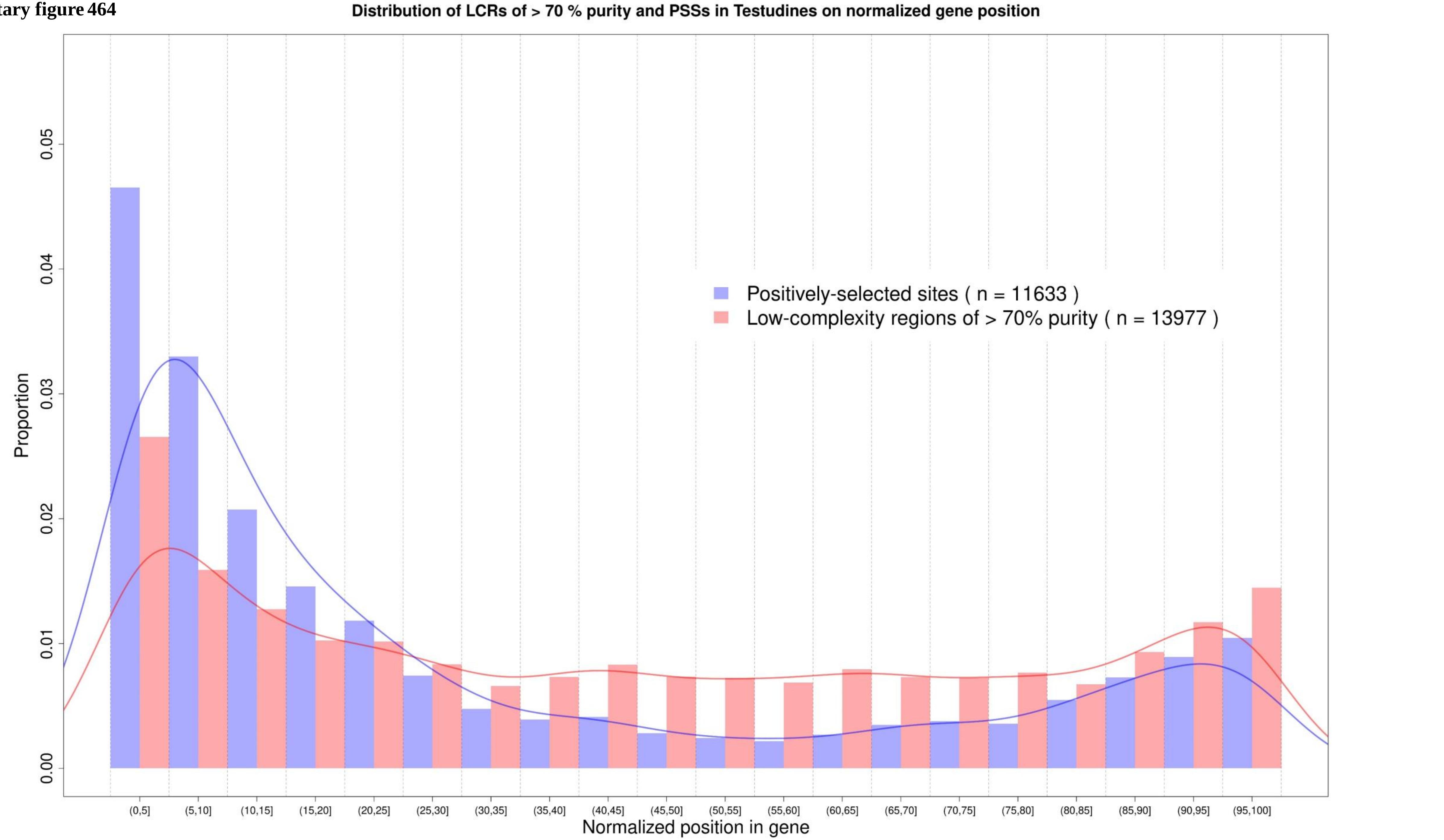

**Supplementary figure 465**

### Distribution of exact LCRs and PSSs in a normalized gene position in all clades

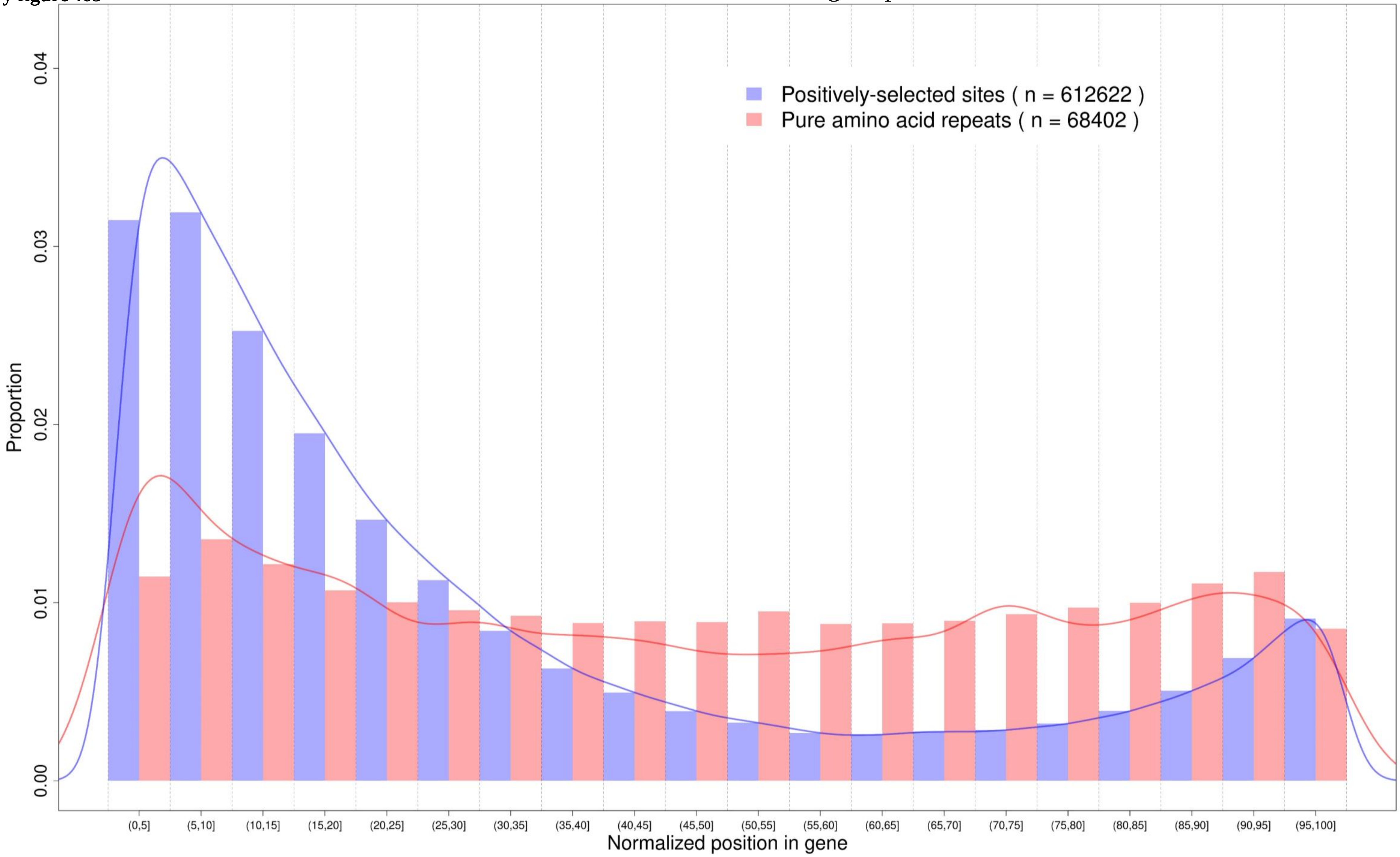

**Supplementary figure 466**

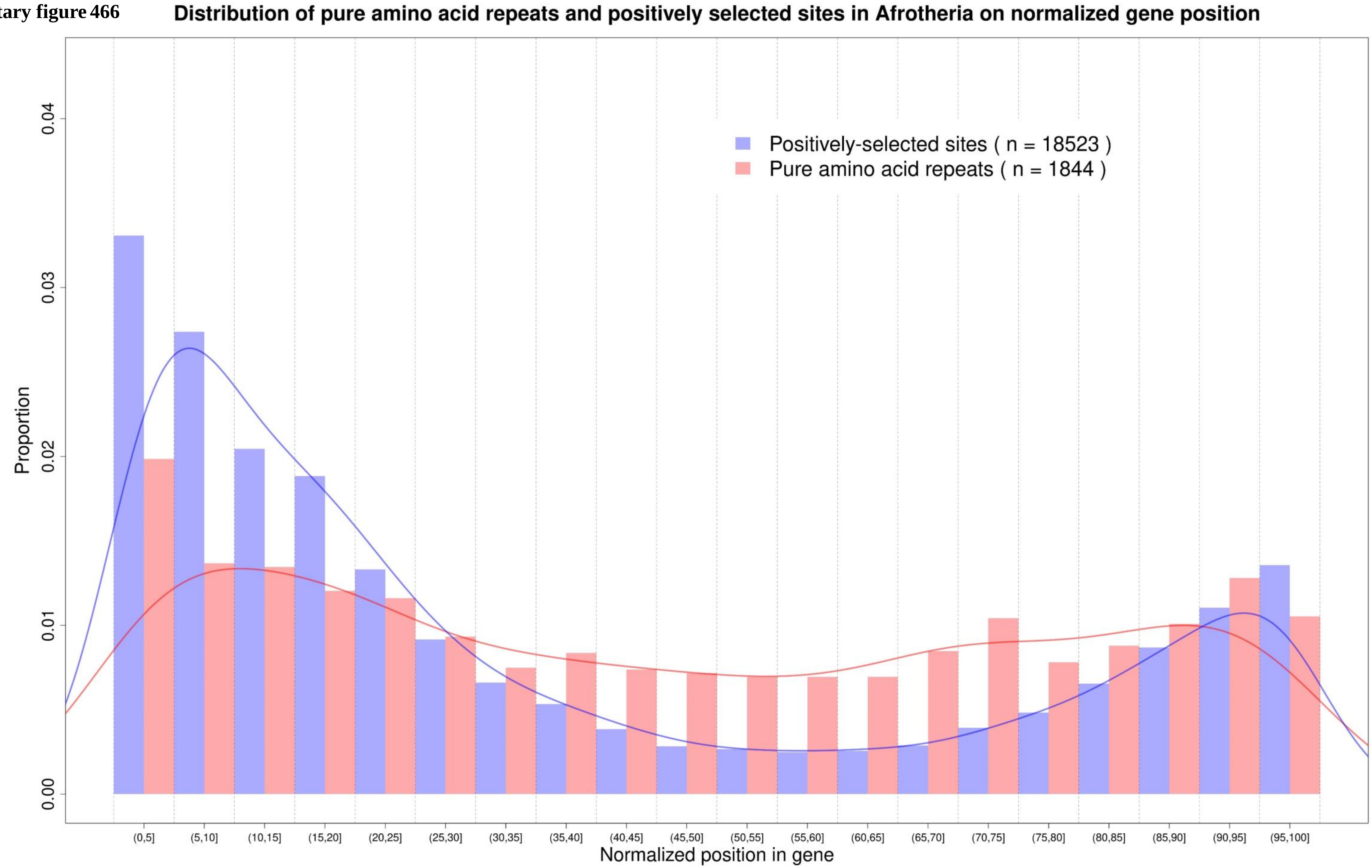

**Supplementary figure 467**

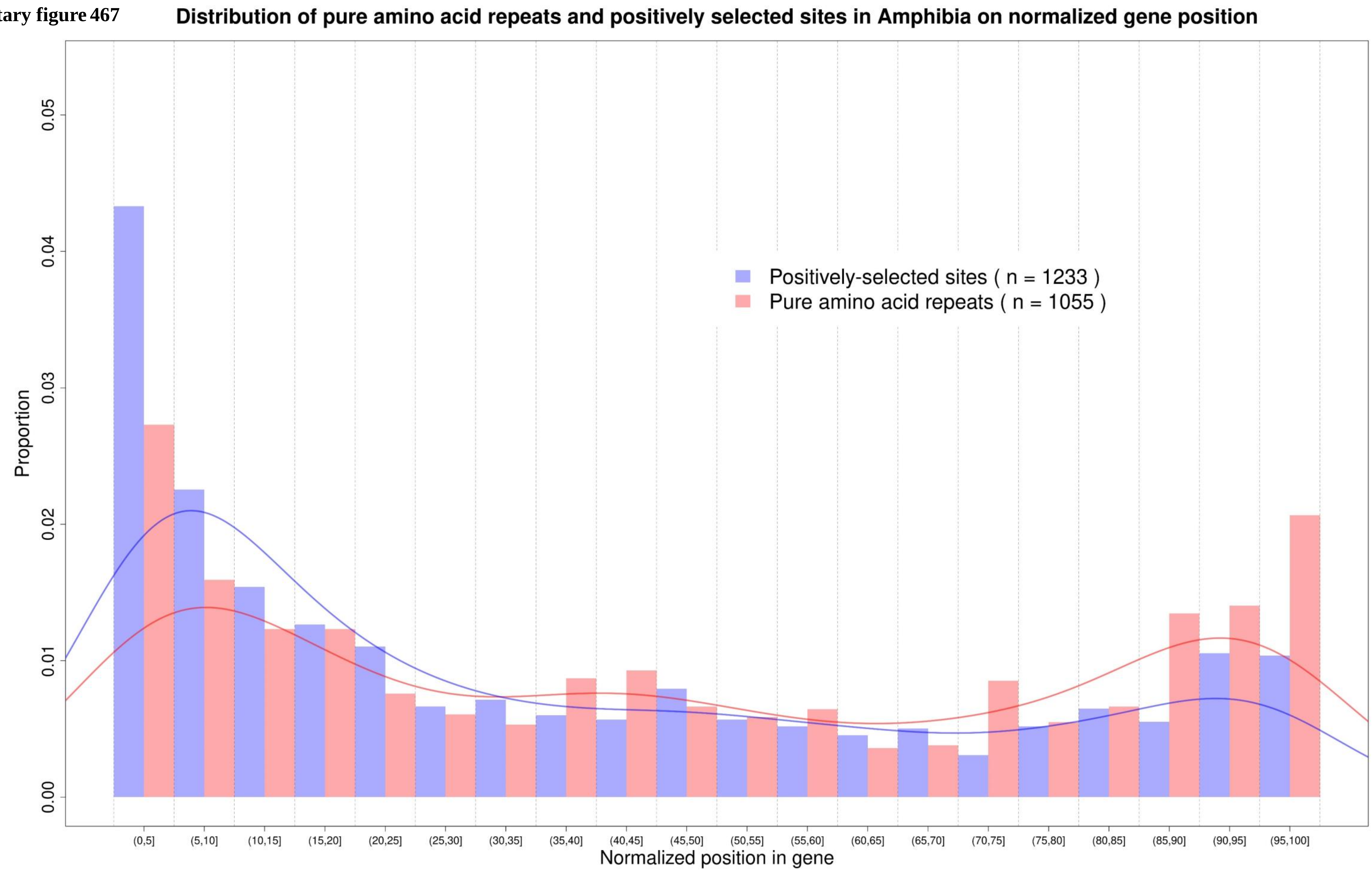

Supplementary figure 468

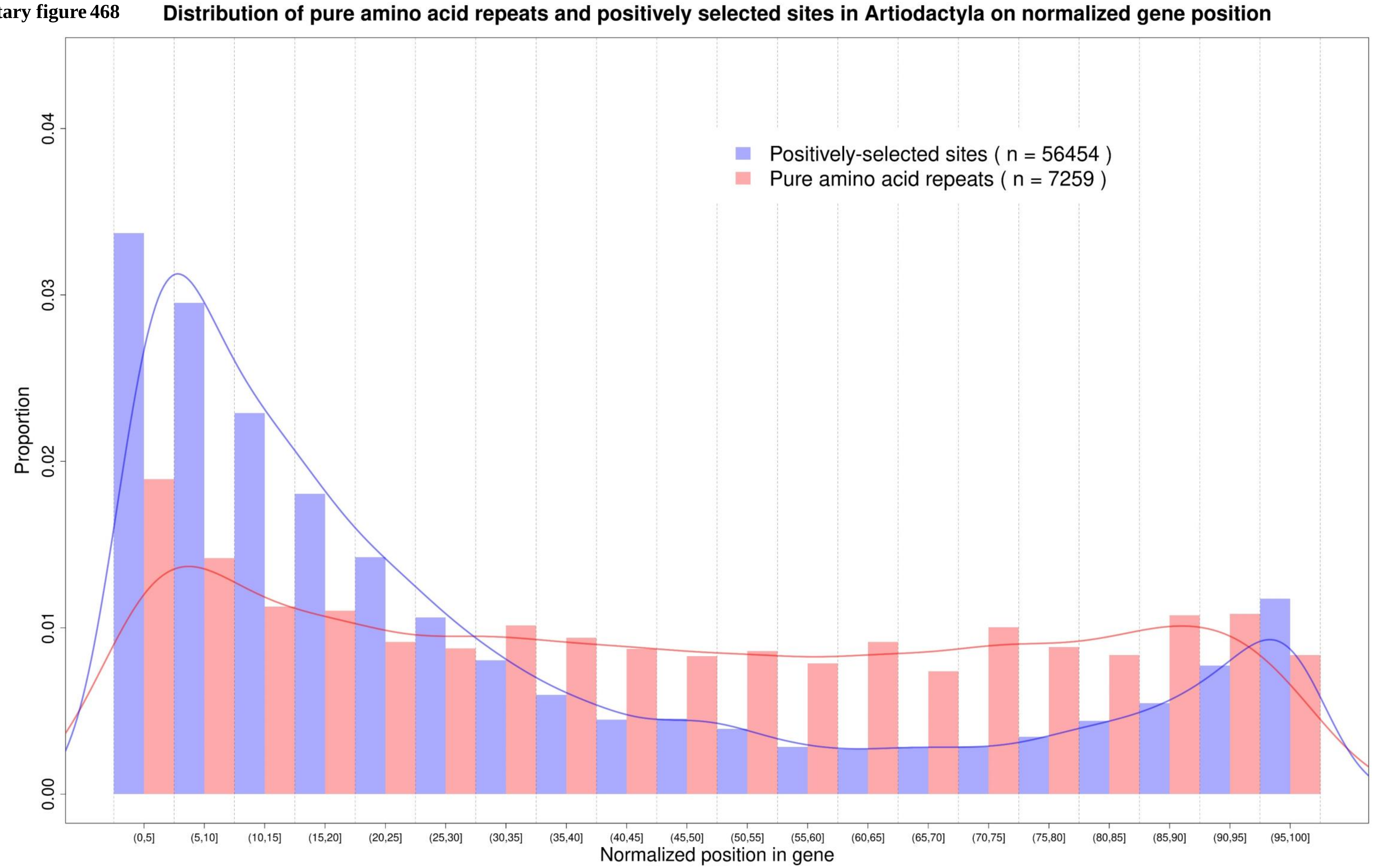

Supplementary figure 469

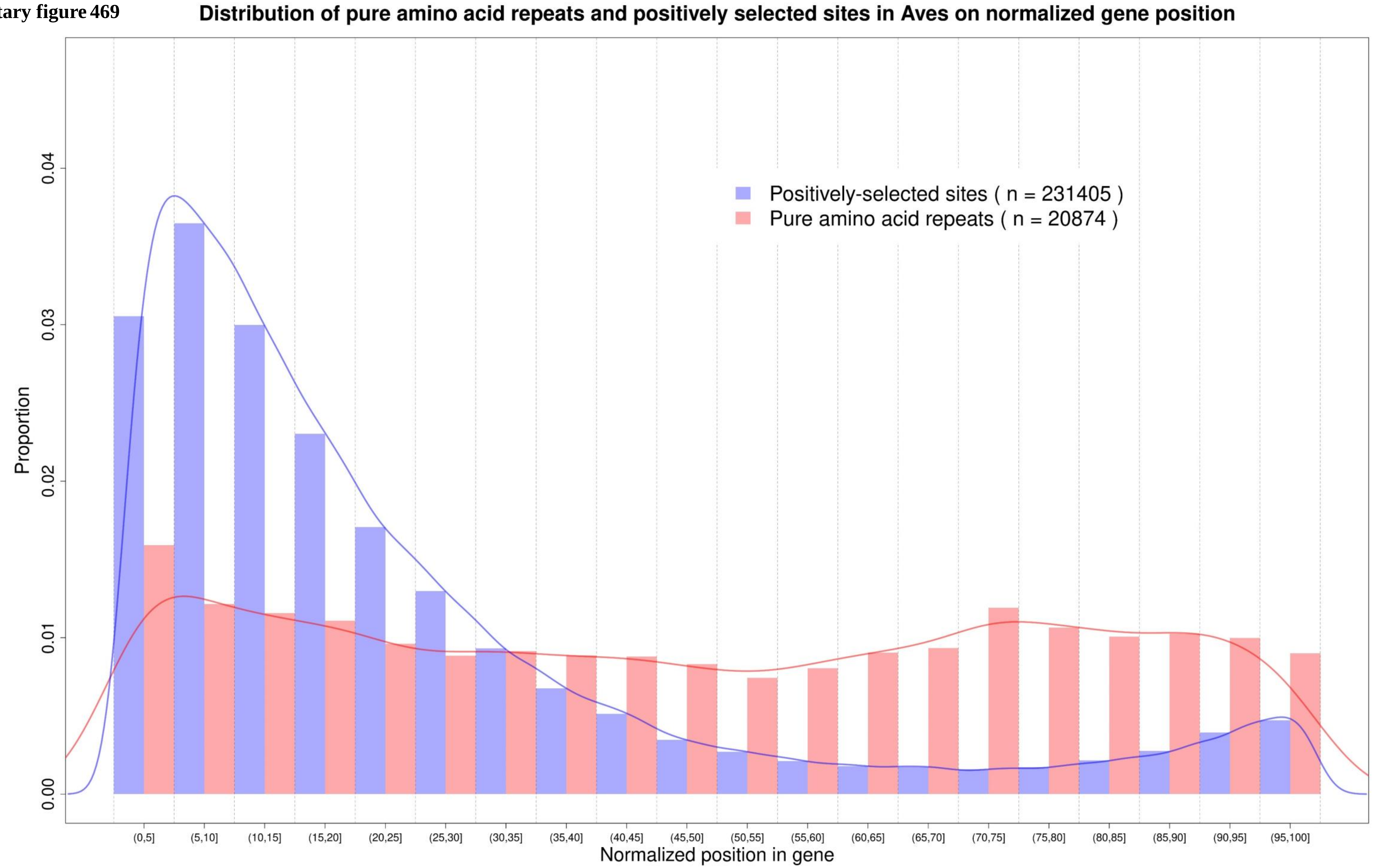

Supplementary figure 470

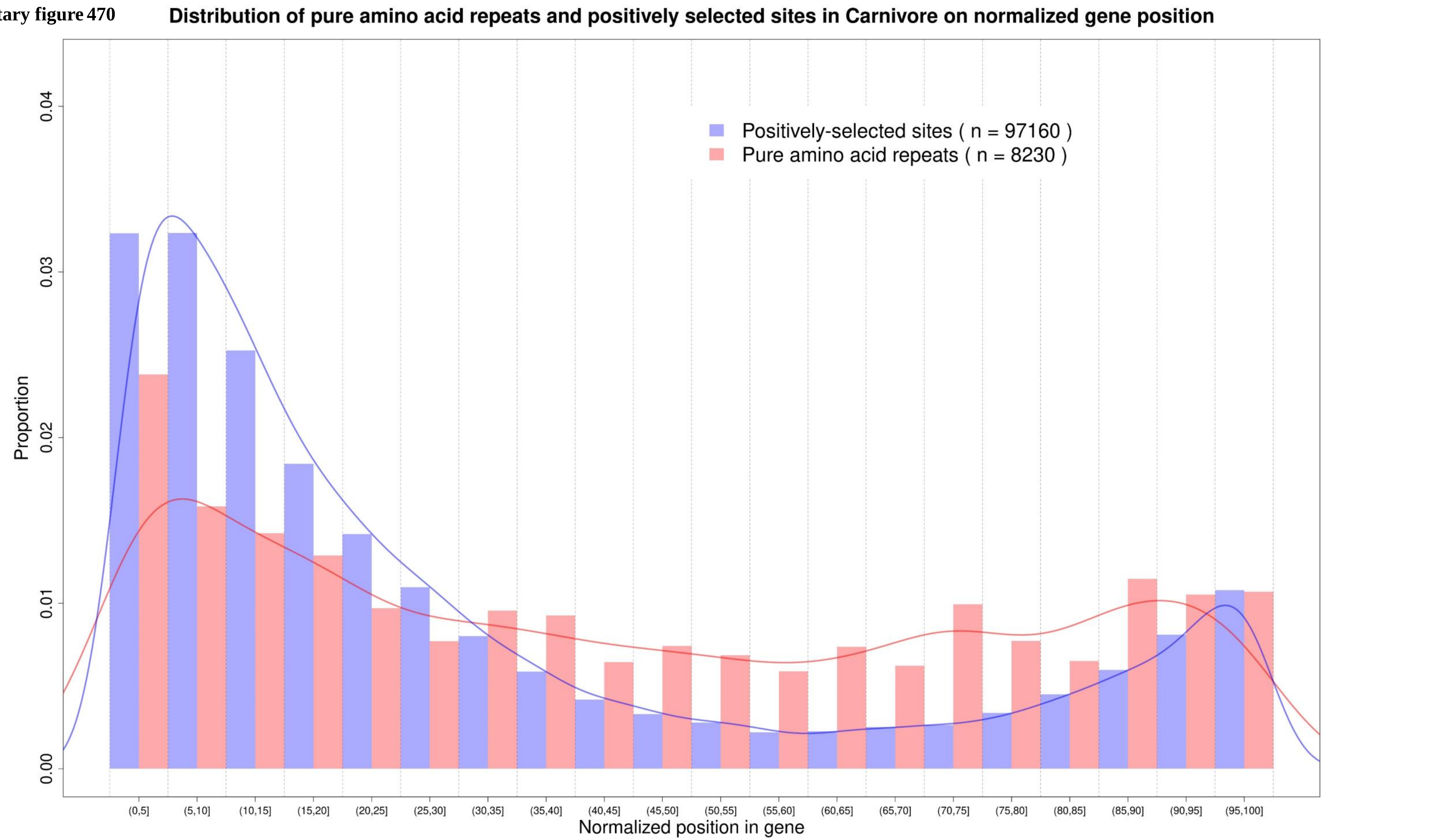

**Supplementary figure 471**

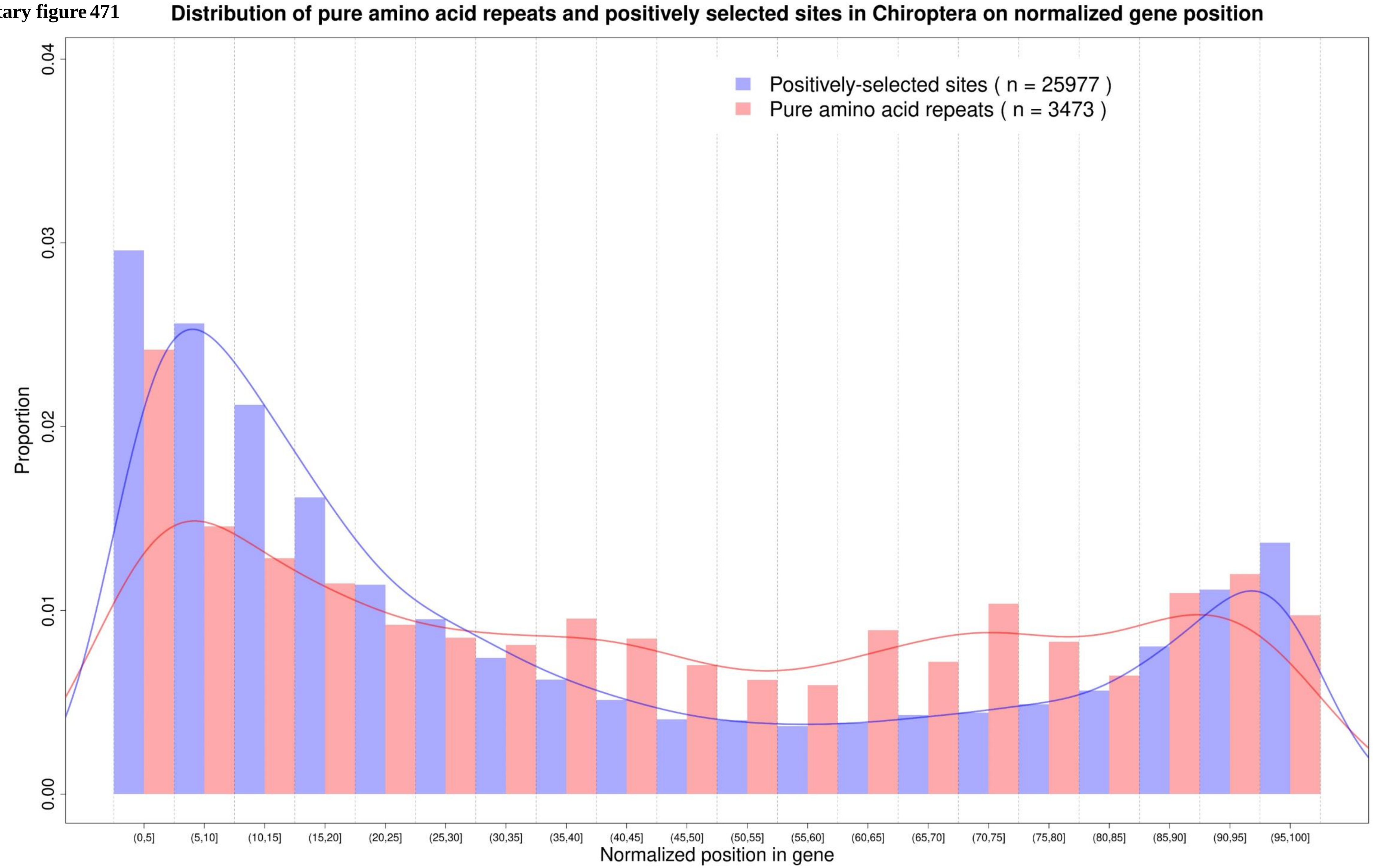

**Supplementary figure 472**

**Distribution of pure amino acid repeats and positively selected sites in Marsupials on normalized gene position**

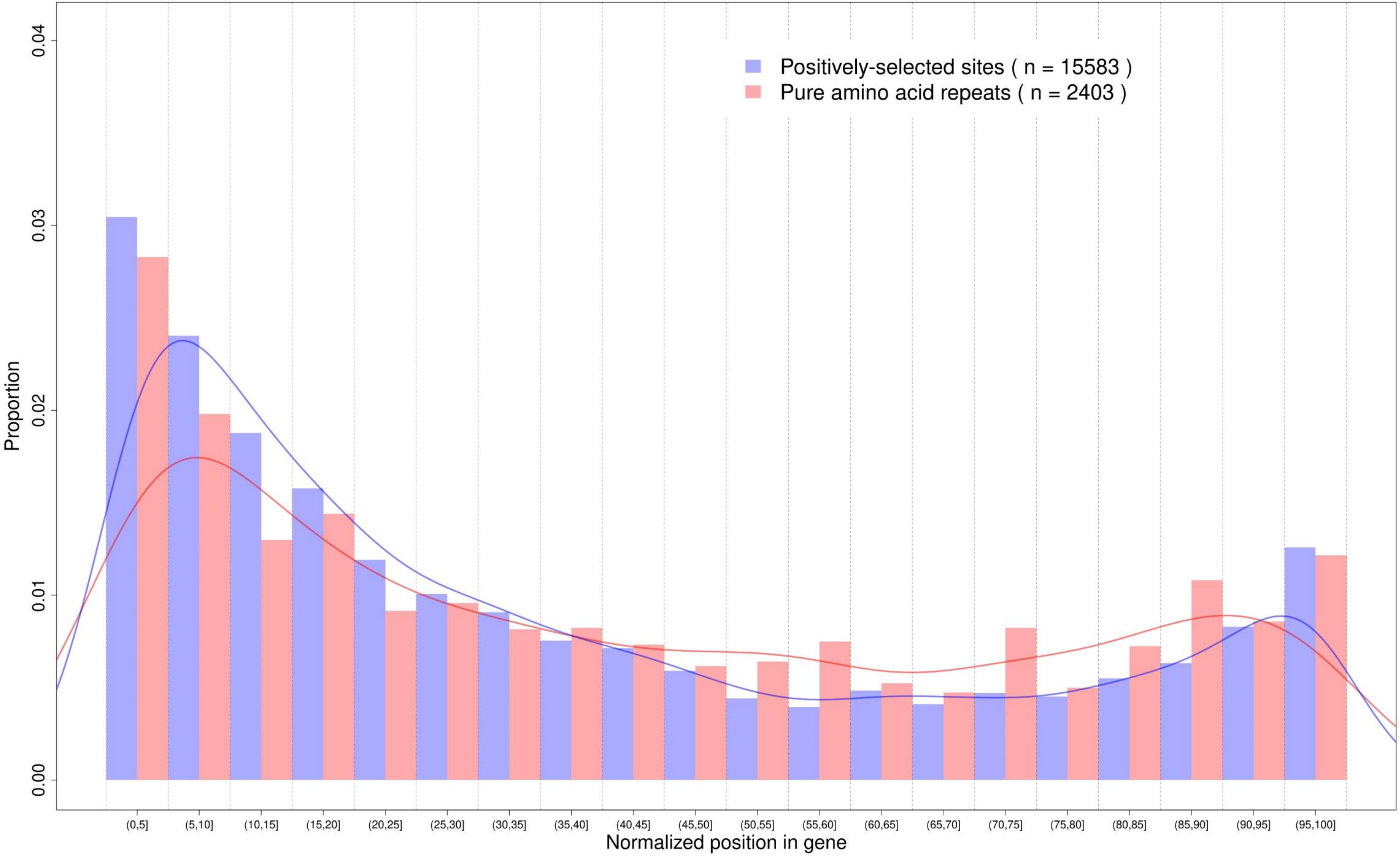

**Supplementary figure 473** **Distribution of pure amino acid repeats and positively selected sites in Perissodactyla on normalized gene position**

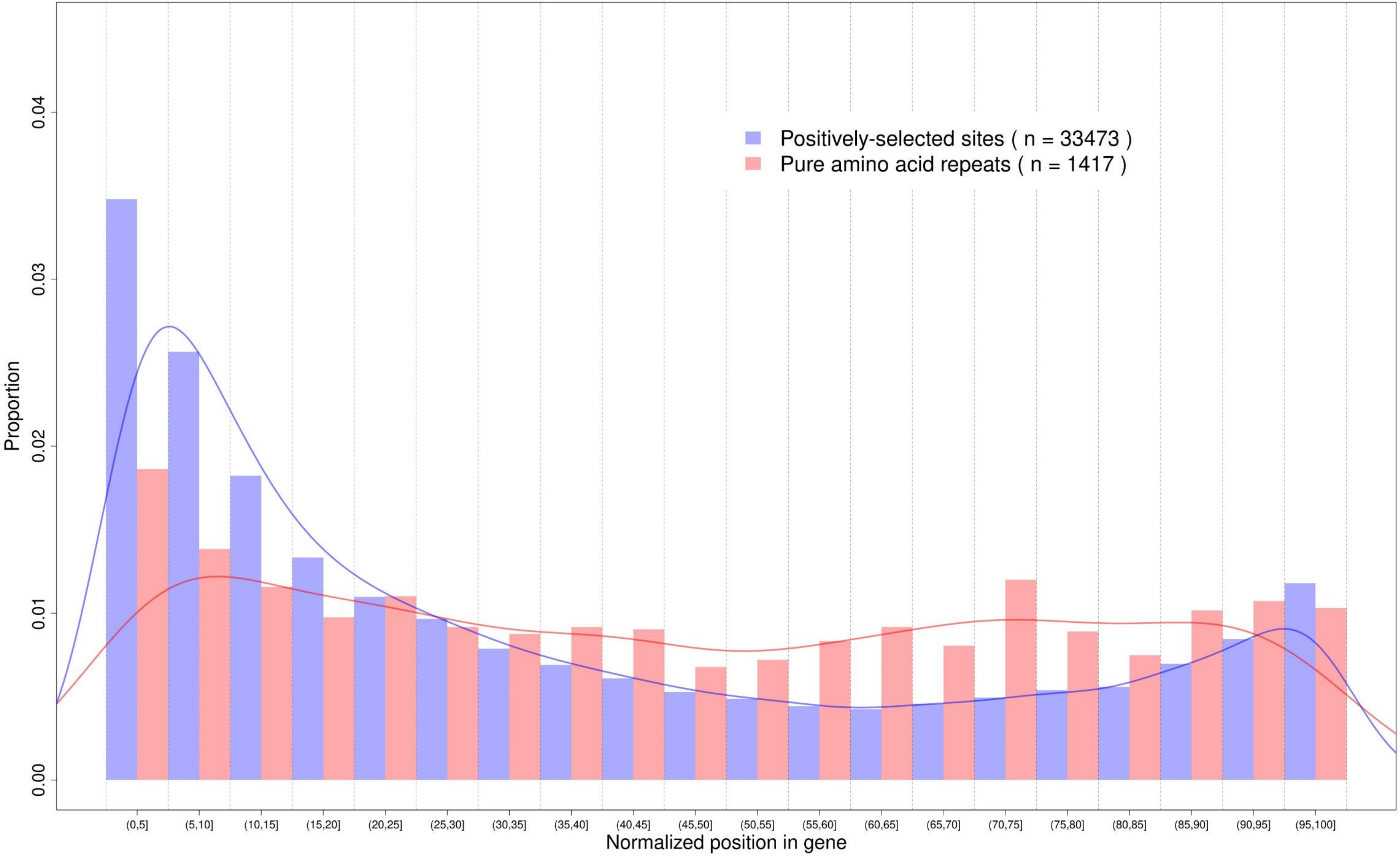

**Supplementary figure 474**

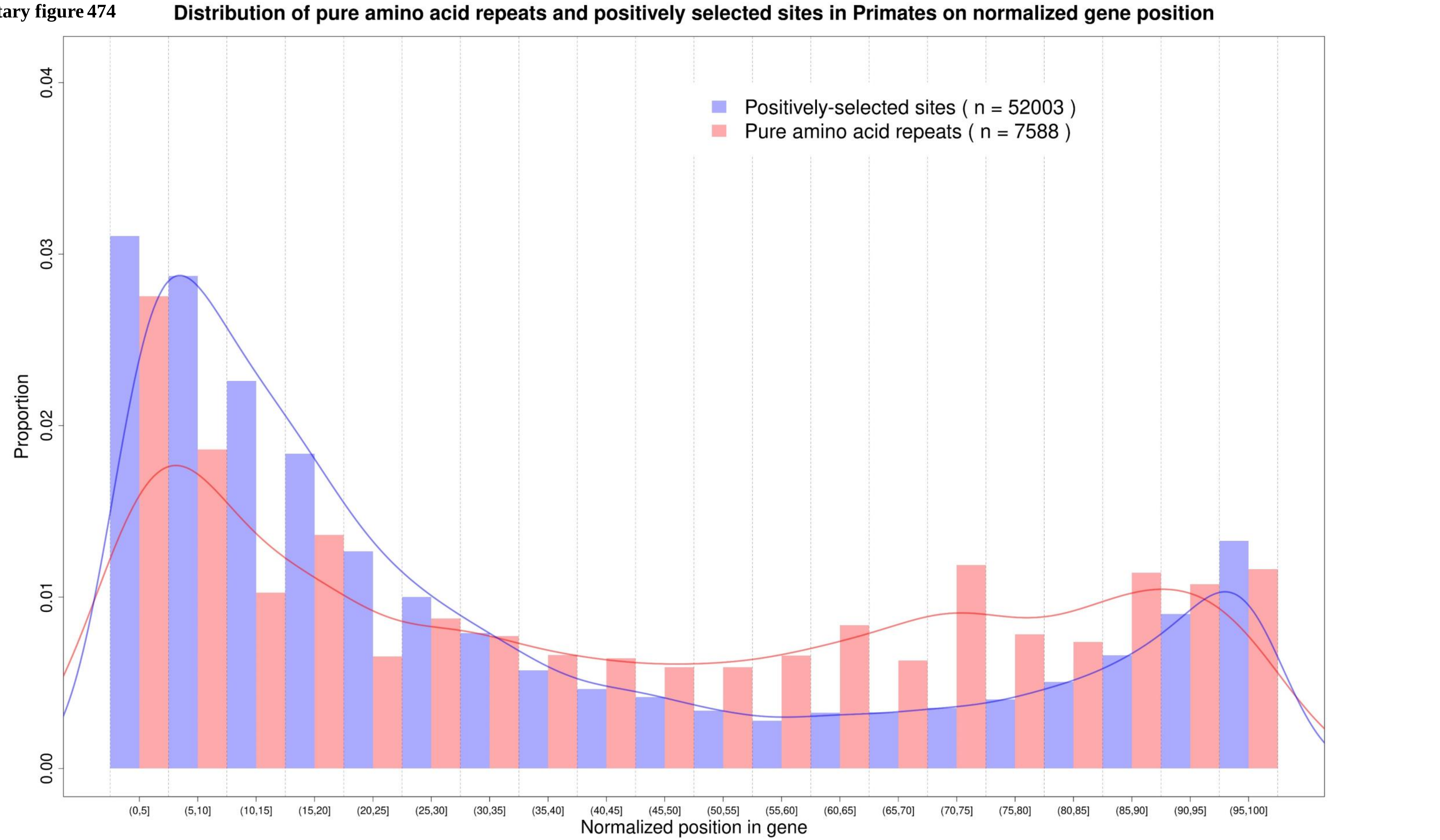

**Supplementary figure 475**

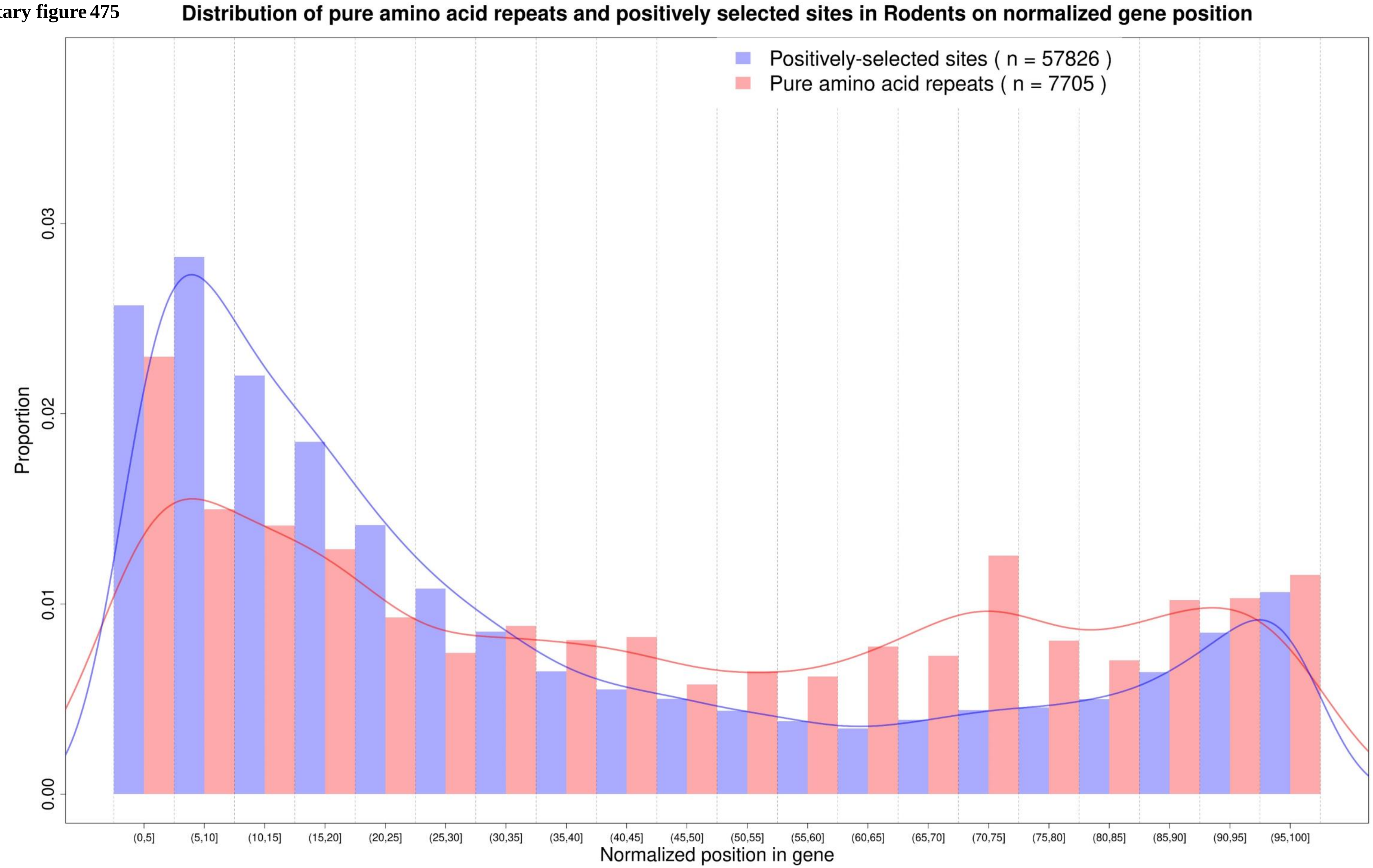

**Supplementary figure 476** **Distribution of pure amino acid repeats and positively selected sites in Squamata on normalized gene position**

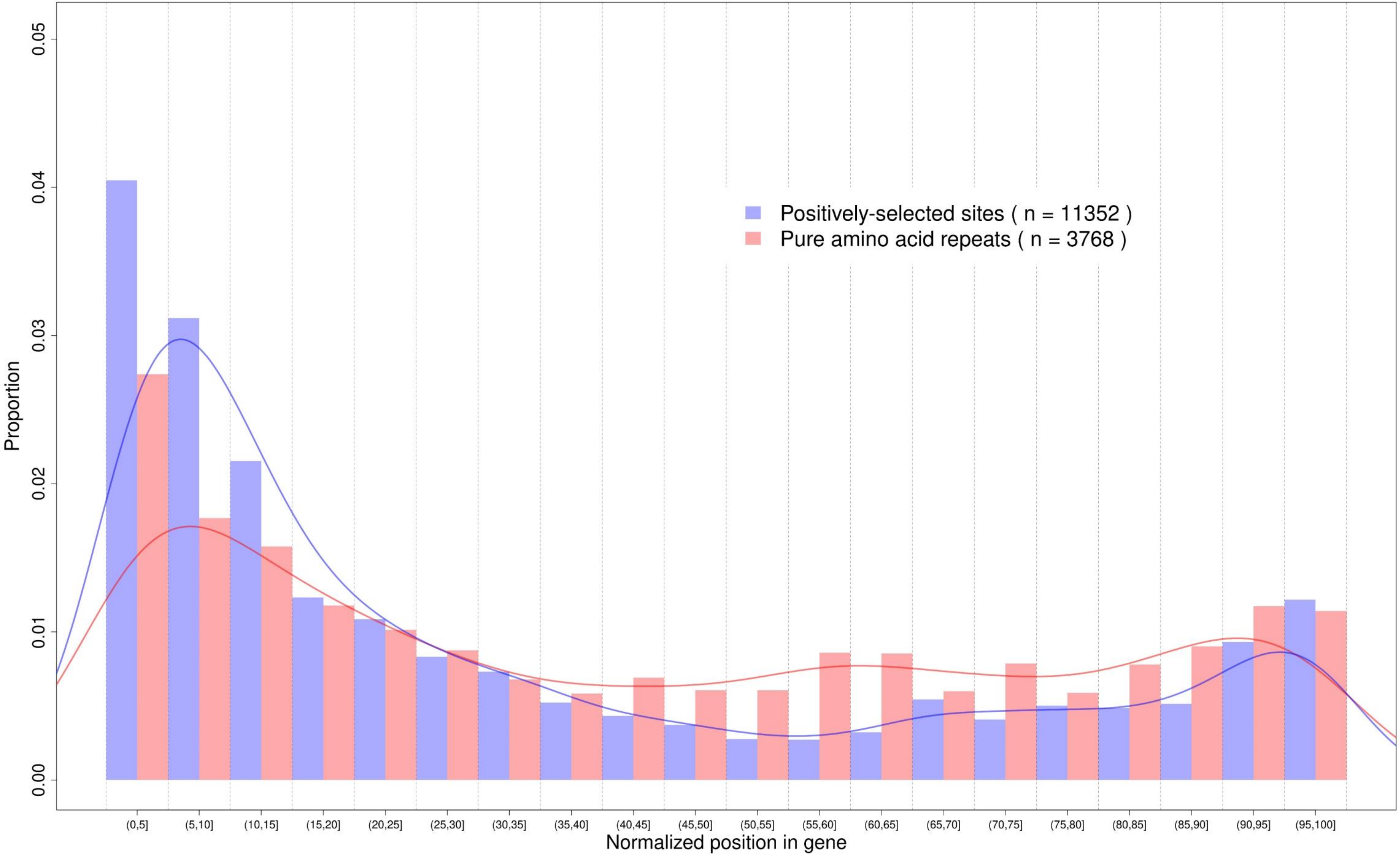

**Supplementary figure 477**

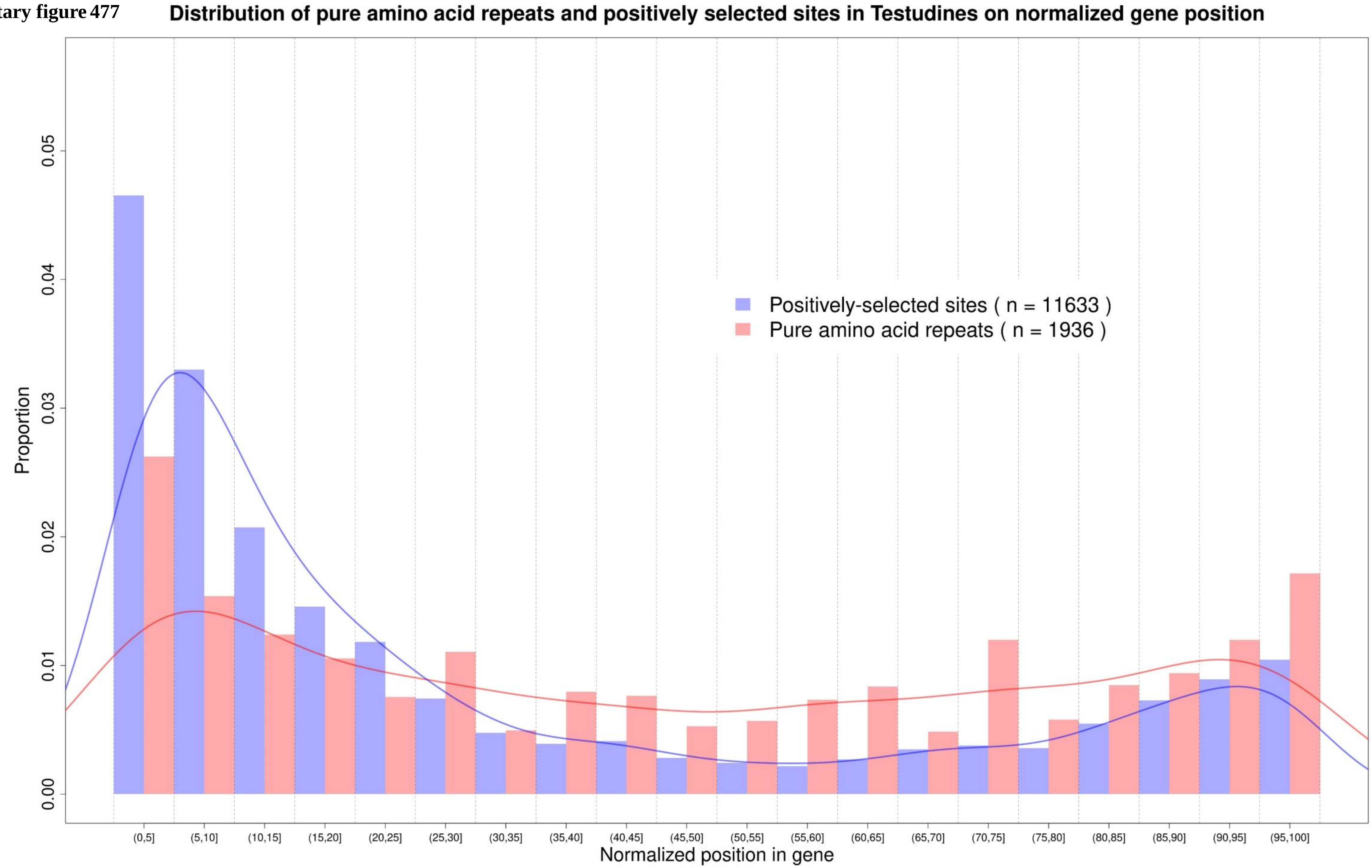

**Supplementary figure 478**

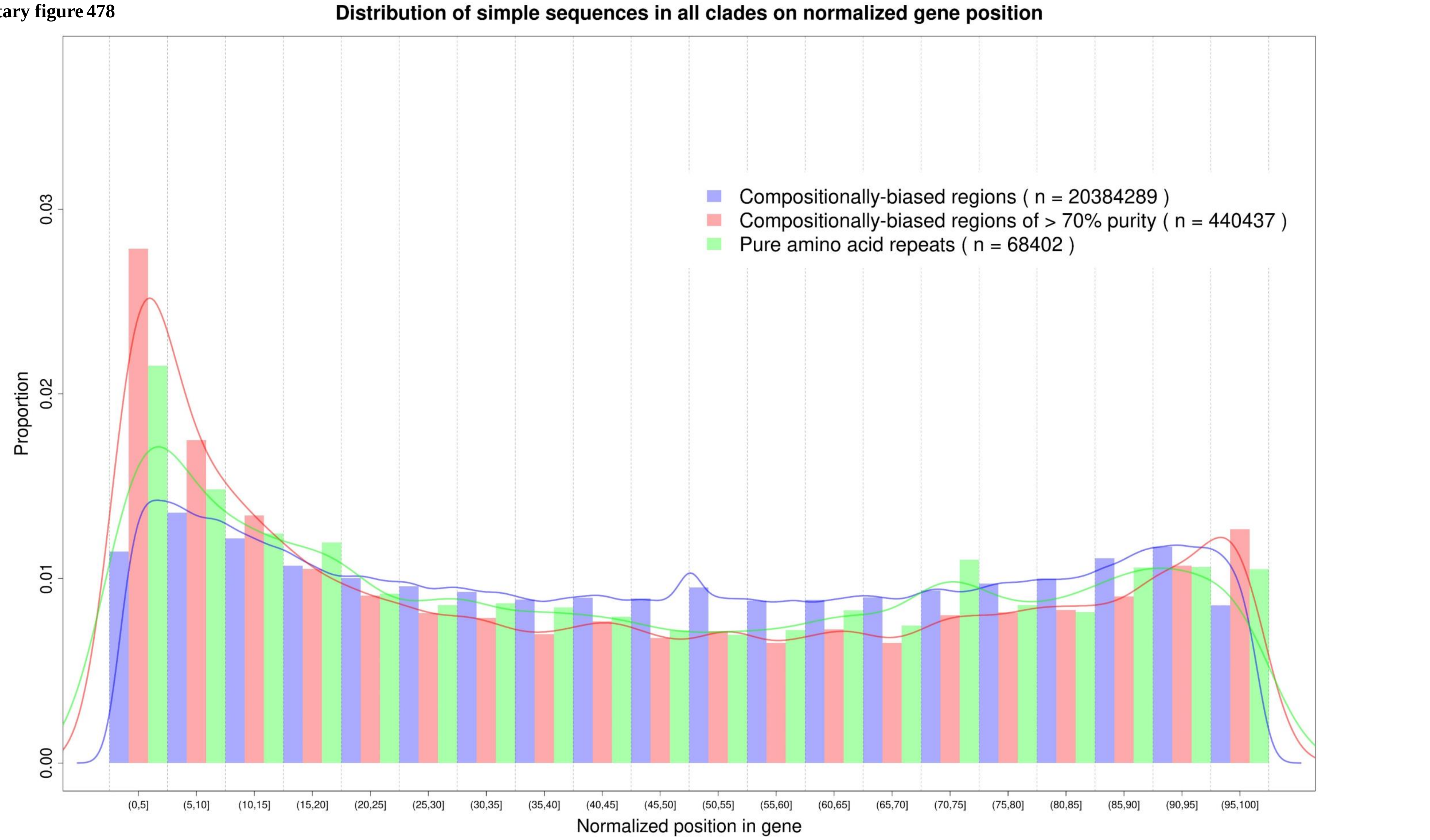

Supplementary figure 479

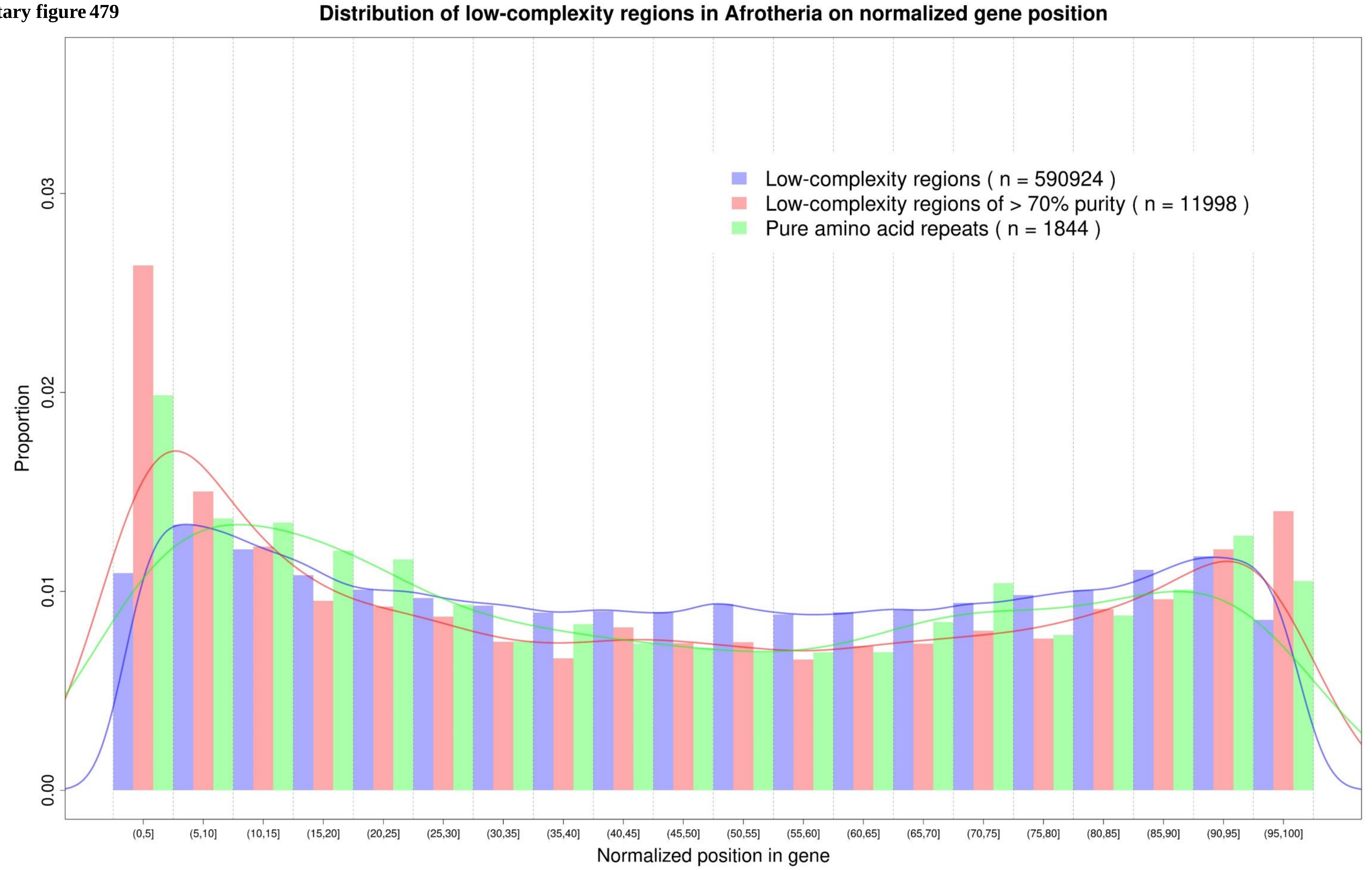

**Supplementary figure 480**

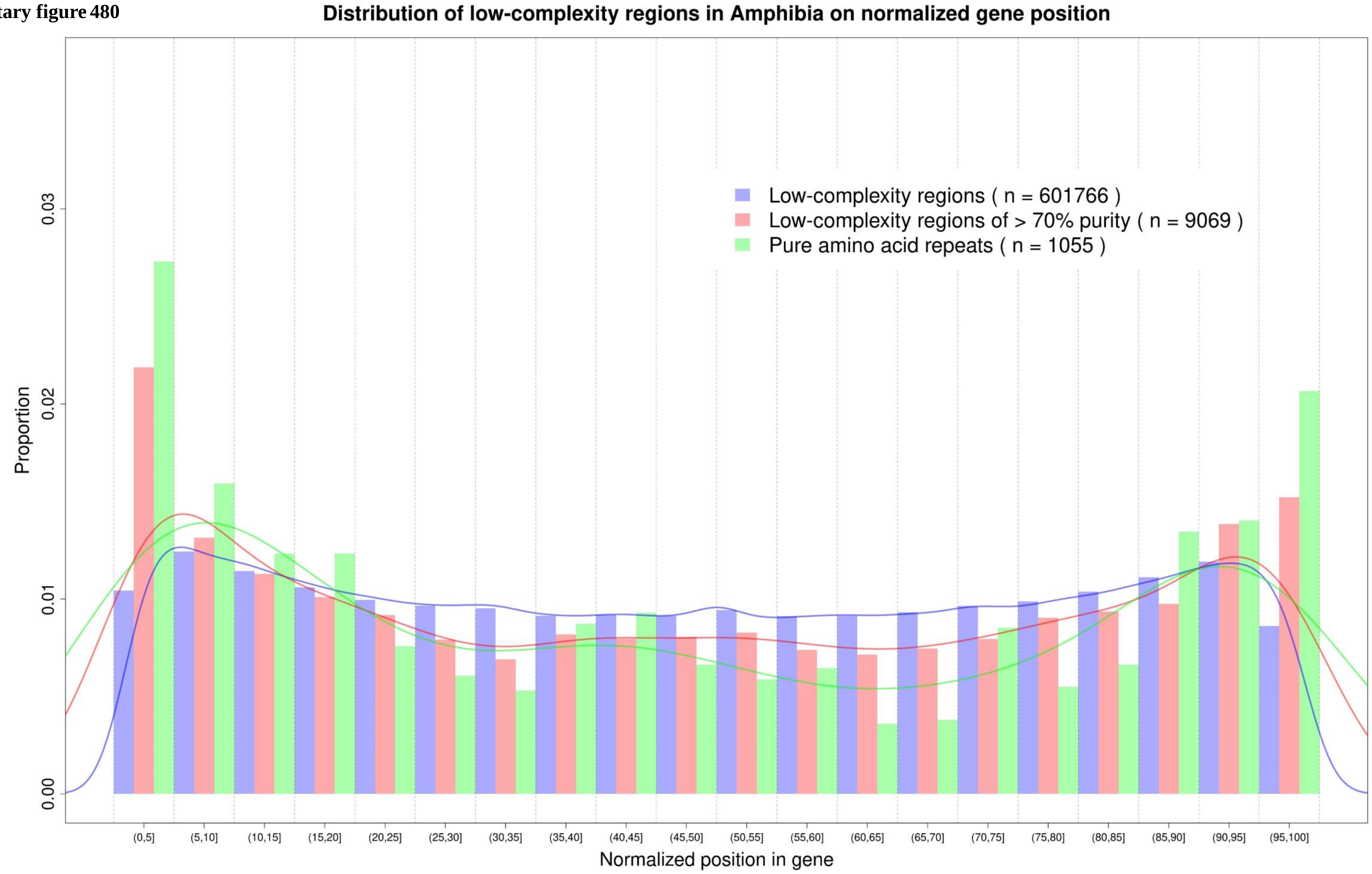

**Supplementary figure 481**

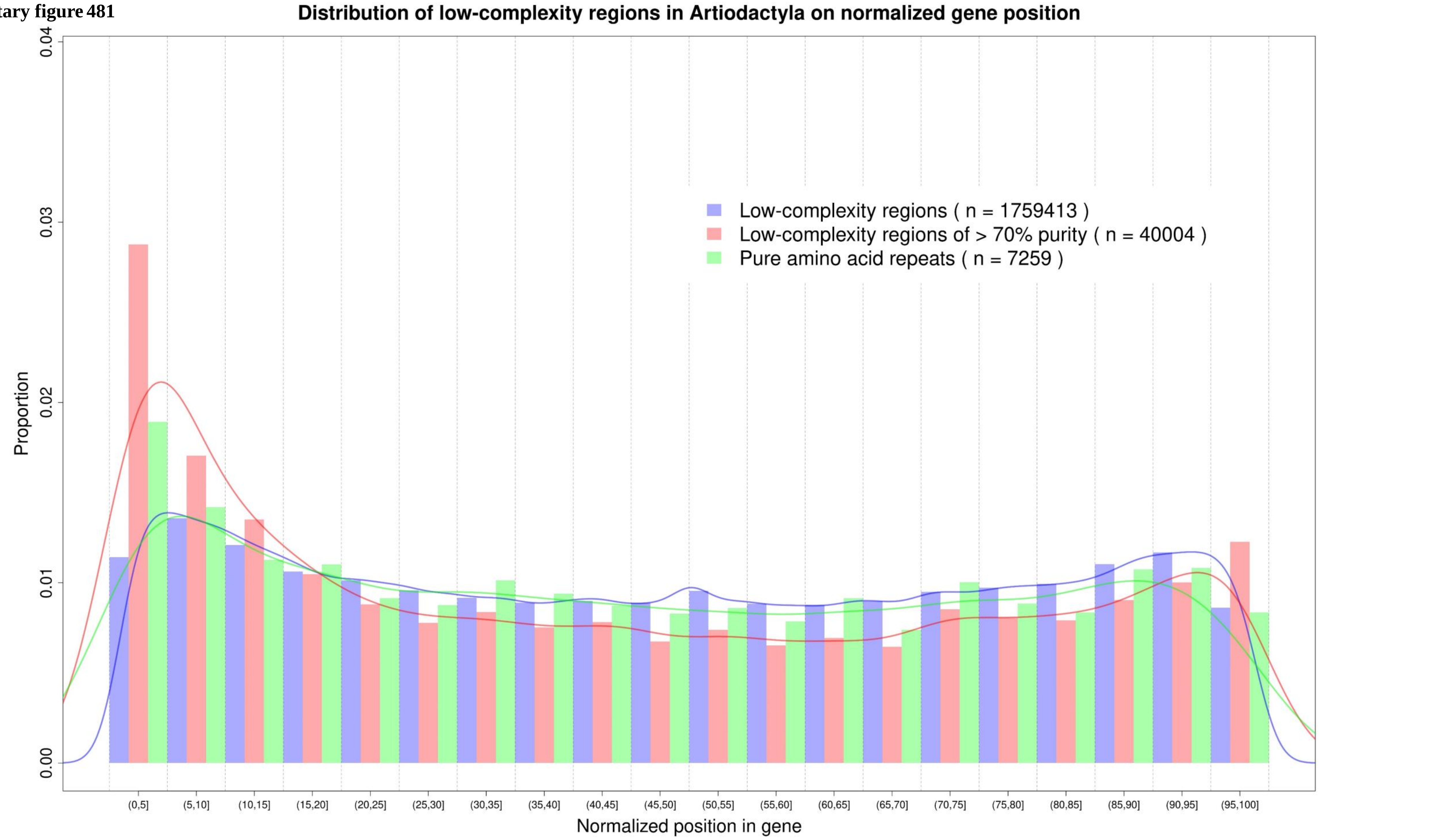

**Supplementary figure 482**

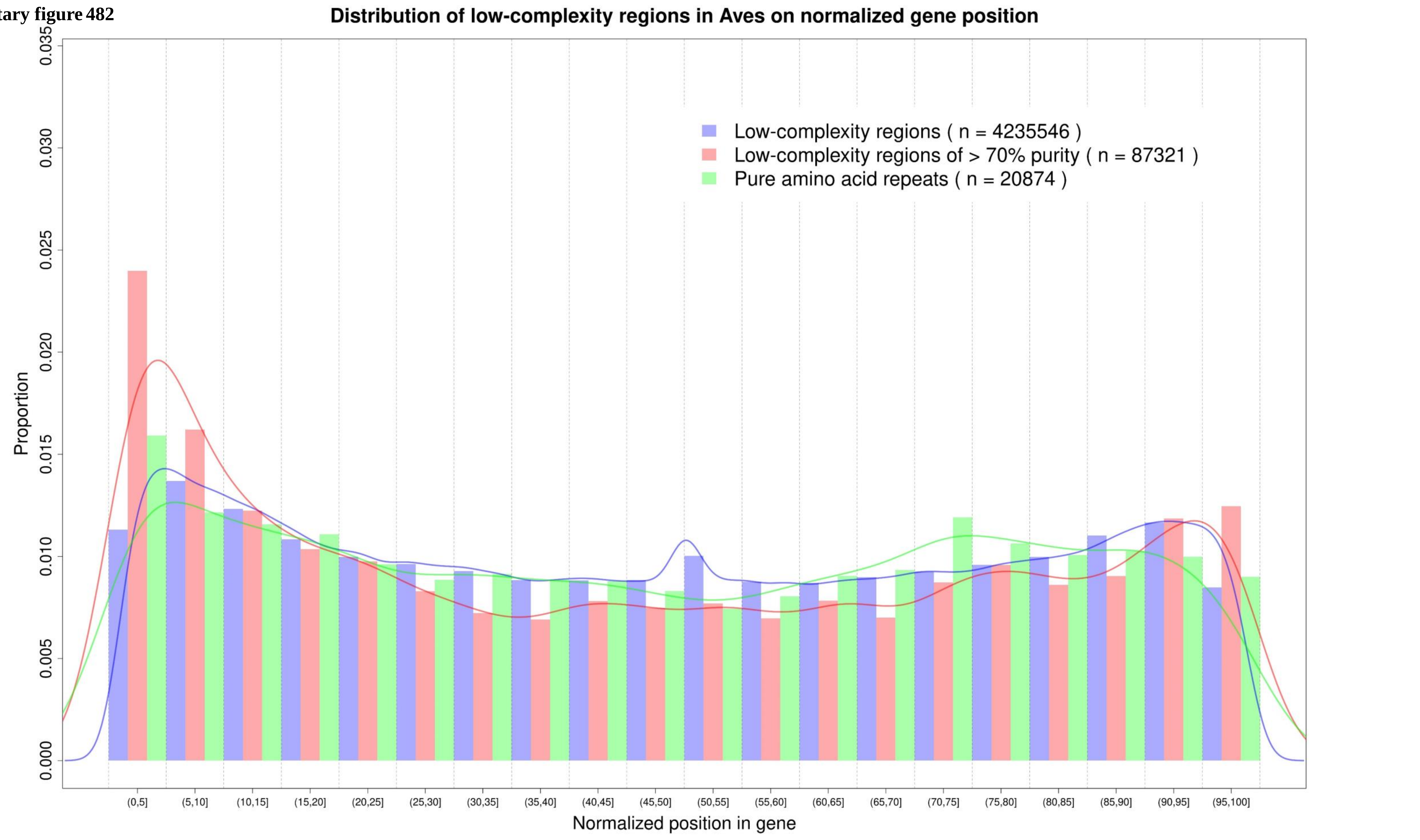

Supplementary figure 483

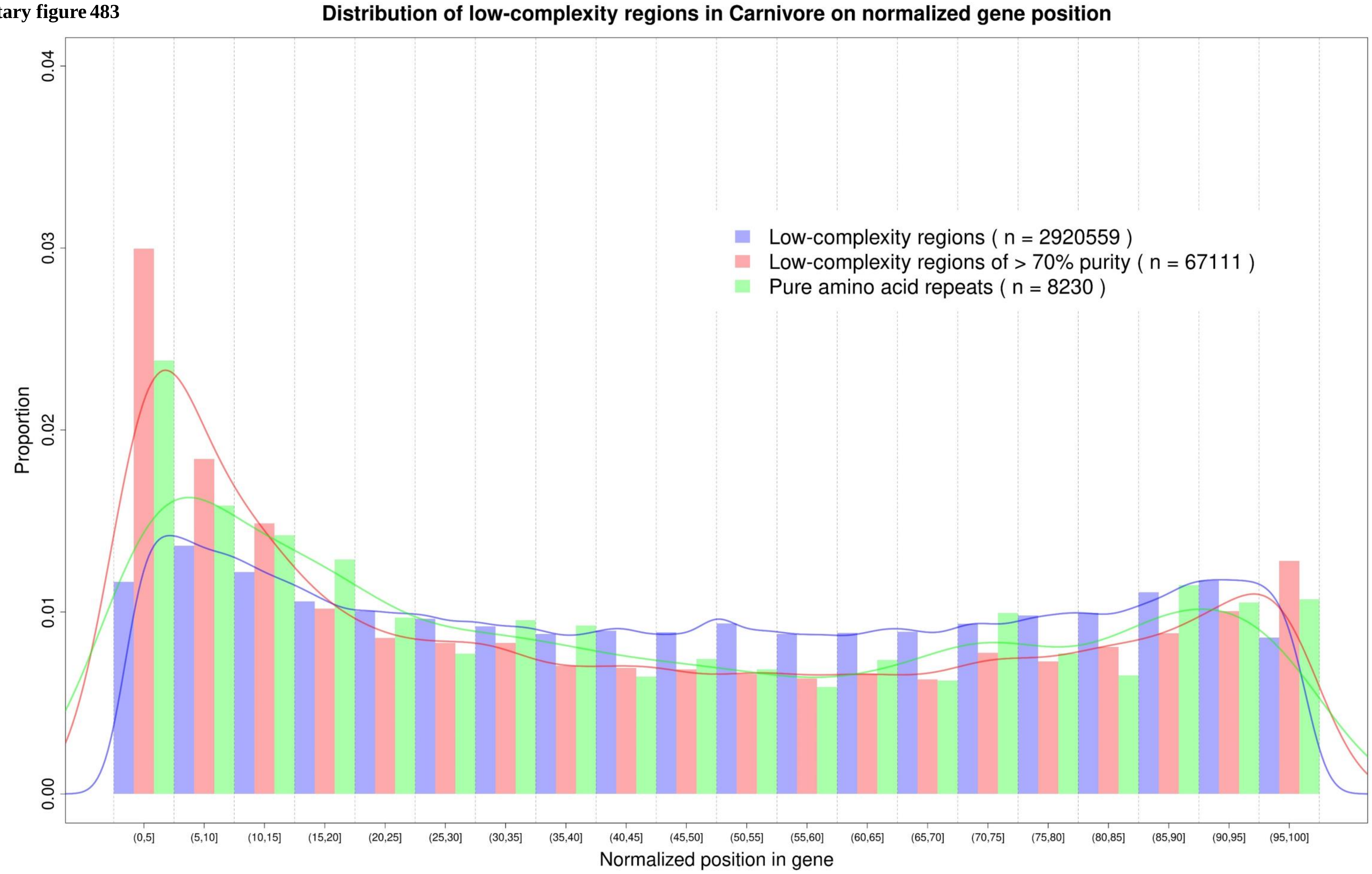

Supplementary figure 484

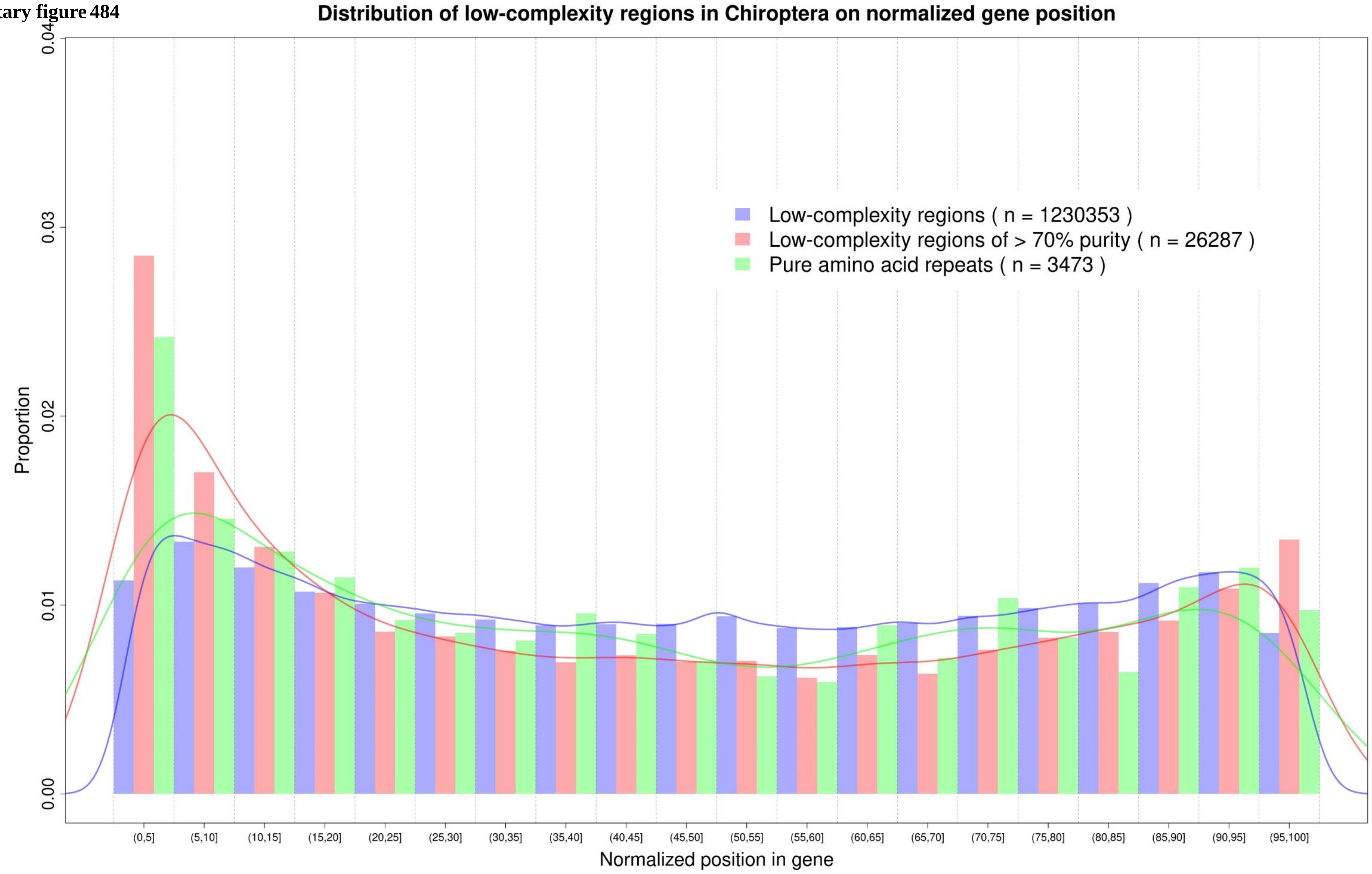

Supplementary figure 485

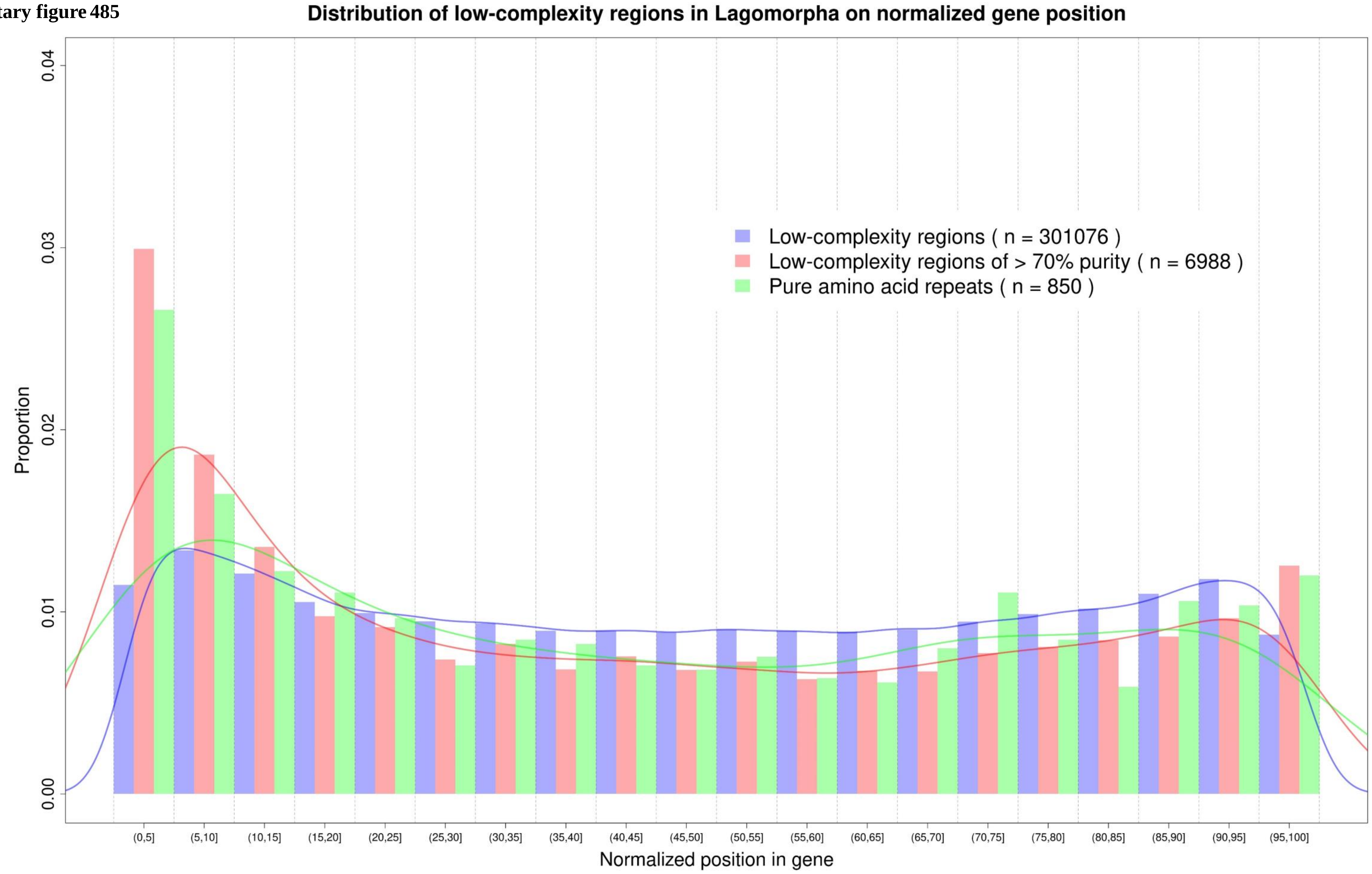

Supplementary figure 486

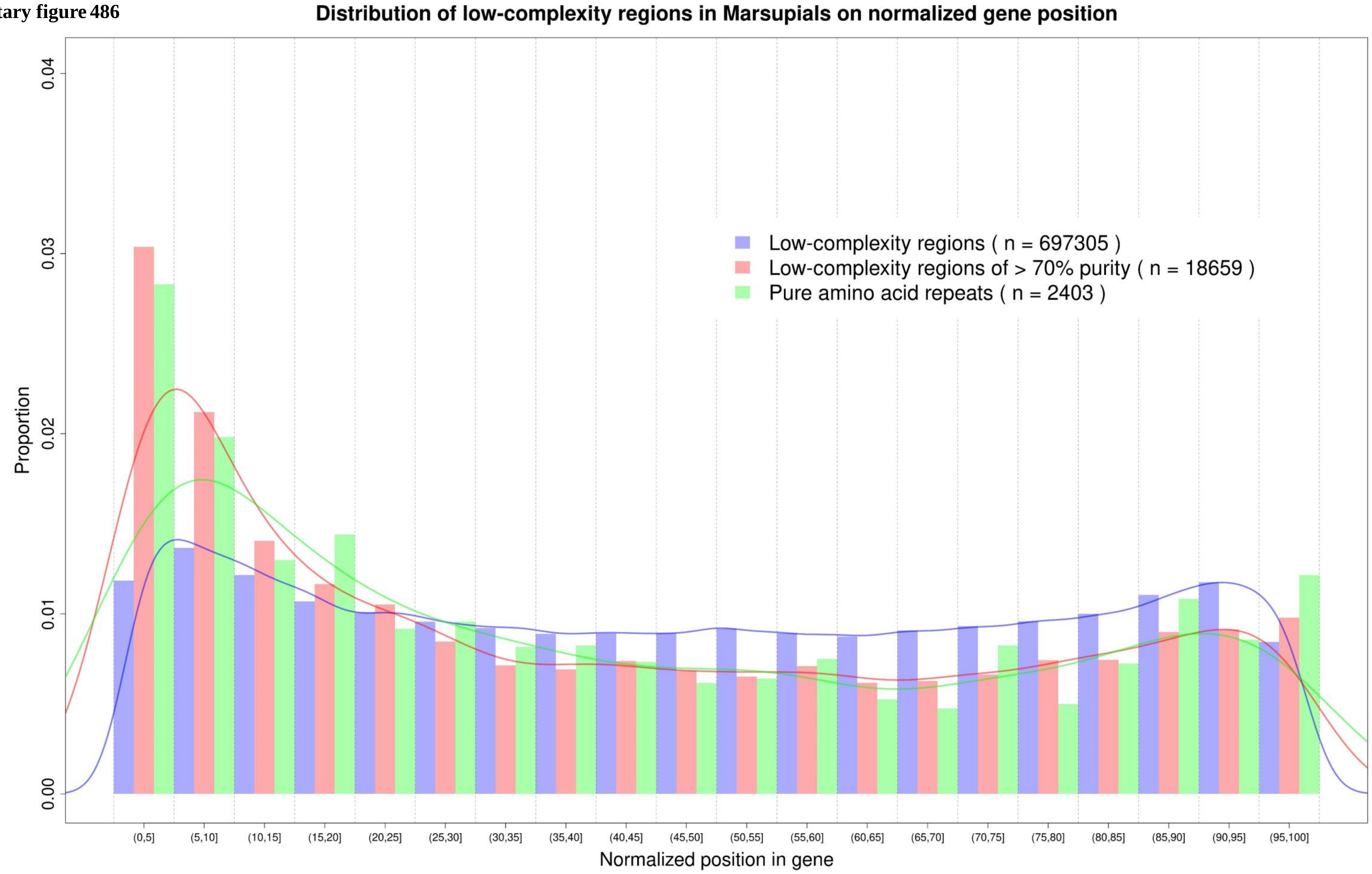

Supplementary figure 487

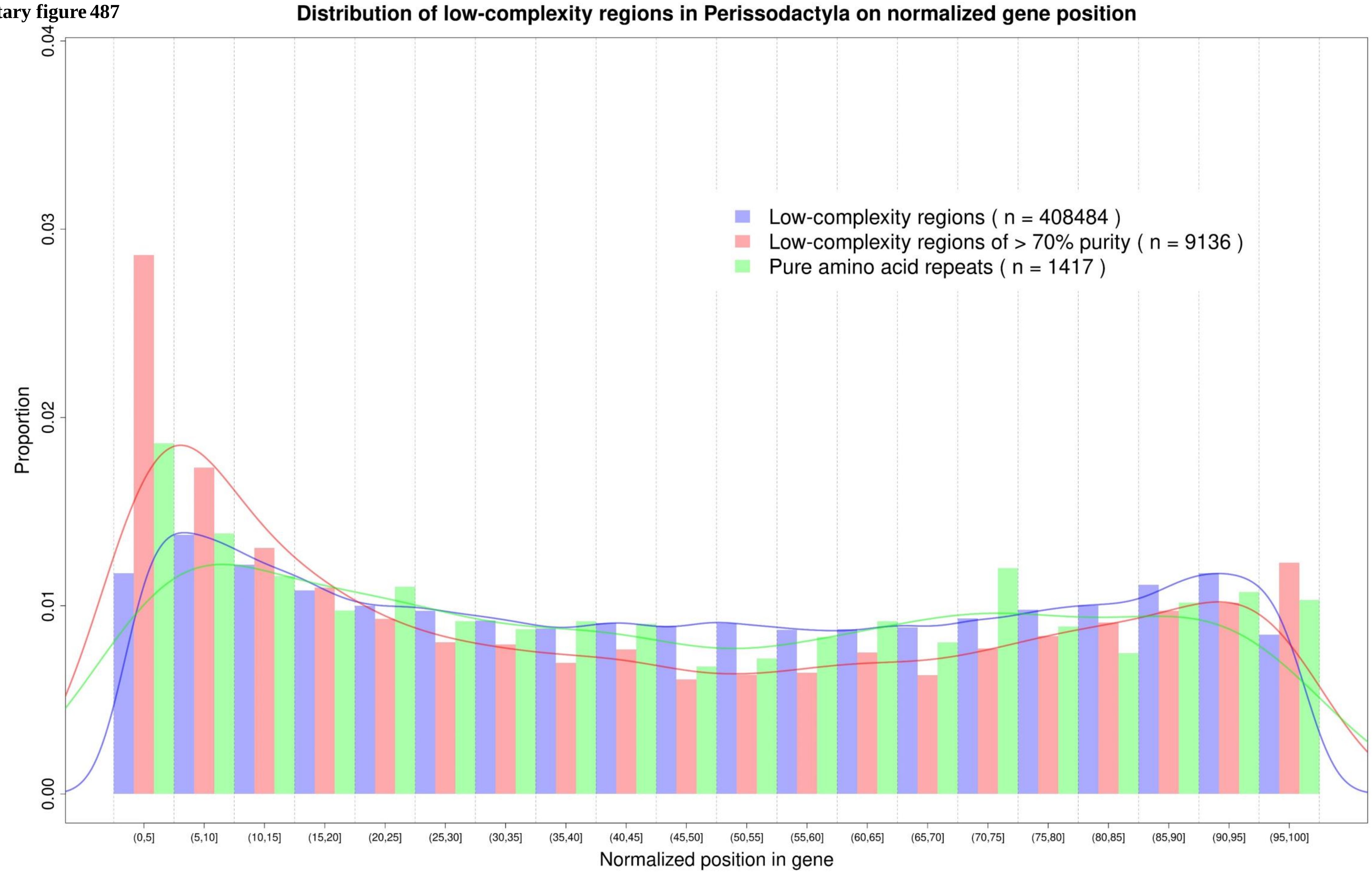

Supplementary figure 488

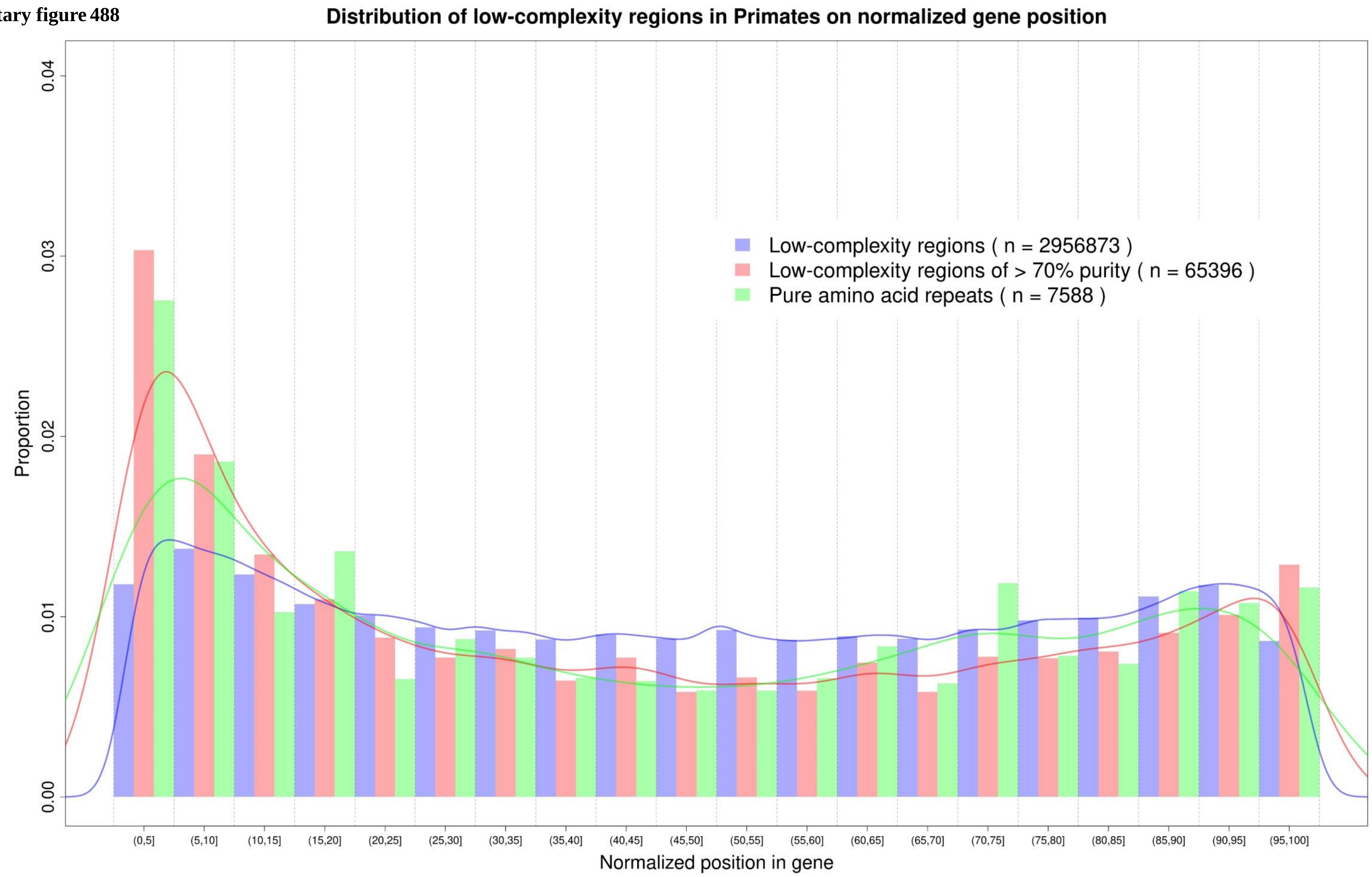

Supplementary figure 489

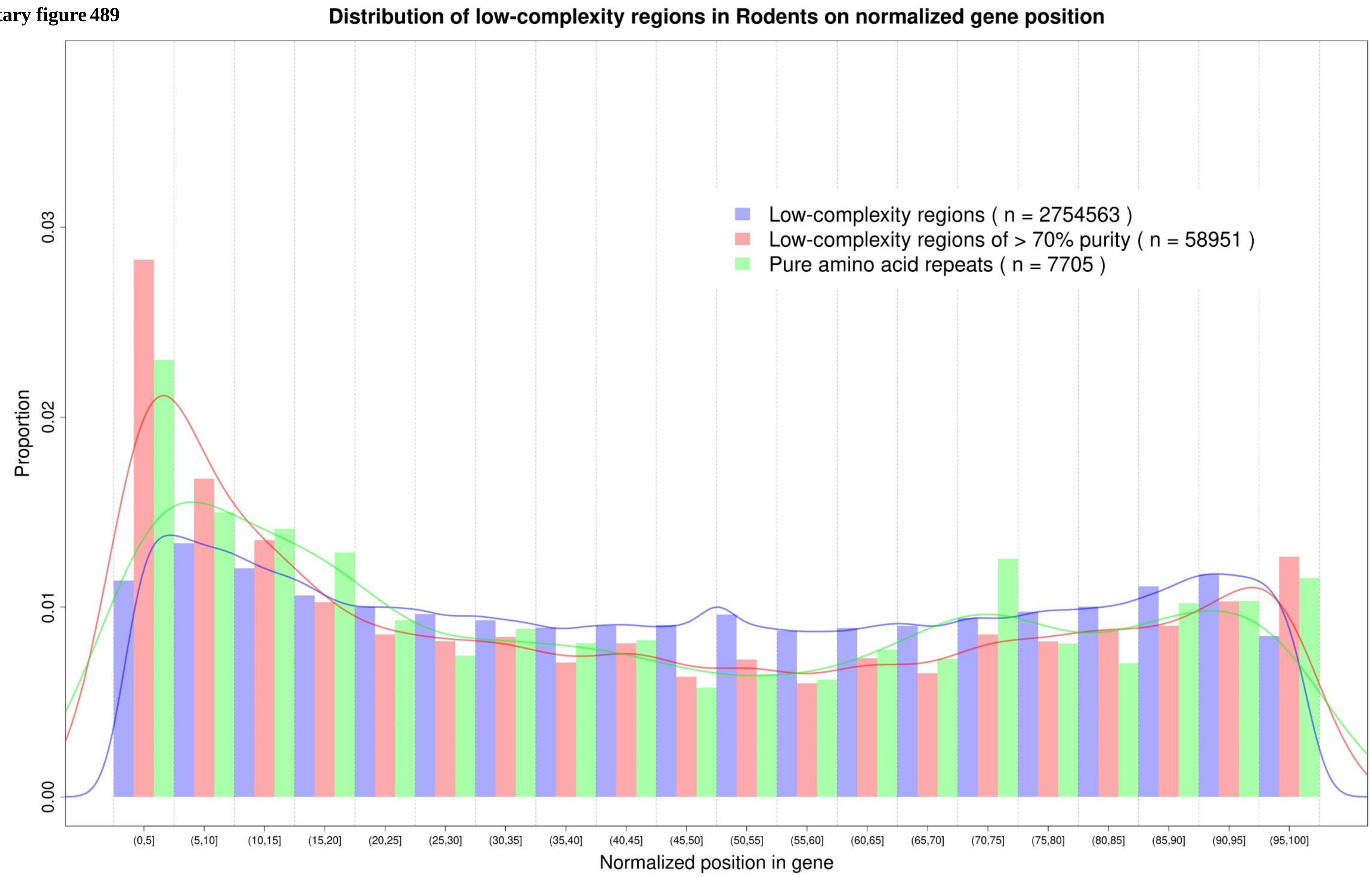

Supplementary figure 490

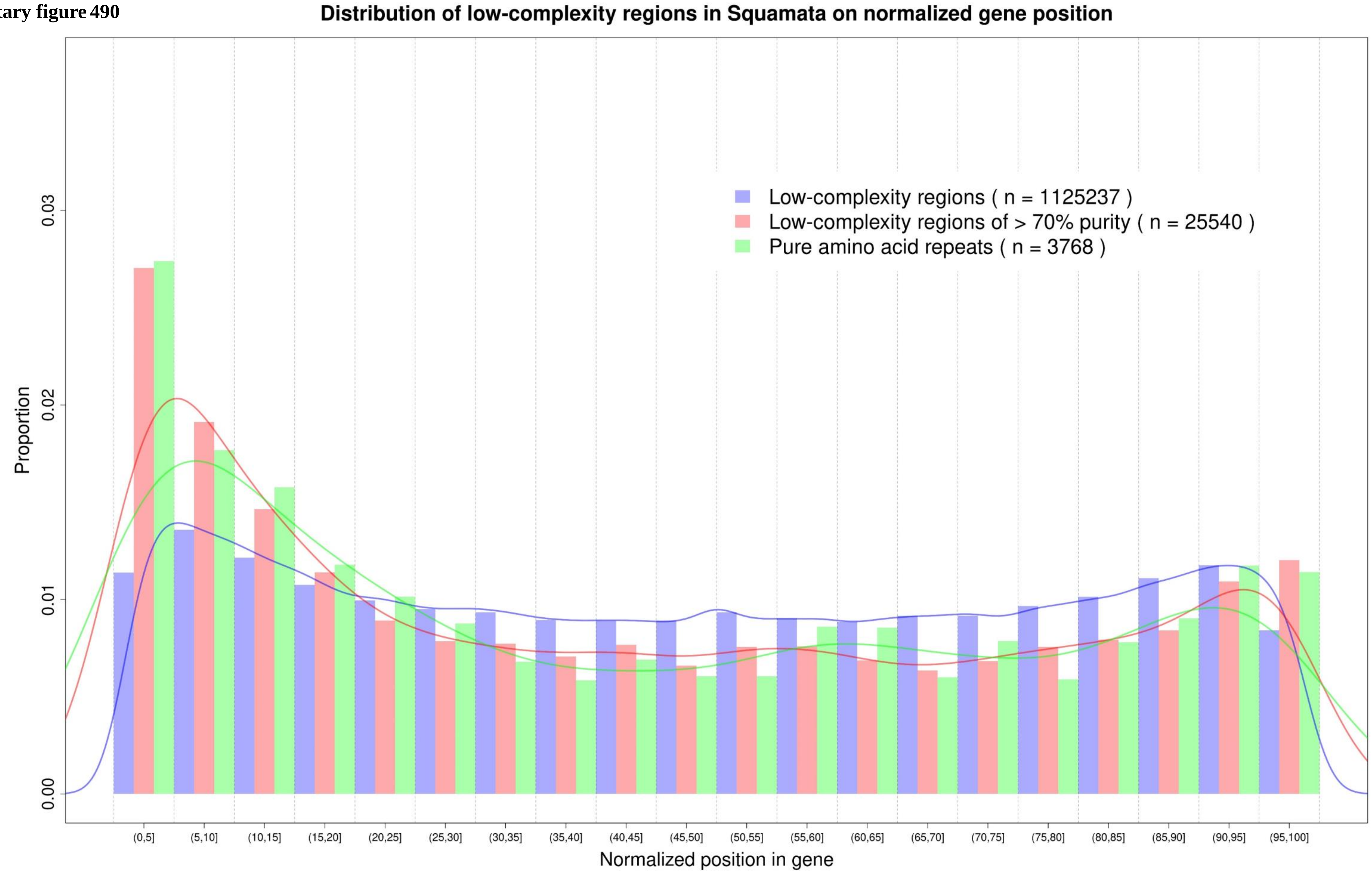

Supplementary figure 491

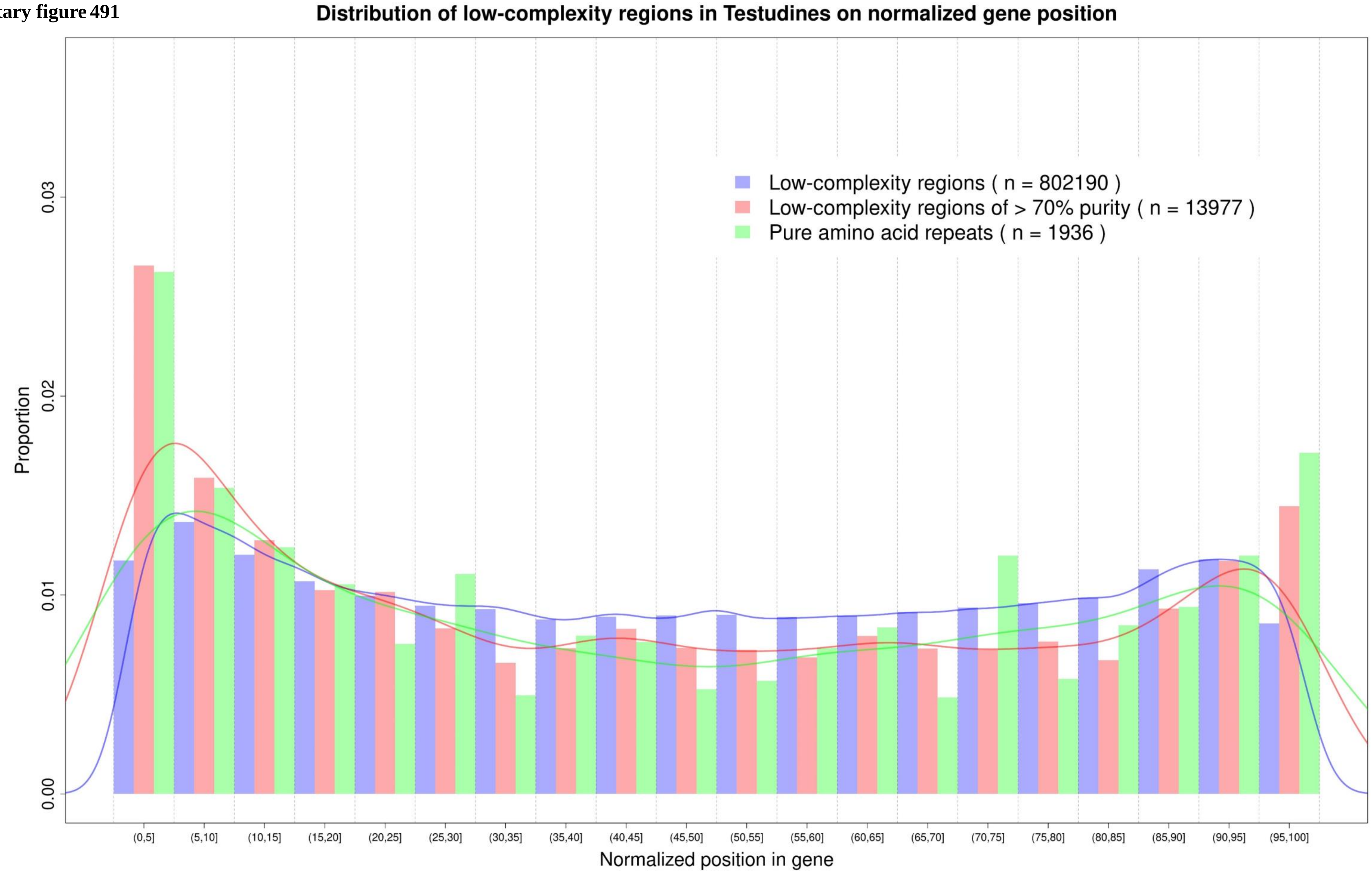

**Supplementary figure 492**

Pairwise correlation of alignment coverage, PSSs and LCRs in all the clades

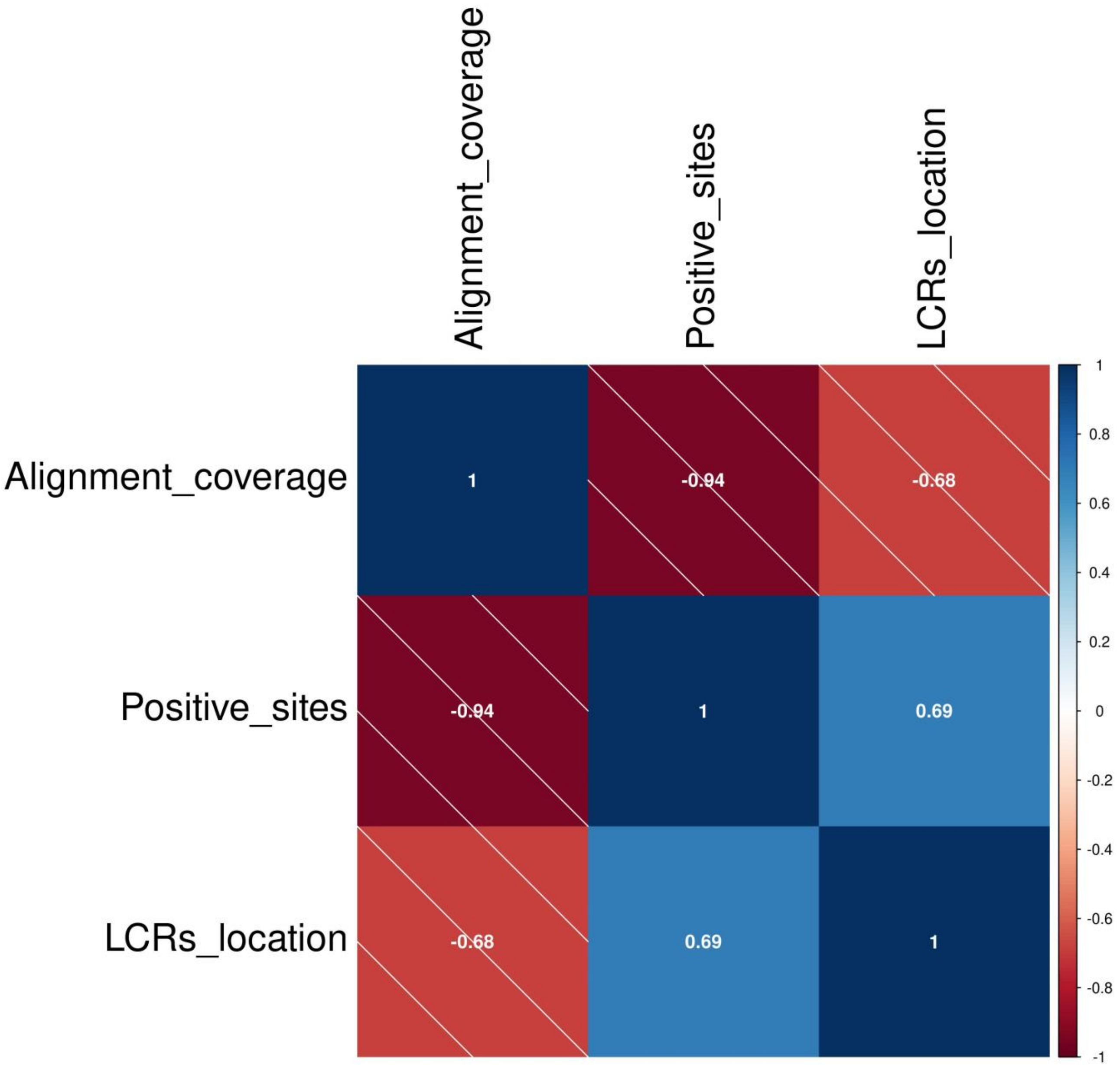

Supplementary figure 493

### Pairwise correlation of alignment coverage, PSSs and LCRs in Afrotheria clade

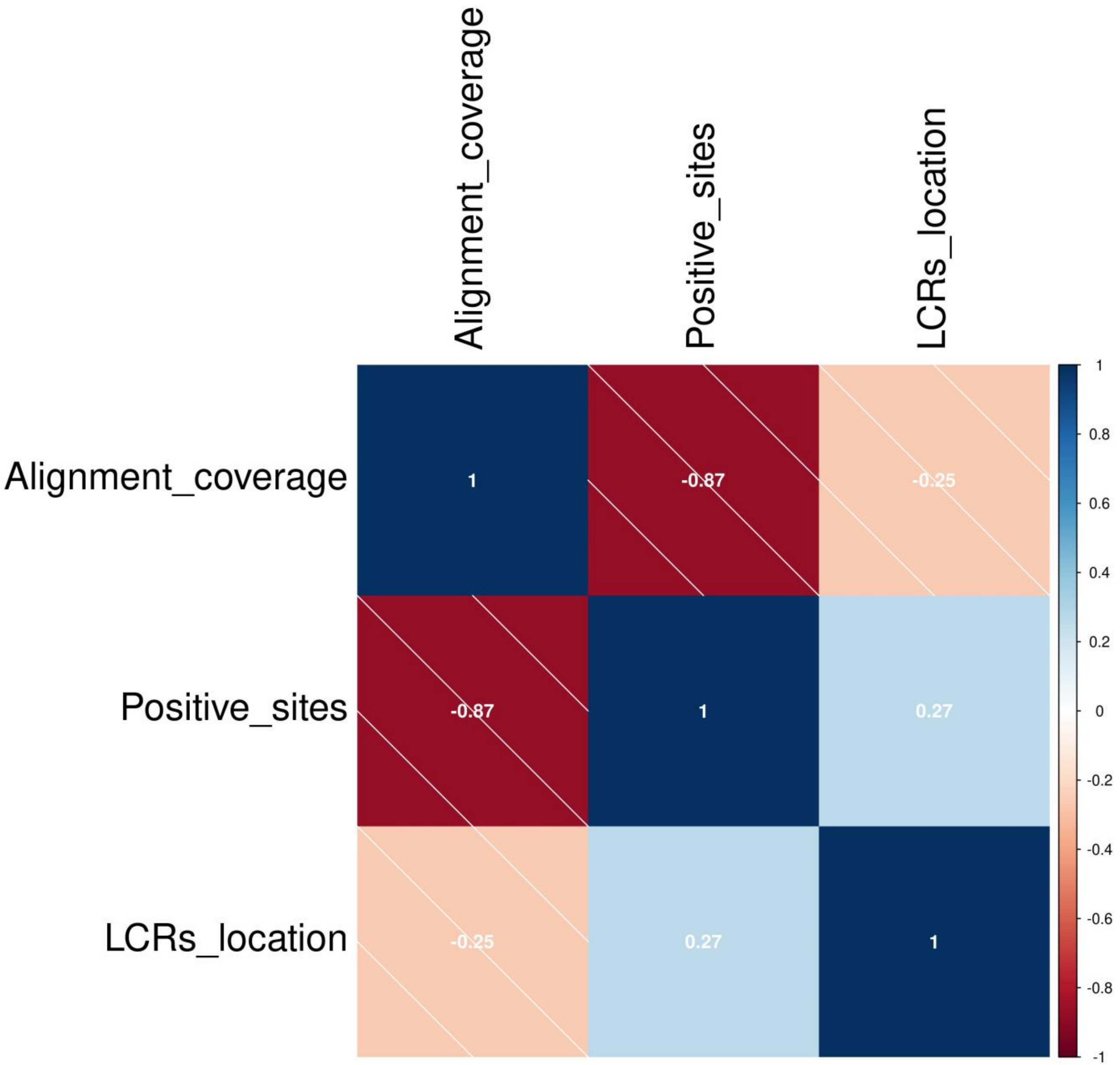

**Supplementary figure 494** Pairwise correlation of alignment coverage, PSSs and LCRs in Amphibia clade

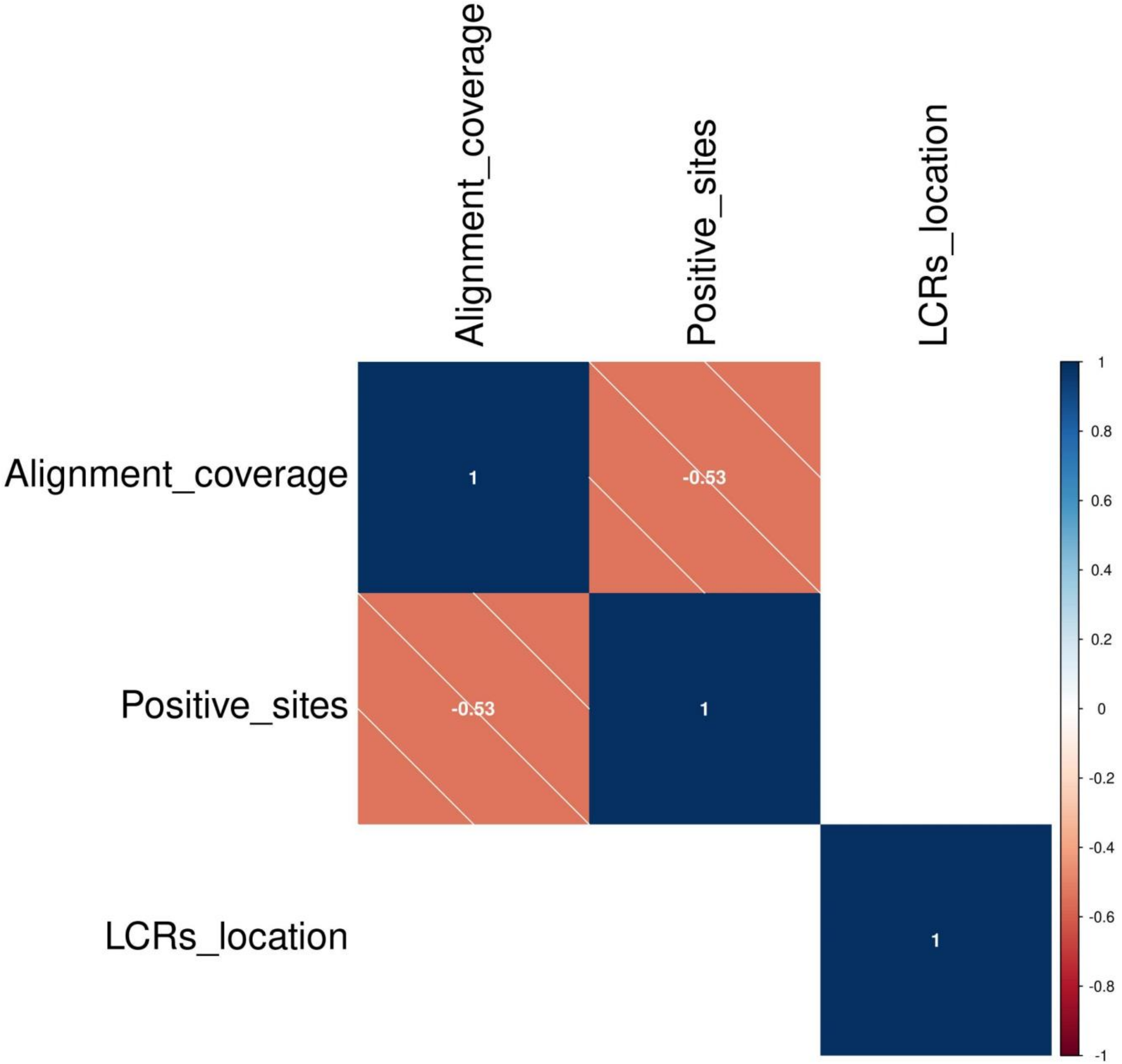

**Supplementary figure 495** Pairwise correlation of alignment coverage, PSSs and LCRs in Artiodactyla clade

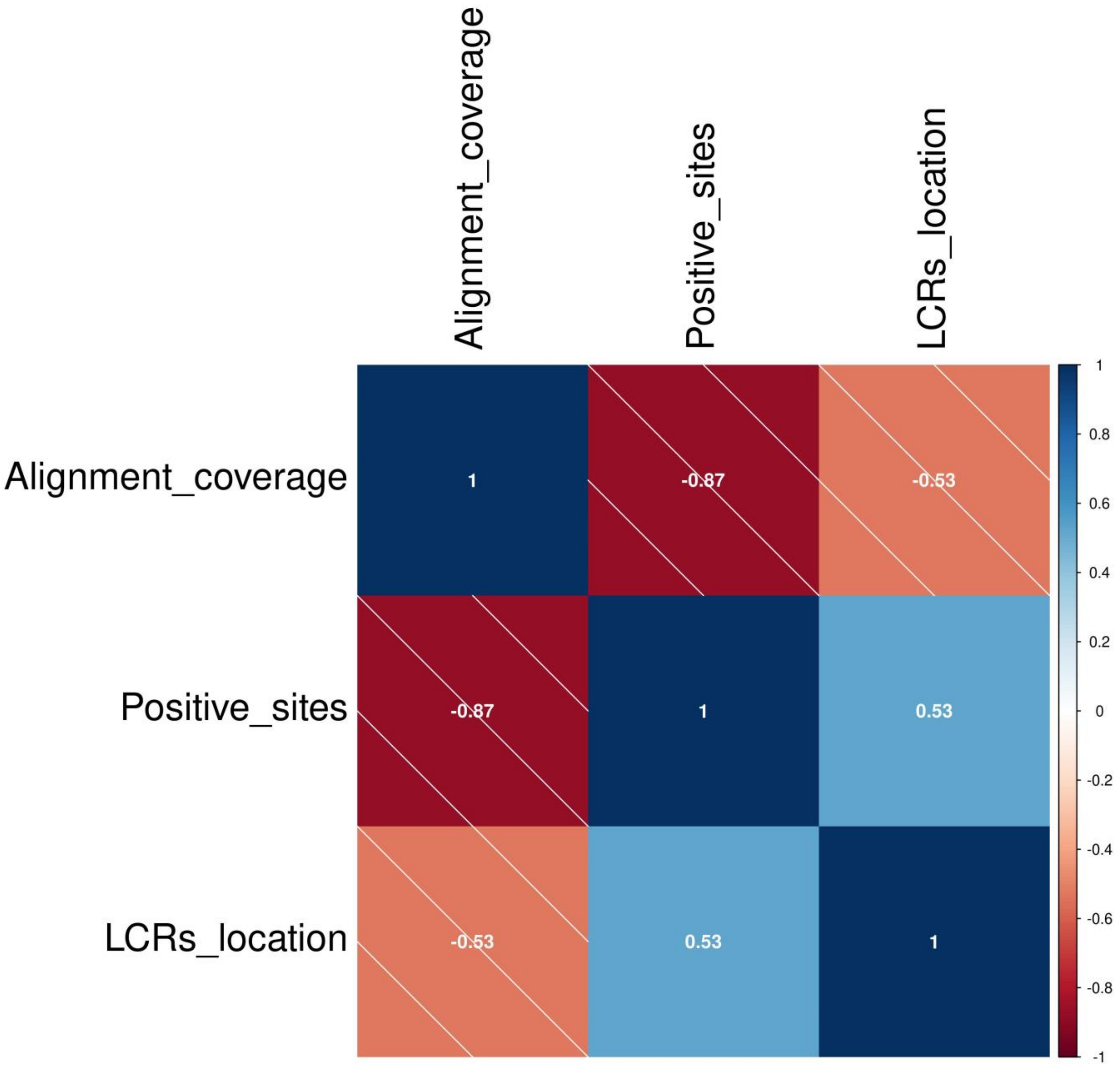

**Supplementary figure 496**

### Pairwise correlation of alignment coverage, PSSs and LCRs in Aves clade

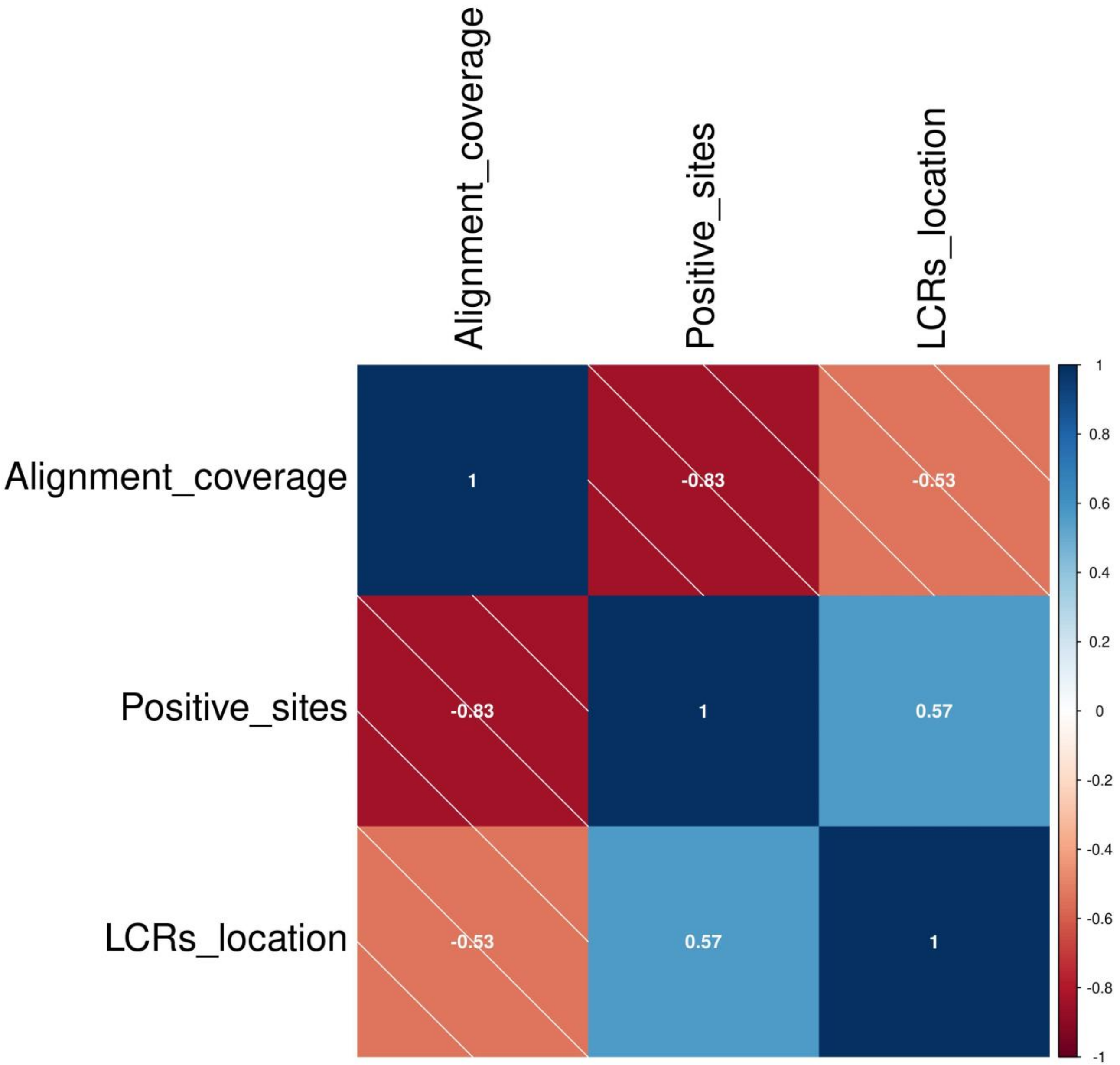

**Supplementary figure 497**

### Pairwise correlation of alignment coverage, PSSs and LCRs in Carnivora clade

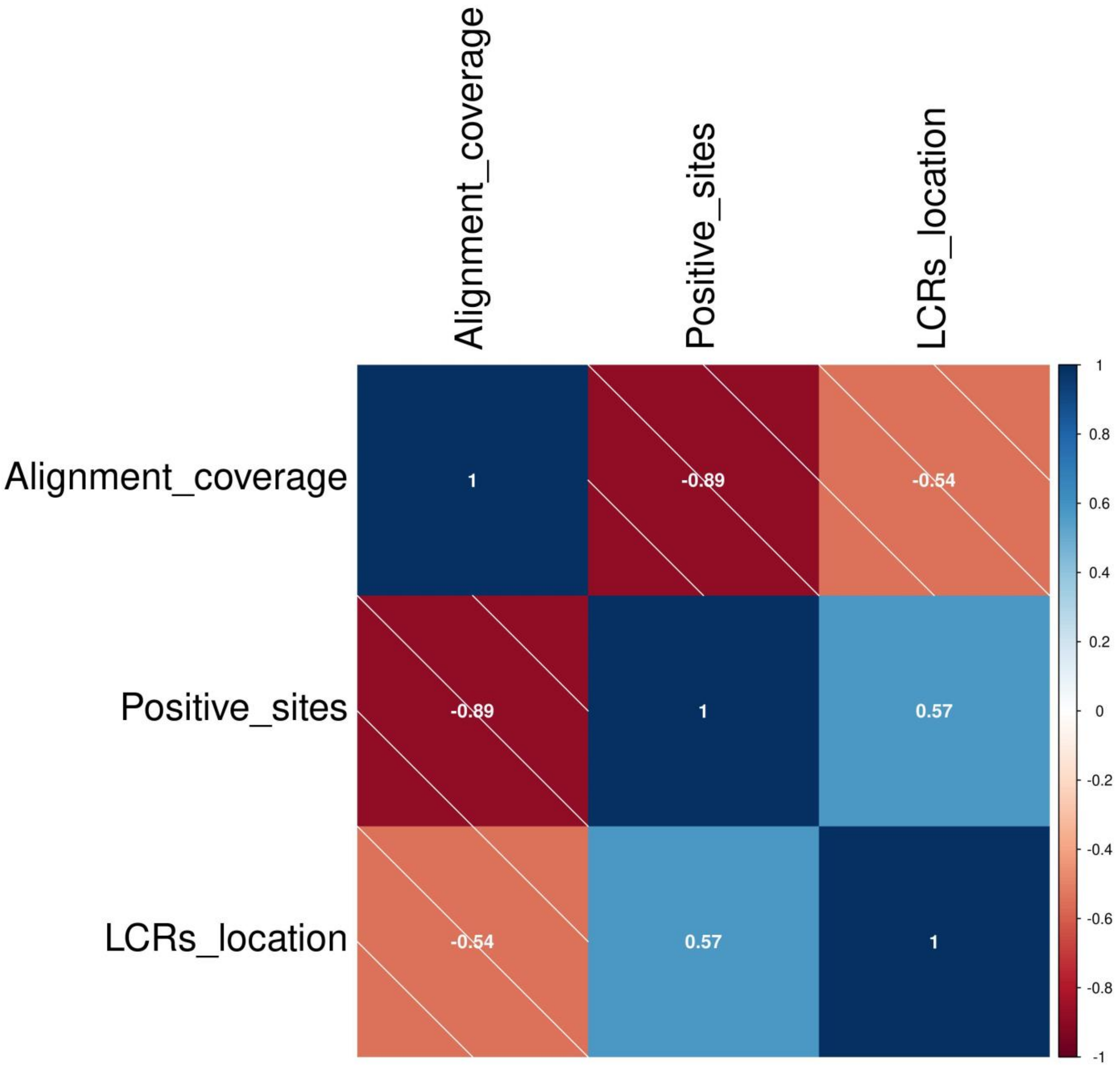

**Supplementary figure 498**

Pairwise correlation of alignment coverage, PSSs and LCRs in Chiroptera clade

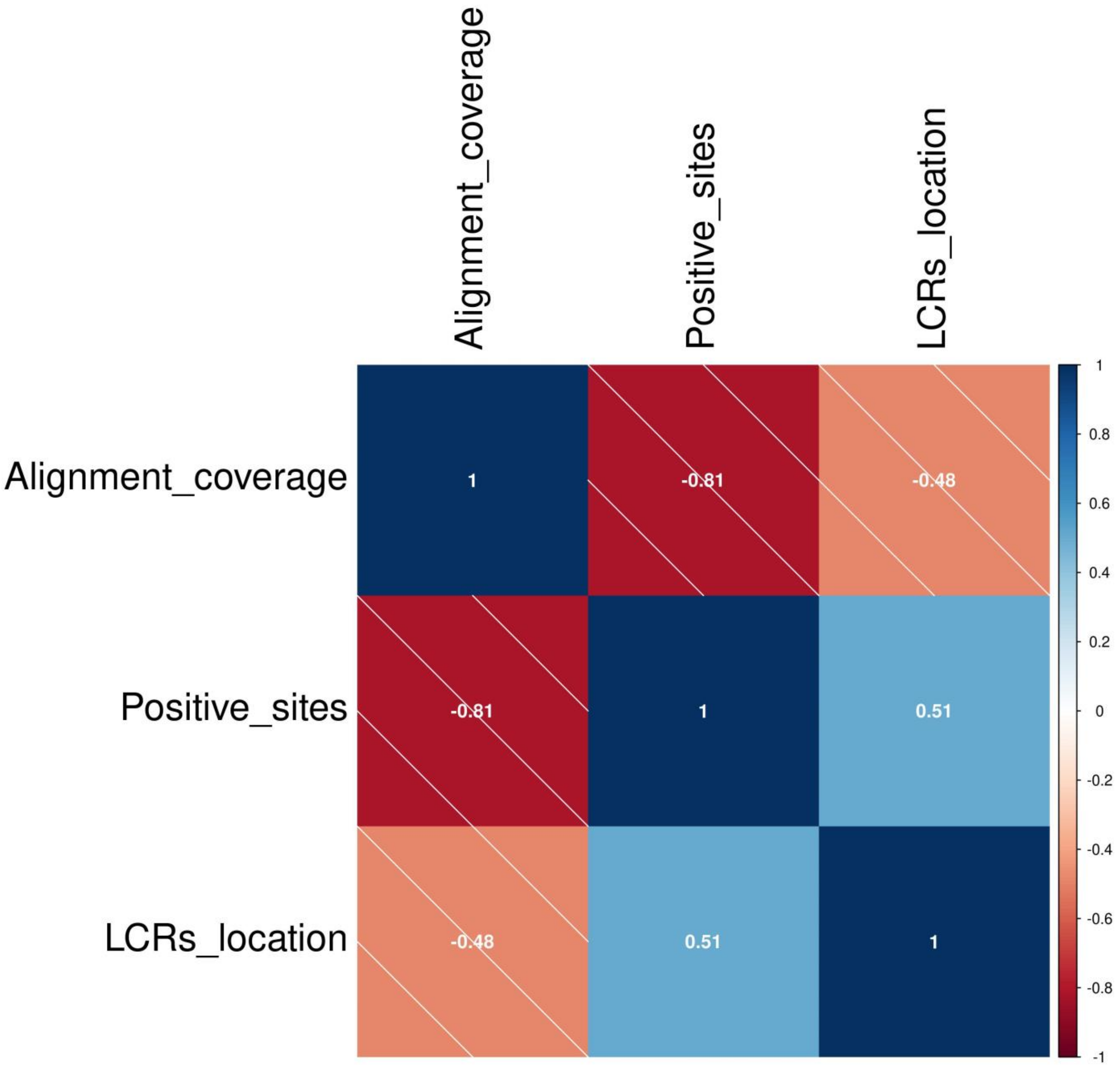

**Supplementary figure 499**

### Pairwise correlation of alignment coverage, PSSs and LCRs in Marsupialia clade

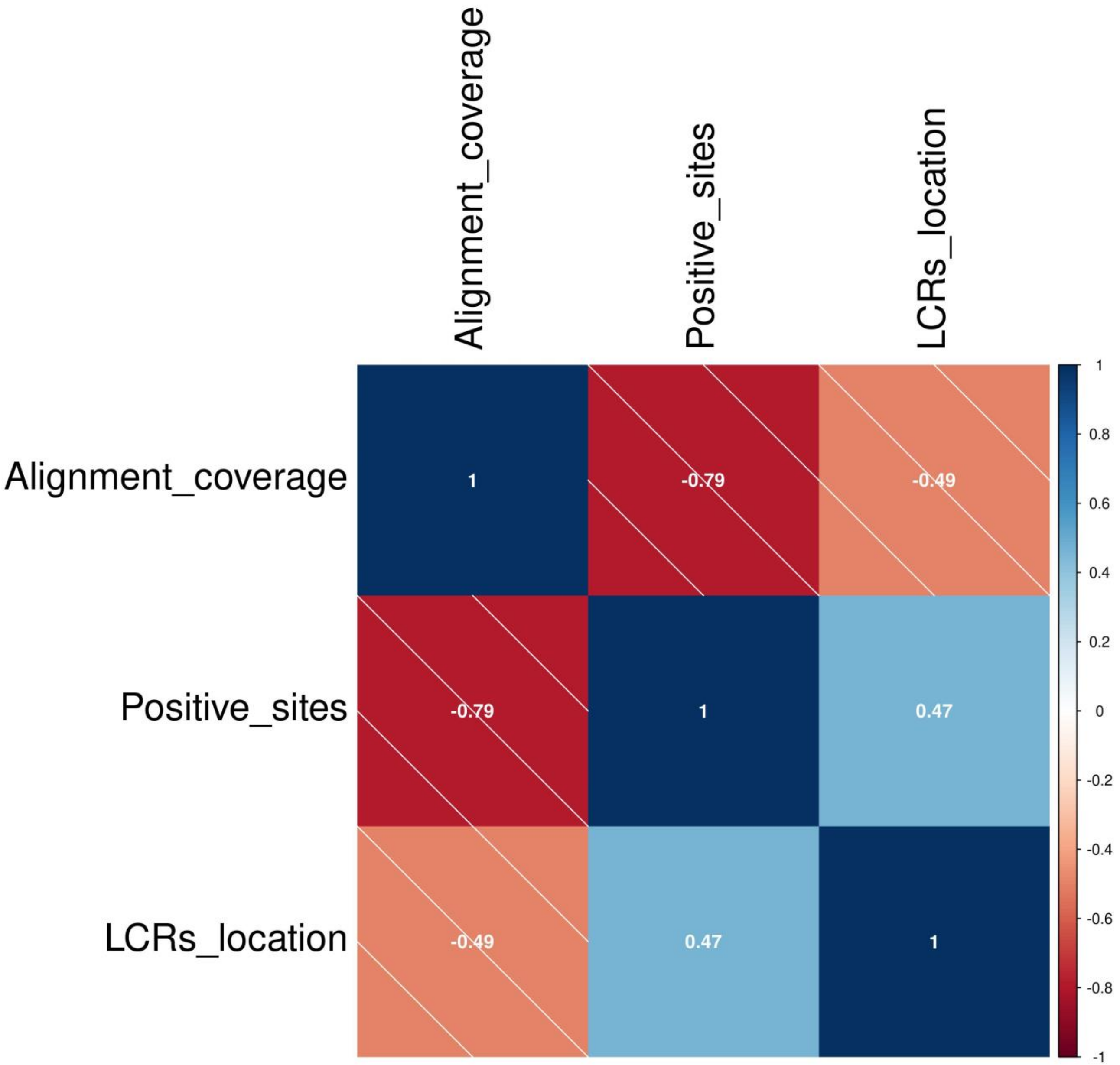

**Supplementary figure 500**

### Pairwise correlation of alignment coverage, PSSs and LCRs in Perissodactyla clade

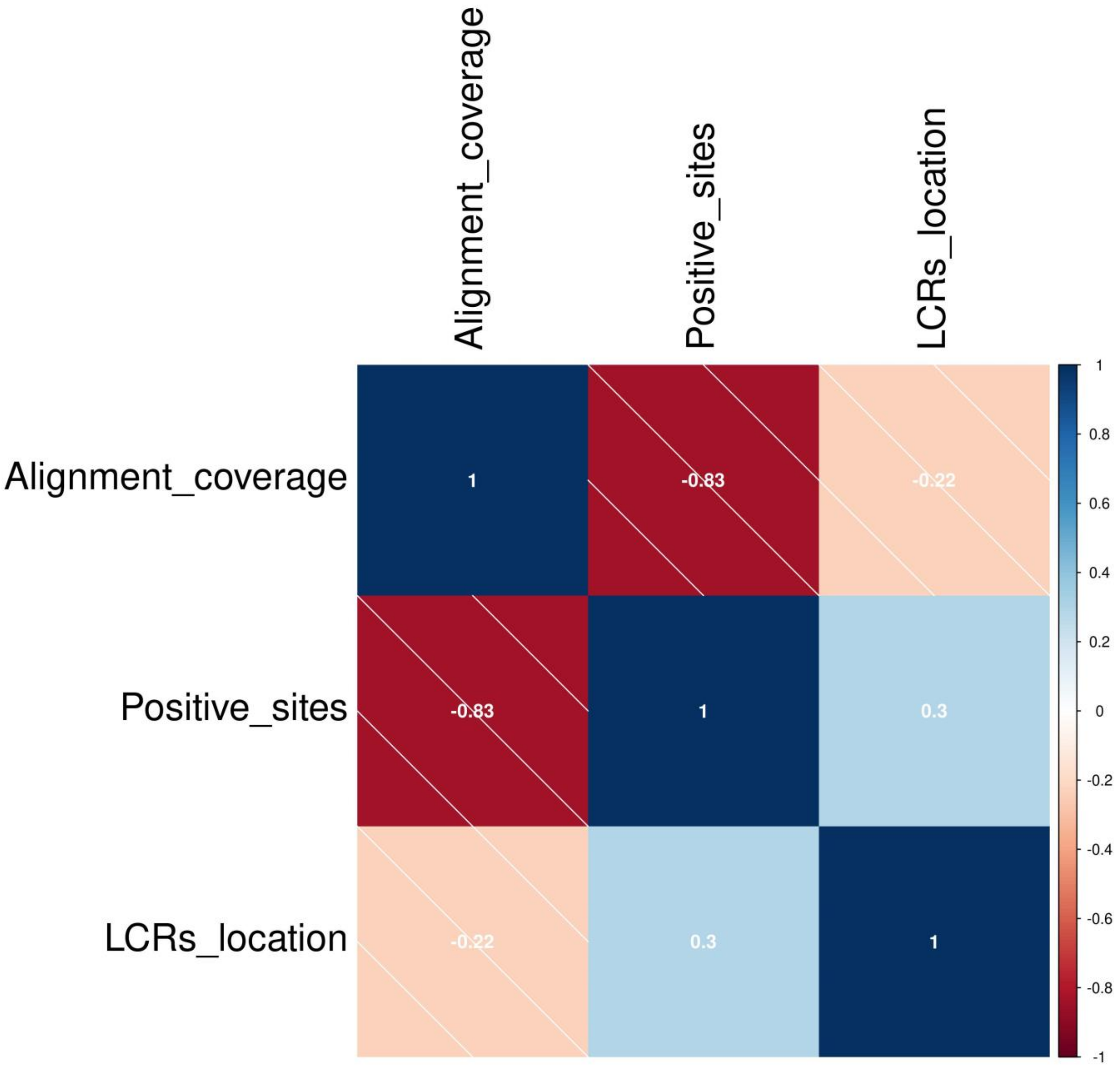

**Supplementary figure 501**

### Pairwise correlation of alignment coverage, PSSs and LCRs in Primates clade

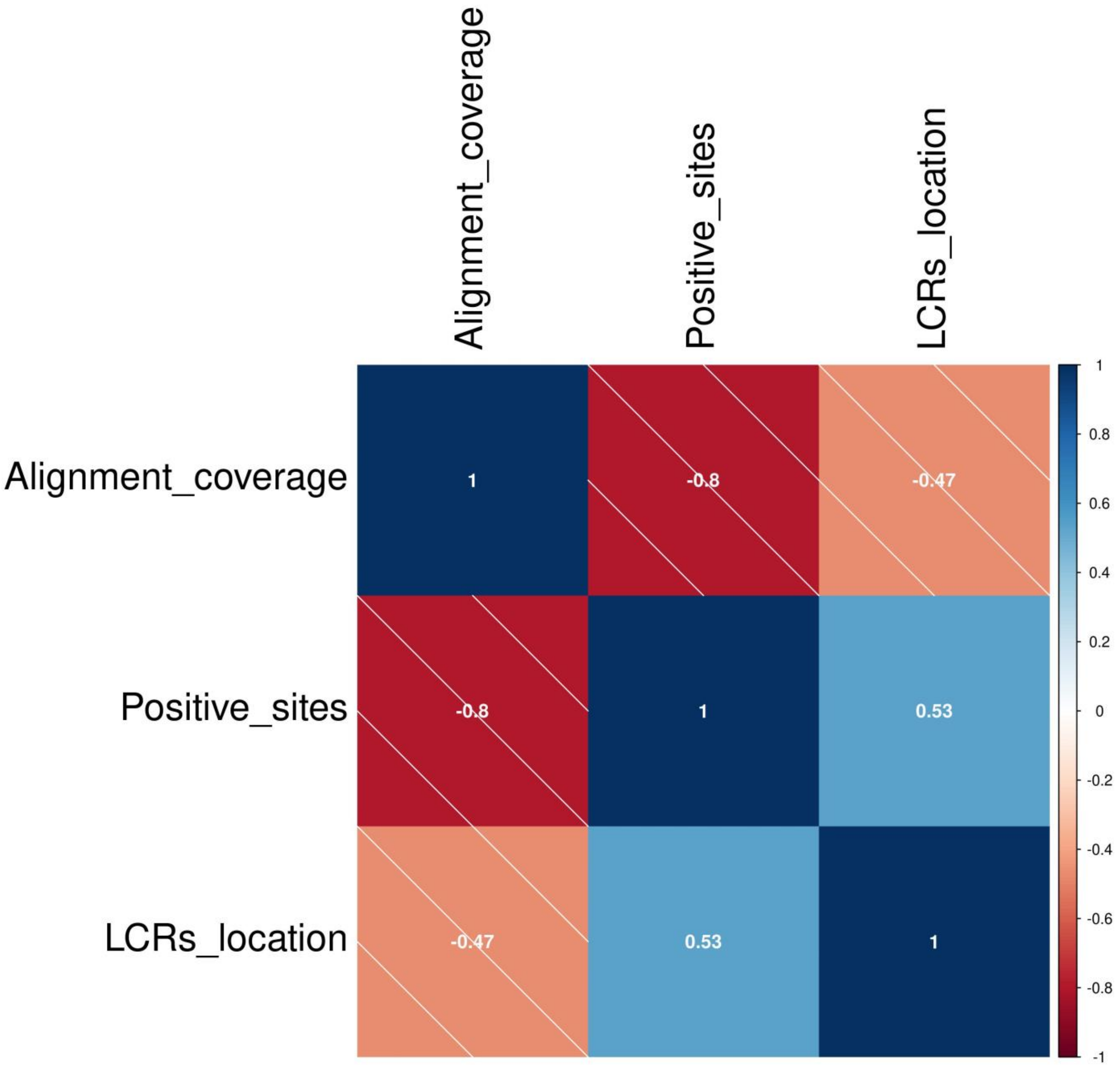

**Supplementary figure 502**

### Pairwise correlation of alignment coverage, PSSs and LCRs in Rodentia clade

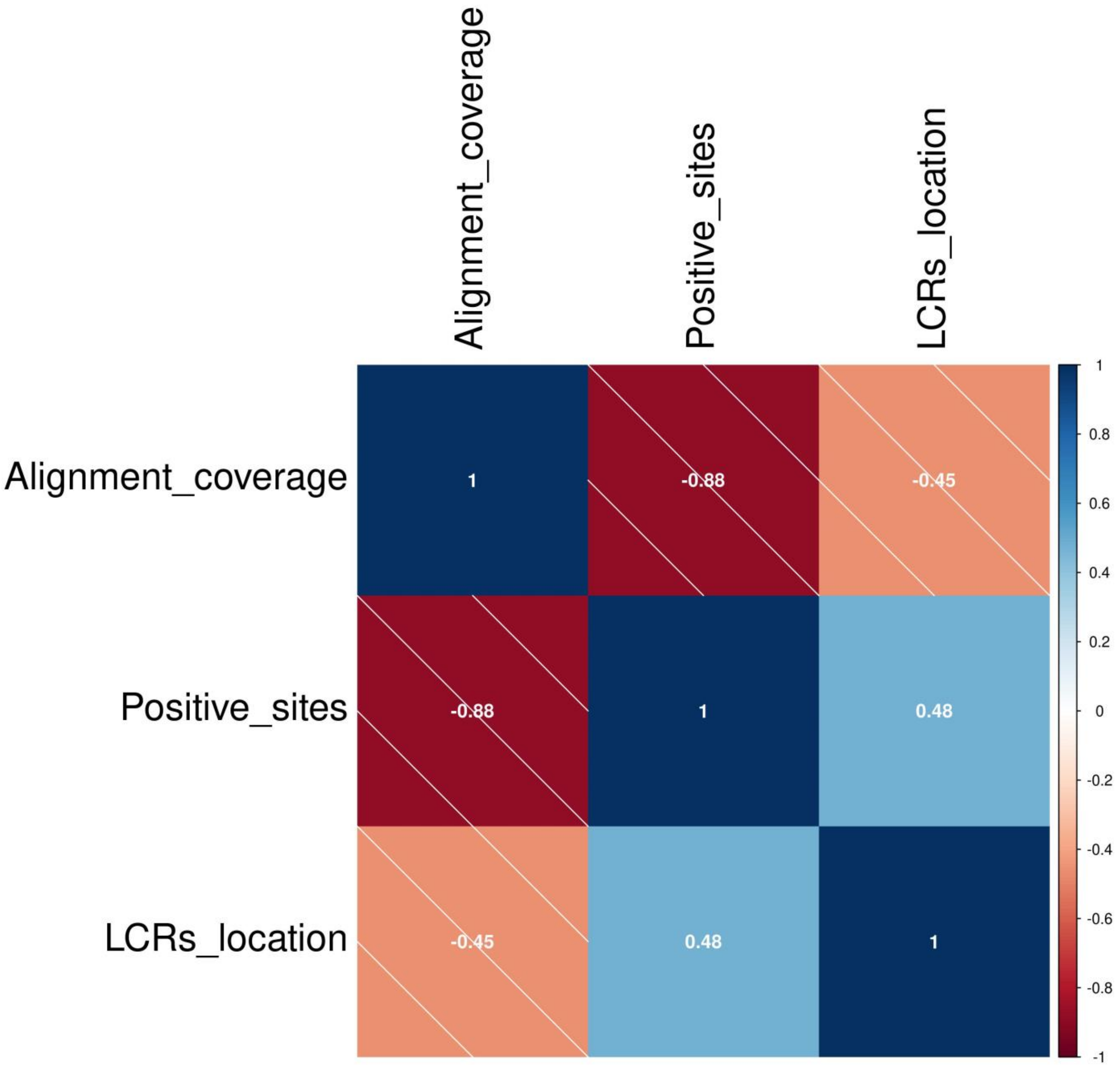

**Supplementary figure 503**

### Pairwise correlation of alignment coverage, PSSs and LCRs in Squamata clade

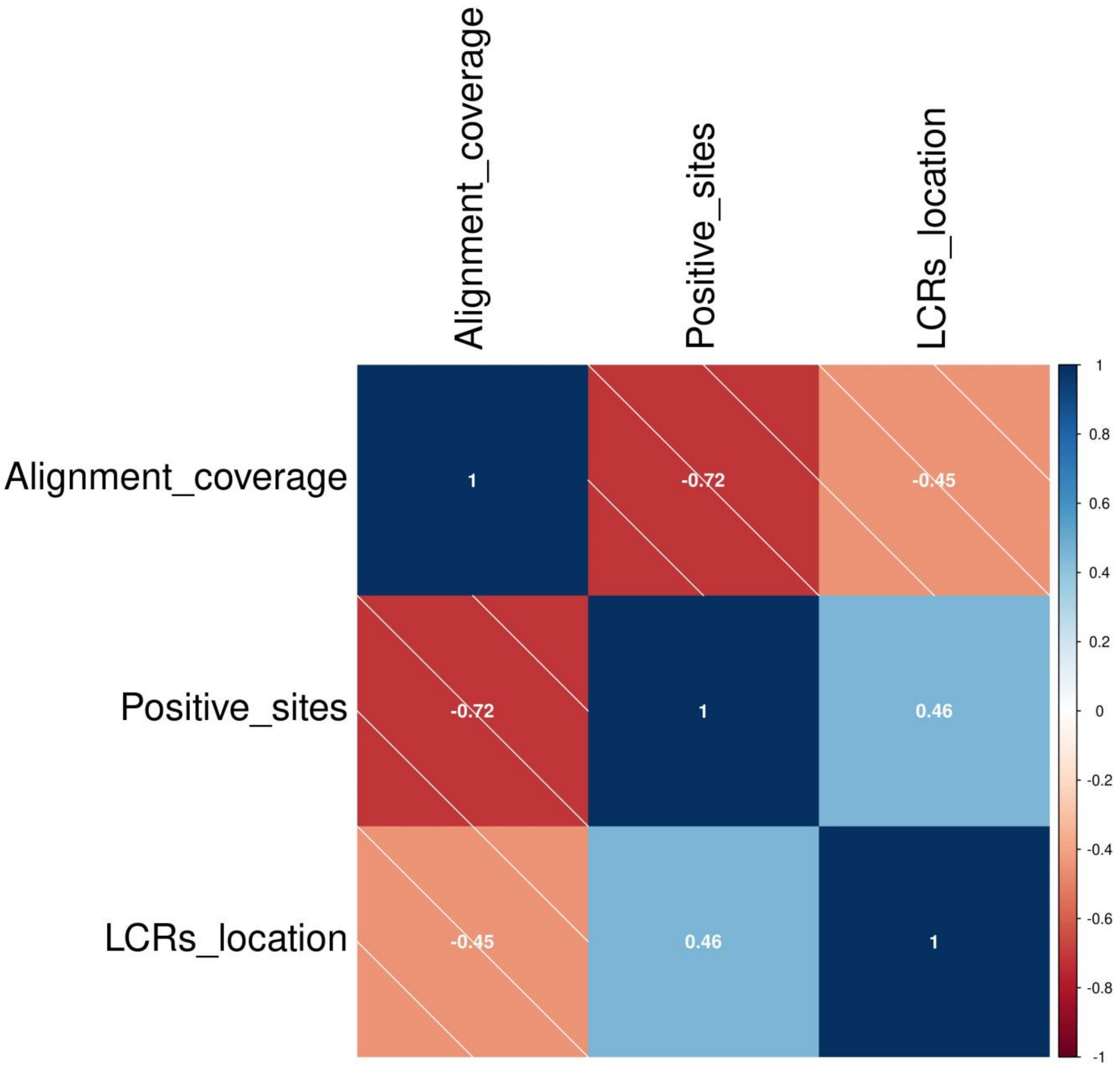

**Supplementary figure 504**

Pairwise correlation of alignment coverage, PSSs and LCRs in Testudines clade

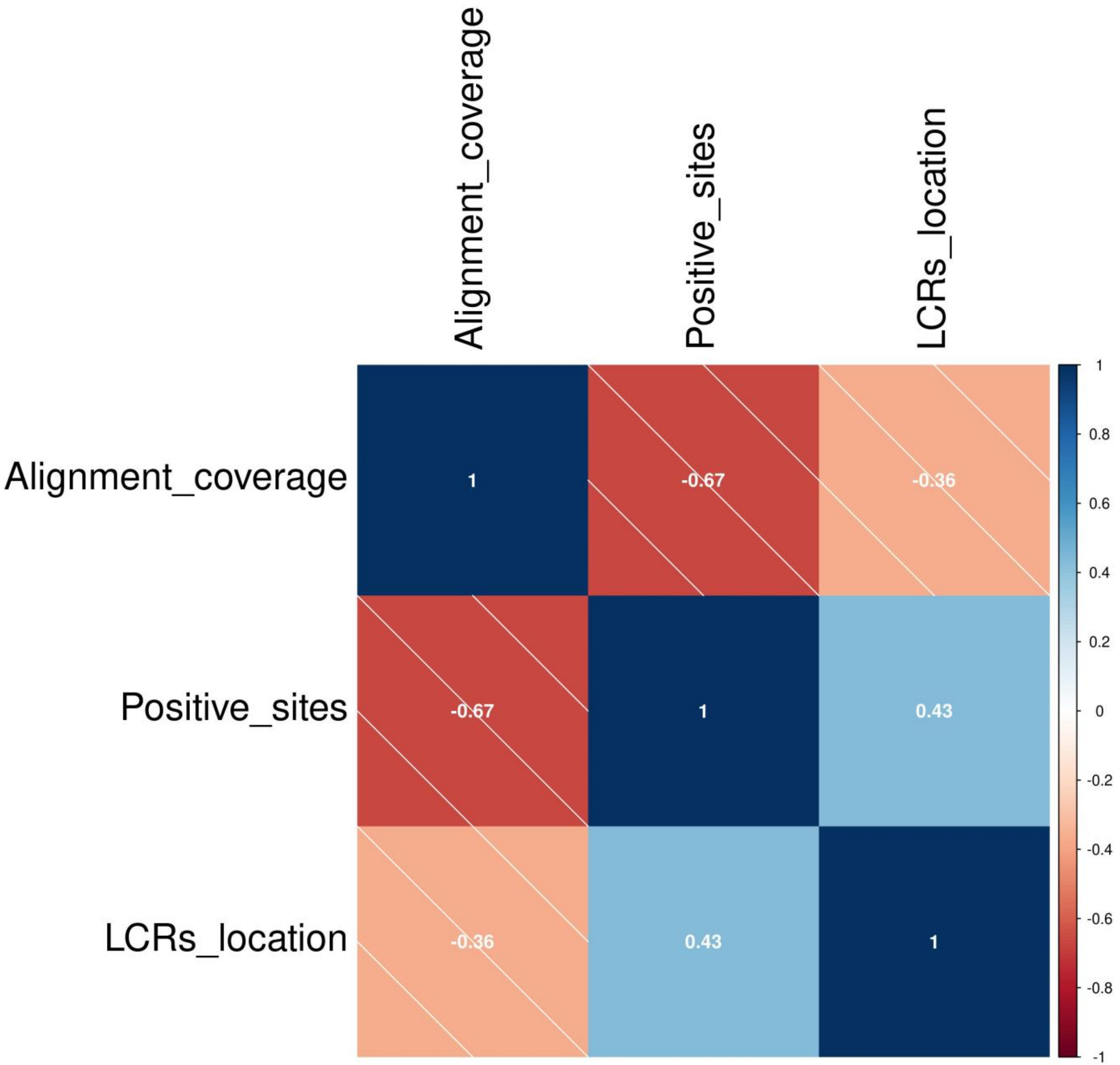

**Supplementary figure 505**

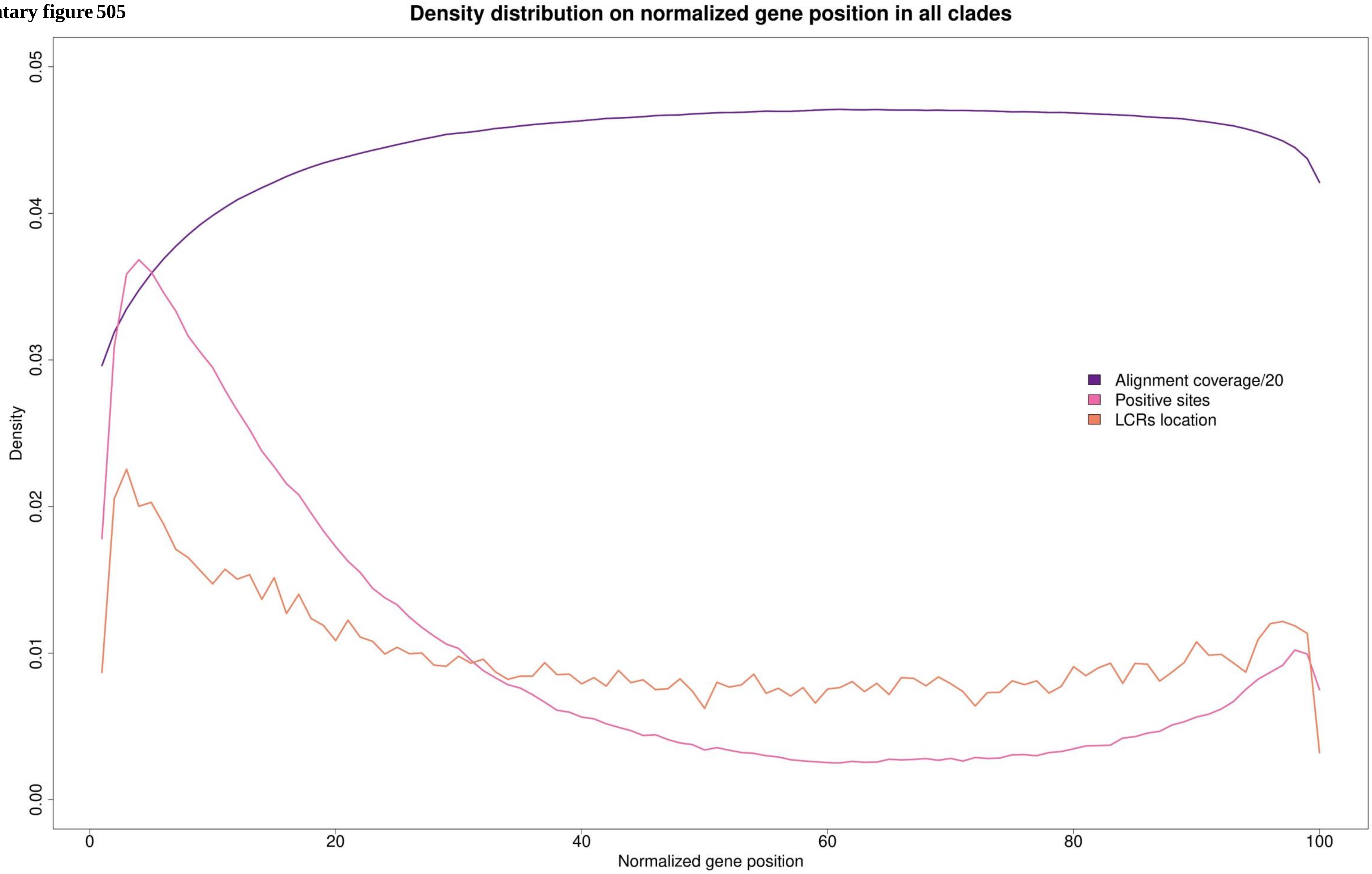

Supplementary figure 506

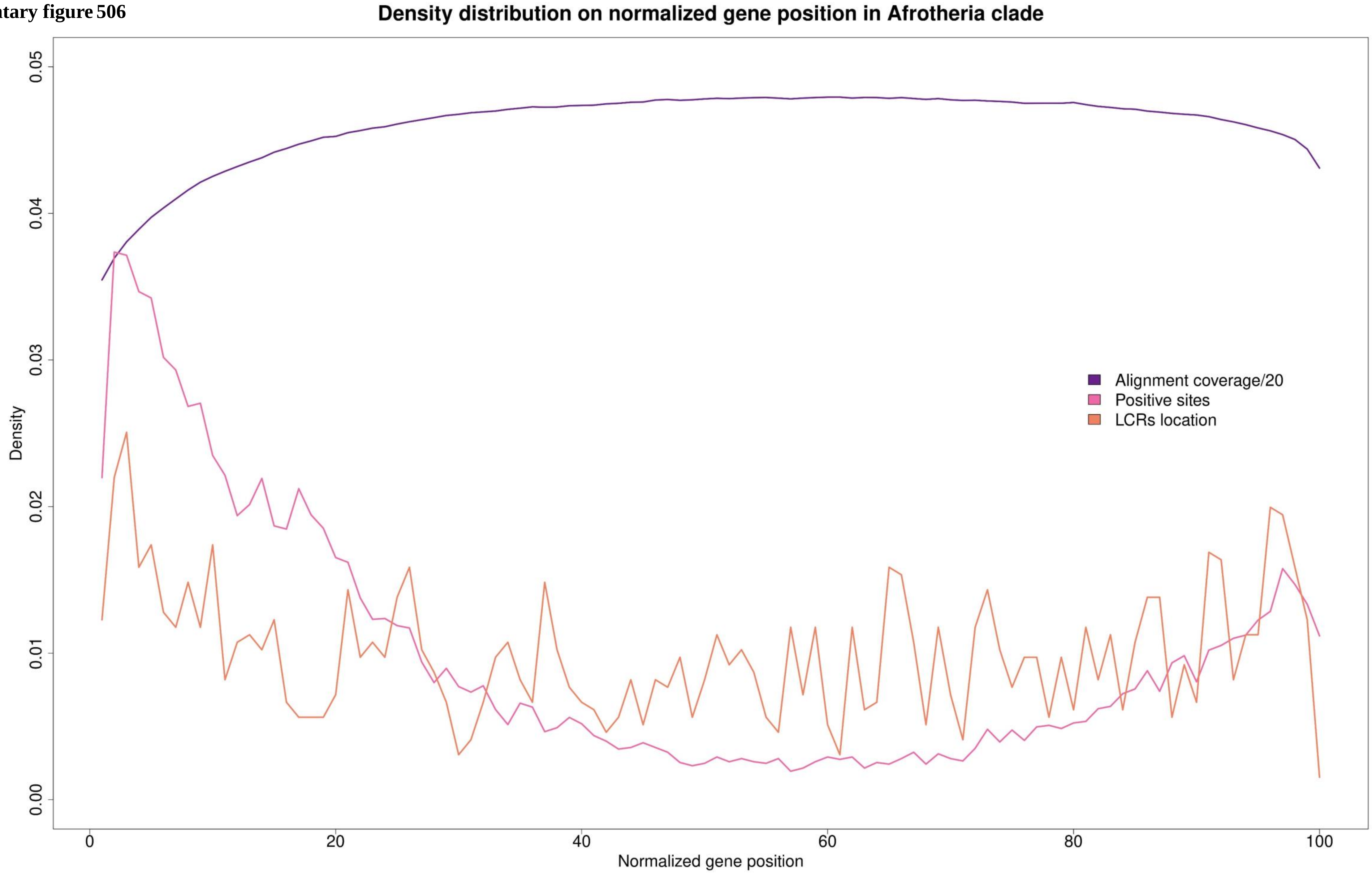

Supplementary figure 507

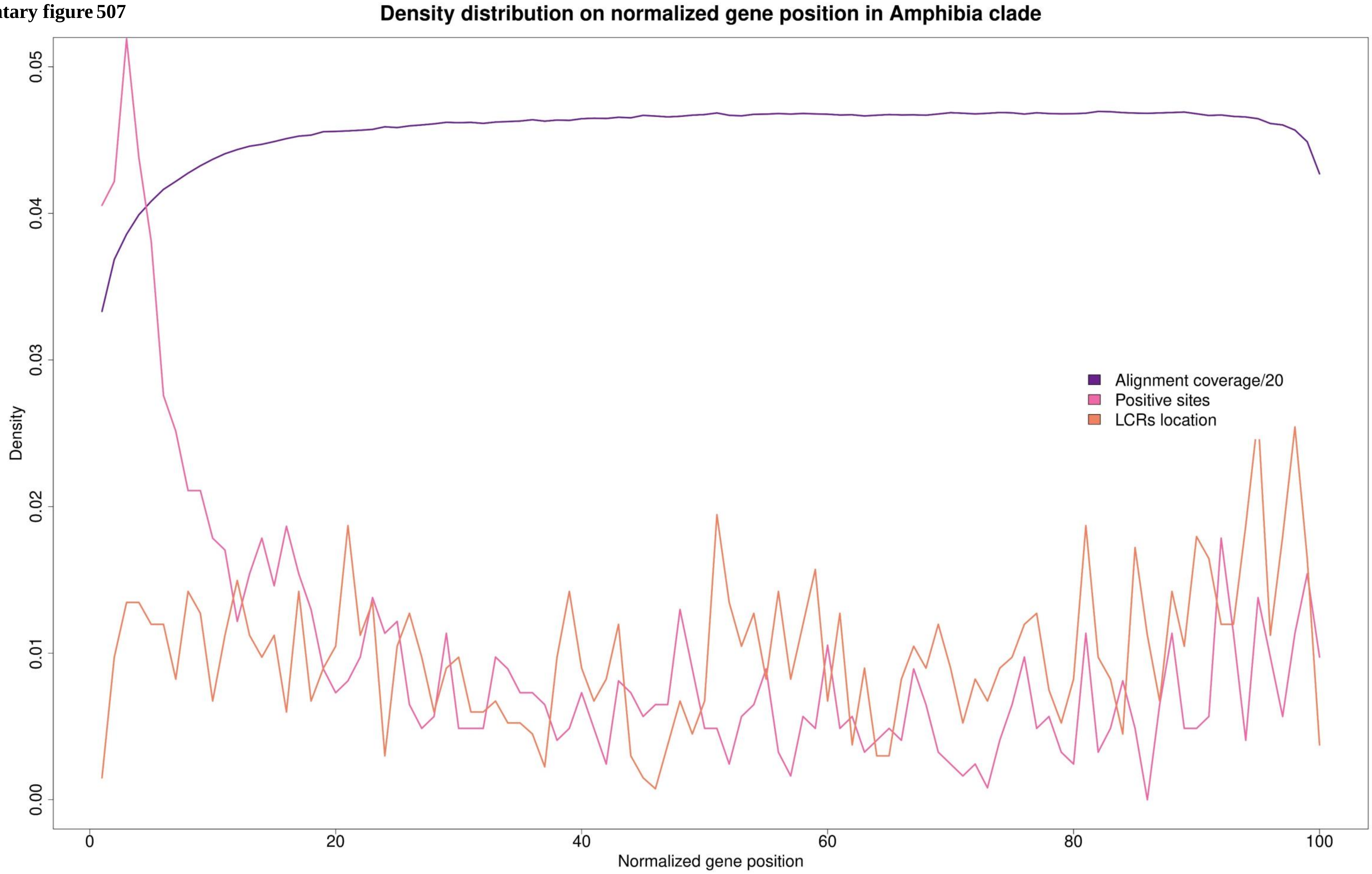

Supplementary figure 508

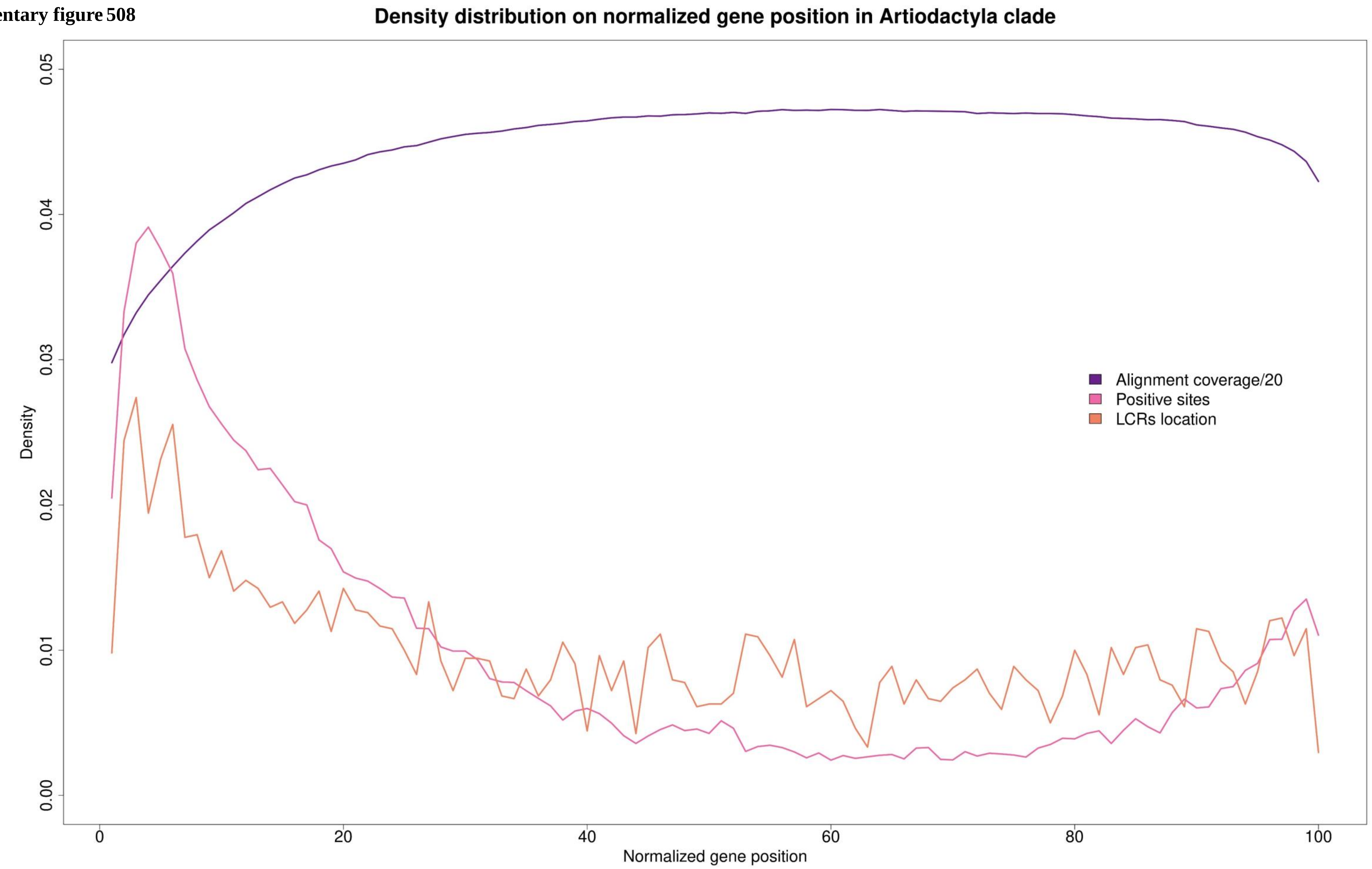

Supplementary figure 509

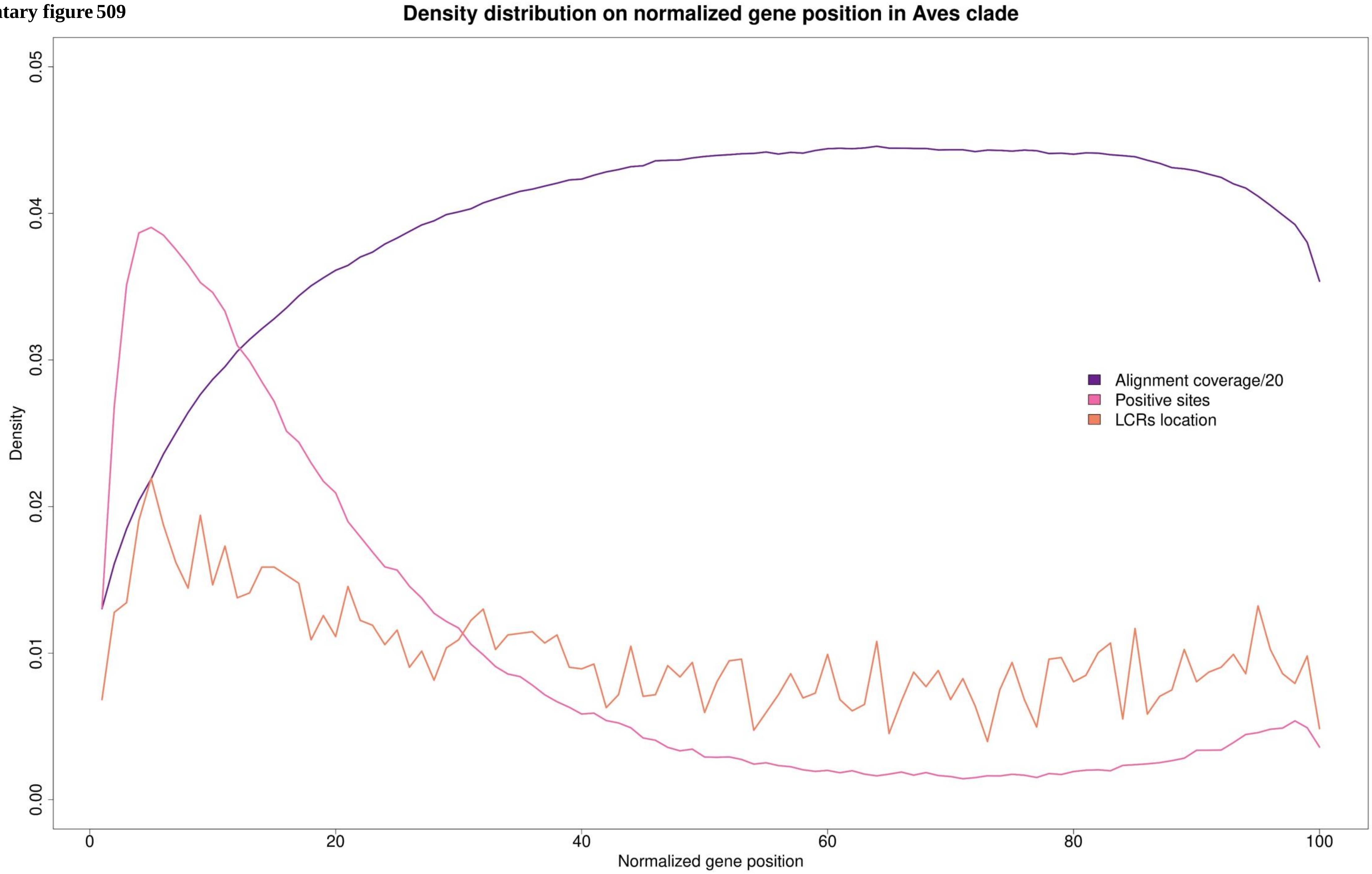

Supplementary figure 510

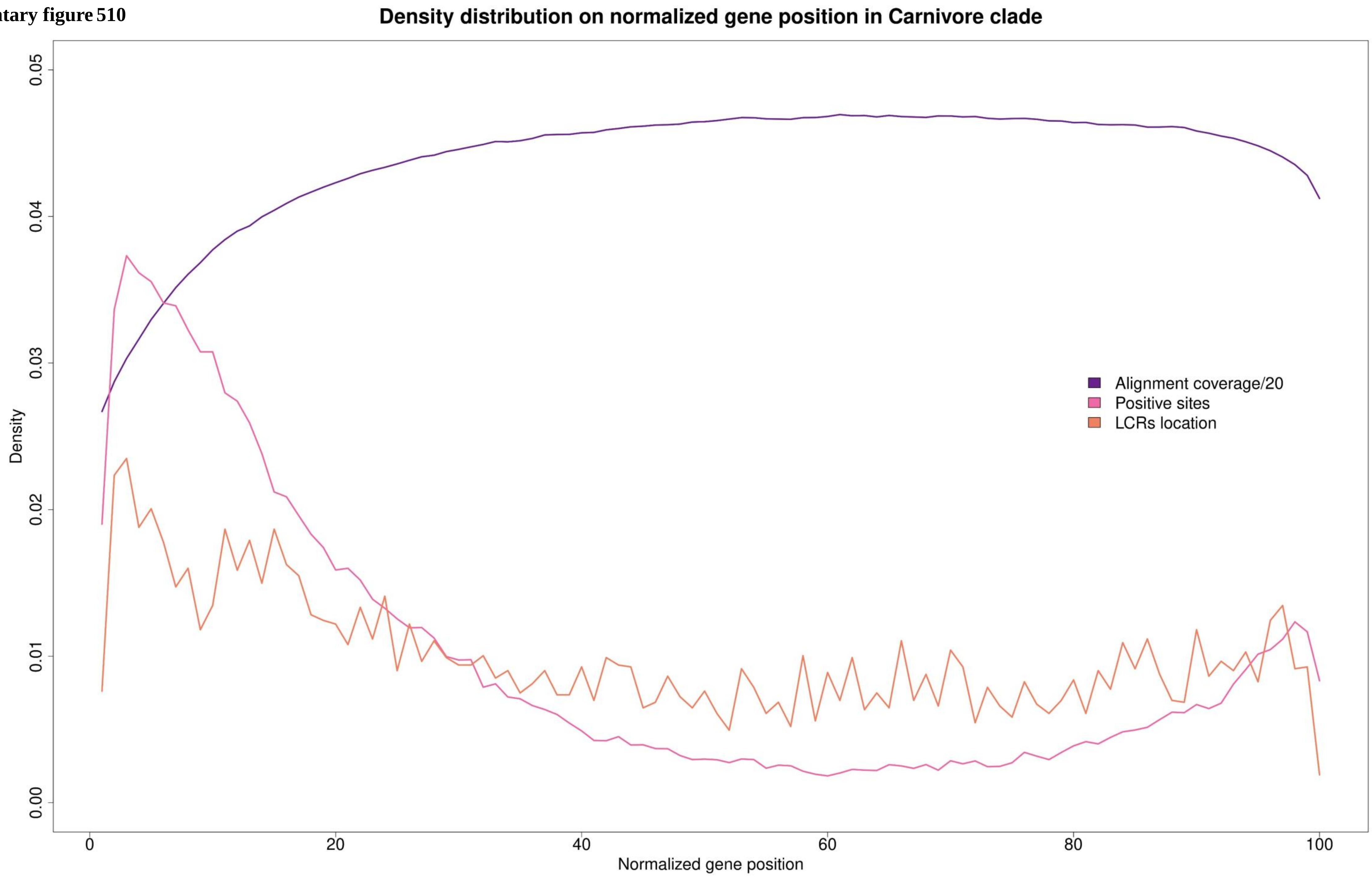

**Supplementary figure 511**

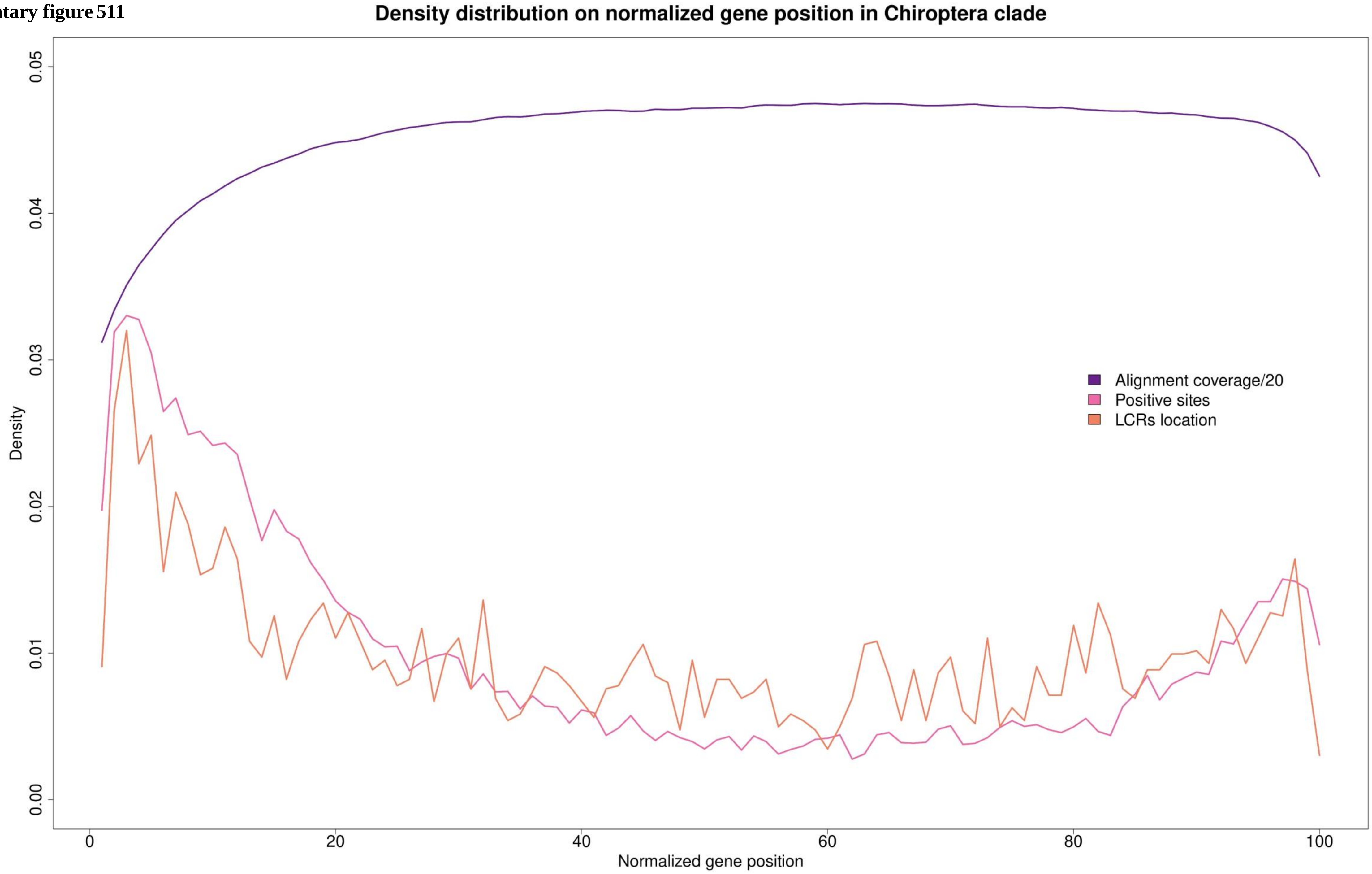

**Supplementary figure 512**

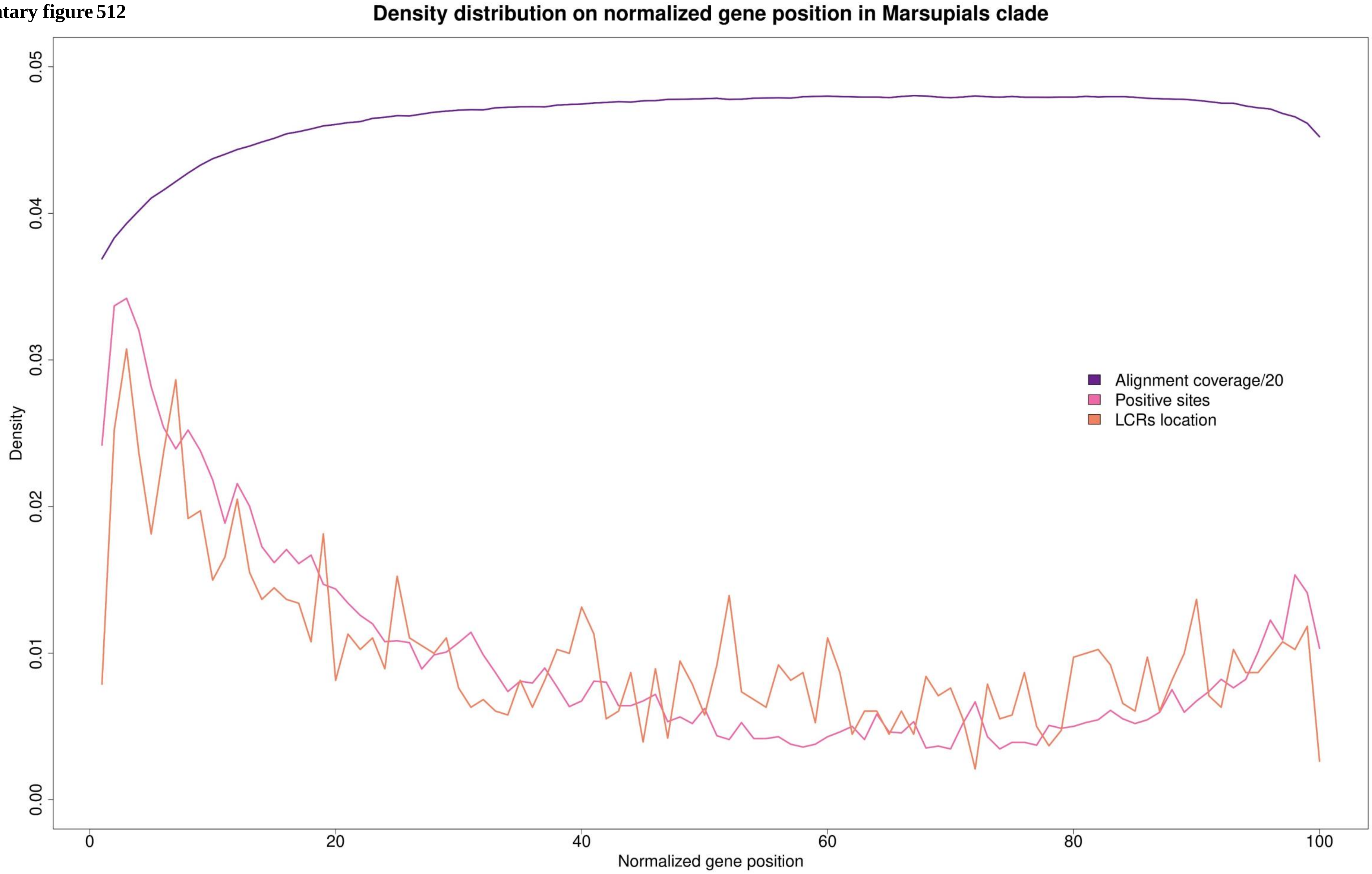

**Supplementary figure 513**

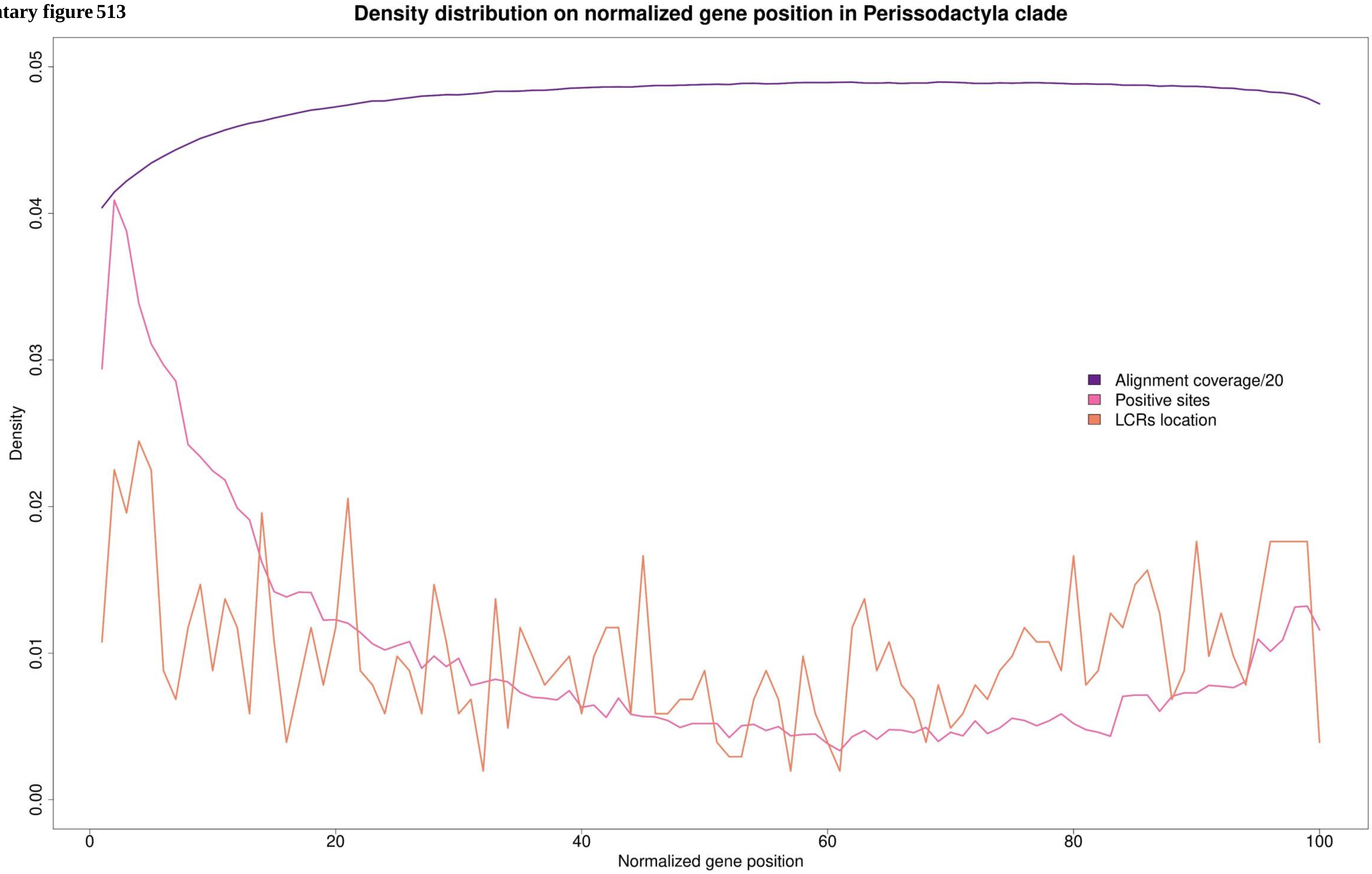

Supplementary figure 514

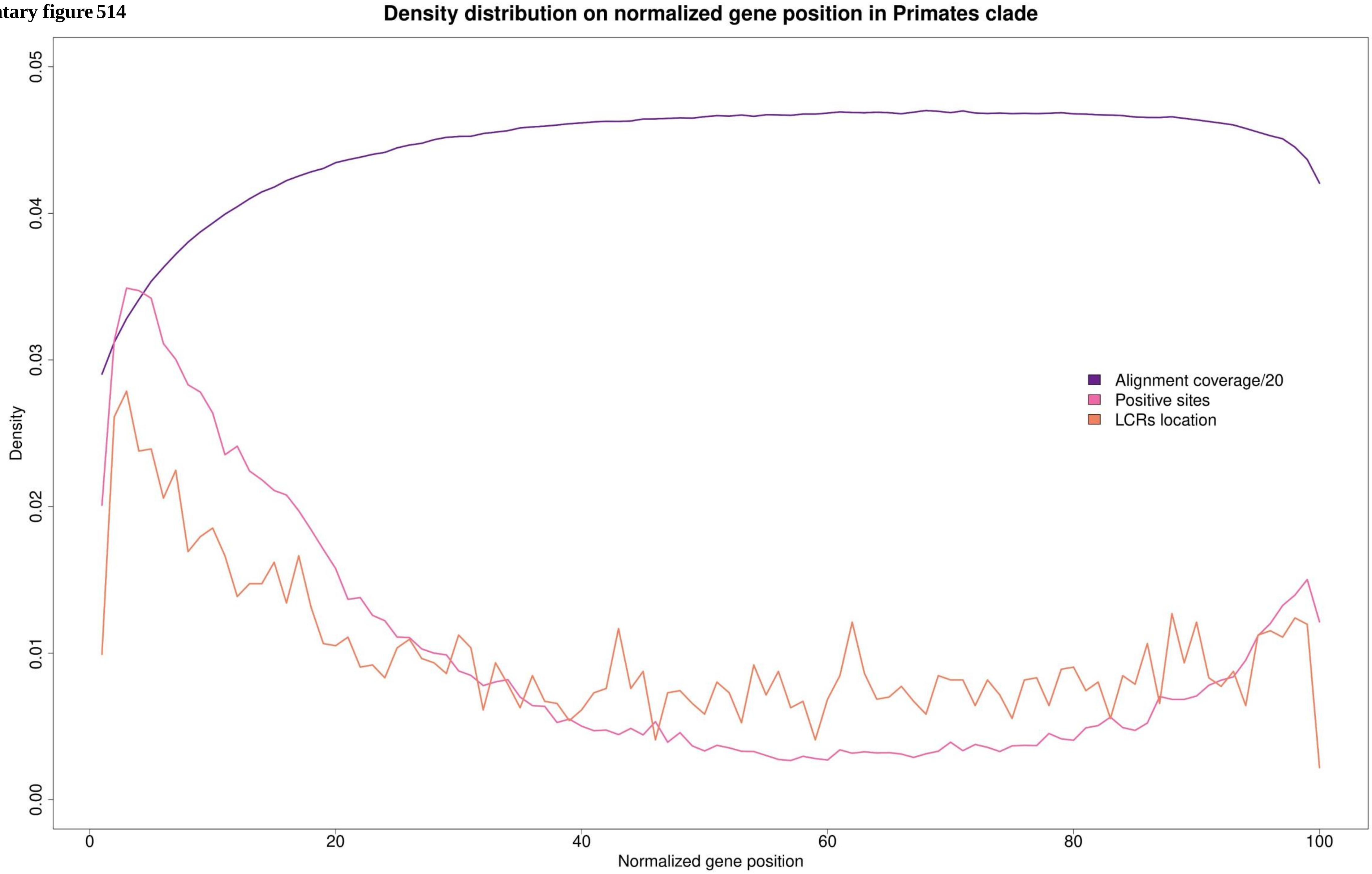

**Supplementary figure 515**

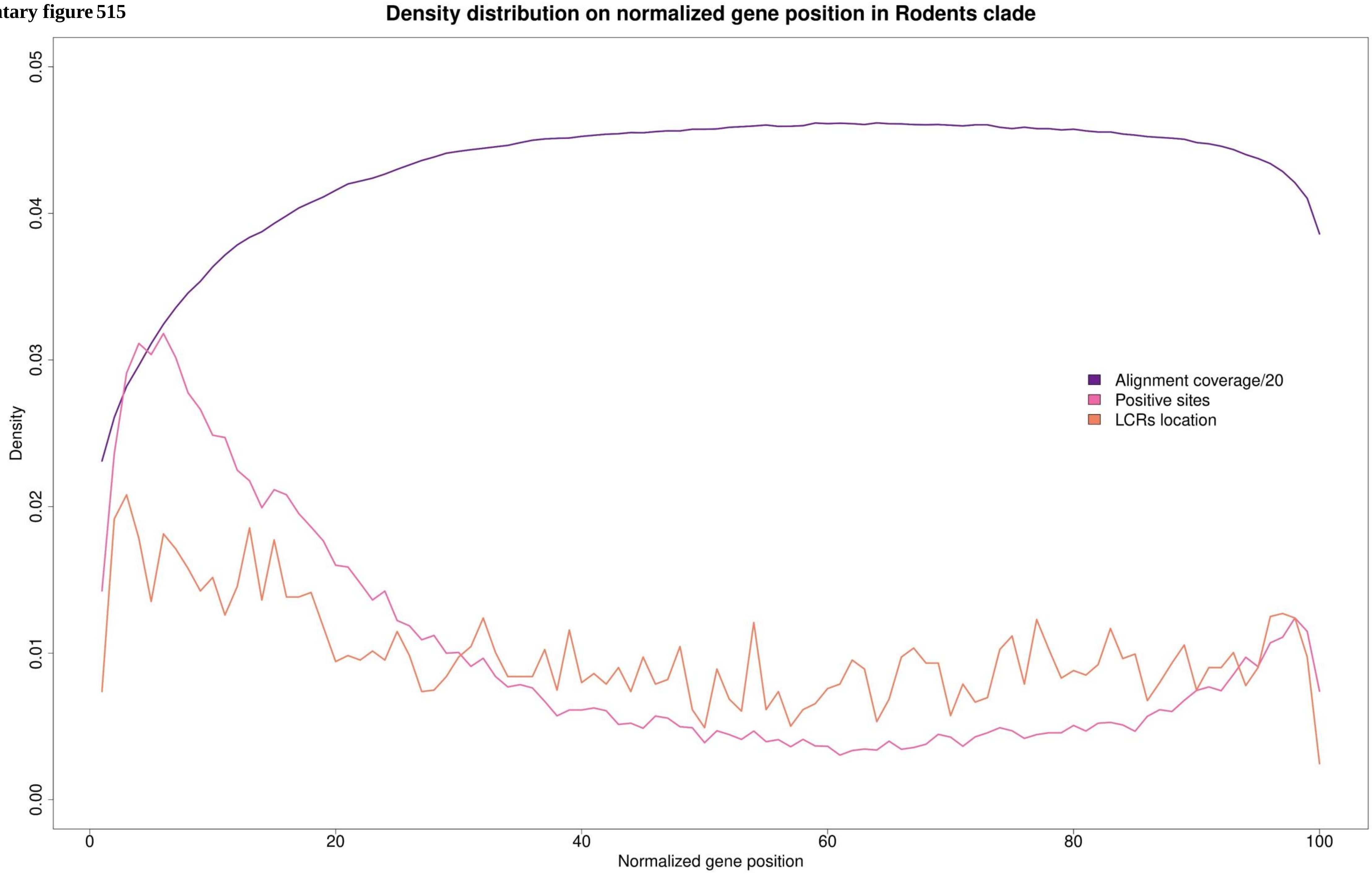

Supplementary figure 516

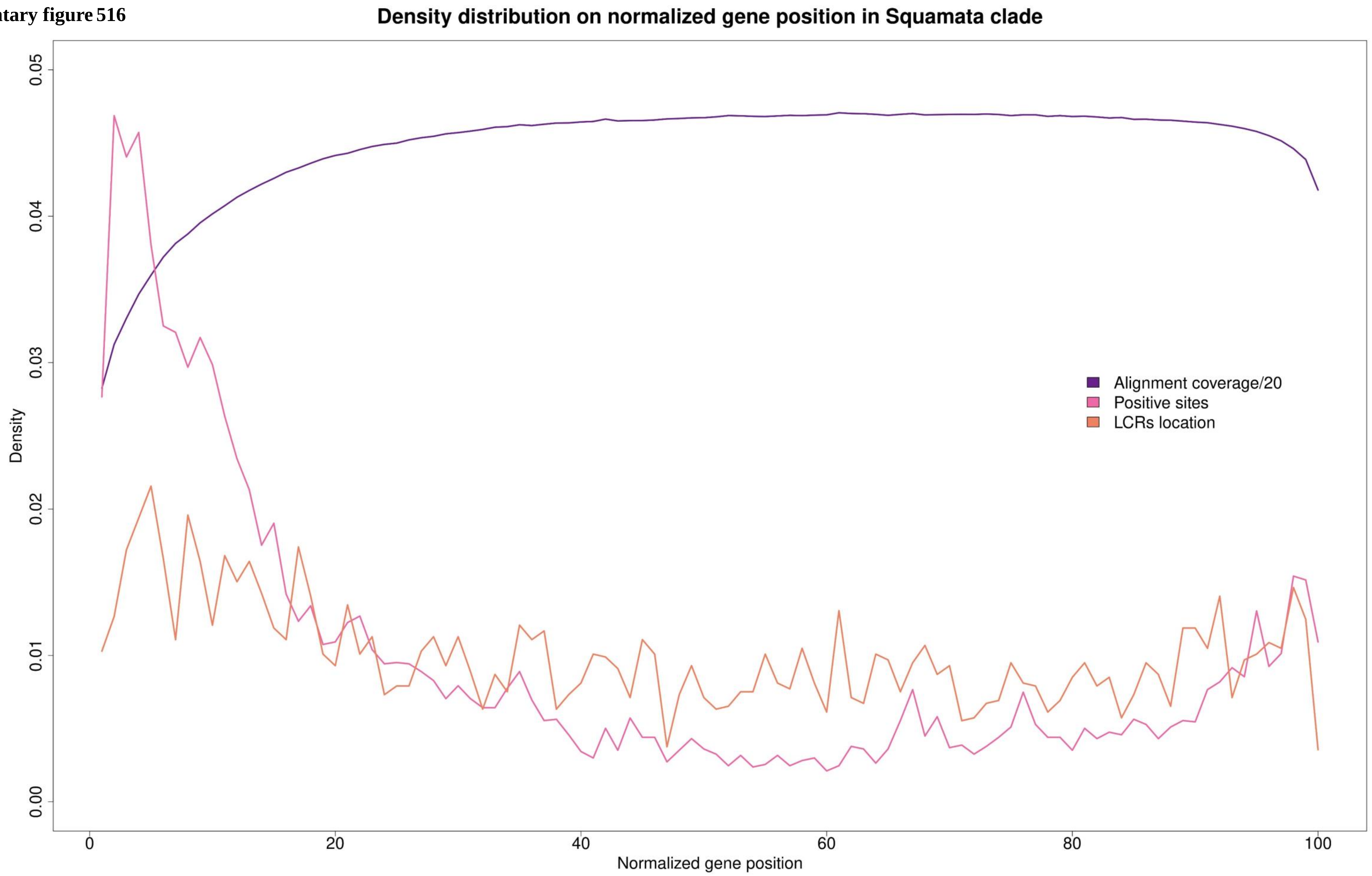

**Supplementary figure 517**

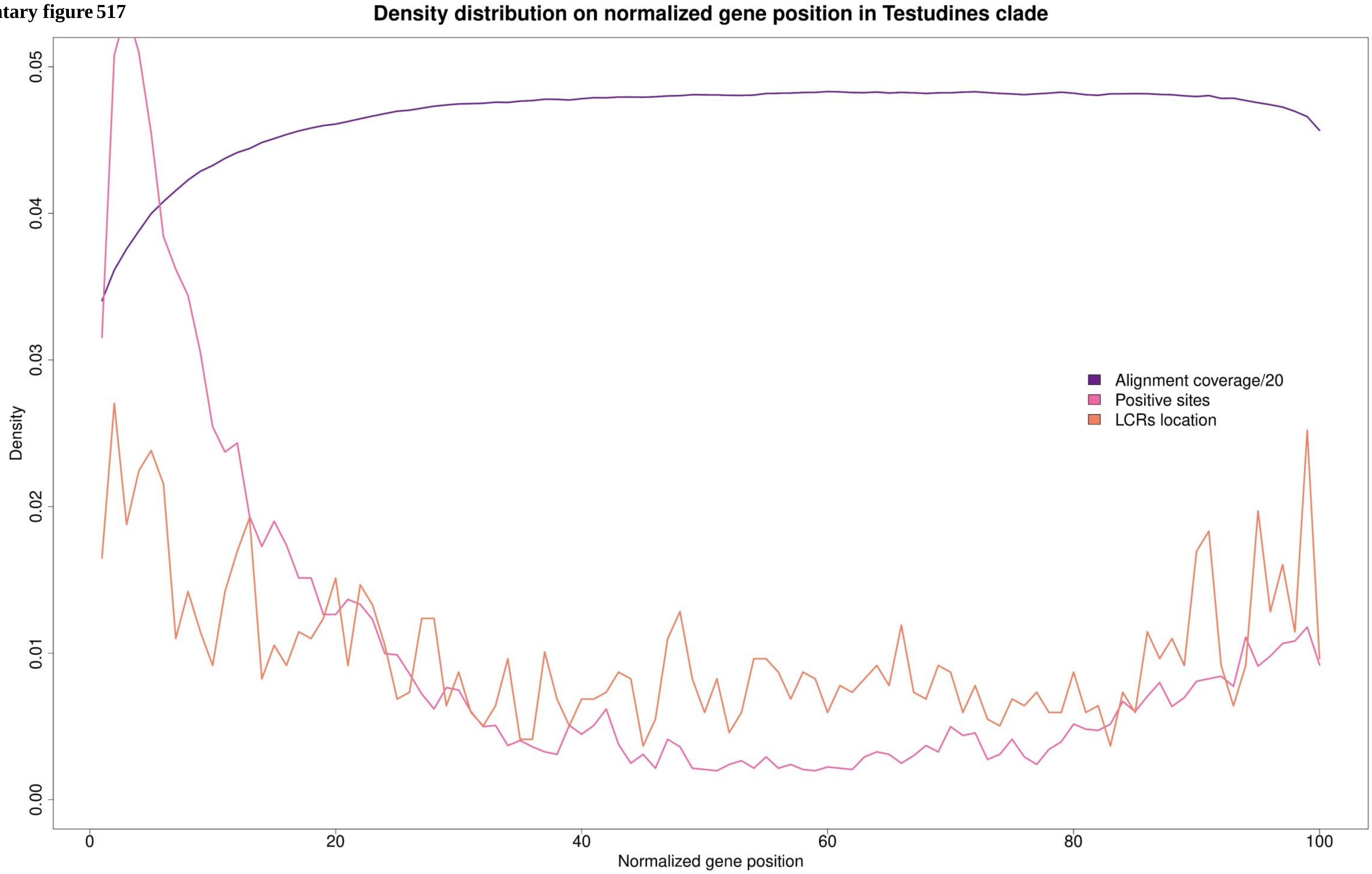

**Supplementary figure 518**

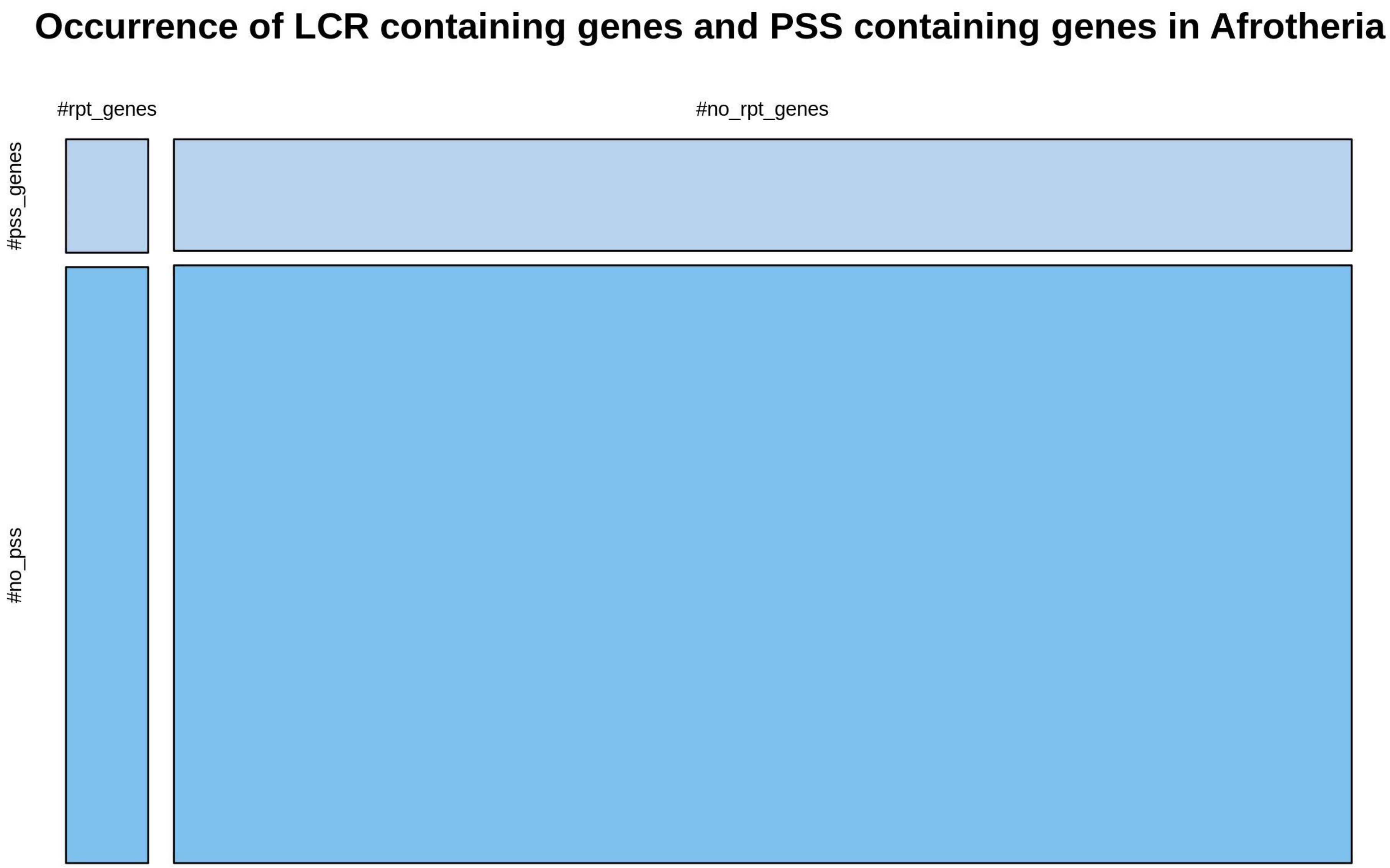

**Supplementary figure 519**

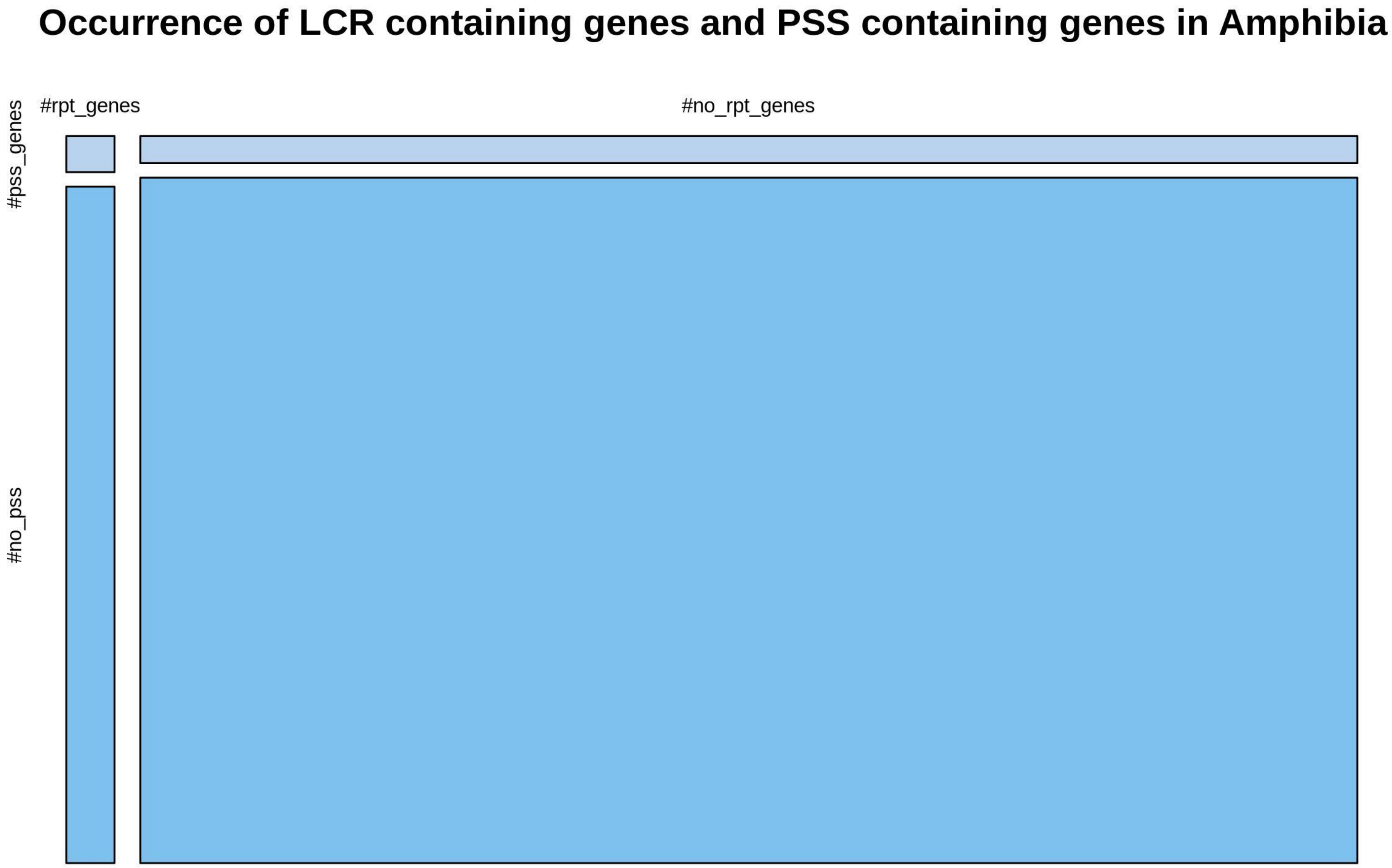

**Supplementary figure 520**

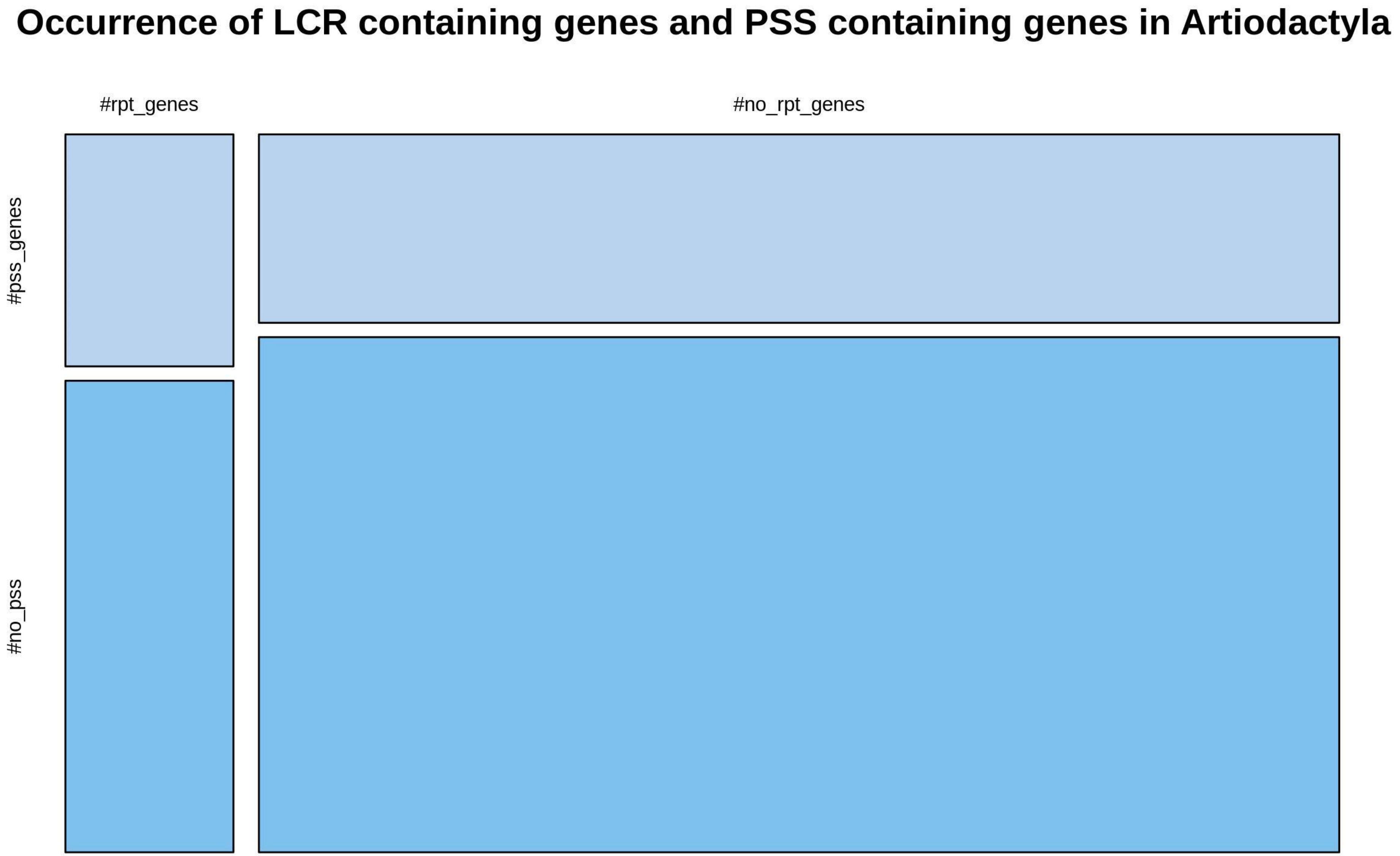

**Supplementary figure 521**

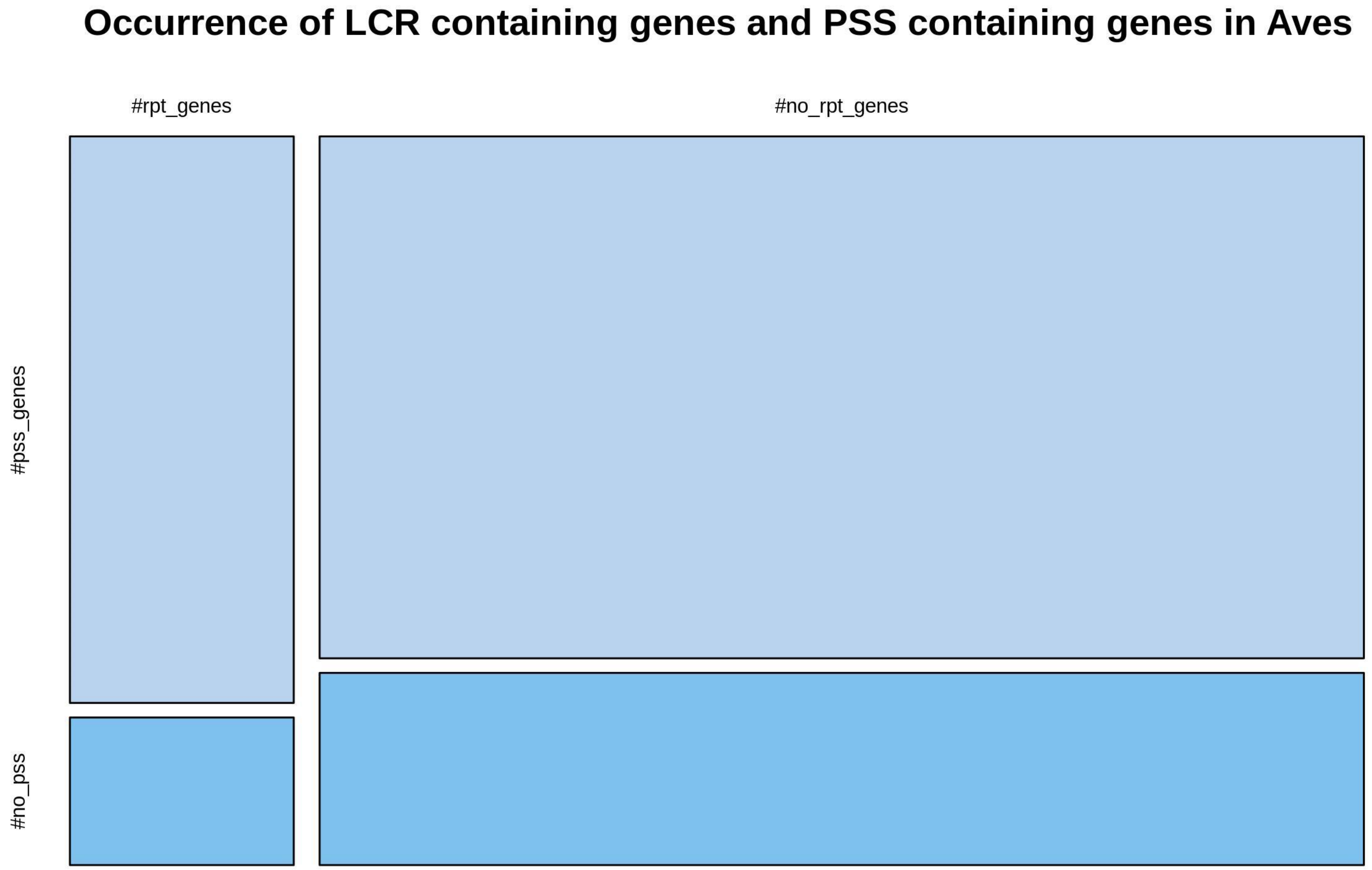

**Supplementary figure 522**

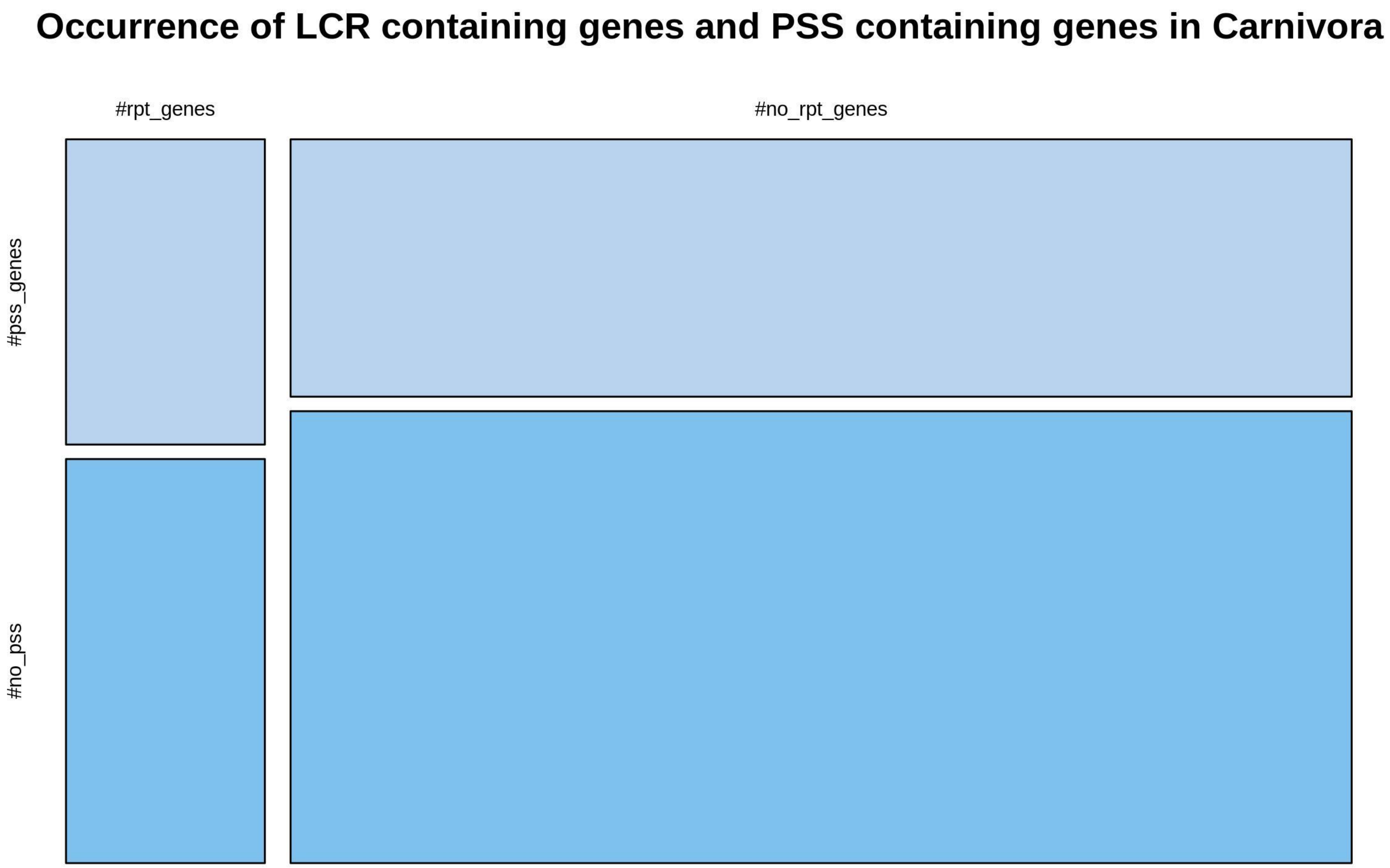

**Supplementary figure 523**

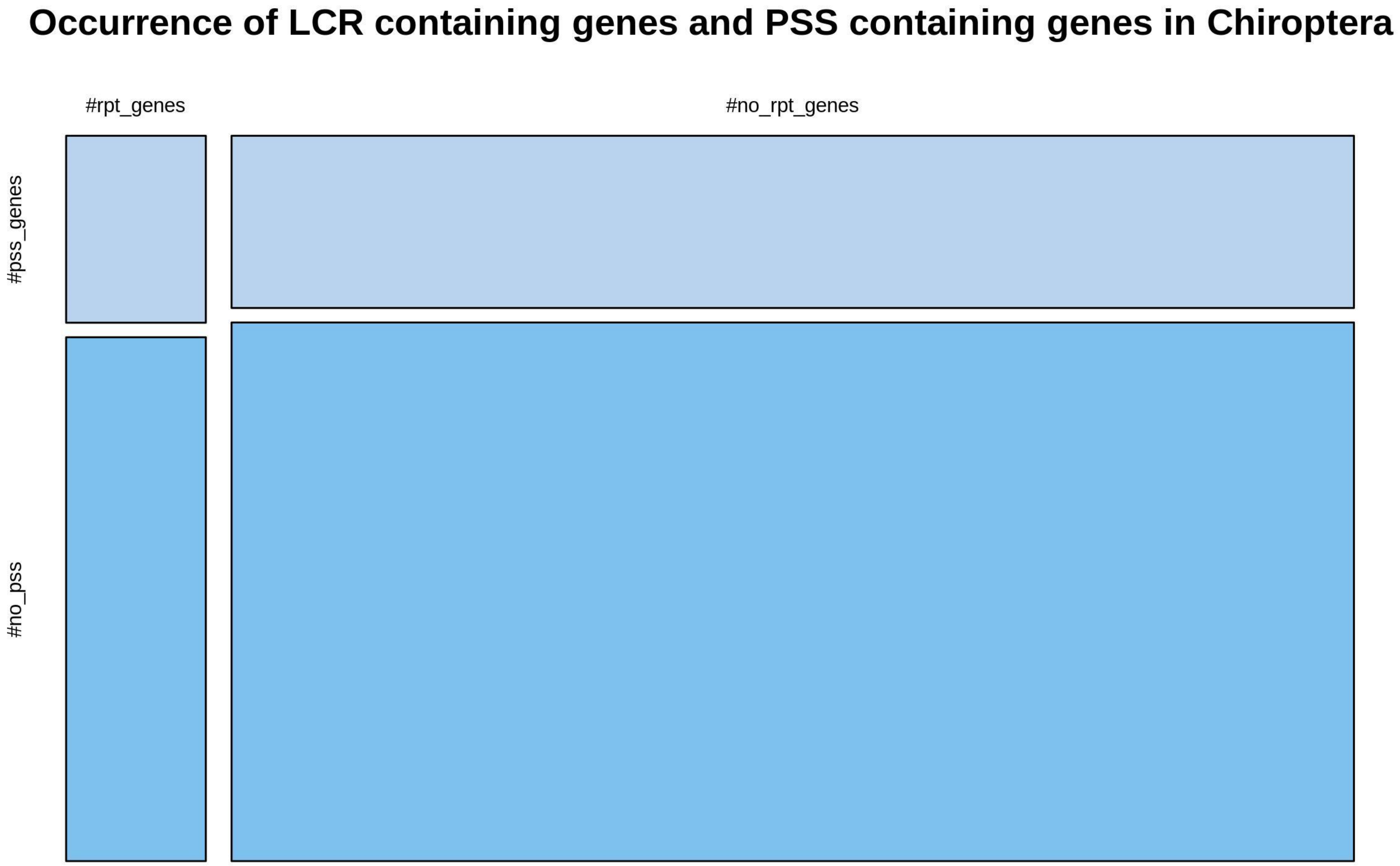

**Supplementary figure 524**

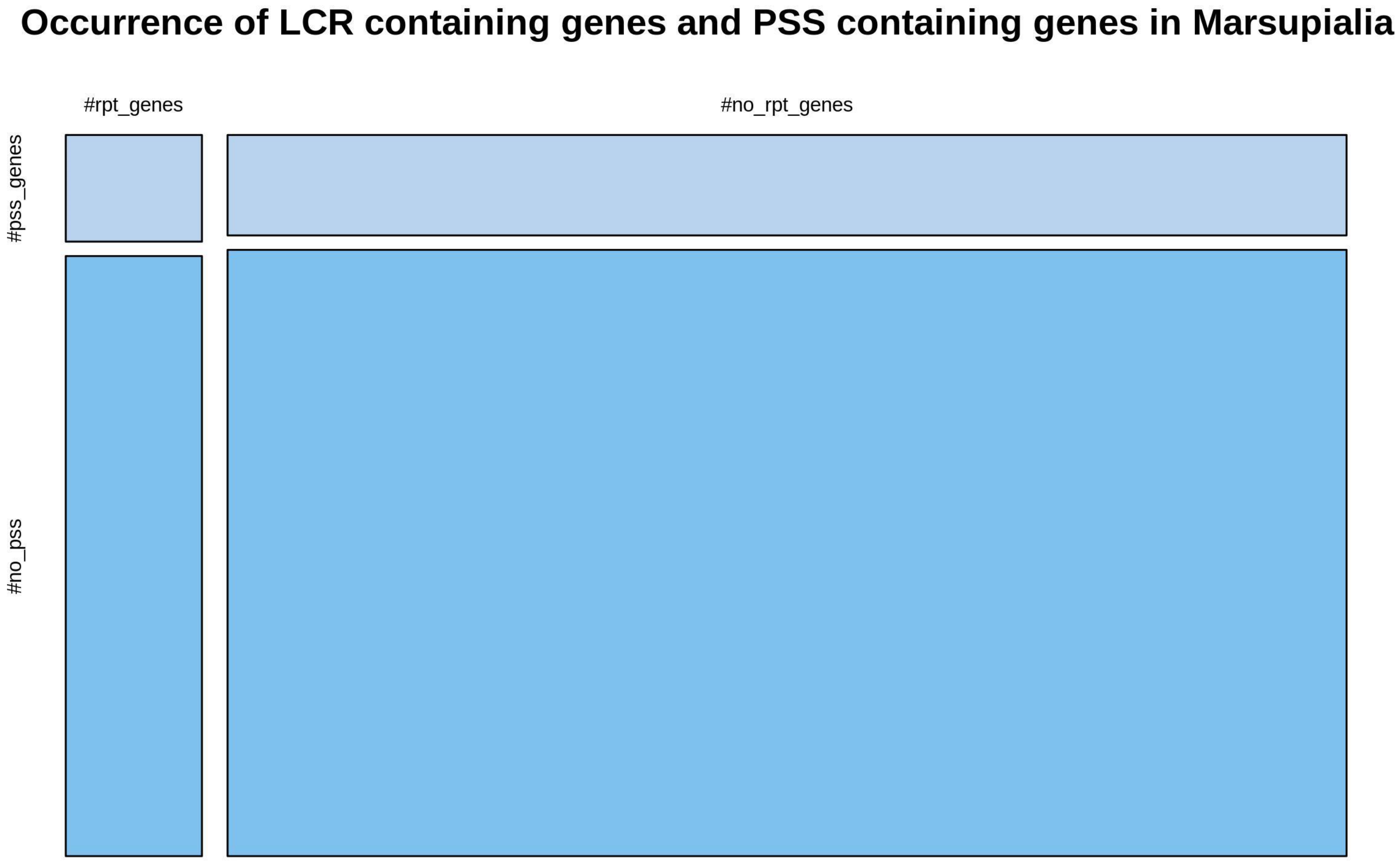

**Supplementary figure 525**

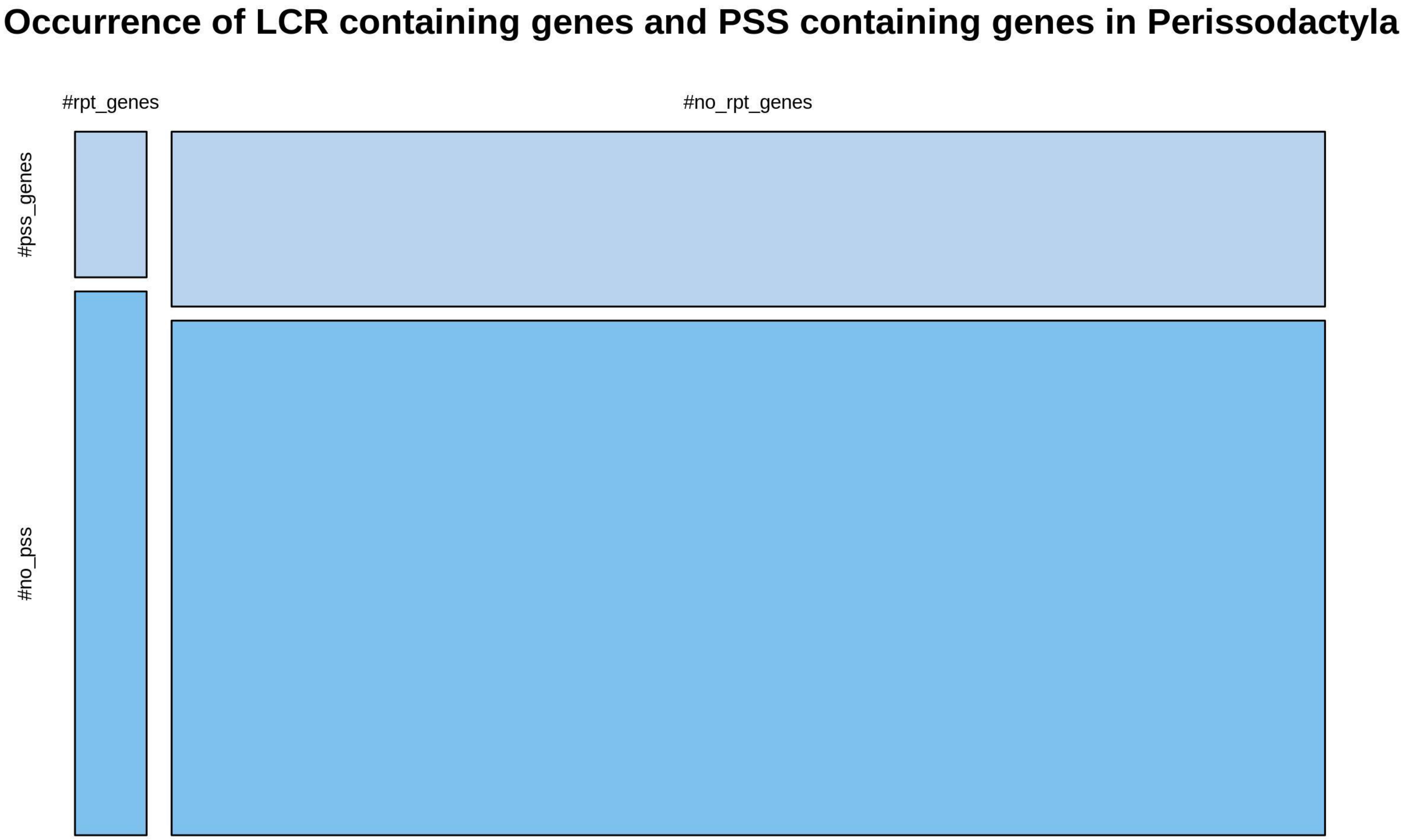

**Supplementary figure 526**

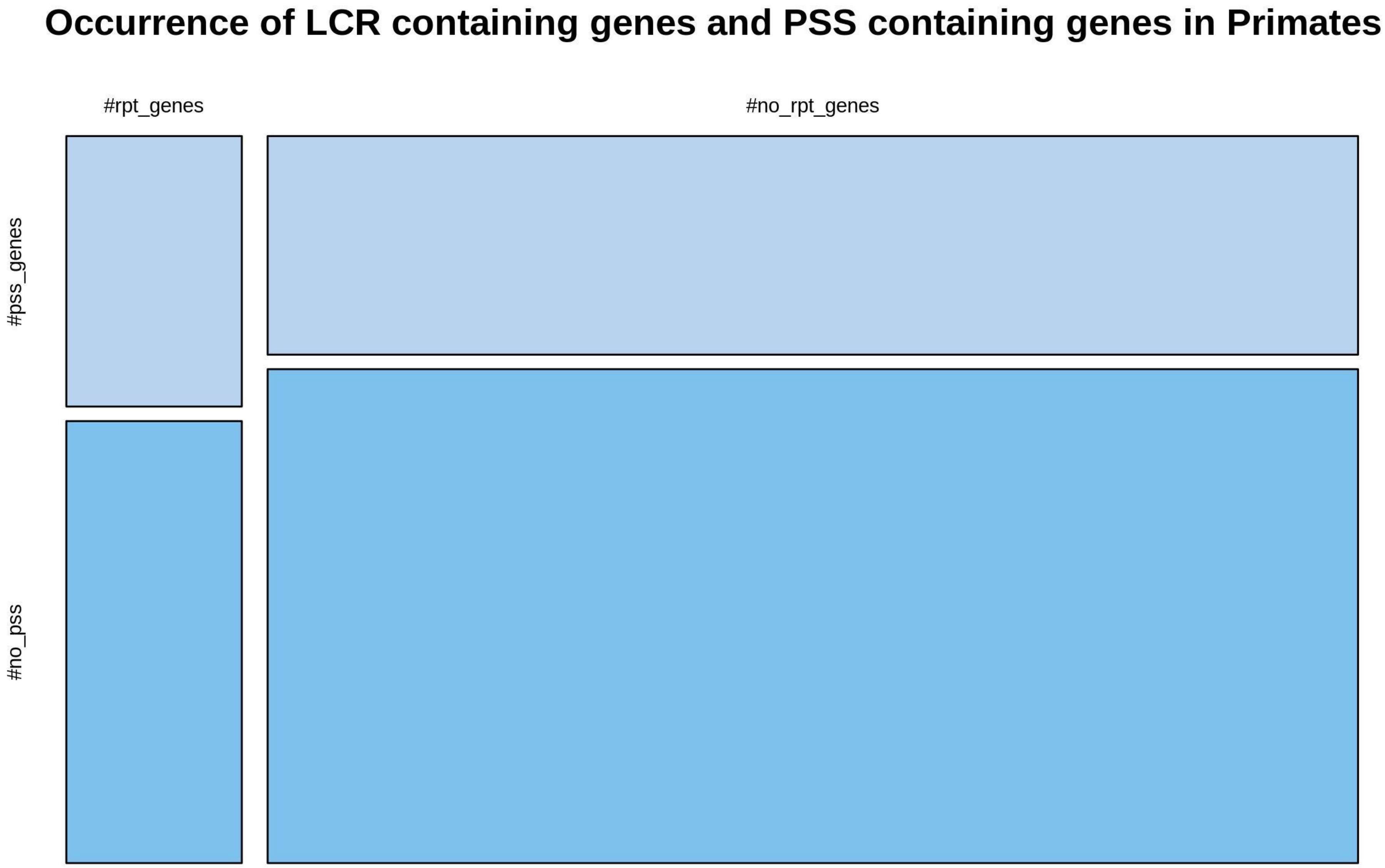

**Supplementary figure 527**

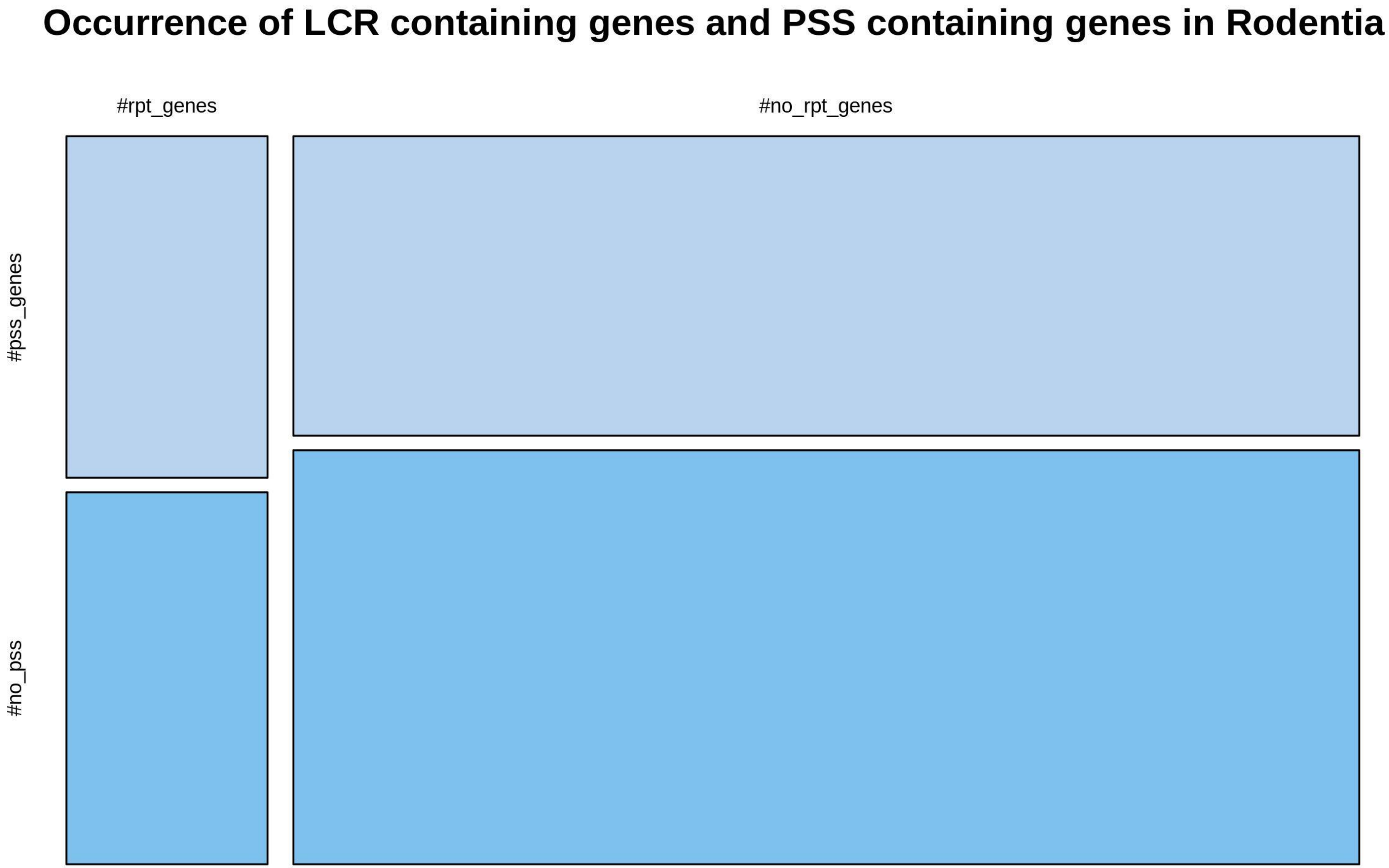

**Supplementary figure 528**

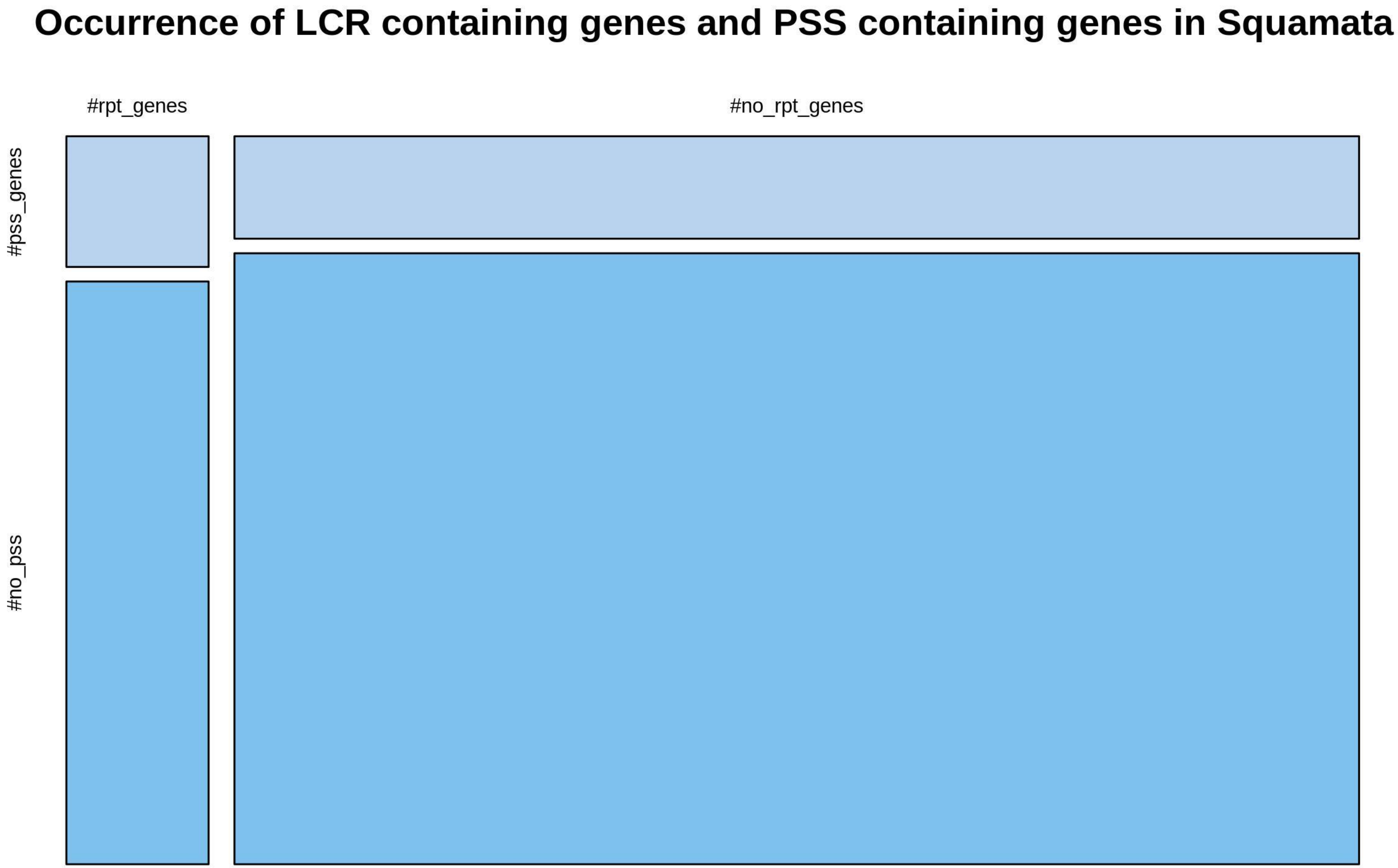

**Supplementary figure 529**

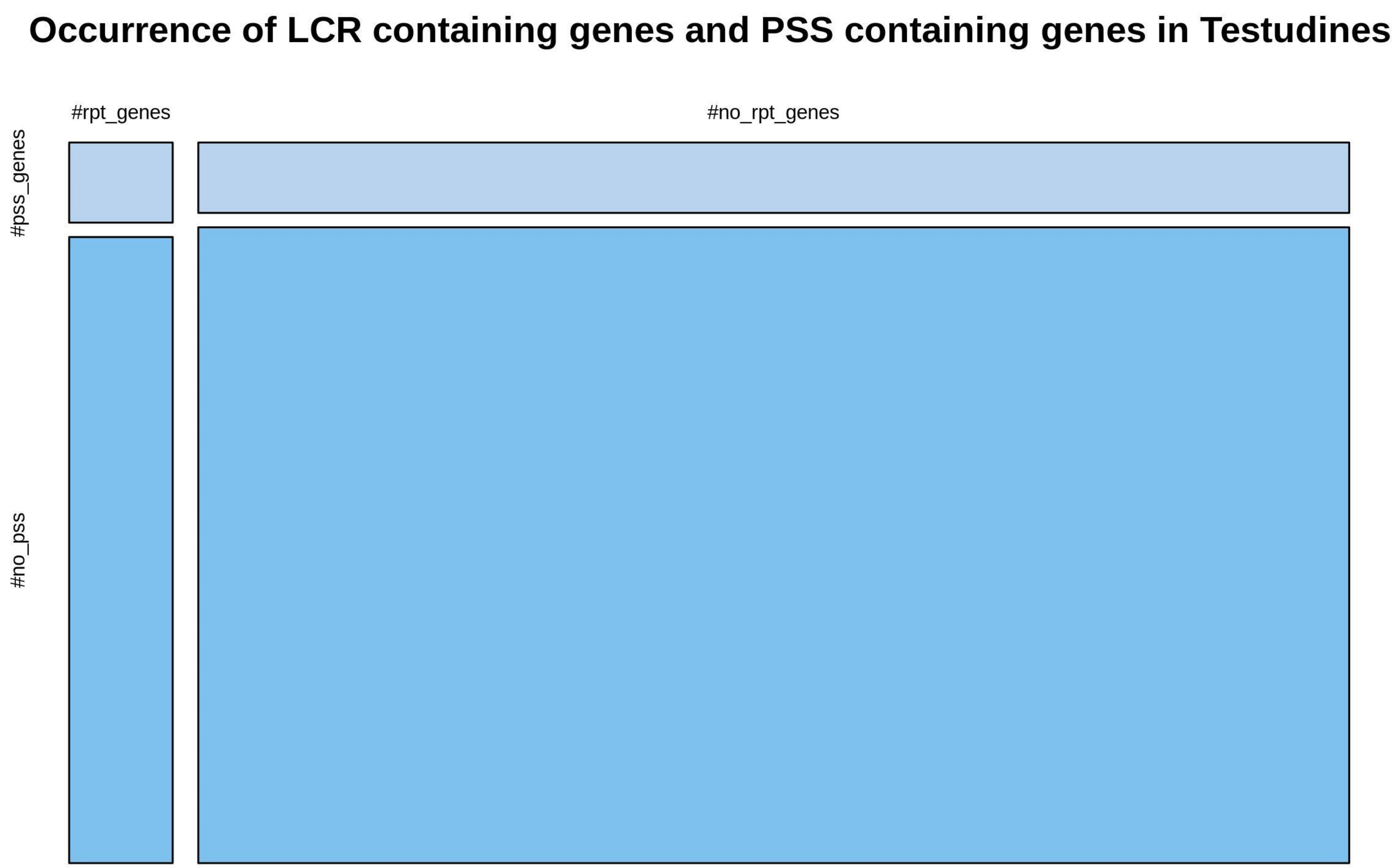

**Supplementary figure 530**

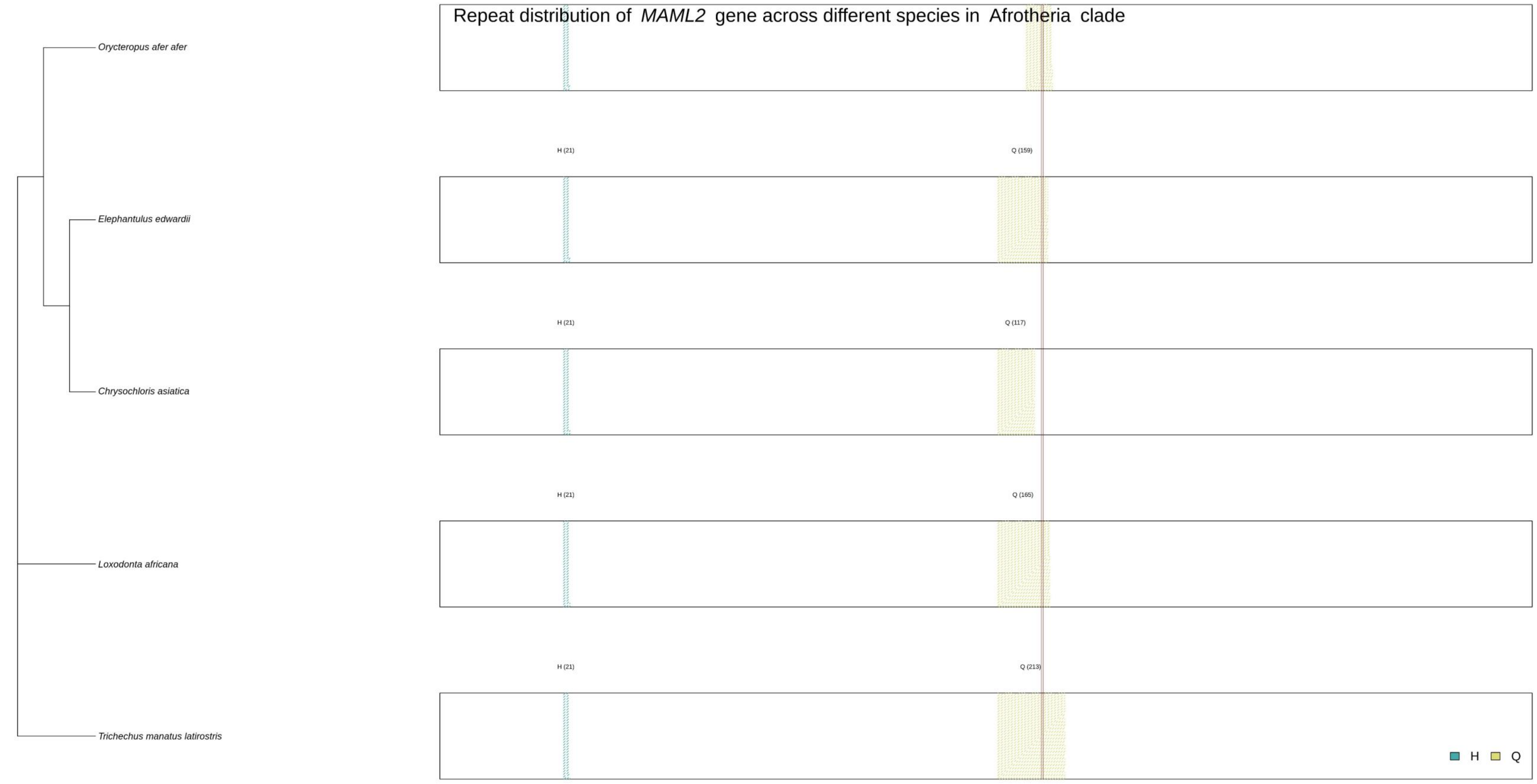

**Supplementary figure 531**

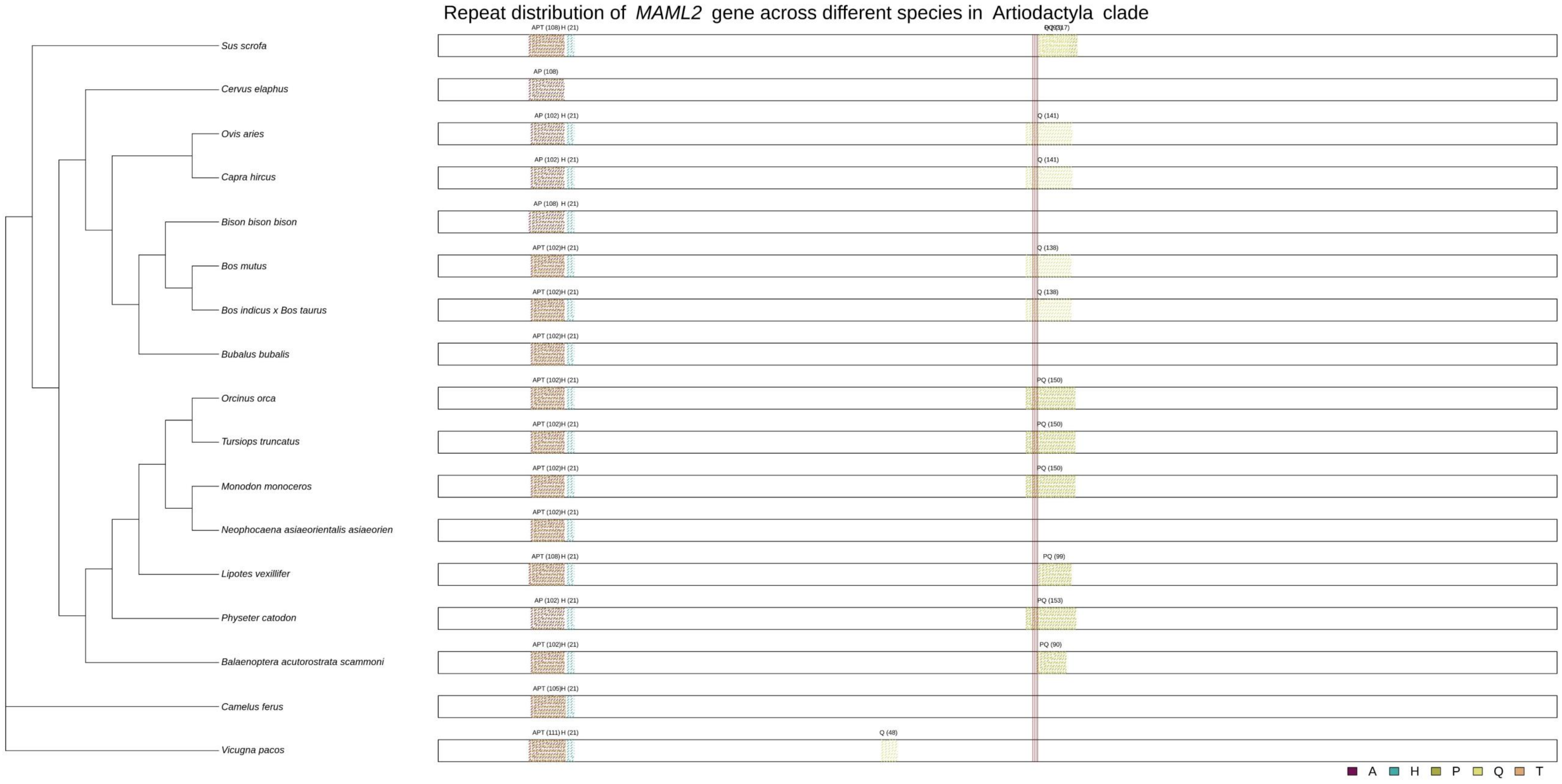

**Supplementary figure 532**

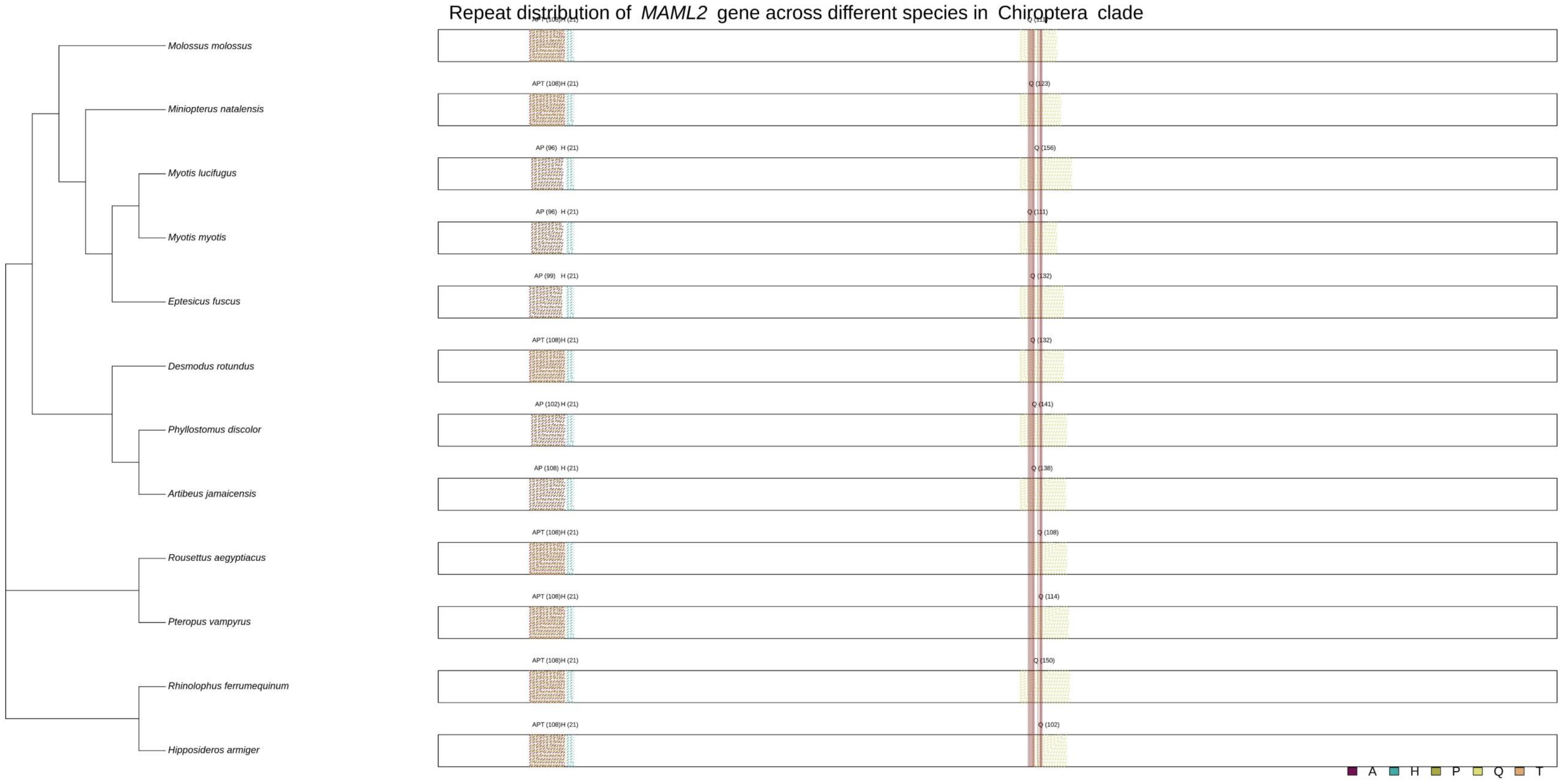

**Supplementary figure 533**

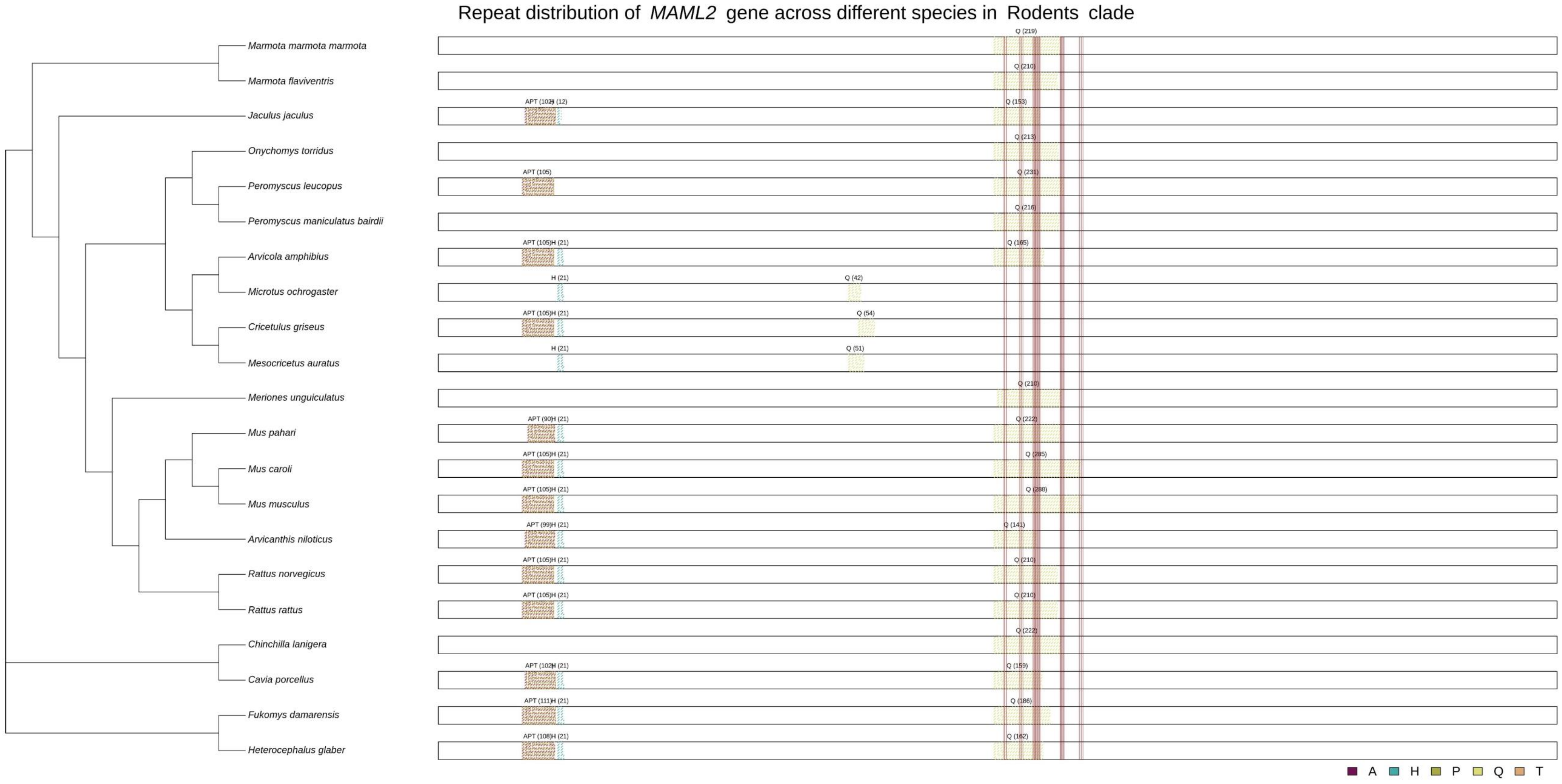
