## Supplementary Text for "Terminal regions of a protein are a hotspot for low complexity regions (LCRs) and selection"

**Dataset preparation**

We downloaded a list of all human protein-coding genes (19619 genes) from Ensembl Biomart release 105 with “Gene_name,” “NCBI_gene_ID,” and “Gene_type” as header (saved as mybiomart_list.txt) (Howe et al. 2021) and filtered out all the mitochondrial, read-through, and LINCs from the list using the Linux command:

grep -v "^MT-" mybiomart_list.txt |awk 'NF==3{print $0}'|grep -v "\bLINC"|grep -v "-"|awk '{print $1,$2}' > final_protein_coding_list

With the above filtering criteria, we made a list of 19124 protein-coding genes and downloaded these protein-coding genes using NCBI datasets (Sayers et al. 2020) command:

for i in `cat final_protein_coding_list|sed '1d'`

do

gene=`echo $i|awk '{print $1}'`

geneid=`echo $i|awk '{print $2}'`

echo $gene $geneid

datasets download ortholog gene-id "$geneid" --filename "$gene".zip

done

NCBI’s Eukaryotic Genome Annotation pipeline calculates ortholog gene groups by comparing the protein sequence similarity and local synteny information in the NCBI Gene dataset. RefSeq genome curators manually review orthologous gene relationships. Orthology is determined by comparing an annotated genome to a reference genome (usually human), and the pairwise orthologs are tracked as a group. The calculation of orthologs is limited to genes in the NCBI Gene database and primarily focuses on vertebrates.

With the following above commands, we successfully retrieved 18586 genes. The downloaded orthologous genes are in compressed “.zip” folder format. We decompressed each folder and saved it according to the gene name as follows:

for i in `ls -1 *zip`

do

gene=`echo $i|sed 's/.zip//g'`

unzip -o “$i”

mv ncbi_dataset/ "$gene"_dataset/

done

mkdir all_datasets

mv *_dataset all_datasets

cd all_datasets

For some genes, the decompression failed, thus making our final list of 18407 genes (**Supplementary Table 2**). The downloaded gene sequences are subsetted into 13 clades ( Afrotheria, Rodentia, Chiroptera, Carnivora, Perissodactyla, Primates, Artiodactyla, Squamata, Testudines, Aves, Lagomorpha, Marsupialia, and Amphibia: 308 species in total, **Supplementary Table 1** ). The species are selected to cover diverse life forms of vertebrate subphylum. Furthermore, the species are grouped according to similar morphology, common ancestry, and similar environmental conditions into class (Aves and Amphibia), order (Primates, Rodentia, Lagomorpha, Chiroptera, Artiodactyla, Perissodactyla, Carnivora, Testudines, and Squamata), infraclass (Marsupialia), and superorder (Afrotheria).

We selected the most similar amino acid sequences for each gene within each clade using the following procedure: (i) We choose all the “NP” denoted sequences for a gene in a clade and align the sequences. Using the aligned sequences, we generate a consensus sequence, (ii) If a species has only one “NP” sequence for a gene, the sequence is considered the final sequence of that species, (iii) If a species has more than one “NP” sequence, we align each sequence with the consensus sequence and select the one with the best per base alignment score, and (iv) if no “NP” sequence exists for a species, all the “XP” sequences of that species for that gene are aligned with the consensus sequence, and the one with best per base alignment score is selected.

A combination of Linux scripts and Python programs achieves the above output.

#The below script is written in python3. Python scripts are indentation sensitive.

##Note: Please save the below command as consen.py

```
import sys
filename=sys.argv[1]
outname=sys.argv[2]
f = open(filename, 'r')
data=f.read()
d=data.replace('\n\n', '@').replace('\n', '.')
ar1 = []
splat = d.split("@")
finalconsen=''
finalnuc=''
def most_frequent(list):
    counter = 0
    num = list[0]
    l=len(list)
    for i in list:
        curr_frequency = list.count(i)
        if(curr_frequency> counter):
            counter = curr_frequency
            num = i
    if counter/l > 0.25:
        return(num)
    else:
        return('-')
for number, paragraph in enumerate(splat, 1):
        ar1 = [paragraph]
        print(number)
        print(ar1)
        newar=ar1[0].split('.')
        if” in newar:
                newar.remove('')
        newar=[ x for x in newar if ' ' not in x ]
        leng=len(newar[0])
        numele=len(newar)
        for pos in range(0,leng):
                consen=[]
                for ele in range(0,numele):
                        obj=newar[ele][pos]
                        consen.append(obj)
                consen=[ x for x in consen if '#' not in x ]
                print(consen)
                print(finalnuc)
                if not consen:
                        consen.append('-')
                finalnuc=most_frequent(consen)
                finalconsen = finalconsen + finalnuc
print(finalconsen)
print(finalconsen, file=open(str(outname)+"_consensus.txt", "w"))
```

####################Python script ends here###################

#####Running the Linux commands to generate the files of similar sequences#######

```
###The script assumes that you are in "all_datasets" directory and have fulfilled all the below mentioned requirements##########
###########################################REQUIREMENTS######################################
##1. list of clade species (with extension *_list) in a folder adjacent to "all_datasets" folder
##2. consen.py: python3 script to make consensus sequence from aligned sequences
##3. needle and muscle program to be installed on your system
##Run it in the folder where all the downloaded datasets are
##all_datasets directory contains all the unzipped directory downloaded from NCBI for protein coding genes with names as : ZZZ3_dataset
##Each directory genename_dataset has a directory “data” which contains files like: data_report.jsonl  dataset_catalog.json  data_table.tsv  gene.fna  protein.faa  rna.fna
##For first objective, we are mainly focussing on file protein.faa for each gene
for dirs in `ls -d all_datasets/*_dataset`
do
cp consen.py "$dirs"/data/consen.py
cp cladelist/* "$dirs"/data/
done
cd all_datasets
ls -d *_dataset > alldatasets
split -d --additional-suffix=_list.txt -l 100 alldatasets
for splits in `ls -1 *_list.txt`
do
for dir in `cat $splits`
do
gene=`echo $dir|sed 's/_dataset//g'`
cd “$dir”/data
cat protein.faa|sed '/>/s/ .*organism=/_/1'|sed '/>/s/].*//g'|sed 's/ /_/g'|awk '{print $1}'|sed '/>/s/$/@/g'|tr -d "\n"|sed 's/>/\n>/g'|sed '1d'|sed 's/@/\n/g' > onelineramino.seqs
grep -A1 ">NP_" onelineramino.seqs|grep -v "\-\-"|grep -v ">"|awk '{print length()}' > length_distribution
grep -A1 ">NP_" onelineramino.seqs|grep -v "\-\-" > npseqs
grep -A1 ">NP_" onelineramino.seqs|grep -v "\-\-"|grep ">"|sed 's/>//g' > npheaders
totalheaders=`wc -l npheaders|awk '{print $1}'`
echo “a=read.table('length_distribution',header=F)" > length.R
echo "b=summary(a)" >> length.R
echo "write.table(b,file='quartile',quote=F,col.names=F,row.names=F)" >> length.R
Rscript length.R
min=`grep "Min." quartile|awk '{print $2}' FS=:`
max=`grep "Max." quartile|awk '{print $2}' FS=:`
first_quart=`grep "1st" quartile|awk '{print $2}' FS=:`
third_quart=`grep "3rd" quartile|awk '{print $2}' FS=:`
rm rangelength.fa
subt=`expr $max - $min`
if [ $totalheaders -le 4 ] || [ $subt -le 40 ]
then
cp npseqs rangelength.fa
else
while read header
do
sequence=`grep -A1 "$header" npseqs|tail -n1`
j=`echo $sequence|wc -c`
echo $first_quart $third_quart $sequence $header
echo $first_quart $third_quart $sequence $header |awk 'length($3) >= $1 && length($3) <= $2 {print $4,$3}' OFS="\n" >> rangelength.fa
done < npheaders
fi
muscle -in rangelength.fa -out rangelengthaligned.aln
sed -e '/>/s/$/#/g' rangelengthaligned.aln|tr -d '\n'|sed -e 's/>/\n>/g' -e 's/#/\n/g'|sed '1d'|grep -v ">" > alignedblock.txt
python3 consen.py alignedblock.txt aminoconsen
sed -i ‘s/-//g’ aminoconsen_consensus.txt
#wc -c aminoconsen_consensus.txt
sed -i ‘1 i\>reference’ aminoconsen_consensus.txt
reflen=`cat aminoconsen_consensus.txt|tail -n1|wc -c`
ref=`cat aminoconsen_consensus.txt|tail -n1`
for clade in `ls *_list`
do
rm -r "$gene"_"$clade"_orf.fa
for i in `cat $clade`
do
score=0
grep “$i” onelineramino.seqs > temp.txt
j=`wc -l temp.txt|awk '{print $1}'`
if [ $j -eq 1 ]
then
accession=`grep “$i” onelineramino.seqs`
seq=`grep -A1 "$accession" onelineramino.seqs|tail -n1`
echo -e "$accession\n$seq" >> "$gene"_"$clade"_orf.fa
elif [ $j -gt 1 ]
then
grep -A1 "$i" onelineramino.seqs > tempmorethanone.fa
leng=`wc -l tempmorethanone.fa|awk '{print $1}'`
for ini in `seq 2 2 $leng`
do
header=`head -n"$ini" tempmorethanone.fa|tail -n2|head -n1`
seq1=`head -n"$ini" tempmorethanone.fa|tail -n1`
seq1len=`echo $seq1|wc -c`
needle -asequence asis:$ref -bsequence asis:$seq1 -gapopen 10.0 -gapextend 0.5 -outfile temprefneedle
sco1=`grep Score temprefneedle |awk '{print $3}'`
q1len=`grep "Length: " temprefneedle|awk '{print $3}'`
score1=`echo $sco1 $q1len|awk '{print $1/$2}'`
echo -e “$seq1\t$score1”
if (( $(echo "$score1 > $score" |bc -l) ))
then
score=$score1
seq=$seq1
seqlen=$seq1len
accession=$header
fi
done
echo -e "$accession\n$seq" >> "$gene"_"$clade"_orf.fa
fi
done
done
rm temp*
cd ../..
done
done

###SAMSN1 and SFTA3 are excluded as SAMSN1 does not have any NM_ sequence for any species and SFTA3 is mentioned as non-coding on NCBI
```

**Low complexity regions identification**

We use fLPS2.0 (Harrison 2021) to identify stretches of low probability in the amino acid sequences. Even though many programs are available to detect low-complexity regions, we selected fLPS2.0 because it is open-source and rapidly identifies LCRs in large datasets. We detect regions of low probability sequences (LPSs) for each amino acid fasta file in a clade-wise manner. Firstly, we calculate the background residue composition for each amino-acid multifasta file using *CompositionMaker,* an accessory program of fLPS2.0. fLPS2.0 identifies LPSs based on the background frequency detected by *CompositionMaker*. Stretches that are longer than four amino acids, have more than 70 % composition, and are composed of less than five different types of amino acids are considered low-complexity regions (LCRs).

#####Script to identify LPSs using fLPS2.0####

```
###running fLPS2.0###
for splits in `ls -1 *_list.txt`
do
for dir in `cat $splits`
do
gene=`echo $dir|sed 's/_dataset//g'`
cd “$dir”/data
for orfs in `ls -1 *_list_orf.fa`
do
prefix=`echo $orfs|cut -f1,2 -d"_"`
CompositionMaker "$orfs"
composfile=`echo "$orfs".COMPOSITION`
fLPS2 -c "$composfile" "$orfs" > "$prefix"_flpsout_withcomposition
done
cd ../..
done
done
cd $curr_dir
mkdir flps_output
cd all_datasets
for dir in `ls -d *_dataset`
do
cp "$dir"/data/*_flpsout_withcomposition ../flps_output
done
cd "$curr_dir"/flps_output
## remove {X} or any repeat with X, select more than 70% and stretch with more than 3aa
#######################################################################
## combine all the composition file to make a single file
rm -v !(*_flpsout_withcomposition)
for files in `ls|grep "flpsout_withcomposition"`
do
gene=`echo $files|cut -f1 -d"_"`
clade=`echo $files|cut -f2 -d"_"`
awk -v g=$gene -v c=$clade '$7>3 && $9 !~ /X/ && ($7/(($6-$5)+1)) > 0.7 {print c,g,$0}' OFS='\t' $files >> morethanseventy_combined
done
```

#######################Linux script ends here######

**Filtering and alignment**

We map all the selected amino-acid sequences to their respective CDS sequence. The **Supplementary Table 2** provides a comprehensive list of gene NCBI accession numbers for all genes**.** We remove all the sequences with incomplete ORF ( absent START codon, absent STOP codon, contains non-nucleotide characters, or sequence not a multiple of three ). If a multifasta file for a gene in a clade contains less than four species, we drop it. Surprisingly, after applying the filtering criteria, the Lagomorpha clade gets entirely filtered out and thus is not included in molecular evolutionary analyses. The remaining files are codon-aligned using the GUIDANCE (Sela et al. 2015) program with MUSCLE (Edgar 2004) aligner with 100 bootstraps.

####Linux commands to achieve the above-mentioned:

```
for dir in `ls -d *_dataset`
do
cd “$dir”/data
rm -f *_similar_length_cds.fa
gene=`echo $dir|cut -f1 -d "_"`
for aa in `ls -1 *_list_orf.fa`
do
  clade=`echo $aa|cut -f2 -d"_"`
  grep ">" "$aa"|sed -e 's/>//g' |cut -f1,2 -d"_" > protein_acc.txt
  grep ">" "$aa"|sed -e 's/>//g' |sed 's/_/\t/2' > pacc_spname.txt
  for pacc in `cat protein_acc.txt`
  do
    spname=`grep "\b$pacc\b" pacc_spname.txt|cut -f2`
    grep "\b$pacc\b" data_table.tsv|cut -f14|sed -e ‘s/:/\t/g’ -e ‘s/-/\t/g’ |awk ‘{print $1,$2-1,$3}’ OFS='\t’ |head -n1 > temp.bed
    cdsacc=`awk '{print $1}' temp.bed`
    bedtools getfasta -fi rna.fna -bed temp.bed |awk '{print $1}' FS=':' > temp.seq
    sed -i “s/$cdsacc/$spname/g” temp.seq
    cat temp.seq >> "$gene"_"$clade"_similar_length_cds.fa
    rm temp.bed temp.seq
  done
 done
cd ../..
done
```

The above script finds the corresponding nucleotide sequence for each amino acid sequence selected in our analysis.

We keep all the CDS multifasta files in one folder and run a Linux command to make the file such that the header will be in one line and the whole sequence of the species in one line.

```
for file in `ls *_similar_length_cds.fa`
do
sed -e '/>/s/$/#/g' -e 's/>/@>/g' “$file”|sed -z 's/\n//g'|sed -e 's/@/\n/g' -e 's/#/\n/g'|sed '1d' > “$file”.txt
done
##removing the sequences which have premature stop codon in between
####we will make use of orthochecker.py for this
##this file takes a multifasta file and output name as input argument and returns the sequences which have proper ORF and exclude those which does not have proper ORF
##please run a command on the multifasta file such that the header will be in one line and the whole sequence of the species will be in one line only
##in the end, it will create a multifasta file with only proper ORFs and trim the STOP codon from the end

####Save the below script as orthochecker.py
import sys
import re
filename=sys.argv[1]
outname=sys.argv[2]
fn = open(filename, 'r')
h = fn.readlines()
for i in range(0,len(h),2):
  seqrange=i+1
  header=h[i]
  line=h[seqrange]
  header=header.strip()
  line=line.strip()
  print(header)
  #print(line)
  #print(len(line))
  if line.startswith("ATG") and len(line)%3==0 and line.endswith((“TAA”, “TAG”, “TGA”)) and re.match(‘^[ATGCN]*$’, line):
    line=line[:-3]
    n=3
    codonlist=[]
    for index in range(0,len(line),n):
      codon=line[index:index+n]
      codonlist.append(codon)
    if ‘TAA’ not in codonlist and ‘TAG’ not in codonlist and ‘TGA’ not in codonlist:
      print(header, file=open(str(outname)+"_orf.txt", "a"))
      print(line, file=open(str(outname)+"_orf.txt", "a"))

ls|grep -v "ortho\|old\|allfiles" > allfiles
for files in `cat allfiles`
do
outname=`echo $files|sed 's/_cds.fa//g'`
python3 orthochecker.py $files $outname
done
mkdir all_orfs
ls|grep -v "ortho\|allfiles\|old" > allfiles
for i in `grep “length_orf” allfiles`
do
cp $i all_orfs/
done
cd all_orfs/
###removing all the files which have less than 4 sequences
find . -type f -exec awk -v x=8 'NR==x{exit 1}' {} \; -exec rm -f {} \;
##the above step leaves 184271 files
```

###Running the alignment on the filtered CDS multifasta files

```
##############################################
guidance=/path/to/guidance.v2.02/www/Guidance
ls > fasta_list
sed -i '/fasta_list/d' fasta_list
for i in `cat fasta_list`
do
j=`echo $i|sed 's/.txt//g'`
perl "$guidance"/guidance.pl --program GUIDANCE --seqFile "$i" --msaProgram MUSCLE --seqType codon --outDir "$j".100_MUSCLE --genCode 1 --bootstraps 100 --proc_num 2
done
```

**Identification of orthologous low-complexity regions**

We can write all possible combinations of a similar LPS region with multiple amino-acid compositions. For example, if an LPS consists of amino acids A, R, and G, we can write it as ARG, AGR, RAG, GRA, RGA, and GAR. We use an awk script (permute.awk) and convert all varying amino-acid compositions to the same combination.

######Save the below script as permute.awk:

```
function permute(s, st,     i, j, n, tmp) {
    n = split(s, item,//)
    if (st > n) {  print s; return }
    for (i=st; i<=n; i++) {
        if (i != st) {
         tmp = item[st]; item[st] = item[i]; item[i] = tmp
         nextstr = item[1]
         for (j=2; j<=n; j++) nextstr = nextstr delim item[j]
        }else {
          nextstr = s
        }
       permute(nextstr, st+1)
       n = split(s, item, //)
   }
}
{ permute($0,1) }
```

#################################

```
###replacing combinations of repeats with one type only###
##replacing all the jumbled LPSs to one only i.e. AC = CA
sed -i ‘s/{//g’ morethanseventy_combined
sed -i ‘s/}//g’ morethanseventy_combined
##removing all the lines which have LPSs of length >4aa
for clades in afrotheria amphibia artiodactyla aves carnivore chiroptera lagomorpha marsupials perissodactyla primates rodents squamata testudines
do
awk 'length($11) < 5 {print $0}' "$clades"_morethanseventy_combined > temp.txt
mv temp.txt "$clades"_morethanseventy_combined
done
#making a file with all unique LPSs
for clades in afrotheria amphibia artiodactyla aves carnivore chiroptera lagomorpha marsupials perissodactyla primates rodents squamata testudines
do
cut -f11 "$clades"_morethanseventy_combined|sort -u > "$clades"_allrepeats
done
for clades in afrotheria amphibia artiodactyla aves carnivore chiroptera lagomorpha marsupials perissodactyla primates rodents squamata testudines
do
files="$clades"_morethanseventy_combined
while [ -s "$clades"_allrepeats ]
do
a=`head -n1 "$clades"_allrepeats`
sed -i "/\b$a\b/d" "$clades"_allrepeats
echo "$a" > "$clades"_permute_file
for newrep in `awk -f permute.awk "$clades"_permute_file`
do
  echo -e "$a\t$newrep"
  awk -v ol=$a -v k=$newrep '$11==k{$11=ol}1' OFS='\t' $files > "$clades"_temp && mv "$clades"_temp $files
  sed -i "/\b$newrep\b/d" "$clades"_allrepeats
done
done
rm "$clades"_allrepeats "$clades"_permute_file
```

The coordinates of LCR stretches are mapped to their respective aligned CDS sequence using a custom script (coordinate_mapper.sh). The stretches with overlap are considered orthologous.

We convert all the aligned CDS multifasta such that the whole aligned sequence for a species in a gene will be in one line.

```
for file in `ls *_similar_length_orf.aln`
do
sed -e '/>/s/$/#/g' -e 's/>/@>/g' “$file”|sed -z 's/\n//g'|sed -e 's/@/\n/g' -e 's/#/\n/g'|sed '1d' > “$file”_oneliner
done

###Save it in a file coordinate_mapper.sh
################coordinate_mapper.sh################
for file in `ls $1`
do
echo $file
gene=`echo $file|cut -d '_' -f1`
clade=`echo $file|cut -d '_' -f2`
repeat=`echo $file|cut -d '_' -f3`
alnfile=`echo "$gene"_"$clade"_similar_length_orf.aln_oneliner`
echo $gene $clade $repeat $alnfile
if [ -f $alnfile ]
then
while read oneline
do
species=`echo $oneline|awk '{print $3}'|cut -f3- -d "_"|sort -u`
(sequ=`sed -e '/>/s/$/#/g' -e 's/>/@>/g' "$gene"_"$clade"_similar_length_orf.aln_oneliner| sed -z 's/\n//g'|sed -e 's/@/\n/g' -e 's/#/\n/g' |grep -A1 "$species"|grep -v "^>" |sed -z 's/\n//g'`
j1=`echo $oneline|awk '{print $4}'`
j2=`echo $oneline|awk '{print $5}'`
len=`echo $sequ|awk '{print length}'`
init=`echo $j1|awk '{print ($1*3) - 2}'`
last=`echo $j2|awk '{print $1*3}'`
a1=0
a2=0
b1=0
b2=0
for char in `echo $sequ|sed 's/.\{1\}/&\t/g'`
do
if [ $a1 -eq $init ]
then
break
elif [ $char != '-' ]
then
a1=`echo $a1|awk '{print $1 + 1}'`
fi
a2=`echo $a2|awk '{print $1+1}'`
done
for char in `echo $sequ|sed 's/.\{1\}/&\t/g'`
do
if [ $b1 -eq $last ]
then
break
elif [ $char != '-' ]
then
b1=`echo $b1|awk '{print $1 + 1}'`
fi
b2=`echo $b2|awk '{print $1+1}'`
done
echo -e “$clade\t$gene\t$species\t$repeat\t$a1\t$b1\t$a2\t$b2”
echo -e "$clade\t$gene\t$species\t$repeat\t$a1\t$b1\t$a2\t$b2" >> "$clade"_"$gene"_"$species"_"$repeat"_coordinates_repeats.txt )&
done < $file
else
echo "$file" >> aln_file_not_present.txt
fi
done
```

############coordinate_mapper.sh ends here#######

```
## making the coordinate files for the respective clade having more than one species for a repeat and saving the rest in “Discarded.txt” file.
rm -f *_aa_coordinates.txt
for clade in `cut -f1 morethanseventy_combined|sort -u`
do
for gene in `cut -f2 "$clade"_morethanseventy_combined|sort -u`
do
for repeat in `awk -v g=$gene '$2==g{print $0}' "$clade"_morethanseventy_combined|awk '{print $11}'|sort -u`
do
awk -v g=$gene '$2==g{print $0}' "$clade"_morethanseventy_combined|awk -v r=$repeat '$11=r{print $3}' > species.list
numsp=`wc -l species.list|awk '{print $1}'`
if [ $numsp -gt 1 ]
then
awk -v g=$gene -v r=$repeat ‘$2==g && $11==r{print $0}’ “$clade”_morethanseventy_combined|awk ‘{print $1,$2,$3,$7,$8,$11}’ OFS=” \t”>> “$gene”_“$clade”_“$repeat” _aa_coordinates.txt
echo "$gene $clade $numsp $repeat"
else
awk -v g=$gene -v r=$repeat ‘$2==g && $11==r{print $0}’ “$clade”_morethanseventy_combined|awk ‘{print $1,$2,$3,$7,$8,$11}’ OFS=” \t”>> Discarded.txt
fi
done
done
done
ls|grep "_aa_coordinates.txt" > allcoordinatefiles
mkdir ../allcoordinateoutputfiles
while read j
do
echo $j
mv "$j" ../allcoordinateoutputfiles
done < allcoordinatefiles
cd ../allcoordinateoutputfiles
find . -name '*_aa_coordinates.txt' -type f | xargs wc -l|awk '$1>3'|cut -f2 -d"/"|grep "_aa_coordinates.txt" > allfileswithmorethanthree
mkdir morethanthreespecies
cut -f1,2 -d"_" allfileswithmorethanthree|sort -u > foralnfiles
###moving aln files to same folder that have more than three species
while read j
do
file=`echo "$j"_similar_length_orf.aln_oneliner`
cp /path/to/alnfiles_oneliner/"$file" morethanthreespecies
done < foralnfiles
###moving coordinate files with more than three lines in them to morethanthreespecies folder
while read j
do
cp “$j” morethanthreespecies
done < allfileswithmorethanthree
cd morethanthreespecies
####since all repeat files with more than three species do not have alignment (probably because some sequences had premature stop codon in between and after excluding them, the number of species in MSA files were less than four), now we are keeping aln files and repeats files in the same folder where both are available
mkdir alnandrepeatcommon
ls|grep aa_coordinates.txt|cut -f1,2 -d_|sort -u > coordfiles
ls|grep aln_oneliner|cut -f1,2 -d_|sort -u > alnfiles
cat coordfiles alnfiles|sort|uniq -c|awk '$1==2{print $0}'
cat coordfiles alnfiles|sort|uniq -c|awk '$1==2{print $2}' > bothfilespresent
while read j
do
echo $j
cp "$j"* alnandrepeatcommon
done < bothfilespresent
cd alnandrepeatcommon
ls|grep "_aa_coordinates.txt" > allcoordinatefiles
split -d --additional-suffix=_coordinate_splits -l 8958 allcoordinatefiles
#####we have moved coordinate_mapper.sh here in this folder. Now we gonna run it.
cp allcoordinatefiles allcoordinatefiles_backup
while [ -s allcoordinatefiles ]
do
j=`ps -ef|grep -c coordinate_mapper.sh`
if [ $j -lt 12 ]
then
i=`head -n1 allcoordinatefiles`
( bash coordinate_mapper.sh “$i”) &
sed -i ‘1d’ allcoordinatefiles
fi
done

###combine all the coordinate output to one file: coordoutcompiled

awk '$5!=0 && $6!=0 && $7!=$8{print $0}' coordoutcompiled > coordinateoutfiltered

while read j
do
echo $j
clade=`echo $j|awk '{print $1}'`
gene=`echo $j|awk '{print $2}'`
species=`echo $j|awk '{print $3}'`
alnlength=`head -n2 alnfiles_oneliner/"$gene"_"$clade"_similar_length_orf.aln_oneliner|tail -n1|awk '{print length}'`
echo "$j $alnlength" >> coordinatecompiledwithalnlength
done < coordinateoutfiltered
```

**Results preparation**

**Violin plots**

To generate a violin plot depicting the length distribution among Low Complexity Regions (LCRs), we first gathered all LCRs. Then we stratified them by the purity (i.e., the ratio of primary residues to total residues in a given stretch) for each clade. Subsequently, we computed each subset’s length distribution and density of LCRs in nucleotides.

A violin plot combines the features of a box plot and a kernel density plot to provide a more detailed and informative view of the data. It consists of a central box representing the data’s interquartile range (IQR) and a “violin” shape on either side of the box representing the kernel density estimate. The width of the violin at any point represents the density of data points at that point. These plots are particularly useful for comparing the distributions of multiple datasets at once, as the shape of each violin reflects the distribution of the corresponding dataset. The violin plot is generated in R (R Core Team 2021) (**see Figure S1-10** and **Fig. 1)**.

**Proportion vs. purity plot**

To comprehensively evaluate the distribution and stratification of Low Complexity Regions (LCRs) within the Tetrapoda clade, we quantified the proportion of LCRs based on the purity of the stretches in the R base (**see Figure S11)**.

**Gene ontology enrichment analyses**

Gene ontology (GO) is a system for categorizing genes and their proteins by their functions, processes, and cellular components in a standardized and hierarchical manner. GO terms provide a normalized vocabulary for describing the functions and features of genes and their products. They can also help analyze large datasets and identify biological processes or pathways associated with a particular phenotype, disease, or experimental condition.

GO enrichment analysis is one such method to identify the involvement of a set of genes in a particular phenotype. A GO enrichment analysis produces a list of GO terms that significantly overrepresent the set of genes of interest, along with statistical measures such as p-values and false discovery rates (FDR) that indicate the significance of the enrichment.

This study performs GO enrichment analysis on the list of genes containing LCRs (>70% purity) using ShinyGO 0.76 (Ge et al. 2020). We analyzed GO for biological processes, molecular functions, and cellular components (**see Figure S12-14, Supplementary Table 4 )**.

**Log-transformed richness plots**

Species available on public datasets have varying levels of assembly and annotations and affect the number of complete ORFs we can recover in a species. The species with fewer complete ORFs or annotated amino acid sequences will show fewer overall repeats detected. We selected the LCRs with a purity of over 70% and pure LCRs and calculated the richness for each species based on their distinct residue composition. We visualized the log(richness) vs. log(number of LCRs) for each species in a clade-wise manner and calculated the slope of the fitted line, which is later visualized in the boxplot (**see Figure S26-45)**.

The richness of LCRs can be biased by the number of LCRs detected in each species. We divide the log(richness) by the log(number of LCRs) in each clade and plot the normalized count (**see Figure S46)**.

**Simpson’s diversity plots**

The Simpson’s Diversity Index is a pivotal metric in quantifying a community’s biodiversity or species richness. It comprehensively accounts for the number of species present and their corresponding relative abundances. The index attains a numerical value between 0 and 1, where higher values signify higher diversity.

To calculate Simpson’s diversity index, first, one calculates the proportional abundance of each species in the community and then calculates the index using the following formula:

D = 1 - ∑(pi)^2

where: D = Simpson’s diversity index

pi = proportional abundance of the ith species

We calculated the number and relative abundance of different amino acid repeats in R. Using this, we calculated the Simpson’s diversity index for each species and used the values for the boxplot. The boxplot shows the distribution of Simpson’s diversity index of all the species in that clade. Similarly, we computed the boxplot using Shannon’s diversity index (**see Figure S47-51)**.

Similarly, we also calculated richness, the slope of richness, and indices of diversity for exact LCRs (LCRs with purity 100%) (**see Figure S52-56)**.

**Stacked barplot**

Each clade can have a clade-specific abundance of particular LCR types. We list each clade’s twenty most abundant LCRs and calculate their proportion. We plot the stacked barplot using the proportion values of these LCRs by coloring each LCR distinctly (**see Fig. 3**). To make the proportion 1 in each clade, we add the proportion of all remaining LCRs. We generate the stacked barplot in R.

**Amino acid LCR distribution in a gene**

For each homopolymer LCR ( i.e., an LCR stretch consisting of only one type of amino acid), we calculate their abundance of occurrence on a normalized gene length. We selected all the LCRs of a particular amino acid from all the clades and converted the mid-position of the LCR to a normalized gene position. For example, if an L LCR stretches from 30 to 60 base-pair positions on a gene length of 1000. The mid-position of the LCR is 45 ( mid of 30 and 60 ) on the gene of length 1000. On normalized gene length (i.e., converting gene length to 100), the LCR’s mid-position becomes 4.5. We bin the normalized LCRs’ position in an interval of five, i.e., all the LCRs’ positions between 0 to 5 will be in the same bin.

We bin LCR positions normalized together, resulting in twenty bins. Next, we calculate the proportion of the LCR in each bin. Assuming a completely random distribution of amino acids, we expect each bin to have a proportion close to 0.05 (or 5%). We consider an LCR over-abundant in a bin if its occurrence exceeds 8% and an under-abundant if it falls below 2%. We calculated the distribution of each amino acid LCR in stratification of purity to better understand the change in selection pressure according to the purity (**see Figure S57-276)**. Similarly, we calculated the clade-specific distribution of LCRs in a normalized gene position (**see Figure S277-419)**.

**Comparison of %GC between genes containing LCRs and genes without LCRs**

Previous studies suggest a higher %GC of gene sequences with LCRs than without on a limited number of genes and species. Here, we calculated the %GC for each gene sequence and sorted the sequences according to “LCR status.” We consider a gene sequence with LCR if the LCR purity is >70%. Using a boxplot, we plotted the distribution of %GC of genes with and without LCRs for each clade (**see Figure S420)**. The distribution of %GC is compared using Wilcoxon’s test in R.

**Molecular evolutionary analyses**

Molecular evolutionary analyses require an aligned CDS multifasta file and its associated species tree. We downloaded the Aves clade’s species tree with 1000 bootstraps from BirdTree (Jetz et al. 2012) and the remaining clades’ species tree from TimeTree (Kumar et al. 2017). We generated a consensus species tree for the avian clade using the sumtree.py script. The species trees downloaded from TimeTree contain internode labels which can potentially create an error during downstream analyses. We removed the internode labels using a custom Linux script (mentioned below). All the species trees are pruned for each multifasta gene file and unrooted using a combination of Linux and R scripts.

```
###The birds’ tree was downloaded from Birdtree.org with 1000 bootstraps and is processed using sumtrees.py
grep “tree_” output.nex |awk '{print $4}' > onlytrees.nwk
sumtrees.py --rooted --ultrametric --ultrametricity-precision 10000 onlytrees.nwk > birds_mrc.tre
cat birds_mrc.tre |sed -e 's/\[[^][]*\]//g' > bird_mrc_tree_cleaned.tre
tail -n3 bird_mrc_tree_cleaned.tre |head -n1|awk '{print $4}' > birds_final.tre
cp birds_final.tre aves_list.nwk
sed -i 's/1.00000000//g' aves_list.nwk
##All the remaining species tree for clades were downloaded from TimeTree.org and are saved as “clade”_list.nwk
##NOTE: if the tree happen to have internode labels (denoted by numbers in single quotes), then remove them as PAML make its own internode labels
for t in `ls *_list.nwk`
do
sed -e "s/'[^()]*'//g" "$t" > temp.nwk
mv temp.nwk "$t"
done

for i in `ls *nwk`
do
j=`echo $i|sed 's/.nwk/_unrooted.nwk/g'`
echo 'library(ape)'  > unroot_tree.r
echo 'a<-read.tree("'$i'")' >> unroot_tree.r
echo 'b<-unroot(a)' >> unroot_tree.r
echo 'write.tree(b,file="'$j'")' >> unroot_tree.r
Rscript unroot_tree.r
done

###save this script as pruning.r
##run the script as follows:
Rscript pruning.r treename listofspeciestokeep.txt outputname
args=commandArgs(trailingOnly=TRUE)
t=args[1]
l=args[2]
o=args[3]
library(phytools)
a<-read.tree(t)
k<-read.table(l)
keep<-as.character(k$V1)
b<-keep.tip(a,keep)
write.tree(b,file=paste(o,".nwk",sep=''))
##########################################the folder “pruning” contains all the alignment files. We are going to prune species tree according to each gene file
ls|grep “_similar_length_orf.aln”> allfiles
##example of file name: A1BG_afrotheria_similar_length_orf.aln
cp allfiles allfiles_backup

for i in `cat allfiles`
o=`echo $i|sed 's/_similar_length_orf.aln//g'`
clade=`echo $i|cut -f2 -d"_"`
t=`ls "$clade"_list_unrooted.nwk`
grep ">" "$i"|sed 's/>//g' > "$o"_list
Rscript pruning.r $t "$o"_list "$o"
done
```

We compare site models M7 and M8 of PAML on aligned gene sequences to detect positively selected sites (PSSs). The sites with posterior probability > 0.95 are considered significant positive sites. Binning is performed on the positive sites based on normalized gene position in bins of 10. Using the branch-free model of PAML, we calculate the ω (dN/dS: non-synonymous substitution rate/synonymous substitution rate) for each lineage.

**Comparison of dN/dS (ω) between genes containing LCRs and genes without LCRs**

To study the effect of selection pressure on the retention/origin of LCR in a gene, we compared the ω values of genes with LCRs and genes without in a stratified manner of purity of LCRs (**see Figure S421-430)**. We subsetted all the lineages’ ω values according to their “LCR status,” compared using boxplot, and used Wilcoxon’s test for the significant difference in the distributions. We also compared the distributions using ω values cutoff of 2 (**see Figure S431-439)**. We show the barplots without outliers for better visualization. The value above the bars shows the p-value of Wilcoxon’s test. The values below the bar represent the number of sequences in that distribution (represented by “n”) and the mean of ω (represented by x-bar).

**Overlapping histograms of PSSs and LCRs**

We binned mid-positions of LCRs in intervals of 10 on a normalized gene position and depicted their frequency using a histogram. We repeated this process for positive sites and overlaid their frequency on the same histogram. To investigate the distribution of LCRs and PSSs, we subsetted the histograms based on clade and purity classes (**see Figure S440-477)**. Specifically, we analyzed the distribution of LCRs in three purity classes: all LCRs, LCRs with a purity greater than 70%, and pure LCRs (**see Figure S478-491)**.

**Correlation of alignment coverage, PSS, and LCR**

We analyzed the correlation between alignment coverage (i.e., the number of nucleotides for each position divided by the total number of sequences considered), positively selected sites, and low complexity regions (LCRs) using corrplots (**see Figure S492-504)**. We performed this analysis at the clade level. To visualize this relationship, we plotted the alignment coverage of aligned files against the normalized gene position and the abundance of LCRs and positively selected sites. To ensure visual consistency, we divided the alignment coverage values by 20 to align the y-axis margin with that of the LCR proportion and positively selected site frequency (**see Figure S505-517)**.

**Mosaic plots**

To gain insight into the co-occurrence of PSSs and LCRs, we investigated the proportion of genes containing PSSs, LCRs, and both. We employed mosaic plots in the R base to analyze these variables at the clade level. Additionally, we utilized Fisher’s test on the contingency table to assess the significance of the observed co-occurrence (**see Figure S518-529)**.

**Bedtools fisher**

We investigated the overlap between PSSs and LCRs at the gene level using bedtools fisher **(Supplementary Table 7)**. This tool allows us to evaluate whether the overlap between two regions in the genome is significant beyond chance. The test evaluates the p-value of significant favor for overlap and avoiding overlap between two regions in a genome. We extended the study clade-wise to understand whether the overlap of PSS and LCR is favored or avoided in a clade **(Supplementary Table 8)**.

**Data and code availability**

All the data and scripts used in generating the results are available on the GitHub link:

<https://github.com/ceglablokdeep/LCRs_global_patterns>

We have uploaded the entire dataset and the necessary scripts to Mendeley datasets, as some files exceed the 25MB limit for uploading on GitHub.
